# Interspecies isobaric labeling-based quantitative proteomics reveals protein changes in the ovary of *Aedes aegypti* co-infected with ZIKV and Wolbachia

**DOI:** 10.1101/2022.03.11.483997

**Authors:** Luis Felipe Costa Ramos, Michele Martins, Jimmy Rodriguez Murillo, Gilberto Barbosa Domont, Danielle Maria Perpétua de Oliveira, Fábio César Sousa Nogueira, Rafael Maciel-de-Freitas, Magno Junqueira

## Abstract

Zika is a vector-borne disease caused by an arbovirus (ZIKV) and overwhelmingly transmitted by *Ae. aegypti*. This disease is linked to adverse fetal outcomes, mostly microcephaly in newborns, and other clinical aspects such as acute febrile illness and neurologic complications, for example, Guillain-Barré syndrome. One of the actual most promising strategies to mitigate arbovirus transmission involves releasing *Ae. aegypti* mosquitoes carrying the maternally inherited endosymbiont bacteria *Wolbachia pipientis*. The presence of Wolbachia is associated with a reduced susceptibility to arboviruses and a fitness cost in mosquito life-history traits as fecundity and fertility. However, the mechanisms by which Wolbachia influences metabolic pathways leading to differences in egg production remains poorly known. To investigate the impact of co-infections on the reproductive tract of the mosquito, we applied an isobaric labeling-based quantitative proteomic strategy to investigate the influence of Wolbachia *w*Mel and ZIKV infection in *Ae. aegypti* ovaries. To the best of our knowledge, this is the most complete proteome of *Ae. aegypti* ovaries reported so far, with a total of 3,913 proteins identified, also,were able to quantify a total of 1,044 Wolbachia proteins in complex sample tissue of *Ae. aegypti* ovary.. Furthermore, we discuss proteins and pathways altered in *Ae. aegypti* during ZIKV infections, Wolbachia infections, co-infection Wolbachia/ZIKV, and compared with no infection, focusing on immune and reproductive aspects of *Ae. aegypti*. The modified aspects were mostly related to the immune priming enhancement by Wolbachia presence and the modulation of the Juvenile Hormone pathway caused by both microorganisms infection.

**Highlights:**

- Proteome changes in *Ae. aegypti*, Wolbachi*a, and* ZIKV interactions
- A great diversity of *Wolbachia* proteins were quantified in *Ae. aegypti* ovary
- Juvenile Hormone pathway is modulated by both infections
- *Wolbachia* enhances *Ae. aegypti* immune priming mechanism
- ZIKV unsettles host immune response by reducing antimicrobial peptides production
- Coinfection triggers oxidative stress and a lack of vitellogenin precursors

## 1 Introduction

Zika virus (ZIKV) is a mosquito-borne arbovirus from the Family Flaviviridae and is related to other arboviruses such as dengue (DENV), yellow fever (YFV), West Nile (WNV), and Japanese encephalitis (JEV) (Zimler et al., 2021). It was originally reported in a primate from the Zika forest in Uganda in 1947 (Dick et al., 1952), but the first large outbreak of ZIKV in humans occurred on the Pacific Island of Yap, in the Federated States of Micronesia in 2007 (Cao-Lormeau et al., 2014). In 2014, ZIKV emerged in the Pacific islands and a few years later invaded the Americas, being firstly identified in 2015 in Bahia state, Brazil (Campos et al., 2015). It became the first major infectious disease linked to adverse fetal outcomes, mostly microcephaly in newborns, and other clinical aspects such as acute febrile illness and neurologic complications, for example, Guillain-Barré syndrome. Such outcomes led the World Health Organization to claim a global public health emergency (Cao-Lormeau et al., 2014). There are no antivirus therapy or vaccines available so far to mitigate ZIKV transmission, i.e., the development of effective control methods targeting the mosquito vector must be encouraged.

The mosquito *Aedes aegypti* is the main vector of ZIKV worldwide (Chouin-Carneiro et al., 2016; Ferreira-de-Brito et al., 2016). During infection in female mosquitoes, the virus can be located in different tissues, but mostly in the gut, salivary glands, and ovaries (Sá-Guimarães et al., 2021). Ovaries infection is of paramount importance to maintain the vertical transmission of arboviruses, even if at low rates (Thangamani et al., 2016; Nag et al., 2021). Therefore, evaluating physiological and molecular changes in mosquito ovaries due to arbovirus infection, especially those affecting life-history traits related to vertical transmission, might provide new insights to design innovative vector control approaches.

Among the strategies to control arbovirus infection in *Ae. aegypti*, the use of the maternally-inherited endosymbiont *Wolbachia pipientis* is proving to be efficient due to a unique combination of phenotypes such as cytoplasmic incompatibility (CI), maternal transmission (MT), and viral-blocking (Moreira et al., 2009; Aliota et al., 2016; Dutra et al., 2016; Nazni et al., 2019; Indriani et al., 2020; Ryan et al., 2020; Mancini et al., 2020; Gesto et al., 2021; Utarini et al., 2021). The bacterium *Wolbachia* is naturally found in >60% of arthropod species, still, no natural infection is observed in *Ae. aegypti*, which led to experimental transinfection by microinjection of different *Wolbachia* strains into mosquito eggs (McMeniman & O’Neil, 2010). After transinfection, *Wolbachia* is currently located in several tissues and organs of *Ae. aegypti* mosquitoes and the mechanisms resulting in viral blocking are under investigation Most likely it involves the activation of the immune system by oxidative stress and down-regulation of host proteins involved in pathways that would help to produce resources essential to the virus life cycle (Ford et al., 2020; Martins et al., 2021; Ogunlade et al., 2021; Pimentel et al., 2021).

Besides its effects on arbovirus blocking, many studies showed that *Wolbachia* presence can present multifold effects on *Ae. aegypti* fitness that could later be an additional hurdle for *Wolbachia* establishment. For instance, *Wolbachia*-infected larvae have more rapid development and higher survivorship (de Oliveira S et al. 2017; Dutra et al. 2016) but reduced adult size (Ross et al., 2014), delayed embryogenic maturation, and *Wolbachia*-infected *Ae. aegypti* females have lower fecundity, fertility rates, quiescent eggs viability, and vector competence, affecting local invasion patterns (Dutra et al. 2015; King et al. 2018; McMeniman & O’Neil, 2010; Farnesi et al., 2019; Garcia et al. 2019; Garcia et al. 2020; Allman et al., 2020; Lau et al., 2021). Therefore, investigating the interaction network involving mosquito vectors, arboviruses and *Wolbachia* could help support the maintenance of a long-term stable blocking phenotype in endemic regions (Edenbourough et al. 2021).

The proteome can be considered more than a simple translation of the protein-coding regions of a genome, as some post-transcriptional and post-translation modifications generate even millions of proteoforms (Smith et al., 2013). Over the last decades, mass spectrometry-based proteomics has emerged as a powerful tool for the identification and quantification of the proteins contained in a biological sample. It has significantly contributed to unravelling many cellular and organism aspects (Schubert et al., 2017), and highlights features and emergent properties of complex systems under different conditions (Cox & Mann, 2011). To overcome the large overall experimental time and sample consumption of proteomics analysis, the isobaric labeling quantitative methods allowed the examination of multiple samples all at once. Moreover, it increases the throughput of quantification by having a higher multiplex capability, which makes it possible to handle several biological replicates, offering statistical robustness (Chen et al., 2022). A diversity of quantitative methods have already been successfully applied in insects to analyze differentially regulated proteins (De Mandal et al., 2020; García-Robles et al., 2020; Serteyn et al., 2020), including a previous study from our group, where it was able to elucidate different mechanisms responses to ZIKV and *Wolbachia* infection in *Ae. aegypti* head and salivary glands (Martins et al., 2021).

In this study, we apply isobaric labeling quantitative mass spectrometry-based proteomics to quantify proteins and identify pathways altered during either ZIKV or *Wolbachia* single infection and co-infection with *Wolbachia/*ZIKV in the *Ae. aegypti* ovaries. We show that it was possible to identify ZIKV peptides and more than 1,000 proteins from *Wolbachia* in mosquitoes’ ovaries. The present study offers a rich resource of data that is helpful to elucidate mechanisms by which *Wolbachia* and ZIKV infection in *Ae. aegypti* can interfere in immune and reproductive aspects.

## 2 Material and methods

### 2.1 Insects

To analyze the *Wolbachia* and ZIKV infection repercussions on *Ae. aegypti* proteome, mosquitoes were used from two different sites in Rio de Janeiro (Rio de Janeiro State, Brazil), with a 13km distance isolated from each other: Porto (22°53′43″ S, 43°11′03″ W) and Tubiacanga(22°47’06”S; 43°13’32”W). Porto is an area populated by mosquitoes without *Wolbachia* infection, i.e., wild type. On the other hand, Tubiacanga is the first site in Latin America where a *Wolbachia* (*w*Mel strain) invasion in *Ae. aegypti* was established, with more than 90% frequency (Garcia et al., 2019). 80 ovitraps were used for eggs collection (Codeço et al., 2015) with a net distance of 25-50 m apart from each other in both sites. Ovitraps were placed over an extensive geographic area to ensure we captured the local *Ae. aegypti* genetic variability, collecting at least 1500 eggs per site. The eggs were hatched and the mosquitoes were maintained at the insectary under a relative humidity of 80 ± 5% and a temperature of 25 ± 3°C, with *ad libitum* access to a 10% sucrose solution. Mosquitoes from the F1 generation were selected for experimental infection.

### 2.2 ZIKV strain and mosquitoes viral infection

*Aedes aegypti* females were orally infected with the ZIKV strain Asian genotype isolated from a patient in Rio de Janeiro (GenBank accession number KU926309). Local wild *Ae. aegypti* populations have high vector competence to this ZIKV strain (Fernandes et al., 2016; da Silveira et al., 2018; Petersen et al., 2018). All the assays were performed with samples containing 3.55 × 10^6^ PFU/ml (Martins et al. 2021). The experimental infection followed the protocol described in detail elsewhere (Martins et al. 2021). Briefly, 6-7 days old *Ae. aegypti* females from each of the two populations (Tubiacanga and Porto) were orally infected through a membrane feeding system (Hemotek, Great Harwood, UK), adapted with a pig-gut covering, which gives access to human blood. The infective blood meals consisted of 1 ml of the supernatant of infected cell culture, 2 ml of washed rabbit erythrocytes, and 0.5 mM of adenosine triphosphate (ATP) as phagostimulant. The same procedure and membrane feeding apparatus were used to feed control mosquitoes, but they received a noninfectious blood meal, with 1 ml of cell culture medium replacing the viral supernatant. After the experimental infection, we had a total of 152 *Ae. aegypti* females.

### 2.3 Ethical approval

*Ae. aegypti* colonies with *Wolbachia* are maintained in the lab by blood-feeding of anonymous donors acquired from the Rio de Janeiro State University blood bank. The blood bags were rejected from the bank due to small blood volume. No information of the donors (including sex, age, and clinical condition) was disclosed. The blood was screened for Dengue virus (DENV) using the Dengue NS1 Ag STRIP (Bio-Rad) before use in the mosquitoes feeding process. The use of human blood was approved by the Fiocruz Ethical Committee (CAAE 53419815.9.0000.5248).

### 2.4 Sample preparation and LC-MS/MS

A total of 152 *Ae. aegypti* mosquitoes were processed, in which 35 females were *Wolbachia*-infected (W), 42 infected exclusively with ZIKV (Z), 39 were co-infected with both *Wolbachia* and ZIKV (WZ), and 36 were non-infected with those microorganisms (A). At 14 days post-infection (dpi), each mosquito ovary was extracted from the body according to the previous method (Charlwood et al., 2018). Protein extraction, digestion, iTRAQ labeling and LC-MS/MS protocol were performed as described previously (Martins et al., 2021). Proteins were extracted by lysis with buffer (7 M urea, 2 M thiourea, 50 mM HEPES pH 8, 75 mM NaCl, 1 mM EDTA, 1 mM PMSF) with the addition of a protease/phosphatase inhibitor cocktail (Roche). Lysates were centrifuged and the supernatants were transferred to new tubes for protein quantification using the Qubit Protein Assay Kit® fluorometric (Invitrogen), following the manufacturer’s instructions. A total of 100 μg of proteins from each conditions were processed. Reduction and alkylation steps were performed using 10 mM dithiothreitol (DTT -GE Healthcare) and 40mM iodoacetamide solution (GE Healthcare), respectively. Samples were diluted 10x with 50 mM HEPES buffer, reducing the concentration of urea/thiourea, and incubated with trypsin (Promega) in a 1:50 (w/w, enzyme/protein) ratio at 37º C for 18 hours. The resulting peptides were desalted with a C-18 macro spin column (Harvard Apparatus) and then vacuum dried.

Peptides were labeledwith isobaric tags for relative and absolute quantitation (iTRAQ) 4-plex (ABSciex), was performed as described in Martins et al. (2021), mixing the tag solutions with sample peptides at room temperature for one hour. Each condition was labeled as follows i) Tag 114 corresponded to sample W (*Wolbachia* infected); ii) Tag 115 to sample A (none infection); iii) Tag 116 to sample WZ (*Wolbachia* and ZIKV co-infection); and iv) Tag 117 to sample Z (ZIKV infection). After the labeling step, samples were combined and vacuum dried for off-line fractionation by HILIC (Hydrophilic interaction liquid chromatography) prior toLC-MS/MS analysis. The dried samples were resuspended in acetonitrile (ACN) 90% / trifluoroacetic acid (TFA) 0.1% and injected into Shimadzu UFLC chromatography using a TSKGel Amide-80 column (15 cm x 2 mm i.d. x 3 μm - Supelco), which was equilibrated using ACN 85%/TFA 0. 1% (phase A). The peptides were eluted in TFA 0.1% (phase B) and every 8 fractions were collected and combined according to the separation and intensity of the peaks. The pools of fractions were dried in a speed vac and resuspended in 0.1% formic acid (FA). LC-MS/MS analysis was performed in an Easy-nLC 1000 coupled to a Q-Exactive Plus mass spectrometer (Thermo Scientific). Ionization was performed in an electrospray source with the acquisition of spectra in positive polarity by data-dependent acquisition (DDA) mode, spray voltage of 2.5 kV, and temperature of 200ºC in the heated capillary. The acquisition was set as follows: full scan or MS1 in the range of 375 - 1800 m/z, resolution of 70,000 (m/z 200), fragmentation of the 10 most intense ions in the HCD collision cell, with standardized collision energy (NCE) of 30, resolution of 17,000 in the acquisition of MS/MS spectra, the first mass of 110 m/z, isolation window of 2.0 m/z and dynamic exclusion of 45 s. A description of the workflow is presented in Figure 1.

**Figure 1:**
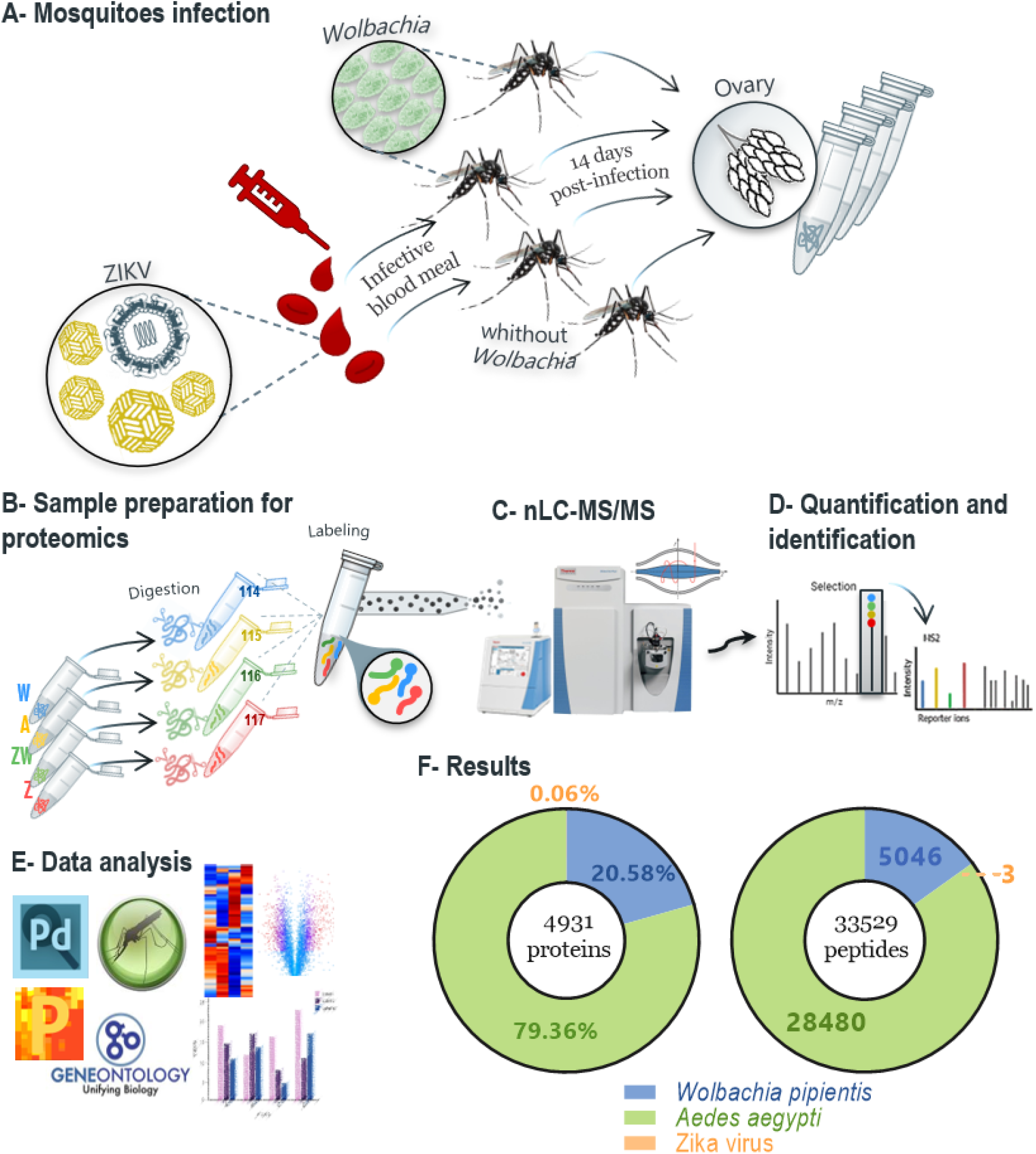
Workflow in the ovary of *Ae. aegypti* females, from four different conditions: A - non-infected mosquitoes; W - *Wolbachia* infected; Z - ZIKV infected; and ZW - *Wolbachia* and ZIKV infection. The peptides were labeled with iTRAQ-4plexand mixed in a 1:1:1:1 ratio; labeled peptides were fractionated offline using HILIC chromatography, and the pool of fractions was analyzed by nLC–MS/MS in Q-Exactive Plus mass spectrometer. Fragmentation scheme shows reporter ions at the low m/z regionused for relative quantification of the peptides/proteins.Finally, we performed statistical analysis in software Perseus and bioinformatics analysis using Vectorbase to enrich proteins differentially regulated for GO and KEGG. A total of 4931 proteins and 33529 peptides were identified including the three organisms.

### 2.5 Data analysis and gene enrichment

The spectra obtained after the LC-MS/MS analyzes were processed in the Proteome Discoverer 2.4 Software (Thermo Scientific), with the Sequest HT search engine against the *Ae. aegypti* (genome version/assembly ID: INSDC: GCA_002204515.1), ZIKV and *Wolbachia* provided by VectorBase (Giraldo-Calderón et al., 2014), ViPR (Pickett et al., 2012), and UniProt (The UniProt Consortium, 2015). 2019), respectively. For the search, the following parameters were used: precursor tolerance of 10 ppm, fragment tolerance of 0.1 Da, tryptic cleavage specificity, two maximum missed cleavage sites allowed, fixed modification of carbamidomethyl (Cys), variable modification of iTRAQ 4-plex (Tyr, Lys and peptide N-terminus), phosphate (Ser, Thr, and Tyr) and oxidation (Met). Peptides with high confidence were selected, and only identifications with q values equal to or less than 0.01 FDR were considered. Target Decoy PSM obtained these values.

Quantitative analysis was performed in Perseus software, version 1.6.12.0 (Tyanova et al., 2016), based on the intensity of the reporter ionsextracted from the MS/MS spectra. All data were transformed into log2 and normalized by subtracting the column median. As a criterion to define the differential proteins, statistical Anova tests were performed between the groups, with a significance level of 0.05 of the p-value. Proteins with p < 0.05 were considered significant with a fold change threshold of 0.5. Enrichment of biological process by gene ontology and metabolic pathways by KEGG for differentially expressed proteins was performed using Fisher’s Exact Test (p<0.05) on the VectorBase website (bit.ly/38OmEX0) (Giraldo-Calderón et al., 2014; Ashburner et al., 2000).

## 3. Results and discussion

### 3.1 Proteome identification and quantification

Isobaric-labeled quantitative proteomics was applied to increase the depth coverage and efficiency of proteomics analysis. By labeling mosquitoes from each of the four experimental groups with iTRAQ 4-plex, samples were multiplexed before off-line fractionation and LC-MS/MS analysis (Figure 1). We identified 4,931 protein groups (Supplementary Table 1), 36,403 peptides, 110733 PSMs and 987,698 MS/MS spectra. The protein search was performed using *Ae. aegypti*, ZIKV, and *Wolbachia* databases simultaneously. Considering all identified proteins, 3,913 belong to *Ae. aegypti*, 1015 to *Wolbachia*, and 3 unique peptides with a total of 3 PSMs to the ZIKV polyprotein.

After checking the quantification using the report ion signals’ abundance (Figure 2A), it is possible to observe that in samples A and Z, the signals’ abundance is very low, showing the signal-to-noise (S/N) behavior of iTRAQ reporter ions for the channels where *Wolbachia* protein signals are not expected. This control is important to show that *Wolbachia* protein identifications were not random matches, but actually, they followed perfectly the quantification trend expected: only relevant signals in the channels with *Wolbachia* infection. Since in W and WZ, where we actually have *Wolbachia* infected samples, there are higher intensity and relevant signals of bacterium proteins revealing the match between identification and quantification pattern of *Wolbachia* proteins.

**Figure 2.**
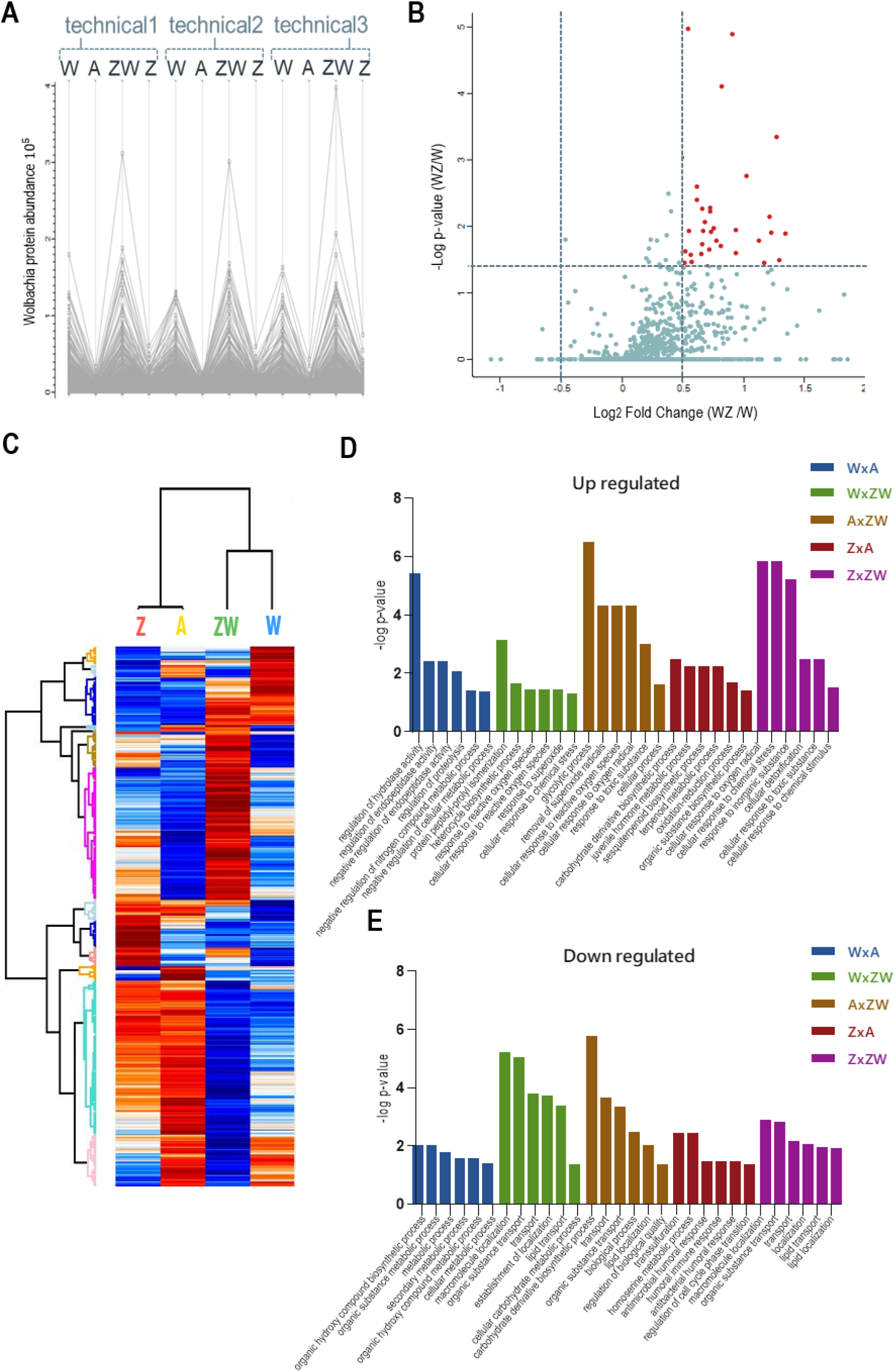
Quantitative proteomics data. A: Abundance chart of the three replicates of identified *Wolbachia pipientis* proteins. High expression is observed in the W and ZW samples, showing that it is possible to compare the expression of proteins by the relative quantification of the iTRAQ; B: Volcano plot of Wolbachia proteins comparing WZ/W. The red dots represent upregulated proteins; C: Heatmap of all Ae aegypti identified proteins that were upregulated (red) or downregulated (blue) during comparison between control, monoinfected and coinfected samples; D:

#### 3.1.1 *Wolbachia* localization in insect ovary and its protein identification

The accumulation of *Wolbachia* in ovaries improves its vertical transmission efficiency through the female germline, suggesting that a balance between vertical transmission and optimum densities is critical for a long-term, stable endosymbiosis (Kaur et al., 2021). Since only 1205 protein-coding genes were reported in the *Wolbachia pipientis* genome (Sinha et al., 2019), we applied an identification strategy using a combined database containing Uniprot proteins addressed to *Wolbachia* species in general. Using this strategy, we were able to identify 1015 unique protein groups from *Wolbachia*. As an additional analysis, we performed a comparison between W and WZ to observe possible *Wolbachia* proteins that might be modulated during ZIKV infection (Figure 2B). Proteins with p < 0.05 were considered significant with a fold change threshold of 0.5. A total of 38 proteins were upregulated in response to virus infection (Table 1), mostly related to catalytic activity, structure activity, metal ion binding, nucleotide binding, RNA and DNA binding. Similar findings were noted after evaluating that the transcriptional response of *Wolbachia* to DENV involves the regulation of *Wolbachia* DNA production and energetic metabolic genes (Leitner et al., 2021).

**Table 1.** *Wolbachia* proteins with significant differences in abundance between W and WZ.

| Protein | Student's<br>T-test<br>p-value<br>WZ_W | Fold<br>change<br>(-log<br>WZ/W) |
| --- | --- | --- |
| Pyruvate, phosphate dikinase | 0.0000105157 | 0.5575689 |
| Leucine--tRNA ligase | 0.0000126546 | 0.9179640 |
| Cytoplasmic incompatibility factor CifA | 0.0000779842 | 0.8362770 |
| Bifunctional DNA-directed RNA polymerase subunit beta-beta' | 0.000444461 | 1.2826614 |
| MgtE intracellular N domain | 0.000908949 | 0.5046193 |
| OmpA family protein | 0.00173934 | 1.0416403 |
| ATP-dependent chaperone ClpB | 0.002478 | 0.6262922 |
| Putative pyruvate, phosphate dikinase regulatory protein | 0.00401212 | 0.6302490 |
| Inositol monophosphatase | 0.00520698 | 0.7422536 |
| Aspartate--tRNA(Asp/Asn) ligase | 0.00534456 | 0.6737426 |
| Malonyl CoA-acyl carrier protein transacylase | 0.00581913 | 0.7404013 |
| Isoleucine--tRNA ligase | 0.00713402 | 1.2248519 |
| Type IV secretion system protein VirB6 | 0.00857789 | 0.6956878 |
| ATP-dependent zinc metalloprotease FtsH | 0.0104455 | 0.7701270 |
| Citrate synthase | 0.0111278 | 0.9538174 |
| Surface antigen, Wsp paralog | 0.0114699 | 0.6838690 |
| Fido domain-containing protein | 0.0115553 | 0.5643422 |
| Porin_4 domain-containing protein | 0.0120586 | 0.7468278 |
| DNA mismatch repair protein MutS | 0.0122699 | 1.2434697 |
| Ankyrin repeat domain protein | 0.0125616 | 1.3596706 |
| Replicative DNA helicase | 0.0159918 | 0.7936010 |
| Hypothetical protein | 0.0160156 | 1.1424662 |
| Outer membrane protein assembly factor BamA | 0.0184722 | 0.6721821 |
| Peptidyl-prolyl cis-trans isomerase | 0.0192305 | 0.8291993 |
| Ribosome-recycling factor | 0.0217489 | 0.7295354 |
| Iron(III) transport system substrate-binding protein | 0.0234158 | 0.5369954 |
| Hypothetical protein | 0.0249885 | 0.9480012 |
| NADH-quinone oxidoreductase subunit C | 0.0257139 | 0.6636648 |
| 50S ribosomal protein L3 | 0.0262632 | 0.5757586 |
| Protein HflC | 0.0318682 | 1.3096765 |
| 50S ribosomal protein L1 | 0.0344079 | 0.5853783 |
| Transcription elongation factor GreA | 0.0344705 | 1.1813854 |
| RmlD_sub_bind domain-containing protein (Fragment) | 0.0349948 | 0.5294895 |
| 50S ribosomal protein L5 | 0.0395048 | 0.7204612 |
| Folate-binding protein YgfZ | 0.0420299 | 0.5038770 |
| ATP synthase gamma chain | 0.0422139 | 0.8000174 |
| Phosphatidylserine decarboxylase proenzyme | 0.0450936 | 1.2252582 |
| Magnesium transporter MgtE | 0.0490137 | 1.1998873 |
| Insulinase family protein | 0.0490153 | 0.5579723 |

One of the positive modulated proteins in WZ compared to W was the Cytoplasmic incompatibility factor (CifA). LePage et al. (2017) concluded that the cytoplasmic incompatibility factor genes enhanced the cytoplasmic incompatibility leading to an embryonic lethality in *D. melanogaster*. The gene additively, through transgenic mechanisms, augments embryonic lethality in crosses between infected males and uninfected females after pioneering genetic studies. The discovery of cifA pioneers genetic studies of prophage WO-induced reproductive manipulations and informs the continuing use of Wolbachia to control dengue and Zika virus transmission to humans.

#### 3.1.2 ZIKV polyprotein peptides were identified in mosquito ovaries

A total of three ZIKV peptides were identified in mosquito ovaries using the Isobaric-labeled quantitative proteomics approach: VPAETLHGTVTVEVQYAGNDGPCKIPVQMAVDMQTLTPVGR, NGGYVSAITQGRREEETPVECFEPSMLK, and DGDIGAVALDYPAGTSGSPILDR. No protein quantification data was obtained despite its identification in ovary tissue. This result leads to two different hypotheses: ZIKV was not located in mosquito ovaries or there was a low amount of ZIKV polyprotein peptides that would be followed by a poor quantification in this tissue.

Vector competence is a key feature to determine the likelihood of pathogen transmission in a given area and has hitherto shown heterogeneous results among field populations (Epelboin et al. 2017; Kaufmann and Kramer 2017; Boyer et al. 2018). ZIKV disseminates through several organs and tissues of mosquitoes, including the ovaries to support further vertical transmission (Campos et al., 2017; Li et al., 2017; González et al. 2019; Lai et al., 2020; Sá-Guimarães et al. 2021). The infection of ovaries by ZIKV is mostly dependent on mosquito genetics, temperature, viral load and the availability of blood meals but peaks around 18 dpi (Thangamani et al., 2016; Nag et al., 2021). Sá-Guimarães et al. (2021) identified ZIKV in *Ae. aegypti* ovaries at 3 dpi after orally challenging insects with a 2.5×10^9^ PFU/ml viral load whereas our ZIKV isolate was 3.55×10^6^ PFU/ml. Intuitively, the higher viral load used by Sá-Guimarães et al. (2021) and the longer incubation period before ovaries dissection by Nag et al. (2021) together might explain partially the lack of ZIKV peptides quantification in *Ae. aegypti* ovaries in our study. Nevertheless, we achieved ZIKV identification in ovaries and revealed that ZIKV mono-infection modulated several proteins in *Ae. aegypti* ovaries that will be further discussed.

### 3.2 Modulated proteins and pathways by infections

A total of 480 proteins from *Ae. aegypti* with significant differences were statistically determined by the ANOVA test. Using the Tukey post-test ANOVA, we defined pairs of proteins with significant differences between the groups (Supplementary Table 1) (Figure 2C). Pathways were enriched using Vectorbase software and the Gene Ontology/KEGG list is represented in Supplementary Table 2. Pathways related to reproductive and immune aspects were the main interest in this data and were the focus of discussion (Figure 2D) and each pathway modulated by discussed proteins is represented in Table 2.

**Table 2:** Biological processes enriched with respect to specific differential proteins: farnesol dehydrogenase, 26S protease regulatory subunits, mevalonate kinase, yellow protein,macroglobulin complement protein, Cu-Zn superoxide dismutase (SOD) proteins and vitellogenin A1.

| farnesol dehydrogenase | 26S protease regulatory subunit |
| --- | --- |
| juvenile hormone metabolic process (GO:0006716) | related to protein catabolic process (GO:0030163) |
| cellular hormone metabolic process (GO:0034754) | regulation of proteolysis (GO:0030162) |
| juvenile hormone biosynthetic process (GO:0006718) | positive regulation of catabolic process (GO:0009896) |
| sesquiterpenoid biosynthetic process (GO:0016106) | positive regulation of cellular protein catabolic process (GO:1903364) |
| terpenoid biosynthetic process (GO:0016114) | positive regulation of proteolysis involved in cellular protein catabolic process (GO:1903052) |
| terpenoid metabolic process (GO:0006721) | positive regulation of protein catabolic process (GO:0045732) |
| sesquiterpenoid metabolic process (GO:0006714) | positive regulation of proteasomal protein catabolic process (GO:1901800) |
| hormone metabolic process (GO:0042445) | positive regulation of cellular catabolic process (GO:0031331) |
| hormone biosynthetic process (GO:0042446) | regulation of proteolysis involved in cellular protein |
|  | catabolic process (GO:1903050) |
| regulation of hormone levels (GO:0010817) | regulation of cellular protein catabolic process (GO:1903362) |
| oxidation-reduction process (GO:0055114) | regulation of proteasomal protein catabolic process (GO:0061136) |
| cellular biosynthetic process (GO:0044249) | positive regulation of proteolysis (GO:0045862) |
| organic substance biosynthetic process (GO:1901576) | regulation of protein catabolic process (GO:0042176) |
| **yellow protein** | macromolecule catabolic process (GO:0009057) |
| organic hydroxy compound biosynthetic process (GO:1901617) | **macroglobulin complement protein** |
| organic substance metabolic process (GO:0071704) | regulation of hydrolase activity (GO:0051336 and GO:0051346) |
| mating behavior (GO:0007617) | regulation of catalytic activity (GO:0050790 and GO:0043086) |
| reproductive behavior (GO:0019098) | regulation of molecular function (GO:0065009 and GO:0044092) |
| melanin biosynthetic process (GO:0042438) | regulation of endopeptidase activity (GO:0052548 and GO:0010951) |
| melanin metabolic process (GO:0006582) | regulation of peptidase activity (GO:0052547 and GO:0010466) |
| metabolic process (GO:0008152) | regulation of proteolysis (GO:0030162 and GO:0045861) |
| mating (GO:0007618) | negative regulation of protein metabolic process (GO:0051248) |
| secondary metabolic process (GO:0019748) | negative regulation of cellular protein metabolic process (GO:0032269) |
| phenol-containing compound biosynthetic process (GO:0046189) | negative regulation of cellular process (GO:0048523) |
| secondary metabolite biosynthetic process (GO:0044550) | negative regulation of nitrogen compound metabolic process (GO:0051172) |
| organic hydroxy compound metabolic process (GO:1901615) | negative regulation of biological process (GO:0048519) |
| behavior (GO:0007610) | negative regulation of cellular metabolic process (GO:0031324) |
| phenol-containing compound metabolic process | **Cu-Zn superoxide dismutase (SOD)** |
| (GO:0018958) | **proteins** |
| cellular metabolic process (GO:0044237) | cellular response to oxygen radical (GO:0071450) |
| **mevalonate kinase protein** | response to reactive oxygen species (GO:0000302) |
| purine nucleotide metabolic process (GO:0006163) | cellular response to superoxide (GO:0071451) |
| purine ribonucleotide metabolic process (GO:0009150) | response to superoxide (GO:0000303) |
| ribose phosphate metabolic process (GO:0019693) | response to oxygen radical (GO:0000305) |
| ribonucleotide metabolic process (GO:0009259) | removal of superoxide radicals (GO:0019430) |
| organic substance catabolic process (GO:1901575) | cellular response to reactive oxygen species (GO:0034614) |
| purine-containing compound metabolic process (GO:0072521) | cellular response to oxidative stress (GO:0034599) |
| nucleotide metabolic process (GO:0009117) | cellular response to chemical stress (GO:0062197) |
| nucleoside phosphate metabolic process (GO:0006753) | response to inorganic substance (GO:0010035) |
| organonitrogen compound metabolic process (GO:1901564) | superoxide metabolic process (GO:0006801) |
| nucleobase-containing small molecule metabolic process (GO:0055086) | reactive oxygen species metabolic process (GO:0072593) |
| organophosphate metabolic process (GO:0019637) | cellular response to oxygen-containing compound (GO:1901701) |
| isoprenoid biosynthetic process (GO:0008299) | response to oxygen-containing compound (GO:1901700) |
| isoprenoid metabolic process (GO:0006720) | metabolic process (GO:0008152) |
| primary metabolic process (GO:0044238) | response to oxidative stress (GO:0006979) |
| organic hydroxy compound biosynthetic process (GO:1901617) | cellular oxidant detoxification (GO:0098869) |
| organic substance metabolic process (GO:0071704) | detoxification (GO:0098754) |
| isopentenyl diphosphate biosynthetic process | cellular detoxification (GO:1990748) |
| mevalonate pathway (GO:0019287) | cellular response to toxic substance (GO:0097237) |
| isopentenyl diphosphate metabolic process (GO:0046490) | response to toxic substance (GO:0009636) |
| isopentenyl diphosphate biosynthetic process (GO:0009240) | oxidation-reduction process (GO:0055114) |
| metabolic process (GO:0008152) | cellular metabolic process (GO:0044237) |
| carbohydrate derivative metabolic process (GO:1901135) | cellular response to chemical stimulus (GO:0070887) |
| organic hydroxy compound metabolic process (GO:1901615) | **vitellogenin A1 protein** |
| nitrogen compound metabolic process (GO:0006807) | macromolecule localization (GO:0033036) |
| sterol biosynthetic process (GO:0016126) | organic substance transport (GO:0071702) |
| sulfur compound metabolic process (GO:0006790) | biological process (GO:0008150) |
| cellular metabolic process (GO:0044237) | transport (GO:0006810) |
| steroid biosynthetic process (GO:0006694) | establishment of localization (GO:0051234) |
| sterol metabolic process (GO:0016125) | localization (GO:0051179) |
| steroid metabolic process (GO:0008202) | lipid transport (GO:0006869) |
| acetyl-CoA metabolic process (GO:0006084) | lipid localization (GO:0010876) |

#### 3.2.1 ZIKV mono-infection controls mosquito machinery to favor its replication and transmission

##### 3.2.1.1 ZIKV infection downregulates antimicrobial peptides and mannose-binding C-type lectin

During the investigation of modulated proteins in ZIKV mono-infection samples compared to control, it was possible to identify attacin (AAEL003389), included in the antimicrobial humoral response (GO:0019730), humoral immune response (GO:0006959), and antibacterial humoral response (GO:0019731) pathways, and gambicin (AAEL004522) downregulation. Both molecules are classified as antimicrobial peptides (AMPs), crucial effectors of the insect’s innate immune system that can provide the first line of defense against a variety of pathogens (Wu et al., 2018). It is widely known that AMPs are normally produced by the Toll/IMD signaling pathway, which is the case of attacins (Buonocore et al., 2021), or can also be produced by JAK-STAT pathway, as gambicin (Zhang et al., 2017). The probable reason that explains its downregulation is the influence of AMPs in virus infection, that difficult DENV and ZIKV establishment in *Ae aegypti* (Xi et al., 2008; Martins et al., 2021). In addition to attacin and gambicin, RpS23: 40S ribosomal protein S23 (AAEL012686) was also downregulated, and it was recently discovered that new AMP functioning as a thev capable of recognizing microorganisms molecules, and an effector, capable of killing the potential pathogens (Ma et al, 2020).

Besides AMPs, a mannose-binding C-type lectin (AAEL000563) was also downregulated. C-type lectins (CTLs) are a family of proteins that contain characteristic modules of carbohydrate recognition domains and play important roles in insect immune responses,such as opsonization, nodule formation, agglutination, encapsulation, melanization, and prophenoloxidase activation (Xia et al., 2018). A more specific function related to mannose-binding lectins is a pattern recognition component of the complement system that binds carbohydrate groups on the surface of microbial pathogens, triggering the lectin activation pathway of complement (Takahashi et al., 2006). It is described that this pathway can neutralize DENV and West Nile Virus (WNV) infection (Fuchs et al., 2011; Avirutnan et al., 2011). Moreover, it was recently described that CTLs can act as a recognition receptor for JAK-STAT immune pathway (Geng et al., 2021), which downregulation may interfere in AMPs production, as gambicin peptide. Those down-regulations combined can facilitate ZIKV infection.

##### 3.2.1.2 ZIKV enhance Juvenile Hormone production and pro-viral host factors for establishing infection

Juvenile Hormone (JH), a representant from a family of sesquiterpenoid hormones in insects, was originally described in *Rhodnius prolixus* as a molecule capable of maintaining the juvenile character of insect larvae to ensure proper metamorphosis timing (Wigglesworth, 1934). However, JHs govern many insects essential aspects of development, metamorphosis and reproduction (Tsang et al., 2020), and its absence in vertebrates may qualify this hormone as a target to control insect pests and disease vectors (Jindra et al., 2015). Its signaling and production pathways comprehend the interaction between insects’ neuronal and fat body organs, and the main effect in female ovaries is the vitellogenin production activation (Santos et al., 2019). Besides, Chang et al. (2021) recently described that JH acts in AMP negative regulation, especially after the post-eclosion phase of the *Ae. aegypti* female gonadotrophic reproductive cycle. In regards to *Ae. aegypti*, it was described that JH analogs enhance ZIKV infection (Alomar et al., 2021). On the other hand, the silencing of ribosomal protein (Rp) genes, responsive to JH, increased ZIKV blocking, once the virus increased global ribosomal activity in the insect (Shi et al, 2021). Taking all of that information into account, it is important to understand JH modulation during ZIKV infection and if *Wolbachia* infection plays any changes as well.

Evaluating our data, the first aspect that will be discussed is ZIKV upregulation in the JH production pathway, as a farnesol dehydrogenase (AAEL017302) was upregulated (Figure 3). Farnesol dehydrogenase was found in *Ae. Aegypti* (Mayoral et al., 2009), and its activity is observed on the second JH branch, oxidizing farnesol to farnesal (Nouzova et al. 2011). It is considered a rate-limiting enzyme and critical in regulating the production of JH in adult mosquitoes (Zifruddin et al., 2021), so its enhancement in ZIKV infection can lead to JH production, as previously described in the literature and exposed in the last paragraph, favoring mosquito reproduction aspects and maybe helping ZIKV vertical and horizontal transmission. Experiments involving farnesol dehydrogenase inhibition showed larvicidal activity and inhibited the ovary growth of female *Ae. albopictus* (Park et al. 2020). Farnesol dehydrogenase modulated several pathways in our analysis: juvenile hormone metabolic process (GO:0006716), cellular hormone metabolic process (GO:0034754), juvenile hormone biosynthetic process (GO:0006718), sesquiterpenoid biosynthetic process (GO:0016106), terpenoid biosynthetic process (GO:0016114), terpenoid metabolic process (GO:0006721), sesquiterpenoid metabolic process (GO:0006714), hormone metabolic process (GO:0042445), hormone biosynthetic process (GO:0042446), regulation of hormone levels (GO:0010817), oxidation-reduction process (GO:0055114), cellular biosynthetic process (GO:0044249) and organic substance biosynthetic process (GO:1901576).

**Figure 3.**
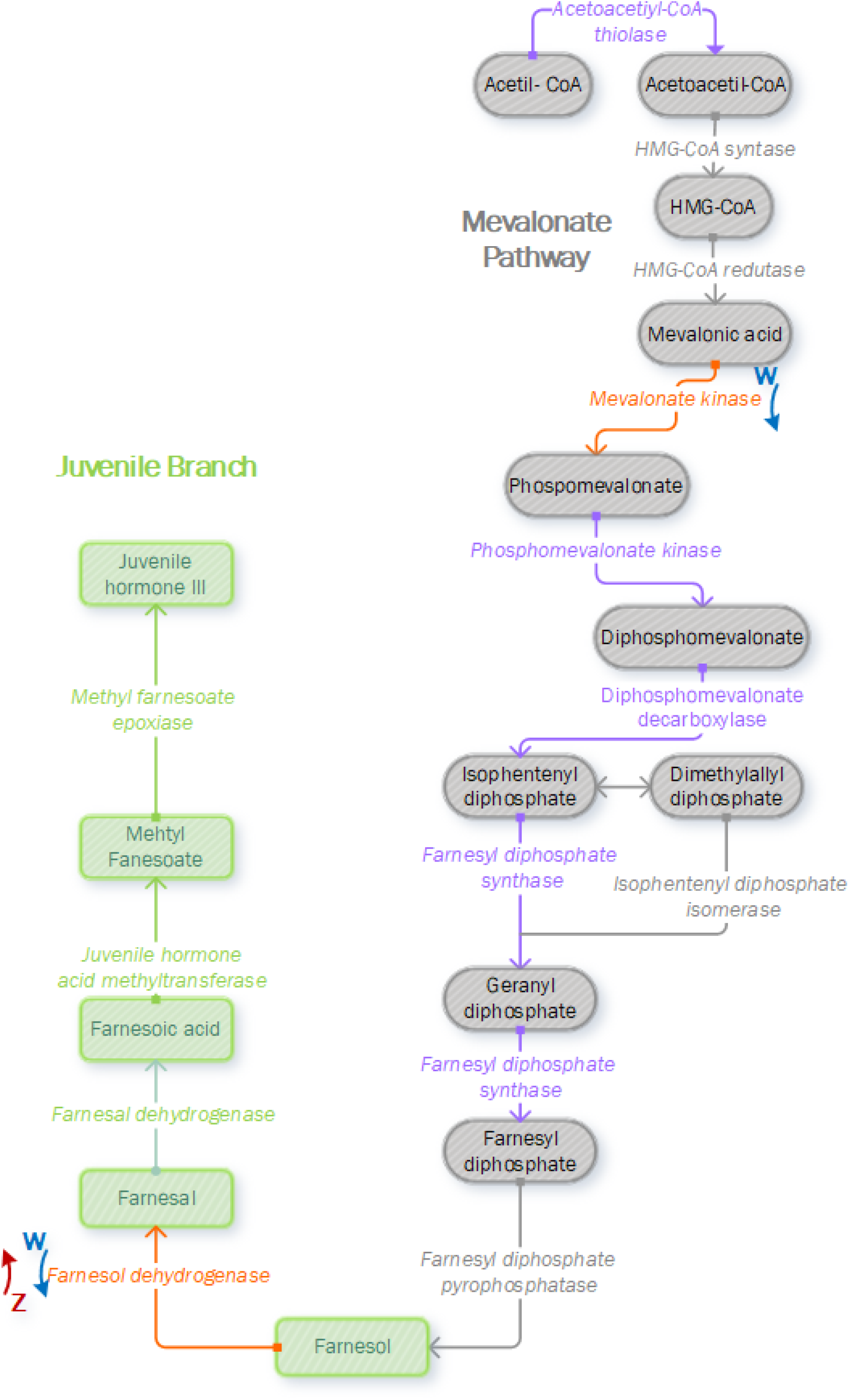
Identification and modulation of proteins that participate in the JH pathway, composed by the mevalonate pathway (early step), represented in gray, and juvenile hormone branch (last step), represented in green (pathway adapted from Nouzova et al., 2011). Enzymes colored in purple were identified in *Ae. aegypti* proteome in this work, and enzymes colored in orange were identified and quantified as modulated during ZIKV or *Wolbachia* infection. Mevalonate kinase and Farnesol dehydrogenase were downregulated in *Wolbachia* infection (blue arrow with W letter) and Farnesol dehydrogenase was upregulated in ZIKV infection (red arrow with Z letter).

ZIKV-capsid interaction with cell host was newly investigated through quantitative label-free proteomics and showed an important contribution of the 26S protease regulatory subunit (Gestuveo et al., 2021), a component of the ubiquitin-proteasome system. We were able to identify two 26S protease regulatory subunit (AAEL002508 and AAEL012943) upregulated, related to protein catabolic process (GO:0030163), regulation of proteolysis (GO:0030162), positive regulation of catabolic process (GO:0009896), positive regulation of cellular protein catabolic process (GO:1903364), positive regulation of proteolysis involved in cellular protein catabolic process (GO:1903052), positive regulation of protein catabolic process (GO:0045732), positive regulation of proteasomal protein catabolic process (GO:1901800), positive regulation of cellular catabolic process (GO:0031331), regulation of proteolysis involved in cellular protein catabolic process (GO:1903050), regulation of cellular protein catabolic process (GO:1903362), regulation of proteasomal protein catabolic process (GO:0061136), positive regulation of proteolysis (GO:0045862), regulation of protein catabolic process (GO:0042176) and macromolecule catabolic process (GO:0009057). It is already described 26S protease regulatory subunit importance in ZIKV infection but the mechanism is currently unknown, but there is a strong relation with ubiquitination processes in other viruses and can help in viral capsid transport to the nucleus, for example (Schneider et al., 2020). Moreover, this ribosomal activity increases the proteolytic activity of the proteasome, which is required for female insect reproduction (Wang et al., 2021), it is sensible to JH and enhances viral replication in *Ae. aegypti* as observed by Shi et al. (2021). Two ribosomal proteins, RpL27a (AAEL013272) and RpL17 (AAEL000180) were also upregulated, related to the organonitrogen compound biosynthetic process (GO:1901566), cellular biosynthetic process (GO:0044249), and organic substance biosynthetic process (GO:1901576).

#### 3.2.2 *Wolbachia* harms mosquito reproductive characteristics but helps its immune system

##### 3.2.2.1 Juvenile Hormone pathway is negatively affected by *Wolbachia* mono-infection

While ZIKV infection induces JH, *Wolbachia* infection downregulates two proteins related to its pathway (Figure 3). One of them is in the same farnesol dehydrogenase (AAEL017302) discussed before, which will affect the second part of JH output cascade. The other protein identified as downregulates is mevalonate kinase (AAEL006435), which modulated pathways are related in Table 2. Mevalonate kinase belongs to the fourth reaction step of the mevalonate pathway, which is responsible for the biosynthesis of many essential molecules important in insect development, reproduction, chemical communication, and defense (Li et al., 2016). Concerning JH, the mevalonate pathway is classified as the early step that forms farnesyl pyrophosphate used on the late step, known as JH branch (Noriega, 2014). Both mevalonate kinase and farnesol dehydrogenase downregulation in *Wolbachia* presence may infer that JH production is negatively affected and this can impact the germline lifecycle from meiosis to gametogenesis, once it was observed that JHs influence embryonic reproductive development (Barton et al., 2021). Both farnesol dehydrogenase and mevalonate kinase have modulated pathways that interact with themselves, and are related to hormone synthesis (Figure 4).

**Figure 4.**
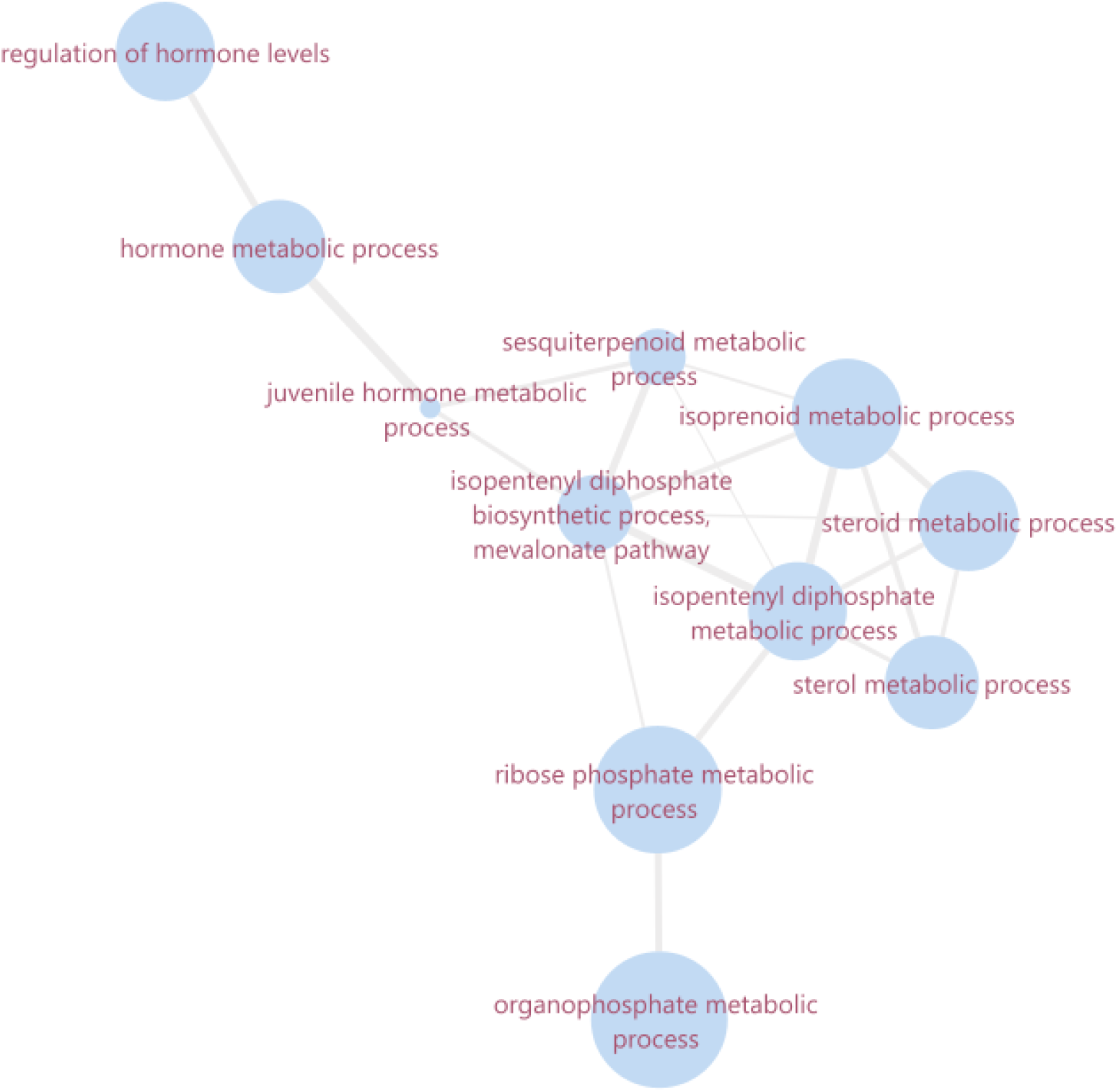
Ontology network of summarized enriched biological processes related to mevalonate kinase and farnesol dehydrogenase. The size of the bubble corresponds to the LogSize value for the GO Term. The overview and the interaction network were obtained in software revigo and cytoscape 3.8.0, respectively.

Allied to this event, a yellow protein was also downregulated. Yellow protein (dopachrome conversion enzyme) is involved in the melanin biosynthetic pathway that significantly accelerates pigmentation reactions in insects and belongs to a rapidly evolving gene family generating functionally diverse paralogs with physiological functions still not understood (Noh et al., 2015). Noh et al. (2020) discovered that two yellow proteins are required for egg desiccation resistance so maybe *Wolbachia* downregulation may decrease eggs resistance leading to a lower reproduction success, which was already described before (McMeniman et al., 2011).

##### 3.2.2.2 Host immune defense is activated by *Wolbachia* monoinfection as a protection barrier to other microorganisms

Microorganisms are important modulators of host phenotype, providing heritable variation upon which natural selection acts (Brinker et al. 2019). Host–parasite interactions represent one of the strongest selection pressures in nature, with considerable impact on the ecology and evolution of parasites and thus on disease epidemiology (Bose and Schulte 2014). The endosymbiotic bacteria *Wolbachia* has been showing an increase in host protection range against pathogens including bacteria, viruses, nematodes and the malaria parasite (Wong et al., 2011). A single mechanism called immune priming might explain this broad-based pathogen protection, in which *Wolbachia* presence upregulates the basal immune response, preparing the insect to defend against subsequent pathogen infection (Ye et al., 2013).

It was observed in our results that macroglobulin complement protein (AAEL001794) was upregulated during *Wolbachia* infection compared to control. Macroglobulin complement-related factor (MCR) is known for being a thioester-containing proteins (TEP) family component in insects, presenting a protease inhibitor activity (Blandin & Levashina, 2004). TEPs were previously described in *Anopheles gambiae* and *Drosophila melanogaster* models, and act in the innate immune response by promoting recruitment of immune cells, phagocytosis, and direct lysis of microbial invaders (Shokal & Eleftherianos, 2017), usually in insects hemolymph (Cheng et al., 2016; Mukherjee et al., 2019). Xiao et al. (2014) performed molecular biology assays evolving MCR in *Ae. aegypti* containing *Wolbachia* during DENV infection, describing that MCR does not directly interact with the flavivirus, requiring a mosquito homologue of scavenger receptor-C (SR-C), which interacts with DENV and MCR simultaneously (*in vitro* and *in vivo*) and this SR-C/MCR axis regulates the expression of antimicrobial peptides (AMPs) consequently increasing anti-DENV immune response.

In addition to MCR, three leucine-rich repeat immune protein (LRRIM) were upregulated during *Wolbachia* infection: LRRIM1, LRRIM8 and LRRIM10A (AAEL012086, AAEL001420, AAEL001401, respectively). LRRIM is evolutionarily conserved among many proteins correlated with innate immunity in an array of organisms, including invertebrates, and it usually forms a disulfide-bridged complex that interacts with the third factor, an TEP, more specifically TEP1, revealing its antimicrobial activity in insect immune response (Waterhouse et al., 2010). More importantly, it was noticed that LRRIM responded to ZIKV infection in *Ae. aegypti* female adults in a transcriptional analysis approach (Zhao et al., 2019; Shi et al., 2021), therefore corroborating with the idea that LRRIM enhancement mediated by *Wolbachia* presence can help in ZIKV block.

A DEAD box ATP-dependent RNA helicase (AAEL008738) was also upregulated during bacterial infection. DEAD-box helicases are a large family of conserved RNA-binding proteins that belong to the broader group of cellular DExD/H helicases, with emerging evidence that it plays a role in the recognition of foreign nucleic acids and the modulation of viral infection (Taschuk & Cherry, 2020). DEAD-box helicases can play an important role in sensing viral infection and directly affecting virus RNA by participating in the RNAi pathway and Toll-like and retinoic acid-inducible gene-I-like receptors signaling pathway, for example (Ahmad & Hu, 2015; Baldaccini & Pfeffer, 2021; Su et al., 2022). However, there is evidence that noncoding subgenomic flavivirus RNA from ZIKV can bind to DEAD/H-box helicase ME31B in *Ae. aegypti* as a way to overcome this defense (Göertz et al., 2019).

### 3.4 Microorganisms co-infection generate oxidative stress and affects yolk component

#### 3.4.1 Immune response mediated by reactive oxygen species and DExD/H helicases

Martins et al (2021) earlier described that *Wolbachia* and ZIKV coinfection increased host cellular aerobic metabolism in order to accumulate reactive oxygen species (ROS) in *Ae. aegypti* head and salivary glands quantitative proteomics. In mosquito ovaries analysis, several pathways related to aerobic metabolism were upregulated, thus corroborating with previous data. The increase in aerobic respiration metabolism already proved to stimulate ROS production in insect cells as part of the host immune response, but it is counterbalanced with antioxidant pathways activation (Zug and Hammerstein, 2015; Matins et al., 2021). ROS accumulation activates the mosquito Toll immune pathway, resulting in the production of AMPs (Pan et al., 2012). Possibly as a result of this ROS amount, Cu-Zn superoxide dismutase (SOD) proteins (AAEL019937 and AAEL025388) were upregulated comparing WZ with W, Z and A. SODs are an important antioxidant enzymes that convert superoxide into oxygen and hydrogen peroxide (Park et al, 2012; Lomate et al., 2015) and it is also related to protecting insects’ ovaries during diapauses (Sim & Denlinger, 2011). Besides ROS-mediated immune response, a DEAD box ATP-dependent RNA helicase (AAEL004978) was upregulated in coinfection. DExD box RNA helicase was already discussed in *Wolbachia* mono-infection upregulations and leads us to believe that it is a *Wolbachia* mediated attempt to overcome ZIKV infection, even though this virus has a way to overcome this immune mechanism (Göertz et al., 2019).

#### 3.4.2 Vitellogenin A1 downregulation during coinfection

During WZ comparison with Z, W and A, it was noticed that vitellogenin A1 (AAEL010434) was downregulated in all analyses, related to macromolecule localization (GO:0033036), organic substance transport (GO:0071702), biological process (GO:0008150), transport (GO:0006810), establishment of localization (GO:0051234), localization (GO:0051179), lipid transport (GO:0006869) and lipid localization (GO:0010876). Vitellogenins are defined as the predominant yolk protein precursors that are produced extraovarially and taken up by the growing oocytes against a concentration gradient (Engelmann, 1979). By having this important role, it is relevant to understand how it behaves during microorganisms infection. Depending on which *Wolbachia* is allocated in the host, the ovary protein level can change, resulting in different protein content in embryos (Christensen et al., 2016), nonetheless there is a report that establish a link between the vitellogenin-related mode of transovarial transmission and efficient maternal transmission of *Wolbachia*, assuming that the bacteria utilized vitellogenin transportation system to enter the insect ovaries (Guo et al., 2018). Vitellogenesis can be directly regulated by insect hormones, including JH (Shapiro & Taylor, 1982; Wu et al., 2021). As it was discussed before, *Wolbachia* decreases enzyme levels of the JH pathway, and it might interfere in vitellogenesis, together with the yellow protein downregulation. On the other hand, female mosquitoes with regular blood feeding presented a high oviposition, probably mediated by nutrition influence in hormones, and it enhanced ZIKV infection (Rocha-Santos et al., 2021). This may lead to comprehension of reproduction–immunity trade-offs in insects. Immune defense and reproduction are physiologically and energetically demanding processes and have been observed to trade off in a diversity of female insects: increased reproductive effort results in reduced immunity, and reciprocally, infection and activation of the immune system reduce reproductive output (Schwenke et al., 2016). Moreover, an endocrine regulation of immunity in insects involving some hormones was already described, including JH, which is characterized as an immune suppressor (Nunes et al., 2021). Our result corroborates with this influence shown in literature, once coinfected mosquitoes have to deal with a major microorganism invasion.

## 4 Conclusions

Proteomics analysis of *Ae. aegypti* ovaries mono-infected with ZIKV or *Wolbachia* highlighted how those microorganisms interplay almost antagonistic responses considering insect reproducibility and immune response particularities. Our data support that ZIKV induces JH production, probably enhancing insect reproduction capability, while *Wolbachia* seems to harm the same hormone pathway and also eggs survival. As for insect innate immunity, *Wolbachia* helps to reinforce *Ae. aegypti* basal features, while ZIKV block los it to facilitate infection. During co-infection, *Wolbachia* helps *Ae. aegypti* to prevent virus infection by stimulating ROS production leading to a Toll-pathway humoral immune response, together with SOD production to control cell homeostasis. Besides, coinfection showed a likely role between the immune system and reproduction features. Finally, this work ought to be an important resource to understand how microorganisms infection can influence *Ae. aegypti* immune response and reproducibility, exposing those findings to the insect research community.

## Availability of raw files

The mass spectrometry proteomics data have been deposited to the ProteomeXchange Consortium via the PRIDE (Perez-Riverol et al., 2022) partner repository with the dataset identifier PXD031728.

**Username:** **Password:** Wj45qw8Q

## Authors Contribution

LFCR, MM and JRM performed the experiments and data analysis. MM produced paper figures. LFCR, MM, JRM, GBD, <u>DMPO</u>, FCSN, RM and MJ wrote the manuscript. GBD, FCSN, RM and MJ provided resources and acquired funding. RM and MJ idealized and coordinated the study. All authors approved the manuscript.

## Acknowledgments

The authors would like to thank Professor Rafael Dias Mesquita, Dr. André Torres, and Dr. Stephanie Serafim de Carvalho (Federal University of Rio de Janeiro) for helpful suggestions and discussions, which improve and strengthen the presentation of this manuscript. We are also grateful to the Laboratório de Apoio ao Desenvolvimento Tecnológico (LADETEC) of the Institute of Chemistry from the Federal University of Rio de Janeiro for providing the infrastructure for the nLC-MS/MS analysis.

## References

Aebersold, R., and Mann, M. (2016). Mass-spectrometric exploration of proteome structure and function. Nature 537, 347–355. doi:10.1038/nature19949.

Ahmad, S., and Hur, S. (2015). Helicases in Antiviral Immunity: Dual Properties as Sensors and Effectors. Trends in Biochemical Sciences 40, 576–585. doi:10.1016/j.tibs.2015.08.001.

Aliota, M. T., Walker, E. C., Yepes, A. U., Velez, I. D., Christensen, B. M., and Osorio, J. E. (2016). The wMel Strain of Wolbachia Reduces Transmission of Chikungunya Virus in Aedes aegypti. PLOS Neglected Tropical Diseases 10, e0004677. doi:10.1371/journal.pntd.0004677.

Allman, M. J., Fraser, J. E., Ritchie, S. A., Joubert, D. A., Simmons, C. P., and Flores, H. A. (2020). Wolbachia’s Deleterious Impact on Aedes aegypti Egg Development: The Potential Role of Nutritional Parasitism. Insects 11, 735. doi:10.3390/insects11110735.

Alomar, A. A., Eastmond, B. H., and Alto, B. W. (2021). Juvenile hormone analog enhances Zika virus infection in Aedes aegypti. Sci Rep 11, 21062. doi:10.1038/s41598-021-00432-1.

Andersen, S. B., Boye, M., Nash, D. R., and Boomsma, J. J. (2012). Dynamic Wolbachia prevalence in Acromyrmex leaf-cutting ants: potential for a nutritional symbiosis: Wolbachia in Acromyrmex ants. Journal of Evolutionary Biology 25, 1340–1350. doi:10.1111/j.1420-9101.2012.02521.x.

Ant, T. H., Herd, C. S., Geoghegan, V., Hoffmann, A. A., and Sinkins, S. P. (2018). The Wolbachia strain wAu provides highly efficient virus transmission blocking in Aedes aegypti. PLoS Pathog 14, e1006815. doi:10.1371/journal.ppat.1006815.

Ashburner, M., Ball, C. A., Blake, J. A., Botstein, D., Butler, H., Cherry, J. M., et al. (2000). Gene Ontology: tool for the unification of biology. Nature Genetics 25, 25–29. doi:10.1038/75556.

Avirutnan, P., Hauhart, R. E., Marovich, M. A., Garred, P., Atkinson, J. P., and Diamond, M. S. (2011). Complement-Mediated Neutralization of Dengue Virus Requires Mannose-Binding Lectin. mBio 2. doi:10.1128/mBio.00276-11.

Avirutnan, P., Hauhart, R. E., Marovich, M. A., Garred, P., Atkinson, J. P., Diamond, M. S. (2011). Complement-mediated neutralization of dengue virus requires mannose-binding lectin. mBio 13, e00276–11. doi: 10.1128/mBio.00276-11.

Axford, J. K., Callahan, A. G., Hoffmann, A. A., Yeap, H. L., and Ross, P. A. (2016). Fitness of wAlbB Wolbachia Infection in Aedes aegypti: Parameter Estimates in an Outcrossed Background and Potential for Population Invasion. The American Journal of Tropical Medicine and Hygiene 94, 507–516. doi:10.4269/ajtmh.15-0608.

Baldaccini, M., and Pfeffer, S. (2021). Untangling the roles of RNA helicases in antiviral innate immunity. PLoS Pathog 17, e1010072. doi:10.1371/journal.ppat.1010072.

Barton, L., Sanny, J., Dawson, E. P., Nouzova, M., Noriega, F. G., Stadtfeld, M., et al. (2021). Bioactive isoprenoids guide migrating germ cells to the embryonic gonad. Developmental Biology doi:10.1101/2021.09.30.462471.

Blandin, S. (2004). Thioester-containing proteins and insect immunity. Molecular Immunology 40, 903–908. doi:10.1016/j.molimm.2003.10.010.

Bose, J., and Schulte, R. D. (2014). Testing GxG interactions between coinfecting microbial parasite genotypes within hosts. Front. Genet. 5. doi:10.3389/fgene.2014.00124.

Boyer, S., Calvez, E., Chouin-Carneiro, T., Diallo, D., and Failloux, A.-B. (2018). An overview of mosquito vectors of Zika virus. Microbes and Infection 20, 646–660. doi:10.1016/j.micinf.2018.01.006.

Brinker, P., Fontaine, M. C., Beukeboom, L. W., and Falcao Salles, J. (2019). Host, Symbionts, and the Microbiome: The Missing Tripartite Interaction. Trends in Microbiology 27, 480–488. doi:10.1016/j.tim.2019.02.002.

Buonocore, F., Fausto, A. M., Della Pelle, G., Roncevic, T., Gerdol, M., and Picchietti, S. (2021). Attacins: A Promising Class of Insect Antimicrobial Peptides. Antibiotics 10, 212. doi:10.3390/antibiotics10020212.

Campos, G., Bandeira, A., and Sardi, S. (2015). Zika Virus Outbreak, Bahia, Brazil. Emerging Infectious Disease journal 21, 1885. doi:10.3201/eid2110.150847.

Campos, S. S., Fernandes, R. S., dos Santos, A. A. C., de Miranda, R. M., Telleria, E. L., Ferreira-de-Brito, A., et al. (2017). Zika virus can be venereally transmitted between Aedes aegypti mosquitoes. Parasites Vectors 10, 605. doi:10.1186/s13071-017-2543-4.

Cao-Lormeau, V. M., Roche, C., Teissier, A., Robin, E., Berry, A. L., Mallet, H. P., et al (2014). Zika virus, French Polynesia, South Pacific, 2013 [letter]. Emerg Infect Dis 20. doi: 10.3201/eid2006.140138.

Chang, M.-M., Wang, Y.-H., Yang, Q.-T., Wang, X.-L., Wang, M., Raikhel, A. S., et al. (2021). Regulation of antimicrobial peptides by juvenile hormone and its receptor, Methoprene-tolerant, in the mosquito Aedes aegypti. Insect Biochemistry and Molecular Biology 128, 103509. doi:10.1016/j.ibmb.2020.103509.

Charlwood J. D., Tomás, E. V. E., Andegiorgish, A. K., Mihreteab, S., LeClair, C. (2018). ‘We like it wet’: a comparison between dissection techniques for the assessment of parity in Anopheles arabiensis and determination of sac stage in mosquitoes alive or dead on collection. PeerJ 6, e5155. doi: 10.7717/peerj.5155.

Chen, X., Sun, Y., Zhang, T., Shu, L., Roepstorff, P., and Yang, F. (2022). Quantitative Proteomics Using Isobaric Labeling: A Practical Guide. Genomics, Proteomics & Bioinformatics, S1672022922000018. doi:10.1016/j.gpb.2021.08.012.

Cheng, G., Liu, Y., Wang, P., and Xiao, X. (2016). Mosquito Defense Strategies against Viral Infection. Trends in Parasitology 32, 177–186. doi:10.1016/j.pt.2015.09.009.

Chouin-Carneiro, T., Vega-Rua, A., Vazeille, M., Yebakima, A., Girod, R., Goindin, D., et al. (2016). Differential Susceptibilities of Aedes aegypti and Aedes albopictus from the Americas to Zika Virus. PLOS Neglected Tropical Diseases 10, e0004543. doi:10.1371/journal.pntd.0004543.

Christensen, S., Pérez Dulzaides, R., Hedrick, V. E., Momtaz, A. J. M. Z., Nakayasu, E. S., Paul, L. N., et al. (2016). Wolbachia Endosymbionts Modify Drosophila Ovary Protein Levels in a Context-Dependent Manner. Appl Environ Microbiol 82, 5354–5363. doi:10.1128/AEM.01255-16.

Codeço, C. T., Lima, A. W. S., Araújo, S. C., Lima, J. B. P., Maciel-de-Freitas, R., Honório, N. A., et al. (2015). Surveillance of Aedes aegypti: Comparison of House Index with Four Alternative Traps. PLOS Neglected Tropical Diseases 9, e0003475. doi:10.1371/journal.pntd.0003475.

Cox, J., and Mann, M. (2011). Quantitative, High-Resolution Proteomics for Data-Driven Systems Biology. Annu. Rev. Biochem. 80, 273–299. doi:10.1146/annurev-biochem-061308-093216.

da Silveira, I. D., Petersen, M. T., Sylvestre, G., Garcia, G. A., David, M. R., Pavan, M. G., et al. (2018). Zika Virus Infection Produces a Reduction on Aedes aegypti Lifespan but No Effects on Mosquito Fecundity and Oviposition Success. Front. Microbiol. 9, 3011. doi:10.3389/fmicb.2018.03011.

da Silveira, I. D., Petersen, M. T., Sylvestre, G., Garcia, G. A., David, M. R., Pavan, M. G., et al. (2018). Zika Virus Infection Produces a Reduction on Aedes aegypti Lifespan but No Effects on Mosquito Fecundity and Oviposition Success. Front. Microbiol. 9. doi:10.3389/fmicb.2018.03011.

Daimon, T., and Shinoda, T. (2013). Function, diversity, and application of insect juvenile hormone epoxidases (CYP15): Insect Juvenile Hormone Epoxidases (CYP15). Biotechnology and Applied Biochemistry 60, 82–91. doi:10.1002/bab.1058.

De Mandal, S., Lin, B., Shi, M., Li, Y., Xu, X., and Jin, F. (2020). iTRAQ-Based Comparative Proteomic Analysis of Larval Midgut From the Beet Armyworm, Spodoptera exigua (Hübner) (Lepidoptera: Noctuidae) Challenged With the Entomopathogenic Bacteria Serratia marcescens. Front. Physiol. 11. doi:10.3389/fphys.2020.00442.

de Oliveira, S., Villela, D. A. M., Dias, F. B. S., Moreira, L. A., and Maciel de Freitas, R. (2017). How does competition among wild type mosquitoes influence the performance of Aedes aegypti and dissemination of Wolbachia pipientis? PLoS Negl Trop Dis 11, e0005947. doi:10.1371/journal.pntd.0005947.

Dick, G. W. A., Kitchen, S. F., and Haddow, A. J. (1952). Zika Virus (I). Isolations and serological specificity. Transactions of The Royal Society of Tropical Medicine and Hygiene 46, 509–520. doi:10.1016/0035-9203(52)90042-4.

Dutra, H. L. C., dos Santos, L. M. B., Caragata, E. P., Silva, J. B. L., Villela, D. A. M., Maciel-de-Freitas, R., et al. (2015). From Lab to Field: The Influence of Urban Landscapes on the Invasive Potential of Wolbachia in Brazilian Aedes aegypti Mosquitoes. PLoS Negl Trop Dis 9, e0003689. doi:10.1371/journal.pntd.0003689.

Dutra, H. L. C., Rocha, M. N., Dias, F. B. S., Mansur, S. B., Caragata, E. P., and Moreira, L. A. (2016). Wolbachia Blocks Currently Circulating Zika Virus Isolates in Brazilian Aedes aegypti Mosquitoes. Cell Host & Microbe 19, 771–774. doi:10.1016/j.chom.2016.04.021.

Edenborough, K. M., Flores, H. A., Simmons, C. P., and Fraser, J. E. (2021). Using Wolbachia to Eliminate Dengue: Will the Virus Fight Back? J Virol 95, e02203–20. doi:10.1128/JVI.02203-20.

Engelmann, F. (1979). “Insect Vitellogenin: Identification, Biosynthesis, and Role in Vitellogenesis,” in Advances in Insect Physiology (Elsevier), 49–108. doi:10.1016/S0065-2806(08)60051-X.

Epelboin, Y., Talaga, S., Epelboin, L., and Dusfour, I. (2017). Zika virus: An updated review of competent or naturally infected mosquitoes. PLoS Negl Trop Dis 11, e0005933. doi:10.1371/journal.pntd.0005933.

Faria, V. G., and Sucena, É. (2013). Wolbachia in the Malpighian Tubules: Evolutionary Dead-End or Adaptation?: Wolbachia IN THE MALPIGHIAN TUBULES. J. Exp. Zool. (Mol. Dev. Evol.) 320, 195–199. doi:10.1002/jez.b.22498.

Farnesi, L. C., Belinato, T. A., Gesto, J. S. M., Martins, A. J., Bruno, R. V., and Moreira, L. A. (2019). Embryonic development and egg viability of wMel-infected Aedes aegypti. Parasites Vectors 12, 211. doi:10.1186/s13071-019-3474-z.

Fernandes, R. S., Campos, S. S., Ferreira-de-Brito, A., Miranda, R. M. de, Silva, K. A. B. da, Castro, M. G. de, et al. (2016). Culex quinquefasciatus from Rio de Janeiro Is Not Competent to Transmit the Local Zika Virus. PLOS Neglected Tropical Diseases 10, e0004993. doi:10.1371/journal.pntd.0004993.

Ferreira-de-Brito, A., Ribeiro, I. P., Miranda, R. M. de, Fernandes, R. S., Campos, S. S., Silva, K. A. B. da, et al. (2016). First detection of natural infection of Aedes aegypti with Zika virus in Brazil and throughout South America. Memórias do Instituto Oswaldo Cruz 111, 655–658. doi:10.1590/0074-02760160332.

Ford, S. A., Albert, I., Allen, S. L., Chenoweth, S. F., Jones, M., Koh, C., et al. (2020). Artificial Selection Finds New Hypotheses for the Mechanism of Wolbachia-Mediated Dengue Blocking in Mosquitoes. Front. Microbiol. 11. doi:10.3389/fmicb.2020.01456.

Fuchs, A., Pinto, A. K., Schwaeble, W. J., and Diamond, M. S. (2011). The lectin pathway of complement activation contributes to protection from West Nile virus infection. Virology 412, 101–109. doi:10.1016/j.virol.2011.01.003.

Garcia, G. A., Hoffmann, A. A., Maciel-de-Freitas, R., and Villela, D. A. M. (2020). Aedes aegypti insecticide resistance underlies the success (and failure) of Wolbachia population replacement. Sci Rep 10, 63. doi:10.1038/s41598-019-56766-4.

Garcia, G. de A., Sylvestre, G., Aguiar, R., Costa, G. B. da, Martins, A. J., Lima, J. B. P., et al. (2019). Matching the genetics of released and local Aedes aegypti populations is critical to assure Wolbachia invasion. PLOS Neglected Tropical Diseases 13, e0007023. doi:10.1371/journal.pntd.0007023.

García-Robles, I., De Loma, J., Capilla, M., Roger, I., Boix-Montesinos, P., Carrión, P., et al. (2020). Proteomic insights into the immune response of the Colorado potato beetle larvae challenged with Bacillus thuringiensis. Developmental & Comparative Immunology 104, 103525. doi:10.1016/j.dci.2019.103525.

Geng, T., Lu, F., Wu, H., Wang, Y., Lou, D., Tu, N., et al. (2021). C-type lectin 5, a novel pattern recognition receptor for the JAK/STAT signaling pathway in Bombyx mori. Journal of Invertebrate Pathology 179, 107473. doi:10.1016/j.jip.2020.107473.

Gesto, J. S. M., Ribeiro, G. S., Rocha, M. N., Dias, F. B. S., Peixoto, J., Carvalho, F. D., et al. (2021). Reduced competence to arboviruses following the sustainable invasion of Wolbachia into native Aedes aegypti from Southeastern Brazil. Sci Rep 11, 10039. doi:10.1038/s41598-021-89409-8.

Gestuveo, R. J., Royle, J., Donald, C. L., Lamont, D. J., Hutchinson, E. C., Merits, A., et al. (2021). Analysis of Zika virus capsid-Aedes aegypti mosquito interactome reveals pro-viral host factors critical for establishing infection. Nat Commun 12, 2766. doi:10.1038/s41467-021-22966-8.

Giraldo-Calderón, G. I., Emrich, S. J., MacCallum, R. M., Maslen, G., Dialynas, E., Topalis, P., Ho, N., Gesing, S., The VectorBase Consortium, Madey, G., Collins, F. H., Lawson, D. (2014). VectorBase: an updated bioinformatics resource for invertebrate vectors and other organisms related with human diseases. Nucleic Acids Research 43, 707–713. doi: 10.1093/nar/gku1117.

Gloria-Soria, A., Chiodo, T. G., and Powell, J. R. (2018). Lack of Evidence for Natural Wolbachia Infections in Aedes aegypti (Diptera: Culicidae). Journal of Medical Entomology. doi:10.1093/jme/tjy084.

Göertz, G. P., van Bree, J. W. M., Hiralal, A., Fernhout, B. M., Steffens, C., Boeren, S., et al. (2019). Subgenomic flavivirus RNA binds the mosquito DEAD/H-box helicase ME31B and determines Zika virus transmission by Aedes aegypti. Proc Natl Acad Sci USA 116, 19136–19144. doi:10.1073/pnas.1905617116.

González, M. A., Pavan, M. G., Fernandes, R. S., Busquets, N., David, M. R., Lourenço-Oliveira, R., et al. (2019). Limited risk of Zika virus transmission by five Aedes albopictus populations from Spain. Parasites Vectors 12, 150. doi:10.1186/s13071-019-3359-1.

Guo, Y., Hoffmann, A. A., Xu, X.-Q., Mo, P.-W., Huang, H.-J., Gong, J.-T., et al. (2018). Vertical Transmission of Wolbachia Is Associated With Host Vitellogenin in Laodelphax striatellus. Front. Microbiol. 9, 2016. doi:10.3389/fmicb.2018.02016.

Hivrale Lomate, P., Sangole, K., and Sunkar, R. (2015). Superoxide dismutase activities in the midgut of Helicoverpa armigera larvae: identification and biochemical properties of a manganese superoxide dismutase. OAIP, 13. doi:10.2147/OAIP.S84053.

Indriani, C., Tantowijoyo, W., Rancès, E., Andari, B., Prabowo, E., Yusdi, D., et al. (2020). Reduced dengue incidence following deployments of Wolbachia-infected Aedes aegypti in Yogyakarta, Indonesia: a quasi-experimental trial using controlled interrupted time series analysis. Gates Open Res 4, 50. doi:10.12688/gatesopenres.13122.1.

Jiggins, F. M. (2017). The spread of Wolbachia through mosquito populations. PLoS Biol 15, e2002780. doi:10.1371/journal.pbio.2002780.

Jindra, M., Bellés, X., and Shinoda, T. (2015). Molecular basis of juvenile hormone signaling. Current Opinion in Insect Science 11, 39–46. doi:10.1016/j.cois.2015.08.004.

Kauffman, E. B., and Kramer, L. D. (2017). Zika Virus Mosquito Vectors: Competence, Biology, and Vector Control. The Journal of Infectious Diseases 216, S976–S990. doi:10.1093/infdis/jix405.

Kaur, R., Shropshire, J. D., Cross, K. L., Leigh, B., Mansueto, A. J., Stewart, V., et al. (2021). Living in the endosymbiotic world of Wolbachia: A centennial review. Cell Host & Microbe 29, 879–893. doi:10.1016/j.chom.2021.03.006.

King, J. G., Souto-Maior, C., Sartori, L. M., Maciel-de-Freitas, R., and Gomes, M. G. M. (2018). Variation in Wolbachia effects on Aedes mosquitoes as a determinant of invasiveness and vectorial capacity. Nat Commun 9, 1483. doi:10.1038/s41467-018-03981-8.

Lai, Z., Zhou, T., Zhou, J., Liu, S., Xu, Y., Gu, J., et al. (2020). Vertical transmission of zika virus in Aedes albopictus. PLoS Negl Trop Dis 14, e0008776. doi:10.1371/journal.pntd.0008776.

Lau, M.-J., Ross, P. A., and Hoffmann, A. A. (2021). Infertility and fecundity loss of Wolbachia-infected Aedes aegypti hatched from quiescent eggs is expected to alter invasion dynamics. PLoS Negl Trop Dis 15, e0009179. doi:10.1371/journal.pntd.0009179.

LePage, D. P., Metcalf, J. A., Bordenstein, S. R., On, J., Perlmutter, J. I., Shropshire, J. D., et al. (2017). Prophage WO genes recapitulate and enhance Wolbachia-induced cytoplasmic incompatibility. Nature 543, 243–247. doi:10.1038/nature21391.

Li, C., Guo, X., Deng, Y., Xing, D., Sun, A., Liu, Q., et al. (2017). Vector competence and transovarial transmission of two Aedes aegypti strains to Zika virus. Emerging Microbes & Infections 6, 1–7. doi:10.1038/emi.2017.8.

Li, Q., Meng, Q.-W., Lü, F.-G., Guo, W.-C., and Li, G.-Q. (2016). Identification of ten mevalonate enzyme-encoding genes and their expression in response to juvenile hormone levels in Leptinotarsa decemlineata (Say). Gene 584, 136–147. doi:10.1016/j.gene.2016.02.023.

Ma, Z., Qu, B., Yao, L., Gao, Z., and Zhang, S. (2020). Identification and functional characterization of ribosomal protein S23 as a new member of antimicrobial protein. Developmental & Comparative Immunology 110, 103730. doi:10.1016/j.dci.2020.103730.

Mancini, M. V., Herd, C. S., Ant, T. H., Murdochy, S. M., and Sinkins, S. P. (2020). Wolbachia strain wAu efficiently blocks arbovirus transmission in Aedes albopictus. PLoS Negl Trop Dis 14, e0007926. doi:10.1371/journal.pntd.0007926.

Manuel, M., Missé, D., and Pompon, J. (2020). Highly Efficient Vertical Transmission for Zika Virus in Aedes aegypti after Long Extrinsic Incubation Time. Pathogens 9, 366. doi:10.3390/pathogens9050366.

Martins, M., Ramos, L. F. C., Murillo, J. R., Torres, A., de Carvalho, S. S., Domont, G. B., et al. (2021). Comprehensive Quantitative Proteome Analysis of Aedes aegypti Identifies Proteins and Pathways Involved in Wolbachia pipientis and Zika Virus Interference Phenomenon. Front. Physiol. 12, 642237. doi:10.3389/fphys.2021.642237.

Mayoral, J. G., Nouzova, M., Navare, A., and Noriega, F. G. (2009). NADP + -dependent farnesol dehydrogenase, a corpora allata enzyme involved in juvenile hormone synthesis. PNAS 106, 21091–21096. doi:10.1073/pnas.0909938106.

McMeniman, C. J., and O’Neill, S. L. (2010). A Virulent Wolbachia Infection Decreases the Viability of the Dengue Vector Aedes aegypti during Periods of Embryonic Quiescence. PLoS Negl Trop Dis 4, e748. doi:10.1371/journal.pntd.0000748.

McMeniman, C. J., Hughes, G. L., and O’Neill, S. L. (2011). A Wolbachia Symbiont in Aedes aegypti Disrupts Mosquito Egg Development to a Greater Extent When Mosquitoes Feed on Nonhuman Versus Human Blood. jnl. med. entom. 48, 76–84. doi:10.1603/ME09188.

Moreira, L. A., Iturbe-Ormaetxe, I., Jeffery, J. A., Lu, G., Pyke, A. T., Hedges, L. M., et al. (2009). A Wolbachia Symbiont in Aedes aegypti Limits Infection with Dengue, Chikungunya, and Plasmodium. Cell 139, 1268–1278. doi:10.1016/j.cell.2009.11.042.

Mukherjee, D., Das, S., Begum, F., Mal, S., and Ray, U. (2019). The Mosquito Immune System and the Life of Dengue Virus: What We Know and Do Not Know. Pathogens 8, 77. doi:10.3390/pathogens8020077.

Nag, D. K., Payne, A. F., Dieme, C., Ciota, A. T., and Kramer, L. D. (2021). Zika virus infects Aedes aegypti ovaries. Virology 561, 58–64. doi:10.1016/j.virol.2021.06.002.

Nazni, W. A., Hoffmann, A. A., NoorAfizah, A., Cheong, Y. L., Mancini, M. V., Golding, N., et al. (2019). Establishment of Wolbachia Strain wAlbB in Malaysian Populations of Aedes aegypti for Dengue Control. Current Biology 29, 4241–4248.e5. doi:10.1016/j.cub.2019.11.007.

Noh, M. Y., Kim, S. H., Gorman, M. J., Kramer, K. J., Muthukrishnan, S., and Arakane, Y. (2020). Yellow-g and Yellow-g2 proteins are required for egg desiccation resistance and temporal pigmentation in the Asian tiger mosquito, Aedes albopictus. Insect Biochemistry and Molecular Biology 122, 103386. doi:10.1016/j.ibmb.2020.103386.

Noh, M. Y., Kramer, K. J., Muthukrishnan, S., Beeman, R. W., Kanost, M. R., and Arakane, Y. (2015). Loss of function of the yellow-e gene causes dehydration-induced mortality of adult Tribolium castaneum. Developmental Biology 399, 315–324. doi:10.1016/j.ydbio.2015.01.009.

Noriega, F. G. (2014). Juvenile Hormone Biosynthesis in Insects: What Is New, What Do We Know, and What Questions Remain? International Scholarly Research Notices 2014, 1–16. doi:10.1155/2014/967361.

Nouzova, M., Edwards, M. J., Mayoral, J. G., and Noriega, F. G. (2011). A coordinated expression of biosynthetic enzymes controls the flux of juvenile hormone precursors in the corpora allata of mosquitoes. Insect Biochemistry and Molecular Biology 41, 660–669. doi:10.1016/j.ibmb.2011.04.008.

Nunes, C., Sucena, É., and Koyama, T. (2021). Endocrine regulation of immunity in insects. FEBS J 288, 3928–3947. doi:10.1111/febs.15581.

Ogunlade, S. T., Meehan, M. T., Adekunle, A. I., Rojas, D. P., Adegboye, O. A., and McBryde, E. S. (2021). A Review: Aedes-Borne Arboviral Infections, Controls and Wolbachia-Based Strategies. Vaccines 9, 32. doi:10.3390/vaccines9010032.

Padilha, K. P., Resck, M. E. B., Cunha, O. A. T. da, Teles-de-Freitas, R., Campos, S. S., Sorgine, M. H. F., et al. (2018). Zika infection decreases Aedes aegypti locomotor activity but does not influence egg production or viability. Mem. Inst. Oswaldo Cruz 113. doi:10.1590/0074-02760180290.

Pan, X., Zhou, G., Wu, J., Bian, G., Lu, P., Raikhel, A. S., et al. (2012a). Wolbachia induces reactive oxygen species (ROS)-dependent activation of the Toll pathway to control dengue virus in the mosquito Aedes aegypti. PNAS 109, E23–E31. doi:10.1073/pnas.1116932108.

Park, D. H., Choi, J. Y., Lee, S.-H., Kim, J. H., Park, M. G., Kim, J. Y., et al. (2020). Mosquito larvicidal activities of farnesol and farnesyl acetate via regulation of juvenile hormone receptor complex formation in Aedes mosquito. Journal of Asia-Pacific Entomology 23, 689–693. doi:10.1016/j.aspen.2020.05.006.

Park, S.-Y., Nair, P. M. G., and Choi, J. (2012). Characterization and expression of superoxide dismutase genes in Chironomus riparius (Diptera, Chironomidae) larvae as a potential biomarker of ecotoxicity. Comparative Biochemistry and Physiology Part C: Toxicology & Pharmacology 156, 187–194. doi:10.1016/j.cbpc.2012.06.003.

Perez-Riverol, Y., Bai, J., Bandla, C., Hewapathirana, S., García-Seisdedos, D., Kamatchinathan, S., Kundu, D., Prakash, A., Frericks-Zipper, A., Eisenacher, M., Walzer, M., Wang, S., Brazma, A., Vizcaíno, J.A. (2022). The PRIDE database resources in 2022: A Hub for mass spectrometry-based proteomics evidences. Nucleic Acids Research 50(D1):D543–D552. doi: 10.1093/nar/gkab1038.

Petersen, L. R., Jamieson, D. J., Powers, A. M., and Honein, M. A. (2016). Zika Virus. N Engl J Med 374, 1552–1563. doi:10.1056/NEJMra1602113.

Petersen, M. T., Silveira, I. D. da, Tátila-Ferreira, A., David, M. R., Chouin-Carneiro, T., Van den Wouwer, L., et al. (2018). The impact of the age of first blood meal and Zika virus infection on Aedes aegypti egg production and longevity. PLoS ONE 13, e0200766. doi:10.1371/journal.pone.0200766.

Petersen, M. T., Silveira, I. D. da, Tátila-Ferreira, A., David, M. R., Chouin-Carneiro, T., Wouwer, L. V. den, et al. (2018). The impact of the age of first blood meal and Zika virus infection on Aedes aegypti egg production and longevity. PLOS ONE 13, e0200766. doi:10.1371/journal.pone.0200766.

Pickett, B. E., Sadat, E. L., Zhang, Y., Noronha, J. M., Squires, R. B., Hunt, V., et al. (2012). ViPR: an open bioinformatics database and analysis resource for virology research. Nucleic Acids Res 40, D593–D598. doi:10.1093/nar/gkr859.

Pietri, J. E., DeBruhl, H., and Sullivan, W. (2016). The rich somatic life of Wolbachia. MicrobiologyOpen 5, 923–936. doi:10.1002/mbo3.390.

Pimentel, A. C., Cesar, C. S., Martins, M., and Cogni, R. (2021). The Antiviral Effects of the Symbiont Bacteria Wolbachia in Insects. Front. Immunol. 11, 626329. doi:10.3389/fimmu.2020.626329.

Ponton, F., Wilson, K., Holmes, A., Raubenheimer, D., Robinson, K. L., and Simpson, S. J. (2015). Macronutrients mediate the functional relationship between Drosophila and Wolbachia. Proc. R. Soc. B. 282, 20142029. doi:10.1098/rspb.2014.2029.

Rauniyar, N., and Yates, J. R. (2014). Isobaric Labeling-Based Relative Quantification in Shotgun Proteomics. Proteome Res 13, 5293–5309. doi: 10.1021/pr500880b

Resck, M. E. B., Padilha, K. P., Cupolillo, A. P., Talyuli, O. A. C., Ferreira-de-Brito, A., Lourenço-de-Oliveira, R., et al. (2020). Unlike Zika, Chikungunya virus interferes in the viability of Aedes aegypti eggs, regardless of females’ age. Sci Rep 10, 13642. doi:10.1038/s41598-020-70367-6.

Rocha-Santos, C., Dutra, A. C. V. P. L., Fróes Santos, R., Cupolillo, C. D. L. S., de Melo Rodovalho, C., Bellinato, D. F., et al. (2021). Effect of Larval Food Availability on Adult Aedes Aegypti (Diptera: Culicidae) Fitness and Susceptibility to Zika Infection. Journal of Medical Entomology 58, 535–547. doi:10.1093/jme/tjaa249.

Romans, P., Tu, Z., Ke, Z., and Hagedorn, H. H. (1995). Analysis of a vitellogenin gene of the mosquito, Aedes aegypti and comparisons to vitellogenins from other organisms. Insect Biochemistry and Molecular Biology 25, 939–958. doi:10.1016/0965-1748(95)00037-V.

Ronquist, F., and Huelsenbeck, J. P. (2003). MrBayes 3: Bayesian phylogenetic inference under mixed models. Bioinformatics 19, 1572–1574. doi:10.1093/bioinformatics/btg180.

Ryan, P. A., Turley, A. P., Wilson, G., Hurst, T. P., Retzki, K., Brown-Kenyon, J., et al. (2020). Establishment of wMel Wolbachia in Aedes aegypti mosquitoes and reduction of local dengue transmission in Cairns and surrounding locations in northern Queensland, Australia. Gates Open Res 3, 1547. doi:10.12688/gatesopenres.13061.2.

Sá-Guimarães, T. E., Salles, T. S., Santos, C. R., Moreira M. F., Souza, W., Caldas, L. A. (2021). Route of Zika virus infection in Aedes aegypti by transmission electron microscopy. BMC Microbiol 21, 300. doi: 10.1186/s12866-021-02366-0.

Santos, C. G., Humann, F. C., and Hartfelder, K. (2019). Juvenile hormone signaling in insect oogenesis. Current Opinion in Insect Science 31, 43–48. doi:10.1016/j.cois.2018.07.010.

Schmid, M. A., Kauffman, E., Payne, A., Harris, E., and Kramer, L. D. (2017). Preparation of Mosquito Salivary Gland Extract and Intradermal Inoculation of Mice. Bio-protocol 7, e2407–e2407.

Schneider, S. M., Lee, B. H., and Nicola, A. V. (2021). Viral entry and the ubiquitin-proteasome system. Cellular Microbiology 23. doi:10.1111/cmi.13276.

Schubert, O. T., Röst, H. L., Collins, B. C., Rosenberger, G., and Aebersold, R. (2017). Quantitative proteomics: challenges and opportunities in basic and applied research. Nat Protoc 12, 1289–1294. doi:10.1038/nprot.2017.040.

Schwenke, R. A., Lazzaro, B. P., and Wolfner, M. F. (2016). Reproduction–Immunity Trade-Offs in Insects. Annu. Rev. Entomol. 61, 239–256. doi:10.1146/annurev-ento-010715-023924.

Searle, B. C., and Yergey, A. L. (2020). An efficient solution for resolving iTRAQ and TMT channel cross-talk. J Mass Spectrom 55, e4354. doi:10.1002/jms.4354.

Serteyn, L., Ponnet, L., Saive, M., Fauconnier, M.-L., and Francis, F. (2020). Changes of feeding behavior and salivary proteome of Brown Marmorated Stink Bug when exposed to insect-induced plant defenses. Arthropod-Plant Interactions 14, 101–112. doi:10.1007/s11829-019-09718-8.

Shapiro, D., and Taylor, J. M. (1982). Steroid Hormone Regulation of Vitellogenin Gene Expressio. Critical Reviews in Biochemistry 12, 187–203. doi:10.3109/10409238209108706.

Shi, Z.-K., Wen, D., Chang, M.-M., Sun, X.-M., Wang, Y.-H., Cheng, C.-H., et al. (2021). Juvenile Hormone-Sensitive Ribosomal Activity Enhances Viral Replication in Aedes aegypti. mSystems 6. doi:10.1128/mSystems.01190-20.

Shokal, U., and Eleftherianos, I. (2017). Evolution and Function of Thioester-Containing Proteins and the Complement System in the Innate Immune Response. Front. Immunol. 8, 759. doi:10.3389/fimmu.2017.00759.

Sim, C., and Denlinger, D. L. (2011). Catalase and superoxide dismutase-2 enhance survival and protect ovaries during overwintering diapause in the mosquito Culex pipiens. Journal of Insect Physiology 57, 628–634. doi:10.1016/j.jinsphys.2011.01.012.

Smith, L., Kelleher, N. & The Consortium for Top Down Proteomics (2013). Proteoform: a single term describing protein complexity. Nat Methods 10, 186–187. doi: 10.1038/nmeth.2369.

Strunov, A., Kiseleva, E., and Gottlieb, Y. (2013). Spatial and temporal distribution of pathogenic Wolbachia strain wMelPop in Drosophila melanogaster central nervous system under different temperature conditions. Journal of Invertebrate Pathology 114, 22–30. doi:10.1016/j.jip.2013.05.001.

Su, C., Tang, Y., and Zheng, C. (2022). DExD/H-box helicases: multifunctional regulators in antiviral innate immunity. Cell. Mol. Life Sci. 79, 2. doi:10.1007/s00018-021-04072-6.

Takahashi, K., Ip, W. E., Michelow, I. C., and Ezekowitz, R. A. B. (2006). The mannose-binding lectin: a prototypic pattern recognition molecule. Current Opinion in Immunology 18, 16–23. doi:10.1016/j.coi.2005.11.014.

Taschuk, F., and Cherry, S. (2020). DEAD-Box Helicases: Sensors, Regulators, and Effectors for Antiviral Defense. Viruses 12, 181. doi:10.3390/v12020181.

Thangamani, S., Huang, J., Hart, C. E., Guzman, H., and Tesh, R. B. (2016). Vertical Transmission of Zika Virus in Aedes aegypti Mosquitoes. The American Journal of Tropical Medicine and Hygiene 95, 1169–1173. doi:10.4269/ajtmh.16-0448.

Tsang, S. S. K., Law, S. T. S., Li, C., Qu, Z., Bendena, W. G., Tobe, S. S., et al. (2020). Diversity of Insect Sesquiterpenoid Regulation. Front. Genet. 11, 1027. doi:10.3389/fgene.2020.01027.

Utarini, A., Indriani, C., Ahmad, R. A., Tantowijoyo, W., Arguni, E., Ansari, M. R., et al. (2021). Efficacy of Wolbachia-Infected Mosquito Deployments for the Control of Dengue. N Engl J Med 384, 2177–2186. doi:10.1056/NEJMoa2030243.

Wang, W., Yang, R.-R., Peng, L.-Y., Zhang, L., Yao, Y.-L., and Bao, Y.-Y. (2021). Proteolytic activity of the proteasome is required for female insect reproduction. Open Biol. 11, rsob.200251, 200251. doi:10.1098/rsob.200251.

Waterhouse, R. M., Povelones, M., and Christophides, G. K. (2010). Sequence-structure-function relations of the mosquito leucine-rich repeat immune proteins. BMC Genomics 11, 531. doi:10.1186/1471-2164-11-531.

Wigglesworth, V. B. (1934). The Physiology of Ecdysis in Rhodnius Prolixus (Hemiptera). II. Factors controlling Moulting and “Metamorphosis”. 33.

Wong, Z. S., Hedges, L. M., Brownlie, J. C., and Johnson, K. N. (2011). Wolbachia-Mediated Antibacterial Protection and Immune Gene Regulation in Drosophila. PLoS ONE 6, e25430. doi:10.1371/journal.pone.0025430.

Wu, Q., Patočka, J., and Kuča, K. (2018). Insect Antimicrobial Peptides, a Mini Review. Toxins 10, 461. doi:10.3390/toxins10110461.

Wu, Z., Yang, L., He, Q., and Zhou, S. (2021). Regulatory Mechanisms of Vitellogenesis in Insects. Front. Cell Dev. Biol. 8, 593613. doi:10.3389/fcell.2020.593613.

Xi, Z., Ramirez, J. L., and Dimopoulos, G. (2008). The Aedes aegypti Toll Pathway Controls Dengue Virus Infection. PLoS Pathog 4, e1000098. doi:10.1371/journal.ppat.1000098.

Xia, X., You, M., Rao, X.-J., and Yu, X.-Q. (2018). Insect C-type lectins in innate immunity. Developmental & Comparative Immunology 83, 70–79. doi:10.1016/j.dci.2017.11.020.

Xiao, X., Liu, Y., Zhang, X., Wang, J., Li, Z., Pang, X., et al. (2014). Complement-Related Proteins Control the Flavivirus Infection of Aedes aegypti by Inducing Antimicrobial Peptides. PLoS Pathog 10, e1004027. doi:10.1371/journal.ppat.1004027.

Ye, Y. H., Woolfit, M., Rancès, E., O’Neill, S. L., and McGraw, E. A. (2013). Wolbachia-Associated Bacterial Protection in the Mosquito Aedes aegypti. PLoS Negl Trop Dis 7, e2362. doi:10.1371/journal.pntd.0002362.

Yeap, H. L., Hoffmann, A. A., Ross, P. A., and Endersby, N. M. (2014). Larval Competition Extends Developmental Time and Decreases Adult Size of wMelPop Wolbachia-Infected Aedes aegypti. The American Journal of Tropical Medicine and Hygiene 91, 198–205. doi:10.4269/ajtmh.13-0576.

Zhang, R., Zhu, Y., Pang, X., Xiao, X., Zhang, R., and Cheng, G. (2017). Regulation of Antimicrobial Peptides in Aedes aegypti Aag2 Cells. Front. Cell. Infect. Microbiol. 7. doi:10.3389/fcimb.2017.00022.

Zhao, L., Alto, B., and Shin, D. (2019). Transcriptional Profile of Aedes aegypti Leucine-Rich Repeat Proteins in Response to Zika and Chikungunya Viruses. IJMS 20, 615. doi:10.3390/ijms20030615.

Zifruddin, A.-N., Mohamad-Khalid, K.-A., Suhaimi, S.-A., Mohamed-Hussein, Z.-A., and Hassan, M. (2021). Molecular characterization and enzyme inhibition studies of NADP+-farnesol dehydrogenase from diamondback moth, Plutella xylostella (Lepidoptera: Plutellidae). Bioscience, Biotechnology, and Biochemistry 85, 1628–1638. doi:10.1093/bbb/zbab072.

Zimler, R. A., Yee, D. A., and Alto, B. W. (2021). Transmission Potential of Zika Virus by Aedes aegypti (Diptera: Culicidae) and Ae. mediovittatus (Diptera: Culicidae) Populations From Puerto Rico. Journal of Medical Entomology 58, 1405–1411. doi:10.1093/jme/tjaa286.

Zug, R., and Hammerstein, P. (2015). Wolbachia and the insect immune system: what reactive oxygen species can tell us about the mechanisms of Wolbachia–host interactions. Front. Microbiol. 6. doi:10.3389/fmicb.2015.01201.

