## Supplementary Table 1 for "Interspecies isobaric labeling-based quantitative proteomics reveals protein changes in the ovary of *Aedes aegypti* co-infected with ZIKV and Wolbachia"

| Proteins U | Checked | Protein FD | Master | Accession | Descriptio | Exp. q-val | Coverage | Contamin | # Peptides | # PSMs | # Unique | # AAs | MW [kDa] | calc. pI |
|---|---|---|---|---|---|---|---|---|---|---|---|---|---|---|
| 4,74077E | FALSE | High | IsMasterP | AAEL0061 | 3:682058 | 0 | 77 | FALSE | 215 | 4390 | 149 | 2142 | 2493 | 718 |
| -2,03596E | FALSE | High | IsMasterP | AAEL0104 | 3:674899 | 0 | 71 | FALSE | 188 | 3426 | 122 | 2148 | 2502 | 708 |
| -8,13987E | FALSE | High | IsMasterP | AAEL0061 | 3:680846 | 0 | 74 | FALSE | 185 | 3161 | 180 | 2087 | 2419 | 686 |
| -5,55874E | FALSE | High | IsMasterP | AAEL0262 | 2:293815 | 0 | 64 | FALSE | 152 | 770 | 56 | 1963 | 2238 | 602 |
| 6,80388E | FALSE | High | IsMasterP | AAEL0065 | 2:594898 | 0 | 76 | FALSE | 48 | 636 | 48 | 471 | 537 | 667 |
| -6,05211E | FALSE | High | IsMasterP | AAEL0182 | 3:192456 | 0 | 46 | FALSE | 153 | 642 | 1 | 4632 | 5150 | 511 |
| -7,57534E | FALSE | High | IsMasterP | AAEL0182 | 3:192453 | 0 | 46 | FALSE | 153 | 640 | 1 | 4583 | 5091 | 510 |
| 3,26303E | FALSE | High | IsMasterP | AAEL0016 | 2:870803 | 0 | 85 | FALSE | 41 | 544 | 10 | 376 | 418 | 539 |
| -4,69234E | FALSE | High | IsMasterP | AAEL0111 | 1:355983 | 0 | 85 | FALSE | 37 | 528 | 8 | 376 | 418 | 548 |
| 2,34399E | FALSE | High | IsMasterP | AAEL0087 | 2:198770 | 0 | 52 | FALSE | 143 | 538 | 143 | 3701 | 4116 | 502 |
| 8,7883E+ | FALSE | High | IsMasterP | AAEL0056 | 2:382064 | 0 | 53 | FALSE | 114 | 558 | 10 | 1940 | 2213 | 599 |
| -5,78764E | FALSE | High | IsMasterP | AAEL0133 | 2:135690 | 0 | 84 | FALSE | 18 | 476 | 18 | 194 | 218 | 841 |
| 4,44007E | FALSE | High | IsMasterP | AAEL0218 | 1:151955 | 0 | 47 | FALSE | 105 | 535 | 1 | 1940 | 2212 | 596 |
| -8,24293E | FALSE | High | IsMasterP | AAEL0150 | 2:191821 | 0 | 66 | FALSE | 154 | 488 | 154 | 2428 | 2795 | 526 |
| -9,20788E | FALSE | High | IsMasterP | AAEL0194 | 3:937492 | 0 | 72 | FALSE | 53 | 468 | 36 | 651 | 711 | 547 |
| -5,29669E | FALSE | High | IsMasterP | AAEL0170 | 2:455769 | 0 | 76 | FALSE | 33 | 406 | 17 | 463 | 504 | 909 |
| 6,35944E | FALSE | High | IsMasterP | AAEL0046 | 1:306245 | 0 | 85 | FALSE | 40 | 407 | 13 | 376 | 418 | 548 |
| -1,14229E | FALSE | High | IsMasterP | AAEL0046 | 1:158961 | 0 | 78 | FALSE | 75 | 382 | 75 | 1006 | 1120 | 952 |
| 1,92628E | FALSE | High | IsMasterP | AAEL0059 | 3:227634 | 0 | 76 | FALSE | 35 | 394 | 2 | 376 | 416 | 548 |
| -8,13535E | FALSE | High | IsMasterP | AAEL0169 | 2:296894 | 0 | 86 | FALSE | 29 | 350 | 29 | 332 | 354 | 818 |
| -3,87532E | FALSE | High | IsMasterP | AAEL0028 | 3:235239 | 0 | 86 | FALSE | 32 | 371 | 30 | 504 | 539 | 512 |
| -2,5179E+ | FALSE | High | IsMasterP | AAEL0036 | 3:653469 | 0 | 59 | FALSE | 79 | 339 | 79 | 1788 | 2005 | 535 |
| -4,33634E | FALSE | High | IsMasterP | AAEL0019 | 3:227586 | 0 | 78 | FALSE | 32 | 334 | 7 | 376 | 416 | 548 |
| -2,86277E | FALSE | High | IsMasterP | AAEL0014 | 2:458208 | 0 | 52 | FALSE | 94 | 334 | 94 | 1969 | 2269 | 548 |
| 4,62578E | FALSE | High | IsMasterP | AAEL0231 | 1:214011 | 0 | 62 | FALSE | 100 | 334 | 100 | 2324 | 2507 | 621 |
| -7,30441E | FALSE | High | IsMasterP | AAEL0058 | 3:218807 | 0 | 55 | FALSE | 105 | 327 | 105 | 2396 | 2763 | 550 |
| -4,15092E | FALSE | High | IsMasterP | AAEL0028 | 3:233420 | 0 | 82 | FALSE | 34 | 322 | 16 | 447 | 501 | 486 |
| 9,18077E | FALSE | High | IsMasterP | AAEL0046 | 1:306225 | 0 | 57 | FALSE | 19 | 307 | 4 | 376 | 417 | 548 |
| 4,60429E | FALSE | High | IsMasterP | AAEL0173 | 3:686148 | 0 | 55 | FALSE | 43 | 350 | 28 | 655 | 722 | 520 |
| 2,83539E | FALSE | High | IsMasterP | AAEL0148 | 2:104793 | 0 | 68 | FALSE | 55 | 332 | 1 | 715 | 815 | 503 |
| 8,20157E | FALSE | High | IsMasterP | AAEL0105 | 2:111977 | 0 | 70 | FALSE | 58 | 318 | 58 | 803 | 888 | 547 |

| | | | | | | | | | | | | | | |
|---|---|---|---|---|---|---|---|---|---|---|---|---|---|---|
| 4,27321E- | FALSE | High | IsMasterP | AAEL0117 | 2:104830 | 0 | 68 | FALSE | 55 | 330 | 1 | 715 | 814 | 503 |
| 8,13359E- | FALSE | High | IsMasterP | AAEL0121 | 2:204340 | 0 | 75 | FALSE | 53 | 343 | 53 | 551 | 594 | 894 |
| -8,39254E | FALSE | High | IsMasterP | AAEL0015 | 2:211071 | 0 | 83 | FALSE | 40 | 299 | 40 | 372 | 405 | 696 |
| 5,9304E+ | FALSE | High | IsMasterP | AAEL0051 | 2:313244 | 0 | 58 | FALSE | 71 | 293 | 71 | 1624 | 1799 | 544 |
| -4,59581E | FALSE | High | IsMasterP | AAEL0029 | 1:261514 | 0 | 83 | FALSE | 44 | 293 | 44 | 516 | 560 | 679 |
| -2,44303E | FALSE | High | IsMasterP | AAEL0115 | 3:948460 | 0 | 78 | FALSE | 49 | 303 | 49 | 574 | 608 | 562 |
| 2,18873E- | FALSE | High | IsMasterP | AAEL0101 | 2:264745 | 0 | 72 | FALSE | 56 | 272 | 56 | 770 | 825 | 925 |
| -5,15546E | FALSE | High | IsMasterP | AAEL0014 | 2:456863 | 0 | 78 | FALSE | 42 | 278 | 42 | 493 | 552 | 578 |
| -3,24124E | FALSE | High | IsMasterP | AAEL0044 | 3:204353 | 0 | 90 | FALSE | 42 | 256 | 42 | 627 | 679 | 701 |
| 6,54759E- | FALSE | High | IsMasterP | AAEL0096 | 1:222740 | 0 | 64 | FALSE | 26 | 266 | 25 | 332 | 372 | 699 |
| -2,94077E | FALSE | High | IsMasterP | AAEL0045 | 2:453152 | 0 | 63 | FALSE | 48 | 261 | 48 | 844 | 943 | 639 |
| -7,9867E+ | FALSE | High | IsMasterP | AAEL0010 | 1:299812 | 0 | 87 | FALSE | 19 | 292 | 15 | 209 | 238 | 607 |
| -7,62628E | FALSE | High | IsMasterP | AAEL0103 | 3:409688 | 0 | 72 | FALSE | 43 | 262 | 43 | 628 | 697 | 947 |
| -7,8963E+ | FALSE | High | IsMasterP | AAEL0109 | 2:151124 | 0 | 66 | FALSE | 70 | 279 | 70 | 883 | 1032 | 558 |
| -6,64454E | FALSE | High | IsMasterP | AAEL0128 | 2:365950 | 0 | 64 | FALSE | 48 | 248 | 48 | 788 | 857 | 844 |
| -2,29934E | FALSE | High | IsMasterP | AAEL0025 | 1:228539 | 0 | 93 | FALSE | 26 | 252 | 26 | 247 | 263 | 634 |
| -1,21129E | FALSE | High | IsMasterP | AAEL0015 | 2:228343 | 0 | 83 | FALSE | 38 | 240 | 38 | 483 | 535 | 557 |
| 5,58173E- | FALSE | High | IsMasterP | AAEL0007 | 2:458823 | 0 | 64 | FALSE | 51 | 240 | 51 | 845 | 969 | 633 |
| 1,32959E- | FALSE | High | IsMasterP | AAEL0096 | 3:209820 | 0 | 65 | FALSE | 58 | 252 | 58 | 1195 | 1321 | 681 |
| -8,58568E | FALSE | High | IsMasterP | AAEL0242 | 2:299315 | 0 | 41 | FALSE | 23 | 219 | 23 | 442 | 477 | 753 |
| -6,09579E | FALSE | High | IsMasterP | AAEL0066 | 3:148473 | 0 | 86 | FALSE | 33 | 249 | 2 | 450 | 499 | 514 |
| 6,82067E- | FALSE | High | IsMasterP | AAEL0057 | 3:114738 | 0 | 78 | FALSE | 33 | 239 | 2 | 364 | 395 | 818 |
| 2,89873E- | FALSE | High | IsMasterP | AAEL0209 | 2:264633 | 0 | 57 | FALSE | 9 | 216 | 9 | 167 | 177 | 882 |
| 5,30445E- | FALSE | High | IsMasterP | AAEL0057 | 3:114738 | 0 | 76 | FALSE | 32 | 233 | 1 | 363 | 391 | 787 |
| 8,97753E- | FALSE | High | IsMasterP | AAEL0132 | 3:280008 | 0 | 85 | FALSE | 32 | 235 | 1 | 450 | 499 | 514 |
| -1,81464E | FALSE | High | IsMasterP | AAEL0073 | 1:259803 | 0 | 69 | FALSE | 25 | 206 | 25 | 221 | 250 | 563 |
| 6,21177E- | FALSE | High | IsMasterP | AAEL0090 | 3:193312 | 0 | 84 | FALSE | 37 | 213 | 36 | 516 | 566 | 806 |
| -1,5116E+ | FALSE | High | IsMasterP | AAEL0006 | 3:395598 | 0 | 81 | FALSE | 46 | 223 | 46 | 494 | 559 | 502 |
| 5,18036E- | FALSE | High | IsMasterP | AAEL0242 | 2:488091 | 0 | 45 | FALSE | 79 | 225 | 79 | 2866 | 3104 | 644 |
| -1,04305E | FALSE | High | IsMasterP | AAEL0087 | 3:132652 | 0 | 89 | FALSE | 38 | 238 | 38 | 614 | 682 | 539 |
| -1,39913E | FALSE | High | IsMasterP | AAEL0028 | 1:108201 | 0 | 76 | FALSE | 36 | 208 | 36 | 532 | 578 | 642 |

| | | | | | | | | | | | | | | |
|---|---|---|---|---|---|---|---|---|---|---|---|---|---|---|
| 4,58692E- | FALSE | High | IsMasterP | AAEL0076 | 1:1349660 | 0 | 82 | FALSE | 39 | 234 | 39 | 534 | 579 | 585 |
| -5,80423E | FALSE | High | IsMasterP | AAEL0194 | 1:2761748 | 0 | 31 | FALSE | 63 | 217 | 63 | 2579 | 2803 | 465 |
| 2,08298E- | FALSE | High | IsMasterP | AAEL0120 | 1:2705567 | 0 | 49 | FALSE | 48 | 218 | 48 | 1040 | 1152 | 560 |
| -8,89119E | FALSE | High | IsMasterP | AAEL0057 | 2:4360936 | 0 | 55 | FALSE | 49 | 212 | 45 | 1080 | 1246 | 665 |
| 9,50397E- | FALSE | High | IsMasterP | AAEL0194 | 3:4017524 | 0 | 89 | FALSE | 17 | 217 | 17 | 196 | 218 | 596 |
| 5,54315E- | FALSE | High | IsMasterP | AAEL0081 | 3:1007637 | 0 | 80 | FALSE | 24 | 222 | 24 | 337 | 353 | 904 |
| 4,98488E- | FALSE | High | IsMasterP | AAEL0014 | 2:4826942 | 0 | 78 | FALSE | 43 | 210 | 43 | 508 | 585 | 492 |
| -4,06E+18 | FALSE | High | IsMasterP | AAEL0091 | 2:9746568 | 0 | 44 | FALSE | 42 | 216 | 42 | 612 | 671 | 677 |
| 4,89867E- | FALSE | High | IsMasterP | AAEL0002 | 2:7536525 | 0 | 64 | FALSE | 52 | 212 | 52 | 921 | 1033 | 739 |
| 3,19953E- | FALSE | High | IsMasterP | AAEL0054 | 1:2460189 | 0 | 67 | FALSE | 40 | 205 | 40 | 570 | 632 | 876 |
| 8,5623E+1 | FALSE | High | IsMasterP | AAEL0073 | 1:2597951 | 0 | 64 | FALSE | 18 | 176 | 18 | 209 | 237 | 597 |
| 2,34093E- | FALSE | High | IsMasterP | AAEL0038 | 3:8514752 | 0 | 87 | FALSE | 19 | 194 | 19 | 171 | 196 | 478 |
| 4,22023E- | FALSE | High | IsMasterP | AAEL0117 | 1:1475106 | 0 | 73 | FALSE | 13 | 193 | 13 | 206 | 223 | 901 |
| -6,58644E | FALSE | High | IsMasterP | AAEL0111 | 3:3791653 | 0 | 77 | FALSE | 29 | 203 | 27 | 258 | 294 | 486 |
| 4,3589E+1 | FALSE | High | IsMasterP | AAEL0069 | 3:3613157 | 0 | 62 | FALSE | 28 | 173 | 28 | 508 | 536 | 683 |
| -4,59993E | FALSE | High | IsMasterP | AAEL0018 | 2:1844319 | 0 | 56 | FALSE | 17 | 162 | 17 | 412 | 458 | 586 |
| 6,0197E+1 | FALSE | High | IsMasterP | AAEL0276 | 3:2509907 | 0 | 30 | FALSE | 86 | 188 | 86 | 4327 | 4958 | 649 |
| 5,14872E- | FALSE | High | IsMasterP | AAEL0196 | 2:6350079 | 0 | 16 | FALSE | 77 | 200 | 77 | 7554 | 8569 | 552 |
| 1,62239E- | FALSE | High | IsMasterP | AAEL0052 | 2:6914971 | 0 | 86 | FALSE | 28 | 155 | 28 | 441 | 455 | 802 |
| -5,48482E | FALSE | High | IsMasterP | AAEL0041 | 2:1471829 | 0 | 79 | FALSE | 26 | 176 | 20 | 448 | 497 | 548 |
| -2,42068E | FALSE | High | IsMasterP | AAEL0023 | 2:3031537 | 0 | 72 | FALSE | 31 | 179 | 31 | 429 | 472 | 900 |
| 3,3813E+1 | FALSE | High | IsMasterP | AAEL0034 | 3:3486673 | 0 | 70 | FALSE | 43 | 171 | 43 | 579 | 663 | 627 |
| -2,16933E | FALSE | High | IsMasterP | AAEL0010 | 1:2998129 | 0 | 65 | FALSE | 13 | 189 | 9 | 211 | 238 | 596 |
| -7,94139E | FALSE | High | IsMasterP | AAEL0088 | 3:2366628 | 0 | 59 | FALSE | 53 | 183 | 53 | 1011 | 1130 | 520 |
| 3,39217E- | FALSE | High | IsMasterP | AAEL0145 | 1:3043970 | 0 | 85 | FALSE | 15 | 167 | 15 | 112 | 113 | 459 |
| 5,56301E- | FALSE | High | IsMasterP | AAEL0057 | 1:2002786 | 0 | 84 | FALSE | 32 | 170 | 32 | 496 | 552 | 557 |
| -1,7889E+ | FALSE | High | IsMasterP | AAEL0067 | 2:8507242 | 0 | 57 | FALSE | 53 | 168 | 5 | 1063 | 1192 | 723 |
| 3,06228E- | FALSE | High | IsMasterP | AAEL0173 | 3:2527040 | 0 | 55 | FALSE | 20 | 187 | 4 | 462 | 505 | 895 |
| 5,43937E- | FALSE | High | IsMasterP | AAEL0025 | 1:2274336 | 0 | 17 | FALSE | 84 | 170 | 84 | 8149 | 9160 | 665 |
| 6,75752E- | FALSE | High | IsMasterP | AAEL0256 | NIGP0100 | 0 | 45 | FALSE | 44 | 171 | 44 | 1107 | 1206 | 441 |
| -2,53694E | FALSE | High | IsMasterP | AAEL0128 | 1:2425762 | 0 | 57 | FALSE | 40 | 188 | 40 | 795 | 911 | 491 |

| | | | | | | | | | | | | | | |
|---|---|---|---|---|---|---|---|---|---|---|---|---|---|---|
| -4,7522E+ | FALSE | High | IsMasterP | AAEL0132 | 2:9471789 | 0 | 80 | FALSE | 22 | 174 | 22 | 198 | 214 | 876 |
| -7,97119E | FALSE | High | IsMasterP | AAEL0063 | 3:2310673 | 0 | 68 | FALSE | 27 | 167 | 27 | 551 | 628 | 632 |
| 8,76374E+ | FALSE | High | IsMasterP | AAEL0173 | 3:3468090 | 0 | 61 | FALSE | 42 | 176 | 42 | 839 | 941 | 573 |
| 9,0673E+1 | FALSE | High | IsMasterP | AAEL0125 | 1:2941531 | 0 | 70 | FALSE | 36 | 159 | 36 | 730 | 792 | 696 |
| 4,76144E+ | FALSE | High | IsMasterP | AAEL0278 | NIGP0100 | 0 | 51 | FALSE | 7 | 132 | 3 | 136 | 154 | 1127 |
| -6,9073E+ | FALSE | High | IsMasterP | AAEL0134 | 2:1592540 | 0 | 81 | FALSE | 40 | 166 | 31 | 504 | 568 | 756 |
| -8,60161E | FALSE | High | IsMasterP | AAEL0091 | 2:3424820 | 0 | 62 | FALSE | 46 | 165 | 46 | 775 | 883 | 671 |
| 2,75923E+ | FALSE | High | IsMasterP | AAEL0094 | 3:3212726 | 0 | 66 | FALSE | 29 | 191 | 6 | 376 | 416 | 558 |
| 5,06528E+ | FALSE | High | IsMasterP | AAEL0059 | 3:2276465 | 0 | 47 | FALSE | 22 | 186 | 1 | 376 | 415 | 548 |
| -6,15204E | FALSE | High | IsMasterP | AAEL0049 | 2:2818069 | 0 | 80 | FALSE | 29 | 159 | 29 | 415 | 438 | 709 |
| 8,50699E+ | FALSE | High | IsMasterP | AAEL0020 | 2:4265535 | 0 | 57 | FALSE | 46 | 169 | 46 | 1009 | 1132 | 733 |
| 6,29035E+ | FALSE | High | IsMasterP | AAEL0068 | 1:8318108 | 0 | 47 | FALSE | 17 | 158 | 17 | 550 | 618 | 619 |
| 4,68175E+ | FALSE | High | IsMasterP | AAEL0045 | 2:3101784 | 0 | 83 | FALSE | 32 | 148 | 32 | 349 | 383 | 796 |
| -3,97127E | FALSE | High | IsMasterP | AAEL0223 | 2:2744089 | 0 | 73 | FALSE | 28 | 158 | 28 | 282 | 307 | 854 |
| -2,53001E | FALSE | High | IsMasterP | AAEL0067 | 2:8507242 | 0 | 56 | FALSE | 49 | 156 | 0 | 1078 | 1210 | 721 |
| 7,98889E+ | FALSE | High | IsMasterP | AAEL0118 | 2:1872359 | 0 | 50 | FALSE | 25 | 158 | 25 | 511 | 563 | 936 |
| 3,79402E+ | FALSE | High | IsMasterP | AAEL0127 | 3:3221304 | 0 | 73 | FALSE | 45 | 175 | 45 | 546 | 595 | 543 |
| -6,58821E | FALSE | High | IsMasterP | AAEL0069 | 3:2795768 | 0 | 77 | FALSE | 32 | 154 | 32 | 531 | 585 | 680 |
| 8,22041E+ | FALSE | High | IsMasterP | AAEL0077 | 2:4703851 | 0 | 71 | FALSE | 23 | 172 | 23 | 333 | 357 | 670 |
| -8,73851E | FALSE | High | IsMasterP | AAEL0067 | 2:8507242 | 0 | 57 | FALSE | 48 | 153 | 1 | 1037 | 1165 | 695 |
| 4,60423E+ | FALSE | High | IsMasterP | AAEL0010 | 1:2100612 | 0 | 59 | FALSE | 27 | 141 | 27 | 407 | 467 | 456 |
| 2,77333E+ | FALSE | High | IsMasterP | AAEL0171 | 3:8891851 | 0 | 55 | FALSE | 30 | 153 | 4 | 601 | 680 | 897 |
| -9,68476E | FALSE | High | IsMasterP | AAEL0067 | 2:8507242 | 0 | 52 | FALSE | 46 | 149 | 1 | 1093 | 1227 | 679 |
| -6,49276E | FALSE | High | IsMasterP | AAEL0130 | 2:2534727 | 0 | 90 | FALSE | 27 | 140 | 26 | 311 | 349 | 788 |
| 5,79271E+ | FALSE | High | IsMasterP | AAEL0275 | 3:3420379 | 0 | 52 | FALSE | 10 | 140 | 9 | 165 | 179 | 1037 |
| -8,88428E | FALSE | High | IsMasterP | AAEL0004 | 3:1852085 | 0 | 81 | FALSE | 33 | 145 | 33 | 510 | 548 | 646 |
| -5,13147E | FALSE | High | IsMasterP | AAEL0070 | 2:2419996 | 0 | 68 | FALSE | 38 | 172 | 38 | 719 | 777 | 818 |
| -2,90708E | FALSE | High | IsMasterP | AAEL0228 | 3:1953027 | 0 | 50 | FALSE | 64 | 168 | 64 | 1681 | 1916 | 577 |
| 8,27417E+ | FALSE | High | IsMasterP | AAEL0171 | 3:8891851 | 0 | 55 | FALSE | 30 | 149 | 4 | 600 | 679 | 897 |
| 1,73215E+ | FALSE | High | IsMasterP | AAEL0044 | 3:1100000 | 0 | 49 | FALSE | 35 | 164 | 33 | 796 | 868 | 727 |
| -8,3995E+ | FALSE | High | IsMasterP | AAEL0068 | 3:5641171 | 0 | 76 | FALSE | 11 | 139 | 11 | 235 | 250 | 595 |

| | | | | | | | | | | | | | | |
|---|---|---|---|---|---|---|---|---|---|---|---|---|---|---|
| 5,97216E- | FALSE | High | IsMasterP | AAEL0004 | 3:1612878 | 0 | 82 | FALSE | 40 | 156 | 40 | 542 | 596 | 655 |
| -6,88662E | FALSE | High | IsMasterP | AAEL0174 | 2:4325922 | 0 | 68 | FALSE | 40 | 148 | 40 | 635 | 737 | 677 |
| 7,83268E- | FALSE | High | IsMasterP | AAEL0149 | 1:2852811 | 0 | 81 | FALSE | 31 | 137 | 31 | 513 | 555 | 740 |
| -2,61732E | FALSE | High | IsMasterP | AAEL0027 | 3:1451843 | 0 | 85 | FALSE | 35 | 166 | 34 | 284 | 324 | 486 |
| 8,73039E- | FALSE | High | IsMasterP | AAEL0150 | 2:3377253 | 0 | 88 | FALSE | 25 | 152 | 25 | 275 | 305 | 632 |
| 5,05349E- | FALSE | High | IsMasterP | AAEL0093 | 2:4048515 | 0 | 95 | FALSE | 24 | 147 | 24 | 272 | 299 | 553 |
| 6,23115E- | FALSE | High | IsMasterP | AAEL0011 | 3:4065087 | 0 | 72 | FALSE | 41 | 136 | 41 | 813 | 901 | 522 |
| -5,19577E | FALSE | High | IsMasterP | AAEL0108 | 3:5204930 | 0 | 69 | FALSE | 27 | 142 | 27 | 315 | 338 | 560 |
| 6,46312E- | FALSE | High | IsMasterP | AAEL0248 | 3:3417219 | 0 | 27 | FALSE | 17 | 174 | 17 | 239 | 264 | 1026 |
| 1,25659E- | FALSE | High | IsMasterP | AAEL0144 | 2:1189369 | 0 | 64 | FALSE | 30 | 154 | 30 | 420 | 455 | 815 |
| 7,19242E- | FALSE | High | IsMasterP | AAEL0073 | 3:1968064 | 0 | 56 | FALSE | 48 | 153 | 48 | 896 | 1036 | 560 |
| -6,31757E | FALSE | High | IsMasterP | AAEL0077 | 2:4702684 | 0 | 74 | FALSE | 31 | 139 | 31 | 533 | 572 | 747 |
| 5,4965E+1 | FALSE | High | IsMasterP | AAEL0106 | 2:1763169 | 0 | 83 | FALSE | 27 | 133 | 27 | 398 | 416 | 854 |
| -7,3684E+ | FALSE | High | IsMasterP | AAEL0052 | 3:8961683 | 0 | 76 | FALSE | 22 | 126 | 22 | 383 | 402 | 596 |
| 6,47907E- | FALSE | High | IsMasterP | AAEL0232 | NIGP0100 | 0 | 29 | FALSE | 42 | 137 | 6 | 1581 | 1781 | 620 |
| -2,22686E | FALSE | High | IsMasterP | AAEL0232 | NIGP0100 | 0 | 59 | FALSE | 33 | 124 | 33 | 844 | 957 | 698 |
| -1,57966E | FALSE | High | IsMasterP | AAEL0124 | 3:1258449 | 0 | 68 | FALSE | 30 | 142 | 30 | 448 | 503 | 544 |
| 6,37079E- | FALSE | High | IsMasterP | AAEL0100 | 3:1068406 | 0 | 75 | FALSE | 38 | 155 | 38 | 546 | 596 | 657 |
| -5,48247E | FALSE | High | IsMasterP | AAEL0082 | 2:5912503 | 0 | 58 | FALSE | 36 | 147 | 36 | 901 | 987 | 616 |
| -7,66298E | FALSE | High | IsMasterP | AAEL0099 | 1:1932537 | 0 | 49 | FALSE | 24 | 148 | 24 | 431 | 489 | 1130 |
| 6,58371E- | FALSE | High | IsMasterP | AAEL0123 | 2:1028683 | 0 | 71 | FALSE | 29 | 138 | 29 | 443 | 480 | 837 |
| 5,40722E- | FALSE | High | IsMasterP | AAEL0197 | 3:8471900 | 0 | 29 | FALSE | 58 | 147 | 58 | 2651 | 2947 | 494 |
| 6,67638E- | FALSE | High | IsMasterP | AAEL0039 | 3:2540176 | 0 | 79 | FALSE | 20 | 130 | 20 | 292 | 315 | 847 |
| 6,59277E- | FALSE | High | IsMasterP | AAEL0059 | 3:1547320 | 0 | 88 | FALSE | 22 | 141 | 22 | 132 | 148 | 714 |
| -8,16906E | FALSE | High | IsMasterP | AAEL0256 | 3:2953881 | 0 | 60 | FALSE | 35 | 137 | 34 | 671 | 726 | 621 |
| 9,18618E- | FALSE | High | IsMasterP | AAEL0070 | 2:1295416 | 0 | 79 | FALSE | 24 | 126 | 24 | 319 | 359 | 549 |
| -1,01714E | FALSE | High | IsMasterP | AAEL0085 | 2:3951151 | 0 | 74 | FALSE | 36 | 145 | 36 | 439 | 504 | 819 |
| -3,13601E | FALSE | High | IsMasterP | AAEL0134 | 2:1993004 | 0 | 59 | FALSE | 34 | 130 | 34 | 685 | 790 | 644 |
| 9,19282E- | FALSE | High | IsMasterP | AAEL0026 | 2:2433659 | 0 | 58 | FALSE | 25 | 115 | 25 | 649 | 726 | 926 |
| -7,77687E | FALSE | High | IsMasterP | AAEL0128 | 1:2425501 | 0 | 74 | FALSE | 36 | 145 | 36 | 590 | 642 | 803 |
| -1,24705E | FALSE | High | IsMasterP | AAEL0066 | 3:2497212 | 0 | 84 | FALSE | 27 | 110 | 27 | 409 | 433 | 837 |

| | | | | | | | | | | | | | | |
|---|---|---|---|---|---|---|---|---|---|---|---|---|---|---|
| 8,7802E+1 | FALSE | High | IsMasterP | AAEL0281 | 1:307321 | 0 | 27 | FALSE | 39 | 132 | 3 | 1656 | 1865 | 624 |
| 3,02583E- | FALSE | High | IsMasterP | AAEL0100 | 2:357540 | 0 | 48 | FALSE | 26 | 129 | 26 | 599 | 672 | 619 |
| -5,55483E | FALSE | High | IsMasterP | AAEL0040 | 3:401671 | 0 | 80 | FALSE | 21 | 125 | 1 | 318 | 356 | 646 |
| -2,78389E | FALSE | High | IsMasterP | AAEL0133 | 2:240486 | 0 | 90 | FALSE | 31 | 136 | 31 | 404 | 455 | 599 |
| -7,71047E | FALSE | High | IsMasterP | AAEL0099 | 3:776875 | 0 | 27 | FALSE | 60 | 134 | 60 | 3343 | 3696 | 766 |
| 1,26635E- | FALSE | High | IsMasterP | AAEL0171 | 3:350829 | 0 | 86 | FALSE | 21 | 153 | 20 | 245 | 267 | 840 |
| 2,21273E- | FALSE | High | IsMasterP | AAEL0068 | 1:530224 | 0 | 85 | FALSE | 27 | 124 | 27 | 323 | 357 | 828 |
| 6,74471E- | FALSE | High | IsMasterP | AAEL0123 | 3:316468 | 0 | 85 | FALSE | 11 | 108 | 11 | 149 | 168 | 422 |
| -7,12415E | FALSE | High | IsMasterP | AAEL0044 | 2:453139 | 0 | 72 | FALSE | 29 | 127 | 29 | 373 | 415 | 758 |
| 6,82713E- | FALSE | High | IsMasterP | AAEL0106 | 3:291643 | 0 | 55 | FALSE | 21 | 107 | 21 | 658 | 719 | 693 |
| -2,01804E | FALSE | High | IsMasterP | AAEL0137 | 1:237623 | 0 | 69 | FALSE | 21 | 116 | 21 | 329 | 344 | 843 |
| -8,97168E | FALSE | High | IsMasterP | AAEL0027 | 3:145020 | 0 | 75 | FALSE | 28 | 133 | 4 | 285 | 329 | 474 |
| -1,14521E | FALSE | High | IsMasterP | AAEL0025 | 1:903624 | 0 | 71 | FALSE | 32 | 115 | 31 | 428 | 477 | 534 |
| 5,43155E- | FALSE | High | IsMasterP | AAEL0104 | 1:204143 | 0 | 54 | FALSE | 25 | 127 | 2 | 552 | 610 | 810 |
| 3,27805E- | FALSE | High | IsMasterP | AAEL0121 | 2:204263 | 0 | 90 | FALSE | 22 | 125 | 22 | 279 | 304 | 652 |
| -3,67569E | FALSE | High | IsMasterP | AAEL0108 | 3:517277 | 0 | 57 | FALSE | 18 | 123 | 18 | 210 | 226 | 970 |
| -8,63805E | FALSE | High | IsMasterP | AAEL0147 | 2:852896 | 0 | 35 | FALSE | 45 | 121 | 45 | 2080 | 2291 | 674 |
| -7,96899E | FALSE | High | IsMasterP | AAEL0139 | 2:332006 | 0 | 71 | FALSE | 20 | 122 | 20 | 354 | 390 | 890 |
| 3,52592E- | FALSE | High | IsMasterP | AAEL0027 | 3:145020 | 0 | 82 | FALSE | 28 | 130 | 3 | 284 | 327 | 479 |
| 3,18151E- | FALSE | High | IsMasterP | AAEL0245 | 2:646866 | 0 | 61 | FALSE | 28 | 124 | 28 | 553 | 626 | 822 |
| 1,70781E- | FALSE | High | IsMasterP | AAEL0017 | 2:443080 | 0 | 83 | FALSE | 38 | 131 | 38 | 543 | 594 | 594 |
| 9,19583E- | FALSE | High | IsMasterP | AAEL0040 | 3:401672 | 0 | 80 | FALSE | 21 | 116 | 1 | 317 | 355 | 619 |
| -6,28637E | FALSE | High | IsMasterP | AAEL0037 | 1:248503 | 0 | 63 | FALSE | 26 | 118 | 26 | 479 | 518 | 809 |
| 7,95926E- | FALSE | High | IsMasterP | AAEL0007 | 3:244989 | 0 | 47 | FALSE | 37 | 120 | 37 | 1088 | 1212 | 534 |
| -7,56687E | FALSE | High | IsMasterP | AAEL0024 | 1:176719 | 0 | 66 | FALSE | 27 | 113 | 26 | 415 | 446 | 860 |
| -6,26933E | FALSE | High | IsMasterP | AAEL0021 | 1:101603 | 0 | 52 | FALSE | 32 | 108 | 32 | 541 | 608 | 892 |
| 1,23793E- | FALSE | High | IsMasterP | AAEL0043 | 2:386664 | 0 | 70 | FALSE | 16 | 113 | 16 | 242 | 269 | 478 |
| 6,62195E- | FALSE | High | IsMasterP | AAEL0260 | 1:235764 | 0 | 54 | FALSE | 27 | 101 | 27 | 778 | 881 | 706 |
| 8,39283E- | FALSE | High | IsMasterP | AAEL0104 | 1:204143 | 0 | 52 | FALSE | 24 | 118 | 1 | 566 | 626 | 784 |
| 1,05564E- | FALSE | High | IsMasterP | AAEL0009 | 3:376987 | 0 | 57 | FALSE | 11 | 98 | 11 | 224 | 246 | 473 |
| 2,56652E- | FALSE | High | IsMasterP | AAEL0075 | 2:217545 | 0 | 62 | FALSE | 24 | 106 | 24 | 605 | 662 | 714 |

| | | | | | | | | | | | | | | |
|---|---|---|---|---|---|---|---|---|---|---|---|---|---|---|
| -5,06788E | FALSE | High | IsMasterP | AAEL0112 | 1:150123 | 0 | 53 | FALSE | 25 | 118 | 25 | 432 | 489 | 768 |
| -6,34524E | FALSE | High | IsMasterP | AAEL0028 | 1:108662 | 0 | 42 | FALSE | 13 | 118 | 13 | 221 | 247 | 935 |
| 4,78587E | FALSE | High | IsMasterP | AAEL0022 | 3:374095 | 0 | 62 | FALSE | 25 | 105 | 25 | 469 | 507 | 900 |
| -4,33422E | FALSE | High | IsMasterP | AAEL0195 | 1:818161 | 0 | 42 | FALSE | 29 | 109 | 29 | 1265 | 1414 | 520 |
| -8,26538E | FALSE | High | IsMasterP | AAEL0000 | 2:184466 | 0 | 42 | FALSE | 38 | 112 | 38 | 1117 | 1223 | 478 |
| 5,68229E | FALSE | High | IsMasterP | AAEL0199 | 3:265360 | 0 | 92 | FALSE | 11 | 97 | 11 | 153 | 154 | 624 |
| -2,62676E | FALSE | High | IsMasterP | AAEL0264 | 2:155094 | 0 | 19 | FALSE | 14 | 103 | 2 | 645 | 705 | 540 |
| -6,8018E+ | FALSE | High | IsMasterP | AAEL0205 | 3:121889 | 0 | 28 | FALSE | 48 | 122 | 48 | 2654 | 2921 | 671 |
| -8,0923E+ | FALSE | High | IsMasterP | AAEL0020 | 2:425558 | 0 | 59 | FALSE | 21 | 116 | 21 | 441 | 480 | 821 |
| 6,22021E | FALSE | High | IsMasterP | AAEL0238 | 1:201253 | 0 | 40 | FALSE | 51 | 111 | 51 | 1638 | 1833 | 841 |
| -2,47942E | FALSE | High | IsMasterP | AAEL0045 | 1:144860 | 0 | 71 | FALSE | 25 | 119 | 25 | 393 | 444 | 742 |
| -6,11414E | FALSE | High | IsMasterP | AAEL0071 | 3:191531 | 0 | 65 | FALSE | 16 | 98 | 16 | 505 | 552 | 506 |
| -2,76875E | FALSE | High | IsMasterP | AAEL0248 | NIGP0100 | 0 | 19 | FALSE | 15 | 114 | 2 | 662 | 733 | 505 |
| -8,64843E | FALSE | High | IsMasterP | AAEL0100 | 2:200456 | 0 | 74 | FALSE | 18 | 104 | 18 | 261 | 288 | 728 |
| -3,08212E | FALSE | High | IsMasterP | AAEL0007 | 2:521070 | 0 | 76 | FALSE | 27 | 109 | 25 | 409 | 458 | 684 |
| 5,10788E | FALSE | High | IsMasterP | AAEL0035 | 2:256239 | 0 | 79 | FALSE | 9 | 90 | 9 | 112 | 115 | 428 |
| 2,70455E | FALSE | High | IsMasterP | AAEL0229 | 1:134422 | 0 | 47 | FALSE | 34 | 116 | 34 | 927 | 1056 | 619 |
| -5,67006E | FALSE | High | IsMasterP | AAEL0169 | 2:133741 | 0 | 46 | FALSE | 23 | 106 | 5 | 749 | 831 | 699 |
| -8,16633E | FALSE | High | IsMasterP | AAEL0229 | 3:341140 | 0 | 43 | FALSE | 11 | 129 | 3 | 169 | 189 | 1046 |
| 9,13505E | FALSE | High | IsMasterP | AAEL0106 | 2:279267 | 0 | 64 | FALSE | 29 | 99 | 29 | 642 | 704 | 493 |
| -4,62444E | FALSE | High | IsMasterP | AAEL0106 | 2:326796 | 0 | 35 | FALSE | 28 | 94 | 28 | 1372 | 1470 | 736 |
| -4,07897E | FALSE | High | IsMasterP | AAEL0196 | 2:136243 | 0 | 58 | FALSE | 26 | 113 | 26 | 498 | 541 | 519 |
| 4,77671E | FALSE | High | IsMasterP | AAEL0255 | 3:196681 | 0 | 70 | FALSE | 22 | 100 | 22 | 410 | 467 | 527 |
| -4,89705E | FALSE | High | IsMasterP | AAEL0154 | 1:224906 | 0 | 59 | FALSE | 33 | 102 | 33 | 633 | 706 | 744 |
| 6,96721E | FALSE | High | IsMasterP | AAEL0093 | 3:380913 | 0 | 68 | FALSE | 18 | 112 | 18 | 333 | 370 | 658 |
| 6,20795E | FALSE | High | IsMasterP | AAEL0081 | 3:929826 | 0 | 61 | FALSE | 21 | 105 | 21 | 503 | 543 | 838 |
| 4,00186E | FALSE | High | IsMasterP | AAEL0117 | 3:376864 | 0 | 49 | FALSE | 20 | 98 | 20 | 604 | 658 | 658 |
| -5,94695E | FALSE | High | IsMasterP | AAEL0024 | 2:302264 | 0 | 86 | FALSE | 18 | 95 | 18 | 206 | 231 | 665 |
| 2,53864E | FALSE | High | IsMasterP | AAEL0093 | 1:199839 | 0 | 55 | FALSE | 30 | 111 | 30 | 557 | 595 | 657 |
| 2,85046E | FALSE | High | IsMasterP | AAEL0169 | 2:133741 | 0 | 46 | FALSE | 24 | 98 | 6 | 747 | 829 | 658 |
| -8,97236E | FALSE | High | IsMasterP | AAEL0093 | 1:199821 | 0 | 69 | FALSE | 18 | 100 | 2 | 265 | 291 | 483 |

| | | | | | | | | | | | | | | |
|---|---|---|---|---|---|---|---|---|---|---|---|---|---|---|
| -6,29316E | FALSE | High | IsMasterP | AAEL0072 | 2:421396 | 0 | 40 | FALSE | 31 | 96 | 31 | 1298 | 1434 | 477 |
| -3,82778E | FALSE | High | IsMasterP | AAEL0049 | 2:281679 | 0 | 45 | FALSE | 22 | 98 | 21 | 638 | 687 | 550 |
| -2,41086E | FALSE | High | IsMasterP | AAEL0140 | 1:238615 | 0 | 63 | FALSE | 24 | 98 | 22 | 489 | 530 | 568 |
| 6,90705E | FALSE | High | IsMasterP | AAEL0123 | 2:111543 | 0 | 74 | FALSE | 12 | 97 | 12 | 168 | 184 | 863 |
| -7,79198E | FALSE | High | IsMasterP | AAEL0018 | 2:276681 | 0 | 78 | FALSE | 18 | 95 | 18 | 267 | 293 | 573 |
| -8,75424E | FALSE | High | IsMasterP | AAEL0039 | 3:268285 | 0 | 73 | FALSE | 15 | 110 | 15 | 148 | 171 | 720 |
| -4,14235E | FALSE | High | IsMasterP | AAEL0149 | 2:145104 | 0 | 79 | FALSE | 12 | 82 | 12 | 156 | 171 | 681 |
| -5,7635E+ | FALSE | High | IsMasterP | AAEL0025 | 1:229081 | 0 | 90 | FALSE | 16 | 83 | 16 | 210 | 228 | 477 |
| -4,83402E | FALSE | High | IsMasterP | AAEL0133 | 2:240430 | 0 | 73 | FALSE | 7 | 98 | 7 | 126 | 137 | 591 |
| -1,49982E | FALSE | High | IsMasterP | AAEL0021 | 1:252337 | 0 | 55 | FALSE | 16 | 101 | 16 | 216 | 242 | 601 |
| 7,17609E | FALSE | High | IsMasterP | AAEL0129 | 2:314312 | 0 | 71 | FALSE | 20 | 90 | 20 | 526 | 579 | 552 |
| -8,90606E | FALSE | High | IsMasterP | AAEL0104 | 1:268456 | 0 | 48 | FALSE | 27 | 103 | 26 | 741 | 801 | 639 |
| -6,54233E | FALSE | High | IsMasterP | AAEL0094 | 3:344871 | 0 | 58 | FALSE | 12 | 88 | 12 | 192 | 221 | 992 |
| -6,64588E | FALSE | High | IsMasterP | AAEL0253 | 1:295494 | 0 | 72 | FALSE | 19 | 101 | 19 | 254 | 274 | 873 |
| 5,29024E | FALSE | High | IsMasterP | AAEL0000 | 3:102899 | 0 | 34 | FALSE | 13 | 100 | 13 | 346 | 393 | 1052 |
| 4,88604E | FALSE | High | IsMasterP | AAEL0004 | 3:187120 | 0 | 58 | FALSE | 25 | 94 | 25 | 593 | 650 | 785 |
| 1,45056E | FALSE | High | IsMasterP | AAEL0011 | 3:164404 | 0 | 56 | FALSE | 24 | 86 | 24 | 521 | 565 | 777 |
| -8,38018E | FALSE | High | IsMasterP | AAEL0057 | 2:437306 | 0 | 48 | FALSE | 29 | 104 | 29 | 758 | 849 | 778 |
| -2,57076E | FALSE | High | IsMasterP | AAEL0019 | 2:193351 | 0 | 73 | FALSE | 23 | 92 | 3 | 351 | 404 | 780 |
| -5,632E+1 | FALSE | High | IsMasterP | AAEL0028 | 1:256024 | 0 | 58 | FALSE | 17 | 83 | 17 | 425 | 476 | 857 |
| 8,87795E | FALSE | High | IsMasterP | AAEL0172 | 3:780794 | 0 | 38 | FALSE | 31 | 96 | 31 | 925 | 1036 | 524 |
| 6,13186E | FALSE | High | IsMasterP | AAEL0055 | 1:873314 | 0 | 65 | FALSE | 20 | 80 | 20 | 280 | 298 | 890 |
| -5,07534E | FALSE | High | IsMasterP | AAEL0125 | 2:238123 | 0 | 68 | FALSE | 15 | 91 | 15 | 260 | 289 | 477 |
| 7,15772E | FALSE | High | IsMasterP | AAEL0047 | 2:288701 | 0 | 58 | FALSE | 27 | 94 | 27 | 499 | 556 | 632 |
| -2,80372E | FALSE | High | IsMasterP | AAEL0131 | 2:188425 | 0 | 94 | FALSE | 22 | 91 | 22 | 326 | 360 | 590 |
| 4,16864E | FALSE | High | IsMasterP | AAEL0041 | 3:401716 | 0 | 75 | FALSE | 20 | 87 | 20 | 318 | 358 | 684 |
| 7,95525E | FALSE | High | IsMasterP | AAEL0037 | 1:248483 | 0 | 51 | FALSE | 10 | 77 | 10 | 174 | 193 | 483 |
| -3,30549E | FALSE | High | IsMasterP | AAEL0142 | 2:406451 | 0 | 60 | FALSE | 25 | 99 | 25 | 487 | 549 | 625 |
| 2,39574E | FALSE | High | IsMasterP | AAEL0075 | 2:327281 | 0 | 67 | FALSE | 14 | 72 | 14 | 211 | 232 | 743 |
| 1,78194E | FALSE | High | IsMasterP | AAEL0068 | 3:369896 | 0 | 65 | FALSE | 26 | 94 | 26 | 564 | 614 | 850 |
| 4,94926E | FALSE | High | IsMasterP | AAEL0093 | 1:551287 | 0 | 68 | FALSE | 12 | 85 | 12 | 255 | 269 | 857 |

| | | | | | | | | | | | | | | |
|---|---|---|---|---|---|---|---|---|---|---|---|---|---|---|
| 3,11772E- | FALSE | High | IsMasterP | AAEL0065 | 2:594793 | 0 | 47 | FALSE | 19 | 87 | 19 | 484 | 553 | 596 |
| -8,24429E | FALSE | High | IsMasterP | AAEL0098 | 3:122234 | 0 | 65 | FALSE | 17 | 71 | 17 | 429 | 483 | 493 |
| 2,85726E- | FALSE | High | IsMasterP | AAEL0042 | 2:276975 | 0 | 60 | FALSE | 11 | 91 | 11 | 109 | 121 | 948 |
| -7,2238E+ | FALSE | High | IsMasterP | AAEL0136 | 1:624688 | 0 | 40 | FALSE | 28 | 93 | 16 | 882 | 998 | 929 |
| 8,64734E- | FALSE | High | IsMasterP | AAEL0027 | 3:145050 | 0 | 45 | FALSE | 25 | 103 | 3 | 424 | 458 | 458 |
| -3,70615E | FALSE | High | IsMasterP | AAEL0055 | 1:241890 | 0 | 65 | FALSE | 19 | 88 | 19 | 421 | 456 | 671 |
| 8,90255E- | FALSE | High | IsMasterP | AAEL0052 | 1:321206 | 0 | 76 | FALSE | 15 | 95 | 15 | 151 | 162 | 1035 |
| 7,57334E- | FALSE | High | IsMasterP | AAEL0068 | 3:565507 | 0 | 44 | FALSE | 15 | 102 | 14 | 327 | 345 | 907 |
| -2,02954E | FALSE | High | IsMasterP | AAEL0229 | 3:382021 | 0 | 36 | FALSE | 33 | 96 | 33 | 1648 | 1795 | 715 |
| 2,86001E- | FALSE | High | IsMasterP | AAEL0064 | 2:438674 | 0 | 64 | FALSE | 10 | 72 | 10 | 228 | 247 | 599 |
| 2,42192E- | FALSE | High | IsMasterP | AAEL0130 | 3:310492 | 0 | 66 | FALSE | 15 | 70 | 15 | 228 | 260 | 547 |
| -8,03076E | FALSE | High | IsMasterP | AAEL0085 | 2:994190 | 0 | 73 | FALSE | 31 | 89 | 30 | 463 | 511 | 796 |
| 1,88313E- | FALSE | High | IsMasterP | AAEL0051 | 1:140765 | 0 | 50 | FALSE | 20 | 80 | 20 | 468 | 525 | 787 |
| -3,66638E | FALSE | High | IsMasterP | AAEL0230 | 1:235737 | 0 | 68 | FALSE | 24 | 97 | 23 | 403 | 455 | 841 |
| 6,41533E- | FALSE | High | IsMasterP | AAEL0096 | 1:222703 | 0 | 67 | FALSE | 22 | 85 | 21 | 340 | 374 | 699 |
| 2,87251E- | FALSE | High | IsMasterP | AAEL0078 | 3:382192 | 0 | 77 | FALSE | 8 | 91 | 8 | 108 | 115 | 818 |
| 1,36239E- | FALSE | High | IsMasterP | AAEL0197 | 3:304098 | 0 | 49 | FALSE | 12 | 86 | 12 | 270 | 301 | 947 |
| 4,37128E- | FALSE | High | IsMasterP | AAEL0113 | 3:140270 | 0 | 69 | FALSE | 23 | 78 | 1 | 324 | 356 | 475 |
| -7,90538E | FALSE | High | IsMasterP | AAEL0018 | 2:276889 | 0 | 65 | FALSE | 28 | 85 | 28 | 698 | 782 | 692 |
| 7,68573E- | FALSE | High | IsMasterP | AAEL0012 | 2:411480 | 0 | 41 | FALSE | 33 | 100 | 33 | 966 | 1081 | 601 |
| 3,05471E- | FALSE | High | IsMasterP | AAEL0019 | 2:193432 | 0 | 82 | FALSE | 17 | 78 | 2 | 211 | 242 | 590 |
| 8,4169E+1 | FALSE | High | IsMasterP | AAEL0113 | 3:140270 | 0 | 67 | FALSE | 23 | 77 | 1 | 325 | 357 | 493 |
| 7,99692E- | FALSE | High | IsMasterP | AAEL0113 | 3:140277 | 0 | 77 | FALSE | 23 | 77 | 1 | 291 | 321 | 487 |
| 7,62467E- | FALSE | High | IsMasterP | AAEL0171 | 2:229895 | 0 | 81 | FALSE | 15 | 79 | 15 | 255 | 270 | 866 |
| -4,99544E | FALSE | High | IsMasterP | AAEL0017 | 2:374707 | 0 | 32 | FALSE | 33 | 92 | 33 | 1384 | 1571 | 679 |
| 2,55001E- | FALSE | High | IsMasterP | AAEL0102 | 3:409279 | 0 | 63 | FALSE | 12 | 87 | 12 | 140 | 153 | 667 |
| 6,58385E- | FALSE | High | IsMasterP | AAEL0290 | 3:235201 | 0 | 62 | FALSE | 21 | 84 | 21 | 305 | 333 | 875 |
| -5,98024E | FALSE | High | IsMasterP | AAEL0211 | NIGP0100 | 0 | 59 | FALSE | 28 | 80 | 28 | 724 | 806 | 647 |
| -2,61116E | FALSE | High | IsMasterP | AAEL0059 | 1:257458 | 0 | 66 | FALSE | 22 | 85 | 22 | 491 | 521 | 572 |
| -2,73515E | FALSE | High | IsMasterP | AAEL0058 | 1:116432 | 0 | 72 | FALSE | 16 | 80 | 16 | 249 | 281 | 844 |
| -5,26281E | FALSE | High | IsMasterP | AAEL0070 | 3:299827 | 0 | 69 | FALSE | 17 | 84 | 17 | 270 | 299 | 827 |

| | | | | | | | | | | | | | | |
|---|---|---|---|---|---|---|---|---|---|---|---|---|---|---|
| 3,82101E- | FALSE | High | IsMasterP | AAEL0098 | 3:1899038 | 0 | 36 | FALSE | 36 | 90 | 36 | 1550 | 1757 | 680 |
| 4,954E+18 | FALSE | High | IsMasterP | AAEL0027 | 3:1450250 | 0 | 22 | FALSE | 23 | 97 | 0 | 891 | 947 | 515 |
| 5,92211E- | FALSE | High | IsMasterP | AAEL0100 | 2:2005877 | 0 | 56 | FALSE | 30 | 80 | 30 | 877 | 977 | 676 |
| -3,28957E | FALSE | High | IsMasterP | AAEL0019 | 2:1931774 | 0 | 45 | FALSE | 22 | 88 | 2 | 554 | 639 | 740 |
| 5,13095E- | FALSE | High | IsMasterP | AAEL0027 | 3:1450250 | 0 | 22 | FALSE | 23 | 96 | 1 | 891 | 946 | 515 |
| -2,76844E | FALSE | High | IsMasterP | AAEL0021 | 1:1022716 | 0 | 75 | FALSE | 26 | 89 | 26 | 398 | 448 | 705 |
| -7,98634E | FALSE | High | IsMasterP | AAEL0198 | 2:3263366 | 0 | 29 | FALSE | 14 | 96 | 2 | 445 | 497 | 483 |
| -5,14478E | FALSE | High | IsMasterP | AAEL0103 | 3:4089805 | 0 | 44 | FALSE | 26 | 88 | 26 | 995 | 1081 | 674 |
| 1,62609E- | FALSE | High | IsMasterP | AAEL0094 | 1:9770190 | 0 | 17 | FALSE | 31 | 94 | 31 | 2224 | 2466 | 668 |
| 5,31966E- | FALSE | High | IsMasterP | AAEL0043 | 3:3656749 | 0 | 61 | FALSE | 17 | 79 | 17 | 354 | 385 | 680 |
| 3,95139E- | FALSE | High | IsMasterP | AAEL0145 | 2:2352182 | 0 | 52 | FALSE | 10 | 88 | 10 | 165 | 176 | 897 |
| -8,61208E | FALSE | High | IsMasterP | AAEL0041 | 2:2559079 | 0 | 57 | FALSE | 20 | 87 | 20 | 331 | 379 | 739 |
| -7,70986E | FALSE | High | IsMasterP | AAEL0012 | 2:6088955 | 0 | 41 | FALSE | 18 | 77 | 18 | 462 | 524 | 774 |
| 2,80693E- | FALSE | High | IsMasterP | AAEL0093 | 1:1998214 | 0 | 38 | FALSE | 17 | 86 | 1 | 445 | 482 | 681 |
| 8,02292E- | FALSE | High | IsMasterP | AAEL0014 | 2:4632062 | 0 | 34 | FALSE | 33 | 82 | 33 | 1246 | 1397 | 637 |
| -3,68401E | FALSE | High | IsMasterP | AAEL0028 | 3:2331842 | 0 | 29 | FALSE | 13 | 98 | 1 | 454 | 508 | 497 |
| -1,22917E | FALSE | High | IsMasterP | AAEL0081 | 3:1457207 | 0 | 59 | FALSE | 18 | 94 | 18 | 206 | 234 | 1071 |
| -1,13823E | FALSE | High | IsMasterP | AAEL0050 | 1:2946322 | 0 | 50 | FALSE | 19 | 92 | 9 | 449 | 505 | 492 |
| -7,23438E | FALSE | High | IsMasterP | AAEL0117 | 3:3670918 | 0 | 80 | FALSE | 19 | 76 | 19 | 293 | 330 | 585 |
| -4,36971E | FALSE | High | IsMasterP | AAEL0106 | 2:1761987 | 0 | 41 | FALSE | 33 | 83 | 33 | 1088 | 1217 | 474 |
| -3,76138E | FALSE | High | IsMasterP | AAEL0205 | 3:1746698 | 0 | 19 | FALSE | 27 | 85 | 27 | 1965 | 2178 | 720 |
| 3,38967E- | FALSE | High | IsMasterP | AAEL0202 | 1:6237353 | 0 | 62 | FALSE | 13 | 88 | 13 | 287 | 314 | 503 |
| -6,25362E | FALSE | High | IsMasterP | AAEL0050 | 2:4443094 | 0 | 59 | FALSE | 21 | 77 | 21 | 500 | 559 | 639 |
| -6,77287E | FALSE | High | IsMasterP | AAEL0052 | 2:1138063 | 0 | 63 | FALSE | 9 | 70 | 9 | 216 | 223 | 493 |
| 7,87859E- | FALSE | High | IsMasterP | AAEL0051 | 2:1729399 | 0 | 88 | FALSE | 11 | 74 | 11 | 136 | 144 | 595 |
| 9,01174E- | FALSE | High | IsMasterP | AAEL0135 | 2:2462414 | 0 | 62 | FALSE | 12 | 67 | 7 | 156 | 180 | 974 |
| 7,08323E- | FALSE | High | IsMasterP | AAEL0129 | 2:2518472 | 0 | 62 | FALSE | 21 | 78 | 21 | 443 | 499 | 559 |
| -5,12322E | FALSE | High | IsMasterP | AAEL0024 | 1:1770320 | 0 | 39 | FALSE | 21 | 94 | 21 | 382 | 459 | 477 |
| -8,75865E | FALSE | High | IsMasterP | AAEL0025 | 1:9039438 | 0 | 65 | FALSE | 21 | 67 | 21 | 573 | 623 | 868 |
| 3,85943E- | FALSE | High | IsMasterP | AAEL0117 | 2:1686855 | 0 | 52 | FALSE | 22 | 80 | 21 | 688 | 787 | 693 |
| -8,74456E | FALSE | High | IsMasterP | AAEL0020 | 1:5823199 | 0 | 65 | FALSE | 15 | 77 | 15 | 159 | 180 | 976 |

| | | | | | | | | | | | | | | |
|---|---|---|---|---|---|---|---|---|---|---|---|---|---|---|
| 5,14503E- | FALSE | High | IsMasterP | AAEL0117 | 3:291101 | 0 | 56 | FALSE | 21 | 80 | 21 | 449 | 488 | 787 |
| 5,43328E- | FALSE | High | IsMasterP | AAEL0010 | 1:299201 | 0 | 55 | FALSE | 28 | 78 | 28 | 624 | 687 | 712 |
| -3,12349E | FALSE | High | IsMasterP | AAEL0088 | 2:241374 | 0 | 55 | FALSE | 19 | 70 | 19 | 398 | 435 | 679 |
| 6,63648E- | FALSE | High | IsMasterP | AAEL0001 | 3:215991 | 0 | 62 | FALSE | 28 | 72 | 28 | 451 | 507 | 507 |
| -4,1952E+ | FALSE | High | IsMasterP | AAEL0009 | 2:104870 | 0 | 45 | FALSE | 14 | 76 | 14 | 261 | 286 | 1080 |
| 6,98948E- | FALSE | High | IsMasterP | AAEL0129 | 3:297242 | 0 | 58 | FALSE | 25 | 70 | 25 | 561 | 632 | 642 |
| -9,16539E | FALSE | High | IsMasterP | AAEL0019 | 2:193432 | 0 | 82 | FALSE | 16 | 71 | 1 | 210 | 241 | 567 |
| -9,20814E | FALSE | High | IsMasterP | AAEL0075 | 2:326933 | 0 | 63 | FALSE | 18 | 67 | 18 | 422 | 464 | 643 |
| -5,99007E | FALSE | High | IsMasterP | AAEL0068 | 2:256451 | 0 | 68 | FALSE | 14 | 81 | 2 | 248 | 282 | 488 |
| -4,63554E | FALSE | High | IsMasterP | AAEL0195 | 2:307390 | 0 | 18 | FALSE | 18 | 73 | 18 | 1235 | 1401 | 658 |
| -4,10655E | FALSE | High | IsMasterP | AAEL0087 | 2:198810 | 0 | 69 | FALSE | 15 | 80 | 15 | 231 | 256 | 679 |
| 6,94816E- | FALSE | High | IsMasterP | AAEL0117 | 2:105789 | 0 | 57 | FALSE | 10 | 67 | 10 | 219 | 232 | 467 |
| 5,29708E- | FALSE | High | IsMasterP | AAEL0099 | 2:379695 | 0 | 60 | FALSE | 20 | 82 | 19 | 488 | 527 | 586 |
| 9,18127E- | FALSE | High | IsMasterP | AAEL0233 | 1:206664 | 0 | 41 | FALSE | 18 | 73 | 18 | 616 | 682 | 610 |
| 2,22539E- | FALSE | High | IsMasterP | AAEL0128 | 1:242520 | 0 | 60 | FALSE | 24 | 78 | 24 | 619 | 677 | 740 |
| -3,89015E | FALSE | High | IsMasterP | AAEL0046 | 1:159001 | 0 | 78 | FALSE | 23 | 70 | 23 | 456 | 501 | 624 |
| -5,54967E | FALSE | High | IsMasterP | AAEL0083 | 1:819965 | 0 | 60 | FALSE | 20 | 78 | 20 | 432 | 476 | 573 |
| 8,63417E- | FALSE | High | IsMasterP | AAEL0139 | 2:965090 | 0 | 63 | FALSE | 14 | 71 | 14 | 324 | 338 | 848 |
| 8,25161E- | FALSE | High | IsMasterP | AAEL0129 | 3:206206 | 0 | 69 | FALSE | 24 | 79 | 24 | 434 | 486 | 564 |
| 8,65233E- | FALSE | High | IsMasterP | AAEL0195 | 1:267632 | 0 | 30 | FALSE | 23 | 80 | 3 | 1065 | 1205 | 620 |
| 3,92523E- | FALSE | High | IsMasterP | AAEL0090 | 2:101731 | 0 | 50 | FALSE | 17 | 70 | 17 | 503 | 574 | 558 |
| -9,34457E | FALSE | High | IsMasterP | AAEL0071 | 3:698688 | 0 | 13 | FALSE | 37 | 78 | 37 | 4684 | 5343 | 635 |
| -6,7019E+ | FALSE | High | IsMasterP | AAEL0195 | 1:267632 | 0 | 30 | FALSE | 21 | 80 | 1 | 995 | 1124 | 521 |
| -2,41287E | FALSE | High | IsMasterP | AAEL0109 | 2:151224 | 0 | 61 | FALSE | 18 | 64 | 18 | 326 | 371 | 562 |
| 7,29684E- | FALSE | High | IsMasterP | AAEL0092 | 2:180827 | 0 | 65 | FALSE | 15 | 82 | 15 | 214 | 245 | 781 |
| 3,37467E- | FALSE | High | IsMasterP | AAEL0134 | 2:159254 | 0 | 59 | FALSE | 21 | 71 | 12 | 505 | 569 | 785 |
| 1,89094E- | FALSE | High | IsMasterP | AAEL0116 | 3:218355 | 0 | 54 | FALSE | 6 | 65 | 6 | 149 | 170 | 1051 |
| 6,89516E- | FALSE | High | IsMasterP | AAEL0064 | 3:216574 | 0 | 71 | FALSE | 15 | 69 | 13 | 377 | 404 | 705 |
| -5,46927E | FALSE | High | IsMasterP | AAEL0115 | 3:312847 | 0 | 40 | FALSE | 11 | 66 | 11 | 446 | 470 | 621 |
| 5,25987E- | FALSE | High | IsMasterP | AAEL0221 | 3:183217 | 0 | 41 | FALSE | 17 | 78 | 17 | 415 | 468 | 1023 |
| -8,17971E | FALSE | High | IsMasterP | AAEL0060 | 3:740427 | 0 | 67 | FALSE | 15 | 67 | 15 | 233 | 257 | 655 |

| | | | | | | | | | | | | | | |
|---|---|---|---|---|---|---|---|---|---|---|---|---|---|---|
| 4,50477E- | FALSE | High | IsMasterP | AAEL0019 | 2:4255327 | 0 | 66 | FALSE | 22 | 75 | 22 | 441 | 487 | 658 |
| -3,09759E | FALSE | High | IsMasterP | AAEL0056 | 3:1601248 | 0 | 57 | FALSE | 24 | 74 | 24 | 512 | 579 | 717 |
| 6,42361E- | FALSE | High | IsMasterP | AAEL0028 | 3:2349633 | 0 | 50 | FALSE | 15 | 68 | 15 | 339 | 380 | 610 |
| 4,24747E- | FALSE | High | IsMasterP | AAEL0126 | 2:2948499 | 0 | 80 | FALSE | 12 | 70 | 12 | 147 | 160 | 615 |
| -3,9771E+ | FALSE | High | IsMasterP | AAEL0059 | 3:1740515 | 0 | 39 | FALSE | 14 | 64 | 14 | 407 | 444 | 441 |
| 5,32957E- | FALSE | High | IsMasterP | AAEL0195 | 2:4371144 | 0 | 39 | FALSE | 21 | 73 | 21 | 667 | 746 | 705 |
| -2,70972E | FALSE | High | IsMasterP | AAEL0253 | NIGP0100 | 0 | 75 | FALSE | 13 | 60 | 13 | 219 | 244 | 825 |
| -6,66803E | FALSE | High | IsMasterP | AAEL0259 | NIGP0100 | 0 | 72 | FALSE | 11 | 72 | 11 | 130 | 152 | 988 |
| -9,04467E | FALSE | High | IsMasterP | AAEL0076 | 2:2508046 | 0 | 53 | FALSE | 18 | 66 | 18 | 486 | 533 | 869 |
| 2,09255E- | FALSE | High | IsMasterP | AAEL0071 | 2:1499707 | 0 | 83 | FALSE | 16 | 77 | 3 | 213 | 234 | 536 |
| 1,19345E- | FALSE | High | IsMasterP | AAEL0233 | 3:3599460 | 0 | 42 | FALSE | 25 | 70 | 25 | 865 | 968 | 609 |
| -9,61174E | FALSE | High | IsMasterP | AAEL0077 | 2:2193461 | 0 | 56 | FALSE | 10 | 78 | 10 | 151 | 176 | 969 |
| -5,9385E+ | FALSE | High | IsMasterP | AAEL0132 | 3:3410311 | 0 | 55 | FALSE | 14 | 65 | 14 | 217 | 246 | 983 |
| -2,25155E | FALSE | High | IsMasterP | AAEL0097 | 1:4146382 | 0 | 50 | FALSE | 20 | 71 | 20 | 616 | 683 | 602 |
| -3,99743E | FALSE | High | IsMasterP | AAEL0194 | 2:6371673 | 0 | 18 | FALSE | 41 | 71 | 41 | 3902 | 4335 | 550 |
| -6,39345E | FALSE | High | IsMasterP | AAEL0043 | 2:3866529 | 0 | 58 | FALSE | 18 | 76 | 18 | 297 | 340 | 976 |
| -6,77877E | FALSE | High | IsMasterP | AAEL0124 | 3:1114086 | 0 | 59 | FALSE | 19 | 68 | 19 | 416 | 468 | 572 |
| -5,48263E | FALSE | High | IsMasterP | AAEL0128 | 2:3352225 | 0 | 54 | FALSE | 18 | 79 | 18 | 262 | 301 | 793 |
| 8,94838E- | FALSE | High | IsMasterP | AAEL0055 | 3:4043619 | 0 | 81 | FALSE | 17 | 61 | 17 | 185 | 197 | 521 |
| 5,56128E- | FALSE | High | IsMasterP | AAEL0088 | 2:2413287 | 0 | 53 | FALSE | 16 | 78 | 16 | 297 | 327 | 888 |
| -5,68032E | FALSE | High | IsMasterP | AAEL0030 | 1:1291102 | 0 | 38 | FALSE | 25 | 70 | 25 | 1024 | 1150 | 503 |
| -5,37691E | FALSE | High | IsMasterP | AAEL0070 | 1:2117067 | 0 | 28 | FALSE | 28 | 74 | 28 | 1133 | 1332 | 866 |
| 5,20883E- | FALSE | High | IsMasterP | AAEL0033 | 3:8136133 | 0 | 46 | FALSE | 9 | 78 | 9 | 134 | 158 | 1095 |
| -7,46264E | FALSE | High | IsMasterP | AAEL0057 | 1:1223858 | 0 | 62 | FALSE | 10 | 64 | 10 | 138 | 159 | 591 |
| -6,54454E | FALSE | High | IsMasterP | AAEL0096 | 2:1698840 | 0 | 29 | FALSE | 17 | 56 | 17 | 1273 | 1339 | 737 |
| -8,82995E | FALSE | High | IsMasterP | AAEL0053 | 3:3093724 | 0 | 58 | FALSE | 19 | 63 | 19 | 422 | 459 | 834 |
| -5,69199E | FALSE | High | IsMasterP | AAEL0100 | 1:1941878 | 0 | 37 | FALSE | 25 | 70 | 25 | 738 | 820 | 891 |
| -9,09009E | FALSE | High | IsMasterP | AAEL0048 | 1:2881966 | 0 | 50 | FALSE | 10 | 74 | 10 | 240 | 263 | 686 |
| 2,52988E- | FALSE | High | IsMasterP | AAEL0048 | 2:1152844 | 0 | 51 | FALSE | 17 | 80 | 16 | 302 | 329 | 966 |
| 1,55325E- | FALSE | High | IsMasterP | AAEL0059 | 3:1385811 | 0 | 50 | FALSE | 17 | 63 | 17 | 482 | 530 | 771 |
| -2,60541E | FALSE | High | IsMasterP | AAEL0118 | 3:1132041 | 0 | 35 | FALSE | 24 | 75 | 24 | 817 | 913 | 624 |

| | | | | | | | | | | | | | | |
|---|---|---|---|---|---|---|---|---|---|---|---|---|---|---|
| -5,59263E | FALSE | High | IsMasterP | AAEL0130 | 1:3173777 | 0 | 42 | FALSE | 19 | 76 | 19 | 378 | 414 | 442 |
| -2,43217E | FALSE | High | IsMasterP | AAEL0076 | 2:4703206 | 0 | 63 | FALSE | 11 | 70 | 11 | 190 | 214 | 992 |
| 7,73305E+ | FALSE | High | IsMasterP | AAEL0016 | 2:3003215 | 0 | 65 | FALSE | 25 | 66 | 25 | 522 | 557 | 725 |
| -2,49831E | FALSE | High | IsMasterP | AAEL0103 | 2:4037797 | 0 | 61 | FALSE | 22 | 68 | 22 | 465 | 518 | 576 |
| 6,69047E+ | FALSE | High | IsMasterP | AAEL0234 | NIGP0100 | 0 | 69 | FALSE | 15 | 69 | 15 | 160 | 176 | 557 |
| 2,86914E+ | FALSE | High | IsMasterP | AAEL0117 | 3:2887793 | 0 | 51 | FALSE | 17 | 76 | 17 | 467 | 516 | 882 |
| 7,9902E+1 | FALSE | High | IsMasterP | AAEL0100 | 3:1069842 | 0 | 47 | FALSE | 17 | 65 | 17 | 437 | 476 | 534 |
| -1,21523E | FALSE | High | IsMasterP | AAEL0134 | 2:3511432 | 0 | 18 | FALSE | 28 | 66 | 28 | 2437 | 2653 | 498 |
| 3,12801E+ | FALSE | High | IsMasterP | AAEL0136 | 2:3343539 | 0 | 50 | FALSE | 8 | 64 | 8 | 219 | 246 | 944 |
| 4,80585E+ | FALSE | High | IsMasterP | AAEL0036 | 3:6572711 | 0 | 44 | FALSE | 22 | 74 | 22 | 541 | 592 | 853 |
| 5,48212E+ | FALSE | High | IsMasterP | AAEL0029 | 1:1083993 | 0 | 46 | FALSE | 28 | 63 | 28 | 1016 | 1113 | 580 |
| 5,05378E+ | FALSE | High | IsMasterP | AAEL0061 | 1:1260304 | 0 | 45 | FALSE | 13 | 61 | 13 | 387 | 418 | 554 |
| -3,06434E | FALSE | High | IsMasterP | AAEL0150 | 2:3377866 | 0 | 66 | FALSE | 13 | 65 | 13 | 140 | 149 | 1083 |
| -3,52541E | FALSE | High | IsMasterP | AAEL0063 | 3:2828219 | 0 | 42 | FALSE | 24 | 65 | 24 | 899 | 992 | 535 |
| -5,59686E | FALSE | High | IsMasterP | AAEL0066 | 3:3532292 | 0 | 82 | FALSE | 21 | 66 | 21 | 356 | 399 | 615 |
| 8,165E+18 | FALSE | High | IsMasterP | AAEL0197 | 2:2321006 | 0 | 31 | FALSE | 15 | 65 | 15 | 645 | 732 | 869 |
| 5,49078E+ | FALSE | High | IsMasterP | AAEL0039 | 3:2674585 | 0 | 41 | FALSE | 22 | 59 | 22 | 833 | 940 | 488 |
| -7,90317E | FALSE | High | IsMasterP | AAEL0080 | 1:6249005 | 0 | 39 | FALSE | 26 | 66 | 13 | 875 | 993 | 931 |
| 5,51219E+ | FALSE | High | IsMasterP | AAEL0027 | 1:2315774 | 0 | 35 | FALSE | 14 | 61 | 14 | 491 | 519 | 922 |
| 4,14028E+ | FALSE | High | IsMasterP | AAEL0052 | 3:9058136 | 0 | 55 | FALSE | 8 | 71 | 8 | 116 | 128 | 969 |
| 2,05874E+ | FALSE | High | IsMasterP | AAEL0132 | 2:3497605 | 0 | 31 | FALSE | 20 | 67 | 20 | 904 | 1001 | 486 |
| -7,98215E | FALSE | High | IsMasterP | AAEL0182 | 3:2340417 | 0 | 8 | FALSE | 37 | 67 | 37 | 9019 | 10101 | 583 |
| 6,86821E+ | FALSE | High | IsMasterP | AAEL0082 | 3:6573829 | 0 | 35 | FALSE | 18 | 66 | 18 | 398 | 422 | 703 |
| -8,55709E | FALSE | High | IsMasterP | AAEL0068 | 2:2564515 | 0 | 54 | FALSE | 13 | 70 | 1 | 248 | 283 | 503 |
| -7,07222E | FALSE | High | IsMasterP | AAEL0053 | 3:2357113 | 0 | 33 | FALSE | 27 | 63 | 27 | 1342 | 1481 | 591 |
| 3,62412E+ | FALSE | High | IsMasterP | AAEL0018 | 2:3893455 | 0 | 55 | FALSE | 14 | 64 | 14 | 400 | 446 | 705 |
| -3,04628E | FALSE | High | IsMasterP | AAEL0255 | 3:1776203 | 0 | 78 | FALSE | 13 | 63 | 13 | 234 | 234 | 853 |
| -6,5062E+ | FALSE | High | IsMasterP | AAEL0057 | 2:4361914 | 0 | 34 | FALSE | 20 | 62 | 20 | 1008 | 1155 | 534 |
| 7,65034E+ | FALSE | High | IsMasterP | AAEL0058 | 2:6633799 | 0 | 53 | FALSE | 26 | 63 | 26 | 964 | 1071 | 652 |
| 8,60821E+ | FALSE | High | IsMasterP | AAEL0020 | 1:5839417 | 0 | 75 | FALSE | 20 | 59 | 20 | 347 | 383 | 470 |
| -8,59734E | FALSE | High | IsMasterP | AAEL0061 | 1:1260863 | 0 | 51 | FALSE | 6 | 53 | 2 | 136 | 153 | 1127 |

| | | | | | | | | | | | | | | |
|---|---|---|---|---|---|---|---|---|---|---|---|---|---|---|
| 8,69723E+ | FALSE | High | IsMasterP | AAEL0068 | 3:4367011 | 0 | 38 | FALSE | 21 | 64 | 21 | 801 | 869 | 721 |
| -3,60599E | FALSE | High | IsMasterP | AAEL0066 | 2:3263181 | 0 | 49 | FALSE | 28 | 67 | 28 | 796 | 874 | 582 |
| -6,49694E | FALSE | High | IsMasterP | AAEL0147 | 2:4645965 | 0 | 44 | FALSE | 29 | 70 | 29 | 966 | 1080 | 793 |
| 8,69859E+ | FALSE | High | IsMasterP | AAEL0081 | 3:1002847 | 0 | 88 | FALSE | 17 | 64 | 17 | 256 | 271 | 771 |
| -9,01333E | FALSE | High | IsMasterP | AAEL0136 | 3:2355373 | 0 | 55 | FALSE | 19 | 66 | 19 | 338 | 381 | 478 |
| -1,86223E | FALSE | High | IsMasterP | AAEL0182 | 2:2093545 | 0 | 35 | FALSE | 15 | 56 | 15 | 692 | 738 | 445 |
| 3,10019E+ | FALSE | High | IsMasterP | AAEL0086 | 2:3214998 | 0 | 66 | FALSE | 9 | 73 | 9 | 119 | 137 | 828 |
| -6,07854E | FALSE | High | IsMasterP | AAEL0005 | 2:1475936 | 0 | 54 | FALSE | 14 | 55 | 14 | 380 | 412 | 494 |
| 2,76604E+ | FALSE | High | IsMasterP | AAEL0028 | 1:2553105 | 0 | 52 | FALSE | 22 | 66 | 22 | 568 | 635 | 828 |
| -6,65335E | FALSE | High | IsMasterP | AAEL0041 | 3:1539812 | 0 | 64 | FALSE | 18 | 60 | 18 | 404 | 438 | 761 |
| 7,29266E+ | FALSE | High | IsMasterP | AAEL0131 | 2:3890512 | 0 | 48 | FALSE | 20 | 67 | 20 | 679 | 761 | 777 |
| 6,93321E+ | FALSE | High | IsMasterP | AAEL0210 | 3:3157502 | 0 | 41 | FALSE | 19 | 70 | 19 | 532 | 599 | 724 |
| -3,22488E | FALSE | High | IsMasterP | AAEL0108 | 3:5216822 | 0 | 50 | FALSE | 9 | 47 | 9 | 393 | 433 | 674 |
| 4,95844E+ | FALSE | High | IsMasterP | AAEL0107 | 2:3053096 | 0 | 91 | FALSE | 14 | 62 | 14 | 106 | 120 | 526 |
| 3,86089E+ | FALSE | High | IsMasterP | AAEL0273 | 2:4604050 | 0 | 23 | FALSE | 27 | 64 | 27 | 1758 | 1958 | 547 |
| 7,83922E+ | FALSE | High | IsMasterP | AAEL0078 | 2:4607315 | 0 | 44 | FALSE | 19 | 60 | 19 | 583 | 639 | 450 |
| 1,34051E+ | FALSE | High | IsMasterP | AAEL0101 | 3:1492604 | 0 | 37 | FALSE | 24 | 62 | 24 | 1102 | 1240 | 473 |
| -3,00944E | FALSE | High | IsMasterP | AAEL0118 | 2:3962271 | 0 | 39 | FALSE | 12 | 53 | 12 | 394 | 425 | 493 |
| -4,85338E | FALSE | High | IsMasterP | AAEL0101 | 3:1493528 | 0 | 41 | FALSE | 13 | 75 | 13 | 275 | 299 | 1032 |
| 2,13174E+ | FALSE | High | IsMasterP | AAEL0092 | 2:1743559 | 0 | 43 | FALSE | 16 | 60 | 16 | 481 | 545 | 639 |
| 5,38373E+ | FALSE | High | IsMasterP | AAEL0122 | 1:1609095 | 0 | 86 | FALSE | 12 | 56 | 12 | 159 | 181 | 459 |
| -3,76326E | FALSE | High | IsMasterP | AAEL0010 | 3:5521880 | 0 | 85 | FALSE | 10 | 71 | 10 | 100 | 107 | 904 |
| 4,13174E+ | FALSE | High | IsMasterP | AAEL0109 | 3:9696429 | 0 | 35 | FALSE | 18 | 62 | 18 | 715 | 802 | 791 |
| -7,46156E | FALSE | High | IsMasterP | AAEL0244 | 1:2854556 | 0 | 21 | FALSE | 21 | 61 | 21 | 1562 | 1715 | 863 |
| 7,1246E+1 | FALSE | High | IsMasterP | AAEL0026 | 2:4649129 | 0 | 63 | FALSE | 22 | 53 | 22 | 544 | 592 | 778 |
| 3,88326E+ | FALSE | High | IsMasterP | AAEL0001 | 3:2123904 | 0 | 36 | FALSE | 26 | 64 | 26 | 910 | 1060 | 628 |
| -6,4647E+ | FALSE | High | IsMasterP | AAEL0138 | 2:1002164 | 0 | 49 | FALSE | 12 | 65 | 12 | 250 | 290 | 529 |
| -2,83565E | FALSE | High | IsMasterP | AAEL0129 | 2:1602407 | 0 | 30 | FALSE | 23 | 65 | 23 | 1066 | 1202 | 680 |
| 1,51773E+ | FALSE | High | IsMasterP | AAEL0041 | 2:1473460 | 0 | 75 | FALSE | 16 | 51 | 14 | 288 | 326 | 559 |
| -1,94222E | FALSE | High | IsMasterP | AAEL0050 | 1:2947351 | 0 | 51 | FALSE | 16 | 66 | 9 | 448 | 505 | 492 |
| -7,31259E | FALSE | High | IsMasterP | AAEL0122 | 1:1466210 | 0 | 39 | FALSE | 20 | 55 | 20 | 737 | 833 | 888 |

| | | | | | | | | | | | | | | |
|---|---|---|---|---|---|---|---|---|---|---|---|---|---|---|
| 7,78445E- | FALSE | High | IsMasterP | AAEL0149 | 2:222780 | 0 | 46 | FALSE | 18 | 64 | 18 | 430 | 457 | 717 |
| 7,06466E- | FALSE | High | IsMasterP | AAEL0170 | 2:213474 | 0 | 54 | FALSE | 10 | 59 | 10 | 257 | 296 | 851 |
| -1,41673E | FALSE | High | IsMasterP | AAEL0136 | 1:155073 | 0 | 80 | FALSE | 14 | 60 | 14 | 130 | 146 | 564 |
| -9,21584E | FALSE | High | IsMasterP | AAEL0098 | 3:121966 | 0 | 38 | FALSE | 22 | 55 | 22 | 798 | 907 | 660 |
| -7,46956E | FALSE | High | IsMasterP | AAEL0115 | 3:281212 | 0 | 54 | FALSE | 21 | 56 | 21 | 496 | 537 | 904 |
| -8,19796E | FALSE | High | IsMasterP | AAEL0036 | 1:189335 | 0 | 65 | FALSE | 11 | 55 | 11 | 327 | 359 | 475 |
| -8,56602E | FALSE | High | IsMasterP | AAEL0065 | 2:133180 | 0 | 53 | FALSE | 15 | 54 | 15 | 336 | 368 | 610 |
| 7,57276E- | FALSE | High | IsMasterP | AAEL0266 | NIGP0100 | 0 | 33 | FALSE | 18 | 58 | 18 | 759 | 812 | 692 |
| -1,50539E | FALSE | High | IsMasterP | AAEL0071 | 2:149970 | 0 | 42 | FALSE | 14 | 60 | 1 | 382 | 423 | 493 |
| 4,5523E+1 | FALSE | High | IsMasterP | AAEL0025 | 1:904900 | 0 | 41 | FALSE | 16 | 62 | 16 | 397 | 444 | 521 |
| -6,32226E | FALSE | High | IsMasterP | AAEL0051 | 2:113794 | 0 | 50 | FALSE | 9 | 41 | 9 | 116 | 128 | 434 |
| -3,13295E | FALSE | High | IsMasterP | AAEL0083 | 2:274104 | 0 | 50 | FALSE | 27 | 65 | 27 | 636 | 714 | 847 |
| -6,8964E+ | FALSE | High | IsMasterP | AAEL0114 | 2:371576 | 0 | 46 | FALSE | 12 | 73 | 12 | 178 | 205 | 1108 |
| 1,41086E- | FALSE | High | IsMasterP | AAEL0010 | 2:101783 | 0 | 65 | FALSE | 14 | 60 | 6 | 340 | 371 | 644 |
| 6,28681E- | FALSE | High | IsMasterP | AAEL0129 | 3:297288 | 0 | 61 | FALSE | 11 | 56 | 11 | 203 | 234 | 530 |
| -2,84914E | FALSE | High | IsMasterP | AAEL0057 | 1:123756 | 0 | 47 | FALSE | 17 | 64 | 17 | 270 | 306 | 1039 |
| -7,80294E | FALSE | High | IsMasterP | AAEL0107 | 3:294224 | 0 | 57 | FALSE | 14 | 64 | 14 | 156 | 173 | 996 |
| 6,3415E+1 | FALSE | High | IsMasterP | AAEL0069 | 2:398786 | 0 | 47 | FALSE | 20 | 65 | 20 | 594 | 663 | 505 |
| 6,15568E- | FALSE | High | IsMasterP | AAEL0174 | 1:241962 | 0 | 47 | FALSE | 10 | 59 | 10 | 176 | 187 | 813 |
| 3,89193E- | FALSE | High | IsMasterP | AAEL0086 | 2:235058 | 0 | 69 | FALSE | 15 | 57 | 13 | 254 | 290 | 577 |
| -1,11963E | FALSE | High | IsMasterP | AAEL0054 | 1:247168 | 0 | 76 | FALSE | 17 | 50 | 13 | 321 | 361 | 511 |
| 7,66765E- | FALSE | High | IsMasterP | AAEL0075 | 1:302208 | 0 | 15 | FALSE | 23 | 62 | 23 | 1892 | 2035 | 674 |
| -3,78061E | FALSE | High | IsMasterP | AAEL0120 | 3:147280 | 0 | 62 | FALSE | 11 | 57 | 11 | 226 | 257 | 614 |
| -4,00235E | FALSE | High | IsMasterP | AAEL0020 | 2:356637 | 0 | 55 | FALSE | 22 | 60 | 22 | 546 | 600 | 701 |
| -1,46508E | FALSE | High | IsMasterP | AAEL0042 | 2:293048 | 0 | 36 | FALSE | 15 | 63 | 2 | 503 | 538 | 890 |
| -7,79158E | FALSE | High | IsMasterP | AAEL0269 | 1:194250 | 0 | 43 | FALSE | 14 | 54 | 14 | 371 | 408 | 769 |
| 8,36132E- | FALSE | High | IsMasterP | AAEL0174 | 1:150839 | 0 | 46 | FALSE | 20 | 57 | 20 | 552 | 616 | 866 |
| 3,84688E- | FALSE | High | IsMasterP | AAEL0047 | 2:290350 | 0 | 52 | FALSE | 21 | 57 | 21 | 630 | 690 | 799 |
| -2,68212E | FALSE | High | IsMasterP | AAEL0004 | 3:186237 | 0 | 48 | FALSE | 15 | 54 | 15 | 396 | 436 | 850 |
| -3,21355E | FALSE | High | IsMasterP | AAEL0046 | 1:306673 | 0 | 46 | FALSE | 20 | 56 | 20 | 590 | 657 | 621 |
| -3,68118E | FALSE | High | IsMasterP | AAEL0143 | 3:289162 | 0 | 63 | FALSE | 12 | 55 | 12 | 205 | 234 | 679 |

| | | | | | | | | | | | | | | |
|---|---|---|---|---|---|---|---|---|---|---|---|---|---|---|
| -9,61457E | FALSE | High | IsMasterP | AAEL0124 | 1:2129188 | 0 | 30 | FALSE | 16 | 55 | 16 | 784 | 879 | 780 |
| 8,70588E+ | FALSE | High | IsMasterP | AAEL0236 | 1:4291674 | 0 | 60 | FALSE | 7 | 60 | 6 | 126 | 132 | 1040 |
| -1,56895E | FALSE | High | IsMasterP | AAEL0060 | 1:1840121 | 0 | 84 | FALSE | 16 | 56 | 16 | 255 | 285 | 681 |
| -9,75095E | FALSE | High | IsMasterP | AAEL0102 | 3:2790075 | 0 | 65 | FALSE | 19 | 59 | 19 | 333 | 337 | 1020 |
| 8,91665E+ | FALSE | High | IsMasterP | AAEL0074 | 2:2913837 | 0 | 75 | FALSE | 11 | 54 | 11 | 153 | 176 | 467 |
| -5,29789E | FALSE | High | IsMasterP | AAEL0034 | 3:3501522 | 0 | 49 | FALSE | 11 | 50 | 11 | 280 | 304 | 676 |
| -6,67759E | FALSE | High | IsMasterP | AAEL0125 | 2:2381354 | 0 | 38 | FALSE | 24 | 58 | 24 | 810 | 918 | 515 |
| 1,06909E+ | FALSE | High | IsMasterP | AAEL0014 | 2:4825682 | 0 | 41 | FALSE | 18 | 50 | 18 | 491 | 567 | 568 |
| 2,1029E+1 | FALSE | High | IsMasterP | AAEL0132 | 2:3994506 | 0 | 32 | FALSE | 17 | 59 | 17 | 674 | 749 | 667 |
| -6,14415E | FALSE | High | IsMasterP | AAEL0137 | 3:2542312 | 0 | 51 | FALSE | 21 | 56 | 21 | 661 | 736 | 766 |
| 7,7417E+1 | FALSE | High | IsMasterP | AAEL0229 | 3:1296799 | 0 | 46 | FALSE | 18 | 52 | 18 | 520 | 601 | 747 |
| -5,0388E+ | FALSE | High | IsMasterP | AAEL0211 | 2:4609135 | 0 | 26 | FALSE | 32 | 55 | 32 | 1869 | 2109 | 540 |
| -6,20442E | FALSE | High | IsMasterP | AAEL0139 | 2:1018258 | 0 | 62 | FALSE | 15 | 55 | 1 | 340 | 373 | 652 |
| 8,32789E+ | FALSE | High | IsMasterP | AAEL0239 | 2:1789887 | 0 | 44 | FALSE | 12 | 58 | 12 | 262 | 296 | 1030 |
| -9,04787E | FALSE | High | IsMasterP | AAEL0281 | 1:1461129 | 0 | 45 | FALSE | 16 | 52 | 14 | 443 | 501 | 834 |
| 6,71265E+ | FALSE | High | IsMasterP | AAEL0130 | 2:2535511 | 0 | 21 | FALSE | 15 | 49 | 15 | 1129 | 1227 | 458 |
| 4,63795E+ | FALSE | High | IsMasterP | AAEL0024 | 3:3381924 | 0 | 40 | FALSE | 21 | 54 | 21 | 726 | 811 | 679 |
| -7,31076E | FALSE | High | IsMasterP | AAEL0156 | 1:1082306 | 0 | 83 | FALSE | 15 | 59 | 15 | 247 | 267 | 843 |
| 8,87031E+ | FALSE | High | IsMasterP | AAEL0110 | 2:2321488 | 0 | 44 | FALSE | 13 | 58 | 13 | 271 | 303 | 501 |
| 3,11989E+ | FALSE | High | IsMasterP | AAEL0072 | 2:4214849 | 0 | 45 | FALSE | 9 | 48 | 9 | 385 | 415 | 475 |
| -5,18075E | FALSE | High | IsMasterP | AAEL0028 | 1:2544717 | 0 | 35 | FALSE | 20 | 56 | 20 | 723 | 819 | 519 |
| 5,98341E+ | FALSE | High | IsMasterP | AAEL0173 | 3:3472765 | 0 | 80 | FALSE | 15 | 56 | 12 | 245 | 265 | 758 |
| 1,29783E+ | FALSE | High | IsMasterP | AAEL0084 | 2:1904524 | 0 | 66 | FALSE | 11 | 39 | 11 | 259 | 290 | 505 |
| -7,14892E | FALSE | High | IsMasterP | AAEL0112 | 2:2263137 | 0 | 33 | FALSE | 18 | 55 | 18 | 836 | 927 | 554 |
| 8,74299E+ | FALSE | High | IsMasterP | AAEL0181 | 3:3666283 | 0 | 60 | FALSE | 6 | 46 | 6 | 142 | 162 | 558 |
| -5,02163E | FALSE | High | IsMasterP | AAEL0012 | 2:4133383 | 0 | 36 | FALSE | 18 | 49 | 18 | 754 | 868 | 728 |
| -4,1903E+ | FALSE | High | IsMasterP | AAEL0173 | 2:4041682 | 0 | 45 | FALSE | 19 | 57 | 19 | 591 | 641 | 547 |
| -8,98726E | FALSE | High | IsMasterP | AAEL0027 | 2:9624842 | 0 | 47 | FALSE | 17 | 48 | 17 | 534 | 598 | 718 |
| 3,64633E+ | FALSE | High | IsMasterP | AAEL0049 | 2:7241999 | 0 | 26 | FALSE | 20 | 56 | 20 | 1061 | 1184 | 475 |
| 5,85669E+ | FALSE | High | IsMasterP | AAEL0255 | 2:2907302 | 0 | 8 | FALSE | 27 | 52 | 27 | 5246 | 5934 | 597 |
| -1,63047E | FALSE | High | IsMasterP | AAEL0081 | 3:5358906 | 0 | 78 | FALSE | 21 | 65 | 21 | 243 | 269 | 961 |

| | | | | | | | | | | | | | | |
|---|---|---|---|---|---|---|---|---|---|---|---|---|---|---|
| 5,21066E- | FALSE | High | IsMasterP | AAEL0122 | 2:145020! | 0 | 32 | FALSE | 22 | 58 | 22 | 1008 | 1154 | 660 |
| -8,76252E | FALSE | High | IsMasterP | AAEL0077 | 3:255431( | 0 | 40 | FALSE | 18 | 54 | 18 | 688 | 801 | 580 |
| -3,05682E | FALSE | High | IsMasterP | AAEL0124 | 1:213297 | 0 | 61 | FALSE | 13 | 52 | 13 | 317 | 350 | 790 |
| -6,16022E | FALSE | High | IsMasterP | AAEL0116 | 3:281587 | 0 | 64 | FALSE | 11 | 50 | 11 | 271 | 292 | 837 |
| -1,52335E | FALSE | High | IsMasterP | AAEL0009 | 3:377081 | 0 | 68 | FALSE | 8 | 48 | 2 | 254 | 284 | 472 |
| 6,53686E- | FALSE | High | IsMasterP | AAEL0064 | 2:414370 | 0 | 30 | FALSE | 23 | 57 | 23 | 1182 | 1347 | 730 |
| -4,04401E | FALSE | High | IsMasterP | AAEL0003 | 3:266486( | 0 | 35 | FALSE | 19 | 56 | 18 | 703 | 797 | 712 |
| -1,53067E | FALSE | High | IsMasterP | AAEL0042 | 2:293048( | 0 | 35 | FALSE | 14 | 57 | 1 | 487 | 522 | 882 |
| -3,56428E | FALSE | High | IsMasterP | AAEL0036 | 2:316610 | 0 | 48 | FALSE | 20 | 55 | 20 | 393 | 457 | 600 |
| -4,06532E | FALSE | High | IsMasterP | AAEL0042 | 2:293502 | 0 | 25 | FALSE | 18 | 52 | 18 | 1153 | 1255 | 739 |
| -3,59501E | FALSE | High | IsMasterP | AAEL0069 | 3:143588 | 0 | 77 | FALSE | 13 | 50 | 13 | 188 | 207 | 812 |
| 1,94444E- | FALSE | High | IsMasterP | AAEL0016 | 2:906965 | 0 | 31 | FALSE | 8 | 49 | 8 | 392 | 437 | 421 |
| -1,44287E | FALSE | High | IsMasterP | AAEL0010 | 3:552083! | 0 | 45 | FALSE | 20 | 49 | 20 | 524 | 585 | 681 |
| -1,88041E | FALSE | High | IsMasterP | AAEL0134 | 1:206424 | 0 | 54 | FALSE | 18 | 51 | 18 | 384 | 432 | 563 |
| -6,2658E+ | FALSE | High | IsMasterP | AAEL0148 | 2:121825 | 0 | 63 | FALSE | 20 | 53 | 20 | 495 | 551 | 878 |
| -4,41772E | FALSE | High | IsMasterP | AAEL0261 | 2:147203 | 0 | 49 | FALSE | 14 | 47 | 10 | 448 | 498 | 559 |
| 7,8133E+ | FALSE | High | IsMasterP | AAEL0253 | 1:177629 | 0 | 49 | FALSE | 23 | 55 | 23 | 679 | 774 | 587 |
| 5,33409E- | FALSE | High | IsMasterP | AAEL0079 | 2:352151 | 0 | 66 | FALSE | 16 | 49 | 16 | 336 | 385 | 610 |
| -9,10547E | FALSE | High | IsMasterP | AAEL0105 | 2:910612 | 0 | 11 | FALSE | 21 | 57 | 21 | 3384 | 3825 | 609 |
| -8,09753E | FALSE | High | IsMasterP | AAEL0047 | 2:379310 | 0 | 68 | FALSE | 12 | 43 | 12 | 311 | 346 | 613 |
| 1,55726E- | FALSE | High | IsMasterP | AAEL0256 | 1:551942 | 0 | 44 | FALSE | 23 | 57 | 18 | 635 | 695 | 599 |
| -2,19513E | FALSE | High | IsMasterP | AAEL0230 | 2:147229 | 0 | 60 | FALSE | 19 | 53 | 19 | 392 | 432 | 684 |
| 8,67112E- | FALSE | High | IsMasterP | AAEL0271 | 1:307820! | 0 | 46 | FALSE | 18 | 50 | 18 | 483 | 540 | 695 |
| -8,87462E | FALSE | High | IsMasterP | AAEL0121 | 2:405952 | 0 | 48 | FALSE | 20 | 56 | 20 | 493 | 563 | 793 |
| 6,54424E- | FALSE | High | IsMasterP | AAEL0049 | 2:281882 | 0 | 25 | FALSE | 21 | 54 | 21 | 1234 | 1386 | 613 |
| 8,49564E- | FALSE | High | IsMasterP | AAEL0027 | 2:144334( | 0 | 53 | FALSE | 15 | 49 | 15 | 387 | 438 | 550 |
| -4,34494E | FALSE | High | IsMasterP | AAEL0245 | 2:127259 | 0 | 17 | FALSE | 15 | 37 | 15 | 1965 | 2277 | 454 |
| -1,00736E | FALSE | High | IsMasterP | AAEL0020 | 1:582764 | 0 | 54 | FALSE | 23 | 54 | 23 | 514 | 580 | 642 |
| 8,68192E- | FALSE | High | IsMasterP | AAEL0205 | 1:273037( | 0 | 11 | FALSE | 27 | 56 | 27 | 3408 | 3899 | 482 |
| -2,47436E | FALSE | High | IsMasterP | AAEL0133 | 2:135754! | 0 | 74 | FALSE | 10 | 38 | 8 | 197 | 219 | 621 |
| 7,14471E- | FALSE | High | IsMasterP | AAEL0065 | 2:133263 | 0 | 55 | FALSE | 10 | 47 | 10 | 153 | 173 | 407 |

| | | | | | | | | | | | | | | |
|---|---|---|---|---|---|---|---|---|---|---|---|---|---|---|
| 3,76841E- | FALSE | High | IsMasterP | AAEL0007 | 3:246025 | 0 | 47 | FALSE | 16 | 54 | 16 | 370 | 420 | 803 |
| 5,53927E- | FALSE | High | IsMasterP | AAEL0099 | 1:307557 | 0 | 31 | FALSE | 20 | 51 | 20 | 1194 | 1291 | 623 |
| 3,47457E- | FALSE | High | IsMasterP | AAEL0129 | 3:206760 | 0 | 53 | FALSE | 19 | 55 | 19 | 407 | 471 | 586 |
| 7,42904E- | FALSE | High | IsMasterP | AAEL0048 | 2:115621 | 0 | 53 | FALSE | 18 | 49 | 18 | 517 | 568 | 516 |
| 2,74138E- | FALSE | High | IsMasterP | AAEL0092 | 3:200124 | 0 | 43 | FALSE | 14 | 45 | 14 | 512 | 559 | 785 |
| -5,73439E | FALSE | High | IsMasterP | AAEL0233 | 1:280191 | 0 | 29 | FALSE | 14 | 58 | 6 | 636 | 700 | 587 |
| 4,48534E- | FALSE | High | IsMasterP | AAEL0057 | 1:121872 | 0 | 40 | FALSE | 16 | 54 | 16 | 660 | 722 | 660 |
| -3,99024E | FALSE | High | IsMasterP | AAEL0003 | 2:367825 | 0 | 35 | FALSE | 19 | 56 | 19 | 560 | 636 | 577 |
| 7,62349E- | FALSE | High | IsMasterP | AAEL0001 | 3:213008 | 0 | 43 | FALSE | 20 | 50 | 20 | 671 | 756 | 711 |
| 7,10981E- | FALSE | High | IsMasterP | AAEL0045 | NIGP0100 | 0 | 84 | FALSE | 10 | 58 | 10 | 83 | 93 | 721 |
| -7,08385E | FALSE | High | IsMasterP | AAEL0147 | 1:456977 | 0 | 40 | FALSE | 18 | 45 | 18 | 619 | 696 | 692 |
| 1,59078E- | FALSE | High | IsMasterP | AAEL0079 | 2:351623 | 0 | 75 | FALSE | 12 | 40 | 12 | 224 | 250 | 711 |
| -7,16507E | FALSE | High | IsMasterP | AAEL0143 | 2:120127 | 0 | 46 | FALSE | 14 | 45 | 14 | 459 | 530 | 564 |
| 2,7304E+1 | FALSE | High | IsMasterP | AAEL0042 | 3:292663 | 0 | 42 | FALSE | 17 | 49 | 17 | 527 | 591 | 660 |
| 2,64064E- | FALSE | High | IsMasterP | AAEL0174 | 1:159466 | 0 | 33 | FALSE | 14 | 52 | 14 | 502 | 554 | 900 |
| 9,2611E+1 | FALSE | High | IsMasterP | AAEL0113 | 1:122104 | 0 | 42 | FALSE | 14 | 47 | 14 | 468 | 528 | 684 |
| -3,91709E | FALSE | High | IsMasterP | AAEL0132 | 3:279908 | 0 | 34 | FALSE | 22 | 50 | 21 | 898 | 1023 | 941 |
| 8,29796E- | FALSE | High | IsMasterP | AAEL0098 | 1:923288 | 0 | 58 | FALSE | 17 | 46 | 17 | 544 | 595 | 828 |
| 3,72071E- | FALSE | High | IsMasterP | AAEL0077 | 2:100865 | 0 | 52 | FALSE | 14 | 42 | 14 | 522 | 583 | 545 |
| -5,28957E | FALSE | High | IsMasterP | AAEL0025 | 1:904713 | 0 | 68 | FALSE | 7 | 41 | 7 | 161 | 170 | 531 |
| 8,6414E+1 | FALSE | High | IsMasterP | AAEL0104 | 1:268145 | 0 | 50 | FALSE | 20 | 55 | 20 | 536 | 613 | 651 |
| -8,75493E | FALSE | High | IsMasterP | AAEL0120 | 1:242496 | 0 | 44 | FALSE | 16 | 47 | 15 | 438 | 491 | 658 |
| 8,7941E+1 | FALSE | High | IsMasterP | AAEL0009 | 3:377081 | 0 | 60 | FALSE | 7 | 45 | 1 | 194 | 222 | 491 |
| 2,26178E- | FALSE | High | IsMasterP | AAEL0041 | 3:403503 | 0 | 56 | FALSE | 8 | 46 | 8 | 157 | 176 | 810 |
| -2,3902E+ | FALSE | High | IsMasterP | AAEL0105 | 3:310086 | 0 | 58 | FALSE | 15 | 49 | 15 | 322 | 350 | 929 |
| -2,46428E | FALSE | High | IsMasterP | AAEL0025 | 1:893401 | 0 | 52 | FALSE | 11 | 53 | 11 | 219 | 254 | 1001 |
| -1,92534E | FALSE | High | IsMasterP | AAEL0077 | 1:712583 | 0 | 75 | FALSE | 11 | 42 | 11 | 182 | 205 | 885 |
| 9,21875E- | FALSE | High | IsMasterP | AAEL0261 | 1:292364 | 0 | 50 | FALSE | 12 | 44 | 11 | 331 | 365 | 526 |
| -8,9604E+ | FALSE | High | IsMasterP | AAEL0124 | 1:123297 | 0 | 41 | FALSE | 14 | 51 | 14 | 509 | 570 | 671 |
| 7,1587E+1 | FALSE | High | IsMasterP | AAEL0033 | 3:804319 | 0 | 44 | FALSE | 21 | 53 | 21 | 574 | 658 | 923 |
| -4,02769E | FALSE | High | IsMasterP | AAEL0049 | 2:259726 | 0 | 73 | FALSE | 16 | 44 | 16 | 276 | 313 | 614 |

| | | | | | | | | | | | | | | |
|---|---|---|---|---|---|---|---|---|---|---|---|---|---|---|
| -1,18736E | FALSE | High | IsMasterP | AAEL0097 | 3:284460 | 0 | 47 | FALSE | 12 | 58 | 12 | 152 | 175 | 1045 |
| -3,22333E | FALSE | High | IsMasterP | AAEL0138 | 1:140471 | 0 | 51 | FALSE | 19 | 51 | 19 | 511 | 575 | 690 |
| -8,93177E | FALSE | High | IsMasterP | AAEL0036 | 2:316596 | 0 | 58 | FALSE | 11 | 45 | 11 | 266 | 282 | 844 |
| -1,85697E | FALSE | High | IsMasterP | AAEL0026 | 3:256575 | 0 | 49 | FALSE | 14 | 46 | 14 | 475 | 500 | 529 |
| 7,76966E | FALSE | High | IsMasterP | AAEL0126 | 3:239371 | 0 | 33 | FALSE | 6 | 44 | 6 | 143 | 161 | 1058 |
| 2,64181E | FALSE | High | IsMasterP | AAEL0153 | 2:112306 | 0 | 57 | FALSE | 17 | 50 | 17 | 378 | 430 | 524 |
| 5,19136E | FALSE | High | IsMasterP | AAEL0232 | 1:147421 | 0 | 23 | FALSE | 16 | 45 | 16 | 1295 | 1465 | 599 |
| -4,41053E | FALSE | High | IsMasterP | AAEL0069 | 2:340721 | 0 | 65 | FALSE | 9 | 37 | 9 | 212 | 238 | 522 |
| -8,22927E | FALSE | High | IsMasterP | AAEL0230 | 3:109880 | 0 | 87 | FALSE | 10 | 39 | 10 | 113 | 124 | 527 |
| -7,35916E | FALSE | High | IsMasterP | AAEL0070 | 3:299807 | 0 | 24 | FALSE | 14 | 51 | 14 | 715 | 747 | 758 |
| 5,23371E | FALSE | High | IsMasterP | AAEL0222 | 3:121415 | 0 | 84 | FALSE | 12 | 45 | 10 | 194 | 216 | 493 |
| -3,92124E | FALSE | High | IsMasterP | AAEL0090 | 2:160646 | 0 | 67 | FALSE | 12 | 44 | 12 | 218 | 246 | 600 |
| -8,09193E | FALSE | High | IsMasterP | AAEL0118 | 2:187278 | 0 | 54 | FALSE | 10 | 43 | 10 | 306 | 332 | 846 |
| 2,8245E+ | FALSE | High | IsMasterP | AAEL0096 | 1:606353 | 0 | 34 | FALSE | 15 | 46 | 15 | 582 | 667 | 629 |
| 7,4528E+ | FALSE | High | IsMasterP | AAEL0061 | 3:142151 | 0 | 21 | FALSE | 15 | 42 | 15 | 1130 | 1251 | 828 |
| 6,82608E | FALSE | High | IsMasterP | AAEL0014 | 2:482770 | 0 | 48 | FALSE | 17 | 54 | 17 | 431 | 491 | 667 |
| -6,22015E | FALSE | High | IsMasterP | AAEL0077 | 3:151720 | 0 | 49 | FALSE | 16 | 41 | 16 | 519 | 554 | 680 |
| -7,92253E | FALSE | High | IsMasterP | AAEL0114 | 3:142717 | 0 | 48 | FALSE | 10 | 40 | 10 | 128 | 147 | 948 |
| -8,45857E | FALSE | High | IsMasterP | AAEL0043 | 3:368069 | 0 | 50 | FALSE | 14 | 51 | 14 | 386 | 442 | 618 |
| -9,10253E | FALSE | High | IsMasterP | AAEL0065 | 2:132675 | 0 | 27 | FALSE | 18 | 50 | 18 | 999 | 1096 | 550 |
| 4,23153E | FALSE | High | IsMasterP | AAEL0276 | 3:752455 | 0 | 12 | FALSE | 13 | 38 | 13 | 2049 | 2211 | 446 |
| 8,83417E | FALSE | High | IsMasterP | AAEL0079 | 1:636146 | 0 | 40 | FALSE | 17 | 48 | 17 | 593 | 701 | 620 |
| -4,17169E | FALSE | High | IsMasterP | AAEL0036 | 2:317115 | 0 | 40 | FALSE | 20 | 53 | 20 | 625 | 702 | 563 |
| -7,00615E | FALSE | High | IsMasterP | AAEL0020 | 2:119972 | 0 | 39 | FALSE | 21 | 52 | 16 | 699 | 754 | 954 |
| -4,00604E | FALSE | High | IsMasterP | AAEL0054 | 1:247250 | 0 | 52 | FALSE | 12 | 40 | 12 | 391 | 442 | 885 |
| 6,79643E | FALSE | High | IsMasterP | AAEL0231 | 2:259455 | 0 | 46 | FALSE | 19 | 47 | 19 | 625 | 681 | 914 |
| -2,75635E | FALSE | High | IsMasterP | AAEL0007 | 2:553584 | 0 | 63 | FALSE | 10 | 46 | 10 | 150 | 171 | 413 |
| -8,06318E | FALSE | High | IsMasterP | AAEL0244 | 3:203214 | 0 | 25 | FALSE | 10 | 56 | 10 | 236 | 271 | 1092 |
| 1,12175E | FALSE | High | IsMasterP | AAEL0274 | 2:187356 | 0 | 24 | FALSE | 18 | 48 | 18 | 1116 | 1274 | 625 |
| 8,01259E | FALSE | High | IsMasterP | AAEL0125 | 2:327970 | 0 | 38 | FALSE | 13 | 54 | 13 | 263 | 303 | 1078 |
| -6,05829E | FALSE | High | IsMasterP | AAEL0264 | 2:155129 | 0 | 43 | FALSE | 11 | 40 | 11 | 390 | 428 | 498 |

| | | | | | | | | | | | | | | |
|---|---|---|---|---|---|---|---|---|---|---|---|---|---|---|
| 2,81992E- | FALSE | High | IsMasterP | AAEL0114 | 3:142679 | 0 | 41 | FALSE | 10 | 46 | 10 | 186 | 215 | 1029 |
| 8,85065E- | FALSE | High | IsMasterP | AAEL0040 | 3:166253 | 0 | 43 | FALSE | 13 | 44 | 13 | 386 | 426 | 550 |
| -4,70468E | FALSE | High | IsMasterP | AAEL0199 | 3:173036 | 0 | 31 | FALSE | 20 | 47 | 20 | 890 | 1007 | 553 |
| 5,56769E- | FALSE | High | IsMasterP | AAEL0061 | 3:307416 | 0 | 56 | FALSE | 17 | 46 | 17 | 389 | 430 | 904 |
| -2,37133E | FALSE | High | IsMasterP | AAEL0224 | 2:688567 | 0 | 52 | FALSE | 16 | 44 | 16 | 385 | 442 | 774 |
| 1,29946E- | FALSE | High | IsMasterP | AAEL0036 | 1:691896 | 0 | 36 | FALSE | 12 | 47 | 12 | 427 | 472 | 799 |
| -2,46609E | FALSE | High | IsMasterP | AAEL0068 | 1:193953 | 0 | 78 | FALSE | 8 | 48 | 8 | 65 | 74 | 1037 |
| 2,83088E- | FALSE | High | IsMasterP | AAEL0079 | 2:351708 | 0 | 73 | FALSE | 14 | 45 | 14 | 222 | 248 | 654 |
| 3,99126E- | FALSE | High | IsMasterP | AAEL0040 | 3:166372 | 0 | 32 | FALSE | 11 | 44 | 10 | 629 | 702 | 548 |
| -7,13796E | FALSE | High | IsMasterP | AAEL0068 | 2:251134 | 0 | 30 | FALSE | 13 | 45 | 13 | 498 | 561 | 887 |
| -3,76381E | FALSE | High | IsMasterP | AAEL0139 | 2:101868 | 0 | 59 | FALSE | 13 | 45 | 2 | 340 | 372 | 651 |
| -1,86302E | FALSE | High | IsMasterP | AAEL0090 | 1:292356 | 0 | 28 | FALSE | 19 | 40 | 19 | 1578 | 1694 | 952 |
| -2,01156E | FALSE | High | IsMasterP | AAEL0009 | 3:380290 | 0 | 13 | FALSE | 17 | 42 | 17 | 2132 | 2426 | 661 |
| -3,31407E | FALSE | High | IsMasterP | AAEL0259 | 1:573018 | 0 | 58 | FALSE | 14 | 41 | 14 | 308 | 346 | 618 |
| 6,77807E- | FALSE | High | IsMasterP | AAEL0080 | 3:132076 | 0 | 50 | FALSE | 17 | 48 | 17 | 455 | 522 | 553 |
| 4,728E+1 | FALSE | High | IsMasterP | AAEL0038 | 3:846252 | 0 | 47 | FALSE | 11 | 52 | 11 | 253 | 277 | 501 |
| 1,47771E- | FALSE | High | IsMasterP | AAEL0016 | 2:299037 | 0 | 49 | FALSE | 14 | 44 | 14 | 417 | 475 | 733 |
| 7,28786E- | FALSE | High | IsMasterP | AAEL0024 | 1:177368 | 0 | 27 | FALSE | 21 | 48 | 21 | 1138 | 1262 | 525 |
| -5,6894E+ | FALSE | High | IsMasterP | AAEL0057 | 2:436115 | 0 | 20 | FALSE | 13 | 42 | 9 | 1004 | 1152 | 658 |
| -9,98442E | FALSE | High | IsMasterP | AAEL0123 | 2:102859 | 0 | 42 | FALSE | 13 | 42 | 13 | 450 | 497 | 515 |
| 8,91664E- | FALSE | High | IsMasterP | AAEL0040 | 1:137417 | 0 | 42 | FALSE | 7 | 44 | 7 | 186 | 198 | 730 |
| -7,17729E | FALSE | High | IsMasterP | AAEL0120 | 3:160802 | 0 | 70 | FALSE | 14 | 50 | 14 | 198 | 220 | 771 |
| 4,45922E- | FALSE | High | IsMasterP | AAEL0058 | 2:661703 | 0 | 33 | FALSE | 6 | 34 | 6 | 195 | 216 | 620 |
| -7,80405E | FALSE | High | IsMasterP | AAEL0201 | 3:315233 | 0 | 52 | FALSE | 14 | 41 | 14 | 381 | 422 | 508 |
| -4,19829E | FALSE | High | IsMasterP | AAEL0273 | 1:644323 | 0 | 57 | FALSE | 11 | 49 | 11 | 156 | 181 | 585 |
| -5,54124E | FALSE | High | IsMasterP | AAEL0086 | 2:245746 | 0 | 70 | FALSE | 8 | 41 | 8 | 90 | 100 | 517 |
| 3,74429E- | FALSE | High | IsMasterP | AAEL0006 | 3:393004 | 0 | 21 | FALSE | 11 | 46 | 11 | 808 | 877 | 446 |
| -8,52042E | FALSE | High | IsMasterP | AAEL0050 | 1:295774 | 0 | 49 | FALSE | 10 | 43 | 9 | 206 | 228 | 530 |
| -8,26569E | FALSE | High | IsMasterP | AAEL0101 | 2:265095 | 0 | 54 | FALSE | 14 | 38 | 13 | 326 | 377 | 646 |
| -1,83354E | FALSE | High | IsMasterP | AAEL0077 | 2:219402 | 0 | 46 | FALSE | 13 | 46 | 13 | 408 | 463 | 582 |
| 5,96826E- | FALSE | High | IsMasterP | AAEL0261 | 1:162003 | 0 | 80 | FALSE | 12 | 39 | 12 | 183 | 209 | 829 |

| | | | | | | | | | | | | | | |
|---|---|---|---|---|---|---|---|---|---|---|---|---|---|---|
| -5,70665E | FALSE | High | IsMasterP | AAEL0130 | 2:253390 | 0 | 62 | FALSE | 16 | 44 | 14 | 309 | 355 | 554 |
| -7,07832E | FALSE | High | IsMasterP | AAEL0175 | 2:587803 | 0 | 12 | FALSE | 11 | 46 | 11 | 386 | 400 | 1064 |
| 5,84255E | FALSE | High | IsMasterP | AAEL0277 | 3:697753 | 0 | 53 | FALSE | 8 | 36 | 8 | 217 | 247 | 592 |
| 6,82006E | FALSE | High | IsMasterP | AAEL0084 | NIGP0100 | 0 | 44 | FALSE | 18 | 47 | 18 | 433 | 490 | 644 |
| -1,45484E | FALSE | High | IsMasterP | AAEL0142 | 3:301554 | 0 | 69 | FALSE | 15 | 45 | 15 | 251 | 276 | 530 |
| 3,04958E | FALSE | High | IsMasterP | AAEL0033 | 3:815137 | 0 | 60 | FALSE | 15 | 40 | 15 | 376 | 427 | 706 |
| 5,78639E | FALSE | High | IsMasterP | AAEL0031 | 1:192133 | 0 | 53 | FALSE | 12 | 37 | 12 | 376 | 417 | 654 |
| -1,4071E+ | FALSE | High | IsMasterP | AAEL0091 | 3:168896 | 0 | 29 | FALSE | 17 | 47 | 17 | 881 | 971 | 563 |
| 2,00958E | FALSE | High | IsMasterP | AAEL0114 | 2:143662 | 0 | 80 | FALSE | 8 | 35 | 8 | 127 | 139 | 718 |
| -2,01316E | FALSE | High | IsMasterP | AAEL0104 | 1:204065 | 0 | 59 | FALSE | 7 | 36 | 7 | 170 | 192 | 473 |
| 6,08465E | FALSE | High | IsMasterP | AAEL0015 | 3:263409 | 0 | 22 | FALSE | 15 | 42 | 15 | 1337 | 1464 | 634 |
| -9,16356E | FALSE | High | IsMasterP | AAEL0108 | 2:127676 | 0 | 60 | FALSE | 15 | 42 | 15 | 265 | 303 | 580 |
| 8,55211E | FALSE | High | IsMasterP | AAEL0197 | 2:690845 | 0 | 51 | FALSE | 15 | 41 | 15 | 520 | 560 | 670 |
| -2,08366E | FALSE | High | IsMasterP | AAEL0094 | 3:338025 | 0 | 54 | FALSE | 13 | 40 | 13 | 401 | 460 | 892 |
| 8,97646E | FALSE | High | IsMasterP | AAEL0147 | 2:324343 | 0 | 52 | FALSE | 13 | 37 | 13 | 421 | 466 | 815 |
| -3,10117E | FALSE | High | IsMasterP | AAEL0116 | 3:281619 | 0 | 37 | FALSE | 17 | 41 | 17 | 775 | 855 | 511 |
| 4,42934E | FALSE | High | IsMasterP | AAEL0105 | 2:910503 | 0 | 58 | FALSE | 10 | 37 | 10 | 262 | 287 | 587 |
| -8,88013E | FALSE | High | IsMasterP | AAEL0060 | 3:743281 | 0 | 32 | FALSE | 14 | 45 | 14 | 636 | 718 | 684 |
| 1,83893E | FALSE | High | IsMasterP | AAEL0221 | NIGP0100 | 0 | 40 | FALSE | 9 | 53 | 1 | 124 | 137 | 1029 |
| 5,14188E | FALSE | High | IsMasterP | AAEL0098 | 1:207255 | 0 | 80 | FALSE | 16 | 46 | 16 | 246 | 277 | 724 |
| -8,79155E | FALSE | High | IsMasterP | AAEL0028 | 1:304883 | 0 | 34 | FALSE | 17 | 45 | 17 | 734 | 824 | 742 |
| -6,66304E | FALSE | High | IsMasterP | AAEL0035 | 3:263598 | 0 | 35 | FALSE | 13 | 36 | 13 | 527 | 584 | 920 |
| 8,33398E | FALSE | High | IsMasterP | AAEL0098 | 3:121945 | 0 | 42 | FALSE | 18 | 47 | 18 | 594 | 672 | 881 |
| 9,08039E | FALSE | High | IsMasterP | AAEL0092 | 2:170797 | 0 | 48 | FALSE | 6 | 31 | 6 | 159 | 173 | 848 |
| -6,28694E | FALSE | High | IsMasterP | AAEL0066 | 2:326276 | 0 | 49 | FALSE | 15 | 41 | 8 | 552 | 601 | 766 |
| 3,36231E | FALSE | High | IsMasterP | AAEL0003 | 2:124233 | 0 | 59 | FALSE | 11 | 44 | 11 | 175 | 192 | 911 |
| -6,56019E | FALSE | High | IsMasterP | AAEL0093 | 3:276095 | 0 | 35 | FALSE | 13 | 41 | 13 | 462 | 516 | 596 |
| 3,75611E | FALSE | High | IsMasterP | AAEL0038 | 3:342753 | 0 | 26 | FALSE | 19 | 40 | 19 | 986 | 1140 | 810 |
| 1,12841E | FALSE | High | IsMasterP | AAEL0173 | 3:347295 | 0 | 50 | FALSE | 10 | 46 | 5 | 245 | 267 | 680 |
| 1,34709E | FALSE | High | IsMasterP | AAEL0070 | 3:170641 | 0 | 32 | FALSE | 18 | 44 | 18 | 886 | 1006 | 535 |
| -1,81714E | FALSE | High | IsMasterP | AAEL0195 | 1:305071 | 0 | 19 | FALSE | 12 | 41 | 12 | 704 | 822 | 555 |

| | | | | | | | | | | | | | | |
|---|---|---|---|---|---|---|---|---|---|---|---|---|---|---|
| -2,06381E | FALSE | High | IsMasterP | AAEL0125 | 2:185971 | 0 | 67 | FALSE | 8 | 36 | 8 | 162 | 179 | 728 |
| -6,07177E | FALSE | High | IsMasterP | AAEL0050 | 3:828204 | 0 | 37 | FALSE | 12 | 48 | 12 | 217 | 251 | 1122 |
| -7,8316E+ | FALSE | High | IsMasterP | AAEL0070 | 3:298555 | 0 | 63 | FALSE | 11 | 33 | 9 | 182 | 207 | 658 |
| -1,78512E | FALSE | High | IsMasterP | AAEL0009 | 2:104487 | 0 | 37 | FALSE | 19 | 43 | 19 | 717 | 807 | 698 |
| -6,74978E | FALSE | High | IsMasterP | AAEL0100 | 2:110797 | 0 | 45 | FALSE | 16 | 38 | 16 | 561 | 608 | 624 |
| 5,19752E- | FALSE | High | IsMasterP | AAEL0148 | 2:104559 | 0 | 54 | FALSE | 9 | 39 | 9 | 175 | 199 | 687 |
| -2,71263E | FALSE | High | IsMasterP | AAEL0108 | 1:546838 | 0 | 23 | FALSE | 7 | 42 | 7 | 207 | 244 | 973 |
| 1,18654E- | FALSE | High | IsMasterP | AAEL0195 | 3:655599 | 0 | 18 | FALSE | 16 | 36 | 16 | 1670 | 1796 | 454 |
| -8,18079E | FALSE | High | IsMasterP | AAEL0250 | 2:168277 | 0 | 42 | FALSE | 14 | 41 | 14 | 429 | 474 | 613 |
| -6,49937E | FALSE | High | IsMasterP | AAEL0065 | 2:593637 | 0 | 66 | FALSE | 12 | 43 | 12 | 128 | 145 | 506 |
| 8,56997E- | FALSE | High | IsMasterP | AAEL0089 | 1:652309 | 0 | 46 | FALSE | 9 | 37 | 9 | 208 | 237 | 496 |
| 7,32389E- | FALSE | High | IsMasterP | AAEL0105 | 1:508738 | 0 | 21 | FALSE | 17 | 45 | 17 | 1181 | 1334 | 601 |
| -8,21204E | FALSE | High | IsMasterP | AAEL0181 | 3:278478 | 0 | 17 | FALSE | 15 | 35 | 15 | 1794 | 1889 | 802 |
| 8,71315E- | FALSE | High | IsMasterP | AAEL0119 | 2:209119 | 0 | 52 | FALSE | 13 | 38 | 13 | 415 | 459 | 635 |
| 9,17061E- | FALSE | High | IsMasterP | AAEL0034 | 1:229156 | 0 | 38 | FALSE | 7 | 41 | 7 | 258 | 290 | 781 |
| 9,58106E- | FALSE | High | IsMasterP | AAEL0104 | 2:446422 | 0 | 50 | FALSE | 13 | 33 | 13 | 512 | 572 | 536 |
| 4,70092E- | FALSE | High | IsMasterP | AAEL0217 | 1:292266 | 0 | 54 | FALSE | 12 | 37 | 12 | 246 | 274 | 689 |
| 4,70486E- | FALSE | High | IsMasterP | AAEL0087 | 3:281090 | 0 | 32 | FALSE | 19 | 41 | 14 | 911 | 988 | 928 |
| -3,40225E | FALSE | High | IsMasterP | AAEL0111 | 2:182351 | 0 | 73 | FALSE | 10 | 35 | 10 | 290 | 309 | 702 |
| 8,04134E- | FALSE | High | IsMasterP | AAEL0000 | 3:104473 | 0 | 49 | FALSE | 12 | 39 | 12 | 332 | 370 | 576 |
| 4,61146E- | FALSE | High | IsMasterP | AAEL0254 | 2:200020 | 0 | 51 | FALSE | 15 | 34 | 15 | 452 | 503 | 887 |
| 4,13063E- | FALSE | High | IsMasterP | AAEL0087 | 3:951651 | 0 | 42 | FALSE | 8 | 36 | 8 | 285 | 315 | 515 |
| 8,45817E- | FALSE | High | IsMasterP | AAEL0101 | 2:264791 | 0 | 50 | FALSE | 12 | 39 | 12 | 370 | 405 | 847 |
| -6,7879E+ | FALSE | High | IsMasterP | AAEL0076 | 1:128739 | 0 | 21 | FALSE | 21 | 35 | 21 | 1847 | 2051 | 538 |
| -7,94876E | FALSE | High | IsMasterP | AAEL0058 | 2:661541 | 0 | 55 | FALSE | 8 | 37 | 8 | 207 | 232 | 736 |
| -4,77301E | FALSE | High | IsMasterP | AAEL0061 | 3:141707 | 0 | 33 | FALSE | 7 | 39 | 7 | 205 | 233 | 595 |
| -7,19928E | FALSE | High | IsMasterP | AAEL0090 | 3:461032 | 0 | 54 | FALSE | 10 | 44 | 10 | 301 | 339 | 576 |
| -5,36386E | FALSE | High | IsMasterP | AAEL0023 | 2:302273 | 0 | 39 | FALSE | 7 | 40 | 7 | 152 | 176 | 1083 |
| 4,99578E- | FALSE | High | IsMasterP | AAEL0139 | 2:305831 | 0 | 53 | FALSE | 21 | 41 | 18 | 541 | 609 | 711 |
| 1,62673E- | FALSE | High | IsMasterP | AAEL0117 | 3:290727 | 0 | 30 | FALSE | 8 | 38 | 8 | 438 | 491 | 609 |
| 3,56796E- | FALSE | High | IsMasterP | AAEL0022 | 2:300314 | 0 | 33 | FALSE | 12 | 41 | 12 | 334 | 367 | 624 |

| | | | | | | | | | | | | | | |
|---|---|---|---|---|---|---|---|---|---|---|---|---|---|---|
| 8,9901E+1 | FALSE | High | IsMasterP | AAEL0018 | 2:274383 | 0 | 50 | FALSE | 15 | 39 | 15 | 409 | 467 | 577 |
| 8,41258E- | FALSE | High | IsMasterP | AAEL0091 | 1:112241 | 0 | 38 | FALSE | 6 | 39 | 6 | 130 | 148 | 999 |
| 3,95967E- | FALSE | High | IsMasterP | AAEL0048 | 2:115712 | 0 | 45 | FALSE | 7 | 30 | 7 | 157 | 181 | 979 |
| 3,68573E- | FALSE | High | IsMasterP | AAEL0127 | 2:162344 | 0 | 44 | FALSE | 9 | 38 | 9 | 241 | 263 | 703 |
| -5,89801E | FALSE | High | IsMasterP | AAEL0209 | NIGP0100 | 0 | 60 | FALSE | 8 | 35 | 8 | 184 | 211 | 736 |
| 1,30337E- | FALSE | High | IsMasterP | AAEL0124 | 3:242489 | 0 | 63 | FALSE | 5 | 29 | 5 | 137 | 145 | 1010 |
| -2,45104E | FALSE | High | IsMasterP | AAEL0052 | 2:692660 | 0 | 84 | FALSE | 5 | 32 | 5 | 144 | 153 | 502 |
| 1,96629E- | FALSE | High | IsMasterP | AAEL0140 | 2:405938 | 0 | 74 | FALSE | 9 | 36 | 9 | 133 | 154 | 639 |
| 2,86021E- | FALSE | High | IsMasterP | AAEL0020 | 2:426569 | 0 | 34 | FALSE | 14 | 37 | 14 | 631 | 692 | 657 |
| -3,91459E | FALSE | High | IsMasterP | AAEL0139 | 3:106460 | 0 | 34 | FALSE | 13 | 39 | 13 | 752 | 859 | 555 |
| 5,71836E- | FALSE | High | IsMasterP | AAEL0091 | 1:112344 | 0 | 67 | FALSE | 8 | 32 | 8 | 162 | 184 | 455 |
| 8,06391E- | FALSE | High | IsMasterP | AAEL0017 | 2:166851 | 0 | 41 | FALSE | 20 | 45 | 17 | 718 | 787 | 967 |
| 7,9134E+1 | FALSE | High | IsMasterP | AAEL0123 | 1:284412 | 0 | 59 | FALSE | 12 | 35 | 12 | 270 | 302 | 624 |
| -5,97085E | FALSE | High | IsMasterP | AAEL0032 | 1:190201 | 0 | 55 | FALSE | 11 | 36 | 11 | 217 | 234 | 505 |
| -6,92361E | FALSE | High | IsMasterP | AAEL0154 | 2:197899 | 0 | 60 | FALSE | 7 | 35 | 7 | 137 | 144 | 724 |
| -4,90424E | FALSE | High | IsMasterP | AAEL0100 | 1:159363 | 0 | 34 | FALSE | 14 | 44 | 14 | 389 | 454 | 648 |
| 8,43261E- | FALSE | High | IsMasterP | AAEL0194 | 3:506440 | 0 | 33 | FALSE | 14 | 43 | 14 | 509 | 572 | 596 |
| 5,54339E- | FALSE | High | IsMasterP | AAEL0023 | 3:375997 | 0 | 43 | FALSE | 14 | 33 | 14 | 394 | 439 | 550 |
| -2,0225E+ | FALSE | High | IsMasterP | AAEL0042 | 2:995423 | 0 | 69 | FALSE | 14 | 40 | 14 | 234 | 266 | 625 |
| -8,29543E | FALSE | High | IsMasterP | AAEL0054 | 1:246219 | 0 | 32 | FALSE | 9 | 35 | 9 | 307 | 345 | 554 |
| -5,16175E | FALSE | High | IsMasterP | AAEL0028 | 1:304488 | 0 | 57 | FALSE | 9 | 37 | 9 | 106 | 116 | 948 |
| -3,76309E | FALSE | High | IsMasterP | AAEL0290 | 2:126273 | 0 | 78 | FALSE | 10 | 39 | 10 | 86 | 100 | 548 |
| 5,7252E+1 | FALSE | High | IsMasterP | AAEL0108 | 3:156709 | 0 | 45 | FALSE | 3 | 25 | 3 | 86 | 100 | 581 |
| -2,47888E | FALSE | High | IsMasterP | AAEL0148 | 2:155770 | 0 | 33 | FALSE | 7 | 31 | 3 | 321 | 364 | 623 |
| 3,46735E- | FALSE | High | IsMasterP | AAEL0125 | 1:285506 | 0 | 49 | FALSE | 15 | 43 | 15 | 408 | 453 | 742 |
| -6,26251E | FALSE | High | IsMasterP | AAEL0004 | 3:183539 | 0 | 10 | FALSE | 18 | 31 | 18 | 4290 | 4625 | 597 |
| 5,5476E+1 | FALSE | High | IsMasterP | AAEL0058 | 3:174875 | 0 | 58 | FALSE | 9 | 35 | 9 | 233 | 252 | 890 |
| -8,43651E | FALSE | High | IsMasterP | AAEL0042 | 1:105224 | 0 | 25 | FALSE | 14 | 32 | 14 | 841 | 931 | 850 |
| -6,57007E | FALSE | High | IsMasterP | AAEL0014 | 2:482906 | 0 | 28 | FALSE | 11 | 45 | 11 | 424 | 480 | 586 |
| 5,63868E- | FALSE | High | IsMasterP | AAEL0183 | 3:109964 | 0 | 37 | FALSE | 21 | 37 | 21 | 830 | 929 | 652 |
| -8,93645E | FALSE | High | IsMasterP | AAEL0090 | 2:396291 | 0 | 19 | FALSE | 13 | 33 | 13 | 1054 | 1204 | 481 |

| | | | | | | | | | | | | | | |
|---|---|---|---|---|---|---|---|---|---|---|---|---|---|---|
| -7,70874E | FALSE | High | IsMasterP | AAEL0104 | 2:446378 | 0 | 42 | FALSE | 13 | 42 | 13 | 450 | 509 | 527 |
| 7,34658E | FALSE | High | IsMasterP | AAEL0104 | 1:204531 | 0 | 33 | FALSE | 11 | 37 | 11 | 368 | 398 | 870 |
| 1,0433E+ | FALSE | High | IsMasterP | AAEL0198 | 2:367568 | 0 | 20 | FALSE | 14 | 37 | 14 | 1105 | 1275 | 571 |
| 4,56155E | FALSE | High | IsMasterP | AAEL0005 | 3:388552 | 0 | 33 | FALSE | 8 | 33 | 8 | 351 | 375 | 768 |
| 8,5597E+ | FALSE | High | IsMasterP | AAEL0115 | 3:313163 | 0 | 19 | FALSE | 10 | 36 | 10 | 884 | 990 | 578 |
| 7,82477E | FALSE | High | IsMasterP | AAEL0052 | 3:309626 | 0 | 60 | FALSE | 18 | 39 | 18 | 475 | 521 | 784 |
| 5,55419E | FALSE | High | IsMasterP | AAEL0046 | 3:246609 | 0 | 64 | FALSE | 7 | 41 | 7 | 115 | 129 | 482 |
| 5,20975E | FALSE | High | IsMasterP | AAEL0019 | 2:426286 | 0 | 40 | FALSE | 8 | 43 | 5 | 139 | 160 | 665 |
| -3,12527E | FALSE | High | IsMasterP | AAEL0132 | 3:280060 | 0 | 41 | FALSE | 12 | 37 | 12 | 282 | 311 | 695 |
| -1,5606E+ | FALSE | High | IsMasterP | AAEL0021 | 2:428659 | 0 | 44 | FALSE | 10 | 32 | 10 | 257 | 292 | 917 |
| -1,51949E | FALSE | High | IsMasterP | AAEL0072 | 3:195539 | 0 | 47 | FALSE | 9 | 39 | 9 | 173 | 199 | 785 |
| 5,96021E | FALSE | High | IsMasterP | AAEL0023 | 2:176069 | 0 | 29 | FALSE | 12 | 38 | 12 | 435 | 510 | 661 |
| 1,76567E | FALSE | High | IsMasterP | AAEL0086 | 2:363057 | 0 | 17 | FALSE | 16 | 38 | 16 | 1484 | 1656 | 644 |
| 5,58723E | FALSE | High | IsMasterP | AAEL0033 | 2:400341 | 0 | 41 | FALSE | 12 | 33 | 12 | 506 | 556 | 775 |
| 9,15843E | FALSE | High | IsMasterP | AAEL0073 | 2:233232 | 0 | 74 | FALSE | 7 | 34 | 7 | 180 | 204 | 680 |
| 8,92761E | FALSE | High | IsMasterP | AAEL0257 | NIGP0100 | 0 | 79 | FALSE | 9 | 29 | 9 | 159 | 181 | 841 |
| -8,78145E | FALSE | High | IsMasterP | AAEL0147 | 2:326650 | 0 | 34 | FALSE | 21 | 41 | 21 | 930 | 1027 | 684 |
| 8,42909E | FALSE | High | IsMasterP | AAEL0134 | 3:153051 | 0 | 39 | FALSE | 14 | 33 | 14 | 713 | 807 | 892 |
| 1,91785E | FALSE | High | IsMasterP | AAEL0070 | 3:298490 | 0 | 22 | FALSE | 13 | 37 | 13 | 945 | 1018 | 822 |
| 8,37156E | FALSE | High | IsMasterP | AAEL0113 | 3:140216 | 0 | 43 | FALSE | 14 | 40 | 14 | 487 | 533 | 618 |
| 6,88937E | FALSE | High | IsMasterP | AAEL0118 | 3:113277 | 0 | 18 | FALSE | 13 | 39 | 13 | 999 | 1134 | 857 |
| -1,74722E | FALSE | High | IsMasterP | AAEL0115 | 2:811888 | 0 | 15 | FALSE | 18 | 40 | 18 | 1929 | 2127 | 643 |
| 3,01579E | FALSE | High | IsMasterP | AAEL0092 | 3:390977 | 0 | 61 | FALSE | 4 | 35 | 4 | 89 | 100 | 986 |
| 7,44101E | FALSE | High | IsMasterP | AAEL0150 | 2:210393 | 0 | 59 | FALSE | 12 | 36 | 12 | 186 | 210 | 566 |
| 9,05105E | FALSE | High | IsMasterP | AAEL0062 | 2:221309 | 0 | 44 | FALSE | 6 | 29 | 6 | 209 | 216 | 660 |
| -6,98705E | FALSE | High | IsMasterP | AAEL0082 | 2:672402 | 0 | 58 | FALSE | 9 | 30 | 9 | 286 | 315 | 592 |
| 3,56346E | FALSE | High | IsMasterP | AAEL0119 | 3:337820 | 0 | 18 | FALSE | 12 | 32 | 12 | 1034 | 1198 | 901 |
| 6,41471E | FALSE | High | IsMasterP | AAEL0085 | 2:157178 | 0 | 29 | FALSE | 16 | 37 | 16 | 811 | 889 | 793 |
| 4,26542E | FALSE | High | IsMasterP | AAEL0113 | 3:201660 | 0 | 37 | FALSE | 18 | 37 | 18 | 782 | 880 | 644 |
| -8,92335E | FALSE | High | IsMasterP | AAEL0044 | 2:473323 | 0 | 29 | FALSE | 20 | 39 | 20 | 933 | 1043 | 734 |
| -8,46071E | FALSE | High | IsMasterP | AAEL0081 | 3:100420 | 0 | 46 | FALSE | 9 | 34 | 9 | 194 | 217 | 898 |

| | | | | | | | | | | | | | | |
|---|---|---|---|---|---|---|---|---|---|---|---|---|---|---|
| 9,1734E+1 | FALSE | High | IsMasterP | AAEL0054 | 2:2956151 | 0 | 50 | FALSE | 6 | 46 | 6 | 70 | 83 | 1005 |
| -2,62426E | FALSE | High | IsMasterP | AAEL0278 | 2:1261415 | 0 | 24 | FALSE | 12 | 37 | 12 | 733 | 814 | 498 |
| 4,51477E- | FALSE | High | IsMasterP | AAEL0086 | 2:3214258 | 0 | 48 | FALSE | 12 | 33 | 2 | 377 | 415 | 844 |
| 3,4745E+1 | FALSE | High | IsMasterP | AAEL0129 | 3:2061692 | 0 | 49 | FALSE | 9 | 40 | 9 | 183 | 210 | 1005 |
| -3,8236E+ | FALSE | High | IsMasterP | AAEL0090 | 1:5879894 | 0 | 43 | FALSE | 15 | 36 | 15 | 504 | 577 | 854 |
| -5,92192E | FALSE | High | IsMasterP | AAEL0208 | 3:6514896 | 0 | 36 | FALSE | 14 | 37 | 14 | 507 | 577 | 742 |
| 7,35879E- | FALSE | High | IsMasterP | AAEL0005 | 3:1630902 | 0 | 40 | FALSE | 11 | 32 | 11 | 442 | 504 | 514 |
| 8,16767E- | FALSE | High | IsMasterP | AAEL0004 | 3:1628550 | 0 | 22 | FALSE | 16 | 37 | 16 | 854 | 955 | 661 |
| 5,53787E- | FALSE | High | IsMasterP | AAEL0073 | 3:1231945 | 0 | 42 | FALSE | 13 | 29 | 13 | 401 | 456 | 633 |
| -5,99633E | FALSE | High | IsMasterP | AAEL0038 | 3:8526269 | 0 | 30 | FALSE | 14 | 35 | 14 | 728 | 812 | 803 |
| 2,39736E- | FALSE | High | IsMasterP | AAEL0093 | 1:5537203 | 0 | 33 | FALSE | 16 | 37 | 16 | 728 | 796 | 742 |
| -2,38443E | FALSE | High | IsMasterP | AAEL0210 | 3:1044065 | 0 | 42 | FALSE | 8 | 38 | 8 | 119 | 134 | 1032 |
| -7,6376E+ | FALSE | High | IsMasterP | AAEL0126 | 2:1879592 | 0 | 55 | FALSE | 10 | 32 | 10 | 272 | 303 | 832 |
| -1,79493E | FALSE | High | IsMasterP | AAEL0131 | 2:1884760 | 0 | 66 | FALSE | 10 | 35 | 10 | 207 | 232 | 657 |
| 3,2198E+1 | FALSE | High | IsMasterP | AAEL0063 | 3:3967776 | 0 | 36 | FALSE | 6 | 33 | 6 | 221 | 248 | 465 |
| -2,77459E | FALSE | High | IsMasterP | AAEL0058 | 1:2239786 | 0 | 45 | FALSE | 17 | 36 | 17 | 477 | 536 | 559 |
| -1,71112E | FALSE | High | IsMasterP | AAEL0004 | 3:1834737 | 0 | 19 | FALSE | 13 | 29 | 13 | 1527 | 1720 | 594 |
| 8,89126E- | FALSE | High | IsMasterP | AAEL0079 | 2:3518546 | 0 | 7 | FALSE | 9 | 36 | 9 | 1805 | 1824 | 550 |
| 3,1365E+1 | FALSE | High | IsMasterP | AAEL0055 | 1:1508864 | 0 | 64 | FALSE | 12 | 38 | 12 | 240 | 265 | 736 |
| -1,75541E | FALSE | High | IsMasterP | AAEL0078 | 2:2725507 | 0 | 25 | FALSE | 15 | 37 | 15 | 944 | 1064 | 884 |
| -8,5116E+ | FALSE | High | IsMasterP | AAEL0094 | 2:1518331 | 0 | 28 | FALSE | 12 | 36 | 12 | 629 | 700 | 734 |
| -4,71655E | FALSE | High | IsMasterP | AAEL0169 | 3:2968040 | 0 | 65 | FALSE | 11 | 32 | 11 | 273 | 301 | 706 |
| -7,17887E | FALSE | High | IsMasterP | AAEL0069 | 1:1573803 | 0 | 24 | FALSE | 18 | 31 | 18 | 1308 | 1491 | 681 |
| 6,49064E- | FALSE | High | IsMasterP | AAEL0135 | 2:1992333 | 0 | 30 | FALSE | 15 | 37 | 13 | 705 | 813 | 743 |
| 4,85696E- | FALSE | High | IsMasterP | AAEL0132 | 3:2799441 | 0 | 38 | FALSE | 22 | 36 | 21 | 945 | 1057 | 966 |
| -8,09093E | FALSE | High | IsMasterP | AAEL0207 | 3:3102186 | 0 | 52 | FALSE | 10 | 40 | 10 | 149 | 171 | 1089 |
| -2,78304E | FALSE | High | IsMasterP | AAEL0106 | 2:1360175 | 0 | 54 | FALSE | 12 | 36 | 11 | 245 | 267 | 761 |
| 1,09245E- | FALSE | High | IsMasterP | AAEL0263 | 3:3371849 | 0 | 14 | FALSE | 14 | 33 | 14 | 1931 | 2018 | 578 |
| 7,94798E- | FALSE | High | IsMasterP | AAEL0117 | 3:2910590 | 0 | 70 | FALSE | 11 | 34 | 9 | 237 | 270 | 569 |
| 2,78149E- | FALSE | High | IsMasterP | AAEL0037 | 1:2479473 | 0 | 48 | FALSE | 13 | 40 | 13 | 371 | 421 | 701 |
| -5,8187E+ | FALSE | High | IsMasterP | AAEL0142 | 3:2953457 | 0 | 28 | FALSE | 9 | 33 | 9 | 567 | 630 | 747 |

| | | | | | | | | | | | | | | |
|---|---|---|---|---|---|---|---|---|---|---|---|---|---|---|
| -1,73759E | FALSE | High | IsMasterP | AAEL0006 | 2:459681 | 0 | 58 | FALSE | 5 | 31 | 5 | 78 | 88 | 742 |
| 2,19629E | FALSE | High | IsMasterP | AAEL0226 | 1:158301 | 0 | 7 | FALSE | 10 | 35 | 6 | 1890 | 2134 | 907 |
| 9,1513E+ | FALSE | High | IsMasterP | AAEL0011 | 2:307745 | 0 | 40 | FALSE | 9 | 32 | 9 | 468 | 510 | 765 |
| 8,31637E | FALSE | High | IsMasterP | AAEL0021 | 1:251225 | 0 | 23 | FALSE | 10 | 30 | 10 | 435 | 481 | 604 |
| -2,40394E | FALSE | High | IsMasterP | AAEL0108 | 3:517719 | 0 | 38 | FALSE | 7 | 32 | 7 | 250 | 293 | 435 |
| 5,37563E | FALSE | High | IsMasterP | AAEL0149 | 3:204254 | 0 | 47 | FALSE | 6 | 39 | 6 | 152 | 170 | 558 |
| -3,40056E | FALSE | High | IsMasterP | AAEL0083 | 1:961733 | 0 | 66 | FALSE | 10 | 32 | 10 | 169 | 190 | 797 |
| -2,58867E | FALSE | High | IsMasterP | AAEL0043 | 3:368097 | 0 | 35 | FALSE | 17 | 36 | 17 | 735 | 837 | 630 |
| -2,29909E | FALSE | High | IsMasterP | AAEL0018 | 2:274542 | 0 | 27 | FALSE | 11 | 31 | 11 | 543 | 609 | 516 |
| -5,58938E | FALSE | High | IsMasterP | AAEL0123 | 1:284553 | 0 | 25 | FALSE | 11 | 35 | 11 | 552 | 630 | 460 |
| -3,2315E+ | FALSE | High | IsMasterP | AAEL0069 | 2:399100 | 0 | 27 | FALSE | 12 | 28 | 12 | 804 | 904 | 610 |
| -5,46793E | FALSE | High | IsMasterP | AAEL0108 | 3:517893 | 0 | 40 | FALSE | 13 | 32 | 13 | 472 | 512 | 917 |
| -2,16073E | FALSE | High | IsMasterP | AAEL0130 | 2:330381 | 0 | 47 | FALSE | 2 | 29 | 1 | 108 | 114 | 718 |
| 1,11619E | FALSE | High | IsMasterP | AAEL0001 | 1:197332 | 0 | 56 | FALSE | 10 | 33 | 10 | 276 | 308 | 572 |
| 7,21194E | FALSE | High | IsMasterP | AAEL0280 | 2:418985 | 0 | 43 | FALSE | 7 | 35 | 7 | 152 | 170 | 1172 |
| 8,37543E | FALSE | High | IsMasterP | AAEL0067 | 2:847675 | 0 | 31 | FALSE | 13 | 33 | 13 | 665 | 718 | 977 |
| -3,8171E+ | FALSE | High | IsMasterP | AAEL0011 | 3:167925 | 0 | 32 | FALSE | 4 | 27 | 4 | 146 | 161 | 740 |
| -7,55604E | FALSE | High | IsMasterP | AAEL0253 | 2:474107 | 0 | 6 | FALSE | 21 | 31 | 21 | 8614 | 9852 | 464 |
| -6,71021E | FALSE | High | IsMasterP | AAEL0142 | 1:838336 | 0 | 58 | FALSE | 13 | 32 | 13 | 326 | 365 | 891 |
| -5,40491E | FALSE | High | IsMasterP | AAEL0069 | 2:398719 | 0 | 40 | FALSE | 10 | 34 | 10 | 284 | 316 | 853 |
| -5,87738E | FALSE | High | IsMasterP | AAEL0002 | 3:265003 | 0 | 39 | FALSE | 9 | 29 | 9 | 336 | 378 | 648 |
| 2,61759E | FALSE | High | IsMasterP | AAEL0226 | 2:131217 | 0 | 14 | FALSE | 16 | 36 | 16 | 2039 | 2244 | 677 |
| -1,07822E | FALSE | High | IsMasterP | AAEL0108 | 2:127687 | 0 | 71 | FALSE | 8 | 29 | 8 | 110 | 123 | 677 |
| -1,0869E+ | FALSE | High | IsMasterP | AAEL0060 | 3:740210 | 0 | 35 | FALSE | 9 | 31 | 9 | 398 | 443 | 660 |
| -3,70791E | FALSE | High | IsMasterP | AAEL0069 | 2:398747 | 0 | 28 | FALSE | 12 | 32 | 12 | 381 | 430 | 706 |
| 7,85261E | FALSE | High | IsMasterP | AAEL0105 | 2:111943 | 0 | 60 | FALSE | 11 | 29 | 11 | 320 | 347 | 539 |
| -3,71357E | FALSE | High | IsMasterP | AAEL0087 | 3:132553 | 0 | 55 | FALSE | 10 | 33 | 10 | 196 | 221 | 572 |
| 3,96951E | FALSE | High | IsMasterP | AAEL0056 | 2:382037 | 0 | 45 | FALSE | 10 | 25 | 10 | 449 | 514 | 534 |
| -6,43531E | FALSE | High | IsMasterP | AAEL0047 | 2:288639 | 0 | 55 | FALSE | 9 | 31 | 9 | 282 | 313 | 895 |
| 8,14993E | FALSE | High | IsMasterP | AAEL0103 | 2:287636 | 0 | 54 | FALSE | 11 | 33 | 11 | 289 | 323 | 863 |
| 6,42191E | FALSE | High | IsMasterP | AAEL0094 | 1:573365 | 0 | 74 | FALSE | 9 | 32 | 9 | 163 | 185 | 506 |

| | | | | | | | | | | | | | | |
|---|---|---|---|---|---|---|---|---|---|---|---|---|---|---|
| 7,27939E- | FALSE | High | IsMasterP | AAEL0042 | 1:104968 | 0 | 58 | FALSE | 7 | 28 | 7 | 132 | 146 | 857 |
| -2,56372E | FALSE | High | IsMasterP | AAEL0081 | 3:108044 | 0 | 42 | FALSE | 13 | 33 | 13 | 291 | 321 | 904 |
| -5,69861E | FALSE | High | IsMasterP | AAEL0067 | 3:168978 | 0 | 47 | FALSE | 10 | 28 | 10 | 358 | 399 | 756 |
| 6,50621E- | FALSE | High | IsMasterP | AAEL0135 | 1:144001 | 0 | 45 | FALSE | 10 | 37 | 10 | 232 | 259 | 853 |
| 9,04363E- | FALSE | High | IsMasterP | AAEL0049 | 2:260868 | 0 | 29 | FALSE | 15 | 29 | 15 | 788 | 891 | 521 |
| -1,73323E | FALSE | High | IsMasterP | AAEL0044 | 3:206730 | 0 | 56 | FALSE | 8 | 34 | 8 | 172 | 195 | 539 |
| 6,73012E- | FALSE | High | IsMasterP | AAEL0087 | 3:280445 | 0 | 23 | FALSE | 16 | 37 | 16 | 972 | 1095 | 524 |
| -1,21717E | FALSE | High | IsMasterP | AAEL0091 | 3:135343 | 0 | 54 | FALSE | 10 | 31 | 10 | 287 | 316 | 521 |
| 7,19378E- | FALSE | High | IsMasterP | AAEL0021 | 3:177829 | 0 | 35 | FALSE | 11 | 34 | 11 | 445 | 494 | 903 |
| -2,28661E | FALSE | High | IsMasterP | AAEL0027 | 3:129324 | 0 | 53 | FALSE | 10 | 28 | 10 | 262 | 303 | 558 |
| 1,65468E- | FALSE | High | IsMasterP | AAEL0130 | 2:337759 | 0 | 17 | FALSE | 13 | 30 | 13 | 1227 | 1390 | 746 |
| -3,15743E | FALSE | High | IsMasterP | AAEL0013 | 2:478534 | 0 | 20 | FALSE | 12 | 34 | 12 | 982 | 1084 | 520 |
| 6,88792E- | FALSE | High | IsMasterP | AAEL0123 | 2:102915 | 0 | 39 | FALSE | 10 | 31 | 10 | 418 | 470 | 596 |
| 2,12178E- | FALSE | High | IsMasterP | AAEL0279 | 3:361349 | 0 | 27 | FALSE | 10 | 27 | 10 | 599 | 671 | 652 |
| 6,38722E- | FALSE | High | IsMasterP | AAEL0009 | 3:379517 | 0 | 31 | FALSE | 14 | 31 | 14 | 562 | 615 | 662 |
| 3,87472E- | FALSE | High | IsMasterP | AAEL0132 | 3:280040 | 0 | 38 | FALSE | 13 | 32 | 13 | 519 | 575 | 587 |
| 2,45855E- | FALSE | High | IsMasterP | AAEL0064 | 3:216530 | 0 | 44 | FALSE | 13 | 30 | 13 | 237 | 272 | 974 |
| -7,12432E | FALSE | High | IsMasterP | AAEL0093 | 3:860924 | 0 | 67 | FALSE | 10 | 26 | 10 | 184 | 208 | 580 |
| 8,63351E- | FALSE | High | IsMasterP | AAEL0198 | 2:367609 | 0 | 37 | FALSE | 12 | 28 | 3 | 557 | 619 | 731 |
| 1,75993E- | FALSE | High | IsMasterP | AAEL0070 | 3:171135 | 0 | 54 | FALSE | 11 | 25 | 11 | 322 | 344 | 749 |
| 4,70873E- | FALSE | High | IsMasterP | AAEL0071 | 3:697811 | 0 | 58 | FALSE | 10 | 34 | 10 | 193 | 207 | 888 |
| -4,48356E | FALSE | High | IsMasterP | AAEL0195 | 2:245788 | 0 | 17 | FALSE | 6 | 22 | 6 | 755 | 829 | 529 |
| 2,61368E- | FALSE | High | IsMasterP | AAEL0070 | 3:299678 | 0 | 37 | FALSE | 8 | 35 | 8 | 359 | 400 | 775 |
| -2,18607E | FALSE | High | IsMasterP | AAEL0085 | 1:638410 | 0 | 28 | FALSE | 18 | 29 | 18 | 981 | 1113 | 595 |
| 7,28466E- | FALSE | High | IsMasterP | AAEL0237 | 3:177575 | 0 | 68 | FALSE | 5 | 32 | 5 | 133 | 138 | 812 |
| 1,64743E- | FALSE | High | IsMasterP | AAEL0116 | 3:214592 | 0 | 23 | FALSE | 7 | 26 | 7 | 534 | 585 | 578 |
| 8,44529E- | FALSE | High | IsMasterP | AAEL0244 | 2:180600 | 0 | 14 | FALSE | 9 | 27 | 9 | 801 | 920 | 807 |
| 8,33577E- | FALSE | High | IsMasterP | AAEL0124 | 1:725017 | 0 | 14 | FALSE | 14 | 28 | 14 | 1334 | 1505 | 648 |
| -4,0211E+ | FALSE | High | IsMasterP | AAEL0105 | 2:367206 | 0 | 68 | FALSE | 7 | 24 | 7 | 87 | 98 | 422 |
| 5,68728E- | FALSE | High | IsMasterP | AAEL0114 | 3:328879 | 0 | 24 | FALSE | 13 | 34 | 13 | 727 | 813 | 587 |
| -1,11721E | FALSE | High | IsMasterP | AAEL0268 | 2:150018 | 0 | 24 | FALSE | 20 | 33 | 20 | 1324 | 1461 | 739 |

| | | | | | | | | | | | | | | |
|---|---|---|---|---|---|---|---|---|---|---|---|---|---|---|
| 3,66458E- | FALSE | High | IsMasterP | AAEL0081 | 3:1025218 | 0 | 21 | FALSE | 13 | 30 | 13 | 884 | 965 | 859 |
| -6,39327E | FALSE | High | IsMasterP | AAEL0148 | 2:1557703 | 0 | 12 | FALSE | 5 | 22 | 1 | 832 | 904 | 802 |
| -3,74465E | FALSE | High | IsMasterP | AAEL0005 | 3:3881371 | 0 | 28 | FALSE | 16 | 31 | 16 | 989 | 1101 | 492 |
| 6,9501E+1 | FALSE | High | IsMasterP | AAEL0013 | 2:4571493 | 0 | 17 | FALSE | 15 | 29 | 15 | 1210 | 1348 | 699 |
| -6,49593E | FALSE | High | IsMasterP | AAEL0054 | 1:2471897 | 0 | 43 | FALSE | 8 | 28 | 6 | 321 | 363 | 538 |
| -7,11554E | FALSE | High | IsMasterP | AAEL0117 | 1:2386294 | 0 | 27 | FALSE | 11 | 34 | 11 | 371 | 409 | 517 |
| -1,45761E | FALSE | High | IsMasterP | AAEL0135 | 1:1440295 | 0 | 45 | FALSE | 10 | 26 | 10 | 417 | 471 | 655 |
| -1,15661E | FALSE | High | IsMasterP | AAEL0086 | 2:3214258 | 0 | 51 | FALSE | 11 | 28 | 1 | 329 | 364 | 644 |
| -2,8846E+ | FALSE | High | IsMasterP | AAEL0033 | 2:4001934 | 0 | 42 | FALSE | 10 | 32 | 10 | 388 | 423 | 496 |
| 9,10853E- | FALSE | High | IsMasterP | AAEL0098 | 1:9242588 | 0 | 6 | FALSE | 20 | 34 | 20 | 4651 | 5184 | 539 |
| 3,49825E- | FALSE | High | IsMasterP | AAEL0013 | 2:2062325 | 0 | 29 | FALSE | 10 | 30 | 10 | 441 | 493 | 649 |
| -6,89405E | FALSE | High | IsMasterP | AAEL0010 | 3:5516219 | 0 | 38 | FALSE | 8 | 27 | 8 | 375 | 413 | 597 |
| 3,63305E- | FALSE | High | IsMasterP | AAEL0000 | 1:2427792 | 0 | 91 | FALSE | 6 | 24 | 6 | 118 | 133 | 512 |
| 6,26327E- | FALSE | High | IsMasterP | AAEL0141 | 2:3553945 | 0 | 44 | FALSE | 12 | 29 | 12 | 360 | 422 | 539 |
| -3,68922E | FALSE | High | IsMasterP | AAEL0245 | 2:1626019 | 0 | 44 | FALSE | 11 | 29 | 11 | 204 | 242 | 1137 |
| 4,56387E- | FALSE | High | IsMasterP | AAEL0024 | 3:3381906 | 0 | 76 | FALSE | 9 | 23 | 9 | 127 | 141 | 618 |
| 7,2651E+1 | FALSE | High | IsMasterP | AAEL0121 | 2:2038623 | 0 | 27 | FALSE | 12 | 29 | 12 | 624 | 694 | 591 |
| -5,13659E | FALSE | High | IsMasterP | AAEL0099 | 1:2072694 | 0 | 62 | FALSE | 9 | 30 | 9 | 119 | 135 | 995 |
| -5,42171E | FALSE | High | IsMasterP | AAEL0116 | 3:2168830 | 0 | 28 | FALSE | 16 | 29 | 16 | 956 | 1047 | 520 |
| -6,39666E | FALSE | High | IsMasterP | AAEL0196 | 2:1042499 | 0 | 27 | FALSE | 13 | 32 | 13 | 704 | 752 | 674 |
| 4,51629E- | FALSE | High | IsMasterP | AAEL0009 | 3:3329940 | 0 | 25 | FALSE | 8 | 26 | 8 | 581 | 638 | 632 |
| -5,8566E+ | FALSE | High | IsMasterP | AAEL0104 | 3:3138071 | 0 | 59 | FALSE | 7 | 27 | 7 | 92 | 105 | 721 |
| 3,89112E- | FALSE | High | IsMasterP | AAEL0003 | 3:9760444 | 0 | 15 | FALSE | 16 | 34 | 16 | 1226 | 1418 | 812 |
| -4,11247E | FALSE | High | IsMasterP | AAEL0008 | 3:3637962 | 0 | 64 | FALSE | 9 | 26 | 9 | 163 | 186 | 500 |
| 1,32546E- | FALSE | High | IsMasterP | AAEL0128 | 2:3354375 | 0 | 41 | FALSE | 12 | 28 | 12 | 454 | 510 | 599 |
| 1,23279E- | FALSE | High | IsMasterP | AAEL0000 | 3:1050981 | 0 | 60 | FALSE | 14 | 32 | 14 | 362 | 396 | 762 |
| -3,69352E | FALSE | High | IsMasterP | AAEL0072 | 3:1962234 | 0 | 10 | FALSE | 12 | 25 | 12 | 1590 | 1800 | 548 |
| -2,84658E | FALSE | High | IsMasterP | AAEL0138 | 1:7190058 | 0 | 53 | FALSE | 8 | 25 | 8 | 228 | 248 | 569 |
| 8,93671E- | FALSE | High | IsMasterP | AAEL0129 | 1:2304134 | 0 | 23 | FALSE | 10 | 27 | 10 | 585 | 660 | 976 |
| 4,73912E- | FALSE | High | IsMasterP | AAEL0212 | 1:2131192 | 0 | 13 | FALSE | 14 | 32 | 14 | 1800 | 1983 | 851 |
| 2,74494E- | FALSE | High | IsMasterP | AAEL0084 | 2:1921843 | 0 | 24 | FALSE | 4 | 25 | 4 | 189 | 220 | 1103 |

| | | | | | | | | | | | | | | |
|---|---|---|---|---|---|---|---|---|---|---|---|---|---|---|
| 2,22049E- | FALSE | High | IsMasterP | AAEL0025 | 1:8931662 | 0 | 66 | FALSE | 5 | 26 | 5 | 93 | 105 | 548 |
| -3,90825E | FALSE | High | IsMasterP | AAEL0029 | 1:1087482 | 0 | 39 | FALSE | 8 | 26 | 8 | 337 | 375 | 511 |
| 2,88728E- | FALSE | High | IsMasterP | AAEL0002 | 3:2649198 | 0 | 10 | FALSE | 9 | 27 | 9 | 1536 | 1708 | 543 |
| -2,62583E | FALSE | High | IsMasterP | AAEL0076 | 2:2507579 | 0 | 37 | FALSE | 13 | 29 | 12 | 646 | 724 | 502 |
| 7,04235E- | FALSE | High | IsMasterP | AAEL0252 | 2:3341331 | 0 | 18 | FALSE | 15 | 29 | 15 | 1393 | 1534 | 552 |
| 6,93552E- | FALSE | High | IsMasterP | AAEL0066 | 3:3532427 | 0 | 54 | FALSE | 9 | 33 | 9 | 124 | 146 | 1035 |
| -5,17194E | FALSE | High | IsMasterP | AAEL0091 | 3:1760262 | 0 | 27 | FALSE | 12 | 28 | 12 | 654 | 718 | 676 |
| -1,12267E | FALSE | High | IsMasterP | AAEL0088 | 3:3372331 | 0 | 37 | FALSE | 12 | 29 | 12 | 483 | 533 | 562 |
| -5,33537E | FALSE | High | IsMasterP | AAEL0058 | 1:2002319 | 0 | 32 | FALSE | 6 | 33 | 6 | 151 | 174 | 1070 |
| 3,42586E- | FALSE | High | IsMasterP | AAEL0017 | 2:4432688 | 0 | 78 | FALSE | 6 | 23 | 6 | 95 | 101 | 465 |
| -9,19585E | FALSE | High | IsMasterP | AAEL0122 | 2:1449983 | 0 | 43 | FALSE | 10 | 28 | 10 | 241 | 282 | 478 |
| 1,00715E- | FALSE | High | IsMasterP | AAEL0137 | 2:9573912 | 0 | 59 | FALSE | 7 | 23 | 7 | 185 | 212 | 931 |
| -2,40451E | FALSE | High | IsMasterP | AAEL0061 | 3:2597690 | 0 | 40 | FALSE | 7 | 26 | 7 | 185 | 198 | 825 |
| 4,61619E- | FALSE | High | IsMasterP | AAEL0206 | 2:1348307 | 0 | 11 | FALSE | 11 | 29 | 11 | 1617 | 1780 | 448 |
| 1,91213E- | FALSE | High | IsMasterP | AAEL0139 | 3:2305839 | 0 | 41 | FALSE | 12 | 31 | 12 | 330 | 380 | 897 |
| 1,75556E- | FALSE | High | IsMasterP | AAEL0000 | 3:1041477 | 0 | 75 | FALSE | 9 | 29 | 9 | 193 | 216 | 484 |
| -2,36517E | FALSE | High | IsMasterP | AAEL0067 | 2:8488592 | 0 | 74 | FALSE | 11 | 29 | 11 | 177 | 201 | 591 |
| 1,58209E- | FALSE | High | IsMasterP | AAEL0142 | 2:3998820 | 0 | 49 | FALSE | 10 | 24 | 10 | 362 | 400 | 597 |
| -7,02811E | FALSE | High | IsMasterP | AAEL0146 | 3:3684811 | 0 | 28 | FALSE | 13 | 31 | 4 | 541 | 624 | 720 |
| 4,59027E- | FALSE | High | IsMasterP | AAEL0121 | 2:2042111 | 0 | 24 | FALSE | 14 | 26 | 14 | 756 | 854 | 960 |
| -4,10346E | FALSE | High | IsMasterP | AAEL0054 | 1:2468369 | 0 | 33 | FALSE | 13 | 28 | 13 | 455 | 489 | 698 |
| -8,04872E | FALSE | High | IsMasterP | AAEL0049 | 2:7239377 | 0 | 35 | FALSE | 7 | 28 | 7 | 213 | 237 | 639 |
| -7,46821E | FALSE | High | IsMasterP | AAEL0049 | 2:2817508 | 0 | 20 | FALSE | 11 | 29 | 11 | 843 | 946 | 714 |
| 5,03817E- | FALSE | High | IsMasterP | AAEL0199 | 1:1396689 | 0 | 13 | FALSE | 9 | 29 | 9 | 1016 | 1118 | 703 |
| 1,7287E+1 | FALSE | High | IsMasterP | AAEL0040 | 1:1354815 | 0 | 61 | FALSE | 9 | 32 | 9 | 194 | 215 | 730 |
| -6,18112E | FALSE | High | IsMasterP | AAEL0198 | 2:3676094 | 0 | 35 | FALSE | 10 | 26 | 1 | 499 | 549 | 647 |
| -2,28296E | FALSE | High | IsMasterP | AAEL0064 | 3:2162887 | 0 | 28 | FALSE | 15 | 26 | 15 | 972 | 1096 | 709 |
| 3,66355E- | FALSE | High | IsMasterP | AAEL0146 | 2:3475971 | 0 | 38 | FALSE | 15 | 26 | 15 | 598 | 657 | 766 |
| 6,76379E- | FALSE | High | IsMasterP | AAEL0088 | 3:3236141 | 0 | 9 | FALSE | 11 | 27 | 11 | 2198 | 2480 | 527 |
| 8,59263E- | FALSE | High | IsMasterP | AAEL0202 | 3:2888422 | 0 | 22 | FALSE | 9 | 27 | 9 | 517 | 563 | 505 |
| 2,94043E- | FALSE | High | IsMasterP | AAEL0105 | 2:2599025 | 0 | 13 | FALSE | 9 | 30 | 9 | 1201 | 1310 | 780 |

| | | | | | | | | | | | | | | |
|---|---|---|---|---|---|---|---|---|---|---|---|---|---|---|
| -3,07994E | FALSE | High | IsMasterP | AAEL0115 | 3:315246! | 0 | 45 | FALSE | 7 | 34 | 7 | 136 | 160 | 1100 |
| 1,02218E- | FALSE | High | IsMasterP | AAEL0117 | NIGP0100 | 0 | 34 | FALSE | 12 | 30 | 12 | 539 | 583 | 724 |
| 8,28782E- | FALSE | High | IsMasterP | AAEL0010 | 3:555227: | 0 | 21 | FALSE | 8 | 23 | 8 | 517 | 527 | 942 |
| -7,17122E | FALSE | High | IsMasterP | AAEL0068 | 2:256719! | 0 | 21 | FALSE | 11 | 24 | 11 | 790 | 865 | 730 |
| 5,67744E- | FALSE | High | IsMasterP | AAEL0070 | 3:299850 | 0 | 14 | FALSE | 7 | 18 | 7 | 621 | 691 | 491 |
| 5,82402E- | FALSE | High | IsMasterP | AAEL0081 | 3:149705( | 0 | 6 | FALSE | 19 | 28 | 19 | 5366 | 5911 | 600 |
| 5,46616E- | FALSE | High | IsMasterP | AAEL0051 | 2:321867! | 0 | 50 | FALSE | 7 | 30 | 7 | 177 | 203 | 944 |
| -1,5162E+ | FALSE | High | IsMasterP | AAEL0114 | 3:328847! | 0 | 48 | FALSE | 5 | 29 | 5 | 112 | 127 | 938 |
| -7,38165E | FALSE | High | IsMasterP | AAEL0040 | 1:137364: | 0 | 43 | FALSE | 11 | 30 | 11 | 403 | 442 | 648 |
| 2,82869E- | FALSE | High | IsMasterP | AAEL0259 | 3:338159! | 0 | 17 | FALSE | 9 | 28 | 9 | 758 | 856 | 522 |
| -6,88987E | FALSE | High | IsMasterP | AAEL0106 | 3:291599: | 0 | 21 | FALSE | 9 | 27 | 9 | 684 | 795 | 664 |
| 5,97226E- | FALSE | High | IsMasterP | AAEL0140 | 1:142958( | 0 | 49 | FALSE | 8 | 27 | 8 | 298 | 331 | 778 |
| -5,55368E | FALSE | High | IsMasterP | AAEL0174 | 3:347304: | 0 | 19 | FALSE | 4 | 28 | 1 | 244 | 265 | 665 |
| -3,62188E | FALSE | High | IsMasterP | AAEL0117 | 3:291059( | 0 | 45 | FALSE | 9 | 26 | 7 | 237 | 270 | 549 |
| 6,00811E- | FALSE | High | IsMasterP | AAEL0090 | 2:270339! | 0 | 32 | FALSE | 19 | 32 | 19 | 725 | 812 | 680 |
| 1,24038E- | FALSE | High | IsMasterP | AAEL0261 | 2:276625( | 0 | 12 | FALSE | 14 | 29 | 14 | 1409 | 1569 | 662 |
| 5,19405E- | FALSE | High | IsMasterP | AAEL0097 | 1:675773: | 0 | 51 | FALSE | 4 | 24 | 4 | 130 | 145 | 539 |
| 1,80646E- | FALSE | High | IsMasterP | AAEL0034 | 2:150864 | 0 | 43 | FALSE | 6 | 24 | 6 | 251 | 277 | 453 |
| 1,29601E- | FALSE | High | IsMasterP | AAEL0290 | 2:303653: | 0 | 14 | FALSE | 5 | 29 | 5 | 660 | 704 | 657 |
| -2,15533E | FALSE | High | IsMasterP | AAEL0122 | 2:335016 | 0 | 59 | FALSE | 7 | 22 | 7 | 235 | 254 | 550 |
| -8,07396E | FALSE | High | IsMasterP | AAEL0148 | 1:149226! | 0 | 36 | FALSE | 5 | 25 | 5 | 205 | 225 | 690 |
| 7,47322E- | FALSE | High | IsMasterP | AAEL0121 | 3:298080! | 0 | 34 | FALSE | 6 | 26 | 6 | 305 | 346 | 642 |
| 3,58434E- | FALSE | High | IsMasterP | AAEL0061 | 3:141764: | 0 | 10 | FALSE | 19 | 31 | 19 | 2657 | 2931 | 630 |
| -6,19371E | FALSE | High | IsMasterP | AAEL0119 | 2:218456! | 0 | 22 | FALSE | 10 | 26 | 10 | 823 | 931 | 668 |
| -3,72993E | FALSE | High | IsMasterP | AAEL0142 | 2:362719 | 0 | 23 | FALSE | 10 | 26 | 10 | 560 | 641 | 903 |
| -3,4399E+ | FALSE | High | IsMasterP | AAEL0028 | 3:235222 | 0 | 31 | FALSE | 4 | 28 | 4 | 115 | 133 | 1074 |
| 8,93061E- | FALSE | High | IsMasterP | AAEL0010 | 3:559336( | 0 | 49 | FALSE | 6 | 23 | 6 | 283 | 313 | 576 |
| -5,81162E | FALSE | High | IsMasterP | AAEL0197 | 2:232116: | 0 | 42 | FALSE | 5 | 21 | 5 | 215 | 232 | 977 |
| -5,64033E | FALSE | High | IsMasterP | AAEL0139 | 3:105826 | 0 | 46 | FALSE | 10 | 26 | 10 | 419 | 438 | 484 |
| -3,3047E+ | FALSE | High | IsMasterP | AAEL0078 | 3:381939: | 0 | 67 | FALSE | 6 | 22 | 6 | 110 | 133 | 728 |
| -1,59972E | FALSE | High | IsMasterP | AAEL0172 | 3:264135! | 0 | 12 | FALSE | 9 | 27 | 9 | 1000 | 1158 | 955 |

| | | | | | | | | | | | | | | |
|---|---|---|---|---|---|---|---|---|---|---|---|---|---|---|
| -1,6429E+ | FALSE | High | IsMasterP | AAEL0008 | 3:1204964 | 0 | 31 | FALSE | 11 | 27 | 11 | 499 | 570 | 591 |
| -8,53225E | FALSE | High | IsMasterP | AAEL0038 | 3:8465654 | 0 | 60 | FALSE | 10 | 28 | 5 | 1142 | 1283 | 796 |
| 5,47808E+ | FALSE | High | IsMasterP | AAEL0248 | 1:8613343 | 0 | 24 | FALSE | 8 | 27 | 8 | 286 | 311 | 600 |
| -4,9855E+ | FALSE | High | IsMasterP | AAEL0081 | 3:5361534 | 0 | 24 | FALSE | 10 | 32 | 10 | 264 | 302 | 1119 |
| -3,63361E | FALSE | High | IsMasterP | AAEL0125 | 1:2852190 | 0 | 29 | FALSE | 9 | 28 | 2 | 536 | 582 | 799 |
| 4,98953E+ | FALSE | High | IsMasterP | AAEL0063 | 3:2309791 | 0 | 37 | FALSE | 6 | 24 | 6 | 336 | 365 | 451 |
| 6,22471E+ | FALSE | High | IsMasterP | AAEL0054 | 1:2471549 | 0 | 42 | FALSE | 9 | 24 | 5 | 321 | 360 | 482 |
| 5,41771E+ | FALSE | High | IsMasterP | AAEL0126 | 1:1460499 | 0 | 15 | FALSE | 12 | 28 | 12 | 1066 | 1117 | 755 |
| -5,35502E | FALSE | High | IsMasterP | AAEL0149 | 2:2228190 | 0 | 29 | FALSE | 13 | 28 | 13 | 694 | 793 | 714 |
| 9,20915E+ | FALSE | High | IsMasterP | AAEL0028 | 1:2560523 | 0 | 34 | FALSE | 9 | 24 | 9 | 496 | 559 | 681 |
| 4,08492E+ | FALSE | High | IsMasterP | AAEL0043 | 3:9304495 | 0 | 36 | FALSE | 6 | 26 | 6 | 148 | 171 | 514 |
| 4,82031E+ | FALSE | High | IsMasterP | AAEL0039 | 3:2669823 | 0 | 16 | FALSE | 7 | 27 | 7 | 649 | 708 | 539 |
| 6,62253E+ | FALSE | High | IsMasterP | AAEL0221 | 1:2183687 | 0 | 14 | FALSE | 12 | 26 | 12 | 1229 | 1422 | 869 |
| 8,12366E+ | FALSE | High | IsMasterP | AAEL0195 | 1:2111789 | 0 | 46 | FALSE | 11 | 23 | 11 | 401 | 434 | 487 |
| 8,83743E+ | FALSE | High | IsMasterP | AAEL0021 | 2:4284055 | 0 | 34 | FALSE | 7 | 26 | 7 | 275 | 310 | 572 |
| 6,52646E+ | FALSE | High | IsMasterP | AAEL0039 | 2:3544893 | 0 | 22 | FALSE | 7 | 26 | 7 | 481 | 532 | 928 |
| 5,75151E+ | FALSE | High | IsMasterP | AAEL0103 | 2:1829785 | 0 | 15 | FALSE | 13 | 29 | 12 | 1268 | 1394 | 673 |
| 8,04325E+ | FALSE | High | IsMasterP | AAEL0174 | 2:3554313 | 0 | 38 | FALSE | 5 | 23 | 5 | 157 | 170 | 819 |
| -1,83959E | FALSE | High | IsMasterP | AAEL0102 | 1:4365207 | 0 | 49 | FALSE | 5 | 22 | 5 | 166 | 196 | 963 |
| 5,03954E+ | FALSE | High | IsMasterP | AAEL0197 | 2:1704224 | 0 | 19 | FALSE | 12 | 22 | 12 | 1212 | 1353 | 563 |
| -6,6189E+ | FALSE | High | IsMasterP | AAEL0017 | 2:1657934 | 0 | 38 | FALSE | 9 | 30 | 9 | 195 | 228 | 1061 |
| 6,04588E+ | FALSE | High | IsMasterP | AAEL0226 | 2:4191528 | 0 | 26 | FALSE | 15 | 27 | 15 | 800 | 905 | 862 |
| -7,28325E | FALSE | High | IsMasterP | AAEL0067 | 2:8373315 | 0 | 37 | FALSE | 9 | 20 | 9 | 325 | 361 | 844 |
| 7,08159E+ | FALSE | High | IsMasterP | AAEL0194 | 2:4562497 | 0 | 25 | FALSE | 10 | 25 | 10 | 688 | 724 | 832 |
| 6,87903E+ | FALSE | High | IsMasterP | AAEL0073 | 3:1231500 | 0 | 50 | FALSE | 7 | 21 | 7 | 233 | 256 | 465 |
| -5,8466E+ | FALSE | High | IsMasterP | AAEL0255 | 1:3058685 | 0 | 29 | FALSE | 16 | 27 | 16 | 811 | 933 | 693 |
| -6,96278E | FALSE | High | IsMasterP | AAEL0151 | 2:1993390 | 0 | 28 | FALSE | 12 | 26 | 10 | 684 | 783 | 706 |
| -6,43268E | FALSE | High | IsMasterP | AAEL0052 | 2:719598[illegible] | 0 | 33 | FALSE | 11 | 29 | 11 | 366 | 420 | 733 |
| 8,50674E+ | FALSE | High | IsMasterP | AAEL0107 | 2:4065445 | 0 | 30 | FALSE | 7 | 27 | 7 | 446 | 491 | 813 |
| -6,37845E | FALSE | High | IsMasterP | AAEL0031 | 2:1918542 | 0 | 39 | FALSE | 10 | 22 | 10 | 424 | 465 | 816 |
| 4,20752E+ | FALSE | High | IsMasterP | AAEL0195 | 2:3806542 | 0 | 15 | FALSE | 9 | 25 | 9 | 844 | 930 | 805 |

| | | | | | | | | | | | | | | |
|---|---|---|---|---|---|---|---|---|---|---|---|---|---|---|
| 5,30907E | FALSE | High | IsMasterP | AAEL0116 | 3:218412 | 0 | 54 | FALSE | 11 | 20 | 11 | 519 | 574 | 503 |
| 1,57873E | FALSE | High | IsMasterP | AAEL0224 | 1:241845 | 0 | 12 | FALSE | 9 | 27 | 9 | 1504 | 1636 | 549 |
| 7,12296E | FALSE | High | IsMasterP | AAEL0197 | 2:351508 | 0 | 22 | FALSE | 14 | 25 | 14 | 1154 | 1249 | 496 |
| -2,59626E | FALSE | High | IsMasterP | AAEL0075 | 2:248940 | 0 | 12 | FALSE | 11 | 26 | 11 | 1386 | 1553 | 667 |
| -8,67693E | FALSE | High | IsMasterP | AAEL0119 | 2:301748 | 0 | 56 | FALSE | 5 | 18 | 5 | 101 | 116 | 721 |
| 8,37747E | FALSE | High | IsMasterP | AAEL0254 | 2:474158 | 0 | 5 | FALSE | 17 | 26 | 17 | 5813 | 6743 | 571 |
| -8,57103E | FALSE | High | IsMasterP | AAEL0234 | 1:111362 | 0 | 77 | FALSE | 12 | 25 | 12 | 227 | 250 | 692 |
| -7,23134E | FALSE | High | IsMasterP | AAEL0251 | 1:716888 | 0 | 30 | FALSE | 4 | 18 | 4 | 277 | 323 | 816 |
| 8,29241E | FALSE | High | IsMasterP | AAEL0056 | 1:473531 | 0 | 30 | FALSE | 9 | 24 | 9 | 412 | 435 | 778 |
| -7,6793E+ | FALSE | High | IsMasterP | AAEL0043 | 3:508851 | 0 | 11 | FALSE | 13 | 24 | 13 | 1642 | 1838 | 506 |
| 1,68082E | FALSE | High | IsMasterP | AAEL0131 | 3:371319 | 0 | 37 | FALSE | 10 | 25 | 10 | 289 | 310 | 888 |
| -8,17328E | FALSE | High | IsMasterP | AAEL0136 | 3:195029 | 0 | 41 | FALSE | 10 | 24 | 10 | 390 | 426 | 835 |
| -3,48183E | FALSE | High | IsMasterP | AAEL0085 | 3:222567 | 0 | 30 | FALSE | 13 | 25 | 12 | 669 | 749 | 718 |
| -4,46441E | FALSE | High | IsMasterP | AAEL0275 | 2:314520 | 0 | 26 | FALSE | 11 | 29 | 11 | 418 | 473 | 534 |
| 5,06581E | FALSE | High | IsMasterP | AAEL0151 | 2:123165 | 0 | 31 | FALSE | 8 | 20 | 8 | 505 | 554 | 662 |
| 2,39667E | FALSE | High | IsMasterP | AAEL0003 | 2:368910 | 0 | 40 | FALSE | 10 | 23 | 10 | 343 | 385 | 597 |
| -1,49842E | FALSE | High | IsMasterP | AAEL0208 | 2:241469 | 0 | 8 | FALSE | 16 | 23 | 16 | 3472 | 3939 | 879 |
| -3,25944E | FALSE | High | IsMasterP | AAEL0182 | 2:327991 | 0 | 14 | FALSE | 17 | 30 | 17 | 1507 | 1709 | 828 |
| 4,53665E | FALSE | High | IsMasterP | AAEL0214 | NIGP0100 | 0 | 11 | FALSE | 7 | 23 | 3 | 955 | 1069 | 717 |
| 5,13705E | FALSE | High | IsMasterP | AAEL0099 | 1:206625 | 0 | 70 | FALSE | 7 | 20 | 7 | 159 | 180 | 679 |
| -1,76285E | FALSE | High | IsMasterP | AAEL0056 | 3:160188 | 0 | 34 | FALSE | 11 | 29 | 11 | 238 | 270 | 885 |
| 8,60962E | FALSE | High | IsMasterP | AAEL0094 | 3:380841 | 0 | 33 | FALSE | 12 | 28 | 12 | 448 | 522 | 618 |
| -6,3413E+ | FALSE | High | IsMasterP | AAEL0086 | 2:214939 | 0 | 37 | FALSE | 9 | 27 | 8 | 354 | 404 | 553 |
| 7,46965E | FALSE | High | IsMasterP | AAEL0107 | 2:305242 | 0 | 19 | FALSE | 9 | 24 | 9 | 612 | 659 | 947 |
| -2,35286E | FALSE | High | IsMasterP | AAEL0031 | 1:190181 | 0 | 60 | FALSE | 9 | 23 | 9 | 165 | 191 | 547 |
| -1,99908E | FALSE | High | IsMasterP | AAEL0030 | 3:399921 | 0 | 44 | FALSE | 6 | 26 | 6 | 95 | 106 | 983 |
| 7,61118E | FALSE | High | IsMasterP | AAEL0146 | 3:368502 | 0 | 19 | FALSE | 10 | 28 | 1 | 541 | 624 | 743 |
| -5,4912E+ | FALSE | High | IsMasterP | AAEL0028 | 1:255059 | 0 | 18 | FALSE | 11 | 26 | 11 | 805 | 938 | 771 |
| -1,08711E | FALSE | High | IsMasterP | AAEL0041 | 1:156739 | 0 | 32 | FALSE | 5 | 20 | 5 | 264 | 284 | 463 |
| 7,28541E | FALSE | High | IsMasterP | AAEL0063 | 1:101377 | 0 | 21 | FALSE | 8 | 20 | 8 | 694 | 770 | 848 |
| -9,13953E | FALSE | High | IsMasterP | AAEL0256 | 1:138065 | 0 | 17 | FALSE | 9 | 21 | 9 | 1109 | 1225 | 677 |

| | | | | | | | | | | | | | | |
|---|---|---|---|---|---|---|---|---|---|---|---|---|---|---|
| 4,1885E+1 | FALSE | High | IsMasterP | AAEL0114 | 3:3290553 | 0 | 50 | FALSE | 5 | 26 | 1 | 136 | 153 | 1127 |
| -7,66022E | FALSE | High | IsMasterP | AAEL0118 | 2:1873031 | 0 | 67 | FALSE | 7 | 24 | 7 | 143 | 163 | 677 |
| 3,14757E+ | FALSE | High | IsMasterP | AAEL0107 | 1:1786671 | 0 | 24 | FALSE | 9 | 24 | 7 | 594 | 656 | 947 |
| 2,058E+18 | FALSE | High | IsMasterP | AAEL0100 | 2:1105776 | 0 | 21 | FALSE | 13 | 22 | 13 | 987 | 1078 | 689 |
| 6,07968E+ | FALSE | High | IsMasterP | AAEL0061 | 3:1405089 | 0 | 32 | FALSE | 8 | 21 | 8 | 410 | 479 | 624 |
| 1,29013E+ | FALSE | High | IsMasterP | AAEL0034 | 3:3482774 | 0 | 16 | FALSE | 7 | 28 | 7 | 578 | 653 | 609 |
| 6,78245E+ | FALSE | High | IsMasterP | AAEL0225 | 1:2832129 | 0 | 51 | FALSE | 10 | 24 | 10 | 336 | 378 | 548 |
| 2,1011E+1 | FALSE | High | IsMasterP | AAEL0016 | 2:8675428 | 0 | 15 | FALSE | 3 | 22 | 3 | 223 | 250 | 477 |
| 7,89356E+ | FALSE | High | IsMasterP | AAEL0060 | 3:3335834 | 0 | 55 | FALSE | 14 | 24 | 14 | 395 | 434 | 938 |
| 2,91934E+ | FALSE | High | IsMasterP | AAEL0277 | 3:2419355 | 0 | 20 | FALSE | 14 | 25 | 14 | 1091 | 1240 | 756 |
| -5,26168E | FALSE | High | IsMasterP | AAEL0138 | 3:2019089 | 0 | 18 | FALSE | 7 | 18 | 7 | 740 | 785 | 711 |
| 6,31937E+ | FALSE | High | IsMasterP | AAEL0107 | 2:3048274 | 0 | 42 | FALSE | 9 | 24 | 9 | 312 | 359 | 605 |
| -2,33587E | FALSE | High | IsMasterP | AAEL0130 | 2:2535148 | 0 | 28 | FALSE | 8 | 21 | 8 | 457 | 522 | 860 |
| 8,3552E+1 | FALSE | High | IsMasterP | AAEL0034 | 3:3501620 | 0 | 54 | FALSE | 10 | 31 | 10 | 148 | 167 | 1023 |
| 5,37366E+ | FALSE | High | IsMasterP | AAEL0044 | 3:2051826 | 0 | 53 | FALSE | 8 | 21 | 8 | 226 | 254 | 832 |
| -1,42621E | FALSE | High | IsMasterP | AAEL0182 | 2:1366353 | 0 | 22 | FALSE | 9 | 21 | 9 | 742 | 865 | 555 |
| 3,48412E+ | FALSE | High | IsMasterP | AAEL0123 | 2:1030437 | 0 | 39 | FALSE | 12 | 23 | 12 | 429 | 468 | 777 |
| -8,60391E | FALSE | High | IsMasterP | AAEL0213 | 1:1217908 | 0 | 35 | FALSE | 5 | 24 | 5 | 208 | 238 | 658 |
| 1,72939E+ | FALSE | High | IsMasterP | AAEL0139 | 3:1061903 | 0 | 23 | FALSE | 8 | 22 | 8 | 583 | 634 | 829 |
| 2,01631E+ | FALSE | High | IsMasterP | AAEL0170 | 2:3059521 | 0 | 17 | FALSE | 10 | 22 | 10 | 997 | 1038 | 426 |
| 1,02686E+ | FALSE | High | IsMasterP | AAEL0197 | 1:5549868 | 0 | 17 | FALSE | 8 | 22 | 8 | 977 | 1088 | 602 |
| -4,85072E | FALSE | High | IsMasterP | AAEL0133 | 3:1731485 | 0 | 38 | FALSE | 10 | 27 | 9 | 381 | 436 | 756 |
| -8,56852E | FALSE | High | IsMasterP | AAEL0039 | 1:2505839 | 0 | 34 | FALSE | 9 | 23 | 9 | 296 | 309 | 1062 |
| -4,7884E+ | FALSE | High | IsMasterP | AAEL0070 | 1:1281471 | 0 | 31 | FALSE | 10 | 23 | 10 | 464 | 525 | 473 |
| 4,28265E+ | FALSE | High | IsMasterP | AAEL0010 | 3:5518706 | 0 | 34 | FALSE | 6 | 22 | 6 | 335 | 367 | 573 |
| 2,71447E+ | FALSE | High | IsMasterP | AAEL0025 | 3:2566241 | 0 | 29 | FALSE | 12 | 22 | 12 | 762 | 839 | 533 |
| -6,30456E | FALSE | High | IsMasterP | AAEL0001 | 3:1096613 | 0 | 47 | FALSE | 10 | 25 | 10 | 273 | 321 | 901 |
| -2,53938E | FALSE | High | IsMasterP | AAEL0072 | 2:2537195 | 0 | 84 | FALSE | 5 | 18 | 5 | 115 | 127 | 545 |
| 5,25648E+ | FALSE | High | IsMasterP | AAEL0050 | 1:2982749 | 0 | 24 | FALSE | 8 | 21 | 8 | 511 | 574 | 629 |
| 8,33498E+ | FALSE | High | IsMasterP | AAEL0040 | 1:1357692 | 0 | 38 | FALSE | 10 | 21 | 10 | 397 | 434 | 481 |
| -8,78731E | FALSE | High | IsMasterP | AAEL0133 | 1:1563671 | 0 | 65 | FALSE | 11 | 21 | 11 | 313 | 355 | 646 |

| | | | | | | | | | | | | | | |
|---|---|---|---|---|---|---|---|---|---|---|---|---|---|---|
| -2,8031E+ | FALSE | High | IsMasterP | AAEL0255 | 2:294856 | 0 | 19 | FALSE | 13 | 23 | 13 | 1012 | 1175 | 944 |
| -2,34726E | FALSE | High | IsMasterP | AAEL0041 | 1:158769 | 0 | 45 | FALSE | 12 | 21 | 12 | 469 | 509 | 749 |
| 4,66432E | FALSE | High | IsMasterP | AAEL0108 | 3:318912 | 0 | 27 | FALSE | 5 | 21 | 5 | 298 | 328 | 541 |
| -2,01278E | FALSE | High | IsMasterP | AAEL0060 | 1:184072 | 0 | 15 | FALSE | 10 | 24 | 10 | 900 | 1040 | 772 |
| 8,81504E | FALSE | High | IsMasterP | AAEL0057 | 2:437262 | 0 | 19 | FALSE | 7 | 24 | 7 | 444 | 515 | 545 |
| -2,35374E | FALSE | High | IsMasterP | AAEL0202 | NIGP0100 | 0 | 14 | FALSE | 5 | 27 | 4 | 226 | 231 | 1059 |
| -5,1501E+ | FALSE | High | IsMasterP | AAEL0007 | 3:242683 | 0 | 38 | FALSE | 10 | 20 | 10 | 411 | 463 | 755 |
| 3,74071E | FALSE | High | IsMasterP | AAEL0054 | 2:296919 | 0 | 29 | FALSE | 9 | 19 | 9 | 665 | 735 | 712 |
| 3,88365E | FALSE | High | IsMasterP | AAEL0069 | 3:143904 | 0 | 44 | FALSE | 10 | 19 | 10 | 368 | 404 | 840 |
| -6,07397E | FALSE | High | IsMasterP | AAEL0035 | 2:256215 | 0 | 35 | FALSE | 9 | 20 | 9 | 513 | 558 | 548 |
| 1,21943E | FALSE | High | IsMasterP | AAEL0058 | 3:220098 | 0 | 11 | FALSE | 12 | 22 | 12 | 2338 | 2551 | 891 |
| -2,93353E | FALSE | High | IsMasterP | AAEL0090 | 2:270330 | 0 | 73 | FALSE | 7 | 19 | 7 | 153 | 168 | 496 |
| 6,1319E+ | FALSE | High | IsMasterP | AAEL0098 | 3:122199 | 0 | 25 | FALSE | 7 | 22 | 7 | 390 | 434 | 529 |
| -8,68117E | FALSE | High | IsMasterP | AAEL0226 | 3:929645 | 0 | 14 | FALSE | 11 | 21 | 5 | 1153 | 1271 | 613 |
| 2,8346E+ | FALSE | High | IsMasterP | AAEL0084 | 2:189815 | 0 | 29 | FALSE | 7 | 23 | 7 | 229 | 257 | 596 |
| -5,22372E | FALSE | High | IsMasterP | AAEL0229 | 1:284636 | 0 | 53 | FALSE | 8 | 26 | 8 | 118 | 135 | 492 |
| -4,67227E | FALSE | High | IsMasterP | AAEL0122 | 3:170268 | 0 | 49 | FALSE | 6 | 18 | 6 | 134 | 150 | 933 |
| 7,26048E | FALSE | High | IsMasterP | AAEL0039 | 3:267056 | 0 | 10 | FALSE | 8 | 25 | 8 | 1027 | 1192 | 788 |
| 1,95568E | FALSE | High | IsMasterP | AAEL0119 | 1:285531 | 0 | 20 | FALSE | 7 | 21 | 7 | 558 | 618 | 526 |
| -7,35286E | FALSE | High | IsMasterP | AAEL0118 | 3:360870 | 0 | 31 | FALSE | 9 | 25 | 9 | 387 | 422 | 533 |
| -8,26076E | FALSE | High | IsMasterP | AAEL0197 | 1:220270 | 0 | 22 | FALSE | 7 | 19 | 7 | 546 | 613 | 765 |
| -1,80412E | FALSE | High | IsMasterP | AAEL0137 | 2:122572 | 0 | 50 | FALSE | 10 | 20 | 10 | 289 | 330 | 544 |
| 5,80303E | FALSE | High | IsMasterP | AAEL0124 | 1:271598 | 0 | 15 | FALSE | 13 | 23 | 13 | 1188 | 1335 | 540 |
| -1,49952E | FALSE | High | IsMasterP | AAEL0020 | 2:426594 | 0 | 20 | FALSE | 8 | 25 | 8 | 665 | 745 | 473 |
| 1,81963E | FALSE | High | IsMasterP | AAEL0111 | 3:194105 | 0 | 24 | FALSE | 9 | 21 | 9 | 567 | 620 | 824 |
| 7,47727E | FALSE | High | IsMasterP | AAEL0138 | 1:263491 | 0 | 31 | FALSE | 9 | 21 | 9 | 490 | 557 | 728 |
| 2,97021E | FALSE | High | IsMasterP | AAEL0054 | 1:804937 | 0 | 8 | FALSE | 8 | 20 | 8 | 1428 | 1633 | 560 |
| -3,11023E | FALSE | High | IsMasterP | AAEL0077 | 3:255581 | 0 | 47 | FALSE | 8 | 25 | 8 | 124 | 136 | 718 |
| -2,73661E | FALSE | High | IsMasterP | AAEL0181 | 2:160663 | 0 | 14 | FALSE | 10 | 22 | 10 | 1113 | 1211 | 928 |
| -4,34741E | FALSE | High | IsMasterP | AAEL0146 | 2:157491 | 0 | 65 | FALSE | 7 | 22 | 7 | 152 | 173 | 560 |
| -5,0954E+ | FALSE | High | IsMasterP | AAEL0173 | 2:300345 | 0 | 11 | FALSE | 11 | 20 | 11 | 1887 | 2085 | 758 |

| | | | | | | | | | | | | | | |
|---|---|---|---|---|---|---|---|---|---|---|---|---|---|---|
| 2,14259E | FALSE | High | IsMasterP | AAEL0033 | 2:461391 | 0 | 74 | FALSE | 5 | 23 | 5 | 127 | 141 | 785 |
| -5,20972E | FALSE | High | IsMasterP | AAEL0113 | 1:165393 | 0 | 44 | FALSE | 8 | 20 | 7 | 331 | 367 | 661 |
| 8,13856E | FALSE | High | IsMasterP | AAEL0076 | 3:347316 | 0 | 18 | FALSE | 3 | 23 | 1 | 245 | 267 | 657 |
| 3,50365E | FALSE | High | IsMasterP | AAEL0018 | 2:275154 | 0 | 45 | FALSE | 8 | 20 | 8 | 165 | 194 | 709 |
| 2,03589E | FALSE | High | IsMasterP | AAEL0169 | 1:209999 | 0 | 16 | FALSE | 10 | 21 | 10 | 715 | 776 | 463 |
| -5,98693E | FALSE | High | IsMasterP | AAEL0138 | 1:178862 | 0 | 63 | FALSE | 5 | 21 | 5 | 86 | 95 | 717 |
| 4,11286E | FALSE | High | IsMasterP | AAEL0128 | 2:105589 | 0 | 47 | FALSE | 6 | 21 | 6 | 221 | 247 | 445 |
| -8,81448E | FALSE | High | IsMasterP | AAEL0035 | 1:693723 | 0 | 35 | FALSE | 7 | 24 | 7 | 151 | 172 | 1051 |
| -2,21994E | FALSE | High | IsMasterP | AAEL0058 | 3:220314 | 0 | 40 | FALSE | 9 | 20 | 9 | 380 | 425 | 873 |
| -2,41701E | FALSE | High | IsMasterP | AAEL0030 | 3:399272 | 0 | 42 | FALSE | 9 | 20 | 9 | 391 | 444 | 756 |
| 2,61274E | FALSE | High | IsMasterP | AAEL0059 | 1:257747 | 0 | 45 | FALSE | 4 | 25 | 4 | 114 | 127 | 870 |
| 3,45302E | FALSE | High | IsMasterP | AAEL0086 | 2:805660 | 0 | 39 | FALSE | 11 | 23 | 11 | 311 | 352 | 535 |
| 4,513E+1 | FALSE | High | IsMasterP | AAEL0028 | 1:255918 | 0 | 31 | FALSE | 5 | 18 | 5 | 359 | 400 | 549 |
| -6,78916E | FALSE | High | IsMasterP | AAEL0113 | 2:331491 | 0 | 51 | FALSE | 10 | 18 | 10 | 422 | 466 | 548 |
| -5,4467E+ | FALSE | High | IsMasterP | AAEL0022 | 2:430736 | 0 | 28 | FALSE | 11 | 17 | 11 | 735 | 803 | 618 |
| 7,6133E+ | FALSE | High | IsMasterP | AAEL0266 | 3:362014 | 0 | 21 | FALSE | 6 | 23 | 6 | 394 | 438 | 623 |
| -8,54498E | FALSE | High | IsMasterP | AAEL0080 | 2:364723 | 0 | 65 | FALSE | 13 | 23 | 13 | 315 | 351 | 627 |
| -7,38394E | FALSE | High | IsMasterP | AAEL0000 | 3:102916 | 0 | 35 | FALSE | 7 | 25 | 7 | 113 | 133 | 1134 |
| 8,45752E | FALSE | High | IsMasterP | AAEL0083 | 2:274184 | 0 | 27 | FALSE | 7 | 24 | 7 | 153 | 173 | 1122 |
| -6,91844E | FALSE | High | IsMasterP | AAEL0042 | 3:117434 | 0 | 27 | FALSE | 11 | 19 | 11 | 695 | 795 | 668 |
| -8,75836E | FALSE | High | IsMasterP | AAEL0103 | 3:409314 | 0 | 22 | FALSE | 8 | 21 | 8 | 534 | 585 | 702 |
| 5,78258E | FALSE | High | IsMasterP | AAEL0212 | 3:103495 | 0 | 19 | FALSE | 11 | 23 | 11 | 679 | 762 | 928 |
| 3,72057E | FALSE | High | IsMasterP | AAEL0262 | 2:174221 | 0 | 24 | FALSE | 5 | 21 | 5 | 261 | 273 | 1078 |
| -7,56648E | FALSE | High | IsMasterP | AAEL0011 | 3:406460 | 0 | 20 | FALSE | 11 | 19 | 11 | 894 | 998 | 640 |
| -5,43034E | FALSE | High | IsMasterP | AAEL0116 | 3:214210 | 0 | 34 | FALSE | 10 | 24 | 10 | 433 | 484 | 625 |
| 8,91487E | FALSE | High | IsMasterP | AAEL0095 | 3:344888 | 0 | 52 | FALSE | 8 | 25 | 8 | 120 | 134 | 1013 |
| -6,67505E | FALSE | High | IsMasterP | AAEL0130 | 1:317072 | 0 | 36 | FALSE | 10 | 17 | 10 | 504 | 556 | 863 |
| -9,39758E | FALSE | High | IsMasterP | AAEL0054 | 1:247122 | 0 | 44 | FALSE | 7 | 21 | 5 | 321 | 357 | 475 |
| 8,04579E | FALSE | High | IsMasterP | AAEL0016 | 1:137424 | 0 | 24 | FALSE | 11 | 20 | 11 | 748 | 831 | 702 |
| -1,06167E | FALSE | High | IsMasterP | AAEL0093 | 1:199746 | 0 | 35 | FALSE | 8 | 27 | 8 | 215 | 243 | 601 |
| 8,76411E | FALSE | High | IsMasterP | AAEL0080 | 3:148505 | 0 | 18 | FALSE | 7 | 20 | 7 | 722 | 832 | 602 |

| | | | | | | | | | | | | | | |
|---|---|---|---|---|---|---|---|---|---|---|---|---|---|---|
| 5,0789E+ | FALSE | High | IsMasterP | AAEL0057 | 3:114091 | 0 | 25 | FALSE | 9 | 21 | 9 | 607 | 668 | 683 |
| 8,38982E | FALSE | High | IsMasterP | AAEL0112 | 3:239485 | 0 | 73 | FALSE | 5 | 19 | 5 | 91 | 108 | 963 |
| 5,35862E | FALSE | High | IsMasterP | AAEL0214 | NIGP0100 | 0 | 50 | FALSE | 9 | 22 | 9 | 177 | 212 | 1061 |
| -7,95575E | FALSE | High | IsMasterP | AAEL0132 | 2:186074 | 0 | 17 | FALSE | 7 | 21 | 7 | 580 | 634 | 544 |
| -7,32859E | FALSE | High | IsMasterP | AAEL0010 | 1:281620 | 0 | 40 | FALSE | 8 | 19 | 8 | 225 | 250 | 553 |
| -4,56921E | FALSE | High | IsMasterP | AAEL0056 | 3:160049 | 0 | 31 | FALSE | 10 | 18 | 10 | 628 | 710 | 646 |
| -3,7654E+ | FALSE | High | IsMasterP | AAEL0138 | 1:103529 | 0 | 21 | FALSE | 8 | 23 | 8 | 413 | 476 | 615 |
| -2,54885E | FALSE | High | IsMasterP | AAEL0130 | 2:349156 | 0 | 33 | FALSE | 6 | 21 | 6 | 204 | 233 | 484 |
| -4,80163E | FALSE | High | IsMasterP | AAEL0034 | 3:349299 | 0 | 35 | FALSE | 10 | 20 | 10 | 389 | 438 | 865 |
| -5,01565E | FALSE | High | IsMasterP | AAEL0197 | 3:369484 | 0 | 37 | FALSE | 8 | 16 | 7 | 490 | 528 | 821 |
| 9,12357E | FALSE | High | IsMasterP | AAEL0195 | 3:143356 | 0 | 7 | FALSE | 7 | 19 | 7 | 1519 | 1664 | 683 |
| 7,85788E | FALSE | High | IsMasterP | AAEL0097 | 3:155608 | 0 | 35 | FALSE | 11 | 23 | 11 | 329 | 370 | 715 |
| 5,20428E | FALSE | High | IsMasterP | AAEL0194 | 2:846183 | 0 | 62 | FALSE | 6 | 17 | 6 | 187 | 212 | 658 |
| -5,26023E | FALSE | High | IsMasterP | AAEL0006 | 3:395933 | 0 | 30 | FALSE | 5 | 18 | 5 | 351 | 392 | 809 |
| 8,17416E | FALSE | High | IsMasterP | AAEL0137 | 3:320125 | 0 | 43 | FALSE | 9 | 20 | 9 | 332 | 395 | 491 |
| -6,70926E | FALSE | High | IsMasterP | AAEL0150 | 1:289815 | 0 | 33 | FALSE | 7 | 19 | 7 | 299 | 342 | 677 |
| 9,01168E | FALSE | High | IsMasterP | AAEL0194 | 3:404619 | 0 | 14 | FALSE | 12 | 20 | 12 | 1199 | 1325 | 714 |
| -7,65671E | FALSE | High | IsMasterP | AAEL0139 | 2:102055 | 0 | 29 | FALSE | 7 | 21 | 7 | 316 | 353 | 503 |
| -8,41985E | FALSE | High | IsMasterP | AAEL0040 | 3:166372 | 0 | 28 | FALSE | 9 | 17 | 8 | 632 | 695 | 654 |
| 8,82305E | FALSE | High | IsMasterP | AAEL0186 | 2:625996 | 0 | 5 | FALSE | 10 | 23 | 10 | 3305 | 3705 | 848 |
| -1,88815E | FALSE | High | IsMasterP | AAEL0273 | 3:250492 | 0 | 30 | FALSE | 6 | 19 | 6 | 295 | 334 | 734 |
| 6,08066E | FALSE | High | IsMasterP | AAEL0012 | 2:413266 | 0 | 39 | FALSE | 3 | 18 | 3 | 123 | 137 | 848 |
| -7,93216E | FALSE | High | IsMasterP | AAEL0003 | 2:368844 | 0 | 34 | FALSE | 6 | 16 | 6 | 437 | 474 | 625 |
| 2,40529E | FALSE | High | IsMasterP | AAEL0057 | 3:204066 | 0 | 9 | FALSE | 11 | 20 | 11 | 2099 | 2306 | 834 |
| 3,90581E | FALSE | High | IsMasterP | AAEL0031 | 1:193103 | 0 | 27 | FALSE | 8 | 21 | 8 | 365 | 421 | 550 |
| -1,08641E | FALSE | High | IsMasterP | AAEL0008 | 3:363783 | 0 | 40 | FALSE | 11 | 21 | 11 | 403 | 455 | 810 |
| -6,02801E | FALSE | High | IsMasterP | AAEL0030 | 3:388147 | 0 | 23 | FALSE | 6 | 20 | 6 | 349 | 382 | 835 |
| -3,69417E | FALSE | High | IsMasterP | AAEL0050 | 1:298089 | 0 | 40 | FALSE | 10 | 20 | 10 | 370 | 406 | 535 |
| 3,89834E | FALSE | High | IsMasterP | AAEL0270 | 2:351649 | 0 | 46 | FALSE | 9 | 21 | 9 | 222 | 247 | 613 |
| -6,14673E | FALSE | High | IsMasterP | AAEL0075 | 2:327264 | 0 | 34 | FALSE | 11 | 20 | 11 | 458 | 514 | 553 |
| 3,44537E | FALSE | High | IsMasterP | AAEL0182 | 1:111377 | 0 | 12 | FALSE | 8 | 18 | 8 | 1521 | 1656 | 781 |

| | | | | | | | | | | | | | |
|---|---|---|---|---|---|---|---|---|---|---|---|---|---|
| -9,17195E | FALSE | High | IsMasterP AAEL0040 | 3:4017353 | 0 | 42 | FALSE | 10 | 20 | 10 | 323 | 371 | 574 |
| 8,15156E+ | FALSE | High | IsMasterP AAEL0270 | 3:2440301 | 0 | 25 | FALSE | 9 | 24 | 8 | 363 | 418 | 1163 |
| -3,22198E | FALSE | High | IsMasterP AAEL0065 | 2:4686282 | 0 | 22 | FALSE | 9 | 19 | 9 | 576 | 661 | 662 |
| -7,1386E+ | FALSE | High | IsMasterP AAEL0073 | 2:3455153 | 0 | 67 | FALSE | 6 | 21 | 6 | 166 | 180 | 554 |
| -5,70807E | FALSE | High | IsMasterP AAEL0147 | 2:3267561 | 0 | 8 | FALSE | 9 | 19 | 9 | 1798 | 1845 | 888 |
| 7,05178E+ | FALSE | High | IsMasterP AAEL0055 | 1:9027989 | 0 | 30 | FALSE | 9 | 22 | 9 | 461 | 512 | 673 |
| 1,09443E+ | FALSE | High | IsMasterP AAEL0145 | 1:2243096 | 0 | 39 | FALSE | 8 | 18 | 8 | 320 | 357 | 892 |
| 4,47627E+ | FALSE | High | IsMasterP AAEL0027 | 3:1299782 | 0 | 35 | FALSE | 9 | 21 | 9 | 409 | 449 | 717 |
| 8,69813E+ | FALSE | High | IsMasterP AAEL0240 | 1:2133406 | 0 | 55 | FALSE | 2 | 15 | 2 | 88 | 99 | 481 |
| -8,33278E | FALSE | High | IsMasterP AAEL0218 | 3:6980432 | 0 | 38 | FALSE | 9 | 19 | 9 | 369 | 418 | 955 |
| -3,89891E | FALSE | High | IsMasterP AAEL0290 | 3:3686120 | 0 | 17 | FALSE | 7 | 24 | 4 | 511 | 588 | 859 |
| 6,3697E+1 | FALSE | High | IsMasterP AAEL0248 | 3:3146644 | 0 | 19 | FALSE | 9 | 18 | 9 | 946 | 1061 | 525 |
| 5,90588E+ | FALSE | High | IsMasterP AAEL0120 | 1:1043741 | 0 | 37 | FALSE | 8 | 19 | 8 | 342 | 405 | 793 |
| 3,64545E+ | FALSE | High | IsMasterP AAEL0124 | 2:2189522 | 0 | 23 | FALSE | 8 | 17 | 8 | 511 | 583 | 623 |
| -2,59978E | FALSE | High | IsMasterP AAEL0040 | 1:1359737 | 0 | 15 | FALSE | 10 | 22 | 10 | 796 | 879 | 473 |
| -3,26165E | FALSE | High | IsMasterP AAEL0140 | 2:1323197 | 0 | 29 | FALSE | 6 | 16 | 6 | 409 | 464 | 534 |
| -4,04285E | FALSE | High | IsMasterP AAEL0057 | 3:1154846 | 0 | 53 | FALSE | 9 | 21 | 9 | 188 | 215 | 868 |
| 8,60376E+ | FALSE | High | IsMasterP AAEL0122 | 2:1450579 | 0 | 40 | FALSE | 11 | 19 | 11 | 465 | 521 | 574 |
| 3,1365E+1 | FALSE | High | IsMasterP AAEL0078 | 3:3815640 | 0 | 16 | FALSE | 8 | 19 | 8 | 661 | 761 | 872 |
| 6,69558E+ | FALSE | High | IsMasterP AAEL0135 | 2:6186352 | 0 | 26 | FALSE | 3 | 16 | 3 | 125 | 136 | 919 |
| 8,96668E+ | FALSE | High | IsMasterP AAEL0091 | 3:1760806 | 0 | 11 | FALSE | 7 | 19 | 7 | 919 | 1027 | 526 |
| 7,36092E+ | FALSE | High | IsMasterP AAEL0250 | 1:1466018 | 0 | 33 | FALSE | 8 | 23 | 8 | 328 | 363 | 530 |
| 4,51689E+ | FALSE | High | IsMasterP AAEL0121 | 2:2745243 | 0 | 18 | FALSE | 8 | 20 | 8 | 712 | 792 | 615 |
| 5,32766E+ | FALSE | High | IsMasterP AAEL0062 | 2:4907414 | 0 | 49 | FALSE | 7 | 15 | 7 | 279 | 293 | 872 |
| -4,08328E | FALSE | High | IsMasterP AAEL0196 | 2:2055348 | 0 | 31 | FALSE | 6 | 16 | 6 | 355 | 396 | 1054 |
| 5,66175E+ | FALSE | High | IsMasterP AAEL0204 | 2:6346905 | 0 | 3 | FALSE | 10 | 21 | 10 | 5515 | 6266 | 497 |
| 4,07442E+ | FALSE | High | IsMasterP AAEL0084 | NIGP0100 | 0 | 38 | FALSE | 7 | 20 | 7 | 200 | 227 | 815 |
| 4,21738E+ | FALSE | High | IsMasterP AAEL0037 | 1:2478678 | 0 | 12 | FALSE | 8 | 23 | 8 | 868 | 983 | 677 |
| -1,00271E | FALSE | High | IsMasterP AAEL0005 | 3:1202484 | 0 | 18 | FALSE | 13 | 20 | 13 | 1298 | 1406 | 578 |
| -7,58501E | FALSE | High | IsMasterP AAEL0071 | 2:1499497 | 0 | 47 | FALSE | 6 | 22 | 6 | 120 | 141 | 976 |
| -2,95734E | FALSE | High | IsMasterP AAEL0025 | 1:2262604 | 0 | 20 | FALSE | 8 | 18 | 8 | 707 | 780 | 526 |

| | | | | | | | | | | | | | | |
|---|---|---|---|---|---|---|---|---|---|---|---|---|---|---|
| 8,18323E- | FALSE | High | IsMasterP | AAEL0147 | 2:169822( | 0 | 26 | FALSE | 9 | 20 | 9 | 537 | 634 | 625 |
| -5,79204E | FALSE | High | IsMasterP | AAEL0044 | 3:204315( | 0 | 25 | FALSE | 7 | 17 | 7 | 477 | 535 | 706 |
| -2,73395E | FALSE | High | IsMasterP | AAEL0123 | 1:284526! | 0 | 47 | FALSE | 9 | 19 | 9 | 300 | 329 | 543 |
| 2,47999E- | FALSE | High | IsMasterP | AAEL0006 | 3:395591: | 0 | 13 | FALSE | 7 | 21 | 7 | 625 | 699 | 585 |
| 6,70995E- | FALSE | High | IsMasterP | AAEL0290 | 3:119541! | 0 | 29 | FALSE | 5 | 20 | 5 | 206 | 236 | 693 |
| 4,63163E- | FALSE | High | IsMasterP | AAEL0095 | 3:223747( | 0 | 45 | FALSE | 6 | 18 | 6 | 219 | 238 | 809 |
| 6,963E+1{ | FALSE | High | IsMasterP | AAEL0073 | 2:233265 | 0 | 37 | FALSE | 7 | 18 | 7 | 329 | 360 | 646 |
| 4,10001E- | FALSE | High | IsMasterP | AAEL0079 | 2:365780 | 0 | 22 | FALSE | 6 | 17 | 6 | 563 | 625 | 408 |
| -7,54232E | FALSE | High | IsMasterP | AAEL0077 | 3:255383( | 0 | 39 | FALSE | 9 | 25 | 9 | 161 | 185 | 1052 |
| -8,32884E | FALSE | High | IsMasterP | AAEL0009 | 3:335494: | 0 | 13 | FALSE | 2 | 15 | 2 | 340 | 376 | 604 |
| -9,16599E | FALSE | High | IsMasterP | AAEL0078 | 2:474316: | 0 | 61 | FALSE | 8 | 17 | 8 | 256 | 274 | 784 |
| -8,53255E | FALSE | High | IsMasterP | AAEL0072 | 3:195590( | 0 | 14 | FALSE | 8 | 19 | 8 | 881 | 980 | 827 |
| 6,00946E- | FALSE | High | IsMasterP | AAEL0040 | 1:135679! | 0 | 46 | FALSE | 5 | 12 | 5 | 332 | 354 | 881 |
| -1,95265E | FALSE | High | IsMasterP | AAEL0059 | 3:173937( | 0 | 39 | FALSE | 4 | 16 | 4 | 174 | 202 | 505 |
| 3,50264E- | FALSE | High | IsMasterP | AAEL0226 | 2:197323: | 0 | 9 | FALSE | 8 | 16 | 8 | 1804 | 1955 | 723 |
| -4,96775E | FALSE | High | IsMasterP | AAEL0150 | 1:153238 | 0 | 59 | FALSE | 3 | 22 | 3 | 95 | 109 | 594 |
| 8,08463E- | FALSE | High | IsMasterP | AAEL0078 | 3:151835: | 0 | 10 | FALSE | 8 | 19 | 8 | 985 | 1077 | 796 |
| -1,82339E | FALSE | High | IsMasterP | AAEL0255 | 2:651807! | 0 | 49 | FALSE | 8 | 18 | 8 | 257 | 289 | 758 |
| 4,31046E- | FALSE | High | IsMasterP | AAEL0081 | 3:107372 | 0 | 67 | FALSE | 4 | 20 | 4 | 89 | 102 | 774 |
| -9,10935E | FALSE | High | IsMasterP | AAEL0127 | 2:756821: | 0 | 11 | FALSE | 8 | 20 | 8 | 1375 | 1477 | 723 |
| 8,99238E- | FALSE | High | IsMasterP | AAEL0182 | 2:231510! | 0 | 9 | FALSE | 9 | 20 | 9 | 1788 | 1950 | 458 |
| 3,52476E- | FALSE | High | IsMasterP | AAEL0058 | 1:225857 | 0 | 30 | FALSE | 8 | 17 | 8 | 358 | 400 | 667 |
| -2,71392E | FALSE | High | IsMasterP | AAEL0122 | 2:384367( | 0 | 10 | FALSE | 10 | 18 | 10 | 1793 | 2059 | 723 |
| -8,25941E | FALSE | High | IsMasterP | AAEL0036 | 3:653118! | 0 | 39 | FALSE | 4 | 14 | 4 | 245 | 261 | 468 |
| -5,13883E | FALSE | High | IsMasterP | AAEL0028 | 1:304725 | 0 | 33 | FALSE | 8 | 17 | 8 | 306 | 348 | 609 |
| -5,81622E | FALSE | High | IsMasterP | AAEL0100 | 3:209206( | 0 | 55 | FALSE | 10 | 16 | 10 | 286 | 325 | 581 |
| -7,44822E | FALSE | High | IsMasterP | AAEL0198 | 2:352692( | 0 | 8 | FALSE | 10 | 18 | 10 | 1953 | 2186 | 524 |
| 2,69472E- | FALSE | High | IsMasterP | AAEL0142 | 1:724906 | 0 | 43 | FALSE | 6 | 14 | 6 | 247 | 264 | 539 |
| 2,66552E- | FALSE | High | IsMasterP | AAEL0044 | 3:206547( | 0 | 25 | FALSE | 5 | 14 | 5 | 346 | 387 | 800 |
| 3,27424E- | FALSE | High | IsMasterP | AAEL0092 | 3:200246: | 0 | 64 | FALSE | 3 | 15 | 3 | 121 | 127 | 850 |
| 9,17248E- | FALSE | High | IsMasterP | AAEL0070 | 1:127944! | 0 | 46 | FALSE | 4 | 14 | 4 | 173 | 195 | 640 |

| | | | | | | | | | | | | | | |
|---|---|---|---|---|---|---|---|---|---|---|---|---|---|---|
| -6,50245E | FALSE | High | IsMasterP | AAEL0139 | 3:230597 | 0 | 42 | FALSE | 4 | 19 | 4 | 112 | 123 | 888 |
| 3,18181E- | FALSE | High | IsMasterP | AAEL0142 | 2:108769 | 0 | 37 | FALSE | 5 | 18 | 5 | 130 | 151 | 1114 |
| -5,89101E | FALSE | High | IsMasterP | AAEL0000 | 3:105057 | 0 | 26 | FALSE | 6 | 16 | 6 | 363 | 399 | 715 |
| 1,26831E- | FALSE | High | IsMasterP | AAEL0023 | 3:375635 | 0 | 32 | FALSE | 6 | 15 | 6 | 327 | 360 | 686 |
| 3,92281E- | FALSE | High | IsMasterP | AAEL0217 | 2:325183 | 0 | 40 | FALSE | 2 | 12 | 2 | 118 | 124 | 775 |
| 3,33462E- | FALSE | High | IsMasterP | AAEL0013 | 3:327097 | 0 | 35 | FALSE | 9 | 17 | 9 | 445 | 514 | 673 |
| 6,64742E- | FALSE | High | IsMasterP | AAEL0100 | 3:112150 | 0 | 13 | FALSE | 7 | 20 | 7 | 609 | 695 | 742 |
| -7,55123E | FALSE | High | IsMasterP | AAEL0146 | 3:368609 | 0 | 18 | FALSE | 7 | 20 | 4 | 535 | 616 | 753 |
| -8,99969E | FALSE | High | IsMasterP | AAEL0008 | 3:332232 | 0 | 41 | FALSE | 8 | 18 | 8 | 328 | 364 | 705 |
| 5,03445E- | FALSE | High | IsMasterP | AAEL0221 | 2:192364 | 0 | 17 | FALSE | 7 | 20 | 1 | 487 | 572 | 988 |
| 7,31687E- | FALSE | High | IsMasterP | AAEL0111 | 3:513275 | 0 | 5 | FALSE | 7 | 16 | 7 | 2141 | 2446 | 591 |
| 3,04298E- | FALSE | High | IsMasterP | AAEL0013 | 3:486966 | 0 | 14 | FALSE | 10 | 17 | 10 | 1073 | 1183 | 939 |
| 2,6285E+1 | FALSE | High | IsMasterP | AAEL0037 | 1:303907 | 0 | 30 | FALSE | 9 | 16 | 9 | 440 | 498 | 681 |
| 4,85728E- | FALSE | High | IsMasterP | AAEL0003 | 2:367910 | 0 | 30 | FALSE | 9 | 18 | 9 | 643 | 677 | 769 |
| 7,62812E- | FALSE | High | IsMasterP | AAEL0081 | 3:146744 | 0 | 16 | FALSE | 7 | 17 | 7 | 650 | 739 | 590 |
| 1,25164E- | FALSE | High | IsMasterP | AAEL0032 | 1:290940 | 0 | 21 | FALSE | 4 | 16 | 4 | 310 | 341 | 516 |
| -7,56197E | FALSE | High | IsMasterP | AAEL0245 | 2:191909 | 0 | 27 | FALSE | 8 | 21 | 8 | 424 | 468 | 911 |
| 5,87774E- | FALSE | High | IsMasterP | AAEL0075 | 2:326973 | 0 | 27 | FALSE | 4 | 13 | 4 | 363 | 411 | 577 |
| 5,85371E- | FALSE | High | IsMasterP | AAEL0225 | 1:305612 | 0 | 8 | FALSE | 7 | 17 | 7 | 1303 | 1473 | 491 |
| 2,46331E- | FALSE | High | IsMasterP | AAEL0033 | 3:808037 | 0 | 31 | FALSE | 12 | 19 | 12 | 661 | 744 | 917 |
| -3,48122E | FALSE | High | IsMasterP | AAEL0257 | 1:144639 | 0 | 45 | FALSE | 4 | 15 | 2 | 180 | 206 | 693 |
| -6,73167E | FALSE | High | IsMasterP | AAEL0013 | 2:569875 | 0 | 26 | FALSE | 5 | 17 | 5 | 204 | 235 | 1083 |
| -1,6852E+ | FALSE | High | IsMasterP | AAEL0105 | 1:301365 | 0 | 36 | FALSE | 9 | 16 | 9 | 487 | 544 | 571 |
| -1,35364E | FALSE | High | IsMasterP | AAEL0044 | 3:212341 | 0 | 49 | FALSE | 6 | 16 | 6 | 182 | 206 | 714 |
| 3,80627E- | FALSE | High | IsMasterP | AAEL0097 | 1:676061 | 0 | 41 | FALSE | 5 | 14 | 5 | 224 | 249 | 487 |
| 4,3497E+1 | FALSE | High | IsMasterP | AAEL0069 | 1:156708 | 0 | 14 | FALSE | 11 | 17 | 11 | 1201 | 1401 | 774 |
| 2,02121E- | FALSE | High | IsMasterP | AAEL0122 | 1:146832 | 0 | 27 | FALSE | 10 | 18 | 10 | 653 | 729 | 576 |
| 3,64086E- | FALSE | High | IsMasterP | AAEL0044 | 3:206509 | 0 | 50 | FALSE | 6 | 21 | 6 | 191 | 223 | 573 |
| 8,29258E- | FALSE | High | IsMasterP | AAEL0202 | NIGP0100 | 0 | 41 | FALSE | 5 | 14 | 5 | 140 | 155 | 847 |
| -6,87895E | FALSE | High | IsMasterP | AAEL0061 | 3:140732 | 0 | 19 | FALSE | 6 | 17 | 6 | 377 | 418 | 831 |
| -6,77743E | FALSE | High | IsMasterP | AAEL0124 | 2:122681 | 0 | 15 | FALSE | 13 | 19 | 13 | 1500 | 1683 | 475 |

| | | | | | | | | | | | | | | |
|---|---|---|---|---|---|---|---|---|---|---|---|---|---|---|
| 2,85552E- | FALSE | High | IsMasterP | AAEL0005 | 3:1247193 | 0 | 27 | FALSE | 7 | 18 | 7 | 329 | 370 | 486 |
| -5,54014E | FALSE | High | IsMasterP | AAEL0028 | 3:2349452 | 0 | 49 | FALSE | 4 | 14 | 4 | 111 | 129 | 1021 |
| 6,296E+18 | FALSE | High | IsMasterP | AAEL0007 | 2:4588626 | 0 | 14 | FALSE | 5 | 17 | 3 | 493 | 538 | 493 |
| -8,30224E | FALSE | High | IsMasterP | AAEL0037 | 1:2037303 | 0 | 9 | FALSE | 6 | 17 | 6 | 923 | 1020 | 681 |
| 4,68512E- | FALSE | High | IsMasterP | AAEL0023 | 3:3741608 | 0 | 55 | FALSE | 9 | 20 | 9 | 220 | 249 | 581 |
| 1,97754E- | FALSE | High | IsMasterP | AAEL0031 | 1:2150439 | 0 | 22 | FALSE | 6 | 14 | 6 | 548 | 616 | 649 |
| 7,44301E- | FALSE | High | IsMasterP | AAEL0105 | 2:9073890 | 0 | 21 | FALSE | 8 | 16 | 8 | 632 | 692 | 627 |
| -5,02022E | FALSE | High | IsMasterP | AAEL0052 | NIGP0100 | 0 | 12 | FALSE | 10 | 18 | 10 | 1310 | 1509 | 756 |
| -4,05899E | FALSE | High | IsMasterP | AAEL0064 | 3:1523447 | 0 | 42 | FALSE | 4 | 13 | 4 | 154 | 167 | 709 |
| 1,22186E- | FALSE | High | IsMasterP | AAEL0080 | 1:6156315 | 0 | 15 | FALSE | 9 | 21 | 9 | 689 | 776 | 810 |
| 8,56827E- | FALSE | High | IsMasterP | AAEL0054 | 3:5988676 | 0 | 22 | FALSE | 5 | 15 | 5 | 365 | 420 | 624 |
| 3,49984E- | FALSE | High | IsMasterP | AAEL0018 | 2:2752742 | 0 | 21 | FALSE | 9 | 18 | 9 | 633 | 718 | 623 |
| 7,41396E- | FALSE | High | IsMasterP | AAEL0001 | 1:1949475 | 0 | 57 | FALSE | 6 | 20 | 6 | 139 | 160 | 979 |
| 3,552E+18 | FALSE | High | IsMasterP | AAEL0256 | 1:6020008 | 0 | 46 | FALSE | 7 | 17 | 7 | 227 | 265 | 904 |
| 9,1714E+1 | FALSE | High | IsMasterP | AAEL0009 | 3:3771175 | 0 | 61 | FALSE | 4 | 19 | 4 | 89 | 100 | 514 |
| -6,87556E | FALSE | High | IsMasterP | AAEL0048 | 1:2885351 | 0 | 12 | FALSE | 4 | 19 | 4 | 485 | 534 | 534 |
| -1,27197E | FALSE | High | IsMasterP | AAEL0122 | 2:1450848 | 0 | 27 | FALSE | 8 | 16 | 8 | 392 | 443 | 524 |
| -4,54888E | FALSE | High | IsMasterP | AAEL0105 | 2:9105251 | 0 | 41 | FALSE | 7 | 17 | 7 | 189 | 219 | 488 |
| 9,15347E- | FALSE | High | IsMasterP | AAEL0038 | 3:3428581 | 0 | 28 | FALSE | 6 | 14 | 6 | 306 | 350 | 897 |
| 6,71739E- | FALSE | High | IsMasterP | AAEL0115 | 2:1231895 | 0 | 16 | FALSE | 6 | 18 | 6 | 341 | 374 | 545 |
| -7,27108E | FALSE | High | IsMasterP | AAEL0173 | 3:2351506 | 0 | 37 | FALSE | 8 | 17 | 8 | 309 | 336 | 976 |
| -4,88749E | FALSE | High | IsMasterP | AAEL0237 | 3:2800734 | 0 | 7 | FALSE | 7 | 18 | 7 | 1459 | 1670 | 714 |
| 2,67662E- | FALSE | High | IsMasterP | AAEL0121 | 2:4059280 | 0 | 26 | FALSE | 4 | 12 | 4 | 239 | 272 | 535 |
| -2,28612E | FALSE | High | IsMasterP | AAEL0007 | 3:2456763 | 0 | 34 | FALSE | 6 | 15 | 6 | 273 | 302 | 580 |
| -3,616E+1 | FALSE | High | IsMasterP | AAEL0127 | 2:9411553 | 0 | 8 | FALSE | 8 | 18 | 8 | 1622 | 1824 | 790 |
| 7,5718E+1 | FALSE | High | IsMasterP | AAEL0084 | NIGP0100 | 0 | 40 | FALSE | 6 | 18 | 6 | 209 | 239 | 954 |
| -4,18097E | FALSE | High | IsMasterP | AAEL0139 | 2:3058310 | 0 | 18 | FALSE | 8 | 16 | 5 | 702 | 782 | 803 |
| -6,33222E | FALSE | High | IsMasterP | AAEL0122 | 1:1465364 | 0 | 36 | FALSE | 4 | 16 | 4 | 126 | 142 | 508 |
| 4,608E+18 | FALSE | High | IsMasterP | AAEL0120 | 3:1473715 | 0 | 31 | FALSE | 8 | 17 | 8 | 415 | 474 | 702 |
| 1,72181E- | FALSE | High | IsMasterP | AAEL0102 | 3:2790462 | 0 | 19 | FALSE | 6 | 15 | 6 | 529 | 570 | 938 |
| -6,18397E | FALSE | High | IsMasterP | AAEL0078 | 2:4605951 | 0 | 33 | FALSE | 6 | 16 | 6 | 214 | 236 | 863 |

| | | | | | | | | | | | | | | |
|---|---|---|---|---|---|---|---|---|---|---|---|---|---|---|
| -6,95601E | FALSE | High | IsMasterP | AAEL0194 | 1:116258: | 0 | 5 | FALSE | 8 | 15 | 8 | 2806 | 3097 | 526 |
| 7,12114E- | FALSE | High | IsMasterP | AAEL0109 | 3:168206 | 0 | 37 | FALSE | 4 | 11 | 4 | 228 | 253 | 515 |
| 5,65842E- | FALSE | High | IsMasterP | AAEL0068 | 3:370351( | 0 | 44 | FALSE | 8 | 19 | 8 | 156 | 169 | 512 |
| 1,72868E- | FALSE | High | IsMasterP | AAEL0133 | 3:173112( | 0 | 31 | FALSE | 4 | 16 | 4 | 235 | 265 | 870 |
| -5,59578E | FALSE | High | IsMasterP | AAEL0136 | 1:267705( | 0 | 7 | FALSE | 8 | 18 | 8 | 1636 | 1767 | 632 |
| -5,45838E | FALSE | High | IsMasterP | AAEL0043 | 3:511146 | 0 | 24 | FALSE | 10 | 19 | 10 | 560 | 639 | 838 |
| 5,1977E+1 | FALSE | High | IsMasterP | AAEL0100 | 1:194273: | 0 | 53 | FALSE | 6 | 16 | 6 | 193 | 219 | 703 |
| 5,90776E- | FALSE | High | IsMasterP | AAEL0016 | 2:300141: | 0 | 20 | FALSE | 8 | 16 | 8 | 641 | 720 | 727 |
| -7,22399E | FALSE | High | IsMasterP | AAEL0093 | 3:851781( | 0 | 19 | FALSE | 7 | 14 | 7 | 635 | 717 | 854 |
| -6,08277E | FALSE | High | IsMasterP | AAEL0024 | 1:987079: | 0 | 10 | FALSE | 9 | 17 | 9 | 1869 | 1919 | 947 |
| 5,261E+18 | FALSE | High | IsMasterP | AAEL0047 | 2:289186: | 0 | 32 | FALSE | 9 | 19 | 9 | 356 | 404 | 815 |
| 5,96759E- | FALSE | High | IsMasterP | AAEL0096 | 3:269102: | 0 | 20 | FALSE | 8 | 19 | 8 | 535 | 614 | 524 |
| 6,26281E- | FALSE | High | IsMasterP | AAEL0131 | 3:843115 | 0 | 38 | FALSE | 7 | 16 | 7 | 250 | 285 | 660 |
| -2,62392E | FALSE | High | IsMasterP | AAEL0083 | 3:413102: | 0 | 30 | FALSE | 8 | 15 | 8 | 431 | 490 | 613 |
| 8,11907E- | FALSE | High | IsMasterP | AAEL0001 | 1:197311 | 0 | 37 | FALSE | 8 | 16 | 8 | 285 | 313 | 692 |
| -3,36035E | FALSE | High | IsMasterP | AAEL0155 | 2:250785( | 0 | 30 | FALSE | 7 | 17 | 7 | 432 | 476 | 799 |
| -3,45539E | FALSE | High | IsMasterP | AAEL0149 | 1:303695 | 0 | 24 | FALSE | 7 | 17 | 7 | 381 | 432 | 526 |
| 1,54398E- | FALSE | High | IsMasterP | AAEL0013 | 2:118115: | 0 | 27 | FALSE | 2 | 15 | 2 | 126 | 139 | 1162 |
| 3,32998E- | FALSE | High | IsMasterP | AAEL0097 | 3:284268: | 0 | 36 | FALSE | 11 | 18 | 2 | 567 | 624 | 712 |
| -1,82628E | FALSE | High | IsMasterP | AAEL0084 | 1:939734 | 0 | 51 | FALSE | 5 | 12 | 5 | 228 | 253 | 652 |
| -6,79307E | FALSE | High | IsMasterP | AAEL0148 | 3:293946( | 0 | 19 | FALSE | 9 | 17 | 9 | 473 | 520 | 774 |
| 7,24744E- | FALSE | High | IsMasterP | AAEL0095 | 3:344952: | 0 | 27 | FALSE | 5 | 15 | 5 | 307 | 330 | 903 |
| -5,15629E | FALSE | High | IsMasterP | AAEL0063 | 1:100826 | 0 | 40 | FALSE | 10 | 16 | 10 | 376 | 415 | 805 |
| 6,79054E- | FALSE | High | IsMasterP | AAEL0003 | 2:124435 | 0 | 31 | FALSE | 10 | 19 | 10 | 448 | 508 | 728 |
| -4,60229E | FALSE | High | IsMasterP | AAEL0055 | 1:906469 | 0 | 21 | FALSE | 8 | 19 | 8 | 476 | 532 | 714 |
| 6,14651E- | FALSE | High | IsMasterP | AAEL0064 | 3:155528: | 0 | 16 | FALSE | 8 | 17 | 8 | 756 | 849 | 667 |
| -1,37554E | FALSE | High | IsMasterP | AAEL0098 | 3:202834 | 0 | 34 | FALSE | 6 | 18 | 6 | 246 | 279 | 957 |
| -2,99479E | FALSE | High | IsMasterP | AAEL0000 | 3:102858 | 0 | 10 | FALSE | 2 | 16 | 2 | 358 | 396 | 530 |
| 4,16029E- | FALSE | High | IsMasterP | AAEL0078 | 1:149140: | 0 | 41 | FALSE | 4 | 14 | 4 | 189 | 206 | 876 |
| 8,61129E- | FALSE | High | IsMasterP | AAEL0002 | 3:215597: | 0 | 52 | FALSE | 4 | 17 | 4 | 112 | 123 | 634 |
| 3,80331E- | FALSE | High | IsMasterP | AAEL0035 | 1:230238 | 0 | 45 | FALSE | 7 | 18 | 7 | 208 | 236 | 623 |

| | | | | | | | | | | | | | | |
|---|---|---|---|---|---|---|---|---|---|---|---|---|---|---|
| 3,87423E- | FALSE | High | IsMasterP | AAEL0131 | 3:3709437 | 0 | 23 | FALSE | 9 | 18 | 9 | 554 | 628 | 670 |
| 4,0901E+1 | FALSE | High | IsMasterP | AAEL0199 | 2:5762370 | 0 | 14 | FALSE | 8 | 16 | 8 | 1050 | 1179 | 816 |
| 7,50486E- | FALSE | High | IsMasterP | AAEL0063 | 3:3964380 | 0 | 27 | FALSE | 10 | 19 | 10 | 503 | 564 | 521 |
| -2,03944E | FALSE | High | IsMasterP | AAEL0127 | 1:2906234 | 0 | 13 | FALSE | 8 | 17 | 8 | 1000 | 1119 | 496 |
| 6,01943E- | FALSE | High | IsMasterP | AAEL0008 | 3:3638327 | 0 | 14 | FALSE | 7 | 14 | 7 | 943 | 1074 | 533 |
| -4,7698E+ | FALSE | High | IsMasterP | AAEL0194 | 3:2759393 | 0 | 6 | FALSE | 8 | 17 | 8 | 2272 | 2546 | 574 |
| 2,64439E- | FALSE | High | IsMasterP | AAEL0035 | 3:2640591 | 0 | 32 | FALSE | 6 | 15 | 6 | 275 | 308 | 605 |
| 7,99934E- | FALSE | High | IsMasterP | AAEL0022 | 2:3085980 | 0 | 26 | FALSE | 8 | 18 | 8 | 433 | 492 | 654 |
| 4,44787E- | FALSE | High | IsMasterP | AAEL0016 | 2:2996641 | 0 | 9 | FALSE | 7 | 16 | 7 | 1284 | 1447 | 829 |
| -8,30431E | FALSE | High | IsMasterP | AAEL0110 | 1:7342259 | 0 | 21 | FALSE | 7 | 15 | 7 | 494 | 550 | 553 |
| 6,45856E- | FALSE | High | IsMasterP | AAEL0002 | 3:2650132 | 0 | 11 | FALSE | 3 | 19 | 3 | 404 | 449 | 724 |
| -7,77972E | FALSE | High | IsMasterP | AAEL0075 | 2:3272199 | 0 | 22 | FALSE | 8 | 16 | 8 | 655 | 744 | 854 |
| -2,10405E | FALSE | High | IsMasterP | AAEL0112 | 2:1797610 | 0 | 12 | FALSE | 8 | 17 | 8 | 1111 | 1215 | 578 |
| -8,40014E | FALSE | High | IsMasterP | AAEL0066 | 2:3250420 | 0 | 25 | FALSE | 7 | 17 | 7 | 374 | 420 | 536 |
| -3,56576E | FALSE | High | IsMasterP | AAEL0227 | 2:1793692 | 0 | 13 | FALSE | 6 | 17 | 6 | 631 | 709 | 783 |
| -8,70598E | FALSE | High | IsMasterP | AAEL0090 | 3:1348680 | 0 | 66 | FALSE | 6 | 16 | 6 | 120 | 134 | 806 |
| -2,66192E | FALSE | High | IsMasterP | AAEL0170 | 2:6799199 | 0 | 18 | FALSE | 6 | 17 | 6 | 456 | 520 | 553 |
| -5,809E+1 | FALSE | High | IsMasterP | AAEL0041 | 3:4017303 | 0 | 34 | FALSE | 7 | 16 | 7 | 320 | 364 | 703 |
| -5,95221E | FALSE | High | IsMasterP | AAEL0075 | 2:3272465 | 0 | 26 | FALSE | 7 | 13 | 7 | 373 | 416 | 818 |
| -1,89745E | FALSE | High | IsMasterP | AAEL0125 | 2:7614577 | 0 | 28 | FALSE | 7 | 17 | 7 | 395 | 447 | 705 |
| -1,93634E | FALSE | High | IsMasterP | AAEL0124 | 3:1114375 | 0 | 26 | FALSE | 5 | 15 | 5 | 360 | 407 | 750 |
| 4,69553E- | FALSE | High | IsMasterP | AAEL0117 | 1:1473482 | 0 | 20 | FALSE | 7 | 17 | 7 | 395 | 447 | 628 |
| -8,18857E | FALSE | High | IsMasterP | AAEL0069 | 3:1431022 | 0 | 45 | FALSE | 10 | 16 | 10 | 394 | 410 | 670 |
| 6,67867E- | FALSE | High | IsMasterP | AAEL0002 | 3:2128666 | 0 | 60 | FALSE | 5 | 15 | 5 | 89 | 93 | 848 |
| -4,00188E | FALSE | High | IsMasterP | AAEL0000 | 3:1020172 | 0 | 26 | FALSE | 7 | 14 | 7 | 437 | 498 | 674 |
| -8,70326E | FALSE | High | IsMasterP | AAEL0081 | 3:7907450 | 0 | 24 | FALSE | 7 | 14 | 7 | 506 | 591 | 559 |
| -1,5671E+ | FALSE | High | IsMasterP | AAEL0270 | 2:4386338 | 0 | 80 | FALSE | 7 | 18 | 7 | 85 | 95 | 635 |
| 3,22007E- | FALSE | High | IsMasterP | AAEL0251 | 2:9327363 | 0 | 8 | FALSE | 9 | 17 | 9 | 1546 | 1766 | 529 |
| -8,94947E | FALSE | High | IsMasterP | AAEL0122 | 1:1517195 | 0 | 9 | FALSE | 7 | 18 | 7 | 947 | 1089 | 628 |
| 5,09246E- | FALSE | High | IsMasterP | AAEL0105 | 1:3013564 | 0 | 27 | FALSE | 6 | 18 | 6 | 293 | 323 | 887 |
| -9,48222E | FALSE | High | IsMasterP | AAEL0049 | 2:2822545 | 0 | 27 | FALSE | 5 | 19 | 5 | 222 | 251 | 709 |

| | | | | | | | | | | | | | | |
|---|---|---|---|---|---|---|---|---|---|---|---|---|---|---|
| -3,01187E | FALSE | High | IsMasterP | AAEL0044 | 3:204396 | 0 | 8 | FALSE | 8 | 18 | 8 | 1020 | 1151 | 478 |
| -7,87388E | FALSE | High | IsMasterP | AAEL0172 | 1:194025 | 0 | 10 | FALSE | 8 | 17 | 8 | 1176 | 1337 | 702 |
| 7,33316E | FALSE | High | IsMasterP | AAEL0060 | 3:307449 | 0 | 15 | FALSE | 7 | 15 | 7 | 750 | 837 | 500 |
| 6,94832E | FALSE | High | IsMasterP | AAEL0069 | 3:170496 | 0 | 16 | FALSE | 5 | 14 | 1 | 471 | 505 | 456 |
| -3,82011E | FALSE | High | IsMasterP | AAEL0065 | 2:469131 | 0 | 36 | FALSE | 6 | 14 | 6 | 278 | 318 | 623 |
| 8,55789E | FALSE | High | IsMasterP | AAEL0109 | 2:108510 | 0 | 29 | FALSE | 7 | 15 | 7 | 521 | 577 | 547 |
| 2,79782E | FALSE | High | IsMasterP | AAEL0148 | 2:257424 | 0 | 40 | FALSE | 8 | 17 | 8 | 280 | 304 | 910 |
| -3,01768E | FALSE | High | IsMasterP | AAEL0067 | 2:851108 | 0 | 58 | FALSE | 6 | 14 | 6 | 165 | 190 | 510 |
| 3,31915E | FALSE | High | IsMasterP | AAEL0073 | 2:345126 | 0 | 14 | FALSE | 3 | 15 | 3 | 342 | 377 | 696 |
| 1,65154E | FALSE | High | IsMasterP | AAEL0041 | 2:146294 | 0 | 29 | FALSE | 9 | 16 | 9 | 422 | 474 | 818 |
| 7,03687E | FALSE | High | IsMasterP | AAEL0023 | 2:302202 | 0 | 25 | FALSE | 8 | 15 | 8 | 503 | 568 | 519 |
| 2,62417E | FALSE | High | IsMasterP | AAEL0198 | 3:301209 | 0 | 35 | FALSE | 9 | 16 | 6 | 357 | 410 | 683 |
| 6,81349E | FALSE | High | IsMasterP | AAEL0047 | 2:378816 | 0 | 20 | FALSE | 6 | 17 | 6 | 433 | 484 | 536 |
| -5,66621E | FALSE | High | IsMasterP | AAEL0170 | 2:179122 | 0 | 12 | FALSE | 8 | 18 | 8 | 728 | 762 | 610 |
| -5,97513E | FALSE | High | IsMasterP | AAEL0255 | 2:468980 | 0 | 20 | FALSE | 7 | 17 | 7 | 576 | 649 | 568 |
| -6,08093E | FALSE | High | IsMasterP | AAEL0011 | 3:407139 | 0 | 67 | FALSE | 6 | 15 | 6 | 127 | 151 | 916 |
| 2,06491E | FALSE | High | IsMasterP | AAEL0238 | 1:306520 | 0 | 6 | FALSE | 10 | 14 | 10 | 4015 | 4220 | 651 |
| 5,26029E | FALSE | High | IsMasterP | AAEL0063 | 3:231626 | 0 | 10 | FALSE | 5 | 15 | 5 | 804 | 922 | 638 |
| 9,15337E | FALSE | High | IsMasterP | AAEL0127 | 3:424021 | 0 | 7 | FALSE | 6 | 19 | 6 | 1251 | 1415 | 583 |
| -6,75531E | FALSE | High | IsMasterP | AAEL0138 | 1:119341 | 0 | 20 | FALSE | 5 | 16 | 5 | 399 | 432 | 582 |
| -3,51085E | FALSE | High | IsMasterP | AAEL0243 | NIGP0100 | 0 | 16 | FALSE | 9 | 14 | 3 | 741 | 830 | 508 |
| -3,22074E | FALSE | High | IsMasterP | AAEL0171 | 2:165800 | 0 | 6 | FALSE | 3 | 15 | 3 | 631 | 687 | 976 |
| 2,57839E | FALSE | High | IsMasterP | AAEL0044 | 3:271068 | 0 | 15 | FALSE | 7 | 19 | 7 | 672 | 729 | 637 |
| -2,71413E | FALSE | High | IsMasterP | AAEL0061 | 3:259777 | 0 | 29 | FALSE | 5 | 17 | 5 | 221 | 258 | 614 |
| -1,73209E | FALSE | High | IsMasterP | AAEL0133 | 3:341427 | 0 | 40 | FALSE | 6 | 16 | 6 | 171 | 182 | 906 |
| 1,58976E | FALSE | High | IsMasterP | AAEL0227 | 2:108064 | 0 | 32 | FALSE | 8 | 18 | 8 | 520 | 584 | 644 |
| 5,92892E | FALSE | High | IsMasterP | AAEL0116 | 3:281478 | 0 | 41 | FALSE | 4 | 15 | 4 | 115 | 118 | 848 |
| -3,94512E | FALSE | High | IsMasterP | AAEL0007 | 3:242885 | 0 | 68 | FALSE | 4 | 11 | 4 | 85 | 92 | 998 |
| 5,691E+1 | FALSE | High | IsMasterP | AAEL0100 | 2:110882 | 0 | 54 | FALSE | 7 | 16 | 7 | 128 | 147 | 517 |
| -6,58666E | FALSE | High | IsMasterP | AAEL0267 | 2:592142 | 0 | 24 | FALSE | 8 | 15 | 8 | 615 | 653 | 881 |
| -8,93091E | FALSE | High | IsMasterP | AAEL0040 | 3:166208 | 0 | 23 | FALSE | 7 | 13 | 7 | 333 | 363 | 910 |

| | | | | | | | | | | | | | | |
|---|---|---|---|---|---|---|---|---|---|---|---|---|---|---|
| -3,74255E | FALSE | High | IsMasterP | AAEL0043 | 3:936933 | 0 | 51 | FALSE | 5 | 15 | 5 | 150 | 170 | 705 |
| -6,40353E | FALSE | High | IsMasterP | AAEL0018 | 2:274295 | 0 | 56 | FALSE | 7 | 16 | 7 | 131 | 152 | 803 |
| -3,16319E | FALSE | High | IsMasterP | AAEL0044 | 2:473315 | 0 | 50 | FALSE | 5 | 16 | 5 | 108 | 118 | 948 |
| -5,40448E | FALSE | High | IsMasterP | AAEL0016 | 2:300456 | 0 | 47 | FALSE | 7 | 17 | 7 | 170 | 194 | 720 |
| 6,5926E+ | FALSE | High | IsMasterP | AAEL0127 | 3:423647 | 0 | 21 | FALSE | 6 | 14 | 6 | 404 | 452 | 648 |
| -4,51561E | FALSE | High | IsMasterP | AAEL0103 | 2:403762 | 0 | 39 | FALSE | 7 | 14 | 7 | 230 | 257 | 652 |
| 1,53009E | FALSE | High | IsMasterP | AAEL0009 | 3:380158 | 0 | 32 | FALSE | 7 | 15 | 7 | 329 | 370 | 470 |
| 1,90402E | FALSE | High | IsMasterP | AAEL0110 | 2:146421 | 0 | 16 | FALSE | 4 | 14 | 3 | 279 | 311 | 611 |
| -1,52603E | FALSE | High | IsMasterP | AAEL0008 | 3:362027 | 0 | 25 | FALSE | 5 | 12 | 5 | 339 | 367 | 857 |
| 1,57863E | FALSE | High | IsMasterP | AAEL0096 | 2:169189 | 0 | 39 | FALSE | 6 | 12 | 6 | 306 | 338 | 699 |
| -5,98624E | FALSE | High | IsMasterP | AAEL0115 | 3:107039 | 0 | 13 | FALSE | 13 | 16 | 13 | 1339 | 1451 | 739 |
| -7,77273E | FALSE | High | IsMasterP | AAEL0275 | 1:119384 | 0 | 9 | FALSE | 8 | 15 | 8 | 1576 | 1794 | 633 |
| -8,79727E | FALSE | High | IsMasterP | AAEL0236 | 3:104320 | 0 | 16 | FALSE | 7 | 15 | 7 | 641 | 720 | 534 |
| 6,78338E | FALSE | High | IsMasterP | AAEL0049 | 2:281781 | 0 | 31 | FALSE | 6 | 14 | 6 | 292 | 320 | 442 |
| -2,16484E | FALSE | High | IsMasterP | AAEL0108 | 3:156746 | 0 | 14 | FALSE | 7 | 17 | 7 | 727 | 810 | 942 |
| -6,00279E | FALSE | High | IsMasterP | AAEL0088 | 2:240808 | 0 | 20 | FALSE | 7 | 17 | 7 | 513 | 587 | 521 |
| 3,34707E | FALSE | High | IsMasterP | AAEL0122 | 3:533803 | 0 | 25 | FALSE | 7 | 14 | 7 | 363 | 408 | 812 |
| 4,24999E | FALSE | High | IsMasterP | AAEL0209 | 2:492423 | 0 | 25 | FALSE | 6 | 17 | 6 | 343 | 381 | 803 |
| -6,22386E | FALSE | High | IsMasterP | AAEL0100 | 3:112258 | 0 | 19 | FALSE | 7 | 16 | 7 | 609 | 689 | 787 |
| -8,05414E | FALSE | High | IsMasterP | AAEL0042 | 1:105060 | 0 | 15 | FALSE | 10 | 14 | 10 | 1242 | 1428 | 866 |
| -3,00976E | FALSE | High | IsMasterP | AAEL0012 | 2:154135 | 0 | 24 | FALSE | 5 | 12 | 5 | 534 | 551 | 885 |
| -5,32798E | FALSE | High | IsMasterP | AAEL0027 | 3:124506 | 0 | 28 | FALSE | 5 | 13 | 5 | 241 | 275 | 797 |
| -3,62325E | FALSE | High | IsMasterP | AAEL0065 | 2:468690 | 0 | 38 | FALSE | 4 | 15 | 4 | 99 | 109 | 964 |
| -5,82922E | FALSE | High | IsMasterP | AAEL0170 | 2:462896 | 0 | 38 | FALSE | 7 | 14 | 7 | 367 | 399 | 690 |
| 5,48307E | FALSE | High | IsMasterP | AAEL0012 | 2:413303 | 0 | 24 | FALSE | 6 | 13 | 6 | 423 | 488 | 574 |
| -2,93653E | FALSE | High | IsMasterP | AAEL0065 | 2:468677 | 0 | 43 | FALSE | 4 | 16 | 4 | 155 | 169 | 712 |
| 5,41027E | FALSE | High | IsMasterP | AAEL0090 | 2:391816 | 0 | 27 | FALSE | 7 | 11 | 7 | 542 | 594 | 728 |
| 7,27291E | FALSE | High | IsMasterP | AAEL0092 | 3:199453 | 0 | 26 | FALSE | 7 | 15 | 4 | 327 | 373 | 633 |
| 5,96991E | FALSE | High | IsMasterP | AAEL0031 | 2:440971 | 0 | 29 | FALSE | 8 | 15 | 8 | 438 | 498 | 948 |
| 3,75532E | FALSE | High | IsMasterP | AAEL0020 | 2:356667 | 0 | 35 | FALSE | 7 | 15 | 7 | 325 | 346 | 762 |
| -6,28672E | FALSE | High | IsMasterP | AAEL0114 | 3:142598 | 0 | 4 | FALSE | 5 | 13 | 5 | 2816 | 3053 | 512 |

| | | | | | | | | | | | | | | |
|---|---|---|---|---|---|---|---|---|---|---|---|---|---|---|
| -3,44985E | FALSE | High | IsMasterP | AAEL0110 | 3:287784 | 0 | 30 | FALSE | 6 | 14 | 6 | 269 | 302 | 887 |
| -4,50187E | FALSE | High | IsMasterP | AAEL0027 | 1:305225 | 0 | 24 | FALSE | 5 | 14 | 5 | 417 | 479 | 854 |
| 7,26348E- | FALSE | High | IsMasterP | AAEL0208 | 2:132859 | 0 | 16 | FALSE | 7 | 15 | 7 | 663 | 728 | 840 |
| 8,58747E- | FALSE | High | IsMasterP | AAEL0026 | 3:256697 | 0 | 29 | FALSE | 6 | 13 | 6 | 414 | 448 | 585 |
| -6,38236E | FALSE | High | IsMasterP | AAEL0084 | 2:190985 | 0 | 67 | FALSE | 6 | 13 | 6 | 152 | 180 | 544 |
| -6,02236E | FALSE | High | IsMasterP | AAEL0051 | 1:168241 | 0 | 29 | FALSE | 3 | 12 | 3 | 196 | 217 | 960 |
| 6,38409E- | FALSE | High | IsMasterP | AAEL0082 | 3:598838 | 0 | 40 | FALSE | 6 | 13 | 6 | 345 | 371 | 587 |
| 5,35024E- | FALSE | High | IsMasterP | AAEL0120 | 2:144273 | 0 | 31 | FALSE | 8 | 13 | 8 | 498 | 565 | 558 |
| -7,37525E | FALSE | High | IsMasterP | AAEL0025 | 3:256859 | 0 | 31 | FALSE | 7 | 13 | 7 | 415 | 453 | 587 |
| -6,57685E | FALSE | High | IsMasterP | AAEL0128 | 1:222036 | 0 | 26 | FALSE | 8 | 15 | 8 | 271 | 305 | 668 |
| 3,71854E- | FALSE | High | IsMasterP | AAEL0031 | 1:119043 | 0 | 36 | FALSE | 3 | 16 | 3 | 147 | 167 | 730 |
| -5,28638E | FALSE | High | IsMasterP | AAEL0145 | 2:325146 | 0 | 39 | FALSE | 2 | 18 | 1 | 122 | 128 | 747 |
| 6,89308E- | FALSE | High | IsMasterP | AAEL0065 | 3:447379 | 0 | 22 | FALSE | 8 | 14 | 8 | 553 | 635 | 597 |
| -8,69981E | FALSE | High | IsMasterP | AAEL0128 | 3:275877 | 0 | 38 | FALSE | 6 | 15 | 6 | 211 | 245 | 907 |
| 2,51745E- | FALSE | High | IsMasterP | AAEL0023 | 2:176164 | 0 | 49 | FALSE | 3 | 14 | 3 | 138 | 160 | 486 |
| -9,02584E | FALSE | High | IsMasterP | AAEL0197 | 3:515922 | 0 | 17 | FALSE | 5 | 15 | 4 | 350 | 383 | 885 |
| 3,71366E- | FALSE | High | IsMasterP | AAEL0124 | 1:123531 | 0 | 27 | FALSE | 7 | 13 | 7 | 370 | 406 | 639 |
| 6,87056E- | FALSE | High | IsMasterP | AAEL0039 | 1:250618 | 0 | 12 | FALSE | 6 | 15 | 6 | 878 | 1013 | 538 |
| 6,80797E- | FALSE | High | IsMasterP | AAEL0170 | 2:391124 | 0 | 21 | FALSE | 4 | 14 | 4 | 193 | 203 | 1032 |
| -2,65137E | FALSE | High | IsMasterP | AAEL0076 | 1:187261 | 0 | 46 | FALSE | 5 | 12 | 5 | 243 | 285 | 432 |
| -3,81973E | FALSE | High | IsMasterP | AAEL0119 | 3:350872 | 0 | 24 | FALSE | 6 | 12 | 6 | 382 | 428 | 914 |
| 3,03225E- | FALSE | High | IsMasterP | AAEL0121 | 2:405960 | 0 | 13 | FALSE | 5 | 15 | 5 | 514 | 573 | 614 |
| 2,43912E- | FALSE | High | IsMasterP | AAEL0112 | 1:215334 | 0 | 32 | FALSE | 6 | 16 | 6 | 224 | 250 | 605 |
| -7,4899E+ | FALSE | High | IsMasterP | AAEL0058 | NIGP0100 | 0 | 19 | FALSE | 5 | 16 | 5 | 344 | 396 | 607 |
| -7,74252E | FALSE | High | IsMasterP | AAEL0040 | 1:135385 | 0 | 49 | FALSE | 6 | 15 | 6 | 149 | 163 | 911 |
| 3,20445E- | FALSE | High | IsMasterP | AAEL0126 | 2:187551 | 0 | 22 | FALSE | 4 | 11 | 4 | 282 | 330 | 847 |
| 4,90487E- | FALSE | High | IsMasterP | AAEL0119 | 3:337658 | 0 | 24 | FALSE | 4 | 13 | 4 | 192 | 217 | 992 |
| -6,18508E | FALSE | High | IsMasterP | AAEL0048 | 2:115636 | 0 | 48 | FALSE | 5 | 16 | 5 | 183 | 216 | 524 |
| 7,27141E- | FALSE | High | IsMasterP | AAEL0063 | 1:100465 | 0 | 27 | FALSE | 5 | 13 | 5 | 307 | 352 | 502 |
| -6,71933E | FALSE | High | IsMasterP | AAEL0094 | 3:344953 | 0 | 31 | FALSE | 7 | 12 | 7 | 289 | 332 | 711 |
| 2,76586E- | FALSE | High | IsMasterP | AAEL0128 | 3:755857 | 0 | 45 | FALSE | 6 | 15 | 6 | 116 | 136 | 952 |

| | | | | | | | | | | | | | | |
|---|---|---|---|---|---|---|---|---|---|---|---|---|---|---|
| -7,83067E | FALSE | High | IsMasterP | AAEL0145 | 1:2243403 | 0 | 12 | FALSE | 4 | 15 | 4 | 465 | 485 | 512 |
| -4,23704E | FALSE | High | IsMasterP | AAEL0007 | 3:2451418 | 0 | 33 | FALSE | 2 | 10 | 2 | 130 | 151 | 456 |
| -6,26343E | FALSE | High | IsMasterP | AAEL0007 | 3:2461894 | 0 | 20 | FALSE | 6 | 12 | 6 | 413 | 454 | 486 |
| 5,05558E+ | FALSE | High | IsMasterP | AAEL0030 | 2:8520417 | 0 | 36 | FALSE | 8 | 12 | 8 | 370 | 405 | 684 |
| 1,86556E+ | FALSE | High | IsMasterP | AAEL0195 | 2:4264048 | 0 | 9 | FALSE | 3 | 13 | 3 | 510 | 580 | 587 |
| -5,3259E+ | FALSE | High | IsMasterP | AAEL0011 | 2:3075452 | 0 | 6 | FALSE | 7 | 14 | 7 | 2422 | 2649 | 662 |
| -6,80899E | FALSE | High | IsMasterP | AAEL0265 | 3:1072422 | 0 | 5 | FALSE | 6 | 12 | 6 | 2054 | 2312 | 502 |
| 8,79502E+ | FALSE | High | IsMasterP | AAEL0107 | 2:3046871 | 0 | 14 | FALSE | 5 | 14 | 5 | 310 | 335 | 667 |
| 7,33349E+ | FALSE | High | IsMasterP | AAEL0001 | 2:1100675 | 0 | 24 | FALSE | 7 | 13 | 7 | 481 | 525 | 938 |
| 4,28301E+ | FALSE | High | IsMasterP | AAEL0237 | NIGP0100 | 0 | 26 | FALSE | 5 | 10 | 1 | 386 | 421 | 459 |
| -9,18842E | FALSE | High | IsMasterP | AAEL0040 | 1:1357091 | 0 | 56 | FALSE | 6 | 12 | 6 | 228 | 261 | 510 |
| 6,80415E+ | FALSE | High | IsMasterP | AAEL0171 | 2:8678552 | 0 | 71 | FALSE | 4 | 13 | 4 | 79 | 90 | 944 |
| -5,3107E+ | FALSE | High | IsMasterP | AAEL0034 | 3:2213796 | 0 | 55 | FALSE | 6 | 15 | 6 | 154 | 174 | 892 |
| -6,02294E | FALSE | High | IsMasterP | AAEL0101 | 2:1619357 | 0 | 33 | FALSE | 2 | 11 | 2 | 149 | 159 | 494 |
| 4,68658E+ | FALSE | High | IsMasterP | AAEL0259 | 3:9707180 | 0 | 24 | FALSE | 4 | 14 | 4 | 208 | 245 | 601 |
| -4,65061E | FALSE | High | IsMasterP | AAEL0108 | 1:2180242 | 0 | 26 | FALSE | 6 | 14 | 5 | 300 | 329 | 974 |
| -3,45038E | FALSE | High | IsMasterP | AAEL0209 | 1:4391880 | 0 | 23 | FALSE | 5 | 15 | 5 | 442 | 503 | 536 |
| 4,68028E+ | FALSE | High | IsMasterP | AAEL0052 | 1:3120332 | 0 | 17 | FALSE | 6 | 11 | 6 | 592 | 655 | 917 |
| 4,00005E+ | FALSE | High | IsMasterP | AAEL0016 | 2:3003279 | 0 | 42 | FALSE | 7 | 14 | 7 | 231 | 257 | 498 |
| -2,00992E | FALSE | High | IsMasterP | AAEL0206 | 2:3448958 | 0 | 6 | FALSE | 8 | 14 | 8 | 2425 | 2706 | 681 |
| -6,18182E | FALSE | High | IsMasterP | AAEL0181 | 3:2212436 | 0 | 11 | FALSE | 8 | 12 | 8 | 1175 | 1302 | 777 |
| -4,46016E | FALSE | High | IsMasterP | AAEL0142 | 1:2857990 | 0 | 20 | FALSE | 11 | 14 | 11 | 822 | 912 | 533 |
| -5,78857E | FALSE | High | IsMasterP | AAEL0015 | 2:2161303 | 0 | 11 | FALSE | 3 | 14 | 3 | 408 | 456 | 931 |
| 9,18797E+ | FALSE | High | IsMasterP | AAEL0048 | 2:1159138 | 0 | 12 | FALSE | 9 | 15 | 9 | 1260 | 1398 | 693 |
| -5,32094E | FALSE | High | IsMasterP | AAEL0263 | NIGP0100 | 0 | 32 | FALSE | 10 | 15 | 1 | 567 | 625 | 725 |
| -3,69858E | FALSE | High | IsMasterP | AAEL0013 | 2:2044890 | 0 | 82 | FALSE | 5 | 14 | 5 | 95 | 107 | 652 |
| -8,88995E | FALSE | High | IsMasterP | AAEL0061 | 3:6856936 | 0 | 13 | FALSE | 8 | 12 | 8 | 1276 | 1442 | 578 |
| -2,45681E | FALSE | High | IsMasterP | AAEL0039 | 3:2539916 | 0 | 31 | FALSE | 10 | 17 | 10 | 353 | 385 | 895 |
| 3,69998E+ | FALSE | High | IsMasterP | AAEL0059 | 3:1739150 | 0 | 54 | FALSE | 5 | 16 | 5 | 118 | 127 | 994 |
| -7,9461E+ | FALSE | High | IsMasterP | AAEL0120 | 2:1866083 | 0 | 18 | FALSE | 6 | 14 | 6 | 510 | 566 | 594 |
| -4,18336E | FALSE | High | IsMasterP | AAEL0021 | 3:1790627 | 0 | 24 | FALSE | 4 | 14 | 4 | 197 | 219 | 1011 |

| | | | | | | | | | | | | | |
|---|---|---|---|---|---|---|---|---|---|---|---|---|---|
| 1,64405E | FALSE | High | IsMasterP | AAEL0272 2:422454 | 0 | 15 | FALSE | 7 | 13 | 7 | 878 | 973 | 559 |
| -6,50418E | FALSE | High | IsMasterP | AAEL0144 1:286793 | 0 | 25 | FALSE | 6 | 12 | 6 | 400 | 457 | 602 |
| 2,78316E | FALSE | High | IsMasterP | AAEL0074 2:134622 | 0 | 37 | FALSE | 4 | 14 | 4 | 139 | 160 | 724 |
| -1,45363E | FALSE | High | IsMasterP | AAEL0175 3:164683 | 0 | 10 | FALSE | 2 | 13 | 2 | 340 | 405 | 714 |
| 6,33487E | FALSE | High | IsMasterP | AAEL0105 2:909892 | 0 | 22 | FALSE | 6 | 14 | 6 | 353 | 417 | 526 |
| -1,42671E | FALSE | High | IsMasterP | AAEL0054 2:296504 | 0 | 27 | FALSE | 7 | 13 | 7 | 463 | 523 | 614 |
| 6,89228E | FALSE | High | IsMasterP | AAEL0031 1:193031 | 0 | 23 | FALSE | 7 | 13 | 7 | 457 | 500 | 720 |
| -4,26921E | FALSE | High | IsMasterP | AAEL0034 3:349263 | 0 | 19 | FALSE | 2 | 11 | 2 | 194 | 217 | 848 |
| -2,59971E | FALSE | High | IsMasterP | AAEL0106 2:135934 | 0 | 33 | FALSE | 10 | 14 | 10 | 491 | 543 | 759 |
| 2,43855E | FALSE | High | IsMasterP | AAEL0142 2:322844 | 0 | 7 | FALSE | 3 | 14 | 3 | 762 | 843 | 497 |
| 7,83606E | FALSE | High | IsMasterP | AAEL0113 3:276458 | 0 | 24 | FALSE | 4 | 19 | 4 | 330 | 364 | 983 |
| -8,40681E | FALSE | High | IsMasterP | AAEL0225 2:149932 | 0 | 13 | FALSE | 5 | 14 | 5 | 625 | 718 | 869 |
| -1,77232E | FALSE | High | IsMasterP | AAEL0249 2:167660 | 0 | 17 | FALSE | 8 | 16 | 8 | 656 | 747 | 702 |
| 8,25302E | FALSE | High | IsMasterP | AAEL0128 3:339280 | 0 | 15 | FALSE | 5 | 14 | 5 | 495 | 545 | 450 |
| 7,39782E | FALSE | High | IsMasterP | AAEL0112 1:223859 | 0 | 14 | FALSE | 7 | 14 | 7 | 607 | 688 | 928 |
| -1,05511E | FALSE | High | IsMasterP | AAEL0030 3:400736 | 0 | 46 | FALSE | 7 | 14 | 3 | 246 | 270 | 581 |
| -5,93023E | FALSE | High | IsMasterP | AAEL0008 3:363635 | 0 | 25 | FALSE | 7 | 12 | 7 | 437 | 490 | 568 |
| 1,31969E | FALSE | High | IsMasterP | AAEL0040 1:137014 | 0 | 25 | FALSE | 1 | 14 | 1 | 59 | 67 | 1043 |
| 5,14027E | FALSE | High | IsMasterP | AAEL0017 2:379131 | 0 | 16 | FALSE | 8 | 13 | 8 | 1056 | 1176 | 445 |
| -5,55531E | FALSE | High | IsMasterP | AAEL0277 NIGP0100 | 0 | 21 | FALSE | 3 | 14 | 2 | 201 | 206 | 1071 |
| 8,99382E | FALSE | High | IsMasterP | AAEL0099 2:931056 | 0 | 49 | FALSE | 3 | 14 | 3 | 82 | 92 | 992 |
| -4,62308E | FALSE | High | IsMasterP | AAEL0083 3:413796 | 0 | 29 | FALSE | 9 | 15 | 9 | 438 | 498 | 680 |
| -3,7804E+ | FALSE | High | IsMasterP | AAEL0015 2:262274 | 0 | 67 | FALSE | 8 | 10 | 8 | 254 | 274 | 482 |
| 1,32161E | FALSE | High | IsMasterP | AAEL0027 2:138160 | 0 | 55 | FALSE | 5 | 9 | 5 | 157 | 172 | 901 |
| 4,93113E | FALSE | High | IsMasterP | AAEL0066 2:325789 | 0 | 8 | FALSE | 2 | 13 | 2 | 349 | 381 | 922 |
| -8,74717E | FALSE | High | IsMasterP | AAEL0078 2:271829 | 0 | 37 | FALSE | 5 | 12 | 5 | 187 | 220 | 512 |
| -6,05806E | FALSE | High | IsMasterP | AAEL0103 3:372303 | 0 | 28 | FALSE | 7 | 14 | 7 | 319 | 343 | 525 |
| 6,34995E | FALSE | High | IsMasterP | AAEL0197 3:369484 | 0 | 44 | FALSE | 2 | 10 | 1 | 94 | 101 | 840 |
| 8,99747E | FALSE | High | IsMasterP | AAEL0117 2:168663 | 0 | 8 | FALSE | 3 | 13 | 2 | 689 | 782 | 637 |
| 5,51616E | FALSE | High | IsMasterP | AAEL0128 1:222074 | 0 | 26 | FALSE | 5 | 15 | 4 | 238 | 274 | 521 |
| 8,80431E | FALSE | High | IsMasterP | AAEL0182 1:294601 | 0 | 7 | FALSE | 5 | 13 | 5 | 930 | 1015 | 629 |

| | | | | | | | | | | | | | | |
|---|---|---|---|---|---|---|---|---|---|---|---|---|---|---|
| -4,52101E | FALSE | High | IsMasterP | AAEL0135 | 2:375534 | 0 | 53 | FALSE | 3 | 13 | 3 | 107 | 117 | 463 |
| 6,96943E | FALSE | High | IsMasterP | AAEL0270 | 2:205130 | 0 | 27 | FALSE | 4 | 11 | 4 | 301 | 325 | 917 |
| -1,01069E | FALSE | High | IsMasterP | AAEL0151 | 2:233026 | 0 | 12 | FALSE | 6 | 16 | 6 | 611 | 696 | 467 |
| -2,11402E | FALSE | High | IsMasterP | AAEL0026 | 3:256668 | 0 | 15 | FALSE | 6 | 13 | 6 | 631 | 694 | 470 |
| 1,62102E | FALSE | High | IsMasterP | AAEL0054 | 1:805240 | 0 | 12 | FALSE | 6 | 12 | 6 | 904 | 1019 | 664 |
| 4,42189E | FALSE | High | IsMasterP | AAEL0060 | 1:184165 | 0 | 32 | FALSE | 4 | 11 | 4 | 174 | 190 | 963 |
| 7,25428E | FALSE | High | IsMasterP | AAEL0019 | 3:226914 | 0 | 15 | FALSE | 4 | 12 | 4 | 481 | 545 | 541 |
| -2,53637E | FALSE | High | IsMasterP | AAEL0171 | 3:221580 | 0 | 27 | FALSE | 6 | 14 | 6 | 227 | 223 | 1095 |
| 6,06022E | FALSE | High | IsMasterP | AAEL0132 | 3:340871 | 0 | 23 | FALSE | 5 | 11 | 5 | 320 | 357 | 843 |
| -6,4324E+ | FALSE | High | IsMasterP | AAEL0000 | 3:105159 | 0 | 27 | FALSE | 6 | 15 | 6 | 360 | 397 | 533 |
| -7,63411E | FALSE | High | IsMasterP | AAEL0040 | 3:166159 | 0 | 10 | FALSE | 7 | 11 | 7 | 1180 | 1341 | 681 |
| 8,38855E | FALSE | High | IsMasterP | AAEL0001 | 3:212830 | 0 | 40 | FALSE | 4 | 10 | 4 | 153 | 176 | 837 |
| 7,85536E | FALSE | High | IsMasterP | AAEL0028 | 1:255908 | 0 | 69 | FALSE | 5 | 16 | 5 | 83 | 94 | 954 |
| 7,90608E | FALSE | High | IsMasterP | AAEL0013 | 3:489633 | 0 | 26 | FALSE | 4 | 10 | 4 | 235 | 261 | 563 |
| -5,26669E | FALSE | High | IsMasterP | AAEL0000 | 3:104360 | 0 | 12 | FALSE | 6 | 13 | 6 | 596 | 656 | 489 |
| -5,52167E | FALSE | High | IsMasterP | AAEL0201 | 2:407754 | 0 | 17 | FALSE | 6 | 14 | 5 | 387 | 438 | 629 |
| 5,67199E | FALSE | High | IsMasterP | AAEL0008 | 3:120508 | 0 | 21 | FALSE | 5 | 17 | 5 | 172 | 189 | 1130 |
| 4,47348E | FALSE | High | IsMasterP | AAEL0098 | 3:186492 | 0 | 30 | FALSE | 2 | 12 | 2 | 128 | 137 | 512 |
| -7,43668E | FALSE | High | IsMasterP | AAEL0014 | 2:263974 | 0 | 4 | FALSE | 7 | 14 | 7 | 3303 | 3666 | 633 |
| 8,09492E | FALSE | High | IsMasterP | AAEL0042 | 2:118670 | 0 | 46 | FALSE | 8 | 13 | 8 | 141 | 172 | 976 |
| 5,5863E+ | FALSE | High | IsMasterP | AAEL0034 | 2:151328 | 0 | 32 | FALSE | 6 | 15 | 6 | 247 | 281 | 487 |
| 1,85497E | FALSE | High | IsMasterP | AAEL0260 | 2:648484 | 0 | 26 | FALSE | 5 | 13 | 5 | 174 | 200 | 527 |
| -7,48194E | FALSE | High | IsMasterP | AAEL0019 | 3:227013 | 0 | 21 | FALSE | 4 | 12 | 4 | 160 | 182 | 643 |
| -8,74648E | FALSE | High | IsMasterP | AAEL0014 | 2:261693 | 0 | 10 | FALSE | 8 | 14 | 8 | 1213 | 1337 | 544 |
| -4,43248E | FALSE | High | IsMasterP | AAEL0091 | 2:418853 | 0 | 22 | FALSE | 8 | 13 | 8 | 520 | 590 | 802 |
| -2,83287E | FALSE | High | IsMasterP | AAEL0199 | 3:453034 | 0 | 14 | FALSE | 8 | 13 | 8 | 891 | 1022 | 549 |
| -5,36503E | FALSE | High | IsMasterP | AAEL0211 | 2:193068 | 0 | 26 | FALSE | 5 | 10 | 5 | 268 | 302 | 923 |
| 4,44749E | FALSE | High | IsMasterP | AAEL0039 | 3:254347 | 0 | 23 | FALSE | 3 | 14 | 3 | 104 | 125 | 1080 |
| -8,57443E | FALSE | High | IsMasterP | AAEL0062 | 3:628702 | 0 | 58 | FALSE | 2 | 11 | 2 | 86 | 93 | 925 |
| -1,10731E | FALSE | High | IsMasterP | AAEL0058 | 1:201231 | 0 | 9 | FALSE | 7 | 15 | 7 | 1227 | 1409 | 762 |
| 2,50205E | FALSE | High | IsMasterP | AAEL0043 | 3:366710 | 0 | 39 | FALSE | 8 | 11 | 8 | 223 | 253 | 802 |

| | | | | | | | | | | | | | | |
|---|---|---|---|---|---|---|---|---|---|---|---|---|---|---|
| 3,641E+18 | FALSE | High | IsMasterP | AAEL0198 | 3:143689 | 0 | 9 | FALSE | 7 | 15 | 7 | 1162 | 1318 | 582 |
| -3,75333E | FALSE | High | IsMasterP | AAEL0041 | 3:153720 | 0 | 30 | FALSE | 4 | 10 | 4 | 272 | 315 | 655 |
| -2,00564E | FALSE | High | IsMasterP | AAEL0147 | 1:360481 | 0 | 34 | FALSE | 3 | 10 | 3 | 201 | 228 | 774 |
| -9,10914E | FALSE | High | IsMasterP | AAEL0005 | 3:388045 | 0 | 39 | FALSE | 3 | 9 | 3 | 180 | 201 | 569 |
| 5,91242E- | FALSE | High | IsMasterP | AAEL0087 | 3:189389 | 0 | 58 | FALSE | 4 | 13 | 4 | 76 | 85 | 884 |
| -6,8431E+ | FALSE | High | IsMasterP | AAEL0077 | 1:718574 | 0 | 41 | FALSE | 3 | 15 | 3 | 90 | 98 | 954 |
| 5,55109E- | FALSE | High | IsMasterP | AAEL0073 | 2:234447 | 0 | 12 | FALSE | 6 | 11 | 6 | 695 | 805 | 492 |
| -1,54348E | FALSE | High | IsMasterP | AAEL0096 | 2:169945 | 0 | 26 | FALSE | 5 | 14 | 5 | 180 | 206 | 848 |
| -5,54891E | FALSE | High | IsMasterP | AAEL0027 | 3:125563 | 0 | 40 | FALSE | 5 | 9 | 5 | 219 | 240 | 680 |
| -4,92035E | FALSE | High | IsMasterP | AAEL0102 | 2:402958 | 0 | 40 | FALSE | 4 | 12 | 4 | 146 | 176 | 941 |
| 3,66432E- | FALSE | High | IsMasterP | AAEL0128 | 1:222054 | 0 | 39 | FALSE | 6 | 12 | 5 | 239 | 272 | 625 |
| -6,85052E | FALSE | High | IsMasterP | AAEL0092 | 2:171962 | 0 | 14 | FALSE | 4 | 10 | 4 | 358 | 395 | 533 |
| 7,0422E+1 | FALSE | High | IsMasterP | AAEL0112 | 2:226716 | 0 | 10 | FALSE | 6 | 12 | 6 | 812 | 918 | 730 |
| 1,73732E- | FALSE | High | IsMasterP | AAEL0227 | 2:147264 | 0 | 28 | FALSE | 5 | 14 | 5 | 199 | 225 | 613 |
| -3,38887E | FALSE | High | IsMasterP | AAEL0076 | 2:470765 | 0 | 14 | FALSE | 6 | 12 | 3 | 647 | 724 | 709 |
| 8,4288E+1 | FALSE | High | IsMasterP | AAEL0053 | 2:271210 | 0 | 25 | FALSE | 6 | 13 | 6 | 332 | 353 | 791 |
| 7,48981E- | FALSE | High | IsMasterP | AAEL0017 | 2:374461 | 0 | 34 | FALSE | 5 | 10 | 5 | 225 | 245 | 750 |
| 6,13699E- | FALSE | High | IsMasterP | AAEL0016 | 2:854021 | 0 | 25 | FALSE | 6 | 10 | 6 | 325 | 374 | 545 |
| -5,4381E+ | FALSE | High | IsMasterP | AAEL0113 | 2:331548 | 0 | 14 | FALSE | 4 | 13 | 4 | 458 | 518 | 812 |
| 6,36769E- | FALSE | High | IsMasterP | AAEL0112 | 1:344710 | 0 | 23 | FALSE | 6 | 12 | 6 | 409 | 464 | 539 |
| -8,87384E | FALSE | High | IsMasterP | AAEL0087 | 3:281179 | 0 | 42 | FALSE | 4 | 14 | 4 | 103 | 119 | 920 |
| 4,14109E- | FALSE | High | IsMasterP | AAEL0136 | 2:324269 | 0 | 16 | FALSE | 4 | 14 | 4 | 255 | 293 | 998 |
| -3,73738E | FALSE | High | IsMasterP | AAEL0126 | 1:146149 | 0 | 15 | FALSE | 6 | 13 | 6 | 552 | 613 | 995 |
| 7,22154E- | FALSE | High | IsMasterP | AAEL0059 | 3:138646 | 0 | 14 | FALSE | 5 | 12 | 5 | 511 | 565 | 684 |
| 2,99209E- | FALSE | High | IsMasterP | AAEL0084 | 2:191814 | 0 | 50 | FALSE | 4 | 10 | 4 | 154 | 176 | 662 |
| -2,65137E | FALSE | High | IsMasterP | AAEL0082 | 3:628315 | 0 | 9 | FALSE | 3 | 9 | 3 | 631 | 696 | 938 |
| -6,33648E | FALSE | High | IsMasterP | AAEL0221 | 2:192364 | 0 | 18 | FALSE | 7 | 15 | 1 | 489 | 574 | 988 |
| 2,75315E- | FALSE | High | IsMasterP | AAEL0063 | 3:231090 | 0 | 30 | FALSE | 7 | 12 | 7 | 334 | 374 | 895 |
| -5,30449E | FALSE | High | IsMasterP | AAEL0045 | 1:144943 | 0 | 14 | FALSE | 8 | 11 | 8 | 956 | 1070 | 609 |
| -5,05008E | FALSE | High | IsMasterP | AAEL0277 | NIGP0100 | 0 | 17 | FALSE | 1 | 9 | 1 | 136 | 149 | 723 |
| 4,65721E- | FALSE | High | IsMasterP | AAEL0030 | 3:400736 | 0 | 41 | FALSE | 7 | 12 | 3 | 246 | 272 | 753 |

| | | | | | | | | | | | | | | |
|---|---|---|---|---|---|---|---|---|---|---|---|---|---|---|
| -3,54075E | FALSE | High | IsMasterP | AAEL0087 | 3:1327323 | 0 | 33 | FALSE | 7 | 14 | 7 | 200 | 228 | 920 |
| 2,56291E+ | FALSE | High | IsMasterP | AAEL0131 | 2:1793341 | 0 | 27 | FALSE | 4 | 13 | 4 | 177 | 201 | 514 |
| 7,07542E+ | FALSE | High | IsMasterP | AAEL0090 | 2:1016834 | 0 | 32 | FALSE | 5 | 14 | 5 | 182 | 210 | 772 |
| 3,54095E+ | FALSE | High | IsMasterP | AAEL0134 | 1:1549066 | 0 | 26 | FALSE | 5 | 11 | 5 | 290 | 323 | 683 |
| -9,33212E | FALSE | High | IsMasterP | AAEL0082 | 2:6725721 | 0 | 56 | FALSE | 3 | 9 | 3 | 136 | 145 | 463 |
| 3,28815E+ | FALSE | High | IsMasterP | AAEL0125 | 2:3280394 | 0 | 39 | FALSE | 8 | 12 | 8 | 288 | 339 | 621 |
| -7,03062E | FALSE | High | IsMasterP | AAEL0227 | NIGP0100 | 0 | 41 | FALSE | 3 | 11 | 3 | 135 | 141 | 970 |
| 3,97252E+ | FALSE | High | IsMasterP | AAEL0046 | 1:1458540 | 0 | 26 | FALSE | 7 | 11 | 7 | 514 | 561 | 762 |
| -8,2817E+ | FALSE | High | IsMasterP | AAEL0109 | 1:1396691 | 0 | 17 | FALSE | 4 | 11 | 4 | 324 | 345 | 614 |
| -8,28419E | FALSE | High | IsMasterP | AAEL0021 | 1:1022335 | 0 | 22 | FALSE | 6 | 13 | 6 | 448 | 493 | 541 |
| 7,85188E+ | FALSE | High | IsMasterP | AAEL0104 | 1:2679712 | 0 | 22 | FALSE | 5 | 14 | 5 | 313 | 352 | 689 |
| -2,18844E | FALSE | High | IsMasterP | AAEL0027 | 3:1293107 | 0 | 8 | FALSE | 5 | 10 | 5 | 995 | 1080 | 989 |
| -9,52952E | FALSE | High | IsMasterP | AAEL0111 | 3:1936108 | 0 | 27 | FALSE | 4 | 12 | 4 | 285 | 318 | 966 |
| 8,19713E+ | FALSE | High | IsMasterP | AAEL0044 | 3:2122800 | 0 | 41 | FALSE | 7 | 14 | 7 | 227 | 265 | 580 |
| 2,94522E+ | FALSE | High | IsMasterP | AAEL0183 | 2:3878376 | 0 | 3 | FALSE | 5 | 11 | 5 | 2438 | 2662 | 644 |
| -5,02747E | FALSE | High | IsMasterP | AAEL0107 | 3:1560366 | 0 | 40 | FALSE | 3 | 9 | 3 | 167 | 192 | 543 |
| 3,03502E+ | FALSE | High | IsMasterP | AAEL0141 | 3:3022274 | 0 | 14 | FALSE | 2 | 11 | 2 | 259 | 277 | 534 |
| 4,44625E+ | FALSE | High | IsMasterP | AAEL0016 | 1:1382962 | 0 | 22 | FALSE | 6 | 14 | 6 | 251 | 284 | 526 |
| -4,53554E | FALSE | High | IsMasterP | AAEL0050 | 1:2986420 | 0 | 29 | FALSE | 5 | 8 | 5 | 287 | 308 | 844 |
| -5,73603E | FALSE | High | IsMasterP | AAEL0137 | 2:3526783 | 0 | 72 | FALSE | 4 | 12 | 4 | 117 | 135 | 554 |
| 1,51497E+ | FALSE | High | IsMasterP | AAEL0110 | 2:1771503 | 0 | 7 | FALSE | 5 | 13 | 5 | 1115 | 1253 | 517 |
| 2,75572E+ | FALSE | High | IsMasterP | AAEL0055 | 3:4051322 | 0 | 19 | FALSE | 5 | 12 | 5 | 367 | 410 | 881 |
| 6,2925E+1 | FALSE | High | IsMasterP | AAEL0024 | 1:1768719 | 0 | 15 | FALSE | 4 | 11 | 4 | 522 | 597 | 802 |
| 6,19748E+ | FALSE | High | IsMasterP | AAEL0052 | 2:6931898 | 0 | 14 | FALSE | 8 | 12 | 8 | 843 | 940 | 664 |
| 6,94325E+ | FALSE | High | IsMasterP | AAEL0217 | NIGP0100 | 0 | 69 | FALSE | 8 | 12 | 8 | 151 | 176 | 917 |
| -3,72528E | FALSE | High | IsMasterP | AAEL0128 | 2:1054406 | 0 | 18 | FALSE | 8 | 9 | 8 | 904 | 998 | 654 |
| 6,97412E+ | FALSE | High | IsMasterP | AAEL0140 | 1:1483330 | 0 | 12 | FALSE | 5 | 9 | 5 | 846 | 965 | 708 |
| 8,60001E+ | FALSE | High | IsMasterP | AAEL0008 | 3:3359356 | 0 | 32 | FALSE | 8 | 11 | 8 | 514 | 564 | 676 |
| -8,56181E | FALSE | High | IsMasterP | AAEL0100 | 3:1120281 | 0 | 21 | FALSE | 8 | 13 | 8 | 375 | 446 | 524 |
| 5,88016E+ | FALSE | High | IsMasterP | AAEL0057 | 2:4373018 | 0 | 9 | FALSE | 2 | 10 | 2 | 402 | 438 | 894 |
| -7,28244E | FALSE | High | IsMasterP | AAEL0120 | 3:1473241 | 0 | 19 | FALSE | 6 | 11 | 6 | 417 | 482 | 606 |

| | | | | | | | | | | | | | | |
|---|---|---|---|---|---|---|---|---|---|---|---|---|---|---|
| -2,99892E | FALSE | High | IsMasterP | AAEL0121 | 2:204221 | 0 | 28 | FALSE | 6 | 11 | 5 | 452 | 493 | 868 |
| 7,32659E | FALSE | High | IsMasterP | AAEL0044 | 2:453197 | 0 | 45 | FALSE | 2 | 9 | 2 | 76 | 93 | 843 |
| 4,25915E | FALSE | High | IsMasterP | AAEL0215 | 1:601675 | 0 | 4 | FALSE | 5 | 11 | 5 | 2004 | 2233 | 643 |
| -6,14158E | FALSE | High | IsMasterP | AAEL0046 | 2:767572 | 0 | 12 | FALSE | 5 | 13 | 5 | 616 | 693 | 574 |
| -7,34E+18 | FALSE | High | IsMasterP | AAEL0090 | 2:411473 | 0 | 25 | FALSE | 4 | 11 | 4 | 189 | 220 | 942 |
| 4,97047E | FALSE | High | IsMasterP | AAEL0018 | 2:375256 | 0 | 19 | FALSE | 4 | 11 | 4 | 318 | 356 | 482 |
| -9,30674E | FALSE | High | IsMasterP | AAEL0083 | 3:224886 | 0 | 6 | FALSE | 4 | 12 | 4 | 975 | 1057 | 952 |
| -4,94349E | FALSE | High | IsMasterP | AAEL0246 | NIGP0100 | 0 | 32 | FALSE | 6 | 12 | 6 | 358 | 381 | 679 |
| -1,5504E+ | FALSE | High | IsMasterP | AAEL0004 | 3:184194 | 0 | 22 | FALSE | 7 | 12 | 7 | 530 | 559 | 683 |
| 4,24266E | FALSE | High | IsMasterP | AAEL0213 | 1:735042 | 0 | 24 | FALSE | 6 | 13 | 6 | 348 | 397 | 494 |
| 8,38301E | FALSE | High | IsMasterP | AAEL0011 | 2:307234 | 0 | 32 | FALSE | 8 | 12 | 8 | 336 | 387 | 508 |
| -7,24343E | FALSE | High | IsMasterP | AAEL0075 | NIGP0100 | 0 | 8 | FALSE | 3 | 11 | 3 | 623 | 728 | 562 |
| 3,12517E | FALSE | High | IsMasterP | AAEL0245 | 2:132653 | 0 | 16 | FALSE | 3 | 10 | 3 | 305 | 350 | 503 |
| 7,0604E+ | FALSE | High | IsMasterP | AAEL0103 | 2:404119 | 0 | 21 | FALSE | 4 | 14 | 4 | 177 | 204 | 1155 |
| 2,2188E+ | FALSE | High | IsMasterP | AAEL0280 | 1:261466 | 0 | 33 | FALSE | 5 | 9 | 5 | 301 | 337 | 454 |
| -8,08942E | FALSE | High | IsMasterP | AAEL0089 | 3:356273 | 0 | 32 | FALSE | 4 | 8 | 4 | 250 | 260 | 803 |
| 4,21865E | FALSE | High | IsMasterP | AAEL0169 | 2:294820 | 0 | 17 | FALSE | 6 | 10 | 6 | 611 | 706 | 506 |
| 4,48822E | FALSE | High | IsMasterP | AAEL0098 | 2:164054 | 0 | 21 | FALSE | 4 | 14 | 4 | 130 | 143 | 805 |
| 2,61949E | FALSE | High | IsMasterP | AAEL0265 | NIGP0100 | 0 | 53 | FALSE | 3 | 10 | 3 | 74 | 81 | 712 |
| -6,02899E | FALSE | High | IsMasterP | AAEL0230 | 3:312596 | 0 | 10 | FALSE | 5 | 9 | 5 | 1004 | 1122 | 638 |
| -3,94733E | FALSE | High | IsMasterP | AAEL0107 | 3:294088 | 0 | 22 | FALSE | 3 | 14 | 3 | 107 | 125 | 999 |
| 6,18144E | FALSE | High | IsMasterP | AAEL0000 | 3:101774 | 0 | 41 | FALSE | 5 | 9 | 5 | 242 | 268 | 600 |
| 1,33592E | FALSE | High | IsMasterP | AAEL0050 | 3:827807 | 0 | 10 | FALSE | 3 | 9 | 3 | 508 | 574 | 775 |
| -6,19499E | FALSE | High | IsMasterP | AAEL0056 | 2:132297 | 0 | 9 | FALSE | 3 | 12 | 3 | 412 | 462 | 549 |
| 7,95754E | FALSE | High | IsMasterP | AAEL0262 | 1:575052 | 0 | 7 | FALSE | 5 | 10 | 5 | 1151 | 1283 | 569 |
| 4,24049E | FALSE | High | IsMasterP | AAEL0101 | 3:148829 | 0 | 18 | FALSE | 3 | 13 | 2 | 208 | 239 | 878 |
| -5,16769E | FALSE | High | IsMasterP | AAEL0148 | 3:322210 | 0 | 75 | FALSE | 3 | 10 | 3 | 67 | 76 | 777 |
| 2,27973E | FALSE | High | IsMasterP | AAEL0099 | 2:929741 | 0 | 15 | FALSE | 4 | 9 | 4 | 546 | 603 | 531 |
| 5,22694E | FALSE | High | IsMasterP | AAEL0013 | 2:205714 | 0 | 19 | FALSE | 7 | 11 | 7 | 574 | 623 | 648 |
| 8,63583E | FALSE | High | IsMasterP | AAEL0219 | 1:215168 | 0 | 5 | FALSE | 5 | 11 | 5 | 1393 | 1584 | 708 |
| 3,31052E | FALSE | High | IsMasterP | AAEL0120 | 3:160446 | 0 | 57 | FALSE | 6 | 12 | 6 | 190 | 216 | 783 |

| | | | | | | | | | | | | | | |
|---|---|---|---|---|---|---|---|---|---|---|---|---|---|---|
| -5,89322E | FALSE | High | IsMasterP | AAEL0063 | 2:140845 | 0 | 15 | FALSE | 5 | 11 | 5 | 640 | 702 | 796 |
| 3,11842E | FALSE | High | IsMasterP | AAEL0232 | 2:279284 | 0 | 21 | FALSE | 3 | 9 | 3 | 194 | 211 | 863 |
| 7,26522E | FALSE | High | IsMasterP | AAEL0052 | 3:888649 | 0 | 12 | FALSE | 6 | 11 | 6 | 735 | 838 | 960 |
| 5,57609E | FALSE | High | IsMasterP | AAEL0140 | 2:406005 | 0 | 11 | FALSE | 7 | 11 | 7 | 772 | 895 | 673 |
| 6,46851E | FALSE | High | IsMasterP | AAEL0145 | 1:245236 | 0 | 19 | FALSE | 7 | 13 | 7 | 543 | 620 | 629 |
| -4,93693E | FALSE | High | IsMasterP | AAEL0014 | 2:478150 | 0 | 9 | FALSE | 4 | 9 | 4 | 856 | 946 | 790 |
| 7,53891E | FALSE | High | IsMasterP | AAEL0074 | 1:216391 | 0 | 40 | FALSE | 3 | 9 | 3 | 189 | 220 | 506 |
| -1,68029E | FALSE | High | IsMasterP | AAEL0083 | 2:274087 | 0 | 19 | FALSE | 4 | 12 | 4 | 305 | 343 | 698 |
| -7,79038E | FALSE | High | IsMasterP | AAEL0120 | 3:147198 | 0 | 28 | FALSE | 6 | 9 | 6 | 556 | 582 | 671 |
| -1,73481E | FALSE | High | IsMasterP | AAEL0197 | 2:443365 | 0 | 9 | FALSE | 4 | 10 | 4 | 857 | 932 | 451 |
| 5,3335E+ | FALSE | High | IsMasterP | AAEL0132 | 2:186023 | 0 | 7 | FALSE | 6 | 13 | 6 | 1296 | 1480 | 524 |
| 2,2681E+ | FALSE | High | IsMasterP | AAEL0056 | 2:382118 | 0 | 12 | FALSE | 4 | 10 | 4 | 369 | 395 | 496 |
| -2,66023E | FALSE | High | IsMasterP | AAEL0011 | 3:407100 | 0 | 30 | FALSE | 5 | 9 | 5 | 344 | 380 | 484 |
| -4,25511E | FALSE | High | IsMasterP | AAEL0112 | 1:223871 | 0 | 15 | FALSE | 3 | 11 | 3 | 400 | 440 | 470 |
| -3,80328E | FALSE | High | IsMasterP | AAEL0102 | 3:279118 | 0 | 28 | FALSE | 4 | 9 | 4 | 192 | 211 | 397 |
| 3,30698E | FALSE | High | IsMasterP | AAEL0014 | 2:264573 | 0 | 24 | FALSE | 4 | 12 | 4 | 303 | 342 | 888 |
| 6,72228E | FALSE | High | IsMasterP | AAEL0135 | 2:246095 | 0 | 12 | FALSE | 7 | 11 | 7 | 1088 | 1204 | 640 |
| 9,00035E | FALSE | High | IsMasterP | AAEL0126 | 1:279337 | 0 | 13 | FALSE | 5 | 13 | 5 | 341 | 394 | 1124 |
| 2,99447E | FALSE | High | IsMasterP | AAEL0011 | 2:307204 | 0 | 16 | FALSE | 6 | 13 | 6 | 324 | 359 | 594 |
| -5,07144E | FALSE | High | IsMasterP | AAEL0111 | 3:193660 | 0 | 53 | FALSE | 5 | 15 | 5 | 100 | 114 | 1052 |
| 3,78469E | FALSE | High | IsMasterP | AAEL0112 | 2:194558 | 0 | 26 | FALSE | 5 | 13 | 5 | 248 | 272 | 859 |
| 3,84312E | FALSE | High | IsMasterP | AAEL0203 | 2:111287 | 0 | 22 | FALSE | 4 | 10 | 4 | 249 | 282 | 646 |
| 4,52832E | FALSE | High | IsMasterP | AAEL0053 | 3:344934 | 0 | 57 | FALSE | 5 | 8 | 5 | 97 | 108 | 655 |
| -5,22873E | FALSE | High | IsMasterP | AAEL0026 | 2:433627 | 0 | 25 | FALSE | 4 | 10 | 4 | 286 | 324 | 834 |
| -2,24763E | FALSE | High | IsMasterP | AAEL0013 | 2:204979 | 0 | 18 | FALSE | 4 | 12 | 4 | 372 | 426 | 496 |
| -3,5255E+ | FALSE | High | IsMasterP | AAEL0044 | 3:109683 | 0 | 15 | FALSE | 5 | 12 | 5 | 508 | 569 | 945 |
| -8,36699E | FALSE | High | IsMasterP | AAEL0122 | 2:384431 | 0 | 14 | FALSE | 5 | 12 | 5 | 478 | 539 | 569 |
| 5,04544E | FALSE | High | IsMasterP | AAEL0029 | 1:261378 | 0 | 23 | FALSE | 7 | 10 | 7 | 421 | 475 | 616 |
| 4,52918E | FALSE | High | IsMasterP | AAEL0006 | 3:395191 | 0 | 30 | FALSE | 4 | 8 | 4 | 282 | 318 | 644 |
| -6,19243E | FALSE | High | IsMasterP | AAEL0123 | 2:102906 | 0 | 66 | FALSE | 5 | 12 | 5 | 124 | 136 | 843 |
| -1,02298E | FALSE | High | IsMasterP | AAEL0129 | 2:124739 | 0 | 55 | FALSE | 4 | 9 | 4 | 163 | 180 | 580 |

| | | | | | | | | | | | | | | |
|---|---|---|---|---|---|---|---|---|---|---|---|---|---|---|
| -5,938E+1 | FALSE | High | IsMasterP | AAEL0235 | 3:3023145 | 0 | 19 | FALSE | 5 | 11 | 5 | 406 | 445 | 455 |
| -5,56524E | FALSE | High | IsMasterP | AAEL0037 | 1:2483851 | 0 | 11 | FALSE | 5 | 10 | 5 | 915 | 1024 | 595 |
| -8,5453E+ | FALSE | High | IsMasterP | AAEL0030 | 2:2133899 | 0 | 15 | FALSE | 6 | 11 | 6 | 723 | 812 | 827 |
| -3,30111E | FALSE | High | IsMasterP | AAEL0103 | 2:1830232 | 0 | 8 | FALSE | 5 | 12 | 4 | 1267 | 1400 | 657 |
| 7,10076E- | FALSE | High | IsMasterP | AAEL0032 | 2:3667056 | 0 | 22 | FALSE | 4 | 9 | 4 | 321 | 354 | 784 |
| -4,12436E | FALSE | High | IsMasterP | AAEL0045 | 2:3100445 | 0 | 31 | FALSE | 3 | 9 | 3 | 159 | 176 | 644 |
| -1,87257E | FALSE | High | IsMasterP | AAEL0026 | 3:2572629 | 0 | 19 | FALSE | 5 | 10 | 5 | 383 | 419 | 488 |
| -8,92492E | FALSE | High | IsMasterP | AAEL0072 | 3:1955801 | 0 | 7 | FALSE | 3 | 12 | 3 | 353 | 406 | 869 |
| -3,07096E | FALSE | High | IsMasterP | AAEL0048 | 2:1157255 | 0 | 51 | FALSE | 5 | 9 | 5 | 123 | 136 | 857 |
| -4,971E+1 | FALSE | High | IsMasterP | AAEL0012 | 2:1530549 | 0 | 13 | FALSE | 5 | 9 | 5 | 592 | 697 | 555 |
| 5,73915E- | FALSE | High | IsMasterP | AAEL0129 | 2:3865635 | 0 | 7 | FALSE | 5 | 10 | 5 | 920 | 998 | 711 |
| 1,2554E+1 | FALSE | High | IsMasterP | AAEL0195 | 2:1387190 | 0 | 12 | FALSE | 8 | 10 | 8 | 1215 | 1351 | 553 |
| -2,77672E | FALSE | High | IsMasterP | AAEL0048 | 1:2884200 | 0 | 22 | FALSE | 6 | 11 | 5 | 442 | 482 | 850 |
| -3,59922E | FALSE | High | IsMasterP | AAEL0195 | 3:1300912 | 0 | 2 | FALSE | 2 | 11 | 2 | 1535 | 1676 | 619 |
| -9,06542E | FALSE | High | IsMasterP | AAEL0170 | 2:2629205 | 0 | 26 | FALSE | 3 | 9 | 3 | 243 | 271 | 1165 |
| -1,08844E | FALSE | High | IsMasterP | AAEL0015 | 2:2241004 | 0 | 55 | FALSE | 3 | 10 | 3 | 75 | 88 | 654 |
| 6,09017E- | FALSE | High | IsMasterP | AAEL0108 | 2:1280469 | 0 | 14 | FALSE | 4 | 9 | 4 | 501 | 542 | 847 |
| 1,8539E+1 | FALSE | High | IsMasterP | AAEL0033 | 3:8003037 | 0 | 10 | FALSE | 5 | 7 | 5 | 1466 | 1541 | 667 |
| -4,0941E+ | FALSE | High | IsMasterP | AAEL0046 | 3:2466320 | 0 | 14 | FALSE | 4 | 10 | 4 | 418 | 481 | 783 |
| -2,84726E | FALSE | High | IsMasterP | AAEL0092 | 2:1126400 | 0 | 27 | FALSE | 4 | 10 | 4 | 258 | 291 | 960 |
| -7,00905E | FALSE | High | IsMasterP | AAEL0000 | 3:1044793 | 0 | 10 | FALSE | 3 | 10 | 3 | 517 | 599 | 829 |
| 9,78871E- | FALSE | High | IsMasterP | AAEL0109 | 2:1084962 | 0 | 38 | FALSE | 5 | 10 | 5 | 140 | 166 | 831 |
| -4,2965E+ | FALSE | High | IsMasterP | AAEL0009 | 3:3801158 | 0 | 17 | FALSE | 2 | 10 | 2 | 215 | 224 | 712 |
| -2,99877E | FALSE | High | IsMasterP | AAEL0105 | 1:3013179 | 0 | 13 | FALSE | 3 | 12 | 3 | 436 | 475 | 630 |
| 7,60688E- | FALSE | High | IsMasterP | AAEL0122 | 1:7690233 | 0 | 35 | FALSE | 4 | 8 | 4 | 225 | 256 | 534 |
| 3,42232E- | FALSE | High | IsMasterP | AAEL0115 | 2:1231676 | 0 | 15 | FALSE | 4 | 9 | 4 | 549 | 606 | 599 |
| 6,33872E- | FALSE | High | IsMasterP | AAEL0186 | MT:2903:3 | 0 | 16 | FALSE | 2 | 8 | 2 | 228 | 265 | 568 |
| 1,88694E- | FALSE | High | IsMasterP | AAEL0231 | 2:3517954 | 0 | 3 | FALSE | 3 | 8 | 3 | 1777 | 1751 | 876 |
| -3,83605E | FALSE | High | IsMasterP | AAEL0241 | 2:1158305 | 0 | 8 | FALSE | 5 | 9 | 5 | 1184 | 1285 | 580 |
| -3,46813E | FALSE | High | IsMasterP | AAEL0171 | 2:3558832 | 0 | 12 | FALSE | 2 | 8 | 2 | 171 | 192 | 884 |
| -6,76289E | FALSE | High | IsMasterP | AAEL0089 | 1:8117864 | 0 | 19 | FALSE | 4 | 7 | 4 | 693 | 732 | 837 |

| | | | | | | | | | | | | | | |
|---|---|---|---|---|---|---|---|---|---|---|---|---|---|---|
| -3,89027E | FALSE | High | IsMasterP | AAEL0266 | 1:142994( | 0 | 6 | FALSE | 5 | 7 | 5 | 1558 | 1784 | 620 |
| -2,19441E | FALSE | High | IsMasterP | AAEL0065 | 2:133173! | 0 | 8 | FALSE | 2 | 11 | 2 | 304 | 337 | 577 |
| -6,24419E | FALSE | High | IsMasterP | AAEL0039 | 3:261632( | 0 | 12 | FALSE | 4 | 12 | 4 | 365 | 388 | 455 |
| 7,39462E- | FALSE | High | IsMasterP | AAEL0197 | 2:315880( | 0 | 6 | FALSE | 5 | 11 | 5 | 1184 | 1249 | 594 |
| -7,63431E | FALSE | High | IsMasterP | AAEL0087 | 3:280414 | 0 | 13 | FALSE | 4 | 10 | 4 | 300 | 338 | 966 |
| -5,9652E+ | FALSE | High | IsMasterP | AAEL0077 | 2:100898( | 0 | 22 | FALSE | 4 | 9 | 4 | 275 | 310 | 759 |
| 5,65547E- | FALSE | High | IsMasterP | AAEL0067 | 2:851589 | 0 | 17 | FALSE | 3 | 8 | 3 | 377 | 424 | 853 |
| -5,28967E | FALSE | High | IsMasterP | AAEL0013 | 1:349331 | 0 | 18 | FALSE | 2 | 9 | 2 | 159 | 166 | 703 |
| 4,60673E- | FALSE | High | IsMasterP | AAEL0236 | 2:863179 | 0 | 26 | FALSE | 2 | 9 | 2 | 149 | 148 | 928 |
| 7,14236E- | FALSE | High | IsMasterP | AAEL0138 | 1:158244 | 0 | 33 | FALSE | 6 | 9 | 6 | 360 | 393 | 695 |
| -4,39791E | FALSE | High | IsMasterP | AAEL0174 | 3:152950 | 0 | 7 | FALSE | 8 | 12 | 8 | 1457 | 1650 | 599 |
| 8,37389E- | FALSE | High | IsMasterP | AAEL0060 | 1:184018 | 0 | 6 | FALSE | 5 | 10 | 5 | 1351 | 1506 | 540 |
| 8,354E+18 | FALSE | High | IsMasterP | AAEL0000 | 1:243039 | 0 | 15 | FALSE | 2 | 7 | 2 | 463 | 528 | 632 |
| -5,6196E+ | FALSE | High | IsMasterP | AAEL0032 | 1:290908 | 0 | 19 | FALSE | 5 | 10 | 5 | 387 | 424 | 788 |
| 7,91679E- | FALSE | High | IsMasterP | AAEL0009 | 2:301817 | 0 | 7 | FALSE | 3 | 11 | 3 | 693 | 776 | 917 |
| 3,44781E- | FALSE | High | IsMasterP | AAEL0025 | 1:901236 | 0 | 18 | FALSE | 4 | 9 | 4 | 349 | 392 | 634 |
| 1,79275E- | FALSE | High | IsMasterP | AAEL0113 | 1:164725 | 0 | 16 | FALSE | 7 | 11 | 7 | 614 | 685 | 857 |
| 1,11043E- | FALSE | High | IsMasterP | AAEL0104 | 3:675966 | 0 | 23 | FALSE | 6 | 9 | 6 | 518 | 585 | 587 |
| -6,10078E | FALSE | High | IsMasterP | AAEL0031 | 1:192086 | 0 | 9 | FALSE | 5 | 12 | 5 | 705 | 799 | 753 |
| -8,30111E | FALSE | High | IsMasterP | AAEL0001 | 3:213029 | 0 | 59 | FALSE | 4 | 12 | 4 | 56 | 66 | 913 |
| -5,17526E | FALSE | High | IsMasterP | AAEL0248 | 2:761719 | 0 | 33 | FALSE | 4 | 10 | 4 | 220 | 231 | 439 |
| -8,8068E+ | FALSE | High | IsMasterP | AAEL0290 | 3:235197 | 0 | 16 | FALSE | 4 | 11 | 4 | 299 | 327 | 744 |
| 9,13923E- | FALSE | High | IsMasterP | AAEL0027 | 2:138307 | 0 | 17 | FALSE | 5 | 11 | 5 | 338 | 384 | 526 |
| -3,312E+1 | FALSE | High | IsMasterP | AAEL0130 | 2:335874 | 0 | 51 | FALSE | 4 | 11 | 4 | 128 | 137 | 615 |
| 2,54734E- | FALSE | High | IsMasterP | AAEL0051 | 2:406344 | 0 | 9 | FALSE | 5 | 8 | 5 | 1019 | 1137 | 505 |
| 1,77369E- | FALSE | High | IsMasterP | AAEL0217 | 3:124920 | 0 | 12 | FALSE | 7 | 10 | 7 | 1430 | 1534 | 629 |
| -2,15113E | FALSE | High | IsMasterP | AAEL0110 | 2:235255 | 0 | 50 | FALSE | 4 | 8 | 4 | 210 | 244 | 615 |
| -2,06405E | FALSE | High | IsMasterP | AAEL0101 | 2:161627 | 0 | 9 | FALSE | 3 | 10 | 3 | 437 | 494 | 831 |
| -1,30444E | FALSE | High | IsMasterP | AAEL0229 | 3:166394 | 0 | 13 | FALSE | 6 | 11 | 6 | 515 | 583 | 676 |
| 6,53396E- | FALSE | High | IsMasterP | AAEL0270 | 2:863725 | 0 | 54 | FALSE | 2 | 7 | 1 | 127 | 126 | 942 |
| -4,65185E | FALSE | High | IsMasterP | AAEL0181 | 3:396674 | 0 | 4 | FALSE | 4 | 13 | 4 | 1079 | 1211 | 545 |

| | | | | | | | | | | | | |
|---|---|---|---|---|---|---|---|---|---|---|---|---|
| -5,79718E | FALSE | High | IsMasterP AAEL0148 3:290867! | 0 | 12 | FALSE | 4 | 9 | 4 | 478 | 536 | 554 |
| 3,59572E- | FALSE | High | IsMasterP AAEL0103 2:182779 | 0 | 13 | FALSE | 4 | 9 | 4 | 521 | 594 | 834 |
| 7,05393E- | FALSE | High | IsMasterP AAEL0077 2:470825( | 0 | 15 | FALSE | 5 | 11 | 5 | 481 | 528 | 554 |
| 7,48922E- | FALSE | High | IsMasterP AAEL0102 2:402977! | 0 | 22 | FALSE | 3 | 9 | 3 | 206 | 224 | 695 |
| -5,64544E | FALSE | High | IsMasterP AAEL0197 1:290197 | 0 | 24 | FALSE | 4 | 7 | 4 | 435 | 456 | 761 |
| -2,42239E | FALSE | High | IsMasterP AAEL0255 1:382447! | 0 | 10 | FALSE | 2 | 12 | 2 | 204 | 242 | 1144 |
| 8,14184E- | FALSE | High | IsMasterP AAEL0067 2:837462 | 0 | 7 | FALSE | 4 | 10 | 4 | 619 | 718 | 655 |
| 3,07942E- | FALSE | High | IsMasterP AAEL0000 3:104849 | 0 | 37 | FALSE | 5 | 10 | 5 | 218 | 248 | 578 |
| -5,61813E | FALSE | High | IsMasterP AAEL0274 2:111068 | 0 | 18 | FALSE | 5 | 9 | 5 | 464 | 523 | 482 |
| 3,69243E- | FALSE | High | IsMasterP AAEL0174 1:130795 | 0 | 8 | FALSE | 1 | 8 | 1 | 262 | 268 | 415 |
| 5,9971E+1 | FALSE | High | IsMasterP AAEL0174 2:330255! | 0 | 32 | FALSE | 1 | 6 | 1 | 97 | 100 | 869 |
| -2,37885E | FALSE | High | IsMasterP AAEL0138 2:427708 | 0 | 28 | FALSE | 4 | 7 | 4 | 269 | 304 | 680 |
| 1,77282E- | FALSE | High | IsMasterP AAEL0037 1:441237 | 0 | 30 | FALSE | 4 | 8 | 4 | 198 | 221 | 668 |
| -8,61917E | FALSE | High | IsMasterP AAEL0026 2:434148 | 0 | 14 | FALSE | 4 | 10 | 4 | 536 | 593 | 646 |
| -5,66264E | FALSE | High | IsMasterP AAEL0081 3:107416 | 0 | 15 | FALSE | 8 | 11 | 8 | 848 | 965 | 825 |
| 3,17215E- | FALSE | High | IsMasterP AAEL0077 2:470247 | 0 | 27 | FALSE | 6 | 8 | 6 | 591 | 649 | 639 |
| 3,57521E- | FALSE | High | IsMasterP AAEL0053 2:257507 | 0 | 26 | FALSE | 4 | 8 | 4 | 331 | 360 | 799 |
| 1,75495E- | FALSE | High | IsMasterP AAEL0204 3:362431 | 0 | 13 | FALSE | 3 | 11 | 3 | 303 | 345 | 529 |
| 8,0517E+1 | FALSE | High | IsMasterP AAEL0078 2:473944 | 0 | 12 | FALSE | 4 | 11 | 4 | 506 | 580 | 540 |
| -6,50511E | FALSE | High | IsMasterP AAEL0139 2:305831( | 0 | 12 | FALSE | 5 | 9 | 2 | 616 | 686 | 853 |
| 1,62272E- | FALSE | High | IsMasterP AAEL0015 2:228209 | 0 | 24 | FALSE | 3 | 6 | 3 | 262 | 297 | 840 |
| -6,94367E | FALSE | High | IsMasterP AAEL0106 2:418134 | 0 | 17 | FALSE | 5 | 8 | 5 | 325 | 362 | 654 |
| -9,97692E | FALSE | High | IsMasterP AAEL0095 3:119590 | 0 | 29 | FALSE | 3 | 8 | 3 | 231 | 264 | 544 |
| -3,30547E | FALSE | High | IsMasterP AAEL0104 3:313577! | 0 | 28 | FALSE | 4 | 9 | 4 | 149 | 174 | 582 |
| -8,30118E | FALSE | High | IsMasterP AAEL0259 1:293582 | 0 | 32 | FALSE | 5 | 10 | 5 | 231 | 271 | 775 |
| 5,58701E- | FALSE | High | IsMasterP AAEL0096 1:222742( | 0 | 7 | FALSE | 6 | 11 | 6 | 1097 | 1259 | 628 |
| -5,76092E | FALSE | High | IsMasterP AAEL0010 1:210106 | 0 | 5 | FALSE | 3 | 9 | 3 | 869 | 942 | 876 |
| 6,56559E- | FALSE | High | IsMasterP AAEL0232 1:987934( | 0 | 12 | FALSE | 3 | 10 | 3 | 392 | 439 | 676 |
| 8,11699E- | FALSE | High | IsMasterP AAEL0111 2:103005( | 0 | 30 | FALSE | 5 | 9 | 5 | 341 | 382 | 501 |
| 9,90687E- | FALSE | High | IsMasterP AAEL0046 1:306215 | 0 | 23 | FALSE | 4 | 8 | 4 | 345 | 380 | 536 |
| 5,62806E- | FALSE | High | IsMasterP AAEL0096 2:169722 | 0 | 7 | FALSE | 3 | 12 | 3 | 558 | 638 | 954 |

| | | | | | | | | | | | | | | |
|---|---|---|---|---|---|---|---|---|---|---|---|---|---|---|
| 5,29393E | FALSE | High | IsMasterP | AAEL0270 | 3:151418 | 0 | 1 | FALSE | 1 | 8 | 1 | 1224 | 1298 | 446 |
| -6,81943E | FALSE | High | IsMasterP | AAEL0230 | 1:288323 | 0 | 7 | FALSE | 4 | 10 | 4 | 882 | 1013 | 784 |
| 1,24203E | FALSE | High | IsMasterP | AAEL0075 | 2:248920 | 0 | 14 | FALSE | 3 | 9 | 3 | 335 | 384 | 615 |
| -7,11323E | FALSE | High | IsMasterP | AAEL0061 | 1:224661 | 0 | 24 | FALSE | 4 | 9 | 4 | 263 | 287 | 595 |
| -3,05377E | FALSE | High | IsMasterP | AAEL0086 | 2:215268 | 0 | 16 | FALSE | 5 | 8 | 5 | 613 | 670 | 541 |
| -6,36979E | FALSE | High | IsMasterP | AAEL0044 | 3:272306 | 0 | 24 | FALSE | 7 | 9 | 7 | 517 | 581 | 664 |
| -5,86066E | FALSE | High | IsMasterP | AAEL0116 | 3:281556 | 0 | 21 | FALSE | 8 | 11 | 8 | 442 | 510 | 535 |
| -3,26017E | FALSE | High | IsMasterP | AAEL0096 | 1:928742 | 0 | 9 | FALSE | 2 | 12 | 2 | 132 | 144 | 1024 |
| 2,73224E | FALSE | High | IsMasterP | AAEL0056 | 1:473265 | 0 | 29 | FALSE | 4 | 10 | 4 | 123 | 143 | 1093 |
| -8,12935E | FALSE | High | IsMasterP | AAEL0131 | 2:179154 | 0 | 24 | FALSE | 2 | 9 | 2 | 203 | 225 | 660 |
| 5,21247E | FALSE | High | IsMasterP | AAEL0049 | 3:323266 | 0 | 6 | FALSE | 1 | 7 | 1 | 273 | 306 | 549 |
| 2,82006E | FALSE | High | IsMasterP | AAEL0052 | 2:229031 | 0 | 27 | FALSE | 5 | 7 | 5 | 335 | 365 | 498 |
| 9,0351E+ | FALSE | High | IsMasterP | AAEL0114 | 2:225313 | 0 | 22 | FALSE | 3 | 9 | 3 | 147 | 153 | 1042 |
| 1,56042E | FALSE | High | IsMasterP | AAEL0039 | 1:250574 | 0 | 56 | FALSE | 6 | 8 | 6 | 133 | 153 | 620 |
| 6,411E+1 | FALSE | High | IsMasterP | AAEL0123 | 3:316628 | 0 | 4 | FALSE | 5 | 9 | 5 | 1865 | 2079 | 686 |
| 3,72964E | FALSE | High | IsMasterP | AAEL0017 | 2:166819 | 0 | 37 | FALSE | 2 | 7 | 2 | 120 | 136 | 514 |
| 6,84345E | FALSE | High | IsMasterP | AAEL0081 | 3:973654 | 0 | 21 | FALSE | 5 | 10 | 5 | 327 | 370 | 689 |
| -4,58651E | FALSE | High | IsMasterP | AAEL0090 | 1:293569 | 0 | 30 | FALSE | 4 | 8 | 4 | 229 | 268 | 799 |
| 8,23904E | FALSE | High | IsMasterP | AAEL0092 | 2:112608 | 0 | 20 | FALSE | 4 | 8 | 4 | 329 | 380 | 885 |
| 8,4457E+ | FALSE | High | IsMasterP | AAEL0170 | 2:182931 | 0 | 12 | FALSE | 4 | 9 | 4 | 539 | 605 | 564 |
| 5,93386E | FALSE | High | IsMasterP | AAEL0150 | 2:210390 | 0 | 25 | FALSE | 5 | 9 | 5 | 307 | 349 | 607 |
| 3,35428E | FALSE | High | IsMasterP | AAEL0040 | 3:166372 | 0 | 13 | FALSE | 4 | 7 | 3 | 625 | 693 | 554 |
| 1,32029E | FALSE | High | IsMasterP | AAEL0072 | 2:312437 | 0 | 26 | FALSE | 5 | 8 | 5 | 304 | 331 | 596 |
| -6,32536E | FALSE | High | IsMasterP | AAEL0205 | 1:324214 | 0 | 7 | FALSE | 4 | 10 | 4 | 901 | 1013 | 615 |
| 5,96156E | FALSE | High | IsMasterP | AAEL0225 | 3:120181 | 0 | 10 | FALSE | 5 | 11 | 5 | 469 | 545 | 828 |
| 5,32889E | FALSE | High | IsMasterP | AAEL0074 | 1:216400 | 0 | 19 | FALSE | 8 | 10 | 8 | 771 | 867 | 667 |
| -4,52498E | FALSE | High | IsMasterP | AAEL0035 | 2:256864 | 0 | 46 | FALSE | 4 | 8 | 4 | 146 | 169 | 725 |
| 8,93369E | FALSE | High | IsMasterP | AAEL0107 | 1:178499 | 0 | 24 | FALSE | 4 | 8 | 4 | 377 | 445 | 736 |
| -3,3026E+ | FALSE | High | IsMasterP | AAEL0121 | 2:188105 | 0 | 18 | FALSE | 5 | 10 | 5 | 347 | 395 | 1005 |
| -6,33342E | FALSE | High | IsMasterP | AAEL0021 | 1:253570 | 0 | 26 | FALSE | 4 | 8 | 4 | 241 | 271 | 521 |
| -6,01725E | FALSE | High | IsMasterP | AAEL0019 | 2:389851 | 0 | 26 | FALSE | 6 | 8 | 6 | 326 | 377 | 604 |

| | | | | | | | | | | | | | | |
|---|---|---|---|---|---|---|---|---|---|---|---|---|---|---|
| -5,07943E | FALSE | High | IsMasterP | AAEL0111 | 2:379703 | 0 | 20 | FALSE | 2 | 8 | 2 | 182 | 205 | 661 |
| 4,44386E | FALSE | High | IsMasterP | AAEL0009 | 3:335515 | 0 | 8 | FALSE | 5 | 10 | 5 | 777 | 854 | 793 |
| -3,89328E | FALSE | High | IsMasterP | AAEL0113 | 3:322160 | 0 | 61 | FALSE | 3 | 9 | 3 | 70 | 79 | 597 |
| -6,08828E | FALSE | High | IsMasterP | AAEL0211 | 1:146717 | 0 | 9 | FALSE | 4 | 8 | 4 | 714 | 789 | 690 |
| -3,17291E | FALSE | High | IsMasterP | AAEL0180 | 2:285143 | 0 | 14 | FALSE | 3 | 7 | 3 | 416 | 468 | 805 |
| 6,16236E | FALSE | High | IsMasterP | AAEL0022 | 2:298936 | 0 | 11 | FALSE | 2 | 8 | 2 | 305 | 340 | 837 |
| 6,79963E | FALSE | High | IsMasterP | AAEL0068 | 3:369360 | 0 | 38 | FALSE | 4 | 8 | 4 | 184 | 202 | 851 |
| -1,75625E | FALSE | High | IsMasterP | AAEL0067 | 2:837116 | 0 | 18 | FALSE | 6 | 10 | 6 | 486 | 539 | 835 |
| -5,38445E | FALSE | High | IsMasterP | AAEL0083 | 1:935056 | 0 | 42 | FALSE | 5 | 9 | 5 | 217 | 243 | 824 |
| -8,50758E | FALSE | High | IsMasterP | AAEL0048 | 2:115450 | 0 | 32 | FALSE | 4 | 9 | 4 | 197 | 216 | 812 |
| 7,49538E | FALSE | High | IsMasterP | AAEL0109 | 3:190430 | 0 | 14 | FALSE | 7 | 11 | 7 | 602 | 686 | 702 |
| -2,421E+1 | FALSE | High | IsMasterP | AAEL0273 | 3:261062 | 0 | 6 | FALSE | 3 | 10 | 3 | 699 | 769 | 520 |
| -4,9322E+ | FALSE | High | IsMasterP | AAEL0066 | 3:352773 | 0 | 17 | FALSE | 4 | 10 | 4 | 307 | 339 | 517 |
| 4,78503E | FALSE | High | IsMasterP | AAEL0043 | 2:387354 | 0 | 16 | FALSE | 5 | 10 | 5 | 495 | 538 | 569 |
| 8,97678E | FALSE | High | IsMasterP | AAEL0002 | 1:659249 | 0 | 24 | FALSE | 6 | 11 | 6 | 288 | 324 | 625 |
| 1,58434E | FALSE | High | IsMasterP | AAEL0025 | 3:256747 | 0 | 19 | FALSE | 5 | 8 | 5 | 428 | 471 | 547 |
| 8,47451E | FALSE | High | IsMasterP | AAEL0019 | 2:194426 | 0 | 29 | FALSE | 5 | 11 | 5 | 182 | 208 | 957 |
| 4,46402E | FALSE | High | IsMasterP | AAEL0006 | 2:285067 | 0 | 7 | FALSE | 6 | 9 | 6 | 1131 | 1292 | 982 |
| -5,08469E | FALSE | High | IsMasterP | AAEL0290 | 3:111468 | 0 | 19 | FALSE | 5 | 10 | 5 | 359 | 402 | 493 |
| 3,7426E+ | FALSE | High | IsMasterP | AAEL0141 | 2:115794 | 0 | 66 | FALSE | 3 | 8 | 3 | 92 | 102 | 454 |
| -5,63251E | FALSE | High | IsMasterP | AAEL0140 | 3:188019 | 0 | 23 | FALSE | 4 | 8 | 4 | 343 | 374 | 679 |
| -4,06131E | FALSE | High | IsMasterP | AAEL0074 | 1:463898 | 0 | 23 | FALSE | 5 | 9 | 5 | 327 | 371 | 686 |
| -8,62269E | FALSE | High | IsMasterP | AAEL0073 | 2:234467 | 0 | 24 | FALSE | 3 | 10 | 3 | 139 | 161 | 747 |
| -3,1928E+ | FALSE | High | IsMasterP | AAEL0032 | 2:366397 | 0 | 30 | FALSE | 3 | 9 | 3 | 271 | 292 | 809 |
| 3,86093E | FALSE | High | IsMasterP | AAEL0023 | 2:175283 | 0 | 20 | FALSE | 4 | 9 | 4 | 210 | 244 | 510 |
| 4,53939E | FALSE | High | IsMasterP | AAEL0056 | 2:381787 | 0 | 15 | FALSE | 3 | 9 | 3 | 389 | 439 | 644 |
| 1,67039E | FALSE | High | IsMasterP | AAEL0057 | 2:294066 | 0 | 11 | FALSE | 3 | 8 | 3 | 462 | 492 | 966 |
| 5,95447E | FALSE | High | IsMasterP | AAEL0115 | 3:106899 | 0 | 3 | FALSE | 6 | 9 | 6 | 2717 | 3007 | 491 |
| 7,66799E | FALSE | High | IsMasterP | AAEL0183 | 2:293598 | 0 | 6 | FALSE | 5 | 10 | 5 | 1197 | 1351 | 607 |
| -8,25296E | FALSE | High | IsMasterP | AAEL0029 | 1:861959 | 0 | 13 | FALSE | 5 | 10 | 5 | 596 | 682 | 553 |
| 2,77942E | FALSE | High | IsMasterP | AAEL0268 | 3:216247 | 0 | 6 | FALSE | 3 | 9 | 3 | 794 | 860 | 897 |

| | | | | | | | | | | | | | | |
|---|---|---|---|---|---|---|---|---|---|---|---|---|---|---|
| 2,87642E- | FALSE | High | IsMasterP | AAEL0035 | 3:2642448 | 0 | 11 | FALSE | 4 | 8 | 4 | 607 | 693 | 521 |
| 7,87359E- | FALSE | High | IsMasterP | AAEL0117 | 3:2887552 | 0 | 10 | FALSE | 5 | 10 | 5 | 666 | 770 | 493 |
| -8,31568E | FALSE | High | IsMasterP | AAEL0043 | 3:3657233 | 0 | 13 | FALSE | 6 | 8 | 6 | 1048 | 1191 | 647 |
| -4,79437E | FALSE | High | IsMasterP | AAEL0267 | 2:2616581 | 0 | 16 | FALSE | 5 | 8 | 5 | 750 | 813 | 637 |
| 1,84341E- | FALSE | High | IsMasterP | AAEL0007 | 2:4587253 | 0 | 20 | FALSE | 4 | 8 | 4 | 279 | 316 | 955 |
| 7,5607E+1 | FALSE | High | IsMasterP | AAEL0126 | 2:1220643 | 0 | 8 | FALSE | 3 | 8 | 3 | 736 | 844 | 724 |
| -1,6484E+ | FALSE | High | IsMasterP | AAEL0008 | 3:3627015 | 0 | 6 | FALSE | 2 | 9 | 2 | 508 | 596 | 679 |
| -2,61175E | FALSE | High | IsMasterP | AAEL0070 | NIGP0100 | 0 | 10 | FALSE | 4 | 9 | 4 | 366 | 413 | 667 |
| 4,88177E- | FALSE | High | IsMasterP | AAEL0014 | 1:1117293 | 0 | 13 | FALSE | 3 | 9 | 3 | 305 | 355 | 596 |
| -9,11781E | FALSE | High | IsMasterP | AAEL0108 | 1:5468388 | 0 | 20 | FALSE | 3 | 9 | 3 | 207 | 246 | 961 |
| 1,5133E+1 | FALSE | High | IsMasterP | AAEL0013 | 2:1181081 | 0 | 14 | FALSE | 3 | 7 | 3 | 230 | 263 | 885 |
| 7,83766E- | FALSE | High | IsMasterP | AAEL0181 | 2:2148967 | 0 | 18 | FALSE | 4 | 10 | 3 | 289 | 329 | 799 |
| 9,23056E- | FALSE | High | IsMasterP | AAEL0091 | 1:1126399 | 0 | 14 | FALSE | 4 | 8 | 4 | 495 | 550 | 619 |
| -8,3685E+ | FALSE | High | IsMasterP | AAEL0002 | 2:2277807 | 0 | 4 | FALSE | 1 | 9 | 1 | 312 | 353 | 863 |
| 4,4383E+1 | FALSE | High | IsMasterP | AAEL0047 | 2:2887160 | 0 | 30 | FALSE | 2 | 7 | 2 | 161 | 179 | 618 |
| -1,30206E | FALSE | High | IsMasterP | AAEL0258 | 3:2642736 | 0 | 21 | FALSE | 3 | 11 | 3 | 159 | 168 | 543 |
| 1,50686E- | FALSE | High | IsMasterP | AAEL0070 | 3:2986076 | 0 | 16 | FALSE | 3 | 7 | 3 | 336 | 369 | 991 |
| 6,14374E- | FALSE | High | IsMasterP | AAEL0110 | 2:2321925 | 0 | 6 | FALSE | 5 | 9 | 5 | 1354 | 1545 | 536 |
| 7,76018E- | FALSE | High | IsMasterP | AAEL0005 | 2:1476223 | 0 | 4 | FALSE | 1 | 8 | 1 | 424 | 476 | 540 |
| 8,95985E- | FALSE | High | IsMasterP | AAEL0133 | 2:1356557 | 0 | 46 | FALSE | 5 | 9 | 5 | 192 | 219 | 640 |
| 5,05471E- | FALSE | High | IsMasterP | AAEL0081 | 3:1472969 | 0 | 20 | FALSE | 2 | 5 | 2 | 321 | 318 | 914 |
| -5,63659E | FALSE | High | IsMasterP | AAEL0019 | 2:4259773 | 0 | 25 | FALSE | 5 | 11 | 2 | 126 | 144 | 898 |
| 3,9912E+1 | FALSE | High | IsMasterP | AAEL0025 | 1:2265531 | 0 | 5 | FALSE | 6 | 9 | 6 | 1212 | 1364 | 674 |
| -8,00093E | FALSE | High | IsMasterP | AAEL0061 | 3:6874343 | 0 | 31 | FALSE | 3 | 10 | 3 | 159 | 184 | 859 |
| 7,20118E- | FALSE | High | IsMasterP | AAEL0086 | 3:1730274 | 0 | 14 | FALSE | 3 | 9 | 3 | 348 | 389 | 684 |
| 1,27798E- | FALSE | High | IsMasterP | AAEL0180 | 1:8848734 | 0 | 6 | FALSE | 5 | 7 | 5 | 1382 | 1604 | 657 |
| -8,30958E | FALSE | High | IsMasterP | AAEL0105 | 2:1231305 | 0 | 23 | FALSE | 2 | 9 | 2 | 125 | 135 | 601 |
| 3,49072E- | FALSE | High | IsMasterP | AAEL0198 | 2:1510177 | 0 | 21 | FALSE | 4 | 8 | 4 | 306 | 350 | 503 |
| 7,61051E- | FALSE | High | IsMasterP | AAEL0082 | 3:6487872 | 0 | 7 | FALSE | 6 | 7 | 6 | 1508 | 1722 | 771 |
| -5,97215E | FALSE | High | IsMasterP | AAEL0196 | 2:4318974 | 0 | 23 | FALSE | 4 | 8 | 4 | 379 | 402 | 708 |
| -6,01675E | FALSE | High | IsMasterP | AAEL0148 | 2:1104758 | 0 | 43 | FALSE | 4 | 8 | 4 | 128 | 143 | 822 |

| | | | | | | | | | | | | | | |
|---|---|---|---|---|---|---|---|---|---|---|---|---|---|---|
| -5,32781E | FALSE | High | IsMasterP | AAEL0117 | 3:288706 | 0 | 7 | FALSE | 4 | 9 | 4 | 848 | 938 | 963 |
| 7,1259E+ | FALSE | High | IsMasterP | AAEL0076 | 2:470366 | 0 | 6 | FALSE | 4 | 7 | 1 | 860 | 985 | 941 |
| 7,47417E | FALSE | High | IsMasterP | AAEL0003 | 2:367796 | 0 | 7 | FALSE | 3 | 9 | 3 | 694 | 779 | 689 |
| 1,55557E | FALSE | High | IsMasterP | AAEL0074 | 1:459360 | 0 | 2 | FALSE | 5 | 8 | 5 | 3644 | 4029 | 928 |
| 3,76567E | FALSE | High | IsMasterP | AAEL0076 | 3:136766 | 0 | 16 | FALSE | 4 | 7 | 4 | 611 | 665 | 642 |
| -4,2162E+ | FALSE | High | IsMasterP | AAEL0101 | 2:264973 | 0 | 18 | FALSE | 4 | 9 | 3 | 319 | 361 | 725 |
| 2,68385E | FALSE | High | IsMasterP | AAEL0142 | 2:399811 | 0 | 22 | FALSE | 4 | 7 | 4 | 277 | 318 | 888 |
| 4,38221E | FALSE | High | IsMasterP | AAEL0212 | 3:263510 | 0 | 7 | FALSE | 4 | 7 | 4 | 1227 | 1357 | 702 |
| 6,82663E | FALSE | High | IsMasterP | AAEL0012 | 2:153030 | 0 | 13 | FALSE | 4 | 8 | 4 | 550 | 596 | 742 |
| -5,43098E | FALSE | High | IsMasterP | AAEL0009 | 3:334479 | 0 | 4 | FALSE | 3 | 9 | 3 | 552 | 608 | 756 |
| 7,90781E | FALSE | High | IsMasterP | AAEL0131 | 1:142885 | 0 | 13 | FALSE | 3 | 7 | 3 | 372 | 425 | 980 |
| 6,29957E | FALSE | High | IsMasterP | AAEL0003 | 2:124628 | 0 | 22 | FALSE | 4 | 8 | 4 | 274 | 300 | 506 |
| 8,16453E | FALSE | High | IsMasterP | AAEL0026 | 3:257164 | 0 | 34 | FALSE | 5 | 10 | 5 | 131 | 141 | 841 |
| -2,05123E | FALSE | High | IsMasterP | AAEL0097 | 1:415359 | 0 | 13 | FALSE | 5 | 10 | 5 | 362 | 424 | 525 |
| 5,44597E | FALSE | High | IsMasterP | AAEL0084 | 1:173611 | 0 | 20 | FALSE | 4 | 8 | 4 | 293 | 311 | 774 |
| 8,41759E | FALSE | High | IsMasterP | AAEL0110 | 2:235607 | 0 | 11 | FALSE | 2 | 6 | 2 | 264 | 299 | 662 |
| -5,72865E | FALSE | High | IsMasterP | AAEL0074 | 2:291355 | 0 | 6 | FALSE | 1 | 9 | 1 | 184 | 209 | 909 |
| -8,36013E | FALSE | High | IsMasterP | AAEL0032 | 2:367519 | 0 | 51 | FALSE | 3 | 9 | 3 | 109 | 122 | 991 |
| -7,17043E | FALSE | High | IsMasterP | AAEL0033 | 3:813529 | 0 | 13 | FALSE | 4 | 8 | 4 | 375 | 434 | 588 |
| -4,9089E+ | FALSE | High | IsMasterP | AAEL0000 | 1:242820 | 0 | 36 | FALSE | 5 | 7 | 5 | 275 | 300 | 512 |
| -7,30994E | FALSE | High | IsMasterP | AAEL0098 | 3:189254 | 0 | 18 | FALSE | 2 | 8 | 2 | 157 | 180 | 461 |
| 6,7089E+ | FALSE | High | IsMasterP | AAEL0088 | 3:132221 | 0 | 25 | FALSE | 3 | 6 | 3 | 208 | 227 | 554 |
| -2,14965E | FALSE | High | IsMasterP | AAEL0093 | 3:337998 | 0 | 17 | FALSE | 2 | 7 | 2 | 303 | 339 | 766 |
| 6,17823E | FALSE | High | IsMasterP | AAEL0035 | 3:180015 | 0 | 15 | FALSE | 5 | 7 | 5 | 505 | 573 | 925 |
| 6,94829E | FALSE | High | IsMasterP | AAEL0198 | 2:333920 | 0 | 13 | FALSE | 6 | 8 | 6 | 783 | 870 | 762 |
| 1,01984E | FALSE | High | IsMasterP | AAEL0056 | 1:473162 | 0 | 17 | FALSE | 4 | 7 | 4 | 421 | 471 | 508 |
| 3,6802E+ | FALSE | High | IsMasterP | AAEL0279 | NIGP0100 | 0 | 55 | FALSE | 3 | 7 | 3 | 95 | 103 | 475 |
| 7,01828E | FALSE | High | IsMasterP | AAEL0196 | 2:174359 | 0 | 12 | FALSE | 4 | 8 | 4 | 577 | 652 | 674 |
| 1,67807E | FALSE | High | IsMasterP | AAEL0217 | 1:303261 | 0 | 16 | FALSE | 6 | 8 | 6 | 484 | 573 | 550 |
| 5,14627E | FALSE | High | IsMasterP | AAEL0048 | 1:288247 | 0 | 22 | FALSE | 5 | 8 | 5 | 448 | 500 | 832 |
| -3,92972E | FALSE | High | IsMasterP | AAEL0034 | 3:348395 | 0 | 4 | FALSE | 3 | 10 | 3 | 776 | 869 | 684 |

| | | | | | | | | | | | | | | |
|---|---|---|---|---|---|---|---|---|---|---|---|---|---|---|
| 5,12408E- | FALSE | High | IsMasterP | AAEL0076 | 1:491123 | 0 | 19 | FALSE | 4 | 6 | 4 | 385 | 441 | 844 |
| 5,42479E- | FALSE | High | IsMasterP | AAEL0235 | NIGP0100 | 0 | 30 | FALSE | 3 | 7 | 3 | 289 | 319 | 464 |
| 8,44275E- | FALSE | High | IsMasterP | AAEL0085 | 3:293463 | 0 | 33 | FALSE | 4 | 8 | 4 | 191 | 214 | 714 |
| 8,17118E- | FALSE | High | IsMasterP | AAEL0058 | 1:225511 | 0 | 25 | FALSE | 5 | 7 | 5 | 353 | 390 | 696 |
| 7,33695E- | FALSE | High | IsMasterP | AAEL0023 | 2:303083 | 0 | 20 | FALSE | 3 | 7 | 3 | 205 | 215 | 1125 |
| 3,66186E- | FALSE | High | IsMasterP | AAEL0055 | 3:43502:4 | 0 | 11 | FALSE | 3 | 7 | 3 | 395 | 456 | 469 |
| 6,55646E- | FALSE | High | IsMasterP | AAEL0104 | 1:268010 | 0 | 40 | FALSE | 4 | 6 | 4 | 201 | 228 | 597 |
| 8,30704E- | FALSE | High | IsMasterP | AAEL0013 | 3:327108 | 0 | 7 | FALSE | 3 | 6 | 3 | 799 | 885 | 658 |
| 8,22298E- | FALSE | High | IsMasterP | AAEL0144 | 3:149517 | 0 | 9 | FALSE | 3 | 10 | 3 | 410 | 471 | 520 |
| -7,62395E | FALSE | High | IsMasterP | AAEL0110 | 3:294582 | 0 | 12 | FALSE | 3 | 7 | 3 | 489 | 542 | 524 |
| -2,65197E | FALSE | High | IsMasterP | AAEL0123 | 2:102748 | 0 | 33 | FALSE | 3 | 7 | 3 | 225 | 251 | 482 |
| -6,53379E | FALSE | High | IsMasterP | AAEL0032 | 3:330386 | 0 | 13 | FALSE | 7 | 7 | 7 | 950 | 1089 | 526 |
| -5,86892E | FALSE | High | IsMasterP | AAEL0106 | 2:279155 | 0 | 18 | FALSE | 2 | 8 | 2 | 179 | 202 | 692 |
| 4,74696E- | FALSE | High | IsMasterP | AAEL0043 | 3:508402 | 0 | 56 | FALSE | 3 | 7 | 3 | 138 | 153 | 448 |
| -1,41555E | FALSE | High | IsMasterP | AAEL0146 | 3:368529 | 0 | 4 | FALSE | 2 | 10 | 1 | 1110 | 1273 | 857 |
| 3,75598E- | FALSE | High | IsMasterP | AAEL0015 | 3:261139 | 0 | 8 | FALSE | 3 | 5 | 3 | 909 | 1009 | 819 |
| 8,11375E- | FALSE | High | IsMasterP | AAEL0006 | 2:285133 | 0 | 18 | FALSE | 3 | 9 | 3 | 217 | 245 | 888 |
| -1,62367E | FALSE | High | IsMasterP | AAEL0013 | 2:447504 | 0 | 18 | FALSE | 8 | 9 | 8 | 533 | 613 | 714 |
| -2,66331E | FALSE | High | IsMasterP | AAEL0057 | 2:437038 | 0 | 38 | FALSE | 4 | 9 | 4 | 172 | 187 | 628 |
| 6,39363E- | FALSE | High | IsMasterP | AAEL0251 | 3:288880 | 0 | 11 | FALSE | 2 | 7 | 2 | 266 | 302 | 752 |
| 5,41169E- | FALSE | High | IsMasterP | AAEL0072 | 1:267314 | 0 | 33 | FALSE | 4 | 8 | 4 | 186 | 212 | 718 |
| 2,18013E- | FALSE | High | IsMasterP | AAEL0000 | 3:104987 | 0 | 17 | FALSE | 5 | 9 | 5 | 372 | 411 | 548 |
| -8,43648E | FALSE | High | IsMasterP | AAEL0089 | 1:811368 | 0 | 6 | FALSE | 2 | 7 | 2 | 559 | 638 | 512 |
| 3,77117E- | FALSE | High | IsMasterP | AAEL0037 | 1:249118 | 0 | 3 | FALSE | 2 | 8 | 2 | 983 | 1118 | 638 |
| -1,6895E+ | FALSE | High | IsMasterP | AAEL0004 | 3:183601 | 0 | 11 | FALSE | 3 | 8 | 3 | 442 | 501 | 764 |
| 1,23349E- | FALSE | High | IsMasterP | AAEL0254 | 3:138451 | 0 | 9 | FALSE | 7 | 9 | 7 | 1388 | 1545 | 825 |
| -5,75195E | FALSE | High | IsMasterP | AAEL0068 | 1:527831 | 0 | 10 | FALSE | 4 | 10 | 4 | 537 | 596 | 563 |
| 7,30133E- | FALSE | High | IsMasterP | AAEL0115 | 3:106633 | 0 | 12 | FALSE | 6 | 9 | 6 | 497 | 560 | 529 |
| -4,69417E | FALSE | High | IsMasterP | AAEL0119 | 3:337695 | 0 | 14 | FALSE | 4 | 10 | 4 | 429 | 473 | 784 |
| 5,11554E- | FALSE | High | IsMasterP | AAEL0013 | 2:205161 | 0 | 20 | FALSE | 2 | 7 | 2 | 188 | 217 | 460 |
| -1,42448E | FALSE | High | IsMasterP | AAEL0135 | 1:144055 | 0 | 4 | FALSE | 1 | 6 | 1 | 449 | 494 | 803 |

| | | | | | | | | | | | | | | |
|---|---|---|---|---|---|---|---|---|---|---|---|---|---|---|
| -4,80432E | FALSE | High | IsMasterP | AAEL0064 | 3:2166642 | 0 | 19 | FALSE | 4 | 7 | 2 | 377 | 404 | 661 |
| -4,5012E+ | FALSE | High | IsMasterP | AAEL0236 | 1:2103450 | 0 | 15 | FALSE | 5 | 7 | 5 | 666 | 774 | 539 |
| 4,90429E- | FALSE | High | IsMasterP | AAEL0267 | 3:2346891 | 0 | 9 | FALSE | 5 | 8 | 5 | 1164 | 1285 | 543 |
| -9,04671E | FALSE | High | IsMasterP | AAEL0005 | 3:3887468 | 0 | 29 | FALSE | 3 | 9 | 3 | 167 | 192 | 559 |
| 5,6699E+1 | FALSE | High | IsMasterP | AAEL0061 | 1:2247763 | 0 | 13 | FALSE | 4 | 7 | 4 | 610 | 669 | 926 |
| -4,63591E | FALSE | High | IsMasterP | AAEL0149 | 3:2042183 | 0 | 25 | FALSE | 4 | 10 | 4 | 174 | 201 | 460 |
| 2,82929E- | FALSE | High | IsMasterP | AAEL0114 | 2:2250892 | 0 | 25 | FALSE | 6 | 7 | 6 | 454 | 501 | 689 |
| 1,24698E- | FALSE | High | IsMasterP | AAEL0097 | 1:4199935 | 0 | 21 | FALSE | 5 | 8 | 5 | 464 | 500 | 706 |
| -4,48075E | FALSE | High | IsMasterP | AAEL0087 | 2:1988347 | 0 | 41 | FALSE | 4 | 8 | 4 | 147 | 161 | 991 |
| 6,47023E- | FALSE | High | IsMasterP | AAEL0037 | 1:2480996 | 0 | 11 | FALSE | 5 | 10 | 5 | 507 | 568 | 595 |
| -6,56673E | FALSE | High | IsMasterP | AAEL0195 | 2:2931811 | 0 | 6 | FALSE | 5 | 8 | 5 | 1244 | 1383 | 454 |
| -8,03032E | FALSE | High | IsMasterP | AAEL0138 | 3:2751339 | 0 | 12 | FALSE | 3 | 9 | 3 | 304 | 353 | 725 |
| 8,99901E- | FALSE | High | IsMasterP | AAEL0010 | 1:2996259 | 0 | 32 | FALSE | 4 | 8 | 4 | 309 | 351 | 714 |
| 2,36869E- | FALSE | High | IsMasterP | AAEL0023 | 1:1194936 | 0 | 22 | FALSE | 5 | 8 | 5 | 404 | 444 | 607 |
| -6,87894E | FALSE | High | IsMasterP | AAEL0122 | 2:1449633 | 0 | 32 | FALSE | 4 | 7 | 4 | 207 | 234 | 508 |
| 9,11924E- | FALSE | High | IsMasterP | AAEL0028 | 3:2350284 | 0 | 13 | FALSE | 4 | 7 | 4 | 557 | 611 | 643 |
| 2,9896E+1 | FALSE | High | IsMasterP | AAEL0217 | 2:2007041 | 0 | 14 | FALSE | 4 | 7 | 4 | 483 | 546 | 736 |
| 2,09762E- | FALSE | High | IsMasterP | AAEL0121 | 2:1436584 | 0 | 28 | FALSE | 4 | 6 | 4 | 231 | 255 | 701 |
| 9,18632E- | FALSE | High | IsMasterP | AAEL0092 | 2:2684722 | 0 | 26 | FALSE | 2 | 9 | 2 | 78 | 88 | 950 |
| -8,93249E | FALSE | High | IsMasterP | AAEL0028 | 1:1082884 | 0 | 9 | FALSE | 3 | 5 | 3 | 832 | 903 | 522 |
| 1,09794E- | FALSE | High | IsMasterP | AAEL0075 | 2:2174735 | 0 | 10 | FALSE | 5 | 8 | 5 | 901 | 1021 | 501 |
| 2,43833E- | FALSE | High | IsMasterP | AAEL0074 | 2:5772584 | 0 | 15 | FALSE | 2 | 8 | 2 | 193 | 215 | 456 |
| 8,41368E- | FALSE | High | IsMasterP | AAEL0003 | 2:1243828 | 0 | 56 | FALSE | 4 | 7 | 4 | 122 | 136 | 497 |
| 4,17619E- | FALSE | High | IsMasterP | AAEL0065 | 2:1333671 | 0 | 26 | FALSE | 4 | 9 | 4 | 160 | 183 | 408 |
| 5,69006E- | FALSE | High | IsMasterP | AAEL0133 | 2:1357362 | 0 | 19 | FALSE | 2 | 7 | 1 | 194 | 215 | 613 |
| 8,89711E- | FALSE | High | IsMasterP | AAEL0108 | 1:2178904 | 0 | 11 | FALSE | 4 | 7 | 4 | 492 | 561 | 496 |
| 1,26918E- | FALSE | High | IsMasterP | AAEL0078 | 2:2726016 | 0 | 7 | FALSE | 5 | 10 | 5 | 827 | 940 | 647 |
| -4,07147E | FALSE | High | IsMasterP | AAEL0199 | 1:2324437 | 0 | 14 | FALSE | 5 | 8 | 5 | 476 | 543 | 794 |
| 3,83989E- | FALSE | High | IsMasterP | AAEL0221 | 2:1342303 | 0 | 13 | FALSE | 4 | 7 | 4 | 620 | 692 | 574 |
| -1,05578E | FALSE | High | IsMasterP | AAEL0290 | 1:8351650 | 0 | 21 | FALSE | 5 | 10 | 3 | 294 | 335 | 698 |
| -7,21609E | FALSE | High | IsMasterP | AAEL0115 | 3:3128266 | 0 | 14 | FALSE | 3 | 7 | 3 | 389 | 434 | 562 |

| | | | | | | | | | | | | | | |
|---|---|---|---|---|---|---|---|---|---|---|---|---|---|---|
| -1,5873E+ | FALSE | High | IsMasterP | AAEL0276 | 1:7538850 | 0 | 8 | FALSE | 3 | 7 | 3 | 366 | 405 | 772 |
| -6,11886E | FALSE | High | IsMasterP | AAEL0021 | 3:1769435 | 0 | 24 | FALSE | 4 | 8 | 4 | 203 | 232 | 780 |
| 5,06723E- | FALSE | High | IsMasterP | AAEL0094 | 2:2465985 | 0 | 5 | FALSE | 4 | 8 | 4 | 998 | 1067 | 654 |
| -1,43729E | FALSE | High | IsMasterP | AAEL0045 | 1:1445341 | 0 | 13 | FALSE | 4 | 9 | 4 | 295 | 338 | 753 |
| -4,53642E | FALSE | High | IsMasterP | AAEL0102 | 2:3412413 | 0 | 12 | FALSE | 3 | 7 | 3 | 386 | 436 | 647 |
| 5,79608E- | FALSE | High | IsMasterP | AAEL0153 | 2:1743363 | 0 | 12 | FALSE | 3 | 7 | 3 | 322 | 357 | 647 |
| -5,13939E | FALSE | High | IsMasterP | AAEL0019 | 3:2270385 | 0 | 4 | FALSE | 5 | 9 | 5 | 1794 | 2068 | 563 |
| 3,26137E- | FALSE | High | IsMasterP | AAEL0096 | 3:2216551 | 0 | 46 | FALSE | 2 | 5 | 2 | 101 | 117 | 488 |
| -3,45252E | FALSE | High | IsMasterP | AAEL0149 | 2:1285282 | 0 | 9 | FALSE | 2 | 5 | 2 | 542 | 581 | 910 |
| 8,65553E- | FALSE | High | IsMasterP | AAEL0173 | 3:1831273 | 0 | 39 | FALSE | 2 | 6 | 2 | 88 | 92 | 703 |
| 6,21103E- | FALSE | High | IsMasterP | AAEL0055 | 3:9059453 | 0 | 2 | FALSE | 1 | 8 | 1 | 880 | 974 | 898 |
| 3,00123E- | FALSE | High | IsMasterP | AAEL0135 | 2:3759393 | 0 | 43 | FALSE | 2 | 8 | 2 | 101 | 112 | 459 |
| -1,04354E | FALSE | High | IsMasterP | AAEL0175 | 1:7377699 | 0 | 6 | FALSE | 4 | 7 | 4 | 905 | 1010 | 862 |
| 1,20485E- | FALSE | High | IsMasterP | AAEL0085 | 2:1572191 | 0 | 22 | FALSE | 4 | 7 | 4 | 236 | 255 | 919 |
| 1,2372E+1 | FALSE | High | IsMasterP | AAEL0067 | 2:8477947 | 0 | 12 | FALSE | 5 | 7 | 5 | 666 | 747 | 658 |
| 9,07969E- | FALSE | High | IsMasterP | AAEL0112 | 2:2260009 | 0 | 22 | FALSE | 4 | 7 | 4 | 315 | 339 | 957 |
| 5,32698E- | FALSE | High | IsMasterP | AAEL0136 | 1:6464666 | 0 | 18 | FALSE | 3 | 6 | 3 | 285 | 292 | 686 |
| -4,89431E | FALSE | High | IsMasterP | AAEL0034 | 2:2163886 | 0 | 14 | FALSE | 4 | 7 | 4 | 432 | 500 | 648 |
| 2,77676E- | FALSE | High | IsMasterP | AAEL0021 | 3:1784418 | 0 | 10 | FALSE | 4 | 7 | 4 | 587 | 648 | 897 |
| 5,42533E- | FALSE | High | IsMasterP | AAEL0080 | 1:6173075 | 0 | 17 | FALSE | 3 | 6 | 3 | 311 | 315 | 775 |
| 3,26902E- | FALSE | High | IsMasterP | AAEL0195 | 3:3179510 | 0 | 10 | FALSE | 3 | 8 | 3 | 387 | 427 | 769 |
| -7,26359E | FALSE | High | IsMasterP | AAEL0027 | 3:1257651 | 0 | 15 | FALSE | 3 | 8 | 3 | 271 | 309 | 637 |
| 5,2295E+1 | FALSE | High | IsMasterP | AAEL0238 | 2:1869261 | 0 | 16 | FALSE | 3 | 8 | 3 | 270 | 285 | 517 |
| 4,61359E- | FALSE | High | IsMasterP | AAEL0194 | 1:4573017 | 0 | 5 | FALSE | 2 | 6 | 2 | 616 | 626 | 815 |
| -5,7618E+ | FALSE | High | IsMasterP | AAEL0124 | 1:1234265 | 0 | 13 | FALSE | 5 | 6 | 5 | 885 | 972 | 692 |
| 5,26767E- | FALSE | High | IsMasterP | AAEL0026 | 2:4648062 | 0 | 4 | FALSE | 3 | 7 | 3 | 1348 | 1487 | 680 |
| 3,58934E- | FALSE | High | IsMasterP | AAEL0021 | 1:1022490 | 0 | 24 | FALSE | 4 | 9 | 3 | 204 | 231 | 778 |
| -2,56419E | FALSE | High | IsMasterP | AAEL0198 | 3:3345589 | 0 | 5 | FALSE | 2 | 6 | 2 | 1016 | 1087 | 887 |
| 8,42775E- | FALSE | High | IsMasterP | AAEL0103 | 2:1829098 | 0 | 7 | FALSE | 1 | 7 | 1 | 215 | 247 | 519 |
| -2,71824E | FALSE | High | IsMasterP | AAEL0224 | 3:2270169 | 0 | 10 | FALSE | 2 | 8 | 2 | 205 | 235 | 939 |
| 5,91353E- | FALSE | High | IsMasterP | AAEL0004 | 3:1628210 | 0 | 23 | FALSE | 3 | 9 | 3 | 160 | 186 | 744 |

| | | | | | | | | | | | | | | |
|---|---|---|---|---|---|---|---|---|---|---|---|---|---|---|
| -4,77471E | FALSE | High | IsMasterP | AAEL0129 | 1:205931 | 0 | 12 | FALSE | 6 | 9 | 6 | 547 | 622 | 705 |
| 1,21618E | FALSE | High | IsMasterP | AAEL0103 | 3:409377 | 0 | 8 | FALSE | 5 | 8 | 5 | 804 | 913 | 925 |
| -5,92109E | FALSE | High | IsMasterP | AAEL0045 | 2:310203 | 0 | 18 | FALSE | 3 | 7 | 3 | 226 | 261 | 954 |
| -3,62652E | FALSE | High | IsMasterP | AAEL0070 | NIGP0100 | 0 | 8 | FALSE | 3 | 7 | 3 | 497 | 576 | 900 |
| 7,16737E | FALSE | High | IsMasterP | AAEL0276 | NIGP0100 | 0 | 71 | FALSE | 2 | 4 | 2 | 99 | 105 | 947 |
| -3,50703E | FALSE | High | IsMasterP | AAEL0058 | 3:218732 | 0 | 7 | FALSE | 4 | 9 | 4 | 623 | 697 | 799 |
| 7,69721E | FALSE | High | IsMasterP | AAEL0136 | 2:334461 | 0 | 4 | FALSE | 2 | 7 | 2 | 588 | 674 | 571 |
| 3,43255E | FALSE | High | IsMasterP | AAEL0011 | 3:168923 | 0 | 9 | FALSE | 4 | 8 | 4 | 594 | 647 | 412 |
| 6,7566E+ | FALSE | High | IsMasterP | AAEL0108 | 3:520920 | 0 | 2 | FALSE | 2 | 6 | 2 | 1345 | 1455 | 884 |
| -5,52144E | FALSE | High | IsMasterP | AAEL0149 | 1:285328 | 0 | 28 | FALSE | 4 | 6 | 4 | 226 | 255 | 752 |
| -5,77052E | FALSE | High | IsMasterP | AAEL0102 | 2:341276 | 0 | 14 | FALSE | 6 | 7 | 6 | 704 | 772 | 706 |
| 4,2778E+ | FALSE | High | IsMasterP | AAEL0020 | 2:356655 | 0 | 9 | FALSE | 4 | 7 | 4 | 668 | 737 | 487 |
| -7,38232E | FALSE | High | IsMasterP | AAEL0023 | 2:303996 | 0 | 11 | FALSE | 2 | 8 | 2 | 214 | 243 | 498 |
| -4,97877E | FALSE | High | IsMasterP | AAEL0142 | 1:272119 | 0 | 10 | FALSE | 2 | 7 | 2 | 304 | 341 | 558 |
| 1,6821E+ | FALSE | High | IsMasterP | AAEL0279 | 1:261425 | 0 | 7 | FALSE | 3 | 5 | 3 | 901 | 1007 | 406 |
| 8,35937E | FALSE | High | IsMasterP | AAEL0095 | 3:344789 | 0 | 13 | FALSE | 5 | 7 | 5 | 538 | 620 | 715 |
| 3,68293E | FALSE | High | IsMasterP | AAEL0033 | 3:808265 | 0 | 8 | FALSE | 3 | 7 | 3 | 541 | 621 | 744 |
| -1,53071E | FALSE | High | IsMasterP | AAEL0290 | 2:369586 | 0 | 25 | FALSE | 5 | 6 | 5 | 307 | 338 | 655 |
| 6,67483E | FALSE | High | IsMasterP | AAEL0024 | 2:779051 | 0 | 22 | FALSE | 2 | 7 | 2 | 129 | 143 | 474 |
| -4,82124E | FALSE | High | IsMasterP | AAEL0060 | 3:737090 | 0 | 13 | FALSE | 3 | 7 | 3 | 355 | 382 | 544 |
| 4,65316E | FALSE | High | IsMasterP | AAEL0001 | 3:215836 | 0 | 30 | FALSE | 4 | 6 | 4 | 293 | 341 | 749 |
| -6,98973E | FALSE | High | IsMasterP | AAEL0276 | 2:199104 | 0 | 10 | FALSE | 2 | 5 | 2 | 476 | 490 | 1045 |
| 6,02334E | FALSE | High | IsMasterP | AAEL0137 | 2:344960 | 0 | 38 | FALSE | 3 | 6 | 3 | 138 | 166 | 505 |
| -7,92613E | FALSE | High | IsMasterP | AAEL0007 | 2:458862 | 0 | 9 | FALSE | 3 | 10 | 1 | 212 | 237 | 733 |
| -9,79868E | FALSE | High | IsMasterP | AAEL0061 | 3:687998 | 0 | 52 | FALSE | 4 | 6 | 4 | 110 | 128 | 602 |
| 1,38797E | FALSE | High | IsMasterP | AAEL0142 | 2:399885 | 0 | 19 | FALSE | 3 | 8 | 3 | 312 | 359 | 548 |
| 2,98027E | FALSE | High | IsMasterP | AAEL0081 | 3:923147 | 0 | 15 | FALSE | 3 | 7 | 3 | 363 | 395 | 909 |
| -8,79399E | FALSE | High | IsMasterP | AAEL0252 | 3:295372 | 0 | 18 | FALSE | 3 | 8 | 3 | 200 | 228 | 539 |
| 7,53378E | FALSE | High | IsMasterP | AAEL0222 | 2:177321 | 0 | 25 | FALSE | 4 | 7 | 4 | 217 | 221 | 725 |
| -5,54584E | FALSE | High | IsMasterP | AAEL0146 | 3:368627 | 0 | 4 | FALSE | 2 | 9 | 1 | 536 | 613 | 777 |
| -8,96249E | FALSE | High | IsMasterP | AAEL0062 | 3:253396 | 0 | 3 | FALSE | 3 | 7 | 3 | 1519 | 1744 | 522 |

| | | | | | | | | | | | | | | |
|---|---|---|---|---|---|---|---|---|---|---|---|---|---|---|
| -8,92355E | FALSE | High | IsMasterP | AAEL0175 | 2:125713 | 0 | 10 | FALSE | 5 | 7 | 5 | 911 | 987 | 493 |
| 3,40918E | FALSE | High | IsMasterP | AAEL0067 | 3:368337 | 0 | 4 | FALSE | 5 | 8 | 5 | 1658 | 1905 | 698 |
| -5,82306E | FALSE | High | IsMasterP | AAEL0113 | 3:201653 | 0 | 7 | FALSE | 2 | 9 | 2 | 363 | 413 | 501 |
| -8,8634E+ | FALSE | High | IsMasterP | AAEL0017 | 2:376433 | 0 | 18 | FALSE | 4 | 8 | 4 | 211 | 235 | 639 |
| -3,40774E | FALSE | High | IsMasterP | AAEL0136 | 1:617815 | 0 | 21 | FALSE | 4 | 7 | 4 | 182 | 206 | 491 |
| 8,77727E | FALSE | High | IsMasterP | AAEL0146 | 3:368585 | 0 | 4 | FALSE | 2 | 9 | 1 | 536 | 610 | 662 |
| -9,12435E | FALSE | High | IsMasterP | AAEL0047 | 2:289213 | 0 | 4 | FALSE | 4 | 7 | 4 | 1524 | 1708 | 739 |
| 3,88659E | FALSE | High | IsMasterP | AAEL0072 | 3:195957 | 0 | 11 | FALSE | 4 | 7 | 4 | 592 | 673 | 692 |
| -9,05289E | FALSE | High | IsMasterP | AAEL0217 | 1:166136 | 0 | 49 | FALSE | 2 | 5 | 2 | 103 | 120 | 545 |
| 8,62398E | FALSE | High | IsMasterP | AAEL0038 | 1:449239 | 0 | 10 | FALSE | 5 | 6 | 5 | 933 | 1023 | 572 |
| -5,34882E | FALSE | High | IsMasterP | AAEL0037 | 1:439529 | 0 | 28 | FALSE | 5 | 7 | 5 | 437 | 496 | 569 |
| -7,73201E | FALSE | High | IsMasterP | AAEL0137 | 2:277647 | 0 | 29 | FALSE | 4 | 7 | 4 | 281 | 314 | 505 |
| 5,10641E | FALSE | High | IsMasterP | AAEL0068 | 3:368438 | 0 | 9 | FALSE | 4 | 9 | 3 | 538 | 616 | 800 |
| 6,42175E | FALSE | High | IsMasterP | AAEL0029 | 3:388746 | 0 | 13 | FALSE | 3 | 6 | 3 | 417 | 454 | 1240 |
| -4,27558E | FALSE | High | IsMasterP | AAEL0056 | 2:381883 | 0 | 23 | FALSE | 3 | 6 | 3 | 184 | 205 | 488 |
| 7,02142E | FALSE | High | IsMasterP | AAEL0035 | 2:126186 | 0 | 3 | FALSE | 4 | 7 | 4 | 2247 | 2603 | 524 |
| 3,65179E | FALSE | High | IsMasterP | AAEL0044 | 3:206533 | 0 | 26 | FALSE | 3 | 7 | 3 | 159 | 173 | 479 |
| -5,3398E+ | FALSE | High | IsMasterP | AAEL0098 | 2:191777 | 0 | 27 | FALSE | 5 | 7 | 5 | 267 | 304 | 911 |
| 1,36026E | FALSE | High | IsMasterP | AAEL0146 | 2:354435 | 0 | 14 | FALSE | 3 | 6 | 3 | 452 | 493 | 750 |
| 3,40201E | FALSE | High | IsMasterP | AAEL0200 | 2:196462 | 0 | 3 | FALSE | 3 | 6 | 3 | 965 | 1055 | 651 |
| -4,61679E | FALSE | High | IsMasterP | AAEL0154 | 3:105080 | 0 | 21 | FALSE | 4 | 5 | 4 | 361 | 401 | 638 |
| -6,74405E | FALSE | High | IsMasterP | AAEL0026 | 2:464892 | 0 | 15 | FALSE | 4 | 7 | 4 | 522 | 593 | 769 |
| -7,52011E | FALSE | High | IsMasterP | AAEL0263 | 2:428799 | 0 | 27 | FALSE | 1 | 6 | 1 | 101 | 103 | 677 |
| -5,35013E | FALSE | High | IsMasterP | AAEL0106 | 3:291668 | 0 | 19 | FALSE | 2 | 6 | 2 | 156 | 178 | 536 |
| 1,21498E | FALSE | High | IsMasterP | AAEL0014 | 2:262347 | 0 | 5 | FALSE | 3 | 8 | 3 | 1053 | 1212 | 583 |
| 9,54833E | FALSE | High | IsMasterP | AAEL0235 | 2:245966 | 0 | 15 | FALSE | 3 | 7 | 3 | 264 | 298 | 895 |
| 2,14954E | FALSE | High | IsMasterP | AAEL0099 | 2:933409 | 0 | 1 | FALSE | 3 | 9 | 3 | 2383 | 2784 | 882 |
| 4,32973E | FALSE | High | IsMasterP | AAEL0038 | 3:854125 | 0 | 11 | FALSE | 2 | 7 | 2 | 418 | 463 | 592 |
| -4,6906E+ | FALSE | High | IsMasterP | AAEL0056 | 2:381777 | 0 | 14 | FALSE | 6 | 8 | 6 | 465 | 514 | 793 |
| -4,5579E+ | FALSE | High | IsMasterP | AAEL0125 | 1:285243 | 0 | 19 | FALSE | 4 | 6 | 4 | 362 | 395 | 876 |
| -7,0733E+ | FALSE | High | IsMasterP | AAEL0134 | 2:199125 | 0 | 33 | FALSE | 3 | 6 | 3 | 171 | 196 | 674 |

| | | | | | | | | | | | | | |
|---|---|---|---|---|---|---|---|---|---|---|---|---|---|
| 4,8756E+1 | FALSE | High | IsMasterP | AAEL0240 3:2373187 | 0 | 7 | FALSE | 3 | 6 | 3 | 590 | 682 | 709 |
| -4,87657E | FALSE | High | IsMasterP | AAEL0280 1:2124619 | 0 | 9 | FALSE | 4 | 7 | 4 | 690 | 794 | 715 |
| -9,18905E | FALSE | High | IsMasterP | AAEL0237 NIGP0100 | 0 | 14 | FALSE | 3 | 6 | 3 | 374 | 419 | 514 |
| 2,1265E+1 | FALSE | High | IsMasterP | AAEL0123 2:1117056 | 0 | 18 | FALSE | 3 | 8 | 3 | 243 | 266 | 597 |
| 1,16667E+ | FALSE | High | IsMasterP | AAEL0129 3:2794033 | 0 | 4 | FALSE | 1 | 7 | 1 | 378 | 428 | 625 |
| -6,16122E | FALSE | High | IsMasterP | AAEL0089 1:8090548 | 0 | 10 | FALSE | 3 | 7 | 3 | 321 | 361 | 1002 |
| -1,61965E | FALSE | High | IsMasterP | AAEL0042 2:2931158 | 0 | 9 | FALSE | 2 | 7 | 2 | 445 | 511 | 557 |
| 4,40621E+ | FALSE | High | IsMasterP | AAEL0264 2:1870810 | 0 | 4 | FALSE | 5 | 8 | 5 | 1851 | 2113 | 630 |
| 3,76417E+ | FALSE | High | IsMasterP | AAEL0029 1:2089617 | 0 | 24 | FALSE | 4 | 8 | 4 | 182 | 208 | 972 |
| 1,09911E+ | FALSE | High | IsMasterP | AAEL0133 3:1001903 | 0 | 13 | FALSE | 3 | 7 | 3 | 240 | 277 | 696 |
| -3,56281E | FALSE | High | IsMasterP | AAEL0097 2:6861615 | 0 | 22 | FALSE | 4 | 9 | 4 | 192 | 214 | 834 |
| 5,94656E+ | FALSE | High | IsMasterP | AAEL0198 2:1215871 | 0 | 16 | FALSE | 3 | 6 | 3 | 336 | 346 | 887 |
| -5,74111E | FALSE | High | IsMasterP | AAEL0052 2:3537213 | 0 | 5 | FALSE | 2 | 6 | 2 | 463 | 509 | 727 |
| -4,62082E | FALSE | High | IsMasterP | AAEL0103 3:3720400 | 0 | 10 | FALSE | 6 | 7 | 6 | 712 | 795 | 605 |
| -7,39042E | FALSE | High | IsMasterP | AAEL0120 2:1397303 | 0 | 4 | FALSE | 2 | 6 | 2 | 691 | 758 | 530 |
| -2,88769E | FALSE | High | IsMasterP | AAEL0128 3:2760001 | 0 | 15 | FALSE | 3 | 8 | 3 | 294 | 330 | 756 |
| 7,73324E+ | FALSE | High | IsMasterP | AAEL0061 3:6956714 | 0 | 10 | FALSE | 3 | 4 | 3 | 582 | 680 | 610 |
| -7,62889E | FALSE | High | IsMasterP | AAEL0137 3:3078708 | 0 | 16 | FALSE | 3 | 5 | 3 | 474 | 519 | 661 |
| -5,48439E | FALSE | High | IsMasterP | AAEL0075 2:2480307 | 0 | 6 | FALSE | 4 | 6 | 4 | 889 | 1011 | 635 |
| -6,34507E | FALSE | High | IsMasterP | AAEL0036 2:3172168 | 0 | 22 | FALSE | 3 | 6 | 3 | 219 | 245 | 469 |
| -2,38623E | FALSE | High | IsMasterP | AAEL0133 3:1733506 | 0 | 12 | FALSE | 4 | 6 | 4 | 571 | 659 | 868 |
| 7,65855E+ | FALSE | High | IsMasterP | AAEL0107 3:3278173 | 0 | 9 | FALSE | 3 | 7 | 1 | 476 | 522 | 824 |
| 7,25603E+ | FALSE | High | IsMasterP | AAEL0096 1:2225943 | 0 | 15 | FALSE | 3 | 6 | 3 | 319 | 347 | 952 |
| 6,99962E+ | FALSE | High | IsMasterP | AAEL0045 1:1446629 | 0 | 6 | FALSE | 3 | 7 | 3 | 713 | 809 | 711 |
| 8,88991E+ | FALSE | High | IsMasterP | AAEL0090 3:1348774 | 0 | 20 | FALSE | 2 | 4 | 2 | 294 | 331 | 718 |
| -8,92316E | FALSE | High | IsMasterP | AAEL0098 3:1872939 | 0 | 7 | FALSE | 4 | 7 | 4 | 813 | 951 | 630 |
| 1,29461E+ | FALSE | High | IsMasterP | AAEL0064 2:4144307 | 0 | 36 | FALSE | 5 | 7 | 5 | 275 | 292 | 829 |
| 8,14599E+ | FALSE | High | IsMasterP | AAEL0117 3:2910959 | 0 | 22 | FALSE | 3 | 6 | 3 | 277 | 300 | 894 |
| 5,48337E+ | FALSE | High | IsMasterP | AAEL0012 2:1519421 | 0 | 29 | FALSE | 5 | 6 | 5 | 249 | 279 | 832 |
| -8,46826E | FALSE | High | IsMasterP | AAEL0175 1:2793231 | 0 | 18 | FALSE | 3 | 6 | 3 | 288 | 317 | 974 |
| -5,63992E | FALSE | High | IsMasterP | AAEL0173 3:4115898 | 0 | 9 | FALSE | 3 | 7 | 1 | 500 | 509 | 994 |

| | | | | | | | | | | | | | | |
|---|---|---|---|---|---|---|---|---|---|---|---|---|---|---|
| 2,49222E- | FALSE | High | IsMasterP | AAEL0139 | 3:2304349 | 0 | 15 | FALSE | 3 | 6 | 3 | 382 | 426 | 679 |
| -8,09478E | FALSE | High | IsMasterP | AAEL0051 | 3:3459812 | 0 | 4 | FALSE | 4 | 7 | 4 | 1381 | 1507 | 816 |
| 5,86629E- | FALSE | High | IsMasterP | AAEL0148 | 2:1527519 | 0 | 20 | FALSE | 4 | 7 | 4 | 276 | 333 | 948 |
| 5,73347E- | FALSE | High | IsMasterP | AAEL0069 | 3:1955160 | 0 | 9 | FALSE | 4 | 6 | 4 | 831 | 963 | 844 |
| 4,12253E- | FALSE | High | IsMasterP | AAEL0067 | 2:8398235 | 0 | 13 | FALSE | 3 | 6 | 3 | 414 | 475 | 470 |
| -5,73855E | FALSE | High | IsMasterP | AAEL0118 | 3:1132332 | 0 | 22 | FALSE | 4 | 7 | 4 | 303 | 345 | 715 |
| 6,27851E- | FALSE | High | IsMasterP | AAEL0145 | 2:4522519 | 0 | 14 | FALSE | 5 | 7 | 5 | 395 | 448 | 901 |
| 8,70084E- | FALSE | High | IsMasterP | AAEL0113 | 1:1208950 | 0 | 19 | FALSE | 4 | 8 | 4 | 252 | 286 | 938 |
| -4,0053E+ | FALSE | High | IsMasterP | AAEL0034 | 2:2163432 | 0 | 11 | FALSE | 4 | 7 | 4 | 430 | 496 | 819 |
| -1,50569E | FALSE | High | IsMasterP | AAEL0236 | 2:3409639 | 0 | 8 | FALSE | 5 | 6 | 5 | 875 | 974 | 717 |
| 1,68712E- | FALSE | High | IsMasterP | AAEL0025 | 3:2565089 | 0 | 14 | FALSE | 2 | 6 | 2 | 303 | 324 | 712 |
| -7,48518E | FALSE | High | IsMasterP | AAEL0232 | 2:2278245 | 0 | 12 | FALSE | 2 | 7 | 2 | 129 | 147 | 992 |
| -4,08514E | FALSE | High | IsMasterP | AAEL0081 | 3:1010383 | 0 | 19 | FALSE | 3 | 6 | 3 | 182 | 205 | 511 |
| 6,79187E- | FALSE | High | IsMasterP | AAEL0220 | 2:4648304 | 0 | 4 | FALSE | 4 | 8 | 4 | 1249 | 1382 | 488 |
| 2,58942E- | FALSE | High | IsMasterP | AAEL0290 | 2:3304110 | 0 | 73 | FALSE | 3 | 6 | 3 | 106 | 112 | 890 |
| 7,42717E- | FALSE | High | IsMasterP | AAEL0083 | 3:4148009 | 0 | 7 | FALSE | 2 | 8 | 2 | 522 | 568 | 878 |
| -7,82695E | FALSE | High | IsMasterP | AAEL0174 | 3:2923104 | 0 | 40 | FALSE | 4 | 5 | 4 | 221 | 253 | 541 |
| -9,06044E | FALSE | High | IsMasterP | AAEL0094 | 1:5742437 | 0 | 18 | FALSE | 5 | 6 | 5 | 330 | 373 | 679 |
| 5,45484E- | FALSE | High | IsMasterP | AAEL0034 | 3:3497761 | 0 | 7 | FALSE | 2 | 7 | 2 | 368 | 413 | 752 |
| -6,89279E | FALSE | High | IsMasterP | AAEL0175 | 3:2243802 | 0 | 11 | FALSE | 2 | 4 | 2 | 513 | 572 | 596 |
| 4,5096E+1 | FALSE | High | IsMasterP | AAEL0143 | 2:1723760 | 0 | 29 | FALSE | 3 | 6 | 3 | 174 | 200 | 514 |
| -6,34844E | FALSE | High | IsMasterP | AAEL0011 | 2:1488931 | 0 | 9 | FALSE | 4 | 6 | 4 | 866 | 994 | 506 |
| -5,40399E | FALSE | High | IsMasterP | AAEL0096 | 3:3664935 | 0 | 10 | FALSE | 4 | 5 | 4 | 570 | 608 | 957 |
| 2,5692E+1 | FALSE | High | IsMasterP | AAEL0028 | 1:2561138 | 0 | 3 | FALSE | 5 | 7 | 5 | 2603 | 2864 | 744 |
| 4,26808E- | FALSE | High | IsMasterP | AAEL0014 | 2:4767847 | 0 | 12 | FALSE | 2 | 7 | 2 | 274 | 300 | 539 |
| 5,57394E- | FALSE | High | IsMasterP | AAEL0140 | 3:1474812 | 0 | 20 | FALSE | 2 | 4 | 2 | 259 | 279 | 535 |
| 8,83527E- | FALSE | High | IsMasterP | AAEL0117 | 3:3564254 | 0 | 4 | FALSE | 3 | 7 | 3 | 1131 | 1274 | 644 |
| 5,67558E- | FALSE | High | IsMasterP | AAEL0142 | 2:1718483 | 0 | 12 | FALSE | 3 | 5 | 3 | 515 | 593 | 671 |
| -7,58667E | FALSE | High | IsMasterP | AAEL0111 | 3:5157220 | 0 | 16 | FALSE | 4 | 7 | 3 | 354 | 386 | 863 |
| 3,56778E- | FALSE | High | IsMasterP | AAEL0113 | 1:1259220 | 0 | 6 | FALSE | 5 | 8 | 5 | 974 | 1102 | 506 |
| 8,92974E- | FALSE | High | IsMasterP | AAEL0142 | 1:2857504 | 0 | 11 | FALSE | 3 | 6 | 3 | 237 | 276 | 601 |

| | | | | | | | | | | | | | | |
|---|---|---|---|---|---|---|---|---|---|---|---|---|---|---|
| -6,6636E+ | FALSE | High | IsMasterP | AAEL0108 | 3:156580 | 0 | 6 | FALSE | 4 | 5 | 4 | 1054 | 1150 | 578 |
| -3,92184E | FALSE | High | IsMasterP | AAEL0173 | 3:411842 | 0 | 11 | FALSE | 3 | 6 | 1 | 512 | 522 | 977 |
| -4,63375E | FALSE | High | IsMasterP | AAEL0268 | 1:278801 | 0 | 8 | FALSE | 2 | 5 | 2 | 488 | 520 | 550 |
| -2,27811E | FALSE | High | IsMasterP | AAEL0022 | 3:375098 | 0 | 6 | FALSE | 1 | 5 | 1 | 265 | 306 | 592 |
| -6,90427E | FALSE | High | IsMasterP | AAEL0018 | 2:375245 | 0 | 52 | FALSE | 4 | 5 | 4 | 128 | 149 | 533 |
| 1,8266E+ | FALSE | High | IsMasterP | AAEL0087 | 3:326006 | 0 | 16 | FALSE | 2 | 6 | 2 | 275 | 314 | 489 |
| 4,09397E- | FALSE | High | IsMasterP | AAEL0021 | 3:176909 | 0 | 18 | FALSE | 5 | 6 | 5 | 451 | 480 | 753 |
| -6,4983E+ | FALSE | High | IsMasterP | AAEL0241 | 2:464665 | 0 | 4 | FALSE | 4 | 7 | 4 | 1720 | 1905 | 821 |
| 6,78659E- | FALSE | High | IsMasterP | AAEL0110 | 2:631361 | 0 | 52 | FALSE | 2 | 5 | 2 | 85 | 94 | 521 |
| 5,87628E- | FALSE | High | IsMasterP | AAEL0008 | 3:335086 | 0 | 9 | FALSE | 3 | 7 | 3 | 535 | 576 | 521 |
| 4,61659E- | FALSE | High | IsMasterP | AAEL0095 | 3:210810 | 0 | 10 | FALSE | 4 | 5 | 4 | 934 | 1055 | 775 |
| 9,68754E- | FALSE | High | IsMasterP | AAEL0000 | 1:242653 | 0 | 5 | FALSE | 4 | 5 | 4 | 2186 | 2302 | 517 |
| -7,53006E | FALSE | High | IsMasterP | AAEL0194 | 1:237656 | 0 | 5 | FALSE | 5 | 7 | 5 | 1533 | 1782 | 571 |
| 2,69857E- | FALSE | High | IsMasterP | AAEL0275 | 3:242792 | 0 | 4 | FALSE | 3 | 6 | 3 | 1436 | 1600 | 676 |
| 1,90546E- | FALSE | High | IsMasterP | AAEL0195 | 3:202578 | 0 | 2 | FALSE | 3 | 6 | 3 | 2596 | 2953 | 621 |
| 8,05798E- | FALSE | High | IsMasterP | AAEL0012 | 2:411489 | 0 | 28 | FALSE | 5 | 7 | 5 | 314 | 351 | 703 |
| 4,14151E- | FALSE | High | IsMasterP | AAEL0084 | 2:190992 | 0 | 22 | FALSE | 2 | 6 | 2 | 125 | 149 | 815 |
| -6,5697E+ | FALSE | High | IsMasterP | AAEL0206 | 3:388809 | 0 | 4 | FALSE | 2 | 6 | 2 | 1002 | 1126 | 484 |
| 9,25785E- | FALSE | High | IsMasterP | AAEL0290 | 2:187003 | 0 | 11 | FALSE | 1 | 5 | 1 | 152 | 179 | 592 |
| -2,66377E | FALSE | High | IsMasterP | AAEL0064 | 3:216516 | 0 | 18 | FALSE | 3 | 7 | 3 | 314 | 349 | 703 |
| 3,44371E- | FALSE | High | IsMasterP | AAEL0070 | NIGP0100 | 0 | 5 | FALSE | 4 | 8 | 4 | 881 | 993 | 695 |
| -4,14997E | FALSE | High | IsMasterP | AAEL0111 | 2:101527 | 0 | 14 | FALSE | 3 | 7 | 2 | 353 | 404 | 620 |
| 3,8282E+ | FALSE | High | IsMasterP | AAEL0095 | 2:356996 | 0 | 6 | FALSE | 1 | 5 | 1 | 281 | 312 | 571 |
| 8,10166E- | FALSE | High | IsMasterP | AAEL0081 | 3:302075 | 0 | 13 | FALSE | 4 | 6 | 4 | 660 | 730 | 828 |
| 3,03148E- | FALSE | High | IsMasterP | AAEL0086 | 2:239676 | 0 | 18 | FALSE | 2 | 6 | 2 | 152 | 173 | 618 |
| -8,37813E | FALSE | High | IsMasterP | AAEL0008 | 3:332923 | 0 | 13 | FALSE | 4 | 7 | 4 | 573 | 629 | 655 |
| 1,65216E- | FALSE | High | IsMasterP | AAEL0053 | 2:370327 | 0 | 23 | FALSE | 4 | 7 | 4 | 318 | 346 | 822 |
| -2,72233E | FALSE | High | IsMasterP | AAEL0195 | 2:211644 | 0 | 11 | FALSE | 6 | 6 | 6 | 804 | 900 | 778 |
| -6,54001E | FALSE | High | IsMasterP | AAEL0074 | 2:134748 | 0 | 41 | FALSE | 3 | 8 | 3 | 64 | 76 | 976 |
| -1,59011E | FALSE | High | IsMasterP | AAEL0062 | 3:244310 | 0 | 11 | FALSE | 3 | 5 | 3 | 565 | 629 | 794 |
| 2,73512E- | FALSE | High | IsMasterP | AAEL0127 | 3:322098 | 0 | 3 | FALSE | 3 | 7 | 3 | 1354 | 1480 | 637 |

| | | | | | | | | | | | | | | |
|---|---|---|---|---|---|---|---|---|---|---|---|---|---|---|
| -6,8676E+ | FALSE | High | IsMasterP | AAEL0075 | NIGP0100 | 0 | 18 | FALSE | 4 | 6 | 4 | 358 | 411 | 810 |
| 6,25696E | FALSE | High | IsMasterP | AAEL0111 | 3:378663 | 0 | 16 | FALSE | 4 | 7 | 4 | 364 | 393 | 695 |
| 8,68883E | FALSE | High | IsMasterP | AAEL0245 | 2:452464 | 0 | 4 | FALSE | 4 | 7 | 4 | 1895 | 2189 | 870 |
| 3,45495E | FALSE | High | IsMasterP | AAEL0040 | 3:166942 | 0 | 14 | FALSE | 4 | 6 | 2 | 307 | 352 | 512 |
| 5,52763E | FALSE | High | IsMasterP | AAEL0278 | 1:574736 | 0 | 17 | FALSE | 4 | 5 | 4 | 531 | 585 | 477 |
| 4,84688E | FALSE | High | IsMasterP | AAEL0228 | 3:295063 | 0 | 19 | FALSE | 4 | 6 | 4 | 261 | 294 | 529 |
| -9,08837E | FALSE | High | IsMasterP | AAEL0104 | 1:278396 | 0 | 6 | FALSE | 4 | 7 | 4 | 1072 | 1192 | 793 |
| 6,1365E+ | FALSE | High | IsMasterP | AAEL0092 | 3:200193 | 0 | 12 | FALSE | 2 | 6 | 2 | 281 | 321 | 558 |
| 1,51753E | FALSE | High | IsMasterP | AAEL0136 | 3:195112 | 0 | 6 | FALSE | 3 | 6 | 3 | 887 | 1000 | 734 |
| -3,62842E | FALSE | High | IsMasterP | AAEL0118 | 3:113291 | 0 | 5 | FALSE | 4 | 7 | 4 | 1121 | 1252 | 711 |
| 9,21581E | FALSE | High | IsMasterP | AAEL0140 | 2:446155 | 0 | 12 | FALSE | 3 | 5 | 3 | 582 | 660 | 592 |
| -5,67206E | FALSE | High | IsMasterP | AAEL0010 | 3:551465 | 0 | 7 | FALSE | 3 | 4 | 3 | 852 | 975 | 890 |
| -8,98241E | FALSE | High | IsMasterP | AAEL0083 | 3:414065 | 0 | 7 | FALSE | 3 | 6 | 3 | 957 | 1059 | 731 |
| 6,72818E | FALSE | High | IsMasterP | AAEL0092 | 2:228427 | 0 | 4 | FALSE | 2 | 6 | 2 | 612 | 673 | 638 |
| 2,50068E | FALSE | High | IsMasterP | AAEL0099 | 1:194137 | 0 | 18 | FALSE | 3 | 5 | 3 | 322 | 359 | 888 |
| -9,08995E | FALSE | High | IsMasterP | AAEL0064 | 3:216387 | 0 | 1 | FALSE | 3 | 6 | 3 | 3432 | 3804 | 627 |
| 4,99152E | FALSE | High | IsMasterP | AAEL0258 | 3:388796 | 0 | 3 | FALSE | 1 | 4 | 1 | 568 | 643 | 951 |
| 8,41604E | FALSE | High | IsMasterP | AAEL0104 | 1:278612 | 0 | 25 | FALSE | 6 | 6 | 6 | 217 | 250 | 627 |
| 7,68517E | FALSE | High | IsMasterP | AAEL0082 | 3:250453 | 0 | 20 | FALSE | 3 | 5 | 3 | 286 | 315 | 479 |
| 6,77636E | FALSE | High | IsMasterP | AAEL0100 | 2:110764 | 0 | 11 | FALSE | 2 | 7 | 2 | 193 | 213 | 680 |
| -4,25683E | FALSE | High | IsMasterP | AAEL0065 | 2:584874 | 0 | 7 | FALSE | 3 | 5 | 3 | 623 | 696 | 607 |
| 5,39814E | FALSE | High | IsMasterP | AAEL0046 | 1:145817 | 0 | 4 | FALSE | 5 | 6 | 5 | 1725 | 1977 | 621 |
| 4,46804E | FALSE | High | IsMasterP | AAEL0256 | 3:192127 | 0 | 4 | FALSE | 3 | 6 | 3 | 1515 | 1713 | 574 |
| -6,88333E | FALSE | High | IsMasterP | AAEL0040 | 1:135748 | 0 | 20 | FALSE | 3 | 7 | 3 | 212 | 236 | 740 |
| 2,60331E | FALSE | High | IsMasterP | AAEL0143 | 1:238746 | 0 | 12 | FALSE | 3 | 6 | 3 | 537 | 611 | 759 |
| -1,89751E | FALSE | High | IsMasterP | AAEL0098 | 3:203118 | 0 | 25 | FALSE | 5 | 6 | 5 | 322 | 364 | 909 |
| 3,58764E | FALSE | High | IsMasterP | AAEL0086 | 2:362958 | 0 | 27 | FALSE | 3 | 5 | 3 | 188 | 215 | 715 |
| 4,55873E | FALSE | High | IsMasterP | AAEL0024 | 1:177142 | 0 | 5 | FALSE | 2 | 5 | 2 | 878 | 952 | 802 |
| -4,00466E | FALSE | High | IsMasterP | AAEL0090 | 2:269458 | 0 | 5 | FALSE | 2 | 4 | 2 | 1103 | 1183 | 926 |
| -4,0046E+ | FALSE | High | IsMasterP | AAEL0004 | 3:183635 | 0 | 6 | FALSE | 2 | 7 | 2 | 216 | 251 | 664 |
| 3,59174E | FALSE | High | IsMasterP | AAEL0042 | 2:118591 | 0 | 17 | FALSE | 4 | 6 | 4 | 190 | 218 | 525 |

| | | | | | | | | | | | | | | |
|---|---|---|---|---|---|---|---|---|---|---|---|---|---|---|
| 2,33251E- | FALSE | High | IsMasterP | AAEL0014 | 2:2621098 | 0 | 5 | FALSE | 2 | 4 | 2 | 722 | 797 | 734 |
| -2,95056E | FALSE | High | IsMasterP | AAEL0151 | 2:1277668 | 0 | 6 | FALSE | 3 | 4 | 3 | 1942 | 2001 | 860 |
| -3,32706E | FALSE | High | IsMasterP | AAEL0083 | 3:4139876 | 0 | 29 | FALSE | 3 | 6 | 3 | 139 | 152 | 787 |
| 6,68975E- | FALSE | High | IsMasterP | AAEL0088 | 2:2413992 | 0 | 22 | FALSE | 4 | 5 | 4 | 313 | 359 | 644 |
| 5,10613E- | FALSE | High | IsMasterP | AAEL0033 | 3:8080294 | 0 | 18 | FALSE | 2 | 6 | 2 | 156 | 176 | 538 |
| -1,14922E | FALSE | High | IsMasterP | AAEL0068 | 1:8334837 | 0 | 3 | FALSE | 1 | 6 | 1 | 510 | 584 | 903 |
| -7,0986E+ | FALSE | High | IsMasterP | AAEL0273 | 1:1847186 | 0 | 24 | FALSE | 5 | 6 | 5 | 271 | 307 | 870 |
| -8,06041E | FALSE | High | IsMasterP | AAEL0024 | 2:7773188 | 0 | 4 | FALSE | 4 | 8 | 4 | 987 | 1098 | 758 |
| 4,75127E- | FALSE | High | IsMasterP | AAEL0137 | 1:2193364 | 0 | 9 | FALSE | 3 | 4 | 3 | 620 | 670 | 894 |
| -8,94137E | FALSE | High | IsMasterP | AAEL0245 | NIGP0100 | 0 | 16 | FALSE | 4 | 5 | 4 | 527 | 578 | 576 |
| -7,03787E | FALSE | High | IsMasterP | AAEL0246 | 3:1049496 | 0 | 7 | FALSE | 2 | 7 | 2 | 398 | 439 | 686 |
| 1,50861E- | FALSE | High | IsMasterP | AAEL0059 | 3:1546684 | 0 | 17 | FALSE | 4 | 7 | 4 | 321 | 347 | 958 |
| -2,14495E | FALSE | High | IsMasterP | AAEL0050 | 2:4460461 | 0 | 7 | FALSE | 1 | 4 | 1 | 355 | 383 | 969 |
| -1,06374E | FALSE | High | IsMasterP | AAEL0109 | 3:1904745 | 0 | 3 | FALSE | 6 | 6 | 6 | 2221 | 2459 | 501 |
| -1,36018E | FALSE | High | IsMasterP | AAEL0049 | 2:2818577 | 0 | 12 | FALSE | 2 | 6 | 2 | 291 | 332 | 544 |
| -7,27823E | FALSE | High | IsMasterP | AAEL0072 | 2:4223877 | 0 | 2 | FALSE | 2 | 4 | 2 | 2387 | 2691 | 739 |
| -4,18589E | FALSE | High | IsMasterP | AAEL0097 | 1:2097347 | 0 | 21 | FALSE | 2 | 4 | 2 | 301 | 341 | 479 |
| -1,02484E | FALSE | High | IsMasterP | AAEL0290 | 1:8348377 | 0 | 17 | FALSE | 4 | 8 | 2 | 294 | 330 | 643 |
| 7,16658E- | FALSE | High | IsMasterP | AAEL0113 | 1:1352344 | 0 | 19 | FALSE | 3 | 5 | 3 | 193 | 217 | 680 |
| -7,82458E | FALSE | High | IsMasterP | AAEL0002 | 1:6560805 | 0 | 10 | FALSE | 4 | 5 | 4 | 736 | 826 | 720 |
| 9,58157E- | FALSE | High | IsMasterP | AAEL0035 | 2:2562656 | 0 | 9 | FALSE | 3 | 5 | 3 | 335 | 377 | 643 |
| -4,12445E | FALSE | High | IsMasterP | AAEL0036 | 2:3167409 | 0 | 10 | FALSE | 3 | 5 | 3 | 500 | 568 | 648 |
| 1,94896E- | FALSE | High | IsMasterP | AAEL0230 | 1:7318510 | 0 | 24 | FALSE | 2 | 4 | 2 | 207 | 221 | 543 |
| -3,18382E | FALSE | High | IsMasterP | AAEL0021 | 3:1785138 | 0 | 25 | FALSE | 2 | 5 | 2 | 151 | 172 | 599 |
| 2,18697E- | FALSE | High | IsMasterP | AAEL0027 | 2:1383188 | 0 | 7 | FALSE | 4 | 6 | 4 | 870 | 980 | 936 |
| -4,11004E | FALSE | High | IsMasterP | AAEL0029 | 1:8604539 | 0 | 13 | FALSE | 4 | 5 | 4 | 443 | 490 | 777 |
| 3,85758E- | FALSE | High | IsMasterP | AAEL0054 | 3:3829964 | 0 | 12 | FALSE | 4 | 6 | 4 | 566 | 633 | 887 |
| -6,38892E | FALSE | High | IsMasterP | AAEL0140 | 1:1429712 | 0 | 9 | FALSE | 2 | 5 | 2 | 422 | 468 | 714 |
| -3,90316E | FALSE | High | IsMasterP | AAEL0246 | 2:2567596 | 0 | 12 | FALSE | 1 | 5 | 1 | 165 | 172 | 885 |
| -7,51822E | FALSE | High | IsMasterP | AAEL0077 | 2:4702826 | 0 | 14 | FALSE | 3 | 7 | 3 | 351 | 390 | 684 |
| 4,1906E+1 | FALSE | High | IsMasterP | AAEL0237 | NIGP0100 | 0 | 5 | FALSE | 2 | 5 | 2 | 904 | 1026 | 555 |

| | | | | | | | | | | | | | | |
|---|---|---|---|---|---|---|---|---|---|---|---|---|---|---|
| 9,18808E | FALSE | High | IsMasterP | AAEL0080 | 2:231101 | 0 | 3 | FALSE | 3 | 8 | 3 | 1064 | 1192 | 497 |
| 2,83991E | FALSE | High | IsMasterP | AAEL0088 | 3:236705 | 0 | 8 | FALSE | 3 | 5 | 3 | 739 | 855 | 923 |
| 3,27988E | FALSE | High | IsMasterP | AAEL0150 | 1:115998 | 0 | 16 | FALSE | 3 | 5 | 3 | 238 | 257 | 796 |
| 1,56174E | FALSE | High | IsMasterP | AAEL0046 | 1:301839 | 0 | 42 | FALSE | 4 | 6 | 4 | 167 | 195 | 914 |
| -7,56405E | FALSE | High | IsMasterP | AAEL0094 | 2:151893 | 0 | 6 | FALSE | 3 | 5 | 3 | 1120 | 1181 | 844 |
| 8,40426E | FALSE | High | IsMasterP | AAEL0004 | 3:184139 | 0 | 20 | FALSE | 4 | 4 | 4 | 476 | 550 | 925 |
| -3,64282E | FALSE | High | IsMasterP | AAEL0024 | 3:338319 | 0 | 4 | FALSE | 2 | 7 | 2 | 585 | 639 | 709 |
| -6,61566E | FALSE | High | IsMasterP | AAEL0078 | 2:461181 | 0 | 3 | FALSE | 1 | 5 | 1 | 537 | 613 | 856 |
| 9,18151E | FALSE | High | IsMasterP | AAEL0138 | 1:358279 | 0 | 7 | FALSE | 4 | 5 | 4 | 785 | 876 | 609 |
| 7,36502E | FALSE | High | IsMasterP | AAEL0148 | 1:278602 | 0 | 14 | FALSE | 4 | 6 | 4 | 513 | 546 | 780 |
| -6,79725E | FALSE | High | IsMasterP | AAEL0014 | 2:456753 | 0 | 29 | FALSE | 1 | 3 | 1 | 108 | 123 | 715 |
| 2,76901E | FALSE | High | IsMasterP | AAEL0002 | 2:754127 | 0 | 17 | FALSE | 2 | 5 | 2 | 172 | 198 | 517 |
| -1,41882E | FALSE | High | IsMasterP | AAEL0033 | 3:340179 | 0 | 44 | FALSE | 2 | 5 | 2 | 94 | 108 | 517 |
| 6,35769E | FALSE | High | IsMasterP | AAEL0219 | 2:428896 | 0 | 13 | FALSE | 1 | 4 | 1 | 126 | 126 | 954 |
| -1,38073E | FALSE | High | IsMasterP | AAEL0010 | 3:555347 | 0 | 6 | FALSE | 4 | 5 | 4 | 1263 | 1272 | 986 |
| -7,76728E | FALSE | High | IsMasterP | AAEL0084 | 1:148979 | 0 | 10 | FALSE | 3 | 5 | 3 | 531 | 584 | 501 |
| -2,71032E | FALSE | High | IsMasterP | AAEL0208 | 2:128407 | 0 | 6 | FALSE | 3 | 5 | 3 | 1264 | 1355 | 577 |
| -2,68163E | FALSE | High | IsMasterP | AAEL0278 | 2:312668 | 0 | 11 | FALSE | 2 | 6 | 2 | 208 | 240 | 539 |
| 5,13417E | FALSE | High | IsMasterP | AAEL0138 | 1:181455 | 0 | 11 | FALSE | 4 | 4 | 4 | 766 | 848 | 897 |
| 7,96017E | FALSE | High | IsMasterP | AAEL0005 | 3:385454 | 0 | 15 | FALSE | 3 | 5 | 3 | 375 | 412 | 657 |
| -8,93016E | FALSE | High | IsMasterP | AAEL0013 | 3:477376 | 0 | 16 | FALSE | 3 | 6 | 3 | 176 | 192 | 955 |
| -3,72801E | FALSE | High | IsMasterP | AAEL0051 | 2:130766 | 0 | 10 | FALSE | 1 | 5 | 1 | 139 | 144 | 507 |
| -1,4984E+ | FALSE | High | IsMasterP | AAEL0098 | 2:164155 | 0 | 3 | FALSE | 2 | 7 | 2 | 652 | 751 | 941 |
| -6,0899E+ | FALSE | High | IsMasterP | AAEL0106 | 2:176242 | 0 | 7 | FALSE | 2 | 7 | 2 | 397 | 459 | 517 |
| -7,10105E | FALSE | High | IsMasterP | AAEL0181 | 2:557401 | 0 | 7 | FALSE | 3 | 7 | 3 | 550 | 620 | 655 |
| 6,74869E | FALSE | High | IsMasterP | AAEL0172 | 2:474324 | 0 | 17 | FALSE | 1 | 4 | 1 | 104 | 116 | 392 |
| 6,25949E | FALSE | High | IsMasterP | AAEL0048 | 3:273428 | 0 | 8 | FALSE | 4 | 6 | 4 | 845 | 925 | 582 |
| 4,76611E | FALSE | High | IsMasterP | AAEL0086 | 2:215358 | 0 | 15 | FALSE | 2 | 5 | 2 | 325 | 357 | 572 |
| -2,86546E | FALSE | High | IsMasterP | AAEL0077 | 3:255334 | 0 | 10 | FALSE | 2 | 5 | 2 | 323 | 384 | 470 |
| -8,75247E | FALSE | High | IsMasterP | AAEL0109 | 3:167733 | 0 | 12 | FALSE | 2 | 7 | 2 | 199 | 223 | 512 |
| 8,67878E | FALSE | High | IsMasterP | AAEL0095 | 3:527709 | 0 | 21 | FALSE | 5 | 6 | 5 | 297 | 345 | 747 |

| | | | | | | | | | | | | | | |
|---|---|---|---|---|---|---|---|---|---|---|---|---|---|---|
| -3,79999E | FALSE | High | IsMasterP | AAEL0195 | 3:121468( | 0 | 7 | FALSE | 3 | 5 | 3 | 666 | 721 | 717 |
| 4,672E+17 | FALSE | High | IsMasterP | AAEL0247 | 1:2035781 | 0 | 3 | FALSE | 2 | 6 | 2 | 1007 | 1138 | 828 |
| -4,10335E | FALSE | High | IsMasterP | AAEL0127 | 1:9312416 | 0 | 24 | FALSE | 2 | 4 | 2 | 178 | 197 | 581 |
| 8,00418E- | FALSE | High | IsMasterP | AAEL0034 | 3:3487209 | 0 | 7 | FALSE | 4 | 4 | 4 | 1182 | 1338 | 846 |
| -3,18646E | FALSE | High | IsMasterP | AAEL0137 | 2:2941127 | 0 | 30 | FALSE | 4 | 6 | 4 | 227 | 253 | 969 |
| 5,41107E- | FALSE | High | IsMasterP | AAEL0093 | 3:8353529 | 0 | 7 | FALSE | 5 | 6 | 5 | 1082 | 1242 | 613 |
| 6,99505E- | FALSE | High | IsMasterP | AAEL0027 | 3:1293035 | 0 | 12 | FALSE | 2 | 5 | 2 | 320 | 336 | 522 |
| -5,27517E | FALSE | High | IsMasterP | AAEL0216 | 1:3078364 | 0 | 3 | FALSE | 3 | 6 | 3 | 1665 | 1811 | 514 |
| -7,24814E | FALSE | High | IsMasterP | AAEL0075 | 1:1986762 | 0 | 25 | FALSE | 3 | 4 | 3 | 222 | 248 | 526 |
| 1,57487E- | FALSE | High | IsMasterP | AAEL0198 | 3:9122615 | 0 | 5 | FALSE | 2 | 4 | 2 | 924 | 1040 | 545 |
| 6,22669E- | FALSE | High | IsMasterP | AAEL0194 | NIGP0100 | 0 | 66 | FALSE | 3 | 4 | 3 | 125 | 131 | 909 |
| 4,8817E+1 | FALSE | High | IsMasterP | AAEL0007 | 2:5538509 | 0 | 5 | FALSE | 2 | 5 | 1 | 671 | 754 | 761 |
| -8,53952E | FALSE | High | IsMasterP | AAEL0007 | 2:4587788 | 0 | 7 | FALSE | 2 | 6 | 2 | 382 | 419 | 500 |
| -9,32817E | FALSE | High | IsMasterP | AAEL0275 | 2:2134928 | 0 | 14 | FALSE | 3 | 4 | 3 | 520 | 597 | 783 |
| 2,57873E- | FALSE | High | IsMasterP | AAEL0013 | 3:4775475 | 0 | 9 | FALSE | 5 | 6 | 5 | 811 | 910 | 909 |
| -3,44536E | FALSE | High | IsMasterP | AAEL0147 | 1:4568143 | 0 | 4 | FALSE | 1 | 6 | 1 | 446 | 515 | 923 |
| -4,18997E | FALSE | High | IsMasterP | AAEL0013 | 2:2062895 | 0 | 18 | FALSE | 3 | 5 | 3 | 390 | 427 | 544 |
| -6,11586E | FALSE | High | IsMasterP | AAEL0058 | 1:2012897 | 0 | 11 | FALSE | 3 | 5 | 3 | 498 | 549 | 505 |
| -4,04202E | FALSE | High | IsMasterP | AAEL0021 | 2:4284528 | 0 | 14 | FALSE | 3 | 5 | 3 | 325 | 371 | 758 |
| 4,40256E- | FALSE | High | IsMasterP | AAEL0069 | 1:1573106 | 0 | 28 | FALSE | 4 | 5 | 4 | 295 | 323 | 569 |
| -2,95678E | FALSE | High | IsMasterP | AAEL0093 | 3:3808639 | 0 | 6 | FALSE | 3 | 6 | 3 | 589 | 664 | 573 |
| 5,14921E- | FALSE | High | IsMasterP | AAEL0035 | 3:2635733 | 0 | 13 | FALSE | 3 | 5 | 3 | 469 | 501 | 887 |
| 8,72482E- | FALSE | High | IsMasterP | AAEL0276 | 1:7538850 | 0 | 7 | FALSE | 2 | 5 | 2 | 364 | 407 | 847 |
| 6,18906E- | FALSE | High | IsMasterP | AAEL0076 | 2:1072082 | 0 | 6 | FALSE | 3 | 4 | 3 | 1130 | 1274 | 516 |
| -4,63904E | FALSE | High | IsMasterP | AAEL0068 | 1:8339241 | 0 | 2 | FALSE | 1 | 5 | 1 | 966 | 1079 | 838 |
| 4,1834E+1 | FALSE | High | IsMasterP | AAEL0086 | 2:4548825 | 0 | 8 | FALSE | 2 | 6 | 2 | 377 | 436 | 637 |
| 5,94254E- | FALSE | High | IsMasterP | AAEL0085 | 2:9922133 | 0 | 8 | FALSE | 3 | 5 | 3 | 646 | 724 | 683 |
| -4,15995E | FALSE | High | IsMasterP | AAEL0136 | 1:4298969 | 0 | 11 | FALSE | 3 | 6 | 3 | 322 | 370 | 468 |
| 4,9214E+1 | FALSE | High | IsMasterP | AAEL0141 | 2:3058001 | 0 | 15 | FALSE | 6 | 6 | 6 | 465 | 526 | 574 |
| -4,00114E | FALSE | High | IsMasterP | AAEL0089 | 1:8092941 | 0 | 6 | FALSE | 2 | 5 | 2 | 441 | 488 | 1033 |
| 2,12414E- | FALSE | High | IsMasterP | AAEL0201 | 3:3119397 | 0 | 24 | FALSE | 2 | 6 | 2 | 123 | 137 | 680 |

| | | | | | | | | | | | | | | |
|---|---|---|---|---|---|---|---|---|---|---|---|---|---|---|
| 4,36092E | FALSE | High | IsMasterP | AAEL0034 | 2:150064 | 0 | 24 | FALSE | 3 | 5 | 3 | 184 | 204 | 771 |
| 4,22862E | FALSE | High | IsMasterP | AAEL0070 | 3:299688 | 0 | 11 | FALSE | 1 | 6 | 1 | 107 | 120 | 904 |
| 3,77061E | FALSE | High | IsMasterP | AAEL0011 | 2:307251 | 0 | 9 | FALSE | 1 | 7 | 1 | 109 | 123 | 1013 |
| 7,67727E | FALSE | High | IsMasterP | AAEL0004 | 3:187741 | 0 | 14 | FALSE | 2 | 6 | 2 | 185 | 213 | 960 |
| -8,53474E | FALSE | High | IsMasterP | AAEL0214 | 1:283465 | 0 | 5 | FALSE | 3 | 6 | 3 | 1115 | 1248 | 614 |
| -5,22755E | FALSE | High | IsMasterP | AAEL0070 | 3:299780 | 0 | 18 | FALSE | 2 | 5 | 2 | 176 | 195 | 865 |
| 2,63584E | FALSE | High | IsMasterP | AAEL0114 | 3:142554 | 0 | 3 | FALSE | 2 | 4 | 2 | 1120 | 1268 | 948 |
| 1,75929E | FALSE | High | IsMasterP | AAEL0246 | 1:189818 | 0 | 19 | FALSE | 4 | 5 | 4 | 318 | 345 | 799 |
| -8,65499E | FALSE | High | IsMasterP | AAEL0076 | 2:470785 | 0 | 8 | FALSE | 4 | 4 | 1 | 951 | 1055 | 600 |
| 8,3417E+ | FALSE | High | IsMasterP | AAEL0068 | 3:370293 | 0 | 10 | FALSE | 2 | 7 | 2 | 300 | 334 | 772 |
| -1,67088E | FALSE | High | IsMasterP | AAEL0231 | 1:155089 | 0 | 41 | FALSE | 3 | 5 | 1 | 150 | 167 | 503 |
| -7,95708E | FALSE | High | IsMasterP | AAEL0075 | 2:217514 | 0 | 15 | FALSE | 2 | 4 | 2 | 239 | 280 | 882 |
| -6,69212E | FALSE | High | IsMasterP | AAEL0012 | 2:413441 | 0 | 13 | FALSE | 3 | 6 | 3 | 408 | 461 | 736 |
| 3,95984E | FALSE | High | IsMasterP | AAEL0064 | 2:439488 | 0 | 3 | FALSE | 2 | 6 | 2 | 813 | 919 | 687 |
| -3,28823E | FALSE | High | IsMasterP | AAEL0044 | 3:205790 | 0 | 14 | FALSE | 3 | 6 | 3 | 215 | 248 | 533 |
| 6,42367E | FALSE | High | IsMasterP | AAEL0139 | 2:965606 | 0 | 8 | FALSE | 4 | 6 | 4 | 852 | 983 | 583 |
| 4,07362E | FALSE | High | IsMasterP | AAEL0260 | 2:141070 | 0 | 3 | FALSE | 4 | 5 | 4 | 2446 | 2748 | 623 |
| -6,09192E | FALSE | High | IsMasterP | AAEL0112 | 2:179711 | 0 | 13 | FALSE | 4 | 5 | 4 | 479 | 533 | 671 |
| -6,66709E | FALSE | High | IsMasterP | AAEL0199 | 2:442012 | 0 | 2 | FALSE | 4 | 6 | 4 | 4866 | 5332 | 803 |
| 9,1735E+ | FALSE | High | IsMasterP | AAEL0043 | 2:387566 | 0 | 20 | FALSE | 2 | 5 | 2 | 211 | 232 | 484 |
| 7,70355E | FALSE | High | IsMasterP | AAEL0090 | 2:402351 | 0 | 13 | FALSE | 3 | 4 | 3 | 491 | 572 | 508 |
| 1,44255E | FALSE | High | IsMasterP | AAEL0222 | 2:135873 | 0 | 31 | FALSE | 3 | 6 | 3 | 84 | 94 | 941 |
| 7,33059E | FALSE | High | IsMasterP | AAEL0045 | 1:144443 | 0 | 4 | FALSE | 3 | 5 | 3 | 1195 | 1335 | 517 |
| -8,05607E | FALSE | High | IsMasterP | AAEL0091 | 2:976723 | 0 | 13 | FALSE | 2 | 5 | 2 | 267 | 300 | 679 |
| -1,1859E+ | FALSE | High | IsMasterP | AAEL0059 | 2:360919 | 0 | 22 | FALSE | 2 | 4 | 2 | 169 | 193 | 794 |
| -6,45097E | FALSE | High | IsMasterP | AAEL0196 | 2:316709 | 0 | 20 | FALSE | 4 | 6 | 4 | 207 | 219 | 1048 |
| -3,01518E | FALSE | High | IsMasterP | AAEL0061 | 3:686954 | 0 | 8 | FALSE | 3 | 5 | 3 | 553 | 625 | 680 |
| 6,31574E | FALSE | High | IsMasterP | AAEL0104 | 1:278641 | 0 | 12 | FALSE | 4 | 6 | 4 | 446 | 503 | 520 |
| 8,75813E | FALSE | High | IsMasterP | AAEL0183 | 3:383356 | 0 | 6 | FALSE | 4 | 5 | 4 | 903 | 994 | 673 |
| 4,2896E+ | FALSE | High | IsMasterP | AAEL0055 | 1:890613 | 0 | 31 | FALSE | 4 | 5 | 4 | 256 | 289 | 699 |
| -2,48497E | FALSE | High | IsMasterP | AAEL0199 | 3:168256 | 0 | 6 | FALSE | 5 | 6 | 5 | 950 | 1066 | 574 |

| | | | | | | | | | | | | | | |
|---|---|---|---|---|---|---|---|---|---|---|---|---|---|---|
| -7,60009E | FALSE | High | IsMasterP | AAEL0064 | 3:159177 | 0 | 16 | FALSE | 2 | 4 | 2 | 150 | 162 | 799 |
| -2,52921E | FALSE | High | IsMasterP | AAEL0056 | 2:131825 | 0 | 8 | FALSE | 2 | 5 | 2 | 573 | 639 | 696 |
| 8,58977E- | FALSE | High | IsMasterP | AAEL0136 | 3:195174 | 0 | 12 | FALSE | 2 | 6 | 1 | 216 | 248 | 929 |
| -9,31443E | FALSE | High | IsMasterP | AAEL0142 | 1:837845 | 0 | 21 | FALSE | 3 | 5 | 3 | 240 | 273 | 774 |
| 4,38644E- | FALSE | High | IsMasterP | AAEL0024 | 1:989432 | 0 | 7 | FALSE | 2 | 5 | 2 | 580 | 642 | 615 |
| 1,62793E- | FALSE | High | IsMasterP | AAEL0235 | 3:121700 | 0 | 13 | FALSE | 2 | 5 | 2 | 219 | 238 | 892 |
| -2,69874E | FALSE | High | IsMasterP | AAEL0040 | 3:166337 | 0 | 5 | FALSE | 2 | 4 | 2 | 644 | 720 | 665 |
| -3,97111E | FALSE | High | IsMasterP | AAEL0035 | 2:256203 | 0 | 6 | FALSE | 4 | 5 | 4 | 777 | 859 | 660 |
| 8,87531E- | FALSE | High | IsMasterP | AAEL0171 | 2:400164 | 0 | 4 | FALSE | 3 | 5 | 3 | 1134 | 1283 | 859 |
| -2,88379E | FALSE | High | IsMasterP | AAEL0103 | 2:404155 | 0 | 23 | FALSE | 3 | 5 | 3 | 200 | 230 | 1056 |
| 6,43E+17 | FALSE | High | IsMasterP | AAEL0008 | 3:334526 | 0 | 8 | FALSE | 3 | 5 | 3 | 550 | 616 | 618 |
| -4,20673E | FALSE | High | IsMasterP | AAEL0045 | 2:310441 | 0 | 9 | FALSE | 3 | 6 | 1 | 476 | 521 | 825 |
| 4,96822E- | FALSE | High | IsMasterP | AAEL0103 | 3:409598 | 0 | 5 | FALSE | 4 | 4 | 4 | 1304 | 1472 | 689 |
| 6,91316E- | FALSE | High | IsMasterP | AAEL0082 | 2:611891 | 0 | 8 | FALSE | 3 | 5 | 3 | 478 | 547 | 935 |
| 9,09436E- | FALSE | High | IsMasterP | AAEL0060 | 3:332956 | 0 | 3 | FALSE | 2 | 5 | 2 | 1032 | 1164 | 844 |
| 1,6637E+1 | FALSE | High | IsMasterP | AAEL0080 | 2:230211 | 0 | 2 | FALSE | 3 | 4 | 3 | 2327 | 2657 | 501 |
| -3,8069E+ | FALSE | High | IsMasterP | AAEL0074 | 1:216416 | 0 | 10 | FALSE | 2 | 6 | 2 | 295 | 341 | 844 |
| 2,31936E- | FALSE | High | IsMasterP | AAEL0064 | 2:438632 | 0 | 9 | FALSE | 2 | 5 | 2 | 397 | 432 | 620 |
| -1,4641E+ | FALSE | High | IsMasterP | AAEL0206 | 1:663882 | 0 | 6 | FALSE | 4 | 5 | 4 | 946 | 1058 | 683 |
| -2,54677E | FALSE | High | IsMasterP | AAEL0170 | 3:186238 | 0 | 4 | FALSE | 1 | 5 | 1 | 359 | 367 | 568 |
| -4,35255E | FALSE | High | IsMasterP | AAEL0026 | 3:257284 | 0 | 7 | FALSE | 2 | 4 | 2 | 445 | 483 | 491 |
| 2,40444E- | FALSE | High | IsMasterP | AAEL0241 | 3:313952 | 0 | 4 | FALSE | 3 | 5 | 3 | 1233 | 1386 | 559 |
| -4,33882E | FALSE | High | IsMasterP | AAEL0086 | 2:454818 | 0 | 22 | FALSE | 4 | 5 | 2 | 298 | 346 | 775 |
| -7,7747E+ | FALSE | High | IsMasterP | AAEL0054 | 1:246236 | 0 | 26 | FALSE | 3 | 5 | 3 | 195 | 214 | 1015 |
| -5,1149E+ | FALSE | High | IsMasterP | AAEL0037 | 2:196081 | 0 | 18 | FALSE | 2 | 5 | 2 | 148 | 170 | 939 |
| 7,01554E- | FALSE | High | IsMasterP | AAEL0090 | 2:269447 | 0 | 9 | FALSE | 1 | 4 | 1 | 162 | 188 | 1023 |
| 8,21804E- | FALSE | High | IsMasterP | AAEL0081 | 3:145846 | 1 | 4 | FALSE | 3 | 6 | 3 | 551 | 632 | 559 |
| 7,12261E- | FALSE | High | IsMasterP | AAEL0280 | NIGP0100 | 1 | 7 | FALSE | 3 | 6 | 3 | 456 | 518 | 601 |
| 6,18277E- | FALSE | High | IsMasterP | AAEL0148 | 2:104872 | 1 | 4 | FALSE | 2 | 7 | 2 | 490 | 567 | 756 |
| -6,19702E | FALSE | High | IsMasterP | AAEL0088 | 3:236778 | 1 | 23 | FALSE | 4 | 5 | 4 | 210 | 245 | 766 |
| 6,20847E- | FALSE | High | IsMasterP | AAEL0022 | 2:430787 | 1 | 28 | FALSE | 3 | 5 | 3 | 143 | 168 | 613 |

| | | | | | | | | | | | | | | |
|---|---|---|---|---|---|---|---|---|---|---|---|---|---|---|
| 5,2071E+1 | FALSE | High | IsMasterP | AAEL0198 | 2:111398( | 1 | 5 | FALSE | 4 | 5 | 4 | 1149 | 1284 | 680 |
| -6,92028E | FALSE | High | IsMasterP | AAEL0043 | 1:235800: | 1 | 4 | FALSE | 2 | 5 | 2 | 788 | 900 | 686 |
| -4,52594E | FALSE | High | IsMasterP | AAEL0125 | 1:285317( | 1 | 5 | FALSE | 1 | 5 | 1 | 252 | 273 | 544 |
| -6,5186E+ | FALSE | High | IsMasterP | AAEL0076 | 3:407295! | 1 | 11 | FALSE | 2 | 5 | 2 | 260 | 286 | 595 |
| -1,48327E | FALSE | High | IsMasterP | AAEL0058 | 1:201571! | 1 | 16 | FALSE | 4 | 4 | 4 | 386 | 424 | 703 |
| -3,38987E | FALSE | High | IsMasterP | AAEL0150 | 2:191816! | 1 | 11 | FALSE | 3 | 5 | 3 | 296 | 325 | 519 |
| -8,4362E+ | FALSE | High | IsMasterP | AAEL0085 | 3:198028: | 1 | 5 | FALSE | 2 | 6 | 2 | 478 | 525 | 788 |
| -7,62447E | FALSE | High | IsMasterP | AAEL0231 | 1:208615! | 1 | 7 | FALSE | 3 | 5 | 3 | 547 | 614 | 730 |
| -1,99545E | FALSE | High | IsMasterP | AAEL0094 | 3:380924: | 1 | 5 | FALSE | 2 | 5 | 2 | 573 | 631 | 787 |
| 5,11396E- | FALSE | High | IsMasterP | AAEL0034 | 3:349859! | 1 | 16 | FALSE | 4 | 6 | 4 | 288 | 331 | 944 |
| 9,30232E- | FALSE | High | IsMasterP | AAEL0122 | 2:332066! | 1 | 35 | FALSE | 2 | 4 | 2 | 103 | 115 | 592 |
| 8,35211E- | FALSE | High | IsMasterP | AAEL0104 | 1:278694! | 1 | 12 | FALSE | 3 | 5 | 3 | 393 | 431 | 777 |
| 5,16294E- | FALSE | High | IsMasterP | AAEL0125 | 2:760728: | 1 | 13 | FALSE | 4 | 6 | 4 | 492 | 552 | 519 |
| 1,13891E- | FALSE | High | IsMasterP | AAEL0105 | 2:314484: | 1 | 6 | FALSE | 2 | 5 | 2 | 401 | 456 | 674 |
| 7,29368E- | FALSE | High | IsMasterP | AAEL0036 | 3:301417- | 1 | 3 | FALSE | 1 | 5 | 1 | 623 | 691 | 583 |
| 7,4638E+1 | FALSE | High | IsMasterP | AAEL0101 | 2:224747! | 1 | 8 | FALSE | 3 | 5 | 3 | 509 | 566 | 544 |
| -7,24529E | FALSE | High | IsMasterP | AAEL0025 | 1:913207! | 1 | 5 | FALSE | 2 | 4 | 2 | 633 | 699 | 505 |
| 5,28841E- | FALSE | High | IsMasterP | AAEL0084 | 1:205387! | 1 | 5 | FALSE | 1 | 5 | 1 | 300 | 331 | 667 |
| 2,6203E+1 | FALSE | High | IsMasterP | AAEL0132 | 2:399434- | 1 | 14 | FALSE | 3 | 4 | 3 | 380 | 417 | 662 |
| 9,01527E- | FALSE | High | IsMasterP | AAEL0135 | 2:246056- | 1 | 10 | FALSE | 2 | 4 | 2 | 400 | 444 | 923 |
| -2,34181E | FALSE | High | IsMasterP | AAEL0067 | 2:847753: | 1 | 16 | FALSE | 3 | 4 | 3 | 580 | 635 | 903 |
| -7,35793E | FALSE | High | IsMasterP | AAEL0004 | 3:183762! | 1 | 1 | FALSE | 1 | 5 | 1 | 993 | 1118 | 592 |
| -8,07951E | FALSE | High | IsMasterP | AAEL0199 | 2:188545: | 1 | 3 | FALSE | 3 | 5 | 3 | 1607 | 1776 | 525 |
| -7,87968E | FALSE | High | IsMasterP | AAEL0131 | 3:294643( | 1 | 2 | FALSE | 3 | 4 | 3 | 2190 | 2391 | 696 |
| -1,70385E | FALSE | High | IsMasterP | AAEL0019 | 2:194470( | 1 | 7 | FALSE | 2 | 4 | 2 | 390 | 454 | 966 |
| -8,89483E | FALSE | High | IsMasterP | AAEL0154 | 2:133162: | 1 | 18 | FALSE | 3 | 4 | 3 | 307 | 333 | 469 |
| -9,05045E | FALSE | High | IsMasterP | AAEL0089 | 1:810528: | 1 | 6 | FALSE | 3 | 5 | 3 | 657 | 745 | 708 |
| 7,15288E- | FALSE | High | IsMasterP | AAEL0229 | 2:191755: | 1 | 8 | FALSE | 4 | 6 | 4 | 708 | 805 | 696 |
| -4,0035E+ | FALSE | High | IsMasterP | AAEL0077 | 2:470306! | 1 | 20 | FALSE | 2 | 4 | 2 | 282 | 328 | 487 |
| 8,17426E- | FALSE | High | IsMasterP | AAEL0076 | 2:250845: | 1 | 36 | FALSE | 3 | 4 | 3 | 151 | 168 | 840 |
| 3,9607E+1 | FALSE | High | IsMasterP | AAEL0170 | 2:309020: | 1 | 3 | FALSE | 3 | 5 | 3 | 1343 | 1512 | 616 |

| | | | | | | | | | | | | | | |
|---|---|---|---|---|---|---|---|---|---|---|---|---|---|---|
| -5,86915E | FALSE | High | IsMasterP | AAEL0086 | 2:2158863 | 1 | 15 | FALSE | 5 | 5 | 5 | 530 | 604 | 655 |
| -3,05541E | FALSE | High | IsMasterP | AAEL0060 | 3:3350150 | 1 | 2 | FALSE | 3 | 5 | 3 | 2192 | 2451 | 730 |
| 3,64593E+ | FALSE | High | IsMasterP | AAEL0225 | 1:1889016 | 1 | 14 | FALSE | 4 | 5 | 4 | 406 | 452 | 693 |
| -6,52587E | FALSE | High | IsMasterP | AAEL0214 | 2:6519509 | 1 | 4 | FALSE | 2 | 5 | 2 | 698 | 805 | 876 |
| -1,72901E | FALSE | High | IsMasterP | AAEL0043 | 1:2358126 | 1 | 9 | FALSE | 4 | 4 | 4 | 791 | 896 | 637 |
| 7,30058E+ | FALSE | High | IsMasterP | AAEL0078 | 3:1516570 | 1 | 5 | FALSE | 2 | 5 | 2 | 664 | 743 | 1021 |
| -5,21331E | FALSE | High | IsMasterP | AAEL0268 | 2:7897845 | 1 | 3 | FALSE | 3 | 5 | 3 | 1280 | 1335 | 989 |
| 2,93983E+ | FALSE | High | IsMasterP | AAEL0110 | 1:7407643 | 1 | 6 | FALSE | 1 | 6 | 1 | 178 | 203 | 846 |
| -7,44916E | FALSE | High | IsMasterP | AAEL0045 | 2:3097792 | 1 | 26 | FALSE | 2 | 6 | 2 | 85 | 91 | 840 |
| 8,38809E+ | FALSE | High | IsMasterP | AAEL0175 | 2:1789247 | 1 | 4 | FALSE | 2 | 4 | 2 | 1128 | 1286 | 1081 |
| 3,2567E+1 | FALSE | High | IsMasterP | AAEL0031 | 3:2698175 | 1 | 3 | FALSE | 1 | 4 | 1 | 693 | 782 | 627 |
| 3,92564E+ | FALSE | High | IsMasterP | AAEL0099 | 2:3342459 | 1 | 8 | FALSE | 1 | 3 | 1 | 359 | 408 | 655 |
| -7,10493E | FALSE | High | IsMasterP | AAEL0093 | 3:8106185 | 1 | 12 | FALSE | 3 | 5 | 3 | 426 | 482 | 715 |
| -4,31455E | FALSE | High | IsMasterP | AAEL0182 | 2:1002849 | 1 | 3 | FALSE | 3 | 6 | 3 | 1394 | 1546 | 624 |
| 8,90262E+ | FALSE | High | IsMasterP | AAEL0076 | 1:1336793 | 2 | 4 | FALSE | 2 | 4 | 2 | 1228 | 1381 | 649 |
| -5,69843E | FALSE | High | IsMasterP | AAEL0143 | 3:1109089 | 2 | 13 | FALSE | 1 | 4 | 1 | 150 | 171 | 860 |
| -8,60735E | FALSE | High | IsMasterP | AAEL0033 | 3:8078870 | 2 | 18 | FALSE | 1 | 3 | 1 | 215 | 236 | 491 |
| -5,79578E | FALSE | High | IsMasterP | AAEL0170 | 2:3930683 | 2 | 10 | FALSE | 3 | 4 | 3 | 553 | 629 | 536 |
| -9,1936E+ | FALSE | High | IsMasterP | AAEL0128 | 2:1055334 | 2 | 20 | FALSE | 2 | 4 | 2 | 291 | 320 | 963 |
| 2,83832E+ | FALSE | High | IsMasterP | AAEL0197 | 1:4739420 | 2 | 9 | FALSE | 3 | 4 | 3 | 666 | 744 | 492 |
| -6,85478E | FALSE | High | IsMasterP | AAEL0053 | 2:3703435 | 2 | 27 | FALSE | 3 | 5 | 3 | 154 | 166 | 994 |
| 7,4749E+1 | FALSE | High | IsMasterP | AAEL0032 | 1:2916576 | 2 | 9 | FALSE | 4 | 5 | 4 | 608 | 683 | 576 |
| -1,64152E | FALSE | High | IsMasterP | AAEL0085 | 3:2226065 | 2 | 1 | FALSE | 2 | 6 | 2 | 2138 | 2395 | 734 |
| 8,75767E+ | FALSE | High | IsMasterP | AAEL0066 | 3:3527725 | 2 | 52 | FALSE | 2 | 3 | 2 | 85 | 99 | 635 |
| -7,53603E | FALSE | High | IsMasterP | AAEL0076 | 3:1374740 | 2 | 3 | FALSE | 1 | 4 | 1 | 546 | 617 | 703 |
| -4,24073E | FALSE | High | IsMasterP | AAEL0152 | 2:7134288 | 2 | 10 | FALSE | 2 | 4 | 2 | 183 | 198 | 766 |
| -6,02328E | FALSE | High | IsMasterP | AAEL0234 | 2:1113282 | 2 | 15 | FALSE | 1 | 3 | 1 | 155 | 179 | 837 |
| 4,15453E+ | FALSE | High | IsMasterP | AAEL0101 | 3:1490820 | 2 | 12 | FALSE | 2 | 5 | 2 | 203 | 226 | 605 |
| -6,26109E | FALSE | High | IsMasterP | AAEL0233 | 3:3654806 | 2 | 12 | FALSE | 1 | 4 | 1 | 155 | 178 | 573 |
| -5,0968E+ | FALSE | High | IsMasterP | AAEL0249 | 2:3290034 | 2 | 42 | FALSE | 2 | 3 | 2 | 144 | 156 | 983 |
| 4,41589E+ | FALSE | High | IsMasterP | AAEL0039 | 3:2539111 | 2 | 9 | FALSE | 1 | 4 | 1 | 205 | 243 | 983 |

| | | | | | | | | | | | | |
|---|---|---|---|---|---|---|---|---|---|---|---|---|
| 1,45367E- | FALSE | High | IsMasterP AAEL0073 2:3429369 | 2 | 15 | FALSE | 2 | 3 | 2 | 297 | 343 | 859 |
| 4,54736E- | FALSE | High | IsMasterP AAEL0064 2:4394229 | 2 | 5 | FALSE | 3 | 4 | 3 | 1160 | 1314 | 664 |
| 8,99353E- | FALSE | High | IsMasterP AAEL0025 1:2264942 | 2 | 5 | FALSE | 4 | 5 | 4 | 1031 | 1203 | 884 |
| -2,95512E | FALSE | High | IsMasterP AAEL0074 3:4082487 | 2 | 1 | FALSE | 1 | 6 | 1 | 1739 | 1908 | 879 |
| 7,92473E- | FALSE | High | IsMasterP AAEL0194 3:1906152 | 2 | 5 | FALSE | 3 | 4 | 3 | 614 | 695 | 594 |
| 6,88902E- | FALSE | High | IsMasterP AAEL0068 2:2561548 | 2 | 15 | FALSE | 3 | 5 | 3 | 290 | 337 | 507 |
| -2,52411E | FALSE | High | IsMasterP AAEL0025 1:2267630 | 2 | 17 | FALSE | 1 | 4 | 1 | 150 | 166 | 388 |
| -5,26385E | FALSE | High | IsMasterP AAEL0081 3:1081808 | 2 | 9 | FALSE | 2 | 4 | 2 | 311 | 346 | 639 |
| -5,95118E | FALSE | High | IsMasterP AAEL0122 2:3842105 | 2 | 3 | FALSE | 2 | 5 | 2 | 770 | 855 | 606 |
| -3,30912E | FALSE | High | IsMasterP AAEL0274 2:2575681 | 2 | 6 | FALSE | 1 | 4 | 1 | 308 | 345 | 1178 |
| 1,85306E- | FALSE | High | IsMasterP AAEL0187 3:3136287 | 2 | 11 | FALSE | 4 | 4 | 4 | 646 | 728 | 491 |
| -4,31134E | FALSE | High | IsMasterP AAEL0145 2:2351792 | 2 | 12 | FALSE | 2 | 5 | 2 | 264 | 299 | 837 |
| 2,20678E- | FALSE | High | IsMasterP AAEL0088 2:1262644 | 2 | 30 | FALSE | 2 | 5 | 2 | 83 | 94 | 873 |
| -1,03629E | FALSE | High | IsMasterP AAEL0251 NIGP0100 | 2 | 5 | FALSE | 3 | 5 | 3 | 758 | 863 | 639 |
| -6,39727E | FALSE | High | IsMasterP AAEL0076 1:1345643 | 2 | 6 | FALSE | 3 | 5 | 3 | 594 | 681 | 875 |
| 4,53827E- | FALSE | High | IsMasterP AAEL0059 1:2577203 | 2 | 12 | FALSE | 4 | 5 | 4 | 405 | 462 | 926 |
| 5,27568E- | FALSE | High | IsMasterP AAEL0006 3:3951445 | 2 | 9 | FALSE | 1 | 5 | 1 | 150 | 171 | 483 |
| -2,79996E | FALSE | High | IsMasterP AAEL0290 2:4384704 | 2 | 6 | FALSE | 2 | 4 | 2 | 526 | 595 | 903 |
| -4,0871E+ | FALSE | High | IsMasterP AAEL0060 3:3361718 | 2 | 2 | FALSE | 3 | 5 | 3 | 2022 | 2288 | 693 |
| -6,10169E | FALSE | High | IsMasterP AAEL0041 2:1463247 | 2 | 16 | FALSE | 2 | 6 | 2 | 76 | 88 | 1115 |
| -2,2806E+ | FALSE | High | IsMasterP AAEL0096 3:2218664 | 2 | 17 | FALSE | 4 | 4 | 4 | 367 | 403 | 668 |
| 2,06949E- | FALSE | High | IsMasterP AAEL0078 3:3811035 | 1 | 8 | FALSE | 1 | 3 | 1 | 265 | 294 | 989 |
| 2,59509E- | FALSE | High | IsMasterP AAEL0110 2:2355533 | 1 | 21 | FALSE | 3 | 4 | 3 | 288 | 326 | 539 |
| 7,72294E- | FALSE | High | IsMasterP AAEL0107 3:3052921 | 1 | 11 | FALSE | 2 | 5 | 2 | 218 | 236 | 394 |
| -1,50666E | FALSE | High | IsMasterP AAEL0035 2:2562985 | 1 | 3 | FALSE | 1 | 5 | 1 | 354 | 398 | 491 |
| 5,22624E- | FALSE | High | IsMasterP AAEL0108 3:5195611 | 1 | 5 | FALSE | 2 | 5 | 2 | 548 | 629 | 489 |
| -8,35868E | FALSE | High | IsMasterP AAEL0105 2:1119635 | 1 | 4 | FALSE | 2 | 6 | 1 | 460 | 513 | 878 |
| 4,26086E- | FALSE | High | IsMasterP AAEL0065 2:5836177 | 1 | 3 | FALSE | 1 | 3 | 1 | 616 | 692 | 517 |
| 5,42457E- | FALSE | High | IsMasterP AAEL0093 3:2974829 | 1 | 3 | FALSE | 3 | 5 | 3 | 1131 | 1281 | 755 |
| 3,33204E- | FALSE | High | IsMasterP AAEL0034 3:3483911 | 1 | 45 | FALSE | 1 | 3 | 1 | 55 | 64 | 925 |
| -1,13093E | FALSE | High | IsMasterP AAEL0217 2:2147995 | 1 | 1 | FALSE | 2 | 4 | 2 | 2426 | 2758 | 733 |

| | | | | | | | | | | | | | | |
|---|---|---|---|---|---|---|---|---|---|---|---|---|---|---|
| 2,5614E+ | FALSE | High | IsMasterP | AAEL0097 | 3:319627 | 1 | 5 | FALSE | 2 | 3 | 2 | 907 | 1014 | 954 |
| -9,00357E | FALSE | High | IsMasterP | AAEL0035 | 2:256271 | 1 | 4 | FALSE | 2 | 5 | 2 | 539 | 612 | 969 |
| 2,38249E | FALSE | High | IsMasterP | AAEL0067 | 3:353440 | 1 | 5 | FALSE | 1 | 3 | 1 | 354 | 403 | 609 |
| -3,35229E | FALSE | High | IsMasterP | AAEL0012 | 2:371126 | 1 | 7 | FALSE | 2 | 3 | 2 | 723 | 806 | 848 |
| 8,40432E | FALSE | High | IsMasterP | AAEL0052 | 2:692764 | 1 | 17 | FALSE | 2 | 4 | 2 | 235 | 255 | 554 |
| -6,02719E | FALSE | High | IsMasterP | AAEL0047 | 2:288613 | 1 | 9 | FALSE | 2 | 4 | 2 | 281 | 312 | 890 |
| 6,69788E | FALSE | High | IsMasterP | AAEL0129 | 2:124948 | 1 | 4 | FALSE | 1 | 6 | 1 | 225 | 249 | 654 |
| -3,68E+18 | FALSE | High | IsMasterP | AAEL0247 | 3:108059 | 1 | 1 | FALSE | 2 | 5 | 2 | 3981 | 4563 | 671 |
| -3,62239E | FALSE | High | IsMasterP | AAEL0083 | 3:224537 | 1 | 13 | FALSE | 3 | 4 | 3 | 358 | 412 | 624 |
| 3,10961E | FALSE | High | IsMasterP | AAEL0048 | 1:288447 | 1 | 8 | FALSE | 2 | 4 | 1 | 439 | 483 | 731 |
| -1,96638E | FALSE | High | IsMasterP | AAEL0052 | 3:895715 | 1 | 8 | FALSE | 1 | 4 | 1 | 211 | 243 | 869 |
| -1,10878E | FALSE | High | IsMasterP | AAEL0138 | 2:100166 | 1 | 15 | FALSE | 2 | 5 | 2 | 217 | 248 | 731 |
| 1,25401E | FALSE | High | IsMasterP | AAEL0093 | 3:380790 | 1 | 16 | FALSE | 3 | 4 | 3 | 321 | 360 | 640 |
| -3,121E+1 | FALSE | High | IsMasterP | AAEL0125 | 2:761132 | 1 | 3 | FALSE | 2 | 4 | 2 | 768 | 857 | 928 |
| 5,93664E | FALSE | High | IsMasterP | AAEL0020 | 2:119988 | 1 | 12 | FALSE | 3 | 4 | 3 | 397 | 443 | 906 |
| 7,47073E | FALSE | High | IsMasterP | AAEL0198 | 2:123230 | 1 | 2 | FALSE | 2 | 5 | 2 | 1510 | 1712 | 555 |
| -3,29585E | FALSE | High | IsMasterP | AAEL0011 | 3:406332 | 1 | 6 | FALSE | 2 | 3 | 2 | 530 | 583 | 690 |
| 2,20502E | FALSE | High | IsMasterP | AAEL0034 | 3:349636 | 1 | 5 | FALSE | 2 | 3 | 2 | 747 | 845 | 461 |
| -4,90192E | FALSE | High | IsMasterP | AAEL0059 | 3:173980 | 1 | 25 | FALSE | 2 | 4 | 2 | 126 | 148 | 914 |
| -7,85833E | FALSE | High | IsMasterP | AAEL0044 | 3:110084 | 1 | 13 | FALSE | 3 | 4 | 3 | 334 | 383 | 980 |
| 4,55773E | FALSE | High | IsMasterP | AAEL0141 | 1:468596 | 1 | 5 | FALSE | 2 | 5 | 2 | 441 | 487 | 684 |
| 3,19048E | FALSE | High | IsMasterP | AAEL0006 | 3:395239 | 1 | 6 | FALSE | 3 | 5 | 3 | 615 | 712 | 625 |
| 7,26837E | FALSE | High | IsMasterP | AAEL0004 | 3:185754 | 1 | 26 | FALSE | 2 | 5 | 2 | 152 | 165 | 488 |
| -3,69012E | FALSE | High | IsMasterP | AAEL0027 | 2:138147 | 1 | 8 | FALSE | 2 | 4 | 2 | 378 | 437 | 872 |
| 6,12369E | FALSE | High | IsMasterP | AAEL0208 | 2:118727 | 1 | 14 | FALSE | 3 | 4 | 3 | 286 | 321 | 947 |
| 6,51203E | FALSE | High | IsMasterP | AAEL0102 | 3:279127 | 1 | 11 | FALSE | 2 | 4 | 2 | 257 | 288 | 933 |
| -2,33671E | FALSE | High | IsMasterP | AAEL0087 | 3:281327 | 1 | 10 | FALSE | 3 | 4 | 3 | 422 | 475 | 683 |
| -9,17823E | FALSE | High | IsMasterP | AAEL0034 | 3:348325 | 1 | 17 | FALSE | 2 | 4 | 2 | 173 | 201 | 785 |
| -2,51408E | FALSE | High | IsMasterP | AAEL0044 | 3:212313 | 1 | 6 | FALSE | 2 | 4 | 2 | 495 | 557 | 522 |
| -1,35504E | FALSE | High | IsMasterP | AAEL0011 | 2:309081 | 1 | 22 | FALSE | 3 | 6 | 3 | 145 | 171 | 894 |
| -2,7621E+ | FALSE | High | IsMasterP | AAEL0027 | 3:129272 | 1 | 25 | FALSE | 1 | 3 | 1 | 103 | 112 | 926 |

| | | | | | | | | | | | | | | |
|---|---|---|---|---|---|---|---|---|---|---|---|---|---|---|
| -3,73972E | FALSE | High | IsMasterP | AAEL0109 | 2:108535 | 1 | 6 | FALSE | 3 | 4 | 3 | 689 | 749 | 764 |
| 2,63465E | FALSE | High | IsMasterP | AAEL0040 | 1:135623 | 1 | 10 | FALSE | 4 | 5 | 4 | 479 | 532 | 775 |
| -8,95087E | FALSE | High | IsMasterP | AAEL0180 | 3:151484 | 2 | 2 | FALSE | 2 | 4 | 2 | 1327 | 1491 | 521 |
| 3,94346E | FALSE | High | IsMasterP | AAEL0085 | 2:157403 | 2 | 3 | FALSE | 1 | 3 | 1 | 698 | 790 | 775 |
| -5,2031E+ | FALSE | High | IsMasterP | AAEL0044 | 3:109707 | 2 | 4 | FALSE | 2 | 4 | 2 | 789 | 888 | 690 |
| 1,07595E | FALSE | High | IsMasterP | AAEL0108 | 3:519166 | 2 | 1 | FALSE | 2 | 5 | 2 | 1542 | 1740 | 890 |
| 4,60429E | FALSE | High | IsMasterP | AAEL0043 | 3:936846 | 2 | 11 | FALSE | 3 | 5 | 3 | 466 | 511 | 829 |
| -2,8661E+ | FALSE | High | IsMasterP | AAEL0087 | 3:281149 | 2 | 10 | FALSE | 3 | 4 | 3 | 416 | 480 | 517 |
| -2,40078E | FALSE | High | IsMasterP | AAEL0067 | 2:839541 | 2 | 16 | FALSE | 2 | 4 | 2 | 257 | 301 | 585 |
| 3,78211E | FALSE | High | IsMasterP | AAEL0142 | 1:838247 | 2 | 12 | FALSE | 3 | 5 | 3 | 362 | 404 | 847 |
| 4,71551E | FALSE | High | IsMasterP | AAEL0213 | 3:319958 | 2 | 33 | FALSE | 4 | 4 | 4 | 211 | 244 | 549 |
| -7,62571E | FALSE | High | IsMasterP | AAEL0052 | 3:905143 | 2 | 4 | FALSE | 1 | 6 | 1 | 241 | 271 | 907 |
| -8,59457E | FALSE | High | IsMasterP | AAEL0027 | 1:304624 | 2 | 13 | FALSE | 4 | 5 | 4 | 457 | 511 | 467 |
| -4,40929E | FALSE | High | IsMasterP | AAEL0136 | 1:235887 | 2 | 9 | FALSE | 1 | 4 | 1 | 175 | 202 | 477 |
| -4,84352E | FALSE | High | IsMasterP | AAEL0057 | 2:437284 | 2 | 5 | FALSE | 2 | 4 | 2 | 603 | 678 | 897 |
| 6,57769E | FALSE | High | IsMasterP | AAEL0057 | 1:122364 | 2 | 14 | FALSE | 1 | 3 | 1 | 176 | 201 | 566 |
| -2,62492E | FALSE | High | IsMasterP | AAEL0113 | 2:926254 | 2 | 10 | FALSE | 3 | 4 | 3 | 437 | 523 | 964 |
| 7,26609E | FALSE | High | IsMasterP | AAEL0229 | 1:148074 | 2 | 6 | FALSE | 1 | 4 | 1 | 219 | 250 | 621 |
| -6,03543E | FALSE | High | IsMasterP | AAEL0095 | 1:397961 | 2 | 3 | FALSE | 1 | 4 | 1 | 895 | 1028 | 859 |
| 6,23287E | FALSE | High | IsMasterP | AAEL0051 | 2:376751 | 2 | 4 | FALSE | 1 | 4 | 1 | 391 | 442 | 488 |
| -6,62975E | FALSE | High | IsMasterP | AAEL0039 | 3:254134 | 2 | 6 | FALSE | 2 | 5 | 1 | 383 | 447 | 784 |
| -7,03434E | FALSE | High | IsMasterP | AAEL0129 | 1:916723 | 2 | 22 | FALSE | 1 | 3 | 1 | 158 | 160 | 856 |
| 5,69253E | FALSE | High | IsMasterP | AAEL0118 | 3:113320 | 2 | 9 | FALSE | 4 | 5 | 4 | 557 | 654 | 749 |
| -1,658E+1 | FALSE | High | IsMasterP | AAEL0243 | 1:201961 | 2 | 2 | FALSE | 2 | 5 | 2 | 1509 | 1687 | 701 |
| 3,44939E | FALSE | High | IsMasterP | AAEL0024 | 3:127454 | 2 | 1 | FALSE | 1 | 4 | 1 | 1172 | 1336 | 679 |
| -1,30195E | FALSE | High | IsMasterP | AAEL0044 | 3:271102 | 2 | 17 | FALSE | 2 | 4 | 2 | 148 | 173 | 932 |
| -4,92487E | FALSE | High | IsMasterP | AAEL0112 | 1:348749 | 2 | 36 | FALSE | 3 | 4 | 3 | 184 | 210 | 665 |
| 8,08527E | FALSE | High | IsMasterP | AAEL0244 | 2:827465 | 2 | 17 | FALSE | 2 | 4 | 2 | 139 | 161 | 507 |
| 5,5594E+ | FALSE | High | IsMasterP | AAEL0202 | 2:122330 | 2 | 3 | FALSE | 1 | 3 | 1 | 539 | 597 | 574 |
| 3,57151E | FALSE | High | IsMasterP | AAEL0096 | 2:169199 | 2 | 8 | FALSE | 3 | 4 | 3 | 704 | 766 | 657 |
| -3,31816E | FALSE | High | IsMasterP | AAEL0106 | 2:136455 | 2 | 22 | FALSE | 1 | 3 | 1 | 78 | 91 | 887 |

| | | | | | | | | | | | | | | |
|---|---|---|---|---|---|---|---|---|---|---|---|---|---|---|
| 1,99668E | FALSE | High | IsMasterP | AAEL0270 | 1:598859 | 2 | 5 | FALSE | 1 | 3 | 1 | 602 | 646 | 730 |
| 2,19898E | FALSE | High | IsMasterP | AAEL0026 | 2:433736 | 2 | 12 | FALSE | 3 | 4 | 3 | 492 | 555 | 581 |
| 6,37409E | FALSE | High | IsMasterP | AAEL0004 | 3:162946 | 2 | 15 | FALSE | 2 | 3 | 2 | 213 | 247 | 832 |
| -1,40745E | FALSE | High | IsMasterP | AAEL0076 | 1:134937 | 2 | 10 | FALSE | 3 | 4 | 3 | 459 | 510 | 890 |
| -6,20215E | FALSE | High | IsMasterP | AAEL0150 | 2:104953 | 2 | 9 | FALSE | 2 | 4 | 2 | 331 | 376 | 891 |
| 1,50398E | FALSE | High | IsMasterP | AAEL0112 | 1:215234 | 2 | 3 | FALSE | 2 | 4 | 2 | 1071 | 1156 | 901 |
| -6,16575E | FALSE | High | IsMasterP | AAEL0198 | 2:865645 | 2 | 9 | FALSE | 1 | 4 | 1 | 183 | 200 | 812 |
| 4,53993E | FALSE | High | IsMasterP | AAEL0132 | 1:323449 | 2 | 28 | FALSE | 3 | 4 | 3 | 178 | 201 | 590 |
| -9,0621E+ | FALSE | High | IsMasterP | AAEL0149 | 3:315711 | 2 | 5 | FALSE | 2 | 4 | 2 | 615 | 706 | 601 |
| 2,91834E | FALSE | High | IsMasterP | AAEL0060 | 3:736754 | 2 | 4 | FALSE | 2 | 3 | 2 | 931 | 1071 | 690 |
| -5,21043E | FALSE | High | IsMasterP | AAEL0006 | 3:395088 | 2 | 12 | FALSE | 2 | 4 | 2 | 219 | 246 | 554 |
| -1,79877E | FALSE | High | IsMasterP | AAEL0010 | 1:299177 | 2 | 11 | FALSE | 1 | 3 | 1 | 212 | 235 | 500 |
| -3,21982E | FALSE | High | IsMasterP | AAEL0029 | 3:361993 | 2 | 17 | FALSE | 2 | 3 | 2 | 216 | 245 | 980 |
| -7,48759E | FALSE | High | IsMasterP | AAEL0062 | 3:628330 | 2 | 6 | FALSE | 3 | 3 | 3 | 1518 | 1642 | 925 |
| 5,16963E | FALSE | High | IsMasterP | AAEL0068 | 2:340357 | 2 | 4 | FALSE | 2 | 3 | 2 | 346 | 393 | 920 |
| 4,29937E | FALSE | High | IsMasterP | AAEL0067 | 2:848044 | 2 | 8 | FALSE | 3 | 3 | 3 | 829 | 923 | 517 |
| -4,9184E+ | FALSE | High | IsMasterP | AAEL0066 | 2:464010 | 2 | 20 | FALSE | 3 | 4 | 3 | 252 | 286 | 844 |
| 1,09583E | FALSE | High | IsMasterP | AAEL0197 | 1:200176 | 2 | 6 | FALSE | 3 | 4 | 3 | 1018 | 1114 | 681 |
| 2,32902E | FALSE | High | IsMasterP | AAEL0014 | 2:264533 | 2 | 4 | FALSE | 3 | 5 | 3 | 887 | 1019 | 580 |
| 1,41018E | FALSE | High | IsMasterP | AAEL0002 | 3:266531 | 2 | 11 | FALSE | 1 | 3 | 1 | 157 | 160 | 897 |
| -5,51674E | FALSE | High | IsMasterP | AAEL0051 | 3:346846 | 2 | 2 | FALSE | 1 | 3 | 1 | 867 | 977 | 601 |
| 1,07963E | FALSE | High | IsMasterP | AAEL0014 | 2:455898 | 2 | 13 | FALSE | 2 | 4 | 2 | 202 | 230 | 670 |
| 2,55428E | FALSE | High | IsMasterP | AAEL0049 | 3:323132 | 2 | 6 | FALSE | 1 | 3 | 1 | 272 | 306 | 552 |
| 7,94359E | FALSE | High | IsMasterP | AAEL0253 | 1:120291 | 2 | 7 | FALSE | 1 | 3 | 1 | 244 | 281 | 766 |
| 5,85338E | FALSE | High | IsMasterP | AAEL0002 | 2:754214 | 2 | 11 | FALSE | 2 | 3 | 2 | 377 | 424 | 511 |
| -6,50307E | FALSE | High | IsMasterP | AAEL0013 | 3:476941 | 2 | 16 | FALSE | 3 | 3 | 3 | 360 | 400 | 664 |
| 1,84013E | FALSE | High | IsMasterP | AAEL0214 | NIGP0100 | 2 | 4 | FALSE | 2 | 4 | 2 | 624 | 716 | 683 |
| 8,634E+1 | FALSE | High | IsMasterP | AAEL0255 | 3:551270 | 2 | 4 | FALSE | 2 | 4 | 2 | 978 | 1095 | 564 |
| 1,93319E | FALSE | High | IsMasterP | AAEL0073 | 2:233852 | 2 | 6 | FALSE | 2 | 4 | 2 | 428 | 483 | 505 |
| -5,96075E | FALSE | High | IsMasterP | AAEL0106 | 2:136023 | 2 | 5 | FALSE | 1 | 3 | 1 | 360 | 389 | 507 |
| -5,3279E+ | FALSE | High | IsMasterP | AAEL0082 | 1:440253 | 2 | 5 | FALSE | 3 | 4 | 3 | 525 | 571 | 521 |

| | | | | | | | | | | | | | | |
|---|---|---|---|---|---|---|---|---|---|---|---|---|---|---|
| -7,11377E | FALSE | High | IsMasterP | AAEL0022 | 3:373251 | 2 | 10 | FALSE | 3 | 4 | 3 | 512 | 576 | 543 |
| -3,25818E | FALSE | High | IsMasterP | AAEL0195 | 2:354607 | 2 | 3 | FALSE | 3 | 4 | 3 | 1466 | 1657 | 557 |
| 2,23725E | FALSE | High | IsMasterP | AAEL0011 | 2:307646 | 2 | 6 | FALSE | 2 | 4 | 2 | 397 | 429 | 785 |
| 8,04522E | FALSE | High | IsMasterP | AAEL0044 | 2:453279 | 2 | 7 | FALSE | 3 | 4 | 3 | 482 | 524 | 769 |
| -5,50228E | FALSE | High | IsMasterP | AAEL0182 | 2:238595 | 2 | 4 | FALSE | 1 | 2 | 1 | 709 | 808 | 587 |
| -1,1564E+ | FALSE | High | IsMasterP | AAEL0085 | 3:222560 | 2 | 10 | FALSE | 2 | 4 | 2 | 239 | 265 | 844 |
| -6,79976E | FALSE | High | IsMasterP | AAEL0214 | 2:272939 | 2 | 7 | FALSE | 3 | 4 | 3 | 710 | 782 | 736 |
| -6,7006E+ | FALSE | High | IsMasterP | AAEL0042 | 1:104562 | 2 | 4 | FALSE | 2 | 3 | 2 | 968 | 1083 | 857 |
| 1,23843E | FALSE | High | IsMasterP | AAEL0226 | 1:644475 | 2 | 5 | FALSE | 2 | 5 | 2 | 596 | 688 | 761 |
| 3,45874E | FALSE | High | IsMasterP | AAEL0138 | 3:170434 | 2 | 5 | FALSE | 2 | 3 | 2 | 718 | 768 | 746 |
| -3,39341E | FALSE | High | IsMasterP | AAEL0031 | NIGP0100 | 2 | 5 | FALSE | 2 | 4 | 2 | 538 | 597 | 841 |
| -4,36881E | FALSE | High | IsMasterP | AAEL0003 | 2:124261 | 2 | 9 | FALSE | 3 | 5 | 3 | 486 | 519 | 878 |
| -6,54941E | FALSE | High | IsMasterP | AAEL0098 | 2:191697 | 2 | 11 | FALSE | 2 | 4 | 2 | 243 | 270 | 555 |
| 8,8313E+ | FALSE | High | IsMasterP | AAEL0097 | 1:272739 | 2 | 29 | FALSE | 2 | 5 | 2 | 83 | 95 | 517 |
| -4,75837E | FALSE | High | IsMasterP | AAEL0175 | 3:664916 | 2 | 48 | FALSE | 2 | 3 | 2 | 129 | 126 | 873 |
| 1,76412E | FALSE | High | IsMasterP | AAEL0198 | 2:148704 | 2 | 1 | FALSE | 4 | 5 | 4 | 5634 | 6457 | 527 |
| -2,06079E | FALSE | High | IsMasterP | AAEL0077 | 2:470545 | 2 | 16 | FALSE | 3 | 3 | 3 | 410 | 467 | 496 |
| -7,37346E | FALSE | High | IsMasterP | AAEL0052 | 2:638065 | 2 | 3 | FALSE | 1 | 3 | 1 | 464 | 512 | 611 |
| -6,03679E | FALSE | High | IsMasterP | AAEL0250 | 2:276442 | 2 | 5 | FALSE | 3 | 4 | 3 | 667 | 740 | 514 |
| -5,58739E | FALSE | High | IsMasterP | AAEL0008 | 3:362040 | 2 | 1 | FALSE | 1 | 3 | 1 | 1464 | 1599 | 473 |
| -4,9842E+ | FALSE | High | IsMasterP | AAEL0046 | 1:158876 | 2 | 8 | FALSE | 2 | 3 | 2 | 606 | 676 | 790 |
| 5,02003E | FALSE | High | IsMasterP | AAEL0170 | 2:473366 | 2 | 2 | FALSE | 2 | 4 | 2 | 1129 | 1280 | 816 |
| -6,20786E | FALSE | High | IsMasterP | AAEL0132 | 2:349702 | 2 | 30 | FALSE | 2 | 4 | 2 | 89 | 100 | 1070 |
| 1,59467E | FALSE | High | IsMasterP | AAEL0070 | 3:170653 | 2 | 3 | FALSE | 1 | 4 | 1 | 452 | 490 | 470 |
| -5,24494E | FALSE | High | IsMasterP | AAEL0251 | 1:183191 | 2 | 4 | FALSE | 3 | 4 | 3 | 1024 | 1105 | 762 |
| 3,42517E | FALSE | High | IsMasterP | AAEL0281 | 2:650668 | 2 | 3 | FALSE | 3 | 4 | 3 | 1284 | 1454 | 585 |
| -3,26667E | FALSE | High | IsMasterP | AAEL0196 | 3:290357 | 2 | 29 | FALSE | 3 | 4 | 3 | 146 | 159 | 1017 |
| 7,3694E+ | FALSE | High | IsMasterP | AAEL0250 | 2:586258 | 2 | 2 | FALSE | 3 | 3 | 3 | 1638 | 1868 | 1096 |
| 4,03812E | FALSE | High | IsMasterP | AAEL0093 | 2:451640 | 2 | 20 | FALSE | 3 | 5 | 3 | 148 | 167 | 929 |
| 2,15779E | FALSE | High | IsMasterP | AAEL0139 | 2:352208 | 2 | 15 | FALSE | 1 | 3 | 1 | 110 | 119 | 529 |
| 3,61008E | FALSE | High | IsMasterP | AAEL0199 | 3:807460 | 2 | 3 | FALSE | 1 | 5 | 1 | 291 | 329 | 623 |

| | | | | | | | | | | | | | | |
|---|---|---|---|---|---|---|---|---|---|---|---|---|---|---|
| 5,62589E | FALSE | High | IsMasterP | AAEL0086 | 1:241340 | 2 | 5 | FALSE | 2 | 4 | 2 | 513 | 557 | 534 |
| -5,82901E | FALSE | High | IsMasterP | AAEL0096 | 3:209656 | 2 | 17 | FALSE | 2 | 3 | 2 | 260 | 283 | 686 |
| -8,23767E | FALSE | High | IsMasterP | AAEL0016 | 2:299344 | 2 | 4 | FALSE | 3 | 3 | 3 | 1347 | 1534 | 718 |
| -3,88284E | FALSE | High | IsMasterP | AAEL0035 | 1:692350 | 2 | 10 | FALSE | 3 | 4 | 3 | 575 | 626 | 639 |
| -5,13682E | FALSE | High | IsMasterP | AAEL0173 | 3:477874 | 2 | 2 | FALSE | 4 | 4 | 4 | 2599 | 2811 | 834 |
| -3,85871E | FALSE | High | IsMasterP | AAEL0149 | 2:172073 | 2 | 3 | FALSE | 2 | 4 | 2 | 750 | 841 | 670 |
| -8,775E+1 | FALSE | High | IsMasterP | AAEL0032 | 2:367236 | 2 | 12 | FALSE | 4 | 4 | 4 | 617 | 710 | 582 |
| -8,0413E+ | FALSE | High | IsMasterP | AAEL0087 | 2:198820 | 2 | 10 | FALSE | 2 | 3 | 2 | 392 | 446 | 895 |
| -7,11487E | FALSE | High | IsMasterP | AAEL0107 | 3:294196 | 2 | 8 | FALSE | 4 | 4 | 4 | 663 | 762 | 616 |
| 3,24493E | FALSE | High | IsMasterP | AAEL0040 | 1:135657 | 2 | 44 | FALSE | 3 | 4 | 3 | 124 | 139 | 677 |
| -1,10395E | FALSE | High | IsMasterP | AAEL0000 | 3:102799 | 2 | 8 | FALSE | 3 | 4 | 3 | 649 | 719 | 687 |
| -5,39569E | FALSE | High | IsMasterP | AAEL0101 | 2:357389 | 2 | 17 | FALSE | 2 | 4 | 2 | 172 | 196 | 567 |
| -4,83405E | FALSE | High | IsMasterP | AAEL0003 | 2:124395 | 2 | 3 | FALSE | 1 | 3 | 1 | 700 | 762 | 916 |
| -2,98537E | FALSE | High | IsMasterP | AAEL0080 | 2:230933 | 2 | 1 | FALSE | 3 | 5 | 3 | 3586 | 3920 | 516 |
| 3,78965E | FALSE | High | IsMasterP | AAEL0077 | 3:133472 | 2 | 9 | FALSE | 2 | 4 | 2 | 388 | 434 | 497 |
| 1,89742E | FALSE | High | IsMasterP | AAEL0023 | 2:304642 | 2 | 5 | FALSE | 2 | 4 | 2 | 531 | 612 | 644 |
| -8,59573E | FALSE | High | IsMasterP | AAEL0118 | 3:113468 | 2 | 15 | FALSE | 2 | 3 | 2 | 302 | 337 | 742 |
| -8,82217E | FALSE | High | IsMasterP | AAEL0075 | 1:302517 | 2 | 7 | FALSE | 2 | 4 | 2 | 361 | 394 | 778 |
| 3,4155E+ | FALSE | High | IsMasterP | AAEL0247 | 2:191379 | 2 | 14 | FALSE | 4 | 4 | 4 | 276 | 289 | 768 |
| -7,50333E | FALSE | High | IsMasterP | AAEL0195 | 1:206920 | 2 | 4 | FALSE | 2 | 3 | 1 | 651 | 725 | 496 |
| -5,08706E | FALSE | High | IsMasterP | AAEL0212 | 1:144836 | 2 | 25 | FALSE | 3 | 3 | 3 | 225 | 253 | 552 |
| 7,56546E | FALSE | High | IsMasterP | AAEL0040 | 3:401682 | 2 | 12 | FALSE | 3 | 4 | 3 | 352 | 405 | 711 |
| -1,23016E | FALSE | High | IsMasterP | AAEL0062 | 3:246587 | 2 | 10 | FALSE | 4 | 4 | 4 | 458 | 522 | 850 |
| -2,77084E | FALSE | High | IsMasterP | AAEL0195 | 3:297330 | 2 | 26 | FALSE | 3 | 3 | 3 | 162 | 184 | 654 |
| -3,26656E | FALSE | High | IsMasterP | AAEL0195 | 2:176442 | 2 | 7 | FALSE | 3 | 4 | 3 | 807 | 905 | 909 |
| 8,40232E | FALSE | High | IsMasterP | AAEL0051 | 3:346789 | 2 | 48 | FALSE | 2 | 4 | 2 | 82 | 95 | 963 |
| -2,26069E | FALSE | High | IsMasterP | AAEL0092 | 2:487234 | 2 | 30 | FALSE | 2 | 4 | 2 | 87 | 99 | 676 |
| -4,39838E | FALSE | High | IsMasterP | AAEL0282 | 3:125720 | 2 | 5 | FALSE | 1 | 4 | 1 | 245 | 272 | 536 |
| 5,62631E | FALSE | High | IsMasterP | AAEL0117 | 1:147407 | 2 | 20 | FALSE | 3 | 3 | 3 | 318 | 361 | 592 |
| -1,25919E | FALSE | High | IsMasterP | AAEL0023 | 1:107104 | 2 | 2 | FALSE | 2 | 4 | 2 | 1926 | 2160 | 654 |
| 6,09196E | FALSE | High | IsMasterP | AAEL0027 | 3:129267 | 2 | 14 | FALSE | 1 | 3 | 1 | 188 | 208 | 473 |

| | | | | | | | | | | | | | | |
|---|---|---|---|---|---|---|---|---|---|---|---|---|---|---|
| -6,33751E | FALSE | High | IsMasterP | AAEL0065 | 2:5932829 | 2 | 8 | FALSE | 1 | 2 | 1 | 393 | 428 | 884 |
| -2,99221E | FALSE | High | IsMasterP | AAEL0088 | 3:1332293 | 2 | 5 | FALSE | 2 | 4 | 2 | 528 | 603 | 637 |
| -4,09403E | FALSE | High | IsMasterP | AAEL0080 | 2:3655393 | 2 | 21 | FALSE | 1 | 3 | 1 | 157 | 175 | 536 |
| -5,59076E | FALSE | High | IsMasterP | AAEL0028 | 3:1293550 | 2 | 7 | FALSE | 2 | 3 | 2 | 418 | 467 | 888 |
| 1,59714E- | FALSE | High | IsMasterP | AAEL0004 | 3:1853168 | 2 | 29 | FALSE | 3 | 5 | 3 | 76 | 84 | 977 |
| -1,05147E | FALSE | High | IsMasterP | AAEL0054 | 2:2957399 | 2 | 3 | FALSE | 1 | 2 | 1 | 783 | 892 | 844 |
| 9,05758E- | FALSE | High | IsMasterP | AAEL0061 | 3:6937154 | 2 | 5 | FALSE | 1 | 3 | 1 | 417 | 486 | 818 |
| 8,42966E- | FALSE | High | IsMasterP | AAEL0273 | 1:1424759 | 2 | 5 | FALSE | 2 | 4 | 2 | 528 | 582 | 897 |
| -5,6876E+ | FALSE | High | IsMasterP | AAEL0061 | 1:1260797 | 2 | 6 | FALSE | 2 | 3 | 2 | 344 | 393 | 673 |
| -3,15105E | FALSE | High | IsMasterP | AAEL0118 | 3:3202613 | 2 | 8 | FALSE | 2 | 3 | 2 | 571 | 654 | 765 |
| 3,17972E- | FALSE | High | IsMasterP | AAEL0194 | 1:1463574 | 2 | 5 | FALSE | 4 | 4 | 4 | 1293 | 1425 | 774 |
| -5,06136E | FALSE | High | IsMasterP | AAEL0102 | 3:2790924 | 2 | 35 | FALSE | 3 | 3 | 3 | 126 | 144 | 790 |
| -5,34769E | FALSE | High | IsMasterP | AAEL0131 | 3:3711667 | 2 | 1 | FALSE | 1 | 3 | 1 | 1694 | 1909 | 717 |
| -8,33733E | FALSE | High | IsMasterP | AAEL0113 | 3:2016812 | 2 | 7 | FALSE | 2 | 3 | 2 | 394 | 451 | 933 |
| 2,43612E- | FALSE | High | IsMasterP | AAEL0084 | 1:9355659 | 2 | 9 | FALSE | 2 | 5 | 2 | 228 | 259 | 554 |
| -8,60192E | FALSE | High | IsMasterP | AAEL0091 | 2:4191416 | 3 | 2 | FALSE | 1 | 4 | 1 | 493 | 573 | 714 |
| -6,62608E | FALSE | High | IsMasterP | AAEL0129 | 2:1249231 | 3 | 2 | FALSE | 1 | 4 | 1 | 609 | 718 | 916 |
| -8,00006E | FALSE | High | IsMasterP | AAEL0037 | 2:1979054 | 3 | 14 | FALSE | 1 | 2 | 1 | 143 | 151 | 784 |
| -6,02453E | FALSE | High | IsMasterP | AAEL0141 | 3:3022719 | 3 | 6 | FALSE | 2 | 3 | 2 | 506 | 549 | 750 |
| 7,90569E- | FALSE | High | IsMasterP | AAEL0117 | 1:1480415 | 3 | 18 | FALSE | 3 | 3 | 3 | 231 | 262 | 615 |
| 9,19978E- | FALSE | High | IsMasterP | AAEL0148 | 3:1976960 | 3 | 9 | FALSE | 2 | 3 | 2 | 359 | 420 | 644 |
| 6,1365E+1 | FALSE | High | IsMasterP | AAEL0100 | 2:2001475 | 3 | 15 | FALSE | 3 | 4 | 3 | 196 | 220 | 426 |
| 1,17341E- | FALSE | High | IsMasterP | AAEL0038 | 3:8464312 | 3 | 23 | FALSE | 1 | 2 | 1 | 79 | 90 | 725 |
| 4,05152E- | FALSE | High | IsMasterP | AAEL0215 | 2:2188932 | 3 | 3 | FALSE | 2 | 3 | 2 | 718 | 827 | 764 |
| 7,86584E- | FALSE | High | IsMasterP | AAEL0084 | 1:5339746 | 3 | 7 | FALSE | 1 | 3 | 1 | 270 | 304 | 903 |
| -3,89545E | FALSE | High | IsMasterP | AAEL0047 | 2:2891776 | 3 | 4 | FALSE | 2 | 3 | 2 | 853 | 950 | 500 |
| 4,40236E- | FALSE | High | IsMasterP | AAEL0052 | 2:6912329 | 3 | 7 | FALSE | 3 | 4 | 3 | 524 | 556 | 479 |
| 2,8652E+1 | FALSE | High | IsMasterP | AAEL0015 | 3:2609116 | 3 | 36 | FALSE | 3 | 3 | 3 | 108 | 121 | 548 |
| 6,65434E- | FALSE | High | IsMasterP | AAEL0138 | 1:1192602 | 3 | 2 | FALSE | 1 | 3 | 1 | 1124 | 1223 | 759 |
| 4,02458E- | FALSE | High | IsMasterP | AAEL0265 | 1:2713553 | 3 | 4 | FALSE | 2 | 4 | 2 | 504 | 585 | 835 |
| -5,02889E | FALSE | High | IsMasterP | AAEL0100 | 3:2091741 | 3 | 8 | FALSE | 2 | 4 | 2 | 372 | 420 | 500 |

| | | | | | | | | | | | | | | |
|---|---|---|---|---|---|---|---|---|---|---|---|---|---|---|
| -5,92999E | FALSE | High | IsMasterP | AAEL0202 | 3:380837 | 3 | 24 | FALSE | 2 | 5 | 2 | 51 | 64 | 1256 |
| -2,46567E | FALSE | High | IsMasterP | AAEL0002 | 3:266478 | 3 | 11 | FALSE | 2 | 3 | 2 | 229 | 262 | 879 |
| -6,97002E | FALSE | High | IsMasterP | AAEL0029 | 3:288890 | 3 | 8 | FALSE | 2 | 3 | 2 | 380 | 427 | 586 |
| 1,01872E | FALSE | High | IsMasterP | AAEL0148 | 1:271403 | 3 | 5 | FALSE | 3 | 4 | 3 | 1013 | 1155 | 762 |
| 1,71563E | FALSE | High | IsMasterP | AAEL0125 | 2:761886 | 3 | 3 | FALSE | 3 | 4 | 3 | 1256 | 1398 | 511 |
| -3,02925E | FALSE | High | IsMasterP | AAEL0198 | 2:578666 | 3 | 1 | FALSE | 1 | 3 | 1 | 1032 | 1090 | 964 |
| -5,96692E | FALSE | High | IsMasterP | AAEL0035 | 3:329815 | 3 | 9 | FALSE | 2 | 4 | 2 | 278 | 310 | 683 |
| -5,04734E | FALSE | High | IsMasterP | AAEL0063 | 3:396822 | 3 | 15 | FALSE | 3 | 3 | 3 | 339 | 394 | 647 |
| 6,64744E | FALSE | High | IsMasterP | AAEL0051 | 2:226546 | 3 | 6 | FALSE | 1 | 3 | 1 | 186 | 213 | 906 |
| 8,67984E | FALSE | High | IsMasterP | AAEL0131 | 2:337829 | 3 | 9 | FALSE | 2 | 4 | 2 | 308 | 340 | 806 |
| 6,83182E | FALSE | High | IsMasterP | AAEL0039 | 1:250266 | 3 | 10 | FALSE | 1 | 2 | 1 | 404 | 459 | 539 |
| 2,48177E | FALSE | High | IsMasterP | AAEL0059 | 1:257447 | 3 | 13 | FALSE | 2 | 4 | 2 | 168 | 194 | 879 |
| 7,03824E | FALSE | High | IsMasterP | AAEL0115 | 3:106814 | 3 | 9 | FALSE | 3 | 3 | 3 | 522 | 599 | 1112 |
| 4,03756E | FALSE | High | IsMasterP | AAEL0013 | 1:845836 | 3 | 4 | FALSE | 1 | 4 | 1 | 231 | 239 | 489 |
| 4,471E+1 | FALSE | High | IsMasterP | AAEL0110 | 1:737685 | 3 | 5 | FALSE | 1 | 3 | 1 | 305 | 342 | 620 |
| 5,49482E | FALSE | High | IsMasterP | AAEL0218 | 2:367463 | 3 | 4 | FALSE | 1 | 4 | 1 | 234 | 254 | 649 |
| 5,53556E | FALSE | High | IsMasterP | AAEL0043 | 3:509442 | 3 | 8 | FALSE | 1 | 2 | 1 | 318 | 369 | 916 |
| 8,75168E | FALSE | High | IsMasterP | AAEL0228 | 1:102068 | 3 | 5 | FALSE | 2 | 4 | 2 | 562 | 640 | 517 |
| -6,65489E | FALSE | High | IsMasterP | AAEL0139 | 2:651063 | 3 | 9 | FALSE | 2 | 3 | 2 | 292 | 336 | 643 |
| -4,88189E | FALSE | High | IsMasterP | AAEL0266 | 3:948618 | 3 | 26 | FALSE | 2 | 3 | 2 | 134 | 158 | 952 |
| 2,48583E | FALSE | High | IsMasterP | AAEL0068 | 3:369643 | 3 | 3 | FALSE | 3 | 4 | 3 | 1021 | 1132 | 793 |
| -3,3669E+ | FALSE | High | IsMasterP | AAEL0050 | 2:444324 | 3 | 5 | FALSE | 3 | 4 | 3 | 826 | 938 | 862 |
| 2,98129E | FALSE | High | IsMasterP | AAEL0086 | 3:172443 | 3 | 6 | FALSE | 1 | 3 | 1 | 367 | 389 | 488 |
| -2,61443E | FALSE | High | IsMasterP | AAEL0125 | 2:328019 | 3 | 27 | FALSE | 4 | 4 | 4 | 135 | 156 | 960 |
| -9,3119E+ | FALSE | High | IsMasterP | AAEL0181 | 2:595404 | 3 | 15 | FALSE | 3 | 4 | 3 | 324 | 368 | 615 |
| 4,74873E | FALSE | High | IsMasterP | AAEL0114 | 2:225131 | 3 | 13 | FALSE | 2 | 4 | 2 | 376 | 422 | 687 |
| 1,56504E | FALSE | High | IsMasterP | AAEL0068 | 1:528465 | 3 | 14 | FALSE | 2 | 3 | 2 | 217 | 240 | 642 |
| -1,48108E | FALSE | High | IsMasterP | AAEL0148 | 1:271328 | 3 | 7 | FALSE | 2 | 3 | 2 | 507 | 579 | 847 |
| 2,4331E+ | FALSE | High | IsMasterP | AAEL0069 | 1:157351 | 3 | 2 | FALSE | 2 | 3 | 2 | 1780 | 2049 | 512 |
| 1,17577E | FALSE | High | IsMasterP | AAEL0054 | 3:383797 | 3 | 2 | FALSE | 1 | 3 | 1 | 1036 | 1163 | 910 |
| -3,257E+1 | FALSE | High | IsMasterP | AAEL0031 | 1:275790 | 3 | 5 | FALSE | 2 | 3 | 2 | 705 | 778 | 689 |

| | | | | | | | | | | | | | | |
|---|---|---|---|---|---|---|---|---|---|---|---|---|---|---|
| -2,21912E | FALSE | High | IsMasterP | AAEL0116 | 3:2190105 | 3 | 14 | FALSE | 1 | 2 | 1 | 158 | 180 | 910 |
| -7,25904E | FALSE | High | IsMasterP | AAEL0274 | 3:1022590 | 3 | 4 | FALSE | 2 | 4 | 2 | 614 | 705 | 891 |
| 5,55567E+ | FALSE | High | IsMasterP | AAEL0146 | 1:2615621 | 3 | 22 | FALSE | 2 | 3 | 2 | 140 | 148 | 805 |
| 1,52986E+ | FALSE | High | IsMasterP | AAEL0252 | 1:2540038 | 3 | 6 | FALSE | 2 | 3 | 2 | 826 | 908 | 862 |
| -1,72577E | FALSE | High | IsMasterP | AAEL0173 | 2:8622879 | 3 | 32 | FALSE | 1 | 3 | 1 | 101 | 99 | 848 |
| -8,50146E | FALSE | High | IsMasterP | AAEL0133 | 2:1357212 | 3 | 12 | FALSE | 2 | 4 | 1 | 195 | 218 | 623 |
| -7,35641E | FALSE | High | IsMasterP | AAEL0032 | 2:3672475 | 3 | 5 | FALSE | 1 | 3 | 1 | 249 | 274 | 649 |
| 7,77381E+ | FALSE | High | IsMasterP | AAEL0048 | 2:7133797 | 3 | 25 | FALSE | 2 | 2 | 2 | 173 | 201 | 573 |
| -9,14327E | FALSE | High | IsMasterP | AAEL0248 | 2:8636632 | 3 | 54 | FALSE | 2 | 3 | 1 | 127 | 127 | 939 |
| -5,40563E | FALSE | High | IsMasterP | AAEL0155 | 1:2187256 | 3 | 6 | FALSE | 2 | 4 | 2 | 578 | 644 | 783 |
| 8,08573E+ | FALSE | High | IsMasterP | AAEL0038 | 1:2031950 | 3 | 10 | FALSE | 4 | 4 | 4 | 459 | 514 | 980 |
| -8,95957E | FALSE | High | IsMasterP | AAEL0121 | 1:1423125 | 3 | 8 | FALSE | 2 | 4 | 2 | 321 | 365 | 522 |
| -5,85326E | FALSE | High | IsMasterP | AAEL0049 | 3:3231432 | 3 | 2 | FALSE | 3 | 4 | 3 | 1639 | 1837 | 721 |
| -1,4391E+ | FALSE | High | IsMasterP | AAEL0076 | 2:2510523 | 3 | 3 | FALSE | 2 | 3 | 2 | 1300 | 1456 | 765 |
| 3,37587E+ | FALSE | High | IsMasterP | AAEL0066 | 3:1481537 | 3 | 15 | FALSE | 1 | 3 | 1 | 111 | 125 | 580 |
| -6,57083E | FALSE | High | IsMasterP | AAEL0040 | 1:1354346 | 3 | 12 | FALSE | 2 | 2 | 2 | 481 | 534 | 701 |
| -1,4574E+ | FALSE | High | IsMasterP | AAEL0120 | 3:1603653 | 3 | 3 | FALSE | 1 | 3 | 1 | 385 | 420 | 501 |
| 9,81695E+ | FALSE | High | IsMasterP | AAEL0268 | NIGP0100 | 3 | 7 | FALSE | 2 | 4 | 2 | 179 | 205 | 744 |
| -2,8374E+ | FALSE | High | IsMasterP | AAEL0222 | 1:1870912 | 3 | 4 | FALSE | 2 | 3 | 2 | 1029 | 1173 | 664 |
| 3,26565E+ | FALSE | High | IsMasterP | AAEL0172 | 2:2488928 | 3 | 36 | FALSE | 2 | 4 | 2 | 64 | 70 | 999 |
| 9,1369E+1 | FALSE | High | IsMasterP | AAEL0096 | 1:6061010 | 3 | 8 | FALSE | 2 | 3 | 2 | 512 | 583 | 935 |
| 2,37528E+ | FALSE | High | IsMasterP | AAEL0197 | 2:8332201 | 3 | 2 | FALSE | 2 | 4 | 2 | 1574 | 1741 | 925 |
| -3,96028E | FALSE | High | IsMasterP | AAEL0203 | 1:3544977 | 3 | 4 | FALSE | 2 | 3 | 2 | 787 | 878 | 564 |
| 1,98373E+ | FALSE | High | IsMasterP | AAEL0138 | 1:7190185 | 3 | 4 | FALSE | 3 | 4 | 3 | 1184 | 1331 | 486 |
| 1,90741E+ | FALSE | High | IsMasterP | AAEL0104 | 1:2684366 | 3 | 8 | FALSE | 2 | 3 | 2 | 319 | 356 | 569 |
| 7,78273E+ | FALSE | High | IsMasterP | AAEL0254 | 3:1476461 | 3 | 5 | FALSE | 1 | 3 | 1 | 282 | 312 | 623 |
| -4,47186E | FALSE | High | IsMasterP | AAEL0086 | 2:2448850 | 3 | 7 | FALSE | 1 | 2 | 1 | 518 | 583 | 658 |
| -6,98023E | FALSE | High | IsMasterP | AAEL0120 | 1:2703905 | 3 | 5 | FALSE | 3 | 4 | 3 | 665 | 736 | 788 |
| 2,20959E+ | FALSE | High | IsMasterP | AAEL0123 | 2:1116004 | 3 | 4 | FALSE | 2 | 4 | 2 | 626 | 690 | 635 |
| 8,34118E+ | FALSE | High | IsMasterP | AAEL0128 | 2:1055982 | 3 | 25 | FALSE | 3 | 3 | 3 | 188 | 210 | 951 |
| 7,93618E+ | FALSE | High | IsMasterP | AAEL0097 | 3:1556212 | 3 | 17 | FALSE | 1 | 2 | 1 | 136 | 155 | 728 |

| | | | | | | | | | | | | | | |
|---|---|---|---|---|---|---|---|---|---|---|---|---|---|---|
| 6,87658E- | FALSE | High | IsMasterP | AAEL0066 | 2:462757 | 3 | 3 | FALSE | 2 | 3 | 2 | 1066 | 1198 | 882 |
| -2,46671E | FALSE | High | IsMasterP | AAEL0083 | 3:224914 | 3 | 1 | FALSE | 2 | 4 | 2 | 2031 | 2337 | 557 |
| 3,65723E- | FALSE | High | IsMasterP | AAEL0109 | 3:168205 | 3 | 6 | FALSE | 1 | 4 | 1 | 220 | 259 | 797 |
| 7,02404E- | FALSE | High | IsMasterP | AAEL0172 | 2:349730 | 3 | 3 | FALSE | 1 | 3 | 1 | 666 | 756 | 772 |
| 5,96831E- | FALSE | High | IsMasterP | AAEL0030 | 1:130714 | 3 | 7 | FALSE | 3 | 4 | 3 | 557 | 626 | 501 |
| -8,77794E | FALSE | High | IsMasterP | AAEL0100 | 3:209077 | 3 | 2 | FALSE | 1 | 2 | 1 | 1193 | 1334 | 553 |
| 6,76284E- | FALSE | High | IsMasterP | AAEL0197 | 2:429783 | 3 | 1 | FALSE | 1 | 2 | 1 | 5536 | 6031 | 609 |
| 8,83999E- | FALSE | High | IsMasterP | AAEL0034 | 2:153672 | 3 | 2 | FALSE | 1 | 2 | 1 | 1187 | 1295 | 715 |
| 1,55925E- | FALSE | High | IsMasterP | AAEL0116 | 3:281604 | 3 | 6 | FALSE | 3 | 3 | 3 | 686 | 780 | 630 |
| 6,26402E- | FALSE | High | IsMasterP | AAEL0123 | 1:303341 | 3 | 6 | FALSE | 3 | 4 | 3 | 547 | 606 | 660 |
| -6,63075E | FALSE | High | IsMasterP | AAEL0077 | 3:255561 | 3 | 14 | FALSE | 2 | 3 | 2 | 232 | 264 | 477 |
| -6,97405E | FALSE | High | IsMasterP | AAEL0122 | 3:534181 | 3 | 7 | FALSE | 2 | 3 | 2 | 1062 | 1188 | 524 |
| -3,86708E | FALSE | High | IsMasterP | AAEL0014 | 2:262698 | 3 | 7 | FALSE | 2 | 3 | 2 | 614 | 687 | 552 |
| -4,69507E | FALSE | High | IsMasterP | AAEL0252 | NIGP0100 | 3 | 7 | FALSE | 2 | 3 | 2 | 605 | 688 | 572 |
| -7,16729E | FALSE | High | IsMasterP | AAEL0140 | 2:159692 | 3 | 29 | FALSE | 2 | 3 | 2 | 87 | 100 | 740 |
| -7,07249E | FALSE | High | IsMasterP | AAEL0067 | 3:240765 | 3 | 7 | FALSE | 2 | 4 | 2 | 375 | 414 | 829 |
| 9,02358E- | FALSE | High | IsMasterP | AAEL0029 | 1:108242 | 3 | 8 | FALSE | 2 | 3 | 2 | 419 | 483 | 572 |
| -1,36761E | FALSE | High | IsMasterP | AAEL0139 | 3:106583 | 3 | 4 | FALSE | 2 | 3 | 2 | 1495 | 1669 | 571 |
| 4,3886E+ | FALSE | High | IsMasterP | AAEL0117 | 3:288827 | 3 | 8 | FALSE | 2 | 3 | 2 | 345 | 398 | 463 |
| 1,44969E- | FALSE | High | IsMasterP | AAEL0074 | 2:126364 | 3 | 5 | FALSE | 1 | 4 | 1 | 213 | 238 | 459 |
| -7,53823E | FALSE | High | IsMasterP | AAEL0114 | 3:328905 | 3 | 4 | FALSE | 1 | 3 | 1 | 410 | 448 | 681 |
| 4,84004E- | FALSE | High | IsMasterP | AAEL0137 | 1:220037 | 3 | 4 | FALSE | 3 | 3 | 3 | 1426 | 1603 | 559 |
| -7,05784E | FALSE | High | IsMasterP | AAEL0197 | 3:339610 | 3 | 3 | FALSE | 1 | 2 | 1 | 993 | 1111 | 696 |
| -1,68892E | FALSE | High | IsMasterP | AAEL0230 | 3:218469 | 3 | 3 | FALSE | 1 | 3 | 1 | 499 | 588 | 567 |
| 9,82868E- | FALSE | High | IsMasterP | AAEL0130 | 2:337809 | 3 | 2 | FALSE | 1 | 3 | 1 | 947 | 1044 | 736 |
| -5,08555E | FALSE | High | IsMasterP | AAEL0085 | 2:156527 | 3 | 3 | FALSE | 1 | 2 | 1 | 574 | 652 | 562 |
| -1,75683E | FALSE | High | IsMasterP | AAEL0005 | 3:164032 | 3 | 5 | FALSE | 2 | 3 | 2 | 441 | 516 | 633 |
| -5,53871E | FALSE | High | IsMasterP | AAEL0254 | 1:140278 | 3 | 3 | FALSE | 1 | 3 | 1 | 1053 | 1180 | 904 |
| 7,77602E- | FALSE | High | IsMasterP | AAEL0068 | 3:411527 | 3 | 6 | FALSE | 1 | 3 | 1 | 263 | 292 | 848 |
| 8,28978E- | FALSE | High | IsMasterP | AAEL0123 | 3:963036 | 3 | 7 | FALSE | 2 | 3 | 2 | 607 | 668 | 497 |
| -7,69215E | FALSE | High | IsMasterP | AAEL0094 | 2:246653 | 3 | 2 | FALSE | 1 | 4 | 1 | 606 | 683 | 554 |

| | | | | | | | | | | | | | |
|---|---|---|---|---|---|---|---|---|---|---|---|---|---|
| 1,632E+18 | FALSE | High | IsMasterP AAEL0038 | 3:846283 | 3 | 11 | FALSE | 3 | 3 | 3 | 435 | 479 | 484 |
| -7,81244E | FALSE | High | IsMasterP AAEL0048 | 2:711010 | 3 | 3 | FALSE | 1 | 3 | 1 | 1008 | 1132 | 634 |
| -8,11596E | FALSE | High | IsMasterP AAEL0194 | 3:100163 | 3 | 6 | FALSE | 1 | 3 | 1 | 426 | 476 | 822 |
| -8,62209E | FALSE | High | IsMasterP AAEL0239 | 1:566754 | 3 | 6 | FALSE | 2 | 3 | 2 | 557 | 623 | 749 |
| -2,07539E | FALSE | High | IsMasterP AAEL0088 | 3:132671 | 3 | 15 | FALSE | 2 | 3 | 2 | 210 | 250 | 695 |
| 1,1404E+1 | FALSE | High | IsMasterP AAEL0248 | 1:295629 | 3 | 2 | FALSE | 1 | 3 | 1 | 564 | 617 | 627 |
| 3,25551E- | FALSE | High | IsMasterP AAEL0019 | 2:195532 | 3 | 15 | FALSE | 2 | 4 | 2 | 188 | 210 | 947 |
| -4,79311E | FALSE | High | IsMasterP AAEL0053 | 2:271105 | 3 | 7 | FALSE | 3 | 3 | 3 | 754 | 839 | 720 |
| 1,97274E- | FALSE | High | IsMasterP AAEL0216 | 1:255969 | 3 | 5 | FALSE | 1 | 3 | 1 | 302 | 333 | 904 |
| 7,44051E- | FALSE | High | IsMasterP AAEL0033 | 2:461630 | 3 | 4 | FALSE | 1 | 2 | 1 | 382 | 429 | 436 |
| 4,33807E- | FALSE | High | IsMasterP AAEL0120 | 2:186685 | 3 | 9 | FALSE | 2 | 3 | 2 | 336 | 394 | 936 |
| -4,06348E | FALSE | High | IsMasterP AAEL0048 | 1:288298 | 3 | 3 | FALSE | 2 | 4 | 2 | 763 | 828 | 743 |
| 6,72248E- | FALSE | High | IsMasterP AAEL0279 | NIGP0100 | 3 | 11 | FALSE | 2 | 3 | 2 | 270 | 310 | 903 |
| -3,01522E | FALSE | High | IsMasterP AAEL0007 | 2:556780 | 3 | 2 | FALSE | 2 | 4 | 2 | 983 | 1135 | 615 |
| -4,55E+18 | FALSE | High | IsMasterP AAEL0108 | 2:383581 | 3 | 8 | FALSE | 2 | 3 | 2 | 306 | 340 | 677 |
| -8,35431E | FALSE | High | IsMasterP AAEL0051 | 2:339180 | 3 | 17 | FALSE | 3 | 3 | 3 | 289 | 332 | 900 |
| 2,94953E- | FALSE | High | IsMasterP AAEL0085 | 2:157255 | 3 | 4 | FALSE | 2 | 3 | 2 | 766 | 867 | 615 |
| 4,67139E- | FALSE | High | IsMasterP AAEL0022 | 3:375983 | 3 | 8 | FALSE | 2 | 3 | 2 | 393 | 437 | 507 |
| 2,52526E- | FALSE | High | IsMasterP AAEL0092 | 2:170758 | 3 | 4 | FALSE | 3 | 3 | 3 | 1152 | 1285 | 885 |
| 7,94172E- | FALSE | High | IsMasterP AAEL0100 | 2:200494 | 3 | 9 | FALSE | 1 | 3 | 1 | 178 | 202 | 526 |
| 1,98521E- | FALSE | High | IsMasterP AAEL0040 | 3:402265 | 3 | 2 | FALSE | 2 | 3 | 2 | 1686 | 1865 | 651 |
| -6,82822E | FALSE | High | IsMasterP AAEL0130 | 2:335656 | 3 | 8 | FALSE | 1 | 2 | 1 | 335 | 378 | 520 |
| -6,81348E | FALSE | High | IsMasterP AAEL0078 | 1:634027 | 3 | 2 | FALSE | 1 | 3 | 1 | 648 | 720 | 547 |
| -4,38296E | FALSE | High | IsMasterP AAEL0196 | 2:240837 | 3 | 4 | FALSE | 1 | 2 | 1 | 584 | 673 | 891 |
| 8,77361E- | FALSE | High | IsMasterP AAEL0020 | 2:426309 | 3 | 4 | FALSE | 2 | 4 | 2 | 470 | 525 | 517 |
| 3,97917E- | FALSE | High | IsMasterP AAEL0081 | 3:945539 | 3 | 39 | FALSE | 2 | 2 | 2 | 122 | 139 | 750 |
| -6,82931E | FALSE | High | IsMasterP AAEL0137 | 2:122552 | 3 | 7 | FALSE | 1 | 4 | 1 | 134 | 151 | 985 |
| 6,8145E+1 | FALSE | High | IsMasterP AAEL0063 | 3:396578 | 3 | 6 | FALSE | 1 | 2 | 1 | 339 | 396 | 926 |
| 2,03872E- | FALSE | High | IsMasterP AAEL0039 | 1:250507 | 3 | 22 | FALSE | 3 | 4 | 3 | 154 | 178 | 692 |
| -2,25435E | FALSE | High | IsMasterP AAEL0029 | 1:859191 | 3 | 10 | FALSE | 2 | 3 | 2 | 194 | 220 | 679 |
| -5,1349E+ | FALSE | High | IsMasterP AAEL0108 | 3:319173 | 3 | 1 | FALSE | 1 | 3 | 1 | 2015 | 2198 | 611 |

| | | | | | | | | | | | | | | |
|---|---|---|---|---|---|---|---|---|---|---|---|---|---|---|
| 7,68901E- | FALSE | High | IsMasterP | AAEL0098 | 3:122211( | 3 | 10 | FALSE | 2 | 3 | 2 | 317 | 345 | 529 |
| 1,50248E- | FALSE | High | IsMasterP | AAEL0094 | 3:380649: | 3 | 4 | FALSE | 3 | 3 | 3 | 1008 | 1168 | 863 |
| 4,75477E- | FALSE | High | IsMasterP | AAEL0110 | 3:287769! | 3 | 7 | FALSE | 2 | 3 | 2 | 321 | 364 | 869 |
| -5,27281E | FALSE | High | IsMasterP | AAEL0045 | 2:338750: | 3 | 2 | FALSE | 1 | 3 | 1 | 487 | 552 | 576 |
| -4,13159E | FALSE | High | IsMasterP | AAEL0173 | 2:329056! | 3 | 7 | FALSE | 2 | 3 | 2 | 407 | 451 | 467 |
| 5,84914E- | FALSE | High | IsMasterP | AAEL0052 | 3:308480: | 3 | 15 | FALSE | 1 | 2 | 1 | 143 | 162 | 888 |
| -1,69797E | FALSE | High | IsMasterP | AAEL0028 | 1:255671( | 3 | 10 | FALSE | 2 | 3 | 2 | 257 | 296 | 982 |
| 4,47537E- | FALSE | High | IsMasterP | AAEL0004 | 3:185351: | 3 | 8 | FALSE | 2 | 2 | 2 | 836 | 905 | 510 |
| 3,73325E- | FALSE | High | IsMasterP | AAEL0267 | 2:183044! | 3 | 3 | FALSE | 2 | 3 | 2 | 1289 | 1416 | 698 |
| 2,94813E- | FALSE | High | IsMasterP | AAEL0020 | 2:423992: | 3 | 9 | FALSE | 3 | 3 | 3 | 522 | 595 | 856 |
| -1,99799E | FALSE | High | IsMasterP | AAEL0131 | 2:374026: | 3 | 18 | FALSE | 2 | 2 | 2 | 313 | 347 | 609 |
| -6,1878E+ | FALSE | High | IsMasterP | AAEL0004 | 3:186858( | 3 | 11 | FALSE | 3 | 3 | 3 | 393 | 454 | 642 |
| 3,38669E- | FALSE | High | IsMasterP | AAEL0264 | 2:330270( | 3 | 30 | FALSE | 2 | 2 | 2 | 187 | 191 | 952 |
| 4,64883E- | FALSE | High | IsMasterP | AAEL0142 | 3:295110: | 3 | 10 | FALSE | 3 | 3 | 3 | 443 | 509 | 531 |
| 3,52146E- | FALSE | High | IsMasterP | AAEL0095 | NIGP0100 | 3 | 8 | FALSE | 2 | 3 | 2 | 317 | 348 | 585 |
| -1,31748E | FALSE | High | IsMasterP | AAEL0204 | 1:307404. | 3 | 11 | FALSE | 2 | 3 | 2 | 264 | 301 | 829 |
| -8,5611E+ | FALSE | High | IsMasterP | AAEL0079 | 2:430934: | 3 | 4 | FALSE | 2 | 3 | 2 | 846 | 937 | 664 |
| 5,79745E- | FALSE | High | IsMasterP | AAEL0080 | 2:365384! | 3 | 6 | FALSE | 2 | 3 | 2 | 548 | 576 | 515 |
| 4,63017E- | FALSE | High | IsMasterP | AAEL0105 | 2:112126: | 3 | 4 | FALSE | 1 | 3 | 1 | 400 | 441 | 536 |
| 8,57896E- | FALSE | High | IsMasterP | AAEL0083 | 3:413897! | 3 | 35 | FALSE | 2 | 2 | 2 | 182 | 196 | 592 |
| 8,01265E- | FALSE | High | IsMasterP | AAEL0227 | 1:454910: | 3 | 16 | FALSE | 1 | 2 | 1 | 184 | 210 | 486 |
| 4,4075E+1 | FALSE | High | IsMasterP | AAEL0066 | 2:462714( | 4 | 7 | FALSE | 1 | 3 | 1 | 200 | 218 | 483 |
| 5,47405E- | FALSE | High | IsMasterP | AAEL0139 | 3:230482! | 4 | 21 | FALSE | 2 | 4 | 2 | 99 | 103 | 1165 |
| 8,75876E- | FALSE | High | IsMasterP | AAEL0139 | 3:230653( | 4 | 1 | FALSE | 1 | 3 | 1 | 1029 | 1168 | 821 |
| -4,59927E | FALSE | High | IsMasterP | AAEL0154 | 2:355583( | 4 | 16 | FALSE | 2 | 4 | 2 | 146 | 163 | 862 |
| 8,55762E- | FALSE | High | IsMasterP | AAEL0233 | 1:790332: | 4 | 6 | FALSE | 2 | 4 | 2 | 470 | 519 | 511 |
| 5,97539E- | FALSE | High | IsMasterP | AAEL0277 | 3:371108! | 4 | 2 | FALSE | 3 | 4 | 3 | 1983 | 2259 | 535 |
| 4,32426E- | FALSE | High | IsMasterP | AAEL0084 | 2:190155( | 4 | 11 | FALSE | 2 | 2 | 2 | 308 | 347 | 522 |
| 6,96272E- | FALSE | High | IsMasterP | AAEL0112 | 2:184493: | 4 | 5 | FALSE | 1 | 3 | 1 | 248 | 277 | 674 |
| -7,91125E | FALSE | High | IsMasterP | AAEL0078 | 2:271832: | 4 | 2 | FALSE | 1 | 2 | 1 | 1373 | 1478 | 634 |
| 6,51536E- | FALSE | High | IsMasterP | AAEL0195 | 1:700599: | 4 | 1 | FALSE | 3 | 3 | 3 | 3324 | 3678 | 560 |

| | | | | | | | | | | | | | | |
|---|---|---|---|---|---|---|---|---|---|---|---|---|---|---|
| -3,71882E | FALSE | High | IsMasterP | AAEL0037 | 2:317058 | 4 | 12 | FALSE | 1 | 2 | 1 | 178 | 212 | 919 |
| 5,70549E | FALSE | High | IsMasterP | AAEL0087 | 3:281231 | 4 | 5 | FALSE | 2 | 3 | 2 | 602 | 685 | 563 |
| -3,81017E | FALSE | High | IsMasterP | AAEL0057 | 2:293960 | 4 | 4 | FALSE | 1 | 3 | 1 | 460 | 512 | 695 |
| -5,91406E | FALSE | High | IsMasterP | AAEL0000 | 1:243009 | 4 | 16 | FALSE | 2 | 2 | 2 | 426 | 489 | 677 |
| -5,58054E | FALSE | High | IsMasterP | AAEL0100 | 1:159421 | 4 | 11 | FALSE | 4 | 4 | 4 | 345 | 390 | 489 |
| 6,56753E | FALSE | High | IsMasterP | AAEL0009 | 3:380214 | 4 | 5 | FALSE | 2 | 3 | 2 | 487 | 525 | 739 |
| -6,00431E | FALSE | High | IsMasterP | AAEL0180 | 3:322613 | 4 | 2 | FALSE | 3 | 3 | 3 | 1987 | 2263 | 596 |
| -2,5238E+ | FALSE | High | IsMasterP | AAEL0172 | 3:136823 | 4 | 1 | FALSE | 1 | 3 | 1 | 2577 | 2746 | 711 |
| -5,25939E | FALSE | High | IsMasterP | AAEL0088 | 2:241343 | 4 | 7 | FALSE | 1 | 3 | 1 | 177 | 204 | 816 |
| -2,72247E | FALSE | High | IsMasterP | AAEL0145 | 1:602270 | 4 | 12 | FALSE | 1 | 2 | 1 | 130 | 151 | 520 |
| 2,85613E | FALSE | High | IsMasterP | AAEL0036 | 1:692456 | 4 | 4 | FALSE | 3 | 3 | 3 | 1326 | 1483 | 681 |
| -3,69387E | FALSE | High | IsMasterP | AAEL0030 | 2:213526 | 4 | 4 | FALSE | 2 | 3 | 2 | 524 | 606 | 806 |
| 5,85736E | FALSE | High | IsMasterP | AAEL0117 | 3:288749 | 4 | 21 | FALSE | 3 | 3 | 3 | 173 | 199 | 554 |
| -6,87933E | FALSE | High | IsMasterP | AAEL0075 | 3:100197 | 4 | 3 | FALSE | 3 | 3 | 3 | 1135 | 1229 | 668 |
| -2,91956E | FALSE | High | IsMasterP | AAEL0148 | 2:103740 | 4 | 11 | FALSE | 2 | 3 | 2 | 191 | 212 | 769 |
| -3,26355E | FALSE | High | IsMasterP | AAEL0017 | 2:443224 | 4 | 6 | FALSE | 3 | 3 | 3 | 769 | 867 | 545 |
| 6,19748E | FALSE | High | IsMasterP | AAEL0047 | 2:378655 | 4 | 12 | FALSE | 2 | 2 | 2 | 496 | 553 | 922 |
| 8,23903E | FALSE | High | IsMasterP | AAEL0131 | 1:142897 | 4 | 7 | FALSE | 1 | 2 | 1 | 286 | 323 | 648 |
| -6,66824E | FALSE | High | IsMasterP | AAEL0126 | 2:990526 | 4 | 11 | FALSE | 2 | 3 | 2 | 334 | 369 | 819 |
| -4,57303E | FALSE | High | IsMasterP | AAEL0046 | 3:247001 | 4 | 6 | FALSE | 1 | 3 | 1 | 204 | 235 | 686 |
| -2,72698E | FALSE | High | IsMasterP | AAEL0086 | 2:321144 | 4 | 8 | FALSE | 2 | 2 | 2 | 823 | 870 | 812 |
| 5,53166E | FALSE | High | IsMasterP | AAEL0276 | 2:341908 | 4 | 1 | FALSE | 1 | 3 | 1 | 940 | 1060 | 489 |
| -7,20267E | FALSE | High | IsMasterP | AAEL0092 | 2:112912 | 4 | 10 | FALSE | 2 | 3 | 2 | 282 | 309 | 942 |
| -4,9549E+ | FALSE | High | IsMasterP | AAEL0126 | 1:279308 | 4 | 16 | FALSE | 2 | 3 | 2 | 257 | 289 | 676 |
| -4,90671E | FALSE | High | IsMasterP | AAEL0088 | 3:133363 | 4 | 19 | FALSE | 2 | 3 | 2 | 189 | 212 | 655 |
| -3,52829E | FALSE | High | IsMasterP | AAEL0058 | 1:201206 | 4 | 6 | FALSE | 2 | 2 | 2 | 1136 | 1246 | 988 |
| 4,48158E | FALSE | High | IsMasterP | AAEL0049 | 2:260779 | 5 | 7 | FALSE | 3 | 3 | 3 | 542 | 620 | 654 |
| 7,67684E | FALSE | High | IsMasterP | AAEL0021 | 2:430623 | 5 | 10 | FALSE | 1 | 3 | 1 | 112 | 126 | 835 |
| 8,64302E | FALSE | High | IsMasterP | AAEL0069 | 2:398837 | 5 | 12 | FALSE | 2 | 3 | 2 | 330 | 368 | 802 |
| -3,1228E+ | FALSE | High | IsMasterP | AAEL0099 | 2:934054 | 5 | 3 | FALSE | 1 | 2 | 1 | 687 | 761 | 671 |
| 6,49868E | FALSE | High | IsMasterP | AAEL0119 | 3:452601 | 5 | 9 | FALSE | 2 | 3 | 2 | 282 | 312 | 531 |

| | | | | | | | | | | | | | | |
|---|---|---|---|---|---|---|---|---|---|---|---|---|---|---|
| 5,09741E | FALSE | High | IsMasterP | AAEL0087 | 3:133161 | 5 | 2 | FALSE | 1 | 3 | 1 | 451 | 515 | 762 |
| 4,66637E | FALSE | High | IsMasterP | AAEL0048 | 3:274601 | 5 | 5 | FALSE | 1 | 2 | 1 | 542 | 598 | 662 |
| 5,3625E+ | FALSE | High | IsMasterP | AAEL0086 | 2:239556 | 5 | 2 | FALSE | 1 | 3 | 1 | 699 | 786 | 831 |
| -1,39252E | FALSE | High | IsMasterP | AAEL0252 | 1:176945 | 5 | 9 | FALSE | 2 | 2 | 2 | 441 | 494 | 502 |
| 4,2827E+ | FALSE | High | IsMasterP | AAEL0050 | 2:445992 | 5 | 4 | FALSE | 2 | 3 | 2 | 575 | 657 | 985 |
| -8,68268E | FALSE | High | IsMasterP | AAEL0087 | 3:132462 | 5 | 3 | FALSE | 1 | 3 | 1 | 365 | 419 | 791 |
| -8,42747E | FALSE | High | IsMasterP | AAEL0008 | 3:362393 | 5 | 20 | FALSE | 2 | 2 | 2 | 306 | 345 | 557 |
| 4,40133E | FALSE | High | IsMasterP | AAEL0037 | 2:197398 | 6 | 17 | FALSE | 2 | 2 | 2 | 195 | 217 | 824 |
| 3,26746E | FALSE | High | IsMasterP | AAEL0183 | 2:407606 | 6 | 3 | FALSE | 1 | 2 | 1 | 983 | 1093 | 862 |
| 6,90252E | FALSE | High | IsMasterP | AAEL0023 | 3:373287 | 5 | 1 | FALSE | 2 | 3 | 2 | 1814 | 2023 | 625 |
| -2,83364E | FALSE | High | IsMasterP | AAEL0072 | 3:390148 | 5 | 1 | FALSE | 1 | 2 | 1 | 3651 | 4063 | 681 |
| 1,97976E | FALSE | High | IsMasterP | AAEL0233 | 1:305243 | 5 | 3 | FALSE | 2 | 3 | 2 | 727 | 829 | 599 |
| -8,95626E | FALSE | High | IsMasterP | AAEL0042 | 2:293458 | 5 | 8 | FALSE | 2 | 3 | 2 | 304 | 350 | 554 |
| -2,52476E | FALSE | High | IsMasterP | AAEL0067 | 2:851214 | 5 | 3 | FALSE | 1 | 2 | 1 | 820 | 898 | 923 |
| -2,20296E | FALSE | High | IsMasterP | AAEL0085 | 2:363272 | 5 | 3 | FALSE | 1 | 2 | 1 | 561 | 615 | 791 |
| 5,60965E | FALSE | High | IsMasterP | AAEL0127 | 1:290596 | 6 | 3 | FALSE | 2 | 3 | 2 | 955 | 1075 | 585 |
| 2,42146E | FALSE | High | IsMasterP | AAEL0001 | 1:194566 | 6 | 24 | FALSE | 2 | 3 | 2 | 124 | 149 | 970 |
| 1,22104E | FALSE | High | IsMasterP | AAEL0012 | 2:152983 | 6 | 10 | FALSE | 1 | 2 | 1 | 143 | 159 | 995 |
| -8,65274E | FALSE | High | IsMasterP | AAEL0218 | 1:176347 | 6 | 11 | FALSE | 3 | 3 | 3 | 423 | 467 | 737 |
| 1,85852E | FALSE | High | IsMasterP | AAEL0043 | 3:938299 | 6 | 8 | FALSE | 2 | 3 | 2 | 369 | 418 | 515 |
| -5,21361E | FALSE | High | IsMasterP | AAEL0031 | 2:253744 | 6 | 11 | FALSE | 1 | 2 | 1 | 156 | 183 | 473 |
| -8,84613E | FALSE | High | IsMasterP | AAEL0033 | 3:793062 | 6 | 4 | FALSE | 1 | 3 | 1 | 265 | 281 | 1032 |
| 4,27749E | FALSE | High | IsMasterP | AAEL0039 | 3:266885 | 6 | 8 | FALSE | 2 | 3 | 2 | 326 | 368 | 809 |
| -2,66992E | FALSE | High | IsMasterP | AAEL0106 | 2:279189 | 6 | 4 | FALSE | 1 | 3 | 1 | 328 | 366 | 926 |
| -8,0395E+ | FALSE | High | IsMasterP | AAEL0143 | 2:238475 | 7 | 8 | FALSE | 2 | 2 | 2 | 481 | 538 | 800 |
| -7,46638E | FALSE | High | IsMasterP | AAEL0092 | 2:268403 | 7 | 6 | FALSE | 2 | 3 | 2 | 470 | 537 | 594 |
| 4,15426E | FALSE | High | IsMasterP | AAEL0047 | 2:290411 | 7 | 5 | FALSE | 1 | 2 | 1 | 327 | 377 | 571 |
| 5,50578E | FALSE | High | IsMasterP | AAEL0051 | 2:172658 | 7 | 3 | FALSE | 1 | 3 | 1 | 356 | 402 | 819 |
| 8,21491E | FALSE | High | IsMasterP | AAEL0020 | 2:356594 | 7 | 4 | FALSE | 3 | 3 | 3 | 1182 | 1334 | 756 |
| -4,32594E | FALSE | High | IsMasterP | AAEL0154 | 1:188978 | 7 | 4 | FALSE | 1 | 3 | 1 | 396 | 433 | 733 |
| 7,91332E | FALSE | High | IsMasterP | AAEL0075 | 2:248893 | 7 | 13 | FALSE | 1 | 2 | 1 | 187 | 203 | 736 |

| | | | | | | | | | | | | | | |
|---|---|---|---|---|---|---|---|---|---|---|---|---|---|---|
| -1,18041E | FALSE | High | IsMasterP | AAEL0070 | 3:2984302 | 7 | 14 | FALSE | 2 | 2 | 2 | 225 | 253 | 701 |
| 6,67142E- | FALSE | High | IsMasterP | AAEL0143 | 3:4469145 | 7 | 2 | FALSE | 1 | 2 | 1 | 734 | 815 | 709 |
| -8,66553E | FALSE | High | IsMasterP | AAEL0068 | 3:3684605 | 7 | 5 | FALSE | 2 | 3 | 2 | 536 | 616 | 800 |
| -5,61917E | FALSE | High | IsMasterP | AAEL0119 | 2:2181006 | 7 | 6 | FALSE | 1 | 3 | 1 | 218 | 249 | 560 |
| 2,774E+18 | FALSE | High | IsMasterP | AAEL0097 | 3:1555825 | 7 | 16 | FALSE | 2 | 2 | 2 | 296 | 331 | 482 |
| 8,00568E- | FALSE | High | IsMasterP | AAEL0085 | 3:2224865 | 7 | 10 | FALSE | 1 | 2 | 1 | 210 | 229 | 652 |
| 2,56164E- | FALSE | High | IsMasterP | AAEL0018 | 2:2767428 | 7 | 9 | FALSE | 1 | 3 | 1 | 171 | 188 | 724 |
| -4,29998E | FALSE | High | IsMasterP | AAEL0140 | 2:1691227 | 7 | 3 | FALSE | 2 | 3 | 2 | 1040 | 1185 | 632 |
| -4,02593E | FALSE | High | IsMasterP | AAEL0013 | 3:4764718 | 7 | 6 | FALSE | 2 | 2 | 2 | 565 | 650 | 910 |
| 9,13878E- | FALSE | High | IsMasterP | AAEL0087 | 3:2804759 | 7 | 8 | FALSE | 2 | 2 | 2 | 403 | 457 | 660 |
| 6,36691E- | FALSE | High | IsMasterP | AAEL0010 | 1:1852202 | 7 | 15 | FALSE | 2 | 3 | 2 | 156 | 167 | 441 |
| 7,06999E- | FALSE | High | IsMasterP | AAEL0009 | 3:3158760 | 7 | 13 | FALSE | 2 | 2 | 2 | 174 | 201 | 847 |
| 9,11335E- | FALSE | High | IsMasterP | AAEL0019 | 2:4239736 | 7 | 5 | FALSE | 2 | 3 | 2 | 526 | 605 | 878 |
| -9,53293E | FALSE | High | IsMasterP | AAEL0063 | 1:1014496 | 7 | 10 | FALSE | 2 | 3 | 2 | 233 | 263 | 790 |
| 5,88283E- | FALSE | High | IsMasterP | AAEL0202 | 1:1119188 | 7 | 1 | FALSE | 1 | 3 | 1 | 2355 | 2609 | 642 |
| -5,22395E | FALSE | High | IsMasterP | AAEL0131 | 2:3891354 | 7 | 14 | FALSE | 2 | 2 | 2 | 425 | 484 | 695 |
| 2,23622E- | FALSE | High | IsMasterP | AAEL0271 | 3:1395000 | 7 | 7 | FALSE | 1 | 2 | 1 | 244 | 269 | 553 |
| -6,23576E | FALSE | High | IsMasterP | AAEL0087 | 3:2810703 | 7 | 3 | FALSE | 1 | 3 | 1 | 391 | 458 | 658 |
| -7,76947E | FALSE | High | IsMasterP | AAEL0013 | 3:4766423 | 7 | 4 | FALSE | 2 | 3 | 2 | 511 | 588 | 701 |
| -4,8508E+ | FALSE | High | IsMasterP | AAEL0274 | 2:2185243 | 7 | 6 | FALSE | 2 | 3 | 2 | 420 | 480 | 854 |
| -6,24854E | FALSE | High | IsMasterP | AAEL0052 | 3:8872371 | 7 | 2 | FALSE | 1 | 2 | 1 | 784 | 853 | 585 |
| 1,7458E+1 | FALSE | High | IsMasterP | AAEL0058 | NIGP0100 | 7 | 9 | FALSE | 1 | 2 | 1 | 161 | 184 | 941 |
| -4,6168E+ | FALSE | High | IsMasterP | AAEL0073 | 1:2717739 | 7 | 5 | FALSE | 2 | 3 | 2 | 475 | 514 | 484 |
| -1,68339E | FALSE | High | IsMasterP | AAEL0036 | 2:3167623 | 7 | 8 | FALSE | 2 | 3 | 2 | 289 | 338 | 824 |
| -6,34927E | FALSE | High | IsMasterP | AAEL0031 | 3:2709580 | 7 | 13 | FALSE | 3 | 3 | 3 | 238 | 273 | 932 |
| 5,43013E- | FALSE | High | IsMasterP | AAEL0005 | 3:3887750 | 7 | 6 | FALSE | 1 | 3 | 1 | 160 | 182 | 488 |
| 4,93194E- | FALSE | High | IsMasterP | AAEL0181 | 3:2761803 | 7 | 4 | FALSE | 2 | 3 | 2 | 966 | 1094 | 550 |
| -6,23881E | FALSE | High | IsMasterP | AAEL0120 | 3:3080021 | 7 | 23 | FALSE | 2 | 2 | 2 | 150 | 164 | 977 |
| -5,0384E+ | FALSE | High | IsMasterP | AAEL0115 | 3:3128392 | 7 | 12 | FALSE | 2 | 2 | 2 | 339 | 387 | 535 |
| 3,55715E- | FALSE | High | IsMasterP | AAEL0208 | NIGP0100 | 7 | 6 | FALSE | 2 | 2 | 2 | 296 | 327 | 939 |
| 4,30157E- | FALSE | High | IsMasterP | AAEL0034 | 2:1532166 | 7 | 6 | FALSE | 2 | 2 | 2 | 759 | 882 | 799 |

| | | | | | | | | | | | | | | |
|---|---|---|---|---|---|---|---|---|---|---|---|---|---|---|
| 4,51902E | FALSE | High | IsMasterP | AAEL0031 | 1:192935 | 7 | 7 | FALSE | 2 | 3 | 2 | 316 | 356 | 547 |
| 7,81401E | FALSE | High | IsMasterP | AAEL0102 | 3:157547 | 7 | 13 | FALSE | 1 | 2 | 1 | 214 | 231 | 615 |
| 2,09919E | FALSE | High | IsMasterP | AAEL0280 | 2:300510 | 7 | 3 | FALSE | 1 | 2 | 1 | 894 | 982 | 370 |
| 8,51697E | FALSE | High | IsMasterP | AAEL0091 | 1:234126 | 7 | 18 | FALSE | 2 | 2 | 2 | 149 | 172 | 543 |
| -7,94864E | FALSE | High | IsMasterP | AAEL0224 | 1:167733 | 7 | 3 | FALSE | 1 | 3 | 1 | 450 | 481 | 856 |
| 7,17556E | FALSE | High | IsMasterP | AAEL0050 | 2:444364 | 7 | 7 | FALSE | 3 | 3 | 3 | 427 | 481 | 671 |
| 7,08615E | FALSE | High | IsMasterP | AAEL0183 | 3:301116 | 7 | 3 | FALSE | 1 | 3 | 1 | 645 | 723 | 812 |
| -8,7E+18 | FALSE | High | IsMasterP | AAEL0036 | 2:317087 | 7 | 5 | FALSE | 2 | 3 | 2 | 722 | 812 | 539 |
| 7,25458E | FALSE | High | IsMasterP | AAEL0043 | 2:293463 | 7 | 5 | FALSE | 2 | 3 | 2 | 385 | 434 | 674 |
| -1,09616E | FALSE | High | IsMasterP | AAEL0114 | 2:143647 | 7 | 8 | FALSE | 1 | 2 | 1 | 180 | 207 | 904 |
| -6,08186E | FALSE | High | IsMasterP | AAEL0097 | 3:155716 | 7 | 4 | FALSE | 3 | 3 | 3 | 807 | 887 | 525 |
| 6,20582E | FALSE | High | IsMasterP | AAEL0002 | 3:216221 | 7 | 8 | FALSE | 2 | 3 | 2 | 297 | 329 | 1007 |
| -3,44648E | FALSE | High | IsMasterP | AAEL0212 | 3:293674 | 7 | 3 | FALSE | 1 | 2 | 1 | 639 | 686 | 929 |
| 1,01018E | FALSE | High | IsMasterP | AAEL0209 | 2:644584 | 7 | 13 | FALSE | 2 | 2 | 2 | 350 | 391 | 533 |
| -2,70061E | FALSE | High | IsMasterP | AAEL0010 | 3:312615 | 7 | 4 | FALSE | 1 | 2 | 1 | 653 | 739 | 936 |
| -4,86219E | FALSE | High | IsMasterP | AAEL0061 | 3:696867 | 7 | 8 | FALSE | 2 | 3 | 2 | 443 | 489 | 708 |
| -4,45215E | FALSE | High | IsMasterP | AAEL0065 | 2:593502 | 7 | 35 | FALSE | 2 | 2 | 2 | 116 | 135 | 825 |
| 6,75972E | FALSE | High | IsMasterP | AAEL0113 | 1:120092 | 7 | 6 | FALSE | 1 | 2 | 1 | 250 | 280 | 796 |
| 3,88768E | FALSE | High | IsMasterP | AAEL0206 | 3:214681 | 7 | 3 | FALSE | 2 | 3 | 2 | 803 | 908 | 769 |
| -6,58115E | FALSE | High | IsMasterP | AAEL0012 | 2:153239 | 7 | 26 | FALSE | 2 | 3 | 2 | 77 | 87 | 972 |
| -2,36836E | FALSE | High | IsMasterP | AAEL0073 | 2:233409 | 7 | 20 | FALSE | 1 | 1 | 1 | 186 | 215 | 624 |
| -2,98994E | FALSE | High | IsMasterP | AAEL0082 | 2:674208 | 7 | 15 | FALSE | 1 | 2 | 1 | 137 | 146 | 470 |
| -2,23011E | FALSE | High | IsMasterP | AAEL0018 | 2:375232 | 7 | 12 | FALSE | 1 | 3 | 1 | 114 | 128 | 957 |
| 4,10654E | FALSE | High | IsMasterP | AAEL0053 | 2:643909 | 7 | 17 | FALSE | 2 | 2 | 2 | 274 | 318 | 600 |
| -2,42317E | FALSE | High | IsMasterP | AAEL0093 | 1:552780 | 7 | 4 | FALSE | 1 | 3 | 1 | 315 | 350 | 638 |
| -4,99763E | FALSE | High | IsMasterP | AAEL0147 | 1:211953 | 7 | 9 | FALSE | 2 | 2 | 2 | 472 | 523 | 520 |
| -3718396 | FALSE | High | IsMasterP | AAEL0281 | NIGP0100 | 7 | 6 | FALSE | 2 | 2 | 2 | 726 | 828 | 503 |
| 7,50943E | FALSE | High | IsMasterP | AAEL0122 | 2:384054 | 7 | 4 | FALSE | 1 | 2 | 1 | 509 | 592 | 787 |
| -3,58246E | FALSE | High | IsMasterP | AAEL0007 | 3:243413 | 7 | 13 | FALSE | 2 | 3 | 1 | 245 | 286 | 1130 |
| 4,46183E | FALSE | High | IsMasterP | AAEL0024 | 1:177333 | 7 | 7 | FALSE | 2 | 2 | 2 | 879 | 984 | 806 |
| 2,59344E | FALSE | High | IsMasterP | AAEL0081 | 3:107633 | 7 | 4 | FALSE | 1 | 3 | 1 | 306 | 343 | 563 |

| | | | | | | | | | | | | | | |
|---|---|---|---|---|---|---|---|---|---|---|---|---|---|---|
| -7,53661E | FALSE | High | IsMasterP | AAEL0036 | 1:699303 | 7 | 18 | FALSE | 2 | 2 | 2 | 312 | 357 | 642 |
| 8,66325E- | FALSE | High | IsMasterP | AAEL0017 | 2:165703 | 7 | 14 | FALSE | 2 | 2 | 2 | 317 | 355 | 628 |
| -2,35357E | FALSE | High | IsMasterP | AAEL0097 | 3:284486 | 7 | 7 | FALSE | 2 | 3 | 2 | 370 | 413 | 580 |
| -2,42675E | FALSE | High | IsMasterP | AAEL0143 | 3:445891 | 7 | 2 | FALSE | 1 | 2 | 1 | 905 | 1058 | 574 |
| 5,14395E- | FALSE | High | IsMasterP | AAEL0246 | 2:448274 | 7 | 4 | FALSE | 2 | 2 | 2 | 1995 | 2191 | 768 |
| 4,95627E- | FALSE | High | IsMasterP | AAEL0002 | 2:753567 | 7 | 6 | FALSE | 1 | 2 | 1 | 223 | 254 | 464 |
| -7,10166E | FALSE | High | IsMasterP | AAEL0021 | 1:243871 | 7 | 30 | FALSE | 2 | 2 | 2 | 115 | 132 | 846 |
| -2,4732E+ | FALSE | High | IsMasterP | AAEL0107 | 2:306346 | 7 | 5 | FALSE | 2 | 2 | 2 | 346 | 399 | 648 |
| 4,60073E- | FALSE | High | IsMasterP | AAEL0001 | 3:213028 | 7 | 7 | FALSE | 1 | 2 | 1 | 171 | 199 | 928 |
| -3,05118E | FALSE | High | IsMasterP | AAEL0100 | 2:358197 | 7 | 11 | FALSE | 2 | 3 | 2 | 238 | 264 | 588 |
| -1,78828E | FALSE | High | IsMasterP | AAEL0202 | 2:111743 | 7 | 3 | FALSE | 1 | 2 | 1 | 484 | 535 | 781 |
| 6,28063E- | FALSE | High | IsMasterP | AAEL0049 | 3:323488 | 8 | 4 | FALSE | 1 | 3 | 1 | 310 | 344 | 557 |
| 4,96492E- | FALSE | High | IsMasterP | AAEL0094 | 2:247108 | 8 | 9 | FALSE | 2 | 3 | 2 | 256 | 293 | 911 |
| -3,90852E | FALSE | High | IsMasterP | AAEL0227 | 2:188083 | 8 | 11 | FALSE | 2 | 3 | 2 | 198 | 225 | 907 |
| 6,61596E- | FALSE | High | IsMasterP | AAEL0074 | 2:134750 | 8 | 6 | FALSE | 1 | 3 | 1 | 240 | 276 | 1017 |
| -5,62691E | FALSE | High | IsMasterP | AAEL0195 | 3:245281 | 8 | 1 | FALSE | 1 | 2 | 1 | 1338 | 1516 | 847 |
| -7,99926E | FALSE | High | IsMasterP | AAEL0009 | 3:380008 | 8 | 9 | FALSE | 2 | 2 | 2 | 534 | 599 | 555 |
| 7,22089E- | FALSE | High | IsMasterP | AAEL0280 | 1:597440 | 8 | 5 | FALSE | 2 | 2 | 2 | 1536 | 1706 | 725 |
| 9,12623E- | FALSE | High | IsMasterP | AAEL0085 | 1:642944 | 8 | 5 | FALSE | 2 | 2 | 2 | 790 | 871 | 540 |
| -7,75017E | FALSE | High | IsMasterP | AAEL0030 | 3:397841 | 8 | 3 | FALSE | 1 | 2 | 1 | 583 | 637 | 702 |
| -7,93636E | FALSE | High | IsMasterP | AAEL0011 | 2:307293 | 8 | 7 | FALSE | 2 | 3 | 2 | 394 | 437 | 632 |
| -4,37993E | FALSE | High | IsMasterP | AAEL0025 | 3:256818 | 8 | 2 | FALSE | 1 | 3 | 1 | 361 | 397 | 543 |
| 6,59094E- | FALSE | High | IsMasterP | AAEL0077 | 1:718815 | 8 | 18 | FALSE | 2 | 2 | 2 | 201 | 234 | 952 |
| -1,91733E | FALSE | High | IsMasterP | AAEL0254 | 3:206234 | 8 | 1 | FALSE | 2 | 2 | 2 | 3914 | 4447 | 780 |
| 2,45246E- | FALSE | High | IsMasterP | AAEL0019 | 3:227087 | 8 | 2 | FALSE | 1 | 2 | 1 | 626 | 708 | 586 |
| -7,01645E | FALSE | High | IsMasterP | AAEL0073 | 3:408309 | 8 | 3 | FALSE | 1 | 2 | 1 | 351 | 384 | 692 |
| 2,0996E+ | FALSE | High | IsMasterP | AAEL0066 | 3:148446 | 8 | 14 | FALSE | 1 | 3 | 1 | 91 | 100 | 958 |
| -1,23686E | FALSE | High | IsMasterP | AAEL0036 | 3:651085 | 8 | 2 | FALSE | 1 | 2 | 1 | 608 | 679 | 787 |
| -2,75785E | FALSE | High | IsMasterP | AAEL0275 | 1:908384 | 8 | 3 | FALSE | 1 | 3 | 1 | 1179 | 1264 | 945 |
| 8,7667E+ | FALSE | High | IsMasterP | AAEL0135 | 1:144023 | 8 | 3 | FALSE | 1 | 2 | 1 | 422 | 487 | 862 |
| 2,50058E- | FALSE | High | IsMasterP | AAEL0245 | 1:307040 | 8 | 2 | FALSE | 2 | 3 | 2 | 1973 | 2164 | 911 |

| | | | | | | | | | | | | | | |
|---|---|---|---|---|---|---|---|---|---|---|---|---|---|---|
| -1,00677E | FALSE | High | IsMasterP | AAEL0249 | 1:611419 | 8 | 6 | FALSE | 2 | 2 | 2 | 547 | 615 | 560 |
| 4,09825E- | FALSE | High | IsMasterP | AAEL0035 | 1:691578 | 8 | 23 | FALSE | 1 | 1 | 1 | 151 | 167 | 881 |
| -9,12176E | FALSE | High | IsMasterP | AAEL0132 | 1:307370 | 8 | 6 | FALSE | 1 | 2 | 1 | 290 | 334 | 951 |
| 5,28355E- | FALSE | High | IsMasterP | AAEL0039 | 3:253907 | 8 | 4 | FALSE | 1 | 3 | 1 | 314 | 340 | 923 |
| 5,7425E+1 | FALSE | High | IsMasterP | AAEL0130 | 1:252678 | 8 | 1 | FALSE | 1 | 3 | 1 | 877 | 1000 | 829 |
| 4,20398E- | FALSE | High | IsMasterP | AAEL0131 | 3:842126 | 8 | 4 | FALSE | 1 | 2 | 1 | 364 | 415 | 885 |
| -9,92954E | FALSE | High | IsMasterP | AAEL0021 | 2:429549 | 8 | 4 | FALSE | 2 | 3 | 2 | 673 | 757 | 655 |
| -6,17714E | FALSE | High | IsMasterP | AAEL0276 | 2:898697 | 8 | 5 | FALSE | 2 | 2 | 2 | 751 | 831 | 481 |
| 3,62779E- | FALSE | High | IsMasterP | AAEL0043 | 3:366697 | 8 | 17 | FALSE | 2 | 2 | 2 | 157 | 179 | 687 |
| 7,91926E- | FALSE | High | IsMasterP | AAEL0253 | 3:109291 | 8 | 6 | FALSE | 1 | 3 | 1 | 200 | 237 | 873 |
| -8,35489E | FALSE | High | IsMasterP | AAEL0054 | 1:788082 | 8 | 4 | FALSE | 1 | 3 | 1 | 366 | 408 | 802 |
| 1,13376E- | FALSE | High | IsMasterP | AAEL0044 | 1:203637 | 8 | 3 | FALSE | 1 | 2 | 1 | 1139 | 1232 | 951 |
| -5,81481E | FALSE | High | IsMasterP | AAEL0030 | 2:355380 | 8 | 10 | FALSE | 2 | 2 | 2 | 327 | 376 | 756 |
| -8,25799E | FALSE | High | IsMasterP | AAEL0195 | 2:453370 | 8 | 5 | FALSE | 2 | 3 | 2 | 404 | 447 | 634 |
| 9,12464E- | FALSE | High | IsMasterP | AAEL0274 | 2:179741 | 8 | 12 | FALSE | 2 | 2 | 2 | 288 | 332 | 982 |
| -1,99546E | FALSE | High | IsMasterP | AAEL0197 | 3:200745 | 8 | 1 | FALSE | 1 | 1 | 1 | 3675 | 4199 | 571 |
| 8,58503E- | FALSE | High | IsMasterP | AAEL0140 | 1:220406 | 8 | 4 | FALSE | 1 | 2 | 1 | 375 | 422 | 662 |
| -7,19445E | FALSE | High | IsMasterP | AAEL0038 | 3:850911 | 8 | 1 | FALSE | 1 | 2 | 1 | 1096 | 1245 | 651 |
| -6,5192E+ | FALSE | High | IsMasterP | AAEL0198 | 2:215460 | 8 | 2 | FALSE | 1 | 3 | 1 | 364 | 421 | 711 |
| -7,16458E | FALSE | High | IsMasterP | AAEL0120 | 2:186377 | 8 | 4 | FALSE | 1 | 3 | 1 | 331 | 355 | 708 |
| -3,80434E | FALSE | High | IsMasterP | AAEL0199 | 2:309059 | 8 | 5 | FALSE | 1 | 2 | 1 | 373 | 419 | 511 |
| -4,24266E | FALSE | High | IsMasterP | AAEL0033 | 3:800079 | 8 | 8 | FALSE | 1 | 2 | 1 | 193 | 227 | 974 |
| -4,00289E | FALSE | High | IsMasterP | AAEL0116 | 3:291704 | 8 | 5 | FALSE | 1 | 2 | 1 | 397 | 444 | 799 |
| 4,91802E- | FALSE | High | IsMasterP | AAEL0028 | 1:305571 | 8 | 6 | FALSE | 1 | 2 | 1 | 233 | 258 | 972 |
| -1,77428E | FALSE | High | IsMasterP | AAEL0107 | 3:156738 | 8 | 17 | FALSE | 1 | 2 | 1 | 141 | 159 | 465 |
| -6,33254E | FALSE | High | IsMasterP | AAEL0252 | 3:268353 | 8 | 7 | FALSE | 2 | 2 | 2 | 522 | 570 | 828 |
| 2,82189E- | FALSE | High | IsMasterP | AAEL0182 | 3:359900 | 8 | 1 | FALSE | 2 | 3 | 2 | 1750 | 1946 | 590 |
| 2,21669E- | FALSE | High | IsMasterP | AAEL0087 | 3:957465 | 8 | 1 | FALSE | 1 | 2 | 1 | 1249 | 1416 | 897 |
| 1,26012E- | FALSE | High | IsMasterP | AAEL0003 | 3:400696 | 8 | 3 | FALSE | 1 | 3 | 1 | 412 | 464 | 840 |
| 7,67915E- | FALSE | High | IsMasterP | AAEL0041 | 3:153796 | 8 | 4 | FALSE | 2 | 2 | 2 | 969 | 1110 | 683 |
| -2,34206E | FALSE | High | IsMasterP | AAEL0199 | 2:199007 | 8 | 5 | FALSE | 1 | 2 | 1 | 253 | 284 | 903 |

| | | | | | | | | | | | | | | |
|---|---|---|---|---|---|---|---|---|---|---|---|---|---|---|
| 5,92156E- | FALSE | High | IsMasterP | AAEL0041 | 2:147387 | 8 | 11 | FALSE | 1 | 2 | 1 | 138 | 158 | 970 |
| 4,78111E- | FALSE | High | IsMasterP | AAEL0198 | 1:643263 | 8 | 6 | FALSE | 2 | 2 | 2 | 592 | 655 | 564 |
| 4,32044E- | FALSE | High | IsMasterP | AAEL0145 | 2:348408 | 8 | 2 | FALSE | 1 | 3 | 1 | 425 | 484 | 693 |
| -6,77857E | FALSE | High | IsMasterP | AAEL0112 | 2:179628 | 8 | 13 | FALSE | 2 | 2 | 2 | 324 | 371 | 926 |
| -4,08433E | FALSE | High | IsMasterP | AAEL0081 | 3:935879 | 8 | 3 | FALSE | 1 | 2 | 1 | 646 | 662 | 1059 |
| 7,82269E- | FALSE | High | IsMasterP | AAEL0046 | 3:247533 | 8 | 3 | FALSE | 1 | 2 | 1 | 1193 | 1289 | 824 |
| 2,23795E- | FALSE | High | IsMasterP | AAEL0199 | 3:276244 | 8 | 2 | FALSE | 2 | 2 | 2 | 1860 | 2130 | 938 |
| -9,33932E | FALSE | High | IsMasterP | AAEL0069 | 3:143898 | 8 | 11 | FALSE | 2 | 2 | 2 | 305 | 348 | 531 |
| 6,83442E- | FALSE | High | IsMasterP | AAEL0110 | 2:631013 | 8 | 5 | FALSE | 1 | 2 | 1 | 358 | 410 | 640 |
| 4,94731E- | FALSE | High | IsMasterP | AAEL0048 | 3:273466 | 8 | 1 | FALSE | 1 | 2 | 1 | 810 | 927 | 639 |
| -3,75536E | FALSE | High | IsMasterP | AAEL0271 | 3:149397 | 8 | 3 | FALSE | 1 | 3 | 1 | 418 | 480 | 560 |
| 6,38474E- | FALSE | High | IsMasterP | AAEL0029 | 3:389055 | 8 | 8 | FALSE | 3 | 3 | 3 | 377 | 426 | 487 |
| 5,95049E- | FALSE | High | IsMasterP | AAEL0069 | 2:340163 | 8 | 10 | FALSE | 1 | 2 | 1 | 122 | 141 | 887 |
| 9,17732E- | FALSE | High | IsMasterP | AAEL0075 | 1:302591 | 8 | 14 | FALSE | 1 | 2 | 1 | 133 | 153 | 592 |
| -3,83463E | FALSE | High | IsMasterP | AAEL0043 | 3:941682 | 8 | 6 | FALSE | 2 | 2 | 2 | 644 | 737 | 604 |
| -2,78588E | FALSE | High | IsMasterP | AAEL0197 | 2:274209 | 8 | 5 | FALSE | 1 | 3 | 1 | 237 | 274 | 1004 |
| -4,16571E | FALSE | High | IsMasterP | AAEL0023 | 3:375065 | 8 | 16 | FALSE | 2 | 2 | 2 | 247 | 283 | 718 |
| 3,44725E- | FALSE | High | IsMasterP | AAEL0015 | 2:206638 | 9 | 2 | FALSE | 2 | 2 | 2 | 1284 | 1434 | 674 |
| 3,76747E- | FALSE | High | IsMasterP | AAEL0195 | 2:161719 | 9 | 3 | FALSE | 1 | 3 | 1 | 272 | 305 | 632 |
| -2,55605E | FALSE | High | IsMasterP | AAEL0041 | 3:401784 | 9 | 2 | FALSE | 1 | 2 | 1 | 1294 | 1459 | 797 |
| -1,4134E+ | FALSE | High | IsMasterP | AAEL0066 | 3:200556 | 9 | 9 | FALSE | 1 | 2 | 1 | 140 | 154 | 599 |
| 1,35299E- | FALSE | High | IsMasterP | AAEL0043 | 1:400629 | 9 | 3 | FALSE | 2 | 2 | 2 | 1065 | 1230 | 903 |
| 7,81472E- | FALSE | High | IsMasterP | AAEL0088 | 2:241819 | 9 | 1 | FALSE | 1 | 3 | 1 | 734 | 807 | 661 |
| -8,11902E | FALSE | High | IsMasterP | AAEL0069 | 3:143066 | 9 | 3 | FALSE | 2 | 2 | 2 | 1106 | 1258 | 657 |
| -7,08707E | FALSE | High | IsMasterP | AAEL0006 | 2:408790 | 9 | 3 | FALSE | 1 | 2 | 1 | 517 | 601 | 596 |
| -3,00003E | FALSE | High | IsMasterP | AAEL0008 | 3:332256 | 9 | 3 | FALSE | 2 | 2 | 2 | 703 | 786 | 569 |
| -2,63198E | FALSE | High | IsMasterP | AAEL0092 | 2:170934 | 9 | 4 | FALSE | 2 | 2 | 2 | 860 | 953 | 515 |
| 4,47629E- | FALSE | High | IsMasterP | AAEL0113 | 1:165568 | 9 | 8 | FALSE | 2 | 2 | 2 | 341 | 403 | 483 |
| -2,49061E | FALSE | High | IsMasterP | AAEL0103 | 2:182936 | 9 | 3 | FALSE | 1 | 2 | 1 | 540 | 612 | 559 |
| 6,62097E- | FALSE | High | IsMasterP | AAEL0175 | 1:989206 | 9 | 5 | FALSE | 1 | 3 | 1 | 199 | 225 | 692 |
| 6,27604E- | FALSE | High | IsMasterP | AAEL0039 | 3:267992 | 9 | 5 | FALSE | 2 | 2 | 2 | 1196 | 1300 | 977 |

| | | | | | | | | | | | | | | |
|---|---|---|---|---|---|---|---|---|---|---|---|---|---|---|
| 3,27379E+ | FALSE | High | IsMasterP | AAEL0071 | 3:1907116 | 9 | 2 | FALSE | 1 | 2 | 1 | 697 | 803 | 900 |
| 1,39735E+ | FALSE | High | IsMasterP | AAEL0170 | 1:2713759 | 9 | 3 | FALSE | 1 | 2 | 1 | 504 | 580 | 689 |
| 3,96805E+ | FALSE | High | IsMasterP | AAEL0054 | 2:2963738 | 9 | 2 | FALSE | 1 | 2 | 1 | 1252 | 1380 | 550 |
| -3,45786E | FALSE | Medium | IsMasterP | AAEL0001 | 1:1973440 | 10 | 5 | FALSE | 1 | 2 | 1 | 286 | 324 | 640 |
| -2,03361E | FALSE | Medium | IsMasterP | AAEL0072 | 3:3902450 | 10 | 5 | FALSE | 2 | 2 | 2 | 561 | 618 | 772 |
| 9,39077E+ | FALSE | Medium | IsMasterP | AAEL0063 | 3:3273097 | 10 | 2 | FALSE | 2 | 2 | 2 | 1474 | 1664 | 611 |
| 8,55601E+ | FALSE | Medium | IsMasterP | AAEL0049 | 2:2812220 | 10 | 3 | FALSE | 2 | 2 | 2 | 1141 | 1287 | 865 |
| -2,472E+1 | FALSE | Medium | IsMasterP | AAEL0023 | 2:3022002 | 10 | 2 | FALSE | 1 | 2 | 1 | 523 | 600 | 580 |
| -6,38734E | FALSE | Medium | IsMasterP | AAEL0052 | 2:4183693 | 10 | 8 | FALSE | 2 | 2 | 2 | 508 | 583 | 497 |
| 7,74076E+ | FALSE | Medium | IsMasterP | AAEL0032 | 1:2916920 | 11 | 5 | FALSE | 1 | 2 | 1 | 712 | 805 | 520 |
| -7,31405E | FALSE | Medium | IsMasterP | AAEL0250 | 2:1559483 | 11 | 8 | FALSE | 1 | 2 | 1 | 229 | 230 | 506 |
| 8,47145E+ | FALSE | Medium | IsMasterP | AAEL0140 | 2:4059580 | 11 | 8 | FALSE | 1 | 2 | 1 | 183 | 209 | 957 |
| -7,13199E | FALSE | Medium | IsMasterP | AAEL0150 | 1:2139608 | 11 | 6 | FALSE | 2 | 2 | 2 | 668 | 759 | 913 |
| -8,43188E | FALSE | Medium | IsMasterP | AAEL0130 | 3:3106234 | 11 | 6 | FALSE | 2 | 2 | 2 | 422 | 479 | 671 |
| -2,70585E | FALSE | Medium | IsMasterP | AAEL0183 | 3:2186446 | 11 | 4 | FALSE | 2 | 2 | 2 | 831 | 939 | 813 |
| -2,47814E | FALSE | Medium | IsMasterP | AAEL0052 | 2:2861630 | 11 | 2 | FALSE | 2 | 2 | 2 | 2050 | 2310 | 693 |
| -9,89976E | FALSE | Medium | IsMasterP | AAEL0290 | 3:1385395 | 11 | 17 | FALSE | 2 | 2 | 2 | 138 | 162 | 498 |
| -9,92745E | FALSE | Medium | IsMasterP | AAEL0261 | 3:1755936 | 11 | 1 | FALSE | 1 | 2 | 1 | 2148 | 2398 | 655 |
| -5,11258E | FALSE | Medium | IsMasterP | AAEL0133 | 1:1928043 | 11 | 3 | FALSE | 2 | 2 | 2 | 1049 | 1167 | 473 |
| 8,27514E+ | FALSE | Medium | IsMasterP | AAEL0082 | 3:6329335 | 11 | 7 | FALSE | 1 | 3 | 1 | 181 | 204 | 646 |
| -5,92965E | FALSE | Medium | IsMasterP | AAEL0091 | 1:2340839 | 11 | 1 | FALSE | 1 | 2 | 1 | 1405 | 1522 | 483 |
| 5,90128E+ | FALSE | Medium | IsMasterP | AAEL0009 | 2:2146235 | 11 | 6 | FALSE | 1 | 2 | 1 | 289 | 319 | 766 |
| -2,58401E | FALSE | Medium | IsMasterP | AAEL0231 | 2:6889763 | 11 | 1 | FALSE | 1 | 3 | 1 | 996 | 1131 | 873 |
| 7,09447E+ | FALSE | Medium | IsMasterP | AAEL0274 | 2:2102335 | 11 | 1 | FALSE | 2 | 2 | 2 | 3145 | 3456 | 882 |
| 4,87123E+ | FALSE | Medium | IsMasterP | AAEL0039 | 3:2541834 | 11 | 11 | FALSE | 1 | 1 | 1 | 300 | 338 | 655 |
| -8,08591E | FALSE | Medium | IsMasterP | AAEL0000 | 3:1056699 | 11 | 5 | FALSE | 1 | 3 | 1 | 379 | 427 | 655 |
| -1,05513E | FALSE | Medium | IsMasterP | AAEL0070 | 3:2986279 | 11 | 3 | FALSE | 1 | 2 | 1 | 631 | 719 | 508 |
| -1,66822E | FALSE | Medium | IsMasterP | AAEL0140 | 3:1473852 | 11 | 1 | FALSE | 2 | 3 | 2 | 2181 | 2493 | 746 |
| -7,40082E | FALSE | Medium | IsMasterP | AAEL0005 | 2:2834205 | 11 | 4 | FALSE | 2 | 2 | 2 | 797 | 912 | 853 |
| -5,28754E | FALSE | Medium | IsMasterP | AAEL0046 | 3:2486598 | 11 | 5 | FALSE | 2 | 2 | 2 | 694 | 792 | 609 |
| 2,99054E+ | FALSE | Medium | IsMasterP | AAEL0198 | 2:3263297 | 11 | 2 | FALSE | 1 | 3 | 1 | 915 | 1071 | 629 |

| | | | | | | | | | | | | | | |
|---|---|---|---|---|---|---|---|---|---|---|---|---|---|---|
| -8,33135E | FALSE | Medium | IsMasterP | AAEL0279 | 1:4555113 | 11 | 6 | FALSE | 1 | 2 | 1 | 241 | 261 | 850 |
| -8,01292E | FALSE | Medium | IsMasterP | AAEL0096 | 3:2693127 | 11 | 1 | FALSE | 2 | 3 | 2 | 1972 | 2271 | 846 |
| 9,2067E+1 | FALSE | Medium | IsMasterP | AAEL0091 | 3:1688155 | 12 | 4 | FALSE | 2 | 2 | 2 | 764 | 824 | 654 |
| -4,40508E | FALSE | Medium | IsMasterP | AAEL0009 | 3:3772560 | 12 | 6 | FALSE | 1 | 3 | 1 | 140 | 155 | 870 |
| -1,2098E+ | FALSE | Medium | IsMasterP | AAEL0004 | 3:1871618 | 12 | 4 | FALSE | 2 | 2 | 2 | 693 | 761 | 925 |
| -7,72327E | FALSE | Medium | IsMasterP | AAEL0032 | 1:2908251 | 12 | 14 | FALSE | 2 | 2 | 2 | 184 | 210 | 596 |
| 3,03556E- | FALSE | Medium | IsMasterP | AAEL0040 | 3:4033527 | 12 | 1 | FALSE | 1 | 2 | 1 | 949 | 1036 | 825 |
| 7,6762E+1 | FALSE | Medium | IsMasterP | AAEL0015 | 2:2037839 | 12 | 2 | FALSE | 1 | 2 | 1 | 1435 | 1658 | 796 |
| -1,70788E | FALSE | Medium | IsMasterP | AAEL0142 | 3:2931900 | 12 | 3 | FALSE | 2 | 2 | 2 | 1152 | 1328 | 737 |
| -1,08801E | FALSE | Medium | IsMasterP | AAEL0017 | 2:4418604 | 12 | 5 | FALSE | 2 | 2 | 2 | 1052 | 1196 | 733 |
| -7,85017E | FALSE | Medium | IsMasterP | AAEL0032 | 1:2909795 | 12 | 1 | FALSE | 2 | 3 | 2 | 1715 | 1900 | 458 |
| -4,98794E | FALSE | Medium | IsMasterP | AAEL0132 | 2:9480716 | 12 | 7 | FALSE | 1 | 2 | 1 | 257 | 274 | 588 |
| 2,19362E- | FALSE | Medium | IsMasterP | AAEL0103 | 2:4038009 | 12 | 5 | FALSE | 2 | 2 | 2 | 557 | 619 | 894 |
| 4,47826E- | FALSE | Medium | IsMasterP | AAEL0014 | 1:2881076 | 12 | 12 | FALSE | 2 | 2 | 1 | 287 | 309 | 834 |
| -7,72527E | FALSE | Medium | IsMasterP | AAEL0143 | 3:8673321 | 12 | 5 | FALSE | 1 | 3 | 1 | 379 | 434 | 848 |
| 1,47032E- | FALSE | Medium | IsMasterP | AAEL0112 | 1:3297010 | 12 | 3 | FALSE | 1 | 2 | 1 | 1224 | 1431 | 815 |
| 6,79615E- | FALSE | Medium | IsMasterP | AAEL0107 | 1:1786188 | 12 | 3 | FALSE | 1 | 3 | 1 | 339 | 376 | 736 |
| 8,98492E- | FALSE | Medium | IsMasterP | AAEL0080 | 1:5184317 | 12 | 7 | FALSE | 1 | 2 | 1 | 296 | 332 | 916 |
| -3,74903E | FALSE | Medium | IsMasterP | AAEL0181 | 3:3148471 | 12 | 2 | FALSE | 1 | 2 | 1 | 619 | 710 | 771 |
| -5,92477E | FALSE | Medium | IsMasterP | AAEL0077 | 3:1334652 | 12 | 11 | FALSE | 2 | 2 | 2 | 307 | 345 | 906 |
| -8,5952E+ | FALSE | Medium | IsMasterP | AAEL0108 | 1:5468388 | 12 | 7 | FALSE | 1 | 2 | 1 | 207 | 244 | 970 |
| -1,00119E | FALSE | Medium | IsMasterP | AAEL0218 | 2:2479367 | 12 | 4 | FALSE | 1 | 2 | 1 | 400 | 458 | 469 |
| 6,34537E- | FALSE | Medium | IsMasterP | AAEL0204 | 2:4722898 | 12 | 2 | FALSE | 1 | 2 | 1 | 591 | 680 | 524 |
| 6,49259E- | FALSE | Medium | IsMasterP | AAEL0135 | 2:8279808 | 12 | 5 | FALSE | 1 | 1 | 1 | 437 | 493 | 925 |
| 8,66515E- | FALSE | Medium | IsMasterP | AAEL0134 | 2:1554229 | 13 | 2 | FALSE | 1 | 2 | 1 | 949 | 1038 | 655 |
| -4,75013E | FALSE | Medium | IsMasterP | AAEL0134 | 1:2065171 | 13 | 4 | FALSE | 2 | 2 | 2 | 679 | 778 | 712 |
| -7,36985E | FALSE | Medium | IsMasterP | AAEL0084 | 1:1728837 | 13 | 3 | FALSE | 1 | 2 | 1 | 449 | 515 | 911 |
| -3,43824E | FALSE | Medium | IsMasterP | AAEL0228 | NIGP0100 | 13 | 19 | FALSE | 2 | 2 | 2 | 230 | 256 | 583 |
| 7,40368E- | FALSE | Medium | IsMasterP | AAEL0096 | 3:2102476 | 13 | 3 | FALSE | 1 | 3 | 1 | 289 | 326 | 835 |
| 6,68444E- | FALSE | Medium | IsMasterP | AAEL0039 | 2:3545086 | 13 | 1 | FALSE | 1 | 2 | 1 | 1730 | 1906 | 648 |
| 2,60237E- | FALSE | Medium | IsMasterP | AAEL0170 | 3:1406230 | 13 | 8 | FALSE | 1 | 2 | 1 | 199 | 235 | 740 |

| | | | | | | | | | | | | | | |
|---|---|---|---|---|---|---|---|---|---|---|---|---|---|---|
| 5,89863E+ | FALSE | Medium | IsMasterP | AAEL0245 | 2:7826482 | 13 | 2 | FALSE | 2 | 2 | 2 | 1441 | 1620 | 909 |
| 2,63154E+ | FALSE | Medium | IsMasterP | AAEL0015 | 2:2617741 | 13 | 2 | FALSE | 2 | 2 | 2 | 1291 | 1420 | 806 |
| 7,41561E+ | FALSE | Medium | IsMasterP | AAEL0036 | 3:3013623 | 13 | 3 | FALSE | 2 | 2 | 2 | 615 | 683 | 628 |
| 5,16932E+ | FALSE | Medium | IsMasterP | AAEL0203 | 2:1246459 | 14 | 3 | FALSE | 1 | 2 | 1 | 668 | 757 | 586 |
| -4,94717E | FALSE | Medium | IsMasterP | AAEL0182 | 1:7402363 | 14 | 3 | FALSE | 1 | 2 | 1 | 532 | 589 | 687 |
| -2,66929E | FALSE | Medium | IsMasterP | AAEL0093 | 1:2001231 | 14 | 3 | FALSE | 2 | 2 | 2 | 1141 | 1276 | 497 |
| -4,4594E+ | FALSE | Medium | IsMasterP | AAEL0047 | 2:2899571 | 14 | 14 | FALSE | 2 | 2 | 2 | 237 | 262 | 560 |
| -1,3579E+ | FALSE | Medium | IsMasterP | AAEL0005 | 3:3888780 | 14 | 3 | FALSE | 1 | 2 | 1 | 560 | 623 | 806 |
| 3,78821E+ | FALSE | Medium | IsMasterP | AAEL0169 | 2:2319313 | 14 | 11 | FALSE | 2 | 2 | 2 | 290 | 334 | 819 |
| 3,35761E+ | FALSE | Medium | IsMasterP | AAEL0089 | 3:3371054 | 14 | 1 | FALSE | 1 | 2 | 1 | 1322 | 1486 | 696 |
| -9,16839E | FALSE | Medium | IsMasterP | AAEL0025 | 1:9129755 | 14 | 9 | FALSE | 1 | 2 | 1 | 148 | 167 | 766 |
| 3,20615E+ | FALSE | Medium | IsMasterP | AAEL0135 | 2:3379136 | 14 | 7 | FALSE | 1 | 2 | 1 | 205 | 238 | 876 |
| 2,08112E+ | FALSE | Medium | IsMasterP | AAEL0044 | 3:2051944 | 14 | 2 | FALSE | 1 | 2 | 1 | 726 | 825 | 724 |
| -9,7828E+ | FALSE | Medium | IsMasterP | AAEL0077 | 3:2555106 | 14 | 1 | FALSE | 1 | 2 | 1 | 2039 | 2262 | 701 |
| 7,64197E+ | FALSE | Medium | IsMasterP | AAEL0122 | 2:3843805 | 14 | 6 | FALSE | 1 | 2 | 1 | 251 | 292 | 958 |
| -3,89004E | FALSE | Medium | IsMasterP | AAEL0004 | 3:1859072 | 14 | 3 | FALSE | 1 | 2 | 1 | 530 | 594 | 743 |
| 4,49915E+ | FALSE | Medium | IsMasterP | AAEL0132 | 1:3019742 | 14 | 6 | FALSE | 2 | 2 | 2 | 497 | 537 | 494 |
| -3,79191E | FALSE | Medium | IsMasterP | AAEL0025 | 1:2265246 | 14 | 4 | FALSE | 1 | 1 | 1 | 1038 | 1171 | 674 |
| 4,84688E+ | FALSE | Medium | IsMasterP | AAEL0002 | 3:2649885 | 14 | 14 | FALSE | 2 | 2 | 2 | 168 | 192 | 923 |
| -3,9931E+ | FALSE | Medium | IsMasterP | AAEL0001 | 1:1949484 | 14 | 2 | FALSE | 1 | 2 | 1 | 765 | 869 | 524 |
| -1,57635E | FALSE | Medium | IsMasterP | AAEL0040 | 3:1659686 | 14 | 4 | FALSE | 2 | 2 | 2 | 636 | 687 | 834 |
| 6,22548E+ | FALSE | Medium | IsMasterP | AAEL0106 | 3:2915235 | 14 | 3 | FALSE | 1 | 2 | 1 | 470 | 528 | 725 |
| -6,24449E | FALSE | Medium | IsMasterP | AAEL0098 | 3:1217507 | 14 | 8 | FALSE | 1 | 1 | 1 | 405 | 452 | 970 |
| -8,62495E | FALSE | Medium | IsMasterP | AAEL0256 | 3:2986982 | 14 | 8 | FALSE | 1 | 1 | 1 | 338 | 373 | 973 |
| 9,1488E+1 | FALSE | Medium | IsMasterP | AAEL0020 | 2:4265873 | 14 | 18 | FALSE | 2 | 2 | 2 | 208 | 238 | 881 |
| -7,51816E | FALSE | Medium | IsMasterP | AAEL0143 | 2:5149619 | 14 | 5 | FALSE | 2 | 2 | 2 | 531 | 599 | 506 |
| -2,47221E | FALSE | Medium | IsMasterP | AAEL0252 | 2:3320753 | 14 | 5 | FALSE | 2 | 2 | 2 | 557 | 614 | 705 |
| -3,51184E | FALSE | Medium | IsMasterP | AAEL0063 | 3:2882263 | 14 | 3 | FALSE | 1 | 2 | 1 | 480 | 546 | 525 |
| 7,93673E+ | FALSE | Medium | IsMasterP | AAEL0095 | 3:3450077 | 14 | 2 | FALSE | 1 | 2 | 1 | 675 | 754 | 660 |
| 5,96373E+ | FALSE | Medium | IsMasterP | AAEL0039 | 1:2505456 | 14 | 5 | FALSE | 2 | 2 | 2 | 437 | 498 | 856 |
| 6,16697E+ | FALSE | Medium | IsMasterP | AAEL0031 | 3:2695668 | 14 | 18 | FALSE | 2 | 2 | 2 | 203 | 228 | 1039 |

| | | | | | | | | | | | | | | |
|---|---|---|---|---|---|---|---|---|---|---|---|---|---|---|
| 8,55227E- | FALSE | Medium | IsMasterP | AAEL0090 | 3:1353919 | 14 | 4 | FALSE | 1 | 2 | 1 | 387 | 450 | 936 |
| -5,23631E | FALSE | Medium | IsMasterP | AAEL0143 | 2:2278572 | 14 | 8 | FALSE | 2 | 2 | 2 | 273 | 300 | 611 |
| -7,73307E | FALSE | Medium | IsMasterP | AAEL0234 | 1:2802106 | 14 | 9 | FALSE | 2 | 2 | 1 | 306 | 355 | 711 |
| 2,99802E- | FALSE | Medium | IsMasterP | AAEL0131 | 2:1885301 | 14 | 1 | FALSE | 2 | 2 | 2 | 1034 | 1170 | 470 |
| -7,25831E | FALSE | Medium | IsMasterP | AAEL0040 | 1:1354098 | 14 | 11 | FALSE | 1 | 2 | 1 | 176 | 197 | 837 |
| 8,38779E- | FALSE | Medium | IsMasterP | AAEL0183 | 3:1838599 | 14 | 1 | FALSE | 1 | 2 | 1 | 2423 | 2676 | 928 |
| 7,68983E- | FALSE | Medium | IsMasterP | AAEL0064 | 2:4139441 | 14 | 6 | FALSE | 1 | 1 | 1 | 376 | 426 | 620 |
| 6,86438E- | FALSE | Medium | IsMasterP | AAEL0046 | 1:1188342 | 14 | 26 | FALSE | 2 | 2 | 2 | 130 | 155 | 625 |
| -1,71201E | FALSE | Medium | IsMasterP | AAEL0147 | 1:2120362 | 14 | 10 | FALSE | 1 | 1 | 1 | 357 | 379 | 517 |
| -1,2072E+ | FALSE | Medium | IsMasterP | AAEL0088 | 3:3375974 | 14 | 33 | FALSE | 2 | 2 | 2 | 107 | 119 | 954 |
| 3,30129E- | FALSE | Medium | IsMasterP | AAEL0143 | 2:2384394 | 14 | 7 | FALSE | 1 | 1 | 1 | 485 | 556 | 526 |
| -2,20223E | FALSE | Medium | IsMasterP | AAEL0262 | NIGP0100 | 14 | 6 | FALSE | 2 | 2 | 2 | 583 | 661 | 870 |
| -6,18403E | FALSE | Medium | IsMasterP | AAEL0143 | 2:2386674 | 14 | 6 | FALSE | 1 | 2 | 1 | 356 | 397 | 737 |
| -2,61317E | FALSE | Medium | IsMasterP | AAEL0045 | 2:3101923 | 14 | 10 | FALSE | 2 | 2 | 2 | 345 | 377 | 802 |
| -5,70621E | FALSE | Medium | IsMasterP | AAEL0049 | 2:2818224 | 14 | 1 | FALSE | 1 | 2 | 1 | 907 | 1027 | 876 |
| 5,97887E- | FALSE | Medium | IsMasterP | AAEL0213 | 2:1218668 | 14 | 1 | FALSE | 2 | 2 | 2 | 3010 | 3356 | 856 |
| -3,52344E | FALSE | Medium | IsMasterP | AAEL0043 | 3:9311097 | 14 | 5 | FALSE | 2 | 2 | 2 | 556 | 628 | 794 |
| 5,20843E- | FALSE | Medium | IsMasterP | AAEL0027 | 3:1448855 | 14 | 1 | FALSE | 1 | 2 | 1 | 1207 | 1338 | 494 |
| 6,86988E- | FALSE | Medium | IsMasterP | AAEL0065 | 3:1511330 | 14 | 12 | FALSE | 1 | 1 | 1 | 225 | 265 | 505 |
| -3,84635E | FALSE | Medium | IsMasterP | AAEL0080 | 1:7425611 | 14 | 1 | FALSE | 1 | 2 | 1 | 1287 | 1483 | 709 |
| -5,82418E | FALSE | Medium | IsMasterP | AAEL0014 | 2:4578562 | 14 | 11 | FALSE | 2 | 2 | 2 | 191 | 217 | 558 |
| 2,78484E- | FALSE | Medium | IsMasterP | AAEL0113 | 1:1363446 | 14 | 4 | FALSE | 1 | 2 | 1 | 296 | 333 | 873 |
| 5,79565E- | FALSE | Medium | IsMasterP | AAEL0053 | 3:3084185 | 14 | 4 | FALSE | 1 | 2 | 1 | 349 | 396 | 951 |
| 6,8398E+1 | FALSE | Medium | IsMasterP | AAEL0013 | 2:4780170 | 14 | 4 | FALSE | 1 | 2 | 1 | 453 | 515 | 578 |
| 3,66521E- | FALSE | Medium | IsMasterP | AAEL0067 | 3:1690063 | 14 | 6 | FALSE | 1 | 2 | 1 | 186 | 211 | 822 |
| -1,67132E | FALSE | Medium | IsMasterP | AAEL0004 | 3:1851953 | 14 | 6 | FALSE | 1 | 2 | 1 | 220 | 254 | 506 |
| -8,15047E | FALSE | Medium | IsMasterP | AAEL0043 | 3:5115612 | 14 | 8 | FALSE | 1 | 1 | 1 | 465 | 550 | 897 |
| -1,01678E | FALSE | Medium | IsMasterP | AAEL0039 | 3:2608025 | 14 | 2 | FALSE | 1 | 2 | 1 | 569 | 639 | 870 |
| 6,06423E- | FALSE | Medium | IsMasterP | AAEL0255 | 1:3039310 | 14 | 3 | FALSE | 2 | 2 | 2 | 783 | 881 | 534 |
| 8,14398E- | FALSE | Medium | IsMasterP | AAEL0053 | 2:3698612 | 14 | 2 | FALSE | 1 | 2 | 1 | 607 | 701 | 827 |
| 3,9631E+1 | FALSE | Medium | IsMasterP | AAEL0086 | 2:4548071 | 14 | 4 | FALSE | 1 | 2 | 1 | 225 | 251 | 500 |

| | | | | | | | | | | | | | | |
|---|---|---|---|---|---|---|---|---|---|---|---|---|---|---|
| 7,95768E- | FALSE | Medium | IsMasterP | AAEL0276 | 2:234419 | 14 | 3 | FALSE | 2 | 2 | 2 | 917 | 1036 | 658 |
| -1,91989E | FALSE | Medium | IsMasterP | AAEL0197 | 3:329149 | 14 | 1 | FALSE | 1 | 2 | 1 | 929 | 1000 | 711 |
| 4,60834E- | FALSE | Medium | IsMasterP | AAEL0194 | 2:176218 | 14 | 1 | FALSE | 1 | 2 | 1 | 1179 | 1333 | 703 |
| -4,68292E | FALSE | Medium | IsMasterP | AAEL0121 | 1:437895 | 14 | 8 | FALSE | 1 | 1 | 1 | 434 | 484 | 991 |
| 5,85894E- | FALSE | Medium | IsMasterP | AAEL0138 | 3:303464 | 14 | 9 | FALSE | 1 | 2 | 1 | 158 | 171 | 587 |
| 6,07236E- | FALSE | Medium | IsMasterP | AAEL0057 | 1:201555 | 14 | 1 | FALSE | 1 | 2 | 1 | 821 | 951 | 1018 |
| -8,83979E | FALSE | Medium | IsMasterP | AAEL0088 | 2:241191 | 14 | 15 | FALSE | 1 | 1 | 1 | 191 | 219 | 483 |
| -4,43933E | FALSE | Medium | IsMasterP | AAEL0010 | 1:299609 | 14 | 2 | FALSE | 1 | 2 | 1 | 648 | 719 | 715 |
| -6,49582E | FALSE | Medium | IsMasterP | AAEL0018 | 2:374754 | 14 | 2 | FALSE | 1 | 2 | 1 | 517 | 598 | 873 |
| 4,78591E- | FALSE | Medium | IsMasterP | AAEL0046 | 3:248464 | 14 | 9 | FALSE | 2 | 2 | 2 | 364 | 397 | 904 |
| -8,47537E | FALSE | Medium | IsMasterP | AAEL0134 | 2:199136 | 14 | 12 | FALSE | 2 | 2 | 2 | 242 | 280 | 989 |
| -8,78465E | FALSE | Medium | IsMasterP | AAEL0104 | 1:204118 | 14 | 3 | FALSE | 1 | 2 | 1 | 369 | 415 | 574 |
| -2,66126E | FALSE | Medium | IsMasterP | AAEL0110 | 2:631918 | 14 | 1 | FALSE | 1 | 2 | 1 | 676 | 759 | 872 |
| 2,68108E- | FALSE | Medium | IsMasterP | AAEL0044 | 3:205249 | 14 | 7 | FALSE | 1 | 1 | 1 | 548 | 623 | 848 |
| 1,31326E- | FALSE | Medium | IsMasterP | AAEL0098 | 3:183667 | 15 | 1 | FALSE | 2 | 2 | 2 | 4613 | 5034 | 502 |
| -2,17184E | FALSE | Medium | IsMasterP | AAEL0181 | 3:105064 | 15 | 4 | FALSE | 1 | 1 | 1 | 589 | 654 | 702 |
| 8,07935E- | FALSE | Medium | IsMasterP | AAEL0025 | 3:121665 | 15 | 2 | FALSE | 2 | 2 | 2 | 1103 | 1272 | 891 |
| 4,86057E- | FALSE | Medium | IsMasterP | AAEL0077 | 2:219754 | 15 | 16 | FALSE | 1 | 1 | 1 | 269 | 300 | 903 |
| -5,431E+1 | FALSE | Medium | IsMasterP | AAEL0056 | 1:473607 | 15 | 4 | FALSE | 1 | 1 | 1 | 660 | 739 | 897 |
| -4,48937E | FALSE | Medium | IsMasterP | AAEL0256 | 3:335531 | 15 | 4 | FALSE | 1 | 1 | 1 | 864 | 1009 | 522 |
| -7,34126E | FALSE | Medium | IsMasterP | AAEL0196 | 2:117671 | 15 | 2 | FALSE | 1 | 1 | 1 | 2012 | 2252 | 825 |
| 6,27236E- | FALSE | Medium | IsMasterP | AAEL0201 | 2:115690 | 15 | 1 | FALSE | 1 | 2 | 1 | 804 | 871 | 983 |
| -1,85754E | FALSE | Medium | IsMasterP | AAEL0083 | 2:274075 | 15 | 10 | FALSE | 1 | 1 | 1 | 403 | 471 | 522 |
| 3,07498E- | FALSE | Medium | IsMasterP | AAEL0087 | 3:326023 | 15 | 2 | FALSE | 1 | 2 | 1 | 554 | 615 | 581 |
| 5,7883E+1 | FALSE | Medium | IsMasterP | AAEL0016 | 2:299392 | 15 | 3 | FALSE | 2 | 2 | 2 | 814 | 951 | 925 |
| 2,09879E- | FALSE | Medium | IsMasterP | AAEL0078 | 3:151685 | 15 | 3 | FALSE | 2 | 2 | 2 | 703 | 773 | 875 |
| -6,56695E | FALSE | Medium | IsMasterP | AAEL0218 | 3:327131 | 15 | 1 | FALSE | 2 | 2 | 2 | 1933 | 2152 | 859 |
| -2,59224E | FALSE | Medium | IsMasterP | AAEL0022 | 2:430692 | 15 | 11 | FALSE | 2 | 2 | 2 | 204 | 230 | 564 |
| -7,57834E | FALSE | Medium | IsMasterP | AAEL0103 | 3:372414 | 15 | 2 | FALSE | 1 | 1 | 1 | 2002 | 2146 | 535 |
| 2,7619E+1 | FALSE | Medium | IsMasterP | AAEL0194 | 2:399797 | 16 | 14 | FALSE | 2 | 2 | 2 | 338 | 355 | 999 |
| -1,09251E | FALSE | Medium | IsMasterP | AAEL0023 | 3:374492 | 16 | 8 | FALSE | 1 | 1 | 1 | 526 | 555 | 876 |

| | | | | | | | | | | | | | | |
|---|---|---|---|---|---|---|---|---|---|---|---|---|---|---|
| -6,97179E | FALSE | Medium | IsMasterP | AAEL0066 | 3:148173 | 16 | 13 | FALSE | 1 | 1 | 1 | 139 | 161 | 479 |
| 1,55173E | FALSE | Medium | IsMasterP | AAEL0274 | 2:197510 | 16 | 1 | FALSE | 1 | 2 | 1 | 1115 | 1249 | 456 |
| 3,97287E | FALSE | Medium | IsMasterP | AAEL0106 | 2:418102 | 16 | 1 | FALSE | 1 | 2 | 1 | 703 | 796 | 643 |
| 8,01555E | FALSE | Medium | IsMasterP | AAEL0124 | 1:213374 | 16 | 3 | FALSE | 1 | 2 | 1 | 294 | 339 | 578 |
| -1,35827E | FALSE | Medium | IsMasterP | AAEL0087 | 3:281117 | 16 | 4 | FALSE | 2 | 2 | 2 | 813 | 928 | 879 |
| -4,04061E | FALSE | Medium | IsMasterP | AAEL0128 | 2:335178 | 16 | 5 | FALSE | 2 | 2 | 2 | 438 | 483 | 821 |
| 3,03743E | FALSE | Medium | IsMasterP | AAEL0000 | 3:102992 | 17 | 6 | FALSE | 1 | 1 | 1 | 407 | 454 | 568 |
| -2,97488E | FALSE | Medium | IsMasterP | AAEL0103 | 3:372081 | 17 | 2 | FALSE | 1 | 2 | 1 | 667 | 758 | 832 |
| -2,71753E | FALSE | Medium | IsMasterP | AAEL0021 | 3:176807 | 17 | 1 | FALSE | 1 | 2 | 1 | 1552 | 1666 | 922 |
| -6,28803E | FALSE | Medium | IsMasterP | AAEL0017 | 2:441768 | 17 | 2 | FALSE | 1 | 2 | 1 | 653 | 721 | 677 |
| -4,11995E | FALSE | Medium | IsMasterP | AAEL0032 | 1:291624 | 17 | 2 | FALSE | 1 | 2 | 1 | 491 | 543 | 577 |
| -6,58252E | FALSE | Medium | IsMasterP | AAEL0032 | 3:340157 | 17 | 5 | FALSE | 2 | 2 | 2 | 673 | 767 | 950 |
| -3,26022E | FALSE | Medium | IsMasterP | AAEL0068 | 3:571062 | 17 | 3 | FALSE | 2 | 2 | 2 | 914 | 1033 | 667 |
| 7,27495E | FALSE | Medium | IsMasterP | AAEL0064 | 2:112889 | 17 | 10 | FALSE | 2 | 2 | 2 | 273 | 311 | 936 |
| -6,8991E+ | FALSE | Medium | IsMasterP | AAEL0215 | 2:126633 | 17 | 4 | FALSE | 1 | 2 | 1 | 265 | 299 | 610 |
| -2,57526E | FALSE | Medium | IsMasterP | AAEL0007 | 2:520897 | 17 | 11 | FALSE | 1 | 2 | 1 | 84 | 102 | 844 |
| -8,57988E | FALSE | Medium | IsMasterP | AAEL0226 | 1:107574 | 17 | 11 | FALSE | 1 | 2 | 1 | 123 | 123 | 591 |
| -3,9355E+ | FALSE | Medium | IsMasterP | AAEL0102 | 3:279006 | 17 | 20 | FALSE | 1 | 1 | 1 | 148 | 168 | 496 |
| -7,54091E | FALSE | Medium | IsMasterP | AAEL0058 | 3:220252 | 17 | 8 | FALSE | 2 | 2 | 2 | 327 | 366 | 714 |
| -1,97382E | FALSE | Medium | IsMasterP | AAEL0035 | 3:180078 | 17 | 1 | FALSE | 1 | 2 | 1 | 943 | 1052 | 487 |
| 7,81169E | FALSE | Medium | IsMasterP | AAEL0053 | 3:310036 | 17 | 2 | FALSE | 1 | 2 | 1 | 829 | 923 | 679 |
| 8,32525E | FALSE | Medium | IsMasterP | AAEL0020 | 2:426390 | 17 | 7 | FALSE | 2 | 2 | 2 | 355 | 410 | 478 |
| -3,53496E | FALSE | Medium | IsMasterP | AAEL0137 | 3:234309 | 17 | 10 | FALSE | 1 | 1 | 1 | 362 | 388 | 643 |
| 4,74666E | FALSE | Medium | IsMasterP | AAEL0112 | 1:150733 | 17 | 2 | FALSE | 1 | 1 | 1 | 1222 | 1321 | 693 |
| 3,07422E | FALSE | Medium | IsMasterP | AAEL0080 | 2:230736 | 17 | 8 | FALSE | 1 | 1 | 1 | 419 | 475 | 913 |
| -1,01532E | FALSE | Medium | IsMasterP | AAEL0107 | 2:406517 | 17 | 1 | FALSE | 1 | 2 | 1 | 893 | 1028 | 683 |
| 4,93302E | FALSE | Medium | IsMasterP | AAEL0253 | NIGP0100 | 17 | 16 | FALSE | 1 | 2 | 1 | 156 | 186 | 854 |
| 5,22463E | FALSE | Medium | IsMasterP | AAEL0022 | 2:295577 | 17 | 4 | FALSE | 2 | 2 | 2 | 555 | 635 | 882 |
| -4,72346E | FALSE | Medium | IsMasterP | AAEL0065 | 2:467986 | 17 | 4 | FALSE | 2 | 2 | 2 | 677 | 765 | 681 |
| 6,85856E | FALSE | Medium | IsMasterP | AAEL0092 | 2:174641 | 17 | 7 | FALSE | 1 | 2 | 1 | 171 | 199 | 658 |
| -3,68103E | FALSE | Medium | IsMasterP | AAEL0250 | 1:491341 | 18 | 1 | FALSE | 1 | 2 | 1 | 979 | 1131 | 642 |

| | | | | | | | | | | | | | | |
|---|---|---|---|---|---|---|---|---|---|---|---|---|---|---|
| 8,86419E- | FALSE | Medium | IsMasterP | AAEL0067 | 1:167615 | 18 | 10 | FALSE | 1 | 1 | 1 | 246 | 283 | 892 |
| 2,60219E- | FALSE | Medium | IsMasterP | AAEL0108 | 1:218355 | 18 | 2 | FALSE | 1 | 2 | 1 | 509 | 571 | 720 |
| 6,27916E- | FALSE | Medium | IsMasterP | AAEL0103 | 3:409333 | 18 | 5 | FALSE | 1 | 1 | 1 | 548 | 617 | 785 |
| -8,99312E | FALSE | Medium | IsMasterP | AAEL0127 | 2:941770 | 18 | 3 | FALSE | 2 | 2 | 2 | 1636 | 1833 | 660 |
| -1,81817E | FALSE | Medium | IsMasterP | AAEL0129 | 3:297140 | 18 | 5 | FALSE | 2 | 2 | 2 | 490 | 526 | 502 |
| -2,02535E | FALSE | Medium | IsMasterP | AAEL0113 | 2:926528 | 18 | 4 | FALSE | 1 | 1 | 1 | 955 | 1069 | 709 |
| 1,86186E- | FALSE | Medium | IsMasterP | AAEL0107 | 3:294215 | 18 | 12 | FALSE | 2 | 2 | 2 | 239 | 276 | 838 |
| 5,69913E- | FALSE | Medium | IsMasterP | AAEL0182 | 2:762626 | 18 | 1 | FALSE | 2 | 2 | 2 | 2577 | 2946 | 824 |
| 4,29513E- | FALSE | Medium | IsMasterP | AAEL0086 | 2:238740 | 18 | 3 | FALSE | 1 | 2 | 1 | 362 | 410 | 787 |
| 8,27968E- | FALSE | Medium | IsMasterP | AAEL0264 | 3:405618 | 18 | 4 | FALSE | 1 | 1 | 1 | 657 | 742 | 637 |
| 2,55229E- | FALSE | Medium | IsMasterP | AAEL0264 | 2:129984 | 18 | 4 | FALSE | 1 | 1 | 1 | 843 | 948 | 935 |
| -6,92429E | FALSE | Medium | IsMasterP | AAEL0060 | 3:315542 | 18 | 4 | FALSE | 2 | 2 | 2 | 526 | 589 | 508 |
| 1,18793E- | FALSE | Medium | IsMasterP | AAEL0085 | 2:362426 | 18 | 3 | FALSE | 1 | 2 | 1 | 323 | 367 | 692 |
| -1,46087E | FALSE | Medium | IsMasterP | AAEL0195 | 2:552626 | 18 | 7 | FALSE | 1 | 1 | 1 | 391 | 433 | 942 |
| -2,09901E | FALSE | Medium | IsMasterP | AAEL0205 | 2:176299 | 18 | 2 | FALSE | 1 | 2 | 1 | 618 | 687 | 477 |
| -3,44763E | FALSE | Medium | IsMasterP | AAEL0248 | 2:149882 | 18 | 0 | FALSE | 1 | 2 | 1 | 2232 | 2416 | 756 |
| -7,92222E | FALSE | Medium | IsMasterP | AAEL0024 | 1:176815 | 18 | 8 | FALSE | 1 | 1 | 1 | 319 | 346 | 477 |
| -5,44285E | FALSE | Medium | IsMasterP | AAEL0073 | 1:259841 | 18 | 3 | FALSE | 1 | 1 | 1 | 709 | 798 | 555 |
| -7,42804E | FALSE | Medium | IsMasterP | AAEL0269 | 2:232418 | 18 | 7 | FALSE | 1 | 2 | 1 | 381 | 424 | 547 |
| -6,15545E | FALSE | Medium | IsMasterP | AAEL0270 | 1:303668 | 18 | 5 | FALSE | 1 | 1 | 1 | 791 | 882 | 1008 |
| -3,68375E | FALSE | Medium | IsMasterP | AAEL0019 | 2:389906 | 18 | 6 | FALSE | 2 | 2 | 2 | 304 | 351 | 799 |
| 1,37404E- | FALSE | Medium | IsMasterP | AAEL0203 | 2:147844 | 18 | 2 | FALSE | 1 | 2 | 1 | 957 | 1058 | 693 |
| 1,66765E- | FALSE | Medium | IsMasterP | AAEL0273 | 1:348850 | 18 | 16 | FALSE | 1 | 1 | 1 | 191 | 199 | 720 |
| 4,18705E- | FALSE | Medium | IsMasterP | AAEL0023 | 2:303124 | 18 | 2 | FALSE | 2 | 2 | 2 | 1289 | 1385 | 731 |
| -2,01836E | FALSE | Medium | IsMasterP | AAEL0140 | 2:342909 | 18 | 6 | FALSE | 2 | 2 | 2 | 360 | 415 | 664 |
| -7,74702E | FALSE | Medium | IsMasterP | AAEL0127 | 1:184967 | 19 | 4 | FALSE | 1 | 1 | 1 | 1285 | 1390 | 885 |
| 1,38861E- | FALSE | Medium | IsMasterP | AAEL0146 | 2:157521 | 19 | 5 | FALSE | 1 | 2 | 1 | 274 | 319 | 848 |
| 1,63182E- | FALSE | Medium | IsMasterP | AAEL0262 | 1:605186 | 19 | 1 | FALSE | 1 | 1 | 1 | 2529 | 2878 | 627 |
| -8,26127E | FALSE | Medium | IsMasterP | AAEL0011 | 3:121366 | 19 | 7 | FALSE | 1 | 1 | 1 | 356 | 413 | 764 |
| -7,65924E | FALSE | Medium | IsMasterP | AAEL0181 | 1:127465 | 19 | 1 | FALSE | 1 | 2 | 1 | 708 | 788 | 686 |
| 4,60321E- | FALSE | Medium | IsMasterP | AAEL0266 | 2:324919 | 19 | 1 | FALSE | 1 | 2 | 1 | 1368 | 1481 | 667 |

| | | | | | | | | | | | | | | |
|---|---|---|---|---|---|---|---|---|---|---|---|---|---|---|
| -8,535E+1 | FALSE | Medium | IsMasterP | AAEL0063 | 1:101487[illegible] | 19 | 4 | FALSE | 1 | 2 | 1 | 397 | 426 | 740 |
| 7,20344E- | FALSE | Medium | IsMasterP | AAEL0015 | 2:222104[illegible] | 19 | 3 | FALSE | 1 | 1 | 1 | 863 | 926 | 498 |
| 7,06681E- | FALSE | Medium | IsMasterP | AAEL0149 | 2:145127[illegible] | 19 | 2 | FALSE | 2 | 2 | 2 | 1308 | 1434 | 679 |
| -8,9191E+ | FALSE | Medium | IsMasterP | AAEL0073 | 2:233829[illegible] | 19 | 2 | FALSE | 1 | 2 | 1 | 525 | 607 | 788 |
| -8,95988E | FALSE | Medium | IsMasterP | AAEL0250 | 2:165692[illegible] | 19 | 2 | FALSE | 1 | 2 | 1 | 518 | 594 | 563 |
| 7,77815E- | FALSE | Medium | IsMasterP | AAEL0246 | 2:118212[illegible] | 19 | 13 | FALSE | 1 | 1 | 1 | 204 | 230 | 474 |
| -7,57038E | FALSE | Medium | IsMasterP | AAEL0116 | 3:214340[illegible] | 19 | 1 | FALSE | 1 | 2 | 1 | 621 | 713 | 660 |
| -4,70266E | FALSE | Medium | IsMasterP | AAEL0266 | 3:121344[illegible] | 19 | 13 | FALSE | 1 | 1 | 1 | 255 | 300 | 778 |
| 2,42768E- | FALSE | Medium | IsMasterP | AAEL0067 | 2:836582[illegible] | 19 | 15 | FALSE | 1 | 1 | 1 | 156 | 177 | 525 |
| 3,77878E- | FALSE | Medium | IsMasterP | AAEL0054 | 2:295787[illegible] | 19 | 6 | FALSE | 2 | 2 | 2 | 329 | 373 | 991 |
| 8,30313E- | FALSE | Medium | IsMasterP | AAEL0105 | 1:509434[illegible] | 19 | 1 | FALSE | 1 | 2 | 1 | 841 | 956 | 802 |
| 6,77009E- | FALSE | Medium | IsMasterP | AAEL0266 | 2:112666[illegible] | 19 | 4 | FALSE | 2 | 2 | 2 | 552 | 621 | 567 |
| -2,10333E | FALSE | Medium | IsMasterP | AAEL0130 | 3:310922[illegible] | 19 | 4 | FALSE | 1 | 2 | 1 | 484 | 545 | 717 |
| -5,61392E | FALSE | Medium | IsMasterP | AAEL0028 | 3:234897[illegible] | 19 | 2 | FALSE | 1 | 2 | 1 | 480 | 555 | 837 |
| -4,15936E | FALSE | Medium | IsMasterP | AAEL0016 | 2:297955[illegible] | 19 | 4 | FALSE | 1 | 1 | 1 | 537 | 590 | 629 |
| -7,97011E | FALSE | Medium | IsMasterP | gb:AVW8[illegible] | ncbiId:AV[illegible] | 19 | 1 | FALSE | 1 | 1 | 1 | 3419 | 3784 | 844 |
| 8,85303E- | FALSE | Medium | IsMasterP | AAEL0002 | 1:560141[illegible] | 19 | 3 | FALSE | 1 | 2 | 1 | 394 | 460 | 872 |
| 5,82107E- | FALSE | Medium | IsMasterP | AAEL0150 | 2:233065[illegible] | 19 | 2 | FALSE | 1 | 1 | 1 | 681 | 747 | 708 |
| -9,02652E | FALSE | Medium | IsMasterP | AAEL0175 | 1:135458[illegible] | 19 | 3 | FALSE | 1 | 2 | 1 | 295 | 316 | 756 |
| -7,49708E | FALSE | Medium | IsMasterP | AAEL0198 | 2:152558[illegible] | 19 | 10 | FALSE | 2 | 2 | 2 | 264 | 290 | 925 |
| -6,42648E | FALSE | Medium | IsMasterP | AAEL0056 | 2:131999[illegible] | 19 | 5 | FALSE | 2 | 2 | 2 | 490 | 560 | 564 |
| -4,19281E | FALSE | Medium | IsMasterP | AAEL0049 | 3:323165[illegible] | 19 | 6 | FALSE | 1 | 2 | 1 | 279 | 307 | 544 |
| 7,79041E- | FALSE | Medium | IsMasterP | AAEL0012 | 1:597071[illegible] | 19 | 3 | FALSE | 1 | 2 | 1 | 318 | 336 | 903 |
| 2,27236E- | FALSE | Medium | IsMasterP | AAEL0101 | 2:264726[illegible] | 19 | 3 | FALSE | 1 | 1 | 1 | 763 | 855 | 627 |
| -2,43584E | FALSE | Medium | IsMasterP | AAEL0081 | 3:302160[illegible] | 19 | 14 | FALSE | 1 | 2 | 1 | 63 | 75 | 705 |
| -7,2443E+ | FALSE | Medium | IsMasterP | AAEL0103 | 3:409181[illegible] | 19 | 5 | FALSE | 1 | 1 | 1 | 348 | 402 | 859 |
| 2,93253E- | FALSE | Medium | IsMasterP | AAEL0093 | 3:755303[illegible] | 19 | 2 | FALSE | 1 | 1 | 1 | 1519 | 1690 | 821 |
| -2,84305E | FALSE | Medium | IsMasterP | AAEL0267 | 3:405043[illegible] | 19 | 2 | FALSE | 2 | 2 | 2 | 1288 | 1448 | 587 |
| -3,37745E | FALSE | Medium | IsMasterP | AAEL0083 | 2:273030[illegible] | 19 | 3 | FALSE | 1 | 2 | 1 | 292 | 311 | 701 |
| 4,3181E+[illegible] | FALSE | Medium | IsMasterP | AAEL0093 | 3:851699[illegible] | 19 | 29 | FALSE | 1 | 1 | 1 | 65 | 78 | 1054 |
| 5,9511E+[illegible] | FALSE | Medium | IsMasterP | AAEL0015 | 2:262289[illegible] | 19 | 5 | FALSE | 1 | 1 | 1 | 688 | 795 | 479 |

| | | | | | | | | | | | | | | |
|---|---|---|---|---|---|---|---|---|---|---|---|---|---|---|
| 2,82007E | FALSE | Medium | IsMasterP | AAEL0087 | 3:280355 | 19 | 1 | FALSE | 1 | 2 | 1 | 1294 | 1443 | 737 |
| -5,71467E | FALSE | Medium | IsMasterP | AAEL0035 | 3:180145 | 19 | 2 | FALSE | 1 | 1 | 1 | 944 | 1065 | 545 |
| 1,53248E | FALSE | Medium | IsMasterP | AAEL0021 | 1:251791 | 19 | 9 | FALSE | 2 | 2 | 2 | 222 | 253 | 614 |
| -9,43377E | FALSE | Medium | IsMasterP | AAEL0131 | 3:304145 | 19 | 3 | FALSE | 1 | 1 | 1 | 1281 | 1458 | 695 |
| 4,84145E | FALSE | Medium | IsMasterP | AAEL0018 | 2:374502 | 19 | 9 | FALSE | 2 | 2 | 2 | 338 | 369 | 903 |
| 1,72047E | FALSE | Medium | IsMasterP | AAEL0081 | 3:108004 | 19 | 2 | FALSE | 1 | 1 | 1 | 899 | 1007 | 547 |
| -3,80686E | FALSE | Medium | IsMasterP | AAEL0025 | 1:893228 | 19 | 15 | FALSE | 1 | 2 | 1 | 115 | 131 | 872 |
| 1,35696E | FALSE | Medium | IsMasterP | AAEL0049 | 2:281384 | 19 | 3 | FALSE | 1 | 2 | 1 | 286 | 320 | 856 |
| 5,73876E | FALSE | Medium | IsMasterP | AAEL0110 | 2:277911 | 19 | 5 | FALSE | 1 | 2 | 1 | 236 | 270 | 639 |
| 3,68396E | FALSE | Medium | IsMasterP | AAEL0149 | 2:172034 | 19 | 7 | FALSE | 1 | 2 | 1 | 342 | 396 | 919 |
| -1,56524E | FALSE | Medium | IsMasterP | AAEL0012 | 2:413375 | 19 | 2 | FALSE | 1 | 2 | 1 | 722 | 820 | 567 |
| 2,98286E | FALSE | Medium | IsMasterP | AAEL0102 | 1:358183 | 19 | 4 | FALSE | 1 | 1 | 1 | 550 | 608 | 585 |
| 1,77632E | FALSE | Medium | IsMasterP | AAEL0181 | 2:282083 | 19 | 0 | FALSE | 1 | 1 | 1 | 4316 | 4936 | 599 |
| 3,67618E | FALSE | Medium | IsMasterP | AAEL0006 | 3:395878 | 19 | 2 | FALSE | 1 | 2 | 1 | 607 | 700 | 731 |
| 1,9978E+ | FALSE | Medium | IsMasterP | AAEL0096 | 2:169099 | 19 | 8 | FALSE | 1 | 1 | 1 | 374 | 428 | 736 |
| 2,16382E | FALSE | Medium | IsMasterP | AAEL0173 | 1:193872 | 19 | 3 | FALSE | 1 | 1 | 1 | 1021 | 1101 | 581 |
| -4,20516E | FALSE | Medium | IsMasterP | AAEL0149 | 2:222841 | 19 | 5 | FALSE | 1 | 1 | 1 | 470 | 523 | 586 |
| 8,86535E | FALSE | Medium | IsMasterP | AAEL0091 | 1:156511 | 20 | 6 | FALSE | 1 | 1 | 1 | 415 | 473 | 683 |
| 7,88093E | FALSE | Medium | IsMasterP | AAEL0265 | 2:416592 | 20 | 5 | FALSE | 1 | 2 | 1 | 287 | 324 | 458 |
| 6,9422E+ | FALSE | Medium | IsMasterP | AAEL0095 | 1:140867 | 20 | 2 | FALSE | 2 | 2 | 2 | 1706 | 1816 | 914 |
| 6,25609E | FALSE | Medium | IsMasterP | AAEL0125 | 2:322751 | 20 | 4 | FALSE | 1 | 1 | 1 | 624 | 688 | 761 |
| 5,96951E | FALSE | Medium | IsMasterP | AAEL0141 | 2:305693 | 20 | 3 | FALSE | 1 | 2 | 1 | 360 | 397 | 702 |
| 5,75841E | FALSE | Medium | IsMasterP | AAEL0021 | 1:253581 | 20 | 9 | FALSE | 1 | 1 | 1 | 263 | 292 | 569 |
| 5,32681E | FALSE | Medium | IsMasterP | AAEL0057 | 1:201861 | 20 | 1 | FALSE | 1 | 2 | 1 | 2193 | 2563 | 1045 |
| 1,08635E | FALSE | Medium | IsMasterP | AAEL0090 | 2:269333 | 20 | 3 | FALSE | 2 | 2 | 2 | 993 | 1130 | 803 |
| 4,46244E | FALSE | Medium | IsMasterP | AAEL0231 | NIGP0100 | 20 | 7 | FALSE | 1 | 2 | 1 | 148 | 167 | 976 |
| 2,87809E | FALSE | Medium | IsMasterP | AAEL0135 | 1:144005 | 20 | 12 | FALSE | 1 | 1 | 1 | 154 | 172 | 972 |
| 3,041E+1 | FALSE | Medium | IsMasterP | AAEL0222 | 1:284443 | 21 | 2 | FALSE | 1 | 2 | 1 | 395 | 391 | 928 |
| 6,35182E | FALSE | Medium | IsMasterP | AAEL0196 | 2:291672 | 21 | 3 | FALSE | 1 | 1 | 1 | 1460 | 1537 | 709 |
| 8,98327E | FALSE | Medium | IsMasterP | AAEL0064 | 3:216330 | 21 | 19 | FALSE | 1 | 1 | 1 | 124 | 137 | 600 |
| -8,28032E | FALSE | Medium | IsMasterP | AAEL0097 | 1:209892 | 21 | 6 | FALSE | 1 | 1 | 1 | 499 | 577 | 723 |

| | | | | | | | | | | | | | | |
|---|---|---|---|---|---|---|---|---|---|---|---|---|---|---|
| 2,12503E- | FALSE | Medium | IsMasterP | AAEL0255 | 1:210637 | 21 | 4 | FALSE | 2 | 2 | 2 | 617 | 715 | 501 |
| 3,12081E- | FALSE | Medium | IsMasterP | AAEL0035 | 2:208744 | 21 | 20 | FALSE | 1 | 1 | 1 | 131 | 149 | 434 |
| 3,62304E- | FALSE | Medium | IsMasterP | AAEL0078 | 2:460795 | 21 | 2 | FALSE | 1 | 1 | 1 | 1168 | 1322 | 747 |
| 3,6807E+ | FALSE | Medium | IsMasterP | AAEL0095 | 3:119568 | 21 | 6 | FALSE | 1 | 2 | 1 | 215 | 243 | 769 |
| 4,76742E- | FALSE | Medium | IsMasterP | AAEL0243 | 1:551536 | 21 | 3 | FALSE | 1 | 1 | 1 | 530 | 561 | 699 |
| 7,75247E- | FALSE | Medium | IsMasterP | AAEL0062 | 2:491032 | 21 | 1 | FALSE | 2 | 2 | 2 | 1470 | 1658 | 677 |
| 9,07095E- | FALSE | Medium | IsMasterP | AAEL0048 | 2:722271 | 21 | 3 | FALSE | 2 | 2 | 2 | 788 | 883 | 635 |
| 8,47262E- | FALSE | Medium | IsMasterP | AAEL0248 | NIGP0100 | 21 | 22 | FALSE | 1 | 1 | 1 | 82 | 96 | 827 |
| -3,33639E | FALSE | Medium | IsMasterP | AAEL0111 | 3:381672 | 22 | 3 | FALSE | 1 | 2 | 1 | 392 | 438 | 699 |
| 3,02908E- | FALSE | Medium | IsMasterP | AAEL0079 | 2:474372 | 22 | 2 | FALSE | 1 | 1 | 1 | 1647 | 1847 | 824 |
| 3,27736E- | FALSE | Medium | IsMasterP | AAEL0132 | 2:399347 | 22 | 1 | FALSE | 1 | 2 | 1 | 1058 | 1177 | 633 |
| -8,35622E | FALSE | Medium | IsMasterP | AAEL0029 | 1:107900 | 22 | 1 | FALSE | 1 | 2 | 1 | 805 | 910 | 796 |
| -5,67916E | FALSE | Medium | IsMasterP | AAEL0229 | 2:342451 | 22 | 4 | FALSE | 1 | 1 | 1 | 398 | 465 | 995 |
| -4,84306E | FALSE | Medium | IsMasterP | AAEL0077 | 1:268902 | 22 | 2 | FALSE | 1 | 2 | 1 | 404 | 442 | 860 |
| -3,10042E | FALSE | Medium | IsMasterP | AAEL0025 | 1:226240 | 22 | 2 | FALSE | 1 | 1 | 1 | 686 | 779 | 511 |
| -5,34351E | FALSE | Medium | IsMasterP | AAEL0092 | 2:112778 | 22 | 7 | FALSE | 1 | 1 | 1 | 500 | 569 | 699 |
| 6,51405E- | FALSE | Medium | IsMasterP | AAEL0120 | 3:160204 | 22 | 6 | FALSE | 1 | 1 | 1 | 267 | 318 | 966 |
| 5,03489E- | FALSE | Medium | IsMasterP | AAEL0217 | 1:242876 | 22 | 3 | FALSE | 1 | 1 | 1 | 1122 | 1201 | 957 |
| 1,70666E- | FALSE | Medium | IsMasterP | AAEL0002 | 3:212976 | 22 | 15 | FALSE | 1 | 1 | 1 | 206 | 234 | 634 |
| -3,31171E | FALSE | Medium | IsMasterP | AAEL0097 | 3:284403 | 22 | 6 | FALSE | 1 | 1 | 1 | 530 | 576 | 819 |
| -8,05674E | FALSE | Medium | IsMasterP | AAEL0110 | 3:294329 | 22 | 7 | FALSE | 1 | 1 | 1 | 606 | 674 | 761 |
| -6,97857E | FALSE | Medium | IsMasterP | AAEL0063 | 3:396831 | 22 | 9 | FALSE | 1 | 1 | 1 | 304 | 354 | 718 |
| -1,93163E | FALSE | Medium | IsMasterP | AAEL0029 | 1:262412 | 22 | 6 | FALSE | 1 | 2 | 1 | 290 | 292 | 922 |
| -7,22599E | FALSE | Medium | IsMasterP | AAEL0145 | 2:332339 | 22 | 5 | FALSE | 1 | 2 | 1 | 175 | 201 | 677 |
| 4,91942E- | FALSE | Medium | IsMasterP | AAEL0019 | 2:425323 | 23 | 4 | FALSE | 1 | 2 | 1 | 253 | 285 | 642 |
| 8,46896E- | FALSE | Medium | IsMasterP | AAEL0062 | 3:149722 | 23 | 3 | FALSE | 1 | 1 | 1 | 1331 | 1439 | 568 |
| 1,71436E- | FALSE | Medium | IsMasterP | AAEL0012 | 2:413589 | 23 | 0 | FALSE | 1 | 2 | 1 | 2168 | 2506 | 890 |
| -3,41822E | FALSE | Medium | IsMasterP | AAEL0244 | 1:222213 | 23 | 3 | FALSE | 1 | 1 | 1 | 666 | 757 | 708 |
| -2,32025E | FALSE | Medium | IsMasterP | AAEL0052 | 3:888938 | 23 | 12 | FALSE | 1 | 1 | 1 | 208 | 235 | 507 |
| 1,39136E- | FALSE | Medium | IsMasterP | AAEL0197 | 3:878602 | 24 | 1 | FALSE | 2 | 2 | 2 | 2718 | 2983 | 869 |
| 2,08357E- | FALSE | Medium | IsMasterP | AAEL0236 | NIGP0100 | 24 | 1 | FALSE | 1 | 2 | 1 | 2102 | 2375 | 562 |

| | | | | | | | | | | | | | | |
|---|---|---|---|---|---|---|---|---|---|---|---|---|---|---|
| -5,04948E | FALSE | Medium | IsMasterP | AAEL0093 | 3:297773( | 24 | 3 | FALSE | 1 | 1 | 1 | 1358 | 1498 | 967 |
| 5,02292E- | FALSE | Medium | IsMasterP | AAEL0199 | 1:455990! | 24 | 0 | FALSE | 1 | 2 | 1 | 2937 | 3358 | 516 |
| -2,74192E | FALSE | Medium | IsMasterP | AAEL0085 | 2:362883: | 24 | 2 | FALSE | 1 | 1 | 1 | 1527 | 1688 | 642 |
| 9,03955E- | FALSE | Medium | IsMasterP | AAEL0015 | 2:208042 | 24 | 2 | FALSE | 1 | 2 | 1 | 516 | 579 | 925 |
| -4,13482E | FALSE | Medium | IsMasterP | AAEL0003 | 3:400031! | 24 | 7 | FALSE | 1 | 1 | 1 | 253 | 292 | 964 |
| -7,31716E | FALSE | Medium | IsMasterP | AAEL0054 | 3:599654! | 24 | 1 | FALSE | 1 | 2 | 1 | 694 | 775 | 692 |
| 8,5457E+: | FALSE | Medium | IsMasterP | AAEL0072 | 3:196397! | 24 | 4 | FALSE | 1 | 1 | 1 | 670 | 748 | 670 |
| -3,40645E | FALSE | Medium | IsMasterP | AAEL0119 | 3:452814: | 24 | 2 | FALSE | 1 | 1 | 1 | 1395 | 1545 | 881 |
| 2,52928E- | FALSE | Medium | IsMasterP | AAEL0011 | 2:309093( | 24 | 3 | FALSE | 1 | 2 | 1 | 553 | 636 | 932 |
| -9,06943E | FALSE | Medium | IsMasterP | AAEL0078 | 1:149109 | 24 | 5 | FALSE | 1 | 2 | 1 | 167 | 185 | 496 |
| -7,84357E | FALSE | Medium | IsMasterP | AAEL0098 | 3:122147: | 24 | 4 | FALSE | 1 | 1 | 1 | 985 | 1096 | 665 |
| 1,37384E- | FALSE | Medium | IsMasterP | AAEL0194 | 3:385078 | 24 | 6 | FALSE | 1 | 1 | 1 | 473 | 515 | 526 |
| 7,65838E- | FALSE | Medium | IsMasterP | AAEL0098 | 3:122448: | 25 | 1 | FALSE | 2 | 2 | 2 | 4792 | 5507 | 585 |
| -7,88355E | FALSE | Medium | IsMasterP | AAEL0135 | 1:144295: | 25 | 12 | FALSE | 1 | 1 | 1 | 213 | 239 | 873 |
| -1,57698E | FALSE | Medium | IsMasterP | AAEL0008 | 3:363965 | 25 | 8 | FALSE | 1 | 1 | 1 | 288 | 322 | 607 |
| 6,24413E- | FALSE | Medium | IsMasterP | AAEL0233 | 2:378244( | 25 | 7 | FALSE | 1 | 1 | 1 | 626 | 728 | 717 |
| 1,18111E- | FALSE | Medium | IsMasterP | AAEL0044 | 3:110080! | 25 | 3 | FALSE | 1 | 2 | 1 | 299 | 340 | 519 |
| 4,00538E- | FALSE | Medium | IsMasterP | AAEL0010 | 3:555574! | 25 | 1 | FALSE | 1 | 2 | 1 | 1269 | 1448 | 613 |
| -4,1368E+ | FALSE | Medium | IsMasterP | AAEL0114 | 2:336007 | 25 | 9 | FALSE | 1 | 1 | 1 | 324 | 363 | 793 |
| 3,44612E- | FALSE | Medium | IsMasterP | AAEL0085 | 2:395278! | 25 | 6 | FALSE | 1 | 1 | 1 | 633 | 697 | 541 |
| 1,63985E- | FALSE | Medium | IsMasterP | AAEL0018 | 2:375516( | 26 | 14 | FALSE | 1 | 1 | 1 | 133 | 147 | 597 |
| -4,51565E | FALSE | Medium | IsMasterP | AAEL0123 | 2:111774! | 26 | 2 | FALSE | 1 | 2 | 1 | 478 | 533 | 530 |
| 6,74085E- | FALSE | Medium | IsMasterP | AAEL0063 | 2:141052 | 26 | 9 | FALSE | 2 | 2 | 2 | 361 | 406 | 777 |
| 8,20376E- | FALSE | Medium | IsMasterP | AAEL0150 | 2:337750! | 27 | 8 | FALSE | 1 | 1 | 1 | 303 | 340 | 625 |
| -9,12408E | FALSE | Medium | IsMasterP | AAEL0120 | 1:209063: | 27 | 1 | FALSE | 1 | 2 | 1 | 610 | 675 | 856 |
| -9,01992E | FALSE | Medium | IsMasterP | AAEL0039 | 3:261487! | 28 | 4 | FALSE | 1 | 1 | 1 | 427 | 475 | 639 |
| 6,0336E+: | FALSE | Medium | IsMasterP | AAEL0131 | 2:373776! | 28 | 6 | FALSE | 1 | 1 | 1 | 542 | 589 | 634 |
| 1,97071E- | FALSE | Medium | IsMasterP | AAEL0082 | 3:249899: | 28 | 4 | FALSE | 1 | 1 | 1 | 995 | 1105 | 605 |
| -5,41851E | FALSE | Medium | IsMasterP | AAEL0066 | 3:352856( | 28 | 1 | FALSE | 1 | 1 | 1 | 1139 | 1304 | 526 |
| -1,32605E | FALSE | Medium | IsMasterP | AAEL0138 | 1:222143( | 28 | 13 | FALSE | 1 | 1 | 1 | 238 | 269 | 686 |
| -6,95862E | FALSE | Medium | IsMasterP | AAEL0025 | 3:257580( | 28 | 3 | FALSE | 1 | 1 | 1 | 1283 | 1453 | 657 |

| | | | | | | | | | | | | | | |
|---|---|---|---|---|---|---|---|---|---|---|---|---|---|---|
| -8,32782E | FALSE | Medium | IsMasterP | AAEL0044 | 2:4522259 | 29 | 8 | FALSE | 1 | 1 | 1 | 446 | 492 | 933 |
| -5,87172E | FALSE | Medium | IsMasterP | AAEL0140 | 1:1429868 | 29 | 13 | FALSE | 1 | 2 | 1 | 76 | 91 | 985 |
| -2,42596E | FALSE | Medium | IsMasterP | AAEL0028 | 1:1086805 | 29 | 11 | FALSE | 1 | 1 | 1 | 209 | 240 | 856 |
| -1,83585E | FALSE | Medium | IsMasterP | AAEL0016 | 2:3000540 | 30 | 3 | FALSE | 1 | 1 | 1 | 1165 | 1268 | 512 |
| -5,13126E | FALSE | Medium | IsMasterP | AAEL0094 | 2:2467336 | 30 | 1 | FALSE | 1 | 1 | 1 | 5680 | 6106 | 809 |
| -3,16928E | FALSE | Medium | IsMasterP | AAEL0196 | 2:3548304 | 31 | 3 | FALSE | 1 | 1 | 1 | 1123 | 1217 | 938 |
| 8,843E+18 | FALSE | Medium | IsMasterP | AAEL0060 | 3:3075610 | 32 | 9 | FALSE | 1 | 1 | 1 | 210 | 236 | 521 |
| -7,16362E | FALSE | Medium | IsMasterP | AAEL0106 | 2:1360704 | 32 | 9 | FALSE | 2 | 2 | 2 | 470 | 556 | 677 |
| 8,60497E | FALSE | Medium | IsMasterP | AAEL0031 | 1:1940431 | 32 | 24 | FALSE | 1 | 1 | 1 | 67 | 81 | 928 |
| 6,94692E | FALSE | Medium | IsMasterP | AAEL0049 | 2:7139623 | 33 | 9 | FALSE | 1 | 1 | 1 | 159 | 180 | 1030 |
| -6,29827E | FALSE | Medium | IsMasterP | AAEL0121 | 2:2248981 | 33 | 14 | FALSE | 1 | 1 | 1 | 279 | 307 | 677 |
| 1,62008E | FALSE | Medium | IsMasterP | AAEL0128 | 1:2221310 | 33 | 13 | FALSE | 1 | 1 | 1 | 237 | 271 | 572 |
| -3,0455E+ | FALSE | Medium | IsMasterP | AAEL0203 | 1:1232504 | 33 | 9 | FALSE | 1 | 1 | 1 | 231 | 257 | 536 |
| 8,29675E | FALSE | Medium | IsMasterP | AAEL0106 | 2:1364823 | 33 | 2 | FALSE | 1 | 1 | 1 | 1295 | 1446 | 846 |
| -7,76727E | FALSE | Medium | IsMasterP | AAEL0143 | 3:7519084 | 33 | 7 | FALSE | 1 | 1 | 1 | 514 | 551 | 583 |
| -2,97512E | FALSE | Medium | IsMasterP | AAEL0034 | 3:3492542 | 33 | 4 | FALSE | 1 | 1 | 1 | 533 | 610 | 586 |
| 5,63521E | FALSE | Medium | IsMasterP | AAEL0079 | 2:4315440 | 34 | 4 | FALSE | 1 | 1 | 1 | 533 | 624 | 766 |
| 2,07144E | FALSE | Medium | IsMasterP | AAEL0090 | 3:4609003 | 34 | 11 | FALSE | 2 | 2 | 2 | 168 | 197 | 885 |
| -5,85273E | FALSE | Medium | IsMasterP | AAEL0094 | 3:3380600 | 34 | 8 | FALSE | 1 | 1 | 1 | 418 | 458 | 496 |
| -5,2272E+ | FALSE | Medium | IsMasterP | AAEL0238 | 2:4534337 | 34 | 28 | FALSE | 1 | 1 | 1 | 101 | 109 | 825 |
| 1,03751E | FALSE | Medium | IsMasterP | AAEL0017 | 2:1667906 | 34 | 1 | FALSE | 1 | 2 | 1 | 786 | 908 | 540 |
| -2,3357E+ | FALSE | Medium | IsMasterP | AAEL0051 | 3:3466626 | 34 | 7 | FALSE | 1 | 1 | 1 | 527 | 594 | 648 |
| -2,82765E | FALSE | Medium | IsMasterP | AAEL0049 | 2:2811934 | 34 | 16 | FALSE | 1 | 1 | 1 | 304 | 325 | 654 |
| -5,97634E | FALSE | Medium | IsMasterP | AAEL0148 | 2:3320194 | 34 | 2 | FALSE | 1 | 1 | 1 | 1111 | 1248 | 783 |
| 6,31413E | FALSE | Medium | IsMasterP | AAEL0083 | 1:8200296 | 34 | 5 | FALSE | 1 | 1 | 1 | 628 | 700 | 829 |
| 1,09587E | FALSE | Medium | IsMasterP | AAEL0138 | 1:3594708 | 34 | 2 | FALSE | 1 | 1 | 1 | 702 | 797 | 766 |
| 6,34771E | FALSE | Medium | IsMasterP | AAEL0213 | 1:4330977 | 34 | 4 | FALSE | 1 | 1 | 1 | 1005 | 1081 | 816 |
| -3,39067E | FALSE | Medium | IsMasterP | AAEL0065 | 2:1331185 | 35 | 7 | FALSE | 1 | 1 | 1 | 300 | 328 | 470 |
| 2,69007E | FALSE | Medium | IsMasterP | AAEL0217 | 3:2610133 | 35 | 3 | FALSE | 1 | 1 | 1 | 1124 | 1260 | 558 |
| -2,3993E+ | FALSE | Medium | IsMasterP | AAEL0035 | 2:2088817 | 35 | 5 | FALSE | 1 | 1 | 1 | 463 | 530 | 634 |
| -6,801E+1 | FALSE | Medium | IsMasterP | AAEL0028 | 1:3053032 | 35 | 24 | FALSE | 1 | 1 | 1 | 84 | 101 | 973 |

| | | | | | | | | | | | | | | |
|---|---|---|---|---|---|---|---|---|---|---|---|---|---|---|
| -5,60249E | FALSE | Medium | IsMasterP | AAEL0094 | 1:981939 | 35 | 11 | FALSE | 1 | 1 | 1 | 196 | 215 | 863 |
| -3,92726E | FALSE | Medium | IsMasterP | AAEL0119 | 1:285550 | 35 | 14 | FALSE | 1 | 1 | 1 | 236 | 265 | 860 |
| 1,12464E | FALSE | Medium | IsMasterP | AAEL0263 | 3:166376 | 35 | 6 | FALSE | 1 | 1 | 1 | 514 | 583 | 950 |
| -6,54747E | FALSE | Medium | IsMasterP | AAEL0099 | 1:308052 | 35 | 18 | FALSE | 1 | 1 | 1 | 155 | 176 | 508 |
| 1,55905E | FALSE | Medium | IsMasterP | AAEL0051 | 1:168480 | 35 | 16 | FALSE | 1 | 1 | 1 | 180 | 204 | 894 |
| 1,521E+1 | FALSE | Medium | IsMasterP | AAEL0001 | 1:194970 | 35 | 1 | FALSE | 1 | 1 | 1 | 1342 | 1492 | 453 |
| 4,8631E+ | FALSE | Medium | IsMasterP | AAEL0087 | 3:958407 | 35 | 2 | FALSE | 1 | 1 | 1 | 783 | 902 | 687 |
| 1,4048E+ | FALSE | Medium | IsMasterP | AAEL0021 | 2:428783 | 35 | 21 | FALSE | 1 | 1 | 1 | 127 | 124 | 649 |
| 4,89371E | FALSE | Medium | IsMasterP | AAEL0196 | 2:347670 | 35 | 1 | FALSE | 1 | 1 | 1 | 2244 | 2415 | 638 |
| 2,77583E | FALSE | Medium | IsMasterP | AAEL0056 | 2:337009 | 35 | 4 | FALSE | 1 | 1 | 1 | 664 | 746 | 1037 |
| -2,13014E | FALSE | Medium | IsMasterP | AAEL0126 | 3:239431 | 35 | 3 | FALSE | 1 | 1 | 1 | 493 | 562 | 973 |
| 1,8756E+ | FALSE | Medium | IsMasterP | AAEL0224 | 2:331107 | 35 | 3 | FALSE | 1 | 1 | 1 | 589 | 694 | 489 |
| -7,71677E | FALSE | Medium | IsMasterP | AAEL0125 | 3:242521 | 35 | 7 | FALSE | 1 | 1 | 1 | 342 | 391 | 597 |
| -5,52647E | FALSE | Medium | IsMasterP | AAEL0043 | 3:367068 | 35 | 4 | FALSE | 1 | 1 | 1 | 370 | 413 | 712 |
| -5,62605E | FALSE | Medium | IsMasterP | AAEL0023 | 2:175240 | 35 | 11 | FALSE | 1 | 1 | 1 | 161 | 183 | 969 |
| -8,72149E | FALSE | Medium | IsMasterP | AAEL0065 | 2:468013 | 35 | 15 | FALSE | 1 | 1 | 1 | 131 | 151 | 800 |
| -5,23302E | FALSE | Medium | IsMasterP | AAEL0058 | NIGP0100 | 35 | 2 | FALSE | 1 | 1 | 1 | 785 | 834 | 856 |
| 4,97991E | FALSE | Medium | IsMasterP | AAEL0102 | 3:157422 | 35 | 3 | FALSE | 1 | 1 | 1 | 462 | 530 | 945 |
| -9,15915E | FALSE | Medium | IsMasterP | AAEL0265 | 3:222537 | 35 | 4 | FALSE | 1 | 1 | 1 | 638 | 743 | 892 |
| -6,91963E | FALSE | Medium | IsMasterP | AAEL0014 | 2:262151 | 36 | 19 | FALSE | 1 | 1 | 1 | 165 | 187 | 822 |
| -5,33633E | FALSE | Medium | IsMasterP | AAEL0100 | 1:193263 | 36 | 12 | FALSE | 1 | 1 | 1 | 213 | 246 | 573 |
| -3,13904E | FALSE | Medium | IsMasterP | AAEL0140 | 3:147461 | 36 | 3 | FALSE | 1 | 1 | 1 | 873 | 970 | 600 |
| -2,39232E | FALSE | Medium | IsMasterP | AAEL0064 | 2:414683 | 36 | 9 | FALSE | 1 | 1 | 1 | 211 | 237 | 903 |
| -7,70688E | FALSE | Medium | IsMasterP | AAEL0109 | 1:140175 | 36 | 3 | FALSE | 1 | 1 | 1 | 815 | 895 | 525 |
| 1,31545E | FALSE | Medium | IsMasterP | AAEL0116 | 3:281544 | 36 | 3 | FALSE | 1 | 1 | 1 | 667 | 746 | 931 |
| 2,18308E | FALSE | Medium | IsMasterP | AAEL0038 | 3:342714 | 36 | 26 | FALSE | 1 | 1 | 1 | 86 | 100 | 399 |
| 8,65937E | FALSE | Medium | IsMasterP | AAEL0090 | 3:135222 | 36 | 7 | FALSE | 1 | 1 | 1 | 355 | 379 | 951 |
| -7,9361E+ | FALSE | Medium | IsMasterP | AAEL0001 | 3:215624 | 36 | 1 | FALSE | 1 | 1 | 1 | 3098 | 3450 | 734 |
| -2,58781E | FALSE | Medium | IsMasterP | AAEL0211 | 1:262542 | 36 | 2 | FALSE | 1 | 1 | 1 | 926 | 1033 | 862 |
| 1,52006E | FALSE | Medium | IsMasterP | AAEL0051 | 1:168496 | 36 | 3 | FALSE | 1 | 1 | 1 | 618 | 705 | 649 |
| -4,01781E | FALSE | Medium | IsMasterP | AAEL0048 | 2:115326 | 36 | 16 | FALSE | 1 | 1 | 1 | 201 | 214 | 926 |

| | | | | | | | | | | | | | | |
|---|---|---|---|---|---|---|---|---|---|---|---|---|---|---|
| 1,71937E | FALSE | Medium | IsMasterP | AAEL0032 | 1:291002 | 36 | 3 | FALSE | 1 | 1 | 1 | 329 | 383 | 756 |
| 1,91163E | FALSE | Medium | IsMasterP | AAEL0063 | 1:101462 | 36 | 5 | FALSE | 1 | 1 | 1 | 405 | 457 | 728 |
| -6,76261E | FALSE | Medium | IsMasterP | AAEL0020 | 3:178410 | 36 | 3 | FALSE | 1 | 1 | 1 | 547 | 618 | 677 |
| 6,26672E | FALSE | Medium | IsMasterP | AAEL0228 | 2:159598 | 36 | 3 | FALSE | 1 | 1 | 1 | 400 | 461 | 913 |
| 5,26164E | FALSE | Medium | IsMasterP | AAEL0120 | 3:160417 | 36 | 5 | FALSE | 1 | 1 | 1 | 585 | 652 | 511 |
| -8,9725E+ | FALSE | Medium | IsMasterP | AAEL0040 | 1:137025 | 36 | 8 | FALSE | 1 | 1 | 1 | 372 | 411 | 671 |
| -4,52261E | FALSE | Medium | IsMasterP | AAEL0078 | 3:382282 | 36 | 2 | FALSE | 1 | 1 | 1 | 825 | 932 | 670 |
| 3,63991E | FALSE | Medium | IsMasterP | AAEL0006 | 3:395846 | 36 | 11 | FALSE | 1 | 1 | 1 | 274 | 301 | 543 |
| 3,40476E | FALSE | Medium | IsMasterP | AAEL0279 | 2:601270 | 36 | 1 | FALSE | 1 | 1 | 1 | 1834 | 2023 | 479 |
| -8,93652E | FALSE | Medium | IsMasterP | AAEL0183 | 3:809698 | 36 | 3 | FALSE | 1 | 1 | 1 | 581 | 624 | 554 |
| -8,39567E | FALSE | Medium | IsMasterP | AAEL0083 | 1:235428 | 36 | 1 | FALSE | 1 | 1 | 1 | 2091 | 2293 | 727 |
| -2,80004E | FALSE | Medium | IsMasterP | AAEL0032 | 2:366610 | 36 | 17 | FALSE | 1 | 1 | 1 | 123 | 132 | 496 |
| -8,49357E | FALSE | Medium | IsMasterP | AAEL0207 | 2:348436 | 36 | 2 | FALSE | 1 | 1 | 1 | 905 | 1032 | 824 |
| 7,71962E | FALSE | Medium | IsMasterP | AAEL0030 | 1:130658 | 36 | 3 | FALSE | 1 | 1 | 1 | 729 | 827 | 497 |
| -7,76957E | FALSE | Medium | IsMasterP | AAEL0056 | 1:732576 | 36 | 7 | FALSE | 1 | 1 | 1 | 417 | 471 | 774 |
| 4,26306E | FALSE | Medium | IsMasterP | AAEL0243 | 3:280059 | 37 | 10 | FALSE | 1 | 1 | 1 | 228 | 238 | 976 |
| -1,6471E+ | FALSE | Medium | IsMasterP | AAEL0107 | 3:305725 | 37 | 3 | FALSE | 1 | 1 | 1 | 422 | 484 | 721 |
| -8,41929E | FALSE | Medium | IsMasterP | AAEL0111 | 3:382071 | 37 | 4 | FALSE | 1 | 1 | 1 | 424 | 482 | 878 |
| 3,29416E | FALSE | Medium | IsMasterP | AAEL0216 | 1:128484 | 37 | 3 | FALSE | 1 | 1 | 1 | 753 | 851 | 712 |
| -8,25475E | FALSE | Medium | IsMasterP | AAEL0115 | 1:120330 | 37 | 10 | FALSE | 1 | 1 | 1 | 147 | 174 | 491 |
| -8,34729E | FALSE | Medium | IsMasterP | AAEL0112 | 2:179641 | 37 | 17 | FALSE | 1 | 1 | 1 | 138 | 156 | 594 |
| -7,25651E | FALSE | Medium | IsMasterP | AAEL0237 | 1:103588 | 37 | 3 | FALSE | 1 | 1 | 1 | 975 | 1079 | 931 |
| 5,40625E | FALSE | Medium | IsMasterP | AAEL0096 | 1:605945 | 37 | 9 | FALSE | 1 | 1 | 1 | 232 | 258 | 835 |
| -2,93333E | FALSE | Medium | IsMasterP | AAEL0108 | 1:550976 | 37 | 4 | FALSE | 1 | 1 | 1 | 456 | 505 | 939 |
| 2,46268E | FALSE | Medium | IsMasterP | AAEL0089 | 1:811532 | 37 | 3 | FALSE | 1 | 1 | 1 | 1010 | 1137 | 815 |
| -9,10099E | FALSE | Medium | IsMasterP | AAEL0007 | 2:554310 | 37 | 2 | FALSE | 1 | 1 | 1 | 1298 | 1403 | 545 |
| -5,24098E | FALSE | Medium | IsMasterP | AAEL0088 | 3:133254 | 37 | 1 | FALSE | 1 | 1 | 1 | 1666 | 1904 | 705 |
| 4,85053E | FALSE | Medium | IsMasterP | AAEL0043 | 3:938803 | 37 | 2 | FALSE | 1 | 1 | 1 | 1154 | 1324 | 743 |
| 6,80836E | FALSE | Medium | IsMasterP | AAEL0196 | 1:764033 | 37 | 3 | FALSE | 1 | 1 | 1 | 574 | 663 | 652 |
| -2,21802E | FALSE | Medium | IsMasterP | AAEL0141 | 3:241952 | 37 | 10 | FALSE | 1 | 1 | 1 | 180 | 202 | 618 |
| -6,50394E | FALSE | Medium | IsMasterP | AAEL0209 | 3:138176 | 37 | 1 | FALSE | 1 | 1 | 1 | 3218 | 3473 | 560 |

| | | | | | | | | | | | | | | |
|---|---|---|---|---|---|---|---|---|---|---|---|---|---|---|
| 8,31879E | FALSE | Medium | IsMasterP | AAEL0089 | 3:504125 | 37 | 5 | FALSE | 1 | 1 | 1 | 551 | 617 | 810 |
| 3,43966E | FALSE | Medium | IsMasterP | AAEL0121 | 2:188049 | 37 | 7 | FALSE | 1 | 1 | 1 | 215 | 233 | 964 |
| -5,66401E | FALSE | Medium | IsMasterP | AAEL0232 | 1:961032 | 37 | 10 | FALSE | 1 | 1 | 1 | 182 | 205 | 868 |
| 8,86104E | FALSE | Medium | IsMasterP | AAEL0101 | 3:148676 | 37 | 1 | FALSE | 1 | 1 | 1 | 840 | 961 | 547 |
| -2,59659E | FALSE | Medium | IsMasterP | AAEL0111 | 3:194119 | 37 | 19 | FALSE | 1 | 1 | 1 | 104 | 125 | 1010 |
| -4,58741E | FALSE | Medium | IsMasterP | AAEL0025 | 1:227035 | 37 | 5 | FALSE | 1 | 1 | 1 | 772 | 864 | 651 |
| 9,15068E | FALSE | Medium | IsMasterP | AAEL0108 | 2:127938 | 37 | 2 | FALSE | 1 | 1 | 1 | 1099 | 1217 | 581 |
| -7,01727E | FALSE | Medium | IsMasterP | AAEL0090 | 3:134923 | 38 | 1 | FALSE | 1 | 1 | 1 | 2104 | 2286 | 484 |
| 8,92451E | FALSE | Medium | IsMasterP | AAEL0091 | 1:112779 | 38 | 6 | FALSE | 1 | 1 | 1 | 245 | 285 | 761 |
| -4,32651E | FALSE | Medium | IsMasterP | AAEL0063 | 3:396615 | 39 | 4 | FALSE | 1 | 1 | 1 | 337 | 399 | 869 |
| -6,15313E | FALSE | Medium | IsMasterP | AAEL0038 | 3:342738 | 39 | 3 | FALSE | 1 | 1 | 1 | 427 | 450 | 950 |
| -7,17106E | FALSE | Medium | IsMasterP | AAEL0072 | 2:422336 | 39 | 6 | FALSE | 1 | 1 | 1 | 254 | 295 | 938 |
| -2,5531E+ | FALSE | Medium | IsMasterP | AAEL0036 | 2:317069 | 39 | 10 | FALSE | 1 | 1 | 1 | 140 | 154 | 1017 |
| 6,02056E | FALSE | Medium | IsMasterP | AAEL0146 | 1:119355 | 40 | 1 | FALSE | 1 | 1 | 1 | 1833 | 2019 | 648 |
| -2,81985E | FALSE | Medium | IsMasterP | AAEL0017 | 2:443126 | 40 | 2 | FALSE | 1 | 1 | 1 | 964 | 1050 | 596 |
| 8,90763E | FALSE | Medium | IsMasterP | AAEL0119 | 1:286326 | 39 | 5 | FALSE | 1 | 1 | 1 | 233 | 264 | 692 |
| -8,99172E | FALSE | Medium | IsMasterP | AAEL0000 | 1:242991 | 39 | 5 | FALSE | 1 | 1 | 1 | 424 | 492 | 648 |
| -2,24443E | FALSE | Medium | IsMasterP | AAEL0098 | 3:188012 | 39 | 6 | FALSE | 1 | 1 | 1 | 318 | 368 | 895 |
| 3,04081E | FALSE | Medium | IsMasterP | AAEL0060 | 3:741312 | 39 | 10 | FALSE | 1 | 1 | 1 | 129 | 149 | 955 |
| 7,33399E | FALSE | Medium | IsMasterP | AAEL0183 | 2:285441 | 40 | 1 | FALSE | 1 | 1 | 1 | 1373 | 1548 | 524 |
| -3,85516E | FALSE | Medium | IsMasterP | AAEL0136 | 2:324308 | 40 | 2 | FALSE | 1 | 1 | 1 | 769 | 851 | 680 |
| 8,89056E | FALSE | Medium | IsMasterP | AAEL0233 | 1:261460 | 40 | 23 | FALSE | 1 | 1 | 1 | 87 | 105 | 850 |
| 1,6781E+ | FALSE | Medium | IsMasterP | AAEL0197 | 3:644278 | 40 | 8 | FALSE | 1 | 1 | 1 | 283 | 299 | 1039 |
| -3,44557E | FALSE | Medium | IsMasterP | AAEL0133 | 2:323939 | 40 | 3 | FALSE | 1 | 1 | 1 | 948 | 1075 | 885 |
| -5,62332E | FALSE | Medium | IsMasterP | AAEL0105 | 2:446248 | 40 | 8 | FALSE | 1 | 1 | 1 | 218 | 240 | 477 |
| -3,16734E | FALSE | Medium | IsMasterP | AAEL0093 | NIGP0100 | 40 | 5 | FALSE | 1 | 1 | 1 | 597 | 667 | 739 |
| -3,41894E | FALSE | Medium | IsMasterP | AAEL0120 | 3:160472 | 40 | 2 | FALSE | 1 | 1 | 1 | 1709 | 1937 | 560 |
| -2,22181E | FALSE | Medium | IsMasterP | AAEL0128 | 1:222220 | 41 | 9 | FALSE | 1 | 1 | 1 | 240 | 278 | 562 |
| 1,82796E | FALSE | Medium | IsMasterP | AAEL0019 | 2:194489 | 41 | 2 | FALSE | 1 | 1 | 1 | 689 | 776 | 758 |
| 5,5807E+ | FALSE | Medium | IsMasterP | AAEL0034 | 3:815324 | 41 | 9 | FALSE | 1 | 1 | 1 | 173 | 190 | 607 |
| -6,09342E | FALSE | Medium | IsMasterP | AAEL0199 | 3:325406 | 41 | 3 | FALSE | 1 | 1 | 1 | 752 | 821 | 749 |

| | | | | | | | | | | | | | | |
|---|---|---|---|---|---|---|---|---|---|---|---|---|---|---|
| 7,30554E | FALSE | Medium | IsMasterP | AAEL0169 | 1:1079429 | 41 | 3 | FALSE | 1 | 1 | 1 | 716 | 801 | 829 |
| 5,58767E | FALSE | Medium | IsMasterP | AAEL0010 | 1:1852198 | 41 | 15 | FALSE | 1 | 1 | 1 | 110 | 125 | 606 |
| 7,023E+18 | FALSE | Medium | IsMasterP | AAEL0004 | 3:1852561 | 41 | 5 | FALSE | 1 | 1 | 1 | 694 | 772 | 711 |
| -7,60229E | FALSE | Medium | IsMasterP | AAEL0254 | 1:2858387 | 42 | 4 | FALSE | 1 | 1 | 1 | 755 | 862 | 898 |
| -3,76974E | FALSE | Medium | IsMasterP | AAEL0198 | 2:2153850 | 42 | 2 | FALSE | 1 | 1 | 1 | 1890 | 2115 | 747 |
| -7,11714E | FALSE | Medium | IsMasterP | AAEL0065 | 2:4686478 | 42 | 2 | FALSE | 1 | 1 | 1 | 938 | 1029 | 715 |
| -9,1752E+ | FALSE | Medium | IsMasterP | AAEL0034 | 3:3486586 | 42 | 3 | FALSE | 1 | 1 | 1 | 600 | 684 | 538 |
| -4,22291E | FALSE | Medium | IsMasterP | AAEL0195 | 3:6794320 | 42 | 2 | FALSE | 1 | 1 | 1 | 1764 | 1982 | 803 |
| -6,71547E | FALSE | Medium | IsMasterP | AAEL0241 | 2:1238117 | 42 | 16 | FALSE | 1 | 1 | 1 | 153 | 166 | 906 |
| -7,34958E | FALSE | Medium | IsMasterP | AAEL0259 | 1:3609070 | 43 | 3 | FALSE | 1 | 1 | 1 | 468 | 522 | 718 |
| 1,5343E+1 | FALSE | Medium | IsMasterP | AAEL0214 | 1:9866970 | 43 | 7 | FALSE | 1 | 1 | 1 | 528 | 610 | 562 |
| -4,31845E | FALSE | Medium | IsMasterP | AAEL0106 | 2:1360153 | 43 | 8 | FALSE | 1 | 1 | 1 | 250 | 267 | 717 |
| 6,10175E | FALSE | Medium | IsMasterP | AAEL0061 | 3:1417216 | 43 | 6 | FALSE | 1 | 1 | 1 | 553 | 615 | 834 |
| -5,01152E | FALSE | Medium | IsMasterP | AAEL0143 | 2:5167157 | 43 | 8 | FALSE | 1 | 1 | 1 | 428 | 497 | 822 |
| 3,88828E | FALSE | Medium | IsMasterP | AAEL0124 | 2:2860373 | 43 | 6 | FALSE | 1 | 1 | 1 | 471 | 541 | 907 |
| -8,97005E | FALSE | Medium | IsMasterP | AAEL0276 | NIGP0100 | 43 | 3 | FALSE | 1 | 1 | 1 | 577 | 655 | 923 |
| -6,78767E | FALSE | Medium | IsMasterP | AAEL0008 | 3:3341649 | 43 | 5 | FALSE | 1 | 1 | 1 | 540 | 592 | 807 |
| 8,22686E | FALSE | Medium | IsMasterP | AAEL0140 | 2:3533532 | 44 | 2 | FALSE | 1 | 1 | 1 | 853 | 956 | 676 |
| -5,055E+1 | FALSE | Medium | IsMasterP | AAEL0091 | 3:1351661 | 44 | 5 | FALSE | 1 | 1 | 1 | 306 | 336 | 948 |
| -5,35969E | FALSE | Medium | IsMasterP | AAEL0126 | 3:2397750 | 44 | 7 | FALSE | 1 | 1 | 1 | 486 | 553 | 506 |
| -7,55814E | FALSE | Medium | IsMasterP | AAEL0031 | 3:1820873 | 44 | 2 | FALSE | 1 | 1 | 1 | 1252 | 1402 | 662 |
| 8,49664E | FALSE | Medium | IsMasterP | AAEL0029 | 1:1081183 | 44 | 2 | FALSE | 1 | 1 | 1 | 916 | 1025 | 599 |
| 5,81006E | FALSE | Medium | IsMasterP | AAEL0116 | 3:1923925 | 44 | 2 | FALSE | 1 | 1 | 1 | 856 | 963 | 824 |
| -5,52452E | FALSE | Medium | IsMasterP | AAEL0049 | 2:2596216 | 44 | 2 | FALSE | 1 | 1 | 1 | 712 | 802 | 652 |
| -7,17812E | FALSE | Medium | IsMasterP | AAEL0016 | 1:1388096 | 44 | 3 | FALSE | 1 | 1 | 1 | 484 | 545 | 571 |
| -8,05405E | FALSE | Medium | IsMasterP | AAEL0274 | 2:1258752 | 44 | 0 | FALSE | 1 | 1 | 1 | 2992 | 3384 | 602 |
| 7,04503E | FALSE | Medium | IsMasterP | AAEL0068 | 3:4311215 | 44 | 7 | FALSE | 1 | 1 | 1 | 243 | 281 | 640 |
| 3,14889E | FALSE | Medium | IsMasterP | AAEL0134 | 2:1992714 | 44 | 2 | FALSE | 1 | 1 | 1 | 685 | 791 | 737 |
| 8,34375E | FALSE | Medium | IsMasterP | AAEL0002 | 3:2664183 | 44 | 1 | FALSE | 1 | 1 | 1 | 1627 | 1738 | 910 |
| -8,79096E | FALSE | Medium | IsMasterP | AAEL0113 | 2:2662447 | 45 | 4 | FALSE | 1 | 1 | 1 | 798 | 911 | 709 |
| -2,08346E | FALSE | Medium | IsMasterP | AAEL0002 | 2:7477206 | 45 | 10 | FALSE | 1 | 1 | 1 | 143 | 168 | 756 |

| | | | | | | | | | | | | | | |
|---|---|---|---|---|---|---|---|---|---|---|---|---|---|---|
| -5,5193E+ | FALSE | Medium | IsMasterP | AAEL0134 | 2:121088 | 45 | 2 | FALSE | 1 | 1 | 1 | 889 | 998 | 619 |
| -8,58692E | FALSE | Medium | IsMasterP | AAEL0024 | 1:176754 | 45 | 2 | FALSE | 1 | 1 | 1 | 791 | 870 | 607 |
| -7,82988E | FALSE | Medium | IsMasterP | AAEL0277 | 3:197830 | 45 | 1 | FALSE | 1 | 1 | 1 | 1567 | 1777 | 592 |
| 2,57294E | FALSE | Medium | IsMasterP | AAEL0099 | 1:206637 | 45 | 3 | FALSE | 1 | 1 | 1 | 686 | 771 | 955 |
| 2,18482E | FALSE | Medium | IsMasterP | AAEL0242 | 1:345375 | 45 | 1 | FALSE | 1 | 1 | 1 | 2676 | 2998 | 609 |
| 5,80791E | FALSE | Medium | IsMasterP | AAEL0138 | 1:358490 | 45 | 8 | FALSE | 1 | 1 | 1 | 358 | 387 | 879 |
| -7,10949E | FALSE | Medium | IsMasterP | AAEL0020 | 2:119994 | 45 | 3 | FALSE | 1 | 1 | 1 | 445 | 504 | 548 |
| -7,49276E | FALSE | Medium | IsMasterP | AAEL0249 | 1:886724 | 45 | 3 | FALSE | 1 | 1 | 1 | 1067 | 1199 | 498 |
| 6,89747E | FALSE | Medium | IsMasterP | AAEL0001 | 1:195025 | 45 | 6 | FALSE | 1 | 1 | 1 | 251 | 291 | 917 |
| -4,1784E+ | FALSE | Medium | IsMasterP | AAEL0107 | 2:305340 | 45 | 5 | FALSE | 1 | 1 | 1 | 298 | 346 | 843 |
| -1,89085E | FALSE | Medium | IsMasterP | AAEL0102 | 2:402957 | 45 | 5 | FALSE | 1 | 1 | 1 | 239 | 271 | 894 |
| -2,61E+18 | FALSE | Medium | IsMasterP | AAEL0128 | 3:417832 | 45 | 8 | FALSE | 1 | 1 | 1 | 416 | 449 | 812 |
| 1,1225E+ | FALSE | Medium | IsMasterP | AAEL0054 | 3:382985 | 45 | 10 | FALSE | 1 | 1 | 1 | 215 | 246 | 587 |
| 1,60309E | FALSE | Medium | IsMasterP | AAEL0232 | 1:914139 | 45 | 5 | FALSE | 1 | 1 | 1 | 345 | 384 | 662 |
| 6,13426E | FALSE | Low | IsMasterP | AAEL0041 | 1:122471 | 60 | 3 | FALSE | 1 | 1 | 1 | 699 | 775 | 980 |
| 5,64011E | FALSE | Low | IsMasterP | AAEL0106 | 1:264259 | 60 | 8 | FALSE | 1 | 1 | 1 | 362 | 407 | 730 |
| -1,00031E | FALSE | Low | IsMasterP | AAEL0077 | 3:152871 | 60 | 6 | FALSE | 1 | 1 | 1 | 497 | 571 | 640 |
| 7,62481E | FALSE | Low | IsMasterP | AAEL0213 | 2:402784 | 61 | 4 | FALSE | 1 | 1 | 1 | 765 | 852 | 822 |
| -1,75029E | FALSE | Low | IsMasterP | AAEL0206 | 1:139861 | 61 | 1 | FALSE | 1 | 1 | 1 | 3337 | 3822 | 633 |
| -6,44828E | FALSE | Low | IsMasterP | AAEL0012 | 2:153078 | 61 | 1 | FALSE | 1 | 1 | 1 | 1144 | 1248 | 743 |
| 6,4224E+ | FALSE | Low | IsMasterP | AAEL0138 | 2:122839 | 61 | 9 | FALSE | 1 | 1 | 1 | 167 | 183 | 960 |
| 8,45504E | FALSE | Low | IsMasterP | AAEL0124 | 1:123549 | 61 | 3 | FALSE | 1 | 1 | 1 | 354 | 388 | 888 |
| 4,31534E | FALSE | Low | IsMasterP | AAEL0267 | 2:400980 | 61 | 4 | FALSE | 1 | 1 | 1 | 955 | 1055 | 862 |
| -4,47396E | FALSE | Low | IsMasterP | AAEL0111 | 3:381886 | 61 | 5 | FALSE | 1 | 1 | 1 | 664 | 738 | 737 |
| 9,17782E | FALSE | Low | IsMasterP | AAEL0078 | 3:381363 | 61 | 3 | FALSE | 1 | 1 | 1 | 903 | 1018 | 683 |
| -5,40555E | FALSE | Low | IsMasterP | AAEL0015 | 2:211668 | 61 | 5 | FALSE | 1 | 1 | 1 | 496 | 567 | 898 |
| -5,25251E | FALSE | Low | IsMasterP | AAEL0115 | 3:312824 | 62 | 9 | FALSE | 1 | 1 | 1 | 414 | 485 | 837 |
| -6,86808E | FALSE | Low | IsMasterP | AAEL0175 | 2:210152 | 62 | 7 | FALSE | 1 | 1 | 1 | 398 | 459 | 720 |
| 1,22615E | FALSE | Low | IsMasterP | AAEL0022 | 2:429716 | 62 | 5 | FALSE | 1 | 1 | 1 | 297 | 335 | 614 |
| 7,41945E | FALSE | Low | IsMasterP | AAEL0067 | 3:368125 | 63 | 2 | FALSE | 1 | 1 | 1 | 542 | 598 | 506 |
| 7,34152E | FALSE | Low | IsMasterP | AAEL0039 | 3:267714 | 63 | 3 | FALSE | 1 | 1 | 1 | 408 | 467 | 547 |

| | | | | | | | | | | | | | | |
|---|---|---|---|---|---|---|---|---|---|---|---|---|---|---|
| 2,97822E | FALSE | Low | IsMasterP | AAEL0119 | 2:301715 | 63 | 4 | FALSE | 1 | 1 | 1 | 318 | 357 | 531 |
| -4,90753E | FALSE | Low | IsMasterP | AAEL0052 | 1:321783 | 63 | 3 | FALSE | 1 | 1 | 1 | 521 | 585 | 838 |
| 3,21342E | FALSE | Low | IsMasterP | AAEL0096 | 2:105751 | 64 | 3 | FALSE | 1 | 1 | 1 | 437 | 496 | 859 |
| -4,86513E | FALSE | Low | IsMasterP | AAEL0011 | 2:409893 | 64 | 2 | FALSE | 1 | 1 | 1 | 1714 | 1935 | 511 |
| 6,62481E | FALSE | Low | IsMasterP | AAEL0150 | 2:121949 | 64 | 2 | FALSE | 1 | 1 | 1 | 655 | 735 | 592 |
| -7,60621E | FALSE | Low | IsMasterP | AAEL0270 | 1:127838 | 64 | 4 | FALSE | 1 | 1 | 1 | 703 | 781 | 637 |
| 4,28298E | FALSE | Low | IsMasterP | AAEL0118 | 2:287911 | 64 | 4 | FALSE | 1 | 1 | 1 | 500 | 578 | 806 |
| -4,68523E | FALSE | Low | IsMasterP | AAEL0075 | 1:198648 | 64 | 2 | FALSE | 1 | 1 | 1 | 745 | 844 | 752 |
| -7,85346E | FALSE | Low | IsMasterP | AAEL0075 | 1:198064 | 64 | 7 | FALSE | 1 | 1 | 1 | 156 | 179 | 497 |
| 1,35808E | FALSE | Low | IsMasterP | AAEL0029 | 1:108768 | 65 | 3 | FALSE | 1 | 1 | 1 | 327 | 375 | 554 |
| 8,47444E | FALSE | Low | IsMasterP | AAEL0259 | 1:433791 | 65 | 1 | FALSE | 1 | 1 | 1 | 1037 | 1121 | 701 |
| 4,90123E | FALSE | Low | IsMasterP | AAEL0134 | 2:159325 | 65 | 3 | FALSE | 1 | 1 | 1 | 563 | 631 | 621 |
| 7,74255E | FALSE | Low | IsMasterP | AAEL0092 | 3:199715 | 65 | 1 | FALSE | 1 | 1 | 1 | 936 | 1043 | 676 |
| -3,76596E | FALSE | Low | IsMasterP | AAEL0124 | 1:213451 | 65 | 0 | FALSE | 1 | 1 | 1 | 4887 | 5392 | 547 |
| 3,51761E | FALSE | Low | IsMasterP | AAEL0090 | 2:269541 | 65 | 2 | FALSE | 1 | 1 | 1 | 1272 | 1435 | 642 |
| 4,66983E | FALSE | Low | IsMasterP | AAEL0061 | 3:141308 | 65 | 5 | FALSE | 1 | 1 | 1 | 389 | 429 | 482 |
| -3,34263E | FALSE | Low | IsMasterP | AAEL0015 | 2:229663 | 65 | 8 | FALSE | 1 | 1 | 1 | 484 | 554 | 753 |
| -1,0659E+ | FALSE | Low | IsMasterP | AAEL0011 | 2:307531 | 65 | 4 | FALSE | 1 | 1 | 1 | 323 | 364 | 895 |
| 3,29598E | FALSE | Low | IsMasterP | AAEL0172 | 3:112545 | 65 | 6 | FALSE | 1 | 1 | 1 | 498 | 574 | 841 |
| -6,8492E+ | FALSE | Low | IsMasterP | AAEL0084 | 1:532649 | 66 | 6 | FALSE | 1 | 1 | 1 | 520 | 570 | 740 |
| -6,50785E | FALSE | Low | IsMasterP | AAEL0250 | 1:145577 | 66 | 10 | FALSE | 1 | 1 | 1 | 304 | 347 | 547 |
| 7,54762E | FALSE | Low | IsMasterP | AAEL0198 | 2:347307 | 66 | 2 | FALSE | 1 | 1 | 1 | 1764 | 1910 | 816 |
| 7,10328E | FALSE | Low | IsMasterP | AAEL0132 | 3:340858 | 66 | 11 | FALSE | 1 | 1 | 1 | 165 | 185 | 715 |
| 7,22636E | FALSE | Low | IsMasterP | AAEL0019 | 3:226851 | 66 | 5 | FALSE | 1 | 1 | 1 | 546 | 618 | 764 |
| 4,54024E | FALSE | Low | IsMasterP | AAEL0057 | 2:336848 | 66 | 8 | FALSE | 1 | 1 | 1 | 383 | 430 | 876 |
| -4,83633E | FALSE | Low | IsMasterP | AAEL0214 | 2:360271 | 66 | 2 | FALSE | 1 | 1 | 1 | 1084 | 1201 | 807 |
| 1,61335E | FALSE | Low | IsMasterP | AAEL0009 | 3:379978 | 66 | 3 | FALSE | 1 | 1 | 1 | 1000 | 1141 | 911 |
| 8,85372E | FALSE | Low | IsMasterP | AAEL0272 | 2:504460 | 66 | 4 | FALSE | 1 | 1 | 1 | 737 | 845 | 623 |
| -8,81559E | FALSE | Low | IsMasterP | AAEL0086 | 3:172551 | 66 | 5 | FALSE | 1 | 1 | 1 | 249 | 280 | 588 |
| 7,46201E | FALSE | Low | IsMasterP | AAEL0120 | 2:186696 | 66 | 4 | FALSE | 1 | 1 | 1 | 330 | 389 | 931 |
| 4,2193E+ | FALSE | Low | IsMasterP | AAEL0062 | 1:224639 | 67 | 3 | FALSE | 1 | 1 | 1 | 761 | 857 | 812 |

| | | | | | | | | | | | | | | |
|---|---|---|---|---|---|---|---|---|---|---|---|---|---|---|
| 1,95419E+ | FALSE | Low | IsMasterP | AAEL0080 | 2:2307588 | 67 | 1 | FALSE | 1 | 1 | 1 | 1669 | 1879 | 545 |
| -8,90549E | FALSE | Low | IsMasterP | AAEL0231 | 1:6868452 | 68 | 1 | FALSE | 1 | 1 | 1 | 927 | 1008 | 717 |
| 3,30991E+ | FALSE | Low | IsMasterP | AAEL0080 | 1:1542445 | 68 | 5 | FALSE | 1 | 1 | 1 | 255 | 288 | 960 |
| -3,58454E | FALSE | Low | IsMasterP | AAEL0197 | 3:3327329 | 68 | 8 | FALSE | 1 | 1 | 1 | 429 | 447 | 599 |
| -2,55404E | FALSE | Low | IsMasterP | AAEL0077 | 2:4704335 | 68 | 1 | FALSE | 1 | 1 | 1 | 2911 | 3183 | 553 |
| 7,93186E+ | FALSE | Low | IsMasterP | AAEL0092 | 2:2684446 | 68 | 4 | FALSE | 1 | 1 | 1 | 322 | 378 | 477 |
| 2,33161E+ | FALSE | Low | IsMasterP | AAEL0111 | 2:2318677 | 68 | 2 | FALSE | 1 | 1 | 1 | 696 | 804 | 637 |
| -5,05832E | FALSE | Low | IsMasterP | AAEL0001 | 3:1090236 | 69 | 1 | FALSE | 1 | 1 | 1 | 1687 | 1920 | 655 |
| -4,10868E | FALSE | Low | IsMasterP | AAEL0197 | 1:3077968 | 69 | 2 | FALSE | 1 | 1 | 1 | 2354 | 2538 | 869 |
| -8,99242E | FALSE | Low | IsMasterP | AAEL0037 | 2:1973944 | 69 | 2 | FALSE | 1 | 1 | 1 | 685 | 770 | 957 |
| -8,9997E+ | FALSE | Low | IsMasterP | AAEL0196 | 2:1743755 | 69 | 4 | FALSE | 1 | 1 | 1 | 551 | 626 | 670 |
| 2,90496E+ | FALSE | Low | IsMasterP | AAEL0001 | 2:1096830 | 69 | 9 | FALSE | 1 | 1 | 1 | 180 | 202 | 576 |
| -6,65133E | FALSE | Low | IsMasterP | AAEL0011 | 2:4114721 | 69 | 4 | FALSE | 1 | 1 | 1 | 383 | 418 | 803 |
| -7,7207E+ | FALSE | Low | IsMasterP | AAEL0138 | 3:2018632 | 69 | 3 | FALSE | 1 | 1 | 1 | 567 | 644 | 835 |
| 1,54206E+ | FALSE | Low | IsMasterP | AAEL0074 | 2:2913209 | 69 | 12 | FALSE | 1 | 1 | 1 | 113 | 135 | 809 |
| -7,76859E | FALSE | Low | IsMasterP | AAEL0195 | 1:8094731 | 69 | 2 | FALSE | 1 | 1 | 1 | 2162 | 2379 | 638 |
| 7,83359E+ | FALSE | Low | IsMasterP | AAEL0084 | 1:5430960 | 69 | 2 | FALSE | 1 | 1 | 1 | 811 | 903 | 950 |
| 3,58386E+ | FALSE | Low | IsMasterP | AAEL0048 | 2:1151296 | 69 | 10 | FALSE | 1 | 1 | 1 | 330 | 355 | 832 |
| 3,11035E+ | FALSE | Low | IsMasterP | AAEL0078 | 2:4609610 | 69 | 5 | FALSE | 1 | 1 | 1 | 386 | 431 | 613 |
| 6,34858E+ | FALSE | Low | IsMasterP | AAEL0215 | 1:7917387 | 69 | 1 | FALSE | 1 | 1 | 1 | 3028 | 3369 | 555 |
| 1,79882E+ | FALSE | Low | IsMasterP | AAEL0078 | 1:1032346 | 69 | 6 | FALSE | 1 | 1 | 1 | 219 | 249 | 955 |
| 3,48793E+ | FALSE | Low | IsMasterP | AAEL0198 | 2:3001697 | 69 | 2 | FALSE | 1 | 1 | 1 | 1607 | 1758 | 695 |
| -7,26778E | FALSE | Low | IsMasterP | AAEL0145 | 2:1095718 | 70 | 5 | FALSE | 1 | 1 | 1 | 582 | 660 | 898 |
| -4,89075E | FALSE | Low | IsMasterP | AAEL0000 | 3:1049825 | 70 | 3 | FALSE | 1 | 1 | 1 | 391 | 428 | 681 |
| -5,91721E | FALSE | Low | IsMasterP | AAEL0250 | 2:1817834 | 70 | 3 | FALSE | 1 | 1 | 1 | 413 | 467 | 793 |
| -2,90385E | FALSE | Low | IsMasterP | AAEL0147 | 2:3242954 | 70 | 7 | FALSE | 1 | 1 | 1 | 174 | 188 | 931 |
| -4,8523E+ | FALSE | Low | IsMasterP | AAEL0235 | 2:3129073 | 70 | 1 | FALSE | 1 | 1 | 1 | 2101 | 2347 | 756 |
| 4,39031E+ | FALSE | Low | IsMasterP | AAEL0084 | 2:3951515 | 71 | 14 | FALSE | 1 | 1 | 1 | 120 | 136 | 628 |
| 7,64042E+ | FALSE | Low | IsMasterP | AAEL0181 | 1:1986983 | 71 | 4 | FALSE | 1 | 1 | 1 | 531 | 610 | 750 |
| -2,82162E | FALSE | Low | IsMasterP | AAEL0091 | 3:3909151 | 74 | 4 | FALSE | 1 | 1 | 1 | 630 | 703 | 875 |
| -1,78664E | FALSE | Low | IsMasterP | AAEL0265 | 2:3548287 | 74 | 3 | FALSE | 1 | 1 | 1 | 893 | 983 | 655 |

| | | | | | | | | | | | | | | |
|---|---|---|---|---|---|---|---|---|---|---|---|---|---|---|
| 3,028E+1 | FALSE | Low | IsMasterP | AAEL0204 | 1:286974 | 74 | 12 | FALSE | 1 | 1 | 1 | 184 | 207 | 835 |
| 8,78827E | FALSE | Low | IsMasterP | AAEL0281 | 1:311977 | 75 | 6 | FALSE | 1 | 1 | 1 | 400 | 443 | 680 |
| 6,73056E | FALSE | Low | IsMasterP | AAEL0049 | 3:322744 | 75 | 4 | FALSE | 1 | 1 | 1 | 603 | 667 | 557 |
| 8,68981E | FALSE | Low | IsMasterP | AAEL0115 | 3:106873 | 75 | 5 | FALSE | 1 | 1 | 1 | 380 | 419 | 479 |
| -3,49048E | FALSE | Low | IsMasterP | AAEL0131 | 3:304446 | 76 | 6 | FALSE | 1 | 1 | 1 | 549 | 608 | 689 |
| -1,21872E | FALSE | Low | IsMasterP | AAEL0023 | 2:175154 | 76 | 1 | FALSE | 1 | 1 | 1 | 1746 | 1989 | 616 |
| -3,74035E | FALSE | Low | IsMasterP | AAEL0002 | 2:743947 | 76 | 3 | FALSE | 1 | 1 | 1 | 310 | 346 | 533 |
| 5,16315E | FALSE | Low | IsMasterP | AAEL0151 | 2:199329 | 76 | 4 | FALSE | 1 | 1 | 1 | 476 | 547 | 802 |
| 6,80173E | FALSE | Low | IsMasterP | AAEL0108 | 1:550207 | 76 | 2 | FALSE | 1 | 1 | 1 | 1183 | 1285 | 878 |
| 8,11269E | FALSE | Low | IsMasterP | AAEL0136 | 1:618295 | 76 | 6 | FALSE | 1 | 1 | 1 | 325 | 353 | 683 |
| -2,34036E | FALSE | Low | IsMasterP | AAEL0229 | 2:787390 | 77 | 2 | FALSE | 1 | 1 | 1 | 725 | 803 | 961 |
| -8,95824E | FALSE | Low | IsMasterP | AAEL0022 | 3:374121 | 77 | 19 | FALSE | 1 | 1 | 1 | 74 | 86 | 955 |
| -8,30652E | FALSE | Low | IsMasterP | AAEL0004 | 3:161434 | 77 | 3 | FALSE | 1 | 1 | 1 | 566 | 648 | 487 |
| 8,7808E+ | FALSE | Low | IsMasterP | AAEL0241 | 2:255373 | 77 | 2 | FALSE | 1 | 1 | 1 | 1270 | 1421 | 606 |
| -8,74635E | FALSE | Low | IsMasterP | AAEL0003 | 3:400069 | 77 | 1 | FALSE | 1 | 1 | 1 | 1922 | 2156 | 686 |
| -1,77652E | FALSE | Low | IsMasterP | AAEL0061 | 3:687452 | 77 | 6 | FALSE | 1 | 1 | 1 | 302 | 338 | 488 |
| 7,8211E+ | FALSE | Low | IsMasterP | AAEL0195 | 3:372493 | 78 | 13 | FALSE | 1 | 1 | 1 | 91 | 105 | 815 |
| 1,5758E+ | FALSE | Low | IsMasterP | AAEL0071 | 3:190836 | 77 | 3 | FALSE | 1 | 1 | 1 | 545 | 615 | 800 |
| 7,90902E | FALSE | Low | IsMasterP | AAEL0066 | 2:463067 | 77 | 15 | FALSE | 1 | 1 | 1 | 99 | 114 | 901 |
| -8,36741E | FALSE | Low | IsMasterP | AAEL0006 | 3:394683 | 78 | 5 | FALSE | 1 | 1 | 1 | 751 | 819 | 843 |
| -4,19114E | FALSE | Low | IsMasterP | AAEL0032 | 2:367177 | 78 | 1 | FALSE | 1 | 1 | 1 | 794 | 906 | 785 |
| -6,9285E+ | FALSE | Low | IsMasterP | AAEL0119 | 3:350913 | 79 | 5 | FALSE | 1 | 1 | 1 | 624 | 722 | 664 |
| 2,90264E | FALSE | Low | IsMasterP | AAEL0290 | 3:342496 | 79 | 18 | FALSE | 1 | 1 | 1 | 91 | 100 | 538 |
| -6,03681E | FALSE | Low | IsMasterP | AAEL0067 | 2:836589 | 79 | 2 | FALSE | 1 | 1 | 1 | 422 | 470 | 945 |
| 5,06191E | FALSE | Low | IsMasterP | AAEL0054 | 1:247298 | 79 | 4 | FALSE | 1 | 1 | 1 | 343 | 393 | 780 |
| 1,38826E | FALSE | Low | IsMasterP | AAEL0003 | 2:368695 | 79 | 6 | FALSE | 1 | 1 | 1 | 361 | 409 | 802 |
| -1,9664E+ | FALSE | Low | IsMasterP | AAEL0028 | 1:304919 | 79 | 2 | FALSE | 1 | 1 | 1 | 776 | 875 | 881 |
| 3,09318E | FALSE | Low | IsMasterP | AAEL0112 | 2:226627 | 80 | 2 | FALSE | 1 | 1 | 1 | 1733 | 1963 | 706 |
| 3,10518E | FALSE | Low | IsMasterP | AAEL0052 | 3:895861 | 80 | 5 | FALSE | 1 | 1 | 1 | 485 | 531 | 574 |
| 2,74518E | FALSE | Low | IsMasterP | AAEL0268 | 1:216532 | 80 | 5 | FALSE | 1 | 1 | 1 | 639 | 707 | 840 |
| 1,03227E | FALSE | Low | IsMasterP | AAEL0027 | 3:145288 | 80 | 5 | FALSE | 1 | 1 | 1 | 262 | 292 | 794 |

| | | | | | | | | | | | | | | |
|---|---|---|---|---|---|---|---|---|---|---|---|---|---|---|
| 1,59758E- | FALSE | Low | IsMasterP | AAEL0025 | 1:8941597 | 80 | 3 | FALSE | 1 | 1 | 1 | 924 | 1038 | 667 |
| -1,09587E | FALSE | Low | IsMasterP | AAEL0098 | 3:2030308 | 80 | 1 | FALSE | 1 | 1 | 1 | 3383 | 3829 | 702 |
| -2,73893E | FALSE | Low | IsMasterP | AAEL0004 | 3:1859937 | 80 | 3 | FALSE | 1 | 1 | 1 | 415 | 480 | 762 |
| 2,94796E- | FALSE | Low | IsMasterP | AAEL0232 | 2:3203539 | 80 | 2 | FALSE | 1 | 1 | 1 | 740 | 806 | 668 |
| -2,73827E | FALSE | Low | IsMasterP | AAEL0001 | 2:1101427 | 80 | 12 | FALSE | 1 | 1 | 1 | 208 | 231 | 740 |
| 7,54428E- | FALSE | Low | IsMasterP | AAEL0114 | 2:1868780 | 81 | 7 | FALSE | 1 | 1 | 1 | 154 | 183 | 517 |
| 8,51694E- | FALSE | Low | IsMasterP | AAEL0116 | 3:2207671 | 81 | 4 | FALSE | 1 | 1 | 1 | 380 | 432 | 465 |
| 3,03203E- | FALSE | Low | IsMasterP | AAEL0015 | 3:2615857 | 82 | 3 | FALSE | 1 | 1 | 1 | 1185 | 1337 | 667 |
| -7,22756E | FALSE | Low | IsMasterP | AAEL0147 | 1:6999868 | 82 | 2 | FALSE | 1 | 1 | 1 | 486 | 536 | 467 |
| 8,49939E- | FALSE | Low | IsMasterP | AAEL0092 | 2:1126107 | 82 | 3 | FALSE | 1 | 1 | 1 | 896 | 1005 | 942 |
| 6,37214E- | FALSE | Low | IsMasterP | AAEL0221 | 2:3269823 | 82 | 1 | FALSE | 1 | 1 | 1 | 4228 | 4760 | 632 |
| -6,3031E+ | FALSE | Low | IsMasterP | AAEL0213 | 3:3988928 | 82 | 5 | FALSE | 1 | 1 | 1 | 663 | 761 | 712 |
| -1,91985E | FALSE | Low | IsMasterP | AAEL0251 | 3:3245393 | 82 | 2 | FALSE | 1 | 1 | 1 | 1460 | 1597 | 671 |
| 3,48375E- | FALSE | Low | IsMasterP | AAEL0122 | 1:1607594 | 82 | 1 | FALSE | 1 | 1 | 1 | 3293 | 3677 | 819 |
| 2,24299E- | FALSE | Low | IsMasterP | AAEL0035 | 2:2559152 | 83 | 4 | FALSE | 1 | 1 | 1 | 873 | 938 | 859 |
| 7,94679E- | FALSE | Low | IsMasterP | AAEL0194 | 1:2071627 | 83 | 8 | FALSE | 1 | 1 | 1 | 401 | 453 | 604 |
| 6,15042E- | FALSE | Low | IsMasterP | AAEL0075 | 2:3268484 | 83 | 5 | FALSE | 1 | 1 | 1 | 327 | 360 | 787 |
| 6,33866E- | FALSE | Low | IsMasterP | AAEL0251 | 2:8713492 | 84 | 3 | FALSE | 1 | 1 | 1 | 859 | 927 | 692 |
| 2,89933E- | FALSE | Low | IsMasterP | AAEL0048 | 3:2734491 | 84 | 11 | FALSE | 1 | 1 | 1 | 160 | 188 | 720 |
| 4,14377E- | FALSE | Low | IsMasterP | AAEL0086 | 2:3629384 | 84 | 6 | FALSE | 1 | 1 | 1 | 587 | 673 | 712 |
| 3,88041E- | FALSE | Low | IsMasterP | AAEL0091 | 2:9739020 | 84 | 3 | FALSE | 1 | 1 | 1 | 1161 | 1286 | 687 |
| 3,75112E- | FALSE | Low | IsMasterP | AAEL0030 | 2:2136054 | 84 | 6 | FALSE | 1 | 1 | 1 | 522 | 602 | 846 |
| 6,70224E- | FALSE | Low | IsMasterP | AAEL0111 | 3:3799640 | 84 | 4 | FALSE | 1 | 1 | 1 | 981 | 1109 | 648 |
| -5,51636E | FALSE | Low | IsMasterP | AAEL0241 | 2:1262535 | 84 | 8 | FALSE | 1 | 1 | 1 | 179 | 203 | 791 |
| -4,42012E | FALSE | Low | IsMasterP | AAEL0071 | 2:3601826 | 84 | 2 | FALSE | 1 | 1 | 1 | 794 | 921 | 869 |
| 2,39168E- | FALSE | Low | IsMasterP | AAEL0071 | 3:6990342 | 84 | 3 | FALSE | 1 | 1 | 1 | 902 | 1035 | 778 |
| 3,54399E- | FALSE | Low | IsMasterP | AAEL0139 | 2:1019219 | 84 | 5 | FALSE | 1 | 1 | 1 | 423 | 454 | 718 |
| -5,3557E+ | FALSE | Low | IsMasterP | AAEL0038 | 3:8468635 | 83 | 1 | FALSE | 1 | 1 | 1 | 1243 | 1412 | 894 |
| -3,2721E+ | FALSE | Low | IsMasterP | AAEL0075 | 1:3024752 | 83 | 3 | FALSE | 1 | 1 | 1 | 403 | 449 | 730 |
| -4,7481E+ | FALSE | Low | IsMasterP | AAEL0072 | 1:2673679 | 83 | 5 | FALSE | 1 | 1 | 1 | 274 | 315 | 592 |
| 1,18601E- | FALSE | Low | IsMasterP | AAEL0221 | 1:1841742 | 83 | 2 | FALSE | 1 | 1 | 1 | 674 | 771 | 544 |

| | | | | | | | | | | | | | | |
|---|---|---|---|---|---|---|---|---|---|---|---|---|---|---|
| -5,17903E | FALSE | Low | IsMasterP | AAEL0148 | 3:3222698 | 83 | 3 | FALSE | 1 | 1 | 1 | 316 | 342 | 477 |
| 4,10928E- | FALSE | Low | IsMasterP | AAEL0197 | 1:1888187 | 83 | 3 | FALSE | 1 | 1 | 1 | 439 | 494 | 623 |
| -8,88922E | FALSE | Low | IsMasterP | AAEL0131 | 2:3889219 | 83 | 4 | FALSE | 1 | 1 | 1 | 272 | 310 | 931 |
| 5,21974E- | FALSE | Low | IsMasterP | AAEL0196 | 1:3991058 | 83 | 2 | FALSE | 1 | 1 | 1 | 819 | 933 | 627 |
| 4,10927E- | FALSE | Low | IsMasterP | AAEL0024 | 2:7770385 | 83 | 7 | FALSE | 1 | 1 | 1 | 331 | 364 | 876 |
| 7,95216E- | FALSE | Low | IsMasterP | AAEL0139 | 3:1042915 | 84 | 2 | FALSE | 1 | 1 | 1 | 633 | 686 | 815 |
| -2,28434E | FALSE | Low | IsMasterP | AAEL0226 | 2:1574636 | 83 | 5 | FALSE | 1 | 1 | 1 | 257 | 279 | 725 |
| -3,65413E | FALSE | Low | IsMasterP | AAEL0086 | 2:2636383 | 83 | 3 | FALSE | 1 | 1 | 1 | 436 | 490 | 720 |
| -5,98373E | FALSE | Low | IsMasterP | AAEL0142 | 1:2941015 | 83 | 8 | FALSE | 1 | 1 | 1 | 230 | 258 | 847 |
| 1,57993E- | FALSE | Low | IsMasterP | AAEL0124 | 3:2904852 | 83 | 4 | FALSE | 1 | 1 | 1 | 515 | 568 | 548 |
| -8,17581E | FALSE | Low | IsMasterP | AAEL0198 | 3:7407126 | 83 | 3 | FALSE | 1 | 1 | 1 | 592 | 635 | 913 |
| -2,98206E | FALSE | Low | IsMasterP | AAEL0009 | 3:3168750 | 83 | 1 | FALSE | 1 | 1 | 1 | 731 | 844 | 479 |
| -5,42364E | FALSE | Low | IsMasterP | AAEL0058 | NIGP0100 | 83 | 2 | FALSE | 1 | 1 | 1 | 934 | 1071 | 543 |
| -3,65479E | FALSE | Low | IsMasterP | AAEL0186 | 1:2413567 | 83 | 2 | FALSE | 1 | 1 | 1 | 1033 | 1114 | 950 |
| -6,76957E | FALSE | Low | IsMasterP | AAEL0008 | 3:1201629 | 83 | 2 | FALSE | 1 | 1 | 1 | 616 | 710 | 725 |
| -8,03015E | FALSE | Low | IsMasterP | AAEL0024 | 1:1773063 | 83 | 1 | FALSE | 1 | 1 | 1 | 882 | 986 | 796 |
| -3,57503E | FALSE | Low | IsMasterP | AAEL0038 | 1:4549174 | 83 | 4 | FALSE | 1 | 1 | 1 | 334 | 377 | 945 |
| -5,21947E | FALSE | Low | IsMasterP | AAEL0072 | 2:4223836 | 83 | 5 | FALSE | 1 | 1 | 1 | 310 | 346 | 954 |
| 2,52869E- | FALSE | Low | IsMasterP | AAEL0036 | 2:3172009 | 83 | 3 | FALSE | 1 | 1 | 1 | 423 | 485 | 860 |
| 5,38569E- | FALSE | Low | IsMasterP | AAEL0260 | 2:2225732 | 83 | 6 | FALSE | 1 | 1 | 1 | 197 | 233 | 714 |
| -3,40285E | FALSE | Low | IsMasterP | AAEL0019 | 2:4246950 | 83 | 9 | FALSE | 1 | 1 | 1 | 121 | 137 | 960 |
| 6,74489E- | FALSE | Low | IsMasterP | AAEL0063 | 1:1015431 | 83 | 3 | FALSE | 1 | 1 | 1 | 437 | 473 | 944 |
| 4,77517E- | FALSE | Low | IsMasterP | AAEL0006 | 2:4586445 | 83 | 1 | FALSE | 1 | 1 | 1 | 1356 | 1477 | 797 |
| -5,52158E | FALSE | Low | IsMasterP | AAEL0033 | 2:4001860 | 83 | 6 | FALSE | 1 | 1 | 1 | 265 | 306 | 687 |
| 5,55949E- | FALSE | Low | IsMasterP | AAEL0086 | 2:3629503 | 83 | 4 | FALSE | 1 | 1 | 1 | 283 | 323 | 911 |
| -8,77703E | FALSE | Low | IsMasterP | AAEL0194 | 1:7208132 | 83 | 2 | FALSE | 1 | 1 | 1 | 522 | 574 | 925 |
| -6,36547E | FALSE | Low | IsMasterP | AAEL0019 | 2:4266014 | 83 | 8 | FALSE | 1 | 1 | 1 | 135 | 157 | 492 |
| 4,81623E- | FALSE | Low | IsMasterP | AAEL0024 | 1:9888585 | 83 | 5 | FALSE | 1 | 1 | 1 | 275 | 303 | 913 |
| 8,24018E- | FALSE | Low | IsMasterP | AAEL0095 | 2:3569731 | 83 | 5 | FALSE | 1 | 1 | 1 | 255 | 284 | 494 |
| 2,02699E- | FALSE | Low | IsMasterP | AAEL0195 | 3:3724924 | 83 | 9 | FALSE | 1 | 1 | 1 | 192 | 211 | 558 |
| -4,15661E | FALSE | Low | IsMasterP | AAEL0054 | 1:2471615 | 83 | 5 | FALSE | 1 | 1 | 1 | 321 | 362 | 505 |

| | | | | | | | | | | | | | | |
|---|---|---|---|---|---|---|---|---|---|---|---|---|---|---|
| 5,43573E- | FALSE | Low | IsMasterP | AAEL0182 | 1:1551724 | 83 | 2 | FALSE | 1 | 1 | 1 | 920 | 1051 | 634 |
| -4,07976E | FALSE | Low | IsMasterP | AAEL0102 | 2:3600115 | 83 | 7 | FALSE | 1 | 1 | 1 | 142 | 155 | 606 |
| 5,23173E- | FALSE | Low | IsMasterP | AAEL0206 | 2:4592629 | 83 | 1 | FALSE | 1 | 1 | 1 | 2444 | 2789 | 696 |
| 3,71226E- | FALSE | Low | IsMasterP | AAEL0073 | 3:1229033 | 83 | 1 | FALSE | 1 | 1 | 1 | 924 | 974 | 891 |
| -1,21878E | FALSE | Low | IsMasterP | AAEL0060 | 3:3151158 | 83 | 3 | FALSE | 1 | 1 | 1 | 378 | 421 | 699 |
| 1,36579E- | FALSE | Low | IsMasterP | AAEL0125 | 3:2423756 | 83 | 1 | FALSE | 1 | 1 | 1 | 1091 | 1212 | 526 |
| -2,40314E | FALSE | Low | IsMasterP | AAEL0018 | 2:2742887 | 83 | 6 | FALSE | 1 | 1 | 1 | 225 | 261 | 484 |
| 2,30986E- | FALSE | Low | IsMasterP | AAEL0198 | 3:3726795 | 83 | 1 | FALSE | 1 | 1 | 1 | 790 | 884 | 644 |
| 3,30501E- | FALSE | Low | IsMasterP | AAEL0136 | 1:6161171 | 83 | 2 | FALSE | 1 | 1 | 1 | 649 | 749 | 797 |
| 1,86979E- | FALSE | Low | IsMasterP | AAEL0027 | 2:1446346 | 83 | 15 | FALSE | 1 | 1 | 1 | 123 | 133 | 472 |
| -1,59319E | FALSE | Low | IsMasterP | AAEL0037 | 1:4421399 | 83 | 2 | FALSE | 1 | 1 | 1 | 552 | 620 | 765 |
| 6,88453E- | FALSE | Low | IsMasterP | AAEL0029 | 1:1086846 | 83 | 2 | FALSE | 1 | 1 | 1 | 659 | 745 | 929 |
| 9,16121E- | FALSE | Low | IsMasterP | AAEL0127 | 3:3220345 | 83 | 2 | FALSE | 1 | 1 | 1 | 619 | 701 | 846 |
| 5,57041E- | FALSE | Low | IsMasterP | AAEL0032 | 2:3667818 | 83 | 8 | FALSE | 1 | 1 | 1 | 144 | 165 | 724 |
| 8,945E+18 | FALSE | Low | IsMasterP | AAEL0195 | 3:1878274 | 83 | 2 | FALSE | 1 | 1 | 1 | 994 | 1099 | 717 |
| -5,05773E | FALSE | Low | IsMasterP | AAEL0034 | 2:1512014 | 83 | 6 | FALSE | 1 | 1 | 1 | 248 | 276 | 449 |
| 5,95401E- | FALSE | Low | IsMasterP | AAEL0054 | 3:6022766 | 83 | 2 | FALSE | 1 | 1 | 1 | 993 | 1131 | 583 |
| -8,37466E | FALSE | Low | IsMasterP | AAEL0013 | 3:4773924 | 83 | 5 | FALSE | 1 | 1 | 1 | 332 | 374 | 524 |
| 3,8557E+1 | FALSE | Low | IsMasterP | AAEL0275 | 2:3630715 | 83 | 2 | FALSE | 1 | 1 | 1 | 762 | 825 | 642 |
| -1,63177E | FALSE | Low | IsMasterP | AAEL0094 | 3:3807037 | 83 | 4 | FALSE | 1 | 1 | 1 | 364 | 389 | 521 |
| 5,40979E- | FALSE | Low | IsMasterP | AAEL0045 | 1:1449014 | 83 | 5 | FALSE | 1 | 1 | 1 | 245 | 273 | 881 |
| -3,62218E | FALSE | Low | IsMasterP | AAEL0075 | 2:1713693 | 83 | 1 | FALSE | 1 | 1 | 1 | 761 | 845 | 846 |
| -8,90341E | FALSE | Low | IsMasterP | AAEL0059 | 2:3609240 | 83 | 4 | FALSE | 1 | 1 | 1 | 393 | 435 | 734 |
| 8,84058E- | FALSE | Low | IsMasterP | AAEL0026 | NIGP0100 | 83 | 2 | FALSE | 1 | 1 | 1 | 479 | 537 | 600 |
| 2,15093E- | FALSE | Low | IsMasterP | AAEL0194 | 3:3393530 | 83 | 7 | FALSE | 1 | 1 | 1 | 182 | 203 | 541 |
| 3,10634E- | FALSE | Low | IsMasterP | AAEL0268 | 2:6487293 | 83 | 4 | FALSE | 1 | 1 | 1 | 339 | 371 | 633 |
| 9,20936E- | FALSE | Low | IsMasterP | AAEL0084 | 1:5329372 | 83 | 1 | FALSE | 1 | 1 | 1 | 1034 | 1159 | 752 |
| 3,9659E+1 | FALSE | Low | IsMasterP | AAEL0271 | 2:3323413 | 83 | 5 | FALSE | 1 | 1 | 1 | 223 | 263 | 1166 |
| -2,56097E | FALSE | Low | IsMasterP | AAEL0072 | 2:4215472 | 83 | 4 | FALSE | 1 | 1 | 1 | 306 | 336 | 868 |
| 3,71442E- | FALSE | Low | IsMasterP | AAEL0129 | 2:1248529 | 83 | 4 | FALSE | 1 | 1 | 1 | 446 | 502 | 642 |
| -7,09357E | FALSE | Low | IsMasterP | AAEL0113 | 1:1239560 | 83 | 2 | FALSE | 1 | 1 | 1 | 682 | 777 | 640 |

| | | | | | | | | | | | | | | |
|---|---|---|---|---|---|---|---|---|---|---|---|---|---|---|
| 1,7208E+1 | FALSE | Low | IsMasterP | AAEL0116 | 3:2815721 | 83 | 6 | FALSE | 1 | 1 | 1 | 199 | 227 | 815 |
| -6,46061E | FALSE | Low | IsMasterP | AAEL0105 | 1:3013326 | 83 | 4 | FALSE | 1 | 1 | 1 | 338 | 383 | 696 |
| -7,47148E | FALSE | Low | IsMasterP | AAEL0078 | 1:1491785 | 83 | 4 | FALSE | 1 | 1 | 1 | 275 | 312 | 670 |
| 3,56477E+ | FALSE | Low | IsMasterP | AAEL0213 | 2:1735346 | 83 | 1 | FALSE | 1 | 1 | 1 | 2177 | 2332 | 832 |
| -3,87932E | FALSE | Low | IsMasterP | AAEL0202 | 1:2673438 | 83 | 3 | FALSE | 1 | 1 | 1 | 393 | 445 | 600 |
| -4,60322E | FALSE | Low | IsMasterP | AAEL0175 | 2:2058275 | 83 | 14 | FALSE | 1 | 1 | 1 | 111 | 123 | 973 |
| 4,23491E+ | FALSE | Low | IsMasterP | AAEL0076 | 3:1810956 | 83 | 6 | FALSE | 1 | 1 | 1 | 140 | 159 | 851 |
| -3,58726E | FALSE | Low | IsMasterP | AAEL0071 | 3:1908056 | 83 | 14 | FALSE | 1 | 1 | 1 | 118 | 140 | 903 |
| -1,8947E+ | FALSE | Low | IsMasterP | AAEL0043 | 3:9436546 | 83 | 2 | FALSE | 1 | 1 | 1 | 524 | 588 | 906 |
| -9,0022E+ | FALSE | Low | IsMasterP | AAEL0010 | 3:5548383 | 83 | 3 | FALSE | 1 | 1 | 1 | 330 | 362 | 885 |
| -7,85804E | FALSE | Low | IsMasterP | AAEL0149 | 1:2860398 | 83 | 1 | FALSE | 1 | 1 | 1 | 750 | 773 | 868 |
| -7,46467E | FALSE | Low | IsMasterP | AAEL0072 | 3:1965494 | 83 | 1 | FALSE | 1 | 1 | 1 | 1115 | 1268 | 530 |
| 1,28905E+ | FALSE | Low | IsMasterP | AAEL0183 | 3:4032918 | 83 | 2 | FALSE | 1 | 1 | 1 | 616 | 671 | 620 |
| -4,45012E | FALSE | Low | IsMasterP | AAEL0067 | 3:2407870 | 83 | 12 | FALSE | 1 | 1 | 1 | 105 | 118 | 742 |
| -1,37393E | FALSE | Low | IsMasterP | AAEL0072 | 3:1963159 | 83 | 2 | FALSE | 1 | 1 | 1 | 507 | 581 | 813 |
| -4,06392E | FALSE | Low | IsMasterP | AAEL0211 | 1:2923845 | 83 | 9 | FALSE | 1 | 1 | 1 | 116 | 131 | 749 |
| -8,39272E | FALSE | Low | IsMasterP | AAEL0100 | NIGP0100 | 83 | 2 | FALSE | 1 | 1 | 1 | 508 | 571 | 701 |
| -4,53941E | FALSE | Low | IsMasterP | AAEL0102 | 3:1575194 | 83 | 2 | FALSE | 1 | 1 | 1 | 570 | 630 | 857 |
| 5,41037E+ | FALSE | Low | IsMasterP | AAEL0120 | 1:2704945 | 83 | 6 | FALSE | 1 | 1 | 1 | 160 | 174 | 835 |
| -1,18763E | FALSE | Low | IsMasterP | AAEL0014 | 2:4619054 | 83 | 7 | FALSE | 1 | 1 | 1 | 145 | 160 | 800 |
| -5,04697E | FALSE | Low | IsMasterP | AAEL0042 | 1:1045995 | 83 | 2 | FALSE | 1 | 1 | 1 | 978 | 1132 | 619 |
| -7,82714E | FALSE | Low | IsMasterP | AAEL0290 | 2:3167866 | 83 | 0 | FALSE | 1 | 1 | 1 | 3355 | 3806 | 731 |
| -3,43274E | FALSE | Low | IsMasterP | AAEL0014 | 1:1114609 | 83 | 2 | FALSE | 1 | 1 | 1 | 502 | 567 | 519 |
| 4,12938E+ | FALSE | Low | IsMasterP | AAEL0205 | 2:3291763 | 83 | 2 | FALSE | 1 | 1 | 1 | 818 | 926 | 573 |
| -6,16323E | FALSE | Low | IsMasterP | AAEL0108 | 2:1281468 | 83 | 3 | FALSE | 1 | 1 | 1 | 379 | 427 | 744 |
| -2,05797E | FALSE | Low | IsMasterP | AAEL0084 | 1:5341893 | 83 | 16 | FALSE | 1 | 1 | 1 | 73 | 86 | 841 |
| 7,54706E+ | FALSE | Low | IsMasterP | AAEL0057 | 2:4373833 | 83 | 4 | FALSE | 1 | 1 | 1 | 366 | 413 | 875 |
| -5,95424E | FALSE | Low | IsMasterP | AAEL0056 | 2:1431692 | 83 | 7 | FALSE | 1 | 1 | 1 | 162 | 189 | 665 |
| 2,41626E+ | FALSE | Low | IsMasterP | AAEL0271 | 3:2917179 | 83 | 1 | FALSE | 1 | 1 | 1 | 1378 | 1532 | 851 |
| 1,59754E+ | FALSE | Low | IsMasterP | AAEL0069 | 1:2956670 | 83 | 3 | FALSE | 1 | 1 | 1 | 417 | 481 | 723 |
| -3,49872E | FALSE | Low | IsMasterP | AAEL0009 | 1:2105571 | 83 | 6 | FALSE | 1 | 1 | 1 | 539 | 604 | 541 |

| | | | | | | | | | | | | | | |
|---|---|---|---|---|---|---|---|---|---|---|---|---|---|---|
| -4,31809E | FALSE | Low | IsMasterP | AAEL0204 | 2:9943472 | 83 | 1 | FALSE | 1 | 1 | 1 | 1162 | 1336 | 545 |
| -4,71886E | FALSE | Low | IsMasterP | AAEL0253 | 1:2786667 | 83 | 1 | FALSE | 1 | 1 | 1 | 3773 | 4264 | 621 |
| -8,06479E | FALSE | Low | IsMasterP | AAEL0143 | 3:4463286 | 83 | 3 | FALSE | 1 | 1 | 1 | 470 | 537 | 848 |
| 3,25506E- | FALSE | Low | IsMasterP | AAEL0073 | 3:1231215 | 83 | 1 | FALSE | 1 | 1 | 1 | 777 | 900 | 739 |
| 1,8671E+1 | FALSE | Low | IsMasterP | AAEL0009 | 3:3355990 | 83 | 13 | FALSE | 1 | 1 | 1 | 92 | 104 | 866 |
| -2,29504E | FALSE | Low | IsMasterP | AAEL0205 | 3:6211789 | 83 | 5 | FALSE | 1 | 1 | 1 | 397 | 445 | 692 |
| 3,50213E- | FALSE | Low | IsMasterP | AAEL0081 | 3:9925829 | 83 | 2 | FALSE | 1 | 1 | 1 | 414 | 452 | 514 |
| -5,87644E | FALSE | Low | IsMasterP | AAEL0104 | 1:2788906 | 83 | 2 | FALSE | 1 | 1 | 1 | 474 | 513 | 831 |
| -8,01978E | FALSE | Low | IsMasterP | AAEL0281 | 1:7319351 | 84 | 12 | FALSE | 1 | 1 | 1 | 130 | 147 | 511 |
| 8,41901E- | FALSE | Low | IsMasterP | AAEL0141 | 2:3554125 | 83 | 3 | FALSE | 1 | 1 | 1 | 554 | 632 | 775 |
| 8,20783E- | FALSE | Low | IsMasterP | AAEL0076 | 1:1422644 | 83 | 4 | FALSE | 1 | 1 | 1 | 248 | 287 | 963 |
| 3,58043E- | FALSE | Low | IsMasterP | AAEL0060 | 2:4726231 | 83 | 6 | FALSE | 1 | 1 | 1 | 190 | 207 | 747 |
| -7,90347E | FALSE | Low | IsMasterP | AAEL0022 | 2:3152461 | 83 | 1 | FALSE | 1 | 1 | 1 | 1408 | 1584 | 843 |
| -6,15905E | FALSE | Low | IsMasterP | AAEL0181 | 3:1418259 | 83 | 1 | FALSE | 1 | 1 | 1 | 1253 | 1435 | 657 |
| 1,99696E- | FALSE | Low | IsMasterP | AAEL0019 | 2:3910306 | 83 | 0 | FALSE | 1 | 1 | 1 | 3132 | 3546 | 643 |
| -1,73692E | FALSE | Low | IsMasterP | AAEL0245 | 3:4094237 | 84 | 2 | FALSE | 1 | 1 | 1 | 699 | 794 | 744 |
| 1,77804E- | FALSE | Low | IsMasterP | AAEL0030 | 2:2119916 | 84 | 2 | FALSE | 1 | 1 | 1 | 550 | 626 | 753 |
| -2,19415E | FALSE | Low | IsMasterP | AAEL0276 | 3:3056750 | 84 | 3 | FALSE | 1 | 1 | 1 | 423 | 462 | 595 |
| -5,629E+1 | FALSE | Low | IsMasterP | AAEL0104 | 3:6761196 | 84 | 2 | FALSE | 1 | 1 | 1 | 673 | 765 | 633 |
| 4,94204E- | FALSE | Low | IsMasterP | AAEL0099 | 2:3796849 | 84 | 3 | FALSE | 1 | 1 | 1 | 337 | 380 | 802 |
| 3,14851E- | FALSE | Low | IsMasterP | AAEL0023 | 2:1752630 | 84 | 2 | FALSE | 1 | 1 | 1 | 711 | 812 | 519 |
| -8,981E+1 | FALSE | Low | IsMasterP | AAEL0084 | 1:5320940 | 84 | 3 | FALSE | 1 | 1 | 1 | 545 | 597 | 676 |
| 4,19311E- | FALSE | Low | IsMasterP | AAEL0132 | 2:1859414 | 84 | 2 | FALSE | 1 | 1 | 1 | 581 | 664 | 796 |
| 8,19926E- | FALSE | Low | IsMasterP | AAEL0077 | 1:2688855 | 85 | 3 | FALSE | 1 | 1 | 1 | 595 | 664 | 715 |
| 6,0267E+1 | FALSE | Low | IsMasterP | AAEL0079 | 2:4313558 | 85 | 1 | FALSE | 1 | 1 | 1 | 914 | 1022 | 654 |
| 7,94522E- | FALSE | Low | IsMasterP | AAEL0277 | 1:8532223 | 85 | 4 | FALSE | 1 | 1 | 1 | 313 | 348 | 840 |
| -2,34879E | FALSE | Low | IsMasterP | AAEL0056 | 2:3819292 | 85 | 2 | FALSE | 1 | 1 | 1 | 477 | 549 | 881 |
| -7,66008E | FALSE | Low | IsMasterP | AAEL0228 | 3:3166608 | 85 | 4 | FALSE | 1 | 1 | 1 | 513 | 580 | 775 |
| 5,64479E- | FALSE | Low | IsMasterP | AAEL0004 | NIGP0100 | 85 | 4 | FALSE | 1 | 1 | 1 | 451 | 522 | 888 |
| 7,60001E- | FALSE | Low | IsMasterP | AAEL0052 | 1:3095806 | 85 | 1 | FALSE | 1 | 1 | 1 | 839 | 927 | 788 |
| -2,09077E | FALSE | Low | IsMasterP | AAEL0045 | 1:1452103 | 85 | 7 | FALSE | 1 | 1 | 1 | 213 | 237 | 464 |

| | | | | | | | | | | | | | | |
|---|---|---|---|---|---|---|---|---|---|---|---|---|---|---|
| -6,12501E | FALSE | Low | IsMasterP | AAEL0240 | 2:2005355 | 85 | 2 | FALSE | 1 | 1 | 1 | 875 | 984 | 923 |
| -7,40853E | FALSE | Low | IsMasterP | AAEL0025 | 3:2569507 | 86 | 9 | FALSE | 1 | 1 | 1 | 111 | 128 | 771 |
| 7,33571E+ | FALSE | Low | IsMasterP | AAEL0137 | 3:3360784 | 86 | 7 | FALSE | 1 | 1 | 1 | 179 | 206 | 680 |
| -5,65952E | FALSE | Low | IsMasterP | AAEL0135 | 2:2460133 | 86 | 2 | FALSE | 1 | 1 | 1 | 779 | 903 | 800 |
| -8,33343E | FALSE | Low | IsMasterP | AAEL0282 | 3:4074150 | 85 | 1 | FALSE | 1 | 1 | 1 | 1745 | 1800 | 891 |
| -9,0323E+ | FALSE | Low | IsMasterP | AAEL0108 | 1:5444162 | 85 | 3 | FALSE | 1 | 1 | 1 | 629 | 720 | 711 |
| 5,03772E+ | FALSE | Low | IsMasterP | AAEL0046 | 1:3072135 | 85 | 3 | FALSE | 1 | 1 | 1 | 437 | 480 | 652 |
| 3,96854E+ | FALSE | Low | IsMasterP | AAEL0051 | 3:3467924 | 85 | 6 | FALSE | 1 | 1 | 1 | 151 | 180 | 955 |
| -2,06098E | FALSE | Low | IsMasterP | AAEL0196 | 2:2258033 | 85 | 1 | FALSE | 1 | 1 | 1 | 2012 | 2275 | 661 |
| 1,4699E+1 | FALSE | Low | IsMasterP | AAEL0094 | 1:9832949 | 85 | 3 | FALSE | 1 | 1 | 1 | 292 | 297 | 884 |
| -5,56766E | FALSE | Low | IsMasterP | AAEL0234 | 2:1345764 | 85 | 1 | FALSE | 1 | 1 | 1 | 750 | 854 | 925 |
| -7,05948E | FALSE | Low | IsMasterP | AAEL0135 | 2:2461275 | 85 | 2 | FALSE | 1 | 1 | 1 | 494 | 551 | 560 |
| 3,11332E+ | FALSE | Low | IsMasterP | AAEL0028 | 1:1081802 | 85 | 6 | FALSE | 1 | 1 | 1 | 179 | 192 | 960 |
| 5,29494E+ | FALSE | Low | IsMasterP | AAEL0105 | 2:2618307 | 85 | 2 | FALSE | 1 | 1 | 1 | 729 | 842 | 531 |
| 8,0843E+1 | FALSE | Low | IsMasterP | AAEL0114 | 2:3743146 | 85 | 10 | FALSE | 1 | 1 | 1 | 105 | 115 | 456 |
| 8,21998E+ | FALSE | Low | IsMasterP | AAEL0186 | MT:11547 | 85 | 5 | FALSE | 1 | 1 | 1 | 313 | 361 | 856 |
| 1,17222E+ | FALSE | Low | IsMasterP | AAEL0106 | 2:1361085 | 85 | 4 | FALSE | 1 | 1 | 1 | 425 | 498 | 840 |
| 8,26331E+ | FALSE | Low | IsMasterP | AAEL0059 | 3:1386107 | 85 | 12 | FALSE | 1 | 1 | 1 | 86 | 102 | 1024 |
| -1,47097E | FALSE | Low | IsMasterP | AAEL0209 | 2:2550346 | 85 | 3 | FALSE | 1 | 1 | 1 | 799 | 878 | 765 |
| 6,99813E+ | FALSE | Low | IsMasterP | AAEL0069 | 3:1437500 | 85 | 4 | FALSE | 1 | 1 | 1 | 283 | 324 | 911 |
| -5,20242E | FALSE | Low | IsMasterP | AAEL0103 | 2:4041472 | 85 | 2 | FALSE | 1 | 1 | 1 | 571 | 642 | 841 |
| -6,24648E | FALSE | Low | IsMasterP | AAEL0014 | 1:1117505 | 85 | 3 | FALSE | 1 | 1 | 1 | 521 | 601 | 865 |
| -1,81206E | FALSE | Low | IsMasterP | AAEL0080 | 2:2306247 | 85 | 1 | FALSE | 1 | 1 | 1 | 1183 | 1341 | 602 |
| -5,00206E | FALSE | Low | IsMasterP | AAEL0139 | 3:1063934 | 85 | 8 | FALSE | 1 | 1 | 1 | 134 | 155 | 966 |
| 3,63609E+ | FALSE | Low | IsMasterP | AAEL0094 | 3:3807930 | 85 | 4 | FALSE | 1 | 1 | 1 | 462 | 526 | 778 |
| 1,02542E+ | FALSE | Low | IsMasterP | AAEL0017 | 2:3778328 | 85 | 5 | FALSE | 1 | 1 | 1 | 453 | 492 | 737 |
| -4,65554E | FALSE | Low | IsMasterP | AAEL0007 | 2:4593917 | 85 | 1 | FALSE | 1 | 1 | 1 | 849 | 942 | 884 |
| -4,95284E | FALSE | Low | IsMasterP | AAEL0093 | 2:4517095 | 85 | 4 | FALSE | 1 | 1 | 1 | 213 | 245 | 926 |
| 7,84853E+ | FALSE | Low | IsMasterP | AAEL0088 | 2:2408963 | 85 | 2 | FALSE | 1 | 1 | 1 | 773 | 864 | 846 |
| 9,52381E+ | FALSE | Low | IsMasterP | AAEL0101 | 3:1492481 | 86 | 6 | FALSE | 1 | 1 | 1 | 163 | 192 | 1059 |
| -3,11511E | FALSE | Low | IsMasterP | AAEL0142 | 2:3315148 | 86 | 1 | FALSE | 1 | 1 | 1 | 1730 | 1985 | 794 |

| | | | | | | | | | | | | | | |
|---|---|---|---|---|---|---|---|---|---|---|---|---|---|---|
| -5,70834E | FALSE | Low | IsMasterP | AAEL0093 | 3:887890( | 86 | 2 | FALSE | 1 | 1 | 1 | 691 | 829 | 597 |
| 8,41326E- | FALSE | Low | IsMasterP | AAEL0016 | 1:138848( | 86 | 11 | FALSE | 1 | 1 | 1 | 140 | 162 | 910 |
| -1,21121E | FALSE | Low | IsMasterP | AAEL0183 | 3:187219( | 86 | 1 | FALSE | 1 | 1 | 1 | 2107 | 2289 | 879 |
| 3,02777E- | FALSE | Low | IsMasterP | AAEL0067 | 3:169205: | 86 | 4 | FALSE | 1 | 1 | 1 | 593 | 663 | 600 |
| 5,74023E- | FALSE | Low | IsMasterP | AAEL0144 | 1:159066: | 88 | 1 | FALSE | 1 | 1 | 1 | 1784 | 1986 | 594 |
| -6,73513E | FALSE | Low | IsMasterP | AAEL0120 | 1:300976: | 88 | 1 | FALSE | 1 | 1 | 1 | 1050 | 1210 | 592 |
| 2,41648E- | FALSE | Low | IsMasterP | AAEL0061 | 1:265979 | 88 | 12 | FALSE | 1 | 1 | 1 | 113 | 132 | 1027 |
| -2,2418E+ | FALSE | Low | IsMasterP | AAEL0024 | 1:177365 | 88 | 12 | FALSE | 1 | 1 | 1 | 141 | 161 | 1024 |
| -8,12765E | FALSE | Low | IsMasterP | AAEL0106 | 1:423593 | 88 | 4 | FALSE | 1 | 1 | 1 | 213 | 243 | 680 |
| -3,64432E | FALSE | Low | IsMasterP | AAEL0096 | 3:209901 | 88 | 3 | FALSE | 1 | 1 | 1 | 395 | 458 | 784 |
| -3,66988E | FALSE | Low | IsMasterP | AAEL0078 | 3:151666 | 88 | 1 | FALSE | 1 | 1 | 1 | 743 | 833 | 815 |
| -7,3819E+ | FALSE | Low | IsMasterP | AAEL0109 | 3:167650 | 88 | 2 | FALSE | 1 | 1 | 1 | 678 | 773 | 543 |
| -5,40417E | FALSE | Low | IsMasterP | AAEL0195 | 3:177415: | 88 | 1 | FALSE | 1 | 1 | 1 | 2785 | 3136 | 620 |
| -8,95292E | FALSE | Low | IsMasterP | AAEL0130 | 1:199396 | 89 | 4 | FALSE | 1 | 1 | 1 | 648 | 727 | 689 |
| 7,41048E- | FALSE | Low | IsMasterP | AAEL0124 | 3:290290 | 89 | 1 | FALSE | 1 | 1 | 1 | 969 | 1084 | 900 |
| -6,13947E | FALSE | Low | IsMasterP | AAEL0197 | 2:347630 | 89 | 3 | FALSE | 1 | 1 | 1 | 597 | 662 | 834 |
| 6,99868E- | FALSE | Low | IsMasterP | AAEL0007 | 2:521254 | 89 | 2 | FALSE | 1 | 1 | 1 | 554 | 636 | 914 |
| -3,17515E | FALSE | Low | IsMasterP | AAEL0111 | 3:193553: | 89 | 6 | FALSE | 1 | 1 | 1 | 432 | 510 | 486 |
| 1,94638E- | FALSE | Low | IsMasterP | AAEL0099 | 3:319318 | 89 | 10 | FALSE | 1 | 1 | 1 | 339 | 383 | 910 |
| 9,19184E- | FALSE | Low | IsMasterP | AAEL0074 | 2:291555 | 90 | 1 | FALSE | 1 | 1 | 1 | 1541 | 1643 | 634 |
| -3,78403E | FALSE | Low | IsMasterP | AAEL0268 | 2:468586: | 90 | 1 | FALSE | 1 | 1 | 1 | 1311 | 1454 | 717 |
| -5,78267E | FALSE | Low | IsMasterP | AAEL0186 | MT:9798:: | 90 | 12 | FALSE | 1 | 1 | 1 | 173 | 202 | 948 |
| -7,17367E | FALSE | Low | IsMasterP | AAEL0198 | 3:310747 | 90 | 1 | FALSE | 1 | 1 | 1 | 2619 | 2938 | 564 |
| -7,4211E+ | FALSE | Low | IsMasterP | A0A5C4TI | Cytochror | 90 | 2 | FALSE | 1 | 1 | 1 | 695 | 783 | 892 |
| 5,17862E- | FALSE | Low | IsMasterP | AAEL0267 | 1:131643 | 90 | 4 | FALSE | 1 | 1 | 1 | 420 | 439 | 869 |
| -5,60365E | FALSE | Low | IsMasterP | AAEL0084 | 1:171511: | 90 | 2 | FALSE | 1 | 1 | 1 | 495 | 551 | 652 |
| -1,91112E | FALSE | Low | IsMasterP | AAEL0134 | 1:206397 | 90 | 2 | FALSE | 1 | 1 | 1 | 366 | 418 | 654 |
| 1,80658E- | FALSE | Low | IsMasterP | AAEL0070 | 2:224446: | 90 | 8 | FALSE | 1 | 1 | 1 | 272 | 285 | 780 |
| 9,12272E- | FALSE | Low | IsMasterP | AAEL0128 | 3:755663 | 90 | 9 | FALSE | 1 | 1 | 1 | 162 | 186 | 676 |
| -5,47674E | FALSE | Low | IsMasterP | AAEL0015 | 2:209540 | 90 | 1 | FALSE | 1 | 1 | 1 | 1905 | 2132 | 535 |
| 3,3324E+: | FALSE | Low | IsMasterP | AAEL0108 | 3:516836 | 90 | 11 | FALSE | 1 | 1 | 1 | 98 | 108 | 689 |

| | | | | | | | | | | | | | | |
|---|---|---|---|---|---|---|---|---|---|---|---|---|---|---|
| -7,82779E | FALSE | Low | IsMasterP | AAEL0037 | 2:1974769 | 91 | 4 | FALSE | 1 | 1 | 1 | 462 | 500 | 730 |
| 1,53629E+ | FALSE | Low | IsMasterP | AAEL0011 | 2:3072007 | 91 | 3 | FALSE | 1 | 1 | 1 | 319 | 371 | 554 |
| -3,80577E | FALSE | Low | IsMasterP | AAEL0064 | 3:2164656 | 91 | 3 | FALSE | 1 | 1 | 1 | 1005 | 1136 | 828 |
| 8,57944E+ | FALSE | Low | IsMasterP | AAEL0126 | 3:2838650 | 91 | 7 | FALSE | 1 | 1 | 1 | 226 | 245 | 473 |
| 4,94789E+ | FALSE | Low | IsMasterP | AAEL0197 | 2:2388344 | 91 | 1 | FALSE | 1 | 1 | 1 | 2256 | 2537 | 680 |
| -8,89902E | FALSE | Low | IsMasterP | AAEL0248 | 2:4288440 | 92 | 20 | FALSE | 1 | 1 | 1 | 133 | 132 | 832 |
| 7,52224E+ | FALSE | Low | IsMasterP | AAEL0084 | 2:3952444 | 92 | 2 | FALSE | 1 | 1 | 1 | 882 | 1034 | 545 |
| 1,21921E+ | FALSE | Low | IsMasterP | AAEL0218 | 2:9867343 | 92 | 2 | FALSE | 1 | 1 | 1 | 1044 | 1102 | 840 |
| -1,48365E | FALSE | Low | IsMasterP | AAEL0174 | 2:2704904 | 92 | 9 | FALSE | 1 | 1 | 1 | 165 | 188 | 932 |
| -4,80824E | FALSE | Low | IsMasterP | AAEL0142 | 1:7235146 | 92 | 9 | FALSE | 1 | 1 | 1 | 142 | 166 | 566 |
| -6,00004E | FALSE | Low | IsMasterP | AAEL0282 | 3:2465286 | 92 | 1 | FALSE | 1 | 1 | 1 | 1941 | 2154 | 906 |
| 4,65353E+ | FALSE | Low | IsMasterP | AAEL0038 | 1:6568693 | 92 | 1 | FALSE | 1 | 1 | 1 | 1625 | 1813 | 834 |
| -5,90739E | FALSE | Low | IsMasterP | AAEL0054 | 3:3832920 | 92 | 2 | FALSE | 1 | 1 | 1 | 601 | 658 | 816 |
| -7,40183E | FALSE | Low | IsMasterP | AAEL0087 | 3:3260055 | 92 | 5 | FALSE | 1 | 1 | 1 | 173 | 200 | 474 |
| -1,40037E | FALSE | Low | IsMasterP | AAEL0269 | 1:1192355 | 92 | 1 | FALSE | 1 | 1 | 1 | 915 | 1058 | 640 |
| -6,39559E | FALSE | Low | IsMasterP | AAEL0098 | 3:1218371 | 92 | 1 | FALSE | 1 | 1 | 1 | 1646 | 1793 | 854 |
| -6,98364E | FALSE | Low | IsMasterP | AAEL0005 | 3:1249979 | 92 | 2 | FALSE | 1 | 1 | 1 | 718 | 800 | 869 |
| 6,56439E+ | FALSE | Low | IsMasterP | AAEL0069 | 2:2419247 | 92 | 2 | FALSE | 1 | 1 | 1 | 859 | 968 | 510 |
| 3,47268E+ | FALSE | Low | IsMasterP | AAEL0204 | 3:8752600 | 92 | 13 | FALSE | 1 | 1 | 1 | 87 | 103 | 872 |
| 2,95657E+ | FALSE | Low | IsMasterP | AAEL0081 | 3:9071099 | 92 | 6 | FALSE | 1 | 1 | 1 | 137 | 154 | 1078 |
| 7,52913E+ | FALSE | Low | IsMasterP | AAEL0002 | 2:2273424 | 93 | 2 | FALSE | 1 | 1 | 1 | 520 | 583 | 791 |
| -5,86333E | FALSE | Low | IsMasterP | AAEL0127 | 3:1117485 | 93 | 3 | FALSE | 1 | 1 | 1 | 288 | 327 | 602 |
| -8,99845E | FALSE | Low | IsMasterP | AAEL0143 | 3:4465360 | 93 | 5 | FALSE | 1 | 1 | 1 | 176 | 192 | 507 |
| -6,82301E | FALSE | Low | IsMasterP | AAEL0233 | 1:8031130 | 93 | 4 | FALSE | 1 | 1 | 1 | 757 | 847 | 736 |
| -3,45454E | FALSE | Low | IsMasterP | AAEL0078 | 3:3822100 | 93 | 3 | FALSE | 1 | 1 | 1 | 540 | 616 | 901 |
| -7,67494E | FALSE | Low | IsMasterP | AAEL0261 | NIGP0100 | 93 | 3 | FALSE | 1 | 1 | 1 | 411 | 474 | 895 |
| 2,36E+18 | FALSE | Low | IsMasterP | AAEL0203 | 1:3030975 | 93 | 10 | FALSE | 1 | 1 | 1 | 185 | 216 | 484 |
| -6,72112E | FALSE | Low | IsMasterP | AAEL0206 | 2:9923190 | 94 | 2 | FALSE | 1 | 1 | 1 | 690 | 777 | 983 |
| 3,2956E+1 | FALSE | Low | IsMasterP | AAEL0227 | 2:2211305 | 94 | 2 | FALSE | 1 | 1 | 1 | 496 | 537 | 783 |
| 4,69324E+ | FALSE | Low | IsMasterP | AAEL0015 | 2:2241227 | 94 | 4 | FALSE | 1 | 1 | 1 | 296 | 332 | 525 |
| 7,52825E+ | FALSE | Low | IsMasterP | AAEL0211 | 3:1706226 | 94 | 3 | FALSE | 1 | 1 | 1 | 677 | 784 | 677 |

| | | | | | | | | | | | | | | |
|---|---|---|---|---|---|---|---|---|---|---|---|---|---|---|
| 7,40394E- | FALSE | Low | IsMasterP | AAEL0072 | 2:4218105 | 95 | 1 | FALSE | 1 | 1 | 1 | 2379 | 2644 | 720 |
| 1,59052E- | FALSE | Low | IsMasterP | AAEL0028 | 3:2349457 | 95 | 5 | FALSE | 1 | 1 | 1 | 192 | 219 | 841 |
| -6,79793E | FALSE | Low | IsMasterP | AAEL0022 | 3:3755719 | 95 | 5 | FALSE | 1 | 1 | 1 | 215 | 243 | 597 |
| 3,72105E- | FALSE | Low | IsMasterP | AAEL0035 | 3:3294040 | 95 | 3 | FALSE | 1 | 1 | 1 | 748 | 858 | 646 |
| -3,40155E | FALSE | Low | IsMasterP | AAEL0130 | 2:2533642 | 95 | 1 | FALSE | 1 | 1 | 1 | 641 | 713 | 567 |
| 5,77172E- | FALSE | Low | IsMasterP | AAEL0051 | 2:1430950 | 95 | 3 | FALSE | 1 | 1 | 1 | 579 | 647 | 714 |
| -4,82589E | FALSE | Low | IsMasterP | AAEL0036 | 2:3171445 | 95 | 12 | FALSE | 1 | 1 | 1 | 110 | 124 | 829 |
| 1,81203E- | FALSE | Low | IsMasterP | AAEL0131 | 2:1804462 | 96 | 4 | FALSE | 1 | 1 | 1 | 478 | 536 | 788 |
| 7,50989E- | FALSE | Low | IsMasterP | AAEL0205 | 1:1958765 | 96 | 2 | FALSE | 1 | 1 | 1 | 1017 | 1125 | 724 |
| -5,95631E | FALSE | Low | IsMasterP | AAEL0066 | 2:2529167 | 96 | 4 | FALSE | 1 | 1 | 1 | 205 | 233 | 483 |
| -4,80088E | FALSE | Low | IsMasterP | AAEL0062 | 3:6250617 | 96 | 5 | FALSE | 1 | 1 | 1 | 498 | 573 | 906 |
| 6,74538E- | FALSE | Low | IsMasterP | AAEL0199 | 2:2051912 | 96 | 1 | FALSE | 1 | 1 | 1 | 2342 | 2649 | 634 |
| 6,14363E- | FALSE | Low | IsMasterP | AAEL0151 | 2:3351327 | 96 | 3 | FALSE | 1 | 1 | 1 | 762 | 864 | 508 |
| -6,89088E | FALSE | Low | IsMasterP | AAEL0195 | 3:2355592 | 96 | 0 | FALSE | 1 | 1 | 1 | 5973 | 6529 | 750 |
| 7,36142E- | FALSE | Low | IsMasterP | AAEL0026 | 3:6660608 | 97 | 2 | FALSE | 1 | 1 | 1 | 329 | 382 | 906 |
| -6,76207E | FALSE | Low | IsMasterP | AAEL0074 | 2:1232220 | 97 | 1 | FALSE | 1 | 1 | 1 | 1804 | 2006 | 652 |
| -3,36122E | FALSE | Low | IsMasterP | AAEL0147 | 1:2349539 | 97 | 3 | FALSE | 1 | 1 | 1 | 388 | 431 | 539 |
| -3,22127E | FALSE | Low | IsMasterP | AAEL0009 | 3:3770692 | 97 | 1 | FALSE | 1 | 1 | 1 | 812 | 902 | 774 |
| 6,94093E- | FALSE | Low | IsMasterP | AAEL0010 | 1:2997035 | 97 | 3 | FALSE | 1 | 1 | 1 | 650 | 727 | 747 |
| 1,7159E+1 | FALSE | Low | IsMasterP | AAEL0236 | 3:3484168 | 97 | 2 | FALSE | 1 | 1 | 1 | 852 | 956 | 835 |
| 9,16784E- | FALSE | Low | IsMasterP | AAEL0003 | 2:3696398 | 97 | 2 | FALSE | 1 | 1 | 1 | 859 | 982 | 721 |
| 5,24158E- | FALSE | Low | IsMasterP | AAEL0082 | 3:2502272 | 98 | 4 | FALSE | 1 | 1 | 1 | 308 | 348 | 846 |
| 1,36664E- | FALSE | Low | IsMasterP | AAEL0001 | 3:1095486 | 98 | 8 | FALSE | 1 | 1 | 1 | 202 | 217 | 879 |
| -3,65845E | FALSE | Low | IsMasterP | AAEL0078 | 1:2896842 | 98 | 2 | FALSE | 1 | 1 | 1 | 601 | 694 | 611 |
| 2,63097E- | FALSE | Low | IsMasterP | AAEL0248 | 2:3190317 | 99 | 2 | FALSE | 1 | 1 | 1 | 1175 | 1295 | 673 |
| -2,04383E | FALSE | Low | IsMasterP | AAEL0266 | 3:1074613 | 99 | 6 | FALSE | 1 | 1 | 1 | 310 | 348 | 749 |
| -1,76216E | FALSE | Low | IsMasterP | AAEL0182 | 3:4879735 | 99 | 1 | FALSE | 1 | 1 | 1 | 1182 | 1337 | 838 |
| 2,14827E- | FALSE | Low | IsMasterP | AAEL0029 | 1:8594666 | 100 | 3 | FALSE | 1 | 1 | 1 | 336 | 391 | 695 |
| -6,84349E | FALSE | Low | IsMasterP | AAEL0053 | 2:2713861 | 100 | 1 | FALSE | 1 | 1 | 1 | 685 | 773 | 543 |
| -6,61777E | FALSE | Low | IsMasterP | AAEL0082 | 3:2505305 | 100 | 4 | FALSE | 1 | 1 | 1 | 302 | 343 | 925 |
| -7,37365E | FALSE | Low | IsMasterP | AAEL0231 | 2:3336834 | 100 | 2 | FALSE | 1 | 1 | 1 | 901 | 1012 | 753 |

| | | | | | | | | | | | | | | |
|---|---|---|---|---|---|---|---|---|---|---|---|---|---|---|
| 6,14812E+ | FALSE | Low | IsMasterP | AAEL0196 | 1:3079688 | 100 | 1 | FALSE | 1 | 1 | 1 | 934 | 1026 | 501 |
| -2,50127E | FALSE | Low | IsMasterP | AAEL0273 | 2:4317458 | 100 | 2 | FALSE | 1 | 1 | 1 | 964 | 1116 | 769 |
| -9,07735E | FALSE | Low | IsMasterP | AAEL0030 | 2:2121522 | 100 | 3 | FALSE | 1 | 1 | 1 | 497 | 553 | 505 |
| -6,57917E | FALSE | Low | IsMasterP | AAEL0029 | 1:8598981 | 101 | 3 | FALSE | 1 | 1 | 1 | 913 | 1034 | 585 |
| 1,80651E+ | FALSE | Low | IsMasterP | AAEL0173 | 2:4006154 | 101 | 6 | FALSE | 1 | 1 | 1 | 318 | 346 | 796 |
| -3,06855E | FALSE | Low | IsMasterP | AAEL0050 | 2:4451744 | 101 | 1 | FALSE | 1 | 1 | 1 | 1274 | 1425 | 815 |
| -5,50377E | FALSE | Low | IsMasterP | AAEL0047 | 2:3781200 | 101 | 1 | FALSE | 1 | 1 | 1 | 2002 | 2258 | 613 |
| -7,27816E | FALSE | Low | IsMasterP | AAEL0195 | 3:3739958 | 102 | 1 | FALSE | 1 | 1 | 1 | 1809 | 1981 | 865 |
| 2,71539E+ | FALSE | Low | IsMasterP | AAEL0020 | 3:1783929 | 102 | 2 | FALSE | 1 | 1 | 1 | 646 | 730 | 597 |
| -2,95684E | FALSE | Low | IsMasterP | AAEL0195 | 2:2443097 | 103 | 3 | FALSE | 1 | 1 | 1 | 659 | 745 | 652 |
| -7,25825E | FALSE | Low | IsMasterP | AAEL0016 | 1:1380031 | 103 | 5 | FALSE | 1 | 1 | 1 | 334 | 377 | 733 |
| -3,35827E | FALSE | Low | IsMasterP | AAEL0114 | 3:3290648 | 103 | 4 | FALSE | 1 | 1 | 1 | 414 | 465 | 787 |
| -1,15887E | FALSE | Low | IsMasterP | AAEL0102 | 3:2230890 | 103 | 2 | FALSE | 1 | 1 | 1 | 697 | 783 | 647 |
| -8,2836E+ | FALSE | Low | IsMasterP | AAEL0258 | 2:4611057 | 103 | 1 | FALSE | 1 | 1 | 1 | 1906 | 2192 | 627 |
| 3,89351E+ | FALSE | Low | IsMasterP | AAEL0124 | 3:1258157 | 103 | 3 | FALSE | 1 | 1 | 1 | 426 | 478 | 1014 |
| -2,03641E | FALSE | Low | IsMasterP | AAEL0001 | 3:2146129 | 103 | 1 | FALSE | 1 | 1 | 1 | 2103 | 2379 | 778 |
| -3,53133E | FALSE | Low | IsMasterP | AAEL0072 | 3:1967340 | 103 | 4 | FALSE | 1 | 1 | 1 | 289 | 339 | 1004 |
| -2,2199E+ | FALSE | Low | IsMasterP | AAEL0062 | 2:2212559 | 103 | 3 | FALSE | 1 | 1 | 1 | 636 | 713 | 979 |
| -6,67788E | FALSE | Low | IsMasterP | AAEL0078 | 2:4743469 | 103 | 3 | FALSE | 1 | 1 | 1 | 521 | 589 | 733 |
| 8,75777E+ | FALSE | Low | IsMasterP | AAEL0141 | 1:2725605 | 103 | 3 | FALSE | 1 | 1 | 1 | 321 | 368 | 605 |
| -8,65705E | FALSE | Low | IsMasterP | AAEL0262 | 3:3967495 | 103 | 1 | FALSE | 1 | 1 | 1 | 1664 | 1820 | 574 |
| -6,92018E | FALSE | Low | IsMasterP | AAEL0264 | 2:2312591 | 104 | 2 | FALSE | 1 | 1 | 1 | 725 | 810 | 778 |
| -3,40812E | FALSE | Low | IsMasterP | AAEL0052 | 3:3102266 | 104 | 3 | FALSE | 1 | 1 | 1 | 531 | 609 | 951 |
| -2,52599E | FALSE | Low | IsMasterP | AAEL0019 | 2:4259454 | 104 | 12 | FALSE | 1 | 1 | 1 | 168 | 190 | 491 |
| -8,26754E | FALSE | Low | IsMasterP | AAEL0226 | 2:2093096 | 104 | 4 | FALSE | 1 | 1 | 1 | 296 | 333 | 655 |
| -8,82305E | FALSE | Low | IsMasterP | AAEL0009 | 3:3802787 | 104 | 3 | FALSE | 1 | 1 | 1 | 741 | 831 | 604 |
| -1,6602E+ | FALSE | Low | IsMasterP | AAEL0092 | 2:2685862 | 104 | 1 | FALSE | 1 | 1 | 1 | 683 | 781 | 869 |
| 4,47552E+ | FALSE | Low | IsMasterP | AAEL0111 | 3:1941399 | 104 | 4 | FALSE | 1 | 1 | 1 | 229 | 254 | 870 |
| -5,5744E+ | FALSE | Low | IsMasterP | AAEL0269 | 3:4231539 | 104 | 1 | FALSE | 1 | 1 | 1 | 1640 | 1839 | 743 |
| 8,57255E+ | FALSE | Low | IsMasterP | AAEL0274 | 1:2455259 | 105 | 4 | FALSE | 1 | 1 | 1 | 554 | 640 | 573 |
| -6,23307E | FALSE | Low | IsMasterP | AAEL0242 | 2:2142078 | 105 | 5 | FALSE | 1 | 1 | 1 | 573 | 655 | 894 |

| | | | | | | | | | | | | | |
|---|---|---|---|---|---|---|---|---|---|---|---|---|---|
| 7,3455E+1 | FALSE | Low | IsMasterP AAEL0119 | 2:2178242 | 105 | 1 | FALSE | 1 | 1 | 1 | 764 | 866 | 635 |
| 2,84523E- | FALSE | Low | IsMasterP AAEL0005 | 3:1628317 | 105 | 1 | FALSE | 1 | 1 | 1 | 917 | 1022 | 611 |
| -8,53597E | FALSE | Low | IsMasterP AAEL0082 | 3:8752639 | 105 | 8 | FALSE | 1 | 1 | 1 | 120 | 135 | 460 |
| 6,86602E- | FALSE | Low | IsMasterP AAEL0273 | 3:2742830 | 105 | 1 | FALSE | 1 | 1 | 1 | 1718 | 1931 | 604 |
| 7,93301E- | FALSE | Low | IsMasterP AAEL0116 | 3:2816911 | 105 | 3 | FALSE | 1 | 1 | 1 | 778 | 877 | 774 |
| 4,86993E- | FALSE | Low | IsMasterP AAEL0197 | 2:6194826 | 105 | 2 | FALSE | 1 | 1 | 1 | 640 | 710 | 933 |
| -3,29491E | FALSE | Low | IsMasterP AAEL0127 | 1:2030282 | 105 | 20 | FALSE | 1 | 1 | 1 | 105 | 113 | 494 |
| -1,35592E | FALSE | Low | IsMasterP AAEL0030 | 2:2126302 | 105 | 4 | FALSE | 1 | 1 | 1 | 370 | 438 | 961 |
| -2,8328E+ | FALSE | Low | IsMasterP AAEL0027 | 1:1774839 | 105 | 6 | FALSE | 1 | 1 | 1 | 141 | 160 | 821 |
| 6,23462E- | FALSE | Low | IsMasterP AAEL0087 | 3:1326792 | 108 | 2 | FALSE | 1 | 1 | 1 | 361 | 400 | 916 |
| -2,21211E | FALSE | Low | IsMasterP AAEL0004 | 3:1836168 | 108 | 2 | FALSE | 1 | 1 | 1 | 392 | 444 | 465 |
| 6,56197E- | FALSE | Low | IsMasterP AAEL0182 | 2:2097687 | 109 | 1 | FALSE | 1 | 1 | 1 | 688 | 771 | 807 |
| -4,83207E | FALSE | Low | IsMasterP AAEL0052 | 1:1814488 | 109 | 1 | FALSE | 1 | 1 | 1 | 2889 | 3301 | 583 |
| -1,13253E | FALSE | Low | IsMasterP AAEL0033 | 3:8132564 | 109 | 1 | FALSE | 1 | 1 | 1 | 1602 | 1804 | 903 |
| -2,71523E | FALSE | Low | IsMasterP AAEL0004 | 3:1628063 | 110 | 2 | FALSE | 1 | 1 | 1 | 946 | 1072 | 687 |
| -8,0155E+ | FALSE | Low | IsMasterP AAEL0109 | 3:1682322 | 110 | 7 | FALSE | 1 | 1 | 1 | 236 | 271 | 933 |
| -2,5482E+ | FALSE | Low | IsMasterP AAEL0211 | 2:1007833 | 110 | 1 | FALSE | 1 | 1 | 1 | 1208 | 1349 | 781 |
| -1,19454E | FALSE | Low | IsMasterP AAEL0095 | 3:3448510 | 110 | 4 | FALSE | 1 | 1 | 1 | 659 | 730 | 740 |
| 6,47396E- | FALSE | Low | IsMasterP AAEL0092 | 3:3909582 | 110 | 2 | FALSE | 1 | 1 | 1 | 892 | 1028 | 673 |
| 8,33081E- | FALSE | Low | IsMasterP AAEL0069 | 3:1437683 | 110 | 4 | FALSE | 1 | 1 | 1 | 307 | 346 | 453 |
| 9,68031E- | FALSE | Low | IsMasterP AAEL0200 | 2:4465403 | 110 | 3 | FALSE | 1 | 1 | 1 | 634 | 736 | 821 |
| 6,35242E- | FALSE | Low | IsMasterP AAEL0195 | 1:2922829 | 110 | 5 | FALSE | 1 | 1 | 1 | 190 | 221 | 923 |
| -8,2123E+ | FALSE | Low | IsMasterP AAEL0230 | 3:2879692 | 110 | 4 | FALSE | 1 | 1 | 1 | 326 | 359 | 768 |
| 4,59284E- | FALSE | Low | IsMasterP AAEL0138 | 1:3585106 | 111 | 4 | FALSE | 1 | 1 | 1 | 187 | 216 | 559 |
| 6,17416E- | FALSE | Low | IsMasterP AAEL0094 | 3:3450204 | 111 | 2 | FALSE | 1 | 1 | 1 | 1543 | 1730 | 837 |
| -2,90539E | FALSE | Low | IsMasterP AAEL0031 | 3:2700726 | 111 | 7 | FALSE | 1 | 1 | 1 | 328 | 362 | 901 |
| 6,70014E- | FALSE | Low | IsMasterP AAEL0080 | 2:2308288 | 111 | 5 | FALSE | 1 | 1 | 1 | 509 | 581 | 571 |
| 7,40278E- | FALSE | Low | IsMasterP AAEL0206 | 3:9498379 | 111 | 1 | FALSE | 1 | 1 | 1 | 1797 | 2015 | 646 |
| -5,5187E+ | FALSE | Low | IsMasterP AAEL0290 | 1:8456804 | 112 | 15 | FALSE | 1 | 1 | 1 | 59 | 62 | 1024 |
| 5,48076E- | FALSE | Low | IsMasterP AAEL0066 | 2:3265141 | 112 | 5 | FALSE | 1 | 1 | 1 | 296 | 337 | 689 |
| 2,82033E- | FALSE | Low | IsMasterP AAEL0052 | 1:3043667 | 112 | 3 | FALSE | 1 | 1 | 1 | 719 | 798 | 482 |

| | | | | | | | | | | | | | | |
|---|---|---|---|---|---|---|---|---|---|---|---|---|---|---|
| 1,51441E- | FALSE | Low | IsMasterP | AAEL0171 | 1:1360014 | 112 | 8 | FALSE | 1 | 1 | 1 | 91 | 106 | 1174 |
| 6,2387E+1 | FALSE | Low | IsMasterP | AAEL0249 | NIGP0100 | 113 | 7 | FALSE | 1 | 1 | 1 | 433 | 495 | 885 |
| 2,42752E- | FALSE | Low | IsMasterP | AAEL0008 | 3:1201349 | 113 | 1 | FALSE | 1 | 1 | 1 | 1880 | 2166 | 647 |
| 5,93942E- | FALSE | Low | IsMasterP | AAEL0144 | 2:3492465 | 113 | 2 | FALSE | 1 | 1 | 1 | 674 | 749 | 721 |
| 4,65922E- | FALSE | Low | IsMasterP | AAEL0059 | 2:3608848 | 113 | 1 | FALSE | 1 | 1 | 1 | 1931 | 2122 | 865 |
| 2,44922E- | FALSE | Low | IsMasterP | AAEL0053 | 3:3474398 | 113 | 2 | FALSE | 1 | 1 | 1 | 387 | 446 | 882 |
| 3,39355E- | FALSE | Low | IsMasterP | AAEL0136 | 3:1952983 | 114 | 6 | FALSE | 1 | 1 | 1 | 357 | 413 | 853 |
| -5,70853E | FALSE | Low | IsMasterP | AAEL0126 | 3:2838385 | 114 | 6 | FALSE | 1 | 1 | 1 | 287 | 322 | 559 |
| -4,5698E+ | FALSE | Low | IsMasterP | AAEL0101 | 2:2647118 | 114 | 4 | FALSE | 1 | 1 | 1 | 269 | 307 | 743 |
| 3,3218E+1 | FALSE | Low | IsMasterP | AAEL0022 | 2:4306353 | 114 | 4 | FALSE | 1 | 1 | 1 | 296 | 329 | 610 |
| -5,36381E | FALSE | Low | IsMasterP | AAEL0180 | 3:1511650 | 114 | 2 | FALSE | 1 | 1 | 1 | 858 | 975 | 813 |
| 3,96851E- | FALSE | Low | IsMasterP | AAEL0273 | 2:4245687 | 115 | 2 | FALSE | 1 | 1 | 1 | 668 | 776 | 730 |
| -5,22685E | FALSE | Low | IsMasterP | AAEL0222 | 2:1196366 | 114 | 2 | FALSE | 1 | 1 | 1 | 496 | 570 | 884 |
| -9,22284E | FALSE | Low | IsMasterP | AAEL0254 | 3:1200538 | 115 | 1 | FALSE | 1 | 1 | 1 | 1504 | 1670 | 840 |
| -1,06254E | FALSE | Low | IsMasterP | AAEL0226 | 3:3277410 | 116 | 4 | FALSE | 1 | 1 | 1 | 283 | 318 | 573 |
| -6,68612E | FALSE | Low | IsMasterP | AAEL0028 | 1:3046835 | 116 | 4 | FALSE | 1 | 1 | 1 | 300 | 336 | 728 |
| -6,34731E | FALSE | Low | IsMasterP | AAEL0056 | 2:1321208 | 116 | 4 | FALSE | 1 | 1 | 1 | 446 | 505 | 768 |
| 4,50919E- | FALSE | Low | IsMasterP | AAEL0152 | 1:4296707 | 116 | 2 | FALSE | 1 | 1 | 1 | 614 | 676 | 961 |
| 5,29059E- | FALSE | Low | IsMasterP | AAEL0144 | 3:1494436 | 116 | 6 | FALSE | 1 | 1 | 1 | 412 | 473 | 583 |
| -8,23024E | FALSE | Low | IsMasterP | AAEL0173 | 3:2190385 | 116 | 5 | FALSE | 1 | 1 | 1 | 268 | 306 | 1078 |
| -7,45022E | FALSE | Low | IsMasterP | AAEL0048 | 1:2870462 | 116 | 1 | FALSE | 1 | 1 | 1 | 1700 | 1864 | 917 |
| -8,49953E | FALSE | Low | IsMasterP | AAEL0126 | 3:2411114 | 116 | 1 | FALSE | 1 | 1 | 1 | 771 | 868 | 677 |
| 6,75309E- | FALSE | Low | IsMasterP | AAEL0115 | 3:1922369 | 116 | 3 | FALSE | 1 | 1 | 1 | 708 | 799 | 790 |
| 4,56214E- | FALSE | Low | IsMasterP | AAEL0100 | 2:1103820 | 116 | 2 | FALSE | 1 | 1 | 1 | 458 | 529 | 695 |
| -6,24857E | FALSE | Low | IsMasterP | AAEL0051 | 2:7571713 | 117 | 0 | FALSE | 1 | 1 | 1 | 2193 | 2428 | 470 |
| -9,20041E | FALSE | Low | IsMasterP | AAEL0133 | 3:4413463 | 117 | 1 | FALSE | 1 | 1 | 1 | 1189 | 1290 | 717 |
| 2,92108E- | FALSE | Low | IsMasterP | AAEL0010 | 3:5584138 | 117 | 2 | FALSE | 1 | 1 | 1 | 1337 | 1469 | 465 |
| 3,17089E- | FALSE | Low | IsMasterP | AAEL0069 | 2:2421077 | 117 | 5 | FALSE | 1 | 1 | 1 | 372 | 431 | 698 |
| -1,28954E | FALSE | Low | IsMasterP | AAEL0068 | 3:3697972 | 117 | 2 | FALSE | 1 | 1 | 1 | 551 | 604 | 765 |
| 6,32992E- | FALSE | Low | IsMasterP | AAEL0121 | 2:2039876 | 117 | 3 | FALSE | 1 | 1 | 1 | 803 | 869 | 698 |
| -6,16727E | FALSE | Low | IsMasterP | AAEL0215 | 1:9018893 | 117 | 1 | FALSE | 1 | 1 | 1 | 1604 | 1774 | 894 |

| | | | | | | | | | | | | | | |
|---|---|---|---|---|---|---|---|---|---|---|---|---|---|---|
| -7,84834E | FALSE | Low | IsMasterP | AAEL0043 | 3:365759: | 117 | 1 | FALSE | 1 | 1 | 1 | 1373 | 1528 | 601 |
| 5,69468E- | FALSE | Low | IsMasterP | AAEL0022 | 2:430716: | 117 | 11 | FALSE | 1 | 1 | 1 | 269 | 311 | 778 |
| 1,30823E- | FALSE | Low | IsMasterP | AAEL0195 | 1:161613: | 117 | 3 | FALSE | 1 | 1 | 1 | 410 | 462 | 834 |
| -7,88462E | FALSE | Low | IsMasterP | AAEL0199 | 3:807590: | 117 | 8 | FALSE | 1 | 1 | 1 | 224 | 251 | 521 |
| -7,27462E | FALSE | Low | IsMasterP | AAEL0140 | 1:148199: | 117 | 5 | FALSE | 1 | 1 | 1 | 261 | 304 | 816 |
| -2,80764E | FALSE | Low | IsMasterP | AAEL0019 | 2:389720: | 117 | 1 | FALSE | 1 | 1 | 1 | 1370 | 1565 | 734 |
| 3,73592E- | FALSE | Low | IsMasterP | AAEL0070 | 3:299826( | 117 | 14 | FALSE | 1 | 1 | 1 | 125 | 134 | 718 |

| Score Seq | # Peptides | Biological | Cellular Co | Molecular | Pfam IDs | Entrez Ge | Ensembl G | Gene Sym | Chromoso | KEGG Path | WikiPathw | Reactome | # Protein | Abundanc |
|---|---|---|---|---|---|---|---|---|---|---|---|---|---|---|
| 2489662 | 215 | transport | | protein bi | Pf00094, I | 5567518 | AAEL0061 | AaeL_AAE | 3 | | | | 0 | 548 |
| 2003379 | 188 | | | | | | | | | | | | 0 | 541 |
| 1824593 | 185 | | | | | | | | | | | | 0 | 570 |
| 442332 | 152 | | | | | | | | | | | | 0 | 774 |
| 417822 | 48 | | | | | | | | | | | | 0 | 655 |
| 374384 | 153 | | | | | | | | | | | | 0 | 1262 |
| 373479 | 153 | | | | | | | | | | | | 0 | |
| 351273 | 41 | | | | | | | | | | | | 0 | 906 |
| 331526 | 37 | | | | | | | | | | | | 0 | 708 |
| 330275 | 143 | | | | | | | | | | | | 0 | 925 |
| 313667 | 114 | | | | | | | | | | | | 0 | 1001 |
| 305149 | 18 | | | | | | | | | | | | 0 | 931 |
| 302882 | 105 | | | | | | | | | | | | 0 | |
| 285236 | 154 | | | | | | | | | | | | 0 | 741 |
| 258918 | 53 | | | | | | | | | | | | 0 | 884 |
| 257587 | 33 | | | | | | | | | | | | 0 | 706 |
| 243571 | 40 | | | | | | | | | | | | 0 | 877 |
| 239810 | 75 | | | RNA bindi | Pf00076, I | 5565266 | AAEL0046 | AaeL_AAE | 1 | | | | 0 | 855 |
| 232660 | 35 | | | | | | | | | | | | 0 | |
| 230604 | 29 | | | | | | | | | | | | 0 | 939 |
| 212521 | 32 | | | | | | | | | | | | 0 | 885 |
| 209906 | 79 | | | | | | | | | | | | 0 | 967 |
| 205428 | 32 | | | | | | | | | | | | 0 | 1213 |
| 203365 | 94 | | | | | | | | | | | | 0 | 698 |
| 203169 | 100 | | | | | | | | | | | | 0 | 866 |
| 199845 | 105 | | | | | | | | | | | | 0 | 742 |
| 199817 | 34 | | | | | | | | | | | | 0 | 742 |
| 198250 | 19 | | | | | | | | | | | | 0 | 180 |
| 193234 | 43 | cell organization and | | catalytic a | Pf00012, I | 23687769 | AAEL0173 | AaeL_AAE | 3 | Protein export; Protein process | | | 2 | 847 |
| 191891 | 55 | | | | | | | | | | | | 0 | 938 |
| 191369 | 58 | | | | | | | | | | | | 0 | 822 |

| | | | |
|---|---|---|---|
| 190353 | 55 | 0 | 988 |
| 187179 | 53 | 0 | 840 |
| 184906 | 40 | 0 | 591 |
| 181752 | 71 | 0 | 917 |
| 170945 | 44 | 0 | 888 |
| 169363 | 49 | 0 | 830 |
| 164292 | 56 | 0 | 685 |
| 162770 | 42 | 0 | 697 |
| 161060 | 42 | 0 | 653 |
| 160619 | 26 | 0 | 824 |
| 160243 | 48 | 0 | 594 |
| 159720 | 19 | 0 | 695 |
| 158691 | 43 | 0 | 880 |
| 158397 | 70 | 0 | 1044 |
| 157905 | 48 | 0 | 686 |
| 156360 | 26 | 0 | 937 |
| 150737 | 38 | 0 | 851 |
| 150038 | 51 | 0 | 456 |
| 148074 | 58 | 0 | 739 |
| 146005 | 23 | 0 | 845 |
| 144178 | 33 | 0 | 608 |
| 141070 | 33 | 0 | 852 |
| 139303 | 9 | 0 | 993 |
| 138227 | 32 | 0 | 1004 |
| 137417 | 32 | 0 | 766 |
| 137137 | 25 | 0 | 823 |
| 136026 | 37 | 0 | 836 |
| 135468 | 46 | 0 | 626 |
| 135022 | 79 | 0 | 851 |
| 132784 | 38 | 0 | 750 |
| 130691 | 36 | 0 | 901 |

| | | | |
|---|---|---|---|
| 130043 | 39 | 0 | 788 |
| 128598 | 63 | 0 | 1023 |
| 128492 | 48 | 0 | 452 |
| 125404 | 49 | 0 | 751 |
| 124454 | 17 | 0 | 890 |
| 124388 | 24 | 0 | 751 |
| 122262 | 43 | 0 | 1420 |
| 120730 | 42 | 0 | 750 |
| 119558 | 52 | 0 | 754 |
| 117939 | 40 | 0 | 814 |
| 117226 | 18 | 0 | 899 |
| 116962 | 19 | 0 | 951 |
| 114095 | 13 | 0 | 1089 |
| 112981 | 29 | 0 | 643 |
| 112627 | 28 | 0 | 856 |
| 111433 | 17 | 0 | 927 |
| 110024 | 86 | 0 | 721 |
| 107779 | 77 | 0 | 858 |
| 107405 | 28 | 0 | 821 |
| 106783 | 26 | 0 | 741 |
| 106258 | 31 | 0 | 783 |
| 105666 | 43 | 0 | 825 |
| 105304 | 13 | 0 | 717 |
| 104428 | 53 | 0 | 537 |
| 104187 | 15 | 0 | 985 |
| 102518 | 32 | 0 | 850 |
| 102111 | 53 | 0 | 613 |
| 101895 | 20 | 0 | 1108 |
| 101616 | 84 | 0 | 781 |
| 100856 | 44 | 0 | 816 |
| 100582 | 40 | 0 | 638 |

| | | | | | | |
|---|---|---|---|---|---|---|
| 100329 | 22 | | | | 0 | 955 |
| 100066 | 27 | | | | 0 | 817 |
| 99876 | 42 | | | | 0 | 766 |
| 99718 | 36 | | | | 0 | 870 |
| 99631 | 7 | | | | 0 | 865 |
| 98641 | 40 | | | | 0 | 745 |
| 98412 | 46 | | | | 0 | 572 |
| 98198 | 29 | | | | 0 | 1009 |
| 97252 | 22 | | | | 0 | |
| 96269 | 29 | | | | 0 | 853 |
| 96146 | 46 | | | | 0 | 936 |
| 95252 | 17 | | | | 0 | 948 |
| 95100 | 32 | | | | 0 | 949 |
| 95019 | 28 | | | | 0 | 668 |
| 94598 | 49 | | | | 0 | |
| 94567 | 25 | | | | 0 | 901 |
| 94010 | 45 | | | | 0 | 783 |
| 93089 | 32 | | | | 0 | 853 |
| 92863 | 23 | | | | 0 | 800 |
| 92436 | 48 | | | | 0 | |
| 92303 | 27 | | | | 0 | 855 |
| 91537 | 30 | | | | 0 | 1215 |
| 90563 | 46 | | | | 0 | |
| 90468 | 27 | | | | 0 | 836 |
| 90334 | 10 | | | | 0 | 755 |
| 90288 | 33 | | | | 0 | 814 |
| 89754 | 38 | | | | 0 | 780 |
| 89347 | 64 | | | | 0 | 699 |
| 88600 | 30 | RNA bindi Pf00076, I 23687536 AAEL0171 AaeL_AAE | | 3 | 0 | 1131 |
| 88108 | 35 | | | | 0 | 967 |
| 88012 | 11 | | | | 0 | 930 |

| | | | |
|---|---|---|---|
| 87878 | 40 | 0 | 760 |
| 87675 | 40 | 0 | 693 |
| 87035 | 31 | 0 | 721 |
| 86894 | 35 | 0 | 864 |
| 86707 | 25 | 0 | 941 |
| 86544 | 24 | 0 | 885 |
| 85976 | 41 | 0 | 779 |
| 85646 | 27 | 0 | 893 |
| 85628 | 17 | 0 | 740 |
| 85405 | 30 | 0 | 923 |
| 84802 | 48 | 0 | 722 |
| 84535 | 31 | 0 | 703 |
| 84291 | 27 | 0 | 808 |
| 84170 | 22 | 0 | 842 |
| 83929 | 42 | 0 | 1071 |
| 83556 | 33 | 0 | 505 |
| 83281 | 30 | 0 | 456 |
| 83105 | 38 | 0 | 759 |
| 82216 | 36 | 0 | 655 |
| 81178 | 24 | 0 | 844 |
| 80956 | 29 | 0 | 755 |
| 80947 | 58 | 0 | 996 |
| 80230 | 20 | 0 | 844 |
| 80055 | 22 | 0 | 962 |
| 79878 | 35 | 0 | 788 |
| 78872 | 24 | 0 | 641 |
| 78056 | 36 | 0 | 663 |
| 77823 | 34 | 0 | 574 |
| 77717 | 25 | 0 | 1073 |
| 77185 | 36 | 0 | 786 |
| 76552 | 27 | 0 | 896 |

| | | | | | | | |
|---|---|---|---|---|---|---|---|
| 75960 | 39 | | | | | 0 | 710 |
| 75771 | 26 | 5572898 | AAEL0100 | AaeL_AAE | 2 | 0 | 869 |
| 75326 | 21 | | | | | 0 | 1125 |
| 74834 | 31 | | | | | 0 | 851 |
| 74193 | 60 | | | | | 0 | 868 |
| 73750 | 21 | | | | | 0 | 653 |
| 73671 | 27 | | | | | 0 | 692 |
| 73575 | 11 | | | | | 0 | 824 |
| 73462 | 29 | | | | | 0 | 876 |
| 72842 | 21 | | | | | 0 | 814 |
| 72386 | 21 | | | | | 0 | 766 |
| 71951 | 28 | | | | | 0 | 809 |
| 71124 | 32 | | | | | 0 | 830 |
| 70984 | 25 | | | | | 0 | 783 |
| 70875 | 22 | | | | | 0 | 970 |
| 70495 | 18 | | | | | 0 | 707 |
| 69960 | 45 | | | | | 0 | 587 |
| 69492 | 20 | | | | | 0 | 793 |
| 69487 | 28 | | | | | 0 | 1032 |
| 69485 | 28 | | | | | 0 | 871 |
| 69483 | 38 | | | | | 0 | 815 |
| 69454 | 21 | | | | | 0 | 1093 |
| 69119 | 26 | | | | | 0 | 850 |
| 68029 | 37 | | | | | 0 | 572 |
| 67133 | 27 | | | | | 0 | 736 |
| 66883 | 32 | | | | | 0 | 953 |
| 66063 | 16 | | | | | 0 | 1009 |
| 66058 | 27 | | | | | 0 | 817 |
| 65869 | 24 | | | | | 0 | |
| 65575 | 11 | | | | | 0 | 725 |
| 65541 | 24 | | | | | 0 | 718 |

| | | | |
|---|---|---|---|
| 65486 | 25 | 0 | 447 |
| 65357 | 13 | 0 | 1067 |
| 65266 | 25 | 0 | 700 |
| 64825 | 29 | 0 | 664 |
| 64706 | 38 | 0 | 1005 |
| 64527 | 11 | 0 | 871 |
| 64449 | 14 | 0 | |
| 64404 | 48 | 0 | 876 |
| 64024 | 21 | 0 | 766 |
| 64023 | 51 | 0 | 716 |
| 63531 | 25 | 0 | 839 |
| 63075 | 16 | 0 | 972 |
| 63066 | 15 | 0 | 858 |
| 62846 | 18 | 0 | 859 |
| 62762 | 27 | 0 | 639 |
| 62738 | 9 | 0 | 878 |
| 62706 | 34 | 0 | 726 |
| 62467 | 23 | 0 | 1021 |
| 62291 | 11 | 0 | 740 |
| 62264 | 29 | 0 | 733 |
| 61730 | 28 | 0 | 778 |
| 61055 | 26 | 0 | 656 |
| 60725 | 22 | 0 | 874 |
| 60558 | 33 | 0 | 639 |
| 60538 | 18 | 0 | 620 |
| 60180 | 21 | 0 | 793 |
| 59998 | 20 | 0 | 704 |
| 59576 | 18 | 0 | 865 |
| 59382 | 30 | 0 | 743 |
| 59107 | 24 | 0 | 1055 |
| 58796 | 18 | 0 | 1022 |

| | | | | | | | | | | | | |
|---|---|---|---|---|---|---|---|---|---|---|---|---|
| 58376 | 31 | | | | | | | | | | 0 | 1037 |
| 58193 | 22 | | | | | | | | | | 0 | 1021 |
| 57974 | 24 | | | | | | | | | | 0 | 850 |
| 57886 | 12 | | | | | | | | | | 0 | 747 |
| 57648 | 18 | | | | | | | | | | 0 | 753 |
| 57497 | 15 | | | | | | | | | | 0 | 873 |
| 57199 | 12 | | | | | | | | | | 0 | 988 |
| 57134 | 16 | | | | | | | | | | 0 | 958 |
| 56995 | 7 | | | | | | | | | | 0 | 831 |
| 56932 | 16 | | | | | | | | | | 0 | 959 |
| 56688 | 20 | | | | | | | | | | 0 | 516 |
| 56611 | 27 | | | | | | | | | | 0 | 1144 |
| 56303 | 12 | | | | | | | | | | 0 | 824 |
| 56270 | 19 | | | | | | | | | | 0 | 878 |
| 55776 | 13 | | | | | | | | | | 0 | 746 |
| 55715 | 25 | | | | | | | | | | 0 | 741 |
| 55193 | 24 | | | | | | | | | | 0 | 981 |
| 55112 | 29 | metabolic | cytoplasm | catalytic a | Pf00458, I | 5566973 | AAEL0057 | AaeL_AAE | 2 | Aminoacyl-tRNA biosynthesis | 1 | 783 |
| 54857 | 23 | | | | | | | | | | 0 | 1112 |
| 54634 | 17 | metabolic | membran | catalytic a | Pf03435, I | 5576348 | AAEL0028 | AaeL_AAE | 1 | | 0 | 489 |
| 54617 | 31 | | | | | | | | | | 0 | 695 |
| 54572 | 20 | | | | | | | | | | 0 | 948 |
| 54501 | 15 | | | | | | | | | | 0 | 822 |
| 54208 | 27 | | | | | | | | | | 0 | 925 |
| 53987 | 22 | | | | | | | | | | 0 | 878 |
| 53974 | 20 | | | | | | | | | | 0 | 782 |
| 53733 | 10 | | | | Pf03066 | 5578927 | AAEL0037 | AaeL_AAE | 1 | | 0 | 985 |
| 53671 | 25 | | | | | | | | | | 0 | 724 |
| 53502 | 14 | | | | | | | | | | 0 | 950 |
| 53428 | 26 | | | | | | | | | | 0 | 1006 |
| 53406 | 12 | | | | | | | | | | 0 | 850 |

| | | | | | | | | | | |
|---|---|---|---|---|---|---|---|---|---|---|
| 53029 | 19 | | | | | | | | 0 | 519 |
| 52987 | 17 | | | | | | | | 0 | 869 |
| 52941 | 11 | | | | | | | | 0 | 1114 |
| 52785 | 28 | | | | | | | | 0 | 840 |
| 52634 | 25 | | | | | | | | 0 | 857 |
| 52564 | 19 | | | | | | | | 0 | 873 |
| 52533 | 15 | | | | | | | | 0 | 870 |
| 52451 | 15 | | | | | | | | 0 | 795 |
| 52145 | 33 | | | | | | | | 0 | 841 |
| 51934 | 10 | | | | | | | | 0 | 1032 |
| 51811 | 15 | | | | | | | | 0 | 925 |
| 51769 | 31 | | | | | | | | 0 | 709 |
| 51636 | 20 | | | | | | | | 0 | 675 |
| 51509 | 24 | | | | | | | | 0 | 872 |
| 51296 | 22 | | | | | | | | 0 | 937 |
| 51258 | 8 | | | | | | | | 0 | 993 |
| 51235 | 12 | | | | | | | | 0 | 742 |
| 51231 | 23 | | | | | | | | 0 | 922 |
| 51147 | 28 | | | | | | | | 0 | 796 |
| 51071 | 33 | | | | | | | | 0 | 597 |
| 50951 | 17 | | | | | | | | 0 | 881 |
| 50941 | 23 | | | | | | | | 0 | |
| 50934 | 23 | | | | | | | | 0 | 963 |
| 50639 | 15 | metabolic process | catalytic a | Pf00106, I | 23687553 | AAEL0171 | AaeL_AAE | 2 | 0 | 794 |
| 50592 | 33 | | | | | | | | 0 | 839 |
| 50529 | 12 | | | | | | | | 0 | 999 |
| 50498 | 21 | | | | | | | | 0 | 942 |
| 50450 | 28 | | | | | | | | 0 | 873 |
| 50449 | 22 | | | | | | | | 0 | 1032 |
| 50378 | 16 | | | | | | | | 0 | 835 |
| 50104 | 17 | | | | | | | | 0 | 940 |

| | | | | | | | |
|---|---|---|---|---|---|---|---|
| 49965 | 36 | | | | | 0 | 596 |
| 49678 | 23 | | | | | 0 | |
| 49591 | 30 | | | | | 0 | 642 |
| 49517 | 22 | | | | | 0 | 777 |
| 49350 | 23 | | | | | 0 | |
| 49311 | 26 | | | | | 0 | 733 |
| 49260 | 14 | | | | | 0 | |
| 49247 | 26 | 5573231 | AAEL0103 | AaeL_AAE | 3 | 0 | 945 |
| 49181 | 31 | | | | | 0 | 651 |
| 48744 | 17 | | | | | 0 | 750 |
| 48646 | 10 | | | | | 0 | 905 |
| 48261 | 20 | | | | | 0 | 841 |
| 48202 | 18 | | | | | 0 | 914 |
| 48145 | 17 | | | | | 0 | |
| 48023 | 33 | | | | | 0 | 802 |
| 47457 | 13 | | | | | 0 | |
| 47400 | 18 | | | | | 0 | 806 |
| 47331 | 19 | | | | | 0 | 706 |
| 47307 | 19 | | | | | 0 | 854 |
| 47287 | 33 | | | | | 0 | 581 |
| 47086 | 27 | | | | | 0 | 825 |
| 46941 | 13 | | | | | 0 | 837 |
| 46659 | 21 | | | | | 0 | 771 |
| 46546 | 9 | 5563800 | AAEL0052 | AaeL_AAE | 2 | 0 | 919 |
| 46472 | 11 | | | | | 0 | 1057 |
| 46240 | 12 | | | | | 0 | 942 |
| 46136 | 21 | | | | | 0 | 498 |
| 45918 | 21 | | | | | 0 | 826 |
| 45826 | 21 | | | | | 0 | 788 |
| 45689 | 22 | | | | | 0 | 593 |
| 45194 | 15 | | | | | 0 | 773 |

| | | | |
|---|---|---|---|
| 45169 | 21 | 0 | 771 |
| 45131 | 28 | 0 | 816 |
| 44963 | 19 | 0 | 790 |
| 44935 | 28 | 0 | 825 |
| 44800 | 14 | 0 | 770 |
| 44641 | 25 | 0 | 606 |
| 44552 | 16 | 0 | 1139 |
| 44517 | 18 | 0 | 834 |
| 44353 | 14 | 0 | 801 |
| 44341 | 18 | 0 | 644 |
| 44216 | 15 | 0 | 876 |
| 44164 | 10 | 0 | 892 |
| 44146 | 20 | 0 | 762 |
| 43843 | 18 | 0 | 826 |
| 43673 | 24 | 0 | 669 |
| 43619 | 23 | 0 | 927 |
| 43478 | 20 | 0 | 643 |
| 43129 | 14 | 0 | 821 |
| 43077 | 24 | 0 | 799 |
| 43009 | 23 | 0 | 459 |
| 42668 | 17 | 0 | 601 |
| 42554 | 37 | 0 | 576 |
| 42515 | 21 | 0 | |
| 42484 | 18 | 0 | 718 |
| 42275 | 15 | 0 | 631 |
| 42213 | 21 | 0 | 933 |
| 42188 | 6 | 0 | 830 |
| 42121 | 15 | 0 | 619 |
| 41999 | 11 | 0 | 894 |
| 41990 | 17 | 0 | 650 |
| 41832 | 15 | 0 | 815 |

| | | | | | | | | |
|---|---|---|---|---|---|---|---|---|
| 41788 | 22 | | | | | | 0 | 887 |
| 41540 | 24 | | | | | | 0 | 585 |
| 41470 | 15 | | | | | | 0 | 786 |
| 41449 | 12 | | | | | | 0 | 949 |
| 41370 | 14 | | | | | | 0 | 682 |
| 41351 | 21 | | | | | | 0 | 559 |
| 41313 | 13 | | | | | | 0 | 920 |
| 41286 | 11 | | | | | | 0 | 831 |
| 41211 | 18 | | | | | | 0 | 698 |
| 41210 | 16 | | | | | | 0 | 923 |
| 41066 | 25 | | | | | | 0 | 619 |
| 41054 | 10 | | | | | | 0 | 978 |
| 40711 | 14 | | | | | | 0 | 596 |
| 40704 | 20 | | | | | | 0 | 711 |
| 40639 | 41 | | | | | | 0 | 854 |
| 40601 | 18 | | | | | | 0 | 698 |
| 40574 | 19 | | | | | | 0 | 667 |
| 39997 | 18 | | | | | | 0 | 870 |
| 39771 | 17 | | 5566701 | AAEL0055 | AaeL_AAE | 3 | 0 | 944 |
| 39731 | 16 | | | | | | 0 | 609 |
| 39714 | 25 | | | | | | 0 | 774 |
| 39479 | 28 | | | | | | 0 | 679 |
| 39440 | 9 | | | | | | 0 | 802 |
| 39419 | 10 | | | | | | 0 | 1059 |
| 39307 | 17 | | | | | | 0 | 1010 |
| 39298 | 19 | | | | | | 0 | 758 |
| 39196 | 25 | | | | | | 0 | 713 |
| 39095 | 10 | Pf03227 | 5565612 | AAEL0048 | AaeL_AAE | 1 | 0 | 702 |
| 39016 | 17 | | | | | | 0 | 460 |
| 39011 | 17 | | | | | | 0 | 506 |
| 38986 | 24 | | | | | | 0 | 655 |

| | | | |
|---|---|---|---|
| 38908 | 19 | 0 | 953 |
| 38907 | 11 | 0 | 872 |
| 38650 | 25 | 0 | 877 |
| 38624 | 22 | 0 | 904 |
| 38589 | 15 | 0 | 797 |
| 38576 | 17 | 0 | 543 |
| 38381 | 17 | 0 | 766 |
| 38289 | 28 | 0 | 839 |
| 38050 | 8 | 0 | 677 |
| 38041 | 22 | 0 | 1098 |
| 37983 | 28 | 0 | 692 |
| 37868 | 13 | 0 | 708 |
| 37734 | 13 | 0 | 751 |
| 37635 | 24 | 0 | 637 |
| 37630 | 21 | 0 | 725 |
| 37460 | 15 | 0 | 965 |
| 37458 | 22 | 0 | 760 |
| 37437 | 26 | 0 | 849 |
| 37295 | 14 | 0 | 824 |
| 37197 | 8 | 0 | 711 |
| 37135 | 20 | 0 | 546 |
| 36757 | 37 | 0 | 797 |
| 36677 | 18 | 0 | 879 |
| 36624 | 13 | 0 | |
| 36610 | 27 | 0 | 531 |
| 36534 | 14 | 0 | 870 |
| 36519 | 13 | 0 | 889 |
| 36485 | 20 | 0 | 982 |
| 36483 | 26 | 0 | 756 |
| 36454 | 20 | 0 | 765 |
| 36444 | 6 | 0 | 834 |

| | | | | | | | | | |
|---|---|---|---|---|---|---|---|---|---|
| 36383 | 21 | | | | | | | 0 | 746 |
| 36354 | 28 | | | | | | | 0 | 821 |
| 36232 | 29 | | | | | | | 0 | 597 |
| 36220 | 17 | | | | | | | 0 | 833 |
| 36157 | 19 | | | | | | | 0 | 817 |
| 36109 | 15 | | | | | | | 0 | 868 |
| 35995 | 9 | | | | | | | 0 | 862 |
| 35890 | 14 | | | | | | | 0 | 704 |
| 35862 | 22 | | | | | | | 0 | 933 |
| 35850 | 18 | | | | | | | 0 | 816 |
| 35731 | 20 | | | | | | | 0 | 917 |
| 35447 | 19 | | | | | | | 0 | 849 |
| 35423 | 9 | | | | | | | 0 | 850 |
| 35353 | 14 | | | | | | | 0 | 671 |
| 35321 | 27 | | | | | | | 0 | 754 |
| 35108 | 19 | | | | | | | 0 | 825 |
| 34951 | 24 | | | | | | | 0 | 462 |
| 34932 | 12 | protein bi | Pf00515, I | 5575512 | AAEL0118 | AaeL_AAE | 2 | 0 | 975 |
| 34904 | 13 | | | | | | | 0 | 637 |
| 34889 | 16 | | | | | | | 0 | 725 |
| 34651 | 12 | | | | | | | 0 | 936 |
| 34451 | 10 | | | | | | | 0 | 1083 |
| 34389 | 18 | | | | | | | 0 | 644 |
| 34372 | 21 | | | | | | | 0 | 897 |
| 34182 | 22 | | | | | | | 0 | 812 |
| 34093 | 26 | | | | | | | 0 | 735 |
| 34074 | 12 | | Pf00226 | 5578812 | AAEL0138 | AaeL_AAE | 2 | 0 | 699 |
| 33918 | 23 | | | | | | | 0 | 463 |
| 33845 | 16 | | | | | | | 0 | 961 |
| 33643 | 16 | | | | | | | 0 | 840 |
| 33641 | 20 | | | | | | | 0 | 1235 |

| | | | | | | | | | |
|---|---|---|---|---|---|---|---|---|---|
| 33630 | 18 | | | | | | | 0 | 1100 |
| 33582 | 10 | | | | | | | 0 | 962 |
| 33559 | 14 | | | | | | | 0 | 1344 |
| 33308 | 22 | | | | | | | 0 | 630 |
| 33015 | 21 | | | 5574939 | AAEL0115 | AaeL_AAE | 3 | 0 | 839 |
| 32886 | 11 | | | | | | | 0 | 1119 |
| 32874 | 15 | | | | | | | 0 | 802 |
| 32857 | 18 | | | | | | | 0 | 886 |
| 32821 | 14 | | | | | | | 0 | |
| 32768 | 16 | | | | | | | 0 | 807 |
| 32736 | 9 | | | 5566109 | AAEL0051 | AaeL_AAE | 2 | 0 | 860 |
| 32731 | 27 | | | | | | | 0 | 631 |
| 32711 | 12 | | | | | | | 0 | 788 |
| 32686 | 14 | | | | | | | 0 | 783 |
| 32582 | 11 | | | | | | | 0 | 713 |
| 32522 | 17 | | | | | | | 0 | 704 |
| 32509 | 14 | | | | | | | 0 | 621 |
| 32495 | 20 | | | | | | | 0 | 548 |
| 32440 | 10 | RNA bindi | Pf00013, I | 23687901 | AAEL0174 | AaeL_AAE | 1 | 0 | 1016 |
| 32425 | 15 | | | | | | | 0 | 834 |
| 32364 | 17 | | | | | | | 0 | 868 |
| 32333 | 23 | | | | | | | 0 | 828 |
| 32191 | 11 | | | | | | | 0 | 830 |
| 32142 | 22 | | | | | | | 0 | 750 |
| 32139 | 15 | | | | | | | 0 | 713 |
| 32084 | 14 | | | | | | | 0 | 711 |
| 32021 | 20 | | | | | | | 0 | 798 |
| 32019 | 21 | | | | | | | 0 | 740 |
| 32018 | 15 | | | | | | | 0 | 850 |
| 31947 | 20 | | | | | | | 0 | 773 |
| 31945 | 12 | | | | | | | 0 | 964 |

| | | | | | | | | | | |
|---|---|---|---|---|---|---|---|---|---|---|
| 31880 | 16 | | | | | | | | 0 | 795 |
| 31834 | 7 | | | | | | | | 0 | 956 |
| 31749 | 16 | | | | | | | | 0 | 986 |
| 31660 | 19 | | | | | | | | 0 | 906 |
| 31649 | 11 | | | | | | | | 0 | 1032 |
| 31633 | 11 | | | | | | | | 0 | 938 |
| 31531 | 24 | | | | | | | | 0 | 624 |
| 31426 | 18 | | | | | | | | 0 | 1116 |
| 31299 | 17 | | | | | | | | 0 | 725 |
| 31257 | 21 | | | | | | | | 0 | 906 |
| 31254 | 18 | | | | | | | | 0 | 904 |
| 31155 | 32 | | | | | | | | 0 | 778 |
| 31149 | 15 | | | | | | | | 0 | 869 |
| 31087 | 12 | | | | | | | | 0 | 653 |
| 30975 | 16 | | | | | | | | 0 | 887 |
| 30907 | 15 | | | | | | | | 0 | 800 |
| 30817 | 21 | | | | | | | | 0 | 742 |
| 30775 | 15 | | | | | | | | 0 | 883 |
| 30734 | 13 | | | | | | | | 0 | 734 |
| 30678 | 9 | | | | | | | | 0 | 772 |
| 30624 | 20 | | | | | | | | 0 | 540 |
| 30564 | 15 | | | | | | | | 0 | 511 |
| 30461 | 11 | | Pf07165 | 5570664 | AAEL0084 | AaeL_AAE | 2 | | 0 | 822 |
| 30411 | 18 | | | | | | | | 0 | 750 |
| 30342 | 6 | | | | | | | | 0 | 1126 |
| 30313 | 18 | | | | | | | | 0 | 604 |
| 30276 | 19 | catalytic a | Pf00009, I | 23687788 | AAEL0173 | AaeL_AAE | 2 | mRNA surveillance pathway | 1 | 669 |
| 30235 | 17 | | | | | | | | 0 | 635 |
| 30167 | 20 | | | | | | | | 0 | 915 |
| 30128 | 27 | | | | | | | | 0 | 983 |
| 30078 | 21 | | | | | | | | 0 | 610 |

| | | | |
|---|---|---|---|
| 29915 | 22 | 0 | 762 |
| 29832 | 18 | 0 | 635 |
| 29781 | 13 | 0 | 847 |
| 29771 | 11 | 0 | 956 |
| 29582 | 8 | 0 | 1002 |
| 29483 | 23 | 0 | 717 |
| 29461 | 19 | 0 | 837 |
| 29434 | 14 | 0 | 906 |
| 29402 | 20 | 0 | 961 |
| 29387 | 18 | 0 | 775 |
| 29362 | 13 | 0 | 891 |
| 29257 | 8 | 0 | 917 |
| 29217 | 20 | 0 | 667 |
| 29195 | 18 | 0 | 818 |
| 29179 | 20 | 0 | 854 |
| 29170 | 14 | 0 | 711 |
| 29167 | 23 | 0 | 463 |
| 29056 | 16 | 0 | 888 |
| 29022 | 21 | 0 | 680 |
| 29022 | 12 | 0 | 871 |
| 29016 | 23 | 0 | 1053 |
| 28783 | 19 | 0 | 885 |
| 28763 | 18 | 0 | 615 |
| 28720 | 20 | 0 | 678 |
| 28716 | 21 | 0 | 511 |
| 28565 | 15 | 0 | 937 |
| 28405 | 15 | 0 | 782 |
| 28383 | 23 | 0 | 729 |
| 28358 | 27 | 0 | 859 |
| 28318 | 10 | 0 | 916 |
| 28309 | 10 | 0 | 702 |

| | | | | | | | | | |
|---|---|---|---|---|---|---|---|---|---|
| 28295 | 16 | | | | | | | 0 | 1014 |
| 28189 | 20 | | | | | | | 0 | 889 |
| 28170 | 19 | | | | | | | 0 | 607 |
| 28092 | 18 | | | | | | | 0 | 831 |
| 28066 | 14 | | | | | | | 0 | 909 |
| 28022 | 14 | | | | | | | 0 | 956 |
| 28014 | 16 | | | | | | | 0 | 756 |
| 27996 | 19 | | | | | | | 0 | 804 |
| 27982 | 20 | | | | | | | 0 | 635 |
| 27979 | 10 | | | | | | | 0 | 1114 |
| 27954 | 18 | | | | | | | 0 | 691 |
| 27943 | 12 | | | | | | | 0 | 617 |
| 27908 | 14 | | | | | | | 0 | 693 |
| 27897 | 17 | | | | | | | 0 | 737 |
| 27792 | 14 | Pf01798, I | 23687841 | AAEL0174 | AaeL_AAE | 1 | Ribosome biogenesis in eukary | 1 | 732 |
| 27634 | 14 | | | | | | | 0 | 626 |
| 27595 | 22 | | | | | | | 0 | 852 |
| 27578 | 17 | | | | | | | 0 | 766 |
| 27513 | 14 | | | | | | | 0 | 806 |
| 27354 | 7 | | | | | | | 0 | 826 |
| 27344 | 20 | | | | | | | 0 | 552 |
| 27275 | 16 | | | | | | | 0 | 717 |
| 27257 | 7 | | | | | | | 0 | 905 |
| 27190 | 8 | | | | | | | 0 | 1000 |
| 27120 | 15 | | | | | | | 0 | 834 |
| 27073 | 11 | | | | | | | 0 | 741 |
| 27037 | 11 | | | | | | | 0 | 739 |
| 27009 | 12 | | | | | | | 0 | 906 |
| 27005 | 14 | | | | | | | 0 | 843 |
| 26966 | 21 | | | | | | | 0 | 577 |
| 26895 | 16 | | | | | | | 0 | 531 |

| | | | |
|---|---|---|---|
| 26837 | 12 | 0 | 797 |
| 26803 | 19 | 0 | 724 |
| 26763 | 11 | 0 | 867 |
| 26760 | 14 | 0 | 1502 |
| 26707 | 6 | 0 | 873 |
| 26678 | 17 | 0 | 926 |
| 26585 | 16 | 0 | 988 |
| 26501 | 9 | 0 | 993 |
| 26495 | 10 | 0 | 718 |
| 26428 | 14 | 0 | 1107 |
| 26403 | 12 | 0 | 1062 |
| 26265 | 12 | 0 | 812 |
| 26228 | 10 | 0 | 527 |
| 26222 | 15 | 0 | 924 |
| 26082 | 15 | 0 | 520 |
| 26017 | 17 | 0 | 1087 |
| 25937 | 16 | 0 | 1166 |
| 25923 | 10 | 0 | 640 |
| 25917 | 14 | 0 | 670 |
| 25890 | 18 | 0 | 430 |
| 25878 | 13 | 0 | 875 |
| 25852 | 17 | 0 | 725 |
| 25779 | 20 | 0 | 580 |
| 25698 | 21 | 0 | 914 |
| 25631 | 12 | 0 | 729 |
| 25584 | 19 | 0 | 483 |
| 25569 | 10 | 0 | 920 |
| 25498 | 10 | 0 | 604 |
| 25480 | 18 | 0 | 609 |
| 25479 | 13 | 0 | 643 |
| 25451 | 11 | 0 | 968 |

| | | | | | | | |
|---|---|---|---|---|---|---|---|
| 25383 | 10 | | | | | 0 | 829 |
| 25376 | 13 | | | | | 0 | 602 |
| 25307 | 20 | | | | | 0 | 1042 |
| 25283 | 17 | | | | | 0 | 808 |
| 25272 | 16 | | | | | 0 | 598 |
| 25153 | 12 | | | | | 0 | 580 |
| 25124 | 8 | | | | | 0 | 961 |
| 25122 | 14 | | | | | 0 | 688 |
| 25081 | 11 | | | | | 0 | 987 |
| 25076 | 13 | | | | | 0 | 1302 |
| 25013 | 13 | | | | | 0 | 1113 |
| 24999 | 19 | | | | | 0 | 883 |
| 24992 | 17 | 5567452 | AAEL0009 | AaeL_AAE | 3 | 0 | 762 |
| 24938 | 14 | | | | | 0 | 917 |
| 24914 | 17 | | | | | 0 | 534 |
| 24877 | 11 | | | | | 0 | 877 |
| 24866 | 14 | | | | | 0 | 861 |
| 24830 | 21 | | | | | 0 | 714 |
| 24793 | 13 | | | | | 0 | 874 |
| 24770 | 13 | | | | | 0 | 770 |
| 24736 | 7 | | | | | 0 | 818 |
| 24692 | 14 | | | | | 0 | 1025 |
| 24600 | 6 | | | | | 0 | 1080 |
| 24556 | 14 | | | | | 0 | 994 |
| 24489 | 11 | | | | | 0 | 947 |
| 24405 | 8 | | | | | 0 | 902 |
| 24285 | 11 | | | | | 0 | 667 |
| 24274 | 10 | | | | | 0 | 540 |
| 24253 | 14 | | | | | 0 | 389 |
| 24233 | 13 | | | | | 0 | 738 |
| 24229 | 12 | | | | | 0 | 928 |

| | | | | | | | | | | | | |
|---|---|---|---|---|---|---|---|---|---|---|---|---|
| 24177 | 16 | | | | | | | | | | 0 | 744 |
| 24146 | 11 | metabolic | ribosome | structural | Pf00276, I | 23687936 | AAEL0175 | AaeL_AAE | 2 | Ribosome | 1 | 700 |
| 24099 | 8 | | | | | | | | | | 0 | 1029 |
| 24075 | 18 | | | | | | | | | | 0 | 790 |
| 24032 | 15 | | | | | | | | | | 0 | 891 |
| 24031 | 15 | | | | | | | | | | 0 | 665 |
| 24028 | 12 | | | | | | | | | | 0 | 969 |
| 24009 | 17 | | | | | | | | | | 0 | 520 |
| 23759 | 8 | | | | | | | | | | 0 | 918 |
| 23754 | 7 | | | | | | | | | | 0 | 901 |
| 23718 | 15 | | | | | | | | | | 0 | 840 |
| 23706 | 15 | | | | | | | | | | 0 | 795 |
| 23704 | 15 | | | | | | | | | | 0 | 930 |
| 23694 | 13 | | | | | | | | | | 0 | 664 |
| 23671 | 13 | | | | | | | | | | 0 | 635 |
| 23586 | 17 | | | | | | | | | | 0 | 377 |
| 23576 | 10 | | | | | | | | | | 0 | 777 |
| 23575 | 14 | | | | | | | | | | 0 | 764 |
| 23559 | 9 | | | | | | | | | | 0 | |
| 23548 | 16 | | | | | | | | | | 0 | 941 |
| 23518 | 17 | | | | | | | | | | 0 | 593 |
| 23499 | 13 | | | | | | | | | | 0 | 879 |
| 23310 | 18 | | | | | | | | | | 0 | 750 |
| 23177 | 6 | | | | | | | | | | 0 | 1093 |
| 23098 | 15 | | | | | | | | | | 0 | 1103 |
| 23090 | 11 | | | | | | | | | | 0 | 1016 |
| 23081 | 13 | | | | | | | | | | 0 | 700 |
| 22971 | 19 | | | | | | | | | | 0 | 766 |
| 22932 | 10 | | | | | | | | | | 0 | 552 |
| 22915 | 18 | | | | | | | | | | 0 | 604 |
| 22813 | 12 | | | | | | | | | | 0 | 1094 |

| | | | | | | | | | | | | |
|---|---|---|---|---|---|---|---|---|---|---|---|---|
| 22809 | 8 | | | | | | | | | | 0 | 1081 |
| 22733 | 12 | | | | | | | | | | 0 | 688 |
| 22733 | 11 | | | | | | | | | | 0 | 452 |
| 22696 | 19 | | | | | | | | | | 0 | 621 |
| 22681 | 16 | | | | | | | | | | 0 | 668 |
| 22677 | 9 | | | | | | | | | | 0 | 972 |
| 22584 | 7 | | | | | | | | | | 0 | 907 |
| 22561 | 16 | | | | | | | | | | 0 | 903 |
| 22474 | 14 | | | | | | | | | | 0 | 725 |
| 22458 | 12 | | | DNA bindi | Pf01984 | 5568113 | AAEL0065 | AaeL_AAE | 2 | | 0 | 1031 |
| 22449 | 9 | | | | | | | | | | 0 | 790 |
| 22377 | 17 | | | | | | | | | | 0 | 757 |
| 22367 | 15 | | | | | | | | | | 0 | 1155 |
| 22364 | 13 | | | | | | | | | | 0 | 777 |
| 22361 | 7 | | | | | | | | | | 0 | 913 |
| 22328 | 13 | metabolic process | | catalytic a | Pf00443, I | 5573452 | AAEL0104 | AaeL_AAE | 2 | | 0 | 659 |
| 22302 | 12 | | | | | | | | | | 0 | 872 |
| 22293 | 19 | | | | | | | | | | 0 | 1003 |
| 22236 | 10 | | | | Pf11901 | 5574458 | AAEL0111 | AaeL_AAE | 2 | | 0 | 425 |
| 22206 | 12 | | | | | | | | | | 0 | 721 |
| 22200 | 15 | | | | | | | | | | 0 | 684 |
| 22184 | 8 | metabolic | cytoplasm | RNA bindi | Pf06003 | 5570968 | AAEL0087 | AaeL_AAE | 3 | RNA transport | 1 | 1078 |
| 22063 | 12 | | | | | | | | | | 0 | 795 |
| 22053 | 21 | | | | | | | | | | 0 | 775 |
| 22036 | 8 | | | | Pf00581 | 5567139 | AAEL0058 | AaeL_AAE | 2 | | 0 | 963 |
| 22011 | 7 | | | | | | | | | | 0 | 777 |
| 21964 | 10 | | | | | | | | | | 0 | 851 |
| 21958 | 7 | | | | | | | | | | 0 | 654 |
| 21938 | 21 | | | | | | | | | | 0 | 809 |
| 21893 | 8 | | | | | | | | | | 0 | 798 |
| 21833 | 12 | | | | | | | | | | 0 | 635 |

| | | | | | | | | | | |
|---|---|---|---|---|---|---|---|---|---|---|
| 21832 | 15 | | | | | | | | 0 | 668 |
| 21818 | 6 | | | | | | | | 0 | 627 |
| 21804 | 7 | | | | | | | | 0 | 914 |
| 21730 | 9 | | | | | | | | 0 | 851 |
| 21689 | 8 | | | | | | | | 0 | 849 |
| 21636 | 5 | | | | | | | | 0 | 952 |
| 21570 | 5 | | catalytic a | Pf11901 | 5566220 | AAEL0052 | AaeL_AAE | 2 | 0 | 500 |
| 21550 | 9 | | | | | | | | 0 | 716 |
| 21523 | 14 | | | | | | | | 0 | 771 |
| 21477 | 13 | metabolic process | catalytic a | Pf01189, I | 5579023 | AAEL0139 | AaeL_AAE | 3 | 0 | 916 |
| 21450 | 8 | | | | | | | | 0 | 854 |
| 21446 | 20 | | | | | | | | 0 | 1017 |
| 21297 | 12 | | | | | | | | 0 | 990 |
| 21292 | 11 | | | | | | | | 0 | 753 |
| 21267 | 7 | | | | | | | | 0 | 1076 |
| 21165 | 14 | | | | | | | | 0 | 641 |
| 21107 | 14 | | | | | | | | 0 | 625 |
| 21059 | 14 | | | | | | | | 0 | 702 |
| 21020 | 14 | | | | | | | | 0 | 670 |
| 21012 | 9 | | | | | | | | 0 | 382 |
| 21003 | 9 | | | | | | | | 0 | 860 |
| 20989 | 10 | | | | | | | | 0 | 880 |
| 20854 | 3 | | | | | | | | 0 | 1090 |
| 20824 | 7 | | | | | | | | 0 | 764 |
| 20815 | 15 | | | | | | | | 0 | 482 |
| 20739 | 18 | | protein binding | | 5577320 | AAEL0004 | AaeL_AAE | 3 | 0 | 932 |
| 20722 | 9 | | | | | | | | 0 | 724 |
| 20706 | 14 | | | | | | | | 0 | 722 |
| 20638 | 11 | | | | | | | | 0 | 1441 |
| 20577 | 21 | | | | | | | | 0 | 741 |
| 20541 | 13 | | | | | | | | 0 | 486 |

| | | | | | | | | | | | | |
|---|---|---|---|---|---|---|---|---|---|---|---|---|
| 20524 | 13 | | | | | | | | | | 0 | 900 |
| 20494 | 11 | | | | | | | | | | 0 | 1505 |
| 20425 | 14 | | | | | | | | | | 0 | 629 |
| 20327 | 8 | regulation of biologi | | enzyme re | Pf05375 | 5563689 | AAEL0005 | AaeL_AAE | 3 | | 0 | 1044 |
| 20322 | 10 | | | | | | | | | | 0 | 278 |
| 20315 | 18 | | | | | | | | | | 0 | 962 |
| 20312 | 7 | | | | | | | | | | 0 | 911 |
| 20310 | 8 | | | | | | | | | | 0 | 921 |
| 20301 | 12 | | | | | | | | | | 0 | 700 |
| 20272 | 10 | | | | | | | | | | 0 | 879 |
| 20245 | 9 | | | | | | | | | | 0 | 713 |
| 20197 | 12 | | | | | | | | | | 0 | 668 |
| 20186 | 16 | | | | | | | | | | 0 | 976 |
| 20152 | 12 | | | | | | | | | | 0 | 935 |
| 20135 | 7 | | | | | | | | | | 0 | 836 |
| 20088 | 9 | | | | | | | | | | 0 | 950 |
| 20075 | 21 | metabolic process | | catalytic a | Pf01262, I | 5565118 | AAEL0147 | AaeL_AAE | 2 | Lysine degradation; Metabolic | 2 | 431 |
| 20042 | 14 | | | | | | | | | | 0 | 696 |
| 20037 | 13 | metabolic | nucleus | catalytic a | Pf00385, I | 5567298 | AAEL0070 | AaeL_AAE | 3 | | 0 | 1103 |
| 19977 | 14 | | | | | | | | | | 0 | 627 |
| 19886 | 13 | | | | | | | | | | 0 | 648 |
| 19864 | 18 | | | | | | | | | | 0 | 552 |
| 19863 | 4 | | membrane | | | 5571609 | AAEL0092 | AaeL_AAE | 3 | | 0 | 796 |
| 19853 | 12 | | | | | | | | | | 0 | 810 |
| 19852 | 6 | | | | | | | | | | 0 | 789 |
| 19812 | 9 | cellular homeostasis;regulation | | | Pf00085, I | 5570396 | AAEL0082 | AaeL_AAE | 2 | | 0 | 1054 |
| 19767 | 12 | | | | | | | | | | 0 | 285 |
| 19753 | 16 | | | | | | | | | | 0 | 569 |
| 19719 | 18 | | | | | | | | | | 0 | 605 |
| 19699 | 20 | | | | | | | | | | 0 | 720 |
| 19664 | 9 | | | protein bi | Pf00536 | 5570231 | AAEL0081 | AaeL_AAE | 3 | | 0 | 887 |

| | | | | | | | | | |
|---|---|---|---|---|---|---|---|---|---|
| 19660 | 6 | | | | | | | 0 | 911 |
| 19584 | 12 | | | | | | | 0 | 675 |
| 19524 | 12 | | | | | | | 0 | 804 |
| 19516 | 9 | | | | | | | 0 | 687 |
| 19504 | 15 | | | | | | | 0 | 481 |
| 19466 | 14 | | | | | | | 0 | 785 |
| 19380 | 11 | | | | | | | 0 | 815 |
| 19356 | 16 | | | | | | | 0 | 681 |
| 19242 | 13 | | Pf03690 | 5569097 | AAEL0073 | AaeL_AAE | 3 | 0 | 830 |
| 19130 | 14 | | | | | | | 0 | 817 |
| 19116 | 16 | | | | | | | 0 | 808 |
| 19080 | 8 | | | | | | | 0 | 736 |
| 19044 | 10 | | | | | | | 0 | 912 |
| 19024 | 10 | | | | | | | 0 | 659 |
| 18975 | 6 | | | | | | | 0 | 802 |
| 18968 | 17 | | | | | | | 0 | 809 |
| 18910 | 13 | | | | | | | 0 | 675 |
| 18888 | 9 | | | | | | | 0 | 928 |
| 18872 | 12 | | | | | | | 0 | 861 |
| 18839 | 15 | | | | | | | 0 | 986 |
| 18810 | 12 | metal ion | Pf12171, I | 5571937 | AAEL0094 | AaeL_AAE | 2 | 0 | 937 |
| 18759 | 11 | | | | | | | 0 | 886 |
| 18749 | 18 | | | | | | | 0 | 804 |
| 18746 | 15 | | | | | | | 0 | 834 |
| 18735 | 22 | | | | | | | 0 | 767 |
| 18697 | 10 | | | | | | | 0 | 728 |
| 18650 | 12 | | | | | | | 0 | 838 |
| 18602 | 14 | | | | | | | 0 | 1075 |
| 18567 | 11 | | | | | | | 0 | 877 |
| 18548 | 13 | | | | | | | 0 | 887 |
| 18524 | 9 | | | | | | | 0 | 643 |

| | | | | | | | | | | | |
|---|---|---|---|---|---|---|---|---|---|---|---|
| 18435 | 5 | | | | | | | | | 0 | 803 |
| 18397 | 10 | | | | | | | | | 0 | 605 |
| 18351 | 9 | | | | | | | | | 0 | 865 |
| 18314 | 10 | | | | | | | | | 0 | 705 |
| 18233 | 7 | | | | | | | | | 0 | 790 |
| 18206 | 6 | | | | | | | | | 0 | 825 |
| 18185 | 10 | | | | | | | | | 0 | 1311 |
| 18136 | 17 | | | | | | | | | 0 | 613 |
| 18096 | 11 | | | | | | | | | 0 | 1007 |
| 18090 | 11 | | | | | | | | | 0 | 950 |
| 18082 | 12 | | | | | | | | | 0 | 615 |
| 17963 | 13 | | | | | | | | | 0 | 531 |
| 17947 | 2 | | | | | | | | | 0 | 1231 |
| 17936 | 10 | | | | | | | | | 0 | 693 |
| 17932 | 7 | | | | | | | | | 0 | 805 |
| 17930 | 13 | | | | 5568303 | AAEL0067 | AaeL_AAE | 2 | | 0 | 963 |
| 17916 | 4 | | | | | | | | | 0 | 1068 |
| 17863 | 21 | | | | | | | | | 0 | 734 |
| 17857 | 13 | | | | | | | | | 0 | 674 |
| 17762 | 10 | | | | 5568624 | AAEL0069 | AaeL_AAE | 2 | | 0 | 1035 |
| 17758 | 9 | | | | | | | | | 0 | 736 |
| 17720 | 16 | | | | | | | | | 0 | 1028 |
| 17700 | 8 | metabolic process | RNA bindi | Pf01253 | 5573994 | AAEL0108 | AaeL_AAE | 2 | RNA transport | 1 | 884 |
| 17654 | 9 | | | | | | | | | 0 | 958 |
| 17616 | 12 | | | | 5568622 | AAEL0069 | AaeL_AAE | 2 | | 0 | 783 |
| 17598 | 11 | | | | | | | | | 0 | 897 |
| 17596 | 10 | | | | | | | | | 0 | 1032 |
| 17566 | 10 | | | | | | | | | 0 | 977 |
| 17493 | 9 | | | | | | | | | 0 | 473 |
| 17469 | 11 | | | | | | | | | 0 | 761 |
| 17393 | 9 | | protein bi | Pf00241 | 5571976 | AAEL0094 | AaeL_AAE | 1 | | 0 | 532 |

| | | | | | | | | | | |
|---|---|---|---|---|---|---|---|---|---|---|
| 17350 | 7 | membrane | | Pf09360, I | 5564392 | AAEL0042 | AaeL_AAE | 1 | 0 | 631 |
| 17340 | 13 | | | | | | | | 0 | 727 |
| 17330 | 10 | | | | | | | | 0 | 848 |
| 17280 | 10 | | | | | | | | 0 | 854 |
| 17218 | 15 | | | | | | | | 0 | 922 |
| 17193 | 8 | | | | | | | | 0 | 625 |
| 17121 | 16 | | | | | | | | 0 | 519 |
| 17089 | 10 | | | | | | | | 0 | 805 |
| 17064 | 11 | | | | | | | | 0 | 692 |
| 17056 | 10 | | | | | | | | 0 | 646 |
| 17047 | 13 | | | | | | | | 0 | 627 |
| 16985 | 12 | | protein bi | Pf12341 | 5570522 | AAEL0013 | AaeL_AAE | 2 | 0 | 709 |
| 16978 | 10 | | | | | | | | 0 | 650 |
| 16972 | 10 | | | | | | | | 0 | 840 |
| 16961 | 14 | | | | | | | | 0 | 574 |
| 16914 | 13 | | | | | | | | 0 | 773 |
| 16903 | 13 | | | | | | | | 0 | 1131 |
| 16885 | 10 | | | | | | | | 0 | 662 |
| 16833 | 12 | | | | | | | | 0 | 1048 |
| 16827 | 11 | | | | | | | | 0 | 817 |
| 16712 | 10 | | | | | | | | 0 | 830 |
| 16680 | 6 | | | | | | | | 0 | 786 |
| 16678 | 8 | | | | | | | | 0 | 849 |
| 16630 | 18 | | | | | | | | 0 | 911 |
| 16620 | 5 | | | | | | | | 0 | 995 |
| 16601 | 7 | | | | | | | | 0 | 924 |
| 16576 | 9 | | | | | | | | 0 | 1236 |
| 16553 | 14 | | | | | | | | 0 | 593 |
| 16549 | 7 | | | | | | | | 0 | 1292 |
| 16521 | 13 | | | | | | | | 0 | 618 |
| 16482 | 20 | | | | | | | | 0 | 758 |

| | | | | | | | | | |
|---|---|---|---|---|---|---|---|---|---|
| 16472 | 13 | | | | | | | 0 | 891 |
| 16467 | 5 | | | | | | | 0 | 968 |
| 16433 | 16 | | | | | | | 0 | 1003 |
| 16421 | 15 | | | | | | | 0 | 917 |
| 16394 | 8 | | | | | | | 0 | 961 |
| 16379 | 11 | | | | | | | 0 | 934 |
| 16361 | 10 | | | | | | | 0 | 615 |
| 16267 | 11 | | | | | | | 0 | |
| 16263 | 10 | | | | | | | 0 | 743 |
| 16196 | 20 | | | | | | | 0 | 662 |
| 16158 | 10 | metabolic process;regulation o | Pf04111, I | 5570325 | AAEL0013 | AaeL_AAE | 2 | 0 | 916 |
| 16143 | 8 | | | | | | | 0 | 1079 |
| 16122 | 6 | | | | | | | 0 | 994 |
| 16095 | 12 | | | | | | | 0 | 743 |
| 16082 | 11 | | | | | | | 0 | 623 |
| 16060 | 9 | | | | | | | 0 | 805 |
| 16051 | 12 | | | | | | | 0 | 682 |
| 16037 | 9 | | | | | | | 0 | 1066 |
| 16005 | 16 | | | | | | | 0 | 794 |
| 16003 | 13 | | | | | | | 0 | 1097 |
| 15982 | 8 | | | | | | | 0 | 764 |
| 15920 | 7 | | | | | | | 0 | 928 |
| 15915 | 16 | | | | | | | 0 | 562 |
| 15857 | 9 | | Pf05907 | 5566864 | AAEL0008 | AaeL_AAE | 3 | 0 | 949 |
| 15843 | 12 | | | | | | | 0 | 794 |
| 15826 | 14 | | | | | | | 0 | 968 |
| 15811 | 12 | | | | | | | 0 | 1323 |
| 15770 | 8 | | | | | | | 0 | 992 |
| 15767 | 10 | | | | | | | 0 | 1104 |
| 15731 | 14 | | | | | | | 0 | 983 |
| 15714 | 4 | | | | | | | 0 | 723 |

| | | | | | | | | |
|---|---|---|---|---|---|---|---|---|
| 15709 | 5 | | | | | | 0 | 1007 |
| 15699 | 8 | | | | | | 0 | 1154 |
| 15597 | 9 | | | | | | 0 | 942 |
| 15580 | 13 | | | | | | 0 | 783 |
| 15574 | 15 | | | | | | 0 | 913 |
| 15568 | 9 | | | | | | 0 | 930 |
| 15544 | 12 | | | | | | 0 | 568 |
| 15541 | 12 | | | | | | 0 | 771 |
| 15513 | 6 | | | | | | 0 | 828 |
| 15484 | 6 | 5572075 | AAEL0017 | AaeL_AAE | 2 | | 0 | 786 |
| 15456 | 10 | | | | | | 0 | 1030 |
| 15454 | 7 | | | | | | 0 | 885 |
| 15423 | 7 | | | | | | 0 | 871 |
| 15371 | 11 | | | | | | 0 | 721 |
| 15364 | 12 | | | | | | 0 | 934 |
| 15349 | 9 | | | | | | 0 | 997 |
| 15341 | 11 | | | | | | 0 | 824 |
| 15339 | 10 | | | | | | 0 | 833 |
| 15333 | 13 | | | | | | 0 | |
| 15332 | 14 | | | | | | 0 | 863 |
| 15320 | 13 | | | | | | 0 | 726 |
| 15316 | 7 | | | | | | 0 | 580 |
| 15308 | 11 | | | | | | 0 | 799 |
| 15306 | 9 | | | | | | 0 | 607 |
| 15306 | 9 | | | | | | 0 | 895 |
| 15223 | 10 | | | | | | 0 | |
| 15181 | 15 | | | | | | 0 | 711 |
| 15116 | 15 | | | | | | 0 | 773 |
| 15073 | 11 | 5571135 | AAEL0088 | AaeL_AAE | 3 | MAPK signaling pathway - fly | 1 | 1050 |
| 15054 | 9 | | | | | | 0 | 1021 |
| 15048 | 9 | | | | | | 0 | 566 |

| | | | | | | | | | |
|---|---|---|---|---|---|---|---|---|---|
| 15032 | 7 | | | | | | | 0 | 903 |
| 15026 | 12 | | | | | | | 0 | 510 |
| 15020 | 8 | | | | | | | 0 | 1071 |
| 14931 | 11 | | | | | | | 0 | 510 |
| 14920 | 7 | | | | | | | 0 | 871 |
| 14796 | 19 | | | | | | | 0 | 542 |
| 14785 | 7 | | | | | | | 0 | 572 |
| 14783 | 5 | | | | | | | 0 | 1588 |
| 14779 | 11 | | | | | | | 0 | 864 |
| 14774 | 9 | | | | | | | 0 | 685 |
| 14761 | 9 | | | | | | | 0 | 473 |
| 14757 | 8 | | | | | | | 0 | 858 |
| 14737 | 4 | | | | | | | 0 | 791 |
| 14674 | 9 | | | | | | | 0 | 672 |
| 14627 | 19 | | | | | | | 0 | 684 |
| 14619 | 14 | | | | | | | 0 | 762 |
| 14613 | 4 | | | | | | | 0 | 1252 |
| 14609 | 6 | | | | | | | 0 | 1252 |
| 14583 | 5 | | | | | | | 0 | 661 |
| 14583 | 7 | | | 5575980 | AAEL0122 | AaeL_AAE | 2 | 0 | 955 |
| 14468 | 5 | catalytic a | Pf01370, I | 5565330 | AAEL0148 | AaeL_AAE | 1 | 0 | 703 |
| 14426 | 6 | | | | | | | 0 | 938 |
| 14415 | 19 | | | | | | | 0 | 494 |
| 14404 | 10 | | | | | | | 0 | 708 |
| 14361 | 10 | | | | | | | 0 | 493 |
| 14360 | 4 | | | | | | | 0 | 911 |
| 14328 | 6 | | | | | | | 0 | 763 |
| 14312 | 5 | | | | | | | 0 | 1014 |
| 14311 | 10 | | | | | | | 0 | 1117 |
| 14298 | 6 | | | | | | | 0 | 925 |
| 14294 | 9 | | | | | | | 0 | 941 |

| | | | | | | | | | | | |
|---|---|---|---|---|---|---|---|---|---|---|---|
| 14252 | 11 | | | | | | | | | 0 | 694 |
| 14238 | 10 | | | | | | | | | 0 | 956 |
| 14221 | 8 | | | | | | | | | 0 | 625 |
| 14209 | 10 | | | | | | | | | 0 | 603 |
| 14204 | 9 | | | | | | | | | 0 | 1164 |
| 14200 | 6 | | | | | | | | | 0 | 1050 |
| 14142 | 9 | | | | | | | | | 0 | 795 |
| 14082 | 12 | | | | | | | | | 0 | 513 |
| 14059 | 13 | | | | | | | | | 0 | 833 |
| 14057 | 9 | | | | | | | | | 0 | 899 |
| 14016 | 6 | | | | | | | | | 0 | 930 |
| 14011 | 7 | | protein bi | Pf04003 | 5579658 | AAEL0039 | AaeL_AAE | 3 | Ribosome biogenesis in eukary | 1 | 755 |
| 13979 | 12 | | | | | | | | | 0 | 901 |
| 13979 | 11 | | | | | | | | | 0 | 886 |
| 13963 | 7 | | | | | | | | | 0 | 921 |
| 13961 | 7 | metabolic nucleus | | | 5563778 | AAEL0039 | AaeL_AAE | 2 | Metabolic pathways; Mismatch | 8 | 897 |
| 13928 | 13 | | | | | | | | | 0 | 648 |
| 13921 | 5 | | metal ion binding | | 23687839 | AAEL0174 | AAEL8011 | 2 | | 0 | 929 |
| 13889 | 5 | | | | | | | | | 0 | 1037 |
| 13887 | 12 | | | | | | | | | 0 | 741 |
| 13886 | 9 | | | | | | | | | 0 | 654 |
| 13879 | 15 | | | | | | | | | 0 | 735 |
| 13839 | 9 | | | | | | | | | 0 | 742 |
| 13807 | 10 | | | | | | | | | 0 | 1359 |
| 13805 | 7 | | | | | | | | | 0 | 901 |
| 13789 | 16 | | | | | | | | | 0 | 678 |
| 13788 | 12 | | | | | | | | | 0 | 786 |
| 13785 | 11 | | | | | | | | | 0 | 627 |
| 13748 | 7 | | | | | | | | | 0 | 770 |
| 13728 | 10 | | | | | | | | | 0 | 887 |
| 13713 | 9 | | | | | | | | | 0 | 853 |

| | | | | | | | | | | | |
|---|---|---|---|---|---|---|---|---|---|---|---|
| 13695 | 11 | | | | | | | | | 0 | 806 |
| 13645 | 9 | | | | | | | | | 0 | 998 |
| 13640 | 14 | | | | | | | | | 0 | 1022 |
| 13575 | 11 | | | | | | | | | 0 | 807 |
| 13550 | 5 | | | | | | | | | 0 | 881 |
| 13546 | 17 | | | | | | | | | 0 | 719 |
| 13530 | 12 | | | | | | | | | 0 | 1087 |
| 13502 | 4 | | | | | | | | | 0 | 801 |
| 13491 | 9 | | | | | | | | | 0 | 800 |
| 13470 | 13 | | | | | | | | | 0 | 742 |
| 13461 | 10 | regulation of biologi | | enzyme re | Pf00595, I | 5577284 | AAEL0131 | AaeL_AAE | 3 | 0 | 935 |
| 13388 | 10 | | | | | | | | | 0 | 848 |
| 13361 | 13 | | | | | | | | | 0 | 636 |
| 13353 | 11 | | | | | | | | | 0 | 767 |
| 13341 | 8 | | | | | | | | | 0 | 1068 |
| 13326 | 10 | | | | | | | | | 0 | 714 |
| 13263 | 16 | | | | | | | | | 0 | 1084 |
| 13255 | 17 | | | | | | | | | 0 | 783 |
| 13253 | 7 | | | | | | | | | 0 | 706 |
| 13249 | 7 | | | | | | | | | 0 | 1066 |
| 13242 | 11 | | | | | | | | | 0 | 598 |
| 13241 | 12 | | | | | | | | | 0 | 915 |
| 13211 | 9 | | | | | | | | | 0 | 491 |
| 13204 | 9 | | | | | | | | | 0 | 958 |
| 13204 | 9 | | | | | | | | | 0 | 1093 |
| 13146 | 6 | | | | | | | | | 0 | 819 |
| 13139 | 10 | | | | | | | | | 0 | |
| 13124 | 11 | | membrane | | | 5576364 | AAEL0028 | AaeL_AAE | 1 | 0 | 614 |
| 13095 | 5 | | | | | | | | | 0 | 731 |
| 13085 | 8 | | | | | | | | | 0 | 624 |
| 13080 | 9 | | | | | | | | | 0 | 643 |

| | | | | | | | | | |
|---|---|---|---|---|---|---|---|---|---|
| 13051 | 5 | | | | | | | 0 | 1545 |
| 13040 | 7 | | | | | | | 0 | 924 |
| 13005 | 9 | | | | | | | 0 | 907 |
| 12978 | 13 | | | | | | | 0 | 918 |
| 12967 | 8 | | | | | | | 0 | 567 |
| 12965 | 7 | | | | | | | 0 | 849 |
| 12960 | 10 | | | | | | | 0 | 793 |
| 12957 | 3 | | | | | | | 0 | 1101 |
| 12950 | 14 | | | | | | | 0 | 562 |
| 12947 | 14 | | | | | | | 0 | 686 |
| 12941 | 7 | | | | | | | 0 | 674 |
| 12923 | 9 | | | | | | | 0 | 603 |
| 12922 | 8 | | | | | | | 0 | 469 |
| 12916 | 10 | | | | | | | 0 | 471 |
| 12829 | 8 | | | | | | | 0 | 963 |
| 12806 | 9 | regulation of biological process | Pf04156, I | 5564411 | AAEL0182 | AaeL_AAE | 2 | 0 | 925 |
| 12795 | 12 | | | | | | | 0 | 1032 |
| 12791 | 5 | | | | | | | 0 | 585 |
| 12781 | 8 | | | | | | | 0 | 780 |
| 12776 | 10 | | | | | | | 0 | 832 |
| 12765 | 8 | | | | | | | 0 | 1243 |
| 12738 | 10 | | | | | | | 0 | 519 |
| 12724 | 9 | | | | | | | 0 | 1164 |
| 12717 | 10 | | | | | | | 0 | 873 |
| 12692 | 6 | | | | | | | 0 | 830 |
| 12688 | 12 | | | | | | | 0 | 1222 |
| 12682 | 10 | | | | | | | 0 | 524 |
| 12664 | 5 | | | | | | | 0 | 996 |
| 12661 | 8 | | | | | | | 0 | 775 |
| 12649 | 10 | | | | | | | 0 | 826 |
| 12618 | 11 | | | | | | | 0 | 763 |

| | | | | | | | | | | |
|---|---|---|---|---|---|---|---|---|---|---|
| 12588 | 13 | | | | | | | | 0 | 473 |
| 12566 | 12 | | | | | | | | 0 | 799 |
| 12557 | 5 | | | | | | | | 0 | 842 |
| 12453 | 10 | protein bi | Pf12569, I | 5567421 | AAEL0060 | AaeL_AAE | 1 | | 0 | 586 |
| 12447 | 7 | | | | | | | | 0 | 617 |
| 12443 | 5 | | | | | | | | 0 | 1338 |
| 12435 | 10 | | | | | | | | 0 | 724 |
| 12415 | 9 | | | | | | | | 0 | 772 |
| 12404 | 10 | | | | | | | | 0 | 833 |
| 12403 | 9 | | | | | | | | 0 | 921 |
| 12399 | 12 | | | | | | | | 0 | 1012 |
| 12388 | 7 | | | | | | | | 0 | 867 |
| 12373 | 7 | | | | | | | | 0 | 550 |
| 12343 | 11 | | | | | | | | 0 | 685 |
| 12246 | 7 | | | | | | | | 0 | 998 |
| 12208 | 8 | | | | | | | | 0 | 857 |
| 12186 | 6 | | | | | | | | 0 | 918 |
| 12185 | 8 | | | | | | | | 0 | 675 |
| 12135 | 7 | | | | | | | | 0 | 897 |
| 12103 | 9 | | | | | | | | 0 | 406 |
| 12095 | 7 | | | | | | | | 0 | 703 |
| 12083 | 10 | | | | | | | | 0 | 1008 |
| 12079 | 13 | protein bi | Pf03177, I | 5576401 | AAEL0124 | AaeL_AAE | 1 | RNA transport | 1 | 631 |
| 12074 | 8 | | Pf05048, I | 5573181 | AAEL0020 | AaeL_AAE | 2 | | 0 | 692 |
| 12065 | 9 | | | | | | | | 0 | 619 |
| 12044 | 9 | | | | | | | | 0 | 940 |
| 11951 | 8 | | | | | | | | 0 | 327 |
| 11944 | 8 | | | | | | | | 0 | 1107 |
| 11923 | 10 | | | | | | | | 0 | 763 |
| 11896 | 7 | | | | | | | | 0 | 912 |
| 11888 | 11 | | | | | | | | 0 | 878 |

| | | | | | | | | | |
|---|---|---|---|---|---|---|---|---|---|
| 11884 | 5 | | | | | | | 0 | 976 |
| 11882 | 8 | | | | | | | 0 | 883 |
| 11881 | 3 | | | | | | | 0 | |
| 11874 | 8 | Pf10249 | 5572650 | AAEL0018 | AaeL_AAE | 2 | Metabolic pathways; Oxidative | 2 | 548 |
| 11869 | 10 | | | | | | | 0 | 915 |
| 11828 | 5 | | | | | | | 0 | 1072 |
| 11821 | 6 | | | | | | | 0 | 766 |
| 11788 | 7 | | | | | | | 0 | 515 |
| 11786 | 9 | | | | | | | 0 | 765 |
| 11785 | 9 | | | | | | | 0 | 623 |
| 11779 | 4 | | | | | | | 0 | 753 |
| 11774 | 11 | | | | | | | 0 | 918 |
| 11761 | 5 | | | | | | | 0 | 983 |
| 11713 | 10 | | | | | | | 0 | 814 |
| 11712 | 11 | | | | | | | 0 | 830 |
| 11707 | 6 | | | | | | | 0 | 672 |
| 11705 | 13 | | | | | | | 0 | 570 |
| 11683 | 7 | | | | | | | 0 | 809 |
| 11674 | 7 | | | | | | | 0 | 803 |
| 11647 | 11 | | | | | | | 0 | 740 |
| 11646 | 8 | | | | | | | 0 | 987 |
| 11639 | 11 | | | | | | | 0 | 758 |
| 11634 | 5 | | | | | | | 0 | 957 |
| 11634 | 11 | | | | | | | 0 | 685 |
| 11611 | 10 | | | | | | | 0 | 519 |
| 11606 | 8 | | | | | | | 0 | 747 |
| 11603 | 10 | | | | | | | 0 | 700 |
| 11599 | 7 | | | | | | | 0 | 1340 |
| 11595 | 11 | | | | | | | 0 | 898 |
| 11531 | 8 | | | | | | | 0 | 561 |
| 11502 | 7 | | | | | | | 0 | 709 |

| | | | | | | | | | | | |
|---|---|---|---|---|---|---|---|---|---|---|---|
| 11481 | 9 | | | | | | | | | 0 | 1286 |
| 11477 | 5 | | | | | | | | | 0 | 1034 |
| 11475 | 9 | | | | | | | | | 0 | 676 |
| 11451 | 7 | | | | | | | | | 0 | 711 |
| 11439 | 8 | | | | | | | | | 0 | 886 |
| 11423 | 10 | | | | | | | | | 0 | 758 |
| 11410 | 8 | | | | | | | | | 0 | 673 |
| 11403 | 6 | | | | | | | | | 0 | 793 |
| 11394 | 10 | | | | | | | | | 0 | 801 |
| 11354 | 8 | | | | | | | | | 0 | 920 |
| 11344 | 7 | | | | | | | | | 0 | 941 |
| 11343 | 11 | | | | | | | | | 0 | 605 |
| 11323 | 6 | | | | | | | | | 0 | 993 |
| 11306 | 5 | | | | | | | | | 0 | 810 |
| 11267 | 9 | | | | | | | | | 0 | 588 |
| 11255 | 7 | | | | | | | | | 0 | 820 |
| 11237 | 12 | | | | | | | | | 0 | 611 |
| 11227 | 7 | | protein bi | Pf00240, I | 5579053 | AAEL0139 | AaeL_AAE | 2 | Protein processing in endoplas | 1 | 946 |
| 11224 | 9 | | | | | | | | | 0 | 644 |
| 11167 | 10 | | | | | | | | | 0 | 1073 |
| 11158 | 6 | | | | | | | | | 0 | 1128 |
| 11147 | 3 | | | | | | | | | 0 | 1011 |
| 11133 | 6 | metabolic process | catalytic a | Pf08240, I | 5575162 | AAEL0003 | AaeL_AAE | 2 | | 0 | 642 |
| 11069 | 11 | | | | | | | | | 0 | 761 |
| 11064 | 8 | | | | | | | | | 0 | 506 |
| 11034 | 11 | | | | | | | | | 0 | 759 |
| 11026 | 6 | | | | | | | | | 0 | 803 |
| 11011 | 10 | | | | | | | | | 0 | 816 |
| 10983 | 9 | | | | | | | | | 0 | 759 |
| 10983 | 11 | | | | | | | | | 0 | 781 |
| 10976 | 8 | | | | | | | | | 0 | 1320 |

| | | | | | | | | | | |
|---|---|---|---|---|---|---|---|---|---|---|
| 10957 | 10 | | | | | | | | 0 | 1005 |
| 10951 | 9 | | | | | | | | 0 | 1137 |
| 10935 | 9 | | | | | | | | 0 | 506 |
| 10915 | 6 | | Pf00581 | 5569037 | AAEL0073 | AaeL_AAE | 2 | | 0 | 832 |
| 10896 | 9 | | | | | | | | 0 | 745 |
| 10894 | 9 | | | | | | | | 0 | 506 |
| 10888 | 8 | | | | | | | | 0 | 636 |
| 10857 | 9 | | | | | | | | 0 | 558 |
| 10838 | 2 | | | | | | | | 0 | 967 |
| 10800 | 9 | | | | | | | | 0 | 546 |
| 10794 | 7 | | | | | | | | 0 | 454 |
| 10788 | 9 | | | | | | | | 0 | 692 |
| 10779 | 8 | | | | | | | | 0 | 505 |
| 10773 | 8 | | | | | | | | 0 | 1023 |
| 10755 | 10 | | | | | | | | 0 | 923 |
| 10728 | 6 | | | | | | | | 0 | 958 |
| 10727 | 9 | | | | | | | | 0 | 787 |
| 10680 | 11 | | | | | | | | 0 | 643 |
| 10679 | 8 | | | | | | | | 0 | 1037 |
| 10664 | 3 | | | | | | | | 0 | 557 |
| 10651 | 7 | | | | | | | | 0 | 659 |
| 10641 | 8 | | | | | | | | 0 | 759 |
| 10633 | 8 | | | | | | | | 0 | 663 |
| 10540 | 7 | | | | | | | | 0 | 856 |
| 10529 | 6 | | | | | | | | 0 | 1064 |
| 10522 | 10 | | | | | | | | 0 | 824 |
| 10511 | 7 | | | | | | | | 0 | 916 |
| 10505 | 8 | | | | | | | | 0 | 589 |
| 10486 | 13 | catalytic a | Pf01968, I | 5564314 | AAEL0005 | AaeL_AAE | 3 | Glutathione metabolism | 1 | 751 |
| 10485 | 6 | | | | | | | | 0 | 750 |
| 10438 | 8 | | | | | | | | 0 | 498 |

| | | | | | | | | | |
|---|---|---|---|---|---|---|---|---|---|
| 10433 | 9 | | | | | | | 0 | 615 |
| 10413 | 7 | | | | | | | 0 | 647 |
| 10411 | 9 | protein binding | | 5576199 | AAEL0123 | AaeL_AAE | 1 | 0 | 736 |
| 10392 | 7 | | | | | | | 0 | 711 |
| 10379 | 5 | | | | | | | 0 | 1043 |
| 10340 | 6 | | | | | | | 0 | 820 |
| 10320 | 7 | | | | | | | 0 | 798 |
| 10316 | 6 | | | | | | | 0 | 937 |
| 10306 | 9 | | | | | | | 0 | 599 |
| 10302 | 2 | | | 5567181 | AAEL0009 | AaeL_AAE | 3 | 0 | 699 |
| 10284 | 8 | | | | | | | 0 | 969 |
| 10268 | 8 | | | | | | | 0 | 676 |
| 10241 | 5 | | | | | | | 0 | 773 |
| 10216 | 4 | | | | | | | 0 | 613 |
| 10125 | 8 | | | | | | | 0 | 894 |
| 10097 | 3 | | | | | | | 0 | 882 |
| 10075 | 8 | | | | | | | 0 | 827 |
| 10062 | 8 | | | | | | | 0 | 614 |
| 10042 | 4 | | | | | | | 0 | 974 |
| 10000 | 8 | | | | | | | 0 | 873 |
| 9988 | 9 | | | | | | | 0 | 839 |
| 9976 | 8 | | | | | | | 0 | 728 |
| 9962 | 10 | | | | | | | 0 | 718 |
| 9950 | 4 | | | | | | | 0 | 861 |
| 9940 | 8 | | | | | | | 0 | 949 |
| 9910 | 10 | | | | | | | 0 | 784 |
| 9899 | 10 | | | | | | | 0 | 881 |
| 9879 | 6 | | | | | | | 0 | 753 |
| 9862 | 5 | | | | | | | 0 | 529 |
| 9840 | 3 | | | 5571752 | AAEL0092 | AaeL_AAE | 3 | 0 | 1191 |
| 9838 | 4 | | Pf00226 | 5568679 | AAEL0070 | AaeL_AAE | 1 | 0 | 937 |

| | | | | | | | | | | |
|---|---|---|---|---|---|---|---|---|---|---|
| 9838 | 4 | | | | | | | | 0 | 946 |
| 9810 | 5 | | | | | | | | 0 | 586 |
| 9801 | 6 | | | | | | | | 0 | 834 |
| 9793 | 6 | | | | | | | | 0 | 511 |
| 9791 | 2 | | | | | | | | 0 | 961 |
| 9790 | 9 | | | Pf10228, I | 5569797 | AAEL0013 | AaeL_AAE | 3 | 0 | 782 |
| 9788 | 7 | | | | | | | | 0 | 625 |
| 9772 | 7 | | | | | | | | 0 | 469 |
| 9769 | 8 | | protein bi | Pf02179 | 5567242 | AAEL0008 | AaeL_AAE | 3 | 0 | 955 |
| 9762 | 7 | | | | | | | | 0 | 750 |
| 9709 | 7 | | | | | | | | 0 | 751 |
| 9706 | 10 | | | | | | | | 0 | 753 |
| 9705 | 9 | | | | | | | | 0 | 872 |
| 9689 | 9 | | | | | | | | 0 | 1210 |
| 9680 | 7 | | | | | | | | 0 | 668 |
| 9679 | 4 | | | | | | | | 0 | 1093 |
| 9663 | 8 | | | | | | | | 0 | 1253 |
| 9643 | 4 | | protein binding | | 5569343 | AAEL0075 | AaeL_AAE | 2 | 0 | 980 |
| 9627 | 7 | | | | | | | | 0 | 926 |
| 9624 | 12 | | | | | | | | 0 | 763 |
| 9622 | 4 | | | | | | | | 0 | 530 |
| 9610 | 5 | | | | | | | | 0 | 866 |
| 9573 | 9 | | | | | | | | 0 | 1027 |
| 9546 | 6 | nucleus | | Pf00385, I | 5564858 | AAEL0044 | AaeL_AAE | 3 | 0 | 1083 |
| 9544 | 5 | | | | | | | | 0 | 1134 |
| 9530 | 11 | | | | | | | | 0 | 896 |
| 9520 | 10 | | | | | | | | 0 | 859 |
| 9512 | 6 | | | | | | | | 0 | 993 |
| 9508 | 5 | | | | | | | | 0 | 896 |
| 9507 | 6 | | | | | | | | 0 | 1179 |
| 9506 | 13 | | | | | | | | 0 | 974 |

| | | | | | | | | | | | |
|---|---|---|---|---|---|---|---|---|---|---|---|
| 9498 | 7 | cellular hc | membrane | | Pf00085 | 5564319 | AAEL0005 | AaeL_AAE | 3 | 0 | 355 |
| 9489 | 4 | | | | | | | | | 0 | 859 |
| 9489 | 5 | | | | | | | | | 0 | 964 |
| 9482 | 6 | | | | | | | | | 0 | 429 |
| 9479 | 9 | | | | | | | | | 0 | 757 |
| 9475 | 6 | | | | | | | | | 0 | 875 |
| 9473 | 8 | | | | | | | | | 0 | 1147 |
| 9456 | 10 | | | | | | | | | 0 | 849 |
| 9448 | 4 | | | | | 5568054 | AAEL0064 | AaeL_AAE | 3 | 0 | 567 |
| 9444 | 9 | | | | | | | | | 0 | 583 |
| 9423 | 5 | | | | | | | | | 0 | 881 |
| 9416 | 9 | | | | | | | | | 0 | 711 |
| 9401 | 6 | | | | | | | | | 0 | 870 |
| 9398 | 7 | | | | | | | | | 0 | 689 |
| 9365 | 4 | | | | | | | | | 0 | 925 |
| 9358 | 4 | | | | | | | | | 0 | 615 |
| 9333 | 8 | | | | | | | | | 0 | 672 |
| 9329 | 7 | metabolic process | | | Pf10185 | 5573465 | AAEL0105 | AaeL_AAE | 2 | 0 | 1178 |
| 9317 | 6 | | | | | | | | | 0 | 1056 |
| 9312 | 6 | | | | | | | | | 0 | 704 |
| 9287 | 8 | | | | | | | | | 0 | 611 |
| 9271 | 7 | | | | | | | | | 0 | 818 |
| 9267 | 4 | | | RNA bindi | Pf00076, I | 5575846 | AAEL0121 | AaeL_AAE | 2 | 0 | 1028 |
| 9264 | 6 | | | | | | | | | 0 | 827 |
| 9255 | 8 | | | | | | | | | 0 | 551 |
| 9245 | 6 | | | | Pf06244 | 5570608; | AAEL0084 | AaeL_AAE | 1 | 0 | 872 |
| 9244 | 8 | | | | | | | | | 0 | 1144 |
| 9232 | 4 | | | | | | | | | 0 | 898 |
| 9196 | 8 | | | | | | | | | 0 | 606 |
| 9189 | 6 | | | | | | | | | 0 | 820 |
| 9186 | 6 | | | | | | | | | 0 | 509 |

| | | | | | | | | | | | |
|---|---|---|---|---|---|---|---|---|---|---|---|
| 9177 | 8 | | | | | | | | | 0 | 775 |
| 9170 | 4 | transport | | Pf07200 | 5574138 | AAEL0109 | AaeL_AAE | 3 | Endocytosis | 1 | 871 |
| 9155 | 8 | | | | | | | | | 0 | 1064 |
| 9150 | 4 | metabolic membran | catalytic a | Pf01121, I | 5577808 | AAEL0133 | AaeL_AAE | 3 | | 0 | 589 |
| 9135 | 8 | | | | | | | | | 0 | 729 |
| 9124 | 10 | | | Pf03828 | 5564549 | AAEL0043 | AaeL_AAE | 3 | | 0 | 723 |
| 9089 | 6 | | | | | | | | | 0 | 572 |
| 9073 | 8 | | protein bi | Pf02239 | 5571653 | AAEL0016 | AaeL_AAE | 2 | | 0 | 850 |
| 9072 | 7 | | | | | | | | | 0 | 801 |
| 9062 | 9 | | | | | | | | | 0 | 874 |
| 9054 | 9 | | | Pf08190 | 5565384 | AAEL0047 | AaeL_AAE | 2 | | 0 | 939 |
| 9048 | 8 | | | | | | | | | 0 | 720 |
| 9041 | 7 | | | | | | | | | 0 | 1079 |
| 9036 | 8 | | | | | | | | | 0 | 1033 |
| 9034 | 8 | | | | | | | | | 0 | 911 |
| 9010 | 7 | | | | | | | | | 0 | 636 |
| 8999 | 7 | | | | | | | | | 0 | 681 |
| 8977 | 2 | | | | | | | | | 0 | 881 |
| 8947 | 11 | | | | | | | | | 0 | 485 |
| 8944 | 5 | | | | | | | | | 0 | 606 |
| 8916 | 9 | | | | | | | | | 0 | 950 |
| 8884 | 5 | | | | | | | | | 0 | 864 |
| 8884 | 10 | | | | | | | | | 0 | 784 |
| 8883 | 10 | | | | | | | | | 0 | 883 |
| 8877 | 8 | | | | | | | | | 0 | 864 |
| 8877 | 8 | | | | | | | | | 0 | 932 |
| 8874 | 6 | | | | | | | | | 0 | 716 |
| 8873 | 2 | | | | | | | | | 0 | 674 |
| 8871 | 4 | | | | | | | | | 0 | 985 |
| 8850 | 4 | | | Pf15430 | 5570040 | AAEL0002 | AaeL_AAE | 3 | | 0 | 1005 |
| 8845 | 7 | | | | | | | | | 0 | 512 |

| | | | | | | | | | | |
|---|---|---|---|---|---|---|---|---|---|---|
| 8842 | 9 | | | | | | | | 0 | 609 |
| 8837 | 8 | | | | | | | | 0 | 741 |
| 8826 | 10 | metabolic process | catalytic a | Pf00704 | 5567905 | AAEL0063 | AaeL_AAE | 3 | 0 | 830 |
| 8822 | 8 | | | | | | | | 0 | 228 |
| 8818 | 7 | | | | | | | | 0 | 981 |
| 8812 | 8 | | | | | | | | 0 | 901 |
| 8797 | 6 | | | | | | | | 0 | 1097 |
| 8795 | 8 | | | | | | | | 0 | 536 |
| 8791 | 7 | | | | | | | | 0 | 718 |
| 8789 | 7 | | | | | | | | 0 | 656 |
| 8773 | 3 | | | | | | | | 0 | 218 |
| 8769 | 8 | | | | | | | | 0 | 669 |
| 8769 | 8 | | | | | | | | 0 | 1064 |
| 8768 | 7 | | | | | | | | 0 | 1132 |
| 8766 | 6 | | | | | | | | 0 | 912 |
| 8763 | 6 | | | | | | | | 0 | 878 |
| 8757 | 6 | | | | | | | | 0 | 741 |
| 8755 | 7 | | | | | | | | 0 | 684 |
| 8746 | 7 | | | | | | | | 0 | 953 |
| 8739 | 7 | | | | | | | | 0 | 792 |
| 8698 | 5 | | | | | | | | 0 | 892 |
| 8692 | 7 | | | | | | | | 0 | 722 |
| 8687 | 10 | | | | | | | | 0 | 841 |
| 8680 | 5 | | protein bi | Pf07648 | 5570015 | AAEL0002 | AaeL_AAE | 3 | 0 | 645 |
| 8645 | 7 | | | | | | | | 0 | 566 |
| 8637 | 7 | | protein bi | Pf00646, I | 5570170 | AAEL0081 | AaeL_AAE | 3 | 0 | 353 |
| 8635 | 7 | | | | | | | | 0 | 1001 |
| 8627 | 9 | | | | | | | | 0 | 688 |
| 8625 | 7 | | | | | | | | 0 | 399 |
| 8603 | 6 | | | | | | | | 0 | 924 |
| 8584 | 5 | | | | | | | | 0 | 776 |

| | | | | | | | | | | |
|---|---|---|---|---|---|---|---|---|---|---|
| 8570 | 8 | | | | | | | | 0 | 561 |
| 8562 | 8 | | | | | | | | 0 | 748 |
| 8555 | 7 | | | | | | | | 0 | 614 |
| 8550 | 5 | | | | | | | | 0 | 892 |
| 8548 | 6 | | | | | | | | 0 | 997 |
| 8542 | 7 | | | | | | | | 0 | 793 |
| 8540 | 8 | | | | | | | | 0 | 840 |
| 8517 | 6 | | | | | | | | 0 | 751 |
| 8512 | 3 | regulation of biologi | RNA bindi | Pf00013, I | 5569043 | AAEL0073 | AaeL_AAE | 2 | 0 | 1335 |
| 8506 | 9 | | | | | | | | 0 | 530 |
| 8504 | 8 | | | Pf05600 | 5574482; | AAEL0023 | AaeL_AAE | 2 | 0 | 751 |
| 8501 | 9 | | | | | | | | 0 | 946 |
| 8484 | 6 | | | | | | | | 0 | 867 |
| 8475 | 8 | | | | | | | | 0 | 1168 |
| 8447 | 7 | | | | | | | | 0 | 513 |
| 8430 | 6 | | | | | | | | 0 | 1082 |
| 8423 | 10 | | | | | | | | 0 | 896 |
| 8420 | 5 | | | | | | | | 0 | 359 |
| 8414 | 6 | | | | | | | | 0 | 819 |
| 8396 | 5 | | | | | | | | 0 | 1266 |
| 8394 | 9 | | | | | | | | 0 | 733 |
| 8371 | 3 | | | | | | | | 0 | 1090 |
| 8369 | 7 | | | | | | | | 0 | 934 |
| 8358 | 5 | | | | | | | | 0 | 634 |
| 8310 | 6 | | | | | | | | 0 | 696 |
| 8309 | 8 | | | | | | | | 0 | 639 |
| 8290 | 4 | | | | | | | | 0 | 1290 |
| 8278 | 4 | | | | | | | | 0 | 833 |
| 8258 | 7 | | | | | | | | 0 | 584 |
| 8234 | 8 | | | | | | | | 0 | 832 |
| 8225 | 7 | | | | | | | | 0 | 1327 |

| | | | | | | | | | | | |
|---|---|---|---|---|---|---|---|---|---|---|---|
| 8222 | 5 | | | | | | | | | 0 | 743 |
| 8216 | 7 | metabolic process | protein bi | Pf01920 | 5572640 | AAEL0018 | AaeL_AAE | 2 | | 0 | 950 |
| 8213 | 5 | | | | | | | | | 0 | 322 |
| 8184 | 7 | | | | | | | | | 0 | 1136 |
| 8158 | 6 | | | | | | | | | 0 | 819 |
| 8157 | 7 | | membrane | Pf03227 | 5573242 | AAEL0103 | AaeL_AAE | 2 | | 0 | 754 |
| 8150 | 7 | | | | | | | | | 0 | 752 |
| 8143 | 4 | | | | | | | | | 0 | 620 |
| 8139 | 5 | | | | | | | | | 0 | 899 |
| 8130 | 6 | | | | | | | | | 0 | 720 |
| 8083 | 13 | | | | | | | | | 0 | 978 |
| 8058 | 8 | | | | | | | | | 0 | 904 |
| 8048 | 7 | | | | | | | | | 0 | 932 |
| 8012 | 6 | | | | | | | | | 0 | 983 |
| 8011 | 7 | | | | | | | | | 0 | 782 |
| 7998 | 7 | | | | | | | | | 0 | 836 |
| 7972 | 7 | | | | | | | | | 0 | 545 |
| 7969 | 6 | | | | | | | | | 0 | 642 |
| 7955 | 7 | | | | | | | | | 0 | 460 |
| 7949 | 10 | | | | | | | | | 0 | 422 |
| 7930 | 5 | | | | | | | | | 0 | 1160 |
| 7922 | 5 | | | | | | | | | 0 | 625 |
| 7922 | 4 | metabolic process;tr | transporte | Pf04718 | 5568078 | AAEL0065 | AaeL_AAE | 2 | Oxidative phosphorylation; Me | 2 | 547 |
| 7896 | 7 | | | | | | | | | 0 | 900 |
| 7879 | 6 | | | | | | | | | 0 | 756 |
| 7862 | 4 | | | | | | | | | 0 | 1044 |
| 7858 | 7 | | | | | | | | | 0 | 982 |
| 7856 | 7 | | | | | | | | | 0 | 771 |
| 7847 | 8 | | | | | | | | | 0 | 677 |
| 7834 | 7 | | | | | | | | | 0 | 928 |
| 7830 | 5 | | | | | | | | | 0 | 809 |

| | | | | | | | | | |
|---|---|---|---|---|---|---|---|---|---|
| 7825 | 6 | | | | | | | 0 | 899 |
| 7823 | 5 | | | | | | | 0 | 893 |
| 7812 | 7 | | | | | | | 0 | 748 |
| 7804 | 6 | | | | | | | 0 | 749 |
| 7789 | 6 | | | | | | | 0 | 630 |
| 7783 | 3 | | | | | | | 0 | 878 |
| 7774 | 6 | | Pf03747 | 5570310 | AAEL0082 | AaeL_AAE | 3 | 0 | 793 |
| 7764 | 8 | | | | | | | 0 | 1020 |
| 7762 | 7 | | | | | | | 0 | 1034 |
| 7750 | 8 | DNA bindi | Pf02201, I | 5576914 | AAEL0128 | AaeL_AAE | 1 | 0 | 1268 |
| 7741 | 3 | | | | | | | 0 | 404 |
| 7741 | 2 | | | | | | | 0 | 1268 |
| 7738 | 8 | | | | | | | 0 | 791 |
| 7736 | 6 | | | | | | | 0 | 501 |
| 7718 | 3 | | | | | | | 0 | 934 |
| 7717 | 5 | | | | | | | 0 | 308 |
| 7712 | 7 | | | | | | | 0 | 357 |
| 7711 | 6 | | | | | | | 0 | 739 |
| 7697 | 4 | RNA bindi | Pf00076, I | 23687450 | AAEL0170 | AaeL_AAE | 2 | 0 | 883 |
| 7682 | 5 | protein bi | Pf05773 | 5569458 | AAEL0076 | AaeL_AAE | 1 | 0 | 715 |
| 7679 | 6 | | | | | | | 0 | 994 |
| 7677 | 5 | | | | | | | 0 | 574 |
| 7661 | 6 | | | | | | | 0 | 988 |
| 7654 | 5 | | | | | | | 0 | 839 |
| 7651 | 6 | | | | | | | 0 | 954 |
| 7638 | 4 | | | | | | | 0 | 704 |
| 7601 | 4 | | | | | | | 0 | 1238 |
| 7576 | 5 | | | | | | | 0 | 840 |
| 7575 | 5 | | | | | | | 0 | 713 |
| 7573 | 7 | | | | | | | 0 | 755 |
| 7566 | 6 | | | | | | | 0 | 592 |

| | | | | | | | | | | | | |
|---|---|---|---|---|---|---|---|---|---|---|---|---|
| 7551 | 4 | | | | | | | | | | 0 | 1092 |
| 7546 | 2 | | | | | | | | | | 0 | 738 |
| 7545 | 6 | | | | | | | | | | 0 | 972 |
| 7529 | 8 | | | | | | | | | | 0 | 837 |
| 7526 | 3 | | | | | | | | | | 0 | 883 |
| 7524 | 7 | | | | | | | | | | 0 | 805 |
| 7517 | 6 | | | | | | | | | | 0 | 738 |
| 7504 | 5 | | membrane | | Pf01299 | 5573863 | AAEL0107 | AaeL_AAE | 2 | Phagosome; Lysosome | 2 | 718 |
| 7503 | 7 | | | | | | | | | | 0 | 877 |
| 7500 | 5 | | | | | | | | | | 0 | 1045 |
| 7496 | 6 | | | protein bi | Pf08513, I | 5564055 | AAEL0040 | AaeL_AAE | 1 | | 0 | 968 |
| 7444 | 4 | metabolic | nucleus;s | RNA bindi | Pf01423 | 23687581 | AAEL0171 | AaeL_AAE | 2 | Spliceosome; RNA degradation | 2 | 991 |
| 7440 | 6 | | | | | | | | | | 0 | 1027 |
| 7438 | 2 | | | | | | | | | | 0 | 709 |
| 7429 | 4 | | | | | | | | | | 0 | 968 |
| 7426 | 6 | | | | | | | | | | 0 | 923 |
| 7402 | 5 | | | | | | | | | | 0 | 875 |
| 7387 | 6 | | | | | | | | | | 0 | 1039 |
| 7385 | 7 | | | | | | | | | | 0 | 928 |
| 7381 | 8 | | | | | | | | | | 0 | 645 |
| 7368 | 8 | | | metal ion binding | | 5572229 | AAEL0181 | AaeL_AAE | 3 | | 0 | 882 |
| 7348 | 11 | | | | | | | | | | 0 | 965 |
| 7345 | 3 | | | | | | | | | | 0 | 830 |
| 7339 | 9 | | | | | | | | | | 0 | 1078 |
| 7333 | 10 | | | | | | | | | | 0 | |
| 7332 | 5 | | | | | | | | | | 0 | 887 |
| 7319 | 8 | | | | | | | | | | 0 | 661 |
| 7317 | 10 | | | | | | | | | | 0 | 689 |
| 7307 | 5 | | | | Pf00226 | 5567162 | AAEL0059 | AaeL_AAE | 3 | | 0 | 870 |
| 7301 | 6 | | | | | | | | | | 0 | 771 |
| 7286 | 4 | | | | | | | | | | 0 | 1147 |

| | | | | | | | | | | | |
|---|---|---|---|---|---|---|---|---|---|---|---|
| 7286 | 7 | | | | | | | | | 0 | 685 |
| 7272 | 6 | | | | | | | | | 0 | 723 |
| 7270 | 4 | | | | | | | | | 0 | 935 |
| 7270 | 2 | | | | | | | | | 0 | 937 |
| 7270 | 6 | | | | | | | | | 0 | 486 |
| 7269 | 7 | | | | | | | | | 0 | 732 |
| 7262 | 7 | | | | | | | | | 0 | 669 |
| 7250 | 2 | | | | | | | | | 0 | 558 |
| 7234 | 10 | | | | | | | | | 0 | 1065 |
| 7234 | 3 | | | | | | | | | 0 | 656 |
| 7221 | 4 | | | | | | | | | 0 | 818 |
| 7218 | 5 | | | | | | | | | 0 | 749 |
| 7218 | 8 | | | | | | | | | 0 | 1035 |
| 7217 | 5 | | | | | | | | | 0 | 921 |
| 7216 | 7 | | | | | | | | | 0 | 1108 |
| 7187 | 7 | | | | | | | | | 0 | 972 |
| 7170 | 7 | | | | | | | | | 0 | 744 |
| 7153 | 1 | metabolic process;tr | catalytic a | Pf04627 | 5564053 | AAEL0040 | AaeL_AAE | 1 | | 0 | 983 |
| 7121 | 8 | | | | | | | | | 0 | 619 |
| 7119 | 3 | | | | | | | | | 0 | 1160 |
| 7106 | 3 | metabolic process;tr | transporte | Pf05680 | 5572696 | AAEL0099 | AaeL_AAE | 2 | Oxidative phosphorylation; Me | 2 | 632 |
| 7099 | 9 | | | | | | | | | 0 | 1097 |
| 7098 | 8 | | | | | | | | | 0 | 1086 |
| 7088 | 5 | | | | | | | | | 0 | 1033 |
| 7075 | 2 | | | | | | | | | 0 | 1226 |
| 7062 | 5 | | | | | | | | | 0 | 735 |
| 7043 | 7 | | | | | | | | | 0 | 822 |
| 7037 | 2 | | | | | | | | | 0 | |
| 7016 | 3 | | | | | | | | | 0 | 643 |
| 7002 | 5 | | | | | | | | | 0 | 1043 |
| 7001 | 5 | | | | | | | | | 0 | 745 |

| | | | | | | | |
|---|---|---|---|---|---|---|---|
| 6997 | 3 | | | | | 0 | 627 |
| 6991 | 4 | | | | | 0 | 740 |
| 6988 | 6 | | | | | 0 | 482 |
| 6984 | 6 | | | | | 0 | 993 |
| 6982 | 6 | | | | | 0 | 494 |
| 6973 | 4 | | | | | 0 | 618 |
| 6972 | 4 | | | | | 0 | 884 |
| 6960 | 6 | | | | | 0 | 818 |
| 6947 | 5 | | | | | 0 | 689 |
| 6931 | 6 | | | | | 0 | 901 |
| 6925 | 7 | | | | | 0 | 856 |
| 6918 | 4 | | | | | 0 | 975 |
| 6901 | 5 | | | | | 0 | 803 |
| 6892 | 4 | | | | | 0 | 760 |
| 6887 | 6 | | | | | 0 | 722 |
| 6883 | 6 | | | | | 0 | 1390 |
| 6878 | 5 | | | | | 0 | 663 |
| 6867 | 2 | 5572523 | AAEL0098 | AaeL_AAE | 3 | 0 | 1019 |
| 6864 | 7 | | | | | 0 | 788 |
| 6847 | 8 | 5564443 | AAEL0042 | AaeL_AAE | 2 | 0 | 1186 |
| 6831 | 6 | | | | | 0 | 1046 |
| 6815 | 5 | | | | | 0 | 845 |
| 6791 | 4 | | | | | 0 | 991 |
| 6790 | 8 | | | | | 0 | 1011 |
| 6774 | 8 | | | | | 0 | 506 |
| 6768 | 8 | | | | | 0 | 972 |
| 6759 | 5 | | | | | 0 | 757 |
| 6757 | 3 | | | | | 0 | 917 |
| 6753 | 2 | | | | | 0 | 637 |
| 6752 | 7 | | | | | 0 | 1017 |
| 6751 | 8 | | | | | 0 | 911 |

| | | | | | | | | | | | |
|---|---|---|---|---|---|---|---|---|---|---|---|
| 6747 | 7 | | | | | | | | | 0 | 901 |
| 6744 | 4 | | | | | | | | | 0 | 773 |
| 6742 | 3 | | | | | | | | | 0 | 781 |
| 6738 | 3 | cell organ | membrane | Pf15454 | 5563671 | AAEL0005 | AaeL_AAE | 3 | mTOR signaling pathway | 1 | 739 |
| 6738 | 4 | | | | | | | | | 0 | 1246 |
| 6731 | 3 | | | | | | | | | 0 | 597 |
| 6705 | 6 | | | | | | | | | 0 | 1022 |
| 6698 | 5 | | | | | | | | | 0 | 706 |
| 6694 | 5 | | | | | | | | | 0 | 913 |
| 6682 | 4 | | | | | | | | | 0 | 726 |
| 6677 | 6 | | | | | | | | | 0 | 426 |
| 6662 | 4 | | | | | | | | | 0 | 647 |
| 6651 | 6 | | | | | | | | | 0 | 1145 |
| 6647 | 5 | | | | | | | | | 0 | 661 |
| 6643 | 6 | | | | | | | | | 0 | 795 |
| 6636 | 6 | | | | | | | | | 0 | 675 |
| 6628 | 5 | | | | | | | | | 0 | 894 |
| 6626 | 6 | | | | | | | | | 0 | 869 |
| 6612 | 4 | | | | | | | | | 0 | 509 |
| 6604 | 6 | | | | | | | | | 0 | 676 |
| 6593 | 4 | | | | | | | | | 0 | 865 |
| 6578 | 4 | | | | | | | | | 0 | 805 |
| 6556 | 6 | | | | | | | | | 0 | 993 |
| 6555 | 5 | | | | | | | | | 0 | 692 |
| 6550 | 4 | | | | | | | | | 0 | 998 |
| 6548 | 3 | | | | | | | | | 0 | 734 |
| 6541 | 7 | | | | | | | | | 0 | |
| 6539 | 7 | | | | | | | | | 0 | 860 |
| 6524 | 8 | | | | | | | | | 0 | 608 |
| 6516 | 1 | | | | | | | | | 0 | 1097 |
| 6516 | 7 | | | | | | | | | 0 | 912 |

| | | | | | | | | | |
|---|---|---|---|---|---|---|---|---|---|
| 6509 | 7 | | | | | | | 0 | 804 |
| 6485 | 4 | | | | | | | 0 | 727 |
| 6463 | 5 | | | | | | | 0 | 740 |
| 6451 | 5 | | | | | | | 0 | 574 |
| 6429 | 3 | | | | | | | 0 | 751 |
| 6415 | 8 | | | | | | | 0 | 790 |
| 6413 | 3 | | | | | | | 0 | 743 |
| 6395 | 7 | | | | | | | 0 | 925 |
| 6393 | 4 | | | | | | | 0 | 1204 |
| 6391 | 6 | | | | | | | 0 | 594 |
| 6382 | 5 | | | | | | | 0 | 660 |
| 6377 | 5 | | | | | | | 0 | 904 |
| 6375 | 4 | | | | | | | 0 | 915 |
| 6358 | 7 | | | | | | | 0 | 1129 |
| 6330 | 5 | | | | | | | 0 | 988 |
| 6320 | 3 | | | | | | | 0 | 808 |
| 6310 | 2 | | | | | | | 0 | 895 |
| 6303 | 6 | | | | | | | 0 | 815 |
| 6302 | 5 | | | | | | | 0 | 1170 |
| 6301 | 4 | | | | | | | 0 | 786 |
| 6297 | 5 | | | | | | | 0 | 502 |
| 6292 | 5 | | | | | | | 0 | 412 |
| 6289 | 4 | | | | | | | 0 | 1184 |
| 6284 | 8 | | | | | | | 0 | 854 |
| 6280 | 8 | | | | | | | 0 | 632 |
| 6266 | 8 | | | | | | | 0 | 840 |
| 6254 | 5 | | | | | | | 0 | 645 |
| 6251 | 8 | | | 5567249 | AAEL0008 | AaeL_AAE | 3 | 0 | 914 |
| 6246 | 8 | protein bi | Pf03234, I | 5572845 | AAEL0100 | AaeL_AAE | 3 | 0 | 719 |
| 6234 | 2 | | | | | | | 0 | 833 |
| 6211 | 6 | | | | | | | 0 | 516 |

| | | | | | | | | | |
|---|---|---|---|---|---|---|---|---|---|
| 6198 | 6 | | | | | | | 0 | 923 |
| 6194 | 2 | | | | | | | 0 | 801 |
| 6190 | 5 | | | | | | | 0 | 542 |
| 6187 | 5 | | | | | | | 0 | 963 |
| 6177 | 4 | | | | | | | 0 | 386 |
| 6171 | 4 | | | | | | | 0 | 1018 |
| 6171 | 4 | | | | | | | 0 | 1219 |
| 6166 | 6 | | | | | | | 0 | 789 |
| 6140 | 7 | metal ion | Pf00569, I | 5577342 | AAEL0004 | AaeL_AAE | 3 | 0 | 886 |
| 6128 | 6 | | | | | | | 0 | 738 |
| 6125 | 8 | | | | | | | 0 | 990 |
| 6121 | 3 | | | | | | | 0 | 869 |
| 6104 | 3 | | | | | | | 0 | 799 |
| 6103 | 4 | | | | | | | 0 | 1182 |
| 6100 | 5 | | | | | | | 0 | 835 |
| 6080 | 4 | | | | | | | 0 | 1472 |
| 6058 | 6 | | | | | | | 0 | 775 |
| 6056 | 4 | | | 5572533 | AAEL0098 | AaeL_AAE | 2 | 0 | 1015 |
| 6004 | 3 | | | | | | | 0 | 918 |
| 5985 | 5 | | | | | | | 0 | 764 |
| 5984 | 3 | | | | | | | 0 | 417 |
| 5980 | 5 | | | | | | | 0 | 782 |
| 5979 | 3 | | | | | | | 0 | 421 |
| 5975 | 3 | | | | | | | 0 | 767 |
| 5958 | 5 | | | | | | | 0 | 814 |
| 5956 | 3 | | | | | | | 0 | 634 |
| 5954 | 3 | | | | | | | 0 | 1045 |
| 5931 | 4 | | | | | | | 0 | 712 |
| 5920 | 7 | | | | | | | 0 | 803 |
| 5919 | 5 | | | | | | | 0 | 815 |
| 5915 | 6 | | | | | | | 0 | 725 |

| | | | | | | | | | | | | | | |
|---|---|---|---|---|---|---|---|---|---|---|---|---|---|---|
| 5908 | 5 | | | | | | | | | | | | 0 | 594 |
| 5901 | 3 | | | | | | | | | | | | 0 | 665 |
| 5898 | 6 | | | | | | | | | | | | 0 | 1253 |
| 5892 | 7 | | | | | | | | | | | | 0 | 1105 |
| 5887 | 7 | | | | | | | | | | | | 0 | 668 |
| 5873 | 4 | | membrane | | Pf03020, I | 5570540 | AAEL0014 | AaeL_AAE | 2 | | | | 0 | 1038 |
| 5867 | 3 | | | | | | | | | | | | 0 | 577 |
| 5857 | 4 | | | | | | | | | | | | 0 | 498 |
| 5837 | 6 | regulatior | nucleus | | Pf04153 | 5575721 | AAEL0120 | AaeL_AAE | 3 | | | | 0 | 1022 |
| 5835 | 4 | | | | | | | | | | | | 0 | 783 |
| 5810 | 6 | | | | | | | | | | | | 0 | 620 |
| 5808 | 4 | | | | | | | | | | | | 0 | 885 |
| 5806 | 5 | | | | | | | | | | | | 0 | 659 |
| 5802 | 3 | | | protein bi | Pf00498, I | 5574618 | AAEL0112 | AaeL_AAE | 1 | | | | 0 | 897 |
| 5797 | 4 | | | | | | | | | | | | 0 | 980 |
| 5797 | 4 | | | | | | | | | | | | 0 | 961 |
| 5795 | 7 | | | | | | | | | | | | 0 | 720 |
| 5794 | 5 | | | | | | | | | | | | 0 | 893 |
| 5784 | 6 | | | | | | | | | | | | 0 | 804 |
| 5780 | 5 | | | | | | | | | | | | 0 | 1047 |
| 5780 | 5 | | | | | | | | | | | | 0 | 464 |
| 5749 | 4 | | | | | | | | | | | | 0 | 975 |
| 5728 | 5 | | | | | | | | | | | | 0 | 1405 |
| 5728 | 4 | | | | | | | | | | | | 0 | 823 |
| 5724 | 4 | | | | | | | | | | | | 0 | 697 |
| 5720 | 5 | | | | | | | | | | | | 0 | 753 |
| 5704 | 5 | | | | | | | | | | | | 0 | 807 |
| 5703 | 7 | | | | | | | | | | | | 0 | 769 |
| 5697 | 4 | | | | | | | | | | | | 0 | 609 |
| 5681 | 5 | | | | | | | | | | | | 0 | 821 |
| 5680 | 4 | | | | | | | | | | | | 0 | 970 |

| | | | | | | | | | | | |
|---|---|---|---|---|---|---|---|---|---|---|---|
| 5651 | 5 | | | | | | | | | 0 | 1058 |
| 5651 | 5 | | | | | | | | | 0 | 904 |
| 5638 | 6 | | | | | | | | | 0 | 1060 |
| 5631 | 5 | | | | | | | | | 0 | 749 |
| 5618 | 4 | | | | | | | | | 0 | 764 |
| 5613 | 3 | | | | 5564989 | AAEL0045 | AaeL_AAE | 2 | | 0 | 974 |
| 5613 | 5 | | | | | | | | | 0 | 1215 |
| 5605 | 3 | | | | | | | | | 0 | 609 |
| 5597 | 5 | | | | | | | | | 0 | 1053 |
| 5596 | 5 | | | | | | | | | 0 | 958 |
| 5596 | 5 | | | | | | | | | 0 | 831 |
| 5586 | 8 | | | | | | | | | 0 | 848 |
| 5584 | 6 | | | | | | | | | 0 | 586 |
| 5563 | 2 | | | | | | | | | 0 | 828 |
| 5560 | 3 | | metal ion | Pf00076, I | 23687502 | AAEL0170 | AaeL_AAE | 2 | Spliceosome | 1 | 1141 |
| 5542 | 3 | | | | | | | | | 0 | 1508 |
| 5513 | 4 | | | | | | | | | 0 | 831 |
| 5498 | 5 | | | | | | | | | 0 | 1165 |
| 5477 | 4 | | | | | | | | | 0 | 903 |
| 5476 | 4 | | | | | | | | | 0 | 949 |
| 5470 | 3 | | | | | | | | | 0 | 624 |
| 5459 | 5 | | | Pf09774 | 5574060 | AAEL0109 | AaeL_AAE | 2 | | 0 | 819 |
| 5430 | 2 | membrane | | Pf10233 | 5567464 | AAEL0009 | AaeL_AAE | 3 | | 0 | 838 |
| 5418 | 3 | | | | | | | | | 0 | 828 |
| 5416 | 4 | | | | | | | | | 0 | 1114 |
| 5411 | 4 | | | | | | | | | 0 | 547 |
| 5405 | 2 | | | | | | | | | 0 | 212 |
| 5404 | 3 | | | | | | | | | 0 | 900 |
| 5399 | 5 | | | | | | | | | 0 | 1005 |
| 5395 | 2 | | | | | | | | | 0 | 828 |
| 5392 | 4 | | | | | | | | | 0 | 1311 |

| | | | | | | | | | | | | |
|---|---|---|---|---|---|---|---|---|---|---|---|---|
| 5391 | 5 | | | | | | | | | | 0 | 663 |
| 5383 | 2 | metabolic | process | catalytic a | Pf00089, I | 5568135 | AAEL0065 | AaeL_AAE | 2 | | 0 | 769 |
| 5373 | 4 | | | | | | | | | | 0 | 750 |
| 5370 | 5 | | | | | | | | | | 0 | 1235 |
| 5369 | 4 | | | | | | | | | | 0 | 488 |
| 5369 | 4 | | | | | | | | | | 0 | 806 |
| 5358 | 3 | | nucleus | | Pf00385, I | 5568287 | AAEL0067 | AaeL_AAE | 2 | | 0 | 1019 |
| 5354 | 2 | | | | | | | | | | 0 | 1781 |
| 5345 | 2 | | | | | | | | | | 0 | 920 |
| 5327 | 6 | | | | | | | | | | 0 | 784 |
| 5325 | 8 | metabolic | membran | catalytic a | Pf00246, I | 23687919 | AAEL0174 | AaeL_AAE | 3 | | 0 | 783 |
| 5322 | 5 | | | | | | | | | | 0 | 639 |
| 5312 | 2 | | | | | | | | | | 0 | 1442 |
| 5310 | 5 | | | | | | | | | | 0 | 1097 |
| 5306 | 3 | | | | Pf10151 | 5567454 | AAEL0009 | AaeL_AAE | 2 | | 0 | 530 |
| 5295 | 4 | | | | | | | | | | 0 | 791 |
| 5292 | 7 | | | | | | | | | | 0 | 730 |
| 5271 | 6 | | | | | | | | | | 0 | 1021 |
| 5254 | 5 | | | | | | | | | | 0 | 624 |
| 5253 | 4 | metabolic | membrane | | Pf05365 | 5570020 | AAEL0001 | AaeL_AAE | 3 | Metabolic pathways; Oxidative | 2 | 837 |
| 5246 | 4 | | | | | | | | | | 0 | 974 |
| 5241 | 4 | | | | | | | | | | 0 | 834 |
| 5241 | 5 | | | RNA bindi | Pf02926 | 5575750 | AAEL0027 | AaeL_AAE | 2 | | 0 | 744 |
| 5237 | 4 | | | | | | | | | | 0 | 1120 |
| 5201 | 5 | | | | | | | | | | 0 | 989 |
| 5200 | 7 | | | | | | | | | | 0 | 899 |
| 5199 | 4 | | | | | | | | | | 0 | 904 |
| 5199 | 3 | | | | | | | | | | 0 | 863 |
| 5185 | 6 | | | | | | | | | | 0 | 955 |
| 5184 | 2 | | | | | | | | | | 0 | |
| 5184 | 4 | | | nucleotid | Pf00176, I | 5567894 | AAEL0181 | AaeL_AAE | 3 | | 0 | 705 |

| | | | | | | | | | | | |
|---|---|---|---|---|---|---|---|---|---|---|---|
| 5163 | 4 | | | | | | | | | 0 | 648 |
| 5161 | 4 | | | | | | | | | 0 | 434 |
| 5160 | 5 | | | | | | | | | 0 | 929 |
| 5159 | 3 | cell organ | chromoso | DNA binding | | 5580037 | AAEL0102 | AaeL_AAE | 2 | 0 | 1101 |
| 5157 | 4 | | | | | | | | | 0 | 1374 |
| 5156 | 2 | | | | | | | | | 0 | 848 |
| 5144 | 4 | | | | | | | | | 0 | 636 |
| 5142 | 5 | | | | | | | | | 0 | 305 |
| 5135 | 5 | | | | | | | | | 0 | 1007 |
| 5134 | 1 | | | | | | | | | 0 | 930 |
| 5131 | 1 | | | | | | | | | 0 | 1103 |
| 5123 | 4 | | | | | | | | | 0 | 720 |
| 5118 | 4 | | | | | | | | | 0 | 1081 |
| 5116 | 4 | | | | | | | | | 0 | 689 |
| 5107 | 8 | | | | | | | | | 0 | 575 |
| 5089 | 6 | | | | | | | | | 0 | 1093 |
| 5080 | 4 | | | | | | | | | 0 | 794 |
| 5063 | 3 | | | | | | | | | 0 | 862 |
| 5059 | 4 | | | | | | | | | 0 | 672 |
| 5052 | 5 | | | | | | | | | 0 | 1005 |
| 5043 | 3 | | | | Pf07051 | 5571267 | AAEL0015 | AaeL_AAE | 2 | 0 | 782 |
| 5039 | 5 | | | | | | | | | 0 | 842 |
| 5037 | 3 | | | | | | | | | 0 | 775 |
| 5035 | 4 | | | | | | | | | 0 | 768 |
| 5029 | 5 | | | | | | | | | 0 | 879 |
| 5021 | 6 | | | | | | | | | 0 | 540 |
| 5018 | 3 | | | | | | | | | 0 | 885 |
| 5009 | 3 | | | | | | | | | 0 | 329 |
| 4997 | 5 | | | | | | | | | 0 | 758 |
| 4992 | 4 | | | | | | | | | 0 | 489 |
| 4958 | 3 | | | | | | | | | 0 | 969 |

| | | | | | | | | | | | | |
|---|---|---|---|---|---|---|---|---|---|---|---|---|
| 4953 | 1 | | | | | | | | | | 0 | 1058 |
| 4939 | 4 | | | | | | | | | | 0 | 1026 |
| 4938 | 3 | | | | | | | | | | 0 | 686 |
| 4938 | 4 | | | | | | | | | | 0 | 1025 |
| 4922 | 5 | | | | | | | | | | 0 | 836 |
| 4909 | 7 | | | | | | | | | | 0 | 615 |
| 4905 | 8 | | | | | | | | | | 0 | 1128 |
| 4899 | 2 | | | | | | | | | | 0 | 1222 |
| 4898 | 4 | | | | | | | | | | 0 | 696 |
| 4889 | 2 | | | | | | | | | | 0 | 904 |
| 4885 | 1 | | | | | | | | | | 0 | 829 |
| 4880 | 5 | | | protein bi | Pf00400, I | 5566139 | AAEL0052 | AaeL_AAE | 2 | RNA degradation | 1 | 903 |
| 4875 | 3 | | | | | | | | | | 0 | 848 |
| 4874 | 6 | | | | | | | | | | 0 | 817 |
| 4869 | 5 | | | | | | | | | | 0 | 812 |
| 4867 | 2 | | | | | | | | | | 0 | 993 |
| 4865 | 5 | | | | | | | | | | 0 | 770 |
| 4860 | 4 | | | | | | | | | | 0 | 808 |
| 4835 | 4 | | | | | | | | | | 0 | 963 |
| 4832 | 4 | | | | | | | | | | 0 | 735 |
| 4826 | 5 | metabolic process | | catalytic a | Pf01026 | 5566073 | AAEL0150 | AaeL_AAE | 2 | | 0 | 980 |
| 4819 | 4 | | | | | | | | | | 0 | |
| 4817 | 5 | | | | | | | | | | 0 | 902 |
| 4794 | 4 | | | | | | | | | | 0 | 816 |
| 4772 | 5 | | | | | | | | | | 0 | 521 |
| 4770 | 8 | | | | | | | | | | 0 | 395 |
| 4766 | 4 | | | | | 5578357 | AAEL0035 | AaeL_AAE | 2 | | 0 | 1010 |
| 4756 | 4 | | endoplasr | protein bi | Pf06400, I | 5573866 | AAEL0107 | AaeL_AAE | 1 | | 0 | 1018 |
| 4750 | 5 | | | | | | | | | | 0 | 915 |
| 4745 | 4 | | | | | | | | | | 0 | 721 |
| 4739 | 6 | | | | | | | | | | 0 | 877 |

| | | | | | | | | | |
|---|---|---|---|---|---|---|---|---|---|
| 4736 | 2 | | | | | | | 0 | 854 |
| 4733 | 5 | | | | | | | 0 | 910 |
| 4722 | 3 | | | | | | | 0 | 622 |
| 4720 | 4 | | | | | | | 0 | 778 |
| 4713 | 3 | | | | | | | 0 | 954 |
| 4712 | 2 | | | | | | | 0 | 754 |
| 4709 | 4 | | | | | | | 0 | 732 |
| 4709 | 6 | | | | | | | 0 | 828 |
| 4708 | 5 | | | | | | | 0 | 986 |
| 4707 | 4 | | | | | | | 0 | 1033 |
| 4707 | 7 | | | | | | | 0 | 583 |
| 4693 | 3 | | | | | | | 0 | 981 |
| 4690 | 4 | | | | | | | 0 | 807 |
| 4680 | 5 | | | | | | | 0 | 592 |
| 4679 | 6 | | | | | | | 0 | 617 |
| 4673 | 5 | | | | | | | 0 | 1030 |
| 4671 | 5 | | | | | | | 0 | 736 |
| 4670 | 6 | | | | | | | 0 | 1133 |
| 4668 | 5 | | | | | | | 0 | 729 |
| 4640 | 3 | | | | | | | 0 | 833 |
| 4635 | 4 | | | | | | | 0 | 539 |
| 4622 | 5 | | | | | | | 0 | 792 |
| 4621 | 3 | | | | | | | 0 | 1000 |
| 4621 | 3 | | | | | | | 0 | 828 |
| 4619 | 4 | | | | | | | 0 | 896 |
| 4607 | 3 | | | | | | | 0 | 689 |
| 4606 | 3 | | | | | | | 0 | 1130 |
| 4603 | 6 | | | | | | | 0 | 894 |
| 4589 | 5 | protein bi | Pf00400 | 5564513 | AAEL0183 | AaeL_AAE | 2 | 0 | 817 |
| 4583 | 5 | | | | | | | 0 | 846 |
| 4583 | 3 | | | | | | | 0 | 924 |

| | | | | | | | | | | |
|---|---|---|---|---|---|---|---|---|---|---|
| 4581 | 4 | | | | | | | | 0 | 811 |
| 4547 | 5 | transport | catalytic a | Pf01496, I | 5576703 | AAEL0117 | AaeL_AAE | 3 | 0 | 994 |
| 4539 | 6 | | | | | | | | 0 | 673 |
| 4528 | 5 | | | | | | | | 0 | 1169 |
| 4518 | 4 | | | | | | | | 0 | 736 |
| 4517 | 3 | | | | | | | | 0 | 1071 |
| 4509 | 2 | | | | | | | | 0 | 655 |
| 4507 | 4 | | | | | | | | 0 | 722 |
| 4499 | 3 | | | | | | | | 0 | 1219 |
| 4495 | 3 | | | | | | | | 0 | 940 |
| 4483 | 3 | | | | | | | | 0 | 1184 |
| 4464 | 4 | metabolic process | catalytic a | Pf00069, I | 5570866 | AAEL0181 | AaeL_AAE | 2 | 0 | 712 |
| 4460 | 4 | | | | | | | | 0 | 846 |
| 4455 | 1 | metabolic process | | Pf08511 | 5572051 | AAEL0002 | AaeL_AAE | 2 | 0 | 659 |
| 4453 | 2 | | | | | | | | 0 | 831 |
| 4451 | 3 | | | | | | | | 0 | 1184 |
| 4437 | 3 | | | | | | | | 0 | 966 |
| 4429 | 5 | | | | | | | | 0 | 817 |
| 4419 | 1 | | | | | | | | 0 | 779 |
| 4411 | 5 | | | | | | | | 0 | 1007 |
| 4405 | 2 | | | | | | | | 0 | 1451 |
| 4399 | 5 | | | | | | | | 0 | 1294 |
| 4394 | 6 | | | | | | | | 0 | 947 |
| 4393 | 3 | | | | | | | | 0 | 1277 |
| 4379 | 3 | | | | | | | | 0 | 800 |
| 4375 | 5 | | | | | | | | 0 | 908 |
| 4374 | 2 | | | | | | | | 0 | 836 |
| 4374 | 4 | | | | | | | | 0 | 815 |
| 4374 | 6 | | | | | | | | 0 | 779 |
| 4372 | 4 | | | | | | | | 0 | 1191 |
| 4370 | 4 | | | | 5565410 | AAEL0148 | AaeL_AAE | 2 | 0 | 918 |

| | | | | | | | | | | | |
|---|---|---|---|---|---|---|---|---|---|---|---|
| 4368 | 4 | | protein bi | Pf01485, I | 5576092 | AAEL0117 | AaeL_AAE | 3 | | 0 | 718 |
| 4359 | 4 | | | | | | | | | 0 | |
| 4356 | 3 | | | | | | | | | 0 | 521 |
| 4353 | 5 | | | | | | | | | 0 | 1072 |
| 4332 | 4 | | | | | | | | | 0 | 1081 |
| 4329 | 4 | | | | | | | | | 0 | 388 |
| 4327 | 4 | nucleus | | Pf09468 | 5564021 | AAEL0142 | AaeL_AAE | 2 | DNA replication | 1 | 617 |
| 4320 | 4 | | | | | | | | | 0 | 716 |
| 4312 | 4 | | | | | | | | | 0 | 715 |
| 4308 | 3 | | | | | | | | | 0 | 454 |
| 4307 | 3 | | | | | | | | | 0 | 854 |
| 4306 | 4 | | | | | | | | | 0 | 721 |
| 4304 | 5 | | | | | | | | | 0 | 1098 |
| 4297 | 5 | | | | | | | | | 0 | 571 |
| 4291 | 4 | | | | | | | | | 0 | 792 |
| 4289 | 2 | | | Pf01966, I | 5574363 | AAEL0110 | AaeL_AAE | 2 | | 0 | 996 |
| 4287 | 1 | | | | | | | | | 0 | 369 |
| 4280 | 3 | | | | | | | | | 0 | 974 |
| 4273 | 4 | | metal ion | Pf12171, I | 5577997 | AAEL0033 | AaeL_AAE | 3 | | 0 | 937 |
| 4262 | 5 | | | | | | | | | 0 | 803 |
| 4255 | 2 | | | | | | | | | 0 | 391 |
| 4249 | 3 | | | | | | | | | 0 | 347 |
| 4249 | 2 | | | | | | | | | 0 | 514 |
| 4247 | 5 | | | | | | | | | 0 | 682 |
| 4237 | 6 | | | | | | | | | 0 | 984 |
| 4219 | 4 | | | | | | | | | 0 | 1038 |
| 4217 | 3 | | | | | | | | | 0 | 863 |
| 4209 | 4 | | | | | | | | | 0 | 740 |
| 4204 | 6 | | | | | | | | | 0 | |
| 4199 | 5 | | | | | | | | | 0 | 1005 |
| 4199 | 3 | | | | | | | | | 0 | 745 |

| | | | | | | | | | |
|---|---|---|---|---|---|---|---|---|---|
| 4194 | 4 | | | | | | | 0 | 774 |
| 4190 | 3 | | | | | | | 0 | 406 |
| 4189 | 4 | | | | | | | 0 | 629 |
| 4171 | 5 | | | | | | | 0 | 1018 |
| 4167 | 3 | | | | | | | 0 | |
| 4160 | 3 | | | | | | | 0 | 971 |
| 4155 | 4 | | | | | | | 0 | 963 |
| 4155 | 3 | | | | | | | 0 | 873 |
| 4150 | 3 | | | | | | | 0 | 649 |
| 4147 | 3 | | | | | | | 0 | 605 |
| 4140 | 3 | | | | | | | 0 | 1129 |
| 4139 | 7 | | | | | | | 0 | 929 |
| 4136 | 2 | | | 5573649 | AAEL0106 | AaeL_AAE | 2 | 0 | 1052 |
| 4135 | 3 | | Pf10224 | 5564552 | AAEL0043 | AaeL_AAE | 3 | 0 | 1128 |
| 4129 | 2 | | | | | | | 0 | |
| 4127 | 3 | | | | | | | 0 | 1311 |
| 4119 | 3 | | | | | | | 0 | 1076 |
| 4118 | 8 | | | | | | | 0 | 476 |
| 4105 | 4 | | | | | | | 0 | 772 |
| 4103 | 2 | | | | | | | 0 | 784 |
| 4096 | 4 | | | | | | | 0 | 706 |
| 4095 | 5 | | | | | | | 0 | 949 |
| 4089 | 2 | | | | | | | 0 | 769 |
| 4085 | 2 | membran | catalytic activity | 5578921 | AAEL0037 | AaeL_AAE | 1 | 0 | 1008 |
| 4084 | 3 | | | | | | | 0 | 592 |
| 4080 | 7 | | | | | | | 0 | 488 |
| 4080 | 4 | | | | | | | 0 | 570 |
| 4079 | 6 | | | | | | | 0 | 921 |
| 4078 | 4 | | | | | | | 0 | 625 |
| 4075 | 2 | | | 5570344 | AAEL0013 | AaeL_AAE | 2 | 0 | 905 |
| 4065 | 1 | | | | | | | 0 | 549 |

| | | | | | | | | | | | |
|---|---|---|---|---|---|---|---|---|---|---|---|
| 4065 | 4 | | | | | | | | | 0 | |
| 4063 | 5 | | | | | | | | | 0 | 1069 |
| 4056 | 5 | | | | | | | | | 0 | 1165 |
| 4051 | 3 | | | | | | | | | 0 | 1101 |
| 4050 | 4 | | | | | | | | | 0 | 831 |
| 4040 | 4 | | | Pf04969 | 5565702 | AAEL0149 | AaeL_AAE | 3 | Arachidonic acid metabolism; M | 2 | 1013 |
| 4040 | 6 | | | | | | | | | 0 | 876 |
| 4038 | 5 | | | | | | | | | 0 | 851 |
| 4036 | 4 | | | | | | | | | 0 | 1046 |
| 4028 | 5 | metabolic process | catalytic a | Pf01121, I | 5578935 | AAEL0037 | AaeL_AAE | 1 | Metabolic pathways; Pantother | 2 | 701 |
| 4027 | 5 | | | | | | | | | 0 | 767 |
| 4024 | 3 | | | | | | | | | 0 | 677 |
| 4021 | 4 | | | | | | | | | 0 | 870 |
| 4021 | 5 | | | | | | | | | 0 | 849 |
| 4019 | 4 | | | | | | | | | 0 | 855 |
| 4018 | 4 | | | | | | | | | 0 | 944 |
| 4017 | 4 | | | | | | | | | 0 | 851 |
| 4014 | 4 | | | | | | | | | 0 | 792 |
| 4004 | 2 | metabolic membran | catalytic a | Pf02937 | 5571735 | AAEL0092 | AaeL_AAE | 2 | Metabolic pathways; Oxidative | 2 | 613 |
| 4003 | 3 | | | | | | | | | 0 | 786 |
| 3995 | 5 | | | | | | | | | 0 | 726 |
| 3995 | 2 | | | | | | | | | 0 | 784 |
| 3995 | 4 | | | | | | | | | 0 | |
| 3984 | 4 | | | | | | | | | 0 | 1137 |
| 3968 | 2 | | | | | | | | | 0 | 1012 |
| 3962 | 4 | | | | | | | | | 0 | 1087 |
| 3961 | 5 | | | | | | | | | 0 | 498 |
| 3958 | 5 | | | | | | | | | 0 | 927 |
| 3957 | 4 | | | | | | | | | 0 | 1025 |
| 3943 | 5 | | | | | | | | | 0 | 756 |
| 3933 | 3 | | | | | | | | | 0 | |

| | | | | | | | | | | | | | | |
|---|---|---|---|---|---|---|---|---|---|---|---|---|---|---|
| 3930 | 3 | | | | | | | | | | | | 0 | 525 |
| 3926 | 4 | | | | | | | | | | | | 0 | 787 |
| 3917 | 4 | | | | | | | | | | | | 0 | 1014 |
| 3914 | 4 | | | | | | | | | | | | 0 | 587 |
| 3913 | 3 | | | | | | | | | | | | 0 | 646 |
| 3907 | 3 | | | | | | | | | | | | 0 | 761 |
| 3899 | 5 | | | | | | | | | | | | 0 | 953 |
| 3895 | 2 | | | | | | | | | | | | 0 | 985 |
| 3895 | 2 | | | | | | | | | | | | 0 | 1026 |
| 3888 | 2 | regulatior | cytoplasm | | | 23687739 | AAEL0173 | AaeL_AAE | | 3 | | | 0 | 1092 |
| 3883 | 1 | | | | | | | | | | | | 0 | 1132 |
| 3873 | 2 | | | | | | | | | | | | 0 | 846 |
| 3869 | 4 | | | | | | | | | | | | 0 | 620 |
| 3863 | 4 | | | | | | | | | | | | 0 | 1144 |
| 3861 | 5 | | | | | | | | | | | | 0 | 976 |
| 3860 | 4 | | | | | | | | | | | | 0 | 461 |
| 3858 | 3 | | | | | | | | | | | | 0 | 787 |
| 3854 | 4 | | | | | | | | | | | | 0 | 814 |
| 3845 | 4 | | | | | | | | | | | | 0 | 969 |
| 3838 | 3 | | | | | | | | | | | | 0 | 523 |
| 3838 | 3 | | | | | | | | | | | | 0 | 438 |
| 3831 | 3 | | membrane | | Pf05644 | 5575775 | AAEL0027 | AaeL_AAE | | 3 | | | 0 | 743 |
| 3828 | 3 | | | | | | | | | | | | 0 | 793 |
| 3827 | 2 | | | | | | | | | | | | 0 | 831 |
| 3827 | 5 | | | | | | | | | | | | 0 | 1075 |
| 3824 | 3 | | | | | | | | | | | | 0 | 918 |
| 3822 | 4 | | | | | | | | | | | | 0 | 312 |
| 3821 | 2 | | | | | | | | | | | | 0 | 915 |
| 3819 | 1 | | | | | | | | | | | | 0 | 618 |
| 3819 | 2 | | | | | | | | | | | | 0 | 1070 |
| 3815 | 3 | | | | | | | | | | | | 0 | 1063 |

| | | | | | | | | | | | |
|---|---|---|---|---|---|---|---|---|---|---|---|
| 3814 | 6 | | | | | | | | | 0 | 661 |
| 3809 | 5 | | | | | | | | | 0 | 1122 |
| 3801 | 3 | | | | | | | | | 0 | 822 |
| 3800 | 3 | | | | | | | | | 0 | 556 |
| 3799 | 2 | | | | | | | | | 0 | 1006 |
| 3798 | 4 | | | | | | | | | 0 | 669 |
| 3792 | 2 | | | | | | | | | 0 | 934 |
| 3789 | 4 | regulation of biologi | | enzyme re | Pf09184 | 5568555 | AAEL0011 | AaeL_AAE | 3 | 0 | 710 |
| 3784 | 2 | | | | | | | | | 0 | 995 |
| 3779 | 4 | | | | | | | | | 0 | 1146 |
| 3776 | 6 | | | | | | | | | 0 | 818 |
| 3764 | 4 | | | | | | | | | 0 | 922 |
| 3758 | 2 | | | | | | | | | 0 | 748 |
| 3755 | 2 | | | | | | | | | 0 | 692 |
| 3747 | 3 | | | | | | | | | 0 | 1277 |
| 3745 | 5 | | | | | | | | | 0 | 499 |
| 3738 | 3 | | | | | | | | | 0 | 518 |
| 3731 | 5 | | | | | | | | | 0 | 972 |
| 3724 | 2 | | | | | | | | | 0 | 803 |
| 3724 | 3 | | | | Pf13345 | 5567390 | AAEL0060 | AaeL_AAE | 3 | 0 | 1021 |
| 3716 | 4 | | | | | | | | | 0 | 849 |
| 3715 | 2 | | | | | | | | | 0 | 1638 |
| 3708 | 3 | | | | | | | | | 0 | 1071 |
| 3707 | 3 | | | | | | | | | 0 | 685 |
| 3702 | 4 | | | | | | | | | 0 | 841 |
| 3700 | 3 | | nucleus | metal ion | Pf06524 | 5564027 | AAEL0142 | AaeL_AAE | 2 | 0 | 760 |
| 3700 | 3 | | | RNA bindi | Pf00076, I | 5570204 | AAEL0081 | AaeL_AAE | 3 | 0 | 1376 |
| 3699 | 3 | | | | | | | | | 0 | 586 |
| 3698 | 4 | | | | | | | | | 0 | 987 |
| 3691 | 2 | | | | | | | | | 0 | |
| 3689 | 3 | | | | | | | | | 0 | 928 |

| | | | | | | | | | |
|---|---|---|---|---|---|---|---|---|---|
| 3684 | 5 | | | | | | | 0 | 833 |
| 3678 | 5 | | | | | | | 0 | 633 |
| 3672 | 2 | | Pf15328 | 5580159 | AAEL0113 | AaeL_AAE | 3 | 0 | 914 |
| 3661 | 4 | | | | | | | 0 | 997 |
| 3648 | 4 | | | | | | | 0 | 804 |
| 3646 | 2 | | | | | | | 0 | 629 |
| 3639 | 4 | | | | | | | 0 | 507 |
| 3638 | 4 | | | | | | | 0 | 705 |
| 3634 | 2 | | | | | | | 0 | 1493 |
| 3624 | 5 | | | | | | | 0 | 943 |
| 3624 | 5 | | | | | | | 0 | 832 |
| 3621 | 4 | | | 5578589; | AAEL0137 | AaeL_AAE | 2 | 0 | 406 |
| 3617 | 4 | | | | | | | 0 | 451 |
| 3611 | 3 | | | | | | | 0 | 752 |
| 3605 | 3 | | | | | | | 0 | 621 |
| 3601 | 4 | | | | | | | 0 | 897 |
| 3600 | 3 | regulation of biological process | | 5564793 | AAEL0044 | AaeL_AAE | 3 | 0 | 1136 |
| 3598 | 5 | | | | | | | 0 | 855 |
| 3593 | 3 | | | | | | | 0 | 1374 |
| 3590 | 3 | | | | | | | 0 | 455 |
| 3590 | 4 | | | | | | | 0 | 1256 |
| 3589 | 4 | | | | | | | 0 | 563 |
| 3589 | 1 | | | | | | | 0 | 1261 |
| 3578 | 2 | | | | | | | 0 | 668 |
| 3578 | 3 | | | | | | | 0 | 520 |
| 3578 | 3 | | | | | | | 0 | 647 |
| 3578 | 3 | | | | | | | 0 | 501 |
| 3574 | 2 | | | | | | | 0 | 1418 |
| 3574 | 6 | | | | | | | 0 | 453 |
| 3573 | 4 | | | | | | | 0 | 433 |
| 3571 | 3 | | | | | | | 0 | 731 |

| | | | | | | | | |
|---|---|---|---|---|---|---|---|---|
| 3568 | 3 | | | | | | 0 | 738 |
| 3563 | 4 | | | | | | 0 | 884 |
| 3561 | 3 | | | | | | 0 | 1191 |
| 3559 | 3 | | | | | | 0 | 904 |
| 3554 | 1 | | | | | | 0 | 546 |
| 3554 | 3 | Pf05890 | 5571317 | AAEL0089 | AaeL_AAE | 1 | 0 | 960 |
| 3552 | 2 | | | | | | 0 | 846 |
| 3552 | 5 | | | | | | 0 | 852 |
| 3551 | 4 | | | | | | 0 | 808 |
| 3545 | 3 | | | | | | 0 | 1115 |
| 3545 | 4 | | | | | | 0 | 487 |
| 3545 | 3 | | | | | | 0 | 1253 |
| 3533 | 2 | | | | | | 0 | 772 |
| 3530 | 6 | | | | | | 0 | 681 |
| 3526 | 2 | | | | | | 0 | 1274 |
| 3524 | 3 | | | | | | 0 | 876 |
| 3522 | 3 | | | | | | 0 | |
| 3519 | 3 | | | | | | 0 | 758 |
| 3514 | 4 | | | | | | 0 | 931 |
| 3510 | 3 | | | | | | 0 | 784 |
| 3503 | 4 | | | | | | 0 | 601 |
| 3503 | 3 | | | | | | 0 | 270 |
| 3501 | 3 | | | | | | 0 | 489 |
| 3499 | 3 | | | | | | 0 | 601 |
| 3498 | 2 | | | | | | 0 | 930 |
| 3496 | 4 | | | | | | 0 | 618 |
| 3493 | 5 | | | | | | 0 | 930 |
| 3490 | 3 | | | | | | 0 | 893 |
| 3490 | 5 | | | | | | 0 | |
| 3484 | 3 | | | | | | 0 | 605 |
| 3475 | 3 | | 23687805 | AAEL0173 | AaeL_AAE | 3 | 0 | 895 |

| | | | | | | | | | | | | |
|---|---|---|---|---|---|---|---|---|---|---|---|---|
| 3464 | 3 | | | | | | | | | | 0 | 1111 |
| 3459 | 4 | | | | | | | | | | 0 | 847 |
| 3458 | 4 | | | | | | | | | | 0 | 601 |
| 3458 | 4 | | | | | | | | | | 0 | 358 |
| 3455 | 3 | | | | | | | | | | 0 | 918 |
| 3453 | 4 | | | | | | | | | | 0 | 553 |
| 3444 | 5 | regulation | mitochon | protein binding | | 5564707 | AAEL0145 | AaeL_AAE | 2 | | 0 | 638 |
| 3441 | 4 | | | | | | | | | | 0 | 565 |
| 3439 | 4 | | | | | | | | | | 0 | 728 |
| 3438 | 5 | | | | | | | | | | 0 | 520 |
| 3437 | 2 | | | | Pf02893, I | 5575321; | AAEL0025 | AaeL_AAE | 3 | | 0 | 774 |
| 3432 | 2 | | | | | | | | | | 0 | 701 |
| 3425 | 3 | | | | | | | | | | 0 | 504 |
| 3424 | 4 | | | | | | | | | | 0 | 735 |
| 3421 | 3 | | | | | | | | | | 0 | 1161 |
| 3418 | 2 | metabolic process | | RNA bindi | Pf03853, I | 5570555 | AAEL0083 | AaeL_AAE | 3 | | 0 | 756 |
| 3410 | 4 | | | | | | | | | | 0 | 739 |
| 3407 | 5 | | | | | | | | | | 0 | 711 |
| 3406 | 2 | | | | | | | | | | 0 | 785 |
| 3405 | 2 | cell organization and biogenesi | | | Pf01798, I | 23687963 | AAEL0175 | AaeL_AAE | 3 | Spliceosome | 1 | 1322 |
| 3405 | 3 | | | | | | | | | | 0 | 949 |
| 3399 | 4 | | | | | | | | | | 0 | 782 |
| 3398 | 4 | | | | | | | | | | 0 | 1253 |
| 3386 | 5 | | | | | | | | | | 0 | 798 |
| 3380 | 2 | | | | | | | | | | 0 | 615 |
| 3376 | 2 | | | | | | | | | | 0 | 1114 |
| 3364 | 3 | | | | | | | | | | 0 | |
| 3360 | 3 | | | | | | | | | | 0 | 840 |
| 3352 | 4 | | | | | | | | | | 0 | 694 |
| 3350 | 5 | | | | | | | | | | 0 | 636 |
| 3344 | 3 | | | | | | | | | | 0 | 810 |

| | | | | | | | | | | | |
|---|---|---|---|---|---|---|---|---|---|---|---|
| 3337 | 4 | | | | | | | | | 0 | 871 |
| 3330 | 3 | | | | 23687805 | AAEL0173 | AaeL_AAE | 3 | | 0 | |
| 3320 | 2 | | | | | | | | | 0 | 469 |
| 3320 | 1 | | | | | | | | | 0 | 717 |
| 3320 | 4 | | | | | | | | | 0 | 1220 |
| 3320 | 2 | | | | | | | | | 0 | 847 |
| 3316 | 5 | | | | | | | | | 0 | 862 |
| 3315 | 4 | | | | | | | | | 0 | 1156 |
| 3312 | 2 | | | | | | | | | 0 | 877 |
| 3307 | 3 | | | | | | | | | 0 | 327 |
| 3305 | 4 | | | | | | | | | 0 | 1017 |
| 3303 | 4 | | | | | | | | | 0 | 1021 |
| 3289 | 5 | | | | | | | | | 0 | 570 |
| 3288 | 3 | | | | | | | | | 0 | 823 |
| 3288 | 3 | | | | | | | | | 0 | 555 |
| 3269 | 5 | | | | | | | | | 0 | 1078 |
| 3261 | 2 | | | | | | | | | 0 | 568 |
| 3260 | 2 | | | | | | | | | 0 | 861 |
| 3259 | 1 | | | | | | | | | 0 | 1066 |
| 3259 | 3 | | | | | | | | | 0 | 1040 |
| 3256 | 4 | | | | | | | | | 0 | 768 |
| 3255 | 3 | | | | | | | | | 0 | 588 |
| 3252 | 1 | | | | | | | | | 0 | 1166 |
| 3249 | 4 | | | | | | | | | 0 | 400 |
| 3246 | 2 | | | | | | | | | 0 | 682 |
| 3245 | 4 | | | | | | | | | 0 | 409 |
| 3243 | 4 | cell organization and | metal ion | Pf00382, I | 5566397 | AAEL0053 | AaeL_AAE | 2 | Basal transcription factors | 1 | 1037 |
| 3241 | 6 | | | | | | | | | 0 | 866 |
| 3239 | 3 | metabolic process | RNA bindi | Pf04135 | 5569166 | AAEL0074 | AaeL_AAE | 2 | Ribosome biogenesis in eukary | 1 | 842 |
| 3230 | 3 | | | | | | | | | 0 | 721 |
| 3229 | 3 | | | | | | | | | 0 | 760 |

| | | | |
|---|---|---|---|
| 3227 | 4 | 0 | 526 |
| 3225 | 4 | 0 | 694 |
| 3203 | 4 | 0 | 808 |
| 3199 | 4 | 0 | 882 |
| 3197 | 4 | 0 | 579 |
| 3197 | 4 | 0 | 819 |
| 3193 | 4 | 0 | 594 |
| 3189 | 2 | 0 | 887 |
| 3189 | 3 | 0 | 686 |
| 3186 | 4 | 0 | 869 |
| 3184 | 3 | 0 | 745 |
| 3180 | 3 | 0 | 1184 |
| 3175 | 3 | 0 | 776 |
| 3172 | 2 | 0 | 989 |
| 3172 | 3 | 0 | 729 |
| 3171 | 3 | 0 | 656 |
| 3168 | 1 | 0 | 722 |
| 3167 | 6 | 0 | 979 |
| 3163 | 3 | 0 | 1259 |
| 3157 | 2 | 0 | 724 |
| 3155 | 3 | 0 | |
| 3154 | 5 | 0 | |
| 3137 | 3 | 0 | 1032 |
| 3128 | 3 | 0 | 742 |
| 3124 | 3 | 0 | 725 |
| 3122 | 5 | 0 | 576 |
| 3120 | 3 | 0 | 680 |
| 3117 | 2 | 0 | 1383 |
| 3112 | 2 | 0 | 989 |
| 3108 | 2 | 0 | |
| 3104 | 4 | 0 | 1052 |

| | | | |
|---|---|---|---|
| 3103 | 2 | 0 | 1113 |
| 3097 | 3 | 0 | 1495 |
| 3096 | 3 | 0 | 652 |
| 3086 | 4 | 0 | 1117 |
| 3085 | 2 | 0 | 842 |
| 3084 | 1 | 0 | 683 |
| 3084 | 5 | 0 | 723 |
| 3082 | 4 | 0 | 816 |
| 3076 | 3 | 0 | 975 |
| 3075 | 4 | 0 | 697 |
| 3071 | 2 | 0 | 1020 |
| 3064 | 4 | 0 | 473 |
| 3064 | 1 | 0 | 739 |
| 3045 | 6 | 0 | 1066 |
| 3041 | 2 | 0 | 570 |
| 3040 | 2 | 0 | 1343 |
| 3040 | 2 | 0 | 1553 |
| 3039 | 4 | 0 | 832 |
| 3035 | 3 | 0 | 1125 |
| 3028 | 4 | 0 | 803 |
| 3024 | 3 | 0 | 916 |
| 3015 | 3 | 0 | 863 |
| 3014 | 2 | 0 | 1053 |
| 3013 | 2 | 0 | 849 |
| 3010 | 4 | 0 | 919 |
| 3007 | 4 | 0 | 987 |
| 3002 | 4 | 0 | 606 |
| 3000 | 2 | 0 | |
| 2982 | 1 | 0 | 442 |
| 2979 | 3 | 0 | 868 |
| 2973 | 2 | 0 | 907 |

| | | | | | | | | | | | |
|---|---|---|---|---|---|---|---|---|---|---|---|
| 2971 | 3 | | | | | | | | | 0 | 833 |
| 2971 | 3 | | | | | | | | | 0 | 1070 |
| 2964 | 3 | | | | Pf07304 | 5566038 | AAEL0150 | AaeL_AAE | 1 | 0 | 986 |
| 2960 | 4 | | | | | | | | | 0 | 940 |
| 2951 | 3 | | | | | | | | | 0 | 1018 |
| 2937 | 4 | | | | | | | | | 0 | 909 |
| 2931 | 2 | | | | | | | | | 0 | 972 |
| 2929 | 1 | | | | | | | | | 0 | 838 |
| 2929 | 4 | | | | | | | | | 0 | 979 |
| 2924 | 4 | | | | | | | | | 0 | 863 |
| 2920 | 1 | | | | | | | | | 0 | 1283 |
| 2913 | 2 | | | | | 5573777 | AAEL0002 | AaeL_AAE | 2 | 0 | |
| 2911 | 2 | | | | | | | | | 0 | 846 |
| 2907 | 1 | | | | | | | | | 0 | 1093 |
| 2906 | 4 | | | | | | | | | 0 | 1317 |
| 2904 | 3 | | | | | | | | | 0 | 574 |
| 2903 | 3 | | | | | | | | | 0 | 727 |
| 2903 | 2 | | | | | | | | | 0 | 794 |
| 2902 | 4 | | | | | | | | | 0 | 549 |
| 2900 | 3 | | | | | | | | | 0 | |
| 2896 | 3 | | membrane | | Pf04241 | 5570063 | AAEL0013 | AaeL_AAE | 3 | 0 | 625 |
| 2896 | 1 | | | | | 5566110 | AAEL0051 | AaeL_AAE | 2 | 0 | 529 |
| 2892 | 2 | | | | | | | | | 0 | 849 |
| 2889 | 2 | | | | | | | | | 0 | 780 |
| 2887 | 3 | | | | | | | | | 0 | 635 |
| 2886 | 1 | | | | | 23687632 | AAEL0172 | AaeL_AAE | 2 | 0 | 806 |
| 2878 | 4 | | extracellu | protein bi | Pf07174, I | 5565494 | AAEL0048 | AaeL_AAE | 3 | 0 | 866 |
| 2872 | 2 | regulation of biological process | | | Pf07818 | 5570868 | AAEL0086 | AaeL_AAE | 2 | 0 | 967 |
| 2869 | 2 | | | | | | | | | 0 | 803 |
| 2866 | 2 | | | | | | | | | 0 | 940 |
| 2865 | 5 | | | | | | | | | 0 | 798 |

| | | | | | | | | | | |
|---|---|---|---|---|---|---|---|---|---|---|
| 2860 | 3 | | | | | | | | 0 | 971 |
| 2859 | 2 | | | | | | | | 0 | 1009 |
| 2852 | 2 | | | | | | | | 0 | 748 |
| 2851 | 4 | | | | | | | | 0 | 834 |
| 2849 | 4 | | | | | | | | 0 | 772 |
| 2844 | 5 | | | | | | | | 0 | 967 |
| 2840 | 2 | | | | | | | | 0 | 446 |
| 2839 | 3 | | | | | | | | 0 | 625 |
| 2829 | 3 | | | | | | | | 0 | 1001 |
| 2823 | 2 | | | | | | | | 0 | 641 |
| 2818 | 3 | | | | | | | | 0 | 969 |
| 2818 | 2 | | | | | | | | 0 | 552 |
| 2815 | 2 | | | | | | | | 0 | 978 |
| 2814 | 3 | | | | | | | | 0 | |
| 2806 | 5 | | | | | | | | 0 | 1103 |
| 2806 | 1 | | | | | | | | 0 | 1083 |
| 2800 | 3 | | | | | | | | 0 | 977 |
| 2798 | 3 | | | | | | | | 0 | 1036 |
| 2796 | 3 | | | | | | | | 0 | |
| 2795 | 4 | | | | | | | | 0 | 931 |
| 2792 | 3 | | | | | | | | 0 | 934 |
| 2782 | 3 | | | | | | | | 0 | 783 |
| 2775 | 2 | | | | | | | | 0 | 704 |
| 2763 | 3 | | | | | | | | 0 | 1131 |
| 2758 | 1 | | | | | | | | 0 | 801 |
| 2755 | 2 | | | | | | | | 0 | 529 |
| 2754 | 3 | | | | | | | | 0 | |
| 2754 | 3 | | | | | | | | 0 | 1075 |
| 2752 | 6 | | | | | | | | 0 | 757 |
| 2748 | 2 | transport | RNA bindi | Pf00076, I | 5571315 | AAEL0089 | AaeL_AAE | 1 | 0 | 920 |
| 2745 | 2 | | | | | | | | 0 | 1242 |

| | | | | | | | | | | | |
|---|---|---|---|---|---|---|---|---|---|---|---|
| 2745 | 3 | | | | | | | | | 0 | 1302 |
| 2744 | 1 | | | | | | | | | 0 | 666 |
| 2744 | 1 | | | | | | | | | 0 | 926 |
| 2743 | 2 | | | | | | | | | 0 | 661 |
| 2743 | 3 | | | | | | | | | 0 | 957 |
| 2742 | 2 | | | | | | | | | 0 | 1056 |
| 2727 | 2 | | | | | | | | | 0 | 748 |
| 2724 | 4 | | | | | | | | | 0 | 746 |
| 2722 | 4 | | | | | | | | | 0 | |
| 2721 | 2 | metabolic process | | Pf04078 | 5568456 | AAEL0068 | AaeL_AAE | 3 | RNA degradation | 1 | 833 |
| 2714 | 3 | | | | | | | | | 0 | |
| 2712 | 2 | | | | | | | | | 0 | 758 |
| 2711 | 3 | | | | | | | | | 0 | 701 |
| 2703 | 2 | | | | | | | | | 0 | 651 |
| 2701 | 3 | | | | | | | | | 0 | 790 |
| 2700 | 4 | | | | | | | | | 0 | 585 |
| 2699 | 4 | | | | | | | | | 0 | 751 |
| 2691 | 4 | | | | | | | | | 0 | 557 |
| 2690 | 4 | | | | | | | | | 0 | 979 |
| 2686 | 2 | | | | | | | | | 0 | 903 |
| 2685 | 3 | | | | | | | | | 0 | |
| 2685 | 3 | | | | | | | | | 0 | 850 |
| 2672 | 3 | | | | | | | | | 0 | 943 |
| 2671 | 2 | | | | | | | | | 0 | 869 |
| 2659 | 2 | | | | | | | | | 0 | 1075 |
| 2657 | 4 | | | | | | | | | 0 | 1067 |
| 2649 | 3 | | | | | | | | | 0 | |
| 2648 | 4 | | | | | | | | | 0 | 1003 |
| 2643 | 4 | regulation of biologi | protein bi | Pf00514, I | 5566538 | AAEL0183 | AaeL_AAE | 3 | | 0 | 840 |
| 2640 | 4 | | | | | | | | | 0 | 844 |
| 2634 | 5 | | | | | | | | | 0 | 881 |

| | | | | | | | | | | | |
|---|---|---|---|---|---|---|---|---|---|---|---|
| 2634 | 2 | | | | | | | | | 0 | 402 |
| 2630 | 2 | | | | | | | | | 0 | 734 |
| 2627 | 2 | | | | | | | | | 0 | 447 |
| 2623 | 3 | | | | | | | | | 0 | |
| 2619 | 2 | | | | | | | | | 0 | 976 |
| 2618 | 2 | | | | | | | | | 0 | 808 |
| 2615 | 2 | | | | | | | | | 0 | 702 |
| 2614 | 4 | | | | | | | | | 0 | 998 |
| 2612 | 3 | | | | | | | | | 0 | 755 |
| 2604 | 3 | cell organization and | RNA bindi | Pf00472 | 5573252 | AAEL0103 | AaeL_AAE | 2 | | 0 | 845 |
| 2603 | 3 | | | | | | | | | 0 | 863 |
| 2594 | 3 | | | | | | | | | 0 | |
| 2592 | 4 | | | Pf00567 | 5573221 | AAEL0103 | AaeL_AAE | 3 | | 0 | 1074 |
| 2586 | 3 | | | | | | | | | 0 | 827 |
| 2583 | 2 | | | | | | | | | 0 | 748 |
| 2582 | 3 | | | | | | | | | 0 | 1568 |
| 2579 | 2 | | | | | | | | | 0 | |
| 2577 | 2 | | | | | | | | | 0 | 528 |
| 2574 | 4 | | | | | | | | | 0 | 673 |
| 2570 | 1 | | | | | | | | | 0 | 199 |
| 2568 | 2 | | | | | | | | | 0 | 735 |
| 2568 | 3 | | | | | | | | | 0 | 626 |
| 2567 | 4 | | | | | | | | | 0 | |
| 2564 | 3 | | | | | | | | | 0 | 820 |
| 2561 | 2 | | | | | | | | | 0 | |
| 2560 | 1 | | | | | | | | | 0 | 690 |
| 2551 | 3 | | | Pf05182 | 5570138 | AAEL0081 | AaeL_AAE | 3 | mRNA surveillance pathway | 1 | 1052 |
| 2550 | 3 | | | | | | | | | 0 | 900 |
| 2547 | 2 | | | | | | | | | 0 | 641 |
| 2543 | 4 | | | | | | | | | 0 | 1031 |
| 2541 | 3 | | | | | | | | | 0 | 712 |

| | | | | | | | | | | | |
|---|---|---|---|---|---|---|---|---|---|---|---|
| 2534 | 4 | | | | | | | | | 0 | 1008 |
| 2526 | 2 | | | | | | | | | 0 | 862 |
| 2524 | 1 | | | | | | | | | 0 | 811 |
| 2522 | 2 | | | | | | | | | 0 | 825 |
| 2520 | 4 | | | | | | | | | 0 | |
| 2519 | 3 | | | | | | | | | 0 | 908 |
| 2517 | 2 | | | | | | | | | 0 | 788 |
| 2507 | 3 | | | | | | | | | 0 | 594 |
| 2505 | 2 | | | | | | | | | 0 | 691 |
| 2503 | 4 | | | | | | | | | 0 | 766 |
| 2500 | 2 | | | | | | | | | 0 | 1099 |
| 2493 | 3 | | | | | | | | | 0 | 764 |
| 2491 | 4 | | | | | | | | | 0 | 1063 |
| 2475 | 2 | | | | | | | | | 0 | 492 |
| 2475 | 1 | | | | | | | | | 0 | 732 |
| 2472 | 3 | | | | | | | | | 0 | 612 |
| 2471 | 2 | | | | | | | | | 0 | 809 |
| 2469 | 1 | | | | | | | | | 0 | 803 |
| 2468 | 3 | | | | | | | | | 0 | 959 |
| 2447 | 2 | | | | | | | | | 0 | 877 |
| 2443 | 3 | | | | | 5568302 | AAEL0067 | AaeL_AAE | 2 | 0 | 1037 |
| 2442 | 1 | | | | | | | | | 0 | 514 |
| 2442 | 3 | | | | | | | | | 0 | 634 |
| 2435 | 3 | | | | | | | | | 0 | 977 |
| 2431 | 2 | | | | | | | | | 0 | 649 |
| 2429 | 3 | | | | | | | | | 0 | 638 |
| 2428 | 3 | | | | | | | | | 0 | 1477 |
| 2427 | 4 | | | | | | | | | 0 | 876 |
| 2424 | 2 | | | | | | | | | 0 | 841 |
| 2423 | 3 | | | metal ion binding | | 5569505 | AAEL0076 | AaeL_AAE | 2 | 0 | |
| 2422 | 3 | regulatior | extracellu | enzyme re | Pf00207, I | 23687443 | AAEL0170 | AaeL_AAE | 2 | 0 | 1190 |

| | | | | | | | | | |
|---|---|---|---|---|---|---|---|---|---|
| 2422 | 5 | | | | | | | 0 | |
| 2420 | 3 | | | | | | | 0 | |
| 2420 | 4 | | | | | | | 0 | 955 |
| 2413 | 2 | | | | | | | 0 | 918 |
| 2412 | 4 | | | | | | | 0 | 972 |
| 2402 | 2 | | | | | | | 0 | 500 |
| 2401 | 3 | | | | | | | 0 | |
| 2396 | 1 | | | | | | | 0 | 1292 |
| 2394 | 2 | | | | | | | 0 | 833 |
| 2392 | 2 | | | | | | | 0 | |
| 2390 | 1 | membrane | Pf12832 | 5577237 | AAEL0031 | AaeL_AAE | 3 | 0 | 540 |
| 2387 | 1 | | | | | | | 0 | 869 |
| 2385 | 3 | | | | | | | 0 | 1150 |
| 2384 | 3 | | | | | | | 0 | 519 |
| 2375 | 2 | | | | | | | 0 | 779 |
| 2375 | 1 | | | | | | | 0 | 1053 |
| 2375 | 1 | | | | | | | 0 | 1009 |
| 2370 | 3 | | | | | | | 0 | 657 |
| 2360 | 2 | | | | | | | 0 | 384 |
| 2360 | 3 | | | | | | | 0 | 916 |
| 2359 | 3 | | | | | | | 0 | 755 |
| 2356 | 4 | | | | | | | 0 | |
| 2345 | 2 | | | | | | | 0 | 931 |
| 2339 | 2 | | | | | | | 0 | |
| 2331 | 1 | | | | | | | 0 | 814 |
| 2328 | 2 | | | | | | | 0 | 844 |
| 2326 | 1 | | | | | | | 0 | 778 |
| 2323 | 2 | | | | | | | 0 | 812 |
| 2314 | 1 | | | | | | | 0 | 826 |
| 2308 | 2 | | | | | | | 0 | 1017 |
| 2308 | 1 | | Pf09420 | 5563791 | AAEL0039 | AaeL_AAE | 3 | 0 | 995 |

| | | | | | | | | | | |
|---|---|---|---|---|---|---|---|---|---|---|
| 2304 | 2 | | | | | | | | 0 | 858 |
| 2302 | 3 | | | | | | | | 0 | 602 |
| 2296 | 4 | | | | | | | | 0 | 655 |
| 2295 | 1 | | | | | | | | 0 | 1477 |
| 2291 | 3 | | | | | | | | 0 | 955 |
| 2290 | 3 | | | | | | | | 0 | 842 |
| 2285 | 1 | | | | | | | | 0 | |
| 2274 | 2 | | | | | | | | 0 | 920 |
| 2273 | 2 | | | | | | | | 0 | 613 |
| 2271 | 1 | | | | | | | | 0 | 1133 |
| 2259 | 4 | | | | | | | | 0 | 759 |
| 2258 | 2 | | | | | | | | 0 | 806 |
| 2258 | 2 | | | | | | | | 0 | 1024 |
| 2252 | 3 | | | | | | | | 0 | |
| 2244 | 3 | | | | | | | | 0 | 368 |
| 2238 | 4 | | | | | | | | 0 | 720 |
| 2232 | 1 | | | | | | | | 0 | 937 |
| 2217 | 2 | | | | | | | | 0 | 605 |
| 2212 | 3 | | | | | | | | 0 | 969 |
| 2209 | 2 | | | | | | | | 0 | 951 |
| 2208 | 4 | | | | | | | | 0 | |
| 2204 | 1 | | | | | | | | 0 | 989 |
| 2204 | 3 | | | | | | | | 0 | 965 |
| 2202 | 2 | metabolic extracellular | Pf01607 | 5573718 | AAEL0107 | AaeL_AAE | 3 | | 0 | 832 |
| 2199 | 1 | | | | | | | | 0 | 746 |
| 2199 | 2 | | | | | | | | 0 | 829 |
| 2196 | 2 | | | | | | | | 0 | 983 |
| 2192 | 1 | | | | | | | | 0 | 1096 |
| 2186 | 3 | | | | | | | | 0 | 623 |
| 2186 | 1 | | | | | | | | 0 | 553 |
| 2181 | 2 | | | | | | | | 0 | 787 |

| | | | | | | | | | | | | |
|---|---|---|---|---|---|---|---|---|---|---|---|---|
| 2170 | 2 | | | | | | | | | | 0 | |
| 2162 | 2 | | | | | | | | | | 0 | 780 |
| 2162 | 1 | | | | | | | | | | 0 | 502 |
| 2159 | 2 | | | | | | | | | | 0 | 909 |
| 2158 | 2 | | | | | | | | | | 0 | 856 |
| 2153 | 2 | | | | | | | | | | 0 | 754 |
| 2147 | 1 | | | | | | | | | | 0 | 1232 |
| 2146 | 2 | | | | | | | | | | 0 | 569 |
| 2146 | 3 | | | | | | | | | | 0 | 644 |
| 2145 | 2 | | | | | | | | | | 0 | |
| 2143 | 1 | | | | | | | | | | 0 | 609 |
| 2140 | 2 | | | | | | | | | | 0 | 1028 |
| 2139 | 3 | | | | | | | | | | 0 | 772 |
| 2131 | 2 | | | | | | | | | | 0 | 613 |
| 2128 | 3 | | | | | | | | | | 0 | 707 |
| 2124 | 2 | | | | | | | | | | 0 | 969 |
| 2124 | 2 | | | | | | | | | | 0 | 591 |
| 2121 | 2 | | | | | 5578108 | AAEL0034 | AaeL_AAE | 3 | | 0 | |
| 2119 | 2 | | | | | | | | | | 0 | 779 |
| 2116 | 3 | | | | | | | | | | 0 | 693 |
| 2112 | 2 | | | | | | | | | | 0 | 1111 |
| 2111 | 3 | metabolic | membran | catalytic a | Pf00128 | 5565454 | AAEL0006 | AaeL_AAE | 3 | Starch and sucrose metabolism | 3 | 208 |
| 2108 | 2 | | | | | | | | | | 0 | 880 |
| 2107 | 2 | | | | Pf10245 | 5575752 | AAEL0027 | AaeL_AAE | 2 | | 0 | 766 |
| 2106 | 3 | | | | | | | | | | 0 | 879 |
| 2101 | 2 | | | | | | | | | | 0 | 675 |
| 2101 | 3 | | | | | | | | | | 0 | 1074 |
| 2099 | 2 | | | | | 5578080 | AAEL0034 | AaeL_AAE | 3 | | 0 | 991 |
| 2099 | 2 | | | | | | | | | | 0 | 845 |
| 2099 | 3 | | | | | | | | | | 0 | 856 |
| 2097 | 1 | | mitochondrion | | | 5576103 | AAEL0027 | AaeL_AAE | 3 | | 0 | 995 |

| | | | | | | | | | | | |
|---|---|---|---|---|---|---|---|---|---|---|---|
| 2096 | 3 | cell organ | nucleus | DNA bindi | Pf00505 | 5574067 | AAEL0109 | AaeL_AAE | 2 | | 0 | 1097 |
| 2094 | 4 | | | | | | | | | | 0 | 511 |
| 2093 | 2 | | nucleus | | | 5568056 | AAEL0180 | AaeL_AAE | 3 | | 0 | 927 |
| 2092 | 1 | | | | | | | | | | 0 | 1234 |
| 2086 | 2 | | | | | | | | | | 0 | 590 |
| 2084 | 2 | | | | | | | | | | 0 | 941 |
| 2081 | 3 | | | | | | | | | | 0 | 948 |
| 2075 | 3 | cell organization and biogenesi | | | Pf09724 | 5571009 | AAEL0087 | AaeL_AAE | 3 | | 0 | |
| 2075 | 2 | | | | | | | | | | 0 | 850 |
| 2070 | 3 | | | | | | | | | | 0 | 494 |
| 2065 | 4 | | | | | | | | | | 0 | 733 |
| 2061 | 1 | | membrane | | Pf13903 | 5566176 | AAEL0052 | AaeL_AAE | 3 | | 0 | 430 |
| 2060 | 4 | | | | | | | | | | 0 | 970 |
| 2057 | 1 | | | | | | | | | | 0 | 904 |
| 2052 | 2 | | | | | | | | | | 0 | 940 |
| 2051 | 1 | | | | | | | | | | 0 | 1012 |
| 2051 | 3 | | | | | | | | | | 0 | 1007 |
| 2049 | 1 | | | | | | | | | | 0 | 885 |
| 2046 | 1 | | | | | | | | | | 0 | 1151 |
| 2041 | 1 | regulation of biologi | | nucleotid | Pf04670, I | 5566141 | AAEL0051 | AaeL_AAE | 2 | mTOR signaling pathway | 1 | 503 |
| 2040 | 2 | | | | | | | | | | 0 | |
| 2038 | 1 | | | | | | | | | | 0 | 955 |
| 2037 | 4 | | | | | | | | | | 0 | 686 |
| 2033 | 2 | | | | | | | | | | 0 | 646 |
| 2031 | 1 | | | | | | | | | | 0 | 635 |
| 2030 | 2 | | | | | | | | | | 0 | 758 |
| 2026 | 3 | | | | Pf01928 | 5574528 | AAEL0112 | AaeL_AAE | 1 | | 0 | |
| 2022 | 2 | | | | | | | | | | 0 | 1254 |
| 2021 | 1 | | | | | | | | | | 0 | 644 |
| 2019 | 3 | | | | | | | | | | 0 | 921 |
| 2006 | 1 | | | | | | | | | | 0 | 738 |

| | | | | | | | | | | | |
|---|---|---|---|---|---|---|---|---|---|---|---|
| 2005 | 1 | | | | | | | | | 0 | 1003 |
| 2001 | 3 | | | | | | | | | 0 | 842 |
| 1998 | 2 | | | | | | | | | 0 | 792 |
| 1996 | 3 | | | | | | | | | 0 | 723 |
| 1990 | 2 | | | | | | | | | 0 | 581 |
| 1986 | 2 | | | | | | | | | 0 | 1044 |
| 1985 | 1 | | | | | | | | | 0 | 1270 |
| 1981 | 3 | | | | | | | | | 0 | 839 |
| 1979 | 2 | | | | | | | | | 0 | 1288 |
| 1976 | 2 | | | | | | | | | 0 | |
| 1974 | 2 | | | | | | | | | 0 | 629 |
| 1972 | 1 | | | | | | | | | 0 | 916 |
| 1969 | 2 | | | | | | | | | 0 | |
| 1968 | 3 | | | | | | | | | 0 | 269 |
| 1967 | 2 | | | | | | | | | 0 | 516 |
| 1954 | 3 | | | | | | | | | 0 | 998 |
| 1952 | 3 | cell organization and biogenesi | | Pf01172, I | 5568190 | AAEL0066 | AaeL_AAE | 2 | Ribosome biogenesis in eukary | 1 | 898 |
| 1947 | 3 | | | | | | | | | 0 | |
| 1941 | 3 | | | | | | | | | 0 | 933 |
| 1938 | 1 | | | | | | | | | 0 | |
| 1937 | 1 | | | | | | | | | 0 | 991 |
| 1937 | 2 | metabolic process;re | catalytic a | Pf02891, I | 5570516 | AAEL0014 | AaeL_AAE | 2 | | 0 | 833 |
| 1935 | 1 | | | | | | | | | 0 | 922 |
| 1935 | 1 | | | | | | | | | 0 | 983 |
| 1934 | 2 | | | | | | | | | 0 | 678 |
| 1934 | 3 | | | | | | | | | 0 | |
| 1933 | 2 | | | | | | | | | 0 | 784 |
| 1930 | 2 | | | | | | | | | 0 | 722 |
| 1925 | 2 | | | | | | | | | 0 | 867 |
| 1924 | 1 | | | | | | | | | 0 | 320 |
| 1924 | 3 | | | | | | | | | 0 | 336 |

| | | | | | | | | |
|---|---|---|---|---|---|---|---|---|
| 1922 | 3 | | | | | | 0 | 381 |
| 1920 | 3 | | | | | | 0 | |
| 1920 | 2 | | | | | | 0 | 865 |
| 1914 | 3 | | | | | | 0 | 496 |
| 1913 | 1 | | | | | | 0 | |
| 1912 | 2 | | | | | | 0 | 570 |
| 1910 | 3 | | | | | | 0 | 704 |
| 1908 | 2 | | | | | | 0 | 1035 |
| 1904 | 2 | | | | | | 0 | 663 |
| 1903 | 2 | | | | | | 0 | |
| 1903 | 2 | | | | | | 0 | 414 |
| 1900 | 3 | | | | | | 0 | 812 |
| 1895 | 2 | | | | | | 0 | 742 |
| 1893 | 2 | Pf05254 | 5572371 | AAEL0097 | AaeL_AAE | 1 | 0 | 1027 |
| 1890 | 2 | | | | | | 0 | |
| 1890 | 4 | | | | | | 0 | 727 |
| 1887 | 3 | | | | | | 0 | 1310 |
| 1879 | 1 | | | | | | 0 | 767 |
| 1876 | 3 | | | | | | 0 | 941 |
| 1869 | 1 | | | | | | 0 | 1082 |
| 1869 | 2 | | | | | | 0 | |
| 1866 | 2 | | | | | | 0 | 785 |
| 1864 | 2 | | | | | | 0 | |
| 1862 | 1 | | | | | | 0 | 1130 |
| 1861 | 3 | | | | | | 0 | 823 |
| 1858 | 3 | | | | | | 0 | 726 |
| 1857 | 3 | | | | | | 0 | 761 |
| 1856 | 3 | | | | | | 0 | 1200 |
| 1851 | 3 | | | | | | 0 | 794 |
| 1849 | 1 | | | | | | 0 | 1229 |
| 1848 | 1 | | | | | | 0 | 595 |

| | | | | | | | | | | |
|---|---|---|---|---|---|---|---|---|---|---|
| 1847 | 2 | | | | | | | | 0 | 723 |
| 1844 | 2 | | | | | | | | 0 | 849 |
| 1841 | 3 | | | | | | | | 0 | 973 |
| 1837 | 3 | | | | | | | | 0 | 830 |
| 1834 | 4 | | | | | | | | 0 | 640 |
| 1832 | 2 | | | | | | | | 0 | 899 |
| 1830 | 4 | | | | | | | | 0 | 568 |
| 1829 | 2 | | | | | | | | 0 | 521 |
| 1828 | 4 | | | | | | | | 0 | 1103 |
| 1826 | 3 | regulation of biological process | Pf08923 | 5564062 | AAEL0040 | AaeL_AAE | 1 | mTOR signaling pathway | 1 | 989 |
| 1823 | 3 | | | | | | | | 0 | 857 |
| 1822 | 2 | | | | | | | | 0 | 793 |
| 1821 | 1 | protein bi | Pf00397 | 5575136 | AAEL0003 | AaeL_AAE | 2 | | 0 | 1196 |
| 1815 | 3 | | | | | | | | 0 | 765 |
| 1810 | 2 | | | | | | | | 0 | 435 |
| 1808 | 2 | | | | | | | | 0 | 793 |
| 1806 | 2 | | | | | | | | 0 | 977 |
| 1805 | 2 | | | | | | | | 0 | 903 |
| 1804 | 4 | | | | | | | | 0 | 932 |
| 1801 | 2 | | | | | | | | 0 | 808 |
| 1800 | 3 | | | | | | | | 0 | |
| 1800 | 3 | | | | | | | | 0 | 997 |
| 1800 | 4 | | | | | | | | 0 | 729 |
| 1798 | 3 | | | | | | | | 0 | 827 |
| 1794 | 3 | | | | | | | | 0 | 959 |
| 1793 | 2 | | | | | | | | 0 | 467 |
| 1792 | 2 | | | | | | | | 0 | 515 |
| 1791 | 1 | | | | | | | | 0 | 828 |
| 1791 | 3 | | | | | | | | 0 | 827 |
| 1789 | 2 | | | | | | | | 0 | 746 |
| 1786 | 1 | | | | | | | | 0 | 1061 |

| | | | | | | | |
|---|---|---|---|---|---|---|---|
| 1779 | 1 | | | | | 0 | 646 |
| 1778 | 2 | | | | | 0 | 529 |
| 1778 | 1 | | | | | 0 | 769 |
| 1777 | 2 | | | | | 0 | 680 |
| 1776 | 3 | | | | | 0 | 698 |
| 1774 | 1 | | | | | 0 | |
| 1764 | 1 | | | | | 0 | 860 |
| 1761 | 2 | | | | | 0 | 818 |
| 1758 | 2 | | | | | 0 | |
| 1755 | 2 | | | | | 0 | 572 |
| 1753 | 4 | | | | | 0 | 794 |
| 1752 | 3 | | | | | 0 | 829 |
| 1748 | 1 | | | | | 0 | 970 |
| 1742 | 2 | | | | | 0 | 616 |
| 1740 | 2 | | | | | 0 | 1123 |
| 1735 | 1 | | | | | 0 | 498 |
| 1732 | 1 | | | | | 0 | 792 |
| 1730 | 1 | | | | | 0 | |
| 1730 | 2 | | | | | 0 | |
| 1727 | 3 | | | | | 0 | 821 |
| 1724 | 2 | | | | | 0 | 445 |
| 1723 | 3 | | | | | 0 | 911 |
| 1717 | 1 | | | | | 0 | 1669 |
| 1717 | 2 | | | | | 0 | 327 |
| 1716 | 1 | | | | | 0 | 519 |
| 1714 | 2 | | | | | 0 | 958 |
| 1712 | 3 | | | | | 0 | 968 |
| 1711 | 3 | 5571097 | AAEL0015 | AaeL_AAE | 3 | 0 | 682 |
| 1709 | 1 | | | | | 0 | 1139 |
| 1709 | 2 | | | | | 0 | 205 |
| 1706 | 2 | | | | | 0 | |

| | | | | | | | | | | | |
|---|---|---|---|---|---|---|---|---|---|---|---|
| 1705 | 2 | | | | | | | | | 0 | |
| 1705 | 2 | | | | | | | | | 0 | 1233 |
| 1702 | 2 | | | | | | | | | 0 | 750 |
| 1702 | 3 | | | | | | | | | 0 | 733 |
| 1700 | 3 | | | | | | | | | 0 | 1005 |
| 1698 | 1 | | | | | | | | | 0 | 981 |
| 1693 | 2 | | | | | | | | | 0 | 1122 |
| 1691 | 3 | | | | | | | | | 0 | 849 |
| 1691 | 1 | cell organization and biogenesi | | Pf06784, I | 5566140 | AAEL0051 | AaeL_AAE | 2 | | 0 | 585 |
| 1689 | 2 | | | | | | | | | 0 | 1111 |
| 1689 | 1 | | | Pf01636 | 5563704 | AAEL0039 | AaeL_AAE | 1 | | 0 | |
| 1688 | 2 | | | | | | | | | 0 | 1220 |
| 1686 | 3 | | | | | | | | | 0 | 1009 |
| 1683 | 1 | | | | | | | | | 0 | 1171 |
| 1682 | 1 | | | | | | | | | 0 | 1022 |
| 1671 | 1 | | | | | | | | | 0 | 417 |
| 1671 | 1 | | | | | | | | | 0 | |
| 1670 | 2 | | | | | | | | | 0 | 837 |
| 1670 | 2 | | | | | | | | | 0 | |
| 1670 | 2 | | | | | | | | | 0 | 1090 |
| 1668 | 3 | regulation of biologi | protein bi | Pf13646, I | 5568448 | AAEL0068 | AaeL_AAE | 3 | | 0 | 651 |
| 1666 | 3 | | | | | | | | | 0 | |
| 1665 | 1 | metabolic process | catalytic a | Pf03577 | 5570882 | AAEL0086 | AaeL_AAE | 3 | | 0 | 1043 |
| 1664 | 4 | | | Pf14799 | 5580260 | AAEL0125 | AaeL_AAE | 2 | | 0 | |
| 1663 | 3 | | | Pf07159 | 5568120 | AAEL0181 | AaeL_AAE | 2 | | 0 | 801 |
| 1662 | 2 | | | | | | | | | 0 | 781 |
| 1662 | 2 | | | Pf01138, I | 5568404 | AAEL0068 | AaeL_AAE | 1 | RNA degradation | 1 | 968 |
| 1661 | 2 | | | | | | | | | 0 | 329 |
| 1661 | 2 | | | | | | | | | 0 | 1006 |
| 1660 | 1 | | | | | | | | | 0 | 1172 |
| 1656 | 2 | | | | | | | | | 0 | 712 |

| | | | | | | | | | | | | |
|---|---|---|---|---|---|---|---|---|---|---|---|---|
| 1654 | 1 | | | | | | | | | | 0 | 777 |
| 1652 | 2 | | | | | | | | | | 0 | 994 |
| 1652 | 2 | | membrane | | | 5564916 | AAEL0146 | AaeL_AAE | 1 | | 0 | 1108 |
| 1648 | 2 | | | | | | | | | | 0 | 768 |
| 1644 | 1 | | | | | 23687817 | AAEL0173 | AaeL_AAE | 2 | | 0 | |
| 1644 | 2 | | | | | | | | | | 0 | 949 |
| 1640 | 1 | | | | | | | | | | 0 | 429 |
| 1633 | 2 | | | | | 5565660 | AAEL0048 | AaeL_AAE | 2 | | 0 | 968 |
| 1632 | 2 | | | | | | | | | | 0 | |
| 1632 | 2 | | | | | | | | | | 0 | 612 |
| 1631 | 4 | | | | | | | | | | 0 | 709 |
| 1628 | 2 | | | | | | | | | | 0 | 327 |
| 1627 | 3 | | | | | | | | | | 0 | 679 |
| 1626 | 2 | regulation of biologi | | protein bi | Pf13646, I | 5569506 | AAEL0076 | AaeL_AAE | 2 | Longevity regulating pathway - | 2 | |
| 1623 | 1 | | | | | | | | | | 0 | 897 |
| 1620 | 2 | | | | | | | | | | 0 | 749 |
| 1619 | 1 | | | protein bi | Pf00627 | 5575805 | AAEL0120 | AaeL_AAE | 3 | | 0 | 795 |
| 1618 | 2 | | | | | | | | | | 0 | 367 |
| 1615 | 2 | | | | | | | | | | 0 | |
| 1614 | 2 | | | | | | | | | | 0 | 962 |
| 1613 | 2 | | | | | | | | | | 0 | 775 |
| 1610 | 2 | | | | | | | | | | 0 | 1022 |
| 1608 | 2 | | | | | | | | | | 0 | |
| 1606 | 3 | | | | | | | | | | 0 | 890 |
| 1606 | 2 | | | | | | | | | | 0 | 720 |
| 1600 | 1 | | | | | | | | | | 0 | 805 |
| 1599 | 1 | | | | | | | | | | 0 | |
| 1598 | 3 | | | | | | | | | | 0 | 827 |
| 1596 | 2 | | | | | | | | | | 0 | 1457 |
| 1593 | 3 | | | | | | | | | | 0 | 818 |
| 1591 | 1 | | | | | | | | | | 0 | 659 |

| | | | | | | | | | | |
|---|---|---|---|---|---|---|---|---|---|---|
| 1588 | 2 | | | | | | | | 0 | |
| 1586 | 2 | | | | | | | | 0 | 528 |
| 1584 | 1 | | | Pf10217 | 5574139 | AAEL0109 | AaeL_AAE | 3 | 0 | 969 |
| 1584 | 1 | | | | | | | | 0 | 872 |
| 1578 | 3 | | | Pf04969 | 5576878 | AAEL0030 | AaeL_AAE | 1 | 0 | |
| 1578 | 1 | | | | | | | | 0 | 1080 |
| 1575 | 1 | | | | | | | | 0 | |
| 1571 | 1 | | | | | | | | 0 | 674 |
| 1571 | 3 | | | | | | | | 0 | |
| 1571 | 3 | | | | | | | | 0 | |
| 1569 | 2 | | | | | | | | 0 | 807 |
| 1564 | 2 | membrane | | Pf13517 | 5576016 | AAEL0122 | AaeL_AAE | 3 | 0 | |
| 1563 | 2 | | | | | | | | 0 | 845 |
| 1562 | 2 | | | | | | | | 0 | |
| 1562 | 2 | | | | | | | | 0 | 932 |
| 1562 | 2 | | | | | | | | 0 | 849 |
| 1561 | 2 | | | | | | | | 0 | 895 |
| 1560 | 2 | | | | | | | | 0 | 1666 |
| 1560 | 2 | | | | | | | | 0 | 1046 |
| 1559 | 1 | | | | | | | | 0 | 989 |
| 1557 | 1 | | | | | | | | 0 | 797 |
| 1557 | 3 | | | | | | | | 0 | 867 |
| 1556 | 1 | | | | | | | | 0 | 877 |
| 1556 | 1 | | | | | | | | 0 | 661 |
| 1556 | 1 | | | | | | | | 0 | 471 |
| 1553 | 1 | | | | | | | | 0 | 1521 |
| 1552 | 2 | | | | | | | | 0 | |
| 1552 | 1 | | | | | | | | 0 | |
| 1550 | 1 | | protein bi | Pf06229, I | 5568419 | AAEL0068 | AaeL_AAE | 3 | 0 | 971 |
| 1546 | 2 | | | | | | | | 0 | 434 |
| 1541 | 1 | | | | | | | | 0 | 782 |

| | | | | | | | | | |
|---|---|---|---|---|---|---|---|---|---|
| 1540 | 3 | cytoplasm | Pf04194 | 5579169 | AAEL0038 | AaeL_AAE | 3 | 0 | |
| 1537 | 1 | | | | | | | 0 | 1200 |
| 1533 | 1 | | | | | | | 0 | 1014 |
| 1532 | 2 | | | | | | | 0 | |
| 1532 | 2 | | | | | | | 0 | 886 |
| 1531 | 1 | | | | | | | 0 | 673 |
| 1524 | 2 | | | | | | | 0 | 410 |
| 1524 | 3 | | | | | | | 0 | 1014 |
| 1523 | 1 | | | | | | | 0 | |
| 1523 | 1 | | | | | | | 0 | |
| 1522 | 2 | | | | | | | 0 | 1031 |
| 1519 | 2 | | | | | | | 0 | 1058 |
| 1514 | 2 | | | | | | | 0 | 1066 |
| 1513 | 2 | | | | | | | 0 | 1015 |
| 1505 | 2 | | | | | | | 0 | 717 |
| 1504 | 3 | | | | | | | 0 | |
| 1504 | 2 | | | | | | | 0 | |
| 1502 | 2 | | | | | | | 0 | 1286 |
| 1500 | 3 | | | | | | | 0 | |
| 1499 | 1 | | Pf07258 | 5572866 | AAEL0100 | AaeL_AAE | 2 | 0 | 1022 |
| 1498 | 2 | | | | | | | 0 | |
| 1497 | 1 | | | | | | | 0 | 923 |
| 1497 | 1 | | | | | | | 0 | 809 |
| 1496 | 1 | | | | | | | 0 | 640 |
| 1495 | 2 | | | | | | | 0 | 918 |
| 1491 | 2 | | Pf06320 | 5570206 | AAEL0081 | AaeL_AAE | 3 | 0 | 934 |
| 1491 | 1 | | | | | | | 0 | 681 |
| 1487 | 1 | | | | | | | 0 | |
| 1486 | 3 | | | | | | | 0 | 994 |
| 1484 | 2 | | | | | | | 0 | 588 |
| 1483 | 1 | | | | | | | 0 | 1020 |

| | | | | | | | | | |
|---|---|---|---|---|---|---|---|---|---|
| 1483 | 2 | | | | | | | 0 | 746 |
| 1479 | 3 | | | | | | | 0 | 608 |
| 1478 | 2 | | | | | | | 0 | 906 |
| 1476 | 1 | | | | | | | 0 | 880 |
| 1474 | 2 | | | | | | | 0 | 827 |
| 1473 | 1 | | | | | | | 0 | |
| 1471 | 2 | | | | | | | 0 | |
| 1470 | 2 | | | | | | | 0 | |
| 1468 | 2 | | | | | | | 0 | 860 |
| 1464 | 3 | | | | | | | 0 | 1176 |
| 1463 | 2 | | | | | | | 0 | |
| 1463 | 3 | | | | | | | 0 | 393 |
| 1458 | 2 | | | | | | | 0 | 1095 |
| 1457 | 3 | | | | | | | 0 | 766 |
| 1453 | 2 | | | | | | | 0 | 920 |
| 1452 | 2 | | | | | | | 0 | 819 |
| 1451 | 2 | | | | | | | 0 | 646 |
| 1451 | 2 | | | | | | | 0 | |
| 1449 | 1 | | | | | | | 0 | 922 |
| 1448 | 2 | | | | | | | 0 | 807 |
| 1445 | 1 | | | | | | | 0 | |
| 1439 | 1 | | | | | | | 0 | 1016 |
| 1438 | 2 | | | | | | | 0 | |
| 1436 | 1 | | | | | | | 0 | 924 |
| 1436 | 2 | | | | | | | 0 | 671 |
| 1428 | 2 | | | | | | | 0 | 577 |
| 1424 | 3 | | | | | | | 0 | 778 |
| 1421 | 2 | | | | | | | 0 | |
| 1414 | 1 | | | | | | | 0 | 678 |
| 1410 | 1 | protein bi | Pf01398, I | 5569676 | AAEL0078 | AaeL_AAE | 2 | 0 | 1021 |
| 1409 | 3 | | | | | | | 0 | 846 |

| | | | | | | | | | | | |
|---|---|---|---|---|---|---|---|---|---|---|---|
| 1402 | 1 | | | | | | | | | 0 | |
| 1399 | 2 | | | | | | | | | 0 | 626 |
| 1395 | 1 | | | | | | | | | 0 | 788 |
| 1391 | 2 | | | | | | | | | 0 | |
| 1391 | 4 | | | | | | | | | 0 | 1125 |
| 1390 | 2 | | | | | | | | | 0 | 1113 |
| 1387 | 3 | | | | | | | | | 0 | |
| 1384 | 1 | | | | | | | | | 0 | 815 |
| 1383 | 1 | | | Pf05253 | 5571152 | AAEL0088 | AaeL_AAE | 2 | | 0 | 1170 |
| 1376 | 1 | | | | | | | | | 0 | 896 |
| 1375 | 3 | | | | | | | | | 0 | |
| 1374 | 2 | | | | | | | | | 0 | 384 |
| 1371 | 3 | | | Pf05871 | 5575384 | AAEL0117 | AaeL_AAE | 3 | Endocytosis | 1 | 849 |
| 1371 | 3 | | | | | | | | | 0 | 912 |
| 1367 | 2 | | | | | | | | | 0 | 1178 |
| 1365 | 3 | | | | | | | | | 0 | 498 |
| 1365 | 2 | | | | | | | | | 0 | |
| 1361 | 1 | | | | | | | | | 0 | 994 |
| 1360 | 2 | | | | | | | | | 0 | 788 |
| 1358 | 1 | | | | | | | | | 0 | 802 |
| 1356 | 2 | | | | | | | | | 0 | |
| 1353 | 1 | | | | | | | | | 0 | 591 |
| 1347 | 2 | | | | | | | | | 0 | |
| 1345 | 2 | | | | | | | | | 0 | 602 |
| 1345 | 2 | cell organization and biogenesi | | Pf04430 | 5571057 | AAEL0088 | AaeL_AAE | 3 | | 0 | 1047 |
| 1343 | 2 | | | | | | | | | 0 | 1323 |
| 1340 | 3 | | | | | | | | | 0 | |
| 1336 | 1 | | | | | | | | | 0 | 408 |
| 1336 | 2 | | | | | | | | | 0 | 622 |
| 1333 | 1 | regulation of biologi | enzyme re | Pf00023, I | 5572689 | AAEL0099 | AaeL_AAE | 2 | Endocytosis | 1 | 993 |
| 1329 | 2 | | | | | | | | | 0 | 797 |

| | | | | | | | | |
|---|---|---|---|---|---|---|---|---|
| 1328 | 1 | | | | | | 0 | 260 |
| 1328 | 1 | | | | | | 0 | 973 |
| 1327 | 1 | | | | | | 0 | 438 |
| 1326 | 2 | | | | | | 0 | |
| 1323 | 2 | | | | | | 0 | 1314 |
| 1322 | 1 | | 5571088 | AAEL0087 | AaeL_AAE | 3 | 0 | 850 |
| 1318 | 2 | | | | | | 0 | |
| 1314 | 2 | | | | | | 0 | |
| 1313 | 1 | | | | | | 0 | 977 |
| 1312 | 2 | | | | | | 0 | 747 |
| 1312 | 1 | | | | | | 0 | |
| 1312 | 2 | | | | | | 0 | 777 |
| 1310 | 2 | | | | | | 0 | |
| 1306 | 1 | | | | | | 0 | 870 |
| 1305 | 1 | | | | | | 0 | 1087 |
| 1302 | 2 | | | | | | 0 | 373 |
| 1302 | 2 | | | | | | 0 | 783 |
| 1302 | 1 | | 5569606 | AAEL0012 | AaeL_AAE | 2 | 0 | 998 |
| 1299 | 3 | | | | | | 0 | |
| 1295 | 2 | | | | | | 0 | 849 |
| 1294 | 1 | | | | | | 0 | 823 |
| 1293 | 1 | | | | | | 0 | 804 |
| 1292 | 2 | | | | | | 0 | 606 |
| 1292 | 1 | | | | | | 0 | 1040 |
| 1282 | 2 | | | | | | 0 | 611 |
| 1282 | 2 | | | | | | 0 | 610 |
| 1277 | 1 | | | | | | 0 | 575 |
| 1276 | 1 | Pf04818 | 5566031 | AAEL0051 | AaeL_AAE | 2 | 0 | 1137 |
| 1276 | 3 | | | | | | 0 | 457 |
| 1275 | 1 | | | | | | 0 | 854 |
| 1273 | 1 | | | | | | 0 | 801 |

| | | | | | | | | | |
|---|---|---|---|---|---|---|---|---|---|
| 1272 | 2 | | | | | | | 0 | 1121 |
| 1270 | 1 | | | | | | | 0 | 708 |
| 1268 | 2 | | | | | | | 0 | 466 |
| 1263 | 1 | | | | | | | 0 | |
| 1262 | 2 | | | | | | | 0 | 1418 |
| 1261 | 1 | | Pf00169, I | 5570804 | AAEL0085 | AaeL_AAE | 3 | 0 | 692 |
| 1261 | 1 | | | | | | | 0 | |
| 1260 | 2 | | | | | | | 0 | 863 |
| 1259 | 2 | cell organization and biogenesi | Pf03985 | 5570065 | AAEL0013 | AaeL_AAE | 3 | 0 | 843 |
| 1259 | 2 | | | | | | | 0 | |
| 1258 | 2 | | Pf06747 | 5568340; | AAEL0010 | AaeL_AAE | 1 | 0 | 889 |
| 1257 | 2 | | | | | | | 0 | |
| 1254 | 2 | | | | | | | 0 | |
| 1251 | 2 | | | | | | | 0 | 728 |
| 1250 | 1 | | | | | | | 0 | 897 |
| 1248 | 2 | | | | | | | 0 | |
| 1242 | 1 | | | | | | | 0 | 1047 |
| 1242 | 1 | | | | | | | 0 | 768 |
| 1242 | 2 | | Pf15623 | 5570060 | AAEL0013 | AaeL_AAE | 3 | 0 | |
| 1240 | 2 | | | | | | | 0 | |
| 1240 | 1 | | | | | | | 0 | 648 |
| 1239 | 1 | | | | | | | 0 | 659 |
| 1237 | 2 | membrane | | 5569056 | AAEL0073 | AaeL_AAE | 1 | 0 | |
| 1236 | 2 | | | | | | | 0 | 817 |
| 1232 | 3 | | | | | | | 0 | |
| 1230 | 1 | | | | | | | 0 | 989 |
| 1229 | 2 | | | | | | | 0 | 724 |
| 1226 | 2 | | | | | | | 0 | 532 |
| 1226 | 2 | | | | | | | 0 | 811 |
| 1224 | 2 | | | | | | | 0 | |
| 1224 | 2 | | | | | | | 0 | 736 |

| | | | | | | | | | | | | |
|---|---|---|---|---|---|---|---|---|---|---|---|---|
| 1222 | 2 | | | | | | | | | | 0 | 965 |
| 1222 | 1 | | | | | | | | | | 0 | |
| 1219 | 1 | | | | | | | | | | 0 | |
| 1217 | 2 | | | | | | | | | | 0 | 741 |
| 1216 | 1 | | | | | | | | | | 0 | 911 |
| 1216 | 3 | | | | | | | | | | 0 | 887 |
| 1213 | 1 | | | | | | | | | | 0 | 675 |
| 1213 | 2 | | | | | | | | | | 0 | 721 |
| 1211 | 2 | | | | | | | | | | 0 | 695 |
| 1210 | 1 | | | | | | | | | | 0 | |
| 1206 | 3 | | | | | | | | | | 0 | |
| 1204 | 2 | | | | | | | | | | 0 | 832 |
| 1203 | 1 | | | | | | | | | | 0 | |
| 1202 | 2 | | | | | | | | | | 0 | |
| 1200 | 1 | transport | cytoplasm | protein bi | Pf02136, I | 5567843 | AAEL0010 | AaeL_AAE | 3 | mRNA surveillance pathway; Ri | 3 | 941 |
| 1198 | 2 | | | | | | | | | | 0 | 620 |
| 1198 | 2 | | | | | | | | | | 0 | 1359 |
| 1197 | 1 | | | | | | | | | | 0 | 692 |
| 1197 | 2 | | | | | | | | | | 0 | 1169 |
| 1194 | 2 | | | | | 5569590 | AAEL0012 | AaeL_AAE | 2 | | 0 | 1070 |
| 1193 | 1 | | | | | | | | | | 0 | 1168 |
| 1192 | 1 | | | | | | | | | | 0 | |
| 1190 | 1 | cell organization and biogenesi | | | Pf08265 | 5572446 | AAEL0018 | AaeL_AAE | 2 | | 0 | 989 |
| 1190 | 2 | | | | | | | | | | 0 | 912 |
| 1186 | 1 | | | | | | | | | | 0 | 378 |
| 1183 | 2 | | | | | | | | | | 0 | 754 |
| 1182 | 2 | | | | | | | | | | 0 | |
| 1182 | 1 | | | | | | | | | | 0 | 287 |
| 1181 | 2 | | | | | | | | | | 0 | |
| 1180 | 2 | | | | | | | | | | 0 | |
| 1177 | 1 | | | | | | | | | | 0 | 945 |

| | | | | | | | | | | |
|---|---|---|---|---|---|---|---|---|---|---|
| 1173 | 2 | | | | | | | | 0 | |
| 1172 | 2 | | | | | | | | 0 | 1011 |
| 1170 | 2 | | | | | | | | 0 | 1222 |
| 1167 | 1 | membrane | | | 5564282 | AAEL0143 | AaeL_AAE | 3 | 0 | 615 |
| 1167 | 2 | | | | | | | | 0 | |
| 1167 | 1 | | | | | | | | 0 | 1006 |
| 1165 | 2 | | | | | | | | 0 | 1291 |
| 1164 | 2 | | | | | | | | 0 | 887 |
| 1162 | 1 | | | | | | | | 0 | 733 |
| 1160 | 2 | | | | | | | | 0 | |
| 1160 | 1 | | | | | | | | 0 | 766 |
| 1153 | 1 | | | | | | | | 0 | |
| 1152 | 2 | | | | | | | | 0 | |
| 1150 | 2 | | | | | | | | 0 | |
| 1149 | 1 | membrane | | Pf14640 | 5569167 | AAEL0074 | AaeL_AAE | 2 | 0 | 539 |
| 1149 | 1 | | | | | | | | 0 | 542 |
| 1149 | 2 | | | | | | | | 0 | 858 |
| 1148 | 2 | | | | | | | | 0 | |
| 1148 | 2 | | | | | | | | 0 | 787 |
| 1147 | 1 | | | | | | | | 0 | |
| 1145 | 2 | | | | | | | | 0 | 697 |
| 1144 | 1 | | | | | | | | 0 | 1006 |
| 1143 | 2 | | | | | | | | 0 | 1174 |
| 1135 | 2 | | | | | | | | 0 | |
| 1135 | 1 | | | | | | | | 0 | 916 |
| 1130 | 1 | | | | | | | | 0 | |
| 1127 | 1 | | RNA bindi | Pf00076, I | 5568210 | AAEL0066 | AaeL_AAE | 3 | 0 | 721 |
| 1126 | 1 | | | | | | | | 0 | |
| 1125 | 1 | | | | | | | | 0 | 837 |
| 1124 | 1 | | | | | | | | 0 | 1175 |
| 1123 | 2 | | | | | | | | 0 | |

| | | | | | | | | | | |
|---|---|---|---|---|---|---|---|---|---|---|
| 1123 | 2 | | | | | | | | 0 | 1293 |
| 1119 | 1 | | | | | | | | 0 | 828 |
| 1118 | 1 | | | | | | | | 0 | |
| 1117 | 1 | | | | | | | | 0 | 994 |
| 1115 | 1 | | | | | | | | 0 | 1074 |
| 1112 | 1 | | | | | | | | 0 | 690 |
| 1112 | 2 | metabolic process | catalytic a | Pf01564, I | 5573934 | AAEL0021 | AaeL_AAE | 2 | 0 | 656 |
| 1111 | 2 | | | | | | | | 0 | |
| 1104 | 2 | | | | | | | | 0 | |
| 1104 | 1 | | | | | | | | 0 | 1003 |
| 1103 | 1 | | | | | | | | 0 | 839 |
| 1102 | 1 | | | | | | | | 0 | |
| 1102 | 2 | | | | | | | | 0 | 749 |
| 1102 | 2 | | | | | | | | 0 | 784 |
| 1101 | 2 | | | | | | | | 0 | |
| 1100 | 1 | | | | | | | | 0 | |
| 1099 | 1 | | | | | | | | 0 | 902 |
| 1099 | 1 | | | | | | | | 0 | 649 |
| 1098 | 1 | | | | | | | | 0 | 853 |
| 1097 | 1 | | | | | | | | 0 | 701 |
| 1096 | 1 | | | | | | | | 0 | 955 |
| 1096 | 1 | | | | | | | | 0 | |
| 1096 | 1 | | | | | | | | 0 | 1096 |
| 1096 | 1 | | | | | | | | 0 | |
| 1095 | 1 | | | | | | | | 0 | |
| 1095 | 2 | | | | | | | | 0 | 774 |
| 1093 | 2 | | | | | | | | 0 | 743 |
| 1089 | 1 | | | | | | | | 0 | 1008 |
| 1089 | 1 | | | | | | | | 0 | 1012 |
| 1088 | 2 | | | | | | | | 0 | 822 |
| 1088 | 1 | | | | | | | | 0 | 1066 |

| | | | | | | | | | |
|---|---|---|---|---|---|---|---|---|---|
| 1087 | 1 | | Pf09360 | 5564228 | AAEL0041 | AaeL_AAE | 2 | 0 | |
| 1086 | 2 | | | | | | | 0 | |
| 1086 | 1 | | | | | | | 0 | 541 |
| 1086 | 2 | | | | | | | 0 | 1121 |
| 1084 | 1 | | | | | | | 0 | 1265 |
| 1084 | 1 | | | | | | | 0 | 1025 |
| 1083 | 2 | | | | | | | 0 | 982 |
| 1082 | 2 | | | | | | | 0 | 1122 |
| 1081 | 1 | RNA bindi | Pf00013, I | 5574289 | AAEL0110 | AaeL_AAE | 2 | 0 | |
| 1076 | 1 | | | | | | | 0 | 868 |
| 1075 | 1 | | | | | | | 0 | 1001 |
| 1075 | 3 | | | | | | | 0 | |
| 1073 | 1 | | | 5568497 | AAEL0069 | AaeL_AAE | 2 | 0 | 802 |
| 1072 | 1 | | | | | | | 0 | |
| 1068 | 2 | | | | | | | 0 | |
| 1068 | 1 | | | | | | | 0 | 585 |
| 1066 | 2 | | | 5574187 | AAEL0023 | AaeL_AAE | 3 | 0 | |
| 1063 | 2 | | | | | | | 0 | 644 |
| 1062 | 1 | | | | | | | 0 | 864 |
| 1062 | 1 | | | | | | | 0 | 868 |
| 1061 | 1 | | | | | | | 0 | |
| 1059 | 2 | | | | | | | 0 | 311 |
| 1057 | 1 | | | | | | | 0 | 861 |
| 1056 | 2 | | | | | | | 0 | |
| 1054 | 1 | | | | | | | 0 | |
| 1054 | 2 | | | | | | | 0 | 1075 |
| 1054 | 2 | | | | | | | 0 | 702 |
| 1050 | 2 | | | | | | | 0 | 1104 |
| 1050 | 1 | | | | | | | 0 | 640 |
| 1049 | 1 | | | | | | | 0 | 472 |
| 1049 | 2 | | | | | | | 0 | |

| | | | | | | | | | | | | |
|---|---|---|---|---|---|---|---|---|---|---|---|---|
| 1047 | 1 | | | | | | | | | | 0 | 638 |
| 1047 | 1 | metabolic process | | catalytic a | Pf00067 | 23687481 | AAEL0170 | AaeL_AAE | 1 | | 0 | 393 |
| 1046 | 1 | regulatior | nucleus | metal ion | Pf00536, I | 5566512 | AAEL0054 | AaeL_AAE | 2 | | 0 | |
| 1038 | 1 | | | | | | | | | | 0 | |
| 1038 | 2 | | | | | | | | | | 0 | 1117 |
| 1038 | 2 | | | | | | | | | | 0 | 853 |
| 1037 | 2 | | | | | | | | | | 0 | 684 |
| 1037 | 1 | | | protein bi | Pf00023, I | 5574481; | AAEL0023 | AaeL_AAE | 2 | | 0 | 575 |
| 1036 | 2 | | | | | | | | | | 0 | |
| 1034 | 1 | | | | | | | | | | 0 | 418 |
| 1033 | 1 | | | | | | | | | | 0 | 897 |
| 1033 | 1 | | | | | | | | | | 0 | 537 |
| 1033 | 2 | | | | | | | | | | 0 | 815 |
| 1033 | 2 | | | | | | | | | | 0 | 851 |
| 1033 | 2 | | | | | | | | | | 0 | 802 |
| 1032 | 2 | | | | | | | | | | 0 | |
| 1030 | 2 | | | | | | | | | | 0 | |
| 1028 | 1 | | | | | | | | | | 0 | |
| 1027 | 2 | | | | | | | | | | 0 | |
| 1027 | 1 | | | | | | | | | | 0 | 633 |
| 1026 | 1 | | | | | | | | | | 0 | |
| 1025 | 1 | | | | | | | | | | 0 | |
| 1025 | 1 | | | | | | | | | | 0 | 865 |
| 1024 | 2 | | | | | | | | | | 0 | |
| 1024 | 1 | | | | | | | | | | 0 | |
| 1022 | 1 | | | | | | | | | | 0 | 409 |
| 1022 | 1 | | | nucleotid | Pf08585 | 5568693 | AAEL0070 | AaeL_AAE | 3 | Fanconi anemia pathway | 1 | 863 |
| 1022 | 2 | | | | | | | | | | 0 | 686 |
| 1021 | 2 | | | | | | | | | | 0 | 1269 |
| 1021 | 2 | | | | | | | | | | 0 | 957 |
| 1020 | 1 | | | | | | | | | | 0 | 984 |

| | | | | | | | | | | | | |
|---|---|---|---|---|---|---|---|---|---|---|---|---|
| 1019 | 1 | | | | | | | | | | 0 | 869 |
| 1018 | 2 | | | | | | | | | | 0 | 1000 |
| 1015 | 2 | | | | | | | | | | 0 | 1053 |
| 1014 | 1 | regulatior | extracellu | protein bi | Pf00049 | 5567483 | AAEL0009 | AaeL_AAE | 3 | mTOR signaling pathway; Longe | 4 | 667 |
| 1014 | 2 | | | | | | | | | | 0 | 587 |
| 1011 | 2 | | | | | | | | | | 0 | 750 |
| 1007 | 1 | | | | | | | | | | 0 | 1021 |
| 1005 | 1 | | | | | | | | | | 0 | 937 |
| 1004 | 2 | | | | | | | | | | 0 | 880 |
| 1004 | 2 | | | | | | | | | | 0 | 350 |
| 1004 | 2 | | | | | | | | | | 0 | 1049 |
| 1002 | 1 | | | | | | | | | | 0 | |
| 1002 | 2 | | | | | | | | | | 0 | |
| 1001 | 2 | | | | | | | | | | 0 | |
| 1000 | 1 | | | | | | | | | | 0 | 755 |
| 999 | 1 | | | | | | | | | | 0 | |
| 999 | 1 | | | | | | | | | | 0 | 652 |
| 998 | 1 | | | | | | | | | | 0 | 646 |
| 998 | 1 | regulatior | nucleus | DNA bindi | Pf09011, I | 5567819 | AAEL0181 | AaeL_AAE | 3 | | 0 | 587 |
| 996 | 2 | | | | | | | | | | 0 | |
| 996 | 1 | | | | | | | | | | 0 | |
| 995 | 1 | | | | | | | | | | 0 | 1007 |
| 994 | 1 | | | | | | | | | | 0 | 542 |
| 994 | 1 | | | | | | | | | | 0 | |
| 988 | 1 | | | | | | | | | | 0 | |
| 988 | 2 | | | | | | | | | | 0 | 959 |
| 984 | 1 | | | | | | | | | | 0 | |
| 983 | 2 | | | | | | | | | | 0 | 928 |
| 983 | 1 | | | | | | | | | | 0 | 930 |
| 981 | 1 | | | | | | | | | | 0 | 588 |
| 981 | 1 | cell organ | membrane | | Pf07086 | 23687484 | AAEL0170 | AaeL_AAE | 3 | | 0 | |

| | | | | | | | | | | | |
|---|---|---|---|---|---|---|---|---|---|---|---|
| 980 | 2 | | | | | | | | | 0 | 749 |
| 979 | 2 | | | | | | | | | 0 | |
| 978 | 2 | | | | | | | | | 0 | |
| 973 | 1 | | | | | | | | | 0 | 1466 |
| 966 | 1 | | | | | | | | | 0 | 875 |
| 966 | 2 | | | | | | | | | 0 | 728 |
| 965 | 2 | | | | | | | | | 0 | |
| 965 | 1 | | | | | | | | | 0 | |
| 961 | 2 | | | | | | | | | 0 | |
| 954 | 1 | | | | | | | | | 0 | |
| 954 | 1 | | | | | 5574964 | AAEL0025 | AaeL_AAE | 1 | 0 | 639 |
| 953 | 1 | | | | | | | | | 0 | |
| 952 | 1 | | | | | | | | | 0 | |
| 950 | 1 | | | | | | | | | 0 | |
| 950 | 1 | | | | | | | | | 0 | |
| 942 | 1 | | | | | | | | | 0 | 789 |
| 942 | 2 | | | | | | | | | 0 | 802 |
| 937 | 1 | | | | | | | | | 0 | 1826 |
| 936 | 2 | | | | | | | | | 0 | |
| 936 | 1 | | | | | | | | | 0 | |
| 932 | 2 | metabolic process | | catalytic a | Pf00135, I | 5563952 | AAEL0040 | AaeL_AAE | 3 | 0 | 1104 |
| 930 | 1 | | | | | | | | | 0 | |
| 930 | 1 | | | | | | | | | 0 | |
| 929 | 1 | | | | | | | | | 0 | |
| 929 | 2 | | | | | | | | | 0 | |
| 929 | 2 | | membrane | | | 5564097 | AAEL0143 | AaeL_AAE | 2 | 0 | |
| 926 | 2 | | | | | | | | | 0 | 755 |
| 925 | 1 | | | | | | | | | 0 | 648 |
| 925 | 1 | | | | | | | | | 0 | |
| 924 | 2 | | | | | | | | | 0 | 989 |
| 922 | 2 | | | | | | | | | 0 | |

| | | | | | | | | | | |
|---|---|---|---|---|---|---|---|---|---|---|
| 921 | 1 | | | | | | | | 0 | |
| 920 | 2 | metabolic process | catalytic a | Pf00290, I | 5564189 | AAEL0143 | AaeL_AAE | 2 | 0 | 819 |
| 918 | 2 | | | | | | | | 0 | |
| 916 | 2 | | | | | | | | 0 | 731 |
| 916 | 1 | | | | | | | | 0 | 736 |
| 911 | 1 | | | | | | | | 0 | 1004 |
| 910 | 1 | | | | | | | | 0 | |
| 910 | 2 | | | | | | | | 0 | 795 |
| 909 | 1 | | | | | | | | 0 | |
| 909 | 2 | | | Pf01722 | 5571200 | AAEL0088 | AaeL_AAE | 3 | 0 | 1395 |
| 909 | 1 | | | Pf07946 | 5564195 | AAEL0143 | AaeL_AAE | 2 | 0 | 940 |
| 909 | 2 | | | | | | | | 0 | |
| 907 | 1 | | | | | | | | 0 | |
| 905 | 2 | | | | | | | | 0 | 1066 |
| 905 | 1 | | | | | | | | 0 | |
| 904 | 2 | | | | | | | | 0 | 1043 |
| 903 | 2 | | | | | | | | 0 | 393 |
| 902 | 1 | | | | | | | | 0 | 593 |
| 900 | 1 | | | | | | | | 0 | 718 |
| 899 | 1 | | | | | | | | 0 | |
| 898 | 2 | | | | | | | | 0 | 684 |
| 898 | 1 | | | | | | | | 0 | 741 |
| 898 | 1 | | | | | | | | 0 | |
| 896 | 1 | | | | | | | | 0 | |
| 896 | 1 | | | | | | | | 0 | |
| 893 | 1 | | | | | | | | 0 | 985 |
| 892 | 1 | | | | 5564548 | AAEL0043 | AaeL_AAE | 3 | 0 | 1338 |
| 891 | 1 | | | | | | | | 0 | |
| 891 | 2 | | | | | | | | 0 | |
| 888 | 1 | | | | | | | | 0 | 598 |
| 887 | 1 | | | | | | | | 0 | 890 |

| | | | | | | | | |
|---|---|---|---|---|---|---|---|---|
| 883 | 2 | | | | | | 0 | 388 |
| 882 | 1 | | | | | | 0 | 796 |
| 880 | 1 | | | | | | 0 | |
| 878 | 1 | RNA bindi Pf00076, I | 5575824 | AAEL0121 | AaeL_AAE | 1 | 0 | 1208 |
| 875 | 1 | | | | | | 0 | 762 |
| 873 | 1 | | | | | | 0 | |
| 873 | 1 | | | | | | 0 | 912 |
| 873 | 1 | | | | | | 0 | |
| 872 | 1 | | | | | | 0 | |
| 871 | 2 | | | | | | 0 | 604 |
| 871 | 2 | | | | | | 0 | 526 |
| 870 | 1 | | | | | | 0 | |
| 870 | 1 | | | | | | 0 | |
| 866 | 1 | | | | | | 0 | |
| 862 | 2 | | | | | | 0 | |
| 862 | 1 | | | | | | 0 | |
| 861 | 2 | | | | | | 0 | 663 |
| 859 | 1 | | | | | | 0 | |
| 858 | 1 | | | | | | 0 | |
| 857 | 1 | | | | | | 0 | 906 |
| 855 | 1 | | | | | | 0 | |
| 855 | 1 | | | | | | 0 | 816 |
| 852 | 1 | | | | | | 0 | |
| 851 | 1 | | | | | | 0 | |
| 850 | 2 | | | | | | 0 | |
| 847 | 2 | | 5569651 | AAEL0078 | AaeL_AAE | 3 | 0 | 1081 |
| 846 | 2 | | | | | | 0 | 832 |
| 842 | 2 | | | | | | 0 | |
| 841 | 1 | | | | | | 0 | |
| 838 | 2 | | | | | | 0 | 542 |
| 838 | 1 | | | | | | 0 | |

| | | | | | | | | | | | | |
|---|---|---|---|---|---|---|---|---|---|---|---|---|
| 836 | 1 | | | | | | | | | | 0 | |
| 834 | 1 | | | | | | | | | | 0 | 739 |
| 834 | 1 | | | | | | | | | | 0 | |
| 833 | 1 | | | | | | | | | | 0 | |
| 832 | 2 | | | | | | | | | | 0 | 1201 |
| 832 | 2 | | membran | protein bi | Pf00400 | 5576862 | AAEL0128 | AaeL_AAE | 2 | Protein processing in endoplası | 1 | |
| 830 | 1 | | | | | | | | | | 0 | |
| 830 | 1 | transport | | | Pf05667, I | 5573261 | AAEL0103 | AaeL_AAE | 3 | | 0 | 1012 |
| 830 | 1 | | | | | | | | | | 0 | |
| 830 | 1 | | | protein bi | Pf00400 | 5572057 | AAEL0017 | AaeL_AAE | 2 | | 0 | |
| 828 | 1 | | | | | | | | | | 0 | |
| 828 | 2 | | | | | | | | | | 0 | |
| 827 | 2 | | | | | | | | | | 0 | |
| 825 | 2 | | | | | | | | | | 0 | 1255 |
| 825 | 1 | | | | | | | | | | 0 | 765 |
| 825 | 1 | | | | | | | | | | 0 | |
| 824 | 1 | | | | | | | | | | 0 | 761 |
| 823 | 1 | metabolic process | | | Pf08615 | 5573068 | AAEL0102 | AaeL_AAE | 3 | DNA replication | 1 | |
| 823 | 2 | metabolic process | | catalytic a | Pf08241, I | 5579830 | AAEL0058 | AaeL_AAE | 3 | | 0 | 963 |
| 823 | 1 | | | | | | | | | | 0 | 808 |
| 823 | 1 | | | | | | | | | | 0 | 707 |
| 821 | 2 | | | | | | | | | | 0 | 1114 |
| 820 | 1 | | | | | | | | | | 0 | |
| 820 | 1 | | | | | | | | | | 0 | |
| 820 | 1 | | | | Pf04774 | 5569984 | AAEL0080 | AaeL_AAE | 2 | | 0 | 1885 |
| 819 | 1 | | | | | | | | | | 0 | 560 |
| 818 | 1 | | | | | | | | | | 0 | 643 |
| 818 | 2 | | | | Pf04031 | 5574113 | AAEL0022 | AaeL_AAE | 2 | | 0 | 811 |
| 816 | 2 | | | | | | | | | | 0 | 525 |
| 816 | 1 | | | | | | | | | | 0 | 1055 |
| 809 | 1 | | | | | | | | | | 0 | 757 |

| | | | | | | | | | | |
|---|---|---|---|---|---|---|---|---|---|---|
| 808 | 1 | | | | | | | | 0 | |
| 808 | 1 | | | | | | | | 0 | 849 |
| 807 | 1 | | | | | | | | 0 | 1074 |
| 807 | 2 | | | | | | | | 0 | |
| 806 | 2 | | | | | | | | 0 | |
| 805 | 1 | | | | | | | | 0 | |
| 805 | 2 | | | Pf01424 | 5573831 | AAEL0107 | AaeL_AAE | 3 | 0 | |
| 804 | 2 | | | | | | | | 0 | |
| 804 | 1 | metabolic process | catalytic a | Pf00561, I | 5570939 | AAEL0086 | AaeL_AAE | 2 | 0 | 729 |
| 803 | 1 | | | | | | | | 0 | |
| 800 | 1 | | | | | | | | 0 | |
| 799 | 2 | | | | | | | | 0 | |
| 799 | 1 | | | | | | | | 0 | |
| 798 | 1 | | | | | | | | 0 | 906 |
| 798 | 1 | | | | | | | | 0 | 754 |
| 795 | 1 | | | | | | | | 0 | 1146 |
| 794 | 1 | | | | | | | | 0 | |
| 791 | 1 | | | | | | | | 0 | |
| 788 | 1 | | | | | | | | 0 | |
| 788 | 1 | | | | | | | | 0 | |
| 787 | 2 | | | | | | | | 0 | |
| 785 | 1 | | | | | | | | 0 | 719 |
| 781 | 1 | | | | | | | | 0 | |
| 779 | 2 | | | | | | | | 0 | 1018 |
| 778 | 2 | | | | | | | | 0 | |
| 778 | 1 | | | | | | | | 0 | |
| 777 | 1 | | | | | | | | 0 | |
| 776 | 1 | | | | | | | | 0 | |
| 776 | 1 | | | | | | | | 0 | 1125 |
| 774 | 1 | | | | | | | | 0 | 914 |
| 773 | 1 | | | | | | | | 0 | |

| | | | | | | | | | | |
|---|---|---|---|---|---|---|---|---|---|---|
| 773 | 1 | | | | | | | | 0 | 461 |
| 772 | 1 | membrane | Pf05444 | 5571275 | AAEL0015 | AaeL_AAE | 2 | | 0 | |
| 769 | 2 | | | | | | | | 0 | 864 |
| 767 | 1 | | | | | | | | 0 | |
| 766 | 1 | | | | | | | | 0 | 637 |
| 766 | 1 | | | | | | | | 0 | |
| 766 | 1 | | | | | | | | 0 | 958 |
| 765 | 1 | | | | | | | | 0 | |
| 764 | 1 | | | | | | | | 0 | 1034 |
| 764 | 2 | membrane | Pf04756 | 5566520 | AAEL0054 | AaeL_AAE | 2 | Protein processing in endoplasi | 3 | 503 |
| 763 | 1 | | | | | | | | 0 | |
| 762 | 2 | | | | | | | | 0 | |
| 761 | 1 | | | | | | | | 0 | 940 |
| 759 | 1 | | | | | | | | 0 | 853 |
| 759 | 1 | | | | | | | | 0 | |
| 759 | 1 | | | | | | | | 0 | |
| 756 | 1 | protein bi | Pf01399 | 5572030 | AAEL0002 | AaeL_AAE | 1 | | 0 | 475 |
| 754 | 1 | | | | | | | | 0 | |
| 754 | 1 | | | | | | | | 0 | 963 |
| 753 | 2 | | | | | | | | 0 | 887 |
| 751 | 2 | | | | | | | | 0 | |
| 750 | 1 | | | | | | | | 0 | 883 |
| 748 | 1 | | | | | | | | 0 | 827 |
| 748 | 1 | | Pf08318 | 5572921 | AAEL0101 | AaeL_AAE | 2 | | 0 | 915 |
| 746 | 1 | | | | | | | | 0 | |
| 744 | 1 | | | | | | | | 0 | |
| 744 | 1 | | | | | | | | 0 | 884 |
| 741 | 2 | | | | | | | | 0 | |
| 741 | 1 | | | | | | | | 0 | |
| 741 | 1 | | | | | | | | 0 | |
| 739 | 1 | | | | | | | | 0 | |

| | | | | | | | | | | |
|---|---|---|---|---|---|---|---|---|---|---|
| 739 | 1 | | | | | | | | 0 | 1237 |
| 739 | 1 | | | | | | | | 0 | |
| 738 | 2 | | | | | | | | 0 | 694 |
| 737 | 1 | | | | | | | | 0 | |
| 733 | 2 | membrane | | Pf10507 | 5572431 | AAEL0018 | AaeL_AAE | 2 | 0 | |
| 731 | 1 | | | | | | | | 0 | |
| 731 | 1 | | | | | | | | 0 | |
| 730 | 1 | | | | | | | | 0 | |
| 727 | 1 | | | | | | | | 0 | |
| 726 | 1 | | | | | | | | 0 | |
| 726 | 1 | | | | | | | | 0 | 796 |
| 725 | 1 | | | | | | | | 0 | 717 |
| 725 | 1 | | | | | | | | 0 | 591 |
| 725 | 1 | | | | | | | | 0 | 529 |
| 724 | 1 | | | | | | | | 0 | |
| 721 | 1 | | | | | | | | 0 | 756 |
| 721 | 1 | | | | | | | | 0 | |
| 720 | 1 | | | | | | | | 0 | |
| 718 | 1 | | | | | | | | 0 | 1003 |
| 718 | 2 | | | | | | | | 0 | |
| 717 | 1 | | | | | | | | 0 | |
| 717 | 1 | | | | | | | | 0 | 1046 |
| 714 | 1 | cell organization and | metal ion | Pf01106, I | 5580066 | AAEL0021 | AaeL_AAE | 1 | 0 | |
| 713 | 1 | | | | | | | | 0 | 1042 |
| 711 | 2 | | | | | | | | 0 | |
| 710 | 1 | | | | | | | | 0 | |
| 709 | 1 | | | | | | | | 0 | |
| 709 | 1 | | | | | | | | 0 | |
| 708 | 1 | | | | | | | | 0 | |
| 707 | 1 | | | | | | | | 0 | 692 |
| 706 | 1 | | | | | | | | 0 | |

| | | | | | | | | | |
|---|---|---|---|---|---|---|---|---|---|
| 706 | 2 | | | | | | | 0 | 661 |
| 702 | 1 | | | | | | | 0 | 781 |
| 702 | 1 | | | | | | | 0 | |
| 700 | 1 | | | | | | | 0 | |
| 700 | 1 | | | | | | | 0 | |
| 699 | 2 | | | | | | | 0 | 778 |
| 698 | 2 | | | | | | | 0 | |
| 697 | 1 | | | | | | | 0 | |
| 695 | 1 | DNA bindi | Pf04379, I | 5574410 | AAEL0111 | AaeL_AAE | 3 | 0 | 818 |
| 694 | 1 | | | | | | | 0 | |
| 694 | 1 | | | | | | | 0 | 864 |
| 693 | 1 | | | | | | | 0 | 694 |
| 692 | 1 | | | | | | | 0 | 1105 |
| 691 | 1 | | | | | | | 0 | 779 |
| 691 | 1 | | | | | | | 0 | 821 |
| 691 | 1 | | | | | | | 0 | 1243 |
| 691 | 1 | | | | | | | 0 | |
| 690 | 1 | | | | | | | 0 | |
| 690 | 1 | | | | | | | 0 | |
| 687 | 1 | | | | | | | 0 | |
| 686 | 1 | | | | | | | 0 | |
| 686 | 1 | | | | | | | 0 | |
| 686 | 1 | | | | | | | 0 | 1340 |
| 685 | 1 | | | | | | | 0 | |
| 685 | 1 | | | | | | | 0 | 710 |
| 683 | 1 | | | | | | | 0 | |
| 682 | 1 | | | | | | | 0 | |
| 682 | 1 | | | | | | | 0 | 375 |
| 681 | 1 | | | | | | | 0 | |
| 680 | 2 | | | | | | | 0 | 1052 |
| 680 | 1 | | | | | | | 0 | |

| | | | | | | | | | | | |
|---|---|---|---|---|---|---|---|---|---|---|---|
| 679 | 1 | | | | | | | | | 0 | |
| 678 | 1 | | | | | | | | | 0 | 698 |
| 677 | 1 | | | | | | | | | 0 | 641 |
| 676 | 1 | | | | | | | | | 0 | |
| 676 | 1 | | | | | | | | | 0 | |
| 675 | 1 | | | | | | | | | 0 | 826 |
| 675 | 1 | | | | | | | | | 0 | 1075 |
| 674 | 1 | | | | Pf08161 | 5575632 | AAEL0119 | AaeL_AAE | 3 | 0 | |
| 673 | 1 | | | | | | | | | 0 | |
| 673 | 1 | | | | | | | | | 0 | 749 |
| 672 | 1 | | | | | | | | | 0 | |
| 672 | 1 | | | | | | | | | 0 | |
| 672 | 2 | | | | | | | | | 0 | 880 |
| 672 | 1 | | | | | | | | | 0 | |
| 672 | 1 | metabolic process | | catalytic a | Pf00326, I | 5566852 | AAEL0008 | AaeL_AAE | 3 | 0 | 742 |
| 671 | 1 | | | | | | | | | 0 | |
| 671 | 1 | | | | | | | | | 0 | 811 |
| 670 | 1 | | | | | | | | | 0 | 1127 |
| 669 | 1 | | | | | | | | | 0 | |
| 668 | 1 | | | | | | | | | 0 | |
| 668 | 1 | | | | | | | | | 0 | |
| 667 | 1 | | | | | | | | | 0 | |
| 667 | 2 | | | | | | | | | 0 | |
| 665 | 1 | | | | | | | | | 0 | |
| 664 | 1 | | | | | | | | | 0 | 352 |
| 663 | 1 | | | | | | | | | 0 | |
| 662 | 1 | | membrane | | | 5577300 | AAEL0131 | AaeL_AAE | 2 | 0 | |
| 662 | 1 | | | | | | | | | 0 | |
| 661 | 1 | | | | | | | | | 0 | |
| 660 | 1 | | | catalytic a | Pf00583, I | 5578808 | AAEL0138 | AaeL_AAE | 1 | 0 | 1695 |
| 660 | 1 | | | | | | | | | 0 | |

| | | | | | | | | | | | |
|---|---|---|---|---|---|---|---|---|---|---|---|
| 658 | 1 | membrane | | Pf00812 | 5564929 | AAEL0044 | AaeL_AAE | 2 | | 0 | |
| 658 | 1 | | | | | | | | | 0 | 599 |
| 657 | 1 | | | | | | | | | 0 | |
| 656 | 1 | | | | | | | | | 0 | |
| 656 | 1 | | | | | | | | | 0 | 942 |
| 655 | 1 | | | | | | | | | 0 | |
| 654 | 1 | | | | | | | | | 0 | |
| 654 | 2 | | | | | | | | | 0 | |
| 653 | 1 | | | Pf10195 | 5577423 | AAEL0031 | AaeL_AAE | 1 | | 0 | 859 |
| 653 | 1 | cell organization and | RNA bindi | Pf01922 | 5565657 | AAEL0049 | AaeL_AAE | 2 | Protein export | 1 | 848 |
| 653 | 1 | | | | | | | | | 0 | |
| 651 | 1 | | | | | | | | | 0 | |
| 651 | 1 | | | | | | | | | 0 | 1088 |
| 650 | 1 | | | | | | | | | 0 | |
| 650 | 1 | | | | | | | | | 0 | |
| 650 | 1 | | | | | | | | | 0 | |
| 649 | 1 | | | | | | | | | 0 | 772 |
| 648 | 2 | | | | | | | | | 0 | 614 |
| 648 | 1 | | | | | | | | | 0 | |
| 648 | 1 | | | | | | | | | 0 | |
| 646 | 1 | | | | | | | | | 0 | 650 |
| 646 | 1 | | | | | | | | | 0 | |
| 646 | 1 | | | | | | | | | 0 | |
| 646 | 1 | | | | | | | | | 0 | |
| 645 | 1 | | | | | | | | | 0 | 838 |
| 645 | 1 | | | | | | | | | 0 | |
| 645 | 1 | | | | | | | | | 0 | |
| 642 | 1 | | | | | | | | | 0 | |
| 641 | 1 | | | | | | | | | 0 | |
| 641 | 1 | | | | | | | | | 0 | 1037 |
| 641 | 1 | | | | | | | | | 0 | |

| | | | | | | | | | |
|---|---|---|---|---|---|---|---|---|---|
| 641 | 1 | | | | | | | 0 | |
| 640 | 1 | | | | | | | 0 | |
| 640 | 1 | | | | | | | 0 | |
| 639 | 1 | | | | | | | 0 | |
| 638 | 1 | | | | | | | 0 | |
| 637 | 1 | | | | | | | 0 | |
| 627 | 1 | | | | | | | 0 | 1070 |
| 624 | 1 | | | | | | | 0 | |
| 624 | 1 | | | | | | | 0 | |
| 623 | 1 | | | | | | | 0 | |
| 623 | 1 | | | | | | | 0 | |
| 621 | 1 | | | | | | | 0 | 689 |
| 620 | 1 | | | | | | | 0 | |
| 617 | 1 | | | | | | | 0 | |
| 617 | 1 | membrane | Pf06388 | 5574348 | AAEL0023 | AaeL_AAE | 2 | 0 | 504 |
| 617 | 1 | | | | | | | 0 | 1112 |
| 615 | 1 | | Pf03638 | 11068077 | AAEL0058 | LOC110680779 | | 0 | 1472 |
| 613 | 1 | | | | | | | 0 | 861 |
| 613 | 1 | | | | | | | 0 | |
| 606 | 1 | catalytic a | Pf05497 | 5570920 | AAEL0014 | AaeL_AAE | 2 | 0 | 1507 |
| 604 | 1 | | | | | | | 0 | |
| 600 | 1 | | | | | | | 0 | |
| 598 | 1 | | Pf06840 | 5567961 | AAEL0064 | AaeL_AAE | 2 | 0 | |
| 595 | 1 | | | | | | | 0 | |
| 595 | 1 | | | | | | | 0 | 619 |
| 593 | 1 | metal ion | Pf05207 | 5579106 | AAEL0038 | AaeL_AAE | 3 | 0 | 913 |
| 592 | 1 | | | | | | | 0 | |
| 590 | 1 | | | | | | | 0 | |
| 590 | 1 | | | | | | | 0 | |
| 589 | 1 | | | | | | | 0 | |
| 588 | 1 | | Pf11029 | 5565550 | AAEL0048 | AaeL_AAE | 2 | 0 | |

| | | | | | | | | | |
|---|---|---|---|---|---|---|---|---|---|
| 587 | 1 | | | | | | | 0 | |
| 586 | 1 | | | | | | | 0 | |
| 585 | 1 | | | | | | | 0 | |
| 585 | 1 | | | | | | | 0 | |
| 585 | 1 | protein bi | Pf00627 | 5575807 | AAEL0120 | AaeL_AAE | 3 | 0 | |
| 583 | 1 | | | | | | | 0 | |
| 583 | 1 | | | | | | | 0 | 611 |
| 583 | 1 | | | | | | | 0 | |
| 582 | 1 | | | | | | | 0 | 927 |
| 580 | 1 | | Pf06911 | 5578005 | AAEL0183 | AaeL_AAE | 3 | 0 | |
| 579 | 1 | | | | | | | 0 | |
| 579 | 1 | structural | Pf00379 | 5577610 | AAEL0032 | AaeL_AAE | 2 | 0 | |
| 576 | 1 | | | | | | | 0 | |
| 575 | 1 | | | | | | | 0 | |
| 573 | 1 | | | | | | | 0 | |
| 572 | 1 | | | | | | | 0 | |
| 571 | 1 | | | | | | | 0 | 930 |
| 570 | 1 | | | | | | | 0 | |
| 570 | 1 | | | | | | | 0 | |
| 570 | 1 | | | | | | | 0 | |
| 570 | 1 | | | | | | | 0 | 920 |
| 569 | 1 | | | | | | | 0 | 630 |
| 568 | 1 | | | | | | | 0 | |
| 565 | 1 | | | | | | | 0 | |
| 565 | 1 | | | | | | | 0 | |
| 561 | 1 | | | | | | | 0 | |
| 560 | 1 | | | | | | | 0 | 748 |
| 559 | 1 | | | | | | | 0 | 1039 |
| 558 | 1 | | | | | | | 0 | |
| 558 | 1 | | | | | | | 0 | 407 |
| 557 | 1 | | | | | | | 0 | |

| | | | | | | | | | | |
|---|---|---|---|---|---|---|---|---|---|---|
| 556 | 1 | | | | | | | | 0 | |
| 556 | 1 | | | | | | | | 0 | |
| 553 | 1 | | | | | | | | 0 | 856 |
| 553 | 1 | | | | | | | | 0 | |
| 550 | 1 | ribosome | | Pf14978 | 5574439 | AAEL0111 | AaeL_AAE | 3 | 0 | |
| 548 | 1 | | | | | | | | 0 | 720 |
| 546 | 1 | | | | | | | | 0 | |
| 546 | 1 | | | | | | | | 0 | |
| 545 | 1 | | | | | | | | 0 | |
| 544 | 1 | | | | | | | | 0 | |
| 543 | 1 | | | | | | | | 0 | |
| 543 | 1 | | | | | | | | 0 | |
| 543 | 1 | | | | | | | | 0 | |
| 541 | 1 | | | | | | | | 0 | 774 |
| 540 | 1 | | | | | | | | 0 | |
| 540 | 1 | | | | | | | | 0 | |
| 540 | 1 | | | | | | | | 0 | |
| 540 | 1 | | | | | | | | 0 | |
| 539 | 1 | membrane | | Pf06522 | 5567396 | AAEL0060 | AaeL_AAE | 3 | 0 | |
| 538 | 1 | | | | | | | | 0 | |
| 537 | 1 | | | | | | | | 0 | 1048 |
| 536 | 1 | | | | | | | | 0 | 1145 |
| 535 | 1 | | | | | | | | 0 | |
| 535 | 1 | | | | | | | | 0 | |
| 535 | 1 | | | | | | | | 0 | |
| 534 | 1 | | | | | | | | 0 | 882 |
| 534 | 1 | | | | | | | | 0 | |
| 533 | 1 | | catalytic a | Pf00583, I | 5576921 | AAEL0128 | AaeL_AAE | 1 | 0 | |
| 533 | 1 | | | | | | | | 0 | |
| 533 | 1 | | | | | | | | 0 | 686 |
| 532 | 1 | | | | | | | | 0 | |

| | | | | | | | | | | |
|---|---|---|---|---|---|---|---|---|---|---|
| 531 | 1 | | | | | | | | 0 | |
| 531 | 1 | | | | | | | | 0 | |
| 529 | 1 | | | | | | | | 0 | |
| 527 | 1 | | | | | | | | 0 | |
| 526 | 1 | | | | | | | | 0 | |
| 525 | 1 | | | | | | | | 0 | 794 |
| 525 | 1 | | | | | | | | 0 | 525 |
| 524 | 1 | | | | | | | | 0 | |
| 524 | 1 | | | | | | | | 0 | |
| 524 | 1 | | | | | | | | 0 | |
| 523 | 1 | | | | | | | | 0 | |
| 523 | 1 | | | | | | | | 0 | |
| 522 | 1 | | | | | | | | 0 | |
| 522 | 1 | | | | | | | | 0 | |
| 522 | 1 | | | | | | | | 0 | |
| 521 | 1 | | | | | | | | 0 | |
| 521 | 1 | | | | | | | | 0 | 1670 |
| 520 | 1 | protein binding | | 5579066 | AAEL0140 | AaeL_AAE | 2 | mTOR signaling pathway | 1 | 766 |
| 520 | 1 | | | | | | | | 0 | |
| 520 | 1 | | Pf01636, I | 5576651 | AAEL0126 | AaeL_AAE | 3 | | 0 | |
| 520 | 1 | | | | | | | | 0 | |
| 519 | 1 | | | | | | | | 0 | 755 |
| 519 | 1 | | | | | | | | 0 | 1032 |
| 518 | 1 | | | | | | | | 0 | 873 |
| 518 | 1 | | | | | | | | 0 | |
| 517 | 1 | | | | | | | | 0 | |
| 517 | 1 | | | 5568425 | AAEL0068 | AaeL_AAE | 3 | | 0 | |
| 517 | 1 | | | | | | | | 0 | |
| 517 | 1 | | | | | | | | 0 | |
| 516 | 1 | | | | | | | | 0 | |
| 516 | 1 | | | | | | | | 0 | |

| | | | | | | | | | | | |
|---|---|---|---|---|---|---|---|---|---|---|---|
| 515 | 1 | transport | | Pf04100, I | 5578033 | AAEL0134 | AaeL_AAE | 2 | | 0 | |
| 515 | 1 | | | | | | | | | 0 | 765 |
| 515 | 1 | | | | | | | | | 0 | |
| 515 | 1 | | | | | | | | | 0 | 633 |
| 515 | 1 | | | | | | | | | 0 | |
| 515 | 1 | | | | | | | | | 0 | 1292 |
| 514 | 1 | | | | | | | | | 0 | |
| 514 | 1 | | | | | | | | | 0 | |
| 513 | 1 | | | | | | | | | 0 | |
| 512 | 1 | | | | | | | | | 0 | 525 |
| 512 | 1 | | | | | | | | | 0 | |
| 512 | 1 | | | | | | | | | 0 | 928 |
| 511 | 1 | metabolic process | catalytic a | Pf02902 | 5566543; | AAEL0054 | AaeL_AAE | 3 | | 0 | |
| 510 | 1 | | | | | | | | | 0 | 1052 |
| 510 | 1 | | | | | | | | | 0 | 718 |
| 510 | 1 | | | | | | | | | 0 | |
| 510 | 1 | | | | | | | | | 0 | |
| 509 | 1 | | | | | | | | | 0 | |
| 509 | 1 | | | | | | | | | 0 | |
| 509 | 1 | | | | | | | | | 0 | |
| 508 | 1 | | | | | | | | | 0 | |
| 508 | 1 | | | | | | | | | 0 | |
| 508 | 1 | | | | | | | | | 0 | 1014 |
| 508 | 1 | | | | | | | | | 0 | |
| 507 | 1 | | | | | | | | | 0 | 1333 |
| 507 | 1 | | | Pf11977 | 5571291 | AAEL0015 | AaeL_AAE | 2 | | 0 | |
| 506 | 1 | | protein bi | Pf00646, I | 5574961 | AAEL0115 | AaeL_AAE | 3 | | 0 | |
| 505 | 1 | | | | | | | | | 0 | |
| 504 | 1 | metabolic process | catalytic a | Pf01876 | 5573941 | AAEL0022 | AaeL_AAE | 2 | Ribosome biogenesis in eukary | 2 | 763 |
| 503 | 1 | | | | | | | | | 0 | |
| 503 | 1 | | | | | | | | | 0 | |

| | | | | | | | | | |
|---|---|---|---|---|---|---|---|---|---|
| 502 | 1 | | | | | | | 0 | |
| 502 | 1 | | | | | | | 0 | |
| 501 | 1 | | | 5580029 | AAEL0096 | AaeL_AAE | 2 | 0 | |
| 501 | 1 | | | | | | | 0 | |
| 501 | 1 | | | | | | | 0 | 610 |
| 501 | 1 | | | | | | | 0 | |
| 501 | 1 | | | | | | | 0 | 864 |
| 501 | 1 | | | | | | | 0 | 822 |
| 500 | 1 | | | | | | | 0 | |
| 499 | 1 | | | | | | | 0 | |
| 499 | 1 | | | | | | | 0 | |
| 499 | 1 | | | | | | | 0 | |
| 499 | 1 | | | | | | | 0 | 820 |
| 498 | 1 | | | | | | | 0 | 979 |
| 498 | 1 | | | | | | | 0 | |
| 498 | 1 | | | | | | | 0 | |
| 498 | 1 | | | | | | | 0 | |
| 498 | 1 | | | | | | | 0 | |
| 498 | 1 | | | | | | | 0 | 818 |
| 497 | 1 | | | | | | | 0 | |
| 497 | 1 | | | | | | | 0 | 1052 |
| 497 | 1 | | | | | | | 0 | |
| 496 | 1 | | | | | | | 0 | 659 |
| 496 | 1 | | | | | | | 0 | |
| 495 | 1 | | | | | | | 0 | |
| 495 | 1 | | | | | | | 0 | |
| 495 | 1 | | | | | | | 0 | |
| 495 | 1 | | | | | | | 0 | |
| 495 | 1 | catalytic a | Pf00583 | 5570879 | AAEL0086 | AaeL_AAE | 3 | 0 | 697 |
| 495 | 1 | | | 5575710 | AAEL0120 | AaeL_AAE | 2 | 0 | |
| 494 | 1 | | | | | | | 0 | |

| | | | | | | | | | | |
|---|---|---|---|---|---|---|---|---|---|---|
| 494 | 1 | | | | | | | | 0 | 928 |
| 493 | 1 | | | | | | | | 0 | 898 |
| 493 | 1 | | | | | | | | 0 | |
| 492 | 1 | | | | | | | | 0 | 1444 |
| 492 | 1 | | | | | | | | 0 | |
| 491 | 1 | | | | | | | | 0 | |
| 491 | 1 | | | | | | | | 0 | |
| 491 | 1 | | | | | | | | 0 | |
| 491 | 1 | | | | | | | | 0 | 338 |
| 491 | 1 | | | | | | | | 0 | |
| 491 | 1 | | | | | | | | 0 | |
| 491 | 1 | | | | 5567749 | AAEL0001 | AaeL_AAE | 2 | 0 | |
| 490 | 1 | | | | | | | | 0 | |
| 490 | 1 | | | | | | | | 0 | |
| 490 | 1 | regulation of biological process | | | 5569180 | AAEL0074 | AaeL_AAE | 2 | 0 | 916 |
| 490 | 1 | | | | | | | | 0 | |
| 490 | 1 | | | | | | | | 0 | 718 |
| 490 | 1 | | | | | | | | 0 | |
| 490 | 1 | | | | | | | | 0 | 826 |
| 490 | 1 | | | | | | | | 0 | 1158 |
| 489 | 1 | | | | | | | | 0 | |
| 489 | 1 | | | | | | | | 0 | |
| 489 | 1 | | | | | | | | 0 | |
| 489 | 1 | | | | | | | | 0 | 552 |
| 489 | 1 | | | | | | | | 0 | 582 |
| 488 | 1 | | | | | | | | 0 | |
| 488 | 1 | | | | | | | | 0 | |
| 488 | 1 | | membrane | Pf10270 | 5570695 | AAEL0084 | AaeL_AAE | 2 | 0 | 579 |
| 488 | 1 | | | | | | | | 0 | |
| 487 | 1 | | | | | | | | 0 | |
| 487 | 1 | | | | | | | | 0 | |

| | | | | | | | | | | |
|---|---|---|---|---|---|---|---|---|---|---|
| 487 | 1 | | | | | | | | 0 | |
| 486 | 1 | | | | | | | | 0 | |
| 486 | 1 | | | | | | | | 0 | |
| 486 | 1 | | | | | | | | 0 | |
| 486 | 1 | | | | | | | | 0 | |
| 486 | 1 | | | | | | | | 0 | |
| 486 | 1 | | | | | | | | 0 | |
| 485 | 1 | | | | | | | | 0 | |
| 485 | 1 | | | | | | | | 0 | |
| 485 | 1 | | | | | | | | 0 | |
| 484 | 1 | | | | | | | | 0 | |
| 484 | 1 | cell organ | membrane;mitochor | Pf15061 | 5574196 | AAEL0022 | AaeL_AAE | 3 | 0 | |
| 484 | 1 | | | Pf08216 | 5578457 | AAEL0004 | AaeL_AAE | 3 | 0 | |
| 484 | 1 | | | | | | | | 0 | |
| 484 | 1 | | | | | | | | 0 | |
| 484 | 1 | | | | | | | | 0 | 707 |
| 484 | 1 | | | | | | | | 0 | 638 |
| 484 | 1 | | | | | | | | 0 | 802 |
| 484 | 1 | | | | | | | | 0 | |
| 483 | 1 | | | | | | | | 0 | 303 |
| 483 | 1 | | | | | | | | 0 | 556 |
| 483 | 1 | | | | | | | | 0 | 670 |
| 482 | 1 | | | | | | | | 0 | 564 |
| 482 | 1 | | | | | | | | 0 | |
| 482 | 1 | | | | | | | | 0 | 528 |
| 482 | 1 | | DNA bindi | Pf05485 | 5575165 | AAEL0003 | AaeL_AAE | 2 | 0 | |
| 482 | 1 | | | | | | | | 0 | 377 |
| 481 | 1 | | | | | | | | 0 | 1392 |
| 481 | 1 | | | | | | | | 0 | |
| 481 | 1 | | | | | | | | 0 | |
| 481 | 1 | | | | | | | | 0 | 911 |

| | | | | | | | | | |
|---|---|---|---|---|---|---|---|---|---|
| 481 | 1 | | | | | | | 0 | 1338 |
| 481 | 1 | | | | | | | 0 | |
| 481 | 1 | | | | | | | 0 | 709 |
| 481 | 1 | | | | | | | 0 | |
| 481 | 1 | | Pf07162 | 5567753 | AAEL0001 | AaeL_AAE | 2 | 0 | |
| 480 | 1 | | | | | | | 0 | 1142 |
| 480 | 1 | metal ion | Pf04438 | 5575099 | AAEL0116 | AaeL_AAE | 3 | 0 | |
| 480 | 1 | | | | | | | 0 | |
| 480 | 1 | | | | | | | 0 | 1097 |
| 480 | 1 | | | | | | | 0 | |
| 480 | 1 | | | | | | | 0 | |
| 480 | 1 | | | | | | | 0 | |
| 480 | 1 | | | | | | | 0 | |
| 480 | 1 | | | | | | | 0 | |
| 480 | 1 | | | | | | | 0 | |
| 479 | 1 | | | | | | | 0 | |
| 479 | 1 | | Pf12752, I | 5569309 | AAEL0075 | AaeL_AAE | 2 | 0 | |
| 479 | 1 | | | | | | | 0 | |
| 478 | 1 | | | | | | | 0 | |
| 478 | 1 | | | | | | | 0 | |
| 478 | 1 | | | | | | | 0 | |
| 478 | 1 | | | | | | | 0 | |
| 478 | 1 | | | | | | | 0 | 857 |
| 478 | 1 | | | | | | | 0 | |
| 477 | 1 | | | | | | | 0 | |
| 476 | 1 | | | | | | | 0 | |
| 474 | 1 | | | | | | | 0 | 840 |
| 473 | 1 | | | | | | | 0 | |
| 471 | 1 | | | | | | | 0 | |
| 468 | 1 | | | | | | | 0 | 1096 |
| 468 | 1 | | | | | | | 0 | 525 |

| | | | |
|---|---|---|---|
| 468 | 1 | 0 | |
| 467 | 1 | 0 | |
| 466 | 1 | 0 | |
| 465 | 1 | 0 | 852 |
| 464 | 1 | 0 | 973 |
| 463 | 1 | 0 | 911 |
| 462 | 1 | 0 | 805 |
| 461 | 1 | 0 | |
| 460 | 1 | 0 | |
| 460 | 1 | 0 | 583 |
| 459 | 1 | 0 | |
| 459 | 1 | 0 | 1121 |
| 459 | 1 | 0 | 337 |
| 457 | 1 | 0 | 1017 |
| 454 | 1 | 0 | |
| 453 | 1 | 0 | 642 |
| 453 | 1 | 0 | |
| 449 | 1 | 0 | |
| 448 | 1 | 0 | |
| 448 | 1 | 0 | 650 |
| 447 | 1 | 0 | |
| 446 | 1 | 0 | |
| 446 | 1 | 0 | |
| 445 | 1 | 0 | |
| 444 | 1 | 0 | 645 |
| 444 | 1 | 0 | |
| 443 | 1 | 0 | |
| 443 | 1 | 0 | |
| 441 | 1 | 0 | 743 |
| 441 | 1 | 0 | |
| 440 | 1 | 0 | 987 |

| | | | | | | | | | |
|---|---|---|---|---|---|---|---|---|---|
| 439 | 1 | | | | | | | 0 | |
| 438 | 1 | | | 5573072 | AAEL0102 | AaeL_AAE | 2 | 0 | |
| 438 | 1 | | | | | | | 0 | |
| 438 | 1 | | | | | | | 0 | 841 |
| 436 | 1 | | | | | | | 0 | 909 |
| 435 | 1 | | | | | | | 0 | 943 |
| 434 | 1 | | | | | | | 0 | |
| 433 | 1 | | | | | | | 0 | |
| 433 | 1 | | | | | | | 0 | |
| 432 | 1 | structural | Pf00379 | 5575766 | AAEL0027 | AaeL_AAE | 2 | 0 | |
| 432 | 1 | | | | | | | 0 | 752 |
| 431 | 1 | | | | | | | 0 | |
| 430 | 1 | | | | | | | 0 | |
| 430 | 1 | | | | | | | 0 | |
| 430 | 1 | | | | | | | 0 | |
| 428 | 1 | | | | | | | 0 | 916 |
| 428 | 1 | | | | | | | 0 | |
| 425 | 1 | | | | | | | 0 | |
| 424 | 1 | | | | | | | 0 | 743 |
| 424 | 1 | | | | | | | 0 | 948 |
| 423 | 1 | | | | | | | 0 | 590 |
| 421 | 1 | | | | | | | 0 | 376 |
| 419 | 1 | | | | | | | 0 | |
| 418 | 1 | | | | | | | 0 | 402 |
| 418 | 1 | | | | | | | 0 | 697 |
| 417 | 1 | | | | | | | 0 | 807 |
| 417 | 1 | | | | | | | 0 | |
| 416 | 1 | | | | | | | 0 | |
| 414 | 1 | | | | | | | 0 | |
| 414 | 1 | | | | | | | 0 | |
| 414 | 1 | regulation of biological process | Pf11916, I | 5574743 | AAEL0113 | AaeL_AAE | 1 | 0 | |

| | | | | | | | | | | |
|---|---|---|---|---|---|---|---|---|---|---|
| 414 | 1 | membrane | | Pf04148 | 5569429 | AAEL0116 | AaeL_AAE | 3 | 0 | 616 |
| 411 | 1 | | | | | | | | 0 | |
| 411 | 1 | | | | | | | | 0 | |
| 410 | 1 | | | | | | | | 0 | |
| 409 | 1 | | | | | | | | 0 | |
| 408 | 1 | | | | 23687983 | AAEL0175 | AaeL_AAE | 2 | 0 | 808 |
| 408 | 1 | | | | | | | | 0 | |
| 407 | 1 | | | | | | | | 0 | |
| 407 | 1 | | | | | | | | 0 | |
| 406 | 1 | | | | | | | | 0 | 874 |
| 406 | 1 | | | | | | | | 0 | 769 |
| 406 | 1 | | | | | | | | 0 | 776 |
| 405 | 1 | regulation of biologi | catalytic a | Pf00071, I | 5564132 | AAEL0183 | AaeL_AAE | 3 | 0 | |
| 404 | 1 | | | | 5568393 | AAEL0067 | AaeL_AAE | 3 | 0 | 1221 |
| 404 | 1 | | | | | | | | 0 | |
| 404 | 1 | | | | | | | | 0 | 887 |
| 403 | 1 | | | | | | | | 0 | |
| 402 | 1 | | | | | | | | 0 | |
| 402 | 1 | | | | | | | | 0 | |
| 401 | 1 | | | | | | | | 0 | 736 |
| 401 | 1 | | | | | | | | 0 | |
| 401 | 1 | | | | | | | | 0 | |
| 399 | 1 | | | | | | | | 0 | |
| 398 | 1 | | | | | | | | 0 | |
| 397 | 1 | | | | | | | | 0 | |
| 396 | 1 | | protein bi | Pf00240, I | 5570654 | AAEL0084 | AaeL_AAE | 1 | 0 | 1342 |
| 396 | 1 | | | | | | | | 0 | 812 |
| 396 | 1 | | | | | | | | 0 | |
| 396 | 1 | | | | | | | | 0 | 987 |
| 395 | 1 | | | | | | | | 0 | 696 |
| 395 | 1 | | | | | | | | 0 | |

| | | | | | | | | | | |
|---|---|---|---|---|---|---|---|---|---|---|
| 394 | 1 | | | | | | | | 0 | |
| 393 | 1 | | | | | | | | 0 | |
| 393 | 1 | | metal ion | Pf00643 | 5564280 | AAEL0143 | AaeL_AAE | 3 | 0 | 1105 |
| 390 | 1 | | | | | | | | 0 | |
| 390 | 1 | | | | | | | | 0 | 1028 |
| 389 | 1 | | | | | | | | 0 | 826 |
| 388 | 1 | | | Pf00339, I | 5570228 | AAEL0081 | AaeL_AAE | 3 | 0 | 824 |
| 387 | 1 | | | | | | | | 0 | |
| 386 | 1 | | | | | | | | 0 | |
| 386 | 1 | | metal ion | Pf00643 | 5563854 | AAEL0141 | AaeL_AAE | 2 | 0 | |
| 386 | 1 | | | Pf00076, I | 5569476 | AAEL0076 | AaeL_AAE | 1 | 0 | |
| 385 | 1 | | | | | | | | 0 | 693 |
| 384 | 1 | | | | | | | | 0 | |
| 384 | 1 | | | | | | | | 0 | 768 |
| 384 | 1 | | | | | | | | 0 | |
| 382 | 1 | | | | | | | | 0 | 944 |
| 382 | 1 | | | | | | | | 0 | |
| 381 | 1 | | | | | | | | 0 | |
| 381 | 1 | | | | | | | | 0 | |
| 381 | 1 | | | | | | | | 0 | |
| 380 | 1 | | | | 5574346 | AAEL0023 | AaeL_AAE | 2 | 0 | |
| 380 | 1 | | | | | | | | 0 | |
| 379 | 1 | | | | | | | | 0 | |
| 379 | 1 | | | | | | | | 0 | |
| 379 | 1 | | | | | | | | 0 | |
| 379 | 1 | | | | | | | | 0 | |
| 378 | 1 | | | | | | | | 0 | |
| 378 | 1 | | | | | | | | 0 | 635 |
| 378 | 1 | | | | | | | | 0 | |
| 378 | 1 | membrane | | Pf09779 | 5566213 | AAEL0052 | AaeL_AAE | 1 | 0 | |
| 378 | 1 | | | | | | | | 0 | |

| | | | | | | | | |
|---|---|---|---|---|---|---|---|---|
| 377 | 1 | | | | | | 0 | |
| 377 | 1 | | 5575336 | AAEL0025 | AaeL_AAE | 3 | 0 | |
| 377 | 1 | | | | | | 0 | 635 |
| 376 | 1 | | | | | | 0 | |
| 376 | 1 | | | | | | 0 | 437 |
| 375 | 1 | | | | | | 0 | |
| 375 | 1 | | | | | | 0 | |
| 375 | 1 | | | | | | 0 | 700 |
| 374 | 1 | | | | | | 0 | |
| 373 | 1 | | | | | | 0 | |
| 373 | 1 | | | | | | 0 | 805 |
| 373 | 1 | | 5578152 | AAEL0135 | AaeL_AAE | 2 | 0 | |
| 372 | 1 | Pf04714 | 5576471 | AAEL0028 | AaeL_AAE | 1 | 0 | |
| 371 | 1 | | | | | | 0 | 686 |
| 371 | 1 | | | | | | 0 | |
| 371 | 1 | | | | | | 0 | 466 |
| 370 | 1 | | | | | | 0 | |
| 370 | 1 | Pf05347, I | 5567260 | AAEL0059 | AaeL_AAE | 3 | 0 | |
| 369 | 1 | | | | | | 0 | 669 |
| 369 | 1 | Pf00300 | 5568499 | AAEL0069 | AaeL_AAE | 3 | 0 | 660 |
| 369 | 1 | | | | | | 0 | |
| 369 | 1 | | | | | | 0 | |
| 368 | 1 | | | | | | 0 | |
| 368 | 1 | | | | | | 0 | |
| 368 | 1 | | | | | | 0 | |
| 368 | 1 | | | | | | 0 | |
| 367 | 1 | | | | | | 0 | 909 |
| 367 | 1 | | | | | | 0 | 943 |
| 367 | 1 | | | | | | 0 | |
| 366 | 1 | | | | | | 0 | |
| 366 | 1 | | | | | | 0 | |

| | | | | | | | | | | |
|---|---|---|---|---|---|---|---|---|---|---|
| 366 | 1 | | | | | | | | 0 | 755 |
| 366 | 1 | | | | | | | | 0 | 847 |
| 365 | 1 | | | | | | | | 0 | |
| 364 | 1 | | | | | | | | 0 | |
| 363 | 1 | | | | | | | | 0 | |
| 362 | 1 | | | | | | | | 0 | |
| 362 | 1 | | | | | | | | 0 | |
| 361 | 1 | | | | | | | | 0 | |
| 360 | 1 | | | | | | | | 0 | |
| 360 | 1 | | | | | | | | 0 | 807 |
| 360 | 1 | cell organization and biogenesi | | Pf11719, I | 5569648 | AAEL0078 | AaeL_AAE | 3 | 0 | |
| 359 | 1 | | | | | | | | 0 | |
| 359 | 1 | | | | | | | | 0 | |
| 359 | 1 | | | | | | | | 0 | |
| 358 | 1 | | | | | | | | 0 | |
| 357 | 1 | | | | | | | | 0 | 841 |
| 357 | 1 | | | | | | | | 0 | |
| 356 | 1 | | | | | | | | 0 | |
| 356 | 1 | | | Pf09731 | 5572703 | AAEL0099 | AaeL_AAE | 3 | 0 | |
| 355 | 1 | regulation of biologi | protein bi | Pf00627, I | 5569186 | AAEL0074 | AaeL_AAE | 2 | 0 | |
| 355 | 1 | | | | | | | | 0 | |
| 355 | 1 | | | | | | | | 0 | |
| 355 | 1 | | | | | | | | 0 | 802 |
| 355 | 1 | | | | | | | | 0 | |
| 355 | 1 | | | | | | | | 0 | |
| 354 | 1 | | | | | | | | 0 | |
| 354 | 1 | | | | | | | | 0 | |
| 354 | 1 | | | | | | | | 0 | |
| 354 | 1 | | | | | | | | 0 | |
| 354 | 1 | | | | | | | | 0 | |
| 352 | 1 | | | | | | | | 0 | |

| | | | | | | | | | | | |
|---|---|---|---|---|---|---|---|---|---|---|---|
| 352 | 1 | | | | | | | | | 0 | |
| 352 | 1 | | | | | | | | | 0 | 798 |
| 351 | 1 | | | | | | | | | 0 | |
| 351 | 1 | metabolic extracellular | | | 5576626 | AAEL0126 | AaeL_AAE | 3 | | 0 | |
| 351 | 1 | | | | | | | | | 0 | |
| 351 | 1 | | | | | | | | | 0 | |
| 350 | 1 | | | | | | | | | 0 | |
| 350 | 1 | | | | | | | | | 0 | |
| 350 | 1 | | | | | | | | | 0 | 1235 |
| 350 | 1 | | | | | | | | | 0 | |
| 350 | 1 | | | | | | | | | 0 | |
| 350 | 1 | | | | | | | | | 0 | 744 |
| 349 | 1 | | | | | | | | | 0 | |
| 349 | 1 | | | | | | | | | 0 | |
| 349 | 1 | | | | | | | | | 0 | |
| 349 | 1 | | | | | | | | | 0 | |
| 349 | 1 | | | | | | | | | 0 | |
| 348 | 1 | | | | | | | | | 0 | |
| 348 | 1 | | | | | | | | | 0 | |
| 348 | 1 | | | | | | | | | 0 | |
| 347 | 1 | | | | | | | | | 0 | 710 |
| 347 | 1 | metabolic process | catalytic a | Pf08241, I | 5576784 | AAEL0127 | AaeL_AAE | 3 | Glycine, serine and threonine n | 1 | |
| 346 | 1 | | | | | | | | | 0 | |
| 346 | 1 | | | | | | | | | 0 | |
| 345 | 1 | | | | | | | | | 0 | 1212 |
| 345 | 1 | | | | | | | | | 0 | 389 |
| 345 | 1 | | | | | | | | | 0 | |
| 344 | 1 | | | | | | | | | 0 | |
| 344 | 1 | | | | | | | | | 0 | 405 |
| 344 | 1 | | | | | | | | | 0 | 751 |
| 344 | 1 | | | | | | | | | 0 | |

| | | | | | | | | | | | |
|---|---|---|---|---|---|---|---|---|---|---|---|
| 344 | 1 | | | | | | | | | 0 | 813 |
| 344 | 1 | | | | | | | | | 0 | |
| 344 | 1 | | | | | | | | | 0 | |
| 343 | 1 | | | | | | | | | 0 | |
| 343 | 1 | | | | | | | | | 0 | 808 |
| 343 | 1 | | | | | | | | | 0 | |
| 343 | 1 | | | Pf03660 | 5578758 | AAEL0036 | AaeL_AAE | 2 | Spliceosome | 1 | |
| 342 | 1 | | | | | | | | | 0 | |
| 342 | 1 | | | | | | | | | 0 | |
| 341 | 1 | | protein bi | Pf05773 | 5568224 | AAEL0066 | AaeL_AAE | 2 | | 0 | |
| 341 | 1 | | | | | | | | | 0 | |
| 341 | 1 | | | | | | | | | 0 | |
| 341 | 1 | | | | | | | | | 0 | 770 |
| 340 | 1 | | | | | | | | | 0 | |
| 340 | 1 | | | | | | | | | 0 | |
| 340 | 1 | | | | | | | | | 0 | |
| 339 | 1 | | | | | | | | | 0 | |
| 339 | 1 | cellular homeostasis | protein bi | Pf01436, I | 5567489 | AAEL0009 | AaeL_AAE | 3 | | 0 | 645 |
| 339 | 1 | | | | | | | | | 0 | 619 |
| 338 | 1 | | | | | | | | | 0 | |
| 338 | 1 | | | | | | | | | 0 | |
| 338 | 1 | | | | | | | | | 0 | 462 |
| 337 | 1 | | | | | | | | | 0 | |
| 337 | 1 | | | | | | | | | 0 | |
| 336 | 1 | | | | | | | | | 0 | |
| 335 | 1 | | | | | | | | | 0 | |
| 335 | 1 | | | | | | | | | 0 | 544 |
| 334 | 1 | | | | | | | | | 0 | |
| 334 | 1 | | | | | | | | | 0 | |
| 334 | 1 | | | | | | | | | 0 | |
| 334 | 1 | | | | | | | | | 0 | |

| | | | |
|---|---|---|---|
| 334 | 1 | 0 | |
| 333 | 1 | 0 | 270 |
| 333 | 1 | 0 | |
| 333 | 1 | 0 | |
| 333 | 1 | 0 | |
| 333 | 1 | 0 | |
| 333 | 1 | 0 | |
| 332 | 1 | 0 | 901 |
| 332 | 1 | 0 | |
| 332 | 1 | 0 | 685 |
| 332 | 1 | 0 | |
| 332 | 1 | 0 | |
| 332 | 1 | 0 | |
| 331 | 1 | 0 | 613 |
| 331 | 1 | 0 | |
| 331 | 1 | 0 | |
| 331 | 1 | 0 | |
| 331 | 1 | 0 | |
| 331 | 1 | 0 | |
| 331 | 1 | 0 | |
| 331 | 1 | 0 | 814 |
| 330 | 1 | 0 | |
| 330 | 1 | 0 | |
| 330 | 1 | 0 | |
| 330 | 1 | 0 | |
| 329 | 1 | 0 | 801 |
| 329 | 1 | 0 | |
| 329 | 1 | 0 | 1079 |
| 329 | 1 | 0 | 889 |
| 329 | 1 | 0 | |
| 329 | 1 | 0 | |

| | | | | | | | | | | | | |
|---|---|---|---|---|---|---|---|---|---|---|---|---|
| 329 | 1 | | | | | | | | | | 0 | 565 |
| 329 | 1 | | | | | | | | | | 0 | |
| 329 | 1 | | | | | | | | | | 0 | 1154 |
| 329 | 1 | | | | | | | | | | 0 | 609 |
| 328 | 1 | | | | | | | | | | 0 | |
| 328 | 1 | | | | | | | | | | 0 | |
| 328 | 1 | | | | | | | | | | 0 | 779 |
| 328 | 1 | | | | | | | | | | 0 | 966 |
| 328 | 1 | | | | | | | | | | 0 | 278 |
| 328 | 1 | metabolic process | | catalytic a | Pf05132 | 5571078 | AAEL0087 | AaeL_AAE | 3 | Purine metabolism; RNA polym | 4 | 834 |
| 328 | 1 | | | protein bi | Pf00856 | 5577326 | AAEL0004 | AaeL_AAE | 3 | | 0 | |
| 327 | 1 | | | | | | | | | | 0 | |
| 327 | 1 | | | | | | | | | | 0 | |
| 327 | 1 | | | | | | | | | | 0 | 703 |
| 327 | 1 | | | | | | | | | | 0 | |
| 327 | 1 | | membrane | | Pf00226 | 5574141 | AAEL0109 | AaeL_AAE | 3 | | 0 | |
| 326 | 1 | | | | | | | | | | 0 | |
| 326 | 1 | | | | | | | | | | 0 | |
| 326 | 1 | | | | | | | | | | 0 | |
| 326 | 1 | | | | | | | | | | 0 | |
| 326 | 1 | | | | | | | | | | 0 | 639 |
| 326 | 1 | | | | | | | | | | 0 | |
| 326 | 1 | | | | | | | | | | 0 | |
| 325 | 1 | | | | | | | | | | 0 | |
| 325 | 1 | | | | | | | | | | 0 | |
| 325 | 1 | | | | | | | | | | 0 | |
| 325 | 1 | | | | | | | | | | 0 | |
| 325 | 1 | | | | | | | | | | 0 | |
| 325 | 1 | | | | | | | | | | 0 | 663 |
| 325 | 1 | | | catalytic a | Pf00583, I | 5568231 | AAEL0066 | AaeL_AAE | 2 | | 0 | |
| 324 | 1 | | | | | | | | | | 0 | 858 |

| | | | | | | | | | | |
|---|---|---|---|---|---|---|---|---|---|---|
| 324 | 1 | | | | | | | | 0 | |
| 324 | 1 | | | | | | | | 0 | |
| 324 | 1 | | | | | | | | 0 | |
| 324 | 1 | | | | | | | | 0 | |
| 324 | 1 | | | Pf12371 | 5567315 | AAEL0059 | AaeL_AAE | 2 | 0 | |
| 324 | 1 | | RNA bindi | Pf00013, I | 5566377 | AAEL0053 | AaeL_AAE | 3 | 0 | |
| 323 | 1 | metabolic process | | Pf02033 | 5578294 | AAEL0136 | AaeL_AAE | 3 | 0 | |
| 323 | 1 | metabolic extracellular | | Pf01607 | 5576627 | AAEL0126 | AaeL_AAE | 3 | 0 | |
| 323 | 1 | cell organization and biogenesi | | Pf07542 | 5572922 | AAEL0101 | AaeL_AAE | 2 | 0 | |
| 323 | 1 | | | | | | | | 0 | |
| 323 | 1 | | metal ion | Pf13815 | 11067983 | AAEL0180 | LOC11067 | 3 | 0 | |
| 323 | 1 | | | | | | | | 0 | |
| 322 | 1 | | | | | | | | 0 | |
| 322 | 1 | | | | | | | | 0 | |
| 322 | 1 | | | | | | | | 0 | 586 |
| 322 | 1 | | | | | | | | 0 | |
| 322 | 1 | | | | | | | | 0 | 644 |
| 322 | 1 | | | | | | | | 0 | 601 |
| 321 | 1 | | | | | | | | 0 | |
| 321 | 1 | | | | 23687752 | AAEL0173 | AaeL_AAE | 3 | 0 | |
| 321 | 1 | | | | | | | | 0 | |
| 321 | 1 | | | | | | | | 0 | |
| 321 | 1 | | | | | | | | 0 | |
| 321 | 1 | | | | | | | | 0 | |
| 321 | 1 | | | | | | | | 0 | |
| 321 | 1 | | | | | | | | 0 | |
| 321 | 1 | | | | | | | | 0 | |
| 321 | 1 | | | | | | | | 0 | 551 |
| 321 | 1 | | | | | | | | 0 | |
| 320 | 1 | | | | | | | | 0 | 861 |
| 320 | 1 | | | | | | | | 0 | |

| | | | | | | | | |
|---|---|---|---|---|---|---|---|---|
| 320 | 1 | | | | | | | 0 |
| 320 | 1 | | | | | | | 0 |
| 320 | 1 | | | | | | | 0 |
| 320 | 1 | | | | | | | 0 |
| 320 | 1 | | | | | | | 0 |
| 320 | 1 | | | | | | | 0 |
| 320 | 1 | membrane | Pf03647, I | 5568715 | AAEL0070 | AaeL_AAE | 3 | 0 |

| Abundanc | Abundanc | Abundanc | Abundanc | Abundanc | Abundanc | Abundanc | Abundanc | Abundanc | Abundanc | Abundanc | Abundanc | Abundanc | Abundanc | Abundanc |
|---|---|---|---|---|---|---|---|---|---|---|---|---|---|---|
| 455 | 754 | 553 | 450 | 744 | 550 | 447 | 749 | 1878776 | 1561008 | 3430753 | 2587307 | 2008930 | 1632349 | 3631399 |
| 413 | 568 | 539 | 421 | 550 | 548 | 416 | 560 | 1398156 | 1066936 | 2583306 | 1466986 | 1413952 | 1104533 | 2623091 |
| 610 | 1030 | 538 | 577 | 992 | 555 | 591 | 1022 | 1841657 | 1970594 | 3232696 | 3328676 | 1746177 | 1871860 | 3243593 |
| 609 | 832 | 786 | 609 | 824 | 790 | 615 | 818 | 274289 | 215734 | 354293 | 294820 | 229823 | 178054 | 292287 |
| 544 | 845 | 669 | 556 | 862 | 648 | 538 | 844 | 441226 | 366572 | 673811 | 569484 | 455543 | 378144 | 680508 |
| 523 | 841 | 1221 | 559 | 850 | 1238 | 519 | 825 | 3825 | 1586 | 3031 | 2549 | 5980 | 2738 | 4897 |
|  |  | 1135 | 780 | 1059 | 962 | 715 | 962 |  |  |  |  | 1706 | 1172 | 1503 |
| 635 | 910 | 913 | 656 | 898 | 903 | 651 | 918 | 118897 | 83369 | 131304 | 119471 | 128022 | 92003 | 140248 |
| 587 | 950 | 782 | 632 | 906 | 816 | 673 | 915 | 47729 | 39616 | 67457 | 64110 | 51024 | 41222 | 65217 |
| 687 | 1035 | 922 | 680 | 1033 | 874 | 639 | 1015 | 524177 | 389439 | 566860 | 586970 | 474851 | 350561 | 515174 |
| 697 | 3476 | 1175 | 927 | 4254 | 986 | 765 | 2536 | 10313 | 7180 | 10307 | 35827 | 12352 | 9743 | 10513 |
| 855 | 1267 | 948 | 853 | 1264 | 926 | 858 | 1286 | 275531 | 253069 | 296057 | 375139 | 347847 | 313306 | 367101 |
|  |  |  |  |  | 1167 | 869 | 1662 |  |  |  |  |  |  |  |
| 531 | 881 | 786 | 569 | 896 | 768 | 559 | 911 | 428200 | 306658 | 577858 | 509293 | 445837 | 322842 | 567373 |
| 816 | 1139 | 859 | 787 | 1168 | 925 | 855 | 1174 | 269166 | 248211 | 304334 | 346654 | 257994 | 236416 | 300279 |
| 574 | 858 | 718 | 574 | 853 | 662 | 526 | 830 | 146984 | 119509 | 208147 | 178656 | 140878 | 112578 | 196268 |
| 668 | 989 | 857 | 669 | 995 | 820 | 637 | 966 | 90323 | 68834 | 103031 | 101906 | 75870 | 59227 | 88562 |
| 546 | 816 | 842 | 527 | 791 | 854 | 510 | 796 | 258964 | 165376 | 302916 | 247109 | 287627 | 179890 | 341504 |
|  |  | 999 | 772 | 1063 | 772 | 476 | 628 |  |  |  |  | 3491 | 2700 | 3495 |
| 955 | 1339 | 963 | 1051 | 1402 | 849 | 857 | 1305 | 192595 | 195928 | 205117 | 274578 | 242675 | 264832 | 252088 |
| 728 | 1187 | 902 | 760 | 1209 | 881 | 742 | 1196 | 226704 | 186403 | 256025 | 303842 | 182434 | 153778 | 202289 |
| 695 | 1023 | 940 | 674 | 1009 | 938 | 665 | 1022 | 294422 | 211771 | 304570 | 311454 | 320057 | 229329 | 340457 |
| 903 | 5774 | 991 | 709 | 2416 | 1089 | 762 | 4194 | 18780 | 13984 | 15488 | 89428 | 33850 | 24224 | 34162 |
| 424 | 778 | 718 | 464 | 798 | 698 | 446 | 778 | 270965 | 164453 | 388251 | 302051 | 295778 | 191270 | 412190 |
| 506 | 849 | 846 | 467 | 856 | 839 | 478 | 841 | 290945 | 169806 | 335879 | 285208 | 358450 | 197966 | 423518 |
| 548 | 888 | 731 | 540 | 865 | 726 | 533 | 879 | 281159 | 207798 | 379112 | 336473 | 343367 | 253825 | 469779 |
| 626 | 1009 | 714 | 588 | 980 | 759 | 652 | 1010 | 129139 | 108977 | 174130 | 175710 | 130462 | 107433 | 182825 |
| 279 | 761 | 281 | 376 | 797 | 341 | 371 | 793 | 2551 | 3962 | 14179 | 10794 | 3655 | 4891 | 13021 |
| 685 | 1058 | 823 | 634 | 1048 | 849 | 661 | 978 | 239398 | 193523 | 282719 | 299004 | 166291 | 128146 | 202118 |
| 806 | 910 | 976 | 892 | 1334 | 1050 | 966 | 1264 | 1234 | 1060 | 1315 | 1197 | 1051 | 960 | 1077 |
| 674 | 1024 | 841 | 694 | 1035 | 852 | 697 | 1043 | 210985 | 172917 | 256728 | 262864 | 237475 | 196133 | 282467 |

| | | | | | | | | | | | | | | |
|---|---|---|---|---|---|---|---|---|---|---|---|---|---|---|
| 850 | 1017 | | | | | | | 916 | 788 | 927 | 943 | | | |
| 704 | 1130 | 799 | 676 | 1079 | 793 | 653 | 1077 | 229877 | 192477 | 273585 | 309265 | 209096 | 176918 | 261760 |
| 510 | 1037 | 600 | 505 | 1037 | 639 | 565 | 1048 | 212855 | 183685 | 360022 | 373454 | 208484 | 175734 | 347677 |
| 656 | 1010 | 848 | 617 | 973 | 830 | 608 | 991 | 271146 | 193989 | 295559 | 298600 | 231567 | 168535 | 273096 |
| 790 | 1336 | 872 | 708 | 1242 | 935 | 866 | 1352 | 227267 | 202035 | 255882 | 341774 | 238614 | 193727 | 273684 |
| 754 | 1172 | 806 | 720 | 1113 | 821 | 735 | 1162 | 237006 | 215276 | 285459 | 334447 | 206375 | 184542 | 256204 |
| 502 | 828 | 689 | 507 | 836 | 683 | 495 | 836 | 231165 | 169393 | 337618 | 279594 | 216895 | 159414 | 314715 |
| 530 | 946 | 740 | 583 | 940 | 741 | 589 | 967 | 202230 | 153879 | 290094 | 274370 | 185639 | 146424 | 250997 |
| 630 | 1003 | 645 | 596 | 1027 | 686 | 639 | 1038 | 140105 | 135300 | 214720 | 215316 | 149185 | 137833 | 231271 |
| 759 | 1242 | 770 | 709 | 1177 | 815 | 761 | 1241 | 191629 | 176387 | 232447 | 288764 | 159132 | 146643 | 206745 |
| 526 | 716 | 584 | 488 | 702 | 638 | 562 | 737 | 195162 | 172941 | 328694 | 235387 | 184328 | 154157 | 315736 |
| 611 | 1165 | 707 | 651 | 1171 | 720 | 657 | 1121 | 148952 | 131005 | 214393 | 249780 | 118990 | 109559 | 168371 |
| 767 | 1182 | 870 | 762 | 1163 | 909 | 805 | 1215 | 216243 | 188597 | 245860 | 290520 | 187830 | 164547 | 215944 |
| 688 | 899 | 1016 | 742 | 914 | 982 | 688 | 908 | 242176 | 159585 | 231991 | 208613 | 220439 | 161015 | 217049 |
| 486 | 791 | 702 | 504 | 809 | 737 | 526 | 825 | 177242 | 125474 | 258427 | 204410 | 199141 | 143020 | 283519 |
| 885 | 1146 | 929 | 898 | 1159 | 869 | 833 | 1123 | 173116 | 163618 | 184804 | 211747 | 189271 | 183009 | 203734 |
| 792 | 1277 | 833 | 789 | 1355 | 841 | 794 | 1322 | 190166 | 176884 | 223450 | 285309 | 172369 | 163397 | 207029 |
| 392 | 904 | 437 | 382 | 877 | 452 | 392 | 905 | 164786 | 141387 | 360998 | 326457 | 148112 | 129464 | 339160 |
| 539 | 883 | 701 | 504 | 880 | 672 | 477 | 866 | 254100 | 185299 | 344060 | 303666 | 219184 | 157660 | 312805 |
| 875 | 1301 | 852 | 917 | 1340 | 825 | 840 | 1312 | 171103 | 177190 | 202482 | 263394 | 171008 | 184208 | 200779 |
| 503 | 864 | 591 | 451 | 819 | 542 | 422 | 775 | 12962 | 10718 | 21311 | 18421 | 13467 | 10280 | 22772 |
| 923 | 1503 | 743 | 638 | 1096 | 880 | 1024 | 1595 | 12309 | 13346 | 14452 | 21727 | 13196 | 11328 | 17762 |
| 886 | 1173 | 1010 | 919 | 1220 | 983 | 891 | 1206 | 121287 | 108281 | 122175 | 143315 | 124442 | 113208 | 123239 |
| 803 | 1048 | 1061 | 879 | 1228 | | | | 2991 | 2391 | 2979 | 3122 | 5514 | 4569 | 5197 |
| 465 | 970 | | | | | | | 608 | 369 | 793 | 769 | | | |
| 851 | 1555 | 834 | 848 | 1594 | 812 | 784 | 1495 | 162384 | 167993 | 197423 | 307026 | 176661 | 179561 | 211856 |
| 669 | 1098 | 818 | 646 | 1030 | 829 | 662 | 1060 | 174318 | 139522 | 208410 | 228778 | 171094 | 135134 | 209201 |
| 424 | 875 | 598 | 408 | 845 | 683 | 467 | 861 | 177468 | 120320 | 283711 | 248198 | 211944 | 144629 | 354314 |
| 587 | 877 | 856 | 597 | 901 | 859 | 604 | 887 | 192720 | 132793 | 226411 | 198476 | 193214 | 134711 | 225762 |
| 699 | 898 | 705 | 661 | 880 | 786 | 742 | 930 | 171985 | 160398 | 229367 | 205866 | 218670 | 204907 | 310163 |
| 731 | 1091 | 912 | 741 | 1107 | 893 | 721 | 1065 | 160130 | 129974 | 177768 | 193985 | 122154 | 99315 | 133990 |

| | | | | | | | | | | | | | | |
|---|---|---|---|---|---|---|---|---|---|---|---|---|---|---|
| 720 | 1111 | 828 | 771 | 1161 | 766 | 683 | 1092 | 166987 | 152484 | 211845 | 235366 | 158461 | 147511 | 191373 |
| 774 | 1017 | 995 | 746 | 1011 | 994 | 735 | 995 | 183124 | 138463 | 179002 | 182038 | 222717 | 166996 | 223919 |
| 292 | 768 | 474 | 322 | 796 | 480 | 336 | 801 | 144867 | 93561 | 320245 | 245945 | 141855 | 96173 | 299018 |
| 613 | 1023 | 779 | 673 | 1043 | 792 | 676 | 1030 | 140795 | 114818 | 187406 | 191740 | 148879 | 128464 | 191017 |
| 877 | 1213 | 950 | 926 | 1252 | 949 | 905 | 1253 | 145895 | 143750 | 163873 | 198857 | 183350 | 178797 | 193003 |
| 616 | 989 | 735 | 591 | 974 | 761 | 623 | 998 | 169599 | 139176 | 225908 | 223360 | 161851 | 130065 | 220111 |
| 627 | 787 | 1434 | 615 | 813 | 1395 | 614 | 795 | 302303 | 133565 | 212933 | 167521 | 273401 | 117305 | 190721 |
| 640 | 1031 | 787 | 702 | 1059 | 851 | 766 | 1088 | 150328 | 128319 | 200373 | 206604 | 162057 | 144593 | 205932 |
| 592 | 905 | 691 | 541 | 876 | 719 | 564 | 882 | 137852 | 108148 | 182721 | 165275 | 175278 | 137270 | 253515 |
| 641 | 1134 | 791 | 627 | 1111 | 721 | 556 | 1077 | 185684 | 146323 | 228239 | 258851 | 159541 | 126487 | 201820 |
| 913 | 1483 | 976 | 985 | 1575 | 1036 | 1109 | 1646 | 132684 | 134834 | 147618 | 218991 | 118807 | 119936 | 121783 |
| 909 | 1322 | 929 | 911 | 1347 | 997 | 1000 | 1364 | 201171 | 192230 | 211537 | 279712 | 188869 | 185253 | 203366 |
| 1208 | 1339 | 1011 | 1096 | 1271 | 1008 | 1179 | 1268 | 126942 | 140814 | 116574 | 156062 | 137021 | 148509 | 135496 |
| 583 | 1154 | 641 | 575 | 1098 | 694 | 611 | 1154 | 115073 | 104290 | 178996 | 206540 | 117394 | 105283 | 183240 |
| 653 | 1001 | 875 | 653 | 996 | 825 | 588 | 978 | 145232 | 110899 | 169715 | 169901 | 120794 | 90089 | 138060 |
| 989 | 1471 | 933 | 964 | 1384 | 898 | 941 | 1411 | 100793 | 107532 | 108695 | 159912 | 130706 | 135113 | 140106 |
| 524 | 838 | 712 | 494 | 839 | 725 | 493 | 824 | 150889 | 109603 | 209249 | 175335 | 133869 | 92832 | 187931 |
| 638 | 1078 | 861 | 662 | 1110 | 901 | 679 | 1110 | 149726 | 111456 | 174601 | 188142 | 152995 | 117652 | 177699 |
| 649 | 1079 | 775 | 646 | 1190 | 858 | 748 | 1180 | 93402 | 73855 | 113758 | 122752 | 103510 | 86300 | 133585 |
| 740 | 1222 | 819 | 831 | 1265 | 736 | 751 | 1243 | 81114 | 81033 | 109472 | 133788 | 82306 | 83544 | 100524 |
| 664 | 1097 | 741 | 635 | 1079 | 789 | 682 | 1126 | 179889 | 152562 | 229677 | 251933 | 151460 | 129698 | 204302 |
| 625 | 1036 | 918 | 713 | 1069 | 912 | 695 | 1072 | 123226 | 93393 | 149436 | 154813 | 140244 | 109037 | 152840 |
| 630 | 1060 | 737 | 640 | 1090 | 739 | 655 | 1123 | 64919 | 57112 | 90600 | 96041 | 74448 | 64658 | 100955 |
| 489 | 1023 | 524 | 492 | 1010 | 511 | 455 | 979 | 124675 | 113686 | 232337 | 237797 | 123843 | 116457 | 236524 |
| 911 | 1324 | 1034 | 966 | 1362 | 964 | 902 | 1344 | 82172 | 75975 | 83441 | 110497 | 88663 | 82900 | 85782 |
| 787 | 987 | 858 | 844 | 936 | 810 | 764 | 936 | 99441 | 92051 | 117004 | 115495 | 141375 | 139100 | 164739 |
| 393 | 709 | 714 | 508 | 769 | 652 | 426 | 716 | 6851 | 4396 | 11183 | 7929 | 22270 | 15851 | 31178 |
| 644 | 870 | 1183 | 819 | 913 | 1328 | 729 | 830 | 10733 | 6237 | 9684 | 8425 | 5860 | 4054 | 4953 |
| 526 | 918 | 867 | 571 | 938 | 825 | 545 | 915 | 156755 | 105549 | 200789 | 184340 | 128797 | 84894 | 148548 |
| 625 | 1082 | 855 | 641 | 1099 | 871 | 688 | 1122 | 109394 | 83780 | 134045 | 144971 | 134350 | 100644 | 157093 |
| 461 | 887 | 725 | 502 | 881 | 645 | 469 | 880 | 169891 | 122719 | 266129 | 236185 | 131625 | 91172 | 181535 |

| | | | | | | | | | | | | | | |
|---|---|---|---|---|---|---|---|---|---|---|---|---|---|---|
| 829 | 1298 | 911 | 820 | 1252 | 893 | 766 | 1268 | 134187 | 116492 | 140561 | 182446 | 142266 | 128085 | 156115 |
| 1060 | 1530 | 806 | 1011 | 1523 | 854 | 1094 | 1563 | 106602 | 138408 | 130512 | 199638 | 113832 | 142842 | 141291 |
| 622 | 950 | 785 | 640 | 1001 | 768 | 613 | 967 | 142814 | 116049 | 186488 | 177194 | 146536 | 119394 | 186691 |
| 691 | 1106 | 823 | 653 | 1102 | 850 | 685 | 1111 | 101030 | 80246 | 116153 | 128469 | 129974 | 103157 | 157932 |
| 686 | 1039 | 786 | 592 | 986 | 777 | 593 | 885 | 20246 | 16049 | 23401 | 24309 | 27747 | 20913 | 35324 |
| 696 | 1116 | 632 | 622 | 1134 | 681 | 683 | 1196 | 92521 | 86353 | 124126 | 138480 | 58412 | 57495 | 92473 |
| 538 | 995 | 533 | 503 | 987 | 550 | 516 | 965 | 141793 | 133468 | 248041 | 246825 | 123119 | 116208 | 231046 |
| 914 | 1363 | 1068 | 873 | 1663 | 1050 | 808 | 1971 | 8833 | 7998 | 8751 | 11930 | 7857 | 6424 | 7357 |
| 833 | 1313 | 878 | 872 | 1313 | 837 | 728 | 1189 | 119724 | 117002 | 140432 | 184323 | 119387 | 118522 | 135944 |
| 671 | 1137 | 952 | 651 | 1128 | 949 | 675 | 1135 | 130392 | 93528 | 139354 | 158437 | 101212 | 69206 | 106291 |
| 937 | 1319 | 871 | 874 | 1275 | 912 | 966 | 1383 | 120521 | 119081 | 127139 | 167702 | 115048 | 115510 | 132149 |
| 816 | 1186 | 984 | 865 | 1196 | 942 | 818 | 1221 | 116931 | 100619 | 123242 | 146217 | 119003 | 104709 | 120984 |
| 535 | 940 | 650 | 514 | 949 | 644 | 515 | 947 | 116896 | 93549 | 174947 | 164515 | 141436 | 111698 | 217474 |
| 795 | 1198 | 879 | 809 | 1207 | 924 | 865 | 1294 | 109795 | 96879 | 121860 | 145998 | 110388 | 101542 | 125520 |
| 729 | 1090 | 727 | 627 | 1014 | 771 | 695 | 1094 | 146425 | 136500 | 187119 | 203967 | 130800 | 112739 | 179862 |
| 724 | 1105 | 809 | 686 | 1058 | 826 | 658 | 1039 | 129508 | 110005 | 151910 | 167797 | 117205 | 99384 | 144814 |
| 731 | 1268 | 713 | 664 | 1213 | 730 | 669 | 1200 | 104525 | 95495 | 130641 | 165671 | 132102 | 122856 | 185148 |
| | | 720 | 615 | 985 | 837 | 736 | 1062 | | | | | 3761 | 3210 | 5221 |
| 708 | 1087 | 810 | 660 | 1064 | 813 | 706 | 1119 | 115314 | 95539 | 134930 | 146676 | 116391 | 94826 | 143702 |
| 929 | 1064 | 1308 | 1097 | 959 | 1301 | 779 | 971 | 5293 | 4047 | 4358 | 4636 | 5341 | 4482 | 4084 |
| | | 1050 | 651 | 990 | 899 | 556 | 942 | | | | | 1891 | 1172 | 1800 |
| 747 | 1169 | 851 | 779 | 1187 | 791 | 671 | 1091 | 90072 | 80538 | 107791 | 126034 | 119762 | 109589 | 140759 |
| 592 | 976 | 753 | 629 | 1043 | 824 | 731 | 1033 | 43606 | 34185 | 57734 | 56373 | 48233 | 40278 | 64041 |
| 843 | 1255 | 794 | 784 | 1243 | 831 | 842 | 1283 | 134370 | 139286 | 165171 | 207306 | 109524 | 108049 | 137859 |
| 666 | 1113 | 764 | 603 | 1043 | 827 | 630 | 1068 | 135520 | 115722 | 173844 | 193404 | 162853 | 128489 | 213228 |
| 492 | 856 | 715 | 505 | 888 | 680 | 470 | 849 | 151819 | 106901 | 217120 | 185892 | 119482 | 84409 | 167156 |
| 479 | 584 | 1296 | 756 | 899 | 1189 | 724 | 883 | 8228 | 3487 | 7273 | 4245 | 6570 | 3832 | 5069 |
| 790 | 1125 | 923 | 778 | 1122 | 919 | 735 | 1140 | 106711 | 87113 | 110305 | 124142 | 134537 | 113454 | 145743 |
| 837 | 1168 | 890 | 832 | 1155 | 913 | 816 | 1130 | 76727 | 69029 | 82466 | 96299 | 59951 | 56039 | 67349 |

| | | | | | | | | | | | | | | |
|---|---|---|---|---|---|---|---|---|---|---|---|---|---|---|
| 605 | 1035 | 744 | 598 | 1002 | 746 | 650 | 1053 | 137385 | 109276 | 180743 | 187048 | 125875 | 101277 | 169292 |
| 744 | 1199 | 694 | 763 | 1251 | 649 | 664 | 1182 | 112047 | 120324 | 161740 | 193915 | 100968 | 110972 | 145465 |
| 549 | 924 | 767 | 612 | 964 | 775 | 601 | 982 | 84594 | 64403 | 117326 | 108352 | 83834 | 66904 | 109305 |
| 630 | 1016 | 867 | 667 | 1143 | 869 | 649 | 1054 | 113549 | 82801 | 131461 | 133571 | 128917 | 99169 | 148616 |
| 897 | 1346 | 916 | 856 | 1334 | 931 | 820 | 1238 | 128307 | 122402 | 136413 | 183548 | 104532 | 97619 | 114094 |
| 792 | 1191 | 829 | 718 | 1127 | 922 | 828 | 1182 | 83435 | 74649 | 94240 | 112285 | 87867 | 76191 | 106043 |
| 688 | 1100 | 780 | 701 | 1137 | 761 | 687 | 1122 | 80092 | 70781 | 102849 | 113087 | 88121 | 79207 | 112982 |
| 842 | 1213 | 842 | 801 | 1205 | 847 | 799 | 1145 | 86516 | 81578 | 96867 | 117493 | 86275 | 82065 | 102460 |
| 420 | 900 | 663 | 376 | 842 | 691 | 369 | 820 | 125474 | 71135 | 169548 | 152585 | 132227 | 75092 | 199573 |
| 827 | 1279 | 876 | 790 | 1255 | 873 | 774 | 1264 | 112815 | 101063 | 122268 | 156343 | 103121 | 93053 | 117744 |
| 484 | 967 | 790 | 552 | 987 | 742 | 532 | 983 | 127262 | 85295 | 176287 | 170440 | 134571 | 94065 | 170287 |
| 564 | 997 | 657 | 521 | 969 | 720 | 573 | 988 | 104309 | 83749 | 148408 | 147928 | 84774 | 67214 | 129017 |
| 652 | 1073 | 878 | 720 | 1097 | 842 | 669 | 1069 | 95674 | 77204 | 118472 | 127127 | 112280 | 92137 | 127885 |
| 674 | 1082 | 834 | 675 | 1099 | 839 | 725 | 1184 | 64794 | 51811 | 76912 | 83257 | 75048 | 60707 | 89951 |
| 968 | 1337 | 954 | 628 | 945 | 1006 | 863 | 1270 | 16522 | 14924 | 15422 | 20621 | 10796 | 7103 | 11311 |
| 366 | 810 | 516 | 391 | 788 | 575 | 439 | 832 | 60439 | 43801 | 119743 | 97036 | 92302 | 69965 | 178845 |
| 332 | 855 | 457 | 311 | 848 | 455 | 327 | 883 | 78678 | 57244 | 172641 | 147613 | 83330 | 56779 | 182450 |
| 653 | 1043 | 713 | 621 | 1026 | 730 | 643 | 1034 | 169887 | 146142 | 223742 | 233422 | 136741 | 119042 | 191771 |
| 562 | 914 | 680 | 561 | 869 | 658 | 541 | 874 | 107483 | 92257 | 164157 | 150112 | 135332 | 111686 | 198958 |
| 666 | 864 | 807 | 616 | 835 | 787 | 626 | 854 | 126310 | 99788 | 149741 | 129343 | 111700 | 85299 | 138360 |
| 606 | 914 | 730 | 582 | 882 | 742 | 599 | 892 | 107765 | 86580 | 142793 | 130512 | 112420 | 89585 | 154006 |
| 730 | 1135 | 922 | 670 | 1062 | 922 | 681 | 1103 | 149812 | 109775 | 150438 | 170766 | 150777 | 109606 | 163549 |
| 749 | 1113 | 862 | 771 | 1097 | 817 | 721 | 1044 | 79774 | 70847 | 94531 | 105213 | 79885 | 71465 | 92691 |
| 960 | 1295 | 957 | 884 | 1273 | 836 | 793 | 1190 | 94912 | 94714 | 98671 | 127787 | 89792 | 83003 | 93862 |
| 668 | 1089 | 821 | 701 | 1082 | 864 | 749 | 1122 | 128366 | 108743 | 162889 | 177322 | 124615 | 106450 | 151828 |
| 521 | 896 | 627 | 544 | 919 | 534 | 454 | 878 | 85683 | 69670 | 133761 | 119903 | 60254 | 52198 | 96031 |
| 500 | 910 | 694 | 532 | 976 | 693 | 537 | 1004 | 112352 | 84738 | 169563 | 154235 | 132924 | 101908 | 191584 |
| 533 | 934 | 630 | 564 | 942 | 686 | 620 | 976 | 83498 | 77598 | 145501 | 135947 | 75495 | 67590 | 119905 |
| 944 | 1339 | 1100 | 980 | 1360 | 1042 | 886 | 1272 | 83667 | 73606 | 77980 | 104416 | 77295 | 68879 | 70296 |
| 723 | 1128 | 713 | 684 | 1120 | 755 | 691 | 1122 | 127914 | 117772 | 162785 | 183680 | 104976 | 100790 | 147263 |
| 667 | 1059 | 970 | 765 | 1115 | 937 | 738 | 1105 | 89444 | 66532 | 99820 | 105686 | 82156 | 64792 | 84737 |

| | | | | | | | | | | | | | | |
|---|---|---|---|---|---|---|---|---|---|---|---|---|---|---|
| 302 | 625 | 736 | 340 | 639 | 728 | 285 | 576 | 10307 | 4378 | 14509 | 9065 | 13864 | 6401 | 18824 |
| 751 | 1052 | 952 | 788 | 1047 | 912 | 822 | 1088 | 84571 | 73005 | 97269 | 102350 | 86460 | 71561 | 90811 |
| 1067 | 1336 | 935 | 975 | 1226 | 1030 | 1047 | 1310 | 23937 | 22700 | 21278 | 28418 | 18416 | 19214 | 19696 |
| 806 | 1285 | 796 | 749 | 1274 | 787 | 748 | 1198 | 79571 | 75334 | 93450 | 120090 | 97586 | 91929 | 122656 |
| 514 | 807 | 825 | 530 | 807 | 787 | 492 | 799 | 121179 | 71815 | 139664 | 112655 | 139932 | 90002 | 169659 |
| 540 | 1062 | 728 | 632 | 1192 | 714 | 631 | 1153 | 56233 | 46478 | 86117 | 91415 | 95857 | 83287 | 131753 |
| 739 | 1105 | 658 | 746 | 1139 | 686 | 771 | 1174 | 90901 | 97037 | 131280 | 145013 | 68841 | 78041 | 104682 |
| 715 | 1247 | 862 | 765 | 1278 | 944 | 834 | 1392 | 60015 | 52081 | 72845 | 90873 | 51689 | 45870 | 59967 |
| 668 | 1121 | 861 | 639 | 1129 | 839 | 604 | 1062 | 84386 | 64293 | 96280 | 107921 | 69110 | 51297 | 80261 |
| 718 | 1208 | 749 | 646 | 1134 | 863 | 746 | 1184 | 102334 | 90300 | 125759 | 151855 | 57192 | 49322 | 76369 |
| 594 | 1064 | 844 | 674 | 1127 | 812 | 651 | 1149 | 76123 | 59010 | 99330 | 105671 | 88587 | 70751 | 104938 |
| 634 | 952 | 821 | 609 | 918 | 807 | 599 | 931 | 30370 | 23788 | 37521 | 35711 | 8491 | 6294 | 10337 |
| 673 | 1199 | 847 | 667 | 1214 | 803 | 653 | 1231 | 76015 | 61591 | 91575 | 109758 | 81322 | 64103 | 96036 |
| 438 | 698 | 607 | 326 | 591 | 752 | 357 | 607 | 4963 | 2774 | 6338 | 4423 | 5089 | 2728 | 8378 |
| 861 | 1208 | 1029 | 902 | 1247 | 985 | 866 | 1200 | 96611 | 85763 | 99618 | 120382 | 84471 | 74040 | 82058 |
| 555 | 1007 | 655 | 517 | 973 | 665 | 534 | 979 | 114423 | 89878 | 161907 | 163054 | 68351 | 53965 | 104306 |
| 529 | 871 | 559 | 493 | 835 | 603 | 551 | 889 | 82519 | 74289 | 140465 | 122372 | 62932 | 55488 | 112498 |
| 555 | 947 | 769 | 552 | 931 | 757 | 550 | 980 | 80870 | 56592 | 102035 | 96660 | 89753 | 64402 | 116705 |
| 729 | 1005 | 1080 | 701 | 971 | 1124 | 787 | 1060 | 16484 | 11641 | 15967 | 16049 | 17061 | 11081 | 15804 |
| 733 | 1322 | 837 | 703 | 1278 | 882 | 718 | 1210 | 97374 | 81970 | 111846 | 147851 | 106072 | 89119 | 126757 |
| 731 | 1101 | 764 | 668 | 1045 | 770 | 655 | 1040 | 106705 | 95716 | 130863 | 144094 | 83791 | 73349 | 109728 |
| 1660 | 2042 | 1040 | 1553 | 1967 | 1164 | 1809 | 2199 | 9284 | 14098 | 8492 | 17343 | 3639 | 5436 | 3500 |
| 611 | 940 | 812 | 588 | 907 | 843 | 617 | 970 | 121955 | 87746 | 143525 | 134850 | 79392 | 57513 | 97805 |
| 517 | 836 | 565 | 488 | 827 | 582 | 506 | 831 | 79050 | 71403 | 138192 | 115466 | 86970 | 75113 | 153984 |
| 547 | 966 | 728 | 495 | 865 | 753 | 504 | 907 | 98092 | 72961 | 133319 | 128748 | 108407 | 73706 | 148823 |
| 816 | 1092 | 934 | 781 | 1081 | 1017 | 830 | 1087 | 101529 | 86912 | 106527 | 116336 | 85979 | 71910 | 92029 |
| 1005 | 1370 | 1010 | 996 | 1396 | 888 | 834 | 1162 | 99525 | 99146 | 98684 | 135241 | 84491 | 83399 | 83693 |
| 641 | 1008 | 795 | 630 | 993 | 796 | 626 | 978 | 89952 | 70505 | 110050 | 110952 | 119053 | 94385 | 149768 |
| | | | | | 836 | 421 | 746 | | | | | | | |
| 609 | 1224 | 747 | 632 | 1229 | 732 | 616 | 1226 | 83515 | 70195 | 115192 | 141043 | 91747 | 77681 | 122832 |
| 584 | 1004 | 702 | 553 | 975 | 702 | 569 | 992 | 88565 | 71993 | 123368 | 123907 | 82552 | 65104 | 117661 |

| | | | | | | | | | | | | | | |
|---|---|---|---|---|---|---|---|---|---|---|---|---|---|---|
| 274 | 693 | 523 | 336 | 695 | 498 | 318 | 726 | 93413 | 57160 | 208993 | 144749 | 110516 | 70960 | 211282 |
| 1132 | 1686 | 1029 | 1094 | 1592 | 871 | 899 | 1451 | 100454 | 106615 | 94184 | 158779 | 114893 | 122158 | 111660 |
| 506 | 859 | 711 | 524 | 854 | 662 | 501 | 869 | 80082 | 57860 | 114322 | 98152 | 100397 | 73946 | 141158 |
| 432 | 870 | 624 | 386 | 817 | 664 | 432 | 868 | 92623 | 60300 | 139426 | 121253 | 87826 | 54265 | 140704 |
| 773 | 1091 | 874 | 689 | 1018 | 1024 | 820 | 1154 | 100304 | 77130 | 99793 | 108924 | 106783 | 84139 | 122183 |
| 870 | 1388 | 919 | 916 | 1476 | 879 | 883 | 1413 | 67003 | 66909 | 76920 | 106804 | 55303 | 55093 | 60158 |
| | | 749 | 587 | 974 | | | | | | | | 3619 | 2832 | 4828 |
| 756 | 1203 | 939 | 807 | 1241 | 853 | 747 | 1167 | 99743 | 86100 | 113862 | 137018 | 118035 | 101369 | 125637 |
| 530 | 974 | 759 | 524 | 982 | 735 | 503 | 1017 | 111535 | 77246 | 145619 | 141859 | 104947 | 72396 | 138220 |
| 481 | 795 | 649 | 440 | 762 | 668 | 444 | 752 | 146373 | 98321 | 204293 | 162500 | 82730 | 56121 | 127476 |
| 670 | 1166 | 900 | 703 | 1155 | 818 | 678 | 1181 | 111436 | 89001 | 132744 | 154781 | 101004 | 78950 | 112276 |
| 832 | 1311 | 1145 | 1055 | 1464 | 1082 | 1007 | 1396 | 32988 | 28240 | 33945 | 44490 | 44576 | 41084 | 38928 |
| 409 | 711 | 825 | 419 | 702 | 917 | 555 | 886 | 2384 | 1137 | 2779 | 1977 | 2050 | 1041 | 2486 |
| 741 | 1228 | 866 | 815 | 1286 | 862 | 802 | 1267 | 67698 | 58415 | 78808 | 96751 | 86824 | 81718 | 100207 |
| 513 | 989 | 618 | 524 | 983 | 684 | 556 | 1009 | 66747 | 53543 | 104408 | 103219 | 66913 | 56748 | 108252 |
| 823 | 1437 | 859 | 767 | 1378 | 872 | 808 | 1401 | 37227 | 34900 | 42400 | 60914 | 37274 | 33281 | 43391 |
| 627 | 1047 | 733 | 601 | 981 | 726 | 637 | 1062 | 89927 | 77622 | 123820 | 129674 | 89540 | 73339 | 122122 |
| 1065 | 1343 | 1038 | 1035 | 1342 | 845 | 854 | 1157 | 31338 | 32686 | 30688 | 41203 | 13167 | 13133 | 12689 |
| 586 | 1046 | 776 | 599 | 1054 | 767 | 567 | 1021 | 35959 | 28477 | 48573 | 50804 | 55319 | 42703 | 71324 |
| 705 | 1191 | 732 | 698 | 1178 | 749 | 668 | 1145 | 84111 | 80806 | 114687 | 136558 | 72348 | 68974 | 98813 |
| 692 | 1083 | 790 | 695 | 1070 | 769 | 742 | 1089 | 56136 | 49910 | 72134 | 78144 | 69561 | 61165 | 88033 |
| 592 | 943 | 640 | 577 | 974 | 661 | 638 | 985 | 92782 | 83693 | 141477 | 133452 | 93132 | 83964 | 145427 |
| 738 | 1280 | 945 | 807 | 1332 | 883 | 725 | 1211 | 65821 | 55600 | 75349 | 96409 | 83615 | 71412 | 88459 |
| 286 | 450 | 647 | 311 | 455 | 665 | 282 | 425 | 129578 | 57932 | 202815 | 91291 | 114526 | 55089 | 176935 |
| 437 | 771 | 637 | 455 | 783 | 650 | 511 | 835 | 82844 | 58502 | 133724 | 103103 | 108884 | 77799 | 170992 |
| 678 | 1076 | 811 | 655 | 1060 | 742 | 563 | 1010 | 75990 | 65028 | 95853 | 103101 | 96493 | 77987 | 118981 |
| 611 | 976 | 728 | 660 | 984 | 723 | 665 | 1000 | 59444 | 51600 | 84412 | 82349 | 71234 | 64578 | 97840 |
| 850 | 1402 | 909 | 850 | 1389 | 852 | 826 | 1372 | 59629 | 58534 | 68901 | 96626 | 111251 | 104016 | 122371 |
| 582 | 972 | 721 | 580 | 1035 | 721 | 562 | 984 | 100844 | 79002 | 135791 | 131945 | 96342 | 77463 | 133558 |
| 1069 | 1401 | 921 | 943 | 1305 | 862 | 830 | 1354 | 14711 | 14894 | 13938 | 19526 | 10895 | 11159 | 11833 |
| 870 | 1213 | 961 | 790 | 1130 | 938 | 787 | 1141 | 7298 | 6212 | 7138 | 8658 | 10755 | 8844 | 11196 |

| | | | | | | | | | | | | | | |
|---|---|---|---|---|---|---|---|---|---|---|---|---|---|---|
| 804 | 1109 | 1073 | 804 | 1132 | 1002 | 759 | 1038 | 82723 | 64158 | 79757 | 88412 | 113980 | 85365 | 106211 |
| 902 | 1350 | 992 | 856 | 1289 | 940 | 792 | 1243 | 67344 | 59475 | 65949 | 89036 | 66914 | 57750 | 67488 |
| 663 | 1183 | 802 | 634 | 1190 | 830 | 682 | 1231 | 85216 | 66480 | 100265 | 118639 | 88445 | 70011 | 110345 |
| 664 | 1093 | 856 | 785 | 1122 | 751 | 654 | 1094 | 73543 | 65342 | 98480 | 107644 | 67039 | 61508 | 78361 |
| 700 | 1260 | 687 | 637 | 1224 | 740 | 658 | 1222 | 81665 | 75964 | 108502 | 136704 | 65741 | 60925 | 95639 |
| 757 | 1045 | 869 | 762 | 1005 | 941 | 758 | 1013 | 124158 | 107663 | 142252 | 148659 | 110167 | 96540 | 126720 |
| 862 | 1150 | 990 | 823 | 1135 | 960 | 816 | 1104 | 63510 | 55447 | 64306 | 73975 | 52742 | 43820 | 53249 |
| 748 | 1055 | 906 | 688 | 996 | 967 | 756 | 1138 | 85331 | 66665 | 89096 | 93993 | 86713 | 65848 | 95701 |
| 830 | 1153 | 776 | 768 | 1124 | 799 | 804 | 1133 | 90931 | 90885 | 109457 | 126213 | 69832 | 69113 | 90004 |
| 1005 | 1805 | 949 | 1043 | 1747 | 963 | 1054 | 1932 | 67420 | 70641 | 70286 | 126846 | 85966 | 94492 | 90616 |
| 476 | 901 | 495 | 429 | 860 | 449 | 384 | 834 | 45058 | 41614 | 87401 | 78724 | 60005 | 51993 | 121236 |
| 898 | 1130 | 1256 | 1037 | 1188 | 1189 | 989 | 1191 | 89389 | 70220 | 78168 | 88365 | 92130 | 76117 | 73379 |
| 696 | 1038 | 867 | 716 | 1023 | 866 | 752 | 1091 | 59436 | 50225 | 72138 | 74853 | 58246 | 48110 | 67201 |
| 616 | 964 | 903 | 637 | 966 | 878 | 647 | 977 | 83723 | 58729 | 95316 | 91888 | 60630 | 42794 | 67178 |
| 565 | 812 | 709 | 513 | 789 | 736 | 540 | 807 | 76446 | 57905 | 102538 | 83286 | 86132 | 62259 | 121449 |
| 605 | 959 | 751 | 587 | 990 | 744 | 599 | 981 | 59788 | 48773 | 80639 | 77320 | 71333 | 55766 | 94958 |
| 862 | 1245 | 983 | 838 | 1229 | 1016 | 876 | 1253 | 66458 | 58394 | 67773 | 84362 | 89163 | 76003 | 90683 |
| 667 | 1004 | 769 | 613 | 930 | 734 | 569 | 897 | 77010 | 65562 | 98337 | 98773 | 92298 | 73602 | 119994 |
| 806 | 949 | 1315 | 1049 | 1106 | 1106 | 786 | 805 | 1451 | 1052 | 1305 | 1238 | 2161 | 1724 | 1643 |
| 354 | 647 | 511 | 372 | 655 | 510 | 376 | 662 | 64933 | 46922 | 132682 | 85900 | 60618 | 44166 | 118706 |
| 435 | 792 | 772 | 474 | 822 | 708 | 426 | 814 | 82951 | 51909 | 119279 | 94479 | 89954 | 55268 | 116507 |
| 744 | 1026 | 816 | 612 | 948 | 741 | 507 | 934 | 80496 | 63148 | 84926 | 87170 | 71796 | 53828 | 88004 |
| 803 | 1217 | 863 | 843 | 1265 | 754 | 778 | 1246 | 76275 | 74504 | 92773 | 112937 | 57800 | 56437 | 66967 |
| 694 | 990 | 898 | 685 | 981 | 950 | 746 | 1048 | 84982 | 63772 | 91864 | 90951 | 112297 | 85651 | 125019 |
| 768 | 1248 | 943 | 867 | 1300 | 894 | 829 | 1205 | 80720 | 70572 | 91904 | 114716 | 62070 | 57062 | 65793 |
| 920 | 1353 | 745 | 909 | 1341 | 806 | 930 | 1301 | 46803 | 55060 | 59843 | 80961 | 54358 | 66297 | 72921 |
| 909 | 1259 | 1087 | 997 | 1212 | 944 | 860 | 1215 | 51497 | 47507 | 52255 | 65811 | 53695 | 49243 | 49415 |
| 910 | 1119 | 743 | 924 | 1141 | 765 | 966 | 1162 | 54628 | 68672 | 75428 | 84407 | 79602 | 99007 | 107189 |
| 879 | 1282 | 1007 | 908 | 1284 | 850 | 743 | 1240 | 47369 | 43831 | 49856 | 63925 | 57446 | 51787 | 57050 |
| 909 | 1301 | 963 | 835 | 1263 | 1018 | 887 | 1278 | 64994 | 58744 | 64632 | 84114 | 66419 | 57579 | 68983 |
| 676 | 1102 | 888 | 716 | 1141 | 919 | 753 | 1172 | 55403 | 44094 | 65206 | 71886 | 43871 | 35383 | 49394 |

| | | | | | | | | | | | | | | |
|---|---|---|---|---|---|---|---|---|---|---|---|---|---|---|
| 369 | 697 | 566 | 391 | 730 | 535 | 360 | 699 | 60012 | 42613 | 115543 | 80545 | 67254 | 46428 | 118833 |
| 835 | 1240 | 825 | 802 | 1231 | 847 | 816 | 1268 | 43220 | 41506 | 49737 | 61692 | 42467 | 41306 | 51489 |
| 1148 | 1648 | 1055 | 1055 | 1584 | 997 | 965 | 1469 | 77048 | 79413 | 69147 | 113945 | 48221 | 48194 | 45701 |
| 756 | 1050 | 911 | 836 | 1157 | 896 | 777 | 1022 | 33833 | 30465 | 40277 | 42304 | 30545 | 28047 | 33546 |
| 665 | 902 | 845 | 676 | 917 | 872 | 608 | 867 | 3621 | 2811 | 4226 | 3811 | 4542 | 3632 | 5373 |
| 648 | 1063 | 830 | 652 | 1083 | 831 | 696 | 1116 | 91946 | 68222 | 105293 | 111949 | 69428 | 54569 | 83678 |
| 690 | 952 | 925 | 731 | 994 | 968 | 770 | 991 | 59630 | 47316 | 68560 | 65284 | 59990 | 47415 | 64865 |
| 588 | 873 | 758 | 547 | 893 | 732 | 540 | 876 | 67339 | 49837 | 84723 | 73952 | 79233 | 57192 | 104504 |
| 594 | 951 | 848 | 586 | 950 | 852 | 574 | 927 | 74858 | 52882 | 89025 | 84636 | 79865 | 55150 | 94130 |
| 979 | 1261 | 1121 | 1151 | 1401 | 1060 | 1063 | 1357 | 54841 | 52039 | 53162 | 67020 | 27989 | 28742 | 24978 |
| 840 | 1190 | 976 | 898 | 1255 | 1001 | 932 | 1220 | 33836 | 30723 | 36565 | 43505 | 55086 | 50736 | 56468 |
| 551 | 974 | 701 | 538 | 953 | 707 | 532 | 971 | 58013 | 45122 | 81851 | 79759 | 69092 | 53038 | 98560 |
| 447 | 767 | 709 | 489 | 819 | 768 | 533 | 828 | 42044 | 27801 | 62257 | 47777 | 52630 | 36302 | 74185 |
| 749 | 1213 | 802 | 650 | 1172 | 834 | 675 | 1112 | 79801 | 68478 | 91482 | 110937 | 77412 | 62729 | 96540 |
| 652 | 1062 | 936 | 671 | 1083 | 935 | 630 | 1056 | 68089 | 47410 | 72691 | 77199 | 73546 | 52702 | 78582 |
| 864 | 1186 | 1046 | 840 | 1218 | 1092 | 1031 | 1257 | 66704 | 58035 | 67166 | 79651 | 79418 | 63741 | 75913 |
| 603 | 781 | 793 | 561 | 732 | 788 | 608 | 783 | 55117 | 44828 | 74283 | 57984 | 89142 | 63070 | 112428 |
| 553 | 990 | 949 | 661 | 983 | | | | 346 | 208 | 375 | 371 | 209 | 145 | 220 |
| 738 | 1122 | 792 | 722 | 1106 | 784 | 674 | 1041 | 64148 | 59457 | 80583 | 90413 | 62795 | 57280 | 79328 |
| 495 | 804 | 633 | 520 | 841 | 643 | 527 | 856 | 81748 | 67883 | 137003 | 110197 | 92129 | 75707 | 145659 |
| 807 | 1345 | 916 | 802 | 1226 | 905 | 803 | 1413 | 2549 | 2335 | 2892 | 3892 | 6115 | 5352 | 6673 |
| | | | | | 682 | 578 | 1008 | | | | | | | |
| 403 | 749 | | | | | | | 1617 | 677 | 1679 | 1257 | | | |
| 680 | 1123 | 804 | 660 | 1074 | 818 | 636 | 1039 | 57988 | 49653 | 73021 | 82000 | 48897 | 40124 | 60794 |
| 236 | 417 | 818 | 217 | 388 | 832 | 237 | 386 | 140263 | 39513 | 167142 | 69712 | 126092 | 33487 | 154089 |
| 962 | 1292 | 967 | 939 | 1259 | 1037 | 974 | 1307 | 50858 | 48994 | 50916 | 65795 | 57973 | 56265 | 59952 |
| 697 | 1109 | 838 | 676 | 1130 | 893 | 714 | 1142 | 83032 | 61452 | 88176 | 97746 | 81145 | 65460 | 96853 |
| 616 | 912 | 908 | 682 | 1011 | 953 | 721 | 1011 | 60992 | 43055 | 69884 | 63709 | 82289 | 61846 | 90629 |
| 809 | 1116 | 1014 | 809 | 1117 | 1092 | 897 | 1140 | 70212 | 55070 | 68030 | 75930 | 72193 | 57555 | 71178 |
| 771 | 1250 | 868 | 780 | 1274 | 821 | 731 | 1231 | 59065 | 54555 | 70772 | 88431 | 64242 | 57730 | 73983 |
| 917 | 1380 | 914 | 905 | 1325 | 924 | 884 | 1332 | 82683 | 80614 | 87929 | 121382 | 79233 | 78429 | 86685 |

| | | | | | | | | | | | | | | |
|---|---|---|---|---|---|---|---|---|---|---|---|---|---|---|
| 549 | 970 | 552 | 479 | 881 | 562 | 519 | 927 | 44496 | 41005 | 74624 | 72380 | 59482 | 51606 | 107780 |
| 560 | 1024 | 689 | 568 | 1010 | 646 | 575 | 1046 | 35519 | 30978 | 55332 | 56638 | 70781 | 58375 | 102760 |
| 411 | 625 | 839 | 607 | 831 | 809 | 631 | 770 | 892 | 472 | 1147 | 717 | 1504 | 1089 | 1794 |
| | | | | | 1391 | 793 | 922 | | | | | | | |
| 622 | 1104 | 673 | 530 | 999 | 705 | 576 | 1013 | 88165 | 74776 | 120198 | 132743 | 87759 | 69165 | 130486 |
| | | 810 | 687 | 1119 | 796 | 783 | 1247 | | | | | 5118 | 4342 | 6317 |
| 876 | 1240 | 891 | 779 | 1148 | 911 | 842 | 1253 | 71355 | 66161 | 75530 | 93678 | 47491 | 41533 | 53322 |
| 565 | 955 | 641 | 544 | 921 | 631 | 533 | 918 | 74103 | 64346 | 113881 | 108752 | 72217 | 61312 | 112609 |
| 592 | 983 | 743 | 589 | 965 | 766 | 617 | 982 | 78395 | 61934 | 104555 | 102778 | 63164 | 50083 | 85014 |
| 767 | 1037 | 912 | 764 | 991 | 865 | 757 | 1062 | 55836 | 47327 | 61702 | 64009 | 77557 | 64979 | 85046 |
| 792 | 1324 | 782 | 751 | 1326 | 851 | 801 | 1326 | 54143 | 50993 | 64373 | 85218 | 67715 | 65044 | 86626 |
| 437 | 698 | 1037 | 494 | 706 | 949 | 446 | 706 | 83633 | 39946 | 91502 | 63837 | 50173 | 23890 | 48386 |
| | | | | | 815 | 662 | 941 | | | | | | | |
| 657 | 997 | 928 | 790 | 1054 | 909 | 738 | 1014 | 81327 | 66546 | 101348 | 101052 | 61613 | 52507 | 66427 |
| | | 848 | 656 | 1149 | | | | | | | | 680 | 526 | 802 |
| 643 | 865 | 828 | 627 | 853 | 807 | 635 | 862 | 84030 | 66991 | 104205 | 90089 | 95759 | 72486 | 115672 |
| 590 | 819 | 528 | 528 | 747 | 597 | 547 | 788 | 18882 | 15776 | 26761 | 21910 | 18532 | 18558 | 35116 |
| 666 | 980 | 875 | 680 | 984 | 864 | 654 | 937 | 61583 | 48045 | 72099 | 70633 | 80191 | 62334 | 91651 |
| 523 | 776 | 550 | 479 | 770 | 560 | 477 | 744 | 67547 | 60821 | 116259 | 90174 | 51982 | 45200 | 94451 |
| 674 | 1075 | 863 | 665 | 1074 | 857 | 697 | 1127 | 73527 | 60102 | 89164 | 95881 | 79685 | 61459 | 92373 |
| 731 | 1219 | 826 | 711 | 1180 | 857 | 752 | 1216 | 47059 | 41107 | 56251 | 68564 | 52796 | 45440 | 63896 |
| 566 | 1010 | 717 | 566 | 992 | 712 | 580 | 979 | 49684 | 36457 | 64436 | 65073 | 50436 | 39830 | 70314 |
| 903 | 1259 | 928 | 958 | 1251 | 897 | 987 | 1285 | 32190 | 31641 | 35021 | 44078 | 42982 | 44386 | 46339 |
| 904 | 1362 | 1003 | 898 | 1315 | 946 | 805 | 1289 | 58086 | 49674 | 54948 | 74864 | 30216 | 27054 | 30125 |
| 765 | 1003 | 929 | 760 | 970 | 845 | 687 | 1021 | 30633 | 24891 | 32526 | 32627 | 23416 | 19159 | 25206 |
| 365 | 757 | 579 | 456 | 812 | 529 | 395 | 779 | 53701 | 39297 | 107738 | 81533 | 62944 | 49599 | 108692 |
| 631 | 1026 | 896 | 687 | 1041 | 836 | 612 | 1015 | 63695 | 48700 | 77130 | 79139 | 120391 | 92294 | 134386 |
| 663 | 1076 | 919 | 821 | 1188 | 861 | 686 | 1075 | 63096 | 53117 | 80075 | 86163 | 66078 | 59024 | 71894 |
| 406 | 847 | 584 | 398 | 876 | 519 | 357 | 851 | 45962 | 31477 | 77496 | 65648 | 39304 | 26787 | 67335 |
| 528 | 1003 | 795 | 498 | 994 | 675 | 436 | 924 | 44218 | 30161 | 57173 | 57358 | 72706 | 45488 | 91414 |

| | | | | | | | | | | | | | | |
|---|---|---|---|---|---|---|---|---|---|---|---|---|---|---|
| 519 | 839 | 799 | 571 | 905 | 860 | 573 | 904 | 72259 | 48708 | 93774 | 78642 | 60691 | 43417 | 75994 |
| 504 | 809 | 838 | 536 | 814 | 811 | 499 | 794 | 74102 | 45749 | 90769 | 73397 | 67975 | 43497 | 81154 |
| 644 | 974 | 802 | 649 | 971 | 815 | 653 | 985 | 45728 | 37327 | 57919 | 56429 | 31516 | 25492 | 39300 |
| 708 | 1210 | 815 | 720 | 1189 | 843 | 776 | 1284 | 74781 | 64206 | 90639 | 109657 | 42433 | 37501 | 52059 |
| 586 | 833 | 774 | 500 | 757 | 752 | 588 | 819 | 46866 | 35647 | 60850 | 50670 | 42547 | 27471 | 54949 |
| 630 | 760 | 663 | 712 | 775 | 682 | 688 | 808 | 36837 | 38269 | 60739 | 46166 | 60295 | 64701 | 90931 |
| 1037 | 1488 | 1002 | 1191 | 1536 | | | | 2875 | 2619 | 2525 | 3757 | 1503 | 1787 | 1501 |
| 652 | 1022 | 869 | 666 | 1026 | 864 | 684 | 1083 | 36258 | 28343 | 43463 | 44402 | 46907 | 35930 | 53950 |
| 723 | 1069 | 868 | 687 | 974 | 1100 | 932 | 1057 | 1224 | 1105 | 1528 | 1634 | 23882 | 18899 | 27505 |
| 640 | 1015 | 599 | 602 | 1018 | 603 | 594 | 962 | 44366 | 44091 | 68921 | 69954 | 51206 | 51497 | 85526 |
| 766 | 1221 | 780 | 666 | 1144 | 815 | 706 | 1200 | 59715 | 52217 | 68162 | 83252 | 43261 | 36949 | 55446 |
| 814 | 1262 | 925 | 876 | 1344 | 908 | 822 | 1274 | 62948 | 57423 | 70545 | 89011 | 47302 | 44781 | 51142 |
| 628 | 1227 | 769 | 611 | 1224 | 772 | 603 | 1190 | 72979 | 60088 | 95738 | 117470 | 63272 | 50298 | 82319 |
| 712 | 1048 | 761 | 673 | 1004 | 791 | 695 | 1007 | 52515 | 45305 | 63605 | 66657 | 68163 | 60319 | 89568 |
| 556 | 949 | 647 | 536 | 930 | 659 | 568 | 958 | 75968 | 63110 | 113540 | 107790 | 51908 | 42957 | 80209 |
| 838 | 1284 | 947 | 843 | 1269 | 938 | 842 | 1261 | 37987 | 34355 | 40990 | 52645 | 86901 | 77346 | 91718 |
| 513 | 952 | 690 | 574 | 970 | 725 | 607 | 993 | 77964 | 62203 | 121165 | 115316 | 66784 | 55534 | 96759 |
| 636 | 916 | 793 | 607 | 893 | 731 | 522 | 859 | 51615 | 40005 | 62890 | 57601 | 81757 | 62575 | 103089 |
| 604 | 1107 | 759 | 556 | 1079 | 764 | 595 | 1121 | 74158 | 56101 | 92824 | 102735 | 73851 | 54090 | 97247 |
| 354 | 632 | | | | 442 | 314 | 611 | 3415 | 2629 | 7435 | 4698 | | | |
| 601 | 1052 | 589 | 548 | 966 | 592 | 550 | 969 | 56629 | 56611 | 94164 | 99015 | 40340 | 37542 | 68478 |
| 442 | 708 | 619 | 442 | 716 | 594 | 424 | 693 | 46981 | 36086 | 81581 | 57792 | 60000 | 42767 | 96853 |
| 583 | 1028 | 734 | 590 | 1008 | 705 | 583 | 1064 | 43980 | 35705 | 61251 | 62950 | 53101 | 42674 | 72333 |
| 501 | 863 | 687 | 501 | 824 | 634 | 504 | 844 | 64559 | 51227 | 102310 | 88250 | 60819 | 44364 | 88507 |
| 615 | 867 | 898 | 632 | 906 | 960 | 692 | 924 | 35274 | 23246 | 37810 | 32762 | 26581 | 18711 | 29604 |
| 604 | 984 | 833 | 590 | 989 | 860 | 633 | 1012 | 55483 | 40342 | 66841 | 65738 | 50141 | 35536 | 60220 |
| 611 | 1090 | 595 | 596 | 1087 | 616 | 606 | 1103 | 30512 | 30103 | 49282 | 53710 | 46682 | 46810 | 78515 |
| 762 | 1216 | 1017 | 889 | 1382 | 935 | 743 | 1104 | 25262 | 21526 | 28266 | 34362 | 18100 | 15835 | 17805 |
| 406 | 718 | 662 | 410 | 715 | 642 | 380 | 692 | 75801 | 47396 | 116702 | 83850 | 71474 | 44241 | 107997 |
| 739 | 1240 | 854 | 751 | 1238 | 773 | 660 | 1128 | 51218 | 46452 | 62843 | 77943 | 44787 | 39368 | 52445 |

| | | | | | | | | | | | | | | |
|---|---|---|---|---|---|---|---|---|---|---|---|---|---|---|
| 442 | 861 | 910 | 461 | 878 | 907 | 423 | 862 | 82795 | 41276 | 93344 | 80330 | 90329 | 45757 | 99232 |
| 482 | 960 | 639 | 508 | 965 | 609 | 482 | 949 | 52538 | 43326 | 89832 | 86200 | 66940 | 53246 | 104763 |
| 724 | 1273 | 851 | 704 | 1104 | 848 | 802 | 1282 | 43442 | 40009 | 55239 | 70338 | 56459 | 46671 | 66330 |
| 855 | 1177 | 934 | 848 | 1164 | 910 | 827 | 1191 | 60327 | 54308 | 63542 | 74798 | 81681 | 74118 | 87429 |
| 651 | 1264 | 704 | 681 | 1341 | 800 | 845 | 1319 | 45689 | 43625 | 67016 | 84731 | 43280 | 41887 | 61494 |
| 462 | 725 | 561 | 476 | 759 | 528 | 440 | 717 | 66815 | 55196 | 119526 | 86649 | 49681 | 42156 | 88508 |
| 735 | 1197 | 984 | 750 | 1148 | 1001 | 875 | 1279 | 54998 | 43917 | 59783 | 71568 | 59676 | 45485 | 60653 |
| 611 | 959 | 876 | 644 | 962 | 950 | 737 | 1005 | 50116 | 36857 | 60305 | 57805 | 63929 | 47001 | 72938 |
| 492 | 955 | 710 | 515 | 938 | 655 | 437 | 933 | 62855 | 44289 | 90010 | 85946 | 47724 | 34632 | 67227 |
| 852 | 1542 | 1006 | 882 | 1624 | 902 | 793 | 1400 | 9451 | 8731 | 10243 | 15798 | 8992 | 7883 | 8935 |
| 390 | 811 | 659 | 430 | 852 | 667 | 439 | 833 | 67168 | 42381 | 108584 | 88106 | 60399 | 39350 | 91613 |
| 818 | 1063 | 871 | 735 | 974 | 789 | 632 | 959 | 58928 | 49309 | 60262 | 64067 | 70085 | 59134 | 80442 |
| 523 | 858 | 584 | 502 | 853 | 568 | 467 | 791 | 66845 | 58710 | 112154 | 96221 | 52150 | 44823 | 89303 |
| 697 | 1071 | 762 | 684 | 939 | 787 | 767 | 1073 | 48752 | 47784 | 68546 | 73386 | 49016 | 44011 | 64305 |
| 571 | 863 | 858 | 567 | 881 | 915 | 645 | 930 | 69787 | 46653 | 81677 | 70509 | 66434 | 43909 | 77417 |
| 490 | 775 | 802 | 567 | 837 | 819 | 657 | 852 | 55156 | 38694 | 78968 | 61236 | 56790 | 40154 | 70781 |
| 519 | 985 | 700 | 536 | 1031 | 651 | 511 | 1039 | 53073 | 41330 | 79626 | 78420 | 40423 | 30963 | 57748 |
| 894 | 1394 | 791 | 805 | 1350 | 780 | 825 | 1365 | 56224 | 57759 | 64630 | 90074 | 48714 | 49564 | 61550 |
| 631 | 986 | 992 | 636 | 960 | 1013 | 697 | 1018 | 42368 | 28309 | 44880 | 44235 | 45784 | 29381 | 46171 |
| 394 | 907 | 611 | 414 | 906 | 614 | 417 | 920 | 60466 | 39141 | 99219 | 89988 | 50767 | 34395 | 83074 |
| 642 | 905 | 813 | 642 | 861 | 871 | 698 | 920 | 73395 | 60829 | 94797 | 85804 | 60377 | 47679 | 74229 |
| 551 | 829 | 699 | 546 | 888 | 721 | 567 | 877 | 51199 | 41514 | 75349 | 62472 | 60544 | 47293 | 86673 |
| 630 | 881 | 829 | 627 | 894 | 820 | 638 | 912 | 56276 | 44169 | 70158 | 61814 | 69080 | 52231 | 83305 |
| 753 | 1055 | 1069 | 815 | 1088 | 1104 | 828 | 1085 | 51720 | 36800 | 48843 | 51516 | 53963 | 41143 | 50499 |
| 949 | 1410 | 1127 | 1120 | 1521 | 1093 | 1040 | 1464 | 22557 | 21183 | 22329 | 31474 | 31669 | 31473 | 28106 |
| 665 | 964 | 783 | 625 | 914 | 769 | 647 | 953 | 58654 | 51526 | 77430 | 74631 | 51074 | 40723 | 65198 |
| 557 | 999 | 627 | 456 | 933 | 600 | 426 | 913 | 67577 | 52853 | 94836 | 94744 | 41567 | 30201 | 66252 |
| 376 | 782 | 668 | 351 | 764 | 803 | 440 | 822 | 73476 | 39374 | 104685 | 81883 | 82009 | 43128 | 122812 |
| 308 | 620 | 503 | 343 | 666 | 512 | 335 | 634 | 64458 | 43064 | 140011 | 86778 | 65697 | 44694 | 130489 |
| 389 | 665 | 563 | 501 | 697 | 560 | 450 | 660 | 29121 | 22381 | 57540 | 38275 | 37403 | 33291 | 66431 |
| 580 | 929 | 702 | 645 | 986 | 707 | 640 | 1013 | 55958 | 49534 | 85396 | 79355 | 51528 | 47341 | 73355 |

| | | | | | | | | | | | | | | |
|---|---|---|---|---|---|---|---|---|---|---|---|---|---|---|
| 977 | 1233 | 966 | 958 | 1272 | 873 | 923 | 1219 | 64414 | 66101 | 67626 | 83355 | 48080 | 47709 | 49796 |
| 608 | 911 | 782 | 542 | 874 | 704 | 495 | 835 | 64618 | 45114 | 74142 | 67541 | 63856 | 44274 | 81673 |
| 771 | 1149 | 877 | 766 | 1126 | 901 | 788 | 1121 | 74608 | 65544 | 85045 | 97679 | 53191 | 46453 | 60641 |
| 767 | 1153 | 969 | 796 | 1140 | 905 | 788 | 1145 | 47353 | 40183 | 52385 | 60414 | 56618 | 46488 | 58399 |
| 617 | 1003 | 820 | 670 | 1047 | 765 | 572 | 981 | 63820 | 49410 | 80096 | 80322 | 77205 | 63056 | 94149 |
| 374 | 713 | 481 | 327 | 674 | 550 | 366 | 712 | 49472 | 34073 | 91056 | 64944 | 46614 | 31645 | 96832 |
| 605 | 948 | 791 | 610 | 960 | 826 | 668 | 1017 | 31639 | 24993 | 41327 | 39175 | 73992 | 57042 | 93540 |
| 624 | 972 | 839 | 608 | 951 | 845 | 611 | 941 | 58159 | 43285 | 69339 | 67423 | 48604 | 35195 | 57916 |
| 461 | 846 | 693 | 515 | 911 | 619 | 468 | 894 | 42868 | 29200 | 63320 | 53561 | 19995 | 14871 | 28861 |
| 812 | 1121 | 1030 | 764 | 1105 | 1084 | 849 | 1168 | 84115 | 62210 | 76637 | 85927 | 80112 | 59464 | 77804 |
| 524 | 870 | 673 | 486 | 837 | 646 | 471 | 829 | 48832 | 36945 | 70532 | 61348 | 61275 | 44277 | 91095 |
| 413 | 848 | 757 | 478 | 872 | 723 | 425 | 875 | 60326 | 35243 | 85236 | 72261 | 54147 | 34192 | 71553 |
| 610 | 918 | 788 | 660 | 965 | 721 | 632 | 899 | 45661 | 37063 | 60766 | 55789 | 41654 | 34905 | 52888 |
| 516 | 914 | 626 | 503 | 871 | 658 | 522 | 938 | 46291 | 37484 | 72706 | 66489 | 30596 | 24565 | 48864 |
| 688 | 1161 | 767 | 736 | 1221 | 676 | 685 | 1155 | 41929 | 39796 | 57865 | 67164 | 43504 | 41774 | 56738 |
| 744 | 883 | 1010 | 799 | 900 | 1024 | 812 | 922 | 79792 | 61537 | 82671 | 73026 | 56616 | 44781 | 56077 |
| 442 | 826 | 720 | 451 | 869 | 734 | 476 | 871 | 44056 | 25614 | 57938 | 47872 | 53568 | 33563 | 74408 |
| 777 | 1044 | 860 | 721 | 1000 | 747 | 656 | 960 | 37574 | 34408 | 44263 | 46202 | 12939 | 10845 | 15049 |
| 603 | 895 | 764 | 545 | 890 | 799 | 551 | 912 | 26980 | 19743 | 32746 | 29323 | 41945 | 29926 | 54871 |
| 540 | 941 | 692 | 488 | 923 | 729 | 521 | 903 | 33478 | 25401 | 47062 | 44270 | 39821 | 28060 | 57538 |
| 456 | 738 | 579 | 515 | 785 | 571 | 526 | 830 | 45277 | 37773 | 82853 | 61162 | 46761 | 41575 | 80781 |
| 494 | 776 | 823 | 559 | 811 | 801 | 483 | 749 | 35062 | 21728 | 44000 | 34148 | 52025 | 35347 | 63219 |
| 689 | 996 | 1008 | 807 | 1136 | 1082 | 895 | 1174 | 56056 | 43921 | 63748 | 63470 | 63293 | 50665 | 62813 |
| | | 934 | 401 | 672 | | | | | | | | 7683 | 3303 | 8228 |
| 429 | 842 | 582 | 505 | 859 | 578 | 459 | 859 | 39618 | 32046 | 74628 | 62848 | 46702 | 40474 | 80184 |
| 564 | 889 | 923 | 634 | 884 | 871 | 586 | 856 | 60397 | 39157 | 69442 | 61751 | 43764 | 30038 | 47404 |
| 725 | 1025 | 926 | 708 | 1028 | 917 | 690 | 974 | 36907 | 30087 | 41492 | 42543 | 38602 | 29533 | 41686 |
| 918 | 1226 | 934 | 909 | 1186 | 920 | 875 | 1192 | 36652 | 34268 | 37341 | 45765 | 33763 | 32869 | 36148 |
| 476 | 794 | 776 | 497 | 782 | 722 | 447 | 756 | 55181 | 34766 | 73033 | 58013 | 63994 | 40948 | 82428 |
| 641 | 1317 | 814 | 718 | 1399 | 838 | 710 | 1277 | 32821 | 27517 | 42907 | 56525 | 48963 | 43144 | 60121 |
| 647 | 927 | 957 | 716 | 940 | 1058 | 855 | 1009 | 6461 | 5018 | 7752 | 7185 | 12170 | 9106 | 12711 |

| | | | | | | | | | | | | | | |
|---|---|---|---|---|---|---|---|---|---|---|---|---|---|---|
| 600 | 847 | 744 | 610 | 801 | 778 | 598 | 803 | 36571 | 29407 | 48993 | 41504 | 52253 | 42841 | 70240 |
| 614 | 1105 | 772 | 603 | 1190 | 811 | 651 | 1201 | 49703 | 37168 | 60531 | 66874 | 43358 | 33889 | 56179 |
| 402 | 734 | 633 | 448 | 775 | 617 | 389 | 662 | 46546 | 31297 | 77922 | 57226 | 63732 | 45079 | 100689 |
| 767 | 1377 | 857 | 757 | 1392 | 830 | 751 | 1469 | 43571 | 40116 | 52320 | 72068 | 52291 | 46193 | 60992 |
| 708 | 1112 | 767 | 599 | 949 | 823 | 630 | 972 | 77943 | 67589 | 95431 | 106146 | 55495 | 43368 | 72358 |
| 720 | 1095 | 846 | 721 | 1109 | 872 | 745 | 1115 | 44301 | 36768 | 51060 | 55914 | 31468 | 26803 | 37188 |
| 759 | 1152 | 908 | 796 | 1120 | 900 | 803 | 1183 | 44714 | 39387 | 51887 | 59765 | 57662 | 50584 | 63530 |
| 472 | 968 | 681 | 509 | 994 | 687 | 480 | 951 | 38974 | 26137 | 55338 | 53569 | 29343 | 21942 | 43089 |
| 539 | 717 | 935 | 505 | 687 | 932 | 513 | 712 | 92240 | 53346 | 98908 | 70933 | 89303 | 48205 | 95507 |
| 558 | 975 | 813 | 533 | 909 | 853 | 530 | 901 | 44793 | 30646 | 54886 | 53524 | 44227 | 28997 | 54413 |
| 580 | 863 | 784 | 555 | 853 | 838 | 533 | 835 | 34279 | 21695 | 37391 | 32269 | 28684 | 20312 | 36610 |
| 680 | 927 | 850 | 662 | 938 | 860 | 671 | 927 | 64753 | 51841 | 76244 | 70696 | 54906 | 42726 | 64576 |
| 691 | 1027 | 843 | 651 | 974 | 874 | 710 | 1063 | 23793 | 19358 | 28003 | 28757 | 19563 | 15103 | 23212 |
| 570 | 1206 | 699 | 598 | 1223 | 786 | 752 | 1349 | 53822 | 45712 | 80230 | 96726 | 50770 | 43404 | 72630 |
| 626 | 922 | 793 | 625 | 899 | 798 | 661 | 899 | 56298 | 46793 | 74713 | 68900 | 69988 | 55124 | 88240 |
| 794 | 1197 | 875 | 837 | 1249 | 839 | 845 | 1221 | 40553 | 39042 | 49164 | 58840 | 41051 | 39258 | 46924 |
| 343 | 555 | 433 | 323 | 533 | 560 | 422 | 677 | 32820 | 24382 | 71094 | 39474 | 34558 | 25761 | 79877 |
| 922 | 1131 | 959 | 923 | 1260 | 880 | 820 | 1159 | 33072 | 31282 | 33916 | 38367 | 58518 | 56330 | 61032 |
| 590 | 823 | 643 | 632 | 890 | 582 | 567 | 758 | 63088 | 58487 | 99081 | 81589 | 46592 | 45791 | 72438 |
| 862 | 1374 | 674 | 693 | 1139 | 739 | 861 | 1411 | 35342 | 42014 | 48714 | 66923 | 41929 | 43134 | 62198 |
| 731 | 1027 | 904 | 713 | 1044 | 936 | 746 | 1055 | 87247 | 68127 | 93182 | 95687 | 56709 | 44735 | 62743 |
| 909 | 1322 | 1006 | 836 | 1290 | 1102 | 898 | 1247 | 92953 | 77988 | 85792 | 113411 | 76501 | 63585 | 76057 |
| 407 | 697 | 598 | 366 | 673 | 633 | 380 | 705 | 54776 | 34558 | 84997 | 59246 | 72698 | 44454 | 121515 |
| 731 | 1087 | 869 | 734 | 1090 | 911 | 765 | 1148 | 47910 | 39033 | 53431 | 58097 | 48413 | 40856 | 55688 |
| 677 | 919 | 729 | 609 | 876 | 815 | 696 | 976 | 36771 | 30647 | 45299 | 41625 | 39962 | 33395 | 54817 |
| 555 | 917 | 664 | 494 | 885 | 715 | 542 | 845 | 61452 | 46383 | 83583 | 76636 | 43273 | 32187 | 65214 |
| 686 | 1130 | 696 | 621 | 1101 | 744 | 786 | 1100 | 43479 | 42635 | 62177 | 70229 | 34638 | 30903 | 49801 |
| 450 | 743 | 422 | 428 | 728 | 452 | 431 | 729 | 37277 | 36204 | 80466 | 59786 | 31858 | 32280 | 75464 |
| 803 | 1273 | 961 | 783 | 1263 | 897 | 760 | 1145 | 24437 | 20420 | 25421 | 32359 | 28036 | 22852 | 29185 |
| 606 | 932 | 864 | 620 | 986 | 975 | 789 | 1107 | 20889 | 15089 | 24882 | 23193 | 18229 | 13069 | 21087 |
| 871 | 986 | 1210 | 849 | 940 | 1083 | 737 | 917 | 48773 | 34399 | 39504 | 38957 | 59626 | 41814 | 49259 |

| | | | | | | | | | | | | | | |
|---|---|---|---|---|---|---|---|---|---|---|---|---|---|---|
| 1008 | 1329 | 1097 | 1019 | 1351 | 1168 | 1097 | 1430 | 63376 | 58076 | 57627 | 76603 | 42024 | 39018 | 38297 |
| 869 | 1206 | 1025 | 901 | 1243 | 962 | 863 | 1205 | 64607 | 58342 | 67144 | 80970 | 51332 | 45132 | 50065 |
| 1238 | 1431 | 876 | 753 | 1215 | 1029 | 918 | 1226 | 94130 | 86760 | 70060 | 100272 | 51714 | 44451 | 59047 |
| 528 | 1009 | 643 | 502 | 946 | 707 | 588 | 982 | 39263 | 32875 | 62282 | 62817 | 43417 | 33888 | 67500 |
| 729 | 1138 | 777 | 637 | 1077 | 869 | 785 | 1196 | 37591 | 32660 | 44826 | 51006 | 41668 | 34162 | 53601 |
| 987 | 1268 | 1093 | 1017 | 1251 | 1084 | 1096 | 1234 | 32112 | 28320 | 28697 | 36387 | 37907 | 35273 | 34690 |
| 463 | 870 | 872 | 533 | 908 | 778 | 460 | 859 | 58760 | 33925 | 73285 | 63791 | 44949 | 27504 | 51557 |
| 766 | 1202 | 1024 | 979 | 1365 | 941 | 865 | 1353 | 33224 | 28701 | 37478 | 45040 | 48120 | 45995 | 46979 |
| | | 1007 | 1096 | 1448 | | | | | | | | 2321 | 2525 | 2304 |
| 576 | 895 | 778 | 516 | 827 | 750 | 502 | 817 | 57427 | 40967 | 71152 | 63677 | 52708 | 34927 | 67707 |
| 1192 | 1325 | 864 | 1173 | 1415 | 854 | 1002 | 1241 | 13113 | 18169 | 15243 | 20196 | 14523 | 19714 | 16811 |
| 394 | 696 | 595 | 364 | 649 | 708 | 443 | 753 | 56692 | 35464 | 89905 | 62593 | 60571 | 37038 | 101748 |
| 619 | 788 | 779 | 635 | 780 | 766 | 631 | 988 | 62509 | 49109 | 79374 | 62568 | 59169 | 48247 | 75987 |
| 612 | 886 | 765 | 619 | 886 | 690 | 515 | 834 | 17533 | 13702 | 22405 | 19854 | 13841 | 11201 | 18084 |
| 593 | 1077 | 731 | 607 | 1103 | 681 | 576 | 1116 | 42547 | 35363 | 59665 | 64246 | 54494 | 45248 | 74504 |
| 557 | 855 | 702 | 544 | 816 | 739 | 573 | 837 | 40731 | 32263 | 57874 | 49503 | 45436 | 35190 | 64683 |
| 504 | 893 | 684 | 598 | 927 | 668 | 571 | 934 | 69533 | 56494 | 112030 | 100083 | 59904 | 52308 | 87515 |
| 430 | 710 | 567 | 432 | 735 | 619 | 479 | 802 | 63473 | 49790 | 115732 | 82207 | 29188 | 22218 | 51438 |
| 779 | 1053 | 984 | 732 | 1072 | 1261 | 984 | 1190 | 29750 | 22808 | 29270 | 30830 | 43040 | 32020 | 43719 |
| 850 | 1543 | 773 | 861 | 1574 | 834 | 842 | 1513 | 65887 | 67135 | 79027 | 121956 | 44870 | 49942 | 58019 |
| 613 | 933 | 862 | 632 | 893 | 763 | 537 | 858 | 21040 | 14852 | 24230 | 22604 | 50899 | 37328 | 59041 |
| 724 | 1109 | 867 | 762 | 1125 | 801 | 672 | 970 | 43131 | 37683 | 52080 | 57782 | 41939 | 36887 | 48393 |
| 760 | 1030 | 764 | 720 | 1027 | 823 | 718 | 1027 | 52149 | 47798 | 62857 | 64734 | 41573 | 39197 | 54448 |
| 581 | 1063 | 778 | 572 | 993 | 758 | 631 | 1163 | 45942 | 35614 | 61287 | 65176 | 57565 | 42342 | 73988 |
| 585 | 997 | 687 | 516 | 943 | 640 | 473 | 866 | 4645 | 3807 | 6512 | 6492 | 11303 | 8482 | 16441 |
| 506 | 959 | 806 | 553 | 967 | 773 | 584 | 1029 | 34180 | 24323 | 48097 | 46114 | 38051 | 26129 | 47215 |
| 551 | 936 | 715 | 485 | 901 | 789 | 535 | 895 | 45784 | 31627 | 57369 | 53684 | 49123 | 33328 | 68708 |
| 569 | 925 | 719 | 554 | 912 | 730 | 558 | 934 | 44408 | 34113 | 59997 | 55514 | 49709 | 38284 | 69090 |
| 674 | 998 | 882 | 672 | 1040 | 873 | 655 | 1002 | 52649 | 41755 | 61941 | 61835 | 59299 | 45190 | 67252 |
| 626 | 1005 | 772 | 602 | 982 | 740 | 594 | 973 | 64612 | 52366 | 83615 | 84015 | 31469 | 24552 | 40771 |
| 946 | 1403 | 996 | 929 | 1398 | 1034 | 972 | 1411 | 43901 | 43051 | 45527 | 63863 | 46696 | 43564 | 46897 |

| | | | | | | | | | | | | | | |
|---|---|---|---|---|---|---|---|---|---|---|---|---|---|---|
| 605 | 1064 | 903 | 677 | 1059 | 893 | 644 | 1010 | 41336 | 31451 | 52023 | 55369 | 52011 | 39022 | 57604 |
| 891 | 1189 | 766 | 666 | 1028 | 831 | 704 | 1055 | 49178 | 45837 | 51418 | 61131 | 30484 | 26495 | 39802 |
| 908 | 1234 | 937 | 858 | 1157 | 952 | 874 | 1201 | 46199 | 42525 | 46839 | 57783 | 36994 | 33868 | 39472 |
| 719 | 910 | 984 | 785 | 896 | 877 | 716 | 917 | 63485 | 50326 | 70042 | 63745 | 62295 | 49693 | 63326 |
| 735 | 1134 | 978 | 781 | 1126 | 920 | 705 | 1090 | 49622 | 35345 | 48093 | 54518 | 47685 | 38091 | 48767 |
| 1031 | 1431 | 859 | 913 | 1348 | 885 | 876 | 1283 | 38834 | 42672 | 41393 | 59248 | 31578 | 33549 | 36751 |
| 578 | 1098 | 718 | 671 | 1111 | 650 | 477 | 967 | 28754 | 26643 | 46058 | 50582 | 44278 | 41405 | 61661 |
| 766 | 980 | 1168 | 757 | 993 | 1136 | 697 | 955 | 46810 | 32129 | 41954 | 41099 | 70248 | 45526 | 60124 |
| 629 | 1051 | 672 | 587 | 1035 | 685 | 565 | 1051 | 40767 | 35334 | 56205 | 59060 | 51222 | 44755 | 76279 |
| 728 | 934 | 743 | 578 | 859 | 719 | 536 | 829 | 48644 | 39096 | 53698 | 50129 | 34809 | 27102 | 46857 |
| 703 | 1191 | 861 | 718 | 1219 | 773 | 667 | 1254 | 42852 | 33329 | 47403 | 56476 | 40309 | 33604 | 46813 |
| 622 | 957 | 922 | 804 | 1120 | 786 | 642 | 1013 | 40534 | 32421 | 52120 | 49895 | 56390 | 49172 | 61152 |
| 684 | 939 | 1078 | 751 | 984 | 998 | 772 | 952 | 2688 | 2115 | 3092 | 2903 | 3689 | 2571 | 3423 |
| 437 | 676 | 659 | 439 | 670 | 630 | 428 | 674 | 50531 | 33805 | 77437 | 52333 | 60944 | 40524 | 92416 |
| 476 | 733 | 932 | 522 | 787 | 886 | 531 | 786 | 39193 | 21061 | 44206 | 32396 | 52941 | 29659 | 56801 |
| 507 | 799 | 738 | 525 | 929 | 713 | 505 | 876 | 37403 | 23725 | 46750 | 37375 | 37685 | 26821 | 51070 |
| 571 | 965 | 871 | 705 | 998 | 796 | 634 | 964 | 55275 | 42534 | 74506 | 71883 | 62441 | 50496 | 71653 |
| 810 | 1339 | 840 | 748 | 1233 | 855 | 782 | 1270 | 35090 | 32203 | 39732 | 53207 | 41390 | 36857 | 49269 |
| 636 | 1057 | 729 | 631 | 1130 | 749 | 661 | 1135 | 38627 | 33455 | 52595 | 55617 | 59820 | 51791 | 82103 |
| 654 | 1209 | 779 | 670 | 1181 | 757 | 666 | 1201 | 34812 | 29477 | 45099 | 54543 | 29520 | 25418 | 37911 |
| 533 | 856 | 589 | 648 | 1000 | 608 | 605 | 933 | 52687 | 52043 | 97624 | 83553 | 39611 | 43619 | 67304 |
| 741 | 1868 | 529 | 809 | 1991 | 479 | 748 | 1933 | 24716 | 35887 | 48401 | 90431 | 18391 | 28147 | 34779 |
| 506 | 852 | 859 | 507 | 890 | 1026 | 656 | 951 | 30772 | 18961 | 37458 | 31898 | 31525 | 18623 | 36706 |
| 555 | 945 | 641 | 474 | 855 | 732 | 531 | 904 | 51504 | 38070 | 68643 | 64893 | 35944 | 26600 | 56104 |
| 995 | 1297 | 1096 | 1151 | 1417 | 1084 | 1180 | 1436 | 30915 | 27322 | 27463 | 35630 | 25819 | 27107 | 23548 |
| 471 | 740 | 646 | 541 | 831 | 576 | 439 | 718 | 35230 | 27491 | 58318 | 43127 | 32099 | 26914 | 49718 |
| 493 | 879 | 678 | 466 | 864 | 584 | 456 | 865 | 56966 | 41952 | 85107 | 74844 | 52820 | 36257 | 77855 |
| 462 | 928 | 658 | 473 | 928 | 723 | 503 | 952 | 30935 | 22503 | 48704 | 45196 | 41786 | 30035 | 63526 |
| 779 | 1309 | 906 | 786 | 1239 | 967 | 839 | 1339 | 46638 | 39708 | 50958 | 66715 | 42986 | 37278 | 47453 |
| 662 | 1423 | 972 | 666 | 1320 | 998 | 636 | 1355 | 34863 | 23467 | 35475 | 50491 | 59174 | 40574 | 60905 |
| 471 | 768 | 707 | 565 | 843 | 664 | 505 | 770 | 50003 | 38624 | 81972 | 62976 | 48138 | 38437 | 68085 |

| | | | | | | | | | | | | | | |
|---|---|---|---|---|---|---|---|---|---|---|---|---|---|---|
| 654 | 1036 | 778 | 650 | 1005 | 743 | 604 | 995 | 47254 | 40561 | 62025 | 64281 | 26441 | 22073 | 33966 |
| 486 | 818 | 598 | 431 | 813 | 720 | 561 | 913 | 50602 | 38762 | 79718 | 65189 | 30901 | 22259 | 51657 |
| 839 | 1276 | 809 | 767 | 1231 | 753 | 721 | 1137 | 54670 | 54117 | 64532 | 82320 | 32699 | 31008 | 40408 |
| 812 | 1276 | 964 | 862 | 1265 | 1066 | 1027 | 1487 | 26690 | 22663 | 27906 | 35604 | 51934 | 46430 | 53870 |
| 969 | 1439 | 1218 | 1096 | 1260 | 1036 | 927 | 1394 | 5039 | 4873 | 5027 | 7236 | 1276 | 1148 | 1047 |
| 495 | 828 | 662 | 483 | 811 | 683 | 463 | 755 | 31055 | 21452 | 43298 | 35866 | 56834 | 41469 | 85804 |
| 672 | 1154 | 787 | 626 | 1109 | 816 | 645 | 1096 | 49870 | 40039 | 59613 | 68788 | 46972 | 37332 | 59658 |
| 941 | 1333 | 473 | 328 | 841 | 762 | 639 | 1148 | 866 | 899 | 956 | 1275 | 495 | 343 | 1048 |
| 857 | 1154 | 936 | 796 | 1129 | 923 | 812 | 1084 | 53811 | 47986 | 55984 | 64604 | 59901 | 50931 | 64006 |
| 515 | 815 | 759 | 488 | 785 | 714 | 478 | 800 | 29426 | 19534 | 37949 | 30939 | 30631 | 19667 | 40339 |
| 603 | 922 | 952 | 629 | 906 | 1076 | 636 | 875 | 40946 | 27709 | 45939 | 42348 | 46313 | 30573 | 48628 |
| 769 | 1213 | 1069 | 976 | 1457 | 994 | 970 | 1475 | 31455 | 26382 | 34301 | 41610 | 41953 | 38316 | 39241 |
| 490 | 816 | 768 | 545 | 827 | 675 | 488 | 790 | 48591 | 35692 | 72806 | 59399 | 68119 | 48307 | 88659 |
| 795 | 1167 | 898 | 853 | 1169 | 824 | 868 | 1214 | 43120 | 41903 | 52703 | 61524 | 29674 | 28175 | 33031 |
| 696 | 1023 | 845 | 651 | 994 | 871 | 700 | 1084 | 40410 | 32926 | 47316 | 48395 | 34744 | 26755 | 41097 |
| 860 | 1524 | 810 | 1166 | 2071 | 886 | 979 | 1484 | 9342 | 11297 | 13132 | 20016 | 19511 | 28094 | 24086 |
| 378 | 803 | 490 | 409 | 815 | 541 | 410 | 819 | 33203 | 27110 | 71650 | 57499 | 42892 | 35850 | 87615 |
| 844 | 1263 | 830 | 710 | 1081 | 893 | 840 | 1197 | 33559 | 31893 | 37780 | 47724 | 49741 | 42533 | 59907 |
| 469 | 795 | 693 | 484 | 810 | 672 | 483 | 800 | 24676 | 17036 | 36297 | 28861 | 38743 | 27079 | 55906 |
| 767 | 1206 | 868 | 717 | 1096 | 896 | 802 | 1269 | 26665 | 23483 | 30602 | 36904 | 40622 | 33560 | 46806 |
| 553 | 852 | 1044 | 533 | 804 | 1053 | 535 | 793 | 25289 | 13264 | 24005 | 20453 | 65413 | 33352 | 62627 |
| 739 | 1120 | 897 | 734 | 1111 | 868 | 709 | 1139 | 67522 | 56416 | 76315 | 85505 | 66002 | 53969 | 73562 |
| 479 | 1019 | 607 | 485 | 1016 | 644 | 502 | 1043 | 38860 | 30251 | 63169 | 64345 | 31157 | 24887 | 51330 |
| 558 | 896 | 678 | 540 | 841 | 673 | 534 | 865 | 50590 | 41686 | 74647 | 66896 | 42524 | 33853 | 62687 |
| 496 | 751 | 585 | 469 | 731 | 615 | 543 | 795 | 31398 | 30440 | 61390 | 46102 | 40598 | 32588 | 69428 |
| 843 | 1785 | 853 | 754 | 1574 | 859 | 728 | 1654 | 30561 | 27512 | 32631 | 58263 | 47118 | 41651 | 55256 |
| 574 | 1377 | 895 | 709 | 1500 | 776 | 551 | 1260 | 29223 | 21428 | 37361 | 51445 | 32331 | 25600 | 36106 |
| 507 | 1016 | 704 | 531 | 1004 | 695 | 528 | 1005 | 45101 | 31333 | 61835 | 62813 | 51320 | 38758 | 72929 |
| 845 | 1283 | 953 | 875 | 1313 | 903 | 888 | 1348 | 33068 | 32503 | 38478 | 49378 | 48363 | 44396 | 50760 |
| 825 | 1159 | 964 | 857 | 1105 | 967 | 854 | 1135 | 19413 | 17480 | 21198 | 24562 | 19286 | 17144 | 19998 |
| 576 | 925 | 782 | 647 | 981 | 734 | 599 | 919 | 37834 | 31061 | 53893 | 49834 | 58919 | 48802 | 75388 |

| | | | | | | | | | | | | | | |
|---|---|---|---|---|---|---|---|---|---|---|---|---|---|---|
| 740 | 987 | 1134 | 879 | 1136 | 1069 | 726 | 1036 | 38723 | 28239 | 38185 | 37698 | 54747 | 42444 | 48281 |
| 556 | 912 | 878 | 530 | 882 | 801 | 499 | 786 | 47263 | 29587 | 53191 | 48493 | 57729 | 34848 | 65767 |
| 455 | 1013 | 647 | 452 | 992 | 631 | 437 | 967 | 59311 | 44457 | 97711 | 98965 | 51388 | 35857 | 79380 |
| 712 | 1200 | 815 | 708 | 1183 | 895 | 793 | 1219 | 32609 | 27922 | 39223 | 47081 | 39895 | 34648 | 48968 |
| 733 | 1067 | 967 | 787 | 1158 | 940 | 787 | 1128 | 48782 | 39345 | 53671 | 57267 | 33598 | 27361 | 34762 |
| 2024 | 1920 | 931 | 2280 | 2062 | 816 | 1305 | 1601 | 9110 | 19291 | 9530 | 18294 | 4747 | 11629 | 5101 |
| 569 | 822 | 760 | 579 | 827 | 757 | 570 | 789 | 22411 | 16887 | 29656 | 24380 | 35695 | 27191 | 46944 |
| 797 | 1391 | 914 | 896 | 1417 | 897 | 850 | 1341 | 27510 | 27280 | 34236 | 47630 | 50500 | 49493 | 55238 |
| 449 | 817 | 625 | 461 | 825 | 725 | 555 | 907 | 47777 | 33765 | 75228 | 61441 | 52382 | 38590 | 83762 |
| 1147 | 1398 | 1173 | 1215 | 1389 | 1126 | 1126 | 1247 | 41371 | 42571 | 37125 | 51888 | 24010 | 24874 | 20470 |
| 482 | 842 | 704 | 467 | 815 | 649 | 418 | 772 | 45784 | 31954 | 66280 | 55814 | 62322 | 41339 | 88485 |
| 433 | 844 | 684 | 469 | 860 | 639 | 444 | 817 | 31000 | 21764 | 50265 | 42412 | 16443 | 11265 | 24026 |
| 509 | 896 | 717 | 551 | 981 | 702 | 456 | 854 | 43513 | 31967 | 62816 | 56268 | 32575 | 25069 | 45458 |
| 471 | 792 | 691 | 409 | 759 | 659 | 408 | 750 | 43664 | 27900 | 59229 | 46933 | 47671 | 28241 | 69038 |
| 493 | 839 | 806 | 531 | 843 | 815 | 565 | 865 | 50601 | 34052 | 69128 | 58012 | 49943 | 32910 | 61969 |
| 414 | 865 | 645 | 488 | 950 | 577 | 406 | 937 | 27022 | 17886 | 43167 | 37345 | 36874 | 27897 | 57177 |
| 501 | 900 | 720 | 454 | 862 | 778 | 501 | 873 | 59170 | 34787 | 69418 | 62504 | 50927 | 32101 | 70724 |
| 569 | 930 | 813 | 561 | 942 | 775 | 551 | 885 | 48691 | 36166 | 63528 | 59091 | 54120 | 37296 | 66538 |
| 590 | 903 | 842 | 689 | 1051 | 796 | 710 | 1080 | 21770 | 15928 | 27014 | 24404 | 17966 | 14700 | 21342 |
| 696 | 1270 | 788 | 676 | 1281 | 841 | 743 | 1387 | 28715 | 24181 | 34763 | 44143 | 31423 | 26964 | 39883 |
| 394 | 735 | 589 | 447 | 782 | 581 | 442 | 777 | 40842 | 29149 | 73969 | 54373 | 39471 | 29950 | 67018 |
| 530 | 979 | 748 | 560 | 1102 | 780 | 579 | 1026 | 29685 | 21958 | 41400 | 40512 | 33748 | 25266 | 45130 |
| 488 | 919 | 914 | 492 | 899 | | | | 6790 | 3663 | 7499 | 6894 | 7583 | 4078 | 8295 |
| 1039 | 1463 | 957 | 963 | 1406 | 860 | 814 | 1230 | 19950 | 20716 | 19947 | 29188 | 30458 | 30646 | 31830 |
| 654 | 1065 | 819 | 639 | 1071 | 896 | 699 | 1096 | 34512 | 27072 | 41402 | 44106 | 36754 | 28685 | 44866 |
| 566 | 912 | 681 | 520 | 881 | 690 | 531 | 874 | 31081 | 23756 | 41970 | 38269 | 35838 | 27350 | 52627 |
| 620 | 1207 | 841 | 724 | 1265 | 862 | 716 | 1315 | 19677 | 16498 | 26611 | 32121 | 51707 | 44563 | 61512 |
| 769 | 1067 | 840 | 816 | 1179 | 886 | 855 | 1164 | 44851 | 38063 | 49511 | 52848 | 22716 | 22086 | 27053 |
| 789 | 1293 | 874 | 744 | 1236 | 808 | 766 | 1255 | 44077 | 41277 | 52311 | 67645 | 42260 | 35962 | 48327 |
| 445 | 742 | 638 | 494 | 801 | 642 | 472 | 760 | 27567 | 21228 | 47754 | 35421 | 44747 | 34614 | 70135 |
| 546 | 922 | 495 | 495 | 910 | 553 | 573 | 935 | 30008 | 30855 | 56528 | 52113 | 26392 | 26397 | 53348 |

| | | | | | | | | | | | | | | |
|---|---|---|---|---|---|---|---|---|---|---|---|---|---|---|
| 695 | 1009 | 890 | 805 | 1062 | 802 | 711 | 1067 | 47341 | 41228 | 59364 | 59918 | 47244 | 42720 | 53065 |
| 653 | 1146 | 743 | 654 | 1114 | 761 | 644 | 1109 | 40520 | 36535 | 55993 | 64181 | 47978 | 42289 | 64614 |
| 716 | 1070 | 803 | 595 | 1042 | 834 | 673 | 1144 | 27056 | 22346 | 31204 | 33386 | 20761 | 15384 | 25847 |
| 398 | 662 | 1432 | 339 | 604 | 1273 | 380 | 633 | 62824 | 16633 | 41820 | 27692 | 45970 | 10879 | 32101 |
| 716 | 975 | 875 | 689 | 947 | 937 | 788 | 1037 | 35425 | 29050 | 40564 | 39564 | 34561 | 27223 | 39494 |
| 670 | 937 | 950 | 746 | 1034 | 844 | 578 | 958 | 43029 | 31129 | 46465 | 43555 | 51822 | 40709 | 54538 |
| 692 | 992 | 999 | 736 | 1075 | 952 | 700 | 1007 | 43053 | 30152 | 43592 | 43231 | 41165 | 30338 | 41220 |
| 1090 | 1303 | 930 | 1053 | 1350 | 920 | 982 | 1255 | 32867 | 36095 | 33109 | 43144 | 23487 | 26611 | 25260 |
| 522 | 905 | 773 | 447 | 777 | 687 | 470 | 775 | 38167 | 27766 | 53145 | 48105 | 39370 | 22782 | 50942 |
| 888 | 1102 | 1189 | 970 | 1186 | 1173 | 955 | 1145 | 34581 | 27735 | 31247 | 34419 | 63745 | 52023 | 53619 |
| 1383 | 1883 | 1040 | 1301 | 2045 | 1006 | 1207 | 1900 | 21958 | 28604 | 20676 | 38931 | 23523 | 29425 | 22609 |
| 804 | 1293 | 850 | 816 | 1201 | 891 | 919 | 1207 | 45006 | 44556 | 55395 | 71651 | 37138 | 35632 | 43672 |
| 394 | 812 | 597 | 444 | 928 | 625 | 485 | 943 | 12993 | 9714 | 24657 | 20026 | 27872 | 20732 | 46655 |
| 775 | 1200 | 914 | 800 | 1245 | 864 | 737 | 1216 | 19248 | 16147 | 20832 | 24996 | 34954 | 30604 | 38261 |
| 335 | 831 | 545 | 352 | 836 | 495 | 325 | 831 | 38266 | 24689 | 73597 | 61157 | 24215 | 15661 | 44446 |
| 618 | 899 | 927 | 517 | 833 | 1021 | 608 | 904 | 79068 | 44961 | 72715 | 65361 | 60198 | 33530 | 64912 |
| 1128 | 1191 | 1076 | 924 | 1074 | 1107 | 980 | 1075 | 28261 | 27339 | 24239 | 28869 | 29625 | 25435 | 27525 |
| 528 | 900 | 632 | 492 | 899 | 648 | 528 | 914 | 39425 | 32540 | 61594 | 55423 | 35908 | 27976 | 56815 |
| 546 | 896 | 658 | 528 | 935 | 653 | 511 | 903 | 28734 | 23407 | 42898 | 38418 | 35719 | 28646 | 54258 |
| 248 | 663 | 402 | 255 | 673 | 406 | 236 | 649 | 37641 | 21762 | 87622 | 58126 | 42394 | 26957 | 105568 |
| 465 | 933 | 891 | 518 | 975 | 870 | 495 | 949 | 26261 | 13960 | 30007 | 27994 | 55215 | 32108 | 62003 |
| 507 | 689 | 663 | 447 | 679 | 693 | 467 | 726 | 38106 | 26624 | 52562 | 36204 | 41081 | 27672 | 61928 |
| 466 | 731 | 533 | 425 | 738 | 580 | 482 | 759 | 43727 | 35173 | 75437 | 55177 | 30753 | 24501 | 57693 |
| 628 | 978 | 916 | 600 | 942 | 980 | 706 | 983 | 38227 | 26296 | 41843 | 40922 | 31874 | 20886 | 34813 |
| 515 | 856 | 722 | 495 | 869 | 821 | 529 | 885 | 45778 | 32354 | 62809 | 53749 | 27711 | 18973 | 38367 |
| 313 | 658 | 460 | 308 | 654 | 479 | 278 | 631 | 37307 | 24165 | 77214 | 50825 | 44141 | 29520 | 95897 |
| 600 | 881 | 986 | 632 | 859 | 1148 | 825 | 923 | 40281 | 26263 | 43783 | 38554 | 52780 | 33813 | 53517 |
| 442 | 732 | 652 | 451 | 729 | 617 | 437 | 723 | 28397 | 20800 | 47054 | 34455 | 50734 | 35107 | 77762 |
| 458 | 726 | 620 | 503 | 815 | 642 | 511 | 776 | 32205 | 24220 | 52903 | 38405 | 38314 | 31073 | 61771 |
| 391 | 750 | 623 | 425 | 762 | 596 | 359 | 670 | 78305 | 47592 | 121803 | 91397 | 43170 | 29436 | 69297 |
| 820 | 1438 | 1041 | 1025 | 1653 | 1006 | 909 | 1435 | 21697 | 18388 | 22414 | 32221 | 18532 | 18241 | 17801 |

| | | | | | | | | | | | | | | |
|---|---|---|---|---|---|---|---|---|---|---|---|---|---|---|
| 693 | 981 | 803 | 665 | 946 | 691 | 568 | 857 | 22209 | 18563 | 26775 | 26275 | 30542 | 25302 | 38028 |
| 377 | 802 | 611 | 430 | 882 | 651 | 464 | 884 | 34247 | 21435 | 56853 | 45604 | 23151 | 16283 | 37905 |
| 478 | 777 | 945 | 482 | 792 | 1027 | 506 | 819 | 45849 | 21049 | 44002 | 34180 | 46529 | 23727 | 49218 |
| 656 | 1119 | 872 | 727 | 1177 | 830 | 668 | 1096 | 48864 | 39642 | 60473 | 67674 | 63292 | 52729 | 72555 |
| 472 | 590 | 632 | 542 | 636 | 596 | 490 | 592 | 35984 | 28380 | 60176 | 35508 | 45908 | 39380 | 72671 |
| 479 | 997 | 629 | 501 | 1017 | 748 | 595 | 1051 | 35672 | 29426 | 61459 | 61245 | 21166 | 16863 | 33632 |
| 831 | 1310 | 882 | 780 | 1384 | 1011 | 904 | 1330 | 25588 | 22136 | 26635 | 34881 | 27585 | 24410 | 31286 |
| 506 | 958 | 729 | 572 | 987 | 597 | 427 | 919 | 38085 | 28011 | 55320 | 53013 | 43059 | 33783 | 59064 |
| 771 | 1180 | 984 | 768 | 1210 | 958 | 755 | 1166 | 21738 | 16988 | 22021 | 25991 | 23682 | 18490 | 24060 |
| 1350 | 1253 | 641 | 542 | 920 | 733 | 678 | 1005 | 42707 | 44280 | 32806 | 41110 | 35741 | 30189 | 55736 |
| 704 | 906 | 1146 | 761 | 1010 | 1136 | 823 | 1125 | 2895 | 1832 | 2601 | 2357 | 11377 | 7553 | 9929 |
| 745 | 1068 | 823 | 710 | 1018 | 819 | 672 | 1004 | 36721 | 30957 | 41577 | 44396 | 22859 | 19733 | 27778 |
| 655 | 1063 | 730 | 580 | 978 | 730 | 618 | 986 | 38316 | 32932 | 50315 | 53465 | 35994 | 28594 | 49326 |
| 891 | 1246 | 993 | 953 | 1260 | 1080 | 1065 | 1349 | 47669 | 46321 | 51963 | 64722 | 41775 | 40091 | 42075 |
| 396 | 833 | 496 | 400 | 827 | 566 | 462 | 900 | 30401 | 22569 | 56956 | 47417 | 32918 | 26568 | 66353 |
| 901 | 1373 | 876 | 854 | 1285 | 905 | 983 | 1411 | 30437 | 31271 | 34694 | 47647 | 50448 | 49181 | 57594 |
| 876 | 1513 | 883 | 900 | 1441 | 849 | 860 | 1512 | 25259 | 25692 | 29345 | 44388 | 37831 | 38600 | 42865 |
| 587 | 943 | 643 | 492 | 880 | 657 | 501 | 892 | 39381 | 32349 | 55143 | 51992 | 37140 | 28419 | 57746 |
| 578 | 985 | 972 | 602 | 1043 | 867 | 588 | 957 | 23917 | 15824 | 27371 | 26957 | 16296 | 10092 | 16765 |
| 670 | 1062 | 818 | 656 | 1045 | 745 | 658 | 1089 | 24143 | 21006 | 31366 | 33323 | 30287 | 24314 | 37041 |
| 641 | 1081 | 821 | 631 | 1139 | 773 | 559 | 977 | 34544 | 27067 | 42222 | 45653 | 53106 | 40791 | 64670 |
| 779 | 1174 | 969 | 749 | 1167 | 941 | 686 | 1096 | 64176 | 48807 | 62636 | 73533 | 55366 | 42804 | 57125 |
| 1026 | 1206 | 1040 | 1121 | 1162 | 1091 | 1143 | 1151 | 18492 | 17557 | 17116 | 20640 | 18543 | 19972 | 17821 |
| 686 | 1072 | 1036 | 807 | 1196 | 1039 | 794 | 1153 | 47413 | 32696 | 47680 | 51109 | 31674 | 24666 | 30566 |
| 1272 | 2046 | 909 | 1200 | 2005 | 920 | 1174 | 2067 | 28686 | 38542 | 30303 | 62012 | 34902 | 46053 | 38385 |
| 716 | 1156 | 775 | 614 | 965 | 836 | 667 | 1045 | 52641 | 41773 | 58372 | 67474 | 30879 | 24470 | 39861 |
| 385 | 971 | 698 | 433 | 974 | 761 | 482 | 1048 | 31132 | 17961 | 46696 | 45359 | 35150 | 21794 | 50371 |
| 473 | 614 | 625 | 488 | 666 | 574 | 399 | 677 | 16962 | 14870 | 31425 | 19298 | 19226 | 14997 | 30756 |
| 254 | 738 | 389 | 272 | 742 | 369 | 236 | 701 | 38965 | 25407 | 100170 | 73897 | 31881 | 22330 | 81999 |
| 501 | 892 | 681 | 499 | 912 | 673 | 468 | 872 | 39476 | 26796 | 53520 | 47740 | 39634 | 29064 | 58230 |
| 693 | 1093 | 1052 | 787 | 1179 | 936 | 716 | 1144 | 24863 | 18586 | 26803 | 29299 | 28266 | 21146 | 26861 |

| | | | | | | | | | | | | | | |
|---|---|---|---|---|---|---|---|---|---|---|---|---|---|---|
| 673 | 1054 | 828 | 708 | 1030 | 875 | 801 | 1104 | 40231 | 36417 | 54089 | 57021 | 30503 | 26057 | 36818 |
| 580 | 782 | 770 | 708 | 833 | 773 | 677 | 785 | 44430 | 36831 | 63491 | 49660 | 67504 | 62106 | 87670 |
| 869 | 1296 | 992 | 858 | 1249 | 990 | 834 | 1293 | 21383 | 18075 | 20788 | 26940 | 30827 | 26663 | 31064 |
| 564 | 1076 | 797 | 538 | 1021 | 785 | 522 | 1006 | 51323 | 36653 | 64996 | 69932 | 46164 | 31163 | 57922 |
| 837 | 1272 | 920 | 877 | 1229 | 915 | 851 | 1193 | 27577 | 25905 | 30944 | 39360 | 39886 | 38015 | 43340 |
| 417 | 874 | 675 | 395 | 822 | 694 | 422 | 878 | 38123 | 23936 | 57363 | 50114 | 40748 | 23842 | 60337 |
| 813 | 1121 | 950 | 820 | 1166 | 873 | 726 | 1097 | 51881 | 43542 | 53567 | 60073 | 21679 | 18709 | 22820 |
| 284 | 670 | 535 | 273 | 651 | 543 | 293 | 694 | 32056 | 17514 | 61632 | 41324 | 37993 | 19409 | 70997 |
| 844 | 1263 | 776 | 724 | 1127 | 889 | 857 | 1238 | 22744 | 20895 | 24763 | 31275 | 22352 | 20854 | 28794 |
| 766 | 1189 | 806 | 557 | 1039 | 757 | 622 | 1032 | 29007 | 24639 | 32185 | 38264 | 52420 | 36223 | 65061 |
| 465 | 742 | 787 | 575 | 941 | 835 | 612 | 929 | 46429 | 25667 | 55255 | 41003 | 27651 | 20226 | 35147 |
| 614 | 1162 | 779 | 618 | 1160 | 780 | 644 | 1133 | 40564 | 31345 | 51048 | 59318 | 31682 | 25143 | 40673 |
| 672 | 823 | 827 | 549 | 812 | 887 | 612 | 890 | 32062 | 23186 | 34484 | 28368 | 47144 | 31295 | 56975 |
| 456 | 845 | 644 | 441 | 840 | 600 | 383 | 808 | 14606 | 10048 | 22012 | 18601 | 31477 | 21574 | 48913 |
| 488 | 820 | 667 | 521 | 872 | 615 | 494 | 839 | 24270 | 18651 | 38226 | 31352 | 33673 | 26316 | 50470 |
| 430 | 731 | 382 | 390 | 749 | 302 | 356 | 736 | 18618 | 21232 | 49324 | 36061 | 19241 | 19639 | 50349 |
| 703 | 964 | 810 | 770 | 1006 | 750 | 688 | 956 | 17543 | 15860 | 22569 | 21749 | 27820 | 26451 | 34336 |
| 629 | 922 | 790 | 651 | 949 | 745 | 624 | 991 | 40575 | 33426 | 53117 | 48959 | 21267 | 17523 | 26906 |
| | | | | | 696 | 554 | 1138 | | | | | | | |
| 899 | 1359 | 913 | 897 | 1296 | 1007 | 912 | 1277 | 43184 | 41246 | 45893 | 62353 | 45085 | 44309 | 49377 |
| 466 | 786 | 645 | 506 | 860 | 606 | 461 | 808 | 36418 | 28648 | 61458 | 48282 | 30947 | 24265 | 47963 |
| 678 | 930 | 889 | 709 | 989 | 763 | 576 | 867 | 43453 | 33491 | 49432 | 45970 | 26039 | 20776 | 29291 |
| 548 | 922 | 735 | 531 | 946 | 760 | 554 | 970 | 42115 | 30767 | 56139 | 51757 | 30499 | 22041 | 41522 |
| 930 | 1066 | 992 | 860 | 1087 | 957 | 749 | 945 | 19031 | 16203 | 17416 | 18572 | 14541 | 12617 | 14664 |
| 947 | 1108 | 1097 | 831 | 1075 | 1078 | 799 | 1055 | 5071 | 4355 | 4596 | 5093 | 10594 | 8024 | 9658 |
| 828 | 1201 | 899 | 697 | 1169 | 893 | 718 | 1225 | 43390 | 35353 | 42714 | 51280 | 46532 | 36072 | 51786 |
| 565 | 882 | 624 | 490 | 812 | 637 | 417 | 733 | 28586 | 23081 | 40853 | 36021 | 32909 | 25845 | 52762 |
| 472 | 864 | 728 | 453 | 853 | 819 | 502 | 861 | 43222 | 26650 | 56425 | 48769 | 29162 | 18134 | 40035 |
| 360 | 1006 | 436 | 688 | 943 | 419 | 487 | 1079 | 16860 | 11020 | 30570 | 30748 | 8678 | 13679 | 19887 |
| 500 | 962 | 585 | 489 | 947 | 614 | 477 | 911 | 35972 | 29770 | 59543 | 57305 | 38558 | 32264 | 65947 |
| 839 | 1143 | 1140 | 852 | 1186 | 1150 | 905 | 1200 | 48196 | 36946 | 44061 | 50368 | 27636 | 20661 | 24242 |

| | | | | | | | | | | | | | | |
|---|---|---|---|---|---|---|---|---|---|---|---|---|---|---|
| 973 | 1407 | 1092 | 960 | 1408 | 1036 | 964 | 1474 | 31507 | 28366 | 29150 | 41013 | 30988 | 27240 | 28375 |
| 578 | 847 | 646 | 588 | 830 | 714 | 604 | 854 | 51779 | 43497 | 75246 | 63738 | 24693 | 22448 | 38199 |
| 298 | 553 | 511 | 323 | 562 | 422 | 269 | 520 | 16306 | 10748 | 36089 | 19952 | 24805 | 15692 | 48524 |
| 431 | 835 | 575 | 440 | 876 | 567 | 424 | 830 | 23330 | 16182 | 37561 | 31375 | 34250 | 26204 | 59556 |
| 530 | 906 | 653 | 510 | 882 | 662 | 523 | 928 | 39907 | 31671 | 59706 | 54087 | 41598 | 32496 | 63689 |
| 855 | 1340 | 984 | 882 | 1375 | 966 | 873 | 1362 | 48840 | 42919 | 50221 | 67290 | 44635 | 40007 | 45349 |
| 606 | 812 | 904 | 680 | 872 | 831 | 638 | 866 | 54834 | 36650 | 60490 | 49128 | 55700 | 41854 | 61586 |
| 608 | 1058 | 954 | 654 | 1024 | 854 | 625 | 1045 | 21222 | 14297 | 23503 | 24867 | 18627 | 12781 | 19534 |
| 517 | 867 | 769 | 543 | 897 | 761 | 550 | 896 | 30925 | 22058 | 42682 | 36986 | 31115 | 21951 | 40447 |
| 887 | 1433 | 894 | 752 | 1339 | 879 | 746 | 1339 | 29860 | 25684 | 28951 | 41499 | 40905 | 34415 | 45777 |
| 446 | 802 | 728 | 418 | 804 | 797 | 494 | 919 | 35882 | 20264 | 45399 | 36401 | 38310 | 22001 | 52605 |
| 771 | 1119 | 800 | 803 | 1131 | 777 | 761 | 1103 | 33744 | 34378 | 44565 | 49885 | 31357 | 31465 | 39207 |
| 1152 | 1579 | 1042 | 1024 | 1476 | 1034 | 1087 | 1454 | 15674 | 15634 | 13572 | 21431 | 15781 | 15499 | 15142 |
| 631 | 1054 | 804 | 519 | 906 | 800 | 578 | 913 | 23729 | 19278 | 30536 | 32189 | 32936 | 21261 | 40959 |
| 893 | 1149 | 853 | 805 | 1213 | 855 | 870 | 1273 | 35134 | 34378 | 38501 | 44239 | 19667 | 18569 | 23062 |
| 585 | 1070 | 747 | 649 | 1103 | 773 | 620 | 999 | 12026 | 10680 | 18260 | 19534 | 22950 | 19927 | 30711 |
| 760 | 1272 | 905 | 755 | 1201 | 941 | 816 | 1340 | 23815 | 20758 | 27312 | 34747 | 24426 | 20371 | 26981 |
| 604 | 961 | 944 | 525 | 855 | 920 | 534 | 901 | 25398 | 15296 | 25318 | 24336 | 25719 | 14286 | 27235 |
| 475 | 601 | 390 | 437 | 567 | 354 | 455 | 606 | 25663 | 28687 | 60420 | 36301 | 18173 | 20371 | 46609 |
| 552 | 976 | 713 | 656 | 1100 | 718 | 551 | 906 | 34843 | 26683 | 48320 | 47184 | 24699 | 22719 | 34634 |
| 563 | 705 | 740 | 589 | 752 | 688 | 522 | 720 | 27090 | 22320 | 39631 | 27925 | 39171 | 31163 | 52901 |
| 1072 | 1516 | 1176 | 1183 | 1588 | 1087 | 1056 | 1520 | 22222 | 22104 | 20622 | 31266 | 35083 | 35295 | 29831 |
| 573 | 899 | 730 | 525 | 900 | 759 | 580 | 967 | 22258 | 16056 | 28002 | 25179 | 41205 | 29613 | 56457 |
| 647 | 910 | 788 | 647 | 959 | 639 | 529 | 865 | 23154 | 19306 | 29861 | 27168 | 16065 | 13194 | 20395 |
| 778 | 1102 | 932 | 711 | 1122 | 955 | 783 | 1235 | 23340 | 18863 | 24237 | 26709 | 26672 | 20336 | 28613 |
| 558 | 1068 | 831 | 669 | 1268 | 787 | 659 | 1252 | 44297 | 31779 | 56990 | 60868 | 30569 | 24614 | 36792 |
| 738 | 1106 | 960 | 881 | 1262 | 923 | 892 | 1279 | 39508 | 34260 | 46437 | 51362 | 37657 | 34556 | 39211 |
| 531 | 870 | 648 | 511 | 863 | 600 | 497 | 828 | 34978 | 28421 | 53493 | 46538 | 36073 | 28439 | 55672 |
| 660 | 1079 | 929 | 766 | 1203 | 986 | 897 | 1248 | 38694 | 31544 | 47818 | 51586 | 22911 | 18909 | 24671 |
| 608 | 988 | 748 | 613 | 961 | 770 | 604 | 977 | 34546 | 26324 | 43276 | 42762 | 25842 | 21175 | 34541 |
| 541 | 1199 | 720 | 633 | 1265 | 734 | 611 | 1144 | 28127 | 23973 | 44313 | 53126 | 19447 | 17102 | 27025 |

| | | | | | | | | | | | | | | |
|---|---|---|---|---|---|---|---|---|---|---|---|---|---|---|
| 486 | 969 | 814 | 579 | 1039 | 714 | 554 | 1062 | 41141 | 29930 | 61625 | 59711 | 32051 | 22790 | 39371 |
| 477 | 719 | 569 | 338 | 562 | 691 | 493 | 780 | 22564 | 17147 | 35960 | 25860 | 24686 | 14660 | 43389 |
| 795 | 1148 | 965 | 824 | 1293 | 873 | 717 | 1172 | 10785 | 9381 | 11795 | 13538 | 18442 | 15746 | 19119 |
| 620 | 932 | 850 | 609 | 968 | 841 | 596 | 902 | 39662 | 28904 | 46584 | 43405 | 25136 | 17996 | 29555 |
| 722 | 1252 | 890 | 755 | 1322 | 900 | 765 | 1349 | 35018 | 29798 | 41245 | 51651 | 21877 | 18561 | 24577 |
| 774 | 1003 | 918 | 709 | 968 | 816 | 603 | 896 | 13223 | 10761 | 13895 | 13935 | 22288 | 17229 | 24289 |
| 572 | 745 | 531 | 586 | 729 | 523 | 626 | 789 | 13895 | 15898 | 27772 | 20698 | 12594 | 13911 | 23725 |
| 567 | 1302 | 796 | 633 | 1334 | 656 | 489 | 1220 | 38510 | 30500 | 53816 | 70074 | 43572 | 34659 | 54729 |
| 536 | 835 | 756 | 527 | 840 | 752 | 535 | 841 | 28024 | 19468 | 36333 | 30341 | 41154 | 28726 | 54466 |
| 726 | 930 | 876 | 699 | 892 | 844 | 672 | 873 | 23782 | 18866 | 25975 | 24157 | 27247 | 21752 | 31111 |
| 708 | 1174 | 803 | 732 | 1247 | 750 | 632 | 1192 | 21672 | 17956 | 25363 | 29785 | 28010 | 25519 | 34869 |
| 653 | 820 | 987 | 585 | 802 | 916 | 541 | 796 | 34084 | 21866 | 33499 | 27482 | 31197 | 18506 | 31620 |
| 871 | 1130 | 818 | 694 | 1050 | 803 | 686 | 1077 | 34820 | 30638 | 35181 | 39759 | 20863 | 17705 | 25494 |
| 604 | 1069 | 845 | 754 | 1136 | 732 | 545 | 939 | 36614 | 29378 | 48608 | 51951 | 25052 | 22354 | 29653 |
| 1194 | 1470 | 1085 | 1217 | 1439 | 961 | 1048 | 1357 | 32208 | 35732 | 29920 | 43973 | 24438 | 27414 | 22533 |
| 512 | 855 | 658 | 561 | 909 | 644 | 514 | 871 | 30858 | 24650 | 48125 | 41137 | 27435 | 23420 | 41718 |
| 528 | 808 | 601 | 488 | 777 | 660 | 528 | 773 | 40965 | 34595 | 65508 | 52961 | 32994 | 26784 | 54915 |
| 393 | 729 | 717 | 386 | 669 | 695 | 368 | 666 | 31087 | 17389 | 44266 | 32254 | 30733 | 16538 | 42875 |
| 719 | 972 | 703 | 740 | 1033 | 740 | 705 | 976 | 24453 | 26225 | 36492 | 35475 | 22804 | 24014 | 32460 |
| 313 | 525 | 455 | 334 | 521 | 519 | 378 | 585 | 23859 | 19529 | 62398 | 32731 | 30323 | 22282 | 66654 |
| 581 | 1081 | 862 | 630 | 1048 | 1141 | 1022 | 1364 | 34129 | 23070 | 39688 | 42907 | 64502 | 47165 | 74815 |
| 855 | 1230 | 1044 | 1020 | 1252 | 869 | 825 | 1235 | 37348 | 36262 | 42430 | 52210 | 39999 | 39086 | 38313 |
| 935 | 1422 | 1101 | 1053 | 1511 | 1123 | 994 | 1361 | 6877 | 5900 | 6311 | 8976 | 29238 | 27976 | 26566 |
| 395 | 658 | 775 | 325 | 589 | 810 | 500 | 740 | 16501 | 8542 | 21602 | 14206 | 20914 | 8779 | 26992 |
| 414 | 854 | 524 | 444 | 833 | 537 | 449 | 889 | 23901 | 20497 | 49540 | 42296 | 31165 | 26400 | 59463 |
| 745 | 1028 | 900 | 808 | 1055 | 950 | 908 | 1108 | 9192 | 7352 | 9863 | 10142 | 21850 | 19603 | 24270 |
| 610 | 992 | 736 | 646 | 1076 | 799 | 627 | 994 | 10543 | 8886 | 14564 | 14446 | 19958 | 17522 | 27118 |
| 611 | 934 | 717 | 553 | 921 | 775 | 607 | 971 | 23292 | 19721 | 32261 | 30139 | 25625 | 19768 | 35745 |
| 882 | 1022 | 1126 | 677 | 866 | 1155 | 685 | 902 | 54270 | 33219 | 37672 | 38494 | 43740 | 26325 | 38862 |
| 487 | 784 | 717 | 419 | 726 | 707 | 496 | 789 | 19754 | 12988 | 26642 | 20896 | 38087 | 22250 | 53083 |
| 478 | 665 | 529 | 440 | 702 | 543 | 444 | 664 | 5288 | 5204 | 10882 | 7235 | 21591 | 17943 | 40790 |

| | | | | | | | | | | | | | | |
|---|---|---|---|---|---|---|---|---|---|---|---|---|---|---|
| 774 | 1194 | 986 | 856 | 1259 | 1003 | 902 | 1260 | 45934 | 39522 | 51051 | 60977 | 42779 | 37117 | 43372 |
| 1362 | 1216 | 1104 | 948 | 1089 | 1370 | 1254 | 1188 | 42508 | 38453 | 28236 | 34341 | 21727 | 18664 | 19685 |
| 485 | 738 | 604 | 464 | 712 | 670 | 513 | 745 | 37635 | 29020 | 59865 | 44192 | 35613 | 27336 | 58963 |
| 864 | 1029 | 1156 | 950 | 1039 | 1134 | 989 | 1035 | 33538 | 27777 | 32134 | 33065 | 35201 | 28939 | 30454 |
| 387 | 647 | 317 | 374 | 607 | 286 | 390 | 605 | 22085 | 30737 | 79362 | 51350 | 14076 | 16651 | 44470 |
| 676 | 1140 | 1043 | 734 | 1212 | 922 | 628 | 1103 | 42292 | 29692 | 43954 | 50105 | 40648 | 28605 | 38984 |
| 951 | 1476 | 856 | 925 | 1445 | 979 | 998 | 1383 | 25374 | 26481 | 27856 | 41117 | 27159 | 29348 | 31733 |
| 942 | 1292 | 1006 | 1041 | 1342 | 806 | 838 | 1121 | 35238 | 36047 | 38266 | 49447 | 50285 | 52032 | 50002 |
| 608 | 1147 | 736 | 608 | 1126 | 692 | 601 | 1144 | 42571 | 36970 | 60778 | 69690 | 27747 | 22928 | 37708 |
| 819 | 1246 | 765 | 643 | 1051 | 818 | 763 | 1224 | 35081 | 32690 | 39922 | 49756 | 23937 | 20101 | 31273 |
| 505 | 950 | 792 | 570 | 968 | 776 | 569 | 978 | 34683 | 24566 | 48632 | 46205 | 48167 | 34650 | 60827 |
| 558 | 845 | 605 | 469 | 770 | 641 | 511 | 819 | 26940 | 22526 | 40338 | 34101 | 46773 | 36247 | 77253 |
| 508 | 753 | 888 | 460 | 765 | 852 | 438 | 738 | 47036 | 24474 | 48178 | 36261 | 28505 | 14776 | 32116 |
| 686 | 954 | 891 | 731 | 912 | 940 | 810 | 1022 | 24433 | 17924 | 26133 | 24936 | 39041 | 32015 | 43795 |
| 676 | 1315 | 904 | 696 | 1249 | 940 | 758 | 1297 | 16154 | 13070 | 19330 | 25411 | 21376 | 16461 | 23651 |
| 743 | 1022 | 800 | 597 | 926 | 863 | 660 | 969 | 19314 | 15101 | 20335 | 20785 | 27036 | 20182 | 33811 |
| 371 | 579 | 305 | 320 | 556 | 418 | 398 | 632 | 22322 | 19175 | 51731 | 29943 | 20006 | 20988 | 65584 |
| 605 | 1134 | 653 | 508 | 963 | 675 | 560 | 1035 | 32931 | 28650 | 47336 | 53681 | 11445 | 8907 | 17529 |
| 962 | 1268 | 1112 | 990 | 1309 | 1105 | 970 | 1279 | 26600 | 23210 | 24116 | 30580 | 23493 | 20902 | 21119 |
| 429 | 796 | 599 | 454 | 834 | 614 | 492 | 869 | 31453 | 21499 | 50141 | 39931 | 34082 | 25825 | 56903 |
| 411 | 607 | 672 | 446 | 631 | 684 | 411 | 567 | 33137 | 20979 | 51099 | 31036 | 32607 | 21624 | 48508 |
| 406 | 891 | 615 | 389 | 764 | 637 | 441 | 850 | 16108 | 11844 | 29198 | 26011 | 34580 | 21884 | 56268 |
| 667 | 1110 | 830 | 722 | 1102 | 891 | 690 | 1194 | 35956 | 30120 | 45145 | 50112 | 20392 | 17748 | 24571 |
| 788 | 1263 | 852 | 918 | 1462 | 916 | 923 | 1433 | 37197 | 36211 | 45935 | 57999 | 29433 | 31697 | 34531 |
| 886 | 1085 | 782 | 833 | 1145 | 714 | 815 | 1067 | 24147 | 27110 | 30589 | 33190 | 10251 | 10925 | 13108 |
| 1003 | 1223 | 915 | 729 | 1121 | 923 | 621 | 966 | 27742 | 26391 | 26323 | 32183 | 40374 | 32170 | 44129 |
| 257 | 569 | 293 | 274 | 571 | 208 | 198 | 474 | 13773 | 12427 | 48330 | 27484 | 16817 | 15745 | 57478 |
| 392 | 715 | 665 | 441 | 738 | 571 | 387 | 699 | 26998 | 18608 | 47462 | 33927 | 30513 | 20255 | 45902 |
| 469 | 821 | 646 | 467 | 758 | 684 | 544 | 850 | 36293 | 28094 | 59955 | 49229 | 29828 | 21560 | 46142 |
| 526 | 900 | 681 | 523 | 877 | 772 | 545 | 927 | 32197 | 23487 | 44694 | 40227 | 42814 | 32851 | 62840 |
| 882 | 1383 | 890 | 829 | 1283 | 952 | 999 | 1496 | 37987 | 37779 | 42821 | 59239 | 19950 | 18573 | 22408 |

| | | | | | | | | | | | | | | |
|---|---|---|---|---|---|---|---|---|---|---|---|---|---|---|
| 708 | 1079 | 944 | 819 | 1156 | 929 | 769 | 1058 | 39209 | 30472 | 43055 | 46436 | 31734 | 27528 | 33620 |
| 566 | 1043 | 665 | 540 | 1106 | 711 | 575 | 1039 | 27019 | 22664 | 40017 | 41730 | 26617 | 21639 | 40053 |
| 521 | 767 | 928 | 596 | 704 | 761 | 523 | 857 | 9870 | 6391 | 12276 | 9415 | 4909 | 3151 | 5289 |
| 425 | 770 | 719 | 476 | 827 | 613 | 443 | 825 | 47793 | 29553 | 69571 | 53542 | 42135 | 27874 | 58608 |
| 268 | 974 | 443 | 232 | 961 | 541 | 294 | 981 | 27336 | 15239 | 56808 | 55307 | 24671 | 12886 | 55638 |
| 574 | 987 | 831 | 561 | 992 | 841 | 604 | 998 | 38760 | 28343 | 49351 | 48722 | 29583 | 19966 | 35587 |
| 451 | 830 | 802 | 437 | 808 | 814 | 462 | 828 | 15238 | 8430 | 18706 | 15521 | 16708 | 9114 | 20843 |
| 454 | 752 | 680 | 483 | 783 | 661 | 485 | 796 | 29595 | 19741 | 43446 | 32674 | 36193 | 25681 | 53193 |
| 679 | 1078 | 834 | 706 | 1043 | 878 | 732 | 1033 | 11733 | 9596 | 14142 | 15239 | 27244 | 23050 | 32670 |
| 517 | 1220 | 716 | 463 | 1248 | 704 | 492 | 1306 | 13856 | 8757 | 16953 | 20683 | 23769 | 15389 | 33220 |
| 713 | 1087 | 774 | 713 | 1134 | 733 | 638 | 1043 | 25695 | 22666 | 31794 | 34572 | 34030 | 31354 | 43958 |
| 573 | 966 | 747 | 665 | 972 | 779 | 658 | 991 | 16196 | 12610 | 21997 | 21245 | 33355 | 29666 | 44644 |
| 774 | 1084 | 771 | 677 | 1030 | 729 | 591 | 980 | 27632 | 23439 | 30296 | 32853 | 16474 | 14453 | 21358 |
| 500 | 832 | 652 | 513 | 856 | 666 | 513 | 825 | 42448 | 32202 | 64460 | 53628 | 34022 | 26790 | 52212 |
| 685 | 1180 | 675 | 556 | 1125 | 751 | 743 | 1151 | 38023 | 32457 | 47392 | 55935 | 22545 | 18568 | 33406 |
| 632 | 1068 | 721 | 575 | 942 | 637 | 562 | 1086 | 33804 | 26425 | 41799 | 44642 | 37034 | 29534 | 51393 |
| 409 | 679 | 607 | 346 | 637 | 687 | 402 | 660 | 21204 | 12846 | 31413 | 21318 | 18487 | 10550 | 30460 |
| 527 | 863 | 929 | 543 | 877 | 947 | 563 | 882 | 40359 | 22910 | 43473 | 37533 | 38082 | 22261 | 40994 |
| 641 | 1058 | 900 | 636 | 1019 | 845 | 648 | 1132 | 23839 | 17766 | 27702 | 29312 | 28740 | 20332 | 31947 |
| 748 | 1019 | 936 | 682 | 962 | 957 | 683 | 955 | 31220 | 23668 | 31648 | 32253 | 35792 | 26093 | 38251 |
| 462 | 721 | 964 | 513 | 762 | 909 | 527 | 704 | 35227 | 17362 | 37584 | 27107 | 38680 | 20581 | 40138 |
| 457 | 748 | 934 | 417 | 703 | 937 | 428 | 727 | 34001 | 17526 | 38357 | 28682 | 21930 | 9789 | 23467 |
| 585 | 1004 | 694 | 461 | 912 | 629 | 370 | 831 | 32645 | 23761 | 40596 | 40756 | 23008 | 15287 | 33140 |
| 446 | 854 | 842 | 516 | 946 | 745 | 468 | 929 | 29266 | 15651 | 35102 | 29984 | 20652 | 12648 | 24520 |
| 489 | 787 | 752 | 426 | 719 | 687 | 396 | 714 | 34471 | 21964 | 44930 | 35354 | 49491 | 28022 | 65827 |
| 569 | 861 | 710 | 546 | 860 | 727 | 566 | 897 | 24596 | 19228 | 33777 | 29086 | 40866 | 31419 | 57549 |
| 930 | 1422 | 1010 | 1187 | 1675 | 1012 | 1244 | 1652 | 29459 | 32706 | 35164 | 50017 | 22013 | 25862 | 21791 |
| 831 | 1181 | 1035 | 882 | 1220 | 1060 | 939 | 1306 | 21450 | 16596 | 19961 | 23566 | 26638 | 22691 | 25738 |
| 755 | 4695 | 876 | 594 | 4285 | 767 | 483 | 3021 | 12381 | 10651 | 14114 | 66263 | 17265 | 11711 | 19709 |
| 773 | 1151 | 937 | 751 | 1068 | 887 | 758 | 1074 | 11176 | 9743 | 12605 | 14509 | 35836 | 28702 | 38240 |
| 503 | 987 | 604 | 471 | 826 | 592 | 478 | 943 | 23978 | 18743 | 37270 | 36785 | 18253 | 14236 | 30212 |

| | | | | | | | | | | | | | | |
|---|---|---|---|---|---|---|---|---|---|---|---|---|---|---|
| 669 | 1102 | 886 | 726 | 1127 | 821 | 646 | 1171 | 35850 | 29863 | 44660 | 49208 | 21026 | 17237 | 23740 |
| 425 | 845 | 533 | 398 | 914 | 585 | 410 | 879 | 14709 | 10331 | 24301 | 20528 | 12738 | 9530 | 23921 |
| 821 | 1256 | 925 | 824 | 1206 | 883 | 774 | 1151 | 14962 | 14201 | 17292 | 21726 | 12316 | 10971 | 13318 |
| 460 | 892 | 750 | 576 | 957 | 871 | 642 | 979 | 22418 | 14609 | 31787 | 28348 | 24313 | 18683 | 32435 |
| 708 | 1117 | 816 | 655 | 1036 | 830 | 662 | 1096 | 31831 | 28519 | 40284 | 44985 | 29601 | 23773 | 36287 |
| 691 | 1056 | 818 | 682 | 1147 | 827 | 678 | 970 | 13849 | 11602 | 16786 | 17731 | 21745 | 18128 | 26576 |
| 941 | 892 | 1194 | 819 | 878 | 1271 | 838 | 903 | 44802 | 32150 | 34168 | 30490 | 34637 | 23764 | 29013 |
| 435 | 715 | 619 | 442 | 712 | 574 | 455 | 760 | 19780 | 14035 | 32289 | 23074 | 30967 | 22079 | 50009 |
| 646 | 908 | 976 | 489 | 777 | 1067 | 591 | 877 | 32884 | 21103 | 32668 | 29674 | 17736 | 8881 | 18172 |
| 723 | 1155 | 860 | 668 | 1105 | 877 | 675 | 1101 | 25480 | 19384 | 26827 | 30986 | 34291 | 26638 | 39864 |
| 384 | 756 | 634 | 392 | 731 | 768 | 427 | 819 | 26945 | 16814 | 43788 | 33085 | 24142 | 14920 | 38090 |
| 355 | 618 | 561 | 394 | 650 | 634 | 437 | 699 | 25340 | 16970 | 47736 | 29513 | 26518 | 18631 | 47284 |
| 1136 | 1676 | 1207 | 983 | 1541 | 1230 | 924 | 1149 | 2467 | 2276 | 2005 | 3360 | 4382 | 3569 | 3630 |
| 630 | 1195 | 666 | 583 | 1029 | 733 | 751 | 1296 | 21263 | 19331 | 30675 | 36665 | 18068 | 15811 | 27111 |
| 706 | 918 | 806 | 719 | 943 | 743 | 592 | 905 | 23412 | 20545 | 29096 | 26704 | 31461 | 28060 | 39017 |
| 663 | 817 | 970 | 657 | 909 | 935 | 621 | 784 | 49078 | 33779 | 50949 | 41611 | 24346 | 16475 | 25087 |
| 720 | 1042 | 1024 | 651 | 984 | 965 | 591 | 889 | 21429 | 14444 | 20073 | 20909 | 20494 | 13027 | 20019 |
| 419 | 758 | 765 | 465 | 750 | 741 | 400 | 715 | 10400 | 5937 | 14172 | 10736 | 28098 | 17063 | 36710 |
| 489 | 769 | 596 | 502 | 810 | 641 | 493 | 773 | 25413 | 18419 | 37693 | 29001 | 34839 | 29353 | 58419 |
| 489 | 848 | 1015 | 471 | 832 | 982 | 487 | 807 | 44957 | 21242 | 43417 | 36831 | 46229 | 21459 | 45566 |
| 567 | 977 | 895 | 625 | 999 | 738 | 590 | 1010 | 23881 | 18406 | 32433 | 31672 | 20704 | 14454 | 23131 |
| 710 | 1033 | 997 | 705 | 1230 | 1192 | 823 | 1381 | 30773 | 21237 | 29921 | 30918 | 27539 | 19479 | 27633 |
| 669 | 1031 | 876 | 677 | 1009 | 934 | 699 | 1087 | 25215 | 19088 | 28531 | 29416 | 24055 | 18596 | 27448 |
| 607 | 749 | 985 | 674 | 869 | 939 | 570 | 762 | 44636 | 28305 | 46596 | 34881 | 22252 | 15225 | 22583 |
| 593 | 957 | 762 | 566 | 1023 | 781 | 593 | 997 | 31245 | 23669 | 39925 | 38211 | 15392 | 11438 | 20192 |
| 698 | 964 | 991 | 893 | 1164 | 867 | 668 | 1015 | 17205 | 13394 | 19186 | 18493 | 17984 | 16207 | 18156 |
| 899 | 1233 | 1072 | 864 | 1147 | 1024 | 835 | 1184 | 30767 | 26802 | 29817 | 36778 | 37515 | 30225 | 35001 |
| 643 | 931 | 955 | 767 | 1070 | 951 | 768 | 1085 | 18794 | 12367 | 19245 | 17908 | 15475 | 12420 | 16200 |
| 378 | 818 | 583 | 442 | 898 | 473 | 367 | 906 | 25872 | 20707 | 54719 | 44745 | 17980 | 13649 | 30848 |
| 656 | 1134 | 913 | 764 | 1136 | 815 | 703 | 1171 | 26287 | 22668 | 34541 | 39173 | 27292 | 22845 | 29895 |
| 491 | 809 | 545 | 563 | 839 | 543 | 573 | 813 | 24758 | 22820 | 46510 | 37608 | 16836 | 17377 | 30867 |

| | | | | | | | | | | | | | | |
|---|---|---|---|---|---|---|---|---|---|---|---|---|---|---|
| 494 | 878 | 748 | 584 | 914 | 583 | 399 | 796 | 25817 | 20221 | 40894 | 35886 | 29578 | 23087 | 39566 |
| 647 | 1248 | 833 | 722 | 1165 | 812 | 712 | 1168 | 13101 | 11661 | 18011 | 22477 | 34117 | 29563 | 40943 |
| 730 | 1170 | 836 | 752 | 1097 | 845 | 790 | 1214 | 20011 | 17209 | 23587 | 27604 | 17159 | 15433 | 20536 |
| 649 | 1053 | 829 | 641 | 1047 | 814 | 608 | 1038 | 33672 | 25583 | 39419 | 41516 | 36926 | 28583 | 44564 |
| 653 | 1025 | 922 | 627 | 993 | 895 | 639 | 988 | 20513 | 14518 | 22242 | 22796 | 22751 | 15478 | 24666 |
| 481 | 1049 | 686 | 530 | 1008 | 647 | 466 | 980 | 31270 | 24070 | 50027 | 52491 | 17782 | 13727 | 25907 |
| 472 | 755 | 495 | 450 | 742 | 500 | 438 | 703 | 25931 | 23587 | 49956 | 37728 | 26867 | 24407 | 54227 |
| 654 | 1125 | 851 | 695 | 1200 | 743 | 577 | 1053 | 21009 | 17079 | 26113 | 29370 | 23040 | 18809 | 27069 |
| 446 | 630 | 707 | 428 | 606 | 590 | 305 | 505 | 29802 | 19219 | 43047 | 27110 | 22869 | 13843 | 32369 |
| 472 | 924 | 730 | 565 | 1144 | 591 | 440 | 1009 | 12209 | 8917 | 18894 | 17452 | 17053 | 13186 | 23357 |
| 480 | 738 | 647 | 513 | 748 | 601 | 445 | 742 | 20477 | 15673 | 32652 | 24103 | 20878 | 16553 | 32248 |
| 636 | 1075 | 765 | 679 | 1085 | 736 | 634 | 1039 | 19061 | 17093 | 26894 | 28900 | 20534 | 18210 | 26829 |
| 507 | 857 | 744 | 572 | 907 | 663 | 530 | 869 | 23917 | 18656 | 36773 | 31513 | 20200 | 15525 | 27139 |
| 704 | 1230 | 800 | 722 | 1198 | 844 | 751 | 1329 | 7060 | 5918 | 8409 | 10345 | 19510 | 17605 | 24380 |
| 354 | 774 | 627 | 406 | 825 | 592 | 406 | 820 | 15704 | 9668 | 27338 | 21155 | 25032 | 16201 | 39905 |
| 599 | 960 | 795 | 533 | 881 | 736 | 497 | 915 | 28544 | 22105 | 36906 | 35420 | 29802 | 19999 | 37487 |
| 828 | 1142 | 1146 | 878 | 1084 | 1170 | 865 | 1094 | 48807 | 35728 | 43146 | 49271 | 15072 | 11542 | 13148 |
| 419 | 781 | 571 | 416 | 797 | 567 | 407 | 789 | 26158 | 16551 | 39535 | 30893 | 17858 | 13004 | 31256 |
| 713 | 851 | 916 | 504 | 688 | 1083 | 784 | 908 | 2626 | 1786 | 2505 | 2133 | 6066 | 3336 | 6621 |
| 612 | 848 | 653 | 477 | 828 | 709 | 488 | 789 | 9924 | 7433 | 12149 | 10300 | 16577 | 12099 | 25381 |
| 708 | 1136 | 965 | 759 | 1090 | 939 | 756 | 1007 | 27280 | 23277 | 32867 | 37335 | 55294 | 43488 | 57322 |
| 561 | 791 | 771 | 569 | 792 | 796 | 632 | 993 | 7676 | 5472 | 9760 | 7722 | 9785 | 7224 | 12699 |
| 792 | 1223 | 808 | 706 | 1090 | 829 | 818 | 1271 | 27203 | 25359 | 32029 | 39166 | 26471 | 23134 | 32767 |
| 549 | 876 | 891 | 453 | 793 | 920 | 561 | 884 | 24077 | 14501 | 26422 | 23142 | 29979 | 15243 | 33632 |
| 791 | 1108 | 1009 | 834 | 1056 | 1017 | 829 | 1082 | 24122 | 19193 | 24253 | 26871 | 18757 | 15507 | 18586 |
| 844 | 1306 | 897 | 781 | 1279 | 1022 | 991 | 1425 | 12769 | 11673 | 13822 | 18054 | 7693 | 6695 | 8573 |
| 483 | 702 | 1227 | 470 | 705 | 1242 | 486 | 660 | 16397 | 6401 | 13265 | 9311 | 41495 | 15907 | 33816 |
| 349 | 704 | 619 | 367 | 715 | 692 | 459 | 754 | 26891 | 15818 | 45366 | 31936 | 23718 | 14086 | 38330 |
| 1236 | 1469 | 978 | 902 | 1341 | 1116 | 1022 | 1446 | 15205 | 14548 | 11766 | 17284 | 24439 | 22541 | 24986 |
| 471 | 866 | 677 | 516 | 830 | 648 | 537 | 877 | 27782 | 21164 | 44920 | 38878 | 22691 | 17304 | 33520 |
| 501 | 1004 | 812 | 565 | 974 | 669 | 486 | 898 | 42690 | 28224 | 56331 | 56573 | 24421 | 16985 | 30082 |

| | | | | | | | | | | | | | | |
|---|---|---|---|---|---|---|---|---|---|---|---|---|---|---|
| 585 | 789 | 873 | 560 | 789 | 936 | 635 | 794 | 24228 | 15905 | 27187 | 21441 | 38757 | 24848 | 44370 |
| 563 | 843 | | | | | | | 2012 | 1170 | 2079 | 1753 | | | |
| 755 | 1050 | 924 | 677 | 970 | 932 | 792 | 1132 | 29853 | 22453 | 29750 | 31234 | 26639 | 19540 | 28843 |
| 635 | 1002 | 964 | 680 | 1028 | 783 | 568 | 945 | 18721 | 12963 | 20406 | 20455 | 29964 | 21156 | 31098 |
| 472 | 708 | 947 | 450 | 669 | 865 | 388 | 600 | 21626 | 10622 | 22514 | 15933 | 19499 | 9274 | 20591 |
| 892 | 1307 | 986 | 952 | 1391 | 950 | 943 | 1385 | 27353 | 26121 | 29293 | 38299 | 15544 | 15012 | 15769 |
| 530 | 1007 | 528 | 459 | 928 | 571 | 495 | 964 | 16470 | 14187 | 26783 | 26960 | 14311 | 12450 | 27105 |
| | | 843 | 622 | 899 | 777 | 541 | 799 | | | | | 1768 | 1305 | 2097 |
| 725 | 1008 | 771 | 804 | 1064 | 717 | 738 | 1052 | 33589 | 32781 | 45216 | 45598 | 24925 | 26006 | 32326 |
| 312 | 733 | 632 | 322 | 726 | 666 | 328 | 725 | 27868 | 13142 | 42116 | 30868 | 29870 | 15216 | 47280 |
| 817 | 1213 | 968 | 914 | 1264 | 841 | 732 | 1122 | 28895 | 25753 | 31537 | 38244 | 32424 | 30604 | 33482 |
| 1023 | 1407 | 1171 | 1163 | 1546 | 1034 | 1054 | 1494 | 21903 | 20769 | 20301 | 28570 | 14441 | 14343 | 12330 |
| 1015 | 1389 | 1079 | 1042 | 1438 | 1176 | 1189 | 1496 | 28074 | 28653 | 28238 | 39211 | 16091 | 15547 | 14915 |
| 676 | 1116 | 896 | 810 | 1141 | 820 | 723 | 1179 | 14744 | 13419 | 19841 | 22136 | 21681 | 19585 | 24192 |
| 455 | 715 | 751 | 608 | 856 | 648 | 470 | 751 | 22924 | 16744 | 36794 | 26295 | 18630 | 15081 | 24796 |
| 674 | 904 | 902 | 792 | 1032 | 914 | 765 | 966 | 12382 | 10369 | 15389 | 13911 | 14719 | 12924 | 16320 |
| 377 | 673 | 556 | 290 | 613 | 573 | 308 | 637 | 12277 | 6790 | 18014 | 12132 | 30508 | 15885 | 54843 |
| 901 | 1240 | 1052 | 930 | 1217 | 949 | 842 | 1198 | 11862 | 10024 | 11128 | 13801 | 21834 | 19298 | 20759 |
| 579 | 974 | 752 | 538 | 930 | 770 | 552 | 949 | 22640 | 16523 | 28521 | 27772 | 35823 | 25641 | 47659 |
| 1154 | 1696 | 1046 | 1200 | 1683 | 1017 | 1400 | 1981 | 31993 | 33657 | 29170 | 49466 | 16869 | 19351 | 16133 |
| 637 | 1226 | 773 | 627 | 1186 | 721 | 599 | 1237 | 9317 | 7769 | 12192 | 14952 | 11649 | 9444 | 15072 |
| 906 | 1529 | 1000 | 1005 | 1540 | 912 | 924 | 1484 | 21138 | 20647 | 22790 | 34844 | 18262 | 18347 | 18261 |
| 412 | 750 | 526 | 357 | 716 | 633 | 422 | 755 | 31534 | 23140 | 56097 | 42088 | 32109 | 21821 | 61093 |
| 1029 | 1480 | 983 | 1027 | 1450 | 928 | 1075 | 1549 | 25301 | 27432 | 26654 | 39452 | 26760 | 27960 | 27221 |
| 676 | 1078 | 791 | 624 | 1070 | 749 | 623 | 1082 | 15392 | 13120 | 19397 | 20911 | 14162 | 11175 | 17914 |
| 807 | 1174 | 917 | 751 | 1077 | 999 | 810 | 1086 | 36191 | 30182 | 37379 | 43882 | 31898 | 26124 | 34802 |
| 888 | 6273 | 1265 | 1096 | 6438 | 904 | 672 | 4729 | 12988 | 8719 | 9814 | 61570 | 8349 | 7230 | 6599 |
| 995 | 1439 | 831 | 712 | 1087 | 907 | 735 | 1039 | 16364 | 16418 | 16497 | 23746 | 24668 | 21159 | 29700 |
| 632 | 766 | 837 | 551 | 678 | 978 | 599 | 705 | 30198 | 17300 | 27363 | 20959 | 13603 | 8949 | 16252 |
| 903 | 1288 | 1043 | 996 | 1391 | 1103 | 1054 | 1470 | 23359 | 21458 | 23765 | 30610 | 26738 | 25552 | 25644 |
| 566 | 880 | 717 | 576 | 899 | 689 | 548 | 857 | 19168 | 15006 | 26507 | 23331 | 15022 | 12062 | 20957 |

| | | | | | | | | | | | | | | |
|---|---|---|---|---|---|---|---|---|---|---|---|---|---|---|
| 986 | 1287 | 1107 | 1062 | 1396 | 972 | 846 | 1325 | 22279 | 21810 | 22116 | 28459 | 17507 | 16806 | 15820 |
| 1041 | 1185 | 1012 | 946 | 1211 | 1139 | 1044 | 1283 | 11215 | 10120 | 9719 | 11517 | 16824 | 15719 | 16618 |
| 745 | 1049 | 792 | 623 | 918 | 795 | 692 | 1066 | 22978 | 18158 | 24382 | 25588 | 15169 | 11928 | 19157 |
| 662 | 1051 | 779 | 710 | 1154 | 741 | 632 | 1065 | 25603 | 21639 | 32685 | 34352 | 36285 | 33102 | 46600 |
| 502 | 862 | 925 | 480 | 871 | 899 | 514 | 901 | 24785 | 13643 | 27153 | 23399 | 30585 | 15865 | 33078 |
| 819 | 1076 | 926 | 777 | 1091 | 976 | 926 | 1108 | 25504 | 22451 | 27423 | 29511 | 24228 | 20329 | 26158 |
| 482 | 954 | 646 | 478 | 916 | 700 | 601 | 955 | 20053 | 17012 | 35277 | 33642 | 22790 | 16843 | 35259 |
| 549 | 758 | 743 | 553 | 765 | 782 | 602 | 794 | 29167 | 20756 | 37826 | 28659 | 34502 | 25682 | 46452 |
| 634 | 897 | 784 | 605 | 832 | 814 | 645 | 877 | 9289 | 7112 | 11218 | 10062 | 14634 | 11302 | 18665 |
| 586 | 943 | 789 | 653 | 904 | 908 | 793 | 916 | 12880 | 9608 | 16388 | 15457 | 22371 | 18504 | 28355 |
| 760 | 1017 | 993 | 708 | 967 | 1007 | 680 | 1040 | 35356 | 26094 | 34312 | 34899 | 27469 | 19577 | 27668 |
| 652 | 1003 | 868 | 618 | 934 | 872 | 628 | 1040 | 22557 | 16624 | 25497 | 25579 | 13326 | 9496 | 15361 |
| 703 | 1140 | 901 | 755 | 1149 | 1130 | 952 | 1365 | 13908 | 11232 | 15973 | 18206 | 9912 | 8313 | 11004 |
| 577 | 971 | 789 | 720 | 1063 | 601 | 454 | 817 | 33657 | 26943 | 46664 | 45324 | 44511 | 40602 | 56404 |
| 699 | 1066 | 942 | 701 | 1064 | 1095 | 832 | 1176 | 36148 | 27058 | 38708 | 41268 | 33679 | 25069 | 35767 |
| 811 | 1133 | 1047 | 852 | 1208 | 1040 | 865 | 1107 | 21479 | 17475 | 21541 | 24408 | 28179 | 22928 | 26907 |
| 674 | 1239 | 944 | 845 | 1409 | 754 | 659 | 1230 | 38440 | 31447 | 46662 | 57805 | 24333 | 21768 | 25767 |
| 772 | 1109 | 842 | 669 | 1124 | 764 | 674 | 1122 | 25390 | 23507 | 30469 | 33796 | 16191 | 12871 | 19236 |
| | | 278 | 123 | 595 | 431 | 221 | 574 | | | | | 4137 | 1837 | 14899 |
| 617 | 936 | 1093 | 749 | 1001 | 966 | 578 | 797 | 13761 | 9843 | 15941 | 14926 | 41443 | 28386 | 37912 |
| 524 | 1007 | 863 | 662 | 1004 | 810 | 644 | 1018 | 26613 | 19228 | 36664 | 36912 | 29512 | 22646 | 34199 |
| 321 | 597 | 488 | 266 | 570 | 460 | 237 | 547 | 27194 | 15055 | 46867 | 27959 | 26572 | 14472 | 54402 |
| 569 | 824 | 822 | 571 | 840 | 719 | 528 | 823 | 19133 | 13635 | 23949 | 19737 | 18766 | 13033 | 22817 |
| 324 | 707 | 647 | 338 | 744 | 730 | 434 | 798 | 15970 | 8526 | 26295 | 18588 | 13332 | 6955 | 20593 |
| 765 | 1121 | 838 | 752 | 1075 | 956 | 786 | 1128 | 34864 | 29794 | 38961 | 43687 | 19198 | 17225 | 22903 |
| | | 1493 | 1103 | 1007 | 1277 | 1023 | 928 | | | | | 3671 | 2712 | 2458 |
| 422 | 847 | 742 | 446 | 800 | 826 | 493 | 845 | 20710 | 12307 | 29144 | 24673 | 25211 | 15144 | 33956 |
| 469 | 841 | 775 | 513 | 850 | 760 | 490 | 823 | 30740 | 18654 | 39749 | 33419 | 31573 | 20919 | 40755 |
| 1004 | 1363 | 928 | 866 | 1431 | 1006 | 913 | 1317 | 24988 | 23876 | 23789 | 32414 | 30226 | 28206 | 32579 |
| 955 | 1300 | 986 | 968 | 1398 | 1056 | 910 | 1302 | 31072 | 29068 | 30425 | 39546 | 24607 | 24150 | 24951 |
| 312 | 689 | 612 | 384 | 658 | 550 | 325 | 665 | 9650 | 5317 | 17035 | 11736 | 6399 | 4013 | 10454 |

| | | | | | | | | | | | | | | |
|---|---|---|---|---|---|---|---|---|---|---|---|---|---|---|
| 698 | 876 | 979 | 896 | 980 | 827 | 734 | 904 | 24743 | 19130 | 27392 | 23989 | 43049 | 39415 | 43970 |
| 419 | 637 | 455 | 421 | 662 | 319 | 330 | 559 | 25041 | 20567 | 49128 | 31277 | 24536 | 22718 | 53915 |
| 1016 | 1473 | 1119 | 1059 | 1387 | 1000 | 877 | 1311 | 9184 | 8711 | 8576 | 12629 | 9205 | 8704 | 8222 |
| 406 | 716 | 635 | 432 | 724 | 563 | 454 | 751 | 20166 | 16073 | 39551 | 28327 | 17967 | 12242 | 28307 |
| 697 | 1201 | 871 | 653 | 1128 | 830 | 642 | 1209 | 13315 | 10659 | 15285 | 18355 | 13967 | 10464 | 16031 |
| 478 | 766 | 645 | 562 | 778 | 688 | 460 | 712 | 14926 | 13158 | 27522 | 21073 | 14804 | 12899 | 22948 |
| 404 | 781 | 492 | 315 | 745 | 525 | 369 | 790 | 22264 | 15727 | 38942 | 30429 | 28026 | 17935 | 57017 |
| 1329 | 1103 | 1397 | 1230 | 1023 | 1366 | 1112 | 1009 | 23254 | 19467 | 14645 | 16155 | 35170 | 30958 | 25176 |
| 906 | 1176 | 864 | 956 | 1182 | 801 | 842 | 1112 | 26847 | 28167 | 31087 | 36572 | 23433 | 25901 | 27107 |
| 511 | 828 | 598 | 429 | 791 | 556 | 364 | 729 | 19334 | 14429 | 28223 | 23357 | 15366 | 11033 | 25696 |
| 391 | 666 | 437 | 365 | 655 | 422 | 359 | 649 | 24933 | 20635 | 52745 | 35128 | 15991 | 13349 | 36579 |
| 696 | 1179 | 803 | 650 | 1191 | 973 | 821 | 1361 | 17607 | 14285 | 20513 | 24194 | 10284 | 8328 | 12812 |
| 707 | 1265 | 808 | 703 | 1183 | 792 | 698 | 1369 | 4904 | 4384 | 6204 | 7846 | 1394 | 1214 | 1726 |
| 597 | 2309 | 706 | 746 | 2489 | 699 | 650 | 2397 | 11802 | 10494 | 17567 | 40563 | 14734 | 15561 | 20869 |
| 376 | 725 | 759 | 504 | 862 | 655 | 379 | 712 | 29110 | 15993 | 42587 | 30865 | 16910 | 11231 | 22293 |
| 635 | 967 | 745 | 643 | 952 | 735 | 639 | 983 | 18476 | 15400 | 24249 | 23453 | 23869 | 20594 | 32049 |
| 1551 | 1284 | 1159 | 1230 | 1303 | 1100 | 1221 | 1350 | 29800 | 36928 | 23807 | 30570 | 20429 | 21678 | 17628 |
| 1240 | 1920 | 1126 | 1123 | 1578 | 1199 | 1168 | 1641 | 11156 | 11048 | 8910 | 17106 | 18596 | 18537 | 16511 |
| 384 | 813 | 744 | 349 | 767 | 778 | 291 | 516 | 5236 | 3040 | 7922 | 6438 | 8096 | 3792 | 10877 |
| 829 | 1142 | 918 | 766 | 1093 | 1022 | 916 | 1234 | 9964 | 8650 | 10438 | 11925 | 21129 | 17637 | 23015 |
| 599 | 980 | 758 | 593 | 953 | 772 | 640 | 1024 | 21944 | 18684 | 31218 | 30591 | 25903 | 20240 | 34154 |
| 775 | 1307 | 990 | 853 | 1273 | 965 | 793 | 1296 | 16577 | 13693 | 17676 | 23106 | 17137 | 14765 | 17317 |
| 418 | 702 | 589 | 456 | 685 | 551 | 440 | 713 | 20031 | 16935 | 40536 | 28450 | 13894 | 10755 | 23574 |
| 426 | 907 | 707 | 432 | 937 | 694 | 457 | 887 | 17040 | 10259 | 24081 | 21850 | 24030 | 14697 | 33984 |
| 245 | 690 | 505 | 248 | 670 | 524 | 250 | 681 | 21389 | 10646 | 43399 | 29935 | 19099 | 9386 | 37790 |
| 743 | 980 | 1122 | 969 | 1040 | 902 | 715 | 975 | 41333 | 33689 | 45360 | 44461 | 15411 | 13317 | 13740 |
| 613 | 1031 | 822 | 660 | 1067 | 785 | 585 | 986 | 19770 | 15875 | 25906 | 26710 | 23203 | 18632 | 28214 |
| 844 | 922 | 1159 | 989 | 1069 | 1064 | 885 | 1037 | 9509 | 7908 | 9374 | 8639 | 10008 | 8545 | 8637 |
| 1081 | 1401 | 1181 | 1219 | 1594 | 1144 | 1003 | 1284 | 16466 | 15933 | 14738 | 20645 | 10308 | 10634 | 8727 |
| 812 | 1417 | 870 | 795 | 1330 | 902 | 832 | 1351 | 22992 | 20187 | 24866 | 35233 | 28988 | 26459 | 33302 |
| 653 | 830 | 869 | 620 | 813 | 901 | 632 | 821 | 24412 | 16934 | 25939 | 21524 | 17671 | 12594 | 20327 |

| | | | | | | | | | | | | | | |
|---|---|---|---|---|---|---|---|---|---|---|---|---|---|---|
| 470 | 774 | 702 | 434 | 750 | 725 | 488 | 792 | 28347 | 19197 | 40849 | 31610 | 39472 | 24404 | 56247 |
| 698 | 4699 | 1227 | 724 | 6256 | 1043 | 754 | 5434 | 2977 | 2174 | 3115 | 14638 | 8373 | 4944 | 6825 |
| 391 | 742 | 546 | 321 | 669 | 578 | 358 | 677 | 24577 | 15366 | 39328 | 29200 | 12777 | 7508 | 23386 |
| 436 | 771 | 717 | 522 | 803 | 735 | 557 | 839 | 22441 | 16242 | 37230 | 28700 | 27799 | 20227 | 38781 |
| 466 | 717 | | | | | | | 2653 | 1062 | 2278 | 1633 | | | |
| 745 | 4206 | 1028 | 674 | 4661 | 957 | 669 | 5822 | 5314 | 3773 | 5062 | 21291 | 7519 | 4927 | 7311 |
| 821 | 1046 | 777 | 621 | 947 | 848 | 685 | 978 | 12090 | 12484 | 15202 | 15898 | 13069 | 10441 | 16824 |
| 396 | 732 | 563 | 415 | 751 | 474 | 369 | 698 | 22250 | 17157 | 43337 | 31731 | 32670 | 24055 | 57977 |
| 685 | 1134 | 845 | 648 | 1051 | 875 | 697 | 1095 | 24233 | 19921 | 29088 | 32998 | 18569 | 14231 | 21970 |
| 721 | 1064 | 753 | 651 | 1012 | 857 | 685 | 1082 | 33602 | 26965 | 37379 | 39762 | 18529 | 16000 | 24594 |
| 707 | 1003 | 883 | 686 | 1003 | 839 | 665 | 1000 | 28058 | 21339 | 30172 | 30277 | 29729 | 23106 | 33659 |
| 541 | 929 | 818 | 610 | 1139 | 765 | 550 | 962 | 10673 | 7649 | 14128 | 13127 | 17783 | 13259 | 21747 |
| 646 | 884 | 1163 | 871 | 953 | 978 | 710 | 812 | 7412 | 5318 | 8231 | 7276 | 11803 | 8839 | 10152 |
| 760 | 1084 | 853 | 699 | 1036 | 824 | 682 | 1048 | 13157 | 11280 | 14846 | 16095 | 22496 | 18440 | 26376 |
| 730 | 1066 | 935 | 773 | 1034 | 1002 | 811 | 1054 | 23272 | 18431 | 25255 | 26933 | 33976 | 28099 | 36340 |
| 646 | 1065 | 776 | 696 | 1131 | 711 | 638 | 1106 | 18755 | 13519 | 20911 | 22266 | 19107 | 17144 | 24635 |
| 769 | 1153 | 744 | 653 | 1096 | 730 | 714 | 1081 | 15266 | 18098 | 23544 | 27136 | 24548 | 21559 | 32992 |
| 908 | 1245 | 919 | 825 | 1258 | 1041 | 927 | 1254 | 13473 | 13179 | 14508 | 18069 | 11274 | 10118 | 12262 |
| 939 | 1023 | 759 | 699 | 853 | 950 | 894 | 1097 | 28596 | 25905 | 27575 | 28209 | 6287 | 5787 | 8281 |
| 446 | 749 | 707 | 415 | 727 | 694 | 476 | 751 | 24584 | 14807 | 33191 | 24847 | 21362 | 12556 | 30235 |
| 518 | 785 | 530 | 426 | 800 | 550 | 400 | 705 | 33818 | 26748 | 51684 | 40547 | 17513 | 14070 | 33030 |
| 459 | 764 | 850 | 648 | 839 | 716 | 477 | 775 | 23975 | 14984 | 32623 | 24912 | 22372 | 17040 | 26310 |
| 553 | 975 | 965 | 817 | 1096 | 804 | 473 | 732 | 11066 | 8248 | 14918 | 14542 | 6192 | 5242 | 6414 |
| 1188 | 1222 | 1327 | 1135 | 1199 | 1290 | 1197 | 1196 | 30507 | 26654 | 22443 | 27421 | 29773 | 25454 | 22431 |
| 844 | 1313 | 857 | 892 | 1390 | 805 | 872 | 1418 | 30461 | 28530 | 33791 | 44379 | 18652 | 19413 | 21765 |
| 555 | 882 | 703 | 586 | 912 | 701 | 602 | 895 | 23323 | 19091 | 34377 | 30332 | 26936 | 22454 | 38300 |
| 491 | 874 | 807 | 539 | 895 | 746 | 441 | 773 | 17749 | 11094 | 22577 | 19738 | 11569 | 7736 | 14340 |
| 555 | 1126 | 601 | 590 | 1184 | 626 | 607 | 1129 | 24418 | 21600 | 38917 | 43816 | 23692 | 23258 | 39396 |
| 520 | 776 | 684 | 378 | 647 | 829 | 530 | 816 | 18745 | 12660 | 24343 | 18900 | 10218 | 5644 | 14932 |
| 733 | 1098 | 828 | 734 | 1180 | 871 | 713 | 1044 | 25968 | 21449 | 29268 | 32146 | 13376 | 11856 | 16160 |
| 747 | 1102 | 821 | 765 | 1163 | 978 | 977 | 1384 | 20782 | 18195 | 24368 | 26862 | 14094 | 13123 | 17158 |

| | | | | | | | | | | | | | | |
|---|---|---|---|---|---|---|---|---|---|---|---|---|---|---|
| 530 | 886 | 788 | 516 | 904 | 760 | 454 | 816 | 15637 | 10274 | 19396 | 17177 | 13592 | 8891 | 17245 |
| 837 | 1091 | 954 | 784 | 1127 | 1009 | 878 | 1072 | 13226 | 11086 | 13249 | 14451 | 13634 | 11206 | 14287 |
| 889 | 1254 | 913 | 806 | 1252 | 894 | 772 | 1125 | 20790 | 18083 | 20342 | 25513 | 21284 | 18791 | 23317 |
| 544 | 922 | 775 | 519 | 927 | 846 | 561 | 907 | 25975 | 17501 | 32178 | 29674 | 12876 | 8628 | 16621 |
| 649 | 1169 | 806 | 637 | 1290 | 900 | 659 | 1186 | 17182 | 12659 | 19505 | 22796 | 15255 | 12061 | 18938 |
| 586 | 898 | 745 | 588 | 1019 | 807 | 709 | 1075 | 18263 | 14888 | 25411 | 22816 | 15016 | 11855 | 20168 |
| 929 | 1171 | 1029 | 904 | 1175 | 1078 | 867 | 1113 | 14775 | 12627 | 13589 | 15911 | 20116 | 17656 | 19541 |
| 544 | 858 | 775 | 527 | 841 | 753 | 517 | 847 | 19619 | 13324 | 24490 | 21017 | 15602 | 10617 | 20131 |
| 737 | 1127 | 886 | 777 | 1337 | 869 | 813 | 1298 | 14893 | 13720 | 18617 | 20984 | 24983 | 21928 | 28205 |
| 599 | 929 | 702 | 512 | 937 | 780 | 542 | 976 | 21134 | 17069 | 28499 | 26485 | 19817 | 14457 | 28216 |
| 699 | 1017 | 969 | 722 | 1137 | 984 | 698 | 1108 | 28050 | 20992 | 30011 | 30510 | 22771 | 16970 | 23512 |
| 599 | 1035 | 840 | 547 | 903 | 853 | 636 | 1027 | 25144 | 17763 | 29646 | 30684 | 23563 | 15350 | 28064 |
| 614 | 919 | 767 | 701 | 1075 | 527 | 519 | 808 | 14819 | 14314 | 23311 | 21425 | 14211 | 12992 | 18524 |
| 582 | 857 | 794 | 664 | 939 | 762 | 541 | 859 | 31795 | 24140 | 41453 | 35511 | 42111 | 35247 | 53055 |
| 784 | 1049 | 985 | 714 | 919 | 1094 | 747 | 956 | 18700 | 13727 | 17504 | 18359 | 27038 | 19603 | 27460 |
| 688 | 1190 | 699 | 604 | 1123 | 622 | 555 | 1127 | 20869 | 20099 | 29217 | 34771 | 27003 | 23341 | 38613 |
| 692 | 918 | 989 | 585 | 815 | 1010 | 604 | 928 | 22987 | 14676 | 21201 | 19460 | 14416 | 8525 | 14578 |
| 570 | 860 | 845 | 555 | 802 | 821 | 574 | 875 | 32917 | 23951 | 42049 | 36141 | 36498 | 23949 | 43186 |
| 569 | 1019 | 692 | 637 | 1275 | 664 | 580 | 1228 | 2645 | 2134 | 3749 | 3821 | 1578 | 1452 | 2279 |
| 916 | 1118 | 1036 | 881 | 1292 | 932 | 787 | 1170 | 14774 | 12695 | 13856 | 15495 | 20535 | 17460 | 19815 |
| 418 | 842 | 567 | 378 | 841 | 483 | 331 | 810 | 28904 | 20196 | 48320 | 40682 | 24685 | 16449 | 43537 |
| 675 | 968 | 910 | 659 | 980 | 805 | 554 | 870 | 26540 | 19570 | 29009 | 28082 | 26286 | 19049 | 28896 |
| 330 | 763 | 425 | 285 | 723 | 473 | 300 | 761 | 18770 | 12606 | 38214 | 29176 | 7829 | 5262 | 18438 |
| 748 | 961 | 938 | 703 | 844 | 956 | 679 | 832 | 10826 | 8454 | 11306 | 10870 | 14565 | 10913 | 15532 |
| 983 | 1314 | 993 | 855 | 1271 | 942 | 815 | 1335 | 30855 | 27758 | 28238 | 37109 | 22839 | 19663 | 23005 |
| 630 | 1030 | 824 | 671 | 1093 | 859 | 649 | 1084 | 17060 | 13135 | 20836 | 21461 | 15310 | 12472 | 18590 |
| | | 363 | 273 | 558 | | | | | | | | 2483 | 1867 | 6837 |
| 388 | 853 | 552 | 336 | 749 | 633 | 379 | 881 | 15157 | 9573 | 24688 | 21071 | 8051 | 4902 | 14596 |
| 517 | 886 | 676 | 504 | 877 | 608 | 434 | 770 | 24143 | 17061 | 33026 | 29256 | 7353 | 5483 | 10875 |
| 374 | 698 | 613 | 306 | 663 | 612 | 336 | 654 | 16536 | 9919 | 26492 | 18493 | 16365 | 8173 | 26682 |
| 386 | 702 | 656 | 402 | 744 | 701 | 359 | 678 | 10585 | 6354 | 16468 | 11561 | 20184 | 12367 | 30753 |

| | | | | | | | | | | | | | | |
|---|---|---|---|---|---|---|---|---|---|---|---|---|---|---|
| 1490 | 1245 | 872 | 694 | 1138 | 1371 | 1133 | 1090 | 292 | 281 | 189 | 235 | 1489 | 1186 | 1708 |
| 824 | 1181 | 930 | 863 | 1164 | 955 | 824 | 1062 | 16221 | 14474 | 17564 | 20742 | 30963 | 28759 | 33310 |
| 434 | 663 | 878 | 396 | 574 | 1089 | 452 | 629 | 28306 | 13561 | 31224 | 20690 | 14516 | 6549 | 16541 |
| 881 | 1223 | 811 | 704 | 1121 | 926 | 772 | 1090 | 16830 | 16150 | 18323 | 22412 | 13573 | 11778 | 16736 |
| 496 | 723 | 440 | 352 | 596 | 393 | 316 | 569 | 21234 | 18565 | 37438 | 27061 | 10261 | 8206 | 23295 |
| 697 | 1144 | 923 | 720 | 1195 | 814 | 655 | 1190 | 23391 | 19193 | 27544 | 31501 | 14991 | 11693 | 16244 |
| 731 | 1172 | 775 | 743 | 1222 | 783 | 682 | 1138 | 25292 | 23332 | 31906 | 37403 | 16890 | 16181 | 21785 |
| 795 | 1011 | 1002 | 778 | 1015 | 989 | 744 | 1023 | 12591 | 9087 | 11431 | 11555 | 18212 | 14141 | 18172 |
| 352 | 787 | 607 | 383 | 774 | 701 | 463 | 851 | 17663 | 11067 | 31405 | 24705 | 28423 | 17923 | 46790 |
| 454 | 792 | 807 | 489 | 831 | 816 | 568 | 847 | 20234 | 13393 | 29499 | 23359 | 39037 | 23654 | 48372 |
| 494 | 898 | 837 | 686 | 1004 | 782 | 620 | 985 | 7836 | 5741 | 11624 | 10439 | 7040 | 5770 | 8413 |
| 466 | 913 | 555 | 405 | 860 | 603 | 414 | 873 | 19662 | 15207 | 32616 | 29785 | 12432 | 9078 | 22391 |
| 204 | 537 | 556 | 325 | 670 | 387 | 231 | 595 | 24381 | 10609 | 52027 | 27932 | 18858 | 11014 | 33910 |
| 272 | 583 | 414 | 253 | 589 | 502 | 295 | 604 | 33043 | 19077 | 70085 | 40845 | 11114 | 6786 | 26868 |
| 707 | 1060 | 878 | 627 | 1113 | 917 | 750 | 1126 | 35937 | 26383 | 37319 | 39572 | 13213 | 9438 | 15056 |
| 669 | 1028 | 881 | 683 | 1027 | 975 | 652 | 942 | 21195 | 15331 | 22904 | 23555 | 24414 | 18933 | 27710 |
| 694 | 987 | 949 | 658 | 943 | 932 | 681 | 938 | 29630 | 19936 | 28722 | 28344 | 20669 | 14339 | 21790 |
| 423 | 743 | 539 | 384 | 706 | 548 | 402 | 721 | 11908 | 8611 | 20349 | 15121 | 13608 | 9688 | 25241 |
| 470 | 845 | 734 | 440 | 792 | 779 | 510 | 860 | 19220 | 11571 | 24636 | 20808 | 14535 | 8725 | 19812 |
| 443 | 851 | 786 | 581 | 1028 | 866 | 546 | 947 | 14548 | 7745 | 17485 | 14886 | 13545 | 10018 | 17238 |
| 767 | 1005 | 1237 | 896 | 1000 | 1164 | 799 | 994 | 25607 | 15804 | 20607 | 20709 | 21954 | 15894 | 17744 |
| 310 | 507 | 646 | 560 | 893 | 758 | 596 | 867 | 37004 | 22072 | 71264 | 36147 | 9409 | 8148 | 14553 |
| 988 | 1256 | 1347 | 1068 | 1134 | 1325 | 1106 | 1207 | 14602 | 12395 | 12542 | 15752 | 26060 | 20662 | 19345 |
| 621 | 1009 | 808 | 571 | 981 | 768 | 653 | 1071 | 14752 | 10494 | 16905 | 17059 | 18964 | 13393 | 23468 |
| 703 | 944 | 652 | 566 | 933 | 856 | 663 | 927 | 22349 | 18926 | 26917 | 25419 | 7345 | 6378 | 11267 |
| 352 | 509 | 1247 | 254 | 450 | 1221 | 318 | 540 | 26188 | 7540 | 21426 | 10903 | 26980 | 5494 | 21640 |
| 373 | 658 | 531 | 384 | 672 | 526 | 370 | 678 | 19354 | 13756 | 36908 | 24295 | 19523 | 14111 | 36785 |
| 927 | 1391 | 983 | 950 | 1277 | 897 | 808 | 1161 | 7906 | 7356 | 7936 | 11042 | 18646 | 18019 | 18976 |
| 602 | 882 | 779 | 579 | 900 | 684 | 540 | 871 | 13174 | 10225 | 16994 | 14996 | 15991 | 11879 | 20520 |
| 842 | 1216 | 971 | 962 | 1337 | 900 | 1046 | 1390 | 12505 | 12747 | 15134 | 18401 | 16472 | 16318 | 16970 |
| 774 | 1303 | 749 | 754 | 1284 | 770 | 775 | 1327 | 16042 | 16258 | 21017 | 27382 | 21052 | 21189 | 28091 |

| | | | | | | | | | | | | | | |
|---|---|---|---|---|---|---|---|---|---|---|---|---|---|---|
| 266 | 570 | 487 | 310 | 579 | 496 | 276 | 569 | 18544 | 10411 | 39203 | 22345 | 14831 | 9439 | 30445 |
| 571 | 913 | 850 | 623 | 813 | 807 | 579 | 913 | 13997 | 9997 | 17522 | 15998 | 14238 | 10447 | 16756 |
| 535 | 4124 | 779 | 516 | 3003 | 792 | 537 | 2533 | 6935 | 4405 | 8241 | 33989 | 8792 | 5816 | 11279 |
| 490 | 750 | 519 | 449 | 721 | 580 | 502 | 757 | 27086 | 22654 | 46239 | 34679 | 12406 | 10747 | 23912 |
| 480 | 920 | 538 | 391 | 865 | 615 | 401 | 930 | 9550 | 7435 | 15484 | 14251 | 14908 | 10834 | 27715 |
| 1233 | 988 | 1464 | 1438 | 1025 | 1278 | 1172 | 970 | 20603 | 18988 | 15397 | 15208 | 34348 | 33725 | 23456 |
| 573 | 1103 | 866 | 734 | 1275 | 837 | 677 | 1227 | 18742 | 14833 | 25887 | 28550 | 21787 | 18468 | 25165 |
| 649 | 1021 | 863 | 804 | 1117 | 671 | 566 | 919 | 7085 | 5953 | 9178 | 9367 | 20639 | 19224 | 23922 |
| 711 | 1169 | 863 | 746 | 1193 | 870 | 680 | 1051 | 12580 | 10748 | 15108 | 17666 | 15500 | 13409 | 17971 |
| 839 | 1201 | 943 | 877 | 1180 | 909 | 730 | 1023 | 10828 | 9857 | 11755 | 14114 | 7617 | 7088 | 8080 |
| 887 | 1260 | 1044 | 867 | 1206 | 1171 | 1039 | 1258 | 24193 | 21217 | 23913 | 30124 | 14068 | 11675 | 13471 |
| 712 | 1096 | 780 | 615 | 1018 | 886 | 763 | 1115 | 21952 | 18027 | 25324 | 27752 | 20400 | 16070 | 26143 |
| 345 | 894 | 632 | 404 | 924 | 654 | 505 | 987 | 20020 | 12570 | 36412 | 32561 | 22708 | 14539 | 35948 |
| 455 | 808 | 725 | 395 | 745 | 936 | 472 | 843 | 14546 | 9655 | 21231 | 17148 | 7285 | 3974 | 10048 |
| 746 | 1058 | 1025 | 774 | 1090 | 992 | 748 | 1084 | 34088 | 25453 | 34140 | 36107 | 13482 | 10180 | 13159 |
| 676 | 1240 | 983 | 864 | 1332 | 1012 | 938 | 1405 | 15356 | 12114 | 17925 | 22222 | 9063 | 7970 | 9220 |
| 828 | 1046 | 911 | 745 | 1058 | 924 | 691 | 1049 | 9420 | 8489 | 10258 | 10726 | 20765 | 16972 | 22788 |
| 477 | 727 | 699 | 417 | 645 | 633 | 423 | 663 | 23407 | 16532 | 34693 | 25214 | 26027 | 15531 | 37215 |
| 649 | 986 | 860 | 665 | 1010 | 824 | 611 | 991 | 17799 | 12873 | 19833 | 19553 | 15513 | 11988 | 18032 |
| 276 | 615 | 461 | 309 | 672 | 426 | 289 | 638 | 10255 | 6970 | 25238 | 15518 | 17538 | 11776 | 38052 |
| 564 | 1025 | 900 | 731 | 964 | 780 | 596 | 994 | 20315 | 16303 | 28893 | 29625 | 10025 | 8144 | 11134 |
| 816 | 1109 | 800 | 686 | 976 | 783 | 605 | 1056 | 17601 | 14262 | 17468 | 19364 | 12270 | 10526 | 15337 |
| 474 | 852 | 647 | 518 | 862 | 717 | 619 | 977 | 19555 | 14686 | 30984 | 26392 | 8993 | 7200 | 13911 |
| 772 | 1246 | 678 | 672 | 1205 | 617 | 617 | 1138 | 16007 | 17845 | 23123 | 28803 | 22199 | 21986 | 32729 |
| 463 | 761 | 618 | 497 | 784 | 623 | 449 | 713 | 15709 | 11740 | 25374 | 19321 | 11354 | 9129 | 18369 |
| 571 | 767 | 729 | 435 | 706 | 743 | 500 | 763 | 18486 | 11233 | 19666 | 15077 | 21384 | 12742 | 29314 |
| 387 | 685 | 283 | 351 | 650 | 405 | 422 | 728 | 5037 | 5961 | 15407 | 10553 | 6860 | 8504 | 24198 |
| 1118 | 1649 | 1166 | 1222 | 1642 | 1099 | 949 | 1307 | 21509 | 21726 | 19437 | 32051 | 30941 | 32408 | 26529 |
| 624 | 960 | 809 | 683 | 1118 | 773 | 599 | 932 | 24372 | 19924 | 31932 | 30659 | 10098 | 8533 | 12486 |
| 713 | 1221 | 701 | 446 | 949 | 724 | 512 | 1024 | 34612 | 27072 | 37964 | 46371 | 16509 | 10495 | 23557 |
| 593 | 846 | 912 | 634 | 876 | 809 | 586 | 833 | 16743 | 11308 | 19067 | 16123 | 12984 | 9034 | 14240 |

| | | | | | | | | | | | | | | |
|---|---|---|---|---|---|---|---|---|---|---|---|---|---|---|
| 875 | 1164 | 907 | 808 | 1076 | 1098 | 1014 | 1274 | 18828 | 16891 | 19300 | 22469 | 20050 | 17864 | 22118 |
| 695 | 1259 | 899 | 692 | 1258 | 832 | 655 | 1186 | 10124 | 7969 | 11462 | 14426 | 8824 | 6793 | 9812 |
| 286 | 795 | 537 | 292 | 780 | 718 | 439 | 823 | 18886 | 9852 | 34434 | 27384 | 16193 | 8805 | 30137 |
| 858 | 1184 | 892 | 758 | 1178 | 952 | 870 | 1168 | 13498 | 12648 | 14748 | 17456 | 26173 | 22223 | 29336 |
| 1252 | 1459 | 1189 | 1391 | 1496 | 915 | 1081 | 1476 | 23668 | 27647 | 22087 | 32227 | 26954 | 31543 | 22669 |
| 637 | 1119 | 767 | 617 | 1081 | 947 | 849 | 1243 | 18572 | 15447 | 24241 | 27133 | 13208 | 10634 | 17225 |
| 401 | 739 | 421 | 305 | 684 | 443 | 330 | 689 | 14644 | 11406 | 28425 | 21004 | 22421 | 16230 | 53282 |
| 649 | 1025 | 734 | 512 | 892 | 684 | 527 | 898 | 14529 | 12324 | 19003 | 19484 | 12718 | 8873 | 17318 |
| 447 | 889 | 628 | 473 | 846 | 567 | 424 | 805 | 16673 | 11960 | 26782 | 23812 | 20010 | 15090 | 31887 |
| 328 | 865 | 715 | 346 | 925 | 651 | 321 | 935 | 32302 | 14062 | 42896 | 37090 | 32824 | 15884 | 45883 |
| 712 | 1088 | 1014 | 821 | 1171 | 951 | 673 | 1057 | 17578 | 13645 | 19156 | 20847 | 11746 | 9513 | 11586 |
| 1048 | 1465 | 940 | 1003 | 1392 | 931 | 847 | 1227 | 19544 | 20835 | 19876 | 29115 | 21023 | 22441 | 22364 |
| 636 | 1042 | 835 | 702 | 1126 | 869 | 663 | 1066 | 23365 | 18266 | 28713 | 29906 | 19621 | 16484 | 23486 |
| 770 | 1180 | 908 | 790 | 1155 | 898 | 729 | 1247 | 14722 | 13664 | 17742 | 20930 | 8008 | 6964 | 8817 |
| 571 | 963 | 691 | 560 | 959 | 666 | 560 | 941 | 12560 | 10667 | 18683 | 17995 | 7287 | 5908 | 10541 |
| 604 | 914 | 499 | 542 | 831 | 491 | 461 | 810 | 16510 | 17481 | 28954 | 26471 | 20727 | 22512 | 41566 |
| 595 | 895 | 949 | 808 | 1011 | 661 | 452 | 804 | 21745 | 16004 | 26877 | 24067 | 14608 | 12441 | 15400 |
| 636 | 869 | 831 | 682 | 892 | 654 | 528 | 805 | 31275 | 24773 | 38929 | 33811 | 31386 | 25757 | 37765 |
| 562 | 893 | 655 | 539 | 905 | 637 | 585 | 948 | 6656 | 5054 | 8990 | 8026 | 18219 | 15009 | 27819 |
| 801 | 1064 | 952 | 766 | 1087 | 1023 | 826 | 1122 | 11875 | 9643 | 12036 | 12810 | 29108 | 23397 | 30561 |
| 466 | 807 | 750 | 479 | 840 | 770 | 459 | 788 | 30650 | 18855 | 40419 | 32606 | 11381 | 7273 | 15169 |
| 828 | 903 | 1066 | 887 | 887 | 984 | 776 | 825 | 20989 | 18176 | 21941 | 19805 | 17143 | 14264 | 16084 |
| 393 | 746 | 649 | 502 | 708 | 574 | 434 | 699 | 15391 | 8827 | 22476 | 16768 | 24576 | 19019 | 37862 |
| 445 | 873 | 544 | 398 | 824 | 561 | 445 | 886 | 9200 | 7876 | 17713 | 15464 | 21124 | 15465 | 38839 |
| 632 | 959 | 792 | 701 | 1007 | 902 | 833 | 935 | 25737 | 21776 | 34436 | 33026 | 14485 | 12828 | 18291 |
| 486 | 916 | 763 | 491 | 887 | 697 | 494 | 911 | 18760 | 13040 | 26811 | 24569 | 17535 | 11275 | 22971 |
| 737 | 901 | 1281 | 766 | 910 | 1210 | 703 | 937 | 9581 | 5269 | 7153 | 6444 | 21596 | 12904 | 16855 |
| 643 | 940 | 738 | 526 | 743 | 721 | 490 | 777 | 25705 | 18388 | 28615 | 26885 | 12839 | 9156 | 17394 |
| 392 | 664 | 537 | 377 | 644 | 433 | 302 | 598 | 22103 | 15441 | 39434 | 26180 | 19869 | 13941 | 37027 |
| 451 | 756 | 742 | 432 | 776 | 779 | 481 | 786 | 11502 | 7315 | 16211 | 12259 | 10659 | 6200 | 14363 |

| | | | | | | | | | | | | | | |
|---|---|---|---|---|---|---|---|---|---|---|---|---|---|---|
| 477 | 739 | 1270 | 486 | 740 | 1278 | 433 | 738 | 21849 | 8113 | 16991 | 12558 | 35457 | 13577 | 27923 |
| 830 | 1335 | 1024 | 817 | 1246 | 1056 | 971 | 1231 | 30368 | 24372 | 29374 | 39229 | 23140 | 18466 | 22593 |
| 502 | 775 | 725 | 616 | 840 | 674 | 518 | 764 | 14957 | 11099 | 22129 | 17159 | 29347 | 24925 | 40470 |
| 589 | 862 | 723 | 618 | 906 | 837 | 567 | 952 | 17466 | 14461 | 24556 | 21175 | 9482 | 8110 | 13124 |
| 599 | 892 | 1089 | 700 | 926 | 1022 | 674 | 866 | 16685 | 11287 | 18836 | 16796 | 21006 | 13499 | 19284 |
| 550 | 790 | 978 | 709 | 1022 | 773 | 608 | 936 | 18611 | 13501 | 24558 | 19411 | 10579 | 7666 | 10816 |
| 305 | 636 | 708 | 411 | 762 | 771 | 430 | 731 | 27530 | 12455 | 40878 | 26000 | 17713 | 10265 | 25004 |
| 487 | 893 | 812 | 555 | 1050 | 739 | 491 | 879 | 20217 | 12413 | 25480 | 22747 | 14943 | 10217 | 18411 |
| 630 | 1132 | 778 | 623 | 1145 | 803 | 650 | 1183 | 10867 | 8548 | 13560 | 15351 | 15090 | 12081 | 19393 |
| 692 | 880 | 1051 | 873 | 1071 | 1200 | 791 | 905 | 8508 | 6403 | 9252 | 8144 | 3243 | 2692 | 3084 |
| 675 | 927 | 1020 | 742 | 955 | 1003 | 710 | 969 | 6050 | 4340 | 6429 | 5960 | 13419 | 9766 | 13156 |
| 514 | 1028 | 608 | 528 | 1097 | 692 | 666 | 1090 | 17854 | 15190 | 29525 | 30361 | 6848 | 5949 | 11272 |
| 825 | 1152 | 845 | 717 | 1243 | 896 | 731 | 1165 | 10731 | 8912 | 10804 | 12447 | 7231 | 6136 | 8554 |
| 742 | 1015 | 639 | 735 | 1058 | 687 | 685 | 962 | 10097 | 9250 | 12472 | 12654 | 8177 | 9405 | 12798 |
| 531 | 1243 | 597 | 519 | 1155 | 633 | 540 | 1221 | 16199 | 14638 | 27558 | 34246 | 12997 | 11307 | 21767 |
| 705 | 1025 | 804 | 657 | 957 | 682 | 523 | 912 | 15892 | 13665 | 19386 | 19862 | 31480 | 25702 | 39149 |
| 449 | 831 | 540 | 389 | 820 | 598 | 412 | 834 | 11219 | 8246 | 18356 | 15248 | 15844 | 11414 | 29353 |
| 846 | 1267 | 807 | 679 | 1183 | 715 | 652 | 1170 | 19905 | 17787 | 21032 | 26646 | 24094 | 20280 | 29859 |
| 725 | 828 | 556 | 564 | 818 | 797 | 572 | 791 | 8108 | 9138 | 12599 | 10433 | 12598 | 12792 | 22663 |
| 779 | 1055 | 913 | 726 | 983 | 710 | 534 | 784 | 11041 | 8015 | 10291 | 10854 | 23845 | 18963 | 26127 |
| 965 | 1240 | 1148 | 938 | 1256 | 1096 | 992 | 1278 | 17955 | 15359 | 15923 | 19745 | 25639 | 20955 | 22333 |
| 926 | 1516 | 1018 | 944 | 1529 | 973 | 921 | 1488 | 22126 | 20266 | 21888 | 33193 | 24425 | 22644 | 23997 |
| 491 | 791 | 612 | 506 | 827 | 577 | 472 | 833 | 10159 | 7774 | 15826 | 12516 | 3019 | 2496 | 4936 |
| 454 | 758 | 649 | 402 | 637 | 606 | 449 | 749 | 17221 | 10265 | 22625 | 17144 | 13881 | 8597 | 21403 |
| 395 | 1018 | 591 | 441 | 982 | 508 | 397 | 991 | 14633 | 11429 | 28932 | 29461 | 13835 | 10325 | 23393 |
| 639 | 1013 | 774 | 614 | 979 | 662 | 543 | 871 | 22324 | 18801 | 29411 | 29790 | 17318 | 13740 | 22387 |
| 633 | 1173 | 809 | 641 | 1158 | 780 | 637 | 1191 | 29742 | 23473 | 37056 | 43456 | 13735 | 10892 | 16984 |
| 669 | 1133 | 804 | 725 | 1210 | 818 | 801 | 1378 | 19438 | 15948 | 23824 | 27002 | 24952 | 22498 | 31035 |
| 580 | 989 | 769 | 629 | 1096 | 851 | 680 | 1045 | 18028 | 13780 | 23750 | 23479 | 20746 | 16976 | 26974 |
| 649 | 1123 | 732 | 601 | 1089 | 767 | 643 | 1145 | 22611 | 18796 | 28957 | 32507 | 19508 | 16016 | 26648 |
| 1345 | 1745 | 966 | 922 | 1412 | 918 | 628 | 992 | 14452 | 14729 | 10953 | 19117 | 8678 | 8289 | 8987 |

| | | | | | | | | | | | | | | |
|---|---|---|---|---|---|---|---|---|---|---|---|---|---|---|
| 553 | 852 | 906 | 644 | 956 | 869 | 592 | 863 | 16603 | 9131 | 16517 | 14077 | 18855 | 13410 | 20816 |
| 859 | 1108 | 1040 | 781 | 1033 | 1158 | 850 | 1047 | 27571 | 20843 | 24258 | 26869 | 25185 | 18924 | 24217 |
| 413 | 594 | 565 | 501 | 659 | 625 | 540 | 699 | 4962 | 4044 | 9799 | 5817 | 6315 | 5599 | 11186 |
| 532 | 942 | 820 | 545 | 929 | 763 | 490 | 880 | 13873 | 8861 | 16665 | 15695 | 16209 | 10766 | 19757 |
| 553 | 907 | 884 | 456 | 697 | 770 | 595 | 979 | 20377 | 15131 | 27360 | 24818 | 9531 | 4917 | 10779 |
| 432 | 690 | 506 | 482 | 661 | 457 | 450 | 668 | 9927 | 8484 | 19626 | 13551 | 20133 | 19173 | 39789 |
| 375 | 756 | 630 | 387 | 733 | 583 | 359 | 723 | 23508 | 13850 | 36943 | 27919 | 9907 | 6084 | 15727 |
| 478 | 842 | 606 | 539 | 906 | 592 | 486 | 907 | 15833 | 13562 | 28359 | 23866 | 13600 | 12087 | 22442 |
| 706 | 1226 | 1079 | 691 | 1139 | 859 | 823 | 1378 | 11324 | 8260 | 11706 | 14354 | 10285 | 6585 | 9530 |
| 392 | 670 | 762 | 562 | 766 | 613 | 418 | 680 | 11597 | 8315 | 21224 | 14214 | 13484 | 9936 | 17690 |
| 333 | 938 | 468 | 338 | 897 | 359 | 260 | 862 | 6775 | 4968 | 14934 | 14015 | 6199 | 4479 | 13254 |
| 509 | 822 | 589 | 397 | 740 | 762 | 558 | 873 | 17202 | 12640 | 24848 | 20419 | 17391 | 11718 | 29508 |
| 369 | 626 | 549 | 392 | 658 | 598 | 476 | 685 | 14089 | 10290 | 27904 | 17471 | 15661 | 11173 | 28516 |
| 631 | 953 | 1047 | 687 | 955 | 977 | 631 | 964 | 6677 | 4116 | 6528 | 6223 | 21669 | 14210 | 20688 |
| 782 | 1225 | 949 | 818 | 1223 | 982 | 827 | 1304 | 22510 | 19081 | 24394 | 29880 | 23876 | 20581 | 25157 |
| 483 | 839 | 1045 | 474 | 951 | 905 | 434 | 781 | 21848 | 11013 | 22817 | 19142 | 17581 | 7965 | 16817 |
| 647 | 1243 | 783 | 643 | 1213 | 864 | 729 | 1262 | 22678 | 18654 | 28833 | 35847 | 23581 | 19381 | 30119 |
| 595 | 896 | 764 | 629 | 960 | 648 | 519 | 848 | 14999 | 13886 | 23330 | 20912 | 12493 | 10298 | 16362 |
| 875 | 1033 | 1058 | 927 | 1085 | 958 | 814 | 995 | 24026 | 20260 | 23160 | 23920 | 25008 | 21915 | 23633 |
| 365 | 689 | 643 | 455 | 788 | 706 | 476 | 820 | 5396 | 3537 | 9683 | 6673 | 10620 | 7512 | 16505 |
| 507 | 837 | 533 | 426 | 696 | 778 | 600 | 895 | 15083 | 11608 | 22885 | 19164 | 6851 | 5470 | 12848 |
| 629 | 1014 | 746 | 573 | 964 | 819 | 653 | 1031 | 7884 | 6543 | 10393 | 10538 | 11130 | 8550 | 14922 |
| 474 | 883 | 790 | 593 | 1015 | 717 | 508 | 921 | 8870 | 6342 | 13381 | 11816 | 12946 | 9719 | 16392 |
| 698 | 990 | 871 | 664 | 1026 | 896 | 725 | 965 | 10681 | 8716 | 12484 | 12356 | 7615 | 5806 | 8738 |
| 894 | 1116 | 852 | 731 | 1005 | 1030 | 907 | 1107 | 9217 | 7747 | 8666 | 9671 | 3512 | 3014 | 4123 |
| 538 | 907 | 847 | 527 | 826 | 906 | 604 | 877 | 15574 | 10165 | 18903 | 17154 | 10085 | 6267 | 11900 |
| 594 | 932 | 855 | 584 | 899 | 877 | 585 | 962 | 28119 | 18230 | 30687 | 28593 | 18186 | 12418 | 21274 |
| 390 | 706 | 457 | 286 | 645 | 538 | 344 | 702 | 21698 | 14350 | 36826 | 25982 | 20150 | 12590 | 44070 |
| 659 | 967 | 744 | 698 | 963 | 667 | 640 | 957 | 19522 | 17141 | 26005 | 25150 | 22131 | 20768 | 29734 |
| 594 | 895 | 878 | 689 | 960 | 920 | 730 | 961 | 12447 | 9864 | 16605 | 14860 | 24184 | 18975 | 27534 |
| 344 | 660 | 650 | 395 | 754 | 480 | 342 | 633 | 7628 | 5270 | 15311 | 10105 | 21502 | 13079 | 33101 |

| | | | | | | | | | | | | | | |
|---|---|---|---|---|---|---|---|---|---|---|---|---|---|---|
| 472 | 779 | 639 | 486 | 848 | 597 | 480 | 822 | 13566 | 10410 | 22064 | 17185 | 15418 | 11734 | 24142 |
| 619 | 1040 | 657 | 631 | 1161 | 648 | 481 | 952 | 7272 | 6954 | 11241 | 11695 | 5640 | 5409 | 8578 |
| 710 | 1082 | 739 | 674 | 1066 | 768 | 683 | 1027 | 10892 | 10504 | 14793 | 15999 | 17297 | 15760 | 23399 |
| 552 | 913 | 747 | 569 | 910 | 715 | 491 | 802 | 21914 | 16993 | 30812 | 28133 | 14063 | 10708 | 18816 |
| 945 | 1306 | 990 | 824 | 1223 | 923 | 819 | 1189 | 15403 | 13950 | 14764 | 19278 | 23899 | 19895 | 24148 |
| 620 | 1179 | 520 | 441 | 806 | 746 | 621 | 1210 | 17366 | 13125 | 21180 | 24968 | 10073 | 8547 | 19381 |
| 797 | 1270 | 868 | 823 | 1276 | 828 | 708 | 1143 | 9782 | 9768 | 12263 | 15573 | 19630 | 18601 | 22612 |
| 967 | 1397 | 882 | 949 | 1367 | 923 | 941 | 1362 | 18683 | 19279 | 19931 | 27838 | 12107 | 13021 | 13726 |
| 488 | 849 | 696 | 534 | 813 | 875 | 813 | 858 | 17033 | 13873 | 28419 | 24140 | 19799 | 15185 | 28433 |
| 647 | 871 | 786 | 700 | 884 | 757 | 596 | 832 | 8064 | 7468 | 11541 | 10051 | 9812 | 8741 | 12482 |
| 947 | 1294 | 875 | 806 | 1091 | 932 | 826 | 1177 | 8328 | 8137 | 8591 | 11115 | 8962 | 8256 | 10244 |
| 472 | 947 | 618 | 411 | 831 | 830 | 625 | 976 | 9073 | 6332 | 13429 | 12716 | 11851 | 7887 | 19187 |
| 680 | 829 | 1245 | 1074 | 1196 | 1435 | 1271 | 1449 | 2730 | 2403 | 3532 | 2930 | 6675 | 5755 | 5360 |
| 474 | 843 | 508 | 423 | 742 | 568 | 448 | 787 | 16198 | 12530 | 26411 | 22272 | 7485 | 6236 | 14728 |
| 772 | 1246 | 942 | 762 | 1143 | 870 | 569 | 893 | 8067 | 6967 | 9024 | 11239 | 16159 | 13083 | 17160 |
| 950 | 1291 | 832 | 942 | 1341 | 825 | 905 | 1318 | 16889 | 18178 | 19140 | 24712 | 14599 | 16541 | 17554 |
| 691 | 1402 | 914 | 884 | 1397 | 921 | 825 | 1443 | 12634 | 10556 | 15276 | 21419 | 6419 | 6208 | 7021 |
| 398 | 707 | 629 | 406 | 709 | 663 | 468 | 757 | 19751 | 12821 | 32181 | 22765 | 23808 | 15357 | 37827 |
| 824 | 1171 | 906 | 763 | 1189 | 1018 | 851 | 1287 | 21322 | 18032 | 21889 | 25643 | 13058 | 10999 | 14418 |
| 722 | 1042 | 963 | 702 | 1023 | 925 | 666 | 883 | 4258 | 3520 | 4878 | 5085 | 8655 | 6308 | 8986 |
| 580 | 835 | 829 | 624 | 890 | 606 | 340 | 610 | 10587 | 7321 | 12613 | 10528 | 9361 | 7053 | 11298 |
| 144 | 320 | 757 | 175 | 348 | 764 | 155 | 296 | 16902 | 3349 | 23201 | 7424 | 24122 | 5587 | 31880 |
| 455 | 773 | 718 | 474 | 831 | 679 | 449 | 915 | 17554 | 11119 | 24435 | 18879 | 11657 | 7693 | 16242 |
| 876 | 1288 | 1033 | 1056 | 1442 | 989 | 950 | 1343 | 13348 | 13576 | 15497 | 19956 | 20180 | 20614 | 19529 |
| 850 | 1196 | 802 | 673 | 961 | 1005 | 832 | 1213 | 20688 | 18527 | 21795 | 26065 | 5822 | 4882 | 7255 |
| 709 | 1156 | 761 | 694 | 1105 | 918 | 674 | 1029 | 12530 | 11335 | 15981 | 18482 | 13923 | 12691 | 18290 |
| 802 | 1118 | 786 | 623 | 1035 | 793 | 736 | 1082 | 15140 | 13773 | 17177 | 19204 | 17579 | 13944 | 22377 |
| 615 | 1216 | 709 | 556 | 1129 | 1015 | 1046 | 1345 | 10564 | 8622 | 14024 | 17050 | 12378 | 9705 | 17470 |
| 337 | 751 | 615 | 375 | 822 | 395 | 240 | 640 | 9615 | 6126 | 18177 | 13653 | 9569 | 5828 | 15548 |
| 963 | 1283 | 1314 | 910 | 1185 | 1475 | 917 | 1197 | 10873 | 8793 | 9127 | 11712 | 7542 | 5220 | 5738 |
| 904 | 1235 | 961 | 995 | 1379 | 943 | 811 | 1168 | 13631 | 13164 | 14555 | 17970 | 10119 | 10475 | 10528 |

| | | | | | | | | | | | | | | |
|---|---|---|---|---|---|---|---|---|---|---|---|---|---|---|
| 852 | 1251 | 915 | 823 | 1276 | 845 | 746 | 1313 | 7830 | 7055 | 8276 | 10354 | 12180 | 10965 | 13316 |
| 371 | 750 | 506 | 323 | 710 | 516 | 312 | 736 | 25911 | 16390 | 44201 | 33160 | 17575 | 11228 | 34761 |
| 455 | 811 | 1083 | 523 | 830 | 836 | 456 | 869 | 10366 | 5651 | 12430 | 10078 | 3972 | 1919 | 3667 |
| 374 | 564 | 626 | 459 | 602 | 592 | 406 | 625 | 8677 | 6344 | 16976 | 9576 | 7742 | 5679 | 12363 |
| 829 | 1278 | 1118 | 1069 | 1574 | 1302 | 1334 | 2151 | 3171 | 2737 | 3301 | 4218 | 3253 | 3111 | 2910 |
| 777 | 990 | 789 | 724 | 965 | 867 | 817 | 1061 | 15407 | 15319 | 19707 | 19511 | 12353 | 11347 | 15665 |
| 541 | 777 | 757 | 624 | 856 | 717 | 563 | 806 | 6227 | 5395 | 9965 | 7743 | 8664 | 7145 | 11444 |
| 365 | 931 | 453 | 323 | 782 | 770 | 368 | 927 | 531 | 413 | 1133 | 1054 | 795 | 567 | 1754 |
| 691 | 1263 | 773 | 617 | 975 | 1281 | 1059 | 1320 | 17880 | 12940 | 18728 | 23657 | 14569 | 11622 | 18851 |
| 422 | 688 | 773 | 445 | 749 | 720 | 392 | 697 | 3600 | 2027 | 4801 | 3303 | 8498 | 4897 | 10994 |
| 574 | 905 | 775 | 674 | 922 | 791 | 709 | 865 | 17815 | 13614 | 23719 | 21463 | 6225 | 5414 | 8034 |
| 585 | 945 | 728 | 606 | 921 | 828 | 658 | 945 | 9732 | 7571 | 12932 | 12218 | 17769 | 14796 | 24424 |
| 674 | 1138 | 884 | 706 | 1206 | 888 | 628 | 1056 | 30369 | 23471 | 34843 | 39665 | 14574 | 11648 | 16494 |
| 1140 | 1214 | 1426 | 1292 | 1282 | 1260 | 1150 | 1322 | 15988 | 15053 | 13208 | 16030 | 5349 | 4845 | 3750 |
| 557 | 859 | 716 | 634 | 878 | 693 | 576 | 880 | 14286 | 11914 | 21389 | 18375 | 9254 | 8190 | 12916 |
| 850 | 1196 | 1122 | 828 | 1160 | 1094 | 811 | 1167 | 13165 | 10248 | 12050 | 14412 | 8108 | 5986 | 7228 |
| 991 | 1051 | 1204 | 963 | 990 | 1261 | 1089 | 1107 | 26681 | 21106 | 21290 | 22380 | 19492 | 15591 | 16185 |
| 901 | 1373 | 997 | 932 | 1411 | 1029 | 1057 | 1632 | 15464 | 14217 | 15786 | 21679 | 8901 | 8320 | 8926 |
| 765 | 1176 | 859 | 820 | 1217 | 870 | 826 | 1275 | 10191 | 8413 | 11003 | 12938 | 15376 | 14681 | 17896 |
| 543 | 849 | 811 | 518 | 839 | 845 | 610 | 899 | 8999 | 6405 | 11798 | 10012 | 19574 | 12500 | 24130 |
| 365 | 717 | 517 | 368 | 733 | 529 | 355 | 697 | 3621 | 2496 | 6835 | 4902 | 10328 | 7347 | 19967 |
| 710 | 984 | 892 | 728 | 989 | 895 | 695 | 942 | 14610 | 11973 | 16866 | 16597 | 13969 | 11398 | 15656 |
| 800 | 1074 | 831 | 616 | 984 | 1003 | 756 | 1034 | 13867 | 10809 | 13505 | 14504 | 9297 | 6889 | 11182 |
| 846 | 1211 | 950 | 770 | 1147 | 969 | 761 | 1178 | 17161 | 13403 | 15850 | 19197 | 14576 | 11817 | 15350 |
| 882 | 1268 | 851 | 952 | 1575 | 1163 | 1237 | 1618 | 8864 | 6895 | 7817 | 9910 | 8365 | 9358 | 9833 |
| 569 | 902 | 1004 | 666 | 1010 | 886 | 577 | 922 | 10148 | 6441 | 11321 | 10206 | 15826 | 10491 | 15764 |
| 667 | 1076 | 858 | 659 | 1079 | 817 | 625 | 1068 | 20353 | 15805 | 23688 | 25489 | 12052 | 9255 | 14046 |
| 836 | 1167 | 1012 | 680 | 1029 | 919 | 773 | 1103 | 30613 | 25754 | 30817 | 35953 | 20376 | 13686 | 20140 |
| 749 | 1300 | 1026 | 922 | 1472 | 1039 | 826 | 1332 | 13537 | 11328 | 15115 | 19652 | 13978 | 12569 | 13626 |
| 729 | 1125 | 997 | 555 | 902 | 1021 | 564 | 854 | 22207 | 13735 | 18832 | 21187 | 20673 | 11523 | 20745 |
| 633 | 931 | 831 | 528 | 872 | 930 | 619 | 909 | 31130 | 20242 | 31977 | 29763 | 16794 | 10658 | 20203 |

| | | | | | | | | | | | | | | |
|---|---|---|---|---|---|---|---|---|---|---|---|---|---|---|
| 200 | 532 | 479 | 284 | 615 | 407 | 208 | 549 | 8514 | 4789 | 23953 | 12753 | 16605 | 9825 | 34653 |
| 501 | 922 | 865 | 549 | 986 | 771 | 424 | 956 | 7608 | 4439 | 8861 | 8170 | 6259 | 3970 | 7233 |
| 677 | 1071 | 958 | 747 | 1186 | 993 | 683 | 1137 | 9924 | 6972 | 10298 | 11033 | 10355 | 8079 | 10814 |
| 259 | 596 | 512 | 314 | 673 | 517 | 302 | 629 | 4625 | 2792 | 10770 | 6421 | 9432 | 5781 | 18427 |
| 778 | 1125 | 899 | 972 | 1222 | 772 | 776 | 1122 | 9828 | 10100 | 12989 | 14607 | 24131 | 26106 | 26848 |
| 729 | 892 | 611 | 461 | 714 | 498 | 302 | 632 | 3140 | 2618 | 3590 | 3203 | 10903 | 8219 | 17845 |
| 941 | 972 | 1061 | 892 | 1009 | 1025 | 842 | 1005 | 7372 | 6049 | 6428 | 6246 | 6309 | 5301 | 5946 |
| 650 | 1028 | 746 | 484 | 874 | 812 | 587 | 967 | 16094 | 12324 | 18962 | 19498 | 5620 | 3644 | 7534 |
| 369 | 756 | 548 | 402 | 797 | 643 | 484 | 880 | 3603 | 2345 | 6351 | 4801 | 9878 | 7240 | 18025 |
| 385 | 754 | 618 | 433 | 823 | 619 | 353 | 694 | 16793 | 11080 | 28804 | 21713 | 11442 | 8015 | 18528 |
| 514 | 748 | 1053 | 639 | 970 | 851 | 507 | 721 | 17781 | 10363 | 20174 | 15090 | 14233 | 8634 | 13517 |
| 527 | 880 | 797 | 583 | 964 | 773 | 557 | 956 | 5890 | 4366 | 8284 | 7286 | 12291 | 8997 | 15430 |
| 474 | 892 | 914 | 505 | 889 | 916 | 468 | 907 | 24422 | 13298 | 28072 | 25045 | 18361 | 10142 | 20081 |
| 391 | 771 | 703 | 402 | 813 | 550 | 300 | 709 | 11548 | 6546 | 16752 | 12923 | 26564 | 15187 | 37803 |
| 769 | 1207 | 926 | 790 | 1173 | 920 | 797 | 1268 | 21677 | 18004 | 23427 | 28268 | 30553 | 26062 | 32992 |
| 501 | 790 | 660 | 578 | 811 | 646 | 563 | 900 | 8173 | 6652 | 13280 | 10487 | 6504 | 5696 | 9861 |
| 443 | 760 | 591 | 381 | 732 | 704 | 468 | 724 | 19176 | 12652 | 28546 | 21689 | 8397 | 5412 | 14197 |
| 846 | 1176 | 1033 | 750 | 1049 | 891 | 662 | 1052 | 12824 | 9218 | 10890 | 12809 | 6449 | 4678 | 6240 |
| 601 | 938 | 956 | 541 | 957 | 1138 | 709 | 1090 | 7611 | 4331 | 7210 | 6763 | 7155 | 4049 | 7488 |
| 564 | 842 | 779 | 493 | 820 | 815 | 426 | 821 | 21264 | 17038 | 30217 | 25434 | 22259 | 14086 | 28566 |
| 380 | 613 | 502 | 336 | 596 | 504 | 318 | 595 | 21086 | 13116 | 34483 | 21126 | 6946 | 4643 | 13830 |
| 529 | 768 | 833 | 679 | 883 | 786 | 600 | 859 | 2983 | 1932 | 3648 | 2802 | 22806 | 18591 | 27369 |
| 872 | 1238 | 1020 | 774 | 1128 | 1032 | 804 | 1183 | 10001 | 8481 | 9729 | 12045 | 7827 | 5940 | 7675 |
| 672 | 1012 | 886 | 756 | 1198 | 802 | 718 | 1110 | 13645 | 11080 | 16496 | 16697 | 9561 | 8162 | 10793 |
| 248 | 542 | 511 | 255 | 554 | 501 | 225 | 532 | 10576 | 4767 | 19188 | 10397 | 15616 | 7785 | 30550 |
| 790 | 1189 | 889 | 741 | 1124 | 933 | 784 | 1121 | 9673 | 8758 | 11087 | 13188 | 17887 | 14901 | 20116 |
| 710 | 1032 | 1167 | 628 | 903 | 995 | 601 | 943 | 12206 | 7576 | 10673 | 11016 | 2990 | 1609 | 2562 |
| 638 | 1103 | 932 | 765 | 1058 | 1141 | 916 | 1130 | 7913 | 5620 | 8814 | 9720 | 8430 | 6919 | 9046 |
| 366 | 643 | 605 | 429 | 701 | 594 | 415 | 671 | 10497 | 6336 | 17322 | 11130 | 10488 | 7425 | 17326 |
| 604 | 952 | 795 | 583 | 939 | 718 | 550 | 1037 | 16666 | 12273 | 20319 | 19340 | 9420 | 6914 | 11851 |
| 345 | 695 | 453 | 291 | 538 | 531 | 439 | 673 | 16989 | 11494 | 33363 | 23184 | 15191 | 9751 | 33561 |

| | | | | | | | | | | | | | | |
|---|---|---|---|---|---|---|---|---|---|---|---|---|---|---|
| 594 | 868 | 982 | 794 | 1105 | 746 | 585 | 964 | 17385 | 13336 | 22433 | 19467 | 10905 | 8816 | 11100 |
| 731 | 1315 | 878 | 711 | 1223 | 886 | 745 | 1253 | 8855 | 7435 | 10172 | 13373 | 13091 | 10599 | 14906 |
| 945 | 1403 | 1122 | 999 | 1484 | 1014 | 908 | 1331 | 23686 | 21045 | 22271 | 31243 | 23490 | 20909 | 20930 |
| 399 | 627 | 472 | 333 | 586 | 526 | 355 | 584 | 9137 | 6181 | 15501 | 9712 | 6777 | 4781 | 14351 |
| 432 | 804 | 765 | 511 | 837 | 692 | 435 | 791 | 7914 | 4690 | 10854 | 8728 | 11519 | 7687 | 15055 |
| 544 | 920 | 822 | 648 | 1039 | 867 | 657 | 919 | 8795 | 6617 | 12163 | 11191 | 13617 | 10733 | 16563 |
| 320 | 716 | 592 | 345 | 698 | 530 | 311 | 709 | 12582 | 7033 | 22011 | 15771 | 15728 | 9153 | 26563 |
| 758 | 1143 | 854 | 650 | 939 | 936 | 874 | 1319 | 5688 | 5073 | 6691 | 7646 | 6285 | 4782 | 7357 |
| 793 | 1156 | 736 | 656 | 1034 | 759 | 657 | 1061 | 11943 | 11818 | 14912 | 17236 | 6461 | 5757 | 8783 |
| 644 | 1016 | 838 | 622 | 945 | 845 | 693 | 968 | 16757 | 12350 | 19167 | 19470 | 8581 | 6372 | 10238 |
| 747 | 1157 | 998 | 832 | 1231 | 976 | 758 | 1220 | 25181 | 20049 | 26825 | 31034 | 17898 | 14919 | 17941 |
| 583 | 963 | 739 | 623 | 1005 | 762 | 650 | 960 | 10437 | 8446 | 14488 | 13951 | 22801 | 19215 | 30835 |
| 862 | 1237 | 957 | 858 | 1228 | 980 | 808 | 1160 | 14671 | 11721 | 13591 | 16814 | 11609 | 10416 | 12133 |
| 726 | 1175 | 894 | 631 | 987 | 984 | 678 | 1015 | 15193 | 10670 | 14703 | 17281 | 12752 | 9003 | 14268 |
| 787 | 1169 | 911 | 767 | 1147 | 805 | 683 | 1007 | 16761 | 14482 | 18409 | 21525 | 13740 | 11561 | 15080 |
| 454 | 753 | 710 | 510 | 777 | 682 | 495 | 800 | 10583 | 7563 | 16640 | 12534 | 16456 | 11821 | 23183 |
| 491 | 862 | 675 | 537 | 898 | 630 | 521 | 864 | 15332 | 11056 | 22499 | 19402 | 15223 | 12116 | 22553 |
| 640 | 857 | 958 | 734 | 979 | 1016 | 831 | 930 | 7162 | 5208 | 8134 | 6967 | 6242 | 4782 | 6518 |
| 335 | 864 | 482 | 313 | 751 | 607 | 395 | 867 | 1546 | 1066 | 3184 | 2751 | 2667 | 1733 | 5531 |
| 563 | 1178 | 631 | 528 | 1246 | 703 | 591 | 1190 | 8824 | 8195 | 14557 | 17152 | 3703 | 3098 | 5872 |
| 832 | 1223 | 1051 | 987 | 1503 | 953 | 889 | 1252 | 30198 | 26439 | 31784 | 38887 | 7035 | 6602 | 6692 |
| 798 | 1294 | 898 | 795 | 1349 | 850 | 703 | 1147 | 10594 | 9784 | 12268 | 15875 | 13893 | 12295 | 15463 |
| 573 | 891 | 698 | 524 | 916 | 803 | 723 | 1057 | 7308 | 5338 | 9320 | 8308 | 9082 | 6814 | 13002 |
| 545 | 773 | 781 | 443 | 684 | 810 | 483 | 689 | 16970 | 10470 | 19223 | 14856 | 4942 | 2800 | 6326 |
| 827 | 1267 | 882 | 772 | 1161 | 923 | 892 | 1189 | 15336 | 14677 | 17746 | 22485 | 12128 | 10620 | 13757 |
| 722 | 1067 | 943 | 707 | 1070 | 982 | 797 | 1168 | 16102 | 12473 | 17280 | 18430 | 26244 | 19659 | 27823 |
| 386 | 539 | 722 | 293 | 429 | 731 | 398 | 618 | 15207 | 8210 | 21243 | 11455 | 6422 | 2612 | 8900 |
| 357 | 634 | 695 | 399 | 678 | 732 | 438 | 725 | 8600 | 4561 | 12767 | 8093 | 6871 | 3947 | 9885 |
| 708 | 1149 | 990 | 871 | 1204 | 873 | 736 | 1071 | 5381 | 3863 | 5460 | 6272 | 9497 | 8352 | 9591 |
| 821 | 1445 | 818 | 673 | 1143 | 1044 | 879 | 1540 | 13827 | 11288 | 13753 | 19875 | 17768 | 14612 | 21723 |
| 303 | 588 | 574 | 371 | 643 | 534 | 335 | 611 | 22781 | 13476 | 44515 | 26178 | 23878 | 15458 | 41634 |

| | | | | | | | | | | | | | | |
|---|---|---|---|---|---|---|---|---|---|---|---|---|---|---|
| 483 | 980 | 624 | 412 | 872 | 739 | 585 | 968 | 13541 | 10736 | 22225 | 21788 | 19507 | 12887 | 31245 |
| 626 | 973 | 654 | 534 | 882 | 693 | 537 | 951 | 15053 | 12714 | 20303 | 19751 | 12991 | 10592 | 19853 |
| 634 | 874 | 844 | 666 | 967 | 860 | 688 | 1003 | 27724 | 21188 | 33413 | 29197 | 14648 | 11562 | 17361 |
| 496 | 616 | 297 | 518 | 679 | 289 | 464 | 608 | 3265 | 7100 | 14301 | 8804 | 7074 | 12354 | 23859 |
| 833 | 1194 | 960 | 815 | 1144 | 1028 | 845 | 1195 | 9130 | 7760 | 9311 | 11112 | 15291 | 12977 | 15922 |
| 746 | 1278 | 854 | 691 | 1169 | 850 | 702 | 1192 | 11189 | 9266 | 12421 | 15868 | 13816 | 11186 | 16177 |
| 769 | 1085 | 1075 | 806 | 1116 | 1128 | 806 | 1149 | 19919 | 13965 | 18165 | 19718 | 25260 | 18944 | 23493 |
| 495 | 769 | 663 | 584 | 862 | 573 | 469 | 744 | 8802 | 8123 | 16409 | 12621 | 10184 | 8982 | 15367 |
| 675 | 1040 | 742 | 664 | 1014 | 787 | 652 | 937 | 15970 | 15016 | 22244 | 23138 | 1574 | 1408 | 2121 |
| 497 | 841 | 634 | 503 | 772 | 558 | 477 | 807 | 11664 | 8841 | 17773 | 14954 | 7310 | 5808 | 11538 |
| 257 | 869 | 414 | 348 | 800 | 283 | 277 | 804 | 4701 | 5547 | 21580 | 18763 | 5800 | 4872 | 14012 |
| 493 | 846 | 681 | 493 | 838 | 745 | 571 | 901 | 13982 | 10301 | 20898 | 17681 | 15846 | 11469 | 23259 |
| 1096 | 1347 | 1105 | 955 | 1260 | 1185 | 1267 | 1514 | 14882 | 15337 | 13993 | 18842 | 14354 | 12398 | 12987 |
| 836 | 1057 | 1029 | 786 | 1054 | 1346 | 1059 | 1226 | 26015 | 19211 | 22976 | 24284 | 22080 | 16860 | 21449 |
| 503 | 700 | 1085 | 675 | 899 | 1004 | 622 | 870 | 16181 | 8925 | 17749 | 12421 | 17137 | 10652 | 15792 |
| 785 | 1281 | 881 | 688 | 1110 | 867 | 748 | 1143 | 4381 | 3913 | 4988 | 6390 | 11944 | 9334 | 13560 |
| 663 | 906 | 736 | 709 | 967 | 747 | 646 | 935 | 11443 | 10234 | 15437 | 13992 | 10282 | 9908 | 13966 |
| 743 | 896 | 627 | 717 | 880 | 765 | 765 | 913 | 8924 | 9684 | 13038 | 11679 | 14761 | 16891 | 23560 |
| 805 | 985 | 964 | 765 | 1014 | 1040 | 851 | 1065 | 9481 | 8011 | 9952 | 9799 | 10220 | 8113 | 10604 |
| 726 | 1100 | 874 | 740 | 1047 | 842 | 734 | 1103 | 16001 | 14664 | 20204 | 22215 | 18906 | 16000 | 21631 |
| 955 | 1606 | 938 | 1217 | 1919 | 910 | 1135 | 1856 | 7460 | 7989 | 8365 | 13432 | 6140 | 7972 | 6548 |
| 525 | 769 | 673 | 493 | 764 | 728 | 431 | 671 | 19715 | 14325 | 27304 | 20992 | 18692 | 13687 | 27789 |
| 658 | 994 | 941 | 768 | 989 | 717 | 550 | 852 | 22500 | 17611 | 26752 | 26580 | 7231 | 5904 | 7684 |
| 870 | 1042 | 900 | 1205 | 1580 | 911 | 1200 | 1487 | 6805 | 9169 | 10544 | 10986 | 13892 | 18596 | 15429 |
| 473 | 644 | 370 | 302 | 523 | 468 | 383 | 590 | 7652 | 6390 | 13516 | 8698 | 6008 | 4913 | 16245 |
| 283 | 579 | 371 | 338 | 582 | 410 | 343 | 595 | 4655 | 3725 | 13184 | 7635 | 9647 | 8808 | 26034 |
| 835 | 1281 | 928 | 758 | 1154 | 826 | 603 | 930 | 15974 | 13331 | 15965 | 20453 | 15214 | 12430 | 16402 |
| 441 | 756 | 880 | 592 | 921 | 822 | 552 | 846 | 8519 | 5461 | 12386 | 9366 | 33821 | 22748 | 38451 |
| 436 | 872 | 382 | 389 | 880 | 491 | 442 | 859 | 6698 | 7325 | 16807 | 14652 | 6985 | 7117 | 18300 |
| 757 | 1214 | 828 | 684 | 1141 | 942 | 795 | 1188 | 19499 | 15986 | 21111 | 25622 | 18687 | 15436 | 22555 |
| 637 | 989 | 804 | 674 | 1140 | 821 | 655 | 1075 | 8800 | 7228 | 11340 | 11212 | 11969 | 10039 | 14889 |

| | | | | | | | | | | | | | | |
|---|---|---|---|---|---|---|---|---|---|---|---|---|---|---|
| 510 | 889 | 561 | 507 | 892 | 526 | 564 | 914 | 8802 | 7999 | 15679 | 13940 | 15836 | 14317 | 28222 |
| 559 | 887 | 785 | 613 | 910 | 782 | 591 | 875 | 9648 | 7217 | 12903 | 11442 | 13122 | 10253 | 16713 |
| 546 | 752 | 525 | 493 | 801 | 674 | 574 | 805 | 13586 | 12094 | 22144 | 16660 | 4969 | 4674 | 9473 |
| 797 | 1286 | 902 | 849 | 1270 | 909 | 854 | 1290 | 2138 | 1911 | 2397 | 3083 | 4569 | 4302 | 5066 |
| 649 | 1030 | 1051 | 719 | 1060 | 989 | 640 | 1055 | 24199 | 15756 | 24276 | 25012 | 17453 | 11932 | 16601 |
| 539 | 794 | 792 | 458 | 675 | 918 | 621 | 939 | 5326 | 3620 | 6718 | 5333 | 8584 | 4960 | 10839 |
| 675 | 912 | 785 | 668 | 916 | 741 | 571 | 870 | 23879 | 19188 | 28426 | 25939 | 15932 | 13558 | 20287 |
| 592 | 1036 | 884 | 697 | 1067 | 791 | 586 | 1022 | 6951 | 5480 | 9257 | 9592 | 8718 | 6877 | 9862 |
| 984 | 1085 | 1043 | 787 | 1115 | 1566 | 1111 | 1206 | 19201 | 14155 | 14387 | 15606 | 13683 | 10321 | 13115 |
| 481 | 1062 | 568 | 416 | 937 | 727 | 606 | 1069 | 11307 | 10263 | 21337 | 22659 | 12839 | 9410 | 22617 |
| 585 | 953 | 912 | 691 | 1050 | 912 | 698 | 1070 | 14482 | 11276 | 19277 | 18372 | 12200 | 9253 | 13382 |
| 689 | 1049 | 765 | 531 | 860 | 642 | 447 | 711 | 3796 | 2763 | 4011 | 4208 | 10889 | 7556 | 14229 |
| 942 | 1595 | 861 | 822 | 1324 | 853 | 983 | 1691 | 3942 | 4286 | 4549 | 7254 | 22547 | 21521 | 26186 |
| 1042 | 1311 | 1039 | 937 | 1327 | 982 | 863 | 1221 | 20450 | 18242 | 17514 | 22955 | 16092 | 14515 | 15487 |
| 355 | 669 | 510 | 356 | 700 | 536 | 400 | 728 | 10238 | 7080 | 19953 | 13351 | 5299 | 3704 | 10391 |
| 918 | 1201 | 1091 | 872 | 1181 | 993 | 804 | 1211 | 8235 | 6982 | 7609 | 9137 | 11929 | 9539 | 10936 |
| 685 | 1058 | 974 | 777 | 987 | 986 | 681 | 981 | 13944 | 10651 | 15558 | 16453 | 4357 | 3477 | 4473 |
| 414 | 689 | 429 | 459 | 771 | 369 | 431 | 676 | 8445 | 9743 | 23505 | 16186 | 11763 | 12574 | 27407 |
| 661 | 956 | 800 | 651 | 955 | 861 | 711 | 998 | 9158 | 7386 | 11180 | 10686 | 12676 | 10322 | 15847 |
| 1190 | 1368 | 1341 | 1240 | 1267 | 1245 | 1156 | 1321 | 12765 | 11999 | 10085 | 13795 | 12914 | 11941 | 9628 |
| 442 | 729 | 934 | 650 | 1057 | 857 | 540 | 860 | 7834 | 4727 | 10683 | 7788 | 3689 | 2569 | 3950 |
| 935 | 1094 | 1074 | 889 | 1067 | 1164 | 1004 | 1149 | 11184 | 9598 | 10263 | 11226 | 26150 | 21643 | 24352 |
| 654 | 919 | 886 | 633 | 869 | 937 | 657 | 876 | 22559 | 15786 | 24152 | 22206 | 12522 | 8950 | 14132 |
| 500 | 882 | 687 | 519 | 890 | 616 | 474 | 890 | 20796 | 16416 | 32818 | 28943 | 7775 | 5866 | 11310 |
| 541 | 888 | 851 | 755 | 1037 | 793 | 701 | 1064 | 18621 | 14478 | 26748 | 23756 | 13446 | 11922 | 15800 |
| 465 | 799 | 637 | 427 | 769 | 695 | 513 | 804 | 14693 | 10683 | 22995 | 18367 | 8823 | 5911 | 13856 |
| 1347 | 1743 | 1159 | 1229 | 1651 | 1032 | 1020 | 1425 | 6404 | 6688 | 4963 | 8648 | 5376 | 5700 | 4637 |
| 633 | 873 | 927 | 799 | 1015 | 863 | 802 | 1019 | 7594 | 5776 | 9118 | 7958 | 3952 | 3407 | 4265 |
| 335 | 563 | 562 | 392 | 663 | 538 | 376 | 641 | 14590 | 8377 | 25003 | 14071 | 14621 | 10185 | 26002 |
| 746 | 1288 | 797 | 723 | 1252 | 775 | 674 | 1208 | 13147 | 11779 | 15796 | 20350 | 24348 | 22069 | 30536 |
| 1132 | 1298 | 951 | 717 | 996 | 1071 | 842 | 1195 | 7064 | 6025 | 5324 | 6910 | 11324 | 8537 | 11909 |

| | | | | | | | | | | | | | | |
|---|---|---|---|---|---|---|---|---|---|---|---|---|---|---|
| 585 | 1130 | 683 | 507 | 1031 | 796 | 657 | 1209 | 12614 | 9931 | 16969 | 19170 | 7648 | 5682 | 11204 |
| 924 | 1423 | 938 | 689 | 1172 | 1069 | 856 | 1482 | 23167 | 22523 | 24376 | 34686 | 16924 | 12439 | 18051 |
| 206 | 482 | 269 | 174 | 477 | 333 | 207 | 510 | 12075 | 7715 | 37540 | 18091 | 9309 | 6022 | 34548 |
| 1087 | 1551 | 1024 | 1050 | 1455 | 1279 | 1264 | 1664 | 23307 | 22292 | 20512 | 31815 | 7458 | 7650 | 7286 |
| 666 | 941 | 702 | 523 | 896 | 669 | 524 | 845 | 8051 | 6554 | 9834 | 9252 | 3041 | 2264 | 4330 |
| 607 | 818 | 904 | 739 | 864 | 913 | 797 | 836 | 13373 | 10775 | 17747 | 14520 | 13103 | 10722 | 14502 |
| 611 | 881 | 935 | 615 | 908 | 701 | 557 | 842 | 13567 | 11028 | 18041 | 15901 | 17318 | 11382 | 18515 |
| 269 | 534 | 648 | 304 | 582 | 654 | 270 | 530 | 8866 | 3849 | 14293 | 7629 | 11678 | 5468 | 18014 |
| 675 | 980 | 799 | 674 | 989 | 769 | 700 | 964 | 3424 | 2572 | 3808 | 3733 | 11691 | 9859 | 14635 |
| 569 | 837 | 639 | 438 | 657 | 600 | 444 | 765 | 6764 | 5344 | 9389 | 7854 | 3146 | 2158 | 4923 |
| 584 | 875 | 936 | 501 | 807 | 980 | 574 | 868 | 16081 | 9609 | 16448 | 14393 | 11453 | 6130 | 12235 |
| 603 | 1001 | 895 | 558 | 864 | 845 | 565 | 969 | 7095 | 4737 | 7852 | 7863 | 14463 | 9015 | 16162 |
| 732 | 1118 | 936 | 740 | 1114 | 891 | 702 | 1018 | 14422 | 11330 | 15468 | 17298 | 5828 | 4603 | 6223 |
| 749 | 1093 | 1091 | 874 | 1187 | 1348 | 1046 | 1246 | 15652 | 11932 | 15930 | 17416 | 7484 | 5999 | 6863 |
| 649 | 935 | 752 | 623 | 892 | 686 | 586 | 876 | 16186 | 13440 | 20701 | 19349 | 6421 | 5317 | 8541 |
| 698 | 1015 | 725 | 582 | 919 | 722 | 534 | 802 | 12792 | 10676 | 15304 | 15539 | 10270 | 8250 | 14170 |
| 375 | 635 | 670 | 511 | 775 | 645 | 490 | 745 | 1834 | 1262 | 3366 | 2138 | 10484 | 7986 | 15641 |
| 445 | 836 | 678 | 429 | 799 | 611 | 441 | 871 | 13993 | 9695 | 21785 | 18214 | 18987 | 12019 | 28010 |
| 417 | 707 | 347 | 330 | 605 | 452 | 361 | 565 | 7962 | 7208 | 17297 | 12224 | 7015 | 6679 | 20230 |
| 339 | 618 | 416 | 376 | 636 | 569 | 474 | 671 | 6453 | 5180 | 15277 | 9444 | 3329 | 3014 | 8008 |
| 965 | 1124 | 1007 | 797 | 1041 | 1150 | 952 | 1143 | 13373 | 11119 | 11524 | 12951 | 4761 | 3768 | 4727 |
| 468 | 844 | 779 | 646 | 1013 | 760 | 584 | 973 | 10019 | 7499 | 16035 | 13533 | 15830 | 13135 | 20325 |
| 341 | 716 | 500 | 346 | 704 | 681 | 476 | 779 | 19940 | 12443 | 36473 | 26122 | 4305 | 2978 | 8606 |
| 802 | 1148 | 846 | 729 | 1106 | 832 | 678 | 1064 | 7245 | 6456 | 8050 | 9241 | 9861 | 8498 | 11650 |
| 575 | 1069 | 759 | 586 | 996 | 794 | 529 | 955 | 15434 | 11741 | 20412 | 21825 | 5111 | 3945 | 6732 |
| 530 | 739 | 1112 | 538 | 701 | 1063 | 534 | 753 | 7801 | 3964 | 7475 | 5522 | 17746 | 8579 | 15952 |
| 836 | 1163 | 891 | 786 | 1121 | 838 | 721 | 1147 | 5949 | 5062 | 6057 | 7042 | 11886 | 10486 | 13345 |
| 644 | 1010 | 770 | 617 | 980 | 926 | 761 | 1137 | 8595 | 7186 | 11154 | 11269 | 8837 | 7080 | 11480 |
| 471 | 777 | 754 | 602 | 858 | 573 | 429 | 766 | 6861 | 4779 | 10141 | 7874 | 14966 | 11944 | 19847 |
| 759 | 1064 | 902 | 678 | 1043 | 845 | 650 | 982 | 11413 | 9330 | 12296 | 13085 | 9463 | 7116 | 10493 |
| 652 | 919 | 938 | 707 | 931 | 908 | 746 | 951 | 2488 | 2006 | 3075 | 2825 | 5846 | 4403 | 6232 |

| | | | | | | | | | | | | | | |
|---|---|---|---|---|---|---|---|---|---|---|---|---|---|---|
| 660 | 1076 | 977 | 774 | 1221 | 972 | 710 | 1144 | 11819 | 8676 | 13152 | 14148 | 4072 | 3227 | 4169 |
| 620 | 953 | 870 | 622 | 944 | 813 | 639 | 954 | 10870 | 7549 | 12175 | 11597 | 13329 | 9520 | 15313 |
| 486 | 804 | 763 | 487 | 821 | 792 | 363 | 665 | 16743 | 10883 | 22392 | 17993 | 16747 | 10698 | 21954 |
| 615 | 979 | 776 | 717 | 1139 | 988 | 854 | 1248 | 5113 | 4198 | 6823 | 6678 | 9862 | 9116 | 12715 |
| 434 | 779 | 635 | 405 | 736 | 572 | 347 | 679 | 7012 | 4828 | 11125 | 8668 | 14504 | 9256 | 22858 |
| 699 | 1015 | 983 | 786 | 999 | 876 | 647 | 940 | 6396 | 5094 | 7286 | 7396 | 12536 | 10028 | 12758 |
| 419 | 690 | 798 | 483 | 778 | 740 | 352 | 596 | 10804 | 5704 | 13621 | 9401 | 15305 | 9270 | 19174 |
| 442 | 633 | 1308 | 577 | 776 | 1134 | 418 | 616 | 11092 | 4805 | 10874 | 6887 | 9424 | 4159 | 7203 |
| 680 | 933 | 957 | 627 | 951 | 936 | 616 | 953 | 8078 | 5310 | 7810 | 7286 | 10020 | 6570 | 10472 |
| 1074 | 1083 | 1214 | 977 | 983 | 1112 | 879 | 1021 | 9300 | 7877 | 7333 | 7942 | 21321 | 17160 | 17565 |
| 289 | 802 | 502 | 392 | 884 | 588 | 399 | 865 | 6027 | 4311 | 14922 | 11962 | 10223 | 7983 | 20381 |
| 1371 | 1588 | | | | | | | 1361 | 1471 | 1073 | 1704 | | | |
| 644 | 821 | 626 | 474 | 784 | 549 | 434 | 740 | 11613 | 9456 | 14681 | 12057 | 4672 | 3540 | 7462 |
| 295 | 702 | 494 | 376 | 711 | 515 | 286 | 635 | 23018 | 13564 | 45987 | 32290 | 10870 | 8282 | 22023 |
| 862 | 1312 | 857 | 772 | 1453 | 849 | 742 | 1285 | 10994 | 10140 | 11765 | 15440 | 5649 | 5086 | 6589 |
| 173 | 504 | 337 | 193 | 513 | 382 | 268 | 569 | 11365 | 6384 | 36853 | 18580 | 10872 | 6229 | 32270 |
| 351 | 584 | 372 | 316 | 566 | 357 | 340 | 562 | 5794 | 5690 | 16217 | 9468 | 5262 | 4462 | 14132 |
| 581 | 901 | 699 | 602 | 945 | 786 | 611 | 983 | 6972 | 5479 | 9428 | 8499 | 7335 | 6312 | 10489 |
| 611 | 857 | 1118 | 797 | 1007 | 1151 | 818 | 1014 | 8529 | 5905 | 9660 | 8279 | 15831 | 11283 | 14159 |
| 624 | 1125 | 941 | 627 | 1135 | 832 | 735 | 1217 | 12679 | 11063 | 17728 | 19944 | 7891 | 5257 | 8388 |
| 763 | 1060 | 972 | 777 | 1111 | 990 | 683 | 942 | 8615 | 6612 | 8664 | 9183 | 8275 | 6619 | 8514 |
| 583 | 804 | 575 | 531 | 764 | 625 | 735 | 856 | 9474 | 9624 | 16505 | 13267 | 8491 | 7841 | 14759 |
| 1156 | 1376 | 970 | 1186 | 1291 | 873 | 1073 | 1323 | 11602 | 13584 | 11747 | 16162 | 18028 | 22056 | 18593 |
| 599 | 953 | 891 | 632 | 1057 | 646 | 496 | 857 | 15270 | 10902 | 18200 | 17341 | 15069 | 10691 | 16911 |
| 808 | 1283 | 852 | 546 | 895 | 843 | 624 | 1065 | 14762 | 12498 | 15472 | 19843 | 11013 | 7055 | 12929 |
| 601 | 984 | 707 | 554 | 905 | 561 | 468 | 952 | 14355 | 12259 | 20389 | 20057 | 7175 | 5619 | 10148 |
| 911 | 918 | 1315 | 922 | 870 | 1243 | 942 | 800 | 13594 | 10004 | 10980 | 10078 | 14510 | 10170 | 11034 |
| 641 | 1222 | 895 | 776 | 1507 | 913 | 814 | 1505 | 11807 | 9018 | 14058 | 17176 | 12922 | 11201 | 14438 |
| 528 | 1065 | 802 | 617 | 1225 | 921 | 667 | 1197 | 1853 | 1371 | 2599 | 2767 | 7910 | 6081 | 9864 |
| 632 | 1114 | 1033 | 892 | 1329 | 1150 | 933 | 1086 | 17014 | 14234 | 22520 | 25094 | 7317 | 6318 | 7086 |
| 531 | 777 | 838 | 817 | 804 | 862 | 831 | 889 | 10186 | 9143 | 17212 | 13378 | 9051 | 8832 | 10807 |

| | | | | | | | | | | | | | | |
|---|---|---|---|---|---|---|---|---|---|---|---|---|---|---|
| 1007 | 1352 | 1000 | 885 | 1194 | 1019 | 919 | 1310 | 23560 | 21731 | 21579 | 29174 | 13232 | 11708 | 13226 |
| 629 | 1223 | 834 | 694 | 1240 | 1046 | 817 | 1315 | 5134 | 4376 | 6953 | 8501 | 8403 | 6996 | 10080 |
| 700 | 1050 | 1127 | 789 | 1097 | 871 | 605 | 990 | 5903 | 4253 | 6073 | 6377 | 1441 | 1009 | 1279 |
| 721 | 1056 | 887 | 715 | 1053 | 745 | 615 | 954 | 14212 | 12237 | 16979 | 17929 | 6152 | 4956 | 6934 |
| 772 | 1026 | 771 | 745 | 1001 | 957 | 773 | 991 | 9672 | 8457 | 10960 | 11243 | 9134 | 8825 | 11852 |
| 399 | 593 | 880 | 451 | 675 | 850 | 464 | 720 | 10974 | 5439 | 13636 | 8080 | 15138 | 7750 | 17193 |
| 438 | 795 | 792 | 486 | 837 | 1059 | 659 | 933 | 13286 | 7885 | 18001 | 14309 | 9550 | 5858 | 12056 |
| 525 | 673 | 548 | 362 | 537 | 583 | 398 | 585 | 13411 | 9800 | 18682 | 12567 | 14907 | 9860 | 27209 |
| 505 | 807 | 1003 | 646 | 919 | 1004 | 553 | 877 | 4484 | 2580 | 5112 | 4124 | 14883 | 9580 | 14842 |
| 1045 | 1519 | | | | | | | 564 | 564 | 540 | 820 | | | |
| 979 | 1325 | 870 | 915 | 1167 | 1121 | 882 | 1442 | 20926 | 21164 | 21622 | 28643 | 16493 | 17330 | 18950 |
| 967 | 1341 | 920 | 845 | 1164 | 973 | 914 | 1233 | 8781 | 8573 | 8863 | 11889 | 4734 | 4347 | 5143 |
| 859 | 1191 | 1108 | 856 | 1219 | 1093 | 844 | 1041 | 14579 | 12187 | 14193 | 16907 | 9753 | 7533 | 8799 |
| 887 | 525 | 651 | 946 | 464 | 665 | 886 | 442 | 10212 | 12771 | 14395 | 7555 | 11102 | 16138 | 17060 |
| 769 | 1158 | 963 | 782 | 1139 | 906 | 781 | 1052 | 16463 | 13083 | 17010 | 19700 | 9939 | 8065 | 10316 |
| 605 | 2442 | 859 | 664 | 2858 | 906 | 585 | 3536 | 7827 | 5135 | 8484 | 20714 | 6563 | 5072 | 7636 |
| 681 | 1036 | 768 | 602 | 977 | 822 | 648 | 1051 | 6110 | 4753 | 6982 | 7234 | 14546 | 11396 | 18936 |
| 876 | 1280 | 991 | 863 | 1324 | 1167 | 1014 | 1339 | 14803 | 12483 | 14247 | 18239 | 7550 | 6576 | 7622 |
| 947 | 1531 | 864 | 1034 | 1637 | 925 | 1010 | 1576 | 8428 | 8603 | 9084 | 13911 | 8662 | 10368 | 10023 |
| 499 | 899 | 736 | 483 | 735 | 623 | 513 | 795 | 8243 | 6382 | 12779 | 11492 | 14130 | 9266 | 19202 |
| 604 | 1043 | 927 | 694 | 997 | 947 | 742 | 994 | 11176 | 7651 | 12664 | 13208 | 10041 | 7514 | 10833 |
| 619 | 951 | 826 | 672 | 1027 | 597 | 549 | 1026 | 10962 | 7033 | 11357 | 10803 | 14892 | 12110 | 18031 |
| 709 | 1280 | 827 | 648 | 1218 | 879 | 740 | 1182 | 10465 | 8940 | 12602 | 16126 | 3422 | 2684 | 4139 |
| 593 | 722 | 1030 | 623 | 839 | 897 | 562 | 808 | 12389 | 6815 | 11489 | 8294 | 9463 | 5725 | 9187 |
| | | 766 | 628 | 932 | 682 | 509 | 925 | | | | | 3206 | 2626 | 4184 |
| 755 | 1229 | 870 | 563 | 1072 | 896 | 1048 | 1524 | 24658 | 21010 | 27814 | 34184 | 25562 | 16521 | 29367 |
| 476 | 781 | 746 | 489 | 806 | 807 | 590 | 852 | 4473 | 3224 | 6771 | 5289 | 12195 | 8003 | 16350 |
| 451 | 846 | 698 | 471 | 843 | 677 | 503 | 959 | 12237 | 8018 | 17763 | 15028 | 11112 | 7501 | 15928 |
| 680 | 1033 | 811 | 678 | 1022 | 920 | 735 | 1063 | 6592 | 5149 | 7574 | 7825 | 14123 | 11807 | 17422 |
| 665 | 1080 | 568 | 455 | 908 | 680 | 633 | 1112 | 14627 | 12628 | 18977 | 20488 | 4211 | 3370 | 7411 |
| 1333 | 875 | 986 | 921 | 847 | 849 | 931 | 831 | 6217 | 7230 | 5423 | 4743 | 13951 | 13022 | 14142 |

| | | | | | | | | | | | | | | |
|---|---|---|---|---|---|---|---|---|---|---|---|---|---|---|
| 493 | 769 | 670 | 417 | 684 | 641 | 407 | 685 | 2391 | 1721 | 3489 | 2683 | 13899 | 8634 | 20731 |
| 504 | 886 | 823 | 591 | 1038 | 839 | 608 | 1032 | 10457 | 7290 | 14456 | 12814 | 10693 | 7678 | 12993 |
| 856 | 1202 | 1071 | 919 | 1232 | 1039 | 930 | 1286 | 12386 | 11345 | 13247 | 15919 | 9587 | 8227 | 8948 |
| 755 | 1314 | 963 | 784 | 1323 | 930 | 801 | 1297 | 10636 | 8565 | 11347 | 14910 | 12131 | 9867 | 12591 |
| 328 | 725 | 714 | 385 | 728 | 598 | 442 | 821 | 6582 | 4438 | 13532 | 9815 | 18334 | 9879 | 25682 |
| 549 | 1026 | 923 | 716 | 1057 | 765 | 567 | 1024 | 11554 | 8661 | 15787 | 16203 | 6965 | 5406 | 7550 |
| 599 | 895 | 793 | 647 | 930 | 694 | 582 | 903 | 9997 | 8953 | 14948 | 13382 | 10251 | 8371 | 12933 |
| 297 | 581 | 610 | 364 | 664 | 590 | 421 | 710 | 3957 | 2109 | 7095 | 4123 | 3691 | 2201 | 6054 |
| 723 | 1011 | 1078 | 791 | 1095 | 1075 | 768 | 1100 | 11319 | 7686 | 10624 | 10744 | 16560 | 12147 | 15355 |
| 488 | 760 | 599 | 485 | 723 | 586 | 448 | 704 | 6388 | 4749 | 9737 | 7399 | 4562 | 3692 | 7619 |
| 711 | 1153 | 770 | 698 | 1105 | 829 | 748 | 1098 | 8690 | 7560 | 10627 | 12249 | 8165 | 7401 | 10601 |
| 885 | 1413 | 858 | 940 | 1322 | 973 | 1133 | 1629 | 8672 | 10256 | 11585 | 16372 | 8340 | 9142 | 9725 |
| 763 | 967 | 841 | 569 | 805 | 816 | 600 | 911 | 16772 | 12372 | 16206 | 15667 | 19725 | 13359 | 23465 |
| 736 | 1062 | 1014 | 731 | 1072 | 892 | 652 | 935 | 11002 | 8795 | 11950 | 12697 | 12019 | 8666 | 11848 |
| 681 | 925 | 1171 | 738 | 961 | 1117 | 661 | 907 | 18176 | 11180 | 16409 | 15172 | 8731 | 5503 | 7454 |
| 867 | 1126 | 905 | 1016 | 1345 | 841 | 810 | 1221 | 3451 | 3078 | 3551 | 3999 | 2607 | 2928 | 2882 |
| 636 | 1071 | 846 | 847 | 1257 | 928 | 1032 | 1440 | 8852 | 7574 | 11905 | 12751 | 13051 | 13068 | 15432 |
| 829 | 1242 | 984 | 842 | 1251 | 991 | 859 | 1230 | 7994 | 6744 | 8131 | 10095 | 8094 | 6928 | 8225 |
| 556 | 840 | 779 | 646 | 939 | 573 | 510 | 864 | 13132 | 11800 | 21226 | 17834 | 3842 | 3184 | 4929 |
| 1280 | 1411 | 1202 | 1170 | 1242 | 1041 | 1608 | 1751 | 5913 | 6524 | 5096 | 7190 | 6641 | 6468 | 5526 |
| 455 | 890 | 670 | 470 | 938 | 707 | 559 | 953 | 4340 | 3125 | 6862 | 6108 | 8686 | 6094 | 12972 |
| 821 | 1130 | 966 | 830 | 1175 | 909 | 771 | 1172 | 21324 | 15967 | 19438 | 21959 | 12913 | 11094 | 13363 |
| 1120 | 1420 | 1101 | 1079 | 1195 | 954 | 882 | 1203 | 10716 | 11048 | 9868 | 14011 | 3743 | 3667 | 3399 |
| 837 | 1226 | 864 | 692 | 1065 | 974 | 804 | 1128 | 12726 | 10307 | 12319 | 15105 | 4237 | 3396 | 4904 |
| 672 | 831 | 1262 | 663 | 761 | 1205 | 662 | 770 | 18060 | 9901 | 14733 | 12243 | 9064 | 4763 | 7183 |
| 610 | 1105 | 577 | 462 | 930 | 757 | 610 | 1100 | 5164 | 4287 | 7025 | 7761 | 6502 | 5205 | 11267 |
| 707 | 1187 | 869 | 735 | 1205 | 878 | 755 | 1212 | 17772 | 15295 | 21622 | 25668 | 16966 | 14343 | 19518 |
| | | 1217 | 896 | 1045 | | | | | | | | 4458 | 3283 | 3664 |
| 454 | 793 | | | | 535 | 437 | 874 | 586 | 414 | 911 | 723 | | | |
| 728 | 1043 | 814 | 536 | 796 | 820 | 544 | 808 | 12368 | 8634 | 11856 | 12364 | 13906 | 9159 | 17078 |
| 647 | 949 | 612 | 474 | 818 | 662 | 476 | 852 | 16790 | 14571 | 22522 | 21366 | 14284 | 11066 | 23339 |

| | | | | | | | | | | | | | | |
|---|---|---|---|---|---|---|---|---|---|---|---|---|---|---|
| 1736 | 524 | 723 | 2150 | 564 | 894 | 2188 | 674 | 4240 | 11736 | 6762 | 3541 | 3454 | 10268 | 4776 |
| 492 | 816 | 633 | 395 | 782 | 472 | 263 | 680 | 4602 | 3060 | 6218 | 5075 | 12652 | 7889 | 19982 |
| 340 | 640 | 468 | 259 | 533 | 467 | 275 | 561 | 7136 | 5030 | 14804 | 9476 | 18133 | 10056 | 38759 |
| 793 | 1315 | 1080 | 709 | 1143 | 1034 | 838 | 1350 | 4808 | 3841 | 4844 | 6371 | 14566 | 9563 | 13482 |
| 378 | 638 | 464 | 332 | 544 | 443 | 359 | 581 | 13889 | 10607 | 28087 | 17908 | 7890 | 5653 | 17007 |
| 419 | 714 | 650 | 428 | 705 | 667 | 460 | 732 | 4546 | 3078 | 7354 | 5254 | 13844 | 9111 | 21291 |
| 536 | 824 | 891 | 635 | 945 | 808 | 528 | 820 | 15100 | 9161 | 17082 | 14079 | 7253 | 5167 | 8140 |
| 666 | 954 | 899 | 729 | 950 | 691 | 531 | 836 | 13286 | 10818 | 16232 | 15494 | 13242 | 10739 | 14738 |
| 418 | 636 | 689 | 384 | 625 | 549 | 323 | 642 | 15152 | 9201 | 21987 | 13976 | 10075 | 5612 | 14628 |
| 752 | 1006 | 864 | 687 | 924 | 927 | 800 | 1022 | 12360 | 10323 | 13721 | 13797 | 5931 | 4718 | 6866 |
| 597 | 917 | 913 | 737 | 970 | 967 | 607 | 931 | 2961 | 2063 | 3458 | 3170 | 8511 | 6870 | 9317 |
| 759 | 979 | 711 | 528 | 1013 | 551 | 425 | 854 | 4521 | 3520 | 4639 | 4542 | 1035 | 768 | 1455 |
| 634 | 1075 | 773 | 590 | 1064 | 827 | 641 | 1056 | 12005 | 9466 | 14942 | 16056 | 11763 | 8980 | 15215 |
| 543 | 802 | 833 | 659 | 956 | 847 | 538 | 795 | 5348 | 3819 | 7033 | 5640 | 5824 | 4610 | 6992 |
| 487 | 964 | 714 | 519 | 1002 | 681 | 640 | 1140 | 8741 | 5898 | 12101 | 11665 | 4267 | 3103 | 5975 |
| 280 | 452 | 1271 | 388 | 529 | 1289 | 393 | 568 | 25122 | 5061 | 18079 | 8177 | 12671 | 3866 | 9968 |
| 473 | 752 | 691 | 534 | 772 | 870 | 610 | 835 | 7515 | 5359 | 11332 | 8526 | 12815 | 9903 | 18535 |
| 925 | 1327 | 1073 | 1065 | 1413 | 748 | 1076 | 1020 | 9755 | 8853 | 9575 | 12710 | 4745 | 4710 | 4422 |
| 435 | 692 | 813 | 375 | 618 | 675 | 361 | 580 | 13124 | 7247 | 16649 | 11521 | 11016 | 5079 | 13555 |
| 903 | 7978 | 902 | 643 | 6950 | 1230 | 930 | 9639 | 7421 | 5650 | 6256 | 49909 | 3356 | 2391 | 3719 |
| 1847 | 3231 | 1075 | 1751 | 3163 | 1079 | 1656 | 2929 | 10056 | 17763 | 9618 | 31079 | 14708 | 23966 | 13687 |
| 503 | 755 | 786 | 379 | 651 | 731 | 449 | 723 | 17802 | 10604 | 21079 | 15906 | 13247 | 6378 | 16851 |
| 687 | 1012 | 1019 | 714 | 1009 | 959 | 661 | 1007 | 8838 | 6128 | 8916 | 9025 | 15183 | 10638 | 14896 |
| 796 | 1181 | 995 | 876 | 1233 | 1027 | 928 | 1216 | 21394 | 16831 | 21153 | 24980 | 4452 | 3921 | 4474 |
| 284 | 666 | 450 | 263 | 636 | 333 | 200 | 532 | 11417 | 6411 | 22570 | 15031 | 11519 | 6727 | 25608 |
| 822 | 1055 | 1007 | 888 | 1095 | 848 | 680 | 969 | 11714 | 9909 | 12048 | 12706 | 11725 | 10330 | 11638 |
| 567 | 740 | 700 | 452 | 712 | 711 | 545 | 768 | 3472 | 2602 | 4587 | 3394 | 1077 | 695 | 1538 |
| 707 | 956 | 918 | 746 | 973 | 800 | 635 | 935 | 16795 | 12945 | 18311 | 17501 | 14416 | 11718 | 15710 |
| 474 | 1024 | 526 | 401 | 979 | 647 | 505 | 1075 | 9502 | 7074 | 14911 | 15269 | 5058 | 3852 | 9608 |
| 688 | 985 | 964 | 607 | 825 | 978 | 621 | 951 | 11920 | 8069 | 11723 | 11544 | 10194 | 6426 | 10578 |
| 858 | 1398 | 910 | 798 | 1335 | 1080 | 939 | 1238 | 2007 | 1890 | 2204 | 3080 | 5175 | 4542 | 5690 |

| | | | | | | | | | | | | | | |
|---|---|---|---|---|---|---|---|---|---|---|---|---|---|---|
| 803 | 1205 | 917 | 779 | 1174 | 783 | 689 | 1092 | 13819 | 12316 | 15342 | 18494 | 19867 | 16881 | 21670 |
| 567 | 836 | 753 | 559 | 842 | 776 | 596 | 924 | 9270 | 6800 | 11997 | 10023 | 4793 | 3556 | 6367 |
| 565 | 892 | 692 | 501 | 910 | 662 | 514 | 937 | 4371 | 3161 | 5594 | 4989 | 10557 | 7649 | 15253 |
| 445 | 743 | 882 | 526 | 846 | 828 | 450 | 763 | 3591 | 2161 | 4862 | 3612 | 4866 | 2903 | 5515 |
| 1290 | 1445 | 1053 | 953 | 1449 | 988 | 963 | 1341 | 9704 | 10050 | 7791 | 11256 | 8967 | 8119 | 8518 |
| 527 | 894 | 629 | 539 | 903 | 603 | 491 | 799 | 15897 | 14034 | 26611 | 23793 | 8828 | 7567 | 14027 |
| 1029 | 1647 | 999 | 983 | 1848 | 1223 | 1286 | 1989 | 8309 | 8373 | 8133 | 13395 | 8119 | 7989 | 8128 |
| 337 | 589 | 605 | 320 | 570 | 590 | 313 | 570 | 12703 | 6064 | 17988 | 10603 | 17203 | 9079 | 28415 |
| 771 | 1026 | 938 | 738 | 1091 | 1000 | 831 | 1048 | 9800 | 8283 | 10738 | 11014 | 10383 | 8171 | 11064 |
| 534 | 960 | 649 | 484 | 919 | 604 | 407 | 860 | 16180 | 11898 | 22287 | 21396 | 9135 | 6817 | 14079 |
| 618 | 1093 | 415 | 541 | 921 | 419 | 530 | 919 | 831 | 1205 | 1950 | 2131 | 6556 | 8538 | 15791 |
| 347 | 793 | 723 | 386 | 689 | 798 | 487 | 849 | 10546 | 5653 | 16294 | 12921 | 12710 | 6784 | 17586 |
| 860 | 1060 | 962 | 753 | 1115 | 966 | 748 | 1217 | 10740 | 8064 | 9379 | 9940 | 19300 | 15099 | 20057 |
| 509 | 969 | 837 | 560 | 950 | 790 | 586 | 972 | 9906 | 7629 | 14976 | 14509 | 7753 | 5182 | 9259 |
| 533 | 953 | 939 | 661 | 1054 | 997 | 696 | 1088 | 7677 | 5143 | 9653 | 9203 | 9948 | 7003 | 10598 |
| 512 | 755 | 520 | 410 | 620 | 616 | 468 | 647 | 16516 | 12528 | 24478 | 18472 | 5518 | 4347 | 10613 |
| 585 | 915 | 878 | 645 | 927 | 869 | 602 | 1003 | 2039 | 1335 | 2281 | 2086 | 6906 | 5075 | 7864 |
| 823 | 1307 | 865 | 700 | 1139 | 807 | 643 | 1093 | 3609 | 3419 | 4153 | 5427 | 5192 | 4199 | 6003 |
| 312 | 579 | 457 | 280 | 594 | 430 | 272 | 603 | 4994 | 3064 | 9809 | 5677 | 10840 | 6630 | 23720 |
| 527 | 713 | 606 | 522 | 673 | 742 | 683 | 998 | 9041 | 7054 | 13381 | 9540 | 5174 | 4459 | 8535 |
| 893 | 1304 | 942 | 1071 | 1516 | 925 | 1008 | 1385 | 11747 | 12130 | 13584 | 17708 | 10483 | 11919 | 11133 |
| 599 | 1005 | 944 | 688 | 965 | 873 | 565 | 870 | 22144 | 16470 | 27497 | 27642 | 28487 | 20764 | 30161 |
| 768 | 1200 | 1017 | 781 | 1275 | 1041 | 818 | 1179 | 7481 | 5784 | 7536 | 9042 | 23695 | 18201 | 23292 |
| 506 | 930 | 761 | 539 | 903 | 744 | 516 | 933 | 5406 | 3952 | 7812 | 7263 | 16304 | 11543 | 21422 |
| 1118 | 1464 | 1036 | 1053 | 1407 | 913 | 1033 | 1362 | 11773 | 13189 | 11799 | 17277 | 7289 | 7408 | 7035 |
| 464 | 735 | 829 | 549 | 840 | 756 | 493 | 753 | 4358 | 2756 | 5937 | 4367 | 6499 | 4303 | 7841 |
| | | 940 | 422 | 696 | 921 | 446 | 738 | | | | | 3796 | 1702 | 4036 |
| 698 | 1060 | 840 | 607 | 1092 | 836 | 686 | 1168 | 13936 | 11311 | 16206 | 17174 | 8428 | 6084 | 10030 |
| 342 | 637 | 571 | 393 | 662 | 745 | 517 | 834 | 3619 | 2037 | 5949 | 3788 | 9654 | 6641 | 16894 |
| 478 | 515 | 935 | 556 | 994 | 1207 | 602 | 804 | 16055 | 7000 | 14639 | 7536 | 15945 | 9481 | 17053 |
| 624 | 956 | 838 | 596 | 988 | | | | 7155 | 4894 | 7842 | 7496 | 5332 | 3793 | 6363 |

| | | | | | | | | | | | | | | |
|---|---|---|---|---|---|---|---|---|---|---|---|---|---|---|
| 541 | 985 | 853 | 477 | 831 | 812 | 353 | 677 | 15620 | 10518 | 19439 | 19153 | 12722 | 7103 | 14906 |
| 500 | 733 | 644 | 529 | 850 | 651 | 463 | 773 | 9896 | 6813 | 13615 | 9976 | 8973 | 7375 | 13940 |
| 568 | 951 | 713 | 548 | 890 | 744 | 628 | 983 | 19225 | 14765 | 25994 | 24712 | 10139 | 7782 | 14212 |
| 427 | 856 | 654 | 513 | 896 | 652 | 511 | 896 | 5631 | 4190 | 9817 | 8401 | 7687 | 6032 | 11751 |
| 990 | 447 | 596 | 950 | 443 | 749 | 1204 | 542 | 7458 | 9820 | 9924 | 4434 | 3450 | 5495 | 5787 |
| 688 | 1065 | 696 | 581 | 976 | 903 | 695 | 1014 | 18638 | 16228 | 23580 | 25123 | 4646 | 3875 | 6673 |
| 492 | 669 | | | | 836 | 602 | 907 | 9427 | 6236 | 12683 | 8489 | | | |
| 651 | 966 | 952 | 725 | 1046 | 947 | 669 | 973 | 18568 | 13076 | 20079 | 19402 | 12833 | 9767 | 13481 |
| 1058 | 1236 | 1123 | 886 | 1170 | 1104 | 819 | 1171 | 3731 | 3278 | 3098 | 3830 | 7955 | 6281 | 7086 |
| 403 | 684 | 480 | 323 | 640 | 455 | 184 | 516 | 5017 | 3403 | 8450 | 5783 | 10552 | 7102 | 21970 |
| 506 | 1019 | 750 | 490 | 1014 | 658 | 573 | 1185 | 15058 | 11539 | 22808 | 23239 | 12597 | 8232 | 16793 |
| 783 | 995 | 903 | 816 | 1060 | 884 | 733 | 923 | 3823 | 3315 | 4231 | 4211 | 6940 | 6276 | 7688 |
| 824 | 963 | 733 | 504 | 818 | 800 | 541 | 814 | 6498 | 5852 | 7102 | 6840 | 6555 | 4508 | 8939 |
| 695 | 1025 | 948 | 786 | 1507 | 1014 | 706 | 1152 | 10563 | 6501 | 9353 | 9584 | 14320 | 11876 | 15103 |
| 751 | 953 | 1075 | 811 | 1008 | 990 | 784 | 991 | 16315 | 12404 | 16517 | 15734 | 4132 | 3117 | 3845 |
| 663 | 1106 | 857 | 664 | 1145 | 881 | 655 | 1085 | 8311 | 6819 | 10291 | 11381 | 8964 | 6948 | 10459 |
| 738 | 1090 | 777 | 725 | 1014 | 736 | 618 | 883 | 8202 | 6769 | 9166 | 9990 | 7821 | 7296 | 10064 |
| 474 | 891 | 983 | 540 | 925 | 980 | 572 | 925 | 26223 | 15237 | 32169 | 28653 | 17660 | 9694 | 17965 |
| 828 | 1071 | 1054 | 724 | 947 | 1069 | 778 | 1170 | 5547 | 3925 | 4741 | 5076 | 6072 | 4170 | 5762 |
| 594 | 1004 | 681 | 527 | 968 | 887 | 649 | 1221 | 11945 | 9027 | 15202 | 15265 | 4220 | 3268 | 6199 |
| 261 | 558 | 657 | 211 | 456 | 593 | 334 | 638 | 9800 | 5083 | 19508 | 10892 | 16862 | 5415 | 25649 |
| 254 | 466 | 459 | 255 | 428 | 486 | 371 | 601 | 3886 | 2398 | 9436 | 4396 | 18946 | 10530 | 41298 |
| 877 | 1077 | 771 | 452 | 673 | 617 | 363 | 710 | 10881 | 8057 | 9192 | 9904 | 8774 | 5146 | 11387 |
| 641 | 1183 | 918 | 704 | 1089 | 955 | 776 | 1155 | 6686 | 5016 | 7831 | 9261 | 16591 | 12730 | 18073 |
| 450 | 984 | 676 | 481 | 957 | 746 | 646 | 1158 | 2656 | 1894 | 4204 | 4136 | 24349 | 17335 | 36040 |
| 669 | 896 | 760 | 613 | 909 | 728 | 567 | 811 | 6322 | 5037 | 7530 | 6749 | 10809 | 8721 | 14220 |
| 468 | 761 | 720 | 509 | 759 | 746 | 476 | 699 | 6835 | 4959 | 10595 | 8061 | 4273 | 3019 | 5937 |
| 624 | 996 | 944 | 703 | 1091 | 909 | 642 | 995 | 11564 | 7901 | 12657 | 12602 | 12424 | 9259 | 13162 |
| 528 | 902 | 719 | 475 | 893 | 695 | 514 | 925 | 18567 | 13645 | 25833 | 23294 | 14989 | 9907 | 20849 |
| 542 | 849 | 905 | 673 | 900 | 860 | 603 | 899 | 5936 | 3862 | 7127 | 6052 | 7993 | 5944 | 8834 |
| 341 | 506 | 465 | 313 | 488 | 592 | 405 | 594 | 16839 | 11146 | 32661 | 16516 | 10481 | 7054 | 22536 |

| | | | | | | | | | | | | | | |
|---|---|---|---|---|---|---|---|---|---|---|---|---|---|---|
| 773 | 1203 | 827 | 655 | 1045 | 953 | 829 | 1314 | 3699 | 3095 | 4005 | 4818 | 2423 | 1918 | 2930 |
| 684 | 1543 | 1193 | 1127 | 1370 | 847 | 738 | 1508 | 4854 | 4144 | 6057 | 9343 | 4230 | 3998 | 3547 |
| 396 | 728 | 580 | 479 | 784 | 370 | 320 | 697 | 7416 | 5418 | 13691 | 9972 | 6178 | 5100 | 10654 |
| 758 | 1120 | 687 | 663 | 1218 | 665 | 588 | 1053 | 8645 | 6803 | 8975 | 10053 | 12506 | 12063 | 18204 |
| 160 | 672 | 450 | 234 | 733 | 459 | 257 | 727 | 3368 | 1398 | 8721 | 5864 | 13357 | 6937 | 29680 |
| 854 | 1704 | 1034 | 757 | 1213 | 1042 | 894 | 1578 | 20628 | 17312 | 20271 | 34545 | 16262 | 11908 | 15722 |
| 1051 | 1389 | 1231 | 1098 | 1417 | 921 | 759 | 1152 | 6812 | 5871 | 5589 | 7760 | 8142 | 7262 | 6616 |
| 583 | 916 | 830 | 633 | 1041 | 879 | 779 | 1005 | 5422 | 4005 | 6870 | 6293 | 7279 | 5553 | 8774 |
| 684 | 1200 | 889 | 731 | 1172 | 1033 | 857 | 1311 | 14183 | 10946 | 16013 | 19216 | 11402 | 9371 | 12821 |
| 530 | 762 | 486 | 478 | 653 | 484 | 425 | 626 | 5681 | 4079 | 7699 | 5867 | 6175 | 6080 | 12715 |
| 753 | 1089 | 746 | 487 | 977 | 947 | 722 | 1066 | 8433 | 6418 | 8519 | 9281 | 7711 | 5033 | 10332 |
| 581 | 1038 | 818 | 479 | 860 | 786 | 485 | 851 | 12076 | 8076 | 13891 | 14421 | 5439 | 3185 | 6646 |
| 621 | 883 | 793 | 606 | 933 | 829 | 720 | 894 | 10878 | 8456 | 13606 | 12009 | 13608 | 10411 | 17170 |
| 790 | 969 | 1188 | 815 | 977 | 1075 | 696 | 850 | 25872 | 17284 | 21879 | 21200 | 28208 | 19349 | 23740 |
| 790 | 1249 | 835 | 876 | 1271 | 883 | 764 | 1195 | 7324 | 6924 | 8766 | 10952 | 5181 | 5436 | 6204 |
| 775 | 925 | 1439 | 748 | 1077 | 1297 | 830 | 941 | 9043 | 4763 | 6145 | 5687 | 8231 | 4280 | 5721 |
| 565 | 873 | 1121 | 943 | 1260 | 1265 | 1016 | 1204 | 13197 | 9617 | 17030 | 14873 | 16219 | 13650 | 14471 |
| 777 | 1068 | 961 | 832 | 1047 | 1002 | 819 | 1108 | 10101 | 7733 | 9949 | 10630 | 7622 | 6601 | 7932 |
| 843 | 1321 | 957 | 825 | 1300 | 878 | 769 | 1165 | 5008 | 4600 | 5456 | 7209 | 10872 | 9374 | 11362 |
| 679 | 1043 | 913 | 775 | 1215 | 963 | 789 | 1277 | 10782 | 9587 | 14114 | 14726 | 7746 | 6573 | 8480 |
| 254 | 723 | 461 | 281 | 781 | 485 | 275 | 703 | 7242 | 4401 | 17352 | 12554 | 6398 | 3896 | 13875 |
| 671 | 1131 | 902 | 703 | 1011 | 832 | 690 | 1030 | 12826 | 11005 | 16397 | 18540 | 5901 | 4604 | 6545 |
| 279 | 567 | 454 | 267 | 595 | 488 | 326 | 600 | 4335 | 2868 | 10294 | 5833 | 2407 | 1418 | 5305 |
| 569 | 833 | 721 | 490 | 849 | 804 | 570 | 912 | 12224 | 9075 | 15940 | 13274 | 16129 | 10957 | 22370 |
| 636 | 995 | 862 | 619 | 1038 | 901 | 716 | 1176 | 7124 | 5568 | 8748 | 8707 | 8187 | 5882 | 9497 |
| 441 | 724 | 615 | 427 | 745 | 732 | 536 | 826 | 8431 | 5868 | 13303 | 9630 | 4498 | 3123 | 7310 |
| 882 | 1313 | 980 | 784 | 1226 | 1110 | 1025 | 1346 | 11722 | 9893 | 11220 | 14731 | 13211 | 10577 | 13484 |
| 800 | 1397 | 700 | 738 | 1226 | 796 | 708 | 1190 | 6245 | 7017 | 8769 | 12255 | 5094 | 5374 | 7279 |
| 537 | 753 | 822 | 531 | 818 | 885 | 587 | 860 | 13166 | 8805 | 16399 | 12351 | 9195 | 5941 | 11187 |
| 608 | 820 | 919 | 717 | 974 | 750 | 522 | 789 | 8385 | 6252 | 10287 | 8436 | 8068 | 6300 | 8781 |
| 477 | 863 | 682 | 472 | 812 | 693 | 457 | 813 | 9767 | 6418 | 13467 | 11625 | 13390 | 9274 | 19643 |

| | | | | | | | | | | | | | | |
|---|---|---|---|---|---|---|---|---|---|---|---|---|---|---|
| 403 | 803 | 839 | 750 | 970 | 664 | 497 | 945 | 6948 | 4716 | 11703 | 9403 | 4347 | 3887 | 5181 |
| 439 | 680 | 754 | 568 | 873 | 762 | 519 | 881 | 4889 | 3226 | 7350 | 4997 | 9206 | 6938 | 12210 |
| 1059 | 1266 | 1133 | 996 | 1418 | 1167 | 909 | 1466 | 4430 | 3744 | 3535 | 4474 | 12530 | 11018 | 11059 |
| 689 | 898 | 1116 | 874 | 1103 | 1052 | 672 | 881 | 13360 | 8330 | 12089 | 10858 | 7533 | 5900 | 6749 |
| 386 | 535 | 714 | 467 | 601 | 627 | 318 | 480 | 16158 | 9327 | 24188 | 12951 | 12618 | 8251 | 17685 |
| 771 | 1292 | 806 | 444 | 825 | 1333 | 792 | 1265 | 5587 | 4149 | 5384 | 6955 | 12422 | 6840 | 15412 |
| 603 | 828 | 597 | 611 | 827 | 582 | 640 | 831 | 4970 | 5190 | 8611 | 7129 | 6283 | 6429 | 10523 |
| 291 | 578 | 650 | 393 | 705 | 563 | 322 | 644 | 13592 | 7932 | 27287 | 15785 | 5978 | 3613 | 9203 |
| 938 | 1237 | 1014 | 1053 | 1400 | 1076 | 1017 | 1174 | 6385 | 5860 | 6250 | 7730 | 2202 | 2288 | 2171 |
| 622 | 1170 | 723 | 677 | 1347 | 886 | 799 | 1354 | 15237 | 12108 | 19469 | 22783 | 7999 | 7492 | 11063 |
| 637 | 1153 | 714 | 592 | 1045 | 699 | 615 | 1101 | 2376 | 2442 | 3835 | 4423 | 8038 | 6660 | 11251 |
| 783 | 1208 | 818 | 643 | 1216 | 821 | 655 | 1251 | 6658 | 5894 | 7523 | 9091 | 7156 | 5626 | 8750 |
| 487 | 926 | 815 | 645 | 945 | 765 | 605 | 969 | 5507 | 4068 | 8358 | 7736 | 4051 | 3203 | 4970 |
| 961 | 1438 | 986 | 1124 | 1565 | 1001 | 1045 | 1516 | 10343 | 11076 | 11528 | 16572 | 5241 | 5971 | 5313 |
| 923 | 1286 | 730 | 711 | 1207 | 893 | 847 | 1383 | 5989 | 5640 | 6113 | 7862 | 13099 | 12753 | 17936 |
| 711 | 976 | 956 | 632 | 850 | 1012 | 731 | 927 | 9320 | 6896 | 9697 | 9466 | 9771 | 6454 | 10216 |
| 578 | 816 | 895 | 705 | 985 | 895 | 658 | 984 | 5659 | 4542 | 7863 | 6418 | 10232 | 8059 | 11432 |
| 653 | 1118 | 1169 | 920 | 1084 | 1064 | 814 | 1032 | 2654 | 1941 | 2973 | 3325 | 23711 | 18675 | 20291 |
| 586 | 1012 | 1044 | 723 | 1154 | 828 | 580 | 1018 | 5709 | 4164 | 7104 | 7190 | 4079 | 2825 | 3908 |
| 971 | 1215 | 975 | 829 | 1175 | 921 | 762 | 1161 | 12037 | 11170 | 11499 | 13967 | 16915 | 14375 | 17343 |
| 348 | 594 | 637 | 529 | 758 | 588 | 462 | 694 | 5050 | 3789 | 10886 | 6472 | 12619 | 10484 | 19810 |
| 696 | 1080 | 1141 | 1074 | 1157 | 721 | 460 | 816 | 8931 | 6374 | 9159 | 9896 | 8900 | 8380 | 7799 |
| 1342 | 1443 | 1186 | 1096 | 1317 | 1178 | 1068 | 1394 | 3717 | 3549 | 2645 | 3816 | 7499 | 6931 | 6324 |
| 454 | 742 | 941 | 468 | 705 | 906 | 455 | 686 | 9878 | 5447 | 12004 | 8905 | 14909 | 7413 | 15849 |
| 662 | 1264 | 739 | 490 | 826 | 821 | 545 | 903 | 6469 | 6140 | 9276 | 11725 | 13001 | 8607 | 17584 |
| 468 | 817 | 776 | 204 | 457 | 620 | 386 | 781 | 7891 | 4902 | 10479 | 8562 | 8977 | 2365 | 11570 |
| 612 | 954 | 744 | 596 | 919 | 802 | 589 | 919 | 7766 | 5893 | 9624 | 9180 | 6585 | 5277 | 8853 |
| 847 | 1132 | 923 | 805 | 1055 | 1026 | 856 | 1113 | 7536 | 8303 | 9801 | 11098 | 10816 | 9439 | 11719 |
| 403 | 850 | 388 | 237 | 845 | 626 | 435 | 966 | 802 | 531 | 1317 | 1120 | 5843 | 3564 | 15066 |
| 580 | 1041 | 827 | 601 | 1025 | 799 | 558 | 1014 | 4955 | 3501 | 6036 | 6282 | 13137 | 9535 | 15877 |
| 786 | 1026 | 1070 | 983 | 1188 | 995 | 805 | 984 | 11920 | 9654 | 12285 | 12600 | 4001 | 3677 | 3740 |

| | | | | | | | | | | | | | | |
|---|---|---|---|---|---|---|---|---|---|---|---|---|---|---|
| 902 | 1305 | 1134 | 992 | 1386 | 1096 | 1005 | 1390 | 8532 | 7272 | 8066 | 10528 | 9228 | 8067 | 8135 |
| 653 | 896 | 884 | 606 | 845 | 737 | 568 | 875 | 13620 | 9840 | 15063 | 13493 | 5554 | 3804 | 6280 |
| 598 | 797 | 989 | 496 | 749 | 1067 | 647 | 823 | 7037 | 3968 | 6636 | 5288 | 12509 | 6266 | 12643 |
| 506 | 1036 | 676 | 495 | 954 | 799 | 579 | 1044 | 3429 | 2317 | 4578 | 4742 | 595 | 436 | 881 |
| 578 | 993 | 819 | 589 | 1011 | 780 | 572 | 1020 | 11849 | 8968 | 15508 | 15398 | 6310 | 4538 | 7704 |
| 772 | 1090 | 924 | 775 | 1043 | 1043 | 822 | 1065 | 1332 | 1057 | 1368 | 1491 | 8235 | 6912 | 8915 |
| 613 | 792 | 1061 | 480 | 773 | 1082 | 431 | 720 | 16680 | 8413 | 13727 | 10871 | 7075 | 3200 | 6666 |
| 386 | 792 | 619 | 394 | 816 | 640 | 409 | 815 | 7684 | 4874 | 12620 | 9999 | 9004 | 5735 | 14543 |
| 762 | 1265 | 1031 | 790 | 1192 | 1033 | 784 | 1166 | 4564 | 3302 | 4335 | 5485 | 15612 | 11967 | 15145 |
| 604 | 869 | 1022 | 887 | 1198 | 777 | 552 | 1068 | 7397 | 4663 | 7722 | 6709 | 7740 | 6719 | 7572 |
| 710 | 1138 | 840 | 729 | 1176 | 768 | 656 | 1169 | 9972 | 8524 | 12005 | 13661 | 5741 | 4984 | 6838 |
| 645 | 911 | 785 | 572 | 887 | 756 | 526 | 752 | 1640 | 1248 | 1934 | 1761 | 7742 | 5643 | 9866 |
| 305 | 669 | 506 | 267 | 683 | 607 | 264 | 638 | 5767 | 3000 | 9833 | 6583 | 2499 | 1322 | 4944 |
| 586 | 842 | 792 | 596 | 886 | 750 | 579 | 849 | 11249 | 7953 | 13578 | 11437 | 7758 | 5838 | 9794 |
| 813 | 1132 | 1245 | 901 | 1154 | 1177 | 849 | 1129 | 8247 | 5875 | 7230 | 8185 | 15577 | 11277 | 12513 |
| 1153 | 1477 | 1227 | 834 | 1210 | 1286 | 1001 | 1305 | 13835 | 10573 | 9172 | 13550 | 15562 | 10569 | 12678 |
| 737 | 1058 | 867 | 854 | 1232 | 873 | 881 | 1328 | 2405 | 2134 | 2894 | 3061 | 8840 | 8713 | 10198 |
| 945 | 1255 | 1095 | 851 | 1221 | 1223 | 967 | 1217 | 4297 | 3487 | 3688 | 4630 | 5201 | 4039 | 4748 |
| 573 | 892 | 944 | 598 | 961 | 1010 | 699 | 1359 | 16941 | 10746 | 18762 | 16730 | 8680 | 5495 | 9192 |
| 667 | 961 | 857 | 550 | 890 | 730 | 450 | 828 | 13932 | 9796 | 14685 | 14106 | 9497 | 6092 | 11075 |
| 348 | 800 | 457 | 350 | 771 | 513 | 404 | 833 | 17464 | 9750 | 27982 | 22377 | 2306 | 1764 | 5047 |
| 648 | 1017 | 835 | 694 | 1044 | 890 | 802 | 1107 | 11689 | 9256 | 14278 | 14517 | 8586 | 7140 | 10286 |
| 621 | 930 | 777 | 600 | 868 | 818 | 622 | 888 | 3792 | 2811 | 4525 | 4207 | 3343 | 2581 | 4301 |
| 926 | 1131 | 913 | 919 | 1206 | 885 | 899 | 1129 | 7094 | 7933 | 8563 | 9689 | 10442 | 10509 | 11432 |
| 1075 | 1375 | 1104 | 973 | 1255 | 932 | 840 | 1097 | 5503 | 5312 | 4940 | 6791 | 7602 | 6700 | 6889 |
| 415 | 838 | 592 | 492 | 851 | 608 | 507 | 846 | 3693 | 2799 | 6748 | 5654 | 4178 | 3468 | 7053 |
| 119 | 596 | 397 | 260 | 723 | 294 | 184 | 645 | 1711 | 959 | 8068 | 4810 | 6483 | 4237 | 16310 |
| 536 | 889 | 850 | 553 | 914 | 692 | 354 | 776 | 7076 | 4209 | 7859 | 6989 | 9307 | 6063 | 10955 |
| 618 | 1037 | 940 | 660 | 999 | 975 | 772 | 1071 | 10238 | 6296 | 10185 | 10562 | 10587 | 7436 | 11260 |
| 691 | 918 | 1275 | 962 | 1097 | 847 | 723 | 863 | 1148 | 958 | 1387 | 1273 | 5106 | 3852 | 4004 |
| 839 | 961 | 1000 | 811 | 1086 | 1015 | 767 | 979 | 560 | 358 | 427 | 411 | 9060 | 7350 | 9060 |

| | | | | | | | | | | | | | | |
|---|---|---|---|---|---|---|---|---|---|---|---|---|---|---|
| 467 | 739 | 677 | 492 | 732 | 635 | 393 | 540 | 3732 | 2628 | 5627 | 4158 | 1332 | 967 | 1967 |
| 379 | 662 | 751 | 356 | 681 | 763 | 397 | 654 | 12442 | 6127 | 16175 | 10704 | 8739 | 4146 | 11631 |
| 608 | 910 | 684 | 523 | 820 | 1149 | 935 | 1168 | 5976 | 4843 | 7963 | 7245 | 6152 | 4708 | 8996 |
| 788 | 995 | 832 | 567 | 854 | 1271 | 746 | 892 | 11015 | 7028 | 8919 | 8879 | 8195 | 5583 | 9846 |
| 335 | 663 | 467 | 323 | 693 | 480 | 325 | 700 | 9217 | 6326 | 18894 | 12530 | 7476 | 5179 | 16020 |
| 564 | 912 | 595 | 387 | 627 | 596 | 411 | 681 | 5763 | 4035 | 7150 | 6521 | 5976 | 3888 | 10041 |
| 1067 | 1293 | 938 | 838 | 1177 | 1118 | 1120 | 1422 | 5536 | 5796 | 5433 | 7026 | 5624 | 5022 | 5994 |
| 1173 | 561 | 1512 | 985 | 356 | 1793 | 1175 | 602 | 12060 | 7940 | 6771 | 3797 | 3221 | 2099 | 2131 |
| 804 | 1080 | 1078 | 876 | 1251 | 913 | 732 | 1028 | 8047 | 7039 | 8750 | 9454 | 4127 | 3352 | 3827 |
| 373 | 601 | 691 | 285 | 535 | 681 | 224 | 478 | 17489 | 8315 | 22306 | 13414 | 4345 | 1790 | 6288 |
| 653 | 990 | 707 | 562 | 893 | 684 | 439 | 742 | 10796 | 8995 | 13783 | 13643 | 2077 | 1651 | 2936 |
| 603 | 988 | 689 | 643 | 966 | 823 | 818 | 982 | 5581 | 5267 | 8738 | 8635 | 6593 | 6152 | 9572 |
| 1506 | 1634 | 860 | 727 | 1122 | 812 | 644 | 1135 | 3788 | 3956 | 2626 | 4291 | 4420 | 3740 | 5142 |
| 640 | 985 | 787 | 603 | 907 | 891 | 534 | 850 | 13526 | 7897 | 12332 | 12151 | 2308 | 1770 | 2934 |
| 365 | 728 | 502 | 348 | 688 | 690 | 528 | 780 | 5787 | 3987 | 10926 | 7957 | 4349 | 3018 | 8663 |
| 579 | 919 | 716 | 584 | 932 | 759 | 676 | 1036 | 14044 | 10288 | 17755 | 16325 | 10734 | 8757 | 14998 |
| 525 | 975 | 633 | 426 | 766 | 531 | 406 | 701 | 7777 | 5588 | 10652 | 10381 | 14562 | 9808 | 23001 |
| 978 | 1418 | 891 | 977 | 1499 | 867 | 1048 | 1569 | 11762 | 11272 | 11524 | 16345 | 8332 | 9138 | 9355 |
| 368 | 735 | 585 | 326 | 688 | 580 | 363 | 721 | 9183 | 5424 | 14721 | 10826 | 8054 | 4487 | 13764 |
| 707 | 1215 | 623 | 496 | 1000 | 676 | 570 | 1047 | 10739 | 9076 | 12833 | 15595 | 9545 | 7600 | 15312 |
| 699 | 1026 | 721 | 524 | 858 | 808 | 518 | 900 | 9351 | 6713 | 9597 | 9843 | 3263 | 2373 | 4526 |
| 783 | 1104 | 911 | 730 | 1008 | 777 | 716 | 1102 | 11298 | 10605 | 13549 | 14954 | 7329 | 5873 | 8046 |
| 548 | 902 | 927 | 1155 | 1500 | 841 | 727 | 1037 | 8109 | 5975 | 10905 | 9840 | 1815 | 2261 | 1958 |
| 1147 | 1577 | 955 | 1019 | 1427 | 972 | 1062 | 1463 | 5320 | 5447 | 4750 | 7492 | 9370 | 9995 | 9808 |
| 793 | 1198 | 991 | 857 | 1087 | 982 | 675 | 1021 | 3579 | 2869 | 3618 | 4335 | 9825 | 8496 | 9911 |
| 724 | 1205 | 963 | 863 | 1256 | | | | 1999 | 1609 | 2223 | 2680 | 7613 | 6822 | 7908 |
| 809 | 1145 | 909 | 667 | 982 | 890 | 744 | 1226 | 3390 | 3032 | 3749 | 4291 | 4737 | 3473 | 5210 |
| 536 | 956 | 724 | 508 | 965 | 748 | 512 | 941 | 5291 | 3288 | 6132 | 5862 | 14063 | 9875 | 19423 |
| 800 | 1488 | 876 | 887 | 1465 | 753 | 758 | 1421 | 2736 | 2292 | 2864 | 4262 | 5981 | 6052 | 6827 |
| | | 827 | 602 | 890 | 675 | 505 | 960 | | | | | 465 | 338 | 562 |
| 537 | 876 | 658 | 424 | 713 | 740 | 520 | 871 | 11864 | 9037 | 16838 | 14753 | 7101 | 4579 | 10800 |

| | | | | | | | | | | | | | | |
|---|---|---|---|---|---|---|---|---|---|---|---|---|---|---|
| 493 | 984 | 837 | 635 | 1077 | 871 | 677 | 1143 | 3842 | 2923 | 5928 | 5831 | 11076 | 8401 | 13228 |
| 323 | 838 | 485 | 290 | 753 | 429 | 318 | 849 | 6046 | 4501 | 13946 | 11692 | 6199 | 3705 | 12773 |
| 1206 | 956 | 785 | 1045 | 742 | 961 | 1244 | 1003 | 8815 | 11450 | 9491 | 9076 | 5092 | 6779 | 6485 |
| 1347 | 1263 | 1131 | 1194 | 1195 | 1135 | 1237 | 1342 | 6770 | 8288 | 6152 | 7772 | 11804 | 12453 | 10434 |
| 1432 | 2041 | 1169 | 1102 | 1560 | 1060 | 1082 | 1656 | 3019 | 3146 | 2197 | 4484 | 2630 | 2479 | 2250 |
| 650 | 881 | 825 | 612 | 867 | 860 | 615 | 858 | 5045 | 3866 | 5950 | 5244 | 8520 | 6316 | 10323 |
| 414 | 871 | 683 | 421 | 902 | 599 | 376 | 817 | 4351 | 2828 | 6838 | 5953 | 5014 | 3089 | 7344 |
| 436 | 648 | 319 | 442 | 665 | 208 | 342 | 562 | 4150 | 5930 | 13604 | 8810 | 5708 | 7904 | 17878 |
| 829 | 1195 | 977 | 843 | 1057 | 932 | 782 | 1032 | 7697 | 6340 | 7647 | 9136 | 4336 | 3743 | 4438 |
| 715 | 1194 | 916 | 625 | 1229 | 900 | 656 | 1226 | 10129 | 7780 | 10886 | 12997 | 4561 | 3111 | 4977 |
| 881 | 1159 | 1188 | 1112 | 1513 | 1063 | 919 | 1312 | 6604 | 5274 | 5988 | 6940 | 658 | 616 | 554 |
| 540 | 875 | 696 | 500 | 866 | 776 | 562 | 880 | 1231 | 923 | 1711 | 1498 | 8935 | 6417 | 12837 |
| 867 | 1081 | 716 | 415 | 780 | 1059 | 738 | 953 | 4502 | 3611 | 4164 | 4502 | 1486 | 862 | 2074 |
| 334 | 651 | 700 | 433 | 781 | 724 | 485 | 818 | 6699 | 3248 | 9728 | 6328 | 6403 | 3959 | 9143 |
| 443 | 769 | 403 | 298 | 563 | 365 | 229 | 473 | 11587 | 8926 | 20160 | 15499 | 4427 | 3271 | 10984 |
| 895 | 1112 | 885 | 673 | 978 | | | | 4409 | 3608 | 4033 | 4484 | 11066 | 8414 | 12504 |
| 526 | 845 | 825 | 555 | 1020 | 776 | 495 | 801 | 8715 | 5781 | 10982 | 9281 | 2809 | 1888 | 3405 |
| 970 | 3312 | 917 | 888 | 3229 | 888 | 778 | 3613 | 3644 | 4103 | 4230 | 14009 | 4034 | 3908 | 4400 |
| 493 | 791 | 696 | 548 | 860 | 695 | 532 | 802 | 9703 | 7109 | 14434 | 11424 | 7147 | 5625 | 10264 |
| 882 | 1416 | 1085 | 950 | 1701 | 1050 | 949 | 1667 | 3931 | 3449 | 3912 | 5538 | 3447 | 3018 | 3178 |
| 536 | 982 | 589 | 401 | 690 | 864 | 628 | 934 | 4366 | 2993 | 5580 | 5482 | 3599 | 2452 | 6112 |
| 799 | 1196 | 932 | 738 | 1060 | 679 | 605 | 1061 | 10613 | 10074 | 12605 | 15079 | 9364 | 7420 | 10047 |
| 592 | 733 | 833 | 279 | 396 | 856 | 401 | 560 | 7519 | 5738 | 9697 | 7106 | 15221 | 5107 | 18280 |
| 616 | 965 | 898 | 693 | 1001 | 842 | 625 | 933 | 8709 | 6988 | 11345 | 10954 | 5161 | 3983 | 5749 |
| 592 | 1054 | 638 | 355 | 773 | 685 | 442 | 871 | 5182 | 3489 | 5894 | 6213 | 8546 | 4755 | 13394 |
| 413 | 698 | 503 | 378 | 779 | 519 | 365 | 653 | 1982 | 1516 | 3669 | 2562 | 2027 | 1523 | 4032 |
| 579 | 849 | 1026 | 750 | 1157 | 958 | 701 | 1071 | 8601 | 5627 | 9721 | 8255 | 2805 | 2050 | 2733 |
| 470 | 680 | 285 | 488 | 698 | 323 | 458 | 705 | 6691 | 9555 | 20323 | 13816 | 3703 | 6345 | 13000 |
| 530 | 966 | 759 | 507 | 867 | 1001 | 785 | 1039 | 657 | 459 | 867 | 837 | 6168 | 4126 | 8132 |
| 535 | 910 | 464 | 601 | 963 | 425 | 446 | 901 | 2608 | 2855 | 5335 | 4855 | 1333 | 1727 | 2874 |
| 855 | 903 | 833 | 724 | 939 | 924 | 827 | 929 | 10769 | 9497 | 11108 | 10030 | 11784 | 10243 | 14144 |

| | | | | | | | | | | | | | | |
|---|---|---|---|---|---|---|---|---|---|---|---|---|---|---|
| 442 | 818 | 1073 | 394 | 816 | 1052 | 459 | 866 | 15372 | 6418 | 14534 | 11891 | 16684 | 6122 | 15545 |
| 671 | 1009 | 1050 | 680 | 1048 | 975 | 732 | 1093 | 8238 | 5390 | 8032 | 8104 | 8237 | 5328 | 7841 |
| 551 | 984 | 730 | 630 | 1059 | 865 | 651 | 1167 | 2824 | 2267 | 4117 | 4053 | 2857 | 2467 | 3914 |
| 901 | 1195 | 713 | 598 | 953 | 948 | 822 | 1119 | 3612 | 3175 | 3522 | 4210 | 3132 | 2628 | 4393 |
| 670 | 1027 | 878 | 839 | 1186 | 758 | 586 | 1047 | 9271 | 7435 | 11095 | 11393 | 3427 | 3272 | 3902 |
| 459 | 773 | 796 | 622 | 1040 | 837 | 722 | 986 | 966 | 720 | 1570 | 1215 | 7502 | 5860 | 9421 |
| 1009 | 1204 | 896 | 598 | 957 | 900 | 497 | 824 | 10703 | 9568 | 9487 | 11420 | 22612 | 15097 | 25241 |
| 1011 | 1205 | 1365 | 1222 | 1354 | 1137 | 977 | 1151 | 9019 | 7460 | 7379 | 8889 | 3303 | 2956 | 2419 |
| 449 | 767 | 623 | 443 | 687 | 957 | 697 | 910 | 15978 | 10313 | 22965 | 17625 | 1945 | 1385 | 3123 |
| 756 | 1102 | 921 | 806 | 1145 | 859 | 837 | 1079 | 6861 | 5740 | 7593 | 8368 | 5682 | 4972 | 6167 |
| 605 | 876 | 774 | 507 | 867 | 872 | 594 | 848 | 7448 | 5436 | 8981 | 7868 | 6375 | 4176 | 8236 |
| 731 | 1245 | 926 | 739 | 1098 | | | | 11448 | 9271 | 12676 | 15777 | 3728 | 2975 | 4027 |
| 885 | 1473 | 844 | 826 | 1157 | 855 | 799 | 1131 | 838 | 874 | 988 | 1456 | 5827 | 5701 | 6902 |
| 698 | 1066 | 2343 | 2111 | 2119 | 937 | 733 | 1026 | 8779 | 7507 | 10751 | 11461 | 2130 | 1919 | 909 |
| 590 | 853 | 537 | 400 | 692 | 678 | 482 | 808 | 7694 | 5594 | 9479 | 8087 | 3779 | 2817 | 7042 |
| 923 | 1400 | 876 | 842 | 1358 | 948 | 881 | 1219 | 2463 | 2290 | 2480 | 3472 | 3717 | 3573 | 4244 |
| 559 | 927 | 792 | 602 | 982 | 910 | 649 | 1067 | 9667 | 7017 | 12558 | 11639 | 5195 | 3950 | 6563 |
| 669 | 1144 | 803 | 623 | 1072 | 907 | 748 | 1008 | 4101 | 3392 | 5073 | 5803 | 6441 | 5000 | 8023 |
| 810 | 1009 | 1019 | 744 | 1051 | 891 | 722 | 968 | 3851 | 3240 | 3998 | 4034 | 11573 | 8456 | 11362 |
| 501 | 576 | 683 | 504 | 830 | 655 | 426 | 695 | 850 | 580 | 1156 | 666 | 4403 | 3250 | 6447 |
| 787 | 1046 | 932 | 590 | 975 | 977 | 870 | 1174 | 10746 | 8637 | 10969 | 11477 | 8067 | 5106 | 8659 |
| | | 1018 | 484 | 661 | 629 | 335 | 575 | | | | | 6966 | 3308 | 6842 |
| 738 | 1044 | 1007 | 940 | 1215 | 1235 | 1177 | 1294 | 5391 | 4409 | 5975 | 6239 | 2113 | 1972 | 2097 |
| 543 | 832 | 855 | 530 | 763 | 793 | 559 | 930 | 5187 | 3452 | 6354 | 5285 | 7819 | 4851 | 9150 |
| 355 | 800 | 511 | 386 | 807 | 519 | 368 | 805 | 8363 | 5687 | 16041 | 12835 | 9481 | 7172 | 18559 |
| 298 | 704 | 681 | 508 | 809 | 711 | 586 | 967 | 1131 | 854 | 2860 | 2012 | 13788 | 10292 | 20256 |
| 900 | 1359 | 1062 | 996 | 1518 | 928 | 1023 | 1548 | 11098 | 9893 | 10988 | 14936 | 1366 | 1281 | 1286 |
| 659 | 935 | 1109 | 667 | 883 | 923 | 561 | 802 | 10839 | 7021 | 10649 | 9958 | 3855 | 2317 | 3475 |
| 775 | 1207 | 1068 | 908 | 1235 | 993 | 872 | 1226 | 11660 | 9886 | 12750 | 15395 | 3393 | 2885 | 3178 |
| 481 | 999 | 859 | 678 | 1044 | 745 | 505 | 993 | 12598 | 8414 | 17482 | 17464 | 4874 | 3851 | 5677 |
| 682 | 944 | 733 | 574 | 855 | 941 | 671 | 923 | 6325 | 4917 | 7213 | 6808 | 1703 | 1332 | 2323 |

| | | | | | | | | | | | | | | |
|---|---|---|---|---|---|---|---|---|---|---|---|---|---|---|
| 624 | 982 | 975 | 953 | 1142 | 870 | 828 | 1078 | 5350 | 3911 | 6264 | 6152 | 4663 | 4558 | 4782 |
| 663 | 1030 | 967 | 675 | 1083 | 880 | 604 | 1022 | 8988 | 6546 | 9875 | 10169 | 10758 | 7512 | 11129 |
| 404 | 866 | 687 | 485 | 929 | 582 | 400 | 841 | 8875 | 5760 | 14273 | 12357 | 9503 | 6706 | 13824 |
| 399 | 731 | 809 | 457 | 762 | 1003 | 681 | 916 | 7454 | 3823 | 9580 | 7004 | 10125 | 5727 | 12520 |
| 585 | 779 | 761 | 525 | 714 | 686 | 470 | 590 | 3021 | 1852 | 3165 | 2467 | 8130 | 5615 | 10689 |
| 323 | 608 | 821 | 313 | 625 | 754 | 296 | 592 | 4316 | 1851 | 5726 | 3481 | 7623 | 2912 | 9291 |
| 499 | 654 | 1187 | 925 | 1017 | 1097 | 903 | 963 | 12200 | 8318 | 16678 | 10908 | 5283 | 4117 | 4451 |
| 615 | 1013 | 795 | 625 | 1083 | 847 | 625 | 973 | 3530 | 2620 | 4263 | 4319 | 5351 | 4205 | 6730 |
| 669 | 989 | 827 | 530 | 824 | 713 | 418 | 638 | 7207 | 4892 | 7308 | 7229 | 13454 | 8625 | 16266 |
| 548 | 907 | 932 | 636 | 974 | 1023 | 574 | 915 | 6206 | 3291 | 6010 | 5454 | 11046 | 7535 | 11849 |
| 448 | 679 | 774 | 330 | 579 | 545 | 341 | 538 | 7725 | 5937 | 13259 | 9006 | 17547 | 7490 | 22681 |
| 902 | 1329 | 952 | 824 | 1241 | 814 | 700 | 1050 | 15482 | 14221 | 15774 | 20965 | 12545 | 10859 | 13180 |
| 727 | 1060 | 749 | 575 | 921 | 794 | 687 | 1075 | 5645 | 5085 | 6993 | 7409 | 14263 | 10956 | 19047 |
| 632 | 778 | 525 | 479 | 724 | 673 | 513 | 803 | 4870 | 5199 | 8228 | 6404 | 3131 | 2861 | 5969 |
| 449 | 1099 | 941 | 801 | 1088 | 697 | 582 | 1159 | 5331 | 3874 | 8634 | 9486 | 6754 | 5750 | 7175 |
| 634 | 1037 | 927 | 526 | 948 | 1075 | 574 | 1039 | 8746 | 5389 | 8493 | 8808 | 4590 | 2605 | 4951 |
| 519 | 882 | 791 | 560 | 913 | 879 | 618 | 1019 | 6236 | 4397 | 8477 | 7474 | 14187 | 10038 | 17929 |
| 914 | 1135 | 1180 | 903 | 1182 | 1244 | 948 | 1074 | 17173 | 13861 | 15159 | 17203 | 10916 | 8360 | 9253 |
| 703 | 1378 | 1000 | 988 | 1494 | 929 | 1044 | 1715 | 5612 | 5410 | 7694 | 10604 | 8204 | 8104 | 8202 |
| 680 | 938 | 881 | 788 | 1124 | 813 | 710 | 997 | 1672 | 1365 | 2008 | 1883 | 3502 | 3133 | 3976 |
| 414 | 875 | 576 | 474 | 915 | 643 | 483 | 882 | 10813 | 8317 | 20076 | 17561 | 4687 | 3859 | 8139 |
| 877 | 1288 | 931 | 885 | 1300 | 955 | 949 | 1387 | 3484 | 3857 | 4399 | 5667 | 11692 | 11119 | 12562 |
| 199 | 426 | 1199 | 251 | 469 | 1193 | 234 | 439 | 6662 | 1329 | 6662 | 2838 | 16862 | 3526 | 14068 |
| 1004 | 1055 | 637 | 1032 | 837 | 783 | 1090 | 838 | 2381 | 2885 | 2874 | 3032 | 3013 | 4881 | 4731 |
| 740 | 1205 | 943 | 815 | 1279 | 666 | 566 | 913 | 13181 | 10893 | 14714 | 17728 | 9373 | 8095 | 9935 |
| 623 | 1314 | 805 | 713 | 1283 | 748 | 687 | 1158 | 3288 | 2974 | 4773 | 6273 | 7908 | 7007 | 9828 |
| 835 | 1064 | 1202 | 953 | 1252 | 1281 | 1061 | 1142 | 4953 | 3663 | 4385 | 4665 | 2853 | 2264 | 2374 |
| 796 | 1442 | 980 | 771 | 1321 | 896 | 720 | 1237 | 6077 | 5406 | 6796 | 9796 | 5215 | 4099 | 5319 |
| 604 | 1022 | 795 | 549 | 932 | 872 | 609 | 957 | 13201 | 9764 | 16153 | 16507 | 12582 | 8685 | 15821 |
| 907 | 1690 | 874 | 1091 | 1862 | 880 | 1010 | 1690 | 3353 | 3593 | 3964 | 6697 | 4138 | 5168 | 4736 |
| 723 | 941 | 928 | 645 | 795 | 1047 | 809 | 1086 | 9631 | 7541 | 10425 | 9805 | 2128 | 1480 | 2292 |

| | | | | | | | | | | | | | | |
|---|---|---|---|---|---|---|---|---|---|---|---|---|---|---|
| 569 | 990 | 785 | 549 | 929 | 827 | 626 | 976 | 10257 | 7201 | 12646 | 12518 | 6083 | 4250 | 7748 |
| 896 | 1350 | 855 | 795 | 1329 | 924 | 884 | 1358 | 7265 | 6547 | 7307 | 9862 | 10216 | 9501 | 11955 |
| 501 | 729 | 644 | 481 | 634 | 560 | 493 | 729 | 1332 | 994 | 1981 | 1444 | 10941 | 8180 | 16991 |
| 333 | 626 | 1160 | 426 | 641 | 1085 | 393 | 624 | 14640 | 4175 | 12524 | 7836 | 7509 | 2760 | 6474 |
| 447 | 820 | 679 | 350 | 683 | 742 | 459 | 787 | 4969 | 3020 | 6749 | 5532 | 4818 | 2481 | 7093 |
| 899 | 1126 | 926 | 673 | 1015 | 914 | 565 | 918 | 4964 | 4169 | 4635 | 5220 | 4388 | 3188 | 4737 |
| 444 | 715 | 767 | 545 | 788 | 728 | 536 | 771 | 2070 | 1402 | 3161 | 2259 | 4539 | 3223 | 5916 |
| 496 | 842 | 716 | 520 | 872 | 792 | 560 | 912 | 5776 | 3967 | 8000 | 6739 | 6038 | 4381 | 8433 |
| 1071 | 1333 | 1288 | 1143 | 1481 | 1208 | 1090 | 1427 | 5460 | 4796 | 4479 | 5972 | 7848 | 6967 | 6093 |
| 836 | 4051 | 1144 | 1111 | 7782 | 991 | 947 | 5609 | 3948 | 3508 | 4198 | 17008 | 4717 | 4580 | 4123 |
| 1161 | 1318 | 818 | 681 | 1036 | 1290 | 1138 | 1347 | 3109 | 3049 | 2627 | 3463 | 6346 | 5288 | 7760 |
| 620 | 824 | 726 | 533 | 873 | 731 | 564 | 880 | 1820 | 1584 | 2556 | 2106 | 3314 | 2435 | 4566 |
| 684 | 1092 | 734 | 647 | 1049 | 670 | 516 | 871 | 5913 | 4784 | 6993 | 7637 | 4134 | 3644 | 5628 |
| 455 | 935 | 650 | 441 | 891 | 640 | 473 | 877 | 4999 | 3452 | 7585 | 7091 | 8615 | 5846 | 13246 |
| 722 | 1012 | 917 | 843 | 1338 | 979 | 733 | 1032 | 3175 | 2757 | 3819 | 3866 | 5785 | 5315 | 6306 |
| 774 | 834 | 1254 | 779 | 1023 | 1219 | 757 | 985 | 8274 | 5412 | 6991 | 5830 | 13901 | 8637 | 11089 |
| 1632 | 2446 | 892 | 1509 | 2208 | 1144 | 1746 | 2474 | 4380 | 7399 | 4535 | 11093 | 4198 | 7100 | 4706 |
| 529 | 990 | 974 | 912 | 1362 | 915 | 730 | 1139 | 6785 | 4393 | 8301 | 8216 | 3029 | 2836 | 3110 |
| 628 | 955 | 744 | 514 | 923 | 735 | 512 | 917 | 6556 | 5286 | 8420 | 8038 | 3173 | 2192 | 4264 |
| 708 | 918 | 797 | 578 | 869 | 1120 | 855 | 998 | 9039 | 6357 | 8977 | 8241 | 10505 | 7613 | 13181 |
| 541 | 674 | 1404 | 681 | 738 | 1082 | 681 | 798 | 1605 | 598 | 1106 | 745 | 1123 | 545 | 800 |
| 1466 | 1224 | 1073 | 1198 | 1135 | 789 | 772 | 1315 | 2176 | 2466 | 1682 | 2058 | 3869 | 4322 | 3607 |
| 705 | 1123 | 771 | 460 | 757 | 1001 | 792 | 1126 | 5668 | 4222 | 5987 | 6720 | 8554 | 5104 | 11099 |
| 1225 | 1511 | 1221 | 1105 | 1411 | 1045 | 973 | 1327 | 6047 | 5802 | 4736 | 7157 | 6427 | 5819 | 5265 |
| 542 | 912 | 856 | 653 | 883 | 868 | 667 | 1068 | 1251 | 847 | 1563 | 1426 | 1678 | 1280 | 1960 |
| 531 | 859 | 803 | 483 | 763 | 772 | 467 | 752 | 6118 | 3573 | 6735 | 5788 | 3315 | 1992 | 4127 |
| 645 | 1008 | 759 | 609 | 941 | 742 | 661 | 1052 | 12517 | 9656 | 14977 | 15098 | 8753 | 7028 | 11532 |
| 604 | 858 | 817 | 593 | 854 | 804 | 560 | 836 | 7686 | 5703 | 9434 | 8093 | 7374 | 5351 | 9028 |
| 710 | 930 | 744 | 771 | 919 | 775 | 635 | 862 | 5061 | 4617 | 6501 | 6046 | 4258 | 4409 | 5721 |
| 764 | 1260 | 1026 | 755 | 1163 | 1017 | 765 | 1214 | 5392 | 3460 | 4527 | 5705 | 8695 | 6402 | 8477 |
| 712 | 1094 | 1086 | 974 | 1382 | 948 | 742 | 1054 | 10226 | 7930 | 11141 | 12189 | 2745 | 2463 | 2528 |

| | | | | | | | | | | | | | | |
|---|---|---|---|---|---|---|---|---|---|---|---|---|---|---|
| 682 | 905 | 812 | 721 | 984 | 768 | 735 | 952 | 5021 | 4767 | 6994 | 6331 | 5409 | 4799 | 6661 |
| | | 715 | 449 | 814 | | | | | | | | 2470 | 1550 | 3455 |
| 310 | 728 | 501 | 337 | 634 | 511 | 332 | 635 | 1185 | 705 | 2273 | 1655 | 6512 | 4375 | 12993 |
| 859 | 1312 | 1032 | 778 | 1255 | 1014 | 963 | 1431 | 6470 | 5183 | 6033 | 7918 | 5834 | 4402 | 5655 |
| 719 | 924 | 1119 | 834 | 1165 | 1255 | 772 | 962 | 5230 | 3477 | 4838 | 4472 | 4270 | 3181 | 3816 |
| 321 | 545 | 438 | 362 | 622 | 381 | 347 | 638 | 3184 | 2634 | 8204 | 4467 | 5339 | 4415 | 12201 |
| 477 | 966 | 577 | 446 | 954 | 794 | 618 | 996 | 6109 | 4722 | 9902 | 9570 | 9280 | 7167 | 16085 |
| 563 | 754 | 679 | 538 | 728 | 634 | 522 | 740 | 6139 | 4830 | 8579 | 6472 | 3161 | 2508 | 4659 |
| 584 | 905 | 1073 | 845 | 1287 | 1188 | 1055 | 1354 | 6457 | 5275 | 9032 | 8175 | 4926 | 3883 | 4592 |
| 389 | 986 | 445 | 406 | 1024 | 384 | 311 | 934 | 8784 | 7536 | 19364 | 19091 | 3233 | 2946 | 7262 |
| 615 | 883 | 857 | 602 | 939 | 784 | 557 | 873 | 9087 | 6539 | 10636 | 9390 | 5721 | 4019 | 6672 |
| 486 | 724 | 832 | 642 | 934 | 794 | 424 | 761 | 490 | 330 | 679 | 492 | 3415 | 2634 | 4105 |
| 934 | 1294 | 1150 | 1049 | 1502 | 1301 | 1131 | 1377 | 8362 | 7114 | 7614 | 9850 | 7510 | 6848 | 6530 |
| 527 | 837 | 598 | 630 | 955 | 538 | 534 | 913 | 9798 | 9031 | 17152 | 14363 | 4655 | 4904 | 7781 |
| 600 | 967 | 659 | 486 | 819 | 647 | 500 | 799 | 4227 | 3204 | 5340 | 5162 | 2510 | 1852 | 3808 |
| 1025 | 1307 | 869 | 868 | 1124 | 837 | 792 | 1039 | 1377 | 1417 | 1383 | 1807 | 1499 | 1497 | 1726 |
| 245 | 557 | 307 | 218 | 483 | 380 | 303 | 631 | 6919 | 4602 | 18758 | 10444 | 4294 | 3052 | 13981 |
| 698 | 1055 | 700 | 623 | 1013 | 895 | 683 | 1099 | 1352 | 969 | 1389 | 1465 | 2731 | 2431 | 3903 |
| 751 | 926 | 921 | 694 | 933 | 862 | 684 | 896 | 12444 | 9979 | 13285 | 12304 | 2559 | 1928 | 2779 |
| 481 | 645 | 1013 | 926 | 1376 | 848 | 729 | 1161 | 1930 | 1156 | 2402 | 1549 | 3630 | 3316 | 3582 |
| 467 | 713 | 382 | 483 | 706 | 343 | 408 | 578 | 5090 | 6074 | 13014 | 9280 | 4490 | 5671 | 11746 |
| 355 | 719 | 517 | 486 | 835 | | | | 1900 | 1941 | 5471 | 3932 | 3993 | 3757 | 7727 |
| 332 | 522 | 486 | 287 | 460 | 597 | 360 | 547 | 425 | 275 | 827 | 431 | 3923 | 2314 | 8065 |
| 448 | 739 | 634 | 440 | 662 | 757 | 479 | 683 | 7207 | 4735 | 10563 | 7808 | 5212 | 3618 | 8218 |
| 813 | 1039 | 1009 | 751 | 990 | 1045 | 896 | 1104 | 12218 | 10094 | 12413 | 12900 | 4352 | 3236 | 4312 |
| 1103 | 1513 | 1153 | 1106 | 1294 | 1139 | 806 | 1046 | 6908 | 7340 | 6653 | 10068 | 8720 | 8368 | 7564 |
| 744 | 1257 | 853 | 838 | 1486 | 1094 | 861 | 1242 | 1934 | 1668 | 2240 | 2817 | 1832 | 1799 | 2147 |
| 562 | 950 | 708 | 558 | 918 | 708 | 523 | 880 | 9754 | 7411 | 13187 | 12524 | 6310 | 4972 | 8909 |
| | | 1067 | 1007 | 1365 | 773 | 562 | 923 | | | | | 7074 | 6678 | 6632 |
| 1034 | 1422 | 1022 | 932 | 1236 | 1254 | 1113 | 1868 | 5913 | 6081 | 5881 | 8363 | 6491 | 5919 | 6350 |
| 484 | 747 | 835 | 560 | 814 | 739 | 437 | 740 | 12461 | 8096 | 16719 | 12481 | 17343 | 11633 | 20783 |

| | | | | | | | | | | | | | | |
|---|---|---|---|---|---|---|---|---|---|---|---|---|---|---|
| 622 | 1150 | 808 | 561 | 1066 | 822 | 624 | 1138 | 8177 | 6573 | 10563 | 12148 | 3784 | 2629 | 4683 |
| 260 | 578 | 824 | 470 | 655 | 401 | 232 | 624 | 3190 | 2040 | 7852 | 4542 | 3656 | 2088 | 4439 |
| 444 | 758 | 674 | 475 | 821 | 539 | 356 | 696 | 10891 | 7688 | 17319 | 13128 | 5907 | 4161 | 8766 |
| 535 | 986 | 974 | 666 | 1029 | 939 | 631 | 1053 | 5702 | 2996 | 5598 | 5521 | 2725 | 1862 | 2797 |
| | | 1017 | 801 | 1192 | 944 | 765 | 1133 | | | | | 10437 | 8214 | 10260 |
| 476 | 699 | 901 | 337 | 525 | 901 | 357 | 564 | 17023 | 8349 | 17529 | 12259 | 9515 | 3566 | 10566 |
| 843 | 1339 | 829 | 838 | 1226 | 658 | 608 | 1242 | 4746 | 4154 | 4928 | 6601 | 2797 | 2829 | 3375 |
| 568 | 1033 | 865 | 527 | 1049 | 866 | 567 | 1025 | 5489 | 3570 | 6285 | 6493 | 3018 | 1841 | 3490 |
| 517 | 957 | 804 | 586 | 986 | 721 | 544 | 1006 | 7248 | 5767 | 11161 | 10681 | 14613 | 10645 | 18164 |
| 475 | 856 | 727 | 639 | 929 | 891 | 938 | 1175 | 3597 | 2821 | 5942 | 5087 | 2773 | 2435 | 3812 |
| 918 | 1117 | 1073 | 836 | 1069 | 1234 | 1050 | 1221 | 2086 | 1696 | 1848 | 2065 | 5753 | 4479 | 5360 |
| 749 | 861 | 1136 | 683 | 904 | 877 | 662 | 991 | 2214 | 1785 | 2383 | 2051 | 3149 | 1892 | 2772 |
| 859 | 1151 | 1058 | 923 | 1072 | 1058 | 844 | 1039 | 819 | 668 | 778 | 896 | 2414 | 2105 | 2282 |
| 1051 | 1331 | 1198 | 1323 | 1417 | 1161 | 993 | 1441 | 7405 | 6904 | 6567 | 8743 | 2227 | 2459 | 1859 |

| | | | | | | | | | | | | | | |
|---|---|---|---|---|---|---|---|---|---|---|---|---|---|---|
| 939 | 1091 | 1225 | 858 | 1207 | 1277 | 949 | 1193 | 5264 | 3772 | 4015 | 4380 | 3440 | 2410 | 2809 |
| 596 | 791 | 990 | 560 | 905 | 953 | 563 | 925 | 6191 | 3427 | 5753 | 4553 | 13431 | 7603 | 13572 |
| 266 | 765 | 451 | 258 | 758 | 458 | 214 | 630 | 6435 | 3599 | 13524 | 10341 | 7676 | 4393 | 17031 |
| 415 | 658 | 692 | 466 | 807 | 639 | 407 | 784 | 13779 | 7396 | 17838 | 11732 | 5794 | 3899 | 8373 |
| 411 | 741 | 942 | 594 | 887 | 1019 | 702 | 1007 | 4993 | 2618 | 6368 | 4721 | 3883 | 2450 | 4123 |
| 458 | 771 | 689 | 296 | 573 | 763 | 522 | 905 | 12442 | 8072 | 17610 | 13574 | 7836 | 3367 | 11374 |
| 676 | 1028 | 944 | 642 | 1023 | 788 | 504 | 868 | 17377 | 12380 | 18320 | 18831 | 5164 | 3512 | 5470 |
| 692 | 1135 | 754 | 746 | 1164 | 697 | 490 | 793 | 8348 | 7512 | 10861 | 12333 | 4271 | 4225 | 5666 |
| 754 | 1254 | 917 | 689 | 1369 | 910 | 731 | 1247 | 8393 | 6277 | 8327 | 10443 | 5910 | 4440 | 6446 |
| 490 | 815 | 591 | 516 | 822 | 583 | 486 | 836 | 13340 | 11040 | 22550 | 18382 | 5219 | 4565 | 8839 |
| 371 | 751 | 550 | 426 | 855 | 390 | 278 | 670 | 4040 | 3074 | 8284 | 6219 | 1595 | 1235 | 2899 |
| 401 | 614 | 448 | 384 | 619 | 551 | 429 | 664 | 6028 | 4238 | 10575 | 6489 | 1998 | 1712 | 4457 |
| 844 | 1233 | 847 | 837 | 1420 | 879 | 845 | 1329 | 17038 | 15599 | 18490 | 22806 | 4170 | 4121 | 4925 |
| 440 | 905 | 761 | 585 | 988 | 695 | 548 | 1014 | 7752 | 5458 | 12406 | 11222 | 5765 | 4428 | 7573 |
| 942 | 1459 | 870 | 837 | 1429 | 904 | 740 | 1121 | 1337 | 1391 | 1478 | 2156 | 3259 | 3136 | 3745 |
| 382 | 797 | 529 | 304 | 731 | 478 | 218 | 705 | 1115 | 775 | 2030 | 1617 | 3709 | 2132 | 7013 |

| | | | | | | | | | | | | | | |
|---|---|---|---|---|---|---|---|---|---|---|---|---|---|---|
| | | 710 | 687 | 771 | 855 | 922 | 940 | | | | | 59 | 57 | 83 |
| 789 | 1216 | 1224 | 1041 | 1531 | 1222 | 885 | 1243 | 9123 | 6733 | 8532 | 10374 | 4556 | 3877 | 3723 |
| 946 | 1249 | 1461 | 1236 | 1382 | 1041 | 848 | 1343 | 7267 | 5904 | 6239 | 7790 | 1410 | 1194 | 966 |
| 849 | 1145 | 1017 | 731 | 1198 | 1038 | 812 | 1151 | 8400 | 6475 | 7631 | 8737 | 7477 | 5371 | 7348 |
| 606 | 923 | 544 | 334 | 670 | 433 | 271 | 614 | 2178 | 1587 | 2620 | 2420 | 7421 | 4563 | 13644 |
| 761 | 1286 | 829 | 721 | 1282 | 767 | 632 | 1243 | 4013 | 3016 | 3964 | 5097 | 7090 | 6159 | 8548 |
| 744 | 1097 | 540 | 404 | 634 | 875 | 570 | 961 | 5594 | 4754 | 6388 | 7007 | 1565 | 1169 | 2896 |
| 576 | 876 | 814 | 563 | 865 | 865 | 657 | 929 | 8187 | 5542 | 9624 | 8427 | 14036 | 9704 | 17247 |
| 941 | 1093 | 736 | 733 | 910 | 597 | 563 | 730 | 8086 | 7274 | 7732 | 8447 | 6401 | 6375 | 8692 |
| 504 | 932 | 696 | 507 | 911 | 843 | 690 | 1131 | 10948 | 7870 | 15619 | 14559 | 2430 | 1770 | 3489 |
| 591 | 921 | 693 | 466 | 805 | 640 | 419 | 850 | 13429 | 10345 | 17499 | 16113 | 6122 | 4116 | 8830 |
| 318 | 561 | 693 | 318 | 553 | 674 | 342 | 572 | 9133 | 4283 | 13490 | 7565 | 10909 | 5010 | 15736 |
| 449 | 754 | 922 | 473 | 732 | | | | 4688 | 2422 | 5391 | 4064 | 8920 | 4578 | 9677 |
| 796 | 1102 | 816 | 753 | 1003 | 607 | 908 | 1082 | 7022 | 6582 | 8269 | 9114 | 15255 | 14076 | 18690 |
| 620 | 1067 | 844 | 712 | 1160 | 866 | 662 | 1092 | 9734 | 7064 | 11388 | 12149 | 5145 | 4343 | 6097 |
| 1043 | 1462 | 1090 | 1052 | 1422 | 994 | 1011 | 1491 | 5113 | 5652 | 5418 | 7921 | 3028 | 2921 | 2777 |
| 696 | 900 | 649 | 517 | 800 | 825 | 738 | 906 | 3629 | 2967 | 4262 | 3834 | 5998 | 4784 | 9245 |
| 535 | 1158 | 821 | 509 | 909 | 1012 | 733 | 1257 | 4879 | 3297 | 6164 | 7135 | 5430 | 3370 | 6616 |
| 559 | 927 | 542 | 439 | 845 | 536 | 498 | 861 | 12859 | 11730 | 20966 | 19443 | 13164 | 10668 | 24284 |
| 552 | 907 | 693 | 490 | 824 | 712 | 578 | 856 | 7491 | 5262 | 9533 | 8649 | 1790 | 1264 | 2582 |
| 432 | 780 | 759 | 532 | 821 | 718 | 511 | 817 | 2981 | 1773 | 4106 | 3203 | 3705 | 2598 | 4884 |
| 901 | 1402 | 715 | 887 | 1491 | 770 | 754 | 1179 | 6260 | 7198 | 7990 | 11203 | 3421 | 4241 | 4784 |
| | | 921 | 717 | 1032 | 967 | 794 | 1265 | | | | | 13762 | 10713 | 14937 |
| 732 | 6812 | 631 | 91 | 1713 | 1235 | 859 | 13493 | 5937 | 3822 | 5220 | 35562 | 4318 | 624 | 6843 |
| 785 | 946 | | | | 808 | 644 | 986 | 78 | 61 | 77 | 73 | | | |
| 899 | 1145 | 1416 | 1258 | 1477 | 1238 | 1034 | 1216 | 9213 | 7620 | 8478 | 9707 | 16231 | 14424 | 11465 |
| 352 | 688 | 540 | 439 | 794 | 438 | 385 | 693 | 11800 | 8353 | 23702 | 16309 | 4040 | 3282 | 7483 |
| 805 | 1069 | 1132 | 1131 | 1208 | 1365 | 1490 | 1218 | 6957 | 6040 | 7505 | 8022 | 5075 | 5071 | 4485 |
| 756 | 1146 | 921 | 722 | 1065 | 936 | 597 | 940 | 5175 | 3817 | 5047 | 5786 | 4007 | 3141 | 4353 |
| 558 | 1054 | 951 | 687 | 1128 | 656 | 448 | 1045 | 3761 | 2776 | 4972 | 5241 | 3520 | 2542 | 3700 |
| | | 900 | 782 | 1243 | 823 | 682 | 1041 | | | | | 3983 | 3462 | 4428 |

| | | | | | | | | | | | | | | |
|---|---|---|---|---|---|---|---|---|---|---|---|---|---|---|
| 385 | 776 | 383 | 270 | 698 | 393 | 301 | 750 | 5034 | 3695 | 9586 | 7436 | 2193 | 1547 | 5721 |
| 419 | 752 | 633 | 448 | 829 | 629 | 454 | 895 | 7320 | 3899 | 9299 | 6995 | 8365 | 5919 | 13216 |
| 791 | 1112 | 814 | 635 | 1008 | 676 | 592 | 966 | 11407 | 8897 | 11248 | 12502 | 5602 | 4372 | 6883 |
| 422 | 1008 | 575 | 430 | 892 | 573 | 434 | 1058 | 4966 | 3575 | 8467 | 8533 | 8390 | 6281 | 14601 |
| 971 | 876 | 818 | 880 | 933 | 917 | 772 | 928 | 3148 | 4728 | 4871 | 4266 | 9569 | 10289 | 11695 |
| 769 | 819 | 650 | 556 | 712 | 553 | 502 | 593 | 7945 | 8031 | 10442 | 8556 | 6771 | 5796 | 10417 |
| 783 | 1119 | 909 | 782 | 1131 | 963 | 768 | 1107 | 7137 | 5865 | 7486 | 8376 | 8339 | 7179 | 9178 |
| 833 | 1347 | 1156 | 1048 | 1565 | 1400 | 971 | 1593 | 2964 | 2508 | 3010 | 4055 | 2914 | 2640 | 2520 |
| 756 | 907 | 984 | 732 | 900 | | | | 6173 | 4548 | 6019 | 5459 | 6973 | 5187 | 7085 |
| 1022 | 1030 | 971 | 700 | 961 | 1179 | 1099 | 1163 | 553 | 517 | 506 | 521 | 351 | 253 | 362 |
| 856 | 1433 | 1129 | 869 | 1367 | 1179 | 949 | 1490 | 6613 | 5002 | 5842 | 8371 | 3170 | 2441 | 2808 |
| 882 | 1107 | 1043 | 1392 | 991 | 999 | 1087 | 971 | 2547 | 2656 | 3012 | 3332 | 4126 | 5505 | 3954 |
| 453 | 813 | 643 | 511 | 830 | 759 | 611 | 962 | 3874 | 2829 | 6249 | 5079 | 6982 | 5552 | 10862 |
| 633 | 840 | 1056 | 641 | 846 | 1173 | 755 | 881 | 7642 | 4227 | 6679 | 5610 | 7528 | 4570 | 7129 |
| 668 | 957 | | | | 1007 | 618 | 922 | 6590 | 4507 | 6750 | 6460 | | | |
| 263 | 526 | 619 | 296 | 612 | 507 | 354 | 641 | 3601 | 2059 | 7815 | 4113 | 8034 | 3837 | 12973 |
| 612 | 855 | 690 | 485 | 881 | 699 | 564 | 920 | 5837 | 4542 | 7416 | 6338 | 1625 | 1142 | 2355 |
| 637 | 1060 | 991 | 743 | 888 | 955 | 734 | 793 | 11512 | 9014 | 14143 | 14991 | 3845 | 2882 | 3880 |
| 803 | 1153 | 997 | 880 | 1154 | 918 | 762 | 1061 | 8416 | 6973 | 8688 | 10017 | 4719 | 4165 | 4734 |
| 536 | 670 | 372 | 390 | 552 | 342 | 378 | 533 | 5726 | 5866 | 10941 | 7327 | 2119 | 2223 | 5700 |
| 351 | 701 | 525 | 444 | 840 | 456 | 345 | 682 | 3128 | 2506 | 7135 | 4999 | 2861 | 2425 | 5455 |
| 551 | 816 | 836 | 492 | 826 | 827 | 499 | 844 | 3876 | 2875 | 5217 | 4258 | 5721 | 3367 | 6844 |
| 1016 | 1510 | 929 | 1172 | 1738 | 856 | 995 | 1384 | 6412 | 8215 | 8087 | 12213 | 4582 | 5778 | 4931 |
| 671 | 1123 | 978 | 705 | 1123 | 973 | 629 | 1053 | 2124 | 1714 | 2556 | 2869 | 2598 | 1874 | 2657 |
| 882 | 1100 | 1408 | 1200 | 1345 | 907 | 630 | 999 | 9619 | 7889 | 8949 | 9840 | 2709 | 2309 | 1924 |
| 627 | 919 | 857 | 561 | 911 | 809 | 641 | 1041 | 4592 | 3136 | 5001 | 4597 | 4360 | 2853 | 5086 |
| 190 | 536 | 708 | 494 | 853 | 386 | 227 | 679 | 1523 | 929 | 4882 | 2618 | 6685 | 4666 | 9439 |
| 884 | 1446 | 937 | 796 | 1181 | 988 | 823 | 1185 | 1288 | 1245 | 1408 | 2037 | 4328 | 3675 | 4618 |
| 392 | 766 | 617 | 403 | 796 | 628 | 380 | 717 | 1729 | 1097 | 2799 | 2145 | 2458 | 1605 | 3983 |
| 1133 | 1199 | 1184 | 1072 | 1003 | 1134 | 1086 | 1106 | 3541 | 3747 | 3308 | 3965 | 5914 | 5356 | 4994 |
| 978 | 1462 | 962 | 865 | 1332 | 1027 | 870 | 1267 | 3462 | 3183 | 3256 | 4760 | 10235 | 9209 | 10642 |

| | | | | | | | | | | | | | | |
|---|---|---|---|---|---|---|---|---|---|---|---|---|---|---|
| 494 | 930 | 533 | 395 | 802 | 676 | 510 | 901 | 9293 | 6953 | 14067 | 13079 | 4594 | 3407 | 8622 |
| 698 | 865 | 1006 | 674 | 865 | 978 | 626 | 791 | 11394 | 7083 | 10151 | 8779 | 8590 | 5751 | 8536 |
| 597 | 851 | 870 | 782 | 1037 | 874 | 734 | 1080 | 6965 | 5065 | 8477 | 7210 | 5593 | 5026 | 6428 |
| 542 | 882 | 567 | 494 | 837 | 442 | 415 | 763 | 1953 | 1902 | 3510 | 3096 | 985 | 859 | 1738 |
| 788 | 1160 | 1153 | 899 | 1221 | | | | 2636 | 2064 | 2619 | 3037 | 2052 | 1599 | 1779 |
| 371 | 712 | 712 | 397 | 736 | 628 | 345 | 683 | 3400 | 1884 | 5081 | 3616 | 5224 | 2911 | 7337 |
| 776 | 998 | 878 | 777 | 923 | 922 | 804 | 899 | 3173 | 2634 | 3396 | 3390 | 3323 | 2943 | 3786 |
| 411 | 680 | 863 | 538 | 823 | 937 | 612 | 916 | 5115 | 2966 | 7207 | 4900 | 6855 | 4276 | 7946 |
| 769 | 1075 | 975 | 697 | 1051 | 971 | 707 | 1069 | 7099 | 5483 | 7134 | 7668 | 8252 | 5899 | 8460 |
| 881 | 931 | 1102 | 389 | 609 | 1000 | 653 | 859 | 6586 | 5061 | 5747 | 5353 | 7187 | 2535 | 6520 |
| 652 | 1057 | 936 | 654 | 1161 | 861 | 683 | 1018 | 10122 | 8069 | 12380 | 13092 | 4732 | 3307 | 5057 |
| 929 | 1364 | 1156 | 1451 | 1953 | 1119 | 1608 | 2089 | 5828 | 5876 | 6322 | 8622 | 4715 | 5919 | 4078 |
| 532 | 929 | 892 | 805 | 1234 | 798 | 708 | 1074 | 2346 | 1669 | 3139 | 2915 | 7807 | 7041 | 8750 |
| 447 | 937 | 708 | 448 | 846 | 632 | 429 | 858 | 4324 | 2795 | 6247 | 5856 | 4120 | 2609 | 5821 |
| 1044 | 1049 | 984 | 760 | 866 | 1348 | 1303 | 1131 | 5087 | 4161 | 3984 | 4179 | 4099 | 3166 | 4166 |
| 377 | 810 | 505 | 365 | 797 | 659 | 465 | 822 | 7459 | 5630 | 14954 | 12118 | 7432 | 5367 | 14708 |
| 344 | 755 | 500 | 315 | 741 | 599 | 388 | 794 | 7391 | 4904 | 14256 | 10765 | 3468 | 2184 | 6932 |
| 767 | 1037 | 1299 | 664 | 1091 | 801 | 680 | 977 | 6184 | 4880 | 6359 | 6595 | 3604 | 1842 | 2775 |
| 763 | 1185 | 845 | 852 | 1314 | 755 | 851 | 1236 | 2101 | 1994 | 2615 | 3098 | 7759 | 7823 | 9187 |
| 784 | 1134 | 965 | 710 | 1142 | 886 | 691 | 1075 | 11173 | 8586 | 10947 | 12413 | 10077 | 7413 | 10447 |
| 645 | 1030 | 1084 | 893 | 1239 | 978 | 739 | 1197 | 6698 | 5087 | 7889 | 8125 | 4797 | 3952 | 4427 |
| 1647 | 1784 | 1342 | 1396 | 1376 | 1459 | 1403 | 1405 | 819 | 824 | 500 | 892 | 1904 | 1981 | 1419 |
| 823 | 1108 | 1002 | 812 | 1060 | 1012 | 786 | 1050 | 7760 | 5968 | 7249 | 8032 | 11422 | 9257 | 11402 |
| 473 | 899 | 670 | 452 | 906 | 760 | 575 | 978 | 3027 | 2090 | 4417 | 3973 | 5397 | 3646 | 8060 |
| 621 | 976 | 1016 | 879 | 1361 | 997 | 949 | 1196 | 1570 | 1159 | 1868 | 1824 | 7236 | 6261 | 7120 |
| 698 | 938 | 919 | 729 | 1062 | 832 | 678 | 896 | 4403 | 4044 | 5797 | 5438 | 6819 | 5414 | 7423 |
| 1061 | 1450 | 1436 | 1087 | 1462 | 1156 | 665 | 1288 | 7337 | 5658 | 5333 | 7734 | 7337 | 5554 | 5108 |
| 392 | 706 | 614 | 414 | 716 | 412 | 409 | 598 | 6142 | 4103 | 10477 | 7400 | 6426 | 4337 | 10469 |
| 993 | 756 | 886 | 839 | 872 | 961 | 840 | 856 | 6791 | 6833 | 6878 | 5201 | 5771 | 5469 | 6516 |
| | | 425 | 473 | 1178 | | | | | | | | 1340 | 1492 | 3154 |
| 566 | 861 | 937 | 575 | 868 | 711 | 461 | 765 | 7836 | 4782 | 8448 | 7275 | 9086 | 5578 | 9702 |

| | | | | | | | | | | | | | | |
|---|---|---|---|---|---|---|---|---|---|---|---|---|---|---|
| 574 | 966 | 941 | 608 | 1086 | 1024 | 656 | 995 | 6702 | 4623 | 8050 | 7778 | 2228 | 1440 | 2368 |
| 419 | 662 | 679 | 427 | 777 | 719 | 385 | 627 | 6976 | 4621 | 11021 | 7295 | 5416 | 3408 | 7980 |
| 764 | 1117 | 887 | 811 | 1095 | 946 | 833 | 1108 | 10498 | 8785 | 11492 | 12836 | 6818 | 6236 | 7686 |
| 1149 | 1417 | 947 | 894 | 1179 | 1045 | 1153 | 1456 | 6600 | 7601 | 6617 | 9376 | 8768 | 8282 | 9263 |
| 619 | 1226 | 572 | 426 | 1021 | 631 | 474 | 1039 | 4412 | 3397 | 5486 | 6724 | 4320 | 3219 | 7557 |
| 395 | 803 | | | | | | | 3812 | 2394 | 6064 | 4868 | | | |
| 383 | 815 | 687 | 484 | 861 | 562 | 380 | 761 | 4414 | 3330 | 8704 | 7091 | 1583 | 1115 | 2305 |
| 2149 | 2329 | 831 | 631 | 960 | 779 | 514 | 734 | 777 | 2369 | 1103 | 2567 | 3105 | 2357 | 3739 |
| 1492 | 1779 | 1500 | 1471 | 1671 | 1196 | 1149 | 1322 | 3824 | 3821 | 2561 | 4556 | 4385 | 4299 | 2923 |
| 729 | 1164 | 784 | 558 | 1005 | 1010 | 797 | 1152 | 2693 | 2082 | 2857 | 3326 | 2819 | 2005 | 3595 |
| 741 | 1255 | 909 | 937 | 1334 | 869 | 836 | 1463 | 1356 | 1208 | 1630 | 2046 | 5573 | 5745 | 6130 |
| 493 | 1050 | 454 | 514 | 935 | 520 | 429 | 772 | 2265 | 2749 | 5582 | 5863 | 3640 | 4122 | 8013 |
| 272 | 589 | 550 | 322 | 607 | 432 | 223 | 494 | 3536 | 2130 | 7845 | 4618 | 11005 | 6436 | 20013 |
| 632 | 865 | 797 | 841 | 1079 | 940 | 935 | 1069 | 3086 | 2592 | 4103 | 3550 | 6939 | 7322 | 8707 |
| 405 | 1415 | 597 | 453 | 1351 | 759 | 636 | 1421 | 2048 | 1337 | 3297 | 4664 | 2215 | 1681 | 3710 |
| 842 | 1172 | 997 | 795 | 1072 | 933 | 757 | 1040 | 5169 | 4852 | 5764 | 6757 | 2163 | 1725 | 2170 |
| 912 | 1184 | 989 | 831 | 1006 | 1006 | 916 | 1159 | 8007 | 6430 | 7051 | 8348 | 1887 | 1585 | 1909 |
| 672 | 1100 | 858 | 689 | 1009 | 794 | 595 | 1028 | 6261 | 4925 | 7326 | 8061 | 4473 | 3592 | 5214 |
| 1525 | 1976 | 1569 | 1595 | 2014 | 1101 | 994 | 1360 | 3082 | 3421 | 2243 | 4432 | 7371 | 7494 | 4697 |
| 327 | 800 | 476 | 275 | 807 | 577 | 370 | 851 | 2441 | 1754 | 5368 | 4297 | 9252 | 5353 | 19451 |
| 1255 | 1499 | 1375 | 974 | 1225 | 872 | 665 | 1028 | 4188 | 4187 | 3335 | 4998 | 3623 | 2567 | 2635 |
| 306 | 713 | 478 | 234 | 641 | 615 | 355 | 638 | 7553 | 4112 | 13416 | 9572 | 5089 | 2496 | 10648 |
| 989 | 624 | 1543 | 1599 | 1026 | 1124 | 934 | 994 | 2000 | 1569 | 1586 | 990 | 495 | 513 | 321 |
| 516 | 936 | 713 | 513 | 1092 | 775 | 606 | 1111 | 4078 | 3148 | 6101 | 5712 | 4771 | 3436 | 6694 |
| 337 | 680 | 474 | 370 | 727 | 492 | 373 | 719 | 6706 | 4352 | 12906 | 8772 | 3691 | 2883 | 7788 |
| 429 | 771 | 592 | 400 | 706 | 663 | 466 | 791 | 14400 | 9557 | 22261 | 17168 | 5087 | 3433 | 8588 |
| 422 | 850 | 486 | 369 | 723 | 566 | 450 | 837 | 1872 | 1579 | 3739 | 3177 | 5501 | 4177 | 11326 |
| 624 | 842 | 1350 | 658 | 866 | 1399 | 649 | 907 | 6109 | 2686 | 4307 | 3625 | 7946 | 3875 | 5885 |
| 217 | 666 | 674 | 426 | 745 | 780 | 365 | 716 | 12386 | 5944 | 27369 | 18214 | 8268 | 5224 | 12259 |
| 326 | 470 | 543 | 428 | 567 | 577 | 455 | 718 | 7969 | 5988 | 18385 | 8637 | 3085 | 2434 | 5681 |
| 599 | 1033 | 828 | 661 | 1166 | 725 | 509 | 753 | 1586 | 1300 | 2169 | 2241 | 16477 | 13164 | 19911 |

| | | | | | | | | | | | | | | |
|---|---|---|---|---|---|---|---|---|---|---|---|---|---|---|
| 481 | 633 | 679 | 420 | 569 | 623 | 309 | 520 | 7333 | 4780 | 9931 | 6288 | 6398 | 3958 | 9422 |
| 439 | 782 | 688 | 425 | 780 | 743 | 405 | 794 | 12243 | 6080 | 13850 | 10826 | 4052 | 2505 | 5893 |
| 1138 | 1320 | 1213 | 1374 | 1507 | 1464 | 1432 | 1287 | 2575 | 2459 | 2161 | 2852 | 1195 | 1354 | 986 |
| 915 | 1289 | 725 | 851 | 1274 | 730 | 799 | 1199 | 7601 | 7691 | 8405 | 10836 | 2342 | 2751 | 3231 |
| 382 | 762 | 576 | 406 | 795 | 527 | 392 | 723 | 2493 | 1743 | 4568 | 3482 | 2878 | 2028 | 5000 |
| 772 | 1136 | 1131 | 1027 | 1379 | 1530 | 1038 | 1209 | 7085 | 5699 | 7380 | 8383 | 9540 | 8666 | 8435 |
| 641 | 886 | 669 | 429 | 756 | 666 | 442 | 737 | 7963 | 6031 | 9413 | 8343 | 9380 | 6018 | 14023 |
| 521 | 1010 | 991 | 584 | 960 | 884 | 604 | 924 | 9187 | 5624 | 10788 | 10901 | 9541 | 5624 | 9628 |
| 587 | 977 | 745 | 655 | 1058 | 771 | 527 | 919 | 8014 | 5818 | 9916 | 9689 | 3161 | 2777 | 4242 |
| 788 | 1074 | 1143 | 1027 | 1445 | 1071 | 934 | 1309 | 8585 | 6063 | 7697 | 8266 | 5523 | 4962 | 4832 |
| 372 | 735 | 524 | 302 | 671 | 504 | 320 | 649 | 12218 | 9327 | 25069 | 18415 | 8101 | 4678 | 15471 |
| 1164 | 1565 | 1160 | 1116 | 1528 | 830 | 763 | 1212 | 3423 | 3182 | 2733 | 4278 | 4115 | 3958 | 3547 |
| 551 | 795 | 917 | 664 | 874 | 600 | 324 | 622 | 7400 | 5283 | 9583 | 7620 | 3319 | 2402 | 3620 |
| 406 | 722 | 1019 | 775 | 1184 | 668 | 458 | 780 | 5805 | 3460 | 8529 | 6156 | 2730 | 2076 | 2678 |
| 1492 | 2000 | 1217 | 1417 | 1911 | 718 | 838 | 1114 | 2810 | 3291 | 2205 | 4410 | 2748 | 3198 | 2257 |
| 716 | 1146 | 812 | 622 | 1061 | 830 | 632 | 1007 | 3540 | 2892 | 4039 | 4627 | 4692 | 3594 | 5780 |
| | | 691 | 431 | 810 | 798 | 670 | 1018 | | | | | 3935 | 2450 | 5691 |
| 751 | 953 | 977 | 751 | 981 | 994 | 916 | 1130 | 1885 | 1869 | 2488 | 2371 | 2888 | 2219 | 2957 |
| 846 | 1459 | 606 | 436 | 870 | 623 | 484 | 780 | 1132 | 1028 | 1215 | 1773 | 2061 | 1485 | 3403 |
| 476 | 785 | 824 | 691 | 1094 | 623 | 354 | 589 | 10671 | 6479 | 13612 | 10686 | 3824 | 3209 | 4642 |
| 472 | 689 | 775 | 666 | 830 | 609 | 438 | 702 | 5145 | 4038 | 8560 | 5897 | 3746 | 3219 | 4832 |
| 136 | 511 | | | | 240 | 127 | 564 | 966 | 487 | 3579 | 1831 | | | |
| 284 | 628 | 633 | 447 | 830 | 393 | 248 | 649 | 8604 | 4995 | 17587 | 11049 | 7401 | 5223 | 11685 |
| 397 | 649 | 570 | 394 | 655 | 536 | 351 | 643 | 9024 | 5956 | 15022 | 9742 | 7243 | 5005 | 12717 |
| 911 | 1097 | 1095 | 1022 | 1245 | 1071 | 841 | 1232 | 2173 | 2127 | 2336 | 2564 | 2214 | 2066 | 2022 |
| 515 | 817 | 604 | 425 | 748 | 675 | 530 | 842 | 5402 | 4503 | 8735 | 7132 | 4473 | 3151 | 7411 |
| 683 | 623 | 387 | 1176 | 1074 | 513 | 406 | 648 | 5302 | 3896 | 5704 | 3553 | 2412 | 7325 | 6230 |
| 687 | 1159 | 997 | 804 | 1147 | 1157 | 847 | 1215 | 4809 | 3701 | 5386 | 6244 | 6392 | 5155 | 6412 |
| | | 870 | 766 | 1079 | 724 | 346 | 716 | | | | | 1902 | 1675 | 2187 |
| 323 | 700 | 659 | 398 | 692 | 666 | 396 | 752 | 8324 | 4448 | 13761 | 9627 | 5922 | 3574 | 8986 |
| 922 | 1648 | | | | 879 | 886 | 1518 | 1758 | 1810 | 1963 | 3235 | | | |

| | | | | | | | | | | | | | | |
|---|---|---|---|---|---|---|---|---|---|---|---|---|---|---|
| 951 | 1673 | 1202 | 939 | 1414 | | | | 7103 | 6080 | 6391 | 10694 | 3178 | 2483 | 2644 |
| 752 | 1358 | 774 | 675 | 1167 | | | | 3801 | 3378 | 4490 | 6098 | 9128 | 7960 | 11786 |
| 424 | 696 | 633 | 475 | 712 | 888 | 670 | 939 | 5688 | 4011 | 9461 | 6584 | 3537 | 2657 | 5592 |
| 228 | 480 | 309 | 203 | 485 | 546 | 435 | 740 | 2388 | 1518 | 6670 | 3201 | 3954 | 2597 | 12808 |
| 624 | 895 | | | | 863 | 536 | 759 | 5766 | 3921 | 6284 | 5624 | | | |
| 391 | 824 | 806 | 695 | 1021 | 672 | 372 | 797 | 3434 | 2428 | 6213 | 5121 | 6963 | 6005 | 8642 |
| 379 | 898 | | | | 724 | 501 | 923 | 3361 | 1997 | 5268 | 4733 | | | |
| 354 | 715 | 758 | 515 | 820 | 544 | 367 | 705 | 6435 | 4027 | 11388 | 8144 | 14334 | 9743 | 18917 |
| 500 | 762 | 795 | 542 | 676 | 627 | 365 | 646 | 12970 | 8920 | 17827 | 13585 | 2596 | 1770 | 3266 |
| 323 | 617 | 497 | 321 | 568 | 699 | 426 | 617 | 992 | 616 | 1908 | 1177 | 6520 | 4220 | 13130 |
| 644 | 1166 | 917 | 556 | 946 | 737 | 610 | 1126 | 4660 | 3879 | 6025 | 7022 | 4318 | 2619 | 4711 |
| 541 | 949 | 776 | 691 | 1005 | 769 | 590 | 1011 | 7122 | 5500 | 10161 | 9638 | 4513 | 4022 | 5818 |
| 278 | 562 | 456 | 383 | 591 | 421 | 305 | 583 | 2442 | 1347 | 4846 | 2725 | 2399 | 2010 | 5255 |
| 493 | 726 | 668 | 548 | 839 | 595 | 396 | 755 | 9051 | 6073 | 12316 | 8947 | 5335 | 4380 | 7986 |
| 1079 | 2525 | 908 | 615 | 1570 | 857 | 600 | 1375 | 852 | 792 | 734 | 1853 | 9264 | 6277 | 10203 |
| 558 | 791 | 614 | 455 | 620 | 676 | 488 | 750 | 903 | 666 | 1195 | 946 | 2234 | 1655 | 3638 |
| 306 | 594 | 846 | 731 | 1066 | 991 | 829 | 1224 | 8022 | 3317 | 10854 | 6445 | 2218 | 1918 | 2623 |
| 618 | 935 | 746 | 796 | 1003 | 492 | 372 | 736 | 7642 | 6649 | 10753 | 10049 | 5102 | 5446 | 6841 |
| 389 | 627 | 711 | 344 | 726 | 895 | 501 | 821 | 3897 | 1930 | 4964 | 3111 | 5815 | 2812 | 8179 |
| 1204 | 1243 | | | | 1803 | 1975 | 1487 | 4822 | 4390 | 3646 | 4531 | | | |
| 508 | 606 | 1048 | 506 | 646 | 1095 | 514 | 651 | 12887 | 6902 | 13583 | 8225 | 5005 | 2413 | 4774 |
| 651 | 1065 | 874 | 714 | 1182 | 938 | 778 | 1063 | 4980 | 4149 | 6370 | 6786 | 8093 | 6613 | 9261 |
| 422 | 787 | 874 | 435 | 716 | 1156 | 627 | 905 | 3128 | 1054 | 2497 | 1966 | 749 | 373 | 857 |
| 590 | 893 | 687 | 430 | 752 | 871 | 413 | 673 | 7635 | 5643 | 9569 | 8543 | 2643 | 1656 | 3848 |
| 503 | 905 | 796 | 664 | 1088 | 840 | 734 | 1082 | 5691 | 4651 | 9247 | 8365 | 6398 | 5339 | 8041 |
| 545 | 759 | 1011 | 800 | 1006 | 984 | 772 | 895 | 6169 | 3021 | 5539 | 4203 | 1615 | 1278 | 1598 |
| | | 561 | 467 | 777 | 585 | 429 | 704 | | | | | 3191 | 2656 | 5685 |
| 417 | 809 | 827 | 498 | 790 | 771 | 670 | 858 | 3648 | 1810 | 4343 | 3514 | 6317 | 3804 | 7639 |
| 475 | 1795 | 591 | 368 | 1683 | 622 | 386 | 1518 | 1709 | 1170 | 2464 | 4422 | 5631 | 3509 | 9529 |
| 528 | 961 | 716 | 558 | 908 | 856 | 688 | 1025 | 8041 | 6678 | 12653 | 12158 | 8631 | 6730 | 12053 |
| 616 | 1038 | 871 | 640 | 1009 | 714 | 507 | 967 | 11725 | 8912 | 14476 | 15023 | 4321 | 3176 | 4963 |

| | | | | | | | | | | | | | | |
|---|---|---|---|---|---|---|---|---|---|---|---|---|---|---|
| 571 | 901 | 855 | 501 | 776 | | | | 3959 | 2595 | 4547 | 4097 | 3525 | 2067 | 4122 |
| | | 1251 | 804 | 1174 | | | | | | | | 2509 | 1613 | 2005 |
| 410 | 798 | 480 | 357 | 902 | 612 | 457 | 960 | 821 | 719 | 1752 | 1397 | 1536 | 1144 | 3202 |
| 456 | 887 | 826 | 562 | 881 | 777 | 512 | 932 | 963 | 613 | 1343 | 1191 | 1701 | 1158 | 2059 |
| 929 | 1043 | 1418 | 1172 | 1387 | | | | 3584 | 2731 | 2939 | 3065 | 5254 | 4343 | 3705 |
| 640 | 907 | 1014 | 748 | 1085 | 979 | 730 | 955 | 4756 | 3596 | 5617 | 5093 | 1710 | 1261 | 1687 |
| 615 | 1031 | 971 | 603 | 940 | 827 | 561 | 909 | 9136 | 6515 | 10593 | 10916 | 6433 | 3999 | 6628 |
| 819 | 957 | 1474 | 942 | 925 | 702 | 482 | 835 | 12811 | 9075 | 11081 | 10607 | 4345 | 2775 | 2947 |
| 800 | 1173 | 1025 | 1013 | 1316 | 1687 | 1662 | 1770 | 2573 | 2348 | 2935 | 3443 | 5095 | 5037 | 4972 |
| 548 | 589 | 254 | 582 | 491 | 262 | 598 | 522 | 3343 | 5607 | 10230 | 6022 | 1744 | 3999 | 6875 |
| 879 | 1018 | 823 | 467 | 736 | 730 | 427 | 662 | 3385 | 2925 | 3328 | 3386 | 3540 | 2008 | 4299 |
| 938 | 1294 | | | | 1094 | 1032 | 1114 | 2539 | 2331 | 2487 | 3218 | | | |
| 509 | 810 | 664 | 503 | 818 | 512 | 377 | 739 | 3236 | 2887 | 5674 | 4595 | 8571 | 6489 | 12911 |
| 774 | 999 | 879 | 944 | 1057 | 837 | 773 | 1064 | 9274 | 8719 | 11266 | 11255 | 934 | 1003 | 1063 |
| 389 | 665 | 571 | 433 | 638 | 523 | 368 | 617 | 5694 | 3985 | 10256 | 6819 | 2997 | 2276 | 5252 |
| 978 | 1209 | 864 | 736 | 1127 | 895 | 870 | 1275 | 8981 | 8151 | 8332 | 10072 | 5246 | 4471 | 6074 |
| 358 | 860 | 477 | 295 | 782 | 586 | 332 | 798 | 3073 | 1936 | 5408 | 4649 | 4135 | 2561 | 8669 |
| 452 | 814 | 817 | 350 | 824 | 911 | 569 | 960 | 5619 | 2952 | 6527 | 5313 | 2476 | 1061 | 3031 |
| 932 | 1386 | 937 | 803 | 1108 | | | | 13948 | 12196 | 13087 | 18137 | 3040 | 2605 | 3244 |
| 661 | 985 | 1000 | 689 | 992 | 1044 | 723 | 1099 | 9423 | 5991 | 9061 | 8926 | 13227 | 9110 | 13230 |
| 518 | 1154 | 758 | 514 | 852 | 717 | 570 | 909 | 3802 | 2563 | 4952 | 5713 | 9679 | 6560 | 12763 |
| 404 | 878 | | | | 630 | 531 | 998 | 2322 | 1596 | 3948 | 3465 | | | |
| 859 | 1206 | 958 | 969 | 1418 | 931 | 921 | 1315 | 3521 | 2593 | 3020 | 3641 | 2064 | 2089 | 2155 |
| 200 | 640 | 497 | 314 | 756 | 447 | 245 | 657 | 1588 | 794 | 3969 | 2540 | 4775 | 3016 | 9610 |
| 585 | 844 | 634 | 598 | 849 | 630 | 634 | 929 | 942 | 808 | 1382 | 1166 | 2847 | 2687 | 4493 |
| 376 | 636 | 535 | 441 | 696 | 623 | 553 | 785 | 1899 | 1745 | 4645 | 2955 | 5521 | 4551 | 10313 |
| 616 | 806 | 1180 | 818 | 908 | 929 | 625 | 836 | 5006 | 2975 | 4826 | 3888 | 11382 | 7891 | 9642 |
| 537 | 804 | | | | 1136 | 904 | 1064 | 14073 | 8736 | 16257 | 13075 | | | |
| 657 | 960 | 1497 | 1322 | 1469 | 1133 | 951 | 1234 | 8009 | 6254 | 9513 | 9129 | 5157 | 4552 | 3444 |
| 461 | 763 | | | | 526 | 289 | 670 | 5087 | 3252 | 7054 | 5379 | | | |
| 671 | 1174 | 977 | 864 | 1267 | 679 | 602 | 1175 | 4950 | 4371 | 6514 | 7645 | 5200 | 4602 | 5324 |

| | | | | | | | | | | | | | | |
|---|---|---|---|---|---|---|---|---|---|---|---|---|---|---|
| 365 | 716 | 612 | 432 | 733 | 523 | 404 | 746 | 1637 | 1135 | 3110 | 2226 | 5547 | 3917 | 9066 |
| 683 | 924 | 749 | 607 | 798 | 705 | 681 | 1005 | 5833 | 5742 | 8407 | 7764 | 1560 | 1263 | 2082 |
| 561 | 818 | 682 | 485 | 762 | 728 | 508 | 763 | 5209 | 3615 | 6443 | 5272 | 4677 | 3327 | 6858 |
| 670 | 991 | 949 | 760 | 979 | 734 | 590 | 880 | 4514 | 3429 | 5118 | 5070 | 1705 | 1365 | 1796 |
| 390 | 896 | 663 | 561 | 952 | | | | 1244 | 839 | 2150 | 1927 | 11985 | 10139 | 18084 |
| 794 | 1452 | 870 | 673 | 1165 | 1034 | 952 | 1468 | 1750 | 1698 | 2137 | 3102 | 6866 | 5312 | 7896 |
| 385 | 756 | 461 | 279 | 586 | 603 | 371 | 849 | 7101 | 4600 | 11947 | 9030 | 4644 | 2806 | 10073 |
| 770 | 1907 | 801 | 777 | 1437 | 886 | 738 | 1702 | 2429 | 2109 | 2739 | 5222 | 1936 | 1879 | 2418 |
| 596 | 894 | 1043 | 727 | 975 | 732 | 587 | 1033 | 1243 | 1080 | 1812 | 1619 | 3808 | 2656 | 3651 |
| 651 | 986 | | | | 939 | 733 | 1153 | 7333 | 5499 | 8443 | 8326 | | | |
| 415 | 617 | 732 | 466 | 619 | 703 | 471 | 719 | 3249 | 1810 | 4358 | 2689 | 4224 | 2688 | 5771 |
| 810 | 1117 | 1008 | 747 | 1255 | 1052 | 754 | 1214 | 6782 | 4638 | 5726 | 6398 | 10713 | 7934 | 10626 |
| 588 | 861 | 819 | 735 | 1043 | 744 | 553 | 770 | 965 | 731 | 1243 | 1071 | 6576 | 5905 | 8030 |
| 682 | 1051 | 778 | 456 | 966 | 690 | 419 | 943 | 6825 | 4707 | 6897 | 7249 | 3777 | 2212 | 4852 |
| 486 | 769 | 711 | 473 | 843 | 703 | 389 | 721 | 5834 | 3893 | 8007 | 6161 | 5312 | 3533 | 7466 |
| 561 | 840 | 738 | 572 | 736 | 630 | 420 | 619 | 1719 | 1469 | 2621 | 2202 | 5953 | 4615 | 8062 |
| 555 | 880 | 740 | 540 | 849 | 692 | 607 | 933 | 2742 | 2108 | 3798 | 3344 | 4949 | 3612 | 6684 |
| 806 | 1307 | | | | 907 | 646 | 1158 | 5424 | 4465 | 5538 | 7235 | | | |
| 1359 | 1476 | 1174 | 1020 | 1391 | 1191 | 890 | 1479 | 2420 | 2612 | 1923 | 2837 | 11824 | 10269 | 10072 |
| 670 | 1157 | 896 | 809 | 1200 | 723 | 669 | 1112 | 8680 | 8038 | 11997 | 13876 | 5440 | 4909 | 6070 |
| | | 1094 | 729 | 961 | 1170 | 820 | 1108 | | | | | 9502 | 6332 | 8685 |
| | | 727 | 488 | 645 | 566 | 318 | 477 | | | | | 14164 | 9512 | 19494 |
| 1046 | 1493 | 993 | 968 | 1413 | 1080 | 1082 | 1561 | 6752 | 6846 | 6543 | 9768 | 6506 | 6342 | 6553 |
| 620 | 698 | 1084 | 815 | 695 | 907 | 738 | 739 | 4805 | 4016 | 6477 | 4519 | 15330 | 11526 | 14145 |
| 534 | 1067 | 695 | 532 | 1041 | 636 | 490 | 1000 | 6590 | 4853 | 9084 | 9695 | 5102 | 3900 | 7338 |
| 441 | 812 | 451 | 294 | 596 | | | | 9366 | 7177 | 16268 | 13205 | 4455 | 2902 | 9888 |
| 783 | 990 | 829 | 809 | 1219 | 819 | 689 | 1004 | 825 | 949 | 1213 | 1201 | 1371 | 1338 | 1653 |
| 1603 | 1479 | 1104 | 1407 | 1648 | 923 | 1149 | 1485 | 2932 | 3399 | 2121 | 3137 | 2126 | 2711 | 1927 |
| 563 | 675 | 1364 | 1148 | 1328 | 1417 | 1245 | 1503 | 4942 | 2813 | 4997 | 3375 | 915 | 770 | 671 |
| | | 519 | 331 | 710 | 408 | 214 | 564 | | | | | 6829 | 4346 | 13145 |
| 738 | 743 | 546 | 335 | 686 | 724 | 532 | 865 | 4029 | 2827 | 3829 | 2844 | 12193 | 7477 | 22324 |

| | | | | | | | | | | | | | | |
|---|---|---|---|---|---|---|---|---|---|---|---|---|---|---|
| 920 | 1044 | 666 | 522 | 652 | 1230 | 1007 | 1075 | 2890 | 2391 | 2598 | 2711 | 3705 | 2903 | 5565 |
| 1020 | 1146 | 1534 | 799 | 1062 | | | | 2064 | 1407 | 1380 | 1581 | 689 | 359 | 449 |
| 447 | 923 | 628 | 504 | 1085 | 680 | 513 | 1156 | 9305 | 6377 | 14281 | 13189 | 3116 | 2501 | 4965 |
| 844 | 1180 | 878 | 658 | 959 | 906 | 775 | 1222 | 5214 | 3939 | 4668 | 5508 | 7494 | 5611 | 8532 |
| 533 | 951 | 958 | 621 | 1159 | 964 | 720 | 1272 | 3774 | 2388 | 4481 | 4262 | 4658 | 3018 | 4860 |
| 336 | 573 | 777 | 448 | 663 | 707 | 422 | 603 | 1135 | 558 | 1660 | 951 | 1257 | 724 | 1617 |
| 641 | 1014 | | | | 512 | 464 | 1041 | 5620 | 4979 | 7772 | 7878 | | | |
| 359 | 621 | 654 | 345 | 662 | 742 | 425 | 749 | 6301 | 2774 | 7718 | 4797 | 6213 | 3283 | 9505 |
| 489 | 1071 | | | | 1213 | 839 | 1058 | 159 | 80 | 163 | 174 | | | |
| 585 | 1056 | 776 | 583 | 1002 | 863 | 674 | 967 | 1095 | 918 | 1570 | 1658 | 4067 | 3054 | 5241 |
| 350 | 575 | 808 | 488 | 905 | 896 | 413 | 689 | 9318 | 3199 | 9136 | 5251 | 7842 | 4733 | 9699 |
| 269 | 661 | 350 | 234 | 678 | 412 | 235 | 645 | 7540 | 4286 | 15958 | 10544 | 1607 | 1074 | 4585 |
| 575 | 812 | 650 | 524 | 788 | 793 | 576 | 889 | 1496 | 1164 | 2024 | 1643 | 711 | 573 | 1094 |
| 798 | 1153 | 887 | 572 | 882 | 779 | 800 | 977 | 2076 | 1553 | 1947 | 2245 | 7954 | 5132 | 8966 |
| 313 | 665 | 601 | 345 | 614 | 609 | 369 | 713 | 2649 | 1454 | 4648 | 3092 | 2740 | 1574 | 4558 |
| 1287 | 1513 | 1159 | 936 | 1275 | 904 | 775 | 1087 | 798 | 765 | 595 | 900 | 621 | 502 | 536 |
| 498 | 640 | 1449 | 585 | 910 | 1102 | 691 | 918 | 3542 | 1136 | 2281 | 1459 | 6925 | 2796 | 4777 |
| 516 | 982 | 856 | 624 | 1037 | 823 | 576 | 1056 | 4199 | 2608 | 5050 | 4958 | 11447 | 8347 | 13367 |
| 993 | 1107 | | | | 821 | 647 | 1007 | 3166 | 2795 | 2814 | 3116 | | | |
| 670 | 1043 | 664 | 377 | 681 | 660 | 402 | 709 | 872 | 728 | 1086 | 1133 | 5911 | 3356 | 8900 |
| 861 | 1140 | 882 | 681 | 1103 | 871 | 782 | 1261 | 1666 | 1567 | 1820 | 2074 | 2500 | 1928 | 2833 |
| 794 | 1133 | 859 | 725 | 1301 | 789 | 684 | 1063 | 8258 | 7599 | 9566 | 10841 | 3832 | 3234 | 4462 |
| 860 | 1127 | 1284 | 1155 | 1415 | 1410 | 1243 | 1391 | 6288 | 5134 | 5973 | 6732 | 571 | 513 | 444 |
| 702 | 1088 | | | | 819 | 445 | 817 | 1960 | 1620 | 2308 | 2512 | | | |
| 687 | 1009 | 1199 | 876 | 1109 | 915 | 632 | 992 | 7598 | 5679 | 8270 | 8341 | 746 | 544 | 622 |
| 866 | 1112 | | | | 940 | 946 | 1326 | 4288 | 3761 | 4345 | 4833 | | | |
| 435 | 802 | 718 | 571 | 898 | 451 | 399 | 719 | 4177 | 3001 | 6892 | 5531 | 2332 | 1854 | 3246 |
| | | 1044 | 1048 | 1408 | 1020 | 824 | 1145 | | | | | 841 | 844 | 805 |
| 375 | 695 | 270 | 170 | 505 | | | | 4689 | 3980 | 10618 | 7384 | 5168 | 3257 | 19133 |
| 633 | 929 | 1125 | 937 | 1124 | 881 | 658 | 998 | 3189 | 2325 | 3674 | 3413 | 5263 | 4383 | 4679 |
| 609 | 928 | | | | 1257 | 936 | 969 | 4362 | 2929 | 4811 | 4463 | | | |

| | | | | | | | | | | | | | | |
|---|---|---|---|---|---|---|---|---|---|---|---|---|---|---|
| 693 | 1016 | 924 | 784 | 1078 | 909 | 774 | 1088 | 4210 | 3503 | 5056 | 5138 | 9006 | 7643 | 9751 |
| 625 | 996 | 742 | 956 | 1346 | 876 | 771 | 1154 | 2333 | 1362 | 2180 | 2171 | 1451 | 1871 | 1957 |
| 1165 | 2028 | 879 | 1021 | 1966 | 791 | 660 | 1312 | 2405 | 2840 | 2438 | 4946 | 2098 | 2436 | 2387 |
| 725 | 1157 | 1020 | 676 | 1152 | 826 | 558 | 948 | 6333 | 4885 | 6737 | 7795 | 14511 | 9615 | 14225 |
| 880 | 1179 | 962 | 956 | 1368 | 967 | 926 | 1207 | 2105 | 1819 | 2068 | 2437 | 5522 | 5484 | 5737 |
| 786 | 1054 | 728 | 985 | 1783 | 827 | 947 | 1355 | 2724 | 2356 | 2997 | 3158 | 2143 | 2898 | 2943 |
| 773 | 955 | 934 | 699 | 951 | 934 | 759 | 982 | 5534 | 4404 | 5694 | 5436 | 5688 | 4255 | 6088 |
| 436 | 768 | 775 | 489 | 708 | 842 | 484 | 800 | 1765 | 919 | 2106 | 1617 | 933 | 589 | 1205 |
| 917 | 1494 | 1125 | 1002 | 1324 | 992 | 946 | 1406 | 2476 | 2320 | 2530 | 3779 | 6950 | 6188 | 6177 |
| 684 | 792 | 765 | 537 | 945 | 832 | 570 | 956 | 2135 | 1693 | 2474 | 1960 | 3279 | 2301 | 4288 |
| 1124 | 1372 | 1150 | 1007 | 1290 | 1224 | 1128 | 1306 | 2213 | 1940 | 1725 | 2367 | 1564 | 1370 | 1360 |
| | | 939 | 876 | 1277 | 931 | 870 | 1268 | | | | | 6646 | 6198 | 7075 |
| 802 | 1033 | 1246 | 1040 | 1408 | 1205 | 977 | 1530 | 1256 | 1190 | 1485 | 1534 | 3690 | 3082 | 2963 |
| 515 | 407 | 1175 | 934 | 583 | 1181 | 857 | 685 | 1656 | 780 | 1514 | 616 | 2163 | 1720 | 1841 |
| 1179 | 1283 | 1092 | 1028 | 1175 | 1147 | 1124 | 1783 | 2408 | 2157 | 1829 | 2346 | 5006 | 4715 | 4586 |
| 479 | 735 | | | | 674 | 590 | 845 | 8106 | 6764 | 14122 | 10379 | | | |
| 511 | 906 | 793 | 581 | 862 | 708 | 480 | 766 | 1920 | 1349 | 2640 | 2393 | 3166 | 2322 | 3993 |
| 715 | 1150 | 957 | 869 | 1268 | 960 | 793 | 1272 | 9672 | 8713 | 12183 | 14005 | 1979 | 1797 | 2068 |
| 364 | 693 | 848 | 620 | 904 | | | | 3580 | 2374 | 6517 | 4519 | 1305 | 955 | 1540 |
| | | 530 | 391 | 745 | 519 | 346 | 735 | | | | | 8454 | 6241 | 15961 |
| 437 | 709 | 594 | 432 | 637 | 660 | 435 | 714 | 7225 | 5049 | 11558 | 8191 | 1899 | 1380 | 3198 |
| 699 | 1065 | 573 | 754 | 1138 | 573 | 759 | 1136 | 1212 | 1601 | 2290 | 2439 | 3317 | 4363 | 5785 |
| 524 | 842 | 790 | 545 | 811 | 931 | 538 | 815 | 3326 | 2052 | 3917 | 3296 | 7397 | 5107 | 9369 |
| 205 | 326 | 751 | 176 | 313 | 595 | 169 | 295 | 7535 | 1980 | 9662 | 3153 | 7498 | 1761 | 9984 |
| 450 | 759 | 693 | 560 | 828 | 580 | 445 | 698 | 4900 | 3473 | 7717 | 5854 | 5720 | 4626 | 8256 |
| 606 | 1219 | 853 | 599 | 1232 | | | | 5270 | 3962 | 6536 | 7968 | 7242 | 5091 | 8493 |
| 629 | 963 | 773 | 556 | 869 | | | | 8225 | 5972 | 9500 | 9146 | 5241 | 3769 | 6778 |
| 801 | 1099 | 976 | 795 | 1051 | 1021 | 802 | 1121 | 2907 | 2409 | 3006 | 3303 | 3308 | 2693 | 3389 |
| 548 | 1060 | 669 | 485 | 952 | 815 | 592 | 1044 | 6265 | 4277 | 7802 | 8268 | 1727 | 1251 | 2582 |
| 831 | 1261 | 965 | 846 | 1289 | 867 | 753 | 1165 | 7362 | 6512 | 7834 | 9878 | 10582 | 9276 | 10966 |
| 469 | 1925 | 812 | 490 | 2852 | 758 | 439 | 1834 | 3234 | 1903 | 4054 | 7805 | 6757 | 4076 | 8317 |

| | | | | | | | | | | | | | | |
|---|---|---|---|---|---|---|---|---|---|---|---|---|---|---|
| 563 | 797 | 862 | 463 | 607 | 719 | 494 | 875 | 5009 | 2906 | 5158 | 4111 | 1579 | 848 | 1832 |
| 769 | 972 | 1192 | 892 | 1271 | 1293 | 924 | 1260 | 5933 | 4522 | 5879 | 5712 | 3095 | 2317 | 2597 |
| 655 | 1033 | 1075 | 919 | 1150 | 872 | 716 | 980 | 6556 | 5740 | 8767 | 9053 | 1944 | 1663 | 1810 |
| 466 | 757 | | | | 845 | 584 | 952 | 1558 | 871 | 1868 | 1413 | | | |
| 585 | 858 | 1046 | 741 | 953 | 829 | 546 | 851 | 1834 | 1390 | 2375 | 2037 | 6788 | 4811 | 6492 |
| 588 | 813 | 820 | 633 | 878 | 829 | 653 | 855 | 1896 | 1153 | 1961 | 1593 | 12430 | 9598 | 15159 |
| 299 | 575 | 585 | 425 | 765 | 575 | 451 | 789 | 466 | 312 | 1045 | 601 | 2684 | 1950 | 4589 |
| 475 | 956 | 767 | 565 | 957 | 713 | 600 | 1073 | 4078 | 3097 | 6521 | 6232 | 7137 | 5256 | 9306 |
| 691 | 1048 | | | | 669 | 635 | 979 | 5067 | 3500 | 5064 | 5307 | | | |
| 778 | 616 | 438 | 544 | 642 | | | | 556 | 675 | 868 | 535 | 3443 | 4273 | 7856 |
| 821 | 1041 | 802 | 646 | 874 | | | | 6079 | 5148 | 6273 | 6533 | 969 | 781 | 1208 |
| 632 | 807 | | | | 763 | 716 | 1026 | 565 | 646 | 1023 | 826 | | | |
| 773 | 937 | 1139 | 885 | 1223 | 925 | 697 | 961 | 4902 | 3873 | 5010 | 4696 | 2948 | 2292 | 2589 |
| | | 504 | 336 | 733 | 709 | 559 | 837 | | | | | 3188 | 2128 | 6331 |
| 940 | 1013 | 1092 | 940 | 1050 | 1180 | 909 | 1211 | 7796 | 6640 | 7065 | 7154 | 2043 | 1760 | 1872 |
| 870 | 1068 | 1012 | 912 | 1138 | 1117 | 1010 | 1195 | 1425 | 1146 | 1316 | 1406 | 1817 | 1637 | 1796 |
| 896 | 910 | 667 | 465 | 896 | 705 | 493 | 1001 | 1849 | 1695 | 1892 | 1722 | 3272 | 2284 | 4908 |
| 866 | 1214 | 847 | 760 | 1158 | | | | 8370 | 6997 | 8081 | 9809 | 5505 | 4940 | 6502 |
| | | 718 | 631 | 935 | 498 | 451 | 887 | | | | | 2506 | 2204 | 3492 |
| 805 | 1423 | 788 | 639 | 1145 | 835 | 689 | 1132 | 1020 | 882 | 1096 | 1560 | 4623 | 3745 | 5866 |
| 705 | 1079 | 893 | 701 | 1072 | 1026 | 793 | 1179 | 4573 | 3454 | 4898 | 5284 | 5260 | 4128 | 5892 |
| 530 | 873 | 851 | 567 | 974 | 1077 | 799 | 1004 | 860 | 582 | 1098 | 959 | 523 | 349 | 615 |
| 417 | 970 | 679 | 383 | 852 | 606 | 347 | 828 | 1579 | 935 | 2244 | 2177 | 2081 | 1176 | 3066 |
| 987 | 1140 | 751 | 626 | 1096 | 603 | 609 | 925 | 1539 | 1343 | 1361 | 1552 | 5855 | 4882 | 7801 |
| 538 | 821 | 882 | 555 | 852 | 641 | 668 | 919 | 3463 | 2326 | 4324 | 3551 | 2917 | 1836 | 3307 |
| 223 | 583 | 550 | 412 | 859 | 568 | 324 | 725 | 3914 | 1649 | 7401 | 4312 | 1236 | 927 | 2248 |
| | | 855 | 657 | 840 | 810 | 545 | 810 | | | | | 6646 | 5112 | 7775 |
| 644 | 910 | 1380 | 618 | 906 | 746 | 466 | 746 | 6678 | 4001 | 6213 | 5651 | 2080 | 931 | 1508 |
| 683 | 1141 | 900 | 778 | 1339 | 1087 | 657 | 819 | 5356 | 4832 | 7080 | 8074 | 4202 | 3632 | 4670 |
| 731 | 843 | 877 | 687 | 817 | 947 | 784 | 941 | 5986 | 4754 | 6504 | 5485 | 5133 | 4020 | 5853 |
| 885 | 1225 | 1295 | 1049 | 1356 | 1450 | 861 | 1139 | 4916 | 3502 | 3956 | 4847 | 4950 | 4009 | 3824 |

| | | | | | | | | | | | | | | |
|---|---|---|---|---|---|---|---|---|---|---|---|---|---|---|
| 1593 | 1914 | 953 | 941 | 1324 | 852 | 702 | 1081 | 4088 | 5001 | 3139 | 6008 | 5071 | 5008 | 5321 |
| 563 | 863 | 891 | 852 | 943 | 551 | 412 | 780 | 5191 | 4382 | 7790 | 6726 | 3101 | 2966 | 3480 |
| 671 | 974 | 748 | 618 | 956 | 721 | 604 | 976 | 5983 | 4337 | 6461 | 6290 | 7130 | 5885 | 9527 |
| 418 | 886 | 632 | 430 | 981 | 598 | 457 | 955 | 3357 | 2124 | 5078 | 4497 | 4525 | 3073 | 7154 |
| 878 | 1151 | 740 | 688 | 942 | 896 | 787 | 953 | 3718 | 3411 | 3887 | 4473 | 3266 | 3037 | 4415 |
| 925 | 1420 | 937 | 828 | 1325 | 837 | 645 | 1197 | 2553 | 2236 | 2417 | 3431 | 4950 | 4374 | 5283 |
| 444 | 768 | 808 | 538 | 932 | 735 | 466 | 780 | 1714 | 1018 | 2293 | 1760 | 1281 | 853 | 1586 |
| 471 | 901 | 742 | 443 | 858 | 828 | 491 | 902 | 4844 | 3059 | 6492 | 5852 | 6082 | 3637 | 8202 |
| | | | | | 1101 | 904 | 997 | | | | | | | |
| 672 | 908 | 866 | 680 | 880 | 958 | 768 | 900 | 2865 | 2313 | 3440 | 3124 | 3319 | 2607 | 3830 |
| | | 1197 | 1075 | 1075 | | | | | | | | 1162 | 1044 | 971 |
| 379 | 612 | 761 | 462 | 743 | 757 | 458 | 793 | 5645 | 2823 | 7447 | 4558 | 5495 | 3333 | 7217 |
| 694 | 916 | 750 | 753 | 1005 | 661 | 775 | 912 | 6194 | 6136 | 8840 | 8095 | 6995 | 7018 | 9327 |
| 541 | 763 | 637 | 556 | 806 | 592 | 547 | 786 | 3644 | 3031 | 5603 | 4277 | 2398 | 2093 | 3763 |
| 579 | 960 | 987 | 788 | 1174 | 955 | 772 | 1078 | 4043 | 2962 | 5115 | 4909 | 8406 | 6709 | 8516 |
| 431 | 820 | 637 | 462 | 749 | 567 | 417 | 755 | 2150 | 1583 | 3677 | 3015 | 963 | 699 | 1512 |
| 508 | 794 | 707 | 450 | 743 | 532 | 289 | 664 | 7029 | 4750 | 9353 | 7428 | 1289 | 820 | 1823 |
| 318 | 574 | 586 | 351 | 574 | | | | 1670 | 952 | 2999 | 1722 | 4960 | 2970 | 8469 |
| 477 | 703 | 744 | 713 | 987 | 795 | 659 | 983 | 4557 | 2222 | 4654 | 3272 | 1569 | 1504 | 2109 |
| 684 | 840 | 831 | 695 | 962 | 776 | 694 | 856 | 3906 | 2957 | 4324 | 3630 | 4217 | 3531 | 5077 |
| | | 683 | 474 | 760 | 636 | 443 | 755 | | | | | 2536 | 1762 | 3715 |
| 627 | 900 | 918 | 686 | 932 | 692 | 399 | 845 | 3206 | 2366 | 3773 | 3395 | 5256 | 3924 | 5724 |
| 718 | 1028 | | | | 812 | 621 | 946 | 8733 | 6648 | 9259 | 9522 | | | |
| 822 | 1110 | 894 | 667 | 922 | 901 | 733 | 1001 | 6121 | 5788 | 7042 | 7820 | 2181 | 1628 | 2439 |
| 977 | 1301 | 1200 | 1268 | 1704 | 987 | 740 | 1112 | 3190 | 2898 | 2967 | 3861 | 3816 | 4030 | 3179 |
| 1011 | 1152 | 1210 | 1345 | 1366 | 1283 | 1501 | 1466 | 4952 | 4695 | 4643 | 5346 | 6904 | 7675 | 5705 |
| | | 520 | 279 | 572 | 512 | 328 | 682 | | | | | 7672 | 4122 | 14754 |
| 587 | 957 | 966 | 607 | 995 | 794 | 466 | 831 | 2994 | 1751 | 2985 | 2857 | 2600 | 1634 | 2692 |
| 628 | 965 | 852 | 671 | 1048 | 751 | 547 | 893 | 1406 | 1050 | 1674 | 1615 | 1071 | 844 | 1257 |
| 628 | 1089 | 888 | 746 | 1194 | 918 | 765 | 1153 | 7624 | 5676 | 9030 | 9831 | 1820 | 1529 | 2050 |
| 544 | 819 | 779 | 726 | 1007 | 752 | 600 | 951 | 1003 | 620 | 1138 | 932 | 4243 | 3954 | 5445 |

| | | | | | | | | | | | | | | |
|---|---|---|---|---|---|---|---|---|---|---|---|---|---|---|
| 365 | 593 | 391 | 345 | 551 | 395 | 349 | 571 | 2325 | 2107 | 5780 | 3425 | 3595 | 3173 | 9202 |
| 411 | 680 | 747 | 494 | 765 | 699 | 425 | 678 | 1512 | 847 | 2061 | 1401 | 4998 | 3304 | 6693 |
| 270 | 558 | | | | 404 | 225 | 574 | 3797 | 2297 | 8502 | 4747 | | | |
| | | 598 | 449 | 756 | 592 | 386 | 723 | | | | | 17046 | 12785 | 28485 |
| 721 | 1103 | | | | 1015 | 683 | 1028 | 5830 | 4308 | 5973 | 6589 | | | |
| 557 | 859 | 961 | 668 | 1014 | 961 | 667 | 982 | 3967 | 2735 | 4911 | 4218 | 7502 | 5211 | 7805 |
| 332 | 817 | 805 | 495 | 927 | 805 | 415 | 889 | 4946 | 2339 | 7041 | 5756 | 7280 | 4473 | 9045 |
| 495 | 610 | 721 | 344 | 608 | 714 | 313 | 586 | 2369 | 1176 | 2375 | 1449 | 7671 | 3657 | 10641 |
| 708 | 869 | 742 | 591 | 951 | 471 | 392 | 569 | 593 | 556 | 786 | 683 | 1688 | 1345 | 2275 |
| 518 | 871 | 827 | 549 | 858 | 939 | 533 | 924 | 3484 | 2136 | 4123 | 3590 | 8800 | 5841 | 10642 |
| 931 | 1642 | | | | 876 | 919 | 1554 | 6874 | 7411 | 7965 | 13079 | | | |
| | | 454 | 261 | 621 | | | | | | | | 2132 | 1227 | 4696 |
| 871 | 1164 | | | | 1055 | 645 | 829 | 7841 | 6356 | 7300 | 8497 | | | |
| 636 | 878 | | | | 976 | 447 | 837 | 13660 | 10506 | 16513 | 14492 | | | |
| 581 | 867 | 760 | 520 | 882 | 718 | 448 | 814 | 3234 | 2511 | 4324 | 3749 | 4479 | 3062 | 5891 |
| 1090 | 17227 | 1282 | 1114 | 7183 | | | | 1290 | 897 | 823 | 14174 | 1701 | 1479 | 1327 |
| | | 716 | 554 | 915 | 717 | 526 | 867 | | | | | 2021 | 1562 | 2821 |
| 501 | 807 | 552 | 575 | 903 | 520 | 499 | 807 | 2967 | 2819 | 5625 | 4537 | 3022 | 3150 | 5479 |
| 483 | 925 | 783 | 563 | 944 | 663 | 472 | 900 | 3898 | 2796 | 5790 | 5355 | 2658 | 1910 | 3396 |
| 294 | 464 | 180 | 248 | 442 | 130 | 226 | 352 | 2523 | 3738 | 12700 | 5889 | 1115 | 1533 | 6178 |
| 393 | 683 | 1444 | 836 | 977 | 818 | 413 | 785 | 9946 | 5323 | 13533 | 9239 | 6159 | 3568 | 4266 |
| 410 | 617 | 695 | 530 | 734 | 604 | 481 | 620 | 4876 | 3195 | 7787 | 4804 | 5986 | 4564 | 8608 |
| | | 851 | 702 | 997 | 780 | 670 | 858 | | | | | 4308 | 3554 | 5063 |
| 669 | 992 | 971 | 729 | 985 | 673 | 623 | 843 | 2913 | 2378 | 3553 | 3526 | 4173 | 3130 | 4296 |
| | | 900 | 1077 | 878 | 954 | 1053 | 1015 | | | | | 10175 | 12176 | 11307 |
| 425 | 846 | 725 | 485 | 911 | 695 | 458 | 941 | 5885 | 3626 | 8535 | 7221 | 2165 | 1447 | 2987 |
| 920 | 1263 | 1207 | 1205 | 1547 | 1004 | 1027 | 1356 | 8698 | 7610 | 8268 | 10446 | 5238 | 5228 | 4340 |
| 648 | 1207 | 849 | 549 | 1107 | 968 | 502 | 968 | 7172 | 5164 | 7965 | 9614 | 10813 | 6997 | 12741 |
| 407 | 857 | 591 | 345 | 760 | 633 | 436 | 917 | 6839 | 4350 | 10677 | 9151 | 4287 | 2499 | 7253 |
| 1021 | 1299 | 1021 | 1084 | 1476 | 945 | 832 | 1235 | 10423 | 10320 | 10108 | 13127 | 10191 | 10814 | 9981 |
| 434 | 826 | 617 | 369 | 892 | 852 | 607 | 1052 | 3962 | 2414 | 5562 | 4593 | 2075 | 1241 | 3362 |

| | | | | | | | | | | | | | | |
|---|---|---|---|---|---|---|---|---|---|---|---|---|---|---|
| 845 | 1287 | 895 | 678 | 1085 | | | | 2053 | 1720 | 2036 | 2621 | 5979 | 4530 | 6681 |
| 621 | 890 | 872 | 610 | 926 | 1005 | 725 | 1025 | 4136 | 2977 | 4797 | 4268 | 5932 | 4151 | 6803 |
| 271 | 586 | 888 | 368 | 724 | 953 | 416 | 819 | 2504 | 837 | 3088 | 1809 | 4874 | 2023 | 5492 |
| 747 | 1225 | 834 | 860 | 1451 | 810 | 804 | 1365 | 4180 | 3788 | 5069 | 6210 | 1918 | 1978 | 2298 |
| | | 942 | 688 | 1007 | 761 | 530 | 716 | | | | | 8318 | 6072 | 8830 |
| 803 | 1163 | 767 | 589 | 887 | | | | 4875 | 4310 | 5369 | 6245 | 3074 | 2363 | 4008 |
| 639 | 985 | 863 | 637 | 1042 | 887 | 614 | 879 | 1737 | 1407 | 2203 | 2169 | 2148 | 1586 | 2489 |
| 475 | 871 | 538 | 488 | 950 | 633 | 460 | 804 | 6720 | 5374 | 11306 | 9848 | 2058 | 1865 | 3825 |
| 497 | 830 | 562 | 398 | 818 | 600 | 385 | 735 | 6535 | 4702 | 9457 | 7853 | 4184 | 2963 | 7451 |
| 594 | 907 | 806 | 642 | 1007 | 868 | 603 | 928 | 5678 | 4405 | 7413 | 6724 | 5302 | 4225 | 6577 |
| 802 | 1121 | 1034 | 890 | 1172 | | | | 4287 | 3126 | 3900 | 4372 | 5212 | 4486 | 5040 |
| 636 | 662 | 1036 | 653 | 708 | 999 | 805 | 819 | 1989 | 1656 | 2603 | 1724 | 5195 | 3274 | 5015 |
| 734 | 1068 | 1221 | 851 | 1221 | 1326 | 773 | 1082 | 8755 | 6049 | 8238 | 8794 | 5376 | 3748 | 4404 |
| 257 | 551 | 515 | 262 | 558 | 459 | 215 | 524 | 3560 | 1859 | 7234 | 3986 | 6824 | 3466 | 13242 |
| 520 | 952 | 744 | 480 | 785 | 544 | 448 | 774 | 583 | 414 | 796 | 758 | 2762 | 1781 | 3713 |
| 492 | 891 | 560 | 413 | 632 | 638 | 468 | 761 | 4613 | 3711 | 7538 | 6720 | 586 | 433 | 1047 |
| 645 | 921 | 624 | 399 | 691 | 693 | 526 | 832 | 1908 | 1520 | 2357 | 2170 | 3758 | 2407 | 6027 |
| 737 | 1101 | 762 | 730 | 1073 | 766 | 723 | 1104 | 2595 | 2381 | 3232 | 3557 | 3011 | 2884 | 3952 |
| 580 | 1006 | 922 | 574 | 829 | 834 | 669 | 963 | 1115 | 675 | 1163 | 1170 | 1110 | 691 | 1204 |
| 677 | 896 | 869 | 640 | 888 | 913 | 664 | 908 | 1304 | 1006 | 1487 | 1332 | 2005 | 1475 | 2307 |
| 783 | 1071 | | | | | | | 5066 | 3826 | 4883 | 5230 | | | |
| 486 | 820 | 531 | 467 | 795 | 477 | 394 | 611 | 5374 | 5084 | 10461 | 8579 | 1182 | 1040 | 2227 |
| 396 | 669 | 698 | 389 | 721 | 797 | 505 | 856 | 1777 | 1108 | 2802 | 1873 | 2235 | 1245 | 3202 |
| 683 | 891 | | | | 721 | 467 | 752 | 5695 | 3980 | 5832 | 5193 | | | |
| 372 | 876 | 743 | 473 | 989 | 743 | 455 | 1000 | 2202 | 1262 | 3391 | 2970 | 6557 | 4173 | 8830 |
| 305 | 460 | 815 | 679 | 851 | 916 | 731 | 1005 | 5560 | 2656 | 8713 | 4005 | 903 | 752 | 1107 |
| 884 | 1236 | 810 | 521 | 887 | 1236 | 700 | 1023 | 3585 | 2145 | 2427 | 3000 | 5647 | 3632 | 6970 |
| 722 | 1059 | 799 | 657 | 1035 | 692 | 633 | 935 | 7430 | 6124 | 8483 | 8984 | 4429 | 3642 | 5540 |
| 721 | 1012 | | | | 711 | 623 | 976 | 3360 | 2879 | 3995 | 4043 | | | |
| | | 1119 | 1211 | 1789 | 1204 | 1362 | 1957 | | | | | 5458 | 5906 | 4875 |
| 381 | 455 | 1010 | 431 | 602 | 990 | 411 | 574 | 5702 | 1823 | 4790 | 2181 | 7988 | 3413 | 7911 |

| | | | | | | | | | | | | | | |
|---|---|---|---|---|---|---|---|---|---|---|---|---|---|---|
| | | 613 | 501 | 851 | 983 | 754 | 853 | | | | | 9872 | 8067 | 16095 |
| | | 908 | 637 | 1003 | 823 | 648 | 1058 | | | | | 5129 | 3596 | 5648 |
| 822 | 877 | 718 | 509 | 892 | 833 | 476 | 772 | 2710 | 2330 | 2837 | 2488 | 1265 | 897 | 1762 |
| 697 | 965 | 1001 | 767 | 1040 | 1010 | 747 | 1106 | 4736 | 3595 | 5158 | 4979 | 11821 | 9062 | 11812 |
| 901 | 1355 | | | | 886 | 724 | 1141 | 4506 | 4176 | 4635 | 6278 | | | |
| 411 | 934 | 510 | 366 | 824 | 481 | 332 | 838 | 4814 | 3958 | 9638 | 8998 | 4573 | 3281 | 8969 |
| | | 936 | 754 | 1224 | 970 | 707 | 1058 | | | | | 2698 | 2173 | 2882 |
| 855 | 645 | 1232 | 853 | 640 | 1310 | 839 | 658 | 6945 | 4594 | 5375 | 3465 | 6017 | 4163 | 4884 |
| 2081 | 856 | 823 | 2085 | 860 | 808 | 1969 | 868 | 3920 | 9800 | 4708 | 4032 | 3595 | 9106 | 4368 |
| | | 1137 | 773 | 1066 | 925 | 619 | 922 | | | | | 1238 | 842 | 1089 |
| 390 | 852 | 533 | 327 | 721 | 599 | 383 | 799 | 885 | 638 | 1638 | 1396 | 1024 | 627 | 1920 |
| 561 | 893 | 707 | 484 | 803 | 880 | 611 | 947 | 1731 | 1117 | 1991 | 1778 | 2174 | 1488 | 3075 |
| 806 | 1228 | 830 | 699 | 1120 | 894 | 605 | 1105 | 1480 | 1036 | 1287 | 1580 | 1910 | 1607 | 2301 |
| 367 | 707 | 518 | 364 | 680 | 556 | 398 | 759 | 2700 | 1909 | 5205 | 3681 | 2793 | 1962 | 5394 |
| 462 | 784 | 762 | 488 | 817 | 671 | 369 | 646 | 2635 | 1561 | 3381 | 2651 | 6408 | 4107 | 8410 |
| 777 | 925 | 1108 | 778 | 1048 | 1093 | 800 | 1014 | 2846 | 2100 | 2702 | 2499 | 3415 | 2399 | 3084 |
| 857 | 1091 | | | | 939 | 773 | 1027 | 1599 | 1358 | 1584 | 1728 | | | |
| 107 | 194 | 837 | 625 | 678 | | | | 7095 | 1151 | 10802 | 2094 | 2368 | 1768 | 2830 |
| 301 | 482 | 534 | 425 | 671 | | | | 689 | 540 | 1794 | 865 | 9508 | 7563 | 17802 |
| 631 | 966 | 501 | 320 | 693 | 907 | 771 | 1075 | 8037 | 5538 | 8773 | 8477 | 2898 | 1849 | 5783 |
| 415 | 912 | | | | 950 | 649 | 932 | 3302 | 1817 | 4376 | 3992 | | | |
| | | 837 | 749 | 1149 | 725 | 607 | 1215 | | | | | 5578 | 4990 | 6663 |
| 656 | 819 | 872 | 678 | 1081 | 1059 | 729 | 1347 | 2907 | 2047 | 3121 | 2556 | 3402 | 2645 | 3900 |
| | | 1135 | 1094 | 1516 | 956 | 855 | 1238 | | | | | 3062 | 2950 | 2697 |
| 731 | 1124 | 848 | 593 | 1087 | 876 | 758 | 1251 | 2035 | 1828 | 2499 | 2810 | 1785 | 1250 | 2106 |
| 752 | 1211 | 651 | 569 | 1086 | | | | 11180 | 9960 | 13250 | 16040 | 5099 | 4454 | 7834 |
| 630 | 952 | 824 | 743 | 996 | 858 | 685 | 1052 | 1159 | 939 | 1489 | 1418 | 1681 | 1516 | 2040 |
| 601 | 1135 | 774 | 559 | 1142 | 997 | 732 | 1106 | 6345 | 4697 | 7817 | 8874 | 1681 | 1214 | 2171 |
| 437 | 610 | | | | 885 | 426 | 594 | 6032 | 3195 | 7304 | 4454 | | | |
| 848 | 1223 | | | | 1092 | 961 | 1844 | 2396 | 1998 | 2357 | 2883 | | | |
| 826 | 1209 | 714 | 656 | 1049 | 849 | 785 | 1103 | 1526 | 1266 | 1533 | 1853 | 434 | 399 | 608 |

| | | | | | | | | | | | | | | |
|---|---|---|---|---|---|---|---|---|---|---|---|---|---|---|
| 742 | 919 | 775 | 609 | 781 | | | | 2406 | 2081 | 2803 | 2577 | 4901 | 3849 | 6324 |
| 464 | 778 | | | | 629 | 396 | 664 | 1162 | 897 | 1931 | 1503 | | | |
| 319 | 687 | 1034 | 895 | 1094 | 775 | 411 | 713 | 2820 | 1374 | 4307 | 2958 | 1998 | 1729 | 1932 |
| 565 | 865 | 1064 | 875 | 1193 | 1396 | 599 | 1026 | 1023 | 392 | 693 | 599 | 3607 | 2966 | 3390 |
| 592 | 899 | 880 | 474 | 772 | 657 | 473 | 847 | 6046 | 3749 | 6330 | 5692 | 5265 | 2838 | 5985 |
| 648 | 944 | 826 | 588 | 846 | 896 | 652 | 858 | 7963 | 6133 | 9459 | 8929 | 3656 | 2601 | 4427 |
| | | 1157 | 780 | 1094 | 740 | 532 | 830 | | | | | 1199 | 808 | 1036 |
| 729 | 1050 | 936 | 655 | 1060 | 1177 | 1038 | 1221 | 1557 | 1234 | 1692 | 1777 | 963 | 673 | 1028 |
| 434 | 828 | 544 | 386 | 857 | 478 | 297 | 709 | 6592 | 4671 | 10752 | 8903 | 5906 | 4197 | 10864 |
| 930 | 1207 | 1152 | 969 | 1224 | 1323 | 1055 | 1253 | 6641 | 5447 | 5859 | 7073 | 3593 | 3021 | 3118 |
| 500 | 716 | | | | 1074 | 916 | 1229 | 7322 | 4827 | 9653 | 6914 | | | |
| 503 | 839 | 806 | 503 | 705 | 787 | 558 | 784 | 3855 | 2405 | 4780 | 4009 | 11088 | 6923 | 13753 |
| 616 | 2450 | | | | 1074 | 717 | 2190 | 4053 | 2436 | 3957 | 9693 | | | |
| | | 511 | 353 | 620 | 644 | 439 | 800 | | | | | 3718 | 2569 | 7274 |
| 144 | 455 | 409 | 202 | 545 | 411 | 204 | 576 | 2444 | 958 | 6635 | 3020 | 2597 | 1283 | 6342 |
| 549 | 1008 | 759 | 565 | 1201 | | | | 6649 | 5067 | 9233 | 9309 | 7093 | 5277 | 9340 |
| 740 | 1173 | 881 | 666 | 1112 | 870 | 770 | 1188 | 3912 | 3088 | 4174 | 4894 | 3725 | 2818 | 4230 |
| 468 | 995 | | | | 591 | 467 | 952 | 4835 | 3744 | 7997 | 7958 | | | |
| 875 | 882 | | | | 694 | 538 | 760 | 14373 | 12976 | 14828 | 13075 | | | |
| 863 | 1120 | 831 | 777 | 1037 | 933 | 812 | 1082 | 1959 | 1779 | 2060 | 2308 | 2569 | 2401 | 3092 |
| | | 779 | 609 | 1006 | 599 | 462 | 958 | | | | | 6674 | 5216 | 8569 |
| 772 | 1029 | 844 | 647 | 1000 | 989 | 780 | 941 | 3592 | 2804 | 3630 | 3736 | 1891 | 1451 | 2242 |
| 667 | 924 | 931 | 682 | 1086 | | | | 5315 | 3673 | 5509 | 5091 | 7161 | 5246 | 7695 |
| 428 | 815 | 958 | 463 | 876 | 874 | 440 | 861 | 6649 | 3420 | 7992 | 6516 | 4913 | 2374 | 5128 |
| 702 | 1249 | 736 | 676 | 1249 | 729 | 680 | 1273 | 2833 | 2665 | 3795 | 4740 | 3524 | 3233 | 4786 |
| 685 | 1041 | | | | 955 | 751 | 1067 | 7333 | 6058 | 8844 | 9208 | | | |
| 876 | 1160 | | | | | | | 2583 | 2300 | 2627 | 3046 | | | |
| 716 | 996 | 1270 | 769 | 1161 | 1172 | 745 | 1046 | 2281 | 1489 | 2081 | 2073 | 2562 | 1551 | 2017 |
| 365 | 814 | 696 | 403 | 864 | 713 | 463 | 898 | 5017 | 2942 | 8057 | 6557 | 6092 | 3526 | 8755 |
| 226 | 701 | 590 | 287 | 754 | 926 | 782 | 1213 | 3100 | 1267 | 5609 | 3930 | 1395 | 679 | 2363 |
| 595 | 909 | 792 | 493 | 853 | 693 | 398 | 678 | 1616 | 1222 | 2054 | 1867 | 642 | 399 | 810 |

| | | | | | | | | | | | | | | |
|---|---|---|---|---|---|---|---|---|---|---|---|---|---|---|
| | | 1336 | 1137 | 1205 | 952 | 782 | 963 | | | | | 942 | 802 | 705 |
| 538 | 824 | 770 | 550 | 854 | 810 | 545 | 817 | 3737 | 2579 | 4790 | 3948 | 2260 | 1615 | 2936 |
| 567 | 604 | 509 | 576 | 613 | 576 | 642 | 719 | 1841 | 2079 | 3669 | 2215 | 2452 | 2776 | 4815 |
| 710 | 849 | 968 | 818 | 1068 | | | | 5486 | 4285 | 6034 | 5124 | 728 | 615 | 752 |
| 1257 | 1334 | 998 | 1160 | 1352 | 779 | 1079 | 1109 | 2402 | 3526 | 2805 | 3741 | 3915 | 4553 | 3924 |
| 530 | 928 | 760 | 580 | 1091 | 788 | 548 | 1393 | 5112 | 3593 | 6782 | 6294 | 6587 | 5025 | 8662 |
| 911 | 1158 | 1257 | 909 | 1162 | 1295 | 941 | 1165 | 4458 | 3297 | 3619 | 4192 | 4782 | 3459 | 3806 |
| 516 | 851 | 694 | 634 | 936 | 558 | 479 | 767 | 1233 | 1117 | 2166 | 1842 | 3751 | 3428 | 5406 |
| 546 | 809 | 646 | 630 | 895 | 664 | 502 | 798 | 4809 | 4074 | 7468 | 6044 | 1363 | 1329 | 2110 |
| 353 | 649 | | | | 607 | 354 | 632 | 3231 | 1872 | 5307 | 3445 | | | |
| 967 | 1128 | 1074 | 1091 | 1167 | 1367 | 672 | 923 | 6898 | 6491 | 6713 | 7573 | 5702 | 5792 | 5311 |
| 667 | 1074 | 834 | 602 | 1078 | 764 | 486 | 764 | 3291 | 2843 | 4263 | 4580 | 1168 | 844 | 1401 |
| 414 | 832 | | | | 958 | 481 | 951 | 9326 | 6298 | 15202 | 12650 | | | |
| 433 | 910 | | | | 911 | 687 | 1183 | 2210 | 1353 | 3125 | 2843 | | | |
| 671 | 983 | 816 | 606 | 762 | 1070 | 738 | 1005 | 1802 | 1248 | 1860 | 1828 | 8094 | 6005 | 9916 |
| 456 | 929 | | | | 593 | 414 | 891 | 3227 | 2490 | 5456 | 5068 | | | |
| | | 746 | 285 | 492 | 657 | 265 | 476 | | | | | 5488 | 2097 | 7356 |
| 592 | 904 | 719 | 529 | 970 | 819 | 656 | 1163 | 9471 | 7193 | 12157 | 10988 | 4036 | 2974 | 5617 |
| 418 | 903 | 1161 | 1018 | 1376 | | | | 1812 | 1094 | 2615 | 2361 | 5662 | 4964 | 4877 |
| 844 | 1161 | 1146 | 883 | 1114 | 994 | 903 | 1346 | 4706 | 3575 | 4235 | 4919 | 4840 | 3728 | 4224 |
| 180 | 526 | 142 | 153 | 495 | 180 | 210 | 499 | 2520 | 2188 | 12143 | 6390 | 723 | 778 | 5089 |
| 799 | 1435 | 1016 | 879 | 1283 | 994 | 881 | 1369 | 2204 | 2001 | 2504 | 3593 | 5055 | 4370 | 4973 |
| 572 | 949 | | | | 786 | 568 | 881 | 7127 | 5326 | 9306 | 8831 | | | |
| 532 | 887 | 1058 | 736 | 787 | 1138 | 686 | 821 | 3745 | 2266 | 4259 | 3777 | 8611 | 5992 | 8139 |
| 511 | 1056 | 679 | 599 | 996 | 618 | 509 | 887 | 3761 | 2843 | 5567 | 5881 | 1035 | 913 | 1524 |
| 829 | 998 | 907 | 592 | 929 | 910 | 524 | 922 | 2340 | 1808 | 2179 | 2176 | 4422 | 2885 | 4877 |
| 640 | 1087 | 968 | 583 | 976 | 943 | 760 | 1408 | 1413 | 913 | 1426 | 1551 | 1152 | 694 | 1190 |
| 624 | 969 | 881 | 630 | 947 | | | | 4792 | 3540 | 5672 | 5493 | 5294 | 3784 | 6009 |
| 998 | 1419 | 833 | 819 | 1440 | 711 | 557 | 1285 | 3865 | 4505 | 4516 | 6409 | 7301 | 7180 | 8768 |
| 729 | 1035 | 866 | 540 | 1050 | 1016 | 769 | 1095 | 2675 | 1958 | 2687 | 2782 | 2427 | 1514 | 2803 |

| | | | | | | | | | | | | | | |
|---|---|---|---|---|---|---|---|---|---|---|---|---|---|---|
| 660 | 861 | 944 | 649 | 1068 | 1011 | 798 | 1182 | 1471 | 886 | 1341 | 1154 | 6738 | 4631 | 7137 |
| 356 | 730 | 552 | 465 | 858 | 775 | 581 | 940 | 8457 | 5891 | 16535 | 12078 | 4718 | 3969 | 8540 |
| 719 | 1046 | 1026 | 888 | 1159 | 1102 | 1138 | 1303 | 895 | 694 | 965 | 1010 | 5493 | 4754 | 5354 |
| 560 | 805 | 2499 | 1304 | 628 | | | | 2553 | 1158 | 2068 | 1666 | 3480 | 1816 | 1392 |
| 350 | 692 | 559 | 364 | 847 | 754 | 604 | 959 | 3332 | 1977 | 5650 | 3911 | 7435 | 4840 | 13294 |
| 636 | 851 | 866 | 617 | 838 | 853 | 598 | 811 | 8721 | 5898 | 9271 | 7893 | 6855 | 4886 | 7912 |
| 813 | 1249 | 905 | 809 | 1269 | 927 | 825 | 1229 | 6219 | 5332 | 6559 | 8194 | 4719 | 4218 | 5213 |
| | | 829 | 634 | 985 | 709 | 623 | 1084 | | | | | 2948 | 2254 | 3558 |
| 669 | 880 | 880 | 782 | 1059 | 949 | 734 | 904 | 6092 | 4789 | 7163 | 6300 | 1977 | 1756 | 2246 |
| 271 | 612 | 532 | 316 | 628 | | | | 3667 | 2009 | 7426 | 4543 | 7711 | 4581 | 14495 |
| 477 | 905 | | | | 549 | 400 | 934 | 4618 | 3007 | 6298 | 5697 | | | |
| 284 | 880 | 387 | 271 | 822 | 435 | 282 | 806 | 4487 | 2967 | 10444 | 9189 | 4193 | 2942 | 10841 |
| 833 | 1248 | 1019 | 925 | 1134 | 864 | 741 | 975 | 6495 | 5578 | 6700 | 8360 | 1896 | 1721 | 1861 |
| 981 | 1346 | 886 | 924 | 1327 | 920 | 883 | 1187 | 1223 | 1327 | 1352 | 1821 | 2653 | 2769 | 2996 |
| 592 | 847 | 829 | 500 | 804 | 1003 | 589 | 777 | 7331 | 4617 | 7802 | 6611 | 2851 | 1720 | 3439 |
| 835 | 1249 | 1040 | 762 | 1161 | 1024 | 802 | 1143 | 1402 | 1157 | 1386 | 1731 | 1198 | 878 | 1152 |
| 846 | 989 | 1080 | 936 | 1100 | 707 | 486 | 894 | 3615 | 3037 | 3592 | 3553 | 4701 | 4077 | 4355 |
| 677 | 1024 | 807 | 525 | 751 | | | | 4118 | 3150 | 4655 | 4769 | 1172 | 762 | 1452 |
| 158 | 222 | 1190 | 10 | 10 | 1107 | 10 | 20 | 8952 | 1232 | 7779 | 1730 | 28390 | | 23849 |
| 354 | 649 | 508 | 368 | 600 | 538 | 343 | 728 | 2052 | 1446 | 4080 | 2647 | 760 | 550 | 1497 |
| | | | | | 770 | 522 | 838 | | | | | | | |
| 732 | 1046 | 1188 | 961 | 1014 | 1365 | 1296 | 1220 | 738 | 567 | 773 | 809 | 1252 | 1013 | 1054 |
| 467 | 754 | 695 | 493 | 844 | 812 | 558 | 959 | 4143 | 2822 | 6043 | 4559 | 2901 | 2059 | 4176 |
| 483 | 629 | 736 | 565 | 732 | 665 | 542 | 748 | 1215 | 908 | 1880 | 1182 | 4623 | 3552 | 6283 |
| 546 | 846 | 704 | 582 | 891 | | | | 1809 | 1553 | 2847 | 2410 | 1339 | 1107 | 1901 |
| 537 | 732 | 689 | 526 | 895 | 706 | 458 | 1059 | 897 | 635 | 1182 | 866 | 3324 | 2536 | 4822 |
| | | 1287 | 588 | 806 | 1050 | 623 | 845 | | | | | 12444 | 5683 | 9670 |
| 374 | 790 | 1286 | 350 | 763 | 1303 | 385 | 823 | 4715 | 1408 | 3761 | 2972 | 7022 | 1910 | 5459 |
| 547 | 792 | 712 | 565 | 765 | 673 | 503 | 778 | 2182 | 1855 | 3389 | 2683 | 4739 | 3756 | 6651 |
| 526 | 736 | 1094 | 750 | 987 | | | | 11934 | 6822 | 12965 | 9543 | 358 | 245 | 327 |
| 591 | 1029 | | | | 1010 | 672 | 898 | 3909 | 3129 | 5298 | 5450 | | | |

| | | | | | | | | | | | | | | |
|---|---|---|---|---|---|---|---|---|---|---|---|---|---|---|
| 972 | 1294 | 987 | 885 | 1131 | | | | 1862 | 1804 | 1856 | 2402 | 2166 | 1943 | 2194 |
| 525 | 759 | 649 | 286 | 576 | 931 | 563 | 787 | 3192 | 1988 | 3789 | 2877 | 647 | 285 | 997 |
| 700 | 1111 | | | | 829 | 795 | 1097 | 2937 | 2595 | 3708 | 4118 | | | |
| 603 | 861 | 732 | 697 | 900 | | | | 6600 | 5500 | 9125 | 7860 | 357 | 340 | 487 |
| 369 | 553 | 650 | 455 | 654 | | | | 2196 | 1394 | 3779 | 2092 | 3772 | 2642 | 5801 |
| 1054 | 1475 | 1027 | 972 | 1529 | 1076 | 1170 | 1739 | 1607 | 1623 | 1540 | 2272 | 1426 | 1348 | 1387 |
| 1898 | 1877 | 1299 | 2042 | 2197 | 1515 | 2423 | 2601 | 1022 | 1527 | 805 | 1510 | 1360 | 2139 | 1047 |
| 511 | 958 | 1102 | 654 | 1032 | | | | 5582 | 3397 | 6655 | 6378 | 2537 | 1507 | 2303 |
| 756 | 1049 | 1132 | 605 | 847 | 1157 | 801 | 1054 | 7750 | 4547 | 6015 | 6309 | 4926 | 2632 | 4353 |
| | | 693 | 489 | 872 | 1005 | 756 | 1063 | | | | | 425 | 300 | 612 |
| 406 | 836 | 642 | 416 | 836 | 610 | 389 | 862 | 2451 | 1580 | 3895 | 3257 | 1477 | 957 | 2302 |
| 884 | 1190 | 887 | 732 | 1077 | 944 | 749 | 1114 | 1736 | 1675 | 1895 | 2256 | 1202 | 992 | 1355 |
| | | 927 | 571 | 996 | 1374 | 1103 | 1159 | | | | | 901 | 555 | 972 |
| 214 | 331 | 1679 | 2080 | 1569 | 775 | 689 | 1061 | 1471 | 1172 | 5477 | 1815 | 1036 | 1283 | 617 |
| 283 | 743 | 472 | 361 | 763 | 708 | 513 | 796 | 1634 | 896 | 3167 | 2352 | 1054 | 806 | 2233 |
| 792 | 1057 | | | | 1145 | 900 | 1188 | 5215 | 4142 | 5227 | 5527 | | | |
| 790 | 942 | 790 | 501 | 803 | | | | 1393 | 1226 | 1552 | 1462 | 9555 | 6063 | 12091 |
| | | 926 | 622 | 838 | 1024 | 735 | 1142 | | | | | 1139 | 764 | 1230 |
| 481 | 631 | 839 | 409 | 654 | 1017 | 589 | 745 | 4177 | 2155 | 4476 | 2825 | 5458 | 2656 | 6501 |
| | | 584 | 353 | 652 | | | | | | | | 5205 | 3149 | 8917 |
| 609 | 948 | 1039 | 789 | 1221 | | | | 2743 | 1686 | 2767 | 2624 | 1888 | 1434 | 1817 |
| 492 | 705 | 1074 | 703 | 912 | 1138 | 874 | 1107 | 4939 | 2917 | 5926 | 4176 | 1895 | 1240 | 1765 |
| 992 | 1371 | 966 | 949 | 1350 | 923 | 871 | 1266 | 888 | 955 | 963 | 1320 | 712 | 699 | 737 |
| 623 | 1013 | 1318 | 739 | 948 | 1106 | 650 | 1125 | 1563 | 991 | 1590 | 1612 | 2864 | 1605 | 2173 |
| 478 | 663 | | | | | | | 1727 | 1217 | 2546 | 1688 | | | |
| | | | | | 703 | 442 | 807 | | | | | | | |
| 673 | 985 | 724 | 696 | 1073 | 1040 | 855 | 1134 | 8103 | 6962 | 10341 | 10188 | 2554 | 2459 | 3530 |
| 518 | 934 | 751 | 591 | 1006 | 741 | 597 | 942 | 1767 | 1268 | 2447 | 2286 | 4809 | 3786 | 6406 |
| 429 | 761 | 1030 | 523 | 818 | 1134 | 632 | 879 | 12334 | 6107 | 14225 | 10827 | 9973 | 5063 | 9686 |
| 233 | 687 | | | | 466 | 287 | 726 | 1198 | 872 | 3745 | 2573 | | | |
| 266 | 616 | 497 | 354 | 678 | | | | 4998 | 3954 | 14866 | 9154 | 4748 | 3383 | 9556 |

| | | | | | | | | | | | | | | |
|---|---|---|---|---|---|---|---|---|---|---|---|---|---|---|
| 397 | 678 | | | | 558 | 414 | 552 | 321 | 335 | 843 | 572 | | | |
| | | 681 | 475 | 892 | 699 | 436 | 863 | | | | | 3552 | 2476 | 5214 |
| 725 | 795 | | | | 870 | 558 | 760 | 5278 | 4423 | 6103 | 4853 | | | |
| 297 | 737 | 482 | 282 | 705 | 627 | 337 | 622 | 3247 | 1948 | 6550 | 4828 | 2312 | 1352 | 4797 |
| | | 922 | 547 | 816 | 956 | 556 | 849 | | | | | 2215 | 1315 | 2403 |
| 445 | 893 | 581 | 439 | 836 | | | | 2280 | 1779 | 3999 | 3570 | 8285 | 6261 | 14251 |
| 658 | 1035 | 820 | 735 | 1041 | | | | 10695 | 10000 | 15196 | 15735 | 4993 | 4476 | 6089 |
| 860 | 1129 | | | | 1045 | 805 | 986 | 659 | 547 | 637 | 718 | | | |
| 526 | 850 | 592 | 469 | 799 | 626 | 410 | 732 | 3081 | 2446 | 4647 | 3949 | 3033 | 2401 | 5122 |
| | | 854 | 684 | 1086 | 842 | 698 | 1109 | | | | | 1738 | 1392 | 2035 |
| 274 | 690 | | | | 397 | 269 | 717 | 4234 | 2807 | 10238 | 7064 | | | |
| 733 | 928 | 1012 | 865 | 1164 | 659 | 703 | 934 | 7606 | 6872 | 9370 | 8699 | 1089 | 931 | 1076 |
| 491 | 840 | 576 | 412 | 741 | 652 | 456 | 807 | 1769 | 1171 | 2383 | 2001 | 1535 | 1097 | 2664 |
| 869 | 1043 | 928 | 720 | 992 | 851 | 507 | 768 | 1863 | 1576 | 1814 | 1892 | 2250 | 1746 | 2425 |
| | | 692 | 523 | 831 | 527 | 335 | 689 | | | | | 2406 | 1818 | 3478 |
| 504 | 775 | 831 | 723 | 1062 | 775 | 645 | 1086 | 4192 | 2903 | 5766 | 4466 | 4134 | 3597 | 4975 |
| 711 | 1003 | 1041 | 797 | 1090 | | | | 2130 | 1155 | 1625 | 1630 | 4395 | 3364 | 4221 |
| 532 | 749 | 830 | 559 | 866 | 884 | 589 | 884 | 1528 | 1060 | 1993 | 1493 | 1422 | 958 | 1713 |
| 658 | 904 | 930 | 627 | 926 | 832 | 559 | 882 | 3134 | 2190 | 3330 | 3011 | 1378 | 930 | 1482 |
| 788 | 1071 | 1146 | 896 | 1165 | 1130 | 874 | 1240 | 1086 | 792 | 1004 | 1075 | 1200 | 937 | 1047 |
| | | 911 | 571 | 951 | 962 | 590 | 765 | | | | | 1520 | 953 | 1668 |
| 699 | 1019 | 818 | 694 | 1087 | 944 | 740 | 1091 | 1836 | 1636 | 2339 | 2384 | 2041 | 1734 | 2496 |
| | | 723 | 312 | 720 | 682 | 372 | 780 | | | | | 7712 | 3327 | 10667 |
| 686 | 1333 | 1130 | 640 | 1370 | 1107 | 627 | 1108 | 1759 | 1068 | 1558 | 2077 | 3850 | 2180 | 3408 |
| 614 | 870 | 941 | 696 | 883 | 549 | 474 | 877 | 3049 | 2275 | 3703 | 3223 | 4605 | 3407 | 4895 |
| 479 | 847 | 655 | 491 | 772 | | | | 12157 | 8018 | 16745 | 14181 | 6634 | 4976 | 10125 |
| 583 | 1058 | | | | 717 | 420 | 644 | 1299 | 995 | 1707 | 1806 | | | |
| 1048 | 1611 | 1000 | 747 | 1381 | | | | 9050 | 7905 | 7544 | 12153 | 484 | 362 | 484 |
| 541 | 936 | 751 | 504 | 836 | 993 | 271 | 519 | 12050 | 8210 | 15182 | 14209 | 5537 | 3719 | 7374 |
| 780 | 901 | | | | 1149 | 721 | 737 | 5556 | 3524 | 4519 | 4072 | | | |
| 349 | 683 | 597 | 360 | 709 | 628 | 404 | 686 | 4452 | 2615 | 7484 | 5109 | 4539 | 2742 | 7608 |

| | | | | | | | | | | | | | | |
|---|---|---|---|---|---|---|---|---|---|---|---|---|---|---|
| 368 | 835 | | | | 765 | 466 | 815 | 1534 | 781 | 2122 | 1771 | | | |
| 689 | 1068 | 1096 | 834 | 899 | | | | 739 | 599 | 870 | 929 | 5650 | 4298 | 5155 |
| 777 | 1109 | | | | 1142 | 968 | 1112 | 1674 | 1336 | 1720 | 1907 | | | |
| 814 | 1077 | | | | 774 | 673 | 1005 | 4330 | 4246 | 5219 | 5619 | | | |
| 509 | 697 | 519 | 395 | 734 | | | | 3330 | 2645 | 5200 | 3627 | 2036 | 1547 | 3920 |
| 661 | 1000 | 932 | 704 | 1107 | 949 | 647 | 971 | 2839 | 2087 | 3158 | 3158 | 2783 | 2102 | 2986 |
| 436 | 788 | 723 | 410 | 792 | | | | 5755 | 4423 | 10134 | 7986 | 1711 | 970 | 2366 |
| 270 | 616 | 548 | 336 | 634 | | | | 569 | 294 | 1091 | 672 | 1762 | 1079 | 3214 |
| 1055 | 1291 | 906 | 711 | 1112 | 1001 | 892 | 1242 | 1016 | 972 | 921 | 1189 | 6654 | 5221 | 7346 |
| 826 | 1067 | 1048 | 728 | 1033 | 1075 | 931 | 1050 | 6290 | 5254 | 6362 | 6790 | 4592 | 3190 | 4383 |
| 373 | 543 | 878 | 395 | 507 | 770 | 385 | 702 | 1155 | 502 | 1348 | 732 | 2501 | 1127 | 2850 |
| 777 | 1037 | 826 | 764 | 1081 | 819 | 763 | 1075 | 3911 | 3832 | 4931 | 5114 | 3583 | 3315 | 4340 |
| 1041 | 1247 | 1119 | 928 | 1264 | 1276 | 1141 | 1414 | 893 | 777 | 747 | 931 | 809 | 671 | 723 |
| 620 | 847 | 750 | 583 | 797 | 819 | 636 | 782 | 2525 | 2045 | 3301 | 2797 | 3304 | 2571 | 4405 |
| 263 | 599 | 783 | 553 | 913 | 815 | 574 | 927 | 691 | 417 | 1586 | 950 | 5524 | 3899 | 7051 |
| 560 | 803 | | | | 949 | 698 | 963 | 6793 | 4796 | 8568 | 6882 | | | |
| 645 | 2869 | 993 | 681 | 3343 | 978 | 696 | 952 | 1876 | 1238 | 1920 | 5507 | 2133 | 1463 | 2148 |
| 685 | 1125 | 817 | 675 | 1094 | 844 | 687 | 1066 | 2585 | 1963 | 2864 | 3222 | 3069 | 2535 | 3758 |
| 483 | 849 | | | | 992 | 553 | 866 | 13099 | 6791 | 14059 | 11934 | | | |
| 692 | 1185 | 1013 | 783 | 1212 | | | | 3185 | 2726 | 3940 | 4668 | 2446 | 1890 | 2414 |
| | | 782 | 503 | 729 | 711 | 450 | 775 | | | | | 3493 | 2244 | 4466 |
| 802 | 1170 | 1525 | 1458 | 1631 | 1002 | 789 | 1185 | 2136 | 1718 | 2142 | 2507 | 6828 | 6531 | 4478 |
| 450 | 800 | 776 | 552 | 884 | 904 | 657 | 945 | 4540 | 2804 | 6228 | 4982 | 2583 | 1837 | 3328 |
| 712 | 1110 | | | | 1189 | 1000 | 1205 | 3131 | 2697 | 3787 | 4203 | | | |
| 776 | 1264 | | | | 856 | 789 | 1212 | 4121 | 3333 | 4297 | 5431 | | | |
| 310 | 918 | | | | 464 | 294 | 911 | 5251 | 3488 | 11245 | 10323 | | | |
| 428 | 984 | 530 | 501 | 983 | 603 | 584 | 1051 | 3567 | 2962 | 6924 | 6816 | 6950 | 6573 | 13109 |
| 628 | 915 | 852 | 660 | 929 | | | | 5059 | 3837 | 6107 | 5589 | 2262 | 1754 | 2657 |
| 724 | 1075 | | | | 1126 | 905 | 972 | 3744 | 3279 | 4529 | 4870 | | | |
| 613 | 921 | 705 | 586 | 883 | 504 | 385 | 756 | 3590 | 2951 | 4815 | 4433 | 4571 | 3801 | 6487 |
| 681 | 1062 | 1198 | 724 | 1122 | 1062 | 825 | 1062 | 499 | 320 | 470 | 499 | 545 | 330 | 455 |

| | | | | | | | | | | | | | | |
|---|---|---|---|---|---|---|---|---|---|---|---|---|---|---|
| 344 | 706 | 710 | 409 | 766 | | | | 1311 | 697 | 2028 | 1432 | 421 | 243 | 594 |
| 391 | 642 | 554 | 409 | 609 | 512 | 418 | 597 | 1925 | 1422 | 3636 | 2334 | 629 | 464 | 1136 |
| 501 | 812 | 711 | 461 | 909 | | | | 2607 | 1696 | 3388 | 2749 | 902 | 585 | 1268 |
| 475 | 733 | 1373 | 924 | 1083 | 687 | 495 | 819 | 2710 | 1892 | 3985 | 2922 | 5830 | 3923 | 4248 |
| 714 | 1370 | 956 | 803 | 1262 | 975 | 843 | 1285 | 4531 | 4640 | 6494 | 8893 | 783 | 658 | 819 |
| | | 807 | 619 | 923 | 892 | 750 | 1005 | | | | | 2195 | 1683 | 2720 |
| 672 | 888 | 1025 | 724 | 928 | | | | 7572 | 5922 | 8807 | 7824 | 3630 | 2563 | 3541 |
| 721 | 1138 | 971 | 777 | 1191 | 787 | 550 | 824 | 2445 | 2155 | 2990 | 3403 | 2609 | 2087 | 2688 |
| | | 732 | 625 | 1002 | 736 | 576 | 1012 | | | | | 6445 | 5499 | 8805 |
| 414 | 699 | 598 | 450 | 720 | | | | 1129 | 818 | 1973 | 1380 | 1879 | 1416 | 3144 |
| 481 | 547 | 818 | 542 | 790 | 924 | 650 | 874 | 753 | 456 | 949 | 519 | 4257 | 2820 | 5203 |
| 524 | 818 | 1173 | 810 | 966 | 861 | 626 | 948 | 4626 | 2925 | 5581 | 4564 | 3536 | 2441 | 3014 |
| 893 | 976 | | | | 966 | 798 | 888 | 1650 | 1519 | 1701 | 1660 | | | |
| 408 | 793 | 619 | 399 | 759 | 708 | 368 | 700 | 1870 | 1238 | 3036 | 2408 | 2557 | 1649 | 4130 |
| 898 | 1087 | 1195 | 919 | 1140 | 1384 | 910 | 1371 | 3277 | 2620 | 2917 | 3171 | 3174 | 2440 | 2657 |
| 180 | 541 | 498 | 197 | 555 | 736 | 223 | 544 | 1744 | 630 | 3503 | 1895 | 7458 | 2953 | 14985 |
| 635 | 904 | 844 | 742 | 990 | 747 | 602 | 935 | 3157 | 2532 | 3985 | 3602 | 764 | 672 | 906 |
| | | 1051 | 1038 | 1466 | 1101 | 1014 | 1415 | | | | | 3270 | 3230 | 3112 |
| | | 618 | 497 | 757 | 573 | 435 | 750 | | | | | 3880 | 3121 | 6278 |
| 528 | 937 | 771 | 340 | 540 | | | | 4849 | 3118 | 5905 | 5534 | 4150 | 1832 | 5385 |
| 242 | 1066 | | | | 675 | 394 | 882 | 4994 | 2719 | 11227 | 11972 | | | |
| 735 | 1233 | | | | 802 | 719 | 1225 | 4468 | 3604 | 4906 | 6048 | | | |
| 1437 | 14456 | 1840 | 1578 | 16330 | | | | 1280 | 1102 | 767 | 11085 | 1143 | 980 | 621 |
| 148 | 529 | | | | 524 | 336 | 660 | 1303 | 591 | 3988 | 2109 | | | |
| 358 | 708 | 632 | 406 | 747 | 508 | 368 | 706 | 1285 | 887 | 2475 | 1752 | 709 | 456 | 1122 |
| 894 | 940 | 944 | 865 | 951 | | | | 3101 | 2895 | 3237 | 3042 | 4976 | 4558 | 5270 |
| 686 | 1186 | 771 | 513 | 920 | | | | 2051 | 1452 | 2118 | 2511 | 2387 | 1589 | 3096 |
| 545 | 871 | 1243 | 862 | 1080 | | | | 788 | 630 | 1155 | 1006 | 9303 | 6453 | 7487 |
| 1125 | 1297 | 1093 | 982 | 1325 | 1066 | 993 | 1433 | 1213 | 1198 | 1065 | 1381 | 4161 | 3741 | 3808 |
| 109 | 314 | 127 | 58 | 298 | 149 | 67 | 257 | 701 | 370 | 3413 | 1070 | 2122 | 961 | 16690 |
| | | 622 | 489 | 849 | 707 | 524 | 906 | | | | | 2420 | 1902 | 3889 |

| | | | | | | | | | | | | | | |
|---|---|---|---|---|---|---|---|---|---|---|---|---|---|---|
| | | 1081 | 827 | 1137 | 1071 | 784 | 1131 | | | | | 3764 | 2877 | 3481 |
| 980 | 1083 | 1008 | 823 | 1045 | | | | 1988 | 1580 | 1612 | 1746 | 6634 | 5416 | 6582 |
| 499 | 958 | 674 | 474 | 806 | | | | 1188 | 790 | 1585 | 1519 | 2411 | 1696 | 3579 |
| 368 | 692 | 559 | 316 | 615 | 498 | 255 | 678 | 1965 | 986 | 2681 | 1856 | 536 | 303 | 959 |
| 731 | 1104 | 1024 | 986 | 1413 | | | | 5716 | 4161 | 5690 | 6282 | 1538 | 1480 | 1501 |
| 907 | 1214 | 965 | 766 | 1075 | 1086 | 946 | 1303 | 1427 | 1320 | 1454 | 1765 | 2868 | 2276 | 2971 |
| 878 | 3008 | 756 | 440 | 2236 | 902 | 541 | 4514 | 4022 | 3147 | 3585 | 10785 | 1547 | 901 | 2047 |
| 658 | 1062 | 729 | 687 | 1089 | 710 | 679 | 1064 | 3036 | 2354 | 3576 | 3799 | 3058 | 2884 | 4195 |
| 430 | 954 | 576 | 357 | 952 | 586 | 401 | 894 | 1498 | 1101 | 2561 | 2444 | 1839 | 1140 | 3191 |
| 908 | 1200 | 908 | 690 | 1105 | 953 | 613 | 1031 | 3912 | 3196 | 3521 | 4225 | 2261 | 1719 | 2489 |
| | | 778 | 600 | 896 | | | | | | | | 1386 | 1070 | 1782 |
| 982 | 1487 | 1249 | 1021 | 1308 | 1033 | 797 | 1142 | 5784 | 4657 | 4741 | 7048 | 4064 | 3321 | 3254 |
| 641 | 973 | 885 | 490 | 642 | 901 | 614 | 950 | 2915 | 1853 | 2889 | 2811 | 1344 | 744 | 1518 |
| 651 | 958 | 1276 | 709 | 1066 | 1274 | 734 | 1040 | 9166 | 5096 | 7825 | 7499 | 8270 | 4591 | 6479 |
| 821 | 1157 | 956 | 803 | 1122 | 1012 | 835 | 1165 | 2362 | 1898 | 2311 | 2674 | 2082 | 1749 | 2178 |
| 316 | 590 | 439 | 345 | 645 | 456 | 332 | 606 | 1104 | 837 | 2651 | 1563 | 1135 | 893 | 2585 |
| | | 760 | 503 | 813 | 781 | 622 | 893 | | | | | 1520 | 1006 | 2000 |
| 629 | 991 | 937 | 707 | 1024 | 804 | 675 | 980 | 2440 | 1833 | 2915 | 2890 | 1258 | 950 | 1344 |
| | | 573 | 371 | 788 | 419 | 268 | 720 | | | | | 5583 | 3616 | 9737 |
| 967 | 1410 | 1304 | 1168 | 1699 | | | | 5387 | 4781 | 4942 | 6969 | 5414 | 4849 | 4152 |
| 594 | 807 | 728 | 640 | 806 | 763 | 669 | 877 | 4320 | 3942 | 6637 | 5357 | 2652 | 2332 | 3642 |
| | | 815 | 662 | 994 | 972 | 581 | 894 | | | | | 1909 | 1551 | 2342 |
| 1193 | 1493 | 908 | 1106 | 1309 | | | | 266 | 304 | 255 | 380 | 1834 | 2234 | 2021 |
| | | 1075 | 839 | 1046 | 1007 | 877 | 1325 | | | | | 2457 | 1919 | 2286 |
| 537 | 745 | 466 | 316 | 463 | 575 | 401 | 550 | 1795 | 1202 | 2240 | 1668 | 4388 | 2971 | 9417 |
| 498 | 816 | 808 | 470 | 787 | 670 | 404 | 670 | 5042 | 3213 | 6457 | 5271 | 9911 | 5760 | 12267 |
| 999 | 1307 | | | | 982 | 886 | 1257 | 811 | 836 | 838 | 1094 | | | |
| 183 | 537 | 365 | 381 | 888 | 308 | 194 | 548 | 1454 | 810 | 4420 | 2372 | 4481 | 4673 | 12274 |
| 629 | 1011 | 805 | 526 | 1060 | 882 | 526 | 906 | 2348 | 1469 | 2333 | 2358 | 5667 | 3704 | 7040 |
| 925 | 1252 | 1135 | 907 | 1259 | 1224 | 898 | 1126 | 879 | 693 | 749 | 938 | 829 | 663 | 730 |
| 607 | 917 | 651 | 529 | 893 | | | | 2487 | 2119 | 3494 | 3204 | 1857 | 1511 | 2855 |

| | | | | | | | | | | | | | | |
|---|---|---|---|---|---|---|---|---|---|---|---|---|---|---|
| 633 | 998 | | | | 835 | 623 | 1017 | 2126 | 1732 | 2736 | 2730 | | | |
| 852 | 1145 | | | | 984 | 930 | 1172 | 6199 | 5313 | 6236 | 7137 | | | |
| 880 | 1198 | 1120 | 859 | 1153 | 969 | 754 | 1052 | 2709 | 2152 | 2446 | 2929 | 5174 | 3969 | 4622 |
| 571 | 795 | | | | 671 | 497 | 766 | 2393 | 1781 | 3118 | 2477 | | | |
| | | 932 | 681 | 1018 | 1297 | 1241 | 1240 | | | | | 1282 | 937 | 1375 |
| 826 | 1287 | 1046 | 842 | 1339 | 972 | 776 | 1259 | 3044 | 2650 | 3206 | 4125 | 3953 | 3182 | 3779 |
| 318 | 715 | 436 | 291 | 660 | 446 | 307 | 711 | 918 | 681 | 2140 | 1530 | 983 | 655 | 2255 |
| 865 | 1327 | 1317 | 1155 | 1372 | | | | 2095 | 1873 | 2164 | 2872 | 2095 | 1837 | 1591 |
| | | | 100000 | 100000 | | | | | | | | | 26 | |
| 471 | 917 | | | | 721 | 482 | 879 | 2045 | 1576 | 3342 | 3065 | | | |
| 426 | 604 | 787 | 603 | 809 | 825 | 478 | 924 | 9242 | 5556 | 13028 | 7866 | 3041 | 2329 | 3866 |
| 331 | 518 | 360 | 344 | 494 | 441 | 407 | 613 | 921 | 932 | 2818 | 1460 | 972 | 928 | 2696 |
| 511 | 754 | 558 | 335 | 655 | 705 | 434 | 710 | 1164 | 876 | 1715 | 1292 | 1201 | 721 | 2152 |
| | | 756 | 613 | 883 | 707 | 469 | 976 | | | | | 1800 | 1458 | 2381 |
| 677 | 875 | 890 | 719 | 956 | | | | 4337 | 3273 | 4836 | 4232 | 2240 | 1810 | 2516 |
| 498 | 841 | 1059 | 605 | 853 | | | | 1988 | 1321 | 2652 | 2231 | 2140 | 1222 | 2021 |
| 640 | 1122 | 839 | 662 | 1132 | 788 | 677 | 1053 | 1694 | 1365 | 2131 | 2391 | 2050 | 1617 | 2443 |
| 188 | 519 | 361 | 219 | 502 | 399 | 234 | 556 | 1024 | 525 | 2794 | 1449 | 1116 | 677 | 3090 |
| | | 749 | 369 | 656 | 627 | 587 | 773 | | | | | 1969 | 970 | 2631 |
| 521 | 773 | 1070 | 638 | 798 | 970 | 518 | 712 | 2066 | 1117 | 2146 | 1660 | 6873 | 4101 | 6425 |
| 416 | 754 | 783 | 482 | 762 | | | | 2055 | 1104 | 2651 | 2000 | 4604 | 2835 | 5878 |
| 785 | 913 | 836 | 622 | 916 | 1049 | 760 | 939 | 2945 | 2263 | 2882 | 2630 | 4273 | 3183 | 5113 |
| | | 936 | 785 | 1107 | 820 | 614 | 945 | | | | | 1798 | 1509 | 1922 |
| 635 | 971 | 811 | 534 | 1023 | | | | 10714 | 7648 | 12036 | 11686 | 7197 | 4736 | 8871 |
| 409 | 789 | 1095 | 910 | 1349 | 817 | 647 | 1132 | 3787 | 2151 | 5258 | 4147 | 5943 | 4935 | 5426 |
| 570 | 981 | 813 | 581 | 922 | 809 | 695 | 1052 | 1013 | 717 | 1258 | 1234 | 1138 | 814 | 1400 |
| | | 1192 | 892 | 1210 | 904 | 933 | 798 | | | | | 228 | 171 | 191 |
| 623 | 1085 | 908 | 622 | 1383 | 607 | 443 | 866 | 1205 | 909 | 1458 | 1582 | 6914 | 4737 | 7614 |
| 1351 | 1527 | 1210 | 1209 | 1830 | 1154 | 1224 | 1603 | 1965 | 1822 | 1349 | 2060 | 5198 | 5197 | 4297 |
| 703 | 1213 | | | | 1084 | 741 | 1210 | 1426 | 1225 | 1743 | 2114 | | | |
| 409 | 726 | | | | 675 | 465 | 762 | 1390 | 862 | 2110 | 1532 | | | |

| | | | | | | | | | | | | | | |
|---|---|---|---|---|---|---|---|---|---|---|---|---|---|---|
| | | | | | 807 | 573 | 840 | | | | | | | |
| 492 | 775 | 640 | 478 | 677 | 534 | 498 | 738 | 1934 | 1804 | 3665 | 2839 | 2287 | 1709 | 3573 |
| 760 | 943 | 973 | 725 | 985 | 1220 | 751 | 1020 | 1685 | 1322 | 1740 | 1640 | 2800 | 2085 | 2876 |
| 789 | 968 | 1216 | 1265 | 1021 | | | | 422 | 382 | 484 | 468 | 1464 | 1522 | 1204 |
| | | 745 | 532 | 925 | 817 | 700 | 1237 | | | | | 6755 | 4820 | 9061 |
| 911 | 1117 | | | | 1197 | 1096 | 1122 | 441 | 372 | 408 | 456 | | | |
| | | 1193 | 850 | 971 | 974 | 746 | 659 | | | | | 959 | 683 | 804 |
| 384 | 700 | 584 | 354 | 641 | | | | 2378 | 1354 | 3527 | 2470 | 2296 | 1392 | 3931 |
| | | 790 | 556 | 965 | 811 | 538 | 922 | | | | | 4863 | 3418 | 6153 |
| | | 623 | 503 | 924 | 655 | 548 | 811 | | | | | 5642 | 4562 | 9063 |
| 589 | 1020 | 770 | 556 | 947 | | | | 646 | 471 | 800 | 817 | 685 | 495 | 889 |
| 620 | 941 | 824 | 544 | 870 | | | | 4793 | 3516 | 5670 | 5333 | 2892 | 1910 | 3511 |
| | | 682 | 326 | 648 | 752 | 526 | 741 | | | | | 3047 | 1455 | 4467 |
| 836 | 1707 | 896 | 813 | 1659 | 649 | 501 | 1287 | 1870 | 1677 | 2006 | 3423 | 2225 | 2020 | 2484 |
| 556 | 888 | | | | 1015 | 648 | 869 | 5681 | 3722 | 6695 | 5946 | | | |
| 702 | 933 | | | | 1129 | 779 | 903 | 5415 | 4244 | 6048 | 5643 | | | |
| 788 | 958 | 1245 | 1007 | 1231 | | | | 6023 | 2848 | 3615 | 3462 | 1919 | 1553 | 1542 |
| 972 | 1412 | 1096 | 939 | 1303 | | | | 531 | 493 | 508 | 717 | 3557 | 3046 | 3244 |
| 918 | 1263 | 776 | 756 | 1041 | 1041 | 1051 | 1370 | 3327 | 3088 | 3365 | 4251 | 1428 | 1392 | 1840 |
| 685 | 896 | 875 | 705 | 994 | 910 | 790 | 1059 | 4642 | 3986 | 5823 | 5215 | 3546 | 2860 | 4055 |
| 559 | 861 | | | | | | | 5065 | 3266 | 5841 | 5027 | | | |
| 833 | 894 | | | | 1288 | 935 | 1095 | 340 | 323 | 388 | 347 | | | |
| 523 | 865 | 722 | 486 | 872 | 690 | 495 | 868 | 1524 | 1207 | 2308 | 1996 | 2066 | 1391 | 2861 |
| 200 | 523 | 487 | 186 | 439 | | | | 2698 | 1145 | 5726 | 2998 | 1239 | 473 | 2546 |
| 1170 | 1130 | 1394 | 1020 | 1057 | | | | 6642 | 5112 | 4368 | 4936 | 5572 | 4079 | 3998 |
| | | 451 | 330 | 549 | 423 | 309 | 524 | | | | | 2294 | 1678 | 5086 |
| 696 | 850 | | | | 1067 | 786 | 911 | 3095 | 2218 | 3187 | 2710 | | | |
| 196 | 504 | 643 | 417 | 703 | | | | 395 | 178 | 910 | 459 | 514 | 334 | 800 |
| 532 | 972 | 754 | 568 | 984 | 741 | 500 | 897 | 2398 | 1631 | 3065 | 2978 | 1189 | 896 | 1577 |

| | | | | | | | | | | | | | | |
|---|---|---|---|---|---|---|---|---|---|---|---|---|---|---|
| | | 956 | 836 | 1124 | | | | | | | | 7943 | 6945 | 8310 |
| 983 | 1367 | 1205 | 1002 | 1280 | 1205 | 1014 | 1379 | 5238 | 4291 | 4365 | 5965 | 4938 | 4105 | 4096 |
| 792 | 1117 | 935 | 702 | 975 | | | | 2020 | 1577 | 1992 | 2226 | 3155 | 2368 | 3374 |
| | | 771 | 370 | 637 | 833 | 483 | 789 | | | | | 4677 | 2247 | 6069 |
| 735 | 1077 | | | | 743 | 622 | 1008 | 1989 | 1650 | 2246 | 2418 | | | |
| 458 | 828 | 640 | 478 | 855 | 628 | 507 | 843 | 1820 | 1239 | 2706 | 2240 | 1754 | 1311 | 2743 |
| 181 | 569 | | | | 506 | 279 | 564 | 2044 | 902 | 4990 | 2838 | | | |
| 1005 | 1087 | 746 | 686 | 1104 | 942 | 697 | 1139 | 3409 | 3377 | 3360 | 3654 | 3052 | 2804 | 4089 |
| | | 993 | 808 | 745 | 1023 | 773 | 836 | | | | | 1648 | 1340 | 1659 |
| | | 1132 | 1097 | 1421 | 1091 | 1094 | 1421 | | | | | 3250 | 3149 | 2872 |
| 861 | 955 | 1136 | 907 | 1029 | 1535 | 1473 | 1132 | 2518 | 2103 | 2443 | 2332 | 2160 | 1726 | 1902 |
| 791 | 1200 | | | | 871 | 654 | 1004 | 2186 | 1634 | 2066 | 2478 | | | |
| 985 | 1305 | 912 | 753 | 1080 | 1016 | 925 | 1169 | 2616 | 2417 | 2454 | 3201 | 1456 | 1201 | 1595 |
| 706 | 986 | 912 | 619 | 996 | 979 | 645 | 935 | 1221 | 850 | 1204 | 1186 | 1336 | 907 | 1465 |
| 419 | 786 | 1171 | 742 | 1093 | 1144 | 756 | 1039 | 3133 | 1832 | 4367 | 3431 | 4611 | 2920 | 3937 |
| | | 726 | 491 | 710 | 758 | 421 | 832 | | | | | 3568 | 2413 | 4917 |
| | | 698 | 621 | 898 | 838 | 679 | 907 | | | | | 1233 | 1098 | 1767 |
| 451 | 631 | 1277 | 441 | 725 | 1257 | 506 | 717 | 3774 | 1324 | 2934 | 1852 | 3246 | 1120 | 2541 |
| | | | | | 958 | 722 | 1102 | | | | | | | |
| 631 | 903 | 961 | 618 | 1000 | 1232 | 635 | 964 | 505 | 312 | 495 | 447 | 791 | 508 | 823 |
| | | 467 | 290 | 704 | | | | | | | | 382 | 238 | 819 |
| 514 | 723 | | | | 823 | 506 | 650 | 749 | 417 | 812 | 587 | | | |
| 611 | 785 | 755 | 602 | 1004 | | | | 1773 | 1338 | 2191 | 1720 | 511 | 407 | 676 |
| 608 | 863 | | | | 788 | 767 | 1021 | 1336 | 1269 | 2087 | 1801 | | | |
| 935 | 1487 | 774 | 837 | 1196 | 908 | 640 | 1124 | 4646 | 4729 | 5060 | 7526 | 862 | 932 | 1114 |
| 745 | 1146 | | | | | | | 2685 | 2143 | 2876 | 3297 | | | |
| 558 | 962 | 734 | 556 | 973 | 718 | 491 | 909 | 3763 | 3084 | 5529 | 5318 | 2327 | 1764 | 3170 |
| | | 805 | 389 | 643 | 490 | 308 | 626 | | | | | 4620 | 2232 | 5737 |
| 919 | 1223 | 788 | 740 | 1327 | 1027 | 900 | 1271 | 2368 | 2191 | 2382 | 2915 | 2654 | 2493 | 3368 |
| 449 | 1079 | | | | 653 | 446 | 1010 | 7589 | 5792 | 12898 | 13918 | | | |
| 719 | 1010 | 1320 | 852 | 1044 | 1125 | 750 | 982 | 2208 | 1556 | 2165 | 2187 | 2424 | 1566 | 1837 |

| | | | | | | | | | | | | | | |
|---|---|---|---|---|---|---|---|---|---|---|---|---|---|---|
| 807 | 1405 | | | | 1001 | 778 | 804 | 3365 | 3638 | 4510 | 6339 | | | |
| 464 | 817 | 549 | 518 | 803 | | | | 1743 | 1331 | 2868 | 2344 | 3607 | 3401 | 6569 |
| 967 | 1532 | | | | 565 | 488 | 1187 | 5125 | 5475 | 5659 | 8669 | | | |
| 637 | 1026 | 957 | 697 | 986 | 888 | 611 | 884 | 1558 | 1127 | 1769 | 1816 | 3276 | 2387 | 3424 |
| 496 | 836 | 1486 | 875 | 1180 | | | | 4846 | 2908 | 5860 | 4899 | 2045 | 1203 | 1376 |
| | | 767 | 746 | 953 | | | | | | | | 3430 | 3335 | 4472 |
| | | 689 | 445 | 744 | | | | | | | | 2516 | 1623 | 3649 |
| | | 893 | 711 | 888 | 1032 | 1005 | 975 | | | | | 1664 | 1326 | 1864 |
| 761 | 1120 | | | | 875 | 643 | 957 | 2870 | 2539 | 3336 | 3734 | | | |
| 863 | 1064 | | | | 742 | 548 | 912 | 866 | 635 | 736 | 783 | | | |
| | | 614 | 412 | 702 | 794 | 545 | 817 | | | | | 2173 | 1460 | 3542 |
| 338 | 593 | | | | 698 | 597 | 720 | 1781 | 1533 | 4536 | 2689 | | | |
| 907 | 1346 | | | | 1228 | 910 | 1888 | 1892 | 1568 | 1728 | 2325 | | | |
| 539 | 889 | | | | 698 | 572 | 948 | 13461 | 9482 | 17580 | 15621 | | | |
| 940 | 957 | 900 | 992 | 963 | 739 | 732 | 797 | 2640 | 2699 | 2870 | 2747 | 815 | 898 | 906 |
| 647 | 1009 | | | | 593 | 370 | 694 | 2195 | 1736 | 2682 | 2706 | | | |
| 578 | 682 | 382 | 372 | 603 | 572 | 518 | 641 | 2631 | 2355 | 4073 | 2777 | 1892 | 1843 | 4960 |
| | | 819 | 659 | 933 | 984 | 865 | 1152 | | | | | 4232 | 3406 | 5170 |
| 1056 | 1419 | 1224 | 1711 | 2397 | 1188 | 1621 | 2354 | 2458 | 2813 | 2665 | 3782 | 2634 | 3681 | 2152 |
| 669 | 1220 | | | | 1011 | 807 | 1083 | 81 | 67 | 101 | 123 | | | |
| | | 1004 | 879 | 986 | 955 | 729 | 919 | | | | | 1543 | 1351 | 1537 |
| 952 | 1368 | 951 | 864 | 1325 | 1038 | 937 | 1425 | 3278 | 3073 | 3228 | 4417 | 5688 | 5164 | 5979 |
| | | 481 | 251 | 602 | 466 | 230 | 576 | | | | | 4230 | 2209 | 8799 |
| 697 | 1024 | 868 | 677 | 966 | 820 | 668 | 885 | 3061 | 2310 | 3314 | 3394 | 2551 | 1989 | 2938 |
| 535 | 860 | 645 | 527 | 839 | 700 | 656 | 926 | 4699 | 3743 | 6999 | 6019 | 2603 | 2129 | 4038 |
| 439 | 691 | 562 | 435 | 827 | | | | 7609 | 5787 | 13190 | 9114 | 5333 | 4132 | 9492 |
| 430 | 756 | 764 | 612 | 880 | 618 | 523 | 792 | 3027 | 1675 | 3890 | 2942 | 3872 | 3099 | 5067 |
| | | 419 | 318 | 782 | 1152 | 980 | 1126 | | | | | 754 | 572 | 1802 |
| 369 | 702 | 616 | 368 | 715 | 569 | 390 | 729 | 1174 | 638 | 1732 | 1215 | 1135 | 679 | 1844 |
| 767 | 830 | 1398 | 526 | 553 | | | | 239 | 180 | 235 | 195 | 2224 | 836 | 1591 |
| 556 | 885 | | | | 761 | 539 | 882 | 2069 | 1360 | 2445 | 2164 | | | |

| | | | | | | | | | | | | | | |
|---|---|---|---|---|---|---|---|---|---|---|---|---|---|---|
| | | 902 | 720 | 1046 | | | | | | | | 3428 | 2738 | 3801 |
| 390 | 523 | 674 | 498 | 739 | 588 | 336 | 512 | 691 | 431 | 1105 | 578 | 2194 | 1623 | 3256 |
| 370 | 564 | 991 | 309 | 566 | 997 | 329 | 547 | 241 | 113 | 306 | 172 | 1395 | 435 | 1408 |
| | | 950 | 766 | 992 | 852 | 809 | 1128 | | | | | 952 | 768 | 1002 |
| 757 | 1231 | | | | 908 | 852 | 1187 | 5719 | 3848 | 5081 | 6253 | | | |
| 710 | 924 | 1028 | 747 | 987 | | | | 4623 | 2951 | 4155 | 3839 | 4284 | 3115 | 4169 |
| | | 418 | 195 | 659 | 539 | 299 | 817 | | | | | 4498 | 2094 | 10761 |
| 574 | 986 | 808 | 572 | 952 | 824 | 557 | 942 | 749 | 528 | 919 | 906 | 1240 | 878 | 1535 |
| 1156 | 1122 | 1178 | 1137 | 1174 | 1254 | 1124 | 1261 | 1323 | 1307 | 1130 | 1269 | 1283 | 1238 | 1089 |
| 826 | 1591 | | | | | | | 2657 | 2449 | 2967 | 4721 | | | |
| | | | | | 680 | 573 | 977 | | | | | | | |
| 269 | 673 | | | | | | | 5407 | 3795 | 14094 | 9480 | | | |
| 770 | 1227 | | | | | | | 9666 | 8770 | 11382 | 13967 | | | |
| 765 | 1090 | | | | 700 | 499 | 745 | 3422 | 2870 | 3750 | 4089 | | | |
| 1069 | 1286 | 949 | 753 | 1021 | 548 | 381 | 652 | 2588 | 2348 | 2197 | 2825 | 7053 | 5597 | 7430 |
| 318 | 646 | | | | 502 | 325 | 580 | 4688 | 2994 | 9421 | 6090 | | | |
| | | | | | 1049 | 991 | 1094 | | | | | | | |
| 916 | 1148 | | | | 984 | 923 | 1286 | 2163 | 1994 | 2177 | 2499 | | | |
| 490 | 949 | | | | 869 | 512 | 811 | 4786 | 2978 | 6073 | 5765 | | | |
| 605 | 894 | 808 | 572 | 828 | | | | 2062 | 1555 | 2572 | 2300 | 2474 | 1752 | 3064 |
| | | 995 | 687 | 879 | | | | | | | | 1215 | 838 | 1221 |
| 478 | 928 | 644 | 475 | 853 | 564 | 425 | 863 | 1253 | 1013 | 2121 | 1969 | 1412 | 1040 | 2191 |
| | | 802 | 566 | 984 | 1069 | 931 | 1275 | | | | | 8346 | 5893 | 10413 |
| 651 | 1084 | | | | 633 | 769 | 1541 | 4403 | 4762 | 7313 | 7924 | | | |
| 727 | 1005 | 1039 | 925 | 1519 | 987 | 603 | 1085 | 1380 | 958 | 1317 | 1324 | 2571 | 2290 | 2475 |
| 1281 | 1582 | 1218 | 1095 | 1144 | | | | 535 | 518 | 405 | 640 | 784 | 706 | 644 |
| | | | | | 777 | 481 | 803 | | | | | | | |
| 251 | 649 | 447 | 284 | 657 | 347 | 194 | 580 | 3312 | 2039 | 8111 | 5262 | 3157 | 2008 | 7066 |
| 563 | 801 | | | | 497 | 420 | 723 | 1869 | 1692 | 3006 | 2408 | | | |
| 493 | 802 | 1004 | 577 | 765 | | | | 2916 | 1448 | 2937 | 2356 | 2556 | 1469 | 2545 |
| 610 | 981 | 738 | 561 | 1267 | 964 | 805 | 1211 | 4107 | 3146 | 5156 | 5058 | 1115 | 848 | 1512 |

| | | | | | | | | | | | | | | |
|---|---|---|---|---|---|---|---|---|---|---|---|---|---|---|
| 134 | 438 | 328 | 189 | 526 | 294 | 138 | 483 | 2051 | 1056 | 7889 | 3451 | 2184 | 1260 | 6667 |
| 867 | 996 | | | | | | | 1025 | 914 | 1054 | 1050 | | | |
| 313 | 626 | 517 | 311 | 622 | 558 | 313 | 623 | 2044 | 1458 | 4662 | 2917 | 3512 | 2114 | 6797 |
| | | 1203 | 721 | 1036 | 707 | 638 | 1018 | | | | | 3229 | 1935 | 2684 |
| 755 | 912 | 959 | 573 | 686 | 803 | 532 | 659 | 2121 | 1219 | 1614 | 1472 | 701 | 419 | 731 |
| 657 | 1227 | 851 | 653 | 1212 | 791 | 574 | 1115 | 1750 | 1354 | 2059 | 2527 | 2941 | 2257 | 3455 |
| | | 1299 | 1307 | 1333 | 1104 | 1806 | 1400 | | | | | 694 | 698 | 534 |
| | | 839 | 632 | 1015 | | | | | | | | 1996 | 1504 | 2379 |
| 644 | 869 | | | | 1107 | 601 | 905 | 843 | 556 | 863 | 750 | | | |
| 544 | 1049 | 603 | 483 | 937 | | | | 1178 | 858 | 1578 | 1656 | 1974 | 1583 | 3274 |
| 439 | 604 | | | | 729 | 394 | 629 | 1105 | 625 | 1422 | 859 | | | |
| | | 815 | 649 | 1245 | 711 | 640 | 1284 | | | | | 1325 | 1056 | 1625 |
| 609 | 953 | 748 | 476 | 874 | | | | 3283 | 2295 | 3772 | 3595 | 2675 | 1702 | 3577 |
| 880 | 1042 | 1132 | 927 | 1145 | | | | 5987 | 4846 | 5505 | 5736 | 5028 | 4116 | 4441 |
| 345 | 657 | | | | 468 | 409 | 762 | 2826 | 2614 | 7577 | 4980 | | | |
| 527 | 933 | 853 | 598 | 948 | | | | 10322 | 6949 | 13181 | 12303 | 8043 | 5638 | 9424 |
| 742 | 1033 | | | | 1006 | 815 | 1044 | 2826 | 2102 | 2833 | 2928 | | | |
| | | 925 | 661 | 1041 | 685 | 536 | 1052 | | | | | 4334 | 3096 | 4685 |
| 643 | 1098 | 1033 | 802 | 1120 | 906 | 774 | 1116 | 1573 | 1191 | 1853 | 2034 | 4766 | 3701 | 4615 |
| 682 | 1042 | 873 | 716 | 1148 | | | | 5491 | 4554 | 6675 | 6953 | 2607 | 2138 | 2986 |
| 2395 | 1098 | 830 | 3097 | 1163 | 815 | 2642 | 1043 | 1155 | 3443 | 1437 | 1578 | 1405 | 5245 | 1694 |
| 386 | 771 | 592 | 458 | 903 | 591 | 374 | 531 | 4315 | 2745 | 7119 | 5488 | 2351 | 1818 | 3969 |
| 840 | 1039 | 984 | 777 | 958 | 971 | 840 | 993 | 1799 | 1453 | 1731 | 1798 | 1387 | 1095 | 1410 |
| 509 | 843 | | | | 1077 | 640 | 982 | 2412 | 2007 | 3945 | 3325 | | | |
| 448 | 660 | | | | 605 | 398 | 673 | 3271 | 2402 | 5366 | 3543 | | | |
| 395 | 612 | 662 | 430 | 732 | | | | 1868 | 1282 | 3248 | 1987 | 2347 | 1527 | 3548 |
| 758 | 1094 | 1054 | 768 | 1097 | 1052 | 759 | 1039 | 3373 | 2249 | 2967 | 3245 | 4140 | 3016 | 3927 |
| 327 | 651 | 653 | 574 | 1222 | | | | 1576 | 1128 | 3449 | 2246 | 2064 | 1814 | 3162 |
| 593 | 746 | 822 | 579 | 745 | 854 | 626 | 704 | 1693 | 1175 | 1982 | 1479 | 1589 | 1119 | 1933 |
| 697 | 1035 | 790 | 675 | 1063 | | | | 3649 | 3171 | 4553 | 4713 | 3350 | 2860 | 4240 |

| | | | | | | | | | | | | | | |
|---|---|---|---|---|---|---|---|---|---|---|---|---|---|---|
| 861 | 991 | | | | 951 | 828 | 1165 | 1540 | 1183 | 1374 | 1362 | | | |
| 466 | 724 | 900 | 537 | 792 | | | | 1895 | 1248 | 2678 | 1940 | 1337 | 797 | 1485 |
| 307 | 676 | 375 | 290 | 677 | 404 | 272 | 608 | 2249 | 1485 | 4829 | 3264 | 1509 | 1168 | 4025 |
| | | | | | 958 | 584 | 844 | | | | | | | |
| 1231 | 1314 | | | | 998 | 1048 | 1664 | 1881 | 1633 | 1326 | 1743 | | | |
| 496 | 794 | | | | 717 | 516 | 746 | 736 | 528 | 1063 | 843 | | | |
| | | 863 | 729 | 1074 | 856 | 758 | 1087 | | | | | 1304 | 1102 | 1512 |
| 551 | 895 | 806 | 653 | 991 | 814 | 577 | 1016 | 1924 | 1228 | 2229 | 1994 | 1861 | 1508 | 2308 |
| 671 | 983 | 1159 | 824 | 1380 | | | | 2118 | 1687 | 2513 | 2471 | 2082 | 1480 | 1796 |
| | | 550 | 319 | 704 | 688 | 396 | 789 | | | | | 2485 | 1442 | 4517 |
| 582 | 1062 | 756 | 523 | 733 | | | | 1825 | 1194 | 2052 | 2178 | 6665 | 4614 | 8817 |
| | | 912 | 534 | 1015 | 469 | 247 | 558 | | | | | 4883 | 2860 | 5352 |
| | | 334 | 255 | 531 | 291 | 238 | 421 | | | | | 481 | 367 | 1438 |
| 502 | 827 | 631 | 344 | 714 | | | | 3869 | 2667 | 5317 | 4396 | 1719 | 936 | 2723 |
| 560 | 829 | 1130 | 614 | 838 | 1826 | 924 | 912 | 1817 | 1134 | 2026 | 1679 | 2046 | 1112 | 1810 |
| | | 893 | 679 | 868 | | | | | | | | 3038 | 2311 | 3404 |
| 671 | 671 | | | | | | | 2781 | 1783 | 2658 | 1783 | | | |
| 511 | 1025 | 639 | 428 | 988 | 639 | 436 | 958 | 4066 | 2705 | 5295 | 5430 | 8820 | 5911 | 13806 |
| | | 1014 | 659 | 686 | 1076 | 710 | 783 | | | | | 1524 | 991 | 1503 |
| | | 730 | 430 | 969 | 655 | 421 | 790 | | | | | 3125 | 1842 | 4280 |
| 447 | 867 | | | | 584 | 341 | 666 | 1045 | 721 | 1613 | 1398 | | | |
| 438 | 891 | 715 | 447 | 862 | | | | 4394 | 2915 | 6663 | 5937 | 1943 | 1216 | 2718 |
| | | 1032 | 599 | 969 | 1009 | 571 | 890 | | | | | 5324 | 3091 | 5160 |
| 238 | 434 | 825 | 229 | 446 | 1107 | 430 | 606 | 3416 | 996 | 4181 | 1814 | 4431 | 1229 | 5371 |
| | | 732 | 524 | 805 | 486 | 270 | 600 | | | | | 3901 | 2792 | 5332 |
| 887 | 1133 | 1027 | 772 | 1091 | 1184 | 936 | 1267 | 2327 | 2087 | 2352 | 2666 | 2577 | 1936 | 2509 |
| 239 | 586 | 802 | 315 | 623 | | | | 1485 | 491 | 2052 | 1203 | 5032 | 1976 | 6274 |
| 359 | 713 | | | | | | | 6010 | 4048 | 11289 | 8047 | | | |
| 432 | 788 | 809 | 747 | 1320 | | | | 8244 | 4394 | 10169 | 8017 | 2569 | 2373 | 3175 |
| | | 714 | 578 | 803 | | | | | | | | 4638 | 3757 | 6496 |
| 473 | 658 | | | | 704 | 462 | 780 | 509 | 327 | 692 | 455 | | | |

| | | | | | | | | | | | | | | |
|---|---|---|---|---|---|---|---|---|---|---|---|---|---|---|
| 552 | 898 | 942 | 521 | 788 | 1188 | 846 | 842 | 2641 | 1509 | 2736 | 2457 | 3747 | 2075 | 3980 |
| | | 1116 | 1178 | 1138 | 1186 | 1094 | 1053 | | | | | 567 | 598 | 508 |
| | | 988 | 455 | 707 | | | | | | | | 4038 | 1858 | 4088 |
| 452 | 1027 | | | | 1262 | 1124 | 1442 | 2570 | 1567 | 3469 | 3562 | | | |
| 793 | 1176 | 946 | 839 | 1281 | 994 | 798 | 1148 | 1158 | 1007 | 1270 | 1493 | 1317 | 1167 | 1392 |
| 700 | 1360 | 1298 | 957 | 1160 | | | | 3815 | 3010 | 4300 | 5849 | 10405 | 7672 | 8017 |
| 350 | 664 | 512 | 229 | 600 | | | | 1182 | 614 | 1752 | 1164 | 423 | 190 | 827 |
| 483 | 716 | 786 | 567 | 817 | 773 | 572 | 884 | 5978 | 4004 | 8293 | 5936 | 2839 | 2049 | 3611 |
| 422 | 756 | 1022 | 617 | 1105 | | | | 9505 | 5765 | 13668 | 10328 | 3620 | 2185 | 3543 |
| | | 895 | 747 | 1233 | 890 | 812 | 1121 | | | | | 1944 | 1622 | 2171 |
| | | 819 | 973 | 1384 | 795 | 960 | 1543 | | | | | 1849 | 2196 | 2256 |
| 641 | 858 | | | | 850 | 582 | 915 | 3376 | 2602 | 4059 | 3484 | | | |
| | | 835 | 750 | 1561 | 703 | 420 | 927 | | | | | 1424 | 1280 | 1706 |
| | | 785 | 460 | 944 | 695 | 357 | 767 | | | | | 1513 | 887 | 1927 |
| 655 | 886 | | | | 1028 | 840 | 989 | 1400 | 974 | 1488 | 1319 | | | |
| 535 | 836 | | | | 895 | 674 | 1058 | 1020 | 881 | 1646 | 1377 | | | |
| 1214 | 1282 | 826 | 610 | 1030 | | | | 849 | 759 | 625 | 801 | 4502 | 3324 | 5448 |
| 467 | 818 | | | | 729 | 488 | 855 | 981 | 662 | 1417 | 1159 | | | |
| 956 | 1110 | | | | 1288 | 1182 | 1273 | 2471 | 2021 | 2114 | 2347 | | | |
| 631 | 820 | | | | 1066 | 648 | 945 | 10672 | 6292 | 9974 | 8178 | | | |
| 1044 | 984 | | | | | | | 2195 | 1961 | 1878 | 1848 | | | |
| | | | | | 785 | 1082 | 402 | | | | | | | |
| 794 | 1147 | 1270 | 1112 | 1477 | 1012 | 860 | 1175 | 2089 | 1677 | 2113 | 2423 | 4547 | 3978 | 3579 |
| 754 | 851 | 922 | 569 | 863 | | | | 704 | 582 | 772 | 657 | 2634 | 1624 | 2856 |
| 320 | 433 | 305 | 242 | 348 | 364 | 251 | 412 | 1118 | 944 | 2955 | 1281 | 1449 | 1150 | 4746 |
| 471 | 756 | 1025 | 925 | 1386 | | | | 1638 | 1024 | 2172 | 1642 | 2214 | 1998 | 2160 |
| | | 586 | 520 | 640 | | | | | | | | 1759 | 1560 | 3000 |
| 214 | 819 | 239 | 194 | 851 | | | | 1631 | 1216 | 5676 | 4650 | 2464 | 2005 | 10326 |
| | | 921 | 778 | 1074 | | | | | | | | 1471 | 1243 | 1598 |
| | | | | | 1085 | 939 | 1047 | | | | | | | |
| 673 | 1089 | 944 | 709 | 1173 | 1021 | 771 | 1212 | 1474 | 1050 | 1561 | 1699 | 1754 | 1318 | 1859 |

| | | | | | | | | | | | | | | |
|---|---|---|---|---|---|---|---|---|---|---|---|---|---|---|
| | | | | | 1038 | 785 | 1212 | | | | | | | |
| 1018 | 1597 | | | | 1360 | 1210 | 1580 | 3790 | 3814 | 3747 | 5985 | | | |
| 1166 | 1486 | | | | 809 | 695 | 1063 | 1172 | 1118 | 959 | 1425 | | | |
| 382 | 781 | 640 | 478 | 913 | | | | 2609 | 1623 | 4246 | 3317 | 969 | 724 | 1515 |
| | | 1524 | 1399 | 1326 | | | | | | | | 3209 | 2944 | 2105 |
| 860 | 1357 | | | | 1040 | 840 | 1218 | 2050 | 1753 | 2038 | 2765 | | | |
| 974 | 1101 | | | | 953 | 549 | 813 | 475 | 358 | 368 | 405 | | | |
| 837 | 1335 | 818 | 478 | 942 | | | | 2390 | 2255 | 2694 | 3596 | 1505 | 879 | 1840 |
| 546 | 1030 | | | | 735 | 547 | 1132 | 3310 | 2465 | 4517 | 4651 | | | |
| | | | | | 604 | 555 | 799 | | | | | | | |
| 762 | 1124 | 927 | 867 | 1029 | | | | 2010 | 1999 | 2625 | 2950 | 4268 | 3992 | 4604 |
| | | 831 | 743 | 1161 | 867 | 856 | 1283 | | | | | 3347 | 2991 | 4027 |
| | | 699 | 473 | 737 | 651 | 526 | 770 | | | | | 7734 | 5232 | 11071 |
| | | 812 | 440 | 709 | 713 | 319 | 496 | | | | | 5183 | 2808 | 6387 |
| 432 | 696 | 737 | 834 | 905 | 736 | 571 | 710 | 881 | 706 | 1635 | 1138 | 1134 | 1284 | 1539 |
| 360 | 707 | 482 | 348 | 651 | | | | 2293 | 1521 | 4226 | 2987 | 2120 | 1528 | 4395 |
| 662 | 773 | | | | 576 | 368 | 677 | 325 | 251 | 379 | 293 | | | |
| | | 564 | 522 | 668 | | | | | | | | 2098 | 1941 | 3722 |
| 554 | 901 | | | | 791 | 520 | 734 | 1886 | 1329 | 2398 | 2161 | | | |
| | | 1103 | 1062 | 1552 | 1071 | 938 | 1337 | | | | | 3775 | 3634 | 3421 |
| 555 | 810 | 676 | 472 | 770 | | | | 2031 | 1618 | 2915 | 2361 | 639 | 446 | 944 |
| 361 | 570 | 881 | 384 | 734 | 1112 | 434 | 684 | 3562 | 1279 | 3542 | 2018 | 2779 | 1213 | 3156 |
| 951 | 1206 | | | | 898 | 670 | 968 | 2405 | 1947 | 2048 | 2471 | | | |
| | | | | | 1171 | 897 | 800 | | | | | | | |
| 537 | 778 | | | | 779 | 401 | 619 | 2256 | 1323 | 2463 | 1916 | | | |
| | | 716 | 458 | 857 | 637 | 435 | 754 | | | | | 1343 | 858 | 1875 |
| 502 | 999 | 687 | 522 | 1007 | 714 | 494 | 958 | 3948 | 2748 | 5476 | 5470 | 2596 | 1973 | 3781 |
| | | 1069 | 868 | 1471 | 866 | 643 | 1040 | | | | | 5199 | 4222 | 4862 |
| 665 | 1032 | 867 | 738 | 1142 | | | | 1936 | 1538 | 2312 | 2386 | 3008 | 2559 | 3470 |
| 988 | 927 | | | | 1186 | 1340 | 1278 | 2484 | 2089 | 2114 | 1961 | | | |
| | | | | | 610 | 464 | 1024 | | | | | | | |

| | | | | | | | | | | | | | | |
|---|---|---|---|---|---|---|---|---|---|---|---|---|---|---|
| 743 | 812 | | | | 979 | 523 | 744 | 10438 | 6001 | 8072 | 6556 | | | |
| 756 | 1110 | | | | | | | 2676 | 2443 | 3233 | 3590 | | | |
| | | 459 | 349 | 565 | 426 | 333 | 481 | | | | | 1680 | 1278 | 3659 |
| 831 | 1212 | 1130 | 913 | 1397 | | | | 2025 | 1691 | 2036 | 2468 | 1729 | 1396 | 1529 |
| 1182 | 1570 | 1177 | 1333 | 1773 | 1064 | 1093 | 1483 | 2547 | 2803 | 2371 | 3723 | 2169 | 2457 | 1843 |
| 624 | 1030 | 795 | 452 | 570 | | | | 1963 | 1776 | 2846 | 2932 | 2992 | 1702 | 3765 |
| 467 | 629 | | | | 717 | 429 | 657 | 1346 | 959 | 2053 | 1292 | | | |
| | | 923 | 461 | 741 | 1059 | 787 | 1076 | | | | | 2105 | 1052 | 2282 |
| | | 826 | 911 | 1689 | | | | | | | | 2995 | 3304 | 3628 |
| 1161 | 927 | 932 | 1262 | 811 | 929 | 1186 | 902 | 1713 | 1982 | 1708 | 1583 | 1906 | 2580 | 2045 |
| 487 | 892 | 777 | 483 | 798 | 798 | 511 | 841 | 1215 | 704 | 1448 | 1291 | 2017 | 1255 | 2595 |
| 655 | 862 | 268 | 124 | 466 | | | | 4176 | 3651 | 5577 | 4807 | 377 | 174 | 1405 |
| 720 | 1198 | | | | 730 | 744 | 1193 | 3637 | 3341 | 4638 | 5557 | | | |
| | | | | | 609 | 455 | 844 | | | | | | | |
| | | | | | 765 | 524 | 675 | | | | | | | |
| 801 | 957 | | | | 830 | 593 | 813 | 2276 | 2022 | 2522 | 2413 | | | |
| 485 | 842 | | | | 666 | 467 | 864 | 2290 | 1711 | 3531 | 2973 | | | |
| 887 | 601 | 828 | 925 | 730 | 806 | 847 | 628 | 6254 | 6505 | 7335 | 4405 | 7241 | 8090 | 8744 |
| 380 | 582 | 818 | 482 | 701 | | | | 2410 | 1306 | 3438 | 2002 | 4702 | 2771 | 5748 |
| 573 | 762 | | | | 699 | 449 | 627 | 1463 | 878 | 1531 | 1166 | | | |
| | | 768 | 468 | 943 | 760 | 467 | 827 | | | | | 1100 | 670 | 1432 |
| 1020 | 1354 | 1084 | 1039 | 1356 | | | | 1263 | 1176 | 1153 | 1560 | 2193 | 2101 | 2024 |
| | | 378 | 228 | 606 | 394 | 279 | 699 | | | | | 1323 | 799 | 3500 |
| | | 1589 | 1474 | 1000 | 1709 | 1771 | 1234 | | | | | 2049 | 1902 | 1290 |
| 773 | 1167 | 1201 | 1229 | 1412 | | | | 1568 | 1565 | 2025 | 2362 | 2004 | 2050 | 1668 |
| 531 | 877 | | | | 716 | 513 | 884 | 2974 | 2126 | 4000 | 3509 | | | |
| 750 | 1169 | | | | 1146 | 880 | 1206 | 3646 | 2711 | 3616 | 4226 | | | |
| 782 | 1114 | 995 | 674 | 948 | | | | 5026 | 3885 | 4967 | 5533 | 4554 | 3085 | 4576 |
| 644 | 804 | 643 | 586 | 831 | | | | 1398 | 1096 | 1701 | 1368 | 1749 | 1595 | 2720 |
| 797 | 1213 | | | | 954 | 714 | 1227 | 1512 | 1132 | 1419 | 1722 | | | |

| | | | | | | | | | | | | | | |
|---|---|---|---|---|---|---|---|---|---|---|---|---|---|---|
| | | 718 | 620 | 1094 | 794 | 682 | 1154 | | | | | 4987 | 4306 | 6947 |
| | | 1230 | 916 | 1314 | | | | | | | | 4639 | 3455 | 3771 |
| 448 | 811 | 476 | 410 | 733 | 509 | 403 | 786 | 1131 | 936 | 2090 | 1696 | 1293 | 1112 | 2714 |
| 943 | 974 | 664 | 372 | 743 | | | | 1631 | 1371 | 1455 | 1417 | 4111 | 2306 | 6191 |
| 944 | 1239 | 1159 | 936 | 1240 | | | | 920 | 686 | 727 | 901 | 996 | 804 | 859 |
| 980 | 1324 | | | | | | | 1851 | 1770 | 1806 | 2391 | | | |
| 865 | 1111 | 895 | 778 | 1169 | | | | 6109 | 5376 | 6219 | 6910 | 1170 | 1017 | 1307 |
| 966 | 1220 | 1032 | 784 | 1139 | | | | 3256 | 2801 | 2901 | 3540 | 1262 | 960 | 1224 |
| | | 942 | 743 | 1150 | 1031 | 845 | 1253 | | | | | 1514 | 1195 | 1608 |
| 646 | 1034 | | | | | | | 3832 | 2853 | 4414 | 4564 | | | |
| 600 | 690 | 951 | 580 | 724 | 932 | 566 | 670 | 2896 | 1737 | 2893 | 1996 | 2178 | 1328 | 2289 |
| | | 913 | 799 | 1109 | 815 | 623 | 1142 | | | | | 4110 | 3595 | 4499 |
| 537 | 882 | | | | | | | 6222 | 4165 | 7761 | 6844 | | | |
| | | 1008 | 543 | 881 | 999 | 569 | 817 | | | | | 1286 | 693 | 1276 |
| | | 689 | 215 | 394 | 793 | 633 | 790 | | | | | 3778 | 1181 | 5486 |
| 329 | 748 | 608 | 417 | 769 | 579 | 381 | 813 | 2424 | 1363 | 4142 | 3100 | 1721 | 1181 | 2830 |
| | | 776 | 801 | 1344 | | | | | | | | 1007 | 1039 | 1298 |
| 427 | 698 | | | | 583 | 476 | 819 | 4691 | 3111 | 7289 | 5089 | | | |
| 258 | 501 | | | | 843 | 244 | 504 | 5216 | 1559 | 6038 | 3024 | | | |
| 745 | 924 | | | | 654 | 593 | 776 | 2161 | 1855 | 2490 | 2300 | | | |
| 177 | 636 | | | | | | | 1241 | 707 | 3991 | 2540 | | | |
| 622 | 1095 | 846 | 637 | 1155 | 819 | 615 | 1115 | 2042 | 1475 | 2371 | 2597 | 1935 | 1457 | 2287 |
| | | 1240 | 1030 | 940 | 715 | 411 | 685 | | | | | 1901 | 1579 | 1533 |
| | | 828 | 618 | 867 | 935 | 725 | 973 | | | | | 3820 | 2850 | 4611 |
| 1155 | 1521 | 1163 | 845 | 1180 | | | | 720 | 774 | 670 | 1019 | 1398 | 1016 | 1202 |
| 377 | 675 | | | | | | | 3950 | 2123 | 5625 | 3797 | | | |
| 867 | 1146 | | | | | | | 6365 | 4999 | 5763 | 6604 | | | |
| 524 | 858 | | | | 627 | 533 | 885 | 1807 | 1479 | 2824 | 2424 | | | |
| 307 | 607 | 424 | 289 | 605 | 453 | 264 | 563 | 1535 | 999 | 3253 | 1976 | 1279 | 870 | 3014 |
| | | 1078 | 651 | 940 | | | | | | | | 3586 | 2167 | 3327 |

| | | | | | | | | | | | | | | |
|---|---|---|---|---|---|---|---|---|---|---|---|---|---|---|
| 417 | 910 | | | | 623 | 448 | 880 | 1910 | 1250 | 2993 | 2724 | | | |
| 133 | 356 | 524 | 342 | 586 | | | | 515 | 174 | 1311 | 467 | 243 | 158 | 463 |
| | | 1030 | 811 | 1494 | 904 | 624 | 1093 | | | | | 618 | 487 | 600 |
| | | 843 | 439 | 792 | 1018 | 620 | 921 | | | | | 1633 | 851 | 1938 |
| 660 | 1107 | | | | 982 | 643 | 1104 | 3377 | 1997 | 3024 | 3349 | | | |
| 621 | 975 | 1019 | 662 | 941 | | | | 1290 | 938 | 1511 | 1473 | 2116 | 1375 | 2077 |
| 447 | 731 | 428 | 281 | 635 | | | | 2633 | 1722 | 3851 | 2814 | 1199 | 788 | 2801 |
| 208 | 441 | | | | 376 | 225 | 513 | 1652 | 597 | 2871 | 1266 | | | |
| | | 1274 | 1048 | 1339 | 1091 | 881 | 1133 | | | | | 4631 | 3809 | 3635 |
| 123 | 206 | 355 | 72 | 132 | | | | 2754 | 812 | 6587 | 1354 | 2890 | 584 | 8135 |
| 1262 | 951 | 801 | 1496 | 1046 | | | | 867 | 1220 | 967 | 919 | 400 | 746 | 499 |
| 287 | 617 | 537 | 259 | 616 | | | | 3588 | 1916 | 6685 | 4125 | 2981 | 1439 | 5556 |
| 805 | 1167 | 1224 | 1016 | 1231 | | | | 1463 | 1444 | 1795 | 2095 | 3234 | 2683 | 2641 |
| 611 | 996 | 864 | 566 | 872 | | | | 3765 | 2701 | 4423 | 4405 | 2545 | 1665 | 2945 |
| 550 | 752 | | | | | | | 5333 | 3660 | 6649 | 4997 | | | |
| | | | | | 552 | 286 | 682 | | | | | | | |
| | | | | | 1031 | 1019 | 1318 | | | | | | | |
| | | | | | 728 | 713 | 1125 | | | | | | | |
| 493 | 793 | 723 | 530 | 841 | 473 | 349 | 691 | 1975 | 1539 | 3122 | 2476 | 1760 | 1289 | 2434 |
| | | 858 | 804 | 1145 | 1024 | 892 | 1289 | | | | | 634 | 594 | 738 |
| | | 903 | 684 | 953 | 937 | 723 | 957 | | | | | 2380 | 1803 | 2636 |
| 647 | 960 | 912 | 719 | 971 | 850 | 665 | 940 | 5256 | 3927 | 6073 | 5828 | 3769 | 2971 | 4134 |
| | | 1011 | 892 | 1175 | 646 | 449 | 789 | | | | | 1766 | 1558 | 1748 |
| | | 817 | 557 | 789 | | | | | | | | 1461 | 997 | 1789 |
| 247 | 561 | 426 | 216 | 544 | 395 | 257 | 530 | 4302 | 2596 | 10508 | 5892 | 5848 | 2959 | 13729 |
| 686 | 1165 | 847 | 750 | 1245 | | | | 953 | 758 | 1104 | 1286 | 1040 | 921 | 1228 |
| 494 | 703 | | | | 807 | 577 | 815 | 1867 | 1346 | 2722 | 1914 | | | |
| 1192 | 1171 | | | | 865 | 586 | 781 | 1752 | 1646 | 1381 | 1617 | | | |
| 554 | 941 | 1076 | 618 | 930 | | | | 4740 | 2745 | 4956 | 4662 | 2743 | 1576 | 2549 |
| 756 | 1041 | 989 | 781 | 1089 | 1016 | 802 | 1066 | 4162 | 3198 | 4229 | 4403 | 4304 | 3397 | 4351 |

| | | | | | | | | | | | | | | |
|---|---|---|---|---|---|---|---|---|---|---|---|---|---|---|
| 911 | 1159 | | | | 869 | 847 | 1198 | 840 | 881 | 967 | 1120 | | | |
| 807 | 1160 | 723 | 604 | 1025 | 820 | 602 | 900 | 1794 | 1447 | 1794 | 2082 | 1143 | 955 | 1581 |
| 860 | 1172 | 1088 | 1000 | 940 | | | | 3279 | 2678 | 3115 | 3650 | 890 | 818 | 818 |
| 680 | 1018 | 630 | 753 | 1040 | 669 | 782 | 1099 | 1345 | 1371 | 2017 | 2054 | 1513 | 1808 | 2402 |
| 353 | 831 | | | | 321 | 176 | 535 | 1892 | 1137 | 3223 | 2680 | | | |
| 504 | 1125 | | | | 710 | 342 | 705 | 1245 | 836 | 1660 | 1866 | | | |
| 626 | 956 | | | | 972 | 607 | 838 | 4113 | 2520 | 4029 | 3851 | | | |
| 940 | 1069 | 884 | 760 | 1163 | | | | 1719 | 1724 | 1835 | 1962 | 940 | 808 | 1064 |
| 506 | 658 | | | | 1035 | 507 | 681 | 3138 | 1806 | 3566 | 2347 | | | |
| 257 | 476 | 964 | 741 | 1041 | | | | 1020 | 748 | 2914 | 1386 | 1319 | 1013 | 1367 |
| 959 | 1214 | | | | 591 | 506 | 921 | 2894 | 2645 | 2759 | 3349 | | | |
| | | 632 | 368 | 695 | 804 | 645 | 972 | | | | | 512 | 298 | 810 |
| | | 505 | 255 | 653 | | | | | | | | 1990 | 1006 | 3942 |
| | | 805 | 532 | 828 | | | | | | | | 2922 | 1929 | 3628 |
| 619 | 952 | 700 | 566 | 799 | 675 | 448 | 804 | 1762 | 1443 | 2333 | 2219 | 1362 | 1102 | 1945 |
| 504 | 849 | 799 | 603 | 911 | 743 | 612 | 900 | 2424 | 1873 | 3719 | 3157 | 2750 | 2073 | 3440 |
| 342 | 618 | 592 | 378 | 645 | | | | 1173 | 621 | 1817 | 1122 | 2228 | 1424 | 3765 |
| 411 | 731 | | | | 576 | 362 | 650 | 855 | 599 | 1457 | 1065 | | | |
| | | 761 | 625 | 965 | 1203 | 745 | 1118 | | | | | 4011 | 3296 | 5274 |
| | | 904 | 870 | 2402 | 954 | 817 | 2039 | | | | | 593 | 571 | 656 |
| 1007 | 1359 | | | | 990 | 975 | 1405 | 1234 | 1234 | 1226 | 1667 | | | |
| 548 | 929 | | | | 588 | 411 | 806 | 1838 | 1857 | 3392 | 3151 | | | |
| | | 1038 | 675 | 1071 | | | | | | | | 948 | 616 | 914 |
| | | 967 | 721 | 866 | 937 | 778 | 938 | | | | | 1691 | 1261 | 1748 |
| 812 | 1276 | 1143 | 1014 | 1197 | | | | 1560 | 1320 | 1626 | 2075 | 1917 | 1699 | 1677 |
| | | 918 | 723 | 817 | 907 | 677 | 818 | | | | | 2281 | 1796 | 2485 |
| 526 | 800 | | | | 797 | 583 | 1102 | 1680 | 953 | 1810 | 1448 | | | |
| 507 | 733 | 945 | 509 | 707 | 847 | 478 | 695 | 3931 | 2143 | 4226 | 3099 | 4831 | 2603 | 5112 |
| 503 | 776 | 608 | 511 | 784 | | | | 1509 | 1292 | 2565 | 1991 | 1292 | 1084 | 2124 |
| | | 838 | 787 | 979 | | | | | | | | 3048 | 2861 | 3637 |

| | | | | | | | | | | | | | | |
|---|---|---|---|---|---|---|---|---|---|---|---|---|---|---|
| 552 | 776 | 804 | 349 | 749 | | | | 1406 | 1036 | 1877 | 1457 | 3818 | 1658 | 4747 |
| | | 462 | 424 | 638 | | | | | | | | 3583 | 3291 | 7754 |
| | | 856 | 587 | 794 | | | | | | | | 2966 | 2034 | 3464 |
| 502 | 678 | 1398 | 483 | 743 | | | | 4342 | 1487 | 2963 | 2009 | 4465 | 1543 | 3193 |
| 532 | 1015 | | | | 871 | 668 | 1096 | 854 | 519 | 976 | 991 | | | |
| 645 | 988 | | | | | | | 6939 | 6155 | 9536 | 9426 | | | |
| | | 920 | 863 | 1427 | | | | | | | | 3622 | 3396 | 3936 |
| | | 880 | 734 | 1076 | 1052 | 698 | 1068 | | | | | 562 | 469 | 639 |
| | | | | | 1177 | 651 | 1074 | | | | | | | |
| | | 648 | 547 | 1027 | 810 | 828 | 1217 | | | | | 1213 | 1025 | 1873 |
| 344 | 779 | 632 | 314 | 797 | | | | 3265 | 1755 | 5110 | 3979 | 5070 | 2518 | 8026 |
| | | 708 | 411 | 698 | 643 | 391 | 654 | | | | | 4381 | 2540 | 6185 |
| | | 701 | 381 | 706 | 692 | 359 | 622 | | | | | 850 | 462 | 1213 |
| | | 658 | 421 | 685 | 644 | 438 | 675 | | | | | 2509 | 1607 | 3816 |
| | | 1011 | 699 | 985 | 981 | 669 | 991 | | | | | 3784 | 2617 | 3744 |
| 567 | 1031 | | | | 884 | 571 | 952 | 1329 | 955 | 1684 | 1737 | | | |
| 835 | 1213 | | | | 1149 | 1326 | 1801 | 2279 | 2372 | 2841 | 3447 | | | |
| 2271 | 2150 | | | | | | | 3543 | 4405 | 1940 | 4172 | | | |
| | | 900 | 639 | 903 | | | | | | | | 6034 | 4283 | 6704 |
| | | 674 | 424 | 739 | | | | | | | | 3868 | 2433 | 5740 |
| 660 | 1017 | | | | | | | 2535 | 1517 | 2297 | 2336 | | | |
| | | 693 | 394 | 662 | 687 | 358 | 660 | | | | | 2833 | 1611 | 4091 |
| | | | | | 522 | 568 | 686 | | | | | | | |
| | | | | | 921 | 584 | 855 | | | | | | | |
| | | | | | 981 | 741 | 1031 | | | | | | | |
| | | 697 | 452 | 884 | 876 | 662 | 890 | | | | | 4922 | 3190 | 7063 |
| 456 | 744 | | | | 1249 | 1026 | 1000 | 1277 | 771 | 1691 | 1258 | | | |
| 428 | 841 | 794 | 501 | 890 | | | | 4329 | 2855 | 6677 | 5616 | 1734 | 1095 | 2185 |
| | | 562 | 347 | 572 | 605 | 348 | 589 | | | | | 1830 | 1129 | 3256 |
| 535 | 823 | | | | 815 | 412 | 739 | 4955 | 2680 | 5012 | 4125 | | | |
| | | 764 | 526 | 914 | | | | | | | | 1609 | 1106 | 2104 |

| | | | | | | | | | | | | | | |
|---|---|---|---|---|---|---|---|---|---|---|---|---|---|---|
| 579 | 882 | | | | | | | 7477 | 5283 | 9130 | 8051 | | | |
| | | 761 | 645 | 942 | | | | | | | | 2057 | 1744 | 2703 |
| 500 | 827 | 913 | 901 | 1160 | | | | 2982 | 2039 | 4077 | 3371 | 5589 | 5520 | 6124 |
| 524 | 978 | 475 | 266 | 680 | | | | 1064 | 757 | 1445 | 1413 | 695 | 390 | 1463 |
| 580 | 868 | 985 | 522 | 820 | | | | 1809 | 1045 | 1802 | 1563 | 2098 | 1111 | 2130 |
| | | | | | 848 | 746 | 888 | | | | | | | |
| 572 | 1187 | | | | | | | 3666 | 2634 | 4608 | 5471 | | | |
| | | | | | 1045 | 924 | 934 | | | | | | | |
| 1380 | 1606 | | | | 508 | 432 | 1119 | 4394 | 4348 | 3150 | 5058 | | | |
| 700 | 758 | | | | | | | 1360 | 1013 | 1447 | 1097 | | | |
| | | 503 | 333 | 742 | | | | | | | | 1038 | 687 | 2063 |
| | | | | | 903 | 618 | 886 | | | | | | | |
| 585 | 861 | 751 | 807 | 1070 | | | | 1022 | 561 | 958 | 825 | 101 | 108 | 134 |
| | | 626 | 572 | 806 | 608 | 589 | 803 | | | | | 1942 | 1772 | 3100 |
| 830 | 1098 | | | | | | | 5015 | 3989 | 4807 | 5279 | | | |
| 244 | 467 | 739 | 545 | 815 | | | | 3082 | 1908 | 7834 | 3658 | 2082 | 1535 | 2816 |
| 384 | 731 | | | | 649 | 451 | 816 | 1695 | 1098 | 2859 | 2091 | | | |
| 635 | 1165 | | | | | | | 2417 | 2137 | 3367 | 3923 | | | |
| | | 696 | 508 | 818 | 697 | 453 | 744 | | | | | 1411 | 1029 | 2025 |
| 505 | 688 | 570 | 339 | 711 | | | | 1456 | 1075 | 2127 | 1463 | 2591 | 1540 | 4542 |
| 274 | 543 | 668 | 275 | 507 | | | | 2950 | 1092 | 3982 | 2164 | 4680 | 1927 | 7005 |
| | | 704 | 587 | 873 | 791 | 645 | 866 | | | | | 1756 | 1463 | 2493 |
| | | 990 | 703 | 1135 | 826 | 627 | 1083 | | | | | 1087 | 771 | 1097 |
| | | 619 | 353 | 642 | 591 | 416 | 737 | | | | | 3259 | 1861 | 5267 |
| 725 | 918 | | | | 824 | 451 | 703 | 3027 | 2228 | 3073 | 2820 | | | |
| 862 | 1090 | | | | | | | 1837 | 1183 | 1372 | 1495 | | | |
| | | 1131 | 955 | 1258 | 1084 | 896 | 1238 | | | | | 2571 | 2171 | 2274 |
| | | 708 | 459 | 879 | 663 | 415 | 862 | | | | | 1767 | 1145 | 2494 |
| 448 | 797 | 650 | 548 | 719 | | | | 1862 | 1394 | 3114 | 2483 | 846 | 714 | 1302 |
| 770 | 1708 | 822 | 697 | 1538 | | | | 1572 | 1359 | 1766 | 3017 | 1134 | 963 | 1380 |

| | | | | | | | | | | | | | | |
|---|---|---|---|---|---|---|---|---|---|---|---|---|---|---|
| 425 | 826 | 443 | 380 | 736 | | | | 654 | 715 | 1684 | 1391 | 1649 | 1414 | 3719 |
| 660 | 1027 | 866 | 695 | 1010 | | | | 2948 | 2442 | 3702 | 3802 | 2565 | 2059 | 2962 |
| | | 781 | 661 | 769 | 839 | 669 | 783 | | | | | 1366 | 1155 | 1749 |
| 1153 | 1422 | | | | | | | 1231 | 1175 | 1019 | 1449 | | | |
| 624 | 1041 | 850 | 634 | 987 | | | | 3359 | 2751 | 4409 | 4592 | 3899 | 2909 | 4586 |
| | | 1186 | 787 | 910 | 1243 | 842 | 953 | | | | | 3628 | 2407 | 3059 |
| 1019 | 861 | | | | | | | 1265 | 1412 | 1386 | 1194 | | | |
| | | 1044 | 942 | 1051 | 1149 | 1014 | 1061 | | | | | 1495 | 1348 | 1432 |
| | | 477 | 201 | 421 | 401 | 224 | 449 | | | | | 2340 | 985 | 4909 |
| 589 | 760 | 635 | 599 | 796 | | | | 671 | 654 | 1111 | 844 | 2390 | 2255 | 3764 |
| 314 | 693 | | | | 592 | 433 | 699 | 2296 | 1372 | 4368 | 3027 | | | |
| | | 567 | 266 | 529 | 555 | 263 | 531 | | | | | 2475 | 1162 | 4361 |
| | | 649 | 359 | 720 | 536 | 315 | 730 | | | | | 2283 | 1262 | 3517 |
| | | | | | 1232 | 1123 | 1082 | | | | | | | |
| | | | | | 768 | 581 | 927 | | | | | | | |
| | | 895 | 723 | 926 | | | | | | | | 2431 | 1963 | 2716 |
| 472 | 728 | | | | | | | 4106 | 2923 | 6194 | 4511 | | | |
| | | | | | 818 | 613 | 792 | | | | | | | |
| 892 | 1000 | | | | | | | 375 | 369 | 413 | 413 | | | |
| | | | | | 1127 | 1054 | 1075 | | | | | | | |
| 572 | 808 | 909 | 528 | 820 | | | | 1388 | 973 | 1701 | 1375 | 1432 | 832 | 1574 |
| | | 791 | 667 | 955 | | | | | | | | 1829 | 1544 | 2313 |
| | | 937 | 287 | 697 | 767 | 304 | 647 | | | | | 2636 | 807 | 2812 |
| | | | | | 855 | 656 | 929 | | | | | | | |
| 915 | 1398 | 1073 | 816 | 1119 | | | | 1751 | 1482 | 1620 | 2265 | 2214 | 1683 | 2064 |
| 577 | 839 | | | | 621 | 505 | 796 | 3084 | 2139 | 3709 | 3111 | | | |
| | | | | | 949 | 701 | 982 | | | | | | | |
| | | 387 | 274 | 298 | | | | | | | | 1268 | 900 | 3280 |
| 289 | 627 | | | | 910 | 700 | 918 | 3059 | 1627 | 5641 | 3536 | | | |
| | | 1368 | 1225 | 1068 | | | | | | | | 1007 | 902 | 736 |

| | | | | | | | | | | | | | | |
|---|---|---|---|---|---|---|---|---|---|---|---|---|---|---|
| | | 1078 | 962 | 1227 | | | | | | | | 3589 | 3204 | 3330 |
| 505 | 843 | 703 | 563 | 823 | | | | 2247 | 1534 | 3040 | 2562 | 2877 | 2302 | 4092 |
| | | 502 | 330 | 516 | 533 | 335 | 537 | | | | | 2439 | 1605 | 4862 |
| | | 508 | 449 | 659 | 548 | 484 | 703 | | | | | 6395 | 5645 | 12581 |
| 923 | 1044 | 513 | 362 | 720 | | | | 1996 | 1534 | 1661 | 1734 | 1718 | 1212 | 3348 |
| | | | | | 609 | 322 | 623 | | | | | | | |
| | | 498 | 238 | 573 | | | | | | | | 2349 | 1122 | 4720 |
| 515 | 911 | | | | 892 | 716 | 912 | 540 | 275 | 534 | 486 | | | |
| | | 993 | 545 | 781 | 970 | 645 | 870 | | | | | 1712 | 940 | 1724 |
| | | 946 | 795 | 1172 | 916 | 741 | 1157 | | | | | 1349 | 1133 | 1426 |
| | | 1000 | 488 | 826 | 885 | 557 | 889 | | | | | 2570 | 1253 | 2570 |
| | | 615 | 419 | 774 | | | | | | | | 5032 | 3427 | 8187 |
| | | | | | 445 | 252 | 797 | | | | | | | |
| 818 | 1170 | 877 | 677 | 868 | | | | 1765 | 1150 | 1406 | 1645 | 660 | 509 | 753 |
| 1011 | 1893 | 754 | 722 | 1637 | | | | 310 | 410 | 406 | 769 | 1499 | 1435 | 1989 |
| | | 1047 | 931 | 1509 | 1131 | 1041 | 1603 | | | | | 2176 | 1935 | 2079 |
| 564 | 978 | 774 | 559 | 957 | | | | 3912 | 2898 | 5138 | 5027 | 2478 | 1788 | 3200 |
| | | 1202 | 1385 | 1533 | | | | | | | | 1967 | 2266 | 1636 |
| 659 | 1000 | 1016 | 736 | 1036 | | | | 4040 | 2765 | 4194 | 4194 | 1582 | 1146 | 1556 |
| 663 | 991 | 840 | 793 | 1119 | | | | 1770 | 1453 | 2190 | 2170 | 1976 | 1865 | 2352 |
| 507 | 870 | 664 | 481 | 739 | | | | 924 | 663 | 1307 | 1136 | 907 | 657 | 1365 |
| 1061 | 1129 | | | | 749 | 730 | 1088 | 1539 | 1466 | 1382 | 1560 | | | |
| | | 902 | 770 | 1026 | | | | | | | | 1036 | 884 | 1148 |
| | | | | | 894 | 834 | 1045 | | | | | | | |
| 2079 | 1133 | | | | | | | 2601 | 2869 | 1380 | 1563 | | | |
| 402 | 713 | | | | 583 | 369 | 666 | 3495 | 2510 | 6238 | 4446 | | | |
| 505 | 1217 | | | | | | | 824 | 647 | 1281 | 1560 | | | |
| 618 | 955 | 741 | 640 | 1017 | | | | 510 | 389 | 629 | 600 | 1768 | 1527 | 2385 |
| 299 | 580 | | | | | | | 1982 | 1130 | 3779 | 2190 | | | |
| 715 | 1100 | 920 | 730 | 1005 | | | | 2197 | 1490 | 2083 | 2290 | 2610 | 2070 | 2837 |
| 614 | 979 | 817 | 627 | 965 | | | | 2038 | 1655 | 2693 | 2636 | 1807 | 1386 | 2210 |

| | | | | | | | | | | | | | | |
|---|---|---|---|---|---|---|---|---|---|---|---|---|---|---|
| | | | | | 1006 | 840 | 1011 | | | | | | | |
| 668 | 968 | 839 | 739 | 1123 | | | | 1750 | 1378 | 2062 | 1996 | 2439 | 2147 | 2907 |
| 610 | 946 | | | | | | | 1275 | 725 | 1188 | 1124 | | | |
| | | 401 | 243 | 435 | 884 | 640 | 768 | | | | | 2192 | 1328 | 5462 |
| | | 1114 | 880 | 1228 | 1352 | 746 | 1063 | | | | | 3418 | 2701 | 3069 |
| | | 1112 | 920 | 938 | | | | | | | | 948 | 785 | 852 |
| | | 652 | 478 | 724 | 958 | 768 | 1100 | | | | | 372 | 273 | 570 |
| | | 666 | 377 | 554 | | | | | | | | 892 | 505 | 1339 |
| 415 | 741 | | | | 729 | 385 | 718 | 1410 | 804 | 1935 | 1434 | | | |
| | | | | | 611 | 295 | 586 | | | | | | | |
| | | 637 | 877 | 1035 | | | | | | | | 3313 | 4564 | 5203 |
| | | 626 | 495 | 1104 | 785 | 570 | 1235 | | | | | 2516 | 1987 | 4017 |
| 726 | 930 | | | | | | | 2998 | 2404 | 3310 | 3079 | | | |
| 614 | 1006 | | | | 741 | 583 | 968 | 1975 | 1609 | 2618 | 2634 | | | |
| 859 | 1241 | 1189 | 925 | 1308 | | | | 2852 | 2137 | 2489 | 3089 | 2470 | 1921 | 2078 |
| | | | | | 788 | 565 | 921 | | | | | | | |
| | | | | | 832 | 711 | 993 | | | | | | | |
| | | | | | 1261 | 958 | 1026 | | | | | | | |
| | | 462 | 297 | 579 | | | | | | | | 5176 | 3331 | 11207 |
| 594 | 898 | | | | 734 | 567 | 925 | 818 | 677 | 1139 | 1022 | | | |
| | | | | | 1623 | 1207 | 1186 | | | | | | | |
| 892 | 1193 | | | | | | | 4445 | 3895 | 4368 | 5210 | | | |
| | | 811 | 551 | 923 | 860 | 632 | 994 | | | | | 6326 | 4298 | 7803 |
| | | 801 | 567 | 1027 | 907 | 640 | 1076 | | | | | 3836 | 2715 | 4788 |
| | | 989 | 902 | 1354 | | | | | | | | 660 | 603 | 668 |
| 683 | 851 | | | | | | | 4701 | 2853 | 4179 | 3556 | | | |
| 648 | 905 | 934 | 630 | 934 | | | | 2082 | 1476 | 2277 | 2062 | 2479 | 1671 | 2653 |
| | | 1065 | 912 | 1079 | | | | | | | | 6532 | 5593 | 6133 |

| | | | | | | | | | | | | | | |
|---|---|---|---|---|---|---|---|---|---|---|---|---|---|---|
| 361 | 594 | | | | | | | 3363 | 2631 | 7291 | 4330 | | | |
| | | | | | 590 | 509 | 746 | | | | | | | |
| 591 | 832 | | | | 869 | 551 | 992 | 3538 | 2421 | 4097 | 3410 | | | |
| | | 1125 | 715 | 1153 | 1019 | 678 | 1271 | | | | | 1583 | 1006 | 1407 |
| 456 | 793 | 612 | 535 | 830 | | | | 1598 | 1144 | 2506 | 1987 | 1517 | 1325 | 2479 |
| | | 894 | 782 | 1039 | | | | | | | | 1925 | 1684 | 2154 |
| 886 | 1166 | 1019 | 902 | 1195 | | | | 2338 | 2163 | 2440 | 2846 | 3224 | 2855 | 3164 |
| | | | | | 1195 | 934 | 1087 | | | | | | | |
| 995 | 1154 | | | | | | | 899 | 865 | 869 | 1003 | | | |
| 372 | 699 | 516 | 276 | 641 | | | | 5204 | 3855 | 10353 | 7242 | 3604 | 1929 | 6990 |
| | | 565 | 227 | 488 | 526 | 284 | 526 | | | | | 4072 | 1634 | 7211 |
| 667 | 1088 | 651 | 514 | 866 | | | | 347 | 246 | 369 | 402 | 634 | 501 | 974 |
| 515 | 826 | | | | 825 | 581 | 896 | 2330 | 1407 | 2732 | 2256 | | | |
| | | 584 | 763 | 740 | | | | | | | | 1520 | 1989 | 2606 |
| 319 | 620 | | | | 746 | 490 | 917 | 3165 | 2128 | 6665 | 4133 | | | |
| | | 877 | 656 | 1327 | | | | | | | | 2107 | 1575 | 2402 |
| 458 | 531 | 1010 | 692 | 837 | | | | 3373 | 1604 | 3503 | 1859 | 2470 | 1694 | 2446 |
| 637 | 947 | | | | 452 | 331 | 677 | 2314 | 1663 | 2610 | 2472 | | | |
| | | 512 | 426 | 829 | 497 | 469 | 1249 | | | | | 1441 | 1200 | 2813 |
| 361 | 10 | | | | 895 | 763 | 782 | 30 | 12 | 34 | | | | |
| 727 | 1203 | 675 | 497 | 953 | | | | 3895 | 3425 | 4712 | 5668 | 3162 | 2327 | 4685 |
| 778 | 1059 | | | | | | | 776 | 660 | 848 | 899 | | | |
| | | 789 | 647 | 1246 | 782 | 633 | 1148 | | | | | 1296 | 1063 | 1643 |
| | | | | | 1116 | 556 | 891 | | | | | | | |
| 705 | 777 | | | | | | | 2726 | 2175 | 3084 | 2396 | | | |
| | | 754 | 537 | 749 | | | | | | | | 4997 | 3554 | 6624 |
| | | 1458 | 586 | 907 | 948 | 622 | 1013 | | | | | 4945 | 1987 | 3392 |
| | | | | | 808 | 572 | 988 | | | | | | | |
| | | 1124 | 1055 | 1198 | | | | | | | | 1773 | 1665 | 1578 |

| | | | | | | | | | | | | | | |
|---|---|---|---|---|---|---|---|---|---|---|---|---|---|---|
| 846 | 1132 | | | | 1122 | 801 | 1083 | 3653 | 2499 | 2953 | 3343 | | | |
| | | | | | 886 | 630 | 882 | | | | | | | |
| 560 | 760 | | | | | | | 4684 | 3781 | 6750 | 5133 | | | |
| | | 982 | 772 | 1175 | | | | | | | | 1338 | 1053 | 1363 |
| | | 423 | 179 | 536 | | | | | | | | 3366 | 1424 | 7953 |
| | | | | | 951 | 636 | 1027 | | | | | | | |
| | | 724 | 430 | 638 | 862 | 483 | 740 | | | | | 1008 | 598 | 1392 |
| | | 858 | 637 | 1027 | 879 | 709 | 1177 | | | | | 9114 | 6765 | 10617 |
| | | 1804 | 1503 | 1161 | 1012 | 795 | 814 | | | | | 9025 | 7521 | 5002 |
| 544 | 885 | 1087 | 546 | 835 | | | | 1107 | 756 | 1390 | 1230 | 1637 | 822 | 1506 |
| 428 | 646 | | | | | | | 1152 | 687 | 1607 | 1038 | | | |
| 486 | 860 | | | | | | | 596 | 490 | 1008 | 867 | | | |
| 349 | 650 | | | | 472 | 349 | 611 | 1254 | 828 | 2371 | 1542 | | | |
| 536 | 910 | | | | | | | 423 | 300 | 560 | 510 | | | |
| | | 672 | 369 | 729 | | | | | | | | 1336 | 732 | 1987 |
| | | | | | 1040 | 606 | 829 | | | | | | | |
| 798 | 2224 | | | | 928 | 700 | 2070 | 2048 | 1628 | 2041 | 4540 | | | |
| | | 914 | 952 | 1396 | | | | | | | | 6114 | 6371 | 6689 |
| | | 1647 | 701 | 911 | | | | | | | | 708 | 301 | 430 |
| 751 | 990 | | | | 1096 | 949 | 1501 | 4651 | 3340 | 4446 | 4399 | | | |
| | | 1157 | 616 | 867 | | | | | | | | 3135 | 1669 | 2710 |
| 809 | 1059 | 1016 | 682 | 1154 | | | | 2186 | 1698 | 2098 | 2221 | 1906 | 1280 | 1876 |
| | | 769 | 593 | 917 | 642 | 594 | 898 | | | | | 2220 | 1710 | 2886 |
| | | 332 | 249 | 692 | 277 | 231 | 587 | | | | | 715 | 536 | 2153 |
| | | | | | 984 | 739 | 1004 | | | | | | | |
| | | 1297 | 842 | 930 | 1153 | 685 | 839 | | | | | 8893 | 5773 | 6855 |
| 612 | 839 | | | | | | | 1109 | 981 | 1602 | 1344 | | | |
| | | 823 | 559 | 892 | | | | | | | | 635 | 431 | 771 |

| | | | | | | | | | | | | | | |
|---|---|---|---|---|---|---|---|---|---|---|---|---|---|---|
| 470 | 857 | | | | 1006 | 881 | 1065 | 4397 | 3128 | 6651 | 5701 | | | |
| 536 | 833 | | | | | | | 1338 | 918 | 1712 | 1427 | | | |
| | | 1046 | 733 | 802 | | | | | | | | 654 | 458 | 625 |
| | | 754 | 333 | 417 | 708 | 272 | 352 | | | | | 2398 | 1060 | 3181 |
| | | 946 | 591 | 754 | | | | | | | | 7169 | 4482 | 7579 |
| 676 | 921 | | | | 567 | 559 | 996 | 1418 | 1233 | 1823 | 1679 | | | |
| | | 698 | 498 | 887 | 897 | 614 | 1028 | | | | | 4543 | 3242 | 6506 |
| | | | | | 1005 | 659 | 826 | | | | | | | |
| 563 | 933 | 774 | 568 | 964 | | | | 1818 | 1252 | 2223 | 2073 | 3474 | 2548 | 4489 |
| | | | | | 1063 | 784 | 908 | | | | | | | |
| 681 | 1155 | 818 | 682 | 1077 | | | | 1091 | 859 | 1262 | 1458 | 903 | 754 | 1104 |
| 464 | 836 | 720 | 514 | 825 | | | | 2184 | 1459 | 3146 | 2629 | 1547 | 1104 | 2149 |
| 834 | 971 | | | | | | | 4196 | 3169 | 3799 | 3690 | | | |
| 741 | 1281 | 751 | 646 | 1004 | | | | 3552 | 3379 | 4561 | 5845 | 1402 | 1205 | 1866 |
| 512 | 799 | | | | | | | 2839 | 1772 | 3458 | 2764 | | | |
| 1139 | 955 | | | | | | | 464 | 425 | 373 | 356 | | | |
| | | 807 | 589 | 840 | | | | | | | | 2000 | 1461 | 2479 |
| | | | | | 636 | 486 | 755 | | | | | | | |
| | | 1298 | 1270 | 1250 | | | | | | | | 2322 | 2272 | 1789 |
| | | | | | 903 | 994 | 1142 | | | | | | | |
| 1536 | 1589 | 1386 | 1450 | 1598 | | | | 3388 | 3882 | 2528 | 4015 | 3613 | 3779 | 2607 |
| | | 1435 | 1193 | 1366 | 1507 | 991 | 1147 | | | | | 3913 | 3251 | 2726 |
| 644 | 1103 | | | | 795 | 717 | 1088 | 1555 | 1410 | 2188 | 2414 | | | |
| | | | | | 1154 | 945 | 1160 | | | | | | | |
| | | | | | 610 | 423 | 736 | | | | | | | |
| 275 | 583 | | | | | | | 916 | 673 | 2445 | 1426 | | | |
| | | 1017 | 1087 | 1068 | | | | | | | | 1118 | 1195 | 1099 |
| 1409 | 1993 | 858 | 431 | 833 | | | | 3064 | 4101 | 2912 | 5802 | 5557 | 2789 | 6478 |
| | | 809 | 686 | 804 | 742 | 803 | 866 | | | | | 2400 | 2037 | 2968 |

| | | | | | | | | | | | | | | |
|---|---|---|---|---|---|---|---|---|---|---|---|---|---|---|
| | | | | | 309 | 282 | 382 | | | | | | | |
| 716 | 947 | | | | 698 | 681 | 1035 | 1263 | 1296 | 1809 | 1713 | | | |
| 438 | 703 | | | | | | | 1093 | 747 | 1706 | 1200 | | | |
| | | 566 | 425 | 693 | 534 | 393 | 677 | | | | | 1326 | 996 | 2345 |
| | | | | | 777 | 630 | 868 | | | | | | | |
| 630 | 813 | 837 | 637 | 819 | | | | 2590 | 1975 | 3135 | 2549 | 2201 | 1675 | 2629 |
| 1021 | 1090 | | | | | | | 3070 | 2916 | 2855 | 3112 | | | |
| | | 700 | 507 | 961 | | | | | | | | 2118 | 1534 | 3025 |
| | | 1125 | 1050 | 1234 | 1205 | 1115 | 1212 | | | | | 4414 | 4119 | 3924 |
| 671 | 1333 | | | | 740 | 582 | 1071 | 4292 | 3845 | 5731 | 7637 | | | |
| | | 897 | 795 | 1064 | | | | | | | | 1928 | 1708 | 2149 |
| | | | | | 897 | 677 | 944 | | | | | | | |
| 719 | 1138 | 529 | 522 | 961 | | | | 1365 | 1116 | 1551 | 1766 | 963 | 950 | 1819 |
| | | 663 | 409 | 639 | | | | | | | | 1270 | 783 | 1916 |
| 491 | 789 | | | | | | | 3077 | 2037 | 4146 | 3271 | | | |
| | | | | | 1131 | 987 | 1353 | | | | | | | |
| 250 | 370 | 895 | 291 | 446 | | | | 8437 | 2601 | 10404 | 3849 | 8253 | 2680 | 9218 |
| 809 | 852 | | | | 972 | 764 | 740 | 1331 | 956 | 1181 | 1006 | | | |
| | | 1161 | 1635 | 2190 | | | | | | | | 2474 | 3486 | 2132 |
| | | 800 | 286 | 505 | 816 | 325 | 518 | | | | | 4156 | 1486 | 5199 |
| | | | | | 700 | 350 | 720 | | | | | | | |
| | | | | | 888 | 774 | 1000 | | | | | | | |
| 218 | 642 | 338 | 174 | 646 | | | | 1928 | 1191 | 5475 | 3516 | 2380 | 1224 | 7033 |
| | | | | | 982 | 802 | 1179 | | | | | | | |
| | | | | | 1005 | 699 | 986 | | | | | | | |
| | | 462 | 385 | 716 | | | | | | | | 856 | 714 | 1853 |
| 2867 | 2371 | | | | | | | 10141 | 17149 | 5981 | 14182 | | | |

| | | | | | | | | | | | | | | |
|---|---|---|---|---|---|---|---|---|---|---|---|---|---|---|
| | | 939 | 863 | 910 | | | | | | | | 1577 | 1450 | 1679 |
| 388 | 821 | | | | 553 | 295 | 745 | 990 | 641 | 1652 | 1356 | | | |
| | | 958 | 892 | 1263 | | | | | | | | 2661 | 2479 | 2777 |
| | | | | | 1015 | 996 | 1107 | | | | | | | |
| 855 | 913 | | | | | | | 2725 | 2474 | 2894 | 2641 | | | |
| | | 703 | 458 | 722 | | | | | | | | 1855 | 1207 | 2638 |
| | | | | | 982 | 462 | 737 | | | | | | | |
| 729 | 1318 | | | | | | | 2944 | 2499 | 3428 | 4518 | | | |
| 586 | 939 | | | | | | | 1817 | 1255 | 2142 | 2011 | | | |
| | | | | | 996 | 831 | 1090 | | | | | | | |
| 1014 | 1498 | | | | | | | 1718 | 1600 | 1578 | 2365 | | | |
| | | | | | 822 | 760 | 1091 | | | | | | | |
| 484 | 756 | | | | | | | 723 | 454 | 937 | 708 | | | |
| 381 | 731 | | | | | | | 5685 | 3524 | 9256 | 6766 | | | |
| | | 754 | 782 | 947 | | | | | | | | 1630 | 1690 | 2162 |
| | | 759 | 723 | 809 | | | | | | | | 3302 | 3146 | 4352 |
| 593 | 898 | 653 | 545 | 860 | | | | 1385 | 1264 | 2130 | 1914 | 1644 | 1372 | 2519 |
| | | 1415 | 889 | 982 | | | | | | | | 2671 | 1679 | 1888 |
| | | | | | 601 | 431 | 780 | | | | | | | |
| 268 | 372 | | | | | | | 2084 | 668 | 2488 | 926 | | | |
| | | 896 | 915 | 1466 | | | | | | | | 1258 | 1286 | 1405 |
| | | 738 | 498 | 865 | | | | | | | | 1705 | 1150 | 2310 |
| | | | | | 876 | 790 | 1005 | | | | | | | |
| 779 | 845 | | | | | | | 3264 | 2451 | 3146 | 2658 | | | |
| | | | | | 1012 | 863 | 1129 | | | | | | | |

| | | | | | | | | | | | | | | |
|---|---|---|---|---|---|---|---|---|---|---|---|---|---|---|
| | | | | | 1116 | 760 | 842 | | | | | | | |
| | | 1181 | 1033 | 861 | | | | | | | | 922 | 807 | 781 |
| | | | | | 1352 | 1145 | 1352 | | | | | | | |
| | | 960 | 846 | 1069 | | | | | | | | 1356 | 1195 | 1413 |
| | | 903 | 745 | 1039 | | | | | | | | 2263 | 1867 | 2506 |
| 780 | 937 | | | | | | | 3035 | 2212 | 2837 | 2659 | | | |
| | | | | | 1224 | 1172 | 1331 | | | | | | | |
| | | 999 | 1021 | 1286 | | | | | | | | 2488 | 2543 | 2491 |
| | | 880 | 701 | 952 | | | | | | | | 812 | 646 | 923 |
| | | 1101 | 982 | 1055 | | | | | | | | 1707 | 1522 | 1550 |
| 486 | 904 | | | | | | | 1628 | 1149 | 2363 | 2136 | | | |
| | | 1143 | 924 | 1006 | | | | | | | | 1287 | 1041 | 1126 |
| | | | | | 731 | 444 | 817 | | | | | | | |
| 322 | 630 | | | | | | | 2694 | 1722 | 5342 | 3367 | | | |
| 880 | 1144 | | | | | | | 4143 | 3280 | 3727 | 4262 | | | |
| 1361 | 1159 | | | | | | | 1938 | 1792 | 1317 | 1525 | | | |
| 623 | 917 | | | | | | | 4213 | 3049 | 4896 | 4489 | | | |
| | | | | | 974 | 752 | 902 | | | | | | | |
| 1478 | 1873 | | | | | | | 459 | 450 | 304 | 570 | | | |
| | | 777 | 498 | 696 | | | | | | | | 1626 | 1041 | 2092 |
| | | | | | 603 | 481 | 824 | | | | | | | |
| | | | | | 701 | 505 | 801 | | | | | | | |
| | | 1055 | 594 | 881 | | | | | | | | 4163 | 2342 | 3945 |
| 463 | 685 | | | | | | | 2003 | 1497 | 3236 | 2216 | | | |
| 794 | 1139 | | | | | | | 762 | 663 | 834 | 950 | | | |
| | | | | | 1012 | 908 | 1225 | | | | | | | |
| | | 1006 | 785 | 1075 | | | | | | | | 1117 | 872 | 1111 |
| | | | | | 1064 | 891 | 1133 | | | | | | | |
| | | 498 | 277 | 634 | | | | | | | | 2795 | 1555 | 5614 |
| | | | | | 1450 | 1350 | 1413 | | | | | | | |

| | | | | | | | | | | | | | | |
|---|---|---|---|---|---|---|---|---|---|---|---|---|---|---|
| | | | | | 960 | 647 | 905 | | | | | | | |
| | | | | | 601 | 333 | 556 | | | | | | | |
| | | | | | 769 | 586 | 1106 | | | | | | | |
| | | 928 | 728 | 1187 | | | | | | | | 1413 | 1109 | 1523 |
| | | | | | 662 | 745 | 1170 | | | | | | | |
| 367 | 795 | | | | | | | 1673 | 1005 | 2740 | 2180 | | | |
| | | 1446 | 1601 | 1361 | | | | | | | | 3519 | 3897 | 2434 |
| 523 | 797 | | | | | | | 3441 | 1939 | 3711 | 2958 | | | |
| | | | | | 1141 | 1235 | 1817 | | | | | | | |
| | | 914 | 497 | 789 | | | | | | | | 5586 | 3038 | 6112 |
| | | 2359 | 3145 | 1279 | | | | | | | | 4089 | 5452 | 1734 |
| | | 1200 | 778 | 1056 | | | | | | | | 2435 | 1580 | 2029 |
| | | 871 | 615 | 1326 | | | | | | | | 3360 | 2373 | 3857 |
| 713 | 891 | | | | | | | 2369 | 1817 | 2547 | 2269 | | | |
| | | | | | 706 | 470 | 732 | | | | | | | |
| | | | | | 769 | 314 | 738 | | | | | | | |
| | | | | | 852 | 629 | 948 | | | | | | | |
| 700 | 1034 | | | | | | | 1814 | 1381 | 1971 | 2038 | | | |
| 527 | 584 | | | | | | | 2492 | 2086 | 3955 | 2308 | | | |
| | | 757 | 326 | 583 | | | | | | | | 1067 | 460 | 1410 |
| | | 764 | 664 | 1071 | | | | | | | | 938 | 815 | 1228 |
| | | 956 | 633 | 824 | | | | | | | | 1464 | 969 | 1532 |
| | | 1064 | 820 | 1017 | | | | | | | | 677 | 522 | 637 |
| 456 | 577 | | | | | | | 1767 | 1079 | 2363 | 1364 | | | |
| 1040 | 1349 | | | | | | | 315 | 316 | 304 | 410 | | | |
| | | | | | 620 | 518 | 691 | | | | | | | |
| 178 | 615 | | | | | | | 419 | 183 | 1029 | 633 | | | |
| | | | | | 856 | 424 | 748 | | | | | | | |

| | | | | | | | | | | | | | | |
|---|---|---|---|---|---|---|---|---|---|---|---|---|---|---|
| | | | | | 1164 | 819 | 1073 | | | | | | | |
| 703 | 1015 | | | | | | | 3197 | 2622 | 3732 | 3789 | | | |
| | | | | | 1243 | 951 | 945 | | | | | | | |
| | | | | | 775 | 536 | 891 | | | | | | | |
| 571 | 893 | | | | | | | 1729 | 1372 | 2402 | 2144 | | | |
| | | 776 | 491 | 892 | | | | | | | | 1657 | 1048 | 2135 |
| | | | | | 925 | 674 | 935 | | | | | | | |
| | | | | | 566 | 518 | 703 | | | | | | | |
| | | 778 | 768 | 943 | | | | | | | | 2624 | 2591 | 3374 |
| | | 769 | 595 | 959 | | | | | | | | 3715 | 2876 | 4833 |
| | | 890 | 621 | 955 | | | | | | | | 3833 | 2674 | 4308 |
| 645 | 1071 | | | | | | | 1741 | 1449 | 2248 | 2407 | | | |
| | | 609 | 438 | 748 | | | | | | | | 1596 | 1146 | 2618 |
| | | 857 | 611 | 974 | | | | | | | | 2121 | 1512 | 2474 |
| | | | | | 955 | 648 | 991 | | | | | | | |
| | | | | | 124 | 195 | 445 | | | | | | | |
| | | 645 | 537 | 973 | | | | | | | | 1573 | 1310 | 2441 |
| 563 | 606 | | | | | | | 1210 | 650 | 1155 | 699 | | | |
| 725 | 1019 | | | | | | | 7233 | 4576 | 6315 | 6433 | | | |
| | | 1088 | 1055 | 1173 | | | | | | | | 1980 | 1918 | 1819 |
| | | 873 | 743 | 1178 | | | | | | | | 2778 | 2365 | 3182 |
| | | 992 | 2321 | 944 | | | | | | | | 797 | 1866 | 804 |
| 720 | 1242 | | | | | | | 461 | 376 | 522 | 649 | | | |
| | | | | | 1208 | 1159 | 1359 | | | | | | | |
| 558 | 1018 | | | | | | | 2079 | 1692 | 3031 | 3087 | | | |

| | | | | | | | | | | | | | | |
|---|---|---|---|---|---|---|---|---|---|---|---|---|---|---|
| | | 814 | 645 | 1130 | | | | | | | | 1661 | 1316 | 2039 |
| | | 808 | 737 | 1229 | | | | | | | | 437 | 399 | 541 |
| 682 | 1000 | | | | | | | 2659 | 2285 | 3349 | 3349 | | | |
| 444 | 578 | | | | | | | 963 | 814 | 1834 | 1061 | | | |
| | | | | | 932 | 462 | 618 | | | | | | | |
| | | 942 | 603 | 1036 | | | | | | | | 2612 | 1671 | 2772 |
| | | 178 | 132 | 316 | | | | | | | | 720 | 536 | 4055 |
| | | 1077 | 943 | 1153 | | | | | | | | 4740 | 4150 | 4403 |
| | | 700 | 474 | 838 | | | | | | | | 943 | 639 | 1348 |
| | | | | | 405 | 302 | 709 | | | | | | | |
| 2281 | 1652 | | | | | | | 2516 | 3437 | 1507 | 2489 | | | |
| 581 | 876 | | | | | | | 820 | 622 | 1070 | 938 | | | |
| | | 849 | 712 | 921 | | | | | | | | 4956 | 4159 | 5840 |
| 599 | 967 | | | | | | | 2292 | 1817 | 3035 | 2934 | | | |
| 897 | 895 | | | | | | | 2358 | 2049 | 2285 | 2045 | | | |
| 742 | 833 | | | | | | | 1097 | 932 | 1257 | 1047 | | | |
| | | | | | 880 | 784 | 1035 | | | | | | | |
| | | | | | 752 | 532 | 772 | | | | | | | |
| | | | | | 942 | 756 | 1031 | | | | | | | |
| | | | | | 830 | 579 | 968 | | | | | | | |
| | | | | | 691 | 485 | 911 | | | | | | | |

| | | | | | | | | | | | | | | |
|---|---|---|---|---|---|---|---|---|---|---|---|---|---|---|
| | | 526 | 450 | 1002 | | | | | | | | 218 | 187 | 415 |
| 483 | 793 | | | | | | | 766 | 483 | 1001 | 794 | | | |
| 851 | 492 | | | | | | | 4544 | 6115 | 7181 | 3531 | | | |
| | | | | | 1557 | 1600 | 1962 | | | | | | | |
| 1358 | 1188 | | | | | | | 519 | 546 | 402 | 477 | | | |
| | | 657 | 582 | 904 | | | | | | | | 821 | 727 | 1249 |
| | | 883 | 782 | 1097 | | | | | | | | 547 | 484 | 619 |
| 355 | 818 | | | | | | | 4531 | 3065 | 8637 | 7062 | | | |
| | | 499 | 310 | 579 | | | | | | | | 4927 | 3060 | 9877 |
| 803 | 928 | | | | | | | 1077 | 931 | 1160 | 1077 | | | |
| | | | | | 735 | 470 | 740 | | | | | | | |
| 678 | 1034 | | | | | | | 2045 | 1318 | 1944 | 2011 | | | |
| 470 | 1003 | | | | | | | 141 | 92 | 197 | 197 | | | |
| | | | | | 782 | 630 | 948 | | | | | | | |
| | | 799 | 562 | 1000 | | | | | | | | 604 | 425 | 756 |
| | | 467 | 248 | 687 | | | | | | | | 2734 | 1450 | 5855 |
| | | 858 | 678 | 1084 | | | | | | | | 6486 | 5125 | 7563 |
| 995 | 1267 | | | | | | | 1654 | 1623 | 1632 | 2068 | | | |
| 1266 | 1333 | | | | | | | 840 | 798 | 630 | 840 | | | |
| | | | | | 681 | 849 | 1119 | | | | | | | |
| | | 1189 | 1062 | 1342 | | | | | | | | 3897 | 3480 | 3278 |
| 481 | 890 | | | | | | | 1014 | 639 | 1329 | 1183 | | | |
| | | 1071 | 681 | 867 | | | | | | | | 2106 | 1339 | 1967 |
| | | 786 | 528 | 1208 | | | | | | | | 1265 | 850 | 1610 |

| | | | | | | | | | | | | | | |
|---|---|---|---|---|---|---|---|---|---|---|---|---|---|---|
| | | 1170 | 929 | 1404 | | | | | | | | 1646 | 1308 | 1407 |
| | | 949 | 846 | 1059 | | | | | | | | 1349 | 1202 | 1422 |
| | | 1118 | 815 | 1101 | | | | | | | | 1765 | 1287 | 1579 |
| 456 | 742 | | | | | | | 1156 | 864 | 1895 | 1406 | | | |
| 719 | 1207 | | | | | | | 2894 | 2410 | 3351 | 4044 | | | |
| 629 | 843 | | | | | | | 2007 | 1536 | 2442 | 2060 | | | |
| | | 755 | 538 | 925 | | | | | | | | 2822 | 2013 | 3739 |
| | | | | | 799 | 415 | 726 | | | | | | | |
| | | 554 | 440 | 687 | | | | | | | | 2200 | 1745 | 3969 |
| | | 1163 | 919 | 1205 | | | | | | | | 3581 | 2830 | 3080 |
| 444 | 781 | | | | | | | 980 | 530 | 1195 | 933 | | | |
| 924 | 1368 | | | | | | | 2241 | 2115 | 2288 | 3129 | | | |
| | | | | | 858 | 893 | 1156 | | | | | | | |
| | | | | | 507 | 340 | 626 | | | | | | | |
| 829 | 1268 | | | | | | | 656 | 665 | 802 | 1017 | | | |
| | | | | | 798 | 491 | 744 | | | | | | | |
| 805 | 810 | | | | | | | 1849 | 1415 | 1758 | 1424 | | | |
| | | | | | 1122 | 1072 | 1068 | | | | | | | |
| 516 | 726 | | | | | | | 2376 | 1860 | 3603 | 2615 | | | |
| | | | | | 670 | 447 | 444 | | | | | | | |
| | | 994 | 897 | 1142 | | | | | | | | 1630 | 1471 | 1640 |
| | | 844 | 818 | 979 | | | | | | | | 966 | 936 | 1145 |
| | | | | | 1359 | 976 | 963 | | | | | | | |
| 500 | 827 | | | | | | | 1460 | 1047 | 2094 | 1732 | | | |
| | | 909 | 655 | 964 | | | | | | | | 2241 | 1614 | 2465 |

| | | | | | | | | | | | | | | |
|---|---|---|---|---|---|---|---|---|---|---|---|---|---|---|
| 591 | 703 | | | | | | | 3700 | 2356 | 3989 | 2804 | | | |
| 647 | 849 | | | | | | | 4444 | 3200 | 4949 | 4200 | | | |
| | | | | | 599 | 205 | 454 | | | | | | | |
| 1437 | 1512 | | | | | | | 3284 | 3268 | 2275 | 3438 | | | |
| | | | | | 943 | 749 | 1147 | | | | | | | |
| | | | | | 678 | 462 | 695 | | | | | | | |
| | | | | | 692 | 554 | 953 | | | | | | | |
| 317 | 472 | | | | | | | 1350 | 1265 | 3994 | 1885 | | | |
| | | | | | 776 | 563 | 924 | | | | | | | |
| | | | | | 915 | 944 | 1120 | | | | | | | |
| | | | | | 852 | 504 | 729 | | | | | | | |
| | | 785 | 546 | 996 | | | | | | | | 1100 | 766 | 1402 |
| 769 | 1041 | | | | | | | 1369 | 1150 | 1494 | 1556 | | | |
| 300 | 408 | | | | | | | 735 | 307 | 1023 | 417 | | | |
| 483 | 785 | | | | | | | 1654 | 966 | 2002 | 1571 | | | |
| 863 | 1252 | | | | | | | 1930 | 1438 | 1667 | 2088 | | | |
| | | 1105 | 849 | 1085 | | | | | | | | 2078 | 1597 | 1881 |
| | | 995 | 710 | 1010 | | | | | | | | 1598 | 1140 | 1606 |
| | | 570 | 375 | 597 | | | | | | | | 4764 | 3130 | 8354 |
| 586 | 932 | | | | | | | 2195 | 2330 | 3979 | 3710 | | | |
| 403 | 778 | | | | | | | 1089 | 755 | 1872 | 1456 | | | |
| | | 463 | 278 | 631 | | | | | | | | 1303 | 783 | 2816 |
| 362 | 650 | | | | | | | 661 | 413 | 1141 | 742 | | | |
| | | 636 | 429 | 963 | | | | | | | | 6282 | 4244 | 9882 |

| | | | | | | | | | | | | | | |
|---|---|---|---|---|---|---|---|---|---|---|---|---|---|---|
| | | 541 | 318 | 657 | | | | | | | | 923 | 542 | 1705 |
| | | | | | 360 | 381 | 1020 | | | | | | | |
| | | | | | 857 | 754 | 1129 | | | | | | | |
| | | 634 | 382 | 759 | | | | | | | | 231 | 139 | 365 |
| | | 1061 | 824 | 1264 | | | | | | | | 3710 | 2881 | 3496 |
| | | | | | 664 | 669 | 1450 | | | | | | | |
| | | 673 | 437 | 744 | | | | | | | | 1803 | 1172 | 2678 |
| | | | | | 735 | 634 | 1000 | | | | | | | |
| | | 966 | 756 | 1062 | | | | | | | | 417 | 327 | 432 |
| | | | | | 707 | 475 | 547 | | | | | | | |
| | | 1069 | 834 | 1195 | | | | | | | | 2347 | 1833 | 2196 |
| 671 | 1041 | | | | | | | 437 | 415 | 619 | 644 | | | |
| 436 | 854 | | | | | | | 3378 | 2308 | 5292 | 4517 | | | |
| 507 | 806 | | | | | | | 2626 | 1658 | 3273 | 2637 | | | |
| | | | | | 931 | 774 | 825 | | | | | | | |
| 227 | 605 | | | | | | | 1359 | 1017 | 4482 | 2711 | | | |
| 372 | 763 | | | | | | | 1843 | 1233 | 3312 | 2529 | | | |
| 416 | 697 | | | | | | | 1115 | 692 | 1665 | 1160 | | | |
| 2054 | 580 | | | | | | | 1071 | 3902 | 1899 | 1101 | | | |
| | | 663 | 527 | 901 | | | | | | | | 1546 | 1230 | 2333 |
| 319 | 575 | | | | | | | 856 | 517 | 1619 | 931 | | | |
| | | 1012 | 659 | 908 | | | | | | | | 1801 | 1173 | 1780 |
| 240 | 544 | | | | | | | 1224 | 779 | 3249 | 1768 | | | |
| 1378 | 1350 | | | | | | | 1814 | 1796 | 1303 | 1759 | | | |
| 735 | 962 | | | | | | | 2038 | 1644 | 2237 | 2151 | | | |

| | | | | | | | | | | | | | | |
|---|---|---|---|---|---|---|---|---|---|---|---|---|---|---|
| 1228 | 1231 | | | | | | | 1729 | 1586 | 1292 | 1590 | | | |
| | | 887 | 792 | 973 | | | | | | | | 1717 | 1534 | 1936 |
| 594 | 777 | | | | | | | 1702 | 1427 | 2401 | 1865 | | | |
| | | | | | 911 | 449 | 706 | | | | | | | |
| 647 | 1103 | | | | | | | 1724 | 976 | 1509 | 1664 | | | |
| | | | | | 1124 | 907 | 1444 | | | | | | | |
| | | 737 | 429 | 690 | | | | | | | | 576 | 335 | 781 |
| 902 | 1209 | | | | | | | 1719 | 1413 | 1566 | 1893 | | | |
| | | | | | 984 | 867 | 1194 | | | | | | | |
| | | | | | 1013 | 777 | 973 | | | | | | | |
| | | | | | 966 | 861 | 1105 | | | | | | | |
| | | | | | 468 | 368 | 578 | | | | | | | |
| | | | | | 1034 | 1069 | 1163 | | | | | | | |
| 943 | 1573 | | | | | | | 2907 | 3199 | 3393 | 5338 | | | |
| | | 1218 | 1260 | 1455 | | | | | | | | 2707 | 2800 | 2223 |
| | | | | | 613 | 531 | 1003 | | | | | | | |
| 345 | 663 | | | | | | | 247 | 101 | 293 | 195 | | | |
| | | | | | 672 | 580 | 808 | | | | | | | |
| | | 742 | 636 | 1223 | | | | | | | | 944 | 809 | 1271 |
| 1002 | 1452 | | | | | | | 4227 | 3864 | 3857 | 5601 | | | |
| 303 | 640 | | | | | | | 1419 | 820 | 2705 | 1730 | | | |

| | | | | | | | | | | | | | | |
|---|---|---|---|---|---|---|---|---|---|---|---|---|---|---|
| | | | | | 1000 | 812 | 1199 | | | | | | | |
| | | | | | 833 | 564 | 857 | | | | | | | |
| | | | | | 793 | 647 | 1308 | | | | | | | |
| 604 | 949 | | | | | | | 927 | 657 | 1087 | 1032 | | | |
| 893 | 929 | | | | | | | 3749 | 3441 | 3854 | 3579 | | | |
| 664 | 922 | | | | | | | 5020 | 3655 | 5508 | 5077 | | | |
| 735 | 950 | | | | | | | 1732 | 1581 | 2152 | 2044 | | | |
| | | | | | 794 | 481 | 915 | | | | | | | |
| | | | | | 834 | 825 | 1011 | | | | | | | |
| 347 | 539 | | | | | | | 800 | 476 | 1372 | 740 | | | |
| | | | | | 901 | 688 | 1010 | | | | | | | |
| 844 | 1008 | | | | | | | 3311 | 2492 | 2953 | 2977 | | | |
| 422 | 559 | | | | | | | 173 | 217 | 513 | 287 | | | |
| 819 | 1201 | | | | | | | 314 | 253 | 308 | 370 | | | |
| | | | | | 921 | 656 | 953 | | | | | | | |
| 428 | 796 | | | | | | | 2527 | 1684 | 3934 | 3132 | | | |
| | | | | | 872 | 684 | 925 | | | | | | | |
| | | 869 | 656 | 1064 | | | | | | | | 725 | 547 | 834 |
| 618 | 1317 | | | | | | | 2883 | 2739 | 4435 | 5842 | | | |
| | | 895 | 694 | 1247 | | | | | | | | 2541 | 1970 | 2840 |
| | | | | | 821 | 615 | 1045 | | | | | | | |
| | | | | | 861 | 417 | 735 | | | | | | | |
| | | 784 | 416 | 719 | | | | | | | | 819 | 435 | 1045 |
| 485 | 828 | | | | | | | 2703 | 2031 | 4191 | 3469 | | | |
| | | | | | 699 | 526 | 784 | | | | | | | |
| | | | | | 1098 | 967 | 1452 | | | | | | | |
| | | | | | 441 | 255 | 651 | | | | | | | |
| 514 | 945 | | | | | | | 2342 | 1622 | 3154 | 2981 | | | |
| | | | | | 842 | 559 | 957 | | | | | | | |
| 516 | 815 | | | | | | | 9808 | 5127 | 9940 | 8099 | | | |

| | | | | | | | | | | | | | | |
|---|---|---|---|---|---|---|---|---|---|---|---|---|---|---|
| | | | | | 1469 | 1233 | 1601 | | | | | | | |
| | | 610 | 531 | 897 | | | | | | | | 1716 | 1494 | 2811 |
| | | 730 | 627 | 936 | | | | | | | | 1566 | 1345 | 2147 |
| 659 | 798 | | | | | | | 2299 | 1802 | 2734 | 2181 | | | |
| 410 | 719 | | | | | | | 2129 | 960 | 2342 | 1684 | | | |
| 724 | 1073 | | | | | | | 3268 | 2507 | 3465 | 3718 | | | |
| | | | | | 639 | 470 | 733 | | | | | | | |
| | | | | | 765 | 488 | 822 | | | | | | | |
| | | | | | 803 | 230 | 358 | | | | | | | |
| | | 1086 | 1257 | 1084 | | | | | | | | 502 | 580 | 462 |
| 380 | 588 | | | | | | | 2761 | 1396 | 3671 | 2158 | | | |
| | | 537 | 297 | 671 | | | | | | | | 1436 | 795 | 2676 |
| | | 1039 | 507 | 822 | | | | | | | | 641 | 313 | 617 |
| | | 865 | 603 | 1042 | | | | | | | | 2904 | 2025 | 3358 |
| | | | | | 1149 | 837 | 1040 | | | | | | | |
| 594 | 887 | | | | | | | 3338 | 2166 | 3645 | 3234 | | | |
| | | 415 | 348 | 746 | | | | | | | | 416 | 348 | 1001 |
| | | 584 | 513 | 769 | | | | | | | | 541 | 476 | 927 |
| 539 | 928 | | | | | | | 898 | 652 | 1209 | 1122 | | | |
| 618 | 900 | | | | | | | 4175 | 2721 | 4405 | 3965 | | | |
| 350 | 715 | | | | | | | 3435 | 2040 | 5825 | 4167 | | | |
| 203 | 431 | | | | | | | 3806 | 2060 | 10121 | 4361 | | | |
| | | | | | 753 | 589 | 966 | | | | | | | |
| 353 | 556 | | | | | | | 3455 | 3040 | 8603 | 4787 | | | |
| 461 | 852 | | | | | | | 3890 | 2573 | 5578 | 4752 | | | |
| 551 | 729 | | | | | | | 1483 | 1012 | 1838 | 1340 | | | |
| | | | | | 751 | 290 | 477 | | | | | | | |
| | | | | | 1015 | 751 | 1135 | | | | | | | |
| | | | | | 883 | 596 | 908 | | | | | | | |
| | | | | | 375 | 352 | 834 | | | | | | | |
| | | 695 | 590 | 976 | | | | | | | | 1520 | 1289 | 2186 |

| | | | | | | | | | | | | | | |
|---|---|---|---|---|---|---|---|---|---|---|---|---|---|---|
| 367 | 728 | | | | | | | 1104 | 657 | 1793 | 1305 | | | |
| | | 701 | 493 | 917 | | | | | | | | 4886 | 3435 | 6966 |
| | | 951 | 714 | 1249 | | | | | | | | 3853 | 2895 | 4053 |
| | | | | | 952 | 573 | 882 | | | | | | | |
| | | | | | 929 | 655 | 991 | | | | | | | |
| 600 | 1189 | | | | | | | 1643 | 1219 | 2032 | 2417 | | | |
| | | 1208 | 668 | 796 | | | | | | | | 6111 | 3379 | 5059 |
| | | 673 | 493 | 1031 | | | | | | | | 2367 | 1733 | 3517 |
| | | | | | 561 | 458 | 792 | | | | | | | |
| 469 | 769 | | | | | | | 6401 | 3436 | 7321 | 5628 | | | |
| 622 | 845 | | | | | | | 2400 | 1941 | 3119 | 2637 | | | |
| 530 | 862 | | | | | | | 5755 | 3929 | 7413 | 6391 | | | |
| | | 957 | 696 | 928 | | | | | | | | 4007 | 2916 | 4188 |
| 1123 | 1379 | | | | | | | 1923 | 1768 | 1575 | 2173 | | | |
| | | | | | 861 | 595 | 874 | | | | | | | |
| 752 | 1426 | | | | | | | 4019 | 3405 | 4530 | 6458 | | | |
| | | 1435 | 1048 | 1127 | | | | | | | | 5618 | 4100 | 3914 |
| | | | | | 643 | 374 | 907 | | | | | | | |
| | | 1109 | 1353 | 1512 | | | | | | | | 3680 | 4490 | 3319 |
| 391 | 853 | | | | | | | 4374 | 2323 | 5944 | 5073 | | | |
| | | 701 | 468 | 678 | | | | | | | | 340 | 227 | 485 |
| | | | | | 903 | 392 | 528 | | | | | | | |
| | | | | | 858 | 435 | 705 | | | | | | | |
| | | 1021 | 304 | 584 | | | | | | | | 3171 | 945 | 3107 |
| | | | | | 630 | 408 | 1481 | | | | | | | |
| 1091 | 1329 | | | | | | | 1735 | 1410 | 1292 | 1717 | | | |
| 511 | 762 | | | | | | | 3276 | 2062 | 4037 | 3075 | | | |
| | | 681 | 513 | 625 | | | | | | | | 2235 | 1682 | 3281 |
| 721 | 831 | | | | | | | 697 | 509 | 706 | 586 | | | |
| 615 | 1148 | | | | | | | 1819 | 1607 | 2612 | 2998 | | | |
| | | 1007 | 938 | 1461 | | | | | | | | 1636 | 1524 | 1624 |

| | | | | | | | | | | | | | | |
|---|---|---|---|---|---|---|---|---|---|---|---|---|---|---|
| | | | | | 1016 | 789 | 1070 | | | | | | | |
| 968 | 1203 | | | | | | | 2069 | 1813 | 1873 | 2253 | | | |
| | | 788 | 625 | 927 | | | | | | | | 1242 | 986 | 1577 |
| 799 | 1056 | | | | | | | 2726 | 2117 | 2651 | 2799 | | | |
| 391 | 791 | | | | | | | 3741 | 1772 | 4532 | 3583 | | | |
| 586 | 1036 | | | | | | | 6430 | 4575 | 7806 | 8084 | | | |
| | | | | | 1152 | 1200 | 1145 | | | | | | | |
| | | 955 | 895 | 1176 | | | | | | | | 1357 | 1272 | 1421 |
| | | 1196 | 896 | 1158 | | | | | | | | 1356 | 1016 | 1134 |
| | | 934 | 676 | 890 | | | | | | | | 2874 | 2080 | 3076 |
| 736 | 962 | | | | | | | 3873 | 4117 | 5592 | 5382 | | | |
| | | | | | 744 | 581 | 806 | | | | | | | |
| 652 | 947 | | | | | | | 2503 | 2125 | 3260 | 3088 | | | |
| | | | | | 813 | 597 | 954 | | | | | | | |
| 832 | 1141 | | | | | | | 2802 | 2470 | 2970 | 3389 | | | |
| | | 653 | 454 | 679 | | | | | | | | 2276 | 1581 | 3486 |
| | | 1048 | 825 | 1096 | | | | | | | | 6072 | 4778 | 5792 |
| | | | | | 860 | 548 | 915 | | | | | | | |
| | | 734 | 468 | 806 | | | | | | | | 3817 | 2436 | 5202 |
| | | | | | 707 | 389 | 682 | | | | | | | |
| | | 708 | 793 | 992 | | | | | | | | 197 | 221 | 279 |
| | | | | | 695 | 478 | 733 | | | | | | | |
| | | | | | 498 | 137 | 286 | | | | | | | |
| | | 794 | 581 | 866 | | | | | | | | 1021 | 747 | 1286 |
| | | | | | 881 | 809 | 1273 | | | | | | | |
| | | | | | 362 | 217 | 548 | | | | | | | |
| 449 | 911 | | | | | | | 4063 | 2874 | 6400 | 5829 | | | |
| | | | | | 912 | 671 | 972 | | | | | | | |
| | | 978 | 1004 | 1557 | | | | | | | | 3101 | 3183 | 3170 |
| | | 843 | 500 | 868 | | | | | | | | 4299 | 2549 | 5100 |

| | | | | | | | | | | | | | | |
|---|---|---|---|---|---|---|---|---|---|---|---|---|---|---|
| | | | | | 643 | 481 | 790 | | | | | | | |
| | | | | | 1125 | 1139 | 1852 | | | | | | | |
| 363 | 691 | | | | | | | 388 | 222 | 611 | 422 | | | |
| | | 816 | 455 | 756 | | | | | | | | 1227 | 685 | 1504 |
| 42 | 135 | | | | | | | 3413 | 328 | 7810 | 1051 | | | |
| | | | | | 832 | 498 | 762 | | | | | | | |
| 523 | 1018 | | | | | | | 2983 | 2232 | 4263 | 4341 | | | |
| | | 839 | 575 | 976 | | | | | | | | 3937 | 2699 | 4694 |
| 610 | 898 | | | | | | | 1966 | 1490 | 2443 | 2194 | | | |
| | | 804 | 747 | 1198 | | | | | | | | 2241 | 2080 | 2786 |
| | | | | | 860 | 609 | 991 | | | | | | | |
| 548 | 916 | | | | | | | 1714 | 1369 | 2500 | 2290 | | | |
| | | | | | 1149 | 1078 | 659 | | | | | | | |
| 275 | 563 | | | | | | | 495 | 292 | 1063 | 598 | | | |
| | | 802 | 579 | 847 | | | | | | | | 3070 | 2217 | 3829 |
| | | | | | 748 | 493 | 772 | | | | | | | |
| 472 | 508 | | | | | | | 2213 | 1561 | 3309 | 1682 | | | |
| 424 | 738 | | | | | | | 3566 | 2290 | 5403 | 3987 | | | |
| | | | | | 436 | 274 | 639 | | | | | | | |
| | | | | | 805 | 586 | 826 | | | | | | | |
| | | | | | 796 | 630 | 822 | | | | | | | |
| 610 | 768 | | | | | | | 3027 | 2032 | 3329 | 2558 | | | |
| 597 | 916 | | | | | | | 2283 | 1444 | 2421 | 2217 | | | |
| | | | | | 679 | 534 | 881 | | | | | | | |
| | | 1122 | 719 | 1000 | | | | | | | | 6131 | 3927 | 5465 |
| | | | | | 988 | 886 | 1131 | | | | | | | |

| | | | | | | | | | | | | | | |
|---|---|---|---|---|---|---|---|---|---|---|---|---|---|---|
| 245 | 412 | | | | | | | 6057 | 1964 | 8017 | 3299 | | | |
| 577 | 899 | | | | | | | 1218 | 829 | 1439 | 1294 | | | |
| | | 840 | 644 | 898 | | | | | | | | 1189 | 912 | 1415 |
| | | | | | 736 | 704 | 1314 | | | | | | | |
| | | 536 | 255 | 550 | | | | | | | | 5317 | 2532 | 9912 |
| | | 700 | 439 | 789 | | | | | | | | 2230 | 1397 | 3184 |
| | | | | | 885 | 655 | 1168 | | | | | | | |
| | | 701 | 627 | 1116 | | | | | | | | 4446 | 3975 | 6343 |
| 592 | 1045 | | | | | | | 5888 | 4323 | 7300 | 7631 | | | |
| | | 722 | 533 | 883 | | | | | | | | 2055 | 1515 | 2844 |
| | | 727 | 617 | 970 | | | | | | | | 2883 | 2446 | 3964 |
| | | | | | 598 | 396 | 865 | | | | | | | |
| | | | | | 908 | 1533 | 2174 | | | | | | | |
| | | | | | 889 | 664 | 981 | | | | | | | |
| 534 | 761 | | | | | | | 1068 | 678 | 1270 | 967 | | | |
| | | | | | 1343 | 711 | 904 | | | | | | | |
| | | 658 | 569 | 973 | | | | | | | | 2101 | 1816 | 3194 |
| | | | | | 953 | 602 | 919 | | | | | | | |
| | | | | | 443 | 205 | 503 | | | | | | | |
| | | | | | 460 | 311 | 752 | | | | | | | |
| 520 | 830 | | | | | | | 1698 | 1102 | 2118 | 1758 | | | |
| | | 1133 | 676 | 818 | | | | | | | | 7577 | 4520 | 6685 |
| | | 779 | 565 | 886 | | | | | | | | 1350 | 980 | 1733 |
| | | | | | 784 | 564 | 816 | | | | | | | |
| | | 596 | 472 | 805 | | | | | | | | 847 | 670 | 1421 |
| | | | | | 670 | 431 | 660 | | | | | | | |

| | | | | | | | | | | | | | | |
|---|---|---|---|---|---|---|---|---|---|---|---|---|---|---|
| | | | | | 807 | 538 | 680 | | | | | | | |
| 565 | 862 | | | | | | | 7597 | 5378 | 9515 | 8201 | | | |
| | | | | | 619 | 97 | 208 | | | | | | | |
| | | | | | 989 | 845 | 1085 | | | | | | | |
| 853 | 1006 | | | | | | | 3919 | 2707 | 3174 | 3193 | | | |
| | | 754 | 716 | 1152 | | | | | | | | 1538 | 1461 | 2039 |
| | | 622 | 434 | 771 | | | | | | | | 3244 | 2259 | 5211 |
| 520 | 913 | | | | | | | 1172 | 820 | 1576 | 1438 | | | |
| | | 994 | 821 | 1255 | | | | | | | | 2559 | 2113 | 2573 |
| | | | | | 795 | 698 | 948 | | | | | | | |
| | | 1077 | 793 | 1055 | | | | | | | | 5132 | 3782 | 4767 |
| | | | | | 531 | 282 | 541 | | | | | | | |
| | | | | | 696 | 530 | 918 | | | | | | | |
| | | | | | 973 | 609 | 1003 | | | | | | | |
| | | 782 | 486 | 625 | | | | | | | | 7617 | 4736 | 9742 |
| 449 | 702 | | | | | | | 914 | 578 | 1287 | 903 | | | |
| | | | | | 805 | 582 | 691 | | | | | | | |
| | | | | | 727 | 452 | 760 | | | | | | | |
| 1068 | 1260 | | | | | | | 3967 | 3495 | 3272 | 4124 | | | |
| 255 | 587 | | | | | | | 815 | 533 | 2093 | 1230 | | | |
| | | 708 | 772 | 1073 | | | | | | | | 1840 | 2006 | 2599 |
| | | 931 | 664 | 914 | | | | | | | | 1987 | 1416 | 2134 |
| 357 | 608 | | | | | | | 1014 | 893 | 2504 | 1522 | | | |
| 465 | 834 | | | | | | | 774 | 480 | 1031 | 860 | | | |
| | | 600 | 624 | 1013 | | | | | | | | 4023 | 4183 | 6706 |

| | | | | | | | | | | | | | | |
|---|---|---|---|---|---|---|---|---|---|---|---|---|---|---|
| 644 | 870 | | | | | | | 7756 | 6145 | 9535 | 8294 | | | |
| | | | | | 693 | 530 | 819 | | | | | | | |
| | | 752 | 646 | 949 | | | | | | | | 2148 | 1844 | 2855 |
| 520 | 806 | | | | | | | 5920 | 3812 | 7326 | 5907 | | | |
| | | 923 | 575 | 642 | | | | | | | | 4122 | 2569 | 4467 |
| | | 861 | 818 | 1046 | | | | | | | | 2984 | 2835 | 3467 |
| | | | | | 502 | 332 | 656 | | | | | | | |
| | | | | | 855 | 845 | 1278 | | | | | | | |
| | | 1022 | 744 | 1067 | | | | | | | | 944 | 687 | 924 |
| | | 647 | 292 | 532 | | | | | | | | 2567 | 1159 | 3965 |
| 652 | 1286 | | | | | | | 3471 | 2939 | 4509 | 5797 | | | |
| | | | | | 485 | 169 | 393 | | | | | | | |
| | | 524 | 295 | 634 | | | | | | | | 3567 | 2008 | 6805 |
| | | 1022 | 952 | 1259 | | | | | | | | 3745 | 3487 | 3662 |
| | | 461 | 212 | 394 | | | | | | | | 4104 | 1888 | 8908 |
| 450 | 750 | | | | | | | 863 | 602 | 1337 | 1003 | | | |
| 604 | 1000 | | | | | | | 2635 | 2573 | 4258 | 4258 | | | |
| | | | | | 401 | 344 | 897 | | | | | | | |
| 365 | 617 | | | | | | | 1015 | 800 | 2195 | 1355 | | | |
| | | | | | 670 | 530 | 833 | | | | | | | |
| | | 836 | 197 | 420 | | | | | | | | 9028 | 2131 | 10804 |
| 358 | 518 | | | | | | | 3398 | 2232 | 6244 | 3232 | | | |
| | | 641 | 436 | 747 | | | | | | | | 1825 | 1240 | 2845 |
| | | | | | 859 | 509 | 644 | | | | | | | |
| | | | | | 714 | 339 | 630 | | | | | | | |

| | | | | | | | | | | | | | | |
|---|---|---|---|---|---|---|---|---|---|---|---|---|---|---|
| | | | | | 1017 | 665 | 1088 | | | | | | | |
| 317 | 416 | | | | | | | 274 | 322 | 1016 | 422 | | | |
| | | 718 | 510 | 1009 | | | | | | | | 4046 | 2873 | 5632 |
| | | | | | 874 | 543 | 831 | | | | | | | |
| 574 | 933 | | | | | | | 3210 | 2044 | 3562 | 3325 | | | |
| | | 760 | 475 | 801 | | | | | | | | 3210 | 2006 | 4224 |
| 497 | 1067 | | | | | | | 1631 | 1182 | 2380 | 2541 | | | |
| | | | | | 552 | 10 | 508 | | | | | | | |
| | | | | | 614 | 520 | 1070 | | | | | | | |
| 1842 | 1435 | | | | | | | 2961 | 8902 | 4833 | 6934 | | | |
| | | | | | 422 | 229 | 556 | | | | | | | |
| | | 665 | 596 | 825 | | | | | | | | 2608 | 2336 | 3920 |
| | | 387 | 337 | 769 | | | | | | | | 1132 | 983 | 2921 |
| | | 989 | 946 | 1529 | | | | | | | | 5926 | 5667 | 5991 |
| | | 708 | 437 | 752 | | | | | | | | 1786 | 1104 | 2523 |
| 581 | 1053 | | | | | | | 3889 | 2777 | 4777 | 5029 | | | |
| | | | | | 908 | 988 | 1390 | | | | | | | |
| 628 | 898 | | | | | | | 2680 | 2101 | 3347 | 3004 | | | |
| | | | | | 579 | 378 | 596 | | | | | | | |
| 709 | 1094 | | | | | | | 3591 | 2358 | 3328 | 3641 | | | |
| 611 | 872 | | | | | | | 2143 | 1473 | 2411 | 2102 | | | |

| | | | | | | | | | | | | | | |
|---|---|---|---|---|---|---|---|---|---|---|---|---|---|---|
| 326 | 577 | | | | | | | 2646 | 1528 | 4681 | 2702 | | | |
| 1160 | 1720 | | | | | | | 2208 | 2219 | 1914 | 3293 | | | |
| 598 | 878 | | | | | | | 2073 | 2036 | 3402 | 2986 | | | |
| | | 645 | 653 | 1036 | | | | | | | | 5631 | 5703 | 8729 |
| 982 | 1474 | | | | | | | 2636 | 3321 | 3384 | 4988 | | | |
| 1167 | 1397 | | | | | | | 977 | 1180 | 1011 | 1412 | | | |
| 506 | 588 | | | | | | | 2817 | 5114 | 10117 | 5948 | | | |
| 596 | 1046 | | | | | | | 4173 | 2983 | 5006 | 5237 | | | |
| | | | | | 1030 | 889 | 1084 | | | | | | | |
| | | 947 | 821 | 1391 | | | | | | | | 2375 | 2059 | 2507 |
| 593 | 956 | | | | | | | 1729 | 1458 | 2460 | 2352 | | | |
| | | 921 | 716 | 1246 | | | | | | | | 2999 | 2332 | 3256 |
| | | | | | 795 | 687 | 1073 | | | | | | | |
| 449 | 812 | | | | | | | 735 | 516 | 1150 | 934 | | | |
| | | | | | 932 | 952 | 1681 | | | | | | | |
| | | 1000 | 696 | 1215 | | | | | | | | 6846 | 4762 | 6847 |
| | | | | | 643 | 586 | 1028 | | | | | | | |
| | | 184 | 59 | 126 | | | | | | | | 2225 | 709 | 12095 |
| | | | | | 778 | 525 | 885 | | | | | | | |
| 401 | 608 | | | | | | | 6001 | 3632 | 9054 | 5504 | | | |
| 51 | 139 | | | | | | | 14369 | 862 | 16748 | 2336 | | | |

| | | | | | | | | | | | | | | |
|---|---|---|---|---|---|---|---|---|---|---|---|---|---|---|
| | | 709 | 615 | 878 | | | | | | | | 5298 | 4598 | 7474 |
| | | 507 | 399 | 1465 | | | | | | | | 1572 | 1238 | 3101 |
| | | | | | 528 | 356 | 687 | | | | | | | |
| | | 605 | 303 | 413 | | | | | | | | 3694 | 1854 | 6109 |
| | | | | | 700 | 516 | 904 | | | | | | | |
| | | | | | 1029 | 1510 | 1587 | | | | | | | |
| | | 799 | 574 | 1108 | | | | | | | | 1716 | 1233 | 2146 |
| | | 785 | 792 | 1064 | | | | | | | | 4857 | 4900 | 6185 |
| | | | | | 740 | 490 | 1015 | | | | | | | |
| | | | | | 773 | 530 | 1037 | | | | | | | |
| | | | | | 713 | 496 | 916 | | | | | | | |
| | | | | | 795 | 196 | 295 | | | | | | | |
| 393 | 2057 | | | | | | | 638 | 427 | 1088 | 2238 | | | |
| | | | | | 698 | 426 | 644 | | | | | | | |
| 504 | 939 | | | | | | | 2865 | 2239 | 4446 | 4174 | | | |
| 449 | 734 | | | | | | | 4452 | 3324 | 7404 | 5436 | | | |
| | | 425 | 444 | 742 | | | | | | | | 2761 | 2881 | 6493 |
| | | 615 | 486 | 1068 | | | | | | | | 5463 | 4316 | 8878 |
| | | 608 | 497 | 883 | | | | | | | | 5003 | 4089 | 8234 |
| | | | | | 632 | 381 | 709 | | | | | | | |
| | | | | | 724 | 517 | 790 | | | | | | | |
| | | | | | 690 | 552 | 1015 | | | | | | | |
| | | 541 | 651 | 943 | | | | | | | | 4169 | 5017 | 7703 |
| 381 | 711 | | | | | | | 1569 | 1085 | 2850 | 2026 | | | |
| | | | | | 1034 | 605 | 1010 | | | | | | | |
| 527 | 843 | | | | | | | 1091 | 667 | 1267 | 1068 | | | |

| | | | | | | | | |
|---|---|---|---|---|---|---|---|---|
| | | | 735 | 392 | 1167 | | | |
| 945 | 986 | 1283 | | | | 885 | 924 | 937 |
| | | | 1083 | 834 | 1069 | | | |
| | | | 822 | 616 | 1035 | | | |
| | | | 971 | 768 | 1000 | | | |
| 605 | 365 | 853 | | | | 1497 | 903 | 2474 |

| Abundanc | Abundanc | Abundanc | Abundanc | Abundanc | Found in S | Found in S | Found in S | Found in S | Found in S | Found in S | Found in S | Found in S | Found in S | Found in S |
|---|---|---|---|---|---|---|---|---|---|---|---|---|---|---|
| 2701491 | 2014585 | 1638391 | 3662206 | 2741739 | High | High | High | High | High | High | High | High | High | High |
| 1443138 | 1321140 | 1003151 | 2409983 | 1349233 | High | High | High | High | High | High | High | High | High | High |
| 3218069 | 1602637 | 1704358 | 2886291 | 2948483 | High | High | High | High | High | High | High | High | High | High |
| 240917 | 169962 | 132163 | 215030 | 175972 | High | High | High | High | High | High | High | High | High | High |
| 586702 | 415246 | 344684 | 640510 | 540696 | High | High | High | High | High | High | High | High | High | High |
| 4163 | 1900 | 797 | 1534 | 1265 | High | High | High | High | High | High | High | High | High | High |
| 1592 | 1273 | 946 | 1324 | 1273 | Not Found | Not Found | Not Found | Not Found | High | High | High | High | High | High |
| 125994 | 129198 | 93181 | 143085 | 131368 | High | High | High | High | High | High | High | High | High | High |
| 59077 | 46212 | 38119 | 56637 | 51849 | High | High | High | High | High | High | High | High | High | High |
| 532402 | 417146 | 304834 | 477159 | 484197 | High | High | High | High | High | High | High | High | High | High |
| 44721 | 14460 | 11222 | 14667 | 37188 | High | High | High | High | High | High | High | High | High | High |
| 464137 | 304097 | 281798 | 328340 | 422093 | High | High | High | High | High | High | High | High | High | High |
| | 1057 | 787 | 906 | 1506 | Not Found | Not Found | Not Found | Not Found | Not Found | Not Found | Not Found | Not Found | High | High |
| 508372 | 421201 | 306959 | 548729 | 499911 | High | High | High | High | High | High | High | High | High | High |
| 350656 | 272302 | 251703 | 294485 | 345741 | High | High | High | High | High | High | High | High | High | High |
| 167418 | 134603 | 107013 | 203436 | 168757 | High | High | High | High | High | High | High | High | High | High |
| 88081 | 69909 | 54376 | 85301 | 82413 | High | High | High | High | High | High | High | High | High | High |
| 270168 | 333350 | 199071 | 390460 | 310864 | High | High | High | High | High | High | High | High | High | High |
| 3717 | 3260 | 2011 | 4223 | 2650 | Not Found | Not Found | Not Found | Not Found | High | High | High | High | High | High |
| 353303 | 205479 | 207370 | 242089 | 316017 | High | High | High | High | High | High | High | High | High | High |
| 244553 | 237400 | 199970 | 269569 | 322490 | High | High | High | High | High | High | High | High | High | High |
| 343583 | 299654 | 212328 | 319372 | 326303 | High | High | High | High | High | High | High | High | High | High |
| 82552 | 19382 | 13559 | 17791 | 74612 | High | High | High | High | High | High | High | High | High | High |
| 328943 | 323217 | 206780 | 463273 | 360653 | High | High | High | High | High | High | High | High | High | High |
| 362428 | 328353 | 187222 | 391358 | 329059 | High | High | High | High | High | High | High | High | High | High |
| 406366 | 262987 | 192877 | 362203 | 318208 | High | High | High | High | High | High | High | High | High | High |
| 179143 | 129356 | 111092 | 170456 | 172166 | High | High | High | High | High | High | High | High | High | High |
| 10373 | 4156 | 4525 | 12196 | 9675 | High | High | High | High | High | High | High | High | High | High |
| 211799 | 214391 | 166746 | 252422 | 246824 | High | High | High | High | High | High | High | High | High | High |
| 1437 | 803 | 739 | 765 | 967 | High | High | High | High | High | High | High | High | High | High |
| 292456 | 247765 | 202682 | 290687 | 303171 | High | High | High | High | High | High | High | High | High | High |

| | | | | | | | | | | | | | | |
|---|---|---|---|---|---|---|---|---|---|---|---|---|---|---|
| | | | | | High | High | High | High | Not Founc | Not Founc | Not Founc | Not Founc | Not Founc | Not Founc |
| 282460 | 231323 | 190508 | 291866 | 314308 | High | High | High | High | High | High | High | High | High | High |
| 360618 | 225530 | 199350 | 352677 | 369655 | High | High | High | High | High | High | High | High | High | High |
| 265643 | 248701 | 182269 | 299770 | 297206 | High | High | High | High | High | High | High | High | High | High |
| 339827 | 209865 | 194307 | 224423 | 303498 | High | High | High | High | High | High | High | High | High | High |
| 285088 | 210133 | 188219 | 255937 | 297328 | High | High | High | High | High | High | High | High | High | High |
| 263087 | 209551 | 151979 | 306889 | 256589 | High | High | High | High | High | High | High | High | High | High |
| 235871 | 177945 | 141331 | 240060 | 232117 | High | High | High | High | High | High | High | High | High | High |
| 237444 | 160196 | 149318 | 233610 | 242487 | High | High | High | High | High | High | High | High | High | High |
| 243413 | 157072 | 146501 | 192610 | 238994 | High | High | High | High | High | High | High | High | High | High |
| 221490 | 203489 | 179350 | 318856 | 234848 | High | High | High | High | High | High | High | High | High | High |
| 197120 | 102176 | 93245 | 141878 | 159006 | High | High | High | High | High | High | High | High | High | High |
| 251107 | 210325 | 186267 | 231361 | 281194 | High | High | High | High | High | High | High | High | High | High |
| 198402 | 224410 | 157112 | 228478 | 207370 | High | High | High | High | High | High | High | High | High | High |
| 229254 | 183917 | 131313 | 249449 | 205862 | High | High | High | High | High | High | High | High | High | High |
| 236188 | 157342 | 150828 | 181037 | 203336 | High | High | High | High | High | High | High | High | High | High |
| 280481 | 162001 | 152998 | 192645 | 254612 | High | High | High | High | High | High | High | High | High | High |
| 297340 | 154081 | 133685 | 340771 | 308296 | High | High | High | High | High | High | High | High | High | High |
| 275227 | 181346 | 128755 | 269672 | 233606 | High | High | High | High | High | High | High | High | High | High |
| 269077 | 154948 | 157706 | 187856 | 246447 | High | High | High | High | High | High | High | High | High | High |
| 18643 | 9121 | 7097 | 16824 | 13042 | High | High | High | High | High | High | High | High | High | High |
| 19471 | 12806 | 14895 | 14547 | 23196 | High | High | High | High | High | High | High | High | High | High |
| 150328 | 99036 | 89746 | 100754 | 121538 | High | High | High | High | High | High | High | High | High | High |
| 6383 | | | | | High | High | High | High | High | High | High | High | High | High |
| | | | | | High | High | High | High | Not Founc | Not Founc | Not Founc | Not Founc | Not Founc | Not Founc |
| 337727 | 142539 | 137618 | 175498 | 262454 | High | High | High | High | High | High | High | High | High | High |
| 215527 | 157260 | 125553 | 189666 | 201084 | High | High | High | High | High | High | High | High | High | High |
| 299452 | 200648 | 137220 | 293603 | 252888 | High | High | High | High | High | High | High | High | High | High |
| 203457 | 190503 | 133986 | 221834 | 196753 | High | High | High | High | High | High | High | High | High | High |
| 272877 | 169275 | 159793 | 215289 | 200302 | High | High | High | High | High | High | High | High | High | High |
| 148335 | 159379 | 128672 | 178468 | 190042 | High | High | High | High | High | High | High | High | High | High |

| | | | | | | | | | | | | | | |
|---|---|---|---|---|---|---|---|---|---|---|---|---|---|---|
| 222113 | 186882 | 166518 | 243961 | 266333 | High | High | High | High | High | High | High | High | High | High |
| 226343 | 185306 | 137093 | 186488 | 185621 | High | High | High | High | High | High | High | High | High | High |
| 237974 | 129928 | 90863 | 270435 | 216624 | High | High | High | High | High | High | High | High | High | High |
| 199180 | 142435 | 121602 | 179930 | 185383 | High | High | High | High | High | High | High | High | High | High |
| 241710 | 192612 | 183556 | 202875 | 254132 | High | High | High | High | High | High | High | High | High | High |
| 214393 | 145779 | 119325 | 191567 | 191246 | High | High | High | High | High | High | High | High | High | High |
| 155024 | 244391 | 107644 | 175200 | 139259 | High | High | High | High | High | High | High | High | High | High |
| 218154 | 181752 | 163581 | 213461 | 232260 | High | High | High | High | High | High | High | High | High | High |
| 222065 | 170089 | 133470 | 236681 | 208738 | High | High | High | High | High | High | High | High | High | High |
| 224216 | 126441 | 97383 | 175262 | 188800 | High | High | High | High | High | High | High | High | High | High |
| 191858 | 129275 | 138461 | 124842 | 205549 | High | High | High | High | High | High | High | High | High | High |
| 273982 | 193616 | 194314 | 194292 | 265045 | High | High | High | High | High | High | High | High | High | High |
| 172153 | 122081 | 142779 | 121052 | 153523 | High | High | High | High | High | High | High | High | High | High |
| 201161 | 117530 | 103575 | 169387 | 195519 | High | High | High | High | High | High | High | High | High | High |
| 137565 | 115357 | 82210 | 139805 | 136668 | High | High | High | High | High | High | High | High | High | High |
| 193854 | 138252 | 144829 | 153910 | 217223 | High | High | High | High | High | High | High | High | High | High |
| 157663 | 156940 | 106748 | 216351 | 178336 | High | High | High | High | High | High | High | High | High | High |
| 197233 | 155438 | 117221 | 172608 | 191531 | High | High | High | High | High | High | High | High | High | High |
| 158974 | 90188 | 78620 | 105105 | 124028 | High | High | High | High | High | High | High | High | High | High |
| 127170 | 93941 | 95823 | 127667 | 158727 | High | High | High | High | High | High | High | High | High | High |
| 220368 | 152867 | 132017 | 193660 | 217994 | High | High | High | High | High | High | High | High | High | High |
| 163377 | 177027 | 134999 | 194214 | 208291 | High | High | High | High | High | High | High | High | High | High |
| 110071 | 64051 | 56784 | 86654 | 97342 | High | High | High | High | High | High | High | High | High | High |
| 238920 | 119333 | 106219 | 233431 | 228416 | High | High | High | High | High | High | High | High | High | High |
| 116816 | 66798 | 62459 | 69270 | 93089 | High | High | High | High | High | High | High | High | High | High |
| 154184 | 144423 | 136151 | 178291 | 166865 | High | High | High | High | High | High | High | High | High | High |
| 23978 | 10924 | 7144 | 16765 | 12000 | High | High | High | High | High | High | High | High | High | High |
| 4523 | 9387 | 5151 | 7068 | 5865 | High | High | High | High | High | High | High | High | High | High |
| 139408 | 180751 | 119377 | 219222 | 200544 | High | High | High | High | High | High | High | High | High | High |
| 172569 | 125750 | 99441 | 144455 | 162042 | High | High | High | High | High | High | High | High | High | High |
| 159843 | 147600 | 107320 | 228698 | 201304 | High | High | High | High | High | High | High | High | High | High |

| | | | | | | | | | | | | | | |
|---|---|---|---|---|---|---|---|---|---|---|---|---|---|---|
| 195437 | 153679 | 131682 | 172006 | 218034 | High | High | High | High | High | High | High | High | High | High |
| 215169 | 124984 | 160233 | 146406 | 228771 | High | High | High | High | High | High | High | High | High | High |
| 186826 | 161704 | 129072 | 210428 | 203573 | High | High | High | High | High | High | High | High | High | High |
| 174003 | 110511 | 89076 | 130079 | 144502 | High | High | High | High | High | High | High | High | High | High |
| 34818 | 17096 | 13056 | 22003 | 19472 | High | High | High | High | High | High | High | High | High | High |
| 104896 | 75666 | 75859 | 111071 | 132806 | High | High | High | High | High | High | High | High | High | High |
| 228134 | 115904 | 108706 | 210702 | 203253 | High | High | High | High | High | High | High | High | High | High |
| 12235 | 9707 | 7471 | 9243 | 18213 | High | High | High | High | High | High | High | High | High | High |
| | | | | | High | High | High | High | Not Found | Not Found | Not Found | Not Found | Not Found | Not Found |
| 178482 | 103547 | 90004 | 123660 | 147087 | High | High | High | High | High | High | High | High | High | High |
| 119928 | 132529 | 94288 | 139655 | 158574 | High | High | High | High | High | High | High | High | High | High |
| 168470 | 100574 | 106450 | 110238 | 152488 | High | High | High | High | High | High | High | High | High | High |
| 144664 | 99563 | 86418 | 105695 | 129042 | High | High | High | High | High | High | High | High | High | High |
| 206366 | 109159 | 87206 | 169405 | 160481 | High | High | High | High | High | High | High | High | High | High |
| | | | | | Not Found | Not Found | Not Found | Not Found | Not Found | Not Found | Not Found | Not Found | Not Found | Not Found |
| 151555 | 96648 | 90475 | 104545 | 135257 | High | High | High | High | High | High | High | High | High | High |
| 182317 | 151782 | 136869 | 196924 | 215385 | High | High | High | High | High | High | High | High | High | High |
| 153177 | 125329 | 99921 | 151754 | 157652 | High | High | High | High | High | High | High | High | High | High |
| 224553 | 112285 | 102944 | 153851 | 184613 | High | High | High | High | High | High | High | High | High | High |
| 5145 | 419 | 369 | 501 | 532 | Not Found | Not Found | Not Found | Not Found | High | High | High | High | High | High |
| 152966 | 117359 | 101811 | 144285 | 161521 | High | High | High | High | High | High | High | High | High | High |
| 3918 | 4371 | 2616 | 3360 | 3261 | High | High | High | High | High | High | High | High | High | High |
| 1781 | 3014 | 1864 | 3355 | 3160 | Not Found | Not Found | Not Found | Not Found | High | High | High | High | High | High |
| 167088 | 102228 | 86751 | 129248 | 141035 | High | High | High | High | High | High | High | High | High | High |
| 66803 | 38273 | 33934 | 46427 | 47959 | High | High | High | High | High | High | High | High | High | High |
| 171401 | 129410 | 131048 | 155722 | 199829 | High | High | High | High | High | High | High | High | High | High |
| 222298 | 137997 | 105025 | 166772 | 178174 | High | High | High | High | High | High | High | High | High | High |
| 148398 | 115408 | 79682 | 169628 | 143931 | High | High | High | High | High | High | High | High | High | High |
| 4558 | 7424 | 4522 | 6242 | 5509 | High | High | High | High | High | High | High | High | High | High |
| 163470 | 118688 | 94958 | 129183 | 147325 | High | High | High | High | High | High | High | High | High | High |
| 77808 | 51736 | 46232 | 56666 | 64036 | High | High | High | High | High | High | High | High | High | High |

| | | | | | | | | | | | | | | |
|---|---|---|---|---|---|---|---|---|---|---|---|---|---|---|
| 169711 | 133877 | 116632 | 179524 | 189028 | High | High | High | High | High | High | High | High | High | High |
| 182010 | 87708 | 89748 | 135168 | 159768 | High | High | High | High | High | High | High | High | High | High |
| 105408 | 101733 | 78864 | 131289 | 128863 | High | High | High | High | High | High | High | High | High | High |
| 169895 | 165187 | 123419 | 190138 | 200397 | High | High | High | High | High | High | High | High | High | High |
| 152255 | 130682 | 115074 | 140393 | 173744 | High | High | High | High | High | High | High | High | High | High |
| 119507 | 121963 | 109519 | 132298 | 156437 | High | High | High | High | High | High | High | High | High | High |
| 128461 | 101689 | 91806 | 133597 | 149948 | High | High | High | High | High | High | High | High | High | High |
| 123423 | 97262 | 91818 | 114848 | 131517 | High | High | High | High | High | High | High | High | High | High |
| 168004 | 138958 | 74210 | 201206 | 164979 | High | High | High | High | High | High | High | High | High | High |
| 147790 | 124295 | 110210 | 142353 | 179974 | High | High | High | High | High | High | High | High | High | High |
| 168053 | 133382 | 95677 | 179740 | 176662 | High | High | High | High | High | High | High | High | High | High |
| 125077 | 68086 | 54187 | 94597 | 93497 | High | High | High | High | High | High | High | High | High | High |
| 140230 | 80816 | 64256 | 95994 | 102609 | High | High | High | High | High | High | High | High | High | High |
| 98838 | 72933 | 62982 | 86878 | 102855 | High | High | High | High | High | High | High | High | High | High |
| 10694 | 17869 | 15336 | 17762 | 22563 | High | High | High | High | High | High | High | High | High | High |
| 140882 | 71539 | 54616 | 124365 | 103518 | High | High | High | High | High | High | High | High | High | High |
| 154723 | 69687 | 50038 | 153168 | 135176 | High | High | High | High | High | High | High | High | High | High |
| 196679 | 137279 | 121030 | 188094 | 194415 | High | High | High | High | High | High | High | High | High | High |
| 172920 | 89541 | 73653 | 136034 | 118953 | High | High | High | High | High | High | High | High | High | High |
| 115535 | 112952 | 89848 | 143607 | 122678 | High | High | High | High | High | High | High | High | High | High |
| 135906 | 92183 | 74406 | 124279 | 110798 | High | High | High | High | High | High | High | High | High | High |
| 173714 | 136692 | 100946 | 148249 | 163497 | High | High | High | High | High | High | High | High | High | High |
| 101714 | 115522 | 102032 | 141466 | 147650 | High | High | High | High | High | High | High | High | High | High |
| 119448 | 96155 | 91155 | 114962 | 136797 | High | High | High | High | High | High | High | High | High | High |
| 164263 | 102119 | 88533 | 118180 | 132646 | High | High | High | High | High | High | High | High | High | High |
| 88246 | 48152 | 40881 | 90107 | 79147 | High | High | High | High | High | High | High | High | High | High |
| 186995 | 119994 | 92930 | 173069 | 173835 | High | High | High | High | High | High | High | High | High | High |
| 112960 | 89766 | 81180 | 130887 | 127728 | High | High | High | High | High | High | High | High | High | High |
| 95633 | 82611 | 70240 | 79267 | 100863 | High | High | High | High | High | High | High | High | High | High |
| 164929 | 102806 | 94054 | 136167 | 152744 | High | High | High | High | High | High | High | High | High | High |
| 94506 | 90076 | 70944 | 96102 | 106167 | High | High | High | High | High | High | High | High | High | High |

| | | | | | | | | | | | | | | |
|---|---|---|---|---|---|---|---|---|---|---|---|---|---|---|
| 12021 | 10829 | 4235 | 14879 | 8577 | High | High | High | High | High | High | High | High | High | High |
| 95035 | 103394 | 93103 | 113325 | 123301 | High | High | High | High | High | High | High | High | High | High |
| 24154 | 17053 | 17329 | 16558 | 21689 | High | High | High | High | High | High | High | High | High | High |
| 156305 | 76193 | 72466 | 96847 | 115992 | High | High | High | High | High | High | High | High | High | High |
| 136879 | 132235 | 82735 | 168097 | 134392 | High | High | High | High | High | High | High | High | High | High |
| 157018 | 92278 | 81626 | 129302 | 149048 | High | High | High | High | High | High | High | High | High | High |
| 119217 | 82509 | 92663 | 120197 | 141062 | High | High | High | High | High | High | High | High | High | High |
| 76630 | 75377 | 66542 | 79807 | 111117 | High | High | High | High | High | High | High | High | High | High |
| 90599 | 84688 | 60940 | 100934 | 107231 | High | High | High | High | High | High | High | High | High | High |
| 86631 | 56849 | 49146 | 65859 | 77995 | High | High | High | High | High | High | High | High | High | High |
| 118297 | 90171 | 72272 | 111038 | 127568 | High | High | High | High | High | High | High | High | High | High |
| 9492 | 19633 | 14575 | 24332 | 22645 | High | High | High | High | High | High | High | High | High | High |
| 116630 | 79029 | 64251 | 98397 | 121085 | High | High | High | High | High | High | High | High | High | High |
| 4948 | 15327 | 7270 | 20392 | 12387 | High | High | High | High | High | High | High | High | High | High |
| 102327 | 87632 | 77047 | 88946 | 106762 | High | High | High | High | High | High | High | High | High | High |
| 101466 | 88961 | 71493 | 133759 | 131015 | High | High | High | High | High | High | High | High | High | High |
| 93948 | 86770 | 79278 | 143965 | 127971 | High | High | High | High | High | High | High | High | High | High |
| 108637 | 96449 | 70042 | 127437 | 124826 | High | High | High | High | High | High | High | High | High | High |
| 15348 | 17123 | 11996 | 15239 | 16161 | High | High | High | High | High | High | High | High | High | High |
| 161993 | 95320 | 77628 | 108099 | 130843 | High | High | High | High | High | High | High | High | High | High |
| 114674 | 88236 | 75019 | 114601 | 119150 | High | High | High | High | High | High | High | High | High | High |
| 6885 | 4163 | 6467 | 3575 | 7860 | High | High | High | High | High | High | High | High | High | High |
| 88713 | 100563 | 73567 | 119298 | 115721 | High | High | High | High | High | High | High | High | High | High |
| 127311 | 103366 | 89860 | 177715 | 147763 | High | High | High | High | High | High | High | High | High | High |
| 128668 | 69387 | 46451 | 92179 | 83640 | High | High | High | High | High | High | High | High | High | High |
| 99467 | 65192 | 53149 | 64070 | 69640 | High | High | High | High | High | High | High | High | High | High |
| 116876 | 84967 | 79802 | 95663 | 111197 | High | High | High | High | High | High | High | High | High | High |
| 148762 | 77743 | 61210 | 97709 | 95537 | High | High | High | High | High | High | High | High | High | High |
| | 3033 | 1528 | 3628 | 2707 | Not Founc | Not Founc | Not Founc | Not Founc | Not Founc | Not Founc | Not Founc | Not Founc | High | High |
| 150914 | 77366 | 65060 | 105652 | 129491 | High | High | High | High | High | High | High | High | High | High |
| 114769 | 74858 | 60633 | 106648 | 105767 | High | High | High | High | High | High | High | High | High | High |

| | | | | | | | | | | | | | | |
|---|---|---|---|---|---|---|---|---|---|---|---|---|---|---|
| 146808 | 102369 | 65415 | 205560 | 149329 | High | High | High | High | High | High | High | High | High | High |
| 177769 | 86955 | 89688 | 99810 | 144817 | High | High | High | High | High | High | High | High | High | High |
| 120540 | 43368 | 32854 | 65512 | 56949 | High | High | High | High | High | High | High | High | High | High |
| 115006 | 71520 | 46570 | 107759 | 93486 | High | High | High | High | High | High | High | High | High | High |
| 124346 | 88965 | 71237 | 86840 | 100176 | High | High | High | High | High | High | High | High | High | High |
| 88765 | 49370 | 49548 | 56138 | 79315 | High | High | High | High | High | High | High | High | High | High |
| 4701 | | | | | Not Founc | Not Founc | Not Founc | Not Founc | High | High | High | High | Not Founc | Not Founc |
| 155975 | 106933 | 93563 | 125308 | 146259 | High | High | High | High | High | High | High | High | High | High |
| 135705 | 119522 | 81835 | 162596 | 165363 | High | High | High | High | High | High | High | High | High | High |
| 97085 | 90821 | 60399 | 135959 | 102253 | High | High | High | High | High | High | High | High | High | High |
| 129725 | 82685 | 68534 | 101074 | 119411 | High | High | High | High | High | High | High | High | High | High |
| 57002 | 43367 | 40331 | 40065 | 55926 | High | High | High | High | High | High | High | High | High | High |
| 1744 | 4207 | 2545 | 4587 | 4063 | High | High | High | High | High | High | High | High | High | High |
| 128893 | 83187 | 77386 | 96528 | 122344 | High | High | High | High | High | High | High | High | High | High |
| 106426 | 91055 | 74010 | 133101 | 134317 | High | High | High | High | High | High | High | High | High | High |
| 59801 | 52797 | 48936 | 60576 | 84837 | High | High | High | High | High | High | High | High | High | High |
| 119761 | 85294 | 74843 | 117405 | 124703 | High | High | High | High | High | High | High | High | High | High |
| 17023 | 15524 | 15676 | 18363 | 21246 | High | High | High | High | High | High | High | High | High | High |
| 75145 | 67711 | 50069 | 88291 | 90131 | High | High | High | High | High | High | High | High | High | High |
| 116447 | 75226 | 67075 | 100377 | 114931 | High | High | High | High | High | High | High | High | High | High |
| 94158 | 55951 | 54007 | 72783 | 79249 | High | High | High | High | High | High | High | High | High | High |
| 141640 | 82265 | 79470 | 124542 | 122630 | High | High | High | High | High | High | High | High | High | High |
| 117865 | 54865 | 45054 | 62113 | 75248 | High | High | High | High | High | High | High | High | High | High |
| 80444 | 106196 | 44968 | 159711 | 67928 | High | High | High | High | High | High | High | High | High | High |
| 133821 | 86873 | 68392 | 133714 | 111707 | High | High | High | High | High | High | High | High | High | High |
| 126133 | 84107 | 63872 | 113367 | 114515 | High | High | High | High | High | High | High | High | High | High |
| 96302 | 77352 | 71156 | 107024 | 107019 | High | High | High | High | High | High | High | High | High | High |
| 169996 | 86432 | 83756 | 101455 | 139190 | High | High | High | High | High | High | High | High | High | High |
| 138250 | 93132 | 72652 | 129183 | 127082 | High | High | High | High | High | High | High | High | High | High |
| 15445 | 4687 | 4512 | 5439 | 7362 | High | High | High | High | High | High | High | High | High | High |
| 12657 | 16920 | 14203 | 18038 | 20576 | High | High | High | High | High | High | High | High | High | High |

| | | | | | | | | | | | | | | |
|---|---|---|---|---|---|---|---|---|---|---|---|---|---|---|
| 120219 | 59083 | 44775 | 58981 | 61222 | High | High | High | High | High | High | High | High | High | High |
| 86958 | 67509 | 56912 | 71844 | 89318 | High | High | High | High | High | High | High | High | High | High |
| 131347 | 73631 | 60470 | 88706 | 109183 | High | High | High | High | High | High | High | High | High | High |
| 87926 | 69105 | 60152 | 92022 | 100630 | High | High | High | High | High | High | High | High | High | High |
| 117100 | 65960 | 58663 | 89118 | 108923 | High | High | High | High | High | High | High | High | High | High |
| 127392 | 145647 | 117402 | 154824 | 156806 | High | High | High | High | High | High | High | High | High | High |
| 60456 | 69134 | 58792 | 72007 | 79474 | High | High | High | High | High | High | High | High | High | High |
| 95321 | 79734 | 62315 | 82444 | 93809 | High | High | High | High | High | High | High | High | High | High |
| 101208 | 90200 | 90790 | 112902 | 127908 | High | High | High | High | High | High | High | High | High | High |
| 158344 | 92255 | 101006 | 95849 | 185181 | High | High | High | High | High | High | High | High | High | High |
| 104294 | 50784 | 43443 | 113049 | 94228 | High | High | High | High | High | High | High | High | High | High |
| 87146 | 61522 | 51186 | 51757 | 61665 | High | High | High | High | High | High | High | High | High | High |
| 68758 | 49759 | 43225 | 57482 | 62716 | High | High | High | High | High | High | High | High | High | High |
| 64905 | 59402 | 43772 | 67692 | 66109 | High | High | High | High | High | High | High | High | High | High |
| 95870 | 85988 | 63091 | 116801 | 94274 | High | High | High | High | High | High | High | High | High | High |
| 93987 | 69097 | 55556 | 92812 | 91071 | High | High | High | High | High | High | High | High | High | High |
| 111476 | 74575 | 64289 | 73426 | 91971 | High | High | High | High | High | High | High | High | High | High |
| 111559 | 83855 | 64984 | 114279 | 102559 | High | High | High | High | High | High | High | High | High | High |
| 1817 | 3809 | 2706 | 3444 | 2771 | High | High | High | High | High | High | High | High | High | High |
| 77788 | 68536 | 50514 | 134362 | 88937 | High | High | High | High | High | High | High | High | High | High |
| 95768 | 83731 | 50366 | 118250 | 96315 | High | High | High | High | High | High | High | High | High | High |
| 83419 | 82282 | 56328 | 111092 | 103782 | High | High | High | High | High | High | High | High | High | High |
| 84744 | 54963 | 56727 | 72895 | 90816 | High | High | High | High | High | High | High | High | High | High |
| 122696 | 62572 | 49158 | 65883 | 69057 | High | High | High | High | High | High | High | High | High | High |
| 85533 | 73412 | 68095 | 82146 | 98958 | High | High | High | High | High | High | High | High | High | High |
| 97757 | 62984 | 72685 | 78134 | 101649 | High | High | High | High | High | High | High | High | High | High |
| 59896 | 43304 | 39474 | 45877 | 55737 | High | High | High | High | High | High | High | High | High | High |
| 122321 | 74742 | 94331 | 97676 | 113470 | High | High | High | High | High | High | High | High | High | High |
| 73251 | 51615 | 45086 | 60692 | 75259 | High | High | High | High | High | High | High | High | High | High |
| 87124 | 71020 | 61917 | 69771 | 89147 | High | High | High | High | High | High | High | High | High | High |
| 56345 | 38993 | 31947 | 42441 | 49740 | High | High | High | High | High | High | High | High | High | High |

| | | | | | | | | | | | | | | |
|---|---|---|---|---|---|---|---|---|---|---|---|---|---|---|
| 86725 | 86787 | 58399 | 162241 | 113350 | High | High | High | High | High | High | High | High | High | High |
| 63381 | 44979 | 43317 | 53104 | 67311 | High | High | High | High | High | High | High | High | High | High |
| 72397 | 63581 | 61534 | 63768 | 93648 | High | High | High | High | High | High | High | High | High | High |
| 38823 | 35457 | 30719 | 39556 | 40422 | High | High | High | High | High | High | High | High | High | High |
| 4925 | 5149 | 3587 | 5902 | 5116 | High | High | High | High | High | High | High | High | High | High |
| 90626 | 67028 | 56127 | 80655 | 90026 | High | High | High | High | High | High | High | High | High | High |
| 64456 | 69289 | 55099 | 71601 | 70979 | High | High | High | High | High | High | High | High | High | High |
| 93285 | 66912 | 49371 | 91354 | 80028 | High | High | High | High | High | High | High | High | High | High |
| 89419 | 72108 | 48555 | 84603 | 78430 | High | High | High | High | High | High | High | High | High | High |
| 34992 | 57204 | 57339 | 53957 | 73219 | High | High | High | High | High | High | High | High | High | High |
| 70857 | 52422 | 48803 | 52355 | 63853 | High | High | High | High | High | High | High | High | High | High |
| 93899 | 68934 | 51863 | 97503 | 94631 | High | High | High | High | High | High | High | High | High | High |
| 60783 | 48436 | 33607 | 63098 | 52254 | High | High | High | High | High | High | High | High | High | High |
| 113149 | 67635 | 54730 | 81056 | 90140 | High | High | High | High | High | High | High | High | High | High |
| 85067 | 72146 | 48599 | 77151 | 81501 | High | High | High | High | High | High | High | High | High | High |
| 92493 | 71342 | 67353 | 65317 | 82134 | High | High | High | High | High | High | High | High | High | High |
| 82347 | 98557 | 76063 | 125133 | 97961 | High | High | High | High | High | High | High | High | High | High |
| 217 | | | | | High | High | High | High | High | High | High | High | Not Found | Not Found |
| 87753 | 60673 | 52119 | 77363 | 80568 | High | High | High | High | High | High | High | High | High | High |
| 122500 | 89837 | 73608 | 139769 | 119622 | High | High | High | High | High | High | High | High | High | High |
| 8179 | 2301 | 2041 | 2542 | 3592 | High | High | High | High | High | High | High | High | High | High |
| | 5838 | 4944 | 8558 | 8626 | Not Found | Not Found | Not Found | Not Found | Not Found | Not Found | Not Found | Not Found | High | High |
| | | | | | High | High | High | High | Not Found | Not Found | Not Found | Not Found | Not Found | Not Found |
| 65316 | 38807 | 30149 | 47437 | 49265 | High | High | High | High | High | High | High | High | High | High |
| 59765 | 96616 | 27527 | 116134 | 44771 | High | High | High | High | High | High | High | High | High | High |
| 75491 | 62538 | 58761 | 60303 | 78839 | High | High | High | High | High | High | High | High | High | High |
| 109407 | 88561 | 70830 | 99149 | 113251 | High | High | High | High | High | High | High | High | High | High |
| 91615 | 67939 | 51418 | 71277 | 72034 | High | High | High | High | High | High | High | High | High | High |
| 79502 | 111977 | 92003 | 102545 | 116908 | High | High | High | High | High | High | High | High | High | High |
| 94235 | 49943 | 44470 | 60837 | 74904 | High | High | High | High | High | High | High | High | High | High |
| 114869 | 77908 | 74572 | 84318 | 112273 | High | High | High | High | High | High | High | High | High | High |

| | | | | | | | | | | | | | | |
|---|---|---|---|---|---|---|---|---|---|---|---|---|---|---|
| 94997 | 60973 | 56351 | 108494 | 100584 | High | High | High | High | High | High | High | High | High | High |
| | | | | | Not Found | Not Found | Not Found | Not Found | Not Found | Not Found | Not Found | Not Found | Not Found | Not Found |
| 103820 | 51787 | 46091 | 80219 | 83943 | High | High | High | High | High | High | High | High | High | High |
| 1490 | 481 | 375 | 594 | 457 | High | High | High | High | High | High | High | High | High | High |
| | 6555 | 3740 | 4714 | 4345 | Not Found | Not Found | Not Found | Not Found | Not Found | Not Found | Not Found | Not Found | High | High |
| 130406 | 72147 | 58916 | 102341 | 103665 | High | High | High | High | High | High | High | High | High | High |
| 7066 | 9011 | 8857 | 11318 | 14118 | Not Found | Not Found | Not Found | Not Found | High | High | High | High | High | High |
| 61234 | 48604 | 44904 | 53337 | 66832 | High | High | High | High | High | High | High | High | High | High |
| 103744 | 55864 | 47171 | 88527 | 81289 | High | High | High | High | High | High | High | High | High | High |
| 82024 | 72850 | 58647 | 95101 | 93381 | High | High | High | High | High | High | High | High | High | High |
| 84305 | 65099 | 56986 | 75255 | 79924 | High | High | High | High | High | High | High | High | High | High |
| 114904 | 86130 | 81089 | 101242 | 134212 | High | High | High | High | High | High | High | High | High | High |
| 34139 | 88883 | 41750 | 93632 | 66121 | High | High | High | High | High | High | High | High | High | High |
| | 706 | 573 | 866 | 814 | Not Found | Not Found | Not Found | Not Found | Not Found | Not Found | Not Found | Not Found | High | High |
| 70043 | 56856 | 46161 | 62555 | 63424 | High | High | High | High | High | High | High | High | High | High |
| 922 | | | | | High | High | High | High | High | High | High | High | High | High |
| 98642 | 71283 | 56139 | 88380 | 76145 | High | High | High | High | High | High | High | High | High | High |
| 26248 | 20898 | 19153 | 35012 | 27581 | High | High | High | High | High | High | High | High | High | High |
| 90143 | 71997 | 54500 | 83328 | 78068 | High | High | High | High | High | High | High | High | High | High |
| 72717 | 48351 | 41161 | 86275 | 64207 | High | High | High | High | High | High | High | High | High | High |
| 99245 | 65876 | 53593 | 76837 | 86623 | High | High | High | High | High | High | High | High | High | High |
| 75367 | 55929 | 49070 | 65296 | 79417 | High | High | High | High | High | High | High | High | High | High |
| 69738 | 51048 | 41597 | 71704 | 70184 | High | High | High | High | High | High | High | High | High | High |
| 57955 | 30551 | 33619 | 34077 | 43774 | High | High | High | High | High | High | High | High | High | High |
| 39617 | 60066 | 51086 | 63462 | 81773 | High | High | High | High | High | High | High | High | High | High |
| 24456 | 33635 | 27328 | 39788 | 40630 | High | High | High | High | High | High | High | High | High | High |
| 88254 | 53169 | 39643 | 100488 | 78324 | High | High | High | High | High | High | High | High | High | High |
| 139961 | 127849 | 93626 | 153015 | 155305 | High | High | High | High | High | High | High | High | High | High |
| 85380 | 34666 | 27611 | 40241 | 43258 | High | High | High | High | High | High | High | High | High | High |
| 58992 | 47761 | 32819 | 91968 | 78269 | High | High | High | High | High | High | High | High | High | High |
| 90849 | 52957 | 34233 | 78485 | 72535 | High | High | High | High | High | High | High | High | High | High |

| | | | | | | | | | | | | | | |
|---|---|---|---|---|---|---|---|---|---|---|---|---|---|---|
| 68752 | 70748 | 47160 | 82287 | 74352 | High | High | High | High | High | High | High | High | High | High |
| 66087 | 78580 | 48297 | 96877 | 76902 | High | High | High | High | High | High | High | High | High | High |
| 38174 | 64469 | 51650 | 79146 | 77948 | High | High | High | High | High | High | High | High | High | High |
| 61894 | 80407 | 73998 | 95406 | 122502 | High | High | High | High | High | High | High | High | High | High |
| 41588 | 47713 | 37308 | 63441 | 51932 | High | High | High | High | High | High | High | High | High | High |
| 70516 | 45012 | 45416 | 66039 | 53349 | High | High | High | High | High | High | High | High | High | High |
| 2305 | | | | | High | High | High | High | High | High | High | High | Not Founc | Not Founc |
| 55330 | 53323 | 42217 | 61746 | 66844 | High | High | High | High | High | High | High | High | High | High |
| 26792 | 5321 | 4508 | 4838 | 5114 | High | High | High | High | High | High | High | High | High | High |
| 87061 | 47286 | 46567 | 78437 | 75446 | High | High | High | High | High | High | High | High | High | High |
| 63408 | 62529 | 54178 | 76750 | 92136 | High | High | High | High | High | High | High | High | High | High |
| 68735 | 50993 | 46144 | 56162 | 71551 | High | High | High | High | High | High | High | High | High | High |
| 100791 | 59998 | 46843 | 77727 | 92487 | High | High | High | High | High | High | High | High | High | High |
| 89889 | 49235 | 43264 | 62218 | 62648 | High | High | High | High | High | High | High | High | High | High |
| 74632 | 50997 | 43990 | 77442 | 74226 | High | High | High | High | High | High | High | High | High | High |
| 116377 | 47737 | 42867 | 50888 | 64187 | High | High | High | High | High | High | High | High | High | High |
| 93825 | 93358 | 78221 | 128778 | 127852 | High | High | High | High | High | High | High | High | High | High |
| 92065 | 58198 | 41593 | 79645 | 68424 | High | High | High | High | High | High | High | High | High | High |
| 104967 | 70156 | 54660 | 91837 | 102907 | High | High | High | High | High | High | High | High | High | High |
| | 3379 | 2404 | 7652 | 4675 | High | High | High | High | Not Founc | Not Founc | Not Founc | Not Founc | High | High |
| 66165 | 43292 | 40204 | 73159 | 70919 | High | High | High | High | High | High | High | High | High | High |
| 69327 | 48854 | 34831 | 82185 | 56930 | High | High | High | High | High | High | High | High | High | High |
| | | | | | High | High | High | High | High | High | High | High | High | High |
| 72907 | 55860 | 46169 | 79206 | 84308 | High | High | High | High | High | High | High | High | High | High |
| 72929 | 67292 | 53414 | 106080 | 89493 | High | High | High | High | High | High | High | High | High | High |
| 26812 | 27992 | 20183 | 29162 | 26946 | High | High | High | High | High | High | High | High | High | High |
| 59535 | 37028 | 27250 | 43060 | 43598 | High | High | High | High | High | High | High | High | High | High |
| 85321 | 36912 | 36297 | 59903 | 66059 | High | High | High | High | High | High | High | High | High | High |
| 24604 | 19405 | 15416 | 20755 | 22923 | High | High | High | High | High | High | High | High | High | High |
| 77202 | 63093 | 37372 | 98280 | 67962 | High | High | High | High | High | High | High | High | High | High |
| 64910 | 59663 | 50932 | 77211 | 87075 | High | High | High | High | High | High | High | High | High | High |

| | | | | | | | | | | | | | | |
|---|---|---|---|---|---|---|---|---|---|---|---|---|---|---|
| 87167 | 67437 | 31438 | 74314 | 64069 | High | High | High | High | High | High | High | High | High | High |
| 101061 | 51873 | 41039 | 85154 | 80815 | High | High | High | High | High | High | High | High | High | High |
| 73199 | 48119 | 45541 | 56775 | 72811 | High | High | High | High | High | High | High | High | High | High |
| 101735 | 67299 | 61190 | 73962 | 88105 | High | High | High | High | High | High | High | High | High | High |
| 82458 | 25264 | 26676 | 31584 | 41654 | High | High | High | High | High | High | High | High | High | High |
| 67192 | 58637 | 48811 | 111035 | 79565 | High | High | High | High | High | High | High | High | High | High |
| 69659 | 53170 | 46503 | 53119 | 67948 | High | High | High | High | High | High | High | High | High | High |
| 70131 | 66109 | 51298 | 69579 | 69949 | High | High | High | High | High | High | High | High | High | High |
| 63090 | 33753 | 22500 | 51501 | 48029 | High | High | High | High | High | High | High | High | High | High |
| 14511 | 10638 | 9354 | 11794 | 16509 | High | High | High | High | High | High | High | High | High | High |
| 78062 | 51799 | 34080 | 77605 | 64619 | High | High | High | High | High | High | High | High | High | High |
| 78359 | 61337 | 49170 | 77766 | 74578 | High | High | High | High | High | High | High | High | High | High |
| 76217 | 52164 | 42825 | 91775 | 72609 | High | High | High | High | High | High | High | High | High | High |
| 60405 | 61959 | 60364 | 78689 | 84445 | High | High | High | High | High | High | High | High | High | High |
| 68222 | 65934 | 46474 | 72097 | 67054 | High | High | High | High | High | High | High | High | High | High |
| 59251 | 58199 | 46686 | 71090 | 60583 | High | High | High | High | High | High | High | High | High | High |
| 59557 | 65543 | 51463 | 100721 | 104614 | High | High | High | High | High | High | High | High | High | High |
| 83083 | 34682 | 36661 | 44441 | 60674 | High | High | High | High | High | High | High | High | High | High |
| 44305 | 51238 | 35261 | 50594 | 51487 | High | High | High | High | High | High | High | High | High | High |
| 75302 | 50326 | 34186 | 81932 | 75382 | High | High | High | High | High | High | High | High | High | High |
| 63895 | 45649 | 36547 | 52397 | 48205 | High | High | High | High | High | High | High | High | High | High |
| 76955 | 59199 | 46528 | 82110 | 72022 | High | High | High | High | High | High | High | High | High | High |
| 74454 | 57911 | 45080 | 70637 | 64455 | High | High | High | High | High | High | High | High | High | High |
| 54961 | 48997 | 36739 | 44391 | 48158 | High | High | High | High | High | High | High | High | High | High |
| 42750 | 20641 | 19643 | 18879 | 27641 | High | High | High | High | High | High | High | High | High | High |
| 59572 | 35087 | 29520 | 45613 | 43467 | High | High | High | High | High | High | High | High | High | High |
| 61815 | 38280 | 27179 | 63769 | 58221 | High | High | High | High | High | High | High | High | High | High |
| 93837 | 108435 | 59414 | 135036 | 110988 | High | High | High | High | High | High | High | High | High | High |
| 86913 | 74813 | 48974 | 146241 | 92734 | High | High | High | High | High | High | High | High | High | High |
| 46325 | 60680 | 48729 | 108331 | 71481 | High | High | High | High | High | High | High | High | High | High |
| 72348 | 49853 | 45166 | 70549 | 71455 | High | High | High | High | High | High | High | High | High | High |

| | | | | | | | | | | | | | | |
|---|---|---|---|---|---|---|---|---|---|---|---|---|---|---|
| 63340 | 76424 | 80810 | 87582 | 106786 | High | High | High | High | High | High | High | High | High | High |
| 71414 | 54050 | 38008 | 76825 | 64122 | High | High | High | High | High | High | High | High | High | High |
| 68300 | 38205 | 33398 | 42409 | 47520 | High | High | High | High | High | High | High | High | High | High |
| 66587 | 45881 | 39953 | 50719 | 58075 | High | High | High | High | High | High | High | High | High | High |
| 98588 | 66431 | 49629 | 86786 | 85169 | High | High | High | High | High | High | High | High | High | High |
| 65311 | 71361 | 47534 | 129859 | 92474 | High | High | High | High | High | High | High | High | High | High |
| 89769 | 59564 | 48154 | 72134 | 73387 | High | High | High | High | High | High | High | High | High | High |
| 55104 | 50764 | 36730 | 60109 | 56570 | High | High | High | High | High | High | High | High | High | High |
| 26285 | 37236 | 28173 | 60176 | 53784 | High | High | High | High | High | High | High | High | High | High |
| 85955 | 66006 | 51745 | 60913 | 71156 | High | High | High | High | High | High | High | High | High | High |
| 76225 | 39028 | 28455 | 60391 | 50075 | High | High | High | High | High | High | High | High | High | High |
| 62382 | 58901 | 34626 | 81465 | 71312 | High | High | High | High | High | High | High | High | High | High |
| 51047 | 79920 | 70025 | 110815 | 99672 | High | High | High | High | High | High | High | High | High | High |
| 42581 | 57600 | 45749 | 87576 | 82166 | High | High | High | High | High | High | High | High | High | High |
| 69300 | 30802 | 31245 | 45589 | 52640 | High | High | High | High | High | High | High | High | High | High |
| 50478 | 77559 | 61529 | 75764 | 69827 | High | High | High | High | High | High | High | High | High | High |
| 64659 | 38453 | 24908 | 52366 | 45605 | High | High | High | High | High | High | High | High | High | High |
| 15042 | 26766 | 23516 | 35830 | 34406 | High | High | High | High | High | High | High | High | High | High |
| 48836 | 30501 | 21043 | 38171 | 34823 | High | High | High | High | High | High | High | High | High | High |
| 53118 | 55283 | 39485 | 75799 | 68472 | High | High | High | High | High | High | High | High | High | High |
| 63414 | 49997 | 46069 | 87531 | 72616 | High | High | High | High | High | High | High | High | High | High |
| 51297 | 50199 | 30246 | 62682 | 46928 | High | High | High | High | High | High | High | High | High | High |
| 71362 | 78461 | 64888 | 72540 | 85178 | High | High | High | High | High | High | High | High | High | High |
| 5527 | | | | | Not Founc | Not Founc | Not Founc | Not Founc | High | High | High | High | Not Founc | Not Founc |
| 68844 | 39043 | 30978 | 67528 | 57973 | High | High | High | High | High | High | High | High | High | High |
| 41918 | 55244 | 37181 | 63400 | 54268 | High | High | High | High | High | High | High | High | High | High |
| 42853 | 48831 | 36757 | 53249 | 51847 | High | High | High | High | High | High | High | High | High | High |
| 42878 | 47290 | 44989 | 51420 | 61268 | High | High | High | High | High | High | High | High | High | High |
| 64489 | 51480 | 31891 | 71294 | 53924 | High | High | High | High | High | High | High | High | High | High |
| 84083 | 47208 | 40030 | 56360 | 71993 | High | High | High | High | High | High | High | High | High | High |
| 11953 | 4606 | 3723 | 4354 | 4392 | High | High | High | High | High | High | High | High | High | High |

| | | | | | | | | | | | | | | |
|---|---|---|---|---|---|---|---|---|---|---|---|---|---|---|
| 56288 | 46408 | 35635 | 59613 | 47889 | High | High | High | High | High | High | High | High | High | High |
| 66858 | 48416 | 38857 | 59721 | 71754 | High | High | High | High | High | High | High | High | High | High |
| 77999 | 56695 | 35697 | 91855 | 60839 | High | High | High | High | High | High | High | High | High | High |
| 84902 | 59714 | 54077 | 71961 | 105729 | High | High | High | High | High | High | High | High | High | High |
| 68683 | 74729 | 57212 | 90750 | 88234 | High | High | High | High | High | High | High | High | High | High |
| 41248 | 23284 | 19882 | 26695 | 29770 | High | High | High | High | High | High | High | High | High | High |
| 71165 | 58043 | 51789 | 64484 | 76301 | High | High | High | High | High | High | High | High | High | High |
| 42846 | 23545 | 16446 | 34252 | 32557 | High | High | High | High | High | High | High | High | High | High |
| 65652 | 91655 | 50419 | 98363 | 70011 | High | High | High | High | High | High | High | High | High | High |
| 49438 | 62810 | 39039 | 73638 | 66313 | High | High | High | High | High | High | High | High | High | High |
| 31227 | 59171 | 37679 | 70628 | 58998 | High | High | High | High | High | High | High | High | High | High |
| 60582 | 66506 | 51944 | 77355 | 71703 | High | High | High | High | High | High | High | High | High | High |
| 22617 | 27977 | 22701 | 31995 | 33996 | High | High | High | High | High | High | High | High | High | High |
| 88825 | 60589 | 57969 | 77063 | 103961 | High | High | High | High | High | High | High | High | High | High |
| 79298 | 58843 | 48795 | 73769 | 66307 | High | High | High | High | High | High | High | High | High | High |
| 58597 | 58256 | 58674 | 69404 | 84774 | High | High | High | High | High | High | High | High | High | High |
| 42583 | 39281 | 29558 | 70110 | 47468 | High | High | High | High | High | High | High | High | High | High |
| 76926 | 27855 | 25955 | 31659 | 36707 | High | High | High | High | High | High | High | High | High | High |
| 64502 | 74125 | 72221 | 127431 | 96632 | High | High | High | High | High | High | High | High | High | High |
| 70848 | 40685 | 47403 | 55048 | 77679 | High | High | High | High | High | High | High | High | High | High |
| 65492 | 74901 | 59636 | 79993 | 84403 | High | High | High | High | High | High | High | High | High | High |
| 98149 | 51454 | 41961 | 46711 | 58249 | High | High | High | High | High | High | High | High | High | High |
| 81792 | 56721 | 34041 | 89627 | 63187 | High | High | High | High | High | High | High | High | High | High |
| 60694 | 37625 | 31619 | 41322 | 47440 | High | High | High | High | High | High | High | High | High | High |
| 47997 | 55387 | 47295 | 67973 | 66332 | High | High | High | High | High | High | High | High | High | High |
| 57733 | 62547 | 47419 | 87422 | 73883 | High | High | High | High | High | High | High | High | High | High |
| 54826 | 45911 | 48520 | 61718 | 67909 | High | High | High | High | High | High | High | High | High | High |
| 54940 | 50851 | 48556 | 112613 | 82147 | High | High | High | High | High | High | High | High | High | High |
| 36869 | 40592 | 34400 | 45275 | 51851 | High | High | High | High | High | High | High | High | High | High |
| 20793 | 15074 | 12206 | 15464 | 17118 | High | High | High | High | High | High | High | High | High | High |
| 46312 | 38185 | 26005 | 35268 | 32353 | High | High | High | High | High | High | High | High | High | High |

| | | | | | | | | | | | | | | |
|---|---|---|---|---|---|---|---|---|---|---|---|---|---|---|
| 51750 | 30629 | 28754 | 26220 | 37495 | High | High | High | High | High | High | High | High | High | High |
| 62221 | 60841 | 54542 | 63224 | 76161 | High | High | High | High | High | High | High | High | High | High |
| 71760 | 61889 | 55202 | 60159 | 73736 | High | High | High | High | High | High | High | High | High | High |
| 63888 | 43430 | 36099 | 61398 | 60312 | High | High | High | High | High | High | High | High | High | High |
| 57737 | 33030 | 29836 | 38004 | 45437 | High | High | High | High | High | High | High | High | High | High |
| 43389 | 23693 | 23940 | 21852 | 26967 | High | High | High | High | High | High | High | High | High | High |
| 46794 | 50920 | 30089 | 65436 | 56180 | High | High | High | High | High | High | High | High | High | High |
| 64123 | 33375 | 30697 | 35472 | 47978 | High | High | High | High | High | High | High | High | High | High |
| 3337 | | | | | Not Founc | Not Founc | Not Founc | Not Founc | High | High | High | High | Not Founc | Not Founc |
| 55962 | 56559 | 37863 | 75427 | 61590 | High | High | High | High | High | High | High | High | High | High |
| 23785 | 20812 | 24411 | 24365 | 30237 | High | High | High | High | High | High | High | High | High | High |
| 66035 | 68631 | 42946 | 96949 | 72979 | High | High | High | High | High | High | High | High | High | High |
| 59235 | 52135 | 42963 | 68053 | 67217 | High | High | High | High | High | High | High | High | High | High |
| 16025 | 12221 | 9118 | 17704 | 14770 | High | High | High | High | High | High | High | High | High | High |
| 82163 | 29244 | 24768 | 42969 | 47949 | High | High | High | High | High | High | High | High | High | High |
| 52779 | 50518 | 39202 | 68374 | 57252 | High | High | High | High | High | High | High | High | High | High |
| 81147 | 83180 | 71124 | 124499 | 116296 | High | High | High | High | High | High | High | High | High | High |
| 37808 | 52086 | 40316 | 84210 | 67560 | High | High | High | High | High | High | High | High | High | High |
| 46847 | 26463 | 20639 | 20980 | 24969 | High | High | High | High | High | High | High | High | High | High |
| 91346 | 63514 | 64115 | 76174 | 115287 | High | High | High | High | High | High | High | High | High | High |
| 52717 | 24930 | 17528 | 32662 | 28016 | High | High | High | High | High | High | High | High | High | High |
| 54419 | 47714 | 40002 | 59562 | 57776 | High | High | High | High | High | High | High | High | High | High |
| 55906 | 48953 | 42752 | 59509 | 61140 | High | High | High | High | High | High | High | High | High | High |
| 73483 | 38153 | 31774 | 50318 | 58526 | High | High | High | High | High | High | High | High | High | High |
| 15502 | 3841 | 2837 | 6002 | 5195 | High | High | High | High | High | High | High | High | High | High |
| 45640 | 35804 | 27026 | 46295 | 47643 | High | High | High | High | High | High | High | High | High | High |
| 61901 | 35784 | 24251 | 45338 | 40570 | High | High | High | High | High | High | High | High | High | High |
| 63039 | 48073 | 36731 | 65844 | 61525 | High | High | High | High | High | High | High | High | High | High |
| 69920 | 41207 | 30948 | 47223 | 47333 | High | High | High | High | High | High | High | High | High | High |
| 40033 | 70281 | 56404 | 94954 | 92435 | High | High | High | High | High | High | High | High | High | High |
| 65567 | 31944 | 30052 | 30908 | 43621 | High | High | High | High | High | High | High | High | High | High |

| | | | | | | | | | | | | | | |
|---|---|---|---|---|---|---|---|---|---|---|---|---|---|---|
| 61024 | 34406 | 24812 | 38521 | 38909 | High | High | High | High | High | High | High | High | High | High |
| 40920 | 31061 | 26298 | 37366 | 39436 | High | High | High | High | High | High | High | High | High | High |
| 45661 | 57167 | 52517 | 60079 | 72163 | High | High | High | High | High | High | High | High | High | High |
| 56733 | 30854 | 25196 | 35194 | 32280 | High | High | High | High | High | High | High | High | High | High |
| 54918 | 33131 | 25399 | 36027 | 39283 | High | High | High | High | High | High | High | High | High | High |
| 49551 | 40723 | 40320 | 46034 | 59078 | High | High | High | High | High | High | High | High | High | High |
| 68519 | 30706 | 22531 | 47261 | 45694 | High | High | High | High | High | High | High | High | High | High |
| 59703 | 59501 | 36525 | 52367 | 49999 | High | High | High | High | High | High | High | High | High | High |
| 78961 | 60525 | 49964 | 88386 | 92878 | High | High | High | High | High | High | High | High | High | High |
| 40236 | 39542 | 29510 | 55024 | 45608 | High | High | High | High | High | High | High | High | High | High |
| 57065 | 36231 | 31255 | 46864 | 58781 | High | High | High | High | High | High | High | High | High | High |
| 68462 | 41109 | 33600 | 52302 | 52965 | High | High | High | High | High | High | High | High | High | High |
| 3369 | 2952 | 2285 | 2958 | 2817 | High | High | High | High | High | High | High | High | High | High |
| 61952 | 49896 | 33847 | 79150 | 53338 | High | High | High | High | High | High | High | High | High | High |
| 44680 | 39746 | 23794 | 44843 | 35259 | High | High | High | High | High | High | High | High | High | High |
| 47458 | 41900 | 29695 | 58791 | 51477 | High | High | High | High | High | High | High | High | High | High |
| 71538 | 34731 | 27660 | 43642 | 42053 | High | High | High | High | High | High | High | High | High | High |
| 60752 | 46177 | 42212 | 53988 | 68573 | High | High | High | High | High | High | High | High | High | High |
| 92741 | 39686 | 35023 | 53005 | 60141 | High | High | High | High | High | High | High | High | High | High |
| 44767 | 36221 | 31835 | 47829 | 57433 | High | High | High | High | High | High | High | High | High | High |
| 67326 | 56440 | 56197 | 92851 | 86613 | High | High | High | High | High | High | High | High | High | High |
| 69239 | 22546 | 35163 | 47026 | 90905 | High | High | High | High | High | High | High | High | High | High |
| 32662 | 28862 | 18438 | 28123 | 26737 | High | High | High | High | High | High | High | High | High | High |
| 47956 | 36872 | 26747 | 50383 | 45539 | High | High | High | High | High | High | High | High | High | High |
| 33377 | 22736 | 24736 | 20971 | 30112 | High | High | High | High | High | High | High | High | High | High |
| 41332 | 48823 | 37173 | 84763 | 60902 | High | High | High | High | High | High | High | High | High | High |
| 67236 | 47827 | 37340 | 81935 | 70868 | High | High | High | High | High | High | High | High | High | High |
| 58951 | 43660 | 30401 | 60382 | 57479 | High | High | High | High | High | High | High | High | High | High |
| 58811 | 55425 | 48127 | 57336 | 76786 | High | High | High | High | High | High | High | High | High | High |
| 80389 | 47754 | 30454 | 47847 | 64810 | High | High | High | High | High | High | High | High | High | High |
| 57398 | 58729 | 44651 | 88401 | 68095 | High | High | High | High | High | High | High | High | High | High |

| | | | | | | | | | | | | | | |
|---|---|---|---|---|---|---|---|---|---|---|---|---|---|---|
| 34131 | 48139 | 39099 | 64785 | 64475 | High | High | High | High | High | High | High | High | High | High |
| 42015 | 44414 | 34589 | 61662 | 56303 | High | High | High | High | High | High | High | High | High | High |
| 49722 | 50472 | 48331 | 67036 | 76240 | High | High | High | High | High | High | High | High | High | High |
| 68162 | 52440 | 50500 | 49175 | 73116 | High | High | High | High | High | High | High | High | High | High |
| 1320 | 3910 | 3499 | 3774 | 5261 | High | High | High | High | High | High | High | High | High | High |
| 69563 | 32014 | 21689 | 46875 | 35389 | High | High | High | High | High | High | High | High | High | High |
| 66166 | 35062 | 27714 | 42960 | 47088 | High | High | High | High | High | High | High | High | High | High |
| 881 | 590 | 495 | 775 | 889 | High | High | High | High | High | High | High | High | High | High |
| 72292 | 50648 | 44533 | 54845 | 59477 | High | High | High | High | High | High | High | High | High | High |
| 31664 | 31819 | 21300 | 44559 | 35665 | High | High | High | High | High | High | High | High | High | High |
| 44055 | 62813 | 37132 | 58395 | 51073 | High | High | High | High | High | High | High | High | High | High |
| 57171 | 32549 | 31768 | 32734 | 48279 | High | High | High | High | High | High | High | High | High | High |
| 73316 | 45819 | 33126 | 67885 | 53609 | High | High | High | High | High | High | High | High | High | High |
| 38616 | 24409 | 25713 | 29612 | 35956 | High | High | High | High | High | High | High | High | High | High |
| 40849 | 60030 | 48275 | 68935 | 74705 | High | High | High | High | High | High | High | High | High | High |
| 49883 | 10470 | 11563 | 11812 | 17524 | High | High | High | High | High | High | High | High | High | High |
| 71371 | 37129 | 28162 | 68650 | 56214 | High | High | High | High | High | High | High | High | High | High |
| 64762 | 46257 | 43519 | 51825 | 62042 | High | High | High | High | High | High | High | High | High | High |
| 45289 | 45970 | 33041 | 68450 | 54742 | High | High | High | High | High | High | High | High | High | High |
| 51298 | 32930 | 29502 | 36769 | 46665 | High | High | High | High | High | High | High | High | High | High |
| 50361 | 38934 | 19794 | 36963 | 29323 | High | High | High | High | High | High | High | High | High | High |
| 81762 | 32417 | 26473 | 37358 | 42537 | High | High | High | High | High | High | High | High | High | High |
| 52172 | 37108 | 28946 | 57614 | 60092 | High | High | High | High | High | High | High | High | High | High |
| 52750 | 46529 | 36943 | 69119 | 59809 | High | High | High | High | High | High | High | High | High | High |
| 50786 | 37035 | 32702 | 60263 | 47886 | High | High | High | High | High | High | High | High | High | High |
| 86980 | 38866 | 32926 | 45256 | 74838 | High | High | High | High | High | High | High | High | High | High |
| 54155 | 18300 | 12988 | 23576 | 29707 | High | High | High | High | High | High | High | High | High | High |
| 73226 | 43771 | 33295 | 63003 | 63296 | High | High | High | High | High | High | High | High | High | High |
| 66659 | 30792 | 30292 | 34103 | 45969 | High | High | High | High | High | High | High | High | High | High |
| 22091 | 17275 | 15263 | 17872 | 20278 | High | High | High | High | High | High | High | High | High | High |
| 73973 | 32484 | 26525 | 44281 | 40712 | High | High | High | High | High | High | High | High | High | High |

| | | | | | | | | | | | | | | |
|---|---|---|---|---|---|---|---|---|---|---|---|---|---|---|
| 54827 | 43266 | 29394 | 40487 | 41931 | High | High | High | High | High | High | High | High | High | High |
| 58006 | 48182 | 30028 | 60122 | 47230 | High | High | High | High | High | High | High | High | High | High |
| 78780 | 58778 | 40736 | 93115 | 90041 | High | High | High | High | High | High | High | High | High | High |
| 57904 | 38091 | 33742 | 42548 | 51855 | High | High | High | High | High | High | High | High | High | High |
| 40247 | 30182 | 25280 | 32117 | 36229 | High | High | High | High | High | High | High | High | High | High |
| 10520 | 6606 | 10564 | 8096 | 12960 | High | High | High | High | High | High | High | High | High | High |
| 38822 | 37015 | 27857 | 48891 | 38567 | High | High | High | High | High | High | High | High | High | High |
| 78258 | 36452 | 34530 | 40622 | 54467 | High | High | High | High | High | High | High | High | High | High |
| 69117 | 46216 | 35403 | 63746 | 57787 | High | High | High | High | High | High | High | High | High | High |
| 28432 | 45014 | 44990 | 39960 | 49821 | High | High | High | High | High | High | High | High | High | High |
| 72142 | 32373 | 20834 | 49895 | 38501 | High | High | High | High | High | High | High | High | High | High |
| 20660 | 34941 | 24282 | 54676 | 44696 | High | High | High | High | High | High | High | High | High | High |
| 44587 | 23324 | 15141 | 33209 | 28365 | High | High | High | High | High | High | High | High | High | High |
| 52382 | 54642 | 33852 | 82926 | 62202 | High | High | High | High | High | High | High | High | High | High |
| 52258 | 41262 | 28640 | 50650 | 43818 | High | High | High | High | High | High | High | High | High | High |
| 54312 | 23515 | 16531 | 40749 | 38162 | High | High | High | High | High | High | High | High | High | High |
| 60954 | 33982 | 21880 | 43696 | 38150 | High | High | High | High | High | High | High | High | High | High |
| 62683 | 37246 | 26495 | 48061 | 42541 | High | High | High | High | High | High | High | High | High | High |
| 22440 | 30813 | 27483 | 38728 | 41819 | High | High | High | High | High | High | High | High | High | High |
| 51077 | 35817 | 31636 | 42595 | 59081 | High | High | High | High | High | High | High | High | High | High |
| 52415 | 57572 | 43839 | 99081 | 76973 | High | High | High | High | High | High | High | High | High | High |
| 49723 | 34710 | 25766 | 44504 | 45660 | High | High | High | High | High | High | High | High | High | High |
| 7455 | | | | | High | High | High | High | High | High | High | High | Not Found | Not Found |
| 44758 | 31762 | 30058 | 36922 | 45411 | High | High | High | High | High | High | High | High | High | High |
| 48070 | 52175 | 40705 | 58230 | 63814 | High | High | High | High | High | High | High | High | High | High |
| 46371 | 32231 | 24827 | 46722 | 40825 | High | High | High | High | High | High | High | High | High | High |
| 77824 | 37250 | 30962 | 43236 | 56864 | High | High | High | High | High | High | High | High | High | High |
| 31882 | 21909 | 21131 | 24722 | 28772 | High | High | High | High | High | High | High | High | High | High |
| 59732 | 38686 | 36641 | 47858 | 60069 | High | High | High | High | High | High | High | High | High | High |
| 56184 | 43037 | 31644 | 67022 | 50914 | High | High | High | High | High | High | High | High | High | High |
| 48554 | 26962 | 27954 | 48744 | 45576 | High | High | High | High | High | High | High | High | High | High |

| | | | | | | | | | | | | | | |
|---|---|---|---|---|---|---|---|---|---|---|---|---|---|---|
| 56355 | 34279 | 30414 | 42766 | 45621 | High | High | High | High | High | High | High | High | High | High |
| 71967 | 55957 | 47362 | 73489 | 81483 | High | High | High | High | High | High | High | High | High | High |
| 26945 | 17394 | 14021 | 20846 | 23842 | High | High | High | High | High | High | High | High | High | High |
| 19383 | 39604 | 11816 | 31104 | 19704 | High | High | High | High | High | High | High | High | High | High |
| 37418 | 35714 | 30034 | 38103 | 39512 | High | High | High | High | High | High | High | High | High | High |
| 56366 | 44006 | 30139 | 52123 | 49921 | High | High | High | High | High | High | High | High | High | High |
| 44294 | 22667 | 16657 | 23807 | 23967 | High | High | High | High | High | High | High | High | High | High |
| 34110 | 30227 | 32259 | 32845 | 41229 | High | High | High | High | High | High | High | High | High | High |
| 39573 | 28468 | 19487 | 41454 | 32129 | High | High | High | High | High | High | High | High | High | High |
| 63570 | 34568 | 28138 | 29473 | 33744 | High | High | High | High | High | High | High | High | High | High |
| 46234 | 29987 | 35977 | 29816 | 56652 | High | High | High | High | High | High | High | High | High | High |
| 52458 | 18695 | 19299 | 20989 | 25327 | High | High | High | High | High | High | High | High | High | High |
| 43273 | 22401 | 17412 | 35870 | 33820 | High | High | High | High | High | High | High | High | High | High |
| 47621 | 38004 | 32429 | 43979 | 53483 | High | High | High | High | High | High | High | High | High | High |
| 37140 | 20974 | 13746 | 42356 | 35214 | High | High | High | High | High | High | High | High | High | High |
| 54095 | 67303 | 40111 | 65929 | 59579 | High | High | High | High | High | High | High | High | High | High |
| 29562 | 19822 | 17539 | 17899 | 19237 | High | High | High | High | High | High | High | High | High | High |
| 51075 | 42367 | 34507 | 65401 | 59796 | High | High | High | High | High | High | High | High | High | High |
| 50756 | 31696 | 24798 | 48521 | 43835 | High | High | High | High | High | High | High | High | High | High |
| 71037 | 27352 | 15883 | 67339 | 43714 | High | High | High | High | High | High | High | High | High | High |
| 60459 | 37023 | 21068 | 42561 | 40393 | High | High | High | High | High | High | High | High | High | High |
| 42030 | 44004 | 29662 | 63457 | 46039 | High | High | High | High | High | High | High | High | High | High |
| 42579 | 50801 | 42227 | 87605 | 66535 | High | High | High | High | High | High | High | High | High | High |
| 32795 | 32848 | 23662 | 33505 | 32950 | High | High | High | High | High | High | High | High | High | High |
| 33331 | 29118 | 18786 | 35483 | 31408 | High | High | High | High | High | High | High | High | High | High |
| 62750 | 36488 | 21211 | 76236 | 48117 | High | High | High | High | High | High | High | High | High | High |
| 45948 | 65108 | 46745 | 56694 | 52310 | High | High | High | High | High | High | High | High | High | High |
| 56722 | 47214 | 33439 | 76520 | 55292 | High | High | High | High | High | High | High | High | High | High |
| 50320 | 37624 | 29960 | 58617 | 45479 | High | High | High | High | High | High | High | High | High | High |
| 52780 | 41742 | 25104 | 69986 | 46864 | High | High | High | High | High | High | High | High | High | High |
| 29428 | 32707 | 29554 | 32507 | 46638 | High | High | High | High | High | High | High | High | High | High |

| | | | | | | | | | | | | | | |
|---|---|---|---|---|---|---|---|---|---|---|---|---|---|---|
| 35974 | 42129 | 34632 | 60980 | 52241 | High | High | High | High | High | High | High | High | High | High |
| 33426 | 33582 | 23953 | 51571 | 45572 | High | High | High | High | High | High | High | High | High | High |
| 38958 | 55397 | 27305 | 53922 | 44158 | High | High | High | High | High | High | High | High | High | High |
| 85419 | 32665 | 26319 | 39371 | 43169 | High | High | High | High | High | High | High | High | High | High |
| 46216 | 21350 | 17544 | 35797 | 21180 | High | High | High | High | High | High | High | High | High | High |
| 34191 | 50964 | 40514 | 68098 | 71558 | High | High | High | High | High | High | High | High | High | High |
| 43308 | 32648 | 29196 | 32307 | 42955 | High | High | High | High | High | High | High | High | High | High |
| 58292 | 36346 | 25974 | 60841 | 55882 | High | High | High | High | High | High | High | High | High | High |
| 29116 | 30583 | 24084 | 31914 | 37199 | High | High | High | High | High | High | High | High | High | High |
| 51295 | 20232 | 18724 | 27614 | 27740 | High | High | High | High | High | High | High | High | High | High |
| 10027 | 9866 | 7146 | 8684 | 9768 | High | High | High | High | High | High | High | High | High | High |
| 28274 | 30274 | 24838 | 36980 | 37127 | High | High | High | High | High | High | High | High | High | High |
| 48241 | 9435 | 7996 | 12930 | 12751 | High | High | High | High | High | High | High | High | High | High |
| 52995 | 28363 | 27954 | 26258 | 35418 | High | High | High | High | High | High | High | High | High | High |
| 54858 | 42395 | 34570 | 74849 | 67390 | High | High | High | High | High | High | High | High | High | High |
| 73996 | 27745 | 30130 | 30655 | 43249 | High | High | High | High | High | High | High | High | High | High |
| 61782 | 34676 | 35141 | 40842 | 61770 | High | High | High | High | High | High | High | High | High | High |
| 50830 | 55891 | 42597 | 85041 | 75840 | High | High | High | High | High | High | High | High | High | High |
| 17487 | 8748 | 5938 | 10092 | 9659 | High | High | High | High | High | High | High | High | High | High |
| 38715 | 29507 | 26079 | 39632 | 43164 | High | High | High | High | High | High | High | High | High | High |
| 73652 | 29126 | 21049 | 37683 | 36824 | High | High | High | High | High | High | High | High | High | High |
| 66647 | 66810 | 48728 | 71016 | 77815 | High | High | High | High | High | High | High | High | High | High |
| 20716 | 22730 | 23805 | 20833 | 23972 | High | High | High | High | High | High | High | High | High | High |
| 36545 | 40677 | 31074 | 39159 | 45136 | High | High | High | High | High | High | High | High | High | High |
| 76959 | 36296 | 46310 | 39439 | 81537 | High | High | High | High | High | High | High | High | High | High |
| 38470 | 53422 | 42630 | 63939 | 66803 | High | High | High | High | High | High | High | High | High | High |
| 49067 | 36508 | 23118 | 47989 | 50279 | High | High | High | High | High | High | High | High | High | High |
| 20476 | 27329 | 18985 | 47634 | 32237 | High | High | High | High | High | High | High | High | High | High |
| 60862 | 15794 | 10121 | 42827 | 30006 | High | High | High | High | High | High | High | High | High | High |
| 53095 | 34406 | 23940 | 51100 | 44561 | High | High | High | High | High | High | High | High | High | High |
| 31669 | 24243 | 18542 | 25895 | 29632 | High | High | High | High | High | High | High | High | High | High |

| | | | | | | | | | | | | | | |
|---|---|---|---|---|---|---|---|---|---|---|---|---|---|---|
| 37934 | 32404 | 29665 | 37050 | 40890 | High | High | High | High | High | High | High | High | High | High |
| 73041 | 61767 | 54083 | 79895 | 62682 | High | High | High | High | High | High | High | High | High | High |
| 38791 | 45897 | 38670 | 46363 | 59950 | High | High | High | High | High | High | High | High | High | High |
| 59160 | 40940 | 27261 | 52185 | 52520 | High | High | High | High | High | High | High | High | High | High |
| 53273 | 32723 | 30449 | 35772 | 42686 | High | High | High | High | High | High | High | High | High | High |
| 49589 | 33324 | 20241 | 48019 | 42168 | High | High | High | High | High | High | High | High | High | High |
| 26602 | 32980 | 27419 | 37767 | 41417 | High | High | High | High | High | High | High | High | High | High |
| 46204 | 41628 | 22424 | 76624 | 53171 | High | High | High | High | High | High | High | High | High | High |
| 32445 | 25799 | 24879 | 29030 | 35952 | High | High | High | High | High | High | High | High | High | High |
| 67587 | 31182 | 25616 | 41177 | 42487 | High | High | High | High | High | High | High | High | High | High |
| 33076 | 21986 | 16098 | 26319 | 24448 | High | High | High | High | High | High | High | High | High | High |
| 47180 | 48384 | 39907 | 61993 | 70210 | High | High | High | High | High | High | High | High | High | High |
| 46271 | 36609 | 25270 | 41295 | 36738 | High | High | High | High | High | High | High | High | High | High |
| 41066 | 32498 | 20723 | 54138 | 43742 | High | High | High | High | High | High | High | High | High | High |
| 44012 | 32170 | 25833 | 52298 | 43879 | High | High | High | High | High | High | High | High | High | High |
| 37687 | 12940 | 15234 | 42835 | 31539 | High | High | High | High | High | High | High | High | High | High |
| 34558 | 15729 | 14426 | 20968 | 20038 | High | High | High | High | High | High | High | High | High | High |
| 25547 | 38056 | 31841 | 51067 | 50602 | High | High | High | High | High | High | High | High | High | High |
| | 1786 | 1421 | 2566 | 2920 | Not Founc | Not Founc | Not Founc | Not Founc | Not Founc | Not Founc | Not Founc | Not Founc | High | High |
| 63989 | 34283 | 31027 | 34030 | 43453 | High | High | High | High | High | High | High | High | High | High |
| 41229 | 29609 | 22522 | 48822 | 39425 | High | High | High | High | High | High | High | High | High | High |
| 28970 | 46526 | 35139 | 60990 | 52895 | High | High | High | High | High | High | High | High | High | High |
| 39260 | 31896 | 23238 | 41974 | 40701 | High | High | High | High | High | High | High | High | High | High |
| 15933 | 24362 | 19054 | 25448 | 24052 | High | High | High | High | High | High | High | High | High | High |
| 10383 | 6013 | 4457 | 5577 | 5883 | High | High | High | High | High | High | High | High | High | High |
| 60558 | 48642 | 39075 | 54456 | 66715 | High | High | High | High | High | High | High | High | High | High |
| 42867 | 31420 | 20567 | 49307 | 36138 | High | High | High | High | High | High | High | High | High | High |
| 34161 | 30336 | 18592 | 37050 | 31903 | High | High | High | High | High | High | High | High | High | High |
| 18755 | 11260 | 13091 | 26874 | 28987 | High | High | High | High | High | High | High | High | High | High |
| 62457 | 25943 | 20163 | 42234 | 38471 | High | High | High | High | High | High | High | High | High | High |
| 28741 | 45685 | 35946 | 39710 | 47656 | High | High | High | High | High | High | High | High | High | High |

| | | | | | | | | | | | | | | |
|---|---|---|---|---|---|---|---|---|---|---|---|---|---|---|
| 39948 | 30514 | 28409 | 29463 | 43425 | High | High | High | High | High | High | High | High | High | High |
| 31687 | 42217 | 35690 | 59089 | 50433 | High | High | High | High | High | High | High | High | High | High |
| 27292 | 15214 | 9709 | 36041 | 18729 | High | High | High | High | High | High | High | High | High | High |
| 52159 | 33913 | 25392 | 59849 | 49693 | High | High | High | High | High | High | High | High | High | High |
| 56167 | 30780 | 24323 | 46488 | 43148 | High | High | High | High | High | High | High | High | High | High |
| 62368 | 31580 | 28554 | 32700 | 44547 | High | High | High | High | High | High | High | High | High | High |
| 53706 | 35873 | 27544 | 43172 | 37368 | High | High | High | High | High | High | High | High | High | High |
| 19994 | 39993 | 29269 | 46814 | 48919 | High | High | High | High | High | High | High | High | High | High |
| 36272 | 47947 | 34644 | 63012 | 56450 | High | High | High | High | High | High | High | High | High | High |
| 61277 | 46143 | 39153 | 52467 | 70239 | High | High | High | High | High | High | High | High | High | High |
| 42273 | 27151 | 16844 | 34080 | 31312 | High | High | High | High | High | High | High | High | High | High |
| 44348 | 36400 | 35673 | 46862 | 51703 | High | High | High | High | High | High | High | High | High | High |
| 22354 | 22464 | 23612 | 21721 | 31583 | High | High | High | High | High | High | High | High | High | High |
| 37096 | 35373 | 25588 | 44237 | 40405 | High | High | High | High | High | High | High | High | High | High |
| 27985 | 29504 | 30047 | 34520 | 43951 | High | High | High | High | High | High | High | High | High | High |
| 33889 | 40693 | 32596 | 52613 | 52569 | High | High | High | High | High | High | High | High | High | High |
| 32412 | 26656 | 23116 | 28328 | 37972 | High | High | High | High | High | High | High | High | High | High |
| 23275 | 23606 | 13712 | 25671 | 23125 | High | High | High | High | High | High | High | High | High | High |
| 26448 | 11897 | 15259 | 33570 | 20341 | High | High | High | High | High | High | High | High | High | High |
| 38082 | 26427 | 20253 | 36788 | 33316 | High | High | High | High | High | High | High | High | High | High |
| 39791 | 28375 | 21539 | 41258 | 29699 | High | High | High | High | High | High | High | High | High | High |
| 47370 | 26991 | 26219 | 24826 | 37741 | High | High | High | High | High | High | High | High | High | High |
| 50784 | 22860 | 17473 | 30118 | 29131 | High | High | High | High | High | High | High | High | High | High |
| 19554 | 33983 | 28123 | 53180 | 46027 | High | High | High | High | High | High | High | High | High | High |
| 32095 | 21890 | 17955 | 22932 | 28320 | High | High | High | High | High | High | High | High | High | High |
| 46635 | 33312 | 27876 | 42310 | 52973 | High | High | High | High | High | High | High | High | High | High |
| 49478 | 28799 | 27828 | 31198 | 39900 | High | High | High | High | High | High | High | High | High | High |
| 48057 | 33069 | 27393 | 55136 | 45655 | High | High | High | High | High | High | High | High | High | High |
| 29684 | 43363 | 39439 | 43967 | 54890 | High | High | High | High | High | High | High | High | High | High |
| 33182 | 29028 | 22753 | 37680 | 36820 | High | High | High | High | High | High | High | High | High | High |
| 34183 | 25676 | 21392 | 34994 | 40039 | High | High | High | High | High | High | High | High | High | High |

| | | | | | | | | | | | | | | |
|---|---|---|---|---|---|---|---|---|---|---|---|---|---|---|
| 40914 | 31401 | 24373 | 43997 | 46721 | High | High | High | High | High | High | High | High | High | High |
| 24384 | 18359 | 13096 | 26575 | 20732 | High | High | High | High | High | High | High | High | High | High |
| 24712 | 16889 | 13869 | 19355 | 22694 | High | High | High | High | High | High | High | High | High | High |
| 28606 | 41845 | 29685 | 49782 | 44880 | High | High | High | High | High | High | High | High | High | High |
| 32500 | 44663 | 37985 | 49650 | 66953 | High | High | High | High | High | High | High | High | High | High |
| 23501 | 14098 | 10420 | 17286 | 15489 | High | High | High | High | High | High | High | High | High | High |
| 17294 | 9334 | 11175 | 17841 | 14077 | High | High | High | High | High | High | High | High | High | High |
| 73024 | 39457 | 29390 | 60103 | 73317 | High | High | High | High | High | High | High | High | High | High |
| 45775 | 42089 | 29975 | 55990 | 47103 | High | High | High | High | High | High | High | High | High | High |
| 27760 | 23872 | 19004 | 28283 | 24698 | High | High | High | High | High | High | High | High | High | High |
| 43484 | 25648 | 21611 | 34188 | 40762 | High | High | High | High | High | High | High | High | High | High |
| 25347 | 27297 | 16122 | 29791 | 23709 | High | High | High | High | High | High | High | High | High | High |
| 26756 | 17241 | 14734 | 21466 | 23111 | High | High | High | High | High | High | High | High | High | High |
| 33680 | 26443 | 19683 | 36114 | 33894 | High | High | High | High | High | High | High | High | High | High |
| 32433 | 26956 | 29406 | 28051 | 38077 | High | High | High | High | High | High | High | High | High | High |
| 37914 | 37550 | 29926 | 58264 | 50723 | High | High | High | High | High | High | High | High | High | High |
| 42682 | 46423 | 37152 | 70318 | 54336 | High | High | High | High | High | High | High | High | High | High |
| 28690 | 28496 | 15073 | 40974 | 27305 | High | High | High | High | High | High | High | High | High | High |
| 33526 | 36364 | 34662 | 49160 | 47970 | High | High | High | High | High | High | High | High | High | High |
| 34695 | 25966 | 18927 | 50046 | 29269 | High | High | High | High | High | High | High | High | High | High |
| 78402 | 49244 | 44105 | 43146 | 58854 | High | High | High | High | High | High | High | High | High | High |
| 47961 | 42364 | 40220 | 48777 | 60225 | High | High | High | High | High | High | High | High | High | High |
| 40151 | 17016 | 15058 | 15152 | 20627 | High | High | High | High | High | High | High | High | High | High |
| 15900 | 4425 | 2733 | 5464 | 4044 | High | High | High | High | High | High | High | High | High | High |
| 49561 | 34944 | 29230 | 65091 | 57891 | High | High | High | High | High | High | High | High | High | High |
| 25614 | 14061 | 13440 | 14800 | 16396 | High | High | High | High | High | High | High | High | High | High |
| 29172 | 19438 | 15260 | 24322 | 24174 | High | High | High | High | High | High | High | High | High | High |
| 32924 | 25468 | 19962 | 32879 | 31933 | High | High | High | High | High | High | High | High | High | High |
| 33655 | 61027 | 36193 | 52837 | 47640 | High | High | High | High | High | High | High | High | High | High |
| 38546 | 32893 | 23065 | 46537 | 36703 | High | High | High | High | High | High | High | High | High | High |
| 28640 | 19198 | 15707 | 35365 | 23468 | High | High | High | High | High | High | High | High | High | High |

| | | | | | | | | | | | | | | |
|---|---|---|---|---|---|---|---|---|---|---|---|---|---|---|
| 54614 | 46890 | 42211 | 46772 | 58925 | High | High | High | High | High | High | High | High | High | High |
| 21447 | 52901 | 48433 | 38615 | 45856 | High | High | High | High | High | High | High | High | High | High |
| 41973 | 29127 | 22303 | 43450 | 32387 | High | High | High | High | High | High | High | High | High | High |
| 31648 | 23293 | 20321 | 20549 | 21267 | High | High | High | High | High | High | High | High | High | High |
| 27011 | 18215 | 24840 | 63643 | 38491 | High | High | High | High | High | High | High | High | High | High |
| 47264 | 26085 | 17756 | 28294 | 31212 | High | High | High | High | High | High | High | High | High | High |
| 45866 | 32537 | 33144 | 33222 | 45940 | High | High | High | High | High | High | High | High | High | High |
| 67084 | 26479 | 27525 | 32834 | 36801 | High | High | High | High | High | High | High | High | High | High |
| 42456 | 32166 | 27956 | 46478 | 53176 | High | High | High | High | High | High | High | High | High | High |
| 32881 | 29130 | 27153 | 35607 | 43579 | High | High | High | High | High | High | High | High | High | High |
| 58905 | 30648 | 22451 | 39491 | 38610 | High | High | High | High | High | High | High | High | High | High |
| 59455 | 35411 | 28236 | 55235 | 45214 | High | High | High | High | High | High | High | High | High | High |
| 24579 | 38192 | 19613 | 44821 | 33092 | High | High | High | High | High | High | High | High | High | High |
| 39931 | 29374 | 25316 | 31249 | 31944 | High | High | High | High | High | High | High | High | High | High |
| 29542 | 30550 | 24626 | 32497 | 42159 | High | High | High | High | High | High | High | High | High | High |
| 31311 | 26113 | 19959 | 30250 | 29319 | High | High | High | High | High | High | High | High | High | High |
| 36462 | 29250 | 27861 | 69985 | 44228 | High | High | High | High | High | High | High | High | High | High |
| 16872 | 36222 | 30022 | 53638 | 55510 | High | High | High | High | High | High | High | High | High | High |
| 27637 | 27213 | 23880 | 24622 | 31493 | High | High | High | High | High | High | High | High | High | High |
| 47470 | 30274 | 24220 | 49270 | 42830 | High | High | High | High | High | High | High | High | High | High |
| 30612 | 38497 | 23094 | 56246 | 31902 | High | High | High | High | High | High | High | High | High | High |
| 42965 | 44124 | 30521 | 69245 | 58841 | High | High | High | High | High | High | High | High | High | High |
| 27066 | 20530 | 15885 | 23038 | 27510 | High | High | High | High | High | High | High | High | High | High |
| 50486 | 34251 | 34511 | 37406 | 53619 | High | High | High | High | High | High | High | High | High | High |
| 15005 | 14862 | 16965 | 20821 | 22212 | High | High | High | High | High | High | High | High | High | High |
| 49478 | 40647 | 27361 | 44030 | 42513 | High | High | High | High | High | High | High | High | High | High |
| 32806 | 13156 | 12504 | 63202 | 29984 | High | High | High | High | High | High | High | High | High | High |
| 33864 | 38145 | 25898 | 66839 | 46729 | High | High | High | High | High | High | High | High | High | High |
| 34961 | 20321 | 16141 | 29696 | 25255 | High | High | High | High | High | High | High | High | High | High |
| 55131 | 13912 | 9828 | 18030 | 16713 | High | High | High | High | High | High | High | High | High | High |
| 28743 | 30734 | 32250 | 32276 | 48280 | High | High | High | High | High | High | High | High | High | High |

| | | | | | | | | | | | | | | |
|---|---|---|---|---|---|---|---|---|---|---|---|---|---|---|
| 38874 | 29481 | 24416 | 31742 | 33572 | High | High | High | High | High | High | High | High | High | High |
| 44290 | 24165 | 19551 | 33997 | 35315 | High | High | High | High | High | High | High | High | High | High |
| 3726 | 4194 | 2886 | 5514 | 4724 | High | High | High | High | High | High | High | High | High | High |
| 48481 | 33895 | 24508 | 55265 | 45585 | High | High | High | High | High | High | High | High | High | High |
| 53491 | 15072 | 8186 | 27839 | 27313 | High | High | High | High | High | High | High | High | High | High |
| 35319 | 32288 | 23192 | 38380 | 38312 | High | High | High | High | High | High | High | High | High | High |
| 16850 | 29266 | 16609 | 35938 | 29771 | High | High | High | High | High | High | High | High | High | High |
| 41657 | 19414 | 14239 | 29387 | 23390 | High | High | High | High | High | High | High | High | High | High |
| 34075 | 39529 | 32952 | 45008 | 46481 | High | High | High | High | High | High | High | High | High | High |
| 41444 | 18942 | 13229 | 26889 | 35130 | High | High | High | High | High | High | High | High | High | High |
| 49864 | 29009 | 25251 | 39562 | 41248 | High | High | High | High | High | High | High | High | High | High |
| 43412 | 33163 | 27986 | 42563 | 42169 | High | High | High | High | High | High | High | High | High | High |
| 21991 | 17029 | 13809 | 23374 | 22904 | High | High | High | High | High | High | High | High | High | High |
| 44703 | 27943 | 21526 | 41982 | 34619 | High | High | High | High | High | High | High | High | High | High |
| 37585 | 19510 | 19293 | 25964 | 29886 | High | High | High | High | High | High | High | High | High | High |
| 48391 | 29315 | 25886 | 46045 | 50022 | High | High | High | High | High | High | High | High | High | High |
| 19397 | 10549 | 6175 | 15352 | 10138 | High | High | High | High | High | High | High | High | High | High |
| 35935 | 30307 | 18014 | 32018 | 28225 | High | High | High | High | High | High | High | High | High | High |
| 32550 | 36122 | 27712 | 42748 | 48379 | High | High | High | High | High | High | High | High | High | High |
| 36816 | 43827 | 31293 | 45813 | 43762 | High | High | High | High | High | High | High | High | High | High |
| 30577 | 24672 | 14311 | 27156 | 19105 | High | High | High | High | High | High | High | High | High | High |
| 16492 | 26192 | 11956 | 27948 | 20309 | High | High | High | High | High | High | High | High | High | High |
| 30213 | 19958 | 11750 | 31735 | 26362 | High | High | High | High | High | High | High | High | High | High |
| 23204 | 19339 | 12134 | 25949 | 24111 | High | High | High | High | High | High | High | High | High | High |
| 47351 | 32389 | 18641 | 47128 | 33645 | High | High | High | High | High | High | High | High | High | High |
| 49485 | 30524 | 23766 | 41999 | 37673 | High | High | High | High | High | High | High | High | High | High |
| 36502 | 21397 | 26294 | 21144 | 34934 | High | High | High | High | High | High | High | High | High | High |
| 31412 | 21958 | 19440 | 20709 | 27045 | High | High | High | High | High | High | High | High | High | High |
| 84447 | 13371 | 8421 | 17445 | 52700 | High | High | High | High | High | High | High | High | High | High |
| 40841 | 30998 | 26488 | 34958 | 37560 | High | High | High | High | High | High | High | High | High | High |
| 24969 | 23603 | 19041 | 39874 | 37587 | High | High | High | High | High | High | High | High | High | High |

| | | | | | | | | | | | | | | |
|---|---|---|---|---|---|---|---|---|---|---|---|---|---|---|
| 26766 | 19486 | 15322 | 23726 | 27772 | High | High | High | High | High | High | High | High | High | High |
| 21872 | 12994 | 9116 | 22216 | 19520 | High | High | High | High | High | High | High | High | High | High |
| 16056 | 29574 | 25910 | 33477 | 38534 | High | High | High | High | High | High | High | High | High | High |
| 31039 | 13784 | 10159 | 15823 | 15497 | High | High | High | High | High | High | High | High | High | High |
| 37585 | 42510 | 33902 | 51218 | 56147 | High | High | High | High | High | High | High | High | High | High |
| 30484 | 13766 | 11289 | 16654 | 16158 | High | High | High | High | High | High | High | High | High | High |
| 25465 | 41207 | 27159 | 32411 | 29255 | High | High | High | High | High | High | High | High | High | High |
| 35629 | 30386 | 24072 | 52947 | 40245 | High | High | High | High | High | High | High | High | High | High |
| 14114 | 40864 | 22628 | 38311 | 33594 | High | High | High | High | High | High | High | High | High | High |
| 44058 | 20485 | 15772 | 23370 | 25733 | High | High | High | High | High | High | High | High | High | High |
| 27850 | 15501 | 8610 | 20184 | 16539 | High | High | High | High | High | High | High | High | High | High |
| 30711 | 27892 | 19237 | 43980 | 30727 | High | High | High | High | High | High | High | High | High | High |
| 5595 | 1614 | 1212 | 1312 | 1507 | High | High | High | High | High | High | High | High | High | High |
| 27891 | 10428 | 10693 | 14230 | 18445 | High | High | High | High | High | High | High | High | High | High |
| 36782 | 25823 | 20575 | 34776 | 31474 | High | High | High | High | High | High | High | High | High | High |
| 22816 | 33812 | 22435 | 36156 | 28329 | High | High | High | High | High | High | High | High | High | High |
| 19694 | 32674 | 19988 | 33843 | 30083 | High | High | High | High | High | High | High | High | High | High |
| 27542 | 19367 | 10463 | 26126 | 18683 | High | High | High | High | High | High | High | High | High | High |
| 47312 | 37779 | 29091 | 58977 | 45609 | High | High | High | High | High | High | High | High | High | High |
| 37929 | 43161 | 21401 | 43939 | 35464 | High | High | High | High | High | High | High | High | High | High |
| 23106 | 23597 | 18860 | 31978 | 32305 | High | High | High | High | High | High | High | High | High | High |
| 33983 | 23480 | 16199 | 19692 | 27187 | High | High | High | High | High | High | High | High | High | High |
| 27689 | 20930 | 15664 | 22419 | 24366 | High | High | High | High | High | High | High | High | High | High |
| 19627 | 22224 | 13481 | 23663 | 18024 | High | High | High | High | High | High | High | High | High | High |
| 20663 | 38776 | 29431 | 49669 | 49519 | High | High | High | High | High | High | High | High | High | High |
| 21136 | 19089 | 14701 | 22019 | 22351 | High | High | High | High | High | High | High | High | High | High |
| 40146 | 50505 | 41186 | 49305 | 58370 | High | High | High | High | High | High | High | High | High | High |
| 17327 | 28502 | 23036 | 29978 | 32512 | High | High | High | High | High | High | High | High | High | High |
| 27690 | 28810 | 22346 | 60849 | 55145 | High | High | High | High | High | High | High | High | High | High |
| 33958 | 42064 | 36287 | 51588 | 60426 | High | High | High | High | High | High | High | High | High | High |
| 25905 | 29122 | 30724 | 53592 | 43546 | High | High | High | High | High | High | High | High | High | High |

| | | | | | | | | | | | | | | |
|---|---|---|---|---|---|---|---|---|---|---|---|---|---|---|
| 36159 | 15424 | 10552 | 26439 | 21057 | High | High | High | High | High | High | High | High | High | High |
| 47706 | 43096 | 37795 | 53085 | 61990 | High | High | High | High | High | High | High | High | High | High |
| 22519 | 14673 | 13707 | 17361 | 21083 | High | High | High | High | High | High | High | High | High | High |
| 46672 | 47270 | 35285 | 58048 | 60275 | High | High | High | High | High | High | High | High | High | High |
| 24490 | 26109 | 18640 | 29168 | 28810 | High | High | High | High | High | High | High | High | High | High |
| 26108 | 24943 | 17977 | 38559 | 37793 | High | High | High | High | High | High | High | High | High | High |
| 40213 | 19792 | 17327 | 39558 | 27818 | High | High | High | High | High | High | High | High | High | High |
| 32485 | 29007 | 22508 | 39036 | 41096 | High | High | High | High | High | High | High | High | High | High |
| 19614 | 22654 | 11698 | 38411 | 19383 | High | High | High | High | High | High | High | High | High | High |
| 26731 | 12931 | 9616 | 21873 | 22065 | High | High | High | High | High | High | High | High | High | High |
| 24125 | 14304 | 10603 | 23807 | 17667 | High | High | High | High | High | High | High | High | High | High |
| 29121 | 18397 | 15838 | 24984 | 25946 | High | High | High | High | High | High | High | High | High | High |
| 24621 | 26728 | 21334 | 40289 | 35007 | High | High | High | High | High | High | High | High | High | High |
| 29213 | 13151 | 11709 | 15589 | 20714 | High | High | High | High | High | High | High | High | High | High |
| 32936 | 27976 | 19160 | 47240 | 38745 | High | High | High | High | High | High | High | High | High | High |
| 33039 | 26333 | 17792 | 35801 | 32775 | High | High | High | High | High | High | High | High | High | High |
| 14250 | 24337 | 17995 | 20798 | 22755 | High | High | High | High | High | High | High | High | High | High |
| 24914 | 18819 | 13518 | 33179 | 26173 | High | High | High | High | High | High | High | High | High | High |
| 4553 | 2677 | 1937 | 2471 | 2244 | High | High | High | High | High | High | High | High | High | High |
| 21007 | 25660 | 17675 | 36182 | 28552 | High | High | High | High | High | High | High | High | High | High |
| 62497 | 52144 | 41967 | 55545 | 55949 | High | High | High | High | High | High | High | High | High | High |
| 10063 | 10098 | 8016 | 12687 | 12598 | High | High | High | High | High | High | High | High | High | High |
| 35711 | 18827 | 18591 | 22724 | 28872 | High | High | High | High | High | High | High | High | High | High |
| 26655 | 29935 | 18270 | 32554 | 28786 | High | High | High | High | High | High | High | High | High | High |
| 19621 | 31021 | 25284 | 30496 | 32988 | High | High | High | High | High | High | High | High | High | High |
| 10962 | 20107 | 19504 | 19680 | 28037 | High | High | High | High | High | High | High | High | High | High |
| 23846 | 32442 | 12710 | 26126 | 17253 | High | High | High | High | High | High | High | High | High | High |
| 27423 | 18841 | 12478 | 27213 | 20507 | High | High | High | High | High | High | High | High | High | High |
| 33518 | 9471 | 8672 | 8486 | 12272 | High | High | High | High | High | High | High | High | High | High |
| 27832 | 30557 | 25333 | 47166 | 41347 | High | High | High | High | High | High | High | High | High | High |
| 29301 | 26559 | 19304 | 39714 | 35655 | High | High | High | High | High | High | High | High | High | High |

| | | | | | | | | | | | | | | |
|---|---|---|---|---|---|---|---|---|---|---|---|---|---|---|
| 34991 | 19619 | 13306 | 20953 | 16637 | High | High | High | High | High | High | High | High | High | High |
| | | | | | High | High | High | High | Not Founc | Not Founc | Not Founc | Not Founc | Not Founc | Not Founc |
| 27980 | 19230 | 16355 | 20641 | 23362 | High | High | High | High | High | High | High | High | High | High |
| 31959 | 28863 | 20952 | 36881 | 34868 | High | High | High | High | High | High | High | High | High | High |
| 13777 | 6707 | 3006 | 7756 | 4657 | High | High | High | High | High | High | High | High | High | High |
| 21935 | 21193 | 21040 | 22302 | 30893 | High | High | High | High | High | High | High | High | High | High |
| 25150 | 27475 | 23818 | 48113 | 46372 | High | High | High | High | High | High | High | High | High | High |
| 1886 | 728 | 507 | 938 | 750 | Not Founc | Not Founc | Not Founc | Not Founc | High | High | High | High | High | High |
| 34387 | 22365 | 23023 | 31181 | 32796 | High | High | High | High | High | High | High | High | High | High |
| 34308 | 21676 | 10660 | 32543 | 23586 | High | High | High | High | High | High | High | High | High | High |
| 42315 | 20120 | 17520 | 23932 | 26851 | High | High | High | High | High | High | High | High | High | High |
| 19060 | 18254 | 18611 | 17658 | 26382 | High | High | High | High | High | High | High | High | High | High |
| 21441 | 17068 | 17257 | 14519 | 21713 | High | High | High | High | High | High | High | High | High | High |
| 27594 | 21837 | 19265 | 26632 | 31391 | High | High | High | High | High | High | High | High | High | High |
| 21231 | 27833 | 20202 | 42961 | 32283 | High | High | High | High | High | High | High | High | High | High |
| 16849 | 15071 | 12605 | 16485 | 15925 | High | High | High | High | High | High | High | High | High | High |
| 33622 | 35253 | 18978 | 61525 | 39166 | High | High | High | High | High | High | High | High | High | High |
| 25260 | 24040 | 21337 | 25333 | 30337 | High | High | High | High | High | High | High | High | High | High |
| 44306 | 27376 | 19629 | 35531 | 33725 | High | High | High | High | High | High | High | High | High | High |
| 27152 | 26365 | 36288 | 25917 | 51350 | High | High | High | High | High | High | High | High | High | High |
| 17873 | 10001 | 8316 | 13878 | 17168 | High | High | High | High | High | High | High | High | High | High |
| 28113 | 27719 | 28082 | 30386 | 45100 | High | High | High | High | High | High | High | High | High | High |
| 43737 | 25951 | 17295 | 40970 | 30950 | High | High | High | High | High | High | High | High | High | High |
| 39463 | 13259 | 15358 | 14287 | 22130 | High | High | High | High | High | High | High | High | High | High |
| 19161 | 21481 | 17872 | 28681 | 31040 | High | High | High | High | High | High | High | High | High | High |
| 37495 | 26147 | 21199 | 26182 | 28445 | High | High | High | High | High | High | High | High | High | High |
| 42485 | 12200 | 9068 | 13495 | 63817 | High | High | High | High | High | High | High | High | High | High |
| 32290 | 22056 | 17878 | 24313 | 25269 | High | High | High | High | High | High | High | High | High | High |
| 11023 | 43584 | 26705 | 44557 | 31393 | High | High | High | High | High | High | High | High | High | High |
| 35660 | 25457 | 24322 | 23082 | 33942 | High | High | High | High | High | High | High | High | High | High |
| 18845 | 19202 | 15273 | 27852 | 23862 | High | High | High | High | High | High | High | High | High | High |

| | | | | | | | | | | | | | | |
|---|---|---|---|---|---|---|---|---|---|---|---|---|---|---|
| 22090 | 17868 | 15551 | 18376 | 24341 | High | High | High | High | High | High | High | High | High | High |
| 20125 | 16309 | 14951 | 14317 | 18367 | High | High | High | High | High | High | High | High | High | High |
| 17592 | 9693 | 8438 | 12187 | 12991 | High | High | High | High | High | High | High | High | High | High |
| 53797 | 17532 | 14950 | 23673 | 25222 | High | High | High | High | High | High | High | High | High | High |
| 28812 | 26435 | 15110 | 29389 | 26489 | High | High | High | High | High | High | High | High | High | High |
| 28533 | 14519 | 13773 | 14878 | 16485 | High | High | High | High | High | High | High | High | High | High |
| 32294 | 28474 | 24435 | 40680 | 38862 | High | High | High | High | High | High | High | High | High | High |
| 35550 | 34794 | 26774 | 44482 | 35311 | High | High | High | High | High | High | High | High | High | High |
| 15532 | 11335 | 8991 | 13933 | 12220 | High | High | High | High | High | High | High | High | High | High |
| 25630 | 13085 | 11431 | 14406 | 13196 | High | High | High | High | High | High | High | High | High | High |
| 26768 | 24541 | 16572 | 24373 | 25340 | High | High | High | High | High | High | High | High | High | High |
| 14355 | 25684 | 18504 | 29469 | 30644 | High | High | High | High | High | High | High | High | High | High |
| 12649 | 19702 | 16602 | 17440 | 23803 | High | High | High | High | High | High | High | High | High | High |
| 59941 | 19518 | 14736 | 32456 | 26522 | High | High | High | High | High | High | High | High | High | High |
| 38067 | 34414 | 26153 | 31432 | 36970 | High | High | High | High | High | High | High | High | High | High |
| 32499 | 12599 | 10483 | 12114 | 13406 | High | High | High | High | High | High | High | High | High | High |
| 36318 | 22029 | 19240 | 29199 | 35906 | High | High | High | High | High | High | High | High | High | High |
| 21629 | 16917 | 14922 | 22147 | 24850 | High | High | High | High | High | High | High | High | High | High |
| 8862 | 5980 | 3062 | 13859 | 7951 | Not Founc | Not Founc | Not Founc | Not Founc | High | High | High | High | High | High |
| 37951 | 24379 | 14592 | 25228 | 20114 | High | High | High | High | High | High | High | High | High | High |
| 34340 | 20102 | 15963 | 24806 | 25243 | High | High | High | High | High | High | High | High | High | High |
| 31014 | 21316 | 10991 | 46354 | 25349 | High | High | High | High | High | High | High | High | High | High |
| 19174 | 20948 | 15385 | 29121 | 23979 | High | High | High | High | High | High | High | High | High | High |
| 15321 | 22768 | 13513 | 31168 | 24874 | High | High | High | High | High | High | High | High | High | High |
| 24621 | 24345 | 20003 | 25453 | 28704 | High | High | High | High | High | High | High | High | High | High |
| 2474 | 2314 | 1854 | 1811 | 1681 | Not Founc | Not Founc | Not Founc | Not Founc | High | High | High | High | High | High |
| 27158 | 12128 | 7236 | 14678 | 12400 | High | High | High | High | High | High | High | High | High | High |
| 34651 | 22598 | 14577 | 29732 | 24474 | High | High | High | High | High | High | High | High | High | High |
| 46616 | 34137 | 30976 | 33932 | 44673 | High | High | High | High | High | High | High | High | High | High |
| 34870 | 29493 | 25410 | 27918 | 36341 | High | High | High | High | High | High | High | High | High | High |
| 6875 | 14603 | 8630 | 26530 | 17637 | High | High | High | High | High | High | High | High | High | High |

| | | | | | | | | | | | | | | |
|---|---|---|---|---|---|---|---|---|---|---|---|---|---|---|
| 43108 | 39320 | 34922 | 47546 | 42975 | High | High | High | High | High | High | High | High | High | High |
| 35708 | 13424 | 13879 | 42072 | 23498 | High | High | High | High | High | High | High | High | High | High |
| 11402 | 16989 | 14892 | 16985 | 22265 | High | High | High | High | High | High | High | High | High | High |
| 20487 | 28079 | 22626 | 49879 | 37439 | High | High | High | High | High | High | High | High | High | High |
| 18084 | 14574 | 11270 | 17565 | 21233 | High | High | High | High | High | High | High | High | High | High |
| 17842 | 25849 | 17296 | 37576 | 26743 | High | High | High | High | High | High | High | High | High | High |
| 42480 | 27065 | 19039 | 51552 | 40708 | High | High | High | High | High | High | High | High | High | High |
| 25744 | 33862 | 27549 | 24785 | 25003 | High | High | High | High | High | High | High | High | High | High |
| 32042 | 25876 | 27203 | 32296 | 35929 | High | High | High | High | High | High | High | High | High | High |
| 20315 | 21799 | 14255 | 39197 | 28591 | High | High | High | High | High | High | High | High | High | High |
| 23967 | 17863 | 15188 | 42312 | 27475 | High | High | High | High | High | High | High | High | High | High |
| 15261 | 25512 | 21536 | 26219 | 35693 | High | High | High | High | High | High | High | High | High | High |
| 2041 | 5333 | 4701 | 6732 | 9214 | High | High | High | High | High | High | High | High | High | High |
| 51939 | 10814 | 10057 | 15473 | 37084 | High | High | High | High | High | High | High | High | High | High |
| 19213 | 37634 | 21746 | 57427 | 40898 | High | High | High | High | High | High | High | High | High | High |
| 30523 | 31567 | 27437 | 42960 | 42229 | High | High | High | High | High | High | High | High | High | High |
| 22977 | 15741 | 17480 | 14314 | 19326 | High | High | High | High | High | High | High | High | High | High |
| 26047 | 12026 | 11716 | 10032 | 16465 | High | High | High | High | High | High | High | High | High | High |
| 8348 | 9793 | 3660 | 12594 | 6495 | High | High | High | High | High | High | High | High | High | High |
| 25148 | 22007 | 19708 | 21523 | 26551 | High | High | High | High | High | High | High | High | High | High |
| 32564 | 14874 | 12341 | 19278 | 19740 | High | High | High | High | High | High | High | High | High | High |
| 22045 | 16784 | 13799 | 17401 | 22560 | High | High | High | High | High | High | High | High | High | High |
| 16154 | 29491 | 23515 | 53490 | 38159 | High | High | High | High | High | High | High | High | High | High |
| 31859 | 13766 | 9077 | 19846 | 17605 | High | High | High | High | High | High | High | High | High | High |
| 25304 | 18001 | 8587 | 34373 | 23393 | High | High | High | High | High | High | High | High | High | High |
| 14285 | 32853 | 26038 | 36413 | 35510 | High | High | High | High | High | High | High | High | High | High |
| 30102 | 16367 | 12184 | 20841 | 20548 | High | High | High | High | High | High | High | High | High | High |
| 9233 | 15005 | 12477 | 14100 | 14619 | High | High | High | High | High | High | High | High | High | High |
| 13910 | 17617 | 15454 | 15403 | 19774 | High | High | High | High | High | High | High | High | High | High |
| 44304 | 24846 | 22910 | 27531 | 37181 | High | High | High | High | High | High | High | High | High | High |
| 16534 | 24912 | 17471 | 27662 | 22715 | High | High | High | High | High | High | High | High | High | High |

| | | | | | | | | | | | | | | |
|---|---|---|---|---|---|---|---|---|---|---|---|---|---|---|
| 42174 | 39509 | 26582 | 54465 | 43138 | High | High | High | High | High | High | High | High | High | High |
| 42699 | 6182 | 4470 | 5930 | 32224 | High | High | High | High | High | High | High | High | High | High |
| 15637 | 9873 | 6111 | 17070 | 11557 | High | High | High | High | High | High | High | High | High | High |
| 31153 | 20683 | 15674 | 28143 | 23619 | High | High | High | High | High | High | High | High | High | High |
| | | | | | High | High | High | High | Not Founc | Not Founc | Not Founc | Not Founc | Not Founc | Not Founc |
| 34079 | 4547 | 3179 | 4750 | 27651 | High | High | High | High | High | High | High | High | High | High |
| 15933 | 18661 | 15082 | 22010 | 21523 | High | High | High | High | High | High | High | High | High | High |
| 43552 | 11046 | 8616 | 23324 | 16281 | High | High | High | High | High | High | High | High | High | High |
| 23100 | 31614 | 25193 | 36140 | 39572 | High | High | High | High | High | High | High | High | High | High |
| 24883 | 19510 | 15607 | 22769 | 24639 | High | High | High | High | High | High | High | High | High | High |
| 33748 | 28445 | 22560 | 33911 | 33901 | High | High | High | High | High | High | High | High | High | High |
| 24773 | 7821 | 5622 | 10229 | 9843 | High | High | High | High | High | High | High | High | High | High |
| 9671 | 15545 | 11288 | 15890 | 12904 | High | High | High | High | High | High | High | High | High | High |
| 27314 | 14707 | 12171 | 17848 | 18706 | High | High | High | High | High | High | High | High | High | High |
| 37577 | 36764 | 29737 | 36685 | 38677 | High | High | High | High | High | High | High | High | High | High |
| 27850 | 17114 | 15344 | 24056 | 26597 | High | High | High | High | High | High | High | High | High | High |
| 36172 | 19761 | 19324 | 27061 | 29245 | High | High | High | High | High | High | High | High | High | High |
| 15425 | 14305 | 12733 | 13735 | 17225 | High | High | High | High | High | High | High | High | High | High |
| 7063 | 18679 | 17580 | 19663 | 21570 | High | High | High | High | High | High | High | High | High | High |
| 21977 | 15966 | 10951 | 23010 | 17271 | High | High | High | High | High | High | High | High | High | High |
| 26437 | 26716 | 19455 | 48596 | 34282 | High | High | High | High | High | High | High | High | High | High |
| 22066 | 20388 | 13578 | 28493 | 22071 | High | High | High | High | High | High | High | High | High | High |
| 7033 | 30534 | 17984 | 37992 | 27820 | High | High | High | High | High | High | High | High | High | High |
| 26890 | 8868 | 8230 | 6874 | 8220 | High | High | High | High | High | High | High | High | High | High |
| 30250 | 16559 | 17939 | 20564 | 29156 | High | High | High | High | High | High | High | High | High | High |
| 34929 | 13931 | 11969 | 19866 | 17776 | High | High | High | High | High | High | High | High | High | High |
| 12840 | 16358 | 9676 | 21927 | 16958 | High | High | High | High | High | High | High | High | High | High |
| 46659 | 19719 | 19117 | 31485 | 35560 | High | High | High | High | High | High | High | High | High | High |
| 9654 | 13979 | 8935 | 16856 | 13748 | High | High | High | High | High | High | High | High | High | High |
| 19064 | 16668 | 13655 | 19140 | 19986 | High | High | High | High | High | High | High | High | High | High |
| 19957 | 15420 | 15405 | 15772 | 21835 | High | High | High | High | High | High | High | High | High | High |

| | | | | | | | | | | | | | | |
|---|---|---|---|---|---|---|---|---|---|---|---|---|---|---|
| 15583 | 16356 | 9760 | 21509 | 17544 | High | High | High | High | High | High | High | High | High | High |
| 16104 | 21546 | 18754 | 21353 | 22895 | High | High | High | High | High | High | High | High | High | High |
| 29198 | 18829 | 16268 | 21069 | 23694 | High | High | High | High | High | High | High | High | High | High |
| 15404 | 23591 | 15645 | 27901 | 25319 | High | High | High | High | High | High | High | High | High | High |
| 24428 | 12981 | 9503 | 14430 | 17108 | High | High | High | High | High | High | High | High | High | High |
| 20558 | 18776 | 16483 | 23253 | 24994 | High | High | High | High | High | High | High | High | High | High |
| 22959 | 22831 | 18349 | 21175 | 23558 | High | High | High | High | High | High | High | High | High | High |
| 16933 | 12828 | 8810 | 17028 | 14427 | High | High | High | High | High | High | High | High | High | High |
| 37718 | 17450 | 16325 | 20085 | 26064 | High | High | High | High | High | High | High | High | High | High |
| 26437 | 24636 | 17123 | 31591 | 30838 | High | High | High | High | High | High | High | High | High | High |
| 26739 | 27803 | 19724 | 28261 | 31312 | High | High | High | High | High | High | High | High | High | High |
| 25335 | 15115 | 11267 | 17711 | 18192 | High | High | High | High | High | High | High | High | High | High |
| 19908 | 15607 | 15369 | 29635 | 23942 | High | High | High | High | High | High | High | High | High | High |
| 49830 | 29803 | 21166 | 39096 | 33583 | High | High | High | High | High | High | High | High | High | High |
| 25229 | 23108 | 15768 | 21122 | 20194 | High | High | High | High | High | High | High | High | High | High |
| 43366 | 11915 | 10634 | 19147 | 21575 | High | High | High | High | High | High | High | High | High | High |
| 11877 | 27823 | 16638 | 27534 | 25559 | High | High | High | High | High | High | High | High | High | High |
| 34633 | 30734 | 21506 | 37437 | 32768 | High | High | High | High | High | High | High | High | High | High |
| 2906 | 1264 | 1104 | 1904 | 2339 | High | High | High | High | High | High | High | High | High | High |
| 25592 | 32052 | 27074 | 34391 | 40223 | High | High | High | High | High | High | High | High | High | High |
| 36611 | 23547 | 16145 | 48720 | 39440 | High | High | High | High | High | High | High | High | High | High |
| 28327 | 23184 | 15958 | 28797 | 25060 | High | High | High | High | High | High | High | High | High | High |
| 13324 | 9458 | 5997 | 20010 | 15232 | High | High | High | High | High | High | High | High | High | High |
| 13103 | 20852 | 14809 | 21806 | 18152 | High | High | High | High | High | High | High | High | High | High |
| 29242 | 4085 | 3538 | 4339 | 5792 | High | High | High | High | High | High | High | High | High | High |
| 20312 | 21704 | 16393 | 25265 | 27398 | High | High | High | High | High | High | High | High | High | High |
| 3816 | | | | | Not Founc | Not Founc | Not Founc | Not Founc | High | High | High | High | Not Founc | Not Founc |
| 10939 | 18293 | 10954 | 28911 | 25468 | High | High | High | High | High | High | High | High | High | High |
| 9533 | 12230 | 8727 | 20101 | 15488 | High | High | High | High | High | High | High | High | High | High |
| 17688 | 17195 | 9443 | 28082 | 18379 | High | High | High | High | High | High | High | High | High | High |
| 22866 | 15882 | 8136 | 22645 | 15351 | High | High | High | High | High | High | High | High | High | High |

| | | | | | | | | | | | | | | |
|---|---|---|---|---|---|---|---|---|---|---|---|---|---|---|
| 1943 | 3373 | 2788 | 2461 | 2681 | High | High | High | High | High | High | High | High | High | High |
| 38774 | 21203 | 18288 | 22200 | 23584 | High | High | High | High | High | High | High | High | High | High |
| 9490 | 14486 | 6018 | 13304 | 8364 | High | High | High | High | High | High | High | High | High | High |
| 18767 | 11761 | 9807 | 12703 | 13844 | High | High | High | High | High | High | High | High | High | High |
| 13884 | 13482 | 10847 | 34314 | 19512 | High | High | High | High | High | High | High | High | High | High |
| 19406 | 20145 | 16224 | 24759 | 29460 | High | High | High | High | High | High | High | High | High | High |
| 26624 | 24494 | 21338 | 31300 | 35621 | High | High | High | High | High | High | High | High | High | High |
| 18438 | 25121 | 18893 | 25395 | 25988 | High | High | High | High | High | High | High | High | High | High |
| 36225 | 18165 | 11998 | 25907 | 22053 | High | High | High | High | High | High | High | High | High | High |
| 40187 | 20727 | 14424 | 25392 | 21514 | High | High | High | High | High | High | High | High | High | High |
| 8447 | 11031 | 8747 | 14108 | 13894 | High | High | High | High | High | High | High | High | High | High |
| 19251 | 16015 | 10986 | 26543 | 23175 | High | High | High | High | High | High | High | High | High | High |
| 22720 | 9685 | 5787 | 25016 | 14889 | High | High | High | High | High | High | High | High | High | High |
| 15822 | 28844 | 16980 | 57478 | 34723 | High | High | High | High | High | High | High | High | High | High |
| 16755 | 13573 | 11091 | 14794 | 16658 | High | High | High | High | High | High | High | High | High | High |
| 28458 | 16461 | 11010 | 16889 | 15916 | High | High | High | High | High | High | High | High | High | High |
| 20540 | 20322 | 14836 | 21798 | 20448 | High | High | High | High | High | High | High | High | High | High |
| 17809 | 14963 | 10966 | 27289 | 19677 | High | High | High | High | High | High | High | High | High | High |
| 15694 | 19050 | 12476 | 24454 | 21038 | High | High | High | High | High | High | High | High | High | High |
| 17716 | 12853 | 8102 | 14835 | 14047 | High | High | High | High | High | High | High | High | High | High |
| 17745 | 19719 | 13542 | 16944 | 16850 | High | High | High | High | High | High | High | High | High | High |
| 13000 | 21127 | 16628 | 27878 | 24169 | High | High | High | High | High | High | High | High | High | High |
| 21937 | 35393 | 29530 | 26703 | 32232 | High | High | High | High | High | High | High | High | High | High |
| 23025 | 17119 | 14543 | 22277 | 23857 | High | High | High | High | High | High | High | High | High | High |
| 10510 | 13278 | 10280 | 15517 | 14381 | High | High | High | High | High | High | High | High | High | High |
| 9741 | 51257 | 13336 | 41971 | 22658 | High | High | High | High | High | High | High | High | High | High |
| 24710 | 26885 | 18952 | 51155 | 34690 | High | High | High | High | High | High | High | High | High | High |
| 24225 | 10701 | 9644 | 11930 | 13851 | High | High | High | High | High | High | High | High | High | High |
| 18460 | 13833 | 10916 | 20219 | 17618 | High | High | High | High | High | High | High | High | High | High |
| 22696 | 13948 | 16209 | 15492 | 21535 | High | High | High | High | High | High | High | High | High | High |
| 36082 | 21176 | 21321 | 27513 | 36499 | High | High | High | High | High | High | High | High | High | High |

| | | | | | | | | | | | | | | |
|---|---|---|---|---|---|---|---|---|---|---|---|---|---|---|
| 17636 | 13203 | 7356 | 26644 | 15174 | High | High | High | High | High | High | High | High | High | High |
| 13626 | 21741 | 15603 | 26934 | 24594 | High | High | High | High | High | High | High | High | High | High |
| 33871 | 13944 | 9458 | 17611 | 44602 | High | High | High | High | High | High | High | High | High | High |
| 17244 | 19865 | 17207 | 34258 | 25946 | High | High | High | High | High | High | High | High | High | High |
| 23967 | 21032 | 13718 | 34219 | 31808 | High | High | High | High | High | High | High | High | High | High |
| 24036 | 30725 | 28193 | 24051 | 23336 | High | High | High | High | High | High | High | High | High | High |
| 32084 | 22788 | 18424 | 27228 | 33399 | High | High | High | High | High | High | High | High | High | High |
| 26727 | 23762 | 20052 | 35423 | 32565 | High | High | High | High | High | High | High | High | High | High |
| 21433 | 10296 | 8048 | 11829 | 12429 | High | High | High | High | High | High | High | High | High | High |
| 9532 | 22119 | 17762 | 24346 | 24903 | High | High | High | High | High | High | High | High | High | High |
| 16246 | 7023 | 6229 | 5997 | 7542 | High | High | High | High | High | High | High | High | High | High |
| 26610 | 10989 | 9468 | 12410 | 13841 | High | High | High | High | High | High | High | High | High | High |
| 33213 | 11939 | 9221 | 18254 | 18024 | High | High | High | High | High | High | High | High | High | High |
| 7484 | 5754 | 2904 | 6148 | 5179 | High | High | High | High | High | High | High | High | High | High |
| 14348 | 22340 | 16836 | 22511 | 24394 | High | High | High | High | High | High | High | High | High | High |
| 12282 | 5787 | 5366 | 5719 | 8035 | High | High | High | High | High | High | High | High | High | High |
| 24105 | 7600 | 5681 | 8223 | 8627 | High | High | High | High | High | High | High | High | High | High |
| 23993 | 15046 | 10060 | 23775 | 15757 | High | High | High | High | High | High | High | High | High | High |
| 18218 | 7359 | 5458 | 8933 | 8852 | High | High | High | High | High | High | High | High | High | High |
| 25572 | 22642 | 15374 | 53149 | 33893 | High | High | High | High | High | High | High | High | High | High |
| 10729 | 16727 | 12780 | 21435 | 21299 | High | High | High | High | High | High | High | High | High | High |
| 14966 | 8321 | 6425 | 10626 | 11217 | High | High | High | High | High | High | High | High | High | High |
| 11988 | 19900 | 17186 | 27748 | 27111 | High | High | High | High | High | High | High | High | High | High |
| 39455 | 15866 | 15868 | 25718 | 29279 | High | High | High | High | High | High | High | High | High | High |
| 14396 | 8297 | 5974 | 13319 | 9499 | High | High | High | High | High | High | High | High | High | High |
| 20683 | 27105 | 18258 | 36480 | 27817 | High | High | High | High | High | High | High | High | High | High |
| 15739 | 12072 | 12572 | 29807 | 21693 | High | High | High | High | High | High | High | High | High | High |
| 43573 | 16057 | 13874 | 14613 | 19101 | High | High | High | High | High | High | High | High | High | High |
| 13963 | 19103 | 14806 | 24701 | 23025 | High | High | High | High | High | High | High | High | High | High |
| 22362 | 23731 | 16771 | 32769 | 33541 | High | High | High | High | High | High | High | High | High | High |
| 12471 | 20582 | 14923 | 25450 | 21190 | High | High | High | High | High | High | High | High | High | High |

| | | | | | | | | | | | | | | |
|---|---|---|---|---|---|---|---|---|---|---|---|---|---|---|
| 23795 | 18555 | 17132 | 16896 | 21525 | High | High | High | High | High | High | High | High | High | High |
| 12343 | 10498 | 8269 | 12616 | 14962 | High | High | High | High | High | High | High | High | High | High |
| | | | | | Not Founc | Not Founc | Not Founc | Not Founc | High | High | High | High | Not Founc | Not Founc |
| 23505 | 10341 | 6313 | 14395 | 11849 | High | High | High | High | High | High | High | High | High | High |
| 34548 | 32460 | 29674 | 34105 | 39842 | High | High | High | High | High | High | High | High | High | High |
| 33903 | 11853 | 14005 | 12954 | 19124 | High | High | High | High | High | High | High | High | High | High |
| 18620 | 12387 | 11112 | 13084 | 16263 | High | High | High | High | High | High | High | High | High | High |
| 36464 | 13827 | 10303 | 31190 | 21497 | High | High | High | High | High | High | High | High | High | High |
| 15445 | 15596 | 12003 | 22798 | 20474 | High | High | High | High | High | High | High | High | High | High |
| 26988 | 14977 | 11209 | 26434 | 21271 | High | High | High | High | High | High | High | High | High | High |
| 42442 | 18302 | 9023 | 28132 | 26310 | High | High | High | High | High | High | High | High | High | High |
| 13566 | 38260 | 27089 | 40230 | 42517 | High | High | High | High | High | High | High | High | High | High |
| 31130 | 18890 | 17196 | 20299 | 24904 | High | High | High | High | High | High | High | High | High | High |
| 26450 | 17942 | 13680 | 20648 | 22005 | High | High | High | High | High | High | High | High | High | High |
| 10188 | 13312 | 10801 | 14823 | 18490 | High | High | High | High | High | High | High | High | High | High |
| 10106 | 11683 | 9824 | 17548 | 16521 | High | High | High | High | High | High | High | High | High | High |
| 34551 | 19413 | 18230 | 39504 | 32004 | High | High | High | High | High | High | High | High | High | High |
| 15571 | 8142 | 5564 | 12315 | 9904 | High | High | High | High | High | High | High | High | High | High |
| 33686 | 20812 | 16819 | 31842 | 25636 | High | High | High | High | High | High | High | High | High | High |
| 25168 | 11283 | 10359 | 17708 | 16780 | High | High | High | High | High | High | High | High | High | High |
| 33217 | 13389 | 10813 | 13090 | 14683 | High | High | High | High | High | High | High | High | High | High |
| 12736 | 14544 | 8675 | 18891 | 14893 | High | High | High | High | High | High | High | High | High | High |
| 14267 | 14631 | 11544 | 14877 | 12272 | High | High | High | High | High | High | High | High | High | High |
| 26823 | 13488 | 10204 | 23499 | 16432 | High | High | High | High | High | High | High | High | High | High |
| 31990 | 16481 | 13065 | 29387 | 26033 | High | High | High | High | High | High | High | High | High | High |
| 18422 | 20733 | 19153 | 22995 | 21495 | High | High | High | High | High | High | High | High | High | High |
| 20366 | 15590 | 11066 | 22382 | 20390 | High | High | High | High | High | High | High | High | High | High |
| 15331 | 13228 | 7689 | 10934 | 10241 | High | High | High | High | High | High | High | High | High | High |
| 12918 | 18552 | 12614 | 25732 | 19988 | High | High | High | High | High | High | High | High | High | High |
| 23829 | 14289 | 9962 | 33003 | 19720 | High | High | High | High | High | High | High | High | High | High |
| 11151 | 7751 | 4783 | 9950 | 7818 | High | High | High | High | High | High | High | High | High | High |

| | | | | | | | | | | | | | | |
|---|---|---|---|---|---|---|---|---|---|---|---|---|---|---|
| 20673 | 11858 | 4022 | 9279 | 6849 | High | High | High | High | High | High | High | High | High | High |
| 28157 | 8870 | 8155 | 8396 | 10337 | High | High | High | High | High | High | High | High | High | High |
| 33985 | 26120 | 20081 | 38774 | 29606 | High | High | High | High | High | High | High | High | High | High |
| 11889 | 12704 | 8604 | 15170 | 14440 | High | High | High | High | High | High | High | High | High | High |
| 17850 | 28331 | 18694 | 27732 | 24011 | High | High | High | High | High | High | High | High | High | High |
| 11057 | 9366 | 7369 | 12113 | 11333 | High | High | High | High | High | High | High | High | High | High |
| 19042 | 30081 | 16783 | 39007 | 28520 | High | High | High | High | High | High | High | High | High | High |
| 19328 | 19160 | 12731 | 25934 | 22788 | High | High | High | High | High | High | High | High | High | High |
| 22206 | 14186 | 11476 | 17663 | 20888 | High | High | High | High | High | High | High | High | High | High |
| 3304 | 8867 | 5850 | 7392 | 6688 | High | High | High | High | High | High | High | High | High | High |
| 12565 | 7605 | 5381 | 7580 | 7347 | High | High | High | High | High | High | High | High | High | High |
| 12360 | 18899 | 18186 | 27324 | 29794 | High | High | High | High | High | High | High | High | High | High |
| 10630 | 20333 | 16576 | 22690 | 26436 | High | High | High | High | High | High | High | High | High | High |
| 13544 | 14722 | 14669 | 21418 | 20613 | High | High | High | High | High | High | High | High | High | High |
| 25146 | 14550 | 12412 | 22982 | 28059 | High | High | High | High | High | High | High | High | High | High |
| 37467 | 10374 | 7951 | 15200 | 13858 | High | High | High | High | High | High | High | High | High | High |
| 24073 | 12312 | 8497 | 20603 | 17190 | High | High | High | High | High | High | High | High | High | High |
| 35333 | 13106 | 11947 | 18332 | 21441 | High | High | High | High | High | High | High | High | High | High |
| 18539 | 7248 | 5205 | 9095 | 7193 | High | High | High | High | High | High | High | High | High | High |
| 25686 | 24967 | 18791 | 35170 | 27573 | High | High | High | High | High | High | High | High | High | High |
| 28054 | 17371 | 15721 | 15849 | 20253 | High | High | High | High | High | High | High | High | High | High |
| 36681 | 13669 | 12930 | 14045 | 20894 | High | High | High | High | High | High | High | High | High | High |
| 4081 | 7872 | 6440 | 13640 | 11368 | High | High | High | High | High | High | High | High | High | High |
| 13633 | 14799 | 10973 | 24416 | 18296 | High | High | High | High | High | High | High | High | High | High |
| 22969 | 22597 | 17674 | 44523 | 44110 | High | High | High | High | High | High | High | High | High | High |
| 21922 | 10448 | 8570 | 15783 | 13740 | High | High | High | High | High | High | High | High | High | High |
| 19669 | 14494 | 11831 | 18573 | 22112 | High | High | High | High | High | High | High | High | High | High |
| 37545 | 13873 | 13578 | 16952 | 23351 | High | High | High | High | High | High | High | High | High | High |
| 29554 | 28015 | 22407 | 32933 | 34420 | High | High | High | High | High | High | High | High | High | High |
| 29009 | 16683 | 13989 | 21761 | 24910 | High | High | High | High | High | High | High | High | High | High |
| 12692 | 22798 | 15604 | 24845 | 24635 | High | High | High | High | High | High | High | High | High | High |

| | | | | | | | | | | | | | | |
|---|---|---|---|---|---|---|---|---|---|---|---|---|---|---|
| 19906 | 22058 | 15018 | 25374 | 21888 | High | High | High | High | High | High | High | High | High | High |
| 25015 | 23452 | 17223 | 20261 | 21221 | High | High | High | High | High | High | High | High | High | High |
| 7369 | 20205 | 17460 | 32305 | 22580 | High | High | High | High | High | High | High | High | High | High |
| 18349 | 22877 | 14694 | 29966 | 26372 | High | High | High | High | High | High | High | High | High | High |
| 7513 | 6957 | 5375 | 9033 | 8846 | High | High | High | High | High | High | High | High | High | High |
| 26299 | 14026 | 13811 | 30659 | 20472 | High | High | High | High | High | High | High | High | High | High |
| 11523 | 10647 | 6560 | 18274 | 13220 | High | High | High | High | High | High | High | High | High | High |
| 20332 | 15737 | 12930 | 26585 | 24112 | High | High | High | High | High | High | High | High | High | High |
| 10852 | 6380 | 6112 | 7430 | 10238 | High | High | High | High | High | High | High | High | High | High |
| 13551 | 14498 | 9886 | 23658 | 16092 | High | High | High | High | High | High | High | High | High | High |
| 11888 | 7490 | 5436 | 20870 | 17992 | High | High | High | High | High | High | High | High | High | High |
| 21846 | 10133 | 7410 | 13291 | 11602 | High | High | High | High | High | High | High | High | High | High |
| 18767 | 16164 | 12864 | 27032 | 18506 | High | High | High | High | High | High | High | High | High | High |
| 19765 | 11572 | 7475 | 11840 | 11411 | High | High | High | High | High | High | High | High | High | High |
| 30768 | 11708 | 9865 | 11928 | 15556 | High | High | High | High | High | High | High | High | High | High |
| 15988 | 18001 | 8621 | 19881 | 15534 | High | High | High | High | High | High | High | High | High | High |
| 36536 | 24922 | 21042 | 28849 | 36414 | High | High | High | High | High | High | High | High | High | High |
| 15705 | 14871 | 11919 | 22948 | 19464 | High | High | High | High | High | High | High | High | High | High |
| 25636 | 17919 | 15223 | 18710 | 18613 | High | High | High | High | High | High | High | High | High | High |
| 13013 | 10504 | 7073 | 14868 | 12189 | High | High | High | High | High | High | High | High | High | High |
| 8941 | 5458 | 4206 | 7016 | 6278 | High | High | High | High | High | High | High | High | High | High |
| 14392 | 18724 | 14943 | 22868 | 23584 | High | High | High | High | High | High | High | High | High | High |
| 16635 | 10648 | 7546 | 14857 | 13688 | High | High | High | High | High | High | High | High | High | High |
| 8964 | 12120 | 9810 | 13532 | 13060 | High | High | High | High | High | High | High | High | High | High |
| 4145 | 10537 | 9281 | 10231 | 11326 | High | High | High | High | High | High | High | High | High | High |
| 9824 | 14726 | 9814 | 16253 | 14251 | High | High | High | High | High | High | High | High | High | High |
| 19136 | 14300 | 9540 | 16303 | 15689 | High | High | High | High | High | High | High | High | High | High |
| 28413 | 20261 | 12977 | 37669 | 26445 | High | High | High | High | High | High | High | High | High | High |
| 28626 | 7220 | 6926 | 10822 | 10355 | High | High | High | High | High | High | High | High | High | High |
| 26429 | 15380 | 12215 | 16726 | 16075 | High | High | High | High | High | High | High | High | High | High |
| 24960 | 6394 | 4548 | 13314 | 8426 | High | High | High | High | High | High | High | High | High | High |

| | | | | | | | | | | | | | | |
|---|---|---|---|---|---|---|---|---|---|---|---|---|---|---|
| 20470 | 21912 | 17625 | 36727 | 30202 | High | High | High | High | High | High | High | High | High | High |
| 9960 | 13486 | 10010 | 20824 | 19830 | High | High | High | High | High | High | High | High | High | High |
| 24938 | 10551 | 9380 | 13743 | 14108 | High | High | High | High | High | High | High | High | High | High |
| 17127 | 21135 | 14530 | 29569 | 23714 | High | High | High | High | High | High | High | High | High | High |
| 29540 | 18718 | 16610 | 20286 | 24114 | High | High | High | High | High | High | High | High | High | High |
| 15616 | 11989 | 9983 | 16070 | 19450 | High | High | High | High | High | High | High | High | High | High |
| 28847 | 10898 | 9315 | 13164 | 15046 | High | High | High | High | High | High | High | High | High | High |
| 18767 | 17909 | 18260 | 19413 | 26434 | High | High | High | High | High | High | High | High | High | High |
| 23121 | 13069 | 12143 | 14930 | 12811 | High | High | High | High | High | High | High | High | High | High |
| 11028 | 7958 | 6265 | 10519 | 8754 | High | High | High | High | High | High | High | High | High | High |
| 11177 | 21617 | 19172 | 23198 | 27297 | High | High | High | High | High | High | High | High | High | High |
| 15952 | 18499 | 13932 | 22283 | 21739 | High | High | High | High | High | High | High | High | High | High |
| 6412 | 2086 | 1847 | 1454 | 2107 | High | High | High | High | High | High | High | High | High | High |
| 10934 | 10800 | 8528 | 19022 | 14967 | High | High | High | High | High | High | High | High | High | High |
| 19614 | 8767 | 5732 | 10076 | 9002 | High | High | High | High | High | High | High | High | High | High |
| 23542 | 16542 | 18147 | 20060 | 26439 | High | High | High | High | High | High | High | High | High | High |
| 9812 | 20488 | 18358 | 22254 | 32102 | High | High | High | High | High | High | High | High | High | High |
| 26814 | 18070 | 12744 | 27242 | 20624 | High | High | High | High | High | High | High | High | High | High |
| 17143 | 13429 | 11225 | 13186 | 16972 | High | High | High | High | High | High | High | High | High | High |
| 9193 | 12236 | 8809 | 13226 | 11679 | High | High | High | High | High | High | High | High | High | High |
| 10055 | 22132 | 12396 | 36503 | 22254 | High | High | High | High | High | High | High | High | High | High |
| 11097 | 35157 | 7140 | 46004 | 13622 | High | High | High | High | High | High | High | High | High | High |
| 13497 | 4973 | 3288 | 7326 | 6701 | High | High | High | High | High | High | High | High | High | High |
| 28158 | 17131 | 16452 | 17322 | 23261 | High | High | High | High | High | High | High | High | High | High |
| 6969 | 5869 | 4855 | 5838 | 7080 | High | High | High | High | High | High | High | High | High | High |
| 20215 | 14651 | 10756 | 15954 | 16410 | High | High | High | High | High | High | High | High | High | High |
| 23164 | 11484 | 10660 | 14487 | 15673 | High | High | High | High | High | High | High | High | High | High |
| 19723 | 4609 | 4751 | 4542 | 6112 | High | High | High | High | High | High | High | High | High | High |
| 12784 | 4215 | 2557 | 10671 | 6834 | High | High | High | High | High | High | High | High | High | High |
| 6797 | 5472 | 3402 | 3709 | 4440 | High | High | High | High | High | High | High | High | High | High |
| 14516 | 11686 | 10045 | 12388 | 14463 | High | High | High | High | High | High | High | High | High | High |

| | | | | | | | | | | | | | | |
|---|---|---|---|---|---|---|---|---|---|---|---|---|---|---|
| 16989 | 12921 | 11406 | 15300 | 20082 | High | High | High | High | High | High | High | High | High | High |
| 24668 | 7129 | 4307 | 13815 | 10162 | High | High | High | High | High | High | High | High | High | High |
| 3045 | 14848 | 8103 | 17765 | 15446 | High | High | High | High | High | High | High | High | High | High |
| 7443 | 7498 | 5142 | 12666 | 7917 | High | High | High | High | High | High | High | High | High | High |
| 4581 | 551 | 564 | 423 | 910 | High | High | High | High | High | High | High | High | High | High |
| 15123 | 10258 | 9666 | 11827 | 12553 | High | High | High | High | High | High | High | High | High | High |
| 9800 | 24408 | 19161 | 34044 | 27428 | High | High | High | High | High | High | High | High | High | High |
| 1371 | 6014 | 2872 | 7806 | 7234 | High | High | High | High | High | High | High | High | High | High |
| 18386 | 9613 | 7946 | 7504 | 9907 | High | High | High | High | High | High | High | High | High | High |
| 8231 | 9325 | 5080 | 12944 | 9016 | High | High | High | High | High | High | High | High | High | High |
| 7409 | 3095 | 2774 | 3914 | 3385 | High | High | High | High | High | High | High | High | High | High |
| 22505 | 19157 | 15220 | 23136 | 21862 | High | High | High | High | High | High | High | High | High | High |
| 19884 | 10811 | 7645 | 12170 | 12858 | High | High | High | High | High | High | High | High | High | High |
| 4808 | 16883 | 15406 | 13396 | 17705 | High | High | High | High | High | High | High | High | High | High |
| 11343 | 16928 | 14077 | 24428 | 21489 | High | High | High | High | High | High | High | High | High | High |
| 8381 | 9501 | 7044 | 8686 | 10137 | High | High | High | High | High | High | High | High | High | High |
| 16026 | 23868 | 20622 | 18930 | 20956 | High | High | High | High | High | High | High | High | High | High |
| 12592 | 8822 | 9063 | 8576 | 13994 | High | High | High | High | High | High | High | High | High | High |
| 21778 | 10888 | 10344 | 12515 | 15952 | High | High | High | High | High | High | High | High | High | High |
| 20246 | 28096 | 20276 | 33243 | 29875 | High | High | High | High | High | High | High | High | High | High |
| 14645 | 2415 | 1618 | 4562 | 3179 | High | High | High | High | High | High | High | High | High | High |
| 15486 | 20954 | 16270 | 23421 | 22066 | High | High | High | High | High | High | High | High | High | High |
| 11003 | 12195 | 9193 | 12163 | 12580 | High | High | High | High | High | High | High | High | High | High |
| 17609 | 5370 | 4217 | 5544 | 6530 | High | High | High | High | High | High | High | High | High | High |
| 15485 | 9666 | 10281 | 8309 | 13444 | High | High | High | High | High | High | High | High | High | High |
| 15928 | 9766 | 6367 | 11028 | 10169 | High | High | High | High | High | High | High | High | High | High |
| 15160 | 18422 | 14096 | 22556 | 24093 | High | High | High | High | High | High | High | High | High | High |
| 20716 | 17681 | 14881 | 19243 | 21222 | High | High | High | High | High | High | High | High | High | High |
| 20058 | 20792 | 16531 | 20011 | 26659 | High | High | High | High | High | High | High | High | High | High |
| 18705 | 20496 | 11325 | 20066 | 17129 | High | High | High | High | High | High | High | High | High | High |
| 17623 | 14994 | 9978 | 16130 | 14657 | High | High | High | High | High | High | High | High | High | High |

| | | | | | | | | | | | | | | |
|---|---|---|---|---|---|---|---|---|---|---|---|---|---|---|
| 21318 | 12781 | 6541 | 31371 | 17208 | High | High | High | High | High | High | High | High | High | High |
| 7129 | 6426 | 3535 | 8331 | 7961 | High | High | High | High | High | High | High | High | High | High |
| 12828 | 10889 | 7489 | 10970 | 12471 | High | High | High | High | High | High | High | High | High | High |
| 12408 | 13988 | 8165 | 27051 | 17021 | High | High | High | High | High | High | High | High | High | High |
| 32805 | 13984 | 14054 | 18121 | 20331 | High | High | High | High | High | High | High | High | High | High |
| 12741 | 4432 | 2685 | 8902 | 5628 | High | High | High | High | High | High | High | High | High | High |
| 5999 | 6830 | 5606 | 6662 | 6693 | High | High | High | High | High | High | High | High | High | High |
| 6587 | 11231 | 8114 | 13823 | 13364 | High | High | High | High | High | High | High | High | High | High |
| 14370 | 11557 | 8707 | 17977 | 15811 | High | High | High | High | High | High | High | High | High | High |
| 15256 | 20297 | 11566 | 32792 | 22758 | High | High | High | High | High | High | High | High | High | High |
| 13113 | 14693 | 8762 | 17267 | 12445 | High | High | High | High | High | High | High | High | High | High |
| 14876 | 16800 | 12115 | 21733 | 20778 | High | High | High | High | High | High | High | High | High | High |
| 17847 | 25207 | 12886 | 27525 | 24964 | High | High | High | High | High | High | High | High | High | High |
| 30741 | 15564 | 8486 | 28283 | 20044 | High | High | High | High | High | High | High | High | High | High |
| 38713 | 21906 | 18978 | 23816 | 30196 | High | High | High | High | High | High | High | High | High | High |
| 7998 | 10297 | 8980 | 15949 | 14350 | High | High | High | High | High | High | High | High | High | High |
| 10392 | 21336 | 14179 | 30296 | 21920 | High | High | High | High | High | High | High | High | High | High |
| 6546 | 14614 | 10859 | 16403 | 17264 | High | High | High | High | High | High | High | High | High | High |
| 7167 | 8479 | 5286 | 7453 | 8126 | High | High | High | High | High | High | High | High | High | High |
| 23422 | 18729 | 9787 | 22988 | 18882 | High | High | High | High | High | High | High | High | High | High |
| 8246 | 9544 | 6029 | 18947 | 11269 | High | High | High | High | High | High | High | High | High | High |
| 24168 | 19256 | 14697 | 24505 | 21060 | High | High | High | High | High | High | High | High | High | High |
| 8656 | 8953 | 6979 | 8678 | 10265 | High | High | High | High | High | High | High | High | High | High |
| 12934 | 10094 | 9034 | 12581 | 13966 | High | High | High | High | High | High | High | High | High | High |
| 16926 | 17542 | 7870 | 35047 | 18648 | High | High | High | High | High | High | High | High | High | High |
| 22612 | 13488 | 11335 | 14456 | 16201 | High | High | High | High | High | High | High | High | High | High |
| 2314 | 7183 | 4336 | 7216 | 6804 | High | High | High | High | High | High | High | High | High | High |
| 9569 | 10224 | 8211 | 8959 | 10128 | High | High | High | High | High | High | High | High | High | High |
| 12145 | 8097 | 5649 | 13622 | 9142 | High | High | High | High | High | High | High | High | High | High |
| 11125 | 10907 | 8349 | 15188 | 15755 | High | High | High | High | High | High | High | High | High | High |
| 18049 | 5325 | 4404 | 10022 | 6740 | High | High | High | High | High | High | High | High | High | High |

| | | | | | | | | | | | | | | |
|---|---|---|---|---|---|---|---|---|---|---|---|---|---|---|
| 12264 | 13669 | 10718 | 18315 | 17656 | High | High | High | High | High | High | High | High | High | High |
| 18233 | 13044 | 10963 | 14723 | 18445 | High | High | High | High | High | High | High | High | High | High |
| 31063 | 11708 | 10484 | 11552 | 15375 | High | High | High | High | High | High | High | High | High | High |
| 8417 | 8808 | 5934 | 16734 | 9767 | High | High | High | High | High | High | High | High | High | High |
| 12595 | 21691 | 13647 | 31337 | 24795 | High | High | High | High | High | High | High | High | High | High |
| 17204 | 39090 | 29626 | 45103 | 41464 | High | High | High | High | High | High | High | High | High | High |
| 18543 | 9902 | 5796 | 18665 | 13226 | High | High | High | High | High | High | High | High | High | High |
| 6911 | 14566 | 13607 | 15567 | 20535 | High | High | High | High | High | High | High | High | High | High |
| 9084 | 1483 | 1284 | 1954 | 2074 | High | High | High | High | High | High | High | High | High | High |
| 9673 | 15953 | 13072 | 18869 | 18272 | High | High | High | High | High | High | High | High | High | High |
| 22086 | 20720 | 16089 | 21230 | 25908 | High | High | High | High | High | High | High | High | High | High |
| 30976 | 11178 | 9534 | 14661 | 14072 | High | High | High | High | High | High | High | High | High | High |
| 14900 | 18606 | 15351 | 18988 | 22028 | High | High | High | High | High | High | High | High | High | High |
| 14076 | 20256 | 13952 | 20591 | 20891 | High | High | High | High | High | High | High | High | High | High |
| 17303 | 19617 | 16640 | 24368 | 24544 | High | High | High | High | High | High | High | High | High | High |
| 18010 | 18029 | 13105 | 26451 | 21172 | High | High | High | High | High | High | High | High | High | High |
| 20260 | 6183 | 5109 | 9813 | 8475 | High | High | High | High | High | High | High | High | High | High |
| 6379 | 8972 | 7337 | 8834 | 8213 | High | High | High | High | High | High | High | High | High | High |
| 4155 | 13664 | 8881 | 22500 | 19517 | High | High | High | High | High | High | High | High | High | High |
| 7318 | 8515 | 7152 | 12107 | 14404 | High | High | High | High | High | High | High | High | High | High |
| 10058 | 11192 | 10436 | 11739 | 14699 | High | High | High | High | High | High | High | High | High | High |
| 20855 | 7750 | 6407 | 9118 | 10456 | High | High | High | High | High | High | High | High | High | High |
| 11904 | 11200 | 10094 | 13952 | 14751 | High | High | High | High | High | High | High | High | High | High |
| 4327 | 10988 | 6550 | 13566 | 9345 | High | High | High | High | High | High | High | High | High | High |
| 15971 | 11704 | 11307 | 12680 | 15079 | High | High | High | High | High | High | High | High | High | High |
| 29784 | 12833 | 10421 | 13070 | 15269 | High | High | High | High | High | High | High | High | High | High |
| 3814 | 15378 | 8372 | 21022 | 13002 | High | High | High | High | High | High | High | High | High | High |
| 6706 | 10606 | 6342 | 14492 | 10505 | High | High | High | High | High | High | High | High | High | High |
| 11547 | 6851 | 5771 | 7845 | 8403 | High | High | High | High | High | High | High | High | High | High |
| 24818 | 10955 | 9228 | 10494 | 16157 | High | High | High | High | High | High | High | High | High | High |
| 26755 | 17449 | 10949 | 32680 | 19956 | High | High | High | High | High | High | High | High | High | High |

| | | | | | | | | | | | | | | |
|---|---|---|---|---|---|---|---|---|---|---|---|---|---|---|
| 27233 | 20249 | 16041 | 27418 | 26528 | High | High | High | High | High | High | High | High | High | High |
| 17506 | 1348 | 1044 | 1945 | 1851 | High | High | High | High | High | High | High | High | High | High |
| 16783 | 23231 | 18596 | 27017 | 27095 | High | High | High | High | High | High | High | High | High | High |
| 16200 | 13812 | 22170 | 47813 | 29058 | High | High | High | High | High | High | High | High | High | High |
| 18213 | 18031 | 14822 | 17540 | 20952 | High | High | High | High | High | High | High | High | High | High |
| 18917 | 13865 | 11457 | 16309 | 19444 | High | High | High | High | High | High | High | High | High | High |
| 26222 | 17879 | 12773 | 15855 | 18215 | High | High | High | High | High | High | High | High | High | High |
| 13252 | 7218 | 5905 | 12591 | 9369 | High | High | High | High | High | High | High | High | High | High |
| 2152 | 15461 | 12804 | 19634 | 18394 | High | High | High | High | High | High | High | High | High | High |
| 8907 | 8438 | 7206 | 15117 | 12203 | High | High | High | High | High | High | High | High | High | High |
| 11207 | 3427 | 3357 | 12119 | 9738 | High | High | High | High | High | High | High | High | High | High |
| 19482 | 11994 | 9192 | 16090 | 14502 | High | High | High | High | High | High | High | High | High | High |
| 16369 | 16997 | 18180 | 14347 | 21727 | High | High | High | High | High | High | High | High | High | High |
| 22614 | 7866 | 6190 | 5845 | 7166 | High | High | High | High | High | High | High | High | High | High |
| 14201 | 16621 | 10299 | 16547 | 14400 | High | High | High | High | High | High | High | High | High | High |
| 15049 | 18828 | 16252 | 21716 | 24819 | High | High | High | High | High | High | High | High | High | High |
| 13506 | 19693 | 17032 | 26372 | 24653 | High | High | High | High | High | High | High | High | High | High |
| 20722 | 17727 | 17717 | 23174 | 21157 | High | High | High | High | High | High | High | High | High | High |
| 10754 | 6494 | 5313 | 6247 | 6651 | High | High | High | High | High | High | High | High | High | High |
| 22645 | 9176 | 7997 | 10897 | 12020 | High | High | High | High | High | High | High | High | High | High |
| 12569 | 12285 | 15322 | 13497 | 25054 | High | High | High | High | High | High | High | High | High | High |
| 21233 | 10851 | 6422 | 14895 | 9990 | High | High | High | High | High | High | High | High | High | High |
| 7598 | 8232 | 6312 | 11478 | 9776 | High | High | High | High | High | High | High | High | High | High |
| 24378 | 11412 | 15027 | 12521 | 18615 | High | High | High | High | High | High | High | High | High | High |
| 8500 | 16986 | 13919 | 36301 | 21409 | High | High | High | High | High | High | High | High | High | High |
| 15144 | 7838 | 6548 | 19108 | 11366 | High | High | High | High | High | High | High | High | High | High |
| 18928 | 7337 | 5353 | 8881 | 8257 | High | High | High | High | High | High | High | High | High | High |
| 35416 | 7075 | 4755 | 8608 | 7280 | High | High | High | High | High | High | High | High | High | High |
| 16100 | 15081 | 13583 | 30709 | 26389 | High | High | High | High | High | High | High | High | High | High |
| 25742 | 6611 | 5582 | 7020 | 8341 | High | High | High | High | High | High | High | High | High | High |
| 16980 | 10028 | 8000 | 12212 | 13127 | High | High | High | High | High | High | High | High | High | High |

| | | | | | | | | | | | | | | |
|---|---|---|---|---|---|---|---|---|---|---|---|---|---|---|
| 25163 | 10076 | 10813 | 19156 | 17504 | High | High | High | High | High | High | High | High | High | High |
| 15201 | 13601 | 10274 | 17398 | 15230 | High | High | High | High | High | High | High | High | High | High |
| 7592 | 8974 | 7649 | 13319 | 10724 | High | High | High | High | High | High | High | High | High | High |
| 6433 | 3667 | 3449 | 4037 | 5206 | High | High | High | High | High | High | High | High | High | High |
| 17602 | 9895 | 6408 | 10006 | 10561 | High | High | High | High | High | High | High | High | High | High |
| 7316 | 9001 | 6088 | 9806 | 9207 | High | High | High | High | High | High | High | High | High | High |
| 18582 | 8694 | 6696 | 11729 | 10207 | High | High | High | High | High | High | High | High | High | High |
| 10522 | 10850 | 8037 | 13723 | 14029 | High | High | High | High | High | High | High | High | High | High |
| 14625 | 6441 | 4570 | 4114 | 4959 | High | High | High | High | High | High | High | High | High | High |
| 21196 | 20965 | 17495 | 28856 | 30839 | High | High | High | High | High | High | High | High | High | High |
| 14056 | 13149 | 10062 | 14423 | 15428 | High | High | High | High | High | High | High | High | High | High |
| 12240 | 6808 | 4747 | 10608 | 7541 | High | High | High | High | High | High | High | High | High | High |
| 34675 | 7819 | 9013 | 9165 | 15499 | High | High | High | High | High | High | High | High | High | High |
| 20550 | 13934 | 12249 | 14194 | 17330 | High | High | High | High | High | High | High | High | High | High |
| 7276 | 7655 | 5718 | 14287 | 10397 | High | High | High | High | High | High | High | High | High | High |
| 12912 | 14814 | 11984 | 14912 | 18060 | High | High | High | High | High | High | High | High | High | High |
| 4414 | 7944 | 5491 | 8059 | 7904 | High | High | High | High | High | High | High | High | High | High |
| 21131 | 4375 | 5101 | 11842 | 8003 | High | High | High | High | High | High | High | High | High | High |
| 15131 | 12489 | 10312 | 14499 | 14476 | High | High | High | High | High | High | High | High | High | High |
| 12201 | 18378 | 17065 | 14765 | 19498 | High | High | High | High | High | High | High | High | High | High |
| 4175 | 3611 | 2277 | 4214 | 3624 | High | High | High | High | High | High | High | High | High | High |
| 25977 | 14518 | 12532 | 12476 | 14333 | High | High | High | High | High | High | High | High | High | High |
| 12284 | 12822 | 8998 | 13686 | 11987 | High | High | High | High | High | High | High | High | High | High |
| 10069 | 25163 | 19361 | 40881 | 36369 | High | High | High | High | High | High | High | High | High | High |
| 16382 | 14770 | 13055 | 18618 | 19806 | High | High | High | High | High | High | High | High | High | High |
| 10662 | 12207 | 9014 | 17564 | 14126 | High | High | High | High | High | High | High | High | High | High |
| 7657 | 9767 | 9651 | 9465 | 13487 | High | High | High | High | High | High | High | High | High | High |
| 4329 | 3236 | 3008 | 3751 | 3823 | High | High | High | High | High | High | High | High | High | High |
| 17245 | 14133 | 9870 | 26266 | 16824 | High | High | High | High | High | High | High | High | High | High |
| 38232 | 16868 | 14670 | 21770 | 26293 | High | High | High | High | High | High | High | High | High | High |
| 11856 | 18757 | 14758 | 17519 | 20943 | High | High | High | High | High | High | High | High | High | High |

| | | | | | | | | | | | | | | |
|---|---|---|---|---|---|---|---|---|---|---|---|---|---|---|
| 11546 | 5716 | 4721 | 7185 | 8687 | High | High | High | High | High | High | High | High | High | High |
| 21153 | 44128 | 35312 | 41263 | 61154 | High | High | High | High | High | High | High | High | High | High |
| 16472 | 11350 | 7052 | 34062 | 17381 | High | High | High | High | High | High | High | High | High | High |
| 10598 | 7701 | 7608 | 6021 | 10018 | High | High | High | High | High | High | High | High | High | High |
| 3879 | 8029 | 6288 | 12009 | 10153 | High | High | High | High | High | High | High | High | High | High |
| 12534 | 10373 | 9055 | 11365 | 9501 | High | High | High | High | High | High | High | High | High | High |
| 16806 | 12307 | 9777 | 17563 | 14793 | High | High | High | High | High | High | High | High | High | High |
| 10488 | 12334 | 5090 | 18851 | 9987 | High | High | High | High | High | High | High | High | High | High |
| 14478 | 9393 | 8549 | 12216 | 11781 | High | High | High | High | High | High | High | High | High | High |
| 3233 | 9289 | 6876 | 15479 | 11847 | High | High | High | High | High | High | High | High | High | High |
| 9876 | 7887 | 4619 | 8049 | 6985 | High | High | High | High | High | High | High | High | High | High |
| 13964 | 10629 | 7112 | 12577 | 12189 | High | High | High | High | High | High | High | High | High | High |
| 6930 | 24503 | 19306 | 27501 | 27985 | High | High | High | High | High | High | High | High | High | High |
| 8144 | 3503 | 2719 | 2599 | 3240 | High | High | High | High | High | High | High | High | High | High |
| 7618 | 14333 | 12248 | 20899 | 18315 | High | High | High | High | High | High | High | High | High | High |
| 13020 | 10061 | 7440 | 13940 | 11184 | High | High | High | High | High | High | High | High | High | High |
| 12119 | 15403 | 11702 | 23882 | 17786 | High | High | High | High | High | High | High | High | High | High |
| 22376 | 8556 | 6178 | 14011 | 12210 | High | High | High | High | High | High | High | High | High | High |
| 12238 | 5554 | 4440 | 12284 | 6942 | High | High | High | High | High | High | High | High | High | High |
| 5090 | 13272 | 11059 | 23314 | 15633 | High | High | High | High | High | High | High | High | High | High |
| 4922 | 4144 | 3431 | 3603 | 4118 | High | High | High | High | High | High | High | High | High | High |
| 20594 | 8596 | 6602 | 11310 | 11004 | High | High | High | High | High | High | High | High | High | High |
| 6055 | 4128 | 2886 | 6059 | 4717 | High | High | High | High | High | High | High | High | High | High |
| 12886 | 5704 | 4652 | 6858 | 7298 | High | High | High | High | High | High | High | High | High | High |
| 6704 | 11626 | 7757 | 14651 | 13986 | High | High | High | High | High | High | High | High | High | High |
| 11178 | 22501 | 11293 | 21160 | 15933 | High | High | High | High | High | High | High | High | High | High |
| 14958 | 5278 | 4537 | 6295 | 7221 | High | High | High | High | High | High | High | High | High | High |
| 11252 | 7889 | 6489 | 8523 | 9692 | High | High | High | High | High | High | High | High | High | High |
| 17021 | 4426 | 3314 | 7727 | 5915 | High | High | High | High | High | High | High | High | High | High |
| 10948 | 22816 | 17548 | 26999 | 26522 | High | High | High | High | High | High | High | High | High | High |
| 5799 | 9520 | 7820 | 10484 | 9975 | High | High | High | High | High | High | High | High | High | High |

| | | | | | | | | | | | | | | |
|---|---|---|---|---|---|---|---|---|---|---|---|---|---|---|
| 5090 | 13740 | 10032 | 14135 | 16167 | High | High | High | High | High | High | High | High | High | High |
| 14454 | 4366 | 3436 | 5373 | 5124 | High | High | High | High | High | High | High | High | High | High |
| 18023 | 10848 | 4976 | 13694 | 9113 | High | High | High | High | High | High | High | High | High | High |
| 14483 | 11751 | 10156 | 11892 | 14844 | High | High | High | High | High | High | High | High | High | High |
| 16820 | 5212 | 3159 | 9115 | 6185 | High | High | High | High | High | High | High | High | High | High |
| 12751 | 3821 | 2821 | 4363 | 4102 | High | High | High | High | High | High | High | High | High | High |
| 14917 | 5497 | 2610 | 7425 | 4429 | High | High | High | High | High | High | High | High | High | High |
| 5586 | 10232 | 3771 | 9025 | 5558 | High | High | High | High | High | High | High | High | High | High |
| 9956 | 3649 | 2401 | 3899 | 3715 | High | High | High | High | High | High | High | High | High | High |
| 17259 | 14372 | 11353 | 12921 | 13196 | High | High | High | High | High | High | High | High | High | High |
| 18015 | 17480 | 11874 | 29744 | 25734 | High | High | High | High | High | High | High | High | High | High |
| | | | | | High | High | High | High | Not Founc | Not Founc | Not Founc | Not Founc | Not Founc | Not Founc |
| 5847 | 12634 | 9992 | 23014 | 17019 | High | High | High | High | High | High | High | High | High | High |
| 15668 | 15475 | 8589 | 30028 | 19071 | High | High | High | High | High | High | High | High | High | High |
| 9571 | 11894 | 10398 | 14008 | 18006 | High | High | High | High | High | High | High | High | High | High |
| 16554 | 12426 | 8713 | 32499 | 18489 | High | High | High | High | High | High | High | High | High | High |
| 7996 | 11316 | 10772 | 31684 | 17816 | High | High | High | High | High | High | High | High | High | High |
| 9917 | 14102 | 10959 | 17937 | 17634 | High | High | High | High | High | High | High | High | High | High |
| 14252 | 13909 | 9882 | 12081 | 12248 | High | High | High | High | High | High | High | High | High | High |
| 9522 | 10038 | 8863 | 12064 | 14687 | High | High | High | High | High | High | High | High | High | High |
| 9462 | 16228 | 11197 | 16390 | 15444 | High | High | High | High | High | High | High | High | High | High |
| 11283 | 8626 | 10134 | 13794 | 11814 | High | High | High | High | High | High | High | High | High | High |
| 24001 | 19172 | 23553 | 21958 | 29046 | High | High | High | High | High | High | High | High | High | High |
| 17878 | 19602 | 15049 | 30365 | 26037 | High | High | High | High | High | High | High | High | High | High |
| 11568 | 5786 | 4286 | 6867 | 7311 | High | High | High | High | High | High | High | High | High | High |
| 9187 | 5591 | 4662 | 9967 | 9488 | High | High | High | High | High | High | High | High | High | High |
| 9595 | 6241 | 4734 | 5022 | 4016 | High | High | High | High | High | High | High | High | High | High |
| 21757 | 18940 | 16886 | 20741 | 31220 | High | High | High | High | High | High | High | High | High | High |
| 12080 | 8341 | 6043 | 9058 | 10847 | High | High | High | High | High | High | High | High | High | High |
| 9420 | 4727 | 3836 | 4109 | 4461 | High | High | High | High | High | High | High | High | High | High |
| 8690 | 7285 | 7026 | 8455 | 7517 | High | High | High | High | High | High | High | High | High | High |

| | | | | | | | | | | | | | | |
|---|---|---|---|---|---|---|---|---|---|---|---|---|---|---|
| 15795 | 20506 | 18491 | 20123 | 26354 | High | High | High | High | High | High | High | High | High | High |
| 12498 | 675 | 527 | 645 | 848 | High | High | High | High | High | High | High | High | High | High |
| 1403 | 9692 | 6740 | 11133 | 11021 | High | High | High | High | High | High | High | High | High | High |
| 7303 | 3012 | 2490 | 4045 | 3857 | High | High | High | High | High | High | High | High | High | High |
| 11867 | 5393 | 4355 | 5633 | 5584 | High | High | High | High | High | High | High | High | High | High |
| 11605 | 11730 | 6399 | 13793 | 9926 | High | High | High | High | High | High | High | High | High | High |
| 10087 | 10221 | 6359 | 9653 | 9010 | High | High | High | High | High | High | High | High | High | High |
| 14615 | 13284 | 9061 | 22782 | 13323 | High | High | High | High | High | High | High | High | High | High |
| 13637 | 7998 | 4404 | 7968 | 6988 | High | High | High | High | High | High | High | High | High | High |
| | | | | | High | High | High | High | Not Found | Not Found | Not Found | Not Found | Not Found | Not Found |
| 22116 | 2928 | 2303 | 2612 | 3768 | High | High | High | High | High | High | High | High | High | High |
| 5989 | 6395 | 6002 | 6569 | 8098 | High | High | High | High | High | High | High | High | High | High |
| 10723 | 9975 | 7706 | 9129 | 9500 | High | High | High | High | High | High | High | High | High | High |
| 7915 | 14751 | 19661 | 22182 | 9798 | High | High | High | High | High | High | High | High | High | High |
| 11745 | 3723 | 3212 | 4111 | 4325 | High | High | High | High | High | High | High | High | High | High |
| 21827 | 4870 | 3145 | 5376 | 19010 | High | High | High | High | High | High | High | High | High | High |
| 18494 | 15260 | 12026 | 18564 | 19519 | High | High | High | High | High | High | High | High | High | High |
| 10092 | 10675 | 9276 | 9145 | 12244 | High | High | High | High | High | High | High | High | High | High |
| 16411 | 10233 | 11170 | 11058 | 17431 | High | High | High | High | High | High | High | High | High | High |
| 14119 | 9935 | 8193 | 15959 | 12689 | High | High | High | High | High | High | High | High | High | High |
| 10800 | 6794 | 5324 | 7177 | 7134 | High | High | High | High | High | High | High | High | High | High |
| 18526 | 5948 | 5477 | 9968 | 10223 | High | High | High | High | High | High | High | High | High | High |
| 5041 | 6884 | 5796 | 7830 | 9258 | High | High | High | High | High | High | High | High | High | High |
| 7706 | 20361 | 12755 | 22694 | 18335 | High | High | High | High | High | High | High | High | High | High |
| 3899 | 3382 | 2522 | 4958 | 4586 | Not Found | Not Found | Not Found | Not Found | High | High | High | High | High | High |
| 31470 | 7502 | 8776 | 8371 | 12756 | High | High | High | High | High | High | High | High | High | High |
| 13172 | 5133 | 3751 | 6360 | 5417 | High | High | High | High | High | High | High | High | High | High |
| 13425 | 21670 | 16113 | 32010 | 30700 | High | High | High | High | High | High | High | High | High | High |
| 17810 | 18521 | 14801 | 20142 | 21406 | High | High | High | High | High | High | High | High | High | High |
| 6733 | 8557 | 7965 | 12586 | 13994 | High | High | High | High | High | High | High | High | High | High |
| 11971 | 4370 | 4791 | 5148 | 4280 | High | High | High | High | High | High | High | High | High | High |

| | | | | | | | | | | | | | | |
|---|---|---|---|---|---|---|---|---|---|---|---|---|---|---|
| 14183 | 7667 | 4859 | 11953 | 8190 | High | High | High | High | High | High | High | High | High | High |
| 13484 | 10239 | 7420 | 12208 | 12601 | High | High | High | High | High | High | High | High | High | High |
| 11025 | 9570 | 8566 | 9207 | 11836 | High | High | High | High | High | High | High | High | High | High |
| 16654 | 9532 | 8203 | 10244 | 13290 | High | High | High | High | High | High | High | High | High | High |
| 18693 | 12752 | 9421 | 21320 | 17497 | High | High | High | High | High | High | High | High | High | High |
| 7983 | 13875 | 10292 | 18142 | 18582 | High | High | High | High | High | High | High | High | High | High |
| 12025 | 20985 | 17574 | 30218 | 27281 | High | High | High | High | High | High | High | High | High | High |
| 4020 | 2984 | 2128 | 5059 | 3591 | High | High | High | High | High | High | High | High | High | High |
| 16816 | 9094 | 6501 | 8461 | 9306 | High | High | High | High | High | High | High | High | High | High |
| 5511 | 3227 | 2466 | 5506 | 3877 | High | High | High | High | High | High | High | High | High | High |
| 11716 | 7176 | 6471 | 8657 | 9506 | High | High | High | High | High | High | High | High | High | High |
| 12859 | 8707 | 10143 | 8951 | 14586 | High | High | High | High | High | High | High | High | High | High |
| 18881 | 6604 | 4859 | 8092 | 7376 | High | High | High | High | High | High | High | High | High | High |
| 12698 | 3333 | 2436 | 3737 | 3494 | High | High | High | High | High | High | High | High | High | High |
| 7163 | 14844 | 8784 | 13284 | 12051 | High | High | High | High | High | High | High | High | High | High |
| 3875 | 3209 | 3089 | 3816 | 4659 | High | High | High | High | High | High | High | High | High | High |
| 19392 | 16123 | 17937 | 17375 | 25023 | High | High | High | High | High | High | High | High | High | High |
| 10290 | 12300 | 10656 | 12412 | 15272 | High | High | High | High | High | High | High | High | High | High |
| 4629 | 5388 | 4799 | 9406 | 8131 | High | High | High | High | High | High | High | High | High | High |
| 6863 | 2106 | 3252 | 2023 | 3541 | High | High | High | High | High | High | High | High | High | High |
| 12165 | 11713 | 9256 | 16561 | 15775 | High | High | High | High | High | High | High | High | High | High |
| 15705 | 7119 | 6044 | 7835 | 9185 | High | High | High | High | High | High | High | High | High | High |
| 4060 | 9078 | 8389 | 9512 | 11444 | High | High | High | High | High | High | High | High | High | High |
| 5225 | 5068 | 4185 | 5203 | 5869 | High | High | High | High | High | High | High | High | High | High |
| 5466 | 13089 | 7187 | 10860 | 8366 | High | High | High | High | High | High | High | High | High | High |
| 10478 | 11864 | 9556 | 15665 | 17234 | High | High | High | High | High | High | High | High | High | High |
| 23525 | 8715 | 7494 | 9921 | 12021 | High | High | High | High | High | High | High | High | High | High |
| 3829 | | | | | Not Founc | Not Founc | Not Founc | Not Founc | High | High | High | High | Not Founc | Not Founc |
| | 244 | 200 | 457 | 399 | High | High | High | High | Not Founc | Not Founc | Not Founc | Not Founc | High | High |
| 13592 | 17844 | 11828 | 21748 | 17573 | High | High | High | High | High | High | High | High | High | High |
| 19092 | 5285 | 3801 | 7984 | 6803 | High | High | High | High | High | High | High | High | High | High |

| | | | | | | | | | | | | | | |
|---|---|---|---|---|---|---|---|---|---|---|---|---|---|---|
| 2694 | 3188 | 7800 | 3565 | 2404 | High | High | High | High | High | High | High | High | High | High |
| 15621 | 7884 | 4397 | 16709 | 11355 | High | High | High | High | High | High | High | High | High | High |
| 20678 | 9818 | 5781 | 21044 | 11800 | High | High | High | High | High | High | High | High | High | High |
| 15408 | 19492 | 15799 | 18847 | 25447 | High | High | High | High | High | High | High | High | High | High |
| 9252 | 8913 | 7232 | 20140 | 11697 | High | High | High | High | High | High | High | High | High | High |
| 15003 | 7964 | 5489 | 11945 | 8744 | High | High | High | High | High | High | High | High | High | High |
| 7695 | 16095 | 10512 | 19918 | 16328 | High | High | High | High | High | High | High | High | High | High |
| 13995 | 8603 | 6604 | 12443 | 10406 | High | High | High | High | High | High | High | High | High | High |
| 9146 | 10721 | 6311 | 19516 | 12520 | High | High | High | High | High | High | High | High | High | High |
| 6345 | 12528 | 10812 | 13513 | 13812 | High | High | High | High | High | High | High | High | High | High |
| 9036 | 5272 | 3308 | 5453 | 5076 | High | High | High | High | High | High | High | High | High | High |
| 1473 | 2458 | 1897 | 4459 | 3809 | High | High | High | High | High | High | High | High | High | High |
| 16186 | 15088 | 11697 | 18252 | 19271 | High | High | High | High | High | High | High | High | High | High |
| 6684 | 12849 | 8170 | 15171 | 12066 | High | High | High | High | High | High | High | High | High | High |
| 5986 | 12712 | 11953 | 18664 | 21274 | High | High | High | High | High | High | High | High | High | High |
| 5270 | 21265 | 6491 | 16502 | 9370 | High | High | High | High | High | High | High | High | High | High |
| 14310 | 13751 | 9636 | 15807 | 13191 | High | High | High | High | High | High | High | High | High | High |
| 6249 | 698 | 1003 | 932 | 951 | High | High | High | High | High | High | High | High | High | High |
| 8380 | 6087 | 3252 | 9018 | 5230 | High | High | High | High | High | High | High | High | High | High |
| 25844 | 5037 | 3807 | 4096 | 39483 | High | High | High | High | High | High | High | High | High | High |
| 43290 | 12216 | 18743 | 11320 | 33153 | High | High | High | High | High | High | High | High | High | High |
| 10978 | 5184 | 3183 | 7089 | 5129 | High | High | High | High | High | High | High | High | High | High |
| 15033 | 12093 | 8341 | 12612 | 12697 | High | High | High | High | High | High | High | High | High | High |
| 5516 | 7171 | 6481 | 6983 | 8492 | High | High | High | High | High | High | High | High | High | High |
| 16279 | 11275 | 6755 | 33816 | 17989 | High | High | High | High | High | High | High | High | High | High |
| 12748 | 12959 | 10402 | 15291 | 14820 | High | High | High | High | High | High | High | High | High | High |
| 1095 | 9970 | 7651 | 14027 | 10771 | High | High | High | High | High | High | High | High | High | High |
| 15285 | 11571 | 9195 | 14470 | 13536 | High | High | High | High | High | High | High | High | High | High |
| 9408 | 12284 | 9588 | 18998 | 20423 | High | High | High | High | High | High | High | High | High | High |
| 8723 | 14019 | 8897 | 14327 | 13628 | High | High | High | High | High | High | High | High | High | High |
| 7594 | 12603 | 10958 | 11669 | 14446 | High | High | High | High | High | High | High | High | High | High |

| | | | | | | | | | | | | | | |
|---|---|---|---|---|---|---|---|---|---|---|---|---|---|---|
| 25451 | 8086 | 7117 | 10328 | 11276 | High | High | High | High | High | High | High | High | High | High |
| 5360 | 6367 | 4892 | 8205 | 7582 | High | High | High | High | High | High | High | High | High | High |
| 13873 | 10386 | 8065 | 15684 | 14701 | High | High | High | High | High | High | High | High | High | High |
| 4664 | 4374 | 2378 | 5283 | 4031 | High | High | High | High | High | High | High | High | High | High |
| 12343 | 5261 | 5130 | 5327 | 7141 | High | High | High | High | High | High | High | High | High | High |
| 12668 | 14946 | 12162 | 24788 | 19810 | High | High | High | High | High | High | High | High | High | High |
| 15017 | 6735 | 7078 | 5505 | 10949 | High | High | High | High | High | High | High | High | High | High |
| 16188 | 20555 | 10930 | 34866 | 19879 | High | High | High | High | High | High | High | High | High | High |
| 12076 | 4465 | 3710 | 4466 | 4678 | High | High | High | High | High | High | High | High | High | High |
| 12939 | 7411 | 4995 | 12273 | 10552 | High | High | High | High | High | High | High | High | High | High |
| 14551 | 3895 | 4922 | 9289 | 8534 | High | High | High | High | High | High | High | High | High | High |
| 12117 | 2561 | 1562 | 3209 | 2726 | High | High | High | High | High | High | High | High | High | High |
| 22367 | 20048 | 15517 | 20743 | 25247 | High | High | High | High | High | High | High | High | High | High |
| 8795 | 9662 | 7170 | 12234 | 11890 | High | High | High | High | High | High | High | High | High | High |
| 11168 | 8031 | 5610 | 8058 | 8764 | High | High | High | High | High | High | High | High | High | High |
| 6584 | 3352 | 2545 | 5444 | 3524 | High | High | High | High | High | High | High | High | High | High |
| 7292 | 10294 | 7124 | 11842 | 11877 | High | High | High | High | High | High | High | High | High | High |
| 6837 | 11815 | 9415 | 14646 | 16013 | High | High | High | High | High | High | High | High | High | High |
| 14083 | 9301 | 5886 | 21606 | 13036 | High | High | High | High | High | High | High | High | High | High |
| 5748 | 13945 | 12836 | 18800 | 18767 | High | High | High | High | High | High | High | High | High | High |
| 16874 | 8770 | 9561 | 9484 | 13132 | High | High | High | High | High | High | High | High | High | High |
| 29110 | 17032 | 11027 | 19511 | 16971 | High | High | High | High | High | High | High | High | High | High |
| 29691 | 10917 | 8578 | 10490 | 12366 | High | High | High | High | High | High | High | High | High | High |
| 19336 | 10317 | 7158 | 13876 | 12953 | High | High | High | High | High | High | High | High | High | High |
| 9896 | 3843 | 4349 | 4211 | 5735 | High | High | High | High | High | High | High | High | High | High |
| 6590 | 5455 | 3553 | 7212 | 5434 | High | High | High | High | High | High | High | High | High | High |
| 2809 | 4056 | 1963 | 4402 | 3248 | Not Founc | Not Founc | Not Founc | Not Founc | High | High | High | High | High | High |
| 10948 | 7248 | 5954 | 8674 | 10133 | High | High | High | High | High | High | High | High | High | High |
| 11184 | 6775 | 4698 | 9094 | 7586 | High | High | High | High | High | High | High | High | High | High |
| 16950 | 5648 | 2817 | 4679 | 3762 | High | High | High | High | High | High | High | High | High | High |
| 6287 | | | | | High | High | High | High | High | High | High | High | Not Founc | Not Founc |

| | | | | | | | | | | | | | | |
|---|---|---|---|---|---|---|---|---|---|---|---|---|---|---|
| 12385 | 8710 | 3786 | 10725 | 7264 | High | High | High | High | High | High | High | High | High | High |
| 11847 | 9448 | 6723 | 14513 | 11222 | High | High | High | High | High | High | High | High | High | High |
| 12642 | 8538 | 7211 | 11478 | 11278 | High | High | High | High | High | High | High | High | High | High |
| 10531 | 6804 | 5336 | 10436 | 9348 | High | High | High | High | High | High | High | High | High | High |
| 2563 | 461 | 741 | 615 | 334 | High | High | High | High | High | High | High | High | High | High |
| 6515 | 3049 | 2348 | 3378 | 3426 | High | High | High | High | High | High | High | High | High | High |
| | 6159 | 4431 | 7363 | 6680 | High | High | High | High | Not Founc | Not Founc | Not Founc | Not Founc | High | High |
| 14094 | 7654 | 5404 | 8083 | 7863 | High | High | High | High | High | High | High | High | High | High |
| 8290 | 8978 | 6655 | 8129 | 9517 | High | High | High | High | High | High | High | High | High | High |
| 14070 | 22786 | 9231 | 50115 | 25874 | High | High | High | High | High | High | High | High | High | High |
| 17020 | 9536 | 8311 | 14497 | 17185 | High | High | High | High | High | High | High | High | High | High |
| 8149 | 6386 | 5295 | 7223 | 6665 | High | High | High | High | High | High | High | High | High | High |
| 7309 | 6704 | 4534 | 8378 | 6820 | High | High | High | High | High | High | High | High | High | High |
| 22757 | 6594 | 4586 | 6500 | 7490 | High | High | High | High | High | High | High | High | High | High |
| 3877 | 9503 | 7528 | 9603 | 9513 | High | High | High | High | High | High | High | High | High | High |
| 11979 | 7371 | 5483 | 8365 | 9072 | High | High | High | High | High | High | High | High | High | High |
| 10203 | 8133 | 6831 | 11049 | 9760 | High | High | High | High | High | High | High | High | High | High |
| 16616 | 18807 | 10982 | 19195 | 17754 | High | High | High | High | High | High | High | High | High | High |
| 5456 | 6602 | 4807 | 6176 | 7226 | High | High | High | High | High | High | High | High | High | High |
| 5999 | 3401 | 2486 | 3833 | 4680 | High | High | High | High | High | High | High | High | High | High |
| 11696 | 11363 | 6408 | 19169 | 12231 | High | High | High | High | High | High | High | High | High | High |
| 17688 | 23026 | 17566 | 47406 | 28472 | High | High | High | High | High | High | High | High | High | High |
| 7659 | 5713 | 3359 | 9256 | 6571 | High | High | High | High | High | High | High | High | High | High |
| 19690 | 7182 | 5834 | 7517 | 8685 | High | High | High | High | High | High | High | High | High | High |
| 34488 | 8989 | 7784 | 12055 | 13962 | High | High | High | High | High | High | High | High | High | High |
| 12925 | 3062 | 2383 | 4204 | 3410 | High | High | High | High | High | High | High | High | High | High |
| 4505 | 4546 | 2900 | 6093 | 4260 | High | High | High | High | High | High | High | High | High | High |
| 14355 | 10158 | 7177 | 11172 | 11113 | High | High | High | High | High | High | High | High | High | High |
| 18615 | 11019 | 8159 | 15859 | 14664 | High | High | High | High | High | High | High | High | High | High |
| 7949 | 5232 | 3665 | 6081 | 5469 | High | High | High | High | High | High | High | High | High | High |
| 10996 | 7626 | 5222 | 12888 | 7649 | High | High | High | High | High | High | High | High | High | High |

| | | | | | | | | | | | | | | |
|---|---|---|---|---|---|---|---|---|---|---|---|---|---|---|
| 3061 | 1535 | 1335 | 1610 | 2116 | High | High | High | High | High | High | High | High | High | High |
| 4859 | 5832 | 5087 | 6888 | 10385 | High | High | High | High | High | High | High | High | High | High |
| 8352 | 3317 | 2874 | 8973 | 6258 | High | High | High | High | High | High | High | High | High | High |
| 22165 | 11852 | 10488 | 17829 | 18772 | High | High | High | High | High | High | High | High | High | High |
| 21742 | 6648 | 3720 | 14483 | 10527 | High | High | High | High | High | High | High | High | High | High |
| 19068 | 4635 | 3975 | 4448 | 7016 | High | High | High | High | High | High | High | High | High | High |
| 9372 | 11207 | 9237 | 12169 | 14023 | High | High | High | High | High | High | High | High | High | High |
| 9133 | 2712 | 2405 | 3085 | 3102 | High | High | High | High | High | High | High | High | High | High |
| 15032 | 1878 | 1557 | 1817 | 2383 | High | High | High | High | High | High | High | High | High | High |
| 8304 | 9863 | 8661 | 20361 | 12739 | High | High | High | High | High | High | High | High | High | High |
| 10093 | 12444 | 9491 | 13143 | 14006 | High | High | High | High | High | High | High | High | High | High |
| 5714 | 6227 | 3842 | 7919 | 6735 | High | High | High | High | High | High | High | High | High | High |
| 16021 | 11627 | 10089 | 14020 | 12531 | High | High | High | High | High | High | High | High | High | High |
| 23184 | 10241 | 6630 | 9527 | 8094 | High | High | High | High | High | High | High | High | High | High |
| 7887 | 11663 | 10093 | 13210 | 15781 | High | High | High | High | High | High | High | High | High | High |
| 6164 | 1220 | 780 | 941 | 885 | High | High | High | High | High | High | High | High | High | High |
| 18233 | 2275 | 1826 | 1798 | 2165 | High | High | High | High | High | High | High | High | High | High |
| 8305 | 11282 | 9221 | 11259 | 12470 | High | High | High | High | High | High | High | High | High | High |
| 14772 | 7896 | 6915 | 8989 | 10476 | High | High | High | High | High | High | High | High | High | High |
| 10303 | 12390 | 10152 | 12865 | 16431 | High | High | High | High | High | High | High | High | High | High |
| 10833 | 12291 | 6964 | 25340 | 17817 | High | High | High | High | High | High | High | High | High | High |
| 6618 | 1583 | 1313 | 1904 | 1961 | High | High | High | High | High | High | High | High | High | High |
| 3154 | 6340 | 4235 | 12996 | 7801 | High | High | High | High | High | High | High | High | High | High |
| 18994 | 13608 | 9656 | 16930 | 15446 | High | High | High | High | High | High | High | High | High | High |
| 9855 | 5981 | 4748 | 6635 | 7805 | High | High | High | High | High | High | High | High | High | High |
| 5443 | 10428 | 7631 | 14244 | 11758 | High | High | High | High | High | High | High | High | High | High |
| 16532 | 7995 | 7380 | 7203 | 9694 | High | High | High | High | High | High | High | High | High | High |
| 8925 | 4425 | 3935 | 5559 | 6614 | High | High | High | High | High | High | High | High | High | High |
| 9156 | 3361 | 2229 | 3796 | 3265 | High | High | High | High | High | High | High | High | High | High |
| 8549 | 5814 | 4041 | 7748 | 6112 | High | High | High | High | High | High | High | High | High | High |
| 15950 | 12449 | 8204 | 17970 | 14607 | High | High | High | High | High | High | High | High | High | High |

| | | | | | | | | | | | | | | |
|---|---|---|---|---|---|---|---|---|---|---|---|---|---|---|
| 5028 | 13458 | 10066 | 20262 | 19147 | High | High | High | High | High | High | High | High | High | High |
| 10659 | 1823 | 1242 | 2392 | 2108 | High | High | High | High | High | High | High | High | High | High |
| 15681 | 12398 | 9654 | 10620 | 15574 | High | High | High | High | High | High | High | High | High | High |
| 7441 | 11278 | 7201 | 10718 | 9440 | High | High | High | High | High | High | High | High | High | High |
| 10630 | 12779 | 6491 | 20389 | 9787 | High | High | High | High | High | High | High | High | High | High |
| 12712 | 4545 | 2699 | 3409 | 4313 | High | High | High | High | High | High | High | High | High | High |
| 8707 | 4311 | 4744 | 7409 | 6159 | High | High | High | High | High | High | High | High | High | High |
| 6490 | 11267 | 6440 | 20012 | 12880 | High | High | High | High | High | High | High | High | High | High |
| 3040 | 1939 | 1834 | 1803 | 2117 | High | High | High | High | High | High | High | High | High | High |
| 14899 | 5711 | 5150 | 6444 | 8723 | High | High | High | High | High | High | High | High | High | High |
| 11759 | 4250 | 3742 | 6080 | 6693 | High | High | High | High | High | High | High | High | High | High |
| 10644 | 8655 | 6905 | 10547 | 13198 | High | High | High | High | High | High | High | High | High | High |
| 4696 | 4983 | 3941 | 6514 | 6312 | High | High | High | High | High | High | High | High | High | High |
| 8316 | 6558 | 6841 | 6549 | 9928 | High | High | High | High | High | High | High | High | High | High |
| 21645 | 7646 | 7248 | 8559 | 11836 | High | High | High | High | High | High | High | High | High | High |
| 8686 | 10306 | 7449 | 10189 | 9442 | High | High | High | High | High | High | High | High | High | High |
| 11261 | 7057 | 5193 | 7887 | 7758 | High | High | High | High | High | High | High | High | High | High |
| 21999 | 18275 | 13987 | 17179 | 17725 | High | High | High | High | High | High | High | High | High | High |
| 4510 | 9472 | 6633 | 11439 | 11646 | High | High | High | High | High | High | High | High | High | High |
| 20372 | 17211 | 14233 | 18681 | 21687 | High | High | High | High | High | High | High | High | High | High |
| 15008 | 14448 | 11370 | 24588 | 17057 | High | High | High | High | High | High | High | High | High | High |
| 9022 | 17757 | 11341 | 24644 | 20109 | High | High | High | High | High | High | High | High | High | High |
| 8329 | 6466 | 5862 | 5488 | 7649 | High | High | High | High | High | High | High | High | High | High |
| 11181 | 6276 | 3148 | 6925 | 4753 | High | High | High | High | High | High | High | High | High | High |
| 14526 | 14189 | 9420 | 17286 | 15611 | High | High | High | High | High | High | High | High | High | High |
| 5283 | 4565 | 2838 | 7359 | 5750 | High | High | High | High | High | High | High | High | High | High |
| 8136 | 11203 | 8228 | 13961 | 12830 | High | High | High | High | High | High | High | High | High | High |
| 12361 | 9100 | 7596 | 8868 | 9875 | High | High | High | High | High | High | High | High | High | High |
| 12730 | 4871 | 3382 | 7783 | 7519 | High | High | High | High | High | High | High | High | High | High |
| 16275 | 20760 | 14492 | 25980 | 26337 | High | High | High | High | High | High | High | High | High | High |
| 4444 | 8246 | 6677 | 8291 | 8157 | High | High | High | High | High | High | High | High | High | High |

| | | | | | | | | | | | | | | |
|---|---|---|---|---|---|---|---|---|---|---|---|---|---|---|
| 11274 | 10429 | 9567 | 9515 | 13221 | High | High | High | High | High | High | High | High | High | High |
| 5306 | 966 | 744 | 1311 | 1147 | High | High | High | High | High | High | High | High | High | High |
| 9468 | 6449 | 3915 | 6046 | 4978 | High | High | High | High | High | High | High | High | High | High |
| 840 | 6445 | 4673 | 8065 | 8419 | High | High | High | High | High | High | High | High | High | High |
| 7788 | 11980 | 8790 | 15365 | 15676 | High | High | High | High | High | High | High | High | High | High |
| 9302 | 8116 | 6393 | 7778 | 8286 | High | High | High | High | High | High | High | High | High | High |
| 5150 | 8095 | 3223 | 7479 | 5388 | High | High | High | High | High | High | High | High | High | High |
| 11862 | 9778 | 6249 | 15274 | 12443 | High | High | High | High | High | High | High | High | High | High |
| 18055 | 5399 | 4097 | 5226 | 6092 | High | High | High | High | High | High | High | High | High | High |
| 9068 | 18467 | 13118 | 23761 | 25385 | High | High | High | High | High | High | High | High | High | High |
| 8038 | 6400 | 5471 | 8335 | 9747 | High | High | High | High | High | High | High | High | High | High |
| 8750 | 7013 | 4875 | 9273 | 6975 | High | High | High | High | High | High | High | High | High | High |
| 3378 | 6898 | 3000 | 11360 | 7249 | High | High | High | High | High | High | High | High | High | High |
| 8679 | 8865 | 6842 | 11825 | 10040 | High | High | High | High | High | High | High | High | High | High |
| 14439 | 6162 | 4446 | 5235 | 5908 | High | High | High | High | High | High | High | High | High | High |
| 15335 | 11483 | 8942 | 8931 | 11656 | High | High | High | High | High | High | High | High | High | High |
| 12560 | 8662 | 8734 | 9918 | 13167 | High | High | High | High | High | High | High | High | High | High |
| 5798 | 1788 | 1414 | 1462 | 1779 | High | High | High | High | High | High | High | High | High | High |
| 8837 | 3159 | 2187 | 3129 | 4251 | High | High | High | High | High | High | High | High | High | High |
| 9857 | 4288 | 2642 | 5873 | 4866 | High | High | High | High | High | High | High | High | High | High |
| 3889 | 2610 | 2053 | 5082 | 4231 | High | High | High | High | High | High | High | High | High | High |
| 10743 | 6770 | 6098 | 7607 | 8422 | High | High | High | High | High | High | High | High | High | High |
| 3734 | 3482 | 2649 | 4257 | 3781 | High | High | High | High | High | High | High | High | High | High |
| 13790 | 4539 | 4611 | 5132 | 5791 | High | High | High | High | High | High | High | High | High | High |
| 8644 | 2613 | 2357 | 2804 | 3075 | High | High | High | High | High | High | High | High | High | High |
| 6001 | 2984 | 2488 | 4905 | 4148 | High | High | High | High | High | High | High | High | High | High |
| 11785 | 5829 | 3653 | 19823 | 12788 | High | High | High | High | High | High | High | High | High | High |
| 10018 | 9008 | 4605 | 13015 | 10103 | High | High | High | High | High | High | High | High | High | High |
| 11249 | 6190 | 4903 | 6351 | 6805 | High | High | High | High | High | High | High | High | High | High |
| 4391 | 7438 | 6354 | 8785 | 7584 | High | High | High | High | High | High | High | High | High | High |
| 9844 | 4982 | 3765 | 4909 | 4805 | High | High | High | High | High | High | High | High | High | High |

| | | | | | | | | | | | | | | |
|---|---|---|---|---|---|---|---|---|---|---|---|---|---|---|
| 1440 | 2701 | 1672 | 4254 | 2298 | High | High | High | High | High | High | High | High | High | High |
| 7917 | 4191 | 2183 | 5493 | 3594 | High | High | High | High | High | High | High | High | High | High |
| 7374 | 8238 | 6700 | 7168 | 8369 | High | High | High | High | High | High | High | High | High | High |
| 8404 | 8382 | 4918 | 6594 | 5879 | High | High | High | High | High | High | High | High | High | High |
| 11102 | 6989 | 4725 | 14549 | 10191 | High | High | High | High | High | High | High | High | High | High |
| 6299 | 6551 | 4516 | 10983 | 7485 | High | High | High | High | High | High | High | High | High | High |
| 7052 | 4199 | 4205 | 3756 | 5342 | High | High | High | High | High | High | High | High | High | High |
| 759 | 4005 | 2626 | 2234 | 1345 | High | High | High | High | High | High | High | High | High | High |
| 4789 | 16089 | 12905 | 17631 | 18126 | High | High | High | High | High | High | High | High | High | High |
| 3366 | 5059 | 1664 | 7431 | 3556 | High | High | High | High | High | High | High | High | High | High |
| 2622 | 6030 | 3871 | 8815 | 6540 | High | High | High | High | High | High | High | High | High | High |
| 9250 | 2308 | 2296 | 2806 | 2756 | High | High | High | High | High | High | High | High | High | High |
| 5771 | 4575 | 3634 | 5638 | 6399 | High | High | High | High | High | High | High | High | High | High |
| 2660 | 7457 | 4468 | 8369 | 7113 | High | High | High | High | High | High | High | High | High | High |
| 5959 | 1886 | 1442 | 2733 | 2131 | High | High | High | High | High | High | High | High | High | High |
| 13983 | 7975 | 7098 | 10504 | 10885 | High | High | High | High | High | High | High | High | High | High |
| 17607 | 8396 | 6411 | 15809 | 11084 | High | High | High | High | High | High | High | High | High | High |
| 14022 | 1640 | 1981 | 1891 | 2966 | High | High | High | High | High | High | High | High | High | High |
| 9469 | 6168 | 3855 | 10630 | 7663 | High | High | High | High | High | High | High | High | High | High |
| 15307 | 23760 | 20044 | 35140 | 36786 | High | High | High | High | High | High | High | High | High | High |
| 3885 | 3603 | 2308 | 4457 | 4012 | High | High | High | High | High | High | High | High | High | High |
| 8110 | 12565 | 11581 | 16168 | 17809 | High | High | High | High | High | High | High | High | High | High |
| 2937 | 15782 | 13633 | 18757 | 19460 | High | High | High | High | High | High | High | High | High | High |
| 13995 | 8870 | 9694 | 9125 | 13347 | High | High | High | High | High | High | High | High | High | High |
| 10773 | 11695 | 8039 | 11913 | 12166 | High | High | High | High | High | High | High | High | High | High |
| 9934 | | | | | High | High | High | High | High | High | High | High | High | High |
| 5113 | 1614 | 1350 | 1813 | 2223 | High | High | High | High | High | High | High | High | High | High |
| 18751 | 13304 | 9106 | 17789 | 16748 | High | High | High | High | High | High | High | High | High | High |
| 10002 | 10289 | 10359 | 13667 | 19422 | High | High | High | High | High | High | High | High | High | High |
| 500 | 1130 | 845 | 1674 | 1608 | Not Founc | Not Founc | Not Founc | Not Founc | High | High | High | High | High | High |
| 7699 | 13280 | 9326 | 17937 | 15617 | High | High | High | High | High | High | High | High | High | High |

| | | | | | | | | | | | | | | |
|---|---|---|---|---|---|---|---|---|---|---|---|---|---|---|
| 14252 | 7071 | 5493 | 8117 | 9280 | High | High | High | High | High | High | High | High | High | High |
| 9618 | 7205 | 5335 | 16795 | 14263 | High | High | High | High | High | High | High | High | High | High |
| 4812 | 8610 | 11148 | 8961 | 8988 | High | High | High | High | High | High | High | High | High | High |
| 12473 | 10072 | 10977 | 8873 | 11907 | High | High | High | High | High | High | High | High | High | High |
| 3510 | 9008 | 9195 | 8495 | 14070 | High | High | High | High | High | High | High | High | High | High |
| 8949 | 6163 | 4404 | 7165 | 6150 | High | High | High | High | High | High | High | High | High | High |
| 6624 | 5969 | 3749 | 9962 | 8138 | High | High | High | High | High | High | High | High | High | High |
| 11889 | 287 | 472 | 1379 | 774 | High | High | High | High | High | High | High | High | High | High |
| 4691 | 9777 | 8205 | 10489 | 10824 | High | High | High | High | High | High | High | High | High | High |
| 6119 | 5866 | 4276 | 6520 | 7993 | High | High | High | High | High | High | High | High | High | High |
| 838 | 3297 | 2851 | 3102 | 4070 | High | High | High | High | High | High | High | High | High | High |
| 11111 | 3304 | 2392 | 4257 | 3745 | High | High | High | High | High | High | High | High | High | High |
| 1617 | 10206 | 7115 | 9639 | 9190 | High | High | High | High | High | High | High | High | High | High |
| 7140 | 7090 | 4751 | 9790 | 8008 | High | High | High | High | High | High | High | High | High | High |
| 6180 | 3827 | 2403 | 10487 | 4964 | High | High | High | High | High | High | High | High | High | High |
| 12229 | | | | | High | High | High | High | High | High | High | High | Not Founc | Not Founc |
| 3471 | 6136 | 3915 | 7903 | 6333 | High | High | High | High | High | High | High | High | High | High |
| 14205 | 6055 | 5307 | 6823 | 24648 | High | High | High | High | High | High | High | High | High | High |
| 8831 | 8151 | 6240 | 11731 | 9408 | High | High | High | High | High | High | High | High | High | High |
| 5404 | 2455 | 2218 | 2338 | 3897 | High | High | High | High | High | High | High | High | High | High |
| 4216 | 2049 | 1490 | 2372 | 2215 | High | High | High | High | High | High | High | High | High | High |
| 10652 | 4829 | 4304 | 7111 | 7542 | High | High | High | High | High | High | High | High | High | High |
| 7242 | 8514 | 3992 | 9951 | 5569 | High | High | High | High | High | High | High | High | High | High |
| 5754 | 8626 | 6402 | 10244 | 9562 | High | High | High | High | High | High | High | High | High | High |
| 10348 | 15062 | 9726 | 22001 | 19172 | High | High | High | High | High | High | High | High | High | High |
| 3141 | 10295 | 7235 | 19818 | 12946 | High | High | High | High | High | High | High | High | High | High |
| 3163 | 10841 | 7936 | 11315 | 12117 | High | High | High | High | High | High | High | High | High | High |
| 9075 | 7483 | 10597 | 23142 | 16315 | High | High | High | High | High | High | High | High | High | High |
| 7053 | 2436 | 1911 | 2433 | 2527 | High | High | High | High | High | High | High | High | High | High |
| 2768 | 6808 | 7148 | 16022 | 14428 | High | High | High | High | High | High | High | High | High | High |
| 13282 | 10427 | 9329 | 11284 | 10480 | High | High | High | High | High | High | High | High | High | High |

| | | | | | | | | | | | | | | |
|---|---|---|---|---|---|---|---|---|---|---|---|---|---|---|
| 12681 | 11976 | 5225 | 11387 | 9861 | High | High | High | High | High | High | High | High | High | High |
| 8220 | 782 | 588 | 802 | 877 | High | High | High | High | High | High | High | High | High | High |
| 4144 | 3486 | 2625 | 4032 | 4707 | High | High | High | High | High | High | High | High | High | High |
| 4187 | 9881 | 8565 | 10425 | 11662 | High | High | High | High | High | High | High | High | High | High |
| 4627 | 7076 | 5475 | 9335 | 9776 | High | High | High | High | High | High | High | High | High | High |
| 9802 | 16682 | 14389 | 19927 | 19648 | High | High | High | High | High | High | High | High | High | High |
| 24166 | 13033 | 7188 | 14477 | 11925 | High | High | High | High | High | High | High | High | High | High |
| 3275 | 7014 | 6029 | 6168 | 7100 | High | High | High | High | High | High | High | High | High | High |
| 2145 | 10919 | 7948 | 11406 | 10380 | High | High | High | High | High | High | High | High | High | High |
| 7064 | 5609 | 5466 | 6527 | 7042 | High | High | High | High | High | High | High | High | High | High |
| 7144 | 8478 | 5779 | 9721 | 8243 | High | High | High | High | High | High | High | High | High | High |
| 4421 | | | | | High | High | High | High | High | High | High | High | Not Founc | Not Founc |
| 7988 | 9051 | 8467 | 10592 | 11976 | High | High | High | High | High | High | High | High | High | High |
| 1927 | 6280 | 4911 | 6703 | 6876 | High | High | High | High | High | High | High | High | High | High |
| 4871 | 6069 | 4311 | 8945 | 7226 | High | High | High | High | High | High | High | High | High | High |
| 5763 | 1759 | 1634 | 1855 | 2262 | High | High | High | High | High | High | High | High | High | High |
| 6444 | 6957 | 4963 | 7646 | 8161 | High | High | High | High | High | High | High | High | High | High |
| 8602 | 6010 | 4958 | 6625 | 6679 | High | High | High | High | High | High | High | High | High | High |
| 11938 | 3083 | 2499 | 3461 | 3350 | High | High | High | High | High | High | High | High | High | High |
| 5352 | 4081 | 2653 | 6227 | 4330 | High | High | High | High | High | High | High | High | High | High |
| 8440 | 2898 | 2581 | 2965 | 3480 | High | High | High | High | High | High | High | High | High | High |
| 4523 | 2099 | 1119 | 3335 | 1916 | Not Founc | Not Founc | Not Founc | Not Founc | High | High | High | High | High | High |
| 2548 | 3692 | 3520 | 2990 | 3868 | High | High | High | High | High | High | High | High | High | High |
| 6980 | 5583 | 3935 | 7044 | 6549 | High | High | High | High | High | High | High | High | High | High |
| 14979 | 5550 | 3943 | 10700 | 8618 | High | High | High | High | High | High | High | High | High | High |
| 16387 | 3927 | 3237 | 5521 | 5339 | High | High | High | High | High | High | High | High | High | High |
| 1952 | 3206 | 3534 | 3454 | 5347 | High | High | High | High | High | High | High | High | High | High |
| 3067 | 16986 | 10327 | 18411 | 14761 | High | High | High | High | High | High | High | High | High | High |
| 3924 | 12220 | 10736 | 12313 | 15101 | High | High | High | High | High | High | High | High | High | High |
| 5926 | 11697 | 7930 | 15698 | 15592 | High | High | High | High | High | High | High | High | High | High |
| 1986 | 16869 | 12044 | 17936 | 16560 | High | High | High | High | High | High | High | High | High | High |

| | | | | | | | | | | | | | | |
|---|---|---|---|---|---|---|---|---|---|---|---|---|---|---|
| 5462 | 2317 | 2204 | 2663 | 2871 | High | High | High | High | High | High | High | High | High | High |
| 12058 | 8217 | 5642 | 9340 | 9545 | High | High | High | High | High | High | High | High | High | High |
| 12841 | 8094 | 5561 | 13900 | 11685 | High | High | High | High | High | High | High | High | High | High |
| 9544 | 2649 | 1798 | 2640 | 2417 | High | High | High | High | High | High | High | High | High | High |
| 7628 | 5191 | 3560 | 7571 | 4469 | High | High | High | High | High | High | High | High | High | High |
| 5810 | 6894 | 2703 | 9141 | 5410 | High | High | High | High | High | High | High | High | High | High |
| 4526 | 7918 | 6521 | 7219 | 6951 | High | High | High | High | High | High | High | High | High | High |
| 7289 | 16136 | 11900 | 19041 | 18526 | High | High | High | High | High | High | High | High | High | High |
| 13410 | 7755 | 4544 | 10875 | 6940 | High | High | High | High | High | High | High | High | High | High |
| 11543 | 18411 | 10323 | 17997 | 16475 | High | High | High | High | High | High | High | High | High | High |
| 13143 | 9043 | 5666 | 16598 | 8937 | High | High | High | High | High | High | High | High | High | High |
| 16355 | 18831 | 16186 | 23123 | 24278 | High | High | High | High | High | High | High | High | High | High |
| 17540 | 4677 | 4045 | 5891 | 6334 | High | High | High | High | High | High | High | High | High | High |
| 4320 | 12723 | 9692 | 18906 | 15186 | High | High | High | High | High | High | High | High | High | High |
| 7808 | 19964 | 16679 | 28650 | 33191 | High | High | High | High | High | High | High | High | High | High |
| 4692 | 8941 | 4777 | 8317 | 8645 | High | High | High | High | High | High | High | High | High | High |
| 16363 | 10419 | 7321 | 11855 | 12075 | High | High | High | High | High | High | High | High | High | High |
| 10938 | 5981 | 4561 | 4810 | 5166 | High | High | High | High | High | High | High | High | High | High |
| 12257 | 6045 | 6794 | 6506 | 11159 | High | High | High | High | High | High | High | High | High | High |
| 4471 | 1789 | 1562 | 2200 | 2194 | High | High | High | High | High | High | High | High | High | High |
| 7451 | 2065 | 1550 | 3213 | 2835 | High | High | High | High | High | High | High | High | High | High |
| 16333 | 10452 | 10391 | 10950 | 15188 | High | High | High | High | High | High | High | High | High | High |
| 6599 | 22069 | 4324 | 18496 | 8118 | High | High | High | High | High | High | High | High | High | High |
| 3959 | 6359 | 8856 | 8124 | 6808 | High | High | High | High | High | High | High | High | High | High |
| 12703 | 1741 | 1479 | 2612 | 2384 | High | High | High | High | High | High | High | High | High | High |
| 12606 | 4757 | 4371 | 6362 | 7367 | High | High | High | High | High | High | High | High | High | High |
| 2971 | 11457 | 9485 | 8942 | 10214 | High | High | High | High | High | High | High | High | High | High |
| 7024 | 8770 | 7043 | 9789 | 12113 | High | High | High | High | High | High | High | High | High | High |
| 14750 | 5139 | 3594 | 5896 | 5645 | High | High | High | High | High | High | High | High | High | High |
| 8820 | 3805 | 4365 | 4323 | 7305 | High | High | High | High | High | High | High | High | High | High |
| 1822 | 2213 | 1709 | 2114 | 2295 | High | High | High | High | High | High | High | High | High | High |

| | | | | | | | | | | | | | | |
|---|---|---|---|---|---|---|---|---|---|---|---|---|---|---|
| 7194 | 5772 | 4371 | 6983 | 6815 | High | High | High | High | High | High | High | High | High | High |
| 15885 | 3832 | 3668 | 4149 | 5636 | High | High | High | High | High | High | High | High | High | High |
| 10773 | 7702 | 6786 | 13764 | 10036 | High | High | High | High | High | High | High | High | High | High |
| 4151 | 9753 | 3537 | 8992 | 5607 | High | High | High | High | High | High | High | High | High | High |
| 4844 | 5887 | 3641 | 7937 | 6244 | High | High | High | High | High | High | High | High | High | High |
| 4810 | 1526 | 944 | 1670 | 1533 | High | High | High | High | High | High | High | High | High | High |
| 4662 | 4392 | 3234 | 6036 | 4656 | High | High | High | High | High | High | High | High | High | High |
| 7350 | 14827 | 10480 | 18728 | 17089 | High | High | High | High | High | High | High | High | High | High |
| 9024 | 6841 | 6173 | 5662 | 8081 | High | High | High | High | High | High | High | High | High | High |
| 32084 | 3414 | 3262 | 3446 | 19325 | High | High | High | High | High | High | High | High | High | High |
| 8036 | 1138 | 1004 | 882 | 1189 | High | High | High | High | High | High | High | High | High | High |
| 3988 | 2371 | 1830 | 3245 | 2855 | High | High | High | High | High | High | High | High | High | High |
| 5905 | 4465 | 3443 | 6669 | 5807 | High | High | High | High | High | High | High | High | High | High |
| 11803 | 10351 | 7659 | 16185 | 14192 | High | High | High | High | High | High | High | High | High | High |
| 8438 | 3160 | 2368 | 3228 | 3330 | High | High | High | High | High | High | High | High | High | High |
| 11342 | 9570 | 5944 | 7849 | 7729 | High | High | High | High | High | High | High | High | High | High |
| 10389 | 4938 | 7538 | 4317 | 10679 | High | High | High | High | High | High | High | High | High | High |
| 4236 | 5069 | 4042 | 5539 | 6311 | High | High | High | High | High | High | High | High | High | High |
| 3935 | 2354 | 1639 | 3203 | 2936 | High | High | High | High | High | High | High | High | High | High |
| 11456 | 6614 | 5045 | 5903 | 5890 | High | High | High | High | High | High | High | High | High | High |
| 591 | 2709 | 1705 | 2503 | 1997 | High | High | High | High | High | High | High | High | High | High |
| 4094 | 6948 | 6799 | 8810 | 11585 | High | High | High | High | High | High | High | High | High | High |
| 8404 | 9079 | 7183 | 9074 | 10219 | High | High | High | High | High | High | High | High | High | High |
| 7428 | 10819 | 10071 | 10351 | 13739 | High | High | High | High | High | High | High | High | High | High |
| 1731 | 5793 | 4453 | 6675 | 7132 | High | High | High | High | High | High | High | High | High | High |
| 3148 | 5011 | 3033 | 6494 | 4882 | High | High | High | High | High | High | High | High | High | High |
| 10853 | 1485 | 1322 | 2001 | 2106 | High | High | High | High | High | High | High | High | High | High |
| 7711 | 10211 | 7115 | 12697 | 10619 | High | High | High | High | High | High | High | High | High | High |
| 5259 | 1897 | 1554 | 2449 | 2112 | High | High | High | High | High | High | High | High | High | High |
| 9862 | 7591 | 5705 | 7462 | 9062 | High | High | High | High | High | High | High | High | High | High |
| 3494 | 8087 | 6328 | 8533 | 8994 | High | High | High | High | High | High | High | High | High | High |

| | | | | | | | | | | | | | | |
|---|---|---|---|---|---|---|---|---|---|---|---|---|---|---|
| 6555 | 7137 | 6825 | 9292 | 8842 | High | High | High | High | High | High | High | High | High | High |
| 2813 | | | | | Not Found | Not Found | Not Found | Not Found | High | High | High | High | Not Found | Not Found |
| 8233 | 8850 | 5755 | 17317 | 10990 | High | High | High | High | High | High | High | High | High | High |
| 7095 | 2273 | 2159 | 2242 | 3209 | High | High | High | High | High | High | High | High | High | High |
| 4444 | 8935 | 5496 | 7117 | 6844 | High | High | High | High | High | High | High | High | High | High |
| 7586 | 5646 | 5144 | 14831 | 9468 | High | High | High | High | High | High | High | High | High | High |
| 15349 | 2205 | 1718 | 2777 | 2765 | High | High | High | High | High | High | High | High | High | High |
| 3391 | 1990 | 1638 | 3139 | 2321 | High | High | High | High | High | High | High | High | High | High |
| 5911 | 6202 | 5512 | 5222 | 7069 | High | High | High | High | High | High | High | High | High | High |
| 7437 | 9416 | 7623 | 24511 | 22887 | High | High | High | High | High | High | High | High | High | High |
| 6267 | 4612 | 3278 | 5885 | 5141 | High | High | High | High | High | High | High | High | High | High |
| 3834 | 18266 | 9751 | 23011 | 17507 | High | High | High | High | High | High | High | High | High | High |
| 9805 | 11545 | 10037 | 8877 | 12228 | High | High | High | High | High | High | High | High | High | High |
| 7434 | 16273 | 16141 | 30247 | 27602 | High | High | High | High | High | High | High | High | High | High |
| 3118 | 9487 | 7334 | 14660 | 11718 | High | High | High | High | High | High | High | High | High | High |
| 1939 | 1090 | 1032 | 1303 | 1353 | High | High | High | High | High | High | High | High | High | High |
| 6756 | 2461 | 1958 | 6473 | 4087 | High | High | High | High | High | High | High | High | High | High |
| 3956 | 5752 | 4390 | 6426 | 7065 | High | High | High | High | High | High | High | High | High | High |
| 2593 | 11268 | 8943 | 13077 | 11721 | High | High | High | High | High | High | High | High | High | High |
| 4930 | 9726 | 8357 | 11468 | 13317 | High | High | High | High | High | High | High | High | High | High |
| 8298 | 2582 | 3069 | 7528 | 4347 | High | High | High | High | High | High | High | High | High | High |
| 6451 | | | | | High | High | High | High | High | High | High | High | Not Found | Not Found |
| 3714 | 3694 | 2224 | 6186 | 3383 | High | High | High | High | High | High | High | High | High | High |
| 5437 | 7970 | 5044 | 10528 | 7193 | High | High | High | High | High | High | High | High | High | High |
| 4270 | 7546 | 6469 | 7224 | 7978 | High | High | High | High | High | High | High | High | High | High |
| 9790 | 1860 | 1316 | 1634 | 1709 | High | High | High | High | High | High | High | High | High | High |
| 3190 | 1510 | 1189 | 1380 | 1715 | High | High | High | High | High | High | High | High | High | High |
| 8181 | 6103 | 4507 | 8626 | 7592 | High | High | High | High | High | High | High | High | High | High |
| 9055 | 8880 | 6449 | 11485 | 10597 | Not Found | Not Found | Not Found | Not Found | High | High | High | High | High | High |
| 7850 | 2574 | 2285 | 2053 | 3835 | High | High | High | High | High | High | High | High | High | High |
| 16925 | 17741 | 10485 | 24011 | 17761 | High | High | High | High | High | High | High | High | High | High |

| | | | | | | | | | | | | | | |
|---|---|---|---|---|---|---|---|---|---|---|---|---|---|---|
| 4993 | 4986 | 3787 | 6067 | 6905 | High | High | High | High | High | High | High | High | High | High |
| 2908 | 3318 | 1924 | 8278 | 5167 | High | High | High | High | High | High | High | High | High | High |
| 7198 | 4770 | 3148 | 8851 | 6161 | High | High | High | High | High | High | High | High | High | High |
| 2879 | 3001 | 2017 | 3195 | 3364 | High | High | High | High | High | High | High | High | High | High |
| 12234 | 14943 | 12118 | 15835 | 17936 | Not Founc | Not Founc | Not Founc | Not Founc | High | High | High | High | High | High |
| 5547 | 13531 | 5365 | 15019 | 8471 | High | High | High | High | High | High | High | High | High | High |
| 4138 | 686 | 635 | 1044 | 1296 | High | High | High | High | High | High | High | High | High | High |
| 3661 | 7920 | 5190 | 9150 | 9379 | High | High | High | High | High | High | High | High | High | High |
| 17903 | 7300 | 5515 | 10131 | 10186 | High | High | High | High | High | High | High | High | High | High |
| 3542 | 2133 | 2245 | 2394 | 2813 | High | High | High | High | High | High | High | High | High | High |
| 5728 | 2672 | 2274 | 2165 | 2643 | High | High | High | High | High | High | High | High | High | High |
| 2505 | 10453 | 7898 | 11926 | 11823 | High | High | High | High | High | High | High | High | High | High |
| 2446 | 5650 | 4511 | 5343 | 5553 | High | High | High | High | High | High | High | High | High | High |
| 2633 | 1806 | 1545 | 1556 | 2242 | High | High | High | High | High | High | High | High | High | High |
| | | | | | High | High | High | High | Not Founc | Not Founc | Not Founc | Not Founc | High | High |
| 3390 | 4671 | 3472 | 3658 | 4363 | High | High | High | High | High | High | High | High | High | High |
| 12287 | 15466 | 9142 | 16225 | 15015 | High | High | High | High | High | High | High | High | High | High |
| 12914 | 8127 | 3804 | 17762 | 11198 | High | High | High | High | High | High | High | High | High | High |
| 6760 | 6126 | 3904 | 9589 | 7520 | High | High | High | High | High | High | High | High | High | High |
| 3658 | 3617 | 2489 | 3548 | 3572 | High | High | High | High | High | High | High | High | High | High |
| 6516 | 9730 | 6654 | 12756 | 11546 | High | High | High | High | High | High | High | High | High | High |
| 5597 | 3242 | 2076 | 4116 | 3573 | High | High | High | High | High | High | High | High | High | High |
| 6593 | 2776 | 1949 | 3981 | 3157 | High | High | High | High | High | High | High | High | High | High |
| 8823 | 7800 | 6263 | 8569 | 10685 | High | High | High | High | High | High | High | High | High | High |
| 7269 | 5308 | 4429 | 9109 | 7614 | High | High | High | High | High | High | High | High | High | High |
| 2479 | 4060 | 2899 | 10414 | 6977 | High | High | High | High | High | High | High | High | High | High |
| 2758 | 6705 | 5219 | 12169 | 8075 | High | High | High | High | High | High | High | High | High | High |
| 6993 | 10132 | 9735 | 11520 | 15314 | High | High | High | High | High | High | High | High | High | High |
| 7479 | 10103 | 7956 | 14531 | 14737 | High | High | High | High | High | High | High | High | High | High |
| 5351 | 7393 | 6057 | 8181 | 9169 | High | High | High | High | High | High | High | High | High | High |
| 5129 | 16838 | 7700 | 35259 | 24845 | High | High | High | High | High | High | High | High | High | High |

| | | | | | | | | | | | | | | |
|---|---|---|---|---|---|---|---|---|---|---|---|---|---|---|
| 64 | 759 | 820 | 889 | 835 | Not Found | Not Found | Not Found | Not Found | High | High | High | High | High | High |
| 5700 | 6510 | 4714 | 5327 | 6622 | High | High | High | High | High | High | High | High | High | High |
| 1334 | 6143 | 5003 | 5899 | 7922 | High | High | High | High | High | High | High | High | High | High |
| 8804 | 12382 | 9681 | 11926 | 13727 | High | High | High | High | High | High | High | High | High | High |
| 9146 | 3425 | 2144 | 7901 | 4853 | High | High | High | High | High | High | High | High | High | High |
| 10955 | 4429 | 3648 | 5772 | 7176 | High | High | High | High | High | High | High | High | High | High |
| 1837 | 1531 | 998 | 1751 | 1682 | High | High | High | High | High | High | High | High | High | High |
| 14921 | 2895 | 2199 | 3347 | 3110 | High | High | High | High | High | High | High | High | High | High |
| 7913 | 4311 | 4065 | 7215 | 5268 | High | High | High | High | High | High | High | High | High | High |
| 3178 | 6989 | 5717 | 8287 | 9373 | High | High | High | High | High | High | High | High | High | High |
| 7105 | 2027 | 1327 | 3169 | 2693 | High | High | High | High | High | High | High | High | High | High |
| 8699 | 10521 | 5337 | 15600 | 8925 | High | High | High | High | High | High | High | High | High | High |
| 7081 | | | | | High | High | High | High | High | High | High | High | Not Found | Not Found |
| 18737 | 2533 | 3793 | 4175 | 4519 | High | High | High | High | High | High | High | High | High | High |
| 7070 | 4229 | 3231 | 4883 | 5331 | High | High | High | High | High | High | High | High | High | High |
| 3950 | 5141 | 5230 | 5174 | 7714 | High | High | High | High | High | High | High | High | High | High |
| 7391 | 2798 | 2503 | 3392 | 3072 | High | High | High | High | High | High | High | High | High | High |
| 6015 | 4405 | 3190 | 4351 | 5470 | High | High | High | High | High | High | High | High | High | High |
| 20509 | 9593 | 8907 | 17884 | 15401 | High | High | High | High | High | High | High | High | High | High |
| 2128 | 3363 | 2729 | 4724 | 4044 | High | High | High | High | High | High | High | High | High | High |
| 4008 | 5680 | 4044 | 7914 | 6468 | High | High | High | High | High | High | High | High | High | High |
| 7132 | 9666 | 9473 | 12560 | 14811 | High | High | High | High | High | High | High | High | High | High |
| 15411 | 9058 | 7436 | 9364 | 11846 | Not Found | Not Found | Not Found | Not Found | High | High | High | High | High | High |
| 11719 | 7012 | 4880 | 5680 | 76637 | High | High | High | High | High | High | High | High | High | High |
| | 280 | 223 | 346 | 341 | High | High | High | High | Not Found | Not Found | Not Found | Not Found | High | High |
| 16935 | 6129 | 5120 | 4950 | 6019 | High | High | High | High | High | High | High | High | High | High |
| 5939 | 2140 | 1882 | 4884 | 3383 | High | High | High | High | High | High | High | High | High | High |
| 5416 | 12521 | 13672 | 9175 | 11171 | High | High | High | High | High | High | High | High | High | High |
| 4637 | 3688 | 2351 | 3940 | 3705 | High | High | High | High | High | High | High | High | High | High |
| 4175 | 4276 | 2922 | 6522 | 6814 | High | High | High | High | High | High | High | High | High | High |
| 5504 | 6788 | 5622 | 8243 | 8583 | Not Found | Not Found | Not Found | Not Found | High | High | High | High | High | High |

| | | | | | | | | | | | | | | |
|---|---|---|---|---|---|---|---|---|---|---|---|---|---|---|
| 3996 | 625 | 479 | 1592 | 1194 | High | High | High | High | High | High | High | High | High | High |
| 10955 | 5549 | 4007 | 8821 | 7896 | High | High | High | High | High | High | High | High | High | High |
| 6936 | 6934 | 6072 | 10259 | 9913 | High | High | High | High | High | High | High | High | High | High |
| 13026 | 3180 | 2407 | 5549 | 5871 | High | High | High | High | High | High | High | High | High | High |
| 10908 | 9882 | 8325 | 10780 | 10005 | High | High | High | High | High | High | High | High | High | High |
| 7419 | 5217 | 4736 | 9439 | 5599 | High | High | High | High | High | High | High | High | High | High |
| 10381 | 1978 | 1578 | 2054 | 2274 | High | High | High | High | High | High | High | High | High | High |
| 3944 | 168 | 116 | 120 | 191 | High | High | High | High | High | High | High | High | High | High |
| 6379 | | | | | High | High | High | High | High | High | High | High | Not Founc | Not Founc |
| 348 | 526 | 491 | 446 | 519 | High | High | High | High | High | High | High | High | High | High |
| 3837 | 13067 | 10525 | 11086 | 16520 | High | High | High | High | High | High | High | High | High | High |
| 3920 | 5822 | 6331 | 5826 | 5659 | High | High | High | High | High | High | High | High | High | High |
| 9014 | 4147 | 3337 | 5461 | 5253 | High | High | High | High | High | High | High | High | High | High |
| 6030 | 8487 | 5460 | 7233 | 6374 | High | High | High | High | High | High | High | High | High | High |
| | 12576 | 7716 | 12489 | 11514 | High | High | High | High | Not Founc | Not Founc | Not Founc | Not Founc | High | High |
| 7946 | 3848 | 2688 | 7588 | 4862 | High | High | High | High | High | High | High | High | High | High |
| 2074 | 5152 | 4159 | 7375 | 6789 | High | High | High | High | High | High | High | High | High | High |
| 3447 | 3382 | 2600 | 3543 | 2809 | High | High | High | High | High | High | High | High | High | High |
| 5464 | 5361 | 4448 | 5841 | 6196 | High | High | High | High | High | High | High | High | High | High |
| 3145 | 1686 | 1865 | 4930 | 2628 | High | High | High | High | High | High | High | High | High | High |
| 4584 | 5503 | 4165 | 12073 | 8236 | High | High | High | High | High | High | High | High | High | High |
| 5655 | 3874 | 2334 | 4683 | 3952 | High | High | High | High | High | High | High | High | High | High |
| 8571 | 8634 | 10040 | 10091 | 13967 | High | High | High | High | High | High | High | High | High | High |
| 2984 | 2367 | 1529 | 2431 | 2560 | High | High | High | High | High | High | High | High | High | High |
| 2589 | 2290 | 1589 | 2524 | 2523 | High | High | High | High | High | High | High | High | High | High |
| 4634 | 3027 | 2399 | 3742 | 3896 | High | High | High | High | High | High | High | High | High | High |
| 8052 | 1124 | 661 | 2911 | 1977 | High | High | High | High | High | High | High | High | High | High |
| 5455 | 2700 | 2251 | 2734 | 3241 | High | High | High | High | High | High | High | High | High | High |
| 3172 | 2979 | 1803 | 4744 | 3402 | High | High | High | High | High | High | High | High | High | High |
| 5009 | 6409 | 6137 | 5650 | 6248 | High | High | High | High | High | High | High | High | High | High |
| 14173 | 6548 | 5545 | 6375 | 8076 | High | High | High | High | High | High | High | High | High | High |

| | | | | | | | | | | | | | | |
|---|---|---|---|---|---|---|---|---|---|---|---|---|---|---|
| 6917 | 5404 | 4076 | 7993 | 7201 | High | High | High | High | High | High | High | High | High | High |
| 7381 | 5938 | 3799 | 6072 | 4806 | High | High | High | High | High | High | High | High | High | High |
| 6664 | 7415 | 6220 | 8480 | 9156 | High | High | High | High | High | High | High | High | High | High |
| 1454 | 828 | 778 | 1874 | 1429 | High | High | High | High | High | High | High | High | High | High |
| 2172 | | | | | High | High | High | High | High | High | High | High | Not Founc | Not Founc |
| 5397 | 15442 | 8482 | 24593 | 16791 | High | High | High | High | High | High | High | High | High | High |
| 3495 | 4829 | 4209 | 5238 | 4708 | High | High | High | High | High | High | High | High | High | High |
| 6542 | 7239 | 4725 | 7723 | 7076 | High | High | High | High | High | High | High | High | High | High |
| 8894 | 2513 | 1831 | 2588 | 2766 | High | High | High | High | High | High | High | High | High | High |
| 3969 | 1687 | 1101 | 1687 | 1450 | High | High | High | High | High | High | High | High | High | High |
| 5870 | 8053 | 6386 | 9357 | 9529 | High | High | High | High | High | High | High | High | High | High |
| 7963 | 1032 | 1483 | 923 | 1927 | High | High | High | High | High | High | High | High | High | High |
| 10796 | 4420 | 3922 | 5539 | 5947 | High | High | High | High | High | High | High | High | High | High |
| 4922 | 2132 | 1446 | 3374 | 2895 | High | High | High | High | High | High | High | High | High | High |
| 3610 | 1939 | 1875 | 1438 | 1627 | High | High | High | High | High | High | High | High | High | High |
| 11715 | 11532 | 8141 | 17488 | 14382 | High | High | High | High | High | High | High | High | High | High |
| 5139 | 7302 | 4726 | 12189 | 9680 | High | High | High | High | High | High | High | High | High | High |
| 3029 | 1904 | 1618 | 2378 | 2324 | High | High | High | High | High | High | High | High | High | High |
| 12072 | 4298 | 4844 | 5690 | 7035 | High | High | High | High | High | High | High | High | High | High |
| 11928 | 6905 | 5387 | 7790 | 8377 | High | High | High | High | High | High | High | High | High | High |
| 5484 | 3791 | 2863 | 3875 | 4637 | High | High | High | High | High | High | High | High | High | High |
| 1953 | 2188 | 2103 | 1499 | 2106 | High | High | High | High | High | High | High | High | High | High |
| 12092 | 7130 | 5537 | 7046 | 7395 | High | High | High | High | High | High | High | High | High | High |
| 7299 | 5532 | 4183 | 7279 | 7119 | High | High | High | High | High | High | High | High | High | High |
| 9691 | 2102 | 2001 | 2109 | 2522 | High | High | High | High | High | High | High | High | High | High |
| 7885 | 3473 | 2831 | 4172 | 3737 | High | High | High | High | High | High | High | High | High | High |
| 7465 | 296 | 170 | 256 | 330 | High | High | High | High | High | High | High | High | High | High |
| 7498 | 745 | 738 | 1807 | 1081 | High | High | High | High | High | High | High | High | High | High |
| 5680 | 4916 | 4299 | 5117 | 4379 | High | High | High | High | High | High | High | High | High | High |
| 3716 | | | | | Not Founc | Not Founc | Not Founc | Not Founc | High | High | High | High | Not Founc | Not Founc |
| 8418 | 8489 | 5504 | 11935 | 9133 | High | High | High | High | High | High | High | High | High | High |

| | | | | | | | | | | | | | | |
|---|---|---|---|---|---|---|---|---|---|---|---|---|---|---|
| 2573 | 523 | 335 | 511 | 508 | High | High | High | High | High | High | High | High | High | High |
| 6200 | 9967 | 5334 | 13869 | 8697 | High | High | High | High | High | High | High | High | High | High |
| 8417 | 6344 | 5584 | 6706 | 7432 | High | High | High | High | High | High | High | High | High | High |
| 10920 | 8499 | 9384 | 8137 | 11850 | High | High | High | High | High | High | High | High | High | High |
| 7716 | 8975 | 6743 | 14212 | 14769 | High | High | High | High | High | High | High | High | High | High |
| | | | | | High | High | High | High | Not Founc | Not Founc | Not Founc | Not Founc | Not Founc | Not Founc |
| 1985 | 3442 | 2326 | 6120 | 4655 | High | High | High | High | High | High | High | High | High | High |
| 3588 | 4012 | 2648 | 5150 | 3779 | High | High | High | High | High | High | High | High | High | High |
| 4884 | 6909 | 6637 | 5777 | 7636 | High | High | High | High | High | High | High | High | High | High |
| 3614 | 1515 | 1196 | 1501 | 1728 | High | High | High | High | High | High | High | High | High | High |
| 8178 | 2896 | 2784 | 3331 | 4872 | High | High | High | High | High | High | High | High | High | High |
| 7491 | 1303 | 1076 | 2508 | 1936 | High | High | High | High | High | High | High | High | High | High |
| 12141 | 3301 | 1703 | 7633 | 3773 | High | High | High | High | High | High | High | High | High | High |
| 9393 | 5646 | 5616 | 6008 | 6422 | High | High | High | High | High | High | High | High | High | High |
| 5013 | 5453 | 4563 | 7180 | 10206 | High | High | High | High | High | High | High | High | High | High |
| 2326 | 2370 | 1925 | 2541 | 2643 | High | High | High | High | High | High | High | High | High | High |
| 1920 | 7806 | 7107 | 7760 | 8992 | High | High | High | High | High | High | High | High | High | High |
| 5263 | 12788 | 9576 | 16100 | 16558 | High | High | High | High | High | High | High | High | High | High |
| 9460 | 2968 | 2679 | 2696 | 3668 | High | High | High | High | High | High | High | High | High | High |
| 15699 | 7242 | 4644 | 12551 | 10687 | High | High | High | High | High | High | High | High | High | High |
| 3228 | 3928 | 2995 | 4506 | 4630 | High | High | High | High | High | High | High | High | High | High |
| 6821 | 4534 | 2619 | 7375 | 4702 | High | High | High | High | High | High | High | High | High | High |
| 329 | 3893 | 3233 | 3463 | 3442 | High | High | High | High | High | High | High | High | High | High |
| 7312 | 4416 | 3452 | 5701 | 6336 | High | High | High | High | High | High | High | High | High | High |
| 5659 | 4514 | 3418 | 9166 | 6587 | High | High | High | High | High | High | High | High | High | High |
| 6064 | 3824 | 2686 | 5767 | 4559 | High | High | High | High | High | High | High | High | High | High |
| 8192 | 5138 | 4088 | 9082 | 7606 | High | High | High | High | High | High | High | High | High | High |
| 5097 | 6020 | 2792 | 4302 | 3904 | High | High | High | High | High | High | High | High | High | High |
| 9132 | 4089 | 1913 | 5245 | 3754 | High | High | High | High | High | High | High | High | High | High |
| 3222 | 1260 | 995 | 2185 | 1568 | High | High | High | High | High | High | High | High | High | High |
| 23221 | 4675 | 3281 | 6447 | 4852 | High | High | High | High | High | High | High | High | High | High |

| | | | | | | | | | | | | | | |
|---|---|---|---|---|---|---|---|---|---|---|---|---|---|---|
| 5364 | 2549 | 1265 | 4094 | 2130 | High | High | High | High | High | High | High | High | High | High |
| 4596 | 1410 | 769 | 1898 | 1508 | High | High | High | High | High | High | High | High | High | High |
| 1486 | 2270 | 2221 | 1551 | 1995 | High | High | High | High | High | High | High | High | High | High |
| 4117 | 2952 | 3229 | 4041 | 4847 | High | High | High | High | High | High | High | High | High | High |
| 3973 | 2075 | 1540 | 3934 | 2846 | High | High | High | High | High | High | High | High | High | High |
| 11630 | 1971 | 1337 | 1288 | 1557 | High | High | High | High | High | High | High | High | High | High |
| 10598 | 3559 | 2362 | 5348 | 3944 | High | High | High | High | High | High | High | High | High | High |
| 9246 | 4133 | 2826 | 4677 | 4324 | High | High | High | High | High | High | High | High | High | High |
| 4489 | 12198 | 8344 | 15825 | 14539 | High | High | High | High | High | High | High | High | High | High |
| 6982 | 5784 | 5044 | 5402 | 7073 | High | High | High | High | High | High | High | High | High | High |
| 10375 | 2256 | 1433 | 4480 | 2909 | High | High | High | High | High | High | High | High | High | High |
| 5421 | 8119 | 7463 | 9786 | 11863 | High | High | High | High | High | High | High | High | High | High |
| 3164 | 9062 | 4900 | 15113 | 9406 | High | High | High | High | High | High | High | High | High | High |
| 3170 | 8936 | 6126 | 13381 | 10431 | High | High | High | High | High | High | High | High | High | High |
| 4315 | 3740 | 4365 | 5206 | 5802 | High | High | High | High | High | High | High | High | High | High |
| 6131 | 4270 | 3251 | 5144 | 5177 | High | High | High | High | High | High | High | High | High | High |
| 4609 | 791 | 664 | 992 | 1010 | Not Founc | Not Founc | Not Founc | Not Founc | High | High | High | High | High | High |
| 2899 | 989 | 912 | 995 | 1124 | High | High | High | High | High | High | High | High | High | High |
| 2962 | 4101 | 3192 | 6588 | 5141 | High | High | High | High | High | High | High | High | High | High |
| 5077 | 6767 | 3843 | 10860 | 6395 | High | High | High | High | High | High | High | High | High | High |
| 4011 | 1758 | 1263 | 2884 | 2024 | High | High | High | High | High | High | High | High | High | High |
| | 1259 | 668 | 5251 | 2962 | High | High | High | High | Not Founc | Not Founc | Not Founc | Not Founc | High | High |
| 9702 | 7573 | 4786 | 19267 | 12510 | High | High | High | High | High | High | High | High | High | High |
| 8335 | 7441 | 4866 | 13870 | 8923 | High | High | High | High | High | High | High | High | High | High |
| 2517 | 711 | 558 | 664 | 818 | High | High | High | High | High | High | High | High | High | High |
| 5542 | 6396 | 5018 | 9477 | 7975 | High | High | High | High | High | High | High | High | High | High |
| 6691 | 3612 | 2859 | 7035 | 4562 | High | High | High | High | High | High | High | High | High | High |
| 7354 | 3043 | 2228 | 2630 | 3196 | High | High | High | High | High | High | High | High | High | High |
| 2359 | 7959 | 3805 | 10992 | 7868 | Not Founc | Not Founc | Not Founc | Not Founc | High | High | High | High | High | High |
| 6218 | 3058 | 1819 | 4594 | 3453 | High | High | High | High | High | High | High | High | High | High |
| | 1493 | 1504 | 1699 | 2578 | High | High | High | High | Not Founc | Not Founc | Not Founc | Not Founc | High | High |

| | | | | | | | | | | | | | | |
|---|---|---|---|---|---|---|---|---|---|---|---|---|---|---|
| 3738 | | | | | High | High | High | High | High | High | High | High | Not Founc | Not Founc |
| 13753 | | | | | High | High | High | High | High | High | High | High | Not Founc | Not Founc |
| 3983 | 13107 | 9894 | 14757 | 13862 | High | High | High | High | High | High | High | High | High | High |
| 6218 | 5040 | 4020 | 9235 | 6833 | High | High | High | High | High | High | High | High | High | High |
| | 4785 | 2974 | 5543 | 4204 | High | High | High | High | Not Founc | Not Founc | Not Founc | Not Founc | High | High |
| 8828 | 7847 | 4348 | 11679 | 9305 | High | High | High | High | High | High | High | High | High | High |
| | 12303 | 8521 | 16997 | 15684 | High | High | High | High | Not Founc | Not Founc | Not Founc | Not Founc | High | High |
| 15512 | 7347 | 4948 | 13498 | 9516 | High | High | High | High | High | High | High | High | High | High |
| 2207 | 1042 | 606 | 1660 | 1072 | High | High | High | High | High | High | High | High | High | High |
| 7453 | 2352 | 1434 | 3363 | 2074 | High | High | High | High | High | High | High | High | High | High |
| 4458 | 3771 | 3122 | 5119 | 5762 | High | High | High | High | High | High | High | High | High | High |
| 5849 | 8401 | 6447 | 10931 | 11048 | High | High | High | High | High | High | High | High | High | High |
| 3105 | 1973 | 1430 | 4683 | 2729 | High | High | High | High | High | High | High | High | High | High |
| 6700 | 7924 | 5277 | 13323 | 10058 | High | High | High | High | High | High | High | High | High | High |
| 16017 | 5683 | 3979 | 6632 | 9117 | High | High | High | High | High | High | High | High | High | High |
| 2256 | 2354 | 1697 | 3481 | 2609 | High | High | High | High | High | High | High | High | High | High |
| 2795 | 4976 | 4165 | 5022 | 6148 | High | High | High | High | High | High | High | High | High | High |
| 6864 | 4094 | 3098 | 8322 | 6127 | High | High | High | High | High | High | High | High | High | High |
| 5936 | 4272 | 2389 | 4772 | 3917 | High | High | High | High | High | High | High | High | High | High |
| | 4245 | 4650 | 2354 | 3501 | High | High | High | High | Not Founc | Not Founc | Not Founc | Not Founc | High | High |
| 3083 | 5680 | 2665 | 5188 | 3378 | High | High | High | High | High | High | High | High | High | High |
| 10951 | 3402 | 2821 | 3627 | 3855 | High | High | High | High | High | High | High | High | High | High |
| 614 | 4758 | 2582 | 4116 | 3725 | High | High | High | High | High | High | High | High | High | High |
| 2895 | 7069 | 3353 | 8113 | 5459 | High | High | High | High | High | High | High | High | High | High |
| 8748 | 8516 | 7443 | 10139 | 10966 | High | High | High | High | High | High | High | High | High | High |
| 1608 | 749 | 588 | 761 | 681 | High | High | High | High | High | High | High | High | High | High |
| 4419 | 4008 | 2933 | 6845 | 4821 | High | High | High | High | High | High | High | High | High | High |
| 6031 | 3223 | 2801 | 4183 | 3591 | High | High | High | High | High | High | High | High | High | High |
| 16035 | 8636 | 5359 | 13884 | 21079 | High | High | High | High | High | High | High | High | High | High |
| 10950 | 5175 | 4161 | 6049 | 6201 | High | High | High | High | High | High | High | High | High | High |
| 5007 | 7707 | 5476 | 10792 | 10440 | High | High | High | High | High | High | High | High | High | High |

| | | | | | | | | | | | | | | |
|---|---|---|---|---|---|---|---|---|---|---|---|---|---|---|
| 3199 | | | | | High | High | High | High | High | High | High | High | Not Found | Not Found |
| 2353 | | | | | Not Found | Not Found | Not Found | Not Found | High | High | High | High | Not Found | Not Found |
| 2889 | 4403 | 3286 | 7195 | 6909 | High | High | High | High | High | High | High | High | High | High |
| 1815 | 3246 | 2136 | 4176 | 3892 | High | High | High | High | High | High | High | High | High | High |
| 5137 | | | | | High | High | High | High | High | High | High | High | Not Found | Not Found |
| 1830 | 4979 | 3709 | 5084 | 4857 | High | High | High | High | High | High | High | High | High | High |
| 6233 | 1095 | 743 | 1324 | 1204 | High | High | High | High | High | High | High | High | High | High |
| 2725 | 1693 | 1162 | 2413 | 2016 | High | High | High | High | High | High | High | High | High | High |
| 6545 | 5163 | 5086 | 3060 | 5416 | High | High | High | High | High | High | High | High | High | High |
| 3377 | 1325 | 3025 | 5062 | 2643 | High | High | High | High | High | High | High | High | High | High |
| 3164 | 1324 | 774 | 1815 | 1201 | High | High | High | High | High | High | High | High | High | High |
| | 1130 | 1066 | 1033 | 1150 | High | High | High | High | Not Found | Not Found | Not Found | Not Found | High | High |
| 10561 | 1828 | 1347 | 3570 | 2637 | High | High | High | High | High | High | High | High | High | High |
| 1123 | 2325 | 2147 | 2776 | 2955 | High | High | High | High | High | High | High | High | High | High |
| 3351 | 5074 | 3572 | 9702 | 5987 | High | High | High | High | High | High | High | High | High | High |
| 6843 | 1520 | 1477 | 1699 | 2165 | High | High | High | High | High | High | High | High | High | High |
| 6776 | 7511 | 4257 | 12812 | 10227 | High | High | High | High | High | High | High | High | High | High |
| 2497 | 7245 | 4528 | 7953 | 7636 | High | High | High | High | High | High | High | High | High | High |
| 3594 | | | | | High | High | High | High | High | High | High | High | Not Found | Not Found |
| 13122 | 9709 | 6725 | 9300 | 10217 | High | High | High | High | High | High | High | High | High | High |
| 10878 | 5342 | 4248 | 7449 | 6771 | High | High | High | High | High | High | High | High | High | High |
| | 1966 | 1656 | 3121 | 3116 | High | High | High | High | Not Found | Not Found | Not Found | Not Found | High | High |
| 3056 | 1776 | 1756 | 1906 | 2507 | High | High | High | High | High | High | High | High | High | High |
| 7263 | 2552 | 1401 | 5711 | 3754 | High | High | High | High | High | High | High | High | High | High |
| 3813 | 6208 | 6243 | 9848 | 9153 | High | High | High | High | High | High | High | High | High | High |
| 7180 | 7580 | 6734 | 12173 | 9552 | High | High | High | High | High | High | High | High | High | High |
| 8756 | 3549 | 2386 | 3821 | 3194 | High | High | High | High | High | High | High | High | High | High |
| | 5860 | 4659 | 5156 | 5487 | High | High | High | High | Not Found | Not Found | Not Found | Not Found | High | High |
| 5058 | 8346 | 7004 | 7364 | 9085 | High | High | High | High | High | High | High | High | High | High |
| | 1843 | 1014 | 3507 | 2348 | High | High | High | High | Not Found | Not Found | Not Found | Not Found | High | High |
| 6748 | 4532 | 4018 | 6676 | 7844 | High | High | High | High | High | High | High | High | High | High |

| | | | | | | | | | | | | | | |
|---|---|---|---|---|---|---|---|---|---|---|---|---|---|---|
| 6647 | 5316 | 4107 | 10166 | 7583 | High | High | High | High | High | High | High | High | High | High |
| 1661 | 4914 | 4745 | 6969 | 7002 | High | High | High | High | High | High | High | High | High | High |
| 5224 | 4581 | 3200 | 6293 | 4799 | High | High | High | High | High | High | High | High | High | High |
| 1758 | 1707 | 1374 | 2327 | 2048 | High | High | High | High | High | High | High | High | High | High |
| 17209 | | | | | High | High | High | High | High | High | High | High | Not Found | Not Found |
| 9200 | 4277 | 3936 | 4135 | 6072 | High | High | High | High | High | High | High | High | High | High |
| 5906 | 1348 | 830 | 2237 | 1898 | High | High | High | High | High | High | High | High | High | High |
| 3474 | 3674 | 3060 | 4145 | 7055 | High | High | High | High | High | High | High | High | High | High |
| 3559 | 2225 | 1784 | 3039 | 3140 | High | High | High | High | High | High | High | High | High | High |
| | 8594 | 6706 | 9148 | 10546 | High | High | High | High | Not Found | Not Found | Not Found | Not Found | High | High |
| 3573 | 4953 | 3315 | 7046 | 5068 | High | High | High | High | High | High | High | High | High | High |
| 13337 | 7211 | 5165 | 6854 | 8318 | High | High | High | High | High | High | High | High | High | High |
| 8372 | 3627 | 2698 | 4878 | 3757 | High | High | High | High | High | High | High | High | High | High |
| 4685 | 730 | 443 | 1058 | 998 | High | High | High | High | High | High | High | High | High | High |
| 6294 | 4699 | 2602 | 6683 | 4816 | High | High | High | High | High | High | High | High | High | High |
| 5932 | 1001 | 668 | 1589 | 983 | High | High | High | High | High | High | High | High | High | High |
| 5674 | 1229 | 1079 | 1777 | 1659 | High | High | High | High | High | High | High | High | High | High |
| | 9795 | 6973 | 10795 | 12495 | High | High | High | High | Not Found | Not Found | Not Found | Not Found | High | High |
| 14010 | 11785 | 8802 | 9891 | 14632 | High | High | High | High | High | High | High | High | High | High |
| 7284 | 4040 | 3736 | 5585 | 6210 | High | High | High | High | High | High | High | High | High | High |
| 8342 | 12447 | 8728 | 10640 | 11793 | Not Found | Not Found | Not Found | Not Found | High | High | High | High | High | High |
| 12583 | 5371 | 3015 | 9481 | 4521 | Not Found | Not Found | Not Found | Not Found | High | High | High | High | High | High |
| 9261 | 6461 | 6472 | 5984 | 9338 | High | High | High | High | High | High | High | High | High | High |
| 9835 | 8956 | 7294 | 9877 | 7296 | High | High | High | High | High | High | High | High | High | High |
| 7640 | 3074 | 2372 | 4836 | 4836 | High | High | High | High | High | High | High | High | High | High |
| 5897 | | | | | High | High | High | High | High | High | High | High | Not Found | Not Found |
| 2015 | 9654 | 8124 | 11794 | 11839 | High | High | High | High | High | High | High | High | High | High |
| 3174 | 671 | 835 | 727 | 1079 | High | High | High | High | High | High | High | High | High | High |
| 891 | 4301 | 3777 | 3034 | 4562 | High | High | High | High | High | High | High | High | High | High |
| 9333 | 5794 | 3039 | 14188 | 8003 | Not Found | Not Found | Not Found | Not Found | High | High | High | High | High | High |
| 15313 | 7770 | 5717 | 10739 | 9289 | High | High | High | High | High | High | High | High | High | High |

| | | | | | | | | | | | | | | |
|---|---|---|---|---|---|---|---|---|---|---|---|---|---|---|
| 3628 | 3127 | 2560 | 2541 | 2732 | High | High | High | High | High | High | High | High | High | High |
| 477 | | | | | High | High | High | High | High | High | High | High | Not Found | Not Found |
| 5390 | 4528 | 3417 | 6655 | 7695 | High | High | High | High | High | High | High | High | High | High |
| 8184 | 5184 | 4431 | 5721 | 6993 | High | High | High | High | High | High | High | High | High | High |
| 5632 | 5399 | 4036 | 5603 | 7125 | High | High | High | High | High | High | High | High | High | High |
| 1072 | 1440 | 859 | 2037 | 1229 | High | High | High | High | High | High | High | High | High | High |
| | 9219 | 8354 | 18005 | 18735 | High | High | High | High | Not Found | Not Found | Not Found | Not Found | High | High |
| 6292 | 10102 | 5788 | 13607 | 10191 | High | High | High | High | High | High | High | High | High | High |
| | 7878 | 5451 | 6496 | 6871 | High | High | High | High | Not Found | Not Found | Not Found | Not Found | High | High |
| 5250 | 3094 | 2418 | 3586 | 3469 | High | High | High | High | High | High | High | High | High | High |
| 8778 | 5054 | 2326 | 5639 | 3886 | High | High | High | High | High | High | High | High | High | High |
| 3108 | 3981 | 2272 | 9662 | 6231 | High | High | High | High | High | High | High | High | High | High |
| 862 | 553 | 401 | 697 | 620 | High | High | High | High | High | High | High | High | High | High |
| 7908 | 4125 | 4234 | 5296 | 5176 | High | High | High | High | High | High | High | High | High | High |
| 2798 | 4526 | 2744 | 7429 | 5294 | High | High | High | High | High | High | High | High | High | High |
| 683 | 5051 | 4333 | 5588 | 6075 | High | High | High | High | High | High | High | High | High | High |
| 4346 | 599 | 376 | 544 | 499 | High | High | High | High | High | High | High | High | High | High |
| 13860 | 8538 | 5983 | 10379 | 10965 | High | High | High | High | High | High | High | High | High | High |
| | 6939 | 5466 | 8455 | 8515 | High | High | High | High | Not Found | Not Found | Not Found | Not Found | High | High |
| 6062 | 4767 | 2903 | 7226 | 5123 | High | High | High | High | High | High | High | High | High | High |
| 3124 | 7287 | 6538 | 8362 | 10546 | High | High | High | High | High | High | High | High | High | High |
| 5803 | 5661 | 4912 | 7179 | 7630 | High | High | High | High | High | High | High | High | High | High |
| 629 | 359 | 317 | 255 | 354 | High | High | High | High | High | High | High | High | High | High |
| | 7083 | 3844 | 8645 | 7062 | High | High | High | High | Not Found | Not Found | Not Found | Not Found | High | High |
| 690 | 3351 | 2314 | 3661 | 3633 | High | High | High | High | High | High | High | High | High | High |
| | 9722 | 9782 | 10340 | 13713 | High | High | High | High | Not Found | Not Found | Not Found | Not Found | High | High |
| 2915 | 1219 | 1077 | 2700 | 1941 | High | High | High | High | High | High | High | High | High | High |
| 1134 | 5042 | 4075 | 4944 | 5662 | Not Found | Not Found | Not Found | Not Found | High | High | High | High | High | High |
| 9665 | | | | | High | High | High | High | High | High | High | High | Not Found | Not Found |
| 5258 | 2739 | 2045 | 3109 | 3101 | High | High | High | High | High | High | High | High | High | High |
| | 5703 | 4247 | 4537 | 4396 | High | High | High | High | Not Found | Not Found | Not Found | Not Found | High | High |

| | | | | | | | | | | | | | | |
|---|---|---|---|---|---|---|---|---|---|---|---|---|---|---|
| 10507 | 8488 | 7229 | 9339 | 10157 | High | High | High | High | High | High | High | High | High | High |
| 2633 | 2425 | 2133 | 2768 | 3195 | High | High | High | High | High | High | High | High | High | High |
| 4694 | 6971 | 5814 | 8814 | 11563 | High | High | High | High | High | High | High | High | High | High |
| 16381 | 8769 | 5924 | 10618 | 10062 | High | High | High | High | High | High | High | High | High | High |
| 7849 | 2751 | 2636 | 2846 | 3435 | High | High | High | High | High | High | High | High | High | High |
| 5247 | 3475 | 3977 | 4200 | 5689 | High | High | High | High | High | High | High | High | High | High |
| 5787 | 5811 | 4725 | 6222 | 6111 | High | High | High | High | High | High | High | High | High | High |
| 853 | 4276 | 2458 | 5078 | 4062 | High | High | High | High | High | High | High | High | High | High |
| 8179 | 3587 | 3423 | 3616 | 5086 | High | High | High | High | High | High | High | High | High | High |
| 4051 | 3650 | 2502 | 4387 | 4193 | High | High | High | High | High | High | High | High | High | High |
| 1754 | 1862 | 1716 | 1521 | 1988 | High | High | High | High | High | High | High | High | High | High |
| 9034 | 4925 | 4600 | 5289 | 6708 | Not Founc | Not Founc | Not Founc | Not Founc | High | High | High | High | High | High |
| 4170 | 5141 | 4166 | 4265 | 6525 | High | High | High | High | High | High | High | High | High | High |
| 1073 | 2770 | 2010 | 2345 | 1605 | High | High | High | High | High | High | High | High | High | High |
| 5389 | 1339 | 1312 | 1167 | 2082 | High | High | High | High | High | High | High | High | High | High |
| | 8474 | 7411 | 12571 | 10627 | High | High | High | High | Not Founc | Not Founc | Not Founc | Not Founc | High | High |
| 3443 | 2776 | 1882 | 3923 | 3005 | High | High | High | High | High | High | High | High | High | High |
| 2623 | 1774 | 1465 | 1848 | 2352 | High | High | High | High | High | High | High | High | High | High |
| 1392 | | | | | High | High | High | High | High | High | High | High | Not Founc | Not Founc |
| 11887 | 7195 | 4793 | 13865 | 10184 | Not Founc | Not Founc | Not Founc | Not Founc | High | High | High | High | High | High |
| 2036 | 5510 | 3631 | 8349 | 5965 | High | High | High | High | High | High | High | High | High | High |
| 6581 | 3046 | 4037 | 5320 | 6045 | High | High | High | High | High | High | High | High | High | High |
| 7596 | 1521 | 880 | 1634 | 1332 | High | High | High | High | High | High | High | High | High | High |
| 3121 | 4497 | 1279 | 7557 | 2232 | High | High | High | High | High | High | High | High | High | High |
| 6838 | 4043 | 3104 | 6968 | 4864 | High | High | High | High | High | High | High | High | High | High |
| 10462 | | | | | High | High | High | High | High | High | High | High | Not Founc | Not Founc |
| 5888 | | | | | High | High | High | High | High | High | High | High | Not Founc | Not Founc |
| 3561 | 1833 | 1441 | 1796 | 2013 | High | High | High | High | High | High | High | High | High | High |
| 2459 | 5993 | 4356 | 7354 | 7677 | High | High | High | High | High | High | High | High | High | High |
| 14130 | 3495 | 3033 | 4028 | 4692 | High | High | High | High | High | High | High | High | High | High |
| 23722 | 1775 | 1029 | 2343 | 4296 | High | High | High | High | High | High | High | High | High | High |

| | | | | | | | | | | | | | | |
|---|---|---|---|---|---|---|---|---|---|---|---|---|---|---|
| 1112 | 4606 | 3161 | 6403 | 5604 | High | High | High | High | High | High | High | High | High | High |
| 3302 | 4177 | 2986 | 3232 | 4070 | High | High | High | High | High | High | High | High | High | High |
| 2082 | 1683 | 1381 | 1929 | 1892 | High | High | High | High | High | High | High | High | High | High |
| | 8896 | 6151 | 10530 | 10028 | High | High | High | High | Not Founc | Not Founc | Not Founc | Not Founc | High | High |
| 6188 | 3474 | 2290 | 4191 | 3567 | High | High | High | High | High | High | High | High | High | High |
| 13304 | 2678 | 2110 | 3231 | 2762 | High | High | High | High | High | High | High | High | High | High |
| 3509 | 2227 | 1747 | 3871 | 3055 | High | High | High | High | High | High | High | High | High | High |
| 8903 | 5735 | 4821 | 8041 | 8628 | High | High | High | High | High | High | High | High | High | High |
| | 1522 | 1445 | 2274 | 2226 | High | High | High | High | Not Founc | Not Founc | Not Founc | Not Founc | High | High |
| 5045 | | | | | High | High | High | High | High | High | High | High | Not Founc | Not Founc |
| 1056 | | | | | High | High | High | High | High | High | High | High | Not Founc | Not Founc |
| | 1365 | 1280 | 1789 | 1835 | High | High | High | High | Not Founc | Not Founc | Not Founc | Not Founc | High | High |
| 3166 | 4929 | 3712 | 5327 | 5119 | High | High | High | High | High | High | High | High | High | High |
| 4641 | 4184 | 3299 | 5906 | 4941 | Not Founc | Not Founc | Not Founc | Not Founc | High | High | High | High | High | High |
| 1965 | 5103 | 3929 | 4324 | 5237 | High | High | High | High | High | High | High | High | High | High |
| 2044 | 2029 | 1835 | 1817 | 2170 | High | High | High | High | High | High | High | High | High | High |
| 4396 | 10016 | 7003 | 14211 | 14226 | High | High | High | High | High | High | High | High | High | High |
| 7533 | | | | | High | High | High | High | High | High | High | High | Not Founc | Not Founc |
| 3265 | 3595 | 3260 | 7225 | 6406 | Not Founc | Not Founc | Not Founc | Not Founc | High | High | High | High | High | High |
| 6717 | 3546 | 2925 | 4247 | 4809 | High | High | High | High | High | High | High | High | High | High |
| 6318 | 3761 | 2909 | 3666 | 4324 | High | High | High | High | High | High | High | High | High | High |
| 599 | 5576 | 4138 | 5177 | 5201 | High | High | High | High | High | High | High | High | High | High |
| 2614 | 1999 | 1143 | 3298 | 2730 | High | High | High | High | High | High | High | High | High | High |
| 8554 | 1154 | 1166 | 1915 | 1772 | High | High | High | High | High | High | High | High | High | High |
| 2817 | 364 | 380 | 569 | 523 | High | High | High | High | High | High | High | High | High | High |
| 1932 | 7673 | 4383 | 13512 | 9797 | High | High | High | High | High | High | High | High | High | High |
| 6529 | 8746 | 5886 | 10799 | 8745 | Not Founc | Not Founc | Not Founc | Not Founc | High | High | High | High | High | High |
| 1366 | 2585 | 1614 | 3466 | 2584 | High | High | High | High | High | High | High | High | High | High |
| 6252 | 845 | 511 | 777 | 637 | High | High | High | High | High | High | High | High | High | High |
| 4780 | 3197 | 2648 | 3378 | 3179 | High | High | High | High | High | High | High | High | High | High |
| 5185 | 5680 | 3375 | 3918 | 4461 | High | High | High | High | High | High | High | High | High | High |

| | | | | | | | | | | | | | | |
|---|---|---|---|---|---|---|---|---|---|---|---|---|---|---|
| 7046 | 5742 | 4727 | 6737 | 7279 | High | High | High | High | High | High | High | High | High | High |
| 3282 | 8060 | 6029 | 14624 | 11407 | High | High | High | High | High | High | High | High | High | High |
| 9107 | 3126 | 2620 | 4337 | 4234 | High | High | High | High | High | High | High | High | High | High |
| 7020 | 2562 | 1956 | 4284 | 4093 | High | High | High | High | High | High | High | High | High | High |
| 4158 | 3639 | 3196 | 4062 | 3872 | High | High | High | High | High | High | High | High | High | High |
| 6998 | 4964 | 3825 | 5930 | 7097 | High | High | High | High | High | High | High | High | High | High |
| 1477 | 4369 | 2770 | 5945 | 4636 | High | High | High | High | High | High | High | High | High | High |
| 7035 | 3621 | 2147 | 4373 | 3944 | High | High | High | High | High | High | High | High | High | High |
| | 2801 | 2300 | 2543 | 2535 | Not Found | Not Found | Not Found | Not Found | Not Found | Not Found | Not Found | Not Found | High | High |
| 3371 | 1309 | 1050 | 1367 | 1230 | High | High | High | High | High | High | High | High | High | High |
| 1044 | | | | | Not Found | Not Found | Not Found | Not Found | High | High | High | High | Not Found | Not Found |
| 5361 | 3123 | 1889 | 4127 | 3272 | High | High | High | High | High | High | High | High | High | High |
| 9369 | 2104 | 2467 | 3185 | 2904 | High | High | High | High | High | High | High | High | High | High |
| 3034 | 574 | 531 | 971 | 764 | High | High | High | High | High | High | High | High | High | High |
| 9997 | 5896 | 4767 | 6174 | 6656 | High | High | High | High | High | High | High | High | High | High |
| 1133 | 2792 | 2052 | 4923 | 3718 | High | High | High | High | High | High | High | High | High | High |
| 1354 | 2186 | 1187 | 4113 | 2730 | High | High | High | High | High | High | High | High | High | High |
| 4863 | | | | | High | High | High | High | High | High | High | High | Not Found | Not Found |
| 2081 | 1580 | 1309 | 1987 | 1952 | High | High | High | High | High | High | High | High | High | High |
| 4882 | 6530 | 5837 | 8410 | 7197 | High | High | High | High | High | High | High | High | High | High |
| 2824 | 3849 | 2681 | 6051 | 4569 | Not Found | Not Found | Not Found | Not Found | High | High | High | High | High | High |
| 5335 | 6549 | 3771 | 9461 | 7992 | High | High | High | High | High | High | High | High | High | High |
| | 5014 | 3832 | 6172 | 5840 | High | High | High | High | Not Found | Not Found | Not Found | Not Found | High | High |
| 2248 | 4631 | 3766 | 5140 | 5147 | High | High | High | High | High | High | High | High | High | High |
| 5418 | 1052 | 789 | 1066 | 1186 | High | High | High | High | High | High | High | High | High | High |
| 7792 | 3694 | 4323 | 2880 | 4221 | High | High | High | High | High | High | High | High | High | High |
| 8434 | 8091 | 5185 | 15796 | 10777 | Not Found | Not Found | Not Found | Not Found | High | High | High | High | High | High |
| 2678 | 8755 | 5141 | 11023 | 9156 | High | High | High | High | High | High | High | High | High | High |
| 1318 | 1236 | 900 | 1646 | 1469 | High | High | High | High | High | High | High | High | High | High |
| 2448 | 7175 | 5974 | 7815 | 9008 | High | High | High | High | High | High | High | High | High | High |
| 5483 | 8881 | 7092 | 11814 | 11239 | High | High | High | High | High | High | High | High | High | High |

| | | | | | | | | | | | | | | |
|---|---|---|---|---|---|---|---|---|---|---|---|---|---|---|
| 5070 | 2333 | 2060 | 5899 | 3367 | High | High | High | High | High | High | High | High | High | High |
| 5120 | 1565 | 952 | 2238 | 1517 | High | High | High | High | High | High | High | High | High | High |
| | 3016 | 1683 | 7466 | 4284 | High | High | High | High | High | High | High | High | High | High |
| 21540 | 5676 | 3704 | 9592 | 6937 | Not Founc | Not Founc | Not Founc | Not Founc | High | High | High | High | High | High |
| | 4714 | 3170 | 4642 | 4774 | High | High | High | High | Not Founc | Not Founc | Not Founc | Not Founc | High | High |
| 7915 | 7176 | 4979 | 7470 | 7333 | High | High | High | High | High | High | High | High | High | High |
| 8385 | 5306 | 2734 | 6592 | 5862 | High | High | High | High | High | High | High | High | High | High |
| 6469 | 154 | 67 | 215 | 126 | High | High | High | High | High | High | High | High | High | High |
| 2164 | 5435 | 4527 | 11536 | 6563 | High | High | High | High | High | High | High | High | High | High |
| 9131 | 3192 | 1814 | 3400 | 3140 | High | High | High | High | High | High | High | High | High | High |
| | 5037 | 5283 | 5747 | 8929 | High | High | High | High | Not Founc | Not Founc | Not Founc | Not Founc | High | High |
| 2916 | | | | | Not Founc | Not Founc | Not Founc | Not Founc | High | High | High | High | Not Founc | Not Founc |
| | 4569 | 2797 | 4333 | 3594 | High | High | High | High | Not Founc | Not Founc | Not Founc | Not Founc | High | High |
| | 2317 | 1060 | 2374 | 1987 | High | High | High | High | Not Founc | Not Founc | Not Founc | Not Founc | High | High |
| 5198 | 5475 | 3416 | 7623 | 6208 | High | High | High | High | High | High | High | High | High | High |
| 9530 | | | | | High | High | High | High | High | High | High | High | Not Founc | Not Founc |
| 2582 | 1509 | 1106 | 2104 | 1825 | High | High | High | High | High | High | High | High | High | High |
| 4945 | 2812 | 2699 | 5409 | 4368 | High | High | High | High | High | High | High | High | High | High |
| 3206 | 6257 | 4457 | 9442 | 8496 | High | High | High | High | High | High | High | High | High | High |
| 2733 | 1632 | 2838 | 12531 | 4413 | High | High | High | High | High | High | High | High | High | High |
| 4167 | 2091 | 1057 | 2558 | 2007 | High | High | High | High | High | High | High | High | High | High |
| 6320 | 3852 | 3071 | 6378 | 3952 | High | High | High | High | High | High | High | High | High | High |
| 5048 | 1495 | 1283 | 1915 | 1643 | Not Founc | Not Founc | Not Founc | Not Founc | High | High | High | High | High | High |
| 4231 | 1096 | 1016 | 1629 | 1373 | High | High | High | High | High | High | High | High | High | High |
| 9932 | 9004 | 9944 | 9441 | 9587 | Not Founc | Not Founc | Not Founc | Not Founc | High | High | High | High | High | High |
| 2722 | 2973 | 1958 | 4280 | 4028 | High | High | High | High | High | High | High | High | High | High |
| 6714 | 1793 | 1833 | 1786 | 2422 | High | High | High | High | High | High | High | High | High | High |
| 14110 | 3139 | 1629 | 3242 | 3139 | High | High | High | High | High | High | High | High | High | High |
| 5514 | 4120 | 2838 | 6505 | 5963 | High | High | High | High | High | High | High | High | High | High |
| 14730 | 6063 | 5340 | 6419 | 7925 | High | High | High | High | High | High | High | High | High | High |
| 2999 | 3867 | 2755 | 4541 | 4776 | High | High | High | High | High | High | High | High | High | High |

| | | | | | | | | | | | | | | |
|---|---|---|---|---|---|---|---|---|---|---|---|---|---|---|
| 7249 | | | | | High | High | High | High | High | High | High | High | Not Founc | Not Founc |
| 6303 | 3707 | 2673 | 3688 | 3780 | High | High | High | High | High | High | High | High | High | High |
| 3978 | 8752 | 3824 | 9185 | 7520 | High | High | High | High | High | High | High | High | High | High |
| 3335 | 4820 | 4789 | 5953 | 8126 | High | High | High | High | High | High | High | High | High | High |
| 8890 | 1364 | 949 | 1793 | 1284 | Not Founc | Not Founc | Not Founc | Not Founc | High | High | High | High | High | High |
| 3555 | | | | | High | High | High | High | High | High | High | High | Not Founc | Not Founc |
| 2595 | 4460 | 3084 | 5027 | 4416 | High | High | High | High | High | High | High | High | High | High |
| 3636 | 5093 | 3704 | 8046 | 6465 | High | High | High | High | High | High | High | High | High | High |
| 6096 | 6423 | 4123 | 10707 | 7865 | High | High | High | High | High | High | High | High | High | High |
| 6621 | 3252 | 2258 | 3748 | 3478 | High | High | High | High | High | High | High | High | High | High |
| 5905 | | | | | High | High | High | High | High | High | High | High | Not Founc | Not Founc |
| 3550 | 5798 | 4673 | 5804 | 4751 | High | High | High | High | High | High | High | High | High | High |
| 5376 | 1842 | 1074 | 1389 | 1503 | High | High | High | High | High | High | High | High | High | High |
| 7389 | 7878 | 3688 | 17173 | 9006 | High | High | High | High | High | High | High | High | High | High |
| 2915 | 277 | 228 | 510 | 395 | High | High | High | High | High | High | High | High | High | High |
| 661 | 1559 | 1143 | 2444 | 1860 | High | High | High | High | High | High | High | High | High | High |
| 4166 | 3962 | 3010 | 5719 | 4757 | High | High | High | High | High | High | High | High | High | High |
| 4242 | 2531 | 2389 | 3305 | 3648 | High | High | High | High | High | High | High | High | High | High |
| 999 | 3116 | 2500 | 3735 | 3597 | High | High | High | High | High | High | High | High | High | High |
| 2048 | 3510 | 2553 | 3844 | 3492 | High | High | High | High | High | High | High | High | High | High |
| | | | | | High | High | High | High | High | High | High | High | Not Founc | Not Founc |
| 1770 | 561 | 463 | 1175 | 718 | High | High | High | High | High | High | High | High | High | High |
| 2308 | 7366 | 4671 | 9246 | 7919 | High | High | High | High | High | High | High | High | High | High |
| | 2024 | 1312 | 2807 | 2110 | High | High | High | High | Not Founc | Not Founc | Not Founc | Not Founc | High | High |
| 8733 | 9798 | 6002 | 13184 | 13180 | High | High | High | High | High | High | High | High | High | High |
| 942 | 2367 | 1889 | 2583 | 2596 | High | High | High | High | High | High | High | High | High | High |
| 6181 | 4000 | 2265 | 3235 | 3309 | High | High | High | High | High | High | High | High | High | High |
| 5734 | 2810 | 2570 | 4058 | 3795 | High | High | High | High | High | High | High | High | High | High |
| | 3364 | 2946 | 4732 | 4620 | High | High | High | High | Not Founc | Not Founc | Not Founc | Not Founc | High | High |
| 8721 | 11521 | 13035 | 9571 | 18735 | Not Founc | Not Founc | Not Founc | Not Founc | High | High | High | High | High | High |
| 4760 | 6951 | 2883 | 7022 | 4034 | High | High | High | High | High | High | High | High | High | High |

| | | | | | | | | | | | | | | |
|---|---|---|---|---|---|---|---|---|---|---|---|---|---|---|
| 13693 | 5029 | 3858 | 5114 | 4364 | Not Founc | Not Founc | Not Founc | Not Founc | High | High | High | High | High | High |
| 5663 | 2946 | 2321 | 3581 | 3787 | Not Founc | Not Founc | Not Founc | Not Founc | High | High | High | High | High | High |
| 1573 | 5656 | 3229 | 6790 | 5244 | High | High | High | High | High | High | High | High | High | High |
| 12280 | 3702 | 2739 | 3665 | 4052 | High | High | High | High | High | High | High | High | High | High |
| | 2467 | 2016 | 2783 | 3177 | High | High | High | High | Not Founc | Not Founc | Not Founc | Not Founc | High | High |
| 7387 | 3633 | 2512 | 7558 | 6335 | High | High | High | High | High | High | High | High | High | High |
| 3526 | 8926 | 6505 | 9199 | 9731 | Not Founc | Not Founc | Not Founc | Not Founc | High | High | High | High | High | High |
| 3126 | 6398 | 4100 | 4884 | 3213 | High | High | High | High | High | High | High | High | High | High |
| 3755 | 3695 | 9001 | 4572 | 3970 | High | High | High | High | High | High | High | High | High | High |
| 1161 | 6877 | 4607 | 7437 | 6858 | Not Founc | Not Founc | Not Founc | Not Founc | High | High | High | High | High | High |
| 1384 | 2116 | 1352 | 3531 | 2821 | High | High | High | High | High | High | High | High | High | High |
| 2469 | 2226 | 1545 | 2528 | 2394 | High | High | High | High | High | High | High | High | High | High |
| 2576 | 3060 | 2070 | 3422 | 3782 | High | High | High | High | High | High | High | High | High | High |
| 3667 | 5461 | 3909 | 9821 | 7459 | High | High | High | High | High | High | High | High | High | High |
| 6873 | 2895 | 1594 | 4318 | 2790 | High | High | High | High | High | High | High | High | High | High |
| 3233 | 4747 | 3473 | 4344 | 4404 | High | High | High | High | High | High | High | High | High | High |
| | 4185 | 3444 | 4456 | 4576 | High | High | High | High | Not Founc | Not Founc | Not Founc | Not Founc | High | High |
| 1919 | | | | | High | High | High | High | High | High | High | High | Not Founc | Not Founc |
| 11939 | | | | | High | High | High | High | High | High | High | High | Not Founc | Not Founc |
| 4009 | 2282 | 1939 | 2516 | 2705 | High | High | High | High | High | High | High | High | High | High |
| | 19339 | 13217 | 20352 | 18967 | High | High | High | High | Not Founc | Not Founc | Not Founc | Not Founc | High | High |
| 7656 | 2447 | 2050 | 3376 | 4101 | Not Founc | Not Founc | Not Founc | Not Founc | High | High | High | High | High | High |
| 4216 | 138 | 95 | 130 | 176 | High | High | High | High | High | High | High | High | High | High |
| 4089 | 5352 | 4786 | 5599 | 6933 | Not Founc | Not Founc | Not Founc | Not Founc | High | High | High | High | High | High |
| 2289 | 1432 | 1238 | 1635 | 2044 | High | High | High | High | High | High | High | High | High | High |
| 8508 | | | | | High | High | High | High | High | High | High | High | Not Founc | Not Founc |
| 2032 | 1960 | 1564 | 2283 | 2402 | High | High | High | High | High | High | High | High | High | High |
| 2478 | 1666 | 1224 | 1671 | 1849 | High | High | High | High | High | High | High | High | High | High |
| | 5826 | 2805 | 6583 | 3910 | High | High | High | High | Not Founc | Not Founc | Not Founc | Not Founc | High | High |
| | 628 | 553 | 575 | 1061 | High | High | High | High | Not Founc | Not Founc | Not Founc | Not Founc | High | High |
| 638 | 1702 | 1575 | 2005 | 2212 | High | High | High | High | High | High | High | High | High | High |

| | | | | | | | | | | | | | | |
|---|---|---|---|---|---|---|---|---|---|---|---|---|---|---|
| 4940 | | | | | High | High | High | High | High | High | High | High | Not Founc | Not Founc |
| | 7677 | 4840 | 12213 | 8109 | High | High | High | High | Not Founc | Not Founc | Not Founc | Not Founc | High | High |
| 2114 | 8606 | 4563 | 11100 | 7920 | High | High | High | High | High | High | High | High | High | High |
| 4045 | 1148 | 493 | 823 | 844 | High | High | High | High | High | High | High | High | High | High |
| 4623 | 2733 | 1969 | 4162 | 3524 | High | High | High | High | High | High | High | High | High | High |
| 3747 | 4656 | 3387 | 5197 | 4458 | High | High | High | High | High | High | High | High | High | High |
| 1133 | 2707 | 1947 | 3658 | 3038 | Not Founc | Not Founc | Not Founc | Not Founc | High | High | High | High | High | High |
| 1089 | 9242 | 8151 | 7849 | 9587 | High | High | High | High | High | High | High | High | High | High |
| 9315 | 2395 | 1490 | 5014 | 3553 | High | High | High | High | High | High | High | High | High | High |
| 3817 | 3884 | 3096 | 2935 | 3679 | High | High | High | High | High | High | High | High | High | High |
| | 5891 | 5025 | 5484 | 6739 | High | High | High | High | Not Founc | Not Founc | Not Founc | Not Founc | High | High |
| 9696 | 9051 | 6420 | 11497 | 9011 | High | High | High | High | High | High | High | High | High | High |
| | 9871 | 6586 | 9191 | 20128 | High | High | High | High | Not Founc | Not Founc | Not Founc | Not Founc | High | High |
| 4513 | 4405 | 3005 | 6842 | 5472 | Not Founc | Not Founc | Not Founc | Not Founc | High | High | High | High | High | High |
| 3456 | 5623 | 2794 | 13684 | 7880 | High | High | High | High | High | High | High | High | High | High |
| 11217 | | | | | High | High | High | High | High | High | High | High | Not Founc | Not Founc |
| 4704 | 1870 | 1655 | 2151 | 2556 | High | High | High | High | High | High | High | High | High | High |
| | 3412 | 2696 | 5770 | 5496 | High | High | High | High | Not Founc | Not Founc | Not Founc | Not Founc | High | High |
| | 4072 | 3154 | 5867 | 4461 | High | High | High | High | Not Founc | Not Founc | Not Founc | Not Founc | High | High |
| 3206 | 8853 | 7700 | 9488 | 10269 | High | High | High | High | High | High | High | High | High | High |
| 8619 | 176 | 136 | 294 | 281 | Not Founc | Not Founc | Not Founc | Not Founc | High | High | High | High | High | High |
| 2241 | 4171 | 3292 | 4218 | 3968 | High | High | High | High | High | High | High | High | High | High |
| 8360 | | | | | High | High | High | High | High | High | High | High | Not Founc | Not Founc |
| 4493 | 2434 | 1225 | 2785 | 2398 | High | High | High | High | High | High | High | High | High | High |
| 5977 | 1620 | 1510 | 2221 | 2828 | High | High | High | High | High | High | High | High | High | High |
| | 6273 | 4931 | 6567 | 7006 | High | High | High | High | Not Founc | Not Founc | Not Founc | Not Founc | High | High |
| | | | | | High | High | High | High | Not Founc | Not Founc | Not Founc | Not Founc | Not Founc | Not Founc |
| 2342 | 2473 | 1572 | 2110 | 2207 | High | High | High | High | High | High | High | High | High | High |
| 7561 | 2508 | 1628 | 3518 | 3158 | High | High | High | High | High | High | High | High | High | High |
| 1781 | 838 | 707 | 905 | 1097 | High | High | High | High | High | High | High | High | High | High |
| 691 | 3666 | 2106 | 5288 | 3586 | High | High | High | High | High | High | High | High | High | High |

| | | | | | | | | | | | | | | |
|---|---|---|---|---|---|---|---|---|---|---|---|---|---|---|
| 850 | 5374 | 4413 | 5644 | 5434 | Not Founc | Not Founc | Not Founc | Not Founc | High | High | High | High | High | High |
| 2507 | 4939 | 3323 | 6096 | 4983 | High | High | High | High | High | High | High | High | High | High |
| 2953 | 2212 | 2463 | 3837 | 2759 | High | High | High | High | High | High | High | High | High | High |
| 803 | | | | | High | High | High | High | High | High | High | High | Not Founc | Not Founc |
| 5304 | 3089 | 4280 | 3967 | 4399 | High | High | High | High | High | High | High | High | High | High |
| 9452 | 2832 | 1971 | 3595 | 5007 | High | High | High | High | High | High | High | High | High | High |
| 4423 | 4988 | 3623 | 3852 | 4485 | High | High | High | High | High | High | High | High | High | High |
| 5062 | 2124 | 1825 | 3809 | 2922 | High | High | High | High | High | High | High | High | High | High |
| 1889 | 3998 | 3022 | 6020 | 4806 | High | High | High | High | High | High | High | High | High | High |
| | | | | | Not Founc | Not Founc | Not Founc | Not Founc | High | High | High | High | Not Founc | Not Founc |
| | 2962 | 1729 | 4879 | 3083 | High | High | High | High | Not Founc | Not Founc | Not Founc | Not Founc | High | High |
| 6200 | 2955 | 1452 | 2161 | 1994 | High | High | High | High | High | High | High | High | High | High |
| 1511 | 1819 | 1159 | 2382 | 1819 | High | High | High | High | High | High | High | High | High | High |
| | 1267 | 636 | 1323 | 1257 | High | High | High | High | Not Founc | Not Founc | Not Founc | Not Founc | High | High |
| | 5946 | 4483 | 6529 | 7727 | High | High | High | High | Not Founc | Not Founc | Not Founc | Not Founc | High | High |
| 7557 | 2000 | 1380 | 1869 | 1878 | High | High | High | High | High | High | High | High | High | High |
| | 5057 | 3533 | 8532 | 7606 | High | High | High | High | Not Founc | Not Founc | Not Founc | Not Founc | High | High |
| 3620 | 5420 | 2186 | 8246 | 3925 | Not Founc | Not Founc | Not Founc | Not Founc | High | High | High | High | High | High |
| 5450 | 4728 | 3788 | 5773 | 6714 | High | High | High | High | High | High | High | High | High | High |
| 6709 | | | | | High | High | High | High | High | High | High | High | Not Founc | Not Founc |
| 4707 | 1326 | 1205 | 1335 | 1797 | High | High | High | High | High | High | High | High | High | High |
| 2518 | 1251 | 1464 | 6969 | 3475 | High | High | High | High | High | High | High | High | High | High |
| 6380 | 1832 | 1623 | 1843 | 2523 | High | High | High | High | High | High | High | High | High | High |
| | 6465 | 4669 | 8220 | 7239 | High | High | High | High | Not Founc | Not Founc | Not Founc | Not Founc | High | High |
| 6404 | 8509 | 5130 | 7476 | 6141 | High | High | High | High | High | High | High | High | High | High |
| 1517 | 1494 | 1229 | 2415 | 2142 | High | High | High | High | High | High | High | High | High | High |
| 4532 | 3801 | 2190 | 4176 | 3852 | High | High | High | High | High | High | High | High | High | High |
| 1162 | 3428 | 2765 | 3636 | 5119 | High | High | High | High | High | High | High | High | High | High |
| 5690 | | | | | High | High | High | High | High | High | High | High | Not Founc | Not Founc |
| 12623 | 4026 | 3150 | 5659 | 7271 | High | High | High | High | High | High | High | High | High | High |
| 2943 | 1512 | 1144 | 1488 | 1630 | High | High | High | High | High | High | High | High | High | High |

| | | | | | | | | | | | | | | |
|---|---|---|---|---|---|---|---|---|---|---|---|---|---|---|
| 7623 | 3315 | 2616 | 3279 | 3876 | High | High | High | High | High | High | High | High | High | High |
| 7327 | 4426 | 3320 | 5712 | 5372 | High | High | High | High | High | High | High | High | High | High |
| 6207 | 2329 | 2405 | 2113 | 2754 | High | High | High | High | High | High | High | High | High | High |
| 875 | | | | | High | High | High | High | High | High | High | High | Not Founc | Not Founc |
| 11256 | 15301 | 12257 | 20291 | 19460 | High | High | High | High | High | High | High | High | High | High |
| 6632 | 4275 | 2997 | 5009 | 4060 | High | High | High | High | High | High | High | High | High | High |
| 6617 | 2362 | 2102 | 2547 | 3131 | High | High | High | High | High | High | High | High | High | High |
| 3505 | 4026 | 3540 | 5678 | 6153 | Not Founc | Not Founc | Not Founc | Not Founc | High | High | High | High | High | High |
| 2377 | 1433 | 1108 | 1509 | 1365 | High | High | High | High | High | High | High | High | High | High |
| 9099 | | | | | High | High | High | High | High | High | High | High | Not Founc | Not Founc |
| | 3987 | 2906 | 7265 | 6788 | High | High | High | High | Not Founc | Not Founc | Not Founc | Not Founc | High | High |
| 8906 | 4998 | 3240 | 11478 | 9254 | High | High | High | High | High | High | High | High | High | High |
| 2110 | 1317 | 1131 | 1525 | 1487 | High | High | High | High | High | High | High | High | High | High |
| 3976 | 4151 | 3983 | 4510 | 5355 | High | High | High | High | High | High | High | High | High | High |
| 2764 | 2270 | 1334 | 2264 | 1759 | High | High | High | High | High | High | High | High | High | High |
| 1338 | 1794 | 1405 | 1751 | 2002 | High | High | High | High | High | High | High | High | High | High |
| 4792 | 1407 | 968 | 1991 | 1780 | High | High | High | High | High | High | High | High | High | High |
| 1091 | | | | | High | High | High | High | High | High | High | High | Not Founc | Not Founc |
| 250 | 15702 | | 14186 | 282 | High | High | High | High | High | Not Founc | High | High | High | Not Founc |
| 899 | 654 | 417 | 1216 | 885 | High | High | High | High | High | High | High | High | High | High |
| | 3406 | 2310 | 4425 | 3709 | Not Founc | Not Founc | Not Founc | Not Founc | Not Founc | Not Founc | Not Founc | Not Founc | High | High |
| 1068 | 1465 | 1391 | 1073 | 1309 | High | High | High | High | High | High | High | High | High | High |
| 3524 | 12892 | 8859 | 15877 | 15226 | High | High | High | High | High | High | High | High | High | High |
| 4597 | 4796 | 3908 | 7217 | 5397 | High | High | High | High | High | High | High | High | High | High |
| 1694 | | | | | High | High | High | High | High | High | High | High | Not Founc | Not Founc |
| 4315 | 2051 | 1330 | 2906 | 3078 | High | High | High | High | High | High | High | High | High | High |
| 7798 | 17688 | 10488 | 16846 | 14227 | Not Founc | Not Founc | Not Founc | Not Founc | High | High | High | High | High | High |
| 4167 | 3697 | 1094 | 2837 | 2335 | High | High | High | High | High | High | High | High | High | High |
| 5086 | 1930 | 1445 | 2870 | 2233 | High | High | High | High | High | High | High | High | High | High |
| 323 | | | | | High | High | High | High | High | High | High | High | Not Founc | Not Founc |
| | 13564 | 9022 | 13425 | 12056 | High | High | High | High | Not Founc | Not Founc | Not Founc | Not Founc | High | High |

| | | | | | | | | | | | | | | |
|---|---|---|---|---|---|---|---|---|---|---|---|---|---|---|
| 2481 | | | | | High | High | High | High | High | High | High | High | Not Found | Not Found |
| 574 | 1728 | 1046 | 1857 | 1461 | High | High | High | High | High | High | High | High | High | High |
| | 1693 | 1622 | 2041 | 2239 | High | High | High | High | Not Found | Not Found | Not Found | Not Found | High | High |
| 439 | | | | | High | High | High | High | High | High | High | High | Not Found | Not Found |
| 3793 | | | | | High | High | High | High | High | High | High | High | Not Found | Not Found |
| 2122 | 867 | 943 | 806 | 1402 | High | High | High | High | High | High | High | High | High | High |
| 2301 | 832 | 1331 | 549 | 1428 | High | High | High | High | High | High | High | High | High | High |
| 2377 | | | | | High | High | High | High | High | High | High | High | Not Found | Not Found |
| 3685 | 6552 | 4535 | 5663 | 5968 | High | High | High | High | High | High | High | High | High | High |
| 534 | 2744 | 2064 | 2730 | 2902 | Not Found | Not Found | Not Found | Not Found | High | High | High | High | High | High |
| 1924 | 1667 | 1062 | 2732 | 2355 | High | High | High | High | High | High | High | High | High | High |
| 1460 | 1994 | 1582 | 2113 | 2353 | High | High | High | High | High | High | High | High | High | High |
| 968 | 8085 | 6493 | 5885 | 6820 | Not Found | Not Found | Not Found | Not Found | High | High | High | High | High | High |
| 968 | 2317 | 2062 | 2992 | 3174 | High | High | High | High | High | High | High | High | High | High |
| 1704 | 1545 | 1119 | 2183 | 1738 | High | High | High | High | High | High | High | High | High | High |
| | 2960 | 2327 | 2584 | 3070 | High | High | High | High | Not Found | Not Found | Not Found | Not Found | High | High |
| 9714 | | | | | High | High | High | High | High | High | High | High | Not Found | Not Found |
| 1031 | 4881 | 3501 | 4766 | 5442 | Not Found | Not Found | Not Found | Not Found | High | High | High | High | High | High |
| 4254 | 998 | 578 | 981 | 731 | High | High | High | High | High | High | High | High | High | High |
| 5818 | | | | | Not Found | Not Found | Not Found | Not Found | High | High | High | High | Not Found | Not Found |
| 2219 | | | | | High | High | High | High | High | High | High | High | Not Found | Not Found |
| 1609 | 2028 | 1558 | 1782 | 1973 | High | High | High | High | High | High | High | High | High | High |
| 995 | 670 | 632 | 725 | 918 | High | High | High | High | High | High | High | High | High | High |
| 2060 | 2151 | 1264 | 1944 | 2187 | High | High | High | High | High | High | High | High | High | High |
| | | | | | High | High | High | High | Not Found | Not Found | Not Found | Not Found | Not Found | Not Found |
| | 8825 | 5547 | 12552 | 10128 | Not Found | Not Found | Not Found | Not Found | Not Found | Not Found | Not Found | Not Found | High | High |
| 3789 | 7680 | 6308 | 7381 | 8368 | High | High | High | High | High | High | High | High | High | High |
| 6442 | 2817 | 2268 | 3801 | 3582 | High | High | High | High | High | High | High | High | High | High |
| 7925 | 7165 | 3993 | 6318 | 5551 | High | High | High | High | High | High | High | High | High | High |
| | 3520 | 2167 | 7557 | 5485 | High | High | High | High | Not Found | Not Found | Not Found | Not Found | High | High |
| 6478 | | | | | High | High | High | High | High | High | High | High | Not Found | Not Found |

| | | | | | | | | | | | | | | |
|---|---|---|---|---|---|---|---|---|---|---|---|---|---|---|
| | 6408 | 4759 | 11492 | 6341 | High | High | High | High | Not Found | Not Found | Not Found | Not Found | High | High |
| 4652 | 2931 | 1827 | 4191 | 3616 | Not Found | Not Found | Not Found | Not Found | High | High | High | High | High | High |
| | 5711 | 3667 | 6567 | 4993 | High | High | High | High | Not Found | Not Found | Not Found | Not Found | High | High |
| 3383 | 8116 | 4361 | 12949 | 8060 | High | High | High | High | High | High | High | High | High | High |
| 1962 | 2017 | 1173 | 2109 | 1791 | Not Found | Not Found | Not Found | Not Found | High | High | High | High | High | High |
| 11911 | | | | | High | High | High | High | High | High | High | High | Not Found | Not Found |
| 6337 | | | | | High | High | High | High | High | High | High | High | Not Found | Not Found |
| | 2802 | 2156 | 2680 | 2643 | High | High | High | High | Not Found | Not Found | Not Found | Not Found | High | High |
| 4095 | 7760 | 5083 | 12393 | 9070 | High | High | High | High | High | High | High | High | High | High |
| 2209 | 2557 | 2121 | 3037 | 3369 | Not Found | Not Found | Not Found | Not Found | High | High | High | High | High | High |
| | 2894 | 1963 | 7298 | 5235 | High | High | High | High | Not Found | Not Found | Not Found | Not Found | High | High |
| 1253 | 4710 | 5029 | 7151 | 6677 | High | High | High | High | High | High | High | High | High | High |
| 1974 | 5282 | 3699 | 8103 | 6543 | High | High | High | High | High | High | High | High | High | High |
| 2406 | 1687 | 1005 | 1982 | 1523 | High | High | High | High | High | High | High | High | High | High |
| 2891 | 2447 | 1555 | 4643 | 3201 | Not Found | Not Found | Not Found | Not Found | High | High | High | High | High | High |
| 5283 | 2260 | 1883 | 2918 | 3169 | High | High | High | High | High | High | High | High | High | High |
| 4600 | | | | | High | High | High | High | High | High | High | High | Not Found | Not Found |
| 1484 | 1538 | 1025 | 1739 | 1538 | High | High | High | High | High | High | High | High | High | High |
| 1373 | 2633 | 1770 | 3165 | 2791 | High | High | High | High | High | High | High | High | High | High |
| 1219 | 1039 | 804 | 919 | 1139 | High | High | High | High | High | High | High | High | High | High |
| 1587 | 411 | 252 | 427 | 327 | Not Found | Not Found | Not Found | Not Found | High | High | High | High | High | High |
| 2713 | 1062 | 832 | 1125 | 1227 | High | High | High | High | High | High | High | High | High | High |
| 7684 | 7170 | 3915 | 10510 | 8196 | Not Found | Not Found | Not Found | Not Found | High | High | High | High | High | High |
| 4670 | 1532 | 867 | 1384 | 1534 | High | High | High | High | High | High | High | High | High | High |
| 4323 | 3964 | 3423 | 7224 | 6338 | High | High | High | High | High | High | High | High | High | High |
| 7821 | | | | | High | High | High | High | High | High | High | High | Not Found | Not Found |
| | 11371 | 6661 | 15860 | 10218 | High | High | High | High | Not Found | Not Found | Not Found | Not Found | High | High |
| 669 | | | | | High | High | High | High | High | High | High | High | Not Found | Not Found |
| 6167 | 3731 | 1019 | 3756 | 1949 | High | High | High | High | High | High | High | High | High | High |
| | 1493 | 937 | 1299 | 957 | High | High | High | High | Not Found | Not Found | Not Found | Not Found | High | High |
| 5392 | 2392 | 1539 | 3810 | 2614 | High | High | High | High | High | High | High | High | High | High |

| | | | | | | | | | | | | | | |
|---|---|---|---|---|---|---|---|---|---|---|---|---|---|---|
| | 7178 | 4374 | 9385 | 7645 | High | High | High | High | Not Founc | Not Founc | Not Founc | Not Founc | High | High |
| 4633 | | | | | High | High | High | High | High | High | High | High | Not Founc | Not Founc |
| | 4843 | 4107 | 4242 | 4716 | High | High | High | High | Not Founc | Not Founc | Not Founc | Not Founc | High | High |
| | 1909 | 1659 | 2466 | 2478 | High | High | High | High | Not Founc | Not Founc | Not Founc | Not Founc | High | High |
| 2878 | | | | | High | High | High | High | High | High | High | High | Not Founc | Not Founc |
| 3304 | 5599 | 3815 | 5898 | 5730 | High | High | High | High | High | High | High | High | High | High |
| 1873 | | | | | High | High | High | High | High | High | High | High | Not Founc | Not Founc |
| 2039 | | | | | High | High | High | High | High | High | High | High | Not Founc | Not Founc |
| 8168 | 2189 | 1952 | 2187 | 2716 | High | High | High | High | High | High | High | High | High | High |
| 4530 | 2239 | 1938 | 2082 | 2187 | High | High | High | High | High | High | High | High | High | High |
| 1446 | 2259 | 1131 | 2935 | 2060 | High | High | High | High | High | High | High | High | High | High |
| 4693 | 2961 | 2758 | 3614 | 3887 | High | High | High | High | High | High | High | High | High | High |
| 914 | 614 | 549 | 481 | 681 | High | High | High | High | High | High | High | High | High | High |
| 3510 | 3491 | 2710 | 4263 | 3334 | High | High | High | High | High | High | High | High | High | High |
| 6439 | 4941 | 3481 | 6065 | 5623 | High | High | High | High | High | High | High | High | High | High |
| | 9943 | 7308 | 10475 | 10083 | High | High | High | High | Not Founc | Not Founc | Not Founc | Not Founc | High | High |
| 7182 | 1614 | 1148 | 1650 | 1571 | High | High | High | High | High | High | High | High | High | High |
| 4109 | 1928 | 1568 | 2283 | 2434 | High | High | High | High | High | High | High | High | High | High |
| | 1368 | 763 | 1379 | 1194 | High | High | High | High | Not Founc | Not Founc | Not Founc | Not Founc | High | High |
| 2925 | | | | | High | High | High | High | High | High | High | High | Not Founc | Not Founc |
| 3253 | 2901 | 1836 | 4083 | 3164 | Not Founc | Not Founc | Not Founc | Not Founc | High | High | High | High | High | High |
| 7305 | 6812 | 5363 | 6799 | 8056 | High | High | High | High | High | High | High | High | High | High |
| 2943 | 3065 | 2229 | 3392 | 3204 | High | High | High | High | High | High | High | High | High | High |
| | 1057 | 889 | 889 | 1071 | High | High | High | High | Not Founc | Not Founc | Not Founc | Not Founc | High | High |
| | 3436 | 3169 | 4015 | 4867 | High | High | High | High | Not Founc | Not Founc | Not Founc | Not Founc | High | High |
| | 7697 | 4870 | 16572 | 15094 | High | High | High | High | Not Founc | Not Founc | Not Founc | Not Founc | High | High |
| 12891 | 4042 | 3916 | 6708 | 7053 | High | High | High | High | High | High | High | High | High | High |
| 2468 | | | | | High | High | High | High | High | High | High | High | Not Founc | Not Founc |
| | 2207 | 1773 | 1959 | 1905 | High | High | High | High | Not Founc | Not Founc | Not Founc | Not Founc | High | High |
| 5730 | 1911 | 1460 | 3792 | 2867 | High | High | High | High | High | High | High | High | High | High |
| 511 | 396 | 307 | 373 | 396 | High | High | High | High | High | High | High | High | High | High |

| | | | | | | | | | | | | | | |
|---|---|---|---|---|---|---|---|---|---|---|---|---|---|---|
| 455 | | | | | High | High | High | High | High | High | High | High | Not Founc | Not Founc |
| 692 | 2716 | 2216 | 5307 | 3167 | High | High | High | High | High | High | High | High | High | High |
| 1153 | | | | | High | High | High | High | High | High | High | High | Not Founc | Not Founc |
| 4599 | 1315 | 947 | 1913 | 1567 | High | High | High | High | High | High | High | High | High | High |
| 1034 | 5310 | 4591 | 5448 | 6999 | High | High | High | High | High | High | High | High | High | High |
| 2510 | 2622 | 2204 | 2938 | 2953 | Not Founc | Not Founc | Not Founc | Not Founc | High | High | High | High | High | High |
| 3288 | | | | | High | High | High | High | High | High | High | High | Not Founc | Not Founc |
| 3202 | 4046 | 2827 | 5141 | 4238 | High | High | High | High | High | High | High | High | High | High |
| 8826 | 2435 | 1907 | 3310 | 3350 | Not Founc | Not Founc | Not Founc | Not Founc | High | High | High | High | High | High |
| 2264 | | | | | High | High | High | High | High | High | High | High | Not Founc | Not Founc |
| 4111 | 5473 | 3852 | 5926 | 5178 | High | High | High | High | High | High | High | High | High | High |
| 2911 | 5920 | 4305 | 6877 | 6522 | High | High | High | High | High | High | High | High | High | High |
| | 3025 | 2500 | 3132 | 2782 | High | High | High | High | Not Founc | Not Founc | Not Founc | Not Founc | High | High |
| 3133 | 5923 | 3078 | 8366 | 5859 | High | High | High | High | High | High | High | High | High | High |
| 3028 | 1512 | 994 | 1092 | 1498 | High | High | High | High | High | High | High | High | High | High |
| 8321 | 1627 | 493 | 2211 | 1203 | High | High | High | High | High | High | High | High | High | High |
| 896 | 674 | 543 | 902 | 843 | High | High | High | High | High | High | High | High | High | High |
| 4562 | 4095 | 3768 | 3718 | 5261 | Not Founc | Not Founc | Not Founc | Not Founc | High | High | High | High | High | High |
| 4752 | 7289 | 5529 | 12714 | 9534 | Not Founc | Not Founc | Not Founc | Not Founc | High | High | High | High | High | High |
| 2908 | | | | | High | High | High | High | High | High | High | High | Not Founc | Not Founc |
| | 1861 | 1087 | 2756 | 2431 | High | High | High | High | Not Founc | Not Founc | Not Founc | Not Founc | High | High |
| | 5676 | 5091 | 7078 | 8674 | High | High | High | High | Not Founc | Not Founc | Not Founc | Not Founc | High | High |
| 10141 | | | | | High | High | High | High | High | High | High | High | Not Founc | Not Founc |
| | 3786 | 2429 | 7227 | 4768 | High | High | High | High | Not Founc | Not Founc | Not Founc | Not Founc | High | High |
| 838 | 968 | 701 | 1906 | 1346 | High | High | High | High | High | High | High | High | High | High |
| 5012 | | | | | High | High | High | High | High | High | High | High | Not Founc | Not Founc |
| 2849 | | | | | High | High | High | High | High | High | High | High | Not Founc | Not Founc |
| 8084 | | | | | High | High | High | High | High | High | High | High | Not Founc | Not Founc |
| 5044 | 3071 | 2860 | 2880 | 4126 | High | High | High | High | High | High | High | High | High | High |
| 4967 | 825 | 374 | 5543 | 1422 | High | High | High | High | High | High | High | High | High | High |
| 3304 | 1829 | 1357 | 2589 | 2345 | Not Founc | Not Founc | Not Founc | Not Founc | High | High | High | High | High | High |

| | | | | | | | | | | | | | | |
|---|---|---|---|---|---|---|---|---|---|---|---|---|---|---|
| 3956 | 3338 | 2443 | 3117 | 3526 | Not Founc | Not Founc | Not Founc | Not Founc | High | High | High | High | High | High |
| 6876 | | | | | High | High | High | High | High | High | High | High | Not Founc | Not Founc |
| 2886 | | | | | High | High | High | High | High | High | High | High | Not Founc | Not Founc |
| 590 | 2808 | 1439 | 5640 | 3826 | High | High | High | High | High | High | High | High | High | High |
| 2121 | | | | | High | High | High | High | High | High | High | High | Not Founc | Not Founc |
| 3196 | 3381 | 2943 | 3113 | 4057 | High | High | High | High | High | High | High | High | High | High |
| 4576 | 1381 | 827 | 1530 | 6905 | High | High | High | High | High | High | High | High | High | High |
| 4567 | 617 | 590 | 868 | 924 | High | High | High | High | High | High | High | High | High | High |
| 3038 | 2759 | 1887 | 4706 | 4206 | High | High | High | High | High | High | High | High | High | High |
| 2750 | 1336 | 858 | 1401 | 1445 | High | High | High | High | High | High | High | High | High | High |
| 1596 | | | | | Not Founc | Not Founc | Not Founc | Not Founc | High | High | High | High | Not Founc | Not Founc |
| 4257 | 6910 | 5328 | 6686 | 7637 | High | High | High | High | High | High | High | High | High | High |
| 974 | 2717 | 1850 | 3014 | 2864 | High | High | High | High | High | High | High | High | High | High |
| 6905 | 18305 | 10546 | 14365 | 14936 | High | High | High | High | High | High | High | High | High | High |
| 2443 | 2019 | 1667 | 1996 | 2324 | High | High | High | High | High | High | High | High | High | High |
| 1668 | 2720 | 1977 | 5962 | 3612 | High | High | High | High | High | High | High | High | High | High |
| 1627 | 866 | 690 | 1109 | 990 | Not Founc | Not Founc | Not Founc | Not Founc | High | High | High | High | High | High |
| 1376 | 982 | 824 | 1221 | 1196 | High | High | High | High | High | High | High | High | High | High |
| 7671 | 2392 | 1529 | 5706 | 4108 | Not Founc | Not Founc | Not Founc | Not Founc | High | High | High | High | High | High |
| 7055 | | | | | High | High | High | High | High | High | High | High | Not Founc | Not Founc |
| 2933 | 4805 | 4214 | 6295 | 5521 | High | High | High | High | High | High | High | High | High | High |
| 2329 | 1195 | 714 | 1229 | 1099 | Not Founc | Not Founc | Not Founc | Not Founc | High | High | High | High | High | High |
| 2644 | | | | | High | High | High | High | High | High | High | High | Not Founc | Not Founc |
| 2392 | 5359 | 4668 | 5324 | 7055 | Not Founc | Not Founc | Not Founc | Not Founc | High | High | High | High | High | High |
| 4359 | 1200 | 837 | 2087 | 1148 | High | High | High | High | High | High | High | High | High | High |
| 9656 | 4088 | 2464 | 6106 | 4093 | High | High | High | High | High | High | High | High | High | High |
| | 3155 | 2847 | 3214 | 4040 | High | High | High | High | Not Founc | Not Founc | Not Founc | Not Founc | High | High |
| 10903 | 1686 | 1060 | 5473 | 2999 | High | High | High | High | High | High | High | High | High | High |
| 7460 | 3350 | 1999 | 3797 | 3440 | High | High | High | High | High | High | High | High | High | High |
| 920 | 869 | 638 | 710 | 800 | High | High | High | High | High | High | High | High | High | High |
| 2548 | | | | | High | High | High | High | High | High | High | High | Not Founc | Not Founc |

| | | | | | | | | | | | | | | |
|---|---|---|---|---|---|---|---|---|---|---|---|---|---|---|
| | 2053 | 1532 | 2458 | 2499 | High | High | High | High | Not Founc | Not Founc | Not Founc | Not Founc | High | High |
| | 6396 | 6044 | 6503 | 7619 | High | High | High | High | Not Founc | Not Founc | Not Founc | Not Founc | High | High |
| 5328 | 311 | 242 | 321 | 338 | High | High | High | High | High | High | High | High | High | High |
| | 1530 | 1132 | 2278 | 1745 | High | High | High | High | Not Founc | Not Founc | Not Founc | Not Founc | High | High |
| 1400 | 385 | 368 | 297 | 368 | Not Founc | Not Founc | Not Founc | Not Founc | High | High | High | High | High | High |
| 5061 | 4038 | 3221 | 4154 | 5229 | High | High | High | High | High | High | High | High | High | High |
| 1489 | 1372 | 944 | 3073 | 2184 | High | High | High | High | High | High | High | High | High | High |
| 2183 | | | | | High | High | High | High | High | High | High | High | Not Founc | Not Founc |
| 25 | | | | | Not Founc | Not Founc | Not Founc | Not Founc | Not Founc | High | Not Founc | High | Not Founc | Not Founc |
| | 1901 | 1270 | 2636 | 2317 | High | High | High | High | Not Founc | Not Founc | Not Founc | Not Founc | High | High |
| 3127 | 3011 | 1744 | 3651 | 3374 | High | High | High | High | High | High | High | High | High | High |
| 1332 | 2819 | 2602 | 6392 | 3921 | High | High | High | High | High | High | High | High | High | High |
| 1410 | 4073 | 2506 | 5780 | 4105 | High | High | High | High | High | High | High | High | High | High |
| 2102 | 3495 | 2318 | 4945 | 4825 | Not Founc | Not Founc | Not Founc | Not Founc | High | High | High | High | High | High |
| 2405 | | | | | High | High | High | High | High | High | High | High | Not Founc | Not Founc |
| 1724 | | | | | High | High | High | High | High | High | High | High | Not Founc | Not Founc |
| 2766 | 1831 | 1573 | 2323 | 2445 | High | High | High | High | High | High | High | High | High | High |
| 1550 | 2201 | 1292 | 5516 | 3069 | High | High | High | High | High | High | High | High | High | High |
| 1725 | 1826 | 1711 | 2913 | 2253 | Not Founc | Not Founc | Not Founc | Not Founc | High | High | High | High | High | High |
| 5129 | 2016 | 1078 | 2079 | 1481 | High | High | High | High | High | High | High | High | High | High |
| 4478 | | | | | High | High | High | High | High | High | High | High | Not Founc | Not Founc |
| 4684 | 2566 | 1860 | 2447 | 2297 | High | High | High | High | High | High | High | High | High | High |
| 2127 | 2210 | 1656 | 2697 | 2549 | Not Founc | Not Founc | Not Founc | Not Founc | High | High | High | High | High | High |
| 9072 | | | | | High | High | High | High | High | High | High | High | Not Founc | Not Founc |
| 7319 | 2487 | 1970 | 3044 | 3445 | High | High | High | High | High | High | High | High | High | High |
| 1290 | 1558 | 1338 | 1927 | 2026 | High | High | High | High | High | High | High | High | High | High |
| 231 | 228 | 235 | 252 | 201 | Not Founc | Not Founc | Not Founc | Not Founc | High | High | High | High | High | High |
| 10527 | 2641 | 1929 | 4351 | 3766 | High | High | High | High | High | High | High | High | High | High |
| 7866 | 2339 | 2481 | 2027 | 3250 | High | High | High | High | High | High | High | High | High | High |
| | 9024 | 6169 | 8328 | 10075 | High | High | High | High | Not Founc | Not Founc | Not Founc | Not Founc | High | High |
| | 1995 | 1376 | 2958 | 2255 | High | High | High | High | Not Founc | Not Founc | Not Founc | Not Founc | High | High |

| | | | | | | | | | | | | | | |
|---|---|---|---|---|---|---|---|---|---|---|---|---|---|---|
| | 5039 | 3577 | 6247 | 5249 | Not Found | Not Found | Not Found | Not Found | Not Found | Not Found | Not Found | Not Found | High | High |
| 2418 | 3550 | 3309 | 6648 | 4904 | High | High | High | High | High | High | High | High | High | High |
| 2833 | 2241 | 1380 | 1837 | 1874 | High | High | High | High | High | High | High | High | High | High |
| 1229 | | | | | High | High | High | High | High | High | High | High | Not Found | Not Found |
| 8379 | 3514 | 3012 | 4303 | 5323 | Not Found | Not Found | Not Found | Not Found | High | High | High | High | High | High |
| | 759 | 695 | 634 | 711 | High | High | High | High | Not Found | Not Found | Not Found | Not Found | High | High |
| 780 | 283 | 217 | 291 | 192 | Not Found | Not Found | Not Found | Not Found | High | High | High | High | High | High |
| 2521 | | | | | High | High | High | High | High | High | High | High | Not Found | Not Found |
| 5938 | 3235 | 2146 | 3990 | 3678 | Not Found | Not Found | Not Found | Not Found | High | High | High | High | High | High |
| 8376 | 3268 | 2736 | 4993 | 4047 | Not Found | Not Found | Not Found | Not Found | High | High | High | High | High | High |
| 842 | | | | | High | High | High | High | High | High | High | High | Not Found | Not Found |
| | | | | | High | High | High | High | Not Found | Not Found | Not Found | Not Found | High | High |
| 3054 | | | | | High | High | High | High | High | High | High | High | Not Found | Not Found |
| 2895 | 1049 | 734 | 1395 | 1034 | Not Found | Not Found | Not Found | Not Found | High | High | High | High | High | High |
| 4122 | 2759 | 2133 | 4253 | 5474 | High | High | High | High | High | High | High | High | High | High |
| | 5337 | 3408 | 5261 | 4573 | High | High | High | High | Not Found | Not Found | Not Found | Not Found | High | High |
| | 3331 | 2299 | 2950 | 2664 | High | High | High | High | Not Found | Not Found | Not Found | Not Found | High | High |
| 1898 | | | | | High | High | High | High | High | High | High | High | Not Found | Not Found |
| 4228 | | | | | High | High | High | High | High | High | High | High | Not Found | Not Found |
| 1916 | 2069 | 2088 | 1987 | 2722 | High | High | High | High | High | High | High | High | High | High |
| 4031 | 4474 | 3886 | 4917 | 5207 | High | High | High | High | High | High | High | High | High | High |
| | | | | | High | High | High | High | Not Found | Not Found | Not Found | Not Found | Not Found | Not Found |
| | 720 | 523 | 559 | 613 | High | High | High | High | Not Found | Not Found | Not Found | Not Found | High | High |
| 2496 | 1897 | 1361 | 2748 | 2386 | High | High | High | High | High | High | High | High | High | High |
| 1118 | | | | | High | High | High | High | High | High | High | High | Not Found | Not Found |
| 4227 | | | | | High | High | High | High | High | High | High | High | Not Found | Not Found |
| 2792 | 3225 | 2360 | 7630 | 3995 | Not Found | Not Found | Not Found | Not Found | High | High | High | High | High | High |
| | | | | | Not Found | Not Found | Not Found | Not Found | Not Found | Not Found | Not Found | Not Found | Not Found | Not Found |
| | 4909 | 3614 | 4600 | 4189 | High | High | High | High | Not Found | Not Found | Not Found | Not Found | High | High |
| 562 | | | | | High | High | High | High | High | High | High | High | Not Found | Not Found |
| 1552 | 1261 | 851 | 1701 | 1526 | High | High | High | High | High | High | High | High | High | High |

| | | | | | | | | | | | | | | |
|---|---|---|---|---|---|---|---|---|---|---|---|---|---|---|
| 9338 | | | | | Not Founc | Not Founc | Not Founc | Not Founc | High | High | High | High | Not Founc | Not Founc |
| 5243 | 3826 | 3218 | 3174 | 4376 | High | High | High | High | High | High | High | High | High | High |
| 3290 | | | | | High | High | High | High | High | High | High | High | Not Founc | Not Founc |
| 3866 | 8856 | 5135 | 10625 | 8387 | Not Founc | Not Founc | Not Founc | Not Founc | High | High | High | High | High | High |
| | 6577 | 5510 | 8856 | 8928 | High | High | High | High | Not Founc | Not Founc | Not Founc | Not Founc | High | High |
| 2345 | 1286 | 1039 | 2049 | 1727 | High | High | High | High | High | High | High | High | High | High |
| | 3192 | 1762 | 6305 | 3555 | High | High | High | High | Not Founc | Not Founc | Not Founc | Not Founc | High | High |
| 4512 | 1010 | 747 | 1072 | 1222 | High | High | High | High | High | High | High | High | High | High |
| 1236 | 2373 | 1794 | 2321 | 1941 | Not Founc | Not Founc | Not Founc | Not Founc | High | High | High | High | High | High |
| 4080 | 2512 | 2518 | 2302 | 3270 | Not Founc | Not Founc | Not Founc | Not Founc | High | High | High | High | High | High |
| 1958 | 3174 | 3045 | 2068 | 2341 | High | High | High | High | High | High | High | High | High | High |
| | 1716 | 1287 | 1969 | 1976 | High | High | High | High | Not Founc | Not Founc | Not Founc | Not Founc | High | High |
| 1722 | 2039 | 1857 | 2007 | 2346 | High | High | High | High | High | High | High | High | High | High |
| 1459 | 4208 | 2773 | 4298 | 4018 | High | High | High | High | High | High | High | High | High | High |
| 4305 | 4564 | 3015 | 3988 | 4145 | High | High | High | High | High | High | High | High | High | High |
| 3491 | 3120 | 1733 | 4114 | 3422 | Not Founc | Not Founc | Not Founc | Not Founc | High | High | High | High | High | High |
| 1587 | 2730 | 2213 | 3259 | 2955 | Not Founc | Not Founc | Not Founc | Not Founc | High | High | High | High | High | High |
| 1843 | 6470 | 2604 | 5149 | 3694 | High | High | High | High | High | High | High | High | High | High |
| | 7825 | 5893 | 8167 | 9002 | Not Founc | Not Founc | Not Founc | Not Founc | Not Founc | Not Founc | Not Founc | Not Founc | High | High |
| 823 | 320 | 165 | 260 | 250 | High | High | High | High | High | High | High | High | High | High |
| 577 | | | | | Not Founc | Not Founc | Not Founc | Not Founc | High | High | High | High | Not Founc | Not Founc |
| | 895 | 550 | 1088 | 707 | High | High | High | High | Not Founc | Not Founc | Not Founc | Not Founc | High | High |
| 679 | | | | | High | High | High | High | High | High | High | High | Not Founc | Not Founc |
| | 1658 | 1614 | 2104 | 2148 | High | High | High | High | Not Founc | Not Founc | Not Founc | Not Founc | High | High |
| 1332 | 1193 | 841 | 1314 | 1478 | High | High | High | High | High | High | High | High | High | High |
| | | | | | High | High | High | High | Not Founc | Not Founc | Not Founc | Not Founc | Not Founc | Not Founc |
| 3084 | 2001 | 1370 | 2788 | 2535 | High | High | High | High | High | High | High | High | High | High |
| 3687 | 4298 | 2701 | 8775 | 5492 | Not Founc | Not Founc | Not Founc | Not Founc | High | High | High | High | High | High |
| 4468 | 6053 | 5300 | 5892 | 7488 | High | High | High | High | High | High | High | High | High | High |
| | 1292 | 883 | 1979 | 1998 | High | High | High | High | Not Founc | Not Founc | Not Founc | Not Founc | High | High |
| 1919 | 2711 | 1808 | 2410 | 2366 | High | High | High | High | High | High | High | High | High | High |

| | | | | | | | | | | | | | | |
|---|---|---|---|---|---|---|---|---|---|---|---|---|---|---|
| | 5684 | 4415 | 5679 | 4566 | High | High | High | High | Not Founc | Not Founc | Not Founc | Not Founc | High | High |
| 5272 | | | | | High | High | High | High | High | High | High | High | Not Founc | Not Founc |
| | 2031 | 1754 | 3598 | 4271 | High | High | High | High | Not Founc | Not Founc | Not Founc | Not Founc | High | High |
| 3375 | 1108 | 762 | 1247 | 1103 | High | High | High | High | High | High | High | High | High | High |
| 1624 | | | | | High | High | High | High | High | High | High | High | Not Founc | Not Founc |
| 4261 | | | | | Not Founc | Not Founc | Not Founc | Not Founc | High | High | High | High | Not Founc | Not Founc |
| 2716 | | | | | Not Founc | Not Founc | Not Founc | Not Founc | High | High | High | High | Not Founc | Not Founc |
| 1655 | 510 | 497 | 494 | 482 | Not Founc | Not Founc | Not Founc | Not Founc | High | High | High | High | High | High |
| | 799 | 587 | 914 | 874 | High | High | High | High | Not Founc | Not Founc | Not Founc | Not Founc | High | High |
| | 3640 | 2689 | 4908 | 4475 | High | High | High | High | Not Founc | Not Founc | Not Founc | Not Founc | High | High |
| 2487 | 963 | 661 | 1213 | 991 | Not Founc | Not Founc | Not Founc | Not Founc | High | High | High | High | High | High |
| | 2169 | 1855 | 3108 | 2238 | High | High | High | High | Not Founc | Not Founc | Not Founc | Not Founc | High | High |
| | 1957 | 1450 | 1594 | 3010 | High | High | High | High | Not Founc | Not Founc | Not Founc | Not Founc | High | High |
| | 932 | 763 | 1334 | 1265 | High | High | High | High | Not Founc | Not Founc | Not Founc | Not Founc | High | High |
| 872 | 1693 | 1677 | 2292 | 1827 | High | High | High | High | High | High | High | High | High | High |
| | 8491 | 5293 | 14309 | 9928 | High | High | High | High | Not Founc | Not Founc | Not Founc | Not Founc | High | High |
| 2991 | 1790 | 1622 | 3132 | 2008 | High | High | High | High | High | High | High | High | High | High |
| 4825 | 1427 | 1255 | 1450 | 1671 | Not Founc | Not Founc | Not Founc | Not Founc | High | High | High | High | High | High |
| 5157 | 1994 | 2720 | 1678 | 3951 | High | High | High | High | High | High | High | High | High | High |
| | 2568 | 2049 | 2540 | 2751 | High | High | High | High | Not Founc | Not Founc | Not Founc | Not Founc | High | High |
| 1515 | 1609 | 1228 | 1685 | 1549 | Not Founc | Not Founc | Not Founc | Not Founc | High | High | High | High | High | High |
| 7921 | 7227 | 6522 | 6963 | 9924 | High | High | High | High | High | High | High | High | High | High |
| 5299 | 4541 | 2241 | 9743 | 5615 | Not Founc | Not Founc | Not Founc | Not Founc | High | High | High | High | High | High |
| 2839 | 2173 | 1771 | 2650 | 2344 | High | High | High | High | High | High | High | High | High | High |
| 3387 | 1170 | 1097 | 1672 | 1548 | High | High | High | High | High | High | High | High | High | High |
| 7852 | | | | | High | High | High | High | High | High | High | High | Not Founc | Not Founc |
| 4457 | 1640 | 1389 | 2655 | 2104 | High | High | High | High | High | High | High | High | High | High |
| 1409 | 1191 | 1013 | 1033 | 1164 | Not Founc | Not Founc | Not Founc | Not Founc | High | High | High | High | High | High |
| 1318 | 991 | 679 | 1742 | 1270 | High | High | High | High | High | High | High | High | High | High |
| 880 | | | | | High | High | High | High | High | High | High | High | Not Founc | Not Founc |
| | 1780 | 1261 | 2341 | 2065 | High | High | High | High | Not Founc | Not Founc | Not Founc | Not Founc | High | High |

| | | | | | | | | | | | | | | |
|---|---|---|---|---|---|---|---|---|---|---|---|---|---|---|
| 3977 | | | | | Not Founc | Not Founc | Not Founc | Not Founc | High | High | High | High | Not Founc | Not Founc |
| 2408 | 1858 | 1062 | 3162 | 1618 | High | High | High | High | High | High | High | High | High | High |
| 797 | 1666 | 550 | 1671 | 914 | High | High | High | High | High | High | High | High | High | High |
| 994 | 1600 | 1520 | 1878 | 2119 | Not Founc | Not Founc | Not Founc | Not Founc | High | High | High | High | High | High |
| | 14037 | 13172 | 15468 | 18364 | High | High | High | High | Not Founc | Not Founc | Not Founc | Not Founc | High | High |
| 4114 | | | | | High | High | High | High | High | High | High | High | Not Founc | Not Founc |
| 7096 | 4623 | 2569 | 8581 | 7014 | Not Founc | Not Founc | Not Founc | Not Founc | High | High | High | High | High | High |
| 1461 | 1186 | 802 | 1441 | 1358 | High | High | High | High | High | High | High | High | High | High |
| 1278 | 1332 | 1194 | 1062 | 1339 | High | High | High | High | High | High | High | High | High | High |
| | | | | | High | High | High | High | Not Founc | Not Founc | Not Founc | Not Founc | Not Founc | Not Founc |
| | 3582 | 3016 | 5267 | 5147 | Not Founc | Not Founc | Not Founc | Not Founc | Not Founc | Not Founc | Not Founc | Not Founc | High | High |
| | | | | | High | High | High | High | Not Founc | Not Founc | Not Founc | Not Founc | Not Founc | Not Founc |
| | | | | | High | High | High | High | Not Founc | Not Founc | Not Founc | Not Founc | Not Founc | Not Founc |
| | 552 | 394 | 789 | 588 | High | High | High | High | Not Founc | Not Founc | Not Founc | Not Founc | High | High |
| 7590 | 3545 | 2462 | 6466 | 4216 | High | High | High | High | High | High | High | High | High | High |
| | 3868 | 2501 | 7701 | 4470 | High | High | High | High | Not Founc | Not Founc | Not Founc | Not Founc | High | High |
| | 5772 | 5457 | 5504 | 6019 | Not Founc | Not Founc | Not Founc | Not Founc | Not Founc | Not Founc | Not Founc | Not Founc | High | High |
| | 1862 | 1745 | 1891 | 2433 | High | High | High | High | Not Founc | Not Founc | Not Founc | Not Founc | High | High |
| | 5259 | 3097 | 6048 | 4905 | High | High | High | High | Not Founc | Not Founc | Not Founc | Not Founc | High | High |
| 2537 | | | | | High | High | High | High | High | High | High | High | Not Founc | Not Founc |
| 1073 | | | | | Not Founc | Not Founc | Not Founc | Not Founc | High | High | High | High | Not Founc | Not Founc |
| 1868 | 1047 | 789 | 1856 | 1601 | High | High | High | High | High | High | High | High | High | High |
| 10241 | 2294 | 1998 | 2146 | 2736 | Not Founc | Not Founc | Not Founc | Not Founc | High | High | High | High | High | High |
| | 2618 | 3179 | 4134 | 6372 | High | High | High | High | Not Founc | Not Founc | Not Founc | Not Founc | High | High |
| 3759 | 412 | 252 | 417 | 453 | High | High | High | High | High | High | High | High | High | High |
| 737 | | | | | High | High | High | High | High | High | High | High | Not Founc | Not Founc |
| | 9480 | 5872 | 12197 | 9797 | Not Founc | Not Founc | Not Founc | Not Founc | Not Founc | Not Founc | Not Founc | Not Founc | High | High |
| 4641 | 2663 | 1485 | 7664 | 4442 | High | High | High | High | High | High | High | High | High | High |
| | 942 | 797 | 1896 | 1371 | High | High | High | High | Not Founc | Not Founc | Not Founc | Not Founc | High | High |
| 1948 | | | | | High | High | High | High | High | High | High | High | Not Founc | Not Founc |
| 1916 | 3041 | 2540 | 3154 | 3819 | High | High | High | High | High | High | High | High | High | High |

| | | | | | | | | | | | | | | |
|---|---|---|---|---|---|---|---|---|---|---|---|---|---|---|
| 3506 | 3289 | 1542 | 11188 | 5400 | High | High | High | High | High | High | High | High | High | High |
| | | | | | High | High | High | High | Not Found | Not Found | Not Found | Not Found | Not Found | Not Found |
| 4229 | 2915 | 1636 | 5222 | 3255 | High | High | High | High | High | High | High | High | High | High |
| 2779 | 870 | 784 | 1230 | 1252 | Not Found | Not Found | Not Found | Not Found | High | High | High | High | High | High |
| 502 | 504 | 334 | 627 | 414 | High | High | High | High | High | High | High | High | High | High |
| 4187 | 455 | 330 | 575 | 642 | High | High | High | High | High | High | High | High | High | High |
| 712 | 439 | 717 | 397 | 556 | Not Found | Not Found | Not Found | Not Found | High | High | High | High | High | High |
| 2413 | | | | | Not Found | Not Found | Not Found | Not Found | High | High | High | High | Not Found | Not Found |
| | 996 | 541 | 900 | 815 | High | High | High | High | Not Found | Not Found | Not Found | Not Found | High | High |
| 3066 | | | | | High | High | High | High | High | High | High | High | Not Found | Not Found |
| | | | | | Not Found | Not Found | Not Found | Not Found | Not Found | Not Found | Not Found | Not Found | Not Found | Not Found |
| | 3105 | 1678 | 4257 | 2678 | High | High | High | High | Not Found | Not Found | Not Found | Not Found | High | High |
| 2024 | 3856 | 3469 | 5420 | 6961 | Not Found | Not Found | Not Found | Not Found | High | High | High | High | High | High |
| 3124 | | | | | High | High | High | High | High | High | High | High | Not Found | Not Found |
| 5083 | | | | | High | High | High | High | High | High | High | High | Not Found | Not Found |
| | 3859 | 3371 | 8245 | 6283 | High | High | High | High | Not Found | Not Found | Not Found | Not Found | High | High |
| 8934 | | | | | High | High | High | High | High | High | High | High | Not Found | Not Found |
| | 2703 | 2191 | 2688 | 2808 | High | High | High | High | Not Found | Not Found | Not Found | Not Found | High | High |
| 4880 | 3592 | 2810 | 5241 | 5516 | Not Found | Not Found | Not Found | Not Found | High | High | High | High | High | High |
| 5169 | 1674 | 1430 | 1848 | 2062 | High | High | High | High | High | High | High | High | High | High |
| 3427 | | | | | High | High | High | High | High | High | High | High | Not Found | Not Found |
| 1970 | 1475 | 4783 | 1811 | 1888 | High | High | High | High | High | High | High | High | High | High |
| 3585 | 3932 | 2486 | 6649 | 3532 | High | High | High | High | High | High | High | High | High | High |
| 1351 | 1802 | 1558 | 1855 | 1842 | High | High | High | High | High | High | High | High | High | High |
| | 2093 | 1244 | 1944 | 1909 | High | High | High | High | Not Found | Not Found | Not Found | Not Found | High | High |
| | 3218 | 2120 | 5324 | 3580 | High | High | High | High | Not Found | Not Found | Not Found | Not Found | High | High |
| 2598 | | | | | High | High | High | High | High | High | High | High | Not Found | Not Found |
| 4306 | 3778 | 2726 | 3590 | 3730 | High | High | High | High | High | High | High | High | High | High |
| 3865 | | | | | High | High | High | High | High | High | High | High | Not Found | Not Found |
| 1440 | 1636 | 1199 | 1916 | 1349 | High | High | High | High | High | High | High | High | High | High |
| 4505 | | | | | High | High | High | High | High | High | High | High | Not Found | Not Found |

| | | | | | | | | | | | | | | |
|---|---|---|---|---|---|---|---|---|---|---|---|---|---|---|
| | 1976 | 1719 | 2077 | 2421 | High | High | High | High | Not Found | Not Found | Not Found | Not Found | High | High |
| 1177 | | | | | High | High | High | High | High | High | High | High | Not Found | Not Found |
| 2724 | 1388 | 932 | 3431 | 2084 | High | High | High | High | High | High | High | High | High | High |
| | 9849 | 6002 | 10281 | 8674 | Not Found | Not Found | Not Found | Not Found | Not Found | Not Found | Not Found | Not Found | High | High |
| | 3702 | 3887 | 3710 | 6174 | High | High | High | High | Not Found | Not Found | Not Found | Not Found | High | High |
| | 1830 | 1316 | 2550 | 1903 | High | High | High | High | Not Found | Not Found | Not Found | Not Found | High | High |
| 1623 | 594 | 526 | 694 | 754 | Not Found | Not Found | Not Found | Not Found | High | High | High | High | High | High |
| 2287 | 1759 | 1246 | 2161 | 2196 | High | High | High | High | High | High | High | High | High | High |
| 2479 | | | | | High | High | High | High | High | High | High | High | Not Found | Not Found |
| 3182 | 3570 | 2056 | 5191 | 4093 | Not Found | Not Found | Not Found | Not Found | High | High | High | High | High | High |
| 6460 | | | | | High | High | High | High | High | High | High | High | Not Found | Not Found |
| 5431 | 4621 | 2434 | 9847 | 5499 | Not Found | Not Found | Not Found | Not Found | High | High | High | High | High | High |
| 764 | 1604 | 1311 | 5506 | 2316 | Not Found | Not Found | Not Found | Not Found | High | High | High | High | High | High |
| 1945 | | | | | High | High | High | High | High | High | High | High | Not Found | Not Found |
| 1517 | 3304 | 1672 | 1809 | 1650 | High | High | High | High | High | High | High | High | High | High |
| 2956 | | | | | Not Found | Not Found | Not Found | Not Found | High | High | High | High | Not Found | Not Found |
| | | | | | High | High | High | High | Not Found | Not Found | Not Found | Not Found | Not Found | Not Found |
| 13638 | 4108 | 2806 | 6432 | 6159 | High | High | High | High | High | High | High | High | High | High |
| 1031 | 7126 | 4698 | 6620 | 5187 | Not Found | Not Found | Not Found | Not Found | High | High | High | High | High | High |
| 4148 | 1732 | 1115 | 2646 | 2090 | Not Found | Not Found | Not Found | Not Found | High | High | High | High | High | High |
| | 1844 | 1076 | 3159 | 2105 | High | High | High | High | Not Found | Not Found | Not Found | Not Found | High | High |
| 2344 | | | | | High | High | High | High | High | High | High | High | Not Found | Not Found |
| 4998 | 6667 | 3775 | 6605 | 5877 | Not Found | Not Found | Not Found | Not Found | High | High | High | High | High | High |
| 2396 | 2419 | 939 | 2185 | 1323 | High | High | High | High | High | High | High | High | High | High |
| 4293 | 2300 | 1279 | 4734 | 2839 | Not Found | Not Found | Not Found | Not Found | High | High | High | High | High | High |
| 2737 | 2628 | 2077 | 2219 | 2812 | High | High | High | High | High | High | High | High | High | High |
| 3907 | | | | | High | High | High | High | High | High | High | High | Not Found | Not Found |
| | | | | | High | High | High | High | Not Found | Not Found | Not Found | Not Found | Not Found | Not Found |
| 4190 | | | | | High | High | High | High | High | High | High | High | Not Found | Not Found |
| 5218 | | | | | Not Found | Not Found | Not Found | Not Found | High | High | High | High | Not Found | Not Found |
| | 3979 | 2609 | 5652 | 4406 | High | High | High | High | Not Found | Not Found | Not Found | Not Found | High | High |

| | | | | | | | | | | | | | | |
|---|---|---|---|---|---|---|---|---|---|---|---|---|---|---|
| 3135 | 3200 | 2277 | 2694 | 2269 | High | High | High | High | High | High | High | High | High | High |
| 578 | 648 | 597 | 546 | 575 | Not Found | Not Found | Not Found | Not Found | High | High | High | High | High | High |
| 2889 | | | | | Not Found | Not Found | Not Found | Not Found | High | High | High | High | Not Found | Not Found |
| | 4373 | 3895 | 3465 | 4998 | High | High | High | High | Not Found | Not Found | Not Found | Not Found | High | High |
| 1782 | 1784 | 1432 | 1795 | 2060 | High | High | High | High | High | High | High | High | High | High |
| 9301 | | | | | High | High | High | High | High | High | High | High | Not Found | Not Found |
| 496 | | | | | High | High | High | High | High | High | High | High | Not Found | Not Found |
| 2952 | 1539 | 1139 | 1991 | 1761 | High | High | High | High | High | High | High | High | High | High |
| 3914 | | | | | High | High | High | High | High | High | High | High | Not Found | Not Found |
| 2677 | 1737 | 1585 | 1952 | 2188 | Not Found | Not Found | Not Found | Not Found | High | High | High | High | High | High |
| 3123 | 4036 | 4874 | 5075 | 7830 | Not Found | Not Found | Not Found | Not Found | High | High | High | High | High | High |
| | 6302 | 4318 | 7417 | 6784 | High | High | High | High | Not Found | Not Found | Not Found | Not Found | High | High |
| 2664 | 593 | 354 | 843 | 782 | Not Found | Not Found | Not Found | Not Found | High | High | High | High | High | High |
| 1819 | 2482 | 1276 | 3573 | 2741 | Not Found | Not Found | Not Found | Not Found | High | High | High | High | High | High |
| | 1304 | 1065 | 1268 | 1254 | High | High | High | High | Not Found | Not Found | Not Found | Not Found | High | High |
| | 2872 | 2164 | 3209 | 3395 | High | High | High | High | Not Found | Not Found | Not Found | Not Found | High | High |
| 5610 | | | | | High | High | High | High | High | High | High | High | Not Found | Not Found |
| | 4113 | 2754 | 5644 | 4824 | High | High | High | High | Not Found | Not Found | Not Found | Not Found | High | High |
| | 5020 | 4608 | 3897 | 4960 | High | High | High | High | Not Found | Not Found | Not Found | Not Found | High | High |
| | 3903 | 2373 | 3660 | 3459 | High | High | High | High | Not Found | Not Found | Not Found | Not Found | High | High |
| | | | | | High | High | High | High | Not Found | Not Found | Not Found | Not Found | Not Found | Not Found |
| | 12292 | 16939 | 15660 | 6300 | Not Found | Not Found | Not Found | Not Found | Not Found | Not Found | Not Found | Not Found | High | High |
| 5288 | 1615 | 1373 | 1596 | 1876 | High | High | High | High | High | High | High | High | High | High |
| 2465 | | | | | High | High | High | High | High | High | High | High | Not Found | Not Found |
| 1650 | 1142 | 789 | 3140 | 1295 | High | High | High | High | High | High | High | High | High | High |
| 2993 | | | | | High | High | High | High | High | High | High | High | Not Found | Not Found |
| 1921 | | | | | Not Found | Not Found | Not Found | Not Found | High | High | High | High | High | High |
| 8785 | | | | | High | High | High | High | High | High | High | High | Not Found | Not Found |
| 1716 | | | | | Not Found | Not Found | Not Found | Not Found | High | High | High | High | Not Found | Not Found |
| | 3090 | 2674 | 2849 | 2982 | Not Found | Not Found | Not Found | Not Found | Not Found | Not Found | Not Found | Not Found | High | High |
| 2181 | 1968 | 1487 | 1927 | 2336 | High | High | High | High | High | High | High | High | High | High |

| | | | | | | | | | | | | | | |
|---|---|---|---|---|---|---|---|---|---|---|---|---|---|---|
| | 1990 | 1505 | 1918 | 2324 | Not Founc | Not Founc | Not Founc | Not Founc | Not Founc | Not Founc | Not Founc | Not Founc | High | High |
| | 1027 | 913 | 755 | 1193 | High | High | High | High | Not Founc | Not Founc | Not Founc | Not Founc | High | High |
| | 4984 | 4279 | 6159 | 6546 | High | High | High | High | Not Founc | Not Founc | Not Founc | Not Founc | High | High |
| 1384 | | | | | High | High | High | High | High | High | High | High | Not Founc | Not Founc |
| 2791 | | | | | High | High | High | High | High | High | High | High | Not Founc | Not Founc |
| | 2442 | 1972 | 2348 | 2860 | High | High | High | High | Not Founc | Not Founc | Not Founc | Not Founc | High | High |
| | 3918 | 2257 | 4112 | 3344 | High | High | High | High | Not Founc | Not Founc | Not Founc | Not Founc | High | High |
| 1732 | | | | | High | High | High | High | High | High | High | High | Not Founc | Not Founc |
| | 3342 | 2489 | 4549 | 5152 | High | High | High | High | Not Founc | Not Founc | Not Founc | Not Founc | High | High |
| | 2804 | 2576 | 4644 | 3712 | Not Founc | Not Founc | Not Founc | Not Founc | Not Founc | Not Founc | Not Founc | Not Founc | High | High |
| 4738 | | | | | High | High | High | High | High | High | High | High | Not Founc | Not Founc |
| 4675 | 1398 | 1381 | 1614 | 2070 | Not Founc | Not Founc | Not Founc | Not Founc | High | High | High | High | High | High |
| 8156 | 5669 | 4574 | 8704 | 6703 | Not Founc | Not Founc | Not Founc | Not Founc | High | High | High | High | High | High |
| 4530 | 9474 | 4240 | 13294 | 6590 | Not Founc | Not Founc | Not Founc | Not Founc | High | High | High | High | High | High |
| 1393 | 2512 | 1951 | 3415 | 2425 | High | High | High | High | High | High | High | High | High | High |
| 2860 | | | | | High | High | High | High | High | High | High | High | Not Founc | Not Founc |
| | 3045 | 1945 | 5287 | 3578 | High | High | High | High | Not Founc | Not Founc | Not Founc | Not Founc | High | High |
| 2485 | | | | | High | High | High | High | High | High | High | High | Not Founc | Not Founc |
| | 461 | 303 | 583 | 428 | High | High | High | High | Not Founc | Not Founc | Not Founc | Not Founc | High | High |
| 5309 | 3084 | 2701 | 2880 | 3851 | Not Founc | Not Founc | Not Founc | Not Founc | High | High | High | High | High | High |
| 727 | | | | | High | High | High | High | High | High | High | High | Not Founc | Not Founc |
| 2317 | 4350 | 1699 | 3912 | 2675 | High | High | High | High | High | High | High | High | High | High |
| | 707 | 527 | 787 | 761 | High | High | High | High | Not Founc | Not Founc | Not Founc | Not Founc | High | High |
| | 8400 | 6436 | 7175 | 5741 | Not Founc | Not Founc | Not Founc | Not Founc | Not Founc | Not Founc | Not Founc | Not Founc | High | High |
| | 2631 | 1355 | 3378 | 2093 | High | High | High | High | Not Founc | Not Founc | Not Founc | Not Founc | High | High |
| 1606 | 2016 | 1377 | 3167 | 2387 | Not Founc | Not Founc | Not Founc | Not Founc | High | High | High | High | High | High |
| 3809 | 2734 | 1891 | 3828 | 3668 | High | High | High | High | High | High | High | High | High | High |
| 7152 | 1832 | 1360 | 2116 | 2201 | Not Founc | Not Founc | Not Founc | Not Founc | High | High | High | High | High | High |
| 3963 | | | | | High | High | High | High | High | High | High | High | Not Founc | Not Founc |
| | 1478 | 1670 | 1246 | 1592 | High | High | High | High | Not Founc | Not Founc | Not Founc | Not Founc | High | High |
| | 5523 | 4199 | 9053 | 9272 | Not Founc | Not Founc | Not Founc | Not Founc | Not Founc | Not Founc | Not Founc | Not Founc | High | High |

| | | | | | | | | | | | | | | |
|---|---|---|---|---|---|---|---|---|---|---|---|---|---|---|
| | 2498 | 1334 | 2553 | 1900 | High | High | High | High | Not Founc | Not Founc | Not Founc | Not Founc | High | High |
| | | | | | High | High | High | High | Not Founc | Not Founc | Not Founc | Not Founc | Not Founc | Not Founc |
| 2069 | 1189 | 928 | 2792 | 1343 | Not Founc | Not Founc | Not Founc | Not Founc | High | High | High | High | High | High |
| 2137 | | | | | High | High | High | High | High | High | High | High | Not Founc | Not Founc |
| 3267 | 2494 | 2563 | 2345 | 3477 | High | High | High | High | High | High | High | High | High | High |
| 2146 | | | | | High | High | High | High | High | High | High | High | Not Founc | Not Founc |
| | 1680 | 1005 | 2342 | 1539 | High | High | High | High | Not Founc | Not Founc | Not Founc | Not Founc | High | High |
| 1690 | 1332 | 989 | 1258 | 1353 | Not Founc | Not Founc | Not Founc | Not Founc | High | High | High | High | High | High |
| 6127 | | | | | Not Founc | Not Founc | Not Founc | Not Founc | High | High | High | High | Not Founc | Not Founc |
| 1659 | 1775 | 2266 | 1911 | 1724 | High | High | High | High | High | High | High | High | High | High |
| 2070 | 1070 | 685 | 1341 | 1127 | High | High | High | High | High | High | High | High | High | High |
| | | | | | Not Founc | Not Founc | Not Founc | Not Founc | High | High | High | High | High | High |
| 655 | | | | | High | High | High | High | High | High | High | High | Not Founc | Not Founc |
| | 4695 | 4785 | 6430 | 7668 | High | High | High | High | Not Founc | Not Founc | Not Founc | Not Founc | High | High |
| | 5557 | 4152 | 9118 | 7699 | Not Founc | Not Founc | Not Founc | Not Founc | Not Founc | Not Founc | Not Founc | Not Founc | High | High |
| | 1235 | 846 | 1615 | 1091 | Not Founc | Not Founc | Not Founc | Not Founc | Not Founc | Not Founc | Not Founc | Not Founc | High | High |
| | 1025 | 733 | 1236 | 1005 | High | High | High | High | Not Founc | Not Founc | Not Founc | Not Founc | High | High |
| | 4351 | 3050 | 6532 | 5645 | High | High | High | High | Not Founc | Not Founc | Not Founc | Not Founc | High | High |
| 6382 | 2244 | 2358 | 2784 | 1749 | High | High | High | High | High | High | High | High | High | High |
| 4028 | | | | | High | High | High | High | High | High | High | High | Not Founc | Not Founc |
| | 1492 | 958 | 2136 | 1338 | High | High | High | High | Not Founc | Not Founc | Not Founc | Not Founc | High | High |
| 1350 | 1700 | 1044 | 2238 | 1850 | Not Founc | Not Founc | Not Founc | Not Founc | High | High | High | High | High | High |
| 2744 | | | | | High | High | High | High | High | High | High | High | Not Founc | Not Founc |
| 2122 | 1360 | 963 | 3456 | 2415 | Not Founc | Not Founc | Not Founc | Not Founc | High | High | High | High | High | High |
| 1290 | 1556 | 1612 | 910 | 1123 | Not Founc | Not Founc | Not Founc | Not Founc | High | High | High | High | High | High |
| 2354 | | | | | High | High | High | High | High | High | High | High | Not Founc | Not Founc |
| | 3946 | 2829 | 5514 | 4877 | High | High | High | High | Not Founc | Not Founc | Not Founc | Not Founc | High | High |
| | 2684 | 2062 | 2342 | 2825 | High | High | High | High | Not Founc | Not Founc | Not Founc | Not Founc | High | High |
| 4340 | | | | | High | High | High | High | High | High | High | High | Not Founc | Not Founc |
| 2261 | | | | | High | High | High | High | High | High | High | High | Not Founc | Not Founc |
| | 1791 | 1341 | 1877 | 2302 | High | High | High | High | Not Founc | Not Founc | Not Founc | Not Founc | High | High |

| | | | | | | | | | | | | | | |
|---|---|---|---|---|---|---|---|---|---|---|---|---|---|---|
| 7598 | 4986 | 4282 | 6280 | 7245 | Not Found | Not Found | Not Found | Not Found | High | High | High | High | High | High |
| 4955 | | | | | Not Found | Not Found | Not Found | Not Found | High | High | High | High | Not Found | Not Found |
| 1990 | 1234 | 977 | 2426 | 1907 | High | High | High | High | High | High | High | High | High | High |
| 4601 | | | | | High | High | High | High | High | High | High | High | Not Found | Not Found |
| 1065 | | | | | High | High | High | High | High | High | High | High | Not Found | Not Found |
| | | | | | High | High | High | High | Not Found | Not Found | Not Found | Not Found | Not Found | Not Found |
| 1529 | | | | | High | High | High | High | High | High | High | High | Not Found | Not Found |
| 1394 | | | | | High | High | High | High | High | High | High | High | Not Found | Not Found |
| 1850 | 1988 | 1630 | 1929 | 2417 | Not Found | Not Found | Not Found | Not Found | High | High | High | High | High | High |
| | | | | | High | High | High | High | Not Found | Not Found | Not Found | Not Found | Not Found | Not Found |
| 1658 | 2817 | 1712 | 3024 | 2026 | High | High | High | High | High | High | High | High | High | High |
| 4988 | 5615 | 4292 | 6893 | 7869 | Not Found | Not Found | Not Found | Not Found | High | High | High | High | High | High |
| | | | | | High | High | High | High | Not Found | Not Found | Not Found | Not Found | Not Found | Not Found |
| 1124 | 1911 | 1088 | 1912 | 1561 | Not Found | Not Found | Not Found | Not Found | High | High | High | High | High | High |
| 2160 | 1675 | 1338 | 2113 | 1670 | Not Found | Not Found | Not Found | Not Found | High | High | High | High | High | High |
| 2176 | 1848 | 1216 | 3194 | 2595 | High | High | High | High | High | High | High | High | High | High |
| 1744 | | | | | Not Found | Not Found | Not Found | Not Found | High | High | High | High | Not Found | Not Found |
| | 1536 | 1255 | 2636 | 2159 | High | High | High | High | Not Found | Not Found | Not Found | Not Found | High | High |
| | 3032 | 878 | 3598 | 1813 | High | High | High | High | Not Found | Not Found | Not Found | Not Found | High | High |
| | 2170 | 1968 | 3318 | 2574 | High | High | High | High | Not Found | Not Found | Not Found | Not Found | High | High |
| | | | | | Not Found | Not Found | Not Found | Not Found | Not Found | Not Found | Not Found | Not Found | Not Found | Not Found |
| | | | | | High | High | High | High | Not Found | Not Found | Not Found | Not Found | Not Found | Not Found |
| 2641 | 1724 | 1294 | 2104 | 2346 | High | High | High | High | High | High | High | High | High | High |
| 1440 | 2454 | 1411 | 3433 | 2351 | Not Found | Not Found | Not Found | Not Found | High | High | High | High | High | High |
| 3998 | 2955 | 2291 | 3160 | 3074 | Not Found | Not Found | Not Found | Not Found | High | High | High | High | High | High |
| 1419 | | | | | High | High | High | High | High | High | High | High | Not Found | Not Found |
| | | | | | High | High | High | High | Not Found | Not Found | Not Found | Not Found | Not Found | Not Found |
| | | | | | High | High | High | High | Not Found | Not Found | Not Found | Not Found | Not Found | Not Found |
| | 1504 | 1279 | 2399 | 2123 | High | High | High | High | Not Found | Not Found | Not Found | Not Found | High | High |
| 1823 | 1394 | 813 | 3074 | 1730 | High | High | High | High | High | High | High | High | High | High |
| 3127 | | | | | Not Found | Not Found | Not Found | Not Found | High | High | High | High | High | High |

| | | | | | | | | | | | | | | |
|---|---|---|---|---|---|---|---|---|---|---|---|---|---|---|
| | 1722 | 1239 | 2764 | 2433 | High | High | High | High | Not Founc | Not Founc | Not Founc | Not Founc | High | High |
| 271 | | | | | High | High | High | High | High | High | High | High | Not Founc | Not Founc |
| 897 | 195 | 135 | 216 | 236 | Not Founc | Not Founc | Not Founc | Not Founc | High | High | High | High | High | High |
| 1534 | 1818 | 1108 | 1786 | 1645 | Not Founc | Not Founc | Not Founc | Not Founc | High | High | High | High | High | High |
| | 2033 | 1331 | 2070 | 2285 | High | High | High | High | Not Founc | Not Founc | Not Founc | Not Founc | High | High |
| 1954 | | | | | High | High | High | High | High | High | High | High | Not Founc | Not Founc |
| 1780 | | | | | High | High | High | High | High | High | High | High | Not Founc | Not Founc |
| | 2398 | 1434 | 6381 | 3274 | High | High | High | High | Not Founc | Not Founc | Not Founc | Not Founc | High | High |
| 4867 | 3780 | 3053 | 3465 | 3926 | Not Founc | Not Founc | Not Founc | Not Founc | High | High | High | High | High | High |
| 1077 | | | | | High | High | High | High | High | High | High | High | Not Founc | Not Founc |
| 522 | | | | | High | High | High | High | High | High | High | High | Not Founc | Not Founc |
| 3420 | | | | | High | High | High | High | High | High | High | High | Not Founc | Not Founc |
| 3252 | | | | | High | High | High | High | High | High | High | High | Not Founc | Not Founc |
| 2568 | | | | | High | High | High | High | High | High | High | High | Not Founc | Not Founc |
| | | | | | High | High | High | High | Not Founc | Not Founc | Not Founc | Not Founc | Not Founc | Not Founc |
| | 985 | 510 | 1784 | 1216 | Not Founc | Not Founc | Not Founc | Not Founc | Not Founc | Not Founc | Not Founc | Not Founc | High | High |
| | 7863 | 7777 | 7629 | 10053 | Not Founc | Not Founc | Not Founc | Not Founc | Not Founc | Not Founc | Not Founc | Not Founc | High | High |
| | | | | | Not Founc | Not Founc | Not Founc | Not Founc | Not Founc | Not Founc | Not Founc | Not Founc | High | High |
| | 5510 | 5400 | 7573 | 8523 | Not Founc | Not Founc | Not Founc | Not Founc | Not Founc | Not Founc | Not Founc | Not Founc | High | High |
| 2047 | 2628 | 1938 | 5550 | 3835 | High | High | High | High | High | High | High | High | High | High |
| 845 | 2605 | 2268 | 2544 | 3280 | Not Founc | Not Founc | Not Founc | Not Founc | High | High | High | High | High | High |
| 2512 | 2082 | 1606 | 2222 | 2126 | Not Founc | Not Founc | Not Founc | Not Founc | High | High | High | High | High | High |
| 4015 | 2290 | 1792 | 2695 | 2534 | High | High | High | High | High | High | High | High | High | High |
| 2053 | 1137 | 790 | 1760 | 1388 | Not Founc | Not Founc | Not Founc | Not Founc | High | High | High | High | High | High |
| 1412 | | | | | Not Founc | Not Founc | Not Founc | Not Founc | High | High | High | High | Not Founc | Not Founc |
| 7475 | 3830 | 2490 | 9687 | 5131 | High | High | High | High | High | High | High | High | High | High |
| 1528 | | | | | High | High | High | High | High | High | High | High | Not Founc | Not Founc |
| | 2364 | 1690 | 2928 | 2388 | High | High | High | High | Not Founc | Not Founc | Not Founc | Not Founc | High | High |
| | 4046 | 2742 | 4679 | 3655 | High | High | High | High | Not Founc | Not Founc | Not Founc | Not Founc | High | High |
| 2370 | | | | | High | High | High | High | High | High | High | High | Not Founc | Not Founc |
| 4740 | 3123 | 2466 | 3074 | 3277 | High | High | High | High | High | High | High | High | High | High |

| | | | | | | | | | | | | | | |
|---|---|---|---|---|---|---|---|---|---|---|---|---|---|---|
| | 1171 | 1142 | 1348 | 1615 | High | High | High | High | Not Found | Not Found | Not Found | Not Found | High | High |
| 1620 | 981 | 720 | 1196 | 1076 | High | High | High | High | High | High | High | High | High | High |
| 769 | | | | | High | High | High | High | High | High | High | High | Not Found | Not Found |
| 2499 | 1571 | 1838 | 2350 | 2582 | High | High | High | High | High | High | High | High | High | High |
| | 1407 | 771 | 4381 | 2343 | High | High | High | High | Not Found | Not Found | Not Found | Not Found | High | High |
| | 6002 | 2892 | 8450 | 5959 | High | High | High | High | Not Found | Not Found | Not Found | Not Found | High | High |
| | 1544 | 964 | 1589 | 1331 | High | High | High | High | Not Found | Not Found | Not Found | Not Found | High | High |
| 1237 | | | | | High | High | High | High | High | High | High | High | Not Found | Not Found |
| | 3467 | 1699 | 3348 | 2282 | High | High | High | High | Not Found | Not Found | Not Found | Not Found | High | High |
| 1423 | | | | | High | High | High | High | High | High | High | High | Not Found | Not Found |
| | 1651 | 1415 | 2795 | 2575 | High | High | High | High | Not Found | Not Found | Not Found | Not Found | High | High |
| 563 | 2089 | 1674 | 2597 | 2525 | Not Found | Not Found | Not Found | Not Found | High | High | High | High | High | High |
| 2574 | | | | | High | High | High | High | High | High | High | High | Not Found | Not Found |
| 3003 | | | | | Not Found | Not Found | Not Found | Not Found | High | High | High | High | Not Found | Not Found |
| 1553 | 1877 | 1246 | 2779 | 2233 | High | High | High | High | High | High | High | High | High | High |
| | | | | | High | High | High | High | Not Found | Not Found | Not Found | Not Found | High | High |
| 3135 | 2530 | 2084 | 3407 | 3067 | High | High | High | High | High | High | High | High | High | High |
| 2430 | | | | | High | High | High | High | High | High | High | High | Not Found | Not Found |
| | 861 | 541 | 1495 | 972 | High | High | High | High | Not Found | Not Found | Not Found | Not Found | High | High |
| 5092 | 2043 | 1265 | 1698 | 1899 | Not Found | Not Found | Not Found | Not Found | High | High | High | High | High | High |
| 1576 | 600 | 513 | 628 | 1281 | Not Found | Not Found | Not Found | Not Found | High | High | High | High | High | High |
| | 959 | 944 | 968 | 1361 | High | High | High | High | Not Found | Not Found | Not Found | Not Found | High | High |
| | 1536 | 1073 | 2613 | 2107 | High | High | High | High | Not Found | Not Found | Not Found | Not Found | High | High |
| 978 | | | | | Not Found | Not Found | Not Found | Not Found | High | High | High | High | Not Found | Not Found |
| 1514 | 1580 | 1312 | 1686 | 1583 | Not Found | Not Found | Not Found | Not Found | High | High | High | High | High | High |
| 2007 | | | | | High | High | High | High | High | High | High | High | Not Found | Not Found |
| 2030 | 1908 | 1425 | 2104 | 1721 | Not Found | Not Found | Not Found | Not Found | High | High | High | High | High | High |
| | 1785 | 1306 | 2241 | 2468 | High | High | High | High | Not Found | Not Found | Not Found | Not Found | High | High |
| 3616 | 5325 | 3009 | 6290 | 4373 | High | High | High | High | High | High | High | High | High | High |
| 1664 | | | | | High | High | High | High | High | High | High | High | Not Found | Not Found |
| 3559 | | | | | Not Found | Not Found | Not Found | Not Found | High | High | High | High | Not Found | Not Found |

| | | | | | | | | | | | | | | |
|---|---|---|---|---|---|---|---|---|---|---|---|---|---|---|
| 3554 | | | | | High | High | High | High | High | High | High | High | Not Founc | Not Founc |
| 4948 | | | | | Not Founc | Not Founc | Not Founc | Not Founc | High | High | High | High | Not Founc | Not Founc |
| 2749 | | | | | Not Founc | Not Founc | Not Founc | Not Founc | High | High | High | High | Not Founc | Not Founc |
| 2371 | | | | | High | High | High | High | High | High | High | High | Not Founc | Not Founc |
| | 2587 | 1985 | 2970 | 3254 | High | High | High | High | Not Founc | Not Founc | Not Founc | Not Founc | High | High |
| | | | | | High | High | High | High | Not Founc | Not Founc | Not Founc | Not Founc | Not Founc | Not Founc |
| 5617 | | | | | Not Founc | Not Founc | Not Founc | Not Founc | High | High | High | High | Not Founc | Not Founc |
| 688 | 668 | 443 | 635 | 678 | Not Founc | Not Founc | Not Founc | Not Founc | High | High | High | High | High | High |
| | 551 | 305 | 468 | 503 | Not Founc | Not Founc | Not Founc | Not Founc | Not Founc | Not Founc | Not Founc | Not Founc | High | High |
| 1923 | 638 | 652 | 788 | 958 | Not Founc | Not Founc | Not Founc | Not Founc | High | High | High | High | High | High |
| 6398 | | | | | High | High | High | High | High | High | High | High | Not Founc | Not Founc |
| 4320 | 3911 | 2377 | 6081 | 3977 | Not Founc | Not Founc | Not Founc | Not Founc | High | High | High | High | High | High |
| 857 | 1818 | 944 | 2628 | 1636 | Not Founc | Not Founc | Not Founc | Not Founc | High | High | High | High | High | High |
| 2613 | 2507 | 1704 | 3892 | 2629 | Not Founc | Not Founc | Not Founc | Not Founc | High | High | High | High | High | High |
| 3690 | 6638 | 4526 | 6764 | 6701 | Not Founc | Not Founc | Not Founc | Not Founc | High | High | High | High | High | High |
| | 1140 | 736 | 1289 | 1228 | High | High | High | High | Not Founc | Not Founc | Not Founc | Not Founc | High | High |
| | 2379 | 2745 | 2071 | 3730 | High | High | High | High | Not Founc | Not Founc | Not Founc | Not Founc | High | High |
| | | | | | High | High | High | High | Not Founc | Not Founc | Not Founc | Not Founc | Not Founc | Not Founc |
| 6053 | | | | | Not Founc | Not Founc | Not Founc | Not Founc | High | High | High | High | Not Founc | Not Founc |
| 4244 | | | | | Not Founc | Not Founc | Not Founc | Not Founc | High | High | High | High | Not Founc | Not Founc |
| | | | | | High | High | High | High | Not Founc | Not Founc | Not Founc | Not Founc | Not Founc | Not Founc |
| 2707 | 2708 | 1410 | 3939 | 2602 | Not Founc | Not Founc | Not Founc | Not Founc | High | High | High | High | High | High |
| | 220 | 240 | 422 | 290 | Not Founc | Not Founc | Not Founc | Not Founc | Not Founc | Not Founc | Not Founc | Not Founc | High | High |
| | 1702 | 1079 | 1848 | 1580 | Not Founc | Not Founc | Not Founc | Not Founc | Not Founc | Not Founc | Not Founc | Not Founc | High | High |
| | 5082 | 3837 | 5181 | 5341 | Not Founc | Not Founc | Not Founc | Not Founc | Not Founc | Not Founc | Not Founc | Not Founc | High | High |
| 6242 | 2259 | 1708 | 2580 | 2297 | Not Founc | Not Founc | Not Founc | Not Founc | High | High | High | High | High | High |
| | 2451 | 2013 | 1962 | 1962 | High | High | High | High | Not Founc | Not Founc | Not Founc | Not Founc | High | High |
| 1944 | | | | | High | High | High | High | High | High | High | High | Not Founc | Not Founc |
| 1863 | 843 | 486 | 1394 | 821 | Not Founc | Not Founc | Not Founc | Not Founc | High | High | High | High | High | High |
| | 1953 | 987 | 2397 | 1771 | High | High | High | High | Not Founc | Not Founc | Not Founc | Not Founc | High | High |
| 1922 | | | | | Not Founc | Not Founc | Not Founc | Not Founc | High | High | High | High | Not Founc | Not Founc |

| | | | | | | | | | | | | | | |
|---|---|---|---|---|---|---|---|---|---|---|---|---|---|---|
| | | | | | High | High | High | High | High | High | High | High | Not Founc | Not Founc |
| | | | | | High | High | High | High | Not Founc | Not Founc | Not Founc | Not Founc | Not Founc | Not Founc |
| 2547 | | | | | Not Founc | Not Founc | Not Founc | Not Founc | High | High | High | High | Not Founc | Not Founc |
| 7106 | | | | | High | High | High | High | High | High | High | High | Not Founc | Not Founc |
| 995 | | | | | High | High | High | High | High | High | High | High | Not Founc | Not Founc |
| 1745 | | | | | High | High | High | High | High | High | High | High | Not Founc | Not Founc |
| | 1655 | 1455 | 1950 | 1733 | Not Founc | Not Founc | Not Founc | Not Founc | Not Founc | Not Founc | Not Founc | Not Founc | High | High |
| | | | | | High | High | High | High | Not Founc | Not Founc | Not Founc | Not Founc | High | High |
| | 2199 | 1944 | 2105 | 1967 | Not Founc | Not Founc | Not Founc | Not Founc | Not Founc | Not Founc | Not Founc | Not Founc | High | High |
| | 2980 | 2534 | 5871 | 6571 | High | High | High | High | Not Founc | Not Founc | Not Founc | Not Founc | High | High |
| | | | | | High | High | High | High | Not Founc | Not Founc | Not Founc | Not Founc | Not Founc | Not Founc |
| 1530 | | | | | Not Founc | Not Founc | Not Founc | Not Founc | High | High | High | High | High | High |
| | 1379 | 944 | 1528 | 1353 | Not Founc | Not Founc | Not Founc | Not Founc | Not Founc | Not Founc | Not Founc | Not Founc | High | High |
| 144 | | | | | High | High | High | High | High | High | High | High | Not Founc | Not Founc |
| 2499 | 1628 | 1577 | 2680 | 2151 | Not Founc | Not Founc | Not Founc | Not Founc | High | High | High | High | High | High |
| | | | | | High | High | High | High | Not Founc | Not Founc | Not Founc | Not Founc | Not Founc | Not Founc |
| 2297 | | | | | High | High | High | High | High | High | High | High | Not Founc | Not Founc |
| | 1383 | 960 | 2130 | 1737 | High | High | High | High | Not Founc | Not Founc | Not Founc | Not Founc | High | High |
| | | | | | High | High | High | High | Not Founc | Not Founc | Not Founc | Not Founc | Not Founc | Not Founc |
| 1656 | 1728 | 1123 | 2478 | 1844 | Not Founc | Not Founc | Not Founc | Not Founc | High | High | High | High | High | High |
| 3228 | | | | | High | High | High | High | High | High | High | High | Not Founc | Not Founc |
| 3555 | | | | | High | High | High | High | High | High | High | High | Not Founc | Not Founc |
| 2177 | 4092 | 3334 | 5172 | 4478 | Not Founc | Not Founc | Not Founc | Not Founc | High | High | High | High | High | High |
| 1245 | 1427 | 1083 | 1727 | 1870 | Not Founc | Not Founc | Not Founc | Not Founc | High | High | High | High | High | High |
| 3379 | 2880 | 2026 | 4872 | 3590 | Not Founc | Not Founc | Not Founc | Not Founc | High | High | High | High | High | High |
| | 883 | 484 | 1072 | 754 | High | High | High | High | Not Founc | Not Founc | Not Founc | Not Founc | High | High |
| | | | | | High | High | High | High | Not Founc | Not Founc | Not Founc | Not Founc | Not Founc | Not Founc |
| 2861 | 4016 | 3322 | 3706 | 4587 | Not Founc | Not Founc | Not Founc | Not Founc | High | High | High | High | High | High |
| 2191 | 4894 | 3064 | 7385 | 6368 | Not Founc | Not Founc | Not Founc | Not Founc | High | High | High | High | High | High |
| 936 | | | | | High | High | High | High | High | High | High | High | Not Founc | Not Founc |
| 2123 | | | | | High | High | High | High | High | High | High | High | Not Founc | Not Founc |

| | | | | | | | | | | | | | | |
|---|---|---|---|---|---|---|---|---|---|---|---|---|---|---|
| 2737 | | | | | High | High | High | High | High | High | High | High | Not Found | Not Found |
| 2990 | | | | | High | High | High | High | High | High | High | High | Not Found | Not Found |
| 1345 | 1399 | 1117 | 1669 | 1307 | Not Found | Not Found | Not Found | Not Found | High | High | High | High | High | High |
| | | | | | High | High | High | High | Not Found | Not Found | Not Found | Not Found | Not Found | Not Found |
| 4525 | | | | | High | High | High | High | High | High | High | High | Not Found | Not Found |
| 2783 | 4748 | 3217 | 3820 | 3639 | Not Found | Not Found | Not Found | Not Found | High | High | High | High | High | High |
| | | | | | High | High | High | High | Not Found | Not Found | Not Found | Not Found | Not Found | Not Found |
| 1504 | 1746 | 1540 | 1519 | 1612 | Not Found | Not Found | Not Found | Not Found | High | High | High | High | High | High |
| 2065 | 1709 | 954 | 4264 | 1916 | Not Found | Not Found | Not Found | Not Found | High | High | High | High | High | High |
| 2995 | | | | | High | High | High | High | High | High | High | High | Not Found | Not Found |
| | 2321 | 1696 | 3922 | 2743 | High | High | High | High | Not Found | Not Found | Not Found | Not Found | High | High |
| 2306 | 1647 | 780 | 2970 | 1578 | Not Found | Not Found | Not Found | Not Found | High | High | High | High | High | High |
| 2533 | 2020 | 1187 | 3771 | 2752 | Not Found | Not Found | Not Found | Not Found | High | High | High | High | High | High |
| | 1558 | 1420 | 1265 | 1368 | Not Found | Not Found | Not Found | Not Found | Not Found | Not Found | Not Found | Not Found | High | High |
| | 3544 | 2681 | 4617 | 4282 | Not Found | Not Found | Not Found | Not Found | Not Found | Not Found | Not Found | Not Found | High | High |
| 2514 | | | | | Not Found | Not Found | Not Found | Not Found | High | High | High | High | Not Found | Not Found |
| | | | | | High | High | High | High | Not Found | Not Found | Not Found | Not Found | Not Found | Not Found |
| | | | | | Not Found | Not Found | Not Found | Not Found | Not Found | Not Found | Not Found | Not Found | Not Found | Not Found |
| | 701 | 526 | 857 | 679 | Not Found | Not Found | Not Found | Not Found | Not Found | Not Found | Not Found | Not Found | High | High |
| | | | | | High | High | High | High | Not Found | Not Found | Not Found | Not Found | Not Found | Not Found |
| | 2389 | 2236 | 2121 | 2280 | Not Found | Not Found | Not Found | Not Found | Not Found | Not Found | Not Found | Not Found | High | High |
| 1291 | | | | | High | High | High | High | High | High | High | High | Not Found | Not Found |
| 2210 | | | | | Not Found | Not Found | Not Found | Not Found | High | High | High | High | Not Found | Not Found |
| 1959 | 2256 | 892 | 2940 | 1902 | Not Found | Not Found | Not Found | Not Found | High | High | High | High | High | High |
| | 6852 | 5252 | 8009 | 7438 | Not Found | Not Found | Not Found | Not Found | Not Found | Not Found | Not Found | Not Found | High | High |
| 2310 | | | | | High | High | High | High | High | High | High | High | Not Found | Not Found |
| | 1933 | 1572 | 3113 | 2480 | High | High | High | High | Not Found | Not Found | Not Found | Not Found | High | High |
| | 8588 | 6341 | 9051 | 8888 | Not Found | Not Found | Not Found | Not Found | Not Found | Not Found | Not Found | Not Found | High | High |
| 976 | | | | | Not Found | Not Found | Not Found | Not Found | High | High | High | High | Not Found | Not Found |
| | 2092 | 1611 | 2299 | 2110 | High | High | High | High | Not Found | Not Found | Not Found | Not Found | High | High |
| 786 | | | | | Not Found | Not Found | Not Found | Not Found | High | High | High | High | Not Found | Not Found |

| | | | | | | | | | | | | | | |
|---|---|---|---|---|---|---|---|---|---|---|---|---|---|---|
| 4086 | | | | | Not Found | Not Found | Not Found | Not Found | High | High | High | High | Not Found | Not Found |
| 3369 | | | | | High | High | High | High | High | High | High | High | Not Found | Not Found |
| 2507 | 1907 | 1198 | 3577 | 1919 | Not Found | Not Found | Not Found | Not Found | High | High | High | High | High | High |
| 8290 | 4171 | 3685 | 7608 | 5348 | Not Found | Not Found | Not Found | Not Found | High | High | High | High | High | High |
| 2412 | | | | | High | High | High | High | High | High | High | High | Not Found | Not Found |
| | 5771 | 3051 | 9472 | 5897 | Not Found | Not Found | Not Found | Not Found | Not Found | Not Found | Not Found | Not Found | High | High |
| 2705 | | | | | Not Found | Not Found | Not Found | Not Found | High | High | High | High | Not Found | Not Found |
| | 442 | 355 | 495 | 452 | High | High | High | High | Not Found | Not Found | Not Found | Not Found | High | High |
| 1346 | 3214 | 2138 | 3314 | 2885 | Not Found | Not Found | Not Found | Not Found | High | High | High | High | High | High |
| 1671 | 2755 | 2230 | 3009 | 3482 | Not Found | Not Found | Not Found | Not Found | High | High | High | High | High | High |
| 2124 | 3161 | 1990 | 3570 | 3173 | Not Found | Not Found | Not Found | Not Found | High | High | High | High | High | High |
| 6337 | | | | | Not Found | Not Found | Not Found | Not Found | High | High | High | High | Not Found | Not Found |
| | 1264 | 716 | 2843 | 2265 | Not Found | Not Found | Not Found | Not Found | High | High | High | High | High | High |
| 653 | | | | | High | High | High | High | High | High | High | High | Not Found | Not Found |
| 3256 | | | | | High | High | High | High | High | High | High | High | Not Found | Not Found |
| 3138 | 2121 | 1952 | 1875 | 3007 | Not Found | Not Found | Not Found | Not Found | High | High | High | High | High | High |
| 3063 | | | | | High | High | High | High | High | High | High | High | Not Found | Not Found |
| 2508 | | | | | Not Found | Not Found | Not Found | Not Found | High | High | High | High | Not Found | Not Found |
| 1612 | | | | | High | High | High | High | High | High | High | High | Not Found | Not Found |
| 2633 | | | | | High | High | High | High | High | High | High | High | Not Found | Not Found |
| 1009 | | | | | High | High | High | High | High | High | High | High | Not Found | Not Found |
| | 3943 | 3845 | 5267 | 5731 | High | High | High | High | Not Found | Not Found | Not Found | Not Found | High | High |
| 1178 | | | | | Not Found | Not Found | Not Found | Not Found | High | High | High | High | Not Found | Not Found |
| | 2669 | 2487 | 2984 | 3118 | Not Found | Not Found | Not Found | Not Found | Not Found | Not Found | Not Found | Not Found | High | High |
| | | | | | High | High | High | High | Not Found | Not Found | Not Found | Not Found | Not Found | Not Found |
| | 3854 | 2439 | 6610 | 4401 | High | High | High | High | Not Found | Not Found | Not Found | Not Found | High | High |
| | | | | | High | High | High | High | Not Found | Not Found | Not Found | Not Found | Not Found | Not Found |
| 2427 | | | | | High | High | High | High | High | High | High | High | Not Found | Not Found |
| | | | | | High | High | High | High | Not Found | Not Found | Not Found | Not Found | Not Found | Not Found |
| 2851 | | | | | High | High | High | High | High | High | High | High | Not Found | Not Found |
| 2133 | | | | | High | High | High | High | High | High | High | High | Not Found | Not Found |

| | | | | | | | | | | | | | | |
|---|---|---|---|---|---|---|---|---|---|---|---|---|---|---|
| | 1213 | 1013 | 1206 | 1219 | Not Founc | Not Founc | Not Founc | Not Founc | Not Founc | Not Founc | Not Founc | Not Founc | High | High |
| 3265 | | | | | High | High | High | High | High | High | High | High | Not Founc | Not Founc |
| | | | | | High | High | High | High | Not Founc | Not Founc | Not Founc | Not Founc | Not Founc | Not Founc |
| 2374 | 3121 | 2258 | 3531 | 2713 | Not Founc | Not Founc | Not Founc | Not Founc | High | High | High | High | High | High |
| 3767 | 1821 | 1005 | 1347 | 1433 | Not Founc | Not Founc | Not Founc | Not Founc | High | High | High | High | High | High |
| 800 | | | | | Not Founc | Not Founc | Not Founc | Not Founc | High | High | High | High | Not Founc | Not Founc |
| 413 | 1772 | 1421 | 1850 | 2035 | Not Founc | Not Founc | Not Founc | Not Founc | High | High | High | High | High | High |
| 742 | | | | | Not Founc | Not Founc | Not Founc | Not Founc | High | High | High | High | Not Founc | Not Founc |
| | 1676 | 883 | 2297 | 1650 | High | High | High | High | Not Founc | Not Founc | Not Founc | Not Founc | High | High |
| | 2939 | 1421 | 4812 | 2818 | Not Founc | Not Founc | Not Founc | Not Founc | Not Founc | Not Founc | Not Founc | Not Founc | High | High |
| | | | | | Not Founc | Not Founc | Not Founc | Not Founc | Not Founc | Not Founc | Not Founc | Not Founc | Not Founc | Not Founc |
| 5387 | | | | | Not Founc | Not Founc | Not Founc | Not Founc | High | High | High | High | Not Founc | Not Founc |
| 4434 | 2587 | 1878 | 3297 | 4072 | Not Founc | Not Founc | Not Founc | Not Founc | High | High | High | High | High | High |
| | | | | | High | High | High | High | Not Founc | Not Founc | Not Founc | Not Founc | Not Founc | Not Founc |
| | 2631 | 2070 | 3551 | 3437 | High | High | High | High | Not Founc | Not Founc | Not Founc | Not Founc | High | High |
| 2717 | | | | | High | High | High | High | High | High | High | High | Not Founc | Not Founc |
| | 1936 | 1388 | 2456 | 2262 | Not Founc | Not Founc | Not Founc | Not Founc | Not Founc | Not Founc | Not Founc | Not Founc | High | High |
| | 1761 | 1503 | 2115 | 2100 | Not Founc | Not Founc | Not Founc | Not Founc | Not Founc | Not Founc | Not Founc | Not Founc | High | High |
| | | | | | High | High | High | High | Not Founc | Not Founc | Not Founc | Not Founc | Not Founc | Not Founc |
| | 1008 | 765 | 799 | 820 | Not Founc | Not Founc | Not Founc | Not Founc | Not Founc | Not Founc | Not Founc | Not Founc | High | High |
| 6491 | | | | | Not Founc | Not Founc | Not Founc | Not Founc | High | High | High | High | Not Founc | Not Founc |
| | 1242 | 959 | 1692 | 1565 | High | High | High | High | Not Founc | Not Founc | Not Founc | Not Founc | High | High |
| | 2368 | 1761 | 1459 | 1731 | Not Founc | Not Founc | Not Founc | Not Founc | Not Founc | Not Founc | Not Founc | Not Founc | High | High |
| | | | | | High | High | High | High | Not Founc | Not Founc | Not Founc | Not Founc | Not Founc | Not Founc |
| 7204 | 1195 | 878 | 1390 | 1382 | Not Founc | Not Founc | Not Founc | Not Founc | High | High | High | High | High | High |
| | | | | | High | High | High | High | Not Founc | Not Founc | Not Founc | Not Founc | Not Founc | Not Founc |
| 4918 | 3265 | 2303 | 3601 | 3873 | Not Founc | Not Founc | Not Founc | Not Founc | High | High | High | High | High | High |
| 904 | | | | | Not Founc | Not Founc | Not Founc | Not Founc | High | High | High | High | Not Founc | Not Founc |
| | | | | | High | High | High | High | Not Founc | Not Founc | Not Founc | Not Founc | Not Founc | Not Founc |
| 2478 | | | | | High | High | High | High | High | High | High | High | Not Founc | Not Founc |
| 6621 | | | | | Not Founc | Not Founc | Not Founc | Not Founc | High | High | High | High | Not Founc | Not Founc |

| | | | | | | | | | | | | | | |
|---|---|---|---|---|---|---|---|---|---|---|---|---|---|---|
| | | | | | High | High | High | High | Not Found | Not Found | Not Found | Not Found | Not Found | Not Found |
| | 919 | 793 | 1558 | 1161 | Not Found | Not Found | Not Found | Not Found | Not Found | Not Found | Not Found | Not Found | High | High |
| | 2836 | 1797 | 3263 | 3237 | High | High | High | High | Not Found | Not Found | Not Found | Not Found | High | High |
| 1623 | 3485 | 2318 | 3418 | 4344 | Not Found | Not Found | Not Found | Not Found | High | High | High | High | High | High |
| 2057 | | | | | High | High | High | High | High | High | High | High | Not Found | Not Found |
| 2239 | | | | | Not Found | Not Found | Not Found | Not Found | High | High | High | High | Not Found | Not Found |
| 3781 | | | | | High | High | High | High | High | High | High | High | Not Found | Not Found |
| | 1959 | 1532 | 1640 | 1782 | Not Found | Not Found | Not Found | Not Found | Not Found | Not Found | Not Found | Not Found | High | High |
| | | | | | High | High | High | High | Not Found | Not Found | Not Found | Not Found | Not Found | Not Found |
| 4480 | | | | | High | High | High | High | High | High | High | High | Not Found | Not Found |
| 3518 | 3674 | 1986 | 6990 | 3678 | Not Found | Not Found | Not Found | Not Found | High | High | High | High | High | High |
| | | | | | Not Found | Not Found | Not Found | Not Found | Not Found | Not Found | Not Found | Not Found | Not Found | Not Found |
| 844 | | | | | High | High | High | High | High | High | High | High | Not Found | Not Found |
| | 2066 | 1455 | 2504 | 2243 | High | High | High | High | Not Found | Not Found | Not Found | Not Found | High | High |
| 1929 | | | | | Not Found | Not Found | Not Found | Not Found | High | High | High | High | Not Found | Not Found |
| | | | | | Not Found | Not Found | Not Found | Not Found | Not Found | Not Found | Not Found | Not Found | Not Found | Not Found |
| | 4208 | 2760 | 5638 | 5168 | High | High | High | High | Not Found | Not Found | Not Found | Not Found | High | High |
| 3188 | | | | | Not Found | Not Found | Not Found | Not Found | High | High | High | High | Not Found | Not Found |
| 2046 | | | | | High | High | High | High | High | High | High | High | Not Found | Not Found |
| | 788 | 578 | 1745 | 1182 | High | High | High | High | Not Found | Not Found | Not Found | Not Found | High | High |
| 2332 | 1008 | 952 | 2029 | 2534 | Not Found | Not Found | Not Found | Not Found | High | High | High | High | High | High |
| | 76 | 65 | 85 | 66 | High | High | High | Not Found | Not Found | Not Found | Not Found | Not Found | High | High |
| 4466 | | | | | High | High | High | High | High | High | High | High | Not Found | Not Found |
| | | | | | High | High | High | High | Not Found | Not Found | Not Found | Not Found | Not Found | Not Found |
| 2046 | 1755 | 1421 | 2244 | 2577 | Not Found | Not Found | Not Found | Not Found | High | High | High | High | High | High |
| | 2549 | 1272 | 2285 | 2036 | Not Found | Not Found | Not Found | Not Found | Not Found | Not Found | Not Found | Not Found | High | High |
| | | | | | High | High | High | High | Not Found | Not Found | Not Found | Not Found | Not Found | Not Found |
| 4964 | | | | | Not Found | Not Found | Not Found | Not Found | High | High | High | High | Not Found | Not Found |
| 3076 | 2463 | 1615 | 2597 | 2631 | Not Found | Not Found | Not Found | Not Found | High | High | High | High | High | High |
| | 653 | 462 | 809 | 799 | Not Found | Not Found | Not Found | Not Found | Not Found | Not Found | Not Found | Not Found | High | High |
| 1891 | | | | | Not Found | Not Found | Not Found | Not Found | High | High | High | High | Not Found | Not Found |

| | | | | | | | | | | | | | | |
|---|---|---|---|---|---|---|---|---|---|---|---|---|---|---|
| | 2862 | 2043 | 2550 | 2762 | High | High | High | High | Not Founc | Not Founc | Not Founc | Not Founc | High | High |
| | 3997 | 2843 | 4511 | 3977 | Not Founc | Not Founc | Not Founc | Not Founc | Not Founc | Not Founc | Not Founc | Not Founc | High | High |
| | | | | | High | High | High | High | Not Founc | Not Founc | Not Founc | Not Founc | Not Founc | Not Founc |
| 1601 | | | | | Not Founc | Not Founc | Not Founc | Not Founc | High | High | High | High | Not Founc | Not Founc |
| 4260 | | | | | Not Founc | Not Founc | Not Founc | Not Founc | High | High | High | High | Not Founc | Not Founc |
| | 2784 | 1860 | 2927 | 3006 | Not Founc | Not Founc | Not Founc | Not Founc | Not Founc | Not Founc | Not Founc | Not Founc | High | High |
| 888 | 1067 | 597 | 1237 | 915 | Not Founc | Not Founc | Not Founc | Not Founc | High | High | High | High | High | High |
| 10908 | 6116 | 4933 | 6960 | 8192 | Not Founc | Not Founc | Not Founc | Not Founc | High | High | High | High | High | High |
| 5810 | 6353 | 4988 | 6275 | 5107 | Not Founc | Not Founc | Not Founc | Not Founc | High | High | High | High | High | High |
| | | | | | Not Founc | Not Founc | Not Founc | Not Founc | High | High | High | High | High | High |
| 1258 | | | | | High | High | High | High | High | High | High | High | Not Founc | Not Founc |
| | | | | | High | High | High | High | Not Founc | Not Founc | Not Founc | Not Founc | Not Founc | Not Founc |
| | | | | | High | High | High | High | Not Founc | Not Founc | Not Founc | Not Founc | Not Founc | Not Founc |
| | 1011 | 748 | 2141 | 1308 | High | High | High | High | Not Founc | Not Founc | Not Founc | Not Founc | High | High |
| | | | | | Not Founc | Not Founc | Not Founc | Not Founc | Not Founc | Not Founc | Not Founc | Not Founc | Not Founc | Not Founc |
| | | | | | High | High | High | High | Not Founc | Not Founc | Not Founc | Not Founc | Not Founc | Not Founc |
| 1449 | | | | | Not Founc | Not Founc | Not Founc | Not Founc | High | High | High | High | Not Founc | Not Founc |
| | 1322 | 771 | 1272 | 1055 | Not Founc | Not Founc | Not Founc | Not Founc | Not Founc | Not Founc | Not Founc | Not Founc | High | High |
| | 1469 | 1109 | 1583 | 3277 | High | High | High | High | Not Founc | Not Founc | Not Founc | Not Founc | High | High |
| 9338 | | | | | Not Founc | Not Founc | Not Founc | Not Founc | High | High | High | High | Not Founc | Not Founc |
| 391 | | | | | Not Founc | Not Founc | Not Founc | Not Founc | High | High | High | High | Not Founc | Not Founc |
| | 2893 | 2504 | 2639 | 3961 | High | High | High | High | Not Founc | Not Founc | Not Founc | Not Founc | High | High |
| 2348 | | | | | Not Founc | Not Founc | Not Founc | Not Founc | High | High | High | High | Not Founc | Not Founc |
| 2166 | | | | | High | High | High | High | High | High | High | High | Not Founc | Not Founc |
| 2645 | 925 | 855 | 1440 | 1294 | Not Founc | Not Founc | Not Founc | Not Founc | High | High | High | High | High | High |
| 1490 | 668 | 557 | 2412 | 1416 | Not Founc | Not Founc | Not Founc | Not Founc | High | High | High | High | High | High |
| | 4010 | 3013 | 4076 | 4093 | Not Founc | Not Founc | Not Founc | Not Founc | Not Founc | Not Founc | Not Founc | Not Founc | High | High |
| 6374 | 6836 | 4062 | 5929 | 4977 | Not Founc | Not Founc | Not Founc | Not Founc | High | High | High | High | High | High |
| | | | | | High | High | High | High | Not Founc | Not Founc | Not Founc | Not Founc | Not Founc | Not Founc |
| | | | | | High | High | High | High | Not Founc | Not Founc | Not Founc | Not Founc | Not Founc | Not Founc |
| 688 | | | | | Not Founc | Not Founc | Not Founc | Not Founc | High | High | High | High | Not Founc | Not Founc |

| | | | | | | | | | | | | | | |
|---|---|---|---|---|---|---|---|---|---|---|---|---|---|---|
| | 11165 | 9777 | 11099 | 11826 | High | High | High | High | Not Founc | Not Founc | Not Founc | Not Founc | High | High |
| | | | | | High | High | High | High | Not Founc | Not Founc | Not Founc | Not Founc | Not Founc | Not Founc |
| 502 | | | | | Not Founc | Not Founc | Not Founc | Not Founc | High | High | High | High | Not Founc | Not Founc |
| 1325 | 2270 | 872 | 3207 | 1128 | Not Founc | Not Founc | Not Founc | Not Founc | High | High | High | High | High | High |
| 5716 | | | | | Not Founc | Not Founc | Not Founc | Not Founc | High | High | High | High | Not Founc | Not Founc |
| | 4092 | 4034 | 7211 | 7183 | High | High | High | High | Not Founc | Not Founc | Not Founc | Not Founc | High | High |
| 5768 | 1530 | 1047 | 1705 | 1753 | Not Founc | Not Founc | Not Founc | Not Founc | High | High | High | High | High | High |
| | 7908 | 5187 | 7872 | 6502 | Not Founc | Not Founc | Not Founc | Not Founc | Not Founc | Not Founc | Not Founc | Not Founc | High | High |
| 4329 | | | | | High | High | High | High | High | High | High | High | Not Founc | Not Founc |
| | 2452 | 1809 | 2308 | 2096 | Not Founc | Not Founc | Not Founc | Not Founc | Not Founc | Not Founc | Not Founc | Not Founc | High | High |
| 1190 | | | | | High | High | High | High | High | High | High | High | Not Founc | Not Founc |
| 1773 | | | | | High | High | High | High | High | High | High | High | Not Founc | Not Founc |
| | | | | | High | High | High | High | Not Founc | Not Founc | Not Founc | Not Founc | Not Founc | Not Founc |
| 1874 | | | | | High | High | High | High | High | High | High | High | Not Founc | Not Founc |
| | | | | | High | High | High | High | Not Founc | Not Founc | Not Founc | Not Founc | Not Founc | Not Founc |
| | | | | | High | High | High | High | Not Founc | Not Founc | Not Founc | Not Founc | Not Founc | Not Founc |
| 2082 | | | | | Not Founc | Not Founc | Not Founc | Not Founc | High | High | High | High | Not Founc | Not Founc |
| | | | | | Not Founc | Not Founc | Not Founc | Not Founc | Not Founc | Not Founc | Not Founc | Not Founc | Not Founc | Not Founc |
| | 987 | 754 | 1552 | 1172 | Not Founc | Not Founc | Not Founc | Not Founc | Not Founc | Not Founc | Not Founc | Not Founc | High | High |
| 2235 | | | | | Not Founc | Not Founc | Not Founc | Not Founc | High | High | High | High | Not Founc | Not Founc |
| | | | | | Not Founc | Not Founc | Not Founc | Not Founc | High | High | High | High | Not Founc | Not Founc |
| | 215 | 237 | 238 | 272 | Not Founc | Not Founc | Not Founc | Not Founc | Not Founc | Not Founc | Not Founc | Not Founc | High | High |
| 4166 | | | | | High | High | High | High | High | High | High | High | Not Founc | Not Founc |
| 3724 | 3922 | 2580 | 2602 | 2986 | Not Founc | Not Founc | Not Founc | Not Founc | High | High | High | High | High | High |
| | 1882 | 1697 | 2366 | 2574 | High | High | High | High | Not Founc | Not Founc | Not Founc | Not Founc | High | High |
| | 765 | 627 | 663 | 769 | Not Founc | Not Founc | Not Founc | Not Founc | Not Founc | Not Founc | Not Founc | Not Founc | High | High |
| | 907 | 628 | 1487 | 1095 | Not Founc | Not Founc | Not Founc | Not Founc | Not Founc | Not Founc | Not Founc | Not Founc | High | High |
| | | | | | High | High | High | High | Not Founc | Not Founc | Not Founc | Not Founc | Not Founc | Not Founc |
| 1174 | | | | | Not Founc | Not Founc | Not Founc | Not Founc | High | High | High | High | Not Founc | Not Founc |
| 5393 | | | | | High | High | High | High | High | High | High | High | Not Founc | Not Founc |
| 2386 | 2107 | 2279 | 2839 | 2458 | Not Founc | Not Founc | Not Founc | Not Founc | High | High | High | High | High | High |

| | | | | | | | | | | | | | | |
|---|---|---|---|---|---|---|---|---|---|---|---|---|---|---|
| | 1759 | 1605 | 5691 | 2173 | Not Found | Not Found | Not Found | Not Found | Not Found | Not Found | Not Found | Not Found | High | High |
| | 1320 | 1287 | 1891 | 1956 | High | High | High | High | Not Found | Not Found | Not Found | Not Found | High | High |
| | | | | | High | High | High | High | Not Found | Not Found | Not Found | Not Found | Not Found | Not Found |
| 1624 | 1183 | 871 | 2215 | 1500 | Not Found | Not Found | Not Found | Not Found | High | High | High | High | High | High |
| | 4379 | 3548 | 5634 | 4893 | Not Found | Not Found | Not Found | Not Found | Not Found | Not Found | Not Found | Not Found | High | High |
| 2154 | | | | | High | High | High | High | High | High | High | High | Not Found | Not Found |
| | | | | | High | High | High | High | Not Found | Not Found | Not Found | Not Found | Not Found | Not Found |
| 2907 | | | | | Not Found | Not Found | Not Found | Not Found | High | High | High | High | Not Found | Not Found |
| 4842 | 3707 | 3427 | 3075 | 3728 | Not Found | Not Found | Not Found | Not Found | High | High | High | High | High | High |
| | 4189 | 3296 | 5662 | 6066 | High | High | High | High | Not Found | Not Found | Not Found | Not Found | High | High |
| 2286 | | | | | Not Found | Not Found | Not Found | Not Found | High | High | High | High | Not Found | Not Found |
| | 953 | 720 | 1063 | 1003 | Not Found | Not Found | Not Found | Not Found | Not Found | Not Found | Not Found | Not Found | High | High |
| 1748 | | | | | High | High | High | High | High | High | High | High | Not Found | Not Found |
| 1225 | | | | | Not Found | Not Found | Not Found | Not Found | High | High | High | High | Not Found | Not Found |
| | | | | | High | High | High | High | Not Found | Not Found | Not Found | Not Found | Not Found | Not Found |
| | 2021 | 1762 | 1786 | 2417 | Not Found | Not Found | Not Found | Not Found | Not Found | Not Found | Not Found | Not Found | High | High |
| 4114 | | | | | High | High | High | High | High | High | High | High | Not Found | Not Found |
| | 1669 | 1313 | 1717 | 1272 | High | High | High | High | Not Found | Not Found | Not Found | Not Found | High | High |
| | | | | | Not Found | Not Found | Not Found | Not Found | Not Found | Not Found | Not Found | Not Found | Not Found | Not Found |
| | | | | | Not Found | Not Found | Not Found | Not Found | Not Found | Not Found | Not Found | Not Found | Not Found | Not Found |
| 4669 | | | | | Not Found | Not Found | Not Found | Not Found | High | High | High | High | Not Found | Not Found |
| 2623 | 6655 | 2654 | 8154 | 4226 | Not Found | Not Found | Not Found | Not Found | High | High | High | High | High | High |
| | 1581 | 791 | 2260 | 1628 | Not Found | Not Found | Not Found | Not Found | High | High | High | High | High | High |
| | 2300 | 2002 | 2588 | 2588 | Not Found | Not Found | Not Found | Not Found | Not Found | Not Found | Not Found | Not Found | High | High |
| 4540 | | | | | High | High | High | High | High | High | High | High | Not Found | Not Found |
| | 2857 | 2333 | 2909 | 3429 | Not Found | Not Found | Not Found | Not Found | Not Found | Not Found | Not Found | Not Found | High | High |
| | 3814 | 2654 | 3794 | 3741 | Not Found | Not Found | Not Found | Not Found | Not Found | Not Found | Not Found | Not Found | High | High |
| | | | | | High | High | High | High | Not Found | Not Found | Not Found | Not Found | Not Found | Not Found |
| 1326 | | | | | Not Found | Not Found | Not Found | Not Found | High | High | High | High | Not Found | Not Found |
| | | | | | High | High | High | High | Not Found | Not Found | Not Found | Not Found | Not Found | Not Found |
| | | | | | Not Found | Not Found | Not Found | Not Found | Not Found | Not Found | Not Found | Not Found | Not Found | Not Found |

| | | | | | | | | | | | | | | |
|---|---|---|---|---|---|---|---|---|---|---|---|---|---|---|
| 1528 | | | | | Not Found | Not Found | Not Found | Not Found | High | High | High | High | Not Found | Not Found |
| | 1028 | 549 | 1860 | 1386 | High | High | High | High | Not Found | Not Found | Not Found | Not Found | High | High |
| 3507 | | | | | Not Found | Not Found | Not Found | Not Found | High | High | High | High | Not Found | Not Found |
| | 2383 | 2339 | 2348 | 2599 | Not Found | Not Found | Not Found | Not Found | Not Found | Not Found | Not Found | Not Found | High | High |
| | | | | | High | High | High | High | Not Found | Not Found | Not Found | Not Found | Not Found | Not Found |
| | | | | | Not Found | Not Found | Not Found | Not Found | Not Found | Not Found | Not Found | Not Found | Not Found | Not Found |
| 1905 | | | | | Not Found | Not Found | Not Found | Not Found | High | High | High | High | Not Found | Not Found |
| | 2802 | 1319 | 2854 | 2104 | Not Found | Not Found | Not Found | Not Found | Not Found | Not Found | Not Found | Not Found | High | High |
| | | | | | High | High | High | High | Not Found | Not Found | Not Found | Not Found | Not Found | Not Found |
| | | | | | High | High | High | High | Not Found | Not Found | Not Found | Not Found | Not Found | Not Found |
| | | | | | Not Found | Not Found | Not Found | Not Found | Not Found | Not Found | Not Found | Not Found | Not Found | Not Found |
| | 1591 | 1327 | 1598 | 1743 | Not Found | Not Found | Not Found | Not Found | Not Found | Not Found | Not Found | Not Found | High | High |
| | | | | | High | High | High | High | Not Found | Not Found | Not Found | Not Found | Not Found | Not Found |
| | | | | | Not Found | Not Found | Not Found | Not Found | Not Found | Not Found | Not Found | Not Found | High | High |
| | | | | | High | High | High | High | Not Found | Not Found | Not Found | Not Found | Not Found | Not Found |
| | 1768 | 1635 | 2151 | 2347 | Not Found | Not Found | Not Found | Not Found | Not Found | Not Found | Not Found | Not Found | High | High |
| | | | | | High | High | High | High | Not Found | Not Found | Not Found | Not Found | Not Found | Not Found |
| | | | | | High | High | High | High | Not Found | Not Found | Not Found | Not Found | Not Found | Not Found |
| 2048 | | | | | Not Found | Not Found | Not Found | Not Found | High | High | High | High | Not Found | Not Found |
| 3520 | | | | | Not Found | Not Found | Not Found | Not Found | High | High | High | High | Not Found | Not Found |
| 2167 | | | | | High | High | High | High | High | High | High | High | Not Found | Not Found |
| 1854 | | | | | Not Found | Not Found | Not Found | Not Found | High | High | High | High | Not Found | Not Found |
| | | | | | Not Found | Not Found | Not Found | Not Found | High | High | High | High | Not Found | Not Found |
| | 887 | 636 | 1474 | 1149 | Not Found | Not Found | Not Found | Not Found | Not Found | Not Found | Not Found | Not Found | High | High |
| | | | | | High | High | High | High | Not Found | Not Found | Not Found | Not Found | Not Found | Not Found |
| 2060 | | | | | Not Found | Not Found | Not Found | Not Found | High | High | High | High | Not Found | Not Found |
| | | | | | Not Found | Not Found | Not Found | Not Found | High | High | High | High | Not Found | Not Found |
| 1999 | | | | | Not Found | Not Found | Not Found | Not Found | High | High | High | High | Not Found | Not Found |
| | 2426 | 2188 | 2768 | 2783 | Not Found | Not Found | Not Found | Not Found | Not Found | Not Found | Not Found | Not Found | High | High |
| | | | | | High | High | High | High | Not Found | Not Found | Not Found | Not Found | Not Found | Not Found |
| | 1652 | 1410 | 1633 | 1843 | Not Found | Not Found | Not Found | Not Found | Not Found | Not Found | Not Found | Not Found | High | High |

| | | | | | | | | | | | | | | |
|---|---|---|---|---|---|---|---|---|---|---|---|---|---|---|
| | 1440 | 980 | 1290 | 1087 | Not Found | Not Found | Not Found | Not Found | Not Found | Not Found | Not Found | Not Found | High | High |
| 672 | | | | | Not Found | Not Found | Not Found | Not Found | High | High | High | High | Not Found | Not Found |
| | | | | | Not Found | Not Found | Not Found | Not Found | Not Found | Not Found | Not Found | Not Found | Not Found | Not Found |
| | 948 | 803 | 701 | 949 | Not Found | Not Found | Not Found | Not Found | Not Found | Not Found | Not Found | Not Found | High | High |
| 1510 | | | | | Not Found | Not Found | Not Found | Not Found | High | High | High | High | Not Found | Not Found |
| 2605 | | | | | Not Found | Not Found | Not Found | Not Found | High | High | High | High | Not Found | Not Found |
| | | | | | High | High | High | High | Not Found | Not Found | Not Found | Not Found | Not Found | Not Found |
| | 1489 | 1426 | 1216 | 1619 | Not Found | Not Found | Not Found | Not Found | Not Found | Not Found | Not Found | Not Found | High | High |
| 3202 | | | | | Not Found | Not Found | Not Found | Not Found | High | High | High | High | Not Found | Not Found |
| 878 | | | | | Not Found | Not Found | Not Found | Not Found | High | High | High | High | Not Found | Not Found |
| 1635 | | | | | Not Found | Not Found | Not Found | Not Found | High | High | High | High | Not Found | Not Found |
| | | | | | High | High | High | High | Not Found | Not Found | Not Found | Not Found | Not Found | Not Found |
| 1133 | | | | | Not Found | Not Found | Not Found | Not Found | High | High | High | High | Not Found | Not Found |
| | 2370 | 1438 | 3242 | 2649 | Not Found | Not Found | Not Found | Not Found | Not Found | Not Found | Not Found | Not Found | High | High |
| | | | | | High | High | High | High | Not Found | Not Found | Not Found | Not Found | Not Found | Not Found |
| | | | | | High | High | High | High | Not Found | Not Found | Not Found | Not Found | Not Found | Not Found |
| | | | | | High | High | High | High | Not Found | Not Found | Not Found | Not Found | Not Found | Not Found |
| | | | | | High | High | High | High | Not Found | Not Found | Not Found | Not Found | Not Found | Not Found |
| | 3778 | 2916 | 3878 | 3498 | Not Found | Not Found | Not Found | Not Found | Not Found | Not Found | Not Found | Not Found | High | High |
| | | | | | High | High | High | High | Not Found | Not Found | Not Found | Not Found | Not Found | Not Found |
| 1456 | | | | | Not Found | Not Found | Not Found | Not Found | High | High | High | High | Not Found | Not Found |
| | 2683 | 2143 | 4452 | 3670 | Not Found | Not Found | Not Found | Not Found | Not Found | Not Found | Not Found | Not Found | High | High |
| | 689 | 497 | 983 | 787 | Not Found | Not Found | Not Found | Not Found | Not Found | Not Found | Not Found | Not Found | High | High |
| 3477 | | | | | Not Found | Not Found | Not Found | Not Found | High | High | High | High | Not Found | Not Found |
| | | | | | High | High | High | High | Not Found | Not Found | Not Found | Not Found | Not Found | Not Found |
| | | | | | High | High | High | High | Not Found | Not Found | Not Found | Not Found | Not Found | Not Found |
| | 713 | 640 | 705 | 863 | Not Found | Not Found | Not Found | Not Found | Not Found | Not Found | Not Found | Not Found | High | High |
| 1194 | | | | | Not Found | Not Found | Not Found | Not Found | High | High | High | High | Not Found | Not Found |
| | 1331 | 1115 | 1251 | 1418 | Not Found | Not Found | Not Found | Not Found | Not Found | Not Found | Not Found | Not Found | High | High |
| 3560 | | | | | Not Found | Not Found | Not Found | Not Found | High | High | High | High | Not Found | Not Found |
| | 866 | 806 | 597 | 844 | Not Found | Not Found | Not Found | Not Found | Not Found | Not Found | Not Found | Not Found | High | High |

| | | | | | | | | | | | | | | |
|---|---|---|---|---|---|---|---|---|---|---|---|---|---|---|
| | 4651 | 3137 | 4846 | 4388 | Not Found | Not Found | Not Found | Not Found | Not Found | Not Found | Not Found | Not Found | High | High |
| | | | | | High | High | High | High | Not Found | Not Found | Not Found | Not Found | Not Found | Not Found |
| | 1003 | 555 | 1669 | 928 | Not Found | Not Found | Not Found | Not Found | Not Found | Not Found | Not Found | Not Found | High | High |
| | 1431 | 1090 | 1861 | 2059 | Not Found | Not Found | Not Found | Not Found | Not Found | Not Found | Not Found | Not Found | High | High |
| 1808 | | | | | Not Found | Not Found | Not Found | Not Found | High | High | High | High | Not Found | Not Found |
| | 666 | 750 | 1007 | 1178 | Not Found | Not Found | Not Found | Not Found | Not Found | Not Found | Not Found | Not Found | High | High |
| | | | | | High | High | High | High | Not Found | Not Found | Not Found | Not Found | Not Found | Not Found |
| 3313 | | | | | Not Found | Not Found | Not Found | Not Found | High | High | High | High | Not Found | Not Found |
| | | | | | High | High | High | High | Not Found | Not Found | Not Found | Not Found | Not Found | Not Found |
| | 3644 | 3945 | 3194 | 5803 | Not Found | Not Found | Not Found | Not Found | Not Found | Not Found | Not Found | Not Found | High | High |
| 4819 | | | | | Not Found | Not Found | Not Found | Not Found | High | High | High | High | Not Found | Not Found |
| 2217 | | | | | Not Found | Not Found | Not Found | Not Found | High | High | High | High | Not Found | Not Found |
| 2144 | | | | | Not Found | Not Found | Not Found | Not Found | High | High | High | High | Not Found | Not Found |
| | | | | | Not Found | Not Found | Not Found | Not Found | Not Found | Not Found | Not Found | Not Found | Not Found | Not Found |
| | | | | | Not Found | Not Found | Not Found | Not Found | Not Found | Not Found | Not Found | Not Found | Not Found | Not Found |
| 5115 | | | | | Not Found | Not Found | Not Found | Not Found | High | High | High | High | Not Found | Not Found |
| | | | | | High | High | High | High | Not Found | Not Found | Not Found | Not Found | Not Found | Not Found |
| | 1945 | 1295 | 2757 | 2019 | Not Found | Not Found | Not Found | Not Found | Not Found | Not Found | Not Found | Not Found | High | High |
| | 2103 | 860 | 2735 | 2017 | Not Found | Not Found | Not Found | Not Found | Not Found | Not Found | Not Found | Not Found | High | High |
| | 1916 | 1415 | 2250 | 2134 | Not Found | Not Found | Not Found | Not Found | Not Found | Not Found | Not Found | Not Found | High | High |
| | | | | | High | High | High | High | Not Found | Not Found | Not Found | Not Found | Not Found | Not Found |
| | | | | | High | High | High | High | Not Found | Not Found | Not Found | Not Found | Not Found | Not Found |
| 822 | | | | | Not Found | Not Found | Not Found | Not Found | High | High | High | High | Not Found | Not Found |
| 1315 | | | | | Not Found | Not Found | Not Found | Not Found | High | High | High | High | Not Found | Not Found |
| 1262 | | | | | Not Found | Not Found | Not Found | Not Found | High | High | High | High | Not Found | Not Found |
| 647 | | | | | Not Found | Not Found | Not Found | Not Found | High | High | High | High | Not Found | Not Found |
| | | | | | High | High | High | High | Not Found | Not Found | Not Found | Not Found | Not Found | Not Found |
| | | | | | High | High | High | High | Not Found | Not Found | Not Found | Not Found | Not Found | Not Found |
| | 1337 | 1116 | 2154 | 1488 | Not Found | Not Found | Not Found | Not Found | Not Found | Not Found | Not Found | Not Found | High | High |
| | | | | | High | High | High | High | Not Found | Not Found | Not Found | Not Found | Not Found | Not Found |
| | 896 | 443 | 1047 | 783 | Not Found | Not Found | Not Found | Not Found | Not Found | Not Found | Not Found | Not Found | High | High |

| | | | | | | | | | | | | | | |
|---|---|---|---|---|---|---|---|---|---|---|---|---|---|---|
| | | | | | Not Found | Not Found | Not Found | Not Found | Not Found | Not Found | Not Found | Not Found | Not Found | Not Found |
| | 2471 | 1739 | 2123 | 2277 | Not Found | Not Found | Not Found | Not Found | Not Found | Not Found | Not Found | Not Found | High | High |
| | | | | | High | High | High | High | Not Found | Not Found | Not Found | Not Found | Not Found | Not Found |
| | 3225 | 2468 | 2594 | 2453 | Not Found | Not Found | Not Found | Not Found | Not Found | Not Found | Not Found | Not Found | High | High |
| | 3055 | 2112 | 3940 | 3509 | Not Found | Not Found | Not Found | Not Found | Not Found | Not Found | Not Found | Not Found | High | High |
| | | | | | High | High | High | High | Not Found | Not Found | Not Found | Not Found | Not Found | Not Found |
| | | | | | Not Found | Not Found | Not Found | Not Found | Not Found | Not Found | Not Found | Not Found | Not Found | Not Found |
| 1903 | | | | | Not Found | Not Found | Not Found | Not Found | High | High | High | High | Not Found | Not Found |
| | 1392 | 1013 | 1504 | 1406 | Not Found | Not Found | Not Found | Not Found | Not Found | Not Found | Not Found | Not Found | High | High |
| | 2692 | 2465 | 4758 | 3343 | Not Found | Not Found | Not Found | Not Found | Not Found | Not Found | Not Found | Not Found | High | High |
| 3182 | | | | | Not Found | Not Found | Not Found | Not Found | High | High | High | High | Not Found | Not Found |
| 4636 | | | | | Not Found | Not Found | Not Found | Not Found | High | High | High | High | Not Found | Not Found |
| 4114 | | | | | Not Found | Not Found | Not Found | Not Found | High | High | High | High | Not Found | Not Found |
| | | | | | High | High | High | High | Not Found | Not Found | Not Found | Not Found | Not Found | Not Found |
| 1960 | | | | | Not Found | Not Found | Not Found | Not Found | High | High | High | High | Not Found | Not Found |
| 2411 | | | | | Not Found | Not Found | Not Found | Not Found | High | High | High | High | Not Found | Not Found |
| | | | | | Not Found | Not Found | Not Found | Not Found | Not Found | Not Found | Not Found | Not Found | Not Found | Not Found |
| | 3033 | 2058 | 3177 | 3148 | Not Found | Not Found | Not Found | Not Found | Not Found | Not Found | Not Found | Not Found | High | High |
| | 866 | 1359 | 6958 | 3096 | Not Found | Not Found | Not Found | Not Found | Not Found | Not Found | Not Found | Not Found | High | High |
| 2374 | | | | | Not Found | Not Found | Not Found | Not Found | High | High | High | High | Not Found | Not Found |
| | | | | | High | High | High | High | Not Found | Not Found | Not Found | Not Found | Not Found | Not Found |
| | | | | | High | High | High | High | Not Found | Not Found | Not Found | Not Found | Not Found | Not Found |
| 2134 | | | | | Not Found | Not Found | Not Found | Not Found | High | High | High | High | Not Found | Not Found |
| 3747 | | | | | Not Found | Not Found | Not Found | Not Found | High | High | High | High | Not Found | Not Found |
| 759 | | | | | Not Found | Not Found | Not Found | Not Found | High | High | High | High | Not Found | Not Found |
| | | | | | High | High | High | High | Not Found | Not Found | Not Found | Not Found | Not Found | Not Found |
| | 3641 | 3491 | 3013 | 4093 | Not Found | Not Found | Not Found | Not Found | Not Found | Not Found | Not Found | Not Found | High | High |
| | | | | | Not Found | Not Found | Not Found | Not Found | Not Found | Not Found | Not Found | Not Found | Not Found | Not Found |
| | | | | | Not Found | Not Found | Not Found | Not Found | Not Found | Not Found | Not Found | Not Found | Not Found | Not Found |
| | | | | | High | High | High | High | Not Found | Not Found | Not Found | Not Found | Not Found | Not Found |
| | | | | | Not Found | Not Found | Not Found | Not Found | Not Found | Not Found | Not Found | Not Found | Not Found | Not Found |

| | | | | | | | | | | | | | | |
|---|---|---|---|---|---|---|---|---|---|---|---|---|---|---|
| 2305 | | | | | Not Found | Not Found | Not Found | Not Found | High | High | High | High | Not Found | Not Found |
| 665 | | | | | Not Found | Not Found | Not Found | Not Found | High | High | High | High | Not Found | Not Found |
| | | | | | Not Found | Not Found | Not Found | Not Found | Not Found | Not Found | Not Found | Not Found | Not Found | Not Found |
| | | | | | Not Found | Not Found | Not Found | Not Found | Not Found | Not Found | Not Found | Not Found | Not Found | Not Found |
| | | | | | Not Found | Not Found | Not Found | Not Found | Not Found | Not Found | Not Found | Not Found | Not Found | Not Found |
| | | | | | High | High | High | High | Not Found | Not Found | Not Found | Not Found | Not Found | Not Found |
| | | | | | High | High | High | High | Not Found | Not Found | Not Found | Not Found | Not Found | Not Found |
| | | | | | Not Found | Not Found | Not Found | Not Found | Not Found | Not Found | Not Found | Not Found | Not Found | Not Found |
| | 703 | 348 | 754 | 466 | Not Found | Not Found | Not Found | Not Found | Not Found | Not Found | Not Found | Not Found | High | High |
| 2871 | | | | | Not Found | Not Found | Not Found | Not Found | High | High | High | High | Not Found | Not Found |
| 1281 | | | | | Not Found | Not Found | Not Found | Not Found | High | High | High | High | Not Found | Not Found |
| | | | | | Not Found | Not Found | Not Found | Not Found | Not Found | Not Found | Not Found | Not Found | Not Found | Not Found |
| 5076 | | | | | Not Found | Not Found | Not Found | Not Found | High | High | High | High | Not Found | Not Found |
| 1129 | | | | | Not Found | Not Found | Not Found | Not Found | High | High | High | High | Not Found | Not Found |
| | | | | | Not Found | Not Found | Not Found | Not Found | Not Found | Not Found | Not Found | Not Found | Not Found | Not Found |
| | 2175 | 1619 | 5367 | 3807 | Not Found | Not Found | Not Found | Not Found | Not Found | Not Found | Not Found | Not Found | High | High |
| | | | | | High | High | High | High | Not Found | Not Found | Not Found | Not Found | Not Found | Not Found |
| | | | | | High | High | High | High | Not Found | Not Found | Not Found | Not Found | Not Found | Not Found |
| 5376 | | | | | Not Found | Not Found | Not Found | Not Found | High | High | High | High | Not Found | Not Found |
| | | | | | High | High | High | High | Not Found | Not Found | Not Found | Not Found | Not Found | Not Found |
| | | | | | Not Found | Not Found | Not Found | Not Found | Not Found | Not Found | Not Found | Not Found | Not Found | Not Found |
| | | | | | High | High | High | High | Not Found | Not Found | Not Found | Not Found | Not Found | Not Found |
| | | | | | High | High | High | High | Not Found | Not Found | Not Found | Not Found | Not Found | Not Found |
| | | | | | High | High | High | High | Not Found | Not Found | Not Found | Not Found | Not Found | Not Found |
| | 2915 | 2597 | 3313 | 3428 | Not Found | Not Found | Not Found | Not Found | Not Found | Not Found | Not Found | Not Found | High | High |
| | 2385 | 1687 | 3172 | 2447 | Not Found | Not Found | Not Found | Not Found | Not Found | Not Found | Not Found | Not Found | High | High |
| | 2128 | 1708 | 2258 | 2329 | Not Found | Not Found | Not Found | Not Found | Not Found | Not Found | Not Found | Not Found | High | High |
| | 2103 | 1466 | 2533 | 2451 | Not Found | Not Found | Not Found | Not Found | Not Found | Not Found | Not Found | Not Found | High | High |
| | | | | | Not Found | Not Found | Not Found | Not Found | Not Found | Not Found | Not Found | Not Found | Not Found | Not Found |
| | | | | | High | High | High | High | Not Found | Not Found | Not Found | Not Found | Not Found | Not Found |
| | 1182 | 830 | 1711 | 1558 | Not Found | Not Found | Not Found | Not Found | Not Found | Not Found | Not Found | Not Found | High | High |

| | | | | | | | | | | | | | | |
|---|---|---|---|---|---|---|---|---|---|---|---|---|---|---|
| 416 | | | | | Not Founc | Not Founc | Not Founc | Not Founc | High | High | High | High | Not Founc | Not Founc |
| | | | | | High | High | High | High | Not Founc | Not Founc | Not Founc | Not Founc | Not Founc | Not Founc |
| | | | | | Not Founc | Not Founc | Not Founc | Not Founc | Not Founc | Not Founc | Not Founc | Not Founc | Not Founc | Not Founc |
| | | | | | High | High | High | High | Not Founc | Not Founc | Not Founc | Not Founc | Not Founc | Not Founc |
| | 1999 | 2055 | 1284 | 2520 | Not Founc | Not Founc | Not Founc | Not Founc | Not Founc | Not Founc | Not Founc | Not Founc | High | High |
| | | | | | High | High | High | High | Not Founc | Not Founc | Not Founc | Not Founc | Not Founc | Not Founc |
| | | | | | Not Founc | Not Founc | Not Founc | Not Founc | Not Founc | Not Founc | Not Founc | Not Founc | Not Founc | Not Founc |
| 1129 | | | | | Not Founc | Not Founc | Not Founc | Not Founc | High | High | High | High | Not Founc | Not Founc |
| 680 | | | | | Not Founc | Not Founc | Not Founc | Not Founc | High | High | High | High | Not Founc | Not Founc |
| | | | | | High | High | High | High | Not Founc | Not Founc | Not Founc | Not Founc | Not Founc | Not Founc |
| 5718 | | | | | Not Founc | Not Founc | Not Founc | Not Founc | High | High | High | High | Not Founc | Not Founc |
| | | | | | High | High | High | High | Not Founc | Not Founc | Not Founc | Not Founc | Not Founc | Not Founc |
| | 710 | 454 | 966 | 715 | Not Founc | Not Founc | Not Founc | Not Founc | Not Founc | Not Founc | Not Founc | Not Founc | High | High |
| | | | | | High | High | High | High | Not Founc | Not Founc | Not Founc | Not Founc | Not Founc | Not Founc |
| | | | | | High | High | High | High | Not Founc | Not Founc | Not Founc | Not Founc | Not Founc | Not Founc |
| | | | | | Not Founc | Not Founc | Not Founc | Not Founc | Not Founc | Not Founc | Not Founc | Not Founc | Not Founc | Not Founc |
| | | | | | Not Founc | Not Founc | Not Founc | Not Founc | Not Founc | Not Founc | Not Founc | Not Founc | Not Founc | Not Founc |
| | 1252 | 1010 | 1602 | 1518 | Not Founc | Not Founc | Not Founc | Not Founc | Not Founc | Not Founc | Not Founc | Not Founc | High | High |
| | | | | | Not Founc | Not Founc | Not Founc | Not Founc | High | High | High | High | Not Founc | Not Founc |
| 756 | | | | | Not Founc | Not Founc | Not Founc | Not Founc | High | High | High | High | Not Founc | Not Founc |
| 4024 | | | | | Not Founc | Not Founc | Not Founc | Not Founc | High | High | High | High | Not Founc | Not Founc |
| 8200 | | | | | Not Founc | Not Founc | Not Founc | Not Founc | High | High | High | High | Not Founc | Not Founc |
| | | | | | High | High | High | High | Not Founc | Not Founc | Not Founc | Not Founc | Not Founc | Not Founc |
| | | | | | Not Founc | Not Founc | Not Founc | Not Founc | High | High | High | High | Not Founc | Not Founc |
| | | | | | High | High | High | High | Not Founc | Not Founc | Not Founc | Not Founc | Not Founc | Not Founc |
| | | | | | Not Founc | Not Founc | Not Founc | Not Founc | Not Founc | Not Founc | Not Founc | Not Founc | Not Founc | Not Founc |
| | 444 | 553 | 652 | 729 | Not Founc | Not Founc | Not Founc | Not Founc | Not Founc | Not Founc | Not Founc | Not Founc | High | High |
| 4401 | | | | | Not Founc | Not Founc | Not Founc | Not Founc | High | High | High | High | Not Founc | Not Founc |
| | | | | | High | High | High | High | Not Founc | Not Founc | Not Founc | Not Founc | Not Founc | Not Founc |
| 1705 | | | | | Not Founc | Not Founc | Not Founc | Not Founc | High | High | High | High | Not Founc | Not Founc |
| 1946 | | | | | Not Founc | Not Founc | Not Founc | Not Founc | High | High | High | High | Not Founc | Not Founc |

| | | | | | | | | | | | | | | |
|---|---|---|---|---|---|---|---|---|---|---|---|---|---|---|
| 1975 | | | | | Not Found | Not Found | Not Found | Not Found | High | High | High | High | Not Found | Not Found |
| 1506 | | | | | Not Found | Not Found | Not Found | Not Found | High | High | High | High | Not Found | Not Found |
| 1739 | | | | | Not Found | Not Found | Not Found | Not Found | High | High | High | High | Not Found | Not Found |
| | | | | | Not Found | Not Found | Not Found | Not Found | Not Found | Not Found | Not Found | Not Found | Not Found | Not Found |
| | | | | | High | High | High | High | Not Found | Not Found | Not Found | Not Found | Not Found | Not Found |
| | | | | | Not Found | Not Found | Not Found | Not Found | Not Found | Not Found | Not Found | Not Found | Not Found | Not Found |
| | | | | | High | High | High | High | Not Found | Not Found | Not Found | Not Found | Not Found | Not Found |
| | | | | | High | High | High | High | Not Found | Not Found | Not Found | Not Found | Not Found | Not Found |
| 3457 | | | | | Not Found | Not Found | Not Found | Not Found | High | High | High | High | Not Found | Not Found |
| | 10403 | 5397 | 13013 | 9453 | Not Found | Not Found | Not Found | Not Found | Not Found | Not Found | Not Found | Not Found | High | High |
| 2729 | | | | | Not Found | Not Found | Not Found | Not Found | High | High | High | High | Not Found | Not Found |
| 3711 | | | | | Not Found | Not Found | Not Found | Not Found | High | High | High | High | Not Found | Not Found |
| | | | | | High | High | High | High | Not Found | Not Found | Not Found | Not Found | Not Found | Not Found |
| | | | | | High | High | High | High | Not Found | Not Found | Not Found | Not Found | Not Found | Not Found |
| | 1970 | 2050 | 2295 | 2652 | Not Found | Not Found | Not Found | Not Found | Not Found | Not Found | Not Found | Not Found | High | High |
| | | | | | High | High | High | High | Not Found | Not Found | Not Found | Not Found | Not Found | Not Found |
| | | | | | High | High | High | High | Not Found | Not Found | Not Found | Not Found | Not Found | Not Found |
| | 1927 | 1292 | 3797 | 2378 | Not Found | Not Found | Not Found | Not Found | Not Found | Not Found | Not Found | Not Found | High | High |
| | | | | | High | High | High | High | Not Found | Not Found | Not Found | Not Found | Not Found | Not Found |
| | 2206 | 1358 | 2765 | 2057 | Not Found | Not Found | Not Found | Not Found | Not Found | Not Found | Not Found | Not Found | High | High |
| | | | | | High | High | High | High | Not Found | Not Found | Not Found | Not Found | Not Found | Not Found |
| | 1218 | 1163 | 1085 | 1158 | Not Found | Not Found | Not Found | Not Found | Not Found | Not Found | Not Found | Not Found | High | High |
| | | | | | High | High | High | High | Not Found | Not Found | Not Found | Not Found | Not Found | Not Found |
| | 3158 | 2109 | 4714 | 2095 | Not Found | Not Found | Not Found | Not Found | Not Found | Not Found | Not Found | Not Found | High | High |
| 1873 | | | | | Not Found | Not Found | Not Found | Not Found | High | High | High | High | Not Found | Not Found |
| | | | | | Not Found | Not Found | Not Found | Not Found | Not Found | Not Found | Not Found | Not Found | Not Found | Not Found |
| 1120 | | | | | Not Found | Not Found | Not Found | Not Found | High | High | High | High | Not Found | Not Found |
| | 4419 | 3173 | 3251 | 3131 | Not Found | Not Found | Not Found | Not Found | Not Found | Not Found | Not Found | Not Found | High | High |
| | | | | | High | High | High | High | Not Found | Not Found | Not Found | Not Found | Not Found | Not Found |
| 2376 | | | | | Not Found | Not Found | Not Found | Not Found | High | High | High | High | Not Found | Not Found |
| | | | | | Not Found | Not Found | Not Found | Not Found | Not Found | Not Found | Not Found | Not Found | Not Found | Not Found |

| | | | | | | | | | | | | | | |
|---|---|---|---|---|---|---|---|---|---|---|---|---|---|---|
| | | | | | High | High | High | High | Not Found | Not Found | Not Found | Not Found | Not Found | Not Found |
| | | | | | High | High | High | High | Not Found | Not Found | Not Found | Not Found | Not Found | Not Found |
| | 5139 | 1759 | 8581 | 3894 | Not Found | Not Found | Not Found | Not Found | Not Found | Not Found | Not Found | Not Found | High | High |
| | | | | | High | High | High | High | Not Found | Not Found | Not Found | Not Found | Not Found | Not Found |
| | | | | | Not Found | Not Found | Not Found | Not Found | Not Found | Not Found | Not Found | Not Found | Not Found | Not Found |
| | 1355 | 1075 | 1436 | 1648 | Not Found | Not Found | Not Found | Not Found | Not Found | Not Found | Not Found | Not Found | High | High |
| | 1947 | 1327 | 2871 | 1995 | Not Found | Not Found | Not Found | Not Found | Not Found | Not Found | Not Found | Not Found | High | High |
| | 1117 | 893 | 1613 | 1538 | Not Found | Not Found | Not Found | Not Found | Not Found | Not Found | Not Found | Not Found | High | High |
| | | | | | High | High | High | High | Not Found | Not Found | Not Found | Not Found | Not Found | Not Found |
| | 1494 | 1084 | 1926 | 1779 | Not Found | Not Found | Not Found | Not Found | Not Found | Not Found | Not Found | Not Found | High | High |
| | | | | | Not Found | Not Found | Not Found | Not Found | Not Found | Not Found | Not Found | Not Found | Not Found | Not Found |
| | 1483 | 1529 | 1620 | 1814 | Not Found | Not Found | Not Found | Not Found | Not Found | Not Found | Not Found | Not Found | High | High |
| | 1383 | 817 | 1622 | 1182 | Not Found | Not Found | Not Found | Not Found | Not Found | Not Found | Not Found | Not Found | High | High |
| 1396 | | | | | Not Found | Not Found | Not Found | Not Found | High | High | High | High | Not Found | Not Found |
| | | | | | High | High | High | High | Not Found | Not Found | Not Found | Not Found | Not Found | Not Found |
| | | | | | Not Found | Not Found | Not Found | Not Found | Not Found | Not Found | Not Found | Not Found | Not Found | Not Found |
| | | | | | High | High | High | High | Not Found | Not Found | Not Found | Not Found | Not Found | Not Found |
| | | | | | Not Found | Not Found | Not Found | Not Found | Not Found | Not Found | Not Found | Not Found | High | High |
| | | | | | High | High | High | High | Not Found | Not Found | Not Found | Not Found | Not Found | Not Found |
| | | | | | High | High | High | High | Not Found | Not Found | Not Found | Not Found | Not Found | Not Found |
| 2041 | | | | | Not Found | Not Found | Not Found | Not Found | High | High | High | High | Not Found | Not Found |
| 1622 | | | | | Not Found | Not Found | Not Found | Not Found | High | High | High | High | Not Found | Not Found |
| 4990 | | | | | Not Found | Not Found | Not Found | Not Found | High | High | High | High | Not Found | Not Found |
| | | | | | High | High | High | High | Not Found | Not Found | Not Found | Not Found | Not Found | Not Found |
| | | | | | High | High | High | High | Not Found | Not Found | Not Found | Not Found | Not Found | Not Found |
| 1776 | | | | | Not Found | Not Found | Not Found | Not Found | High | High | High | High | Not Found | Not Found |
| | | | | | Not Found | Not Found | Not Found | Not Found | Not Found | Not Found | Not Found | Not Found | Not Found | Not Found |
| | | | | | High | High | High | High | Not Found | Not Found | Not Found | Not Found | Not Found | Not Found |
| 9519 | | | | | Not Found | Not Found | Not Found | Not Found | High | High | High | High | Not Found | Not Found |
| | | | | | Not Found | Not Found | Not Found | Not Found | Not Found | Not Found | Not Found | Not Found | Not Found | Not Found |
| | | | | | Not Found | Not Found | Not Found | Not Found | Not Found | Not Found | Not Found | Not Found | Not Found | Not Found |

| | | | | | | | | | | | | | | |
|---|---|---|---|---|---|---|---|---|---|---|---|---|---|---|
| 1121 | | | | | Not Found | Not Found | Not Found | Not Found | High | High | High | High | Not Found | Not Found |
| | | | | | High | High | High | High | Not Found | Not Found | Not Found | Not Found | Not Found | Not Found |
| | 362 | 383 | 1005 | 1025 | Not Found | Not Found | Not Found | Not Found | Not Found | Not Found | Not Found | Not Found | High | High |
| | 7233 | 6366 | 8442 | 9528 | Not Found | Not Found | Not Found | Not Found | Not Found | Not Found | Not Found | Not Found | High | High |
| | | | | | Not Found | Not Found | Not Found | Not Found | Not Found | Not Found | Not Found | Not Found | High | High |
| 277 | | | | | Not Found | Not Found | Not Found | Not Found | High | High | High | High | Not Found | Not Found |
| 4420 | | | | | Not Found | Not Found | Not Found | Not Found | High | High | High | High | Not Found | Not Found |
| | | | | | Not Found | Not Found | Not Found | Not Found | Not Found | Not Found | Not Found | Not Found | Not Found | Not Found |
| | | | | | Not Found | Not Found | Not Found | Not Found | Not Found | Not Found | Not Found | Not Found | Not Found | Not Found |
| | 2055 | 2071 | 3097 | 4491 | Not Found | Not Found | Not Found | Not Found | Not Found | Not Found | Not Found | Not Found | High | High |
| 1992 | | | | | Not Found | Not Found | Not Found | Not Found | High | High | High | High | Not Found | Not Found |
| | 2253 | 1944 | 3066 | 3066 | Not Found | Not Found | Not Found | Not Found | Not Found | Not Found | Not Found | Not Found | High | High |
| 459 | | | | | Not Found | Not Found | Not Found | Not Found | High | High | High | High | Not Found | Not Found |
| | 575 | 386 | 814 | 445 | Not Found | Not Found | Not Found | Not Found | Not Found | Not Found | Not Found | Not Found | High | High |
| 2624 | | | | | Not Found | Not Found | Not Found | Not Found | High | High | High | High | Not Found | Not Found |
| | | | | | High | High | High | High | Not Found | Not Found | Not Found | Not Found | Not Found | Not Found |
| | | | | | High | High | High | High | Not Found | Not Found | Not Found | Not Found | Not Found | Not Found |
| | | | | | High | High | High | High | Not Found | Not Found | Not Found | Not Found | Not Found | Not Found |
| | 433 | 360 | 465 | 384 | Not Found | Not Found | Not Found | Not Found | Not Found | Not Found | Not Found | Not Found | High | High |
| | | | | | High | High | High | High | Not Found | Not Found | Not Found | Not Found | Not Found | Not Found |
| | | | | | High | High | High | High | Not Found | Not Found | Not Found | Not Found | Not Found | Not Found |
| | | | | | High | High | High | High | Not Found | Not Found | Not Found | Not Found | Not Found | Not Found |
| | | | | | High | High | High | High | Not Found | Not Found | Not Found | Not Found | Not Found | Not Found |
| 2101 | | | | | Not Found | Not Found | Not Found | Not Found | High | High | High | High | Not Found | Not Found |
| | | | | | High | High | High | High | Not Found | Not Found | Not Found | Not Found | Not Found | Not Found |
| 1616 | | | | | Not Found | Not Found | Not Found | Not Found | High | High | High | High | Not Found | Not Found |
| | | | | | High | High | High | High | Not Found | Not Found | Not Found | Not Found | Not Found | Not Found |
| | | | | | High | High | High | High | Not Found | Not Found | Not Found | Not Found | Not Found | Not Found |
| | | | | | Not Found | Not Found | Not Found | Not Found | Not Found | Not Found | Not Found | Not Found | Not Found | Not Found |
| | | | | | Not Found | Not Found | Not Found | Not Found | Not Found | Not Found | Not Found | Not Found | Not Found | Not Found |
| | | | | | High | High | High | High | Not Found | Not Found | Not Found | Not Found | Not Found | Not Found |

| | | | | | | | | | | | | | | |
|---|---|---|---|---|---|---|---|---|---|---|---|---|---|---|
| | | | | | High | High | High | High | Not Found | Not Found | Not Found | Not Found | Not Found | Not Found |
| 1884 | | | | | Not Found | Not Found | Not Found | Not Found | High | High | High | High | Not Found | Not Found |
| | | | | | High | High | High | High | Not Found | Not Found | Not Found | Not Found | Not Found | Not Found |
| | 995 | 490 | 1092 | 771 | Not Found | Not Found | Not Found | Not Found | Not Found | Not Found | Not Found | Not Found | High | High |
| | | | | | Not Found | Not Found | Not Found | Not Found | Not Found | Not Found | Not Found | Not Found | Not Found | Not Found |
| | | | | | High | High | High | High | Not Found | Not Found | Not Found | Not Found | Not Found | Not Found |
| | 535 | 432 | 476 | 687 | Not Found | Not Found | Not Found | Not Found | Not Found | Not Found | Not Found | Not Found | High | High |
| 539 | | | | | Not Found | Not Found | Not Found | Not Found | High | High | High | High | Not Found | Not Found |
| | | | | | High | High | High | High | Not Found | Not Found | Not Found | Not Found | Not Found | Not Found |
| | | | | | Not Found | Not Found | Not Found | Not Found | Not Found | Not Found | Not Found | Not Found | Not Found | Not Found |
| | | | | | Not Found | Not Found | Not Found | Not Found | Not Found | Not Found | Not Found | Not Found | High | High |
| | | | | | Not Found | Not Found | Not Found | Not Found | Not Found | Not Found | Not Found | Not Found | Not Found | Not Found |
| | | | | | Not Found | Not Found | Not Found | Not Found | Not Found | Not Found | Not Found | Not Found | Not Found | Not Found |
| | | | | | Not Found | Not Found | Not Found | Not Found | Not Found | Not Found | Not Found | Not Found | Not Found | Not Found |
| | | | | | Not Found | Not Found | Not Found | Not Found | High | High | High | High | Not Found | Not Found |
| | 1879 | 1656 | 1911 | 2282 | Not Found | Not Found | Not Found | Not Found | Not Found | Not Found | Not Found | Not Found | High | High |
| | 4382 | 3359 | 4326 | 4210 | Not Found | Not Found | Not Found | Not Found | Not Found | Not Found | Not Found | Not Found | High | High |
| | | | | | Not Found | Not Found | Not Found | Not Found | Not Found | Not Found | Not Found | Not Found | Not Found | Not Found |
| | 1750 | 1560 | 1812 | 2003 | Not Found | Not Found | Not Found | Not Found | Not Found | Not Found | Not Found | Not Found | High | High |
| | 573 | 451 | 1224 | 707 | Not Found | Not Found | Not Found | Not Found | Not Found | Not Found | Not Found | Not Found | High | High |
| | | | | | Not Found | Not Found | Not Found | Not Found | High | High | High | High | Not Found | Not Found |
| | 3884 | 4015 | 3757 | 4370 | Not Found | Not Found | Not Found | Not Found | Not Found | Not Found | Not Found | Not Found | High | High |
| | | | | | High | High | High | High | Not Found | Not Found | Not Found | Not Found | Not Found | Not Found |
| 3233 | | | | | Not Found | Not Found | Not Found | Not Found | High | High | High | High | Not Found | Not Found |
| | 1031 | 895 | 1683 | 1688 | Not Found | Not Found | Not Found | Not Found | Not Found | Not Found | Not Found | Not Found | High | High |
| | | | | | Not Found | Not Found | Not Found | Not Found | High | High | High | High | Not Found | Not Found |
| | | | | | High | High | High | High | Not Found | Not Found | Not Found | Not Found | Not Found | Not Found |
| | 2000 | 1726 | 2977 | 2405 | Not Found | Not Found | Not Found | Not Found | Not Found | Not Found | Not Found | Not Found | High | High |
| 1554 | | | | | Not Found | Not Found | Not Found | Not Found | High | High | High | High | Not Found | Not Found |
| | | | | | High | High | High | High | Not Found | Not Found | Not Found | Not Found | Not Found | Not Found |
| | | | | | High | High | High | High | Not Found | Not Found | Not Found | Not Found | Not Found | Not Found |

| | | | | | | | | | | | | | | |
|---|---|---|---|---|---|---|---|---|---|---|---|---|---|---|
| | 6032 | 4899 | 6032 | 7234 | Not Founc | Not Founc | Not Founc | Not Founc | Not Founc | Not Founc | Not Founc | Not Founc | High | High |
| | 3730 | 2526 | 4476 | 3836 | Not Founc | Not Founc | Not Founc | Not Founc | Not Founc | Not Founc | Not Founc | Not Founc | High | High |
| | 3776 | 3084 | 4763 | 6231 | Not Founc | Not Founc | Not Founc | Not Founc | Not Founc | Not Founc | Not Founc | Not Founc | High | High |
| | | | | | High | High | High | High | Not Founc | Not Founc | Not Founc | Not Founc | Not Founc | Not Founc |
| | | | | | High | High | High | High | Not Founc | Not Founc | Not Founc | Not Founc | Not Founc | Not Founc |
| | | | | | High | High | High | High | Not Founc | Not Founc | Not Founc | Not Founc | Not Founc | Not Founc |
| | | | | | High | High | High | High | Not Founc | Not Founc | Not Founc | Not Founc | Not Founc | Not Founc |
| | 1020 | 618 | 1284 | 1174 | Not Founc | Not Founc | Not Founc | Not Founc | Not Founc | Not Founc | Not Founc | Not Founc | High | High |
| | 511 | 506 | 613 | 620 | Not Founc | Not Founc | Not Founc | Not Founc | Not Founc | Not Founc | Not Founc | Not Founc | High | High |
| | | | | | High | High | High | High | Not Founc | Not Founc | Not Founc | Not Founc | Not Founc | Not Founc |
| | 1501 | 1146 | 1666 | 1683 | Not Founc | Not Founc | Not Founc | Not Founc | Not Founc | Not Founc | Not Founc | Not Founc | High | High |
| | | | | | High | High | High | High | Not Founc | Not Founc | Not Founc | Not Founc | Not Founc | Not Founc |
| | | | | | High | High | High | High | Not Founc | Not Founc | Not Founc | Not Founc | Not Founc | Not Founc |
| | | | | | High | High | High | High | Not Founc | Not Founc | Not Founc | Not Founc | Not Founc | Not Founc |
| | 1232 | 878 | 1338 | 1275 | Not Founc | Not Founc | Not Founc | Not Founc | Not Founc | Not Founc | Not Founc | Not Founc | High | High |
| | | | | | High | High | High | High | Not Founc | Not Founc | Not Founc | Not Founc | Not Founc | Not Founc |
| | 1384 | 1085 | 1587 | 1467 | Not Founc | Not Founc | Not Founc | Not Founc | Not Founc | Not Founc | Not Founc | Not Founc | High | High |
| 887 | | | | | Not Founc | Not Founc | Not Founc | Not Founc | High | High | High | High | Not Founc | Not Founc |
| | | | | | Not Founc | Not Founc | Not Founc | Not Founc | High | High | High | High | Not Founc | Not Founc |
| | | | | | High | High | High | High | Not Founc | Not Founc | Not Founc | Not Founc | Not Founc | Not Founc |
| 3542 | | | | | Not Founc | Not Founc | Not Founc | Not Founc | High | High | High | High | Not Founc | Not Founc |
| | 893 | 670 | 1089 | 1138 | Not Founc | Not Founc | Not Founc | Not Founc | Not Founc | Not Founc | Not Founc | Not Founc | High | High |
| | 4092 | 1982 | 4752 | 3490 | Not Founc | Not Founc | Not Founc | Not Founc | Not Founc | Not Founc | Not Founc | Not Founc | High | High |
| 751 | | | | | Not Founc | Not Founc | Not Founc | Not Founc | High | High | High | High | Not Founc | Not Founc |
| | | | | | High | High | High | High | Not Founc | Not Founc | Not Founc | Not Founc | Not Founc | Not Founc |
| | 3905 | 2939 | 5586 | 4378 | Not Founc | Not Founc | Not Founc | Not Founc | Not Founc | Not Founc | Not Founc | Not Founc | High | High |
| | 2751 | 2424 | 2505 | 3638 | Not Founc | Not Founc | Not Founc | Not Founc | Not Founc | Not Founc | Not Founc | Not Founc | High | High |
| | 679 | 393 | 1539 | 1002 | Not Founc | Not Founc | Not Founc | Not Founc | Not Founc | Not Founc | Not Founc | Not Founc | High | High |
| | | | | | High | High | High | High | Not Founc | Not Founc | Not Founc | Not Founc | Not Founc | Not Founc |
| | 655 | 435 | 778 | 744 | Not Founc | Not Founc | Not Founc | Not Founc | Not Founc | Not Founc | Not Founc | Not Founc | High | High |
| | | | | | High | High | High | High | Not Founc | Not Founc | Not Founc | Not Founc | Not Founc | Not Founc |

| | | | | | | | | | | | | | | |
|---|---|---|---|---|---|---|---|---|---|---|---|---|---|---|
| | 538 | 451 | 366 | 586 | Not Founc | Not Founc | Not Founc | Not Founc | Not Founc | Not Founc | Not Founc | Not Founc | High | High |
| 2522 | | | | | Not Founc | Not Founc | Not Founc | Not Founc | High | High | High | High | Not Founc | Not Founc |
| 2010 | | | | | Not Founc | Not Founc | Not Founc | Not Founc | High | High | High | High | Not Founc | Not Founc |
| | | | | | High | High | High | High | Not Founc | Not Founc | Not Founc | Not Founc | Not Founc | Not Founc |
| | | | | | High | High | High | High | Not Founc | Not Founc | Not Founc | Not Founc | Not Founc | Not Founc |
| | | | | | High | High | High | High | Not Founc | Not Founc | Not Founc | Not Founc | Not Founc | Not Founc |
| | 1183 | 871 | 1852 | 1357 | Not Founc | Not Founc | Not Founc | Not Founc | Not Founc | Not Founc | Not Founc | Not Founc | High | High |
| | 3300 | 2105 | 4315 | 3548 | Not Founc | Not Founc | Not Founc | Not Founc | Not Founc | Not Founc | Not Founc | Not Founc | High | High |
| | 1513 | 434 | 1885 | 675 | Not Founc | Not Founc | Not Founc | Not Founc | Not Founc | Not Founc | Not Founc | Not Founc | High | High |
| 501 | | | | | Not Founc | Not Founc | Not Founc | Not Founc | High | High | High | High | Not Founc | Not Founc |
| | | | | | High | High | High | High | Not Founc | Not Founc | Not Founc | Not Founc | Not Founc | Not Founc |
| 1795 | | | | | Not Founc | Not Founc | Not Founc | Not Founc | High | High | High | High | Not Founc | Not Founc |
| 507 | | | | | Not Founc | Not Founc | Not Founc | Not Founc | High | High | High | High | Not Founc | Not Founc |
| 3498 | | | | | Not Founc | Not Founc | Not Founc | Not Founc | High | High | High | High | Not Founc | Not Founc |
| | 1699 | 1238 | 1478 | 1537 | Not Founc | Not Founc | Not Founc | Not Founc | Not Founc | Not Founc | Not Founc | Not Founc | High | High |
| | | | | | High | High | High | High | Not Founc | Not Founc | Not Founc | Not Founc | Not Founc | Not Founc |
| 746 | | | | | Not Founc | Not Founc | Not Founc | Not Founc | High | High | High | High | Not Founc | Not Founc |
| 713 | | | | | Not Founc | Not Founc | Not Founc | Not Founc | High | High | High | High | Not Founc | Not Founc |
| | | | | | High | High | High | High | Not Founc | Not Founc | Not Founc | Not Founc | Not Founc | Not Founc |
| | | | | | High | High | High | High | Not Founc | Not Founc | Not Founc | Not Founc | Not Founc | Not Founc |
| | | | | | High | High | High | High | Not Founc | Not Founc | Not Founc | Not Founc | Not Founc | Not Founc |
| | | | | | High | High | High | High | Not Founc | Not Founc | Not Founc | Not Founc | Not Founc | Not Founc |
| | 2419 | 1893 | 3213 | 3103 | Not Founc | Not Founc | Not Founc | Not Founc | Not Founc | Not Founc | Not Founc | Not Founc | High | High |
| | | | | | High | High | High | High | Not Founc | Not Founc | Not Founc | Not Founc | Not Founc | Not Founc |
| | | | | | High | High | High | High | Not Founc | Not Founc | Not Founc | Not Founc | Not Founc | Not Founc |
| | | | | | High | High | High | High | Not Founc | Not Founc | Not Founc | Not Founc | Not Founc | Not Founc |
| | 4435 | 1710 | 5904 | 2814 | Not Founc | Not Founc | Not Founc | Not Founc | Not Founc | Not Founc | Not Founc | Not Founc | High | High |
| | 3097 | 2293 | 3052 | 3463 | Not Founc | Not Founc | Not Founc | Not Founc | Not Founc | Not Founc | Not Founc | Not Founc | High | High |
| | 3210 | 2165 | 3634 | 3301 | Not Founc | Not Founc | Not Founc | Not Founc | Not Founc | Not Founc | Not Founc | Not Founc | High | High |
| | 906 | 850 | 2413 | 2013 | Not Founc | Not Founc | Not Founc | Not Founc | Not Founc | Not Founc | Not Founc | Not Founc | High | High |
| 2134 | | | | | Not Founc | Not Founc | Not Founc | Not Founc | High | High | High | High | Not Founc | Not Founc |

| | | | | | | | | | | | | | | |
|---|---|---|---|---|---|---|---|---|---|---|---|---|---|---|
| | | | | | High | High | High | High | Not Found | Not Found | Not Found | Not Found | Not Found | Not Found |
| 6389 | | | | | Not Found | Not Found | Not Found | Not Found | High | High | High | High | Not Found | Not Found |
| 5062 | | | | | Not Found | Not Found | Not Found | Not Found | High | High | High | High | Not Found | Not Found |
| | 417 | 251 | 438 | 387 | Not Found | Not Found | Not Found | Not Found | Not Found | Not Found | Not Found | Not Found | High | High |
| | 3277 | 2309 | 3527 | 3495 | Not Found | Not Found | Not Found | Not Found | Not Found | Not Found | Not Found | Not Found | High | High |
| | | | | | High | High | High | High | Not Found | Not Found | Not Found | Not Found | Not Found | Not Found |
| 4026 | | | | | Not Found | Not Found | Not Found | Not Found | High | High | High | High | Not Found | Not Found |
| 3624 | | | | | Not Found | Not Found | Not Found | Not Found | High | High | High | High | Not Found | Not Found |
| | 620 | 506 | 1106 | 875 | Not Found | Not Found | Not Found | Not Found | Not Found | Not Found | Not Found | Not Found | High | High |
| | | | | | High | High | High | High | Not Found | Not Found | Not Found | Not Found | Not Found | Not Found |
| | | | | | High | High | High | High | Not Found | Not Found | Not Found | Not Found | Not Found | Not Found |
| | | | | | High | High | High | High | Not Found | Not Found | Not Found | Not Found | Not Found | Not Found |
| 3886 | | | | | Not Found | Not Found | Not Found | Not Found | High | High | High | High | Not Found | Not Found |
| | | | | | High | High | High | High | Not Found | Not Found | Not Found | Not Found | Not Found | Not Found |
| | 4847 | 3349 | 5629 | 4918 | Not Found | Not Found | Not Found | Not Found | Not Found | Not Found | Not Found | Not Found | High | High |
| | | | | | High | High | High | High | Not Found | Not Found | Not Found | Not Found | Not Found | Not Found |
| 4412 | | | | | Not Found | Not Found | Not Found | Not Found | High | High | High | High | Not Found | Not Found |
| | 1374 | 799 | 2135 | 1937 | Not Found | Not Found | Not Found | Not Found | Not Found | Not Found | Not Found | Not Found | High | High |
| 5018 | | | | | Not Found | Not Found | Not Found | Not Found | High | High | High | High | Not Found | Not Found |
| | | | | | High | High | High | High | Not Found | Not Found | Not Found | Not Found | Not Found | Not Found |
| 329 | | | | | Not Found | Not Found | Not Found | Not Found | High | High | High | High | Not Found | Not Found |
| | 2212 | 961 | 2450 | 1294 | Not Found | Not Found | Not Found | Not Found | Not Found | Not Found | Not Found | Not Found | High | High |
| | 3061 | 1551 | 3565 | 2513 | Not Found | Not Found | Not Found | Not Found | Not Found | Not Found | Not Found | Not Found | High | High |
| 1814 | | | | | Not Found | Not Found | Not Found | Not Found | High | High | High | High | Not Found | Not Found |
| | 1501 | 971 | 2381 | 3526 | Not Found | Not Found | Not Found | Not Found | Not Found | Not Found | Not Found | Not Found | High | High |
| | | | | | High | High | High | High | Not Found | Not Found | Not Found | Not Found | Not Found | Not Found |
| | | | | | High | High | High | High | Not Found | Not Found | Not Found | Not Found | Not Found | Not Found |
| 2052 | | | | | Not Found | Not Found | Not Found | Not Found | High | High | High | High | Not Found | Not Found |
| | | | | | High | High | High | High | Not Found | Not Found | Not Found | Not Found | Not Found | Not Found |
| | | | | | High | High | High | High | Not Found | Not Found | Not Found | Not Found | Not Found | Not Found |
| 2372 | | | | | Not Found | Not Found | Not Found | Not Found | High | High | High | High | Not Found | Not Found |

| | | | | | | | | | | | | | | |
|---|---|---|---|---|---|---|---|---|---|---|---|---|---|---|
| | 2598 | 2018 | 2557 | 2736 | Not Founc | Not Founc | Not Founc | Not Founc | Not Founc | Not Founc | Not Founc | Not Founc | High | High |
| | | | | | Not Founc | Not Founc | Not Founc | Not Founc | Not Founc | Not Founc | Not Founc | Not Founc | Not Founc | Not Founc |
| | | | | | High | High | High | High | Not Founc | Not Founc | Not Founc | Not Founc | Not Founc | Not Founc |
| 1462 | | | | | Not Founc | Not Founc | Not Founc | Not Founc | High | High | High | High | Not Founc | Not Founc |
| | | | | | High | High | High | High | Not Founc | Not Founc | Not Founc | Not Founc | Not Founc | Not Founc |
| | | | | | High | High | High | High | Not Founc | Not Founc | Not Founc | Not Founc | Not Founc | Not Founc |
| | | | | | High | High | High | High | Not Founc | Not Founc | Not Founc | Not Founc | Not Founc | Not Founc |
| | 3201 | 3334 | 2780 | 3182 | Not Founc | Not Founc | Not Founc | Not Founc | Not Founc | Not Founc | Not Founc | Not Founc | High | High |
| 1671 | | | | | Not Founc | Not Founc | Not Founc | Not Founc | High | High | High | High | Not Founc | Not Founc |
| 1313 | | | | | Not Founc | Not Founc | Not Founc | Not Founc | High | High | High | High | Not Founc | Not Founc |
| 2737 | | | | | Not Founc | Not Founc | Not Founc | Not Founc | High | High | High | High | Not Founc | Not Founc |
| | | | | | High | High | High | High | Not Founc | Not Founc | Not Founc | Not Founc | Not Founc | Not Founc |
| | 1418 | 1106 | 1905 | 1536 | Not Founc | Not Founc | Not Founc | Not Founc | Not Founc | Not Founc | Not Founc | Not Founc | High | High |
| | | | | | High | High | High | High | Not Founc | Not Founc | Not Founc | Not Founc | Not Founc | Not Founc |
| | 1908 | 1402 | 2346 | 2239 | Not Founc | Not Founc | Not Founc | Not Founc | Not Founc | Not Founc | Not Founc | Not Founc | High | High |
| | | | | | High | High | High | High | Not Founc | Not Founc | Not Founc | Not Founc | Not Founc | Not Founc |
| 2366 | | | | | Not Founc | Not Founc | Not Founc | Not Founc | High | High | High | High | Not Founc | Not Founc |
| 6348 | | | | | Not Founc | Not Founc | Not Founc | Not Founc | High | High | High | High | Not Founc | Not Founc |
| | 2180 | 1388 | 2534 | 2318 | Not Founc | Not Founc | Not Founc | Not Founc | Not Founc | Not Founc | Not Founc | Not Founc | High | High |
| 4193 | | | | | Not Founc | Not Founc | Not Founc | Not Founc | High | High | High | High | Not Founc | Not Founc |
| | 1798 | 989 | 2542 | 1733 | Not Founc | Not Founc | Not Founc | Not Founc | Not Founc | Not Founc | Not Founc | Not Founc | High | High |
| 276 | | | | | Not Founc | Not Founc | Not Founc | Not Founc | High | High | High | High | Not Founc | Not Founc |
| | 1353 | 931 | 1947 | 1426 | Not Founc | Not Founc | Not Founc | Not Founc | Not Founc | Not Founc | Not Founc | Not Founc | High | High |
| | 3532 | 971 | 7090 | 2027 | Not Founc | Not Founc | Not Founc | Not Founc | Not Founc | Not Founc | Not Founc | Not Founc | High | High |
| 1114 | | | | | Not Founc | Not Founc | Not Founc | Not Founc | High | High | High | High | Not Founc | Not Founc |
| | 6933 | 6365 | 7870 | 10020 | Not Founc | Not Founc | Not Founc | Not Founc | Not Founc | Not Founc | Not Founc | Not Founc | High | High |
| | 1578 | 946 | 4355 | 2386 | Not Founc | Not Founc | Not Founc | Not Founc | Not Founc | Not Founc | Not Founc | Not Founc | High | High |
| | | | | | High | High | High | High | Not Founc | Not Founc | Not Founc | Not Founc | Not Founc | Not Founc |
| | 3089 | 2274 | 3389 | 3294 | Not Founc | Not Founc | Not Founc | Not Founc | Not Founc | Not Founc | Not Founc | Not Founc | High | High |
| 4937 | | | | | Not Founc | Not Founc | Not Founc | Not Founc | High | High | High | High | Not Founc | Not Founc |
| 4426 | | | | | Not Founc | Not Founc | Not Founc | Not Founc | High | High | High | High | Not Founc | Not Founc |

| | | | | | | | | | | | | | | |
|---|---|---|---|---|---|---|---|---|---|---|---|---|---|---|
| | 3094 | 2317 | 4813 | 3801 | Not Founc | Not Founc | Not Founc | Not Founc | Not Founc | Not Founc | Not Founc | Not Founc | High | High |
| | 817 | 828 | 726 | 1345 | Not Founc | Not Founc | Not Founc | Not Founc | Not Founc | Not Founc | Not Founc | Not Founc | High | High |
| | | | | | High | High | High | High | Not Founc | Not Founc | Not Founc | Not Founc | Not Founc | Not Founc |
| 1137 | | | | | Not Founc | Not Founc | Not Founc | Not Founc | High | High | High | High | Not Founc | Not Founc |
| | | | | | High | High | High | High | Not Founc | Not Founc | Not Founc | Not Founc | Not Founc | Not Founc |
| | | | | | Not Founc | Not Founc | Not Founc | Not Founc | High | High | High | High | Not Founc | Not Founc |
| | 2110 | 1263 | 2535 | 1931 | Not Founc | Not Founc | Not Founc | Not Founc | Not Founc | Not Founc | Not Founc | Not Founc | High | High |
| | | | | | High | High | High | High | Not Founc | Not Founc | Not Founc | Not Founc | Not Founc | Not Founc |
| | | | | | High | High | High | High | Not Founc | Not Founc | Not Founc | Not Founc | Not Founc | Not Founc |
| 4581 | | | | | Not Founc | Not Founc | Not Founc | Not Founc | High | High | High | High | Not Founc | Not Founc |
| | | | | | High | High | High | High | Not Founc | Not Founc | Not Founc | Not Founc | Not Founc | Not Founc |
| 3338 | | | | | Not Founc | Not Founc | Not Founc | Not Founc | High | High | High | High | Not Founc | Not Founc |
| | 3289 | 2331 | 3826 | 3794 | Not Founc | Not Founc | Not Founc | Not Founc | Not Founc | Not Founc | Not Founc | Not Founc | High | High |
| | | | | | High | High | High | High | Not Founc | Not Founc | Not Founc | Not Founc | Not Founc | Not Founc |
| | 3024 | 2837 | 2631 | 1735 | Not Founc | Not Founc | Not Founc | Not Founc | Not Founc | Not Founc | Not Founc | Not Founc | High | High |
| | | | | | High | High | High | High | Not Founc | Not Founc | Not Founc | Not Founc | Not Founc | Not Founc |
| 3243 | | | | | Not Founc | Not Founc | Not Founc | Not Founc | High | High | High | High | Not Founc | Not Founc |
| | 1439 | 948 | 1924 | 1485 | Not Founc | Not Founc | Not Founc | Not Founc | Not Founc | Not Founc | Not Founc | Not Founc | High | High |
| | | | | | High | High | High | High | Not Founc | Not Founc | Not Founc | Not Founc | Not Founc | Not Founc |
| | | | | | High | High | High | High | Not Founc | Not Founc | Not Founc | Not Founc | Not Founc | Not Founc |
| | | | | | High | High | High | High | Not Founc | Not Founc | Not Founc | Not Founc | Not Founc | Not Founc |
| | 1455 | 915 | 3340 | 2133 | Not Founc | Not Founc | Not Founc | Not Founc | Not Founc | Not Founc | Not Founc | Not Founc | High | High |
| | 5282 | 3841 | 6558 | 5416 | Not Founc | Not Founc | Not Founc | Not Founc | Not Founc | Not Founc | Not Founc | Not Founc | High | High |
| | 2471 | 1958 | 3106 | 2553 | Not Founc | Not Founc | Not Founc | Not Founc | Not Founc | Not Founc | Not Founc | Not Founc | High | High |
| | | | | | Not Founc | Not Founc | Not Founc | Not Founc | Not Founc | Not Founc | Not Founc | Not Founc | Not Founc | Not Founc |
| | | | | | Not Founc | Not Founc | Not Founc | Not Founc | Not Founc | Not Founc | Not Founc | Not Founc | High | High |
| | | | | | High | High | High | High | Not Founc | Not Founc | Not Founc | Not Founc | Not Founc | Not Founc |
| | | | | | High | High | High | High | Not Founc | Not Founc | Not Founc | Not Founc | Not Founc | Not Founc |
| | 587 | 462 | 865 | 762 | Not Founc | Not Founc | Not Founc | Not Founc | Not Founc | Not Founc | Not Founc | Not Founc | High | High |
| 5465 | | | | | Not Founc | Not Founc | Not Founc | Not Founc | High | High | High | High | Not Founc | Not Founc |
| | 5449 | 4888 | 5517 | 6242 | Not Founc | Not Founc | Not Founc | Not Founc | Not Founc | Not Founc | Not Founc | Not Founc | High | High |

| | | | | | | | | | | | | | | |
|---|---|---|---|---|---|---|---|---|---|---|---|---|---|---|
| | | | | | High | High | High | High | Not Found | Not Found | Not Found | Not Found | Not Found | Not Found |
| | | | | | High | High | High | High | Not Found | Not Found | Not Found | Not Found | Not Found | Not Found |
| 1271 | | | | | Not Found | Not Found | Not Found | Not Found | High | High | High | High | Not Found | Not Found |
| | | | | | High | High | High | High | Not Found | Not Found | Not Found | Not Found | Not Found | Not Found |
| | 3256 | 3113 | 4422 | 5809 | Not Found | Not Found | Not Found | Not Found | Not Found | Not Found | Not Found | Not Found | High | High |
| 5453 | | | | | Not Found | Not Found | Not Found | Not Found | High | High | High | High | Not Found | Not Found |
| 2511 | | | | | Not Found | Not Found | Not Found | Not Found | High | High | High | High | Not Found | Not Found |
| | 1096 | 811 | 1239 | 1447 | Not Found | Not Found | Not Found | Not Found | Not Found | Not Found | Not Found | Not Found | High | High |
| 7079 | | | | | Not Found | Not Found | Not Found | Not Found | High | High | High | High | Not Found | Not Found |
| | | | | | High | High | High | High | Not Found | Not Found | Not Found | Not Found | Not Found | Not Found |
| 2511 | | | | | Not Found | Not Found | Not Found | Not Found | High | High | High | High | Not Found | Not Found |
| 3846 | | | | | Not Found | Not Found | Not Found | Not Found | High | High | High | High | Not Found | Not Found |
| | 2890 | 1913 | 4830 | 4178 | Not Found | Not Found | Not Found | Not Found | Not Found | Not Found | Not Found | Not Found | High | High |
| | 2191 | 3697 | 2412 | 5243 | Not Found | Not Found | Not Found | Not Found | Not Found | Not Found | Not Found | Not Found | High | High |
| | 3362 | 2509 | 3780 | 3709 | Not Found | Not Found | Not Found | Not Found | Not Found | Not Found | Not Found | Not Found | High | High |
| | | | | | High | High | High | High | Not Found | Not Found | Not Found | Not Found | Not Found | Not Found |
| | 5054 | 2676 | 3762 | 3401 | Not Found | Not Found | Not Found | Not Found | Not Found | Not Found | Not Found | Not Found | High | High |
| | | | | | High | High | High | High | Not Found | Not Found | Not Found | Not Found | Not Found | Not Found |
| 3107 | | | | | Not Found | Not Found | Not Found | Not Found | High | High | High | High | Not Found | Not Found |
| | 2354 | 1485 | 2469 | 2270 | Not Found | Not Found | Not Found | Not Found | Not Found | Not Found | Not Found | Not Found | High | High |
| | 4154 | 1925 | 9377 | 4719 | Not Found | Not Found | Not Found | Not Found | Not Found | Not Found | Not Found | Not Found | High | High |
| | 3017 | 2043 | 6563 | 4937 | Not Found | Not Found | Not Found | Not Found | Not Found | Not Found | Not Found | Not Found | High | High |
| | | | | | High | High | High | High | Not Found | Not Found | Not Found | Not Found | Not Found | Not Found |
| | | | | | Not Found | Not Found | Not Found | Not Found | Not Found | Not Found | Not Found | Not Found | High | High |
| 5466 | | | | | Not Found | Not Found | Not Found | Not Found | High | High | High | High | Not Found | Not Found |
| 1535 | | | | | Not Found | Not Found | Not Found | Not Found | High | High | High | High | Not Found | Not Found |
| | 7170 | 5152 | 9143 | 7458 | Not Found | Not Found | Not Found | Not Found | Not Found | Not Found | Not Found | Not Found | High | High |
| | | | | | High | High | High | High | Not Found | Not Found | Not Found | Not Found | Not Found | Not Found |
| 1144 | | | | | Not Found | Not Found | Not Found | Not Found | High | High | High | High | Not Found | Not Found |
| | 1014 | 653 | 1514 | 1000 | Not Found | Not Found | Not Found | Not Found | Not Found | Not Found | Not Found | Not Found | High | High |
| | | | | | High | High | High | High | Not Found | Not Found | Not Found | Not Found | Not Found | Not Found |

| | | | | | | | | | | | | | | |
|---|---|---|---|---|---|---|---|---|---|---|---|---|---|---|
| | 699 | 466 | 866 | 589 | Not Founc | Not Founc | Not Founc | Not Founc | Not Founc | Not Founc | Not Founc | Not Founc | High | High |
| | | | | | High | High | High | High | Not Founc | Not Founc | Not Founc | Not Founc | Not Founc | Not Founc |
| | | | | | High | High | High | High | Not Founc | Not Founc | Not Founc | Not Founc | Not Founc | Not Founc |
| | 5278 | 823 | 8522 | 1769 | Not Founc | Not Founc | Not Founc | Not Founc | Not Founc | Not Founc | Not Founc | Not Founc | High | High |
| | | | | | High | High | High | High | Not Founc | Not Founc | Not Founc | Not Founc | Not Founc | Not Founc |
| | | | | | Not Founc | Not Founc | Not Founc | Not Founc | Not Founc | Not Founc | Not Founc | Not Founc | Not Founc | Not Founc |
| | 2261 | 1933 | 2288 | 2481 | Not Founc | Not Founc | Not Founc | Not Founc | Not Founc | Not Founc | Not Founc | Not Founc | High | High |
| | | | | | Not Founc | Not Founc | Not Founc | Not Founc | Not Founc | Not Founc | Not Founc | Not Founc | High | High |
| | | | | | High | High | High | High | Not Founc | Not Founc | Not Founc | Not Founc | Not Founc | Not Founc |
| 2348 | | | | | Not Founc | Not Founc | Not Founc | Not Founc | High | High | High | High | Not Founc | Not Founc |
| 4018 | | | | | Not Founc | Not Founc | Not Founc | Not Founc | High | High | High | High | Not Founc | Not Founc |
| | | | | | High | High | High | High | Not Founc | Not Founc | Not Founc | Not Founc | Not Founc | Not Founc |
| | | | | | High | High | High | High | Not Founc | Not Founc | Not Founc | Not Founc | Not Founc | Not Founc |
| 3230 | | | | | Not Founc | Not Founc | Not Founc | Not Founc | High | High | High | High | Not Founc | Not Founc |
| | 1609 | 1413 | 2024 | 1918 | Not Founc | Not Founc | Not Founc | Not Founc | Not Founc | Not Founc | Not Founc | Not Founc | High | High |
| 5028 | | | | | Not Founc | Not Founc | Not Founc | Not Founc | High | High | High | High | Not Founc | Not Founc |
| | 6468 | 3432 | 12186 | 6597 | Not Founc | Not Founc | Not Founc | Not Founc | Not Founc | Not Founc | Not Founc | Not Founc | High | High |
| | 5061 | 3849 | 7267 | 6671 | Not Founc | Not Founc | Not Founc | Not Founc | Not Founc | Not Founc | Not Founc | Not Founc | High | High |
| | 3392 | 2122 | 3486 | 3497 | Not Founc | Not Founc | Not Founc | Not Founc | Not Founc | Not Founc | Not Founc | Not Founc | High | High |
| 6089 | | | | | Not Founc | Not Founc | Not Founc | Not Founc | High | High | High | High | Not Founc | Not Founc |
| | | | | | High | High | High | High | Not Founc | Not Founc | Not Founc | Not Founc | Not Founc | Not Founc |
| | 2377 | 1719 | 2953 | 2041 | Not Founc | Not Founc | Not Founc | Not Founc | Not Founc | Not Founc | Not Founc | Not Founc | High | High |
| | 6474 | 4020 | 8902 | 6763 | Not Founc | Not Founc | Not Founc | Not Founc | Not Founc | Not Founc | Not Founc | Not Founc | High | High |
| | | | | | Not Founc | Not Founc | Not Founc | Not Founc | Not Founc | Not Founc | Not Founc | Not Founc | Not Founc | Not Founc |
| | | | | | High | High | High | High | Not Founc | Not Founc | Not Founc | Not Founc | Not Founc | Not Founc |
| | | | | | High | High | High | High | Not Founc | Not Founc | Not Founc | Not Founc | Not Founc | Not Founc |
| 2787 | | | | | Not Founc | Not Founc | Not Founc | Not Founc | High | High | High | High | Not Founc | Not Founc |
| 1951 | | | | | Not Founc | Not Founc | Not Founc | Not Founc | High | High | High | High | Not Founc | Not Founc |
| | | | | | High | High | High | High | Not Founc | Not Founc | Not Founc | Not Founc | Not Founc | Not Founc |
| | | | | | High | High | High | High | Not Founc | Not Founc | Not Founc | Not Founc | Not Founc | Not Founc |
| 6791 | | | | | Not Founc | Not Founc | Not Founc | Not Founc | High | High | High | High | Not Founc | Not Founc |

| | | | | | | | | | | | | | | |
|---|---|---|---|---|---|---|---|---|---|---|---|---|---|---|
| | | | | | High | High | High | High | Not Founc | Not Founc | Not Founc | Not Founc | Not Founc | Not Founc |
| | 1930 | 1476 | 2787 | 2283 | Not Founc | Not Founc | Not Founc | Not Founc | Not Founc | Not Founc | Not Founc | Not Founc | High | High |
| 2709 | | | | | Not Founc | Not Founc | Not Founc | Not Founc | High | High | High | High | Not Founc | Not Founc |
| | | | | | Not Founc | Not Founc | Not Founc | Not Founc | Not Founc | Not Founc | Not Founc | Not Founc | High | High |
| | | | | | High | High | High | High | Not Founc | Not Founc | Not Founc | Not Founc | Not Founc | Not Founc |
| 2869 | | | | | Not Founc | Not Founc | Not Founc | Not Founc | High | High | High | High | Not Founc | Not Founc |
| 3626 | | | | | Not Founc | Not Founc | Not Founc | Not Founc | High | High | High | High | Not Founc | Not Founc |
| | 1484 | 981 | 2957 | 1941 | Not Founc | Not Founc | Not Founc | Not Founc | Not Founc | Not Founc | Not Founc | Not Founc | High | High |
| | 1842 | 1821 | 2155 | 2755 | Not Founc | Not Founc | Not Founc | Not Founc | Not Founc | Not Founc | Not Founc | Not Founc | High | High |
| 986 | | | | | Not Founc | Not Founc | Not Founc | Not Founc | High | High | High | High | Not Founc | Not Founc |
| | | | | | Not Founc | Not Founc | Not Founc | Not Founc | Not Founc | Not Founc | Not Founc | Not Founc | High | High |
| 2110 | | | | | Not Founc | Not Founc | Not Founc | Not Founc | High | High | High | High | Not Founc | Not Founc |
| | | | | | High | High | High | High | Not Founc | Not Founc | Not Founc | Not Founc | Not Founc | Not Founc |
| | 4256 | 1485 | 8774 | 3445 | Not Founc | Not Founc | Not Founc | Not Founc | Not Founc | Not Founc | Not Founc | Not Founc | High | High |
| 4316 | | | | | Not Founc | Not Founc | Not Founc | Not Founc | High | High | High | High | Not Founc | Not Founc |
| 4609 | | | | | Not Founc | Not Founc | Not Founc | Not Founc | High | High | High | High | Not Founc | Not Founc |
| 3509 | | | | | Not Founc | Not Founc | Not Founc | Not Founc | High | High | High | High | Not Founc | Not Founc |
| | | | | | High | High | High | High | Not Founc | Not Founc | Not Founc | Not Founc | Not Founc | Not Founc |
| | | | | | High | High | High | High | Not Founc | Not Founc | Not Founc | Not Founc | Not Founc | Not Founc |
| | 1667 | 1432 | 4161 | 3734 | Not Founc | Not Founc | Not Founc | Not Founc | Not Founc | Not Founc | Not Founc | Not Founc | High | High |
| | | | | | Not Founc | Not Founc | Not Founc | Not Founc | High | High | High | High | Not Founc | Not Founc |
| | | | | | High | High | High | High | Not Founc | Not Founc | Not Founc | Not Founc | Not Founc | Not Founc |
| | 705 | 557 | 1051 | 876 | Not Founc | Not Founc | Not Founc | Not Founc | Not Founc | Not Founc | Not Founc | Not Founc | High | High |
| 4542 | | | | | Not Founc | Not Founc | Not Founc | Not Founc | High | High | High | High | Not Founc | Not Founc |
| | | | | | Not Founc | Not Founc | Not Founc | Not Founc | Not Founc | Not Founc | Not Founc | Not Founc | High | High |
| | | | | | Not Founc | Not Founc | Not Founc | Not Founc | Not Founc | Not Founc | Not Founc | Not Founc | Not Founc | Not Founc |
| | | | | | High | High | High | High | Not Founc | Not Founc | Not Founc | Not Founc | Not Founc | Not Founc |
| 2126 | | | | | Not Founc | Not Founc | Not Founc | Not Founc | High | High | High | High | Not Founc | Not Founc |
| | 1809 | 1072 | 2105 | 1355 | Not Founc | Not Founc | Not Founc | Not Founc | Not Founc | Not Founc | Not Founc | Not Founc | High | High |
| | 1959 | 930 | 2742 | 1728 | Not Founc | Not Founc | Not Founc | Not Founc | Not Founc | Not Founc | Not Founc | Not Founc | High | High |
| | | | | | High | High | High | High | Not Founc | Not Founc | Not Founc | Not Founc | Not Founc | Not Founc |

| | | | | | | | | | | | | | | |
|---|---|---|---|---|---|---|---|---|---|---|---|---|---|---|
| | 1277 | 835 | 1256 | 1367 | Not Founc | Not Founc | Not Founc | Not Founc | Not Founc | Not Founc | Not Founc | Not Founc | High | High |
| | | | | | High | High | High | High | Not Founc | Not Founc | Not Founc | Not Founc | Not Founc | Not Founc |
| | | | | | Not Founc | Not Founc | Not Founc | Not Founc | High | High | High | High | Not Founc | Not Founc |
| | | | | | Not Founc | Not Founc | Not Founc | Not Founc | High | High | High | High | Not Founc | Not Founc |
| 5684 | | | | | Not Founc | Not Founc | Not Founc | Not Founc | High | High | High | High | Not Founc | Not Founc |
| | | | | | Not Founc | Not Founc | Not Founc | Not Founc | Not Founc | Not Founc | Not Founc | Not Founc | Not Founc | Not Founc |
| | 4779 | 2969 | 5469 | 4548 | Not Founc | Not Founc | Not Founc | Not Founc | Not Founc | Not Founc | Not Founc | Not Founc | High | High |
| | | | | | High | High | High | High | Not Founc | Not Founc | Not Founc | Not Founc | Not Founc | Not Founc |
| 3382 | | | | | Not Founc | Not Founc | Not Founc | Not Founc | High | High | High | High | Not Founc | Not Founc |
| | | | | | High | High | High | High | Not Founc | Not Founc | Not Founc | Not Founc | Not Founc | Not Founc |
| | 20 | | 37 | 19 | Not Founc | Not Founc | Not Founc | Not Founc | Not Founc | Not Founc | Not Founc | Not Founc | High | Not Founc |
| | 3560 | 3016 | 5795 | 6199 | Not Founc | Not Founc | Not Founc | Not Founc | Not Founc | Not Founc | Not Founc | Not Founc | High | High |
| | | | | | Not Founc | Not Founc | Not Founc | Not Founc | Not Founc | Not Founc | Not Founc | Not Founc | Not Founc | Not Founc |
| | | | | | High | High | High | High | Not Founc | Not Founc | Not Founc | Not Founc | Not Founc | Not Founc |
| | 1326 | 718 | 3141 | 1746 | Not Founc | Not Founc | Not Founc | Not Founc | Not Founc | Not Founc | Not Founc | Not Founc | High | High |
| | | | | | Not Founc | Not Founc | Not Founc | Not Founc | Not Founc | Not Founc | Not Founc | Not Founc | Not Founc | Not Founc |
| 3234 | | | | | Not Founc | Not Founc | Not Founc | Not Founc | High | High | High | High | Not Founc | Not Founc |
| 2247 | | | | | Not Founc | Not Founc | Not Founc | Not Founc | High | High | High | High | Not Founc | Not Founc |
| 9160 | | | | | Not Founc | Not Founc | Not Founc | Not Founc | High | High | High | High | Not Founc | Not Founc |
| 1898 | | | | | Not Founc | Not Founc | Not Founc | Not Founc | High | High | High | High | Not Founc | Not Founc |
| | | | | | High | High | High | High | Not Founc | Not Founc | Not Founc | Not Founc | Not Founc | Not Founc |
| | | | | | Not Founc | Not Founc | Not Founc | Not Founc | Not Founc | Not Founc | Not Founc | Not Founc | Not Founc | Not Founc |
| | | | | | Not Founc | Not Founc | Not Founc | Not Founc | High | High | High | High | Not Founc | Not Founc |
| | | | | | Not Founc | Not Founc | Not Founc | Not Founc | Not Founc | Not Founc | Not Founc | Not Founc | Not Founc | Not Founc |
| | 4739 | 5156 | 5219 | 7253 | Not Founc | Not Founc | Not Founc | Not Founc | Not Founc | Not Founc | Not Founc | Not Founc | High | High |
| | | | | | High | High | High | High | Not Founc | Not Founc | Not Founc | Not Founc | Not Founc | Not Founc |
| | 1091 | 712 | 1884 | 1122 | Not Founc | Not Founc | Not Founc | Not Founc | Not Founc | Not Founc | Not Founc | Not Founc | High | High |
| | | | | | High | High | High | High | Not Founc | Not Founc | Not Founc | Not Founc | Not Founc | Not Founc |
| | | | | | High | High | High | High | Not Founc | Not Founc | Not Founc | Not Founc | Not Founc | Not Founc |
| | | | | | Not Founc | Not Founc | Not Founc | Not Founc | High | High | High | High | Not Founc | Not Founc |
| | | | | | High | High | High | High | Not Founc | Not Founc | Not Founc | Not Founc | Not Founc | Not Founc |

| | | | | | | | | | | | | | | |
|---|---|---|---|---|---|---|---|---|---|---|---|---|---|---|
| | | | | | High | High | High | High | Not Found | Not Found | Not Found | Not Found | Not Found | Not Found |
| | | | | | Not Found | Not Found | Not Found | Not Found | Not Found | Not Found | Not Found | Not Found | Not Found | Not Found |
| | | | | | High | High | High | High | Not Found | Not Found | Not Found | Not Found | Not Found | Not Found |
| | | | | | High | High | High | High | Not Found | Not Found | Not Found | Not Found | Not Found | Not Found |
| | | | | | Not Found | Not Found | Not Found | Not Found | Not Found | Not Found | Not Found | Not Found | High | High |
| 9039 | | | | | Not Found | Not Found | Not Found | Not Found | High | High | High | High | Not Found | Not Found |
| | | | | | High | High | High | High | Not Found | Not Found | Not Found | Not Found | Not Found | Not Found |
| | | | | | High | High | High | High | Not Found | Not Found | Not Found | Not Found | Not Found | Not Found |
| | | | | | High | High | High | High | Not Found | Not Found | Not Found | Not Found | Not Found | Not Found |
| | | | | | High | High | High | High | Not Found | Not Found | Not Found | Not Found | Not Found | Not Found |
| | 6144 | 5305 | 5964 | 6467 | Not Found | Not Found | Not Found | Not Found | Not Found | Not Found | Not Found | Not Found | High | High |
| 3487 | | | | | Not Found | Not Found | Not Found | Not Found | High | High | High | High | Not Found | Not Found |
| | | | | | Not Found | Not Found | Not Found | Not Found | Not Found | Not Found | Not Found | Not Found | Not Found | Not Found |
| | | | | | High | High | High | High | Not Found | Not Found | Not Found | Not Found | Not Found | Not Found |
| | | | | | High | High | High | High | Not Found | Not Found | Not Found | Not Found | Not Found | Not Found |
| 4058 | | | | | Not Found | Not Found | Not Found | Not Found | High | High | High | High | Not Found | Not Found |
| | | | | | Not Found | Not Found | Not Found | Not Found | Not Found | Not Found | Not Found | Not Found | Not Found | Not Found |
| | | | | | High | High | High | High | Not Found | Not Found | Not Found | Not Found | Not Found | Not Found |
| | 2205 | 1905 | 2774 | 2975 | Not Found | Not Found | Not Found | Not Found | Not Found | Not Found | Not Found | Not Found | High | High |
| | | | | | Not Found | Not Found | Not Found | Not Found | High | High | High | High | Not Found | Not Found |
| | | | | | High | High | High | High | Not Found | Not Found | Not Found | Not Found | Not Found | Not Found |
| | 2895 | 2955 | 3106 | 5222 | Not Found | Not Found | Not Found | Not Found | Not Found | Not Found | Not Found | Not Found | High | High |
| 8321 | | | | | Not Found | Not Found | Not Found | Not Found | High | High | High | High | Not Found | Not Found |
| | 5168 | 4711 | 8043 | 8269 | Not Found | Not Found | Not Found | Not Found | Not Found | Not Found | Not Found | Not Found | High | High |
| | | | | | Not Found | Not Found | Not Found | Not Found | Not Found | Not Found | Not Found | Not Found | Not Found | Not Found |
| 1520 | | | | | Not Found | Not Found | Not Found | Not Found | High | High | High | High | Not Found | Not Found |
| | | | | | High | High | High | High | Not Found | Not Found | Not Found | Not Found | Not Found | Not Found |
| | 2450 | 1654 | 3151 | 2789 | Not Found | Not Found | Not Found | Not Found | Not Found | Not Found | Not Found | Not Found | High | High |
| | | | | | High | High | High | High | Not Found | Not Found | Not Found | Not Found | Not Found | Not Found |
| | | | | | Not Found | Not Found | Not Found | Not Found | High | High | High | High | Not Found | Not Found |
| | | | | | High | High | High | High | Not Found | Not Found | Not Found | Not Found | Not Found | Not Found |

| | | | | | | | | | | | | | | |
|---|---|---|---|---|---|---|---|---|---|---|---|---|---|---|
| 6559 | | | | | Not Found | Not Found | Not Found | Not Found | High | High | High | High | Not Found | Not Found |
| 4542 | | | | | Not Found | Not Found | Not Found | Not Found | High | High | High | High | Not Found | Not Found |
| | 2219 | 1494 | 4200 | 2887 | Not Found | Not Found | Not Found | Not Found | Not Found | Not Found | Not Found | Not Found | High | High |
| 2523 | | | | | Not Found | Not Found | Not Found | Not Found | High | High | High | High | Not Found | Not Found |
| | 629 | 463 | 898 | 812 | Not Found | Not Found | Not Found | Not Found | Not Found | Not Found | Not Found | Not Found | High | High |
| | 1306 | 1917 | 1269 | 2015 | Not Found | Not Found | Not Found | Not Found | Not Found | Not Found | Not Found | Not Found | High | High |
| 2377 | | | | | Not Found | Not Found | Not Found | Not Found | High | High | High | High | Not Found | Not Found |
| | | | | | Not Found | Not Found | Not Found | Not Found | Not Found | Not Found | Not Found | Not Found | Not Found | Not Found |
| 6581 | | | | | Not Found | Not Found | Not Found | Not Found | High | High | High | High | Not Found | Not Found |
| | 4154 | 2750 | 5617 | 5704 | Not Found | Not Found | Not Found | Not Found | Not Found | Not Found | Not Found | Not Found | High | High |
| | 6693 | 4589 | 8662 | 8985 | Not Found | Not Found | Not Found | Not Found | Not Found | Not Found | Not Found | Not Found | High | High |
| | 2830 | 1970 | 3968 | 3635 | Not Found | Not Found | Not Found | Not Found | Not Found | Not Found | Not Found | Not Found | High | High |
| | | | | | Not Found | Not Found | Not Found | Not Found | Not Found | Not Found | Not Found | Not Found | Not Found | Not Found |
| | 6417 | 1580 | 8068 | 2377 | Not Found | Not Found | Not Found | Not Found | Not Found | Not Found | Not Found | Not Found | High | High |
| | | | | | High | High | High | High | Not Found | Not Found | Not Found | Not Found | Not Found | Not Found |
| | 2339 | 1428 | 3350 | 2158 | Not Found | Not Found | Not Found | Not Found | Not Found | Not Found | Not Found | Not Found | High | High |
| | | | | | High | High | High | High | Not Found | Not Found | Not Found | Not Found | Not Found | Not Found |
| | | | | | High | High | High | High | Not Found | Not Found | Not Found | Not Found | Not Found | Not Found |
| 4815 | | | | | Not Found | Not Found | Not Found | Not Found | High | High | High | High | Not Found | Not Found |
| 9477 | | | | | Not Found | Not Found | Not Found | Not Found | High | High | High | High | Not Found | Not Found |
| 7272 | | | | | Not Found | Not Found | Not Found | Not Found | High | High | High | High | Not Found | Not Found |
| | 2212 | 1334 | 3502 | 2484 | Not Found | Not Found | Not Found | Not Found | Not Found | Not Found | Not Found | Not Found | High | High |
| | | | | | High | High | High | High | Not Found | Not Found | Not Found | Not Found | Not Found | Not Found |
| | 2633 | 1879 | 3636 | 2872 | Not Found | Not Found | Not Found | Not Found | Not Found | Not Found | Not Found | Not Found | High | High |
| | 1459 | 1167 | 2115 | 2148 | Not Found | Not Found | Not Found | Not Found | Not Found | Not Found | Not Found | Not Found | High | High |
| 7264 | | | | | Not Found | Not Found | Not Found | Not Found | High | High | High | High | Not Found | Not Found |
| | | | | | High | High | High | High | Not Found | Not Found | Not Found | Not Found | Not Found | Not Found |
| | | | | | High | High | High | High | Not Found | Not Found | Not Found | Not Found | Not Found | Not Found |
| | 6308 | 3691 | 6099 | 6158 | Not Found | Not Found | Not Found | Not Found | Not Found | Not Found | Not Found | Not Found | High | High |
| | | | | | High | High | High | High | Not Found | Not Found | Not Found | Not Found | Not Found | Not Found |
| | | | | | Not Found | Not Found | Not Found | Not Found | Not Found | Not Found | Not Found | Not Found | Not Found | Not Found |

| | | | | | | | | | | | | | | |
|---|---|---|---|---|---|---|---|---|---|---|---|---|---|---|
| | 2618 | 1397 | 3564 | 4159 | Not Founc | Not Founc | Not Founc | Not Founc | Not Founc | Not Founc | Not Founc | Not Founc | High | High |
| 1202 | | | | | Not Founc | Not Founc | Not Founc | Not Founc | High | High | High | High | Not Founc | Not Founc |
| | 1097 | 845 | 1013 | 1082 | Not Founc | Not Founc | Not Founc | Not Founc | Not Founc | Not Founc | Not Founc | Not Founc | High | High |
| | | | | | Not Founc | Not Founc | Not Founc | Not Founc | Not Founc | Not Founc | Not Founc | Not Founc | Not Founc | Not Founc |
| | 2964 | 2223 | 3608 | 3736 | Not Founc | Not Founc | Not Founc | Not Founc | Not Founc | Not Founc | Not Founc | Not Founc | High | High |
| | 1327 | 1050 | 1367 | 1367 | Not Founc | Not Founc | Not Founc | Not Founc | Not Founc | Not Founc | Not Founc | Not Founc | High | High |
| 2110 | | | | | Not Founc | Not Founc | Not Founc | Not Founc | High | High | High | High | Not Founc | Not Founc |

| Found in S | Found in S | # Protein | Modificati |
|---|---|---|---|
| High | High | 1 | |
| High | High | 1 | |
| High | High | 1 | |
| High | High | 1 | |
| High | High | 1 | |
| High | High | 1 | |
| High | High | 1 | |
| High | High | 1 | |
| High | High | 1 | |
| High | High | 1 | |
| High | High | 1 | |
| High | High | 1 | |
| High | High | 1 | |
| High | High | 1 | |
| High | High | 1 | |
| High | High | 1 | |
| High | High | 1 | |
| High | High | 1 | |
| High | High | 1 | |
| High | High | 1 | |
| High | High | 1 | |
| High | High | 1 | |
| High | High | 1 | |
| High | High | 1 | |
| High | High | 1 | Acetyl [N- |
| High | High | 1 | |
| High | High | 1 | |
| High | High | 1 | |
| High | High | 1 | |
| High | High | 1 | |
| High | High | 1 | |

| | | | |
|---|---|---|---|
| Not Found | Not Found | 1 | |
| High | High | 1 | |
| High | High | 1 | |
| High | High | 1 | |
| High | High | 1 | |
| High | High | 1 | |
| High | High | 1 | |
| High | High | 1 | |
| High | High | 1 | |
| High | High | 1 | |
| High | High | 1 | |
| High | High | 1 | Acetyl [N- |
| High | High | 1 | |
| High | High | 1 | |
| High | High | 1 | |
| High | High | 1 | |
| High | High | 1 | |
| High | High | 1 | |
| High | High | 1 | |
| High | High | 1 | |
| High | High | 1 | |
| High | High | 1 | |
| High | High | 1 | |
| High | High | 1 | |
| Not Found | Not Found | 1 | |
| High | High | 1 | |
| High | High | 1 | |
| High | High | 1 | |
| High | High | 1 | |
| High | High | 1 | |
| High | High | 1 | |

| | | | |
|---|---|---|---|
| High | High | 1 | |
| High | High | 1 | |
| High | High | 1 | |
| High | High | 1 | |
| High | High | 1 | |
| High | High | 1 | |
| High | High | 1 | |
| High | High | 1 | |
| High | High | 1 | |
| High | High | 1 | |
| High | High | 1 | |
| High | High | 1 | |
| High | High | 1 | |
| High | High | 1 | |
| High | High | 1 | |
| High | High | 1 | |
| High | High | 1 | |
| High | High | 1 | Acetyl [N- |
| High | High | 1 | |
| High | High | 1 | |
| High | High | 1 | |
| High | High | 1 | |
| High | High | 1 | Acetyl [N- |
| High | High | 1 | |
| High | High | 1 | Acetyl [N- |
| High | High | 1 | |
| High | High | 1 | |
| High | High | 1 | |
| High | High | 1 | |
| High | High | 1 | |
| High | High | 1 | |

| | | | |
|---|---|---|---|
| High | High | 1 | |
| High | High | 1 | |
| High | High | 1 | |
| High | High | 1 | |
| High | High | 1 | |
| High | High | 1 | Acetyl [N- |
| High | High | 1 | |
| High | High | 1 | |
| Not Founc | Not Founc | 1 | |
| High | High | 1 | |
| High | High | 1 | |
| High | High | 1 | |
| High | High | 1 | |
| High | High | 1 | |
| Not Founc | Not Founc | 1 | |
| High | High | 1 | |
| High | High | 1 | |
| High | High | 1 | |
| High | High | 1 | |
| High | High | 1 | |
| High | High | 1 | |
| High | High | 1 | Acetyl [N- |
| High | High | 1 | |
| High | High | 1 | |
| High | High | 1 | |
| High | High | 1 | |
| High | High | 1 | |
| High | High | 1 | |
| High | High | 1 | Acetyl [N- |
| High | High | 1 | |
| High | High | 1 | |

| | | | |
|---|---|---|---|
| High | High | 1 | |
| High | High | 1 | |
| High | High | 1 | Acetyl [N- |
| High | High | 1 | |
| High | High | 1 | |
| High | High | 1 | |
| High | High | 1 | |
| High | High | 1 | |
| High | High | 1 | |
| High | High | 1 | |
| High | High | 1 | |
| High | High | 1 | |
| High | High | 1 | |
| High | High | 1 | |
| High | High | 1 | |
| High | High | 1 | |
| High | High | 1 | |
| High | High | 1 | Acetyl [N- |
| High | High | 1 | |
| High | High | 1 | |
| High | High | 1 | |
| High | High | 1 | |
| High | High | 1 | |
| High | High | 1 | |
| High | High | 1 | |
| High | High | 1 | |
| High | High | 1 | Acetyl [N- |
| High | High | 1 | |
| High | High | 1 | |
| High | High | 1 | |
| High | High | 1 | |

| | | | |
|---|---|---|---|
| High | High | 1 | |
| High | High | 1 | |
| High | High | 1 | |
| High | High | 1 | |
| High | High | 1 | |
| High | High | 1 | |
| High | High | 1 | |
| High | High | 1 | |
| High | High | 1 | |
| High | High | 1 | |
| High | High | 1 | |
| High | High | 1 | |
| High | High | 1 | |
| High | High | 1 | |
| High | High | 1 | |
| High | High | 1 | |
| High | High | 1 | |
| High | High | 1 | |
| High | High | 1 | |
| High | High | 1 | |
| High | High | 1 | Acetyl [N- |
| High | High | 1 | |
| High | High | 1 | |
| High | High | 1 | |
| High | High | 1 | |
| High | High | 1 | |
| High | High | 1 | |
| High | High | 1 | |
| High | High | 1 | |
| High | High | 1 | |
| High | High | 1 | |

| | | | |
|---|---|---|---|
| High | High | 1 | |
| High | High | 1 | |
| High | High | 1 | |
| High | High | 1 | |
| High | High | 1 | |
| High | High | 1 | |
| Not Founc | Not Founc | 1 | |
| High | High | 1 | |
| High | High | 1 | |
| High | High | 1 | |
| High | High | 1 | |
| High | High | 1 | |
| High | High | 1 | |
| High | High | 1 | |
| High | High | 1 | |
| High | High | 1 | |
| High | High | 1 | |
| High | High | 1 | |
| High | High | 1 | |
| High | High | 1 | |
| High | High | 1 | |
| High | High | 1 | |
| High | High | 1 | |
| High | High | 1 | |
| High | High | 1 | |
| High | High | 1 | |
| High | High | 1 | |
| High | High | 1 | Acetyl [N- |
| High | High | 1 | |
| High | High | 1 | |
| High | High | 1 | |

| | | | |
|---|---|---|---|
| High | High | 1 | |
| High | High | 1 | |
| High | High | 1 | |
| High | High | 1 | |
| High | High | 1 | |
| High | High | 1 | |
| High | High | 1 | |
| High | High | 1 | |
| High | High | 1 | |
| High | High | 1 | |
| High | High | 1 | |
| High | High | 1 | |
| High | High | 1 | |
| High | High | 1 | |
| High | High | 1 | |
| High | High | 1 | |
| High | High | 1 | |
| High | High | 1 | |
| High | High | 1 | |
| High | High | 1 | |
| High | High | 1 | |
| High | High | 1 | |
| High | High | 1 | |
| High | High | 1 | |
| High | High | 1 | |
| High | High | 1 | |
| High | High | 1 | |
| High | High | 1 | |
| High | High | 1 | |
| High | High | 1 | Acetyl [N- |
| High | High | 1 | |

| | | | |
|---|---|---|---|
| High | High | 1 | |
| High | High | 1 | |
| High | High | 1 | |
| High | High | 1 | |
| High | High | 1 | |
| High | High | 1 | |
| High | High | 1 | |
| High | High | 1 | |
| High | High | 1 | |
| High | High | 1 | |
| High | High | 1 | |
| High | High | 1 | |
| High | High | 1 | |
| High | High | 1 | |
| High | High | 1 | |
| High | High | 1 | |
| High | High | 1 | |
| Not Founc | Not Founc | 1 | |
| High | High | 1 | |
| High | High | 1 | |
| High | High | 1 | |
| High | High | 1 | |
| Not Founc | Not Founc | 1 | |
| High | High | 1 | |
| High | High | 1 | |
| High | High | 1 | |
| High | High | 1 | |
| High | High | 1 | |
| High | High | 1 | Acetyl [N- |
| High | High | 1 | |
| High | High | 1 | |

| | | | |
|---|---|---|---|
| High | High | 1 | |
| Not Founc | Not Founc | 1 | |
| High | High | 1 | |
| High | High | 1 | |
| High | High | 1 | |
| High | High | 1 | |
| High | High | 1 | |
| High | High | 1 | |
| High | High | 1 | |
| High | High | 1 | |
| High | High | 1 | |
| High | High | 1 | |
| High | High | 1 | |
| High | High | 1 | |
| High | High | 1 | |
| High | High | 1 | |
| High | High | 1 | |
| High | High | 1 | |
| High | High | 1 | |
| High | High | 1 | |
| High | High | 1 | |
| High | High | 1 | |
| High | High | 1 | |
| High | High | 1 | |
| High | High | 1 | |
| High | High | 1 | |
| High | High | 1 | Acetyl [N- |
| High | High | 1 | |
| High | High | 1 | |
| High | High | 1 | |
| High | High | 1 | |

| | | | |
|---|---|---|---|
| High | High | 1 | |
| High | High | 1 | |
| High | High | 1 | |
| High | High | 1 | |
| High | High | 1 | |
| High | High | 1 | |
| Not Founc | Not Founc | 1 | |
| High | High | 1 | |
| High | High | 1 | |
| High | High | 1 | |
| High | High | 1 | |
| High | High | 1 | |
| High | High | 1 | |
| High | High | 1 | |
| High | High | 1 | |
| High | High | 1 | |
| High | High | 1 | |
| High | High | 1 | |
| High | High | 1 | |
| High | High | 1 | |
| High | High | 1 | |
| High | High | 1 | |
| High | High | 1 | Acetyl [N- |
| High | High | 1 | |
| High | High | 1 | |
| High | High | 1 | Acetyl [N- |
| High | High | 1 | |
| High | High | 1 | |
| High | High | 1 | |
| High | High | 1 | |
| High | High | 1 | |

| | | | |
|---|---|---|---|
| High | High | 1 | |
| High | High | 1 | Acetyl [N- |
| High | High | 1 | |
| High | High | 1 | |
| High | High | 1 | |
| High | High | 1 | |
| High | High | 1 | |
| High | High | 1 | |
| High | High | 1 | |
| High | High | 1 | Acetyl [N- |
| High | High | 1 | |
| High | High | 1 | |
| High | High | 1 | |
| High | High | 1 | |
| High | High | 1 | |
| High | High | 1 | |
| High | High | 1 | |
| High | High | 1 | |
| High | High | 1 | Acetyl [N- |
| High | High | 1 | |
| High | High | 1 | |
| High | High | 1 | |
| High | High | 1 | |
| High | High | 1 | |
| High | High | 1 | |
| High | High | 1 | |
| High | High | 1 | |
| High | High | 1 | |
| High | High | 1 | |
| High | High | 1 | |
| High | High | 1 | |

| | | | |
|---|---|---|---|
| High | High | 1 | |
| High | High | 1 | |
| High | High | 1 | |
| High | High | 1 | |
| High | High | 1 | |
| High | High | 1 | |
| High | High | 1 | |
| High | High | 1 | |
| High | High | 1 | |
| High | High | 1 | |
| High | High | 1 | |
| High | High | 1 | |
| High | High | 1 | |
| High | High | 1 | |
| High | High | 1 | |
| High | High | 1 | |
| High | High | 1 | Acetyl [N- |
| High | High | 1 | |
| High | High | 1 | |
| High | High | 1 | |
| High | High | 1 | Acetyl [N- |
| High | High | 1 | |
| High | High | 1 | |
| Not Founc | Not Founc | 1 | |
| High | High | 1 | |
| High | High | 1 | |
| High | High | 1 | |
| High | High | 1 | |
| High | High | 1 | |
| High | High | 1 | |
| High | High | 1 | |

| | | |
|---|---|---|
| High | High | 1 |
| High | High | 1 |
| High | High | 1 |
| High | High | 1 |
| High | High | 1 |
| High | High | 1 |
| High | High | 1 |
| High | High | 1 |
| High | High | 1 |
| High | High | 1 |
| High | High | 1 |
| High | High | 1 |
| High | High | 1 |
| High | High | 1 |
| High | High | 1 |
| High | High | 1 |
| High | High | 1 |
| High | High | 1 |
| High | High | 1 |
| High | High | 1 |
| High | High | 1 |
| High | High | 1 |
| High | High | 1 |
| High | High | 1 |
| High | High | 1 |
| High | High | 1 |
| High | High | 1 |
| High | High | 1 |
| High | High | 1 |
| High | High | 1 |
| High | High | 1 |

| | | | |
|---|---|---|---|
| High | High | 1 | |
| High | High | 1 | |
| High | High | 1 | Acetyl [N- |
| High | High | 1 | |
| High | High | 1 | |
| High | High | 1 | |
| High | High | 1 | |
| High | High | 1 | |
| Not Founc | Not Founc | 1 | |
| High | High | 1 | |
| High | High | 1 | |
| High | High | 1 | |
| High | High | 1 | |
| High | High | 1 | Acetyl [N- |
| High | High | 1 | |
| High | High | 1 | |
| High | High | 1 | |
| High | High | 1 | |
| High | High | 1 | |
| High | High | 1 | |
| High | High | 1 | |
| High | High | 1 | |
| High | High | 1 | |
| High | High | 1 | |
| High | High | 1 | |
| High | High | 1 | |
| High | High | 1 | |
| High | High | 1 | |
| High | High | 1 | |
| High | High | 1 | |
| High | High | 1 | |

| | | | |
|---|---|---|---|
| High | High | 1 | |
| High | High | 1 | |
| High | High | 1 | |
| High | High | 1 | |
| High | High | 1 | |
| High | High | 1 | |
| High | High | 1 | Acetyl [N- |
| High | High | 1 | |
| High | High | 1 | |
| High | High | 1 | |
| High | High | 1 | |
| High | High | 1 | |
| High | High | 1 | Acetyl [N- |
| High | High | 1 | |
| High | High | 1 | |
| High | High | 1 | |
| High | High | 1 | |
| High | High | 1 | Acetyl [N- |
| High | High | 1 | |
| High | High | 1 | |
| High | High | 1 | |
| High | High | 1 | |
| High | High | 1 | |
| High | High | 1 | |
| High | High | 1 | |
| High | High | 1 | |
| High | High | 1 | |
| High | High | 1 | |
| High | High | 1 | |
| High | High | 1 | |
| High | High | 1 | |

| | | | |
|---|---|---|---|
| High | High | 1 | |
| High | High | 1 | |
| High | High | 1 | |
| High | High | 1 | |
| High | High | 1 | Acetyl [N- |
| High | High | 1 | |
| High | High | 1 | |
| High | High | 1 | |
| High | High | 1 | |
| High | High | 1 | |
| High | High | 1 | |
| High | High | 1 | |
| High | High | 1 | |
| High | High | 1 | |
| High | High | 1 | |
| High | High | 1 | |
| High | High | 1 | |
| High | High | 1 | |
| High | High | 1 | |
| High | High | 1 | |
| High | High | 1 | |
| High | High | 1 | |
| High | High | 1 | |
| High | High | 1 | |
| High | High | 1 | |
| High | High | 1 | |
| High | High | 1 | |
| High | High | 1 | |
| High | High | 1 | |
| High | High | 1 | |
| High | High | 1 | Acetyl [N- |

| | | | |
|---|---|---|---|
| High | High | 1 | |
| High | High | 1 | |
| High | High | 1 | |
| High | High | 1 | |
| High | High | 1 | |
| High | High | 1 | |
| High | High | 1 | |
| High | High | 1 | |
| High | High | 1 | |
| High | High | 1 | Acetyl [N- |
| High | High | 1 | |
| High | High | 1 | |
| High | High | 1 | |
| High | High | 1 | |
| High | High | 1 | |
| High | High | 1 | |
| High | High | 1 | |
| High | High | 1 | |
| High | High | 1 | |
| High | High | 1 | |
| High | High | 1 | |
| High | High | 1 | |
| Not Founc | Not Founc | 1 | Acetyl [N- |
| High | High | 1 | |
| High | High | 1 | |
| High | High | 1 | |
| High | High | 1 | |
| High | High | 1 | |
| High | High | 1 | |
| High | High | 1 | |
| High | High | 1 | |

| | | | |
|---|---|---|---|
| High | High | 1 | |
| High | High | 1 | |
| High | High | 1 | |
| High | High | 1 | |
| High | High | 1 | |
| High | High | 1 | |
| High | High | 1 | |
| High | High | 1 | |
| High | High | 1 | |
| High | High | 1 | |
| High | High | 1 | |
| High | High | 1 | |
| High | High | 1 | |
| High | High | 1 | |
| High | High | 1 | |
| High | High | 1 | |
| High | High | 1 | |
| High | High | 1 | |
| High | High | 1 | |
| High | High | 1 | |
| High | High | 1 | |
| High | High | 1 | |
| High | High | 1 | |
| High | High | 1 | |
| High | High | 1 | |
| High | High | 1 | |
| High | High | 1 | Acetyl [N- |
| High | High | 1 | |
| High | High | 1 | |
| High | High | 1 | |
| High | High | 1 | |

| | | | |
|---|---|---|---|
| High | High | 1 | |
| High | High | 1 | |
| High | High | 1 | |
| High | High | 1 | |
| High | High | 1 | |
| High | High | 1 | |
| High | High | 1 | Acetyl [N- |
| High | High | 1 | |
| High | High | 1 | |
| High | High | 1 | |
| High | High | 1 | Acetyl [N- |
| High | High | 1 | |
| High | High | 1 | |
| High | High | 1 | |
| High | High | 1 | Acetyl [N- |
| High | High | 1 | |
| High | High | 1 | |
| High | High | 1 | |
| High | High | 1 | |
| High | High | 1 | |
| High | High | 1 | |
| High | High | 1 | |
| High | High | 1 | |
| High | High | 1 | |
| High | High | 1 | |
| High | High | 1 | |
| High | High | 1 | |
| High | High | 1 | |
| High | High | 1 | |
| High | High | 1 | |
| High | High | 1 | |

| | | | |
|---|---|---|---|
| High | High | 1 | |
| High | High | 1 | |
| High | High | 1 | |
| High | High | 1 | |
| High | High | 1 | |
| High | High | 1 | Acetyl [N- |
| High | High | 1 | |
| High | High | 1 | |
| High | High | 1 | |
| High | High | 1 | |
| High | High | 1 | |
| High | High | 1 | |
| High | High | 1 | |
| High | High | 1 | |
| High | High | 1 | |
| High | High | 1 | |
| High | High | 1 | |
| High | High | 1 | |
| High | High | 1 | |
| High | High | 1 | |
| High | High | 1 | Acetyl [N- |
| High | High | 1 | |
| High | High | 1 | |
| High | High | 1 | |
| High | High | 1 | |
| High | High | 1 | |
| High | High | 1 | Acetyl [N- |
| High | High | 1 | |
| High | High | 1 | |
| High | High | 1 | |
| High | High | 1 | |

| | | | |
|---|---|---|---|
| High | High | 1 | |
| High | High | 1 | |
| High | High | 1 | |
| High | High | 1 | |
| High | High | 1 | |
| High | High | 1 | |
| High | High | 1 | |
| High | High | 1 | |
| High | High | 1 | |
| High | High | 1 | |
| High | High | 1 | |
| High | High | 1 | |
| High | High | 1 | |
| High | High | 1 | |
| High | High | 1 | |
| High | High | 1 | |
| High | High | 1 | |
| High | High | 1 | |
| High | High | 1 | |
| High | High | 1 | |
| High | High | 1 | |
| High | High | 1 | |
| High | High | 1 | |
| High | High | 1 | |
| High | High | 1 | |
| High | High | 1 | |
| High | High | 1 | Acetyl [N- |
| High | High | 1 | |
| High | High | 1 | |
| High | High | 1 | |
| High | High | 1 | Acetyl [N- |

| | | | |
|---|---|---|---|
| High | High | 1 | |
| High | High | 1 | |
| High | High | 1 | |
| High | High | 1 | |
| High | High | 1 | |
| High | High | 1 | |
| High | High | 1 | |
| High | High | 1 | |
| High | High | 1 | |
| High | High | 1 | |
| High | High | 1 | |
| High | High | 1 | |
| High | High | 1 | |
| High | High | 1 | |
| High | High | 1 | |
| High | High | 1 | |
| High | High | 1 | |
| High | High | 1 | |
| High | High | 1 | Acetyl [N- |
| High | High | 1 | |
| High | High | 1 | |
| High | High | 1 | |
| High | High | 1 | |
| High | High | 1 | |
| High | High | 1 | |
| High | High | 1 | |
| High | High | 1 | |
| High | High | 1 | |
| High | High | 1 | |
| High | High | 1 | |
| High | High | 1 | |

| | | |
|---|---|---|
| High | High | 1 |
| High | High | 1 |
| High | High | 1 |
| High | High | 1 |
| High | High | 1 |
| High | High | 1 |
| High | High | 1 |
| High | High | 1 |
| High | High | 1 |
| High | High | 1 |
| High | High | 1 |
| High | High | 1 |
| High | High | 1 |
| High | High | 1 |
| High | High | 1 |
| High | High | 1 |
| High | High | 1 |
| High | High | 1 |
| High | High | 1 |
| High | High | 1 |
| High | High | 1 |
| High | High | 1 |
| High | High | 1 |
| High | High | 1 |
| High | High | 1 |
| High | High | 1 |
| High | High | 1 |
| High | High | 1 |
| High | High | 1 |
| High | High | 1 |
| High | High | 1 |

| | | | |
|---|---|---|---|
| High | High | 1 | |
| High | High | 1 | |
| High | High | 1 | |
| High | High | 1 | |
| High | High | 1 | |
| High | High | 1 | |
| High | High | 1 | |
| High | High | 1 | |
| High | High | 1 | |
| High | High | 1 | |
| High | High | 1 | Acetyl [N- |
| High | High | 1 | |
| High | High | 1 | |
| High | High | 1 | |
| High | High | 1 | |
| High | High | 1 | |
| High | High | 1 | |
| High | High | 1 | |
| High | High | 1 | |
| High | High | 1 | |
| High | High | 1 | |
| High | High | 1 | |
| High | High | 1 | |
| High | High | 1 | |
| High | High | 1 | |
| High | High | 1 | |
| High | High | 1 | |
| High | High | 1 | |
| High | High | 1 | |
| High | High | 1 | Acetyl [N- |
| High | High | 1 | |

| | | |
|---|---|---|
| High | High | 1 |
| High | High | 1 |
| High | High | 1 |
| High | High | 1 |
| High | High | 1 |
| High | High | 1 |
| High | High | 1 |
| High | High | 1 |
| High | High | 1 |
| High | High | 1 |
| High | High | 1 |
| High | High | 1 |
| High | High | 1 |
| High | High | 1 |
| High | High | 1 |
| High | High | 1 |
| High | High | 1 |
| High | High | 1 |
| High | High | 1 |
| High | High | 1 |
| High | High | 1 |
| High | High | 1 |
| High | High | 1 |
| High | High | 1 |
| High | High | 1 |
| High | High | 1 |
| High | High | 1 |
| High | High | 1 |
| High | High | 1 |
| High | High | 1 |
| High | High | 1 |

| | | | |
|---|---|---|---|
| High | High | 1 | |
| High | High | 1 | |
| High | High | 1 | |
| High | High | 1 | |
| High | High | 1 | |
| High | High | 1 | |
| High | High | 1 | |
| High | High | 1 | |
| High | High | 1 | |
| High | High | 1 | |
| High | High | 1 | |
| High | High | 1 | |
| High | High | 1 | |
| High | High | 1 | Acetyl [N- |
| High | High | 1 | |
| High | High | 1 | |
| High | High | 1 | |
| High | High | 1 | |
| High | High | 1 | |
| High | High | 1 | |
| High | High | 1 | |
| High | High | 1 | |
| High | High | 1 | |
| High | High | 1 | |
| High | High | 1 | |
| High | High | 1 | |
| High | High | 1 | |
| High | High | 1 | |
| High | High | 1 | |
| High | High | 1 | Acetyl [N- |
| High | High | 1 | |

| | | | |
|---|---|---|---|
| High | High | 1 | |
| Not Founc | Not Founc | 1 | |
| High | High | 1 | |
| High | High | 1 | |
| High | High | 1 | |
| High | High | 1 | |
| High | High | 1 | Acetyl [N- |
| High | High | 1 | |
| High | High | 1 | |
| High | High | 1 | |
| High | High | 1 | |
| High | High | 1 | |
| High | High | 1 | |
| High | High | 1 | |
| High | High | 1 | |
| High | High | 1 | |
| High | High | 1 | |
| High | High | 1 | |
| High | High | 1 | |
| High | High | 1 | |
| High | High | 1 | |
| High | High | 1 | |
| High | High | 1 | Acetyl [N- |
| High | High | 1 | |
| High | High | 1 | |
| High | High | 1 | |
| High | High | 1 | |
| High | High | 1 | |
| High | High | 1 | |
| High | High | 1 | |
| High | High | 1 | |

| | | | |
|---|---|---|---|
| High | High | 1 | |
| High | High | 1 | |
| High | High | 1 | |
| High | High | 1 | |
| High | High | 1 | |
| High | High | 1 | |
| High | High | 1 | |
| High | High | 1 | |
| High | High | 1 | |
| High | High | 1 | |
| High | High | 1 | Acetyl [N- |
| High | High | 1 | |
| High | High | 1 | |
| High | High | 1 | |
| High | High | 1 | |
| High | High | 1 | |
| High | High | 1 | |
| High | High | 1 | |
| High | High | 1 | |
| High | High | 1 | |
| High | High | 1 | |
| High | High | 1 | |
| High | High | 1 | |
| High | High | 1 | |
| High | High | 1 | |
| High | High | 1 | |
| High | High | 1 | |
| High | High | 1 | |
| High | High | 1 | |
| High | High | 1 | |
| High | High | 1 | |

| | | | |
|---|---|---|---|
| High | High | 1 | |
| High | High | 1 | |
| High | High | 1 | |
| High | High | 1 | |
| High | High | 1 | |
| High | High | 1 | |
| High | High | 1 | |
| High | High | 1 | |
| High | High | 1 | |
| High | High | 1 | |
| High | High | 1 | |
| High | High | 1 | |
| High | High | 1 | |
| High | High | 1 | |
| High | High | 1 | Acetyl [N- |
| High | High | 1 | |
| High | High | 1 | |
| High | High | 1 | |
| High | High | 1 | |
| High | High | 1 | |
| High | High | 1 | |
| High | High | 1 | |
| High | High | 1 | |
| High | High | 1 | |
| High | High | 1 | |
| High | High | 1 | |
| High | High | 1 | |
| High | High | 1 | |
| High | High | 1 | Acetyl [N- |
| High | High | 1 | |
| High | High | 1 | |

| | | | |
|---|---|---|---|
| High | High | 1 | |
| High | High | 1 | |
| High | High | 1 | |
| High | High | 1 | |
| Not Founc | Not Founc | 1 | |
| High | High | 1 | |
| High | High | 1 | |
| High | High | 1 | |
| High | High | 1 | |
| High | High | 1 | |
| High | High | 1 | |
| High | High | 1 | |
| High | High | 1 | |
| High | High | 1 | |
| High | High | 1 | |
| High | High | 1 | Acetyl [N- |
| High | High | 1 | Acetyl [N- |
| High | High | 1 | |
| High | High | 1 | |
| High | High | 1 | |
| High | High | 1 | |
| High | High | 1 | |
| High | High | 1 | |
| High | High | 1 | |
| High | High | 1 | |
| High | High | 1 | Acetyl [N- |
| High | High | 1 | |
| High | High | 1 | |
| High | High | 1 | |
| High | High | 1 | |
| High | High | 1 | |

| | | |
|---|---|---|
| High | High | 1 |
| High | High | 1 |
| High | High | 1 |
| High | High | 1 |
| High | High | 1 |
| High | High | 1 |
| High | High | 1 |
| High | High | 1 |
| High | High | 1 |
| High | High | 1 |
| High | High | 1 |
| High | High | 1 |
| High | High | 1 |
| High | High | 1 |
| High | High | 1 |
| High | High | 1 |
| High | High | 1 |
| High | High | 1 |
| High | High | 1 |
| High | High | 1 |
| High | High | 1 |
| High | High | 1 |
| High | High | 1 |
| High | High | 1 |
| High | High | 1 |
| High | High | 1 |
| Not Founc | Not Founc | 1 |
| High | High | 1 |
| High | High | 1 |
| High | High | 1 |
| High | High | 1 |

| | | | |
|---|---|---|---|
| High | High | 1 | |
| High | High | 1 | |
| High | High | 1 | |
| High | High | 1 | |
| High | High | 1 | |
| High | High | 1 | |
| High | High | 1 | |
| High | High | 1 | |
| High | High | 1 | |
| High | High | 1 | |
| High | High | 1 | |
| High | High | 1 | |
| High | High | 1 | |
| High | High | 1 | |
| High | High | 1 | |
| High | High | 1 | Acetyl [N- |
| High | High | 1 | |
| High | High | 1 | |
| High | High | 1 | |
| High | High | 1 | |
| High | High | 1 | |
| High | High | 1 | |
| High | High | 1 | |
| High | High | 1 | |
| High | High | 1 | |
| High | High | 1 | |
| High | High | 1 | |
| High | High | 1 | |
| High | High | 1 | |
| High | High | 1 | |
| High | High | 1 | |

| | | |
|---|---|---|
| High | High | 1 |
| High | High | 1 |
| High | High | 1 |
| High | High | 1 |
| High | High | 1 |
| High | High | 1 |
| High | High | 1 |
| High | High | 1 |
| High | High | 1 |
| High | High | 1 |
| High | High | 1 |
| High | High | 1 |
| High | High | 1 |
| High | High | 1 |
| High | High | 1 |
| High | High | 1 |
| High | High | 1 |
| High | High | 1 |
| High | High | 1 |
| High | High | 1 |
| High | High | 1 |
| High | High | 1 |
| High | High | 1 |
| High | High | 1 |
| High | High | 1 |
| High | High | 1 |
| High | High | 1 |
| High | High | 1 |
| High | High | 1 |
| High | High | 1 |
| High | High | 1 |

| | | |
|---|---|---|
| High | High | 1 |
| High | High | 1 |
| Not Found | Not Found | 1 |
| High | High | 1 |
| High | High | 1 |
| High | High | 1 |
| High | High | 1 |
| High | High | 1 |
| High | High | 1 |
| High | High | 1 |
| High | High | 1 |
| High | High | 1 |
| High | High | 1 |
| High | High | 1 |
| High | High | 1 |
| High | High | 1 |
| High | High | 1 |
| High | High | 1 |
| High | High | 1 |
| High | High | 1 |
| High | High | 1 |
| High | High | 1 |
| High | High | 1 |
| High | High | 1 |
| High | High | 1 |
| High | High | 1 |
| High | High | 1 |
| High | High | 1 |
| High | High | 1 |
| High | High | 1 |
| High | High | 1 |

| | | | |
|---|---|---|---|
| High | High | 1 | |
| High | High | 1 | |
| High | High | 1 | |
| High | High | 1 | Acetyl [N- |
| High | High | 1 | |
| High | High | 1 | |
| High | High | 1 | |
| High | High | 1 | |
| High | High | 1 | |
| High | High | 1 | |
| High | High | 1 | |
| High | High | 1 | |
| High | High | 1 | |
| High | High | 1 | |
| High | High | 1 | |
| High | High | 1 | |
| High | High | 1 | |
| High | High | 1 | |
| High | High | 1 | |
| High | High | 1 | |
| High | High | 1 | |
| High | High | 1 | |
| High | High | 1 | |
| High | High | 1 | Acetyl [N- |
| High | High | 1 | |
| High | High | 1 | |
| High | High | 1 | |
| High | High | 1 | |
| High | High | 1 | |
| High | High | 1 | |
| High | High | 1 | |

| | | |
|---|---|---|
| High | High | 1 |
| High | High | 1 |
| High | High | 1 |
| High | High | 1 |
| High | High | 1 |
| High | High | 1 |
| High | High | 1 |
| High | High | 1 |
| High | High | 1 |
| High | High | 1 |
| High | High | 1 |
| High | High | 1 |
| High | High | 1 |
| High | High | 1 |
| High | High | 1 |
| High | High | 1 |
| High | High | 1 |
| High | High | 1 |
| High | High | 1 |
| High | High | 1 |
| High | High | 1 |
| High | High | 1 |
| High | High | 1 |
| High | High | 1 |
| High | High | 1 |
| High | High | 1 |
| High | High | 1 |
| High | High | 1 |
| High | High | 1 |
| High | High | 1 |
| High | High | 1 |

| | | | |
|---|---|---|---|
| High | High | 1 | |
| High | High | 1 | |
| High | High | 1 | |
| High | High | 1 | |
| High | High | 1 | |
| High | High | 1 | |
| High | High | 1 | |
| High | High | 1 | |
| High | High | 1 | |
| High | High | 1 | |
| High | High | 1 | Acetyl [N- |
| High | High | 1 | |
| High | High | 1 | Acetyl [N- |
| High | High | 1 | |
| High | High | 1 | |
| High | High | 1 | |
| High | High | 1 | |
| High | High | 1 | |
| High | High | 1 | Acetyl [N- |
| High | High | 1 | |
| High | High | 1 | |
| High | High | 1 | |
| High | High | 1 | |
| High | High | 1 | |
| High | High | 1 | Acetyl [N- |
| High | High | 1 | |
| High | High | 1 | |
| High | High | 1 | Acetyl [N- |
| High | High | 1 | |
| High | High | 1 | |
| High | High | 1 | |

| | | |
|---|---|---|
| High | High | 1 |
| High | High | 1 |
| High | High | 1 |
| High | High | 1 |
| High | High | 1 |
| High | High | 1 |
| High | High | 1 |
| High | High | 1 |
| High | High | 1 |
| High | High | 1 |
| High | High | 1 |
| High | High | 1 |
| High | High | 1 |
| High | High | 1 |
| High | High | 1 |
| High | High | 1 |
| High | High | 1 |
| High | High | 1 |
| High | High | 1 |
| High | High | 1 |
| High | High | 1 |
| High | High | 1 |
| High | High | 1 |
| High | High | 1 |
| High | High | 1 |
| High | High | 1 |
| High | High | 1 |
| High | High | 1 |
| High | High | 1 |
| High | High | 1 |
| High | High | 1 |

| | | | |
|---|---|---|---|
| High | High | 1 | |
| High | High | 1 | |
| High | High | 1 | Acetyl [N- |
| High | High | 1 | |
| High | High | 1 | |
| High | High | 1 | |
| High | High | 1 | |
| High | High | 1 | |
| High | High | 1 | |
| High | High | 1 | |
| High | High | 1 | |
| High | High | 1 | |
| High | High | 1 | |
| High | High | 1 | |
| High | High | 1 | |
| High | High | 1 | |
| High | High | 1 | |
| High | High | 1 | |
| High | High | 1 | |
| High | High | 1 | |
| High | High | 1 | |
| High | High | 1 | Acetyl [N- |
| High | High | 1 | |
| High | High | 1 | |
| High | High | 1 | |
| High | High | 1 | |
| High | High | 1 | |
| High | High | 1 | |
| High | High | 1 | |
| High | High | 1 | Acetyl [N- |
| High | High | 1 | |

| | | | |
|---|---|---|---|
| High | High | 1 | |
| High | High | 1 | |
| High | High | 1 | |
| High | High | 1 | Acetyl [N- |
| High | High | 1 | |
| High | High | 1 | |
| High | High | 1 | |
| High | High | 1 | |
| High | High | 1 | |
| High | High | 1 | |
| High | High | 1 | |
| High | High | 1 | |
| High | High | 1 | |
| High | High | 1 | |
| High | High | 1 | |
| High | High | 1 | |
| High | High | 1 | Acetyl [N- |
| High | High | 1 | |
| High | High | 1 | |
| High | High | 1 | |
| High | High | 1 | |
| High | High | 1 | |
| High | High | 1 | |
| High | High | 1 | Acetyl [N- |
| High | High | 1 | |
| High | High | 1 | |
| High | High | 1 | |
| High | High | 1 | |
| High | High | 1 | |
| High | High | 1 | |
| High | High | 1 | |

| | | | |
|---|---|---|---|
| High | High | 1 | |
| High | High | 1 | Acetyl [N- |
| High | High | 1 | |
| High | High | 1 | |
| High | High | 1 | |
| High | High | 1 | |
| High | High | 1 | |
| High | High | 1 | |
| High | High | 1 | |
| High | High | 1 | |
| High | High | 1 | |
| High | High | 1 | |
| High | High | 1 | |
| High | High | 1 | |
| High | High | 1 | |
| High | High | 1 | |
| High | High | 1 | |
| High | High | 1 | |
| High | High | 1 | |
| High | High | 1 | |
| High | High | 1 | |
| High | High | 1 | |
| High | High | 1 | |
| High | High | 1 | |
| High | High | 1 | |
| High | High | 1 | |
| High | High | 1 | |
| High | High | 1 | |
| High | High | 1 | |
| High | High | 1 | |
| High | High | 1 | |

| | | | |
|---|---|---|---|
| High | High | 1 | |
| High | High | 1 | |
| High | High | 1 | Acetyl [N- |
| High | High | 1 | |
| High | High | 1 | |
| High | High | 1 | |
| High | High | 1 | |
| High | High | 1 | |
| High | High | 1 | |
| High | High | 1 | |
| High | High | 1 | |
| High | High | 1 | |
| High | High | 1 | |
| High | High | 1 | |
| High | High | 1 | |
| High | High | 1 | Acetyl [N- |
| High | High | 1 | |
| High | High | 1 | |
| High | High | 1 | |
| High | High | 1 | |
| High | High | 1 | |
| High | High | 1 | |
| High | High | 1 | |
| High | High | 1 | |
| High | High | 1 | |
| High | High | 1 | |
| High | High | 1 | |
| High | High | 1 | |
| High | High | 1 | |
| High | High | 1 | |
| High | High | 1 | |

| | | | |
|---|---|---|---|
| High | High | 1 | |
| High | High | 1 | |
| High | High | 1 | |
| High | High | 1 | |
| High | High | 1 | |
| High | High | 1 | |
| High | High | 1 | Acetyl [N- |
| High | High | 1 | |
| High | High | 1 | |
| High | High | 1 | |
| High | High | 1 | |
| High | High | 1 | |
| High | High | 1 | |
| High | High | 1 | |
| High | High | 1 | |
| High | High | 1 | |
| High | High | 1 | |
| High | High | 1 | |
| High | High | 1 | |
| High | High | 1 | |
| High | High | 1 | |
| High | High | 1 | |
| High | High | 1 | |
| High | High | 1 | |
| High | High | 1 | |
| High | High | 1 | |
| High | High | 1 | |
| High | High | 1 | |
| High | High | 1 | |
| High | High | 1 | |
| High | High | 1 | |
| High | High | 1 | |

| | | | |
|---|---|---|---|
| High | High | 1 | |
| High | High | 1 | |
| High | High | 1 | |
| High | High | 1 | |
| High | High | 1 | Acetyl [N- |
| High | High | 1 | |
| High | High | 1 | |
| High | High | 1 | |
| High | High | 1 | |
| High | High | 1 | |
| High | High | 1 | |
| Not Founc | Not Founc | 1 | |
| High | High | 1 | Acetyl [N- |
| High | High | 1 | |
| High | High | 1 | |
| High | High | 1 | |
| High | High | 1 | |
| High | High | 1 | |
| High | High | 1 | |
| High | High | 1 | |
| High | High | 1 | |
| High | High | 1 | |
| High | High | 1 | |
| High | High | 1 | |
| High | High | 1 | |
| High | High | 1 | |
| High | High | 1 | |
| High | High | 1 | |
| High | High | 1 | |
| High | High | 1 | |
| High | High | 1 | |

| | | | |
|---|---|---|---|
| High | High | 1 | |
| High | High | 1 | |
| High | High | 1 | |
| High | High | 1 | Acetyl [N- |
| High | High | 1 | |
| High | High | 1 | |
| High | High | 1 | |
| High | High | 1 | |
| High | High | 1 | |
| Not Founc | Not Founc | 1 | |
| High | High | 1 | |
| High | High | 1 | |
| High | High | 1 | |
| High | High | 1 | |
| High | High | 1 | |
| High | High | 1 | |
| High | High | 1 | |
| High | High | 1 | |
| High | High | 1 | |
| High | High | 1 | |
| High | High | 1 | |
| High | High | 1 | |
| High | High | 1 | |
| High | High | 1 | |
| High | High | 1 | |
| High | High | 1 | |
| High | High | 1 | |
| High | High | 1 | |
| High | High | 1 | |
| High | High | 1 | |
| High | High | 1 | |

| | | | |
|---|---|---|---|
| High | High | 1 | |
| High | High | 1 | |
| High | High | 1 | |
| High | High | 1 | |
| High | High | 1 | |
| High | High | 1 | |
| High | High | 1 | |
| High | High | 1 | |
| High | High | 1 | |
| High | High | 1 | |
| High | High | 1 | |
| High | High | 1 | |
| High | High | 1 | |
| High | High | 1 | |
| High | High | 1 | |
| High | High | 1 | |
| High | High | 1 | |
| High | High | 1 | |
| High | High | 1 | |
| High | High | 1 | |
| High | High | 1 | |
| High | High | 1 | |
| High | High | 1 | Acetyl [N- |
| High | High | 1 | |
| High | High | 1 | |
| High | High | 1 | |
| High | High | 1 | |
| Not Founc | Not Founc | 1 | |
| High | High | 1 | |
| High | High | 1 | |
| High | High | 1 | |

| | | |
|---|---|---|
| High | High | 1 |
| High | High | 1 |
| High | High | 1 |
| High | High | 1 |
| High | High | 1 |
| High | High | 1 |
| High | High | 1 |
| High | High | 1 |
| High | High | 1 |
| High | High | 1 |
| High | High | 1 |
| High | High | 1 |
| High | High | 1 |
| High | High | 1 |
| High | High | 1 |
| High | High | 1 |
| High | High | 1 |
| High | High | 1 |
| High | High | 1 |
| High | High | 1 |
| High | High | 1 |
| High | High | 1 |
| High | High | 1 |
| High | High | 1 |
| High | High | 1 |
| High | High | 1 |
| High | High | 1 |
| High | High | 1 |
| High | High | 1 |
| High | High | 1 |
| High | High | 1 |

| | | |
|---|---|---|
| High | High | 1 |
| High | High | 1 |
| High | High | 1 |
| High | High | 1 |
| High | High | 1 |
| High | High | 1 |
| High | High | 1 |
| High | High | 1 |
| High | High | 1 |
| High | High | 1 |
| High | High | 1 |
| High | High | 1 |
| High | High | 1 |
| High | High | 1 |
| High | High | 1 |
| High | High | 1 |
| High | High | 1 |
| High | High | 1 |
| High | High | 1 |
| High | High | 1 |
| High | High | 1 |
| High | High | 1 |
| High | High | 1 |
| High | High | 1 |
| High | High | 1 |
| High | High | 1 |
| High | High | 1 |
| High | High | 1 |
| High | High | 1 |
| High | High | 1 |
| Not Founc | Not Founc | 1 |

| | | | |
|---|---|---|---|
| High | High | 1 | |
| High | High | 1 | Acetyl [N- |
| High | High | 1 | |
| High | High | 1 | |
| High | High | 1 | |
| High | High | 1 | |
| High | High | 1 | |
| High | High | 1 | |
| High | High | 1 | |
| High | High | 1 | |
| High | High | 1 | |
| High | High | 1 | |
| High | High | 1 | |
| High | High | 1 | |
| High | High | 1 | |
| High | High | 1 | |
| High | High | 1 | |
| High | High | 1 | |
| High | High | 1 | |
| High | High | 1 | Acetyl [N- |
| High | High | 1 | |
| High | High | 1 | |
| High | High | 1 | |
| High | High | 1 | |
| High | High | 1 | |
| High | High | 1 | |
| High | High | 1 | |
| High | High | 1 | |
| High | High | 1 | |
| High | High | 1 | |
| High | High | 1 | |

| | | |
|---|---|---|
| High | High | 1 |
| High | High | 1 |
| High | High | 1 |
| High | High | 1 |
| High | High | 1 |
| High | High | 1 |
| High | High | 1 |
| High | High | 1 |
| High | High | 1 |
| High | High | 1 |
| High | High | 1 |
| High | High | 1 |
| High | High | 1 |
| High | High | 1 |
| High | High | 1 |
| High | High | 1 |
| High | High | 1 |
| High | High | 1 |
| High | High | 1 |
| High | High | 1 |
| High | High | 1 |
| High | High | 1 |
| High | High | 1 |
| High | High | 1 |
| High | High | 1 |
| High | High | 1 |
| High | High | 1 |
| High | High | 1 |
| High | High | 1 |
| High | High | 1 |
| High | High | 1 |

| | | | |
|---|---|---|---|
| High | High | 1 | |
| High | High | 1 | |
| High | High | 1 | |
| High | High | 1 | |
| High | High | 1 | |
| High | High | 1 | |
| High | High | 1 | |
| High | High | 1 | |
| High | High | 1 | |
| High | High | 1 | |
| High | High | 1 | |
| High | High | 1 | |
| High | High | 1 | |
| High | High | 1 | |
| High | High | 1 | |
| High | High | 1 | |
| High | High | 1 | |
| High | High | 1 | |
| High | High | 1 | |
| High | High | 1 | |
| High | High | 1 | Acetyl [N- |
| High | High | 1 | |
| High | High | 1 | |
| High | High | 1 | |
| High | High | 1 | |
| High | High | 1 | |
| High | High | 1 | |
| High | High | 1 | |
| High | High | 1 | |
| High | High | 1 | |
| High | High | 1 | |

| | | |
|---|---|---|
| High | High | 1 |
| High | High | 1 |
| High | High | 1 |
| High | High | 1 |
| High | High | 1 |
| High | High | 1 |
| High | High | 1 |
| High | High | 1 |
| High | High | 1 |
| High | High | 1 |
| High | High | 1 |
| High | High | 1 |
| High | High | 1 |
| High | High | 1 |
| High | High | 1 |
| High | High | 1 |
| High | High | 1 |
| High | High | 1 |
| High | High | 1 |
| High | High | 1 |
| High | High | 1 |
| High | High | 1 |
| High | High | 1 |
| High | High | 1 |
| High | High | 1 |
| High | High | 1 |
| High | High | 1 |
| High | High | 1 |
| High | High | 1 |
| High | High | 1 |
| High | High | 1 |

| | | | |
|---|---|---|---|
| High | High | 1 | |
| High | High | 1 | |
| High | High | 1 | |
| High | High | 1 | |
| High | High | 1 | Acetyl [N- |
| High | High | 1 | |
| High | High | 1 | |
| High | High | 1 | |
| High | High | 1 | |
| High | High | 1 | |
| High | High | 1 | |
| High | High | 1 | |
| High | High | 1 | |
| High | High | 1 | |
| High | High | 1 | |
| High | High | 1 | |
| High | High | 1 | |
| High | High | 1 | |
| High | High | 1 | |
| High | High | 1 | |
| High | High | 1 | |
| High | High | 1 | |
| High | High | 1 | |
| High | High | 1 | |
| High | High | 1 | |
| High | High | 1 | |
| High | High | 1 | |
| High | High | 1 | |
| High | High | 1 | |
| High | High | 1 | |
| High | High | 1 | |

| | | | |
|---|---|---|---|
| High | High | 1 | |
| High | High | 1 | |
| High | High | 1 | |
| High | High | 1 | |
| High | High | 1 | |
| High | High | 1 | |
| High | High | 1 | |
| High | High | 1 | |
| High | High | 1 | |
| High | High | 1 | |
| High | High | 1 | |
| High | High | 1 | |
| High | High | 1 | |
| High | High | 1 | |
| High | High | 1 | |
| Not Founc | Not Founc | 1 | |
| High | High | 1 | |
| High | High | 1 | |
| High | High | 1 | |
| High | High | 1 | |
| High | High | 1 | |
| High | High | 1 | |
| High | High | 1 | |
| High | High | 1 | |
| High | High | 1 | |
| High | High | 1 | Acetyl [N- |
| High | High | 1 | |
| High | High | 1 | |
| High | High | 1 | Acetyl [N- |
| High | High | 1 | Acetyl [N- |
| High | High | 1 | |

| | | | |
|---|---|---|---|
| High | High | 1 | |
| High | High | 1 | |
| High | High | 1 | |
| High | High | 1 | |
| High | High | 1 | |
| High | High | 1 | |
| High | High | 1 | |
| High | High | 1 | |
| High | High | 1 | |
| High | High | 1 | |
| High | High | 1 | |
| Not Found | Not Found | 1 | |
| High | High | 1 | |
| High | High | 1 | |
| High | High | 1 | |
| High | High | 1 | |
| High | High | 1 | |
| High | High | 1 | |
| High | High | 1 | |
| High | High | 1 | |
| High | High | 1 | |
| High | High | 1 | |
| High | High | 1 | |
| High | High | 1 | |
| High | High | 1 | |
| High | High | 1 | |
| High | High | 1 | |
| High | High | 1 | |
| High | High | 1 | Acetyl [N- |
| High | High | 1 | |
| High | High | 1 | |

| | | | |
|---|---|---|---|
| High | High | 1 | |
| High | High | 1 | |
| High | High | 1 | Acetyl [N- |
| High | High | 1 | |
| High | High | 1 | |
| High | High | 1 | |
| High | High | 1 | |
| High | High | 1 | |
| High | High | 1 | |
| High | High | 1 | |
| High | High | 1 | |
| High | High | 1 | |
| High | High | 1 | |
| High | High | 1 | |
| High | High | 1 | |
| High | High | 1 | |
| High | High | 1 | |
| High | High | 1 | |
| High | High | 1 | Acetyl [N- |
| High | High | 1 | |
| High | High | 1 | |
| High | High | 1 | |
| High | High | 1 | |
| High | High | 1 | |
| High | High | 1 | Acetyl [N- |
| High | High | 1 | |
| High | High | 1 | |
| High | High | 1 | |
| High | High | 1 | |
| High | High | 1 | |
| High | High | 1 | |

| | | | |
|---|---|---|---|
| High | High | 1 | |
| High | High | 1 | |
| High | High | 1 | |
| High | High | 1 | |
| High | High | 1 | |
| High | High | 1 | |
| High | High | 1 | |
| High | High | 1 | |
| High | High | 1 | |
| High | High | 1 | |
| High | High | 1 | |
| High | High | 1 | |
| High | High | 1 | |
| High | High | 1 | |
| High | High | 1 | |
| High | High | 1 | |
| High | High | 1 | |
| High | High | 1 | |
| High | High | 1 | |
| High | High | 1 | |
| High | High | 1 | |
| High | High | 1 | |
| High | High | 1 | |
| High | High | 1 | |
| High | High | 1 | Acetyl [N- |
| High | High | 1 | |
| High | High | 1 | |
| High | High | 1 | |
| High | High | 1 | |
| High | High | 1 | |
| High | High | 1 | |

| | | |
|---|---|---|
| High | High | 1 |
| Not Founc | Not Founc | 1 |
| High | High | 1 |
| High | High | 1 |
| High | High | 1 |
| High | High | 1 |
| High | High | 1 |
| High | High | 1 |
| High | High | 1 |
| High | High | 1 |
| High | High | 1 |
| High | High | 1 |
| High | High | 1 |
| High | High | 1 |
| High | High | 1 |
| High | High | 1 |
| High | High | 1 |
| High | High | 1 |
| High | High | 1 |
| High | High | 1 |
| High | High | 1 |
| Not Founc | Not Founc | 1 |
| High | High | 1 |
| High | High | 1 |
| High | High | 1 |
| High | High | 1 |
| High | High | 1 |
| High | High | 1 |
| High | High | 1 |
| High | High | 1 |
| High | High | 1 |

| | | | |
|---|---|---|---|
| High | High | 1 | |
| High | High | 1 | |
| High | High | 1 | |
| High | High | 1 | |
| High | High | 1 | |
| High | High | 1 | |
| High | High | 1 | Acetyl [N- |
| High | High | 1 | |
| High | High | 1 | |
| High | High | 1 | Acetyl [N- |
| High | High | 1 | |
| High | High | 1 | |
| High | High | 1 | |
| High | High | 1 | |
| High | High | 1 | |
| High | High | 1 | |
| High | High | 1 | |
| High | High | 1 | |
| High | High | 1 | |
| High | High | 1 | |
| High | High | 1 | |
| High | High | 1 | |
| High | High | 1 | |
| High | High | 1 | |
| High | High | 1 | |
| High | High | 1 | |
| High | High | 1 | |
| High | High | 1 | |
| High | High | 1 | |
| High | High | 1 | |
| High | High | 1 | |

| | | |
|---|---|---|
| High | High | 1 |
| High | High | 1 |
| High | High | 1 |
| High | High | 1 |
| High | High | 1 |
| High | High | 1 |
| High | High | 1 |
| High | High | 1 |
| High | High | 1 |
| High | High | 1 |
| High | High | 1 |
| High | High | 1 |
| Not Founc | Not Founc | 1 |
| High | High | 1 |
| High | High | 1 |
| High | High | 1 |
| High | High | 1 |
| High | High | 1 |
| High | High | 1 |
| High | High | 1 |
| High | High | 1 |
| High | High | 1 |
| High | High | 1 |
| High | High | 1 |
| High | High | 1 |
| High | High | 1 |
| High | High | 1 |
| High | High | 1 |
| High | High | 1 |
| High | High | 1 |
| High | High | 1 |

| | | | |
|---|---|---|---|
| High | High | 1 | |
| High | High | 1 | |
| High | High | 1 | |
| High | High | 1 | |
| High | High | 1 | |
| High | High | 1 | |
| High | High | 1 | |
| High | High | 1 | |
| Not Founc | Not Founc | 1 | |
| High | High | 1 | Acetyl [N- |
| High | High | 1 | |
| High | High | 1 | |
| High | High | 1 | |
| High | High | 1 | |
| High | High | 1 | |
| High | High | 1 | |
| High | High | 1 | |
| High | High | 1 | |
| High | High | 1 | |
| High | High | 1 | |
| High | High | 1 | |
| High | High | 1 | |
| High | High | 1 | |
| High | High | 1 | |
| High | High | 1 | |
| High | High | 1 | |
| High | High | 1 | |
| High | High | 1 | |
| High | High | 1 | |
| High | High | 1 | |
| High | High | 1 | |

| | | | |
|---|---|---|---|
| High | High | 1 | |
| High | High | 1 | |
| High | High | 1 | |
| High | High | 1 | |
| Not Founc | Not Founc | 1 | |
| High | High | 1 | |
| High | High | 1 | |
| High | High | 1 | |
| High | High | 1 | |
| High | High | 1 | |
| High | High | 1 | |
| High | High | 1 | |
| High | High | 1 | |
| High | High | 1 | |
| High | High | 1 | |
| High | High | 1 | |
| High | High | 1 | |
| High | High | 1 | |
| High | High | 1 | |
| High | High | 1 | |
| High | High | 1 | |
| High | High | 1 | |
| High | High | 1 | |
| High | High | 1 | Acetyl [N- |
| High | High | 1 | |
| High | High | 1 | |
| High | High | 1 | |
| High | High | 1 | Acetyl [N- |
| High | High | 1 | |
| Not Founc | Not Founc | 1 | |
| High | High | 1 | |

| | | | |
|---|---|---|---|
| High | High | 1 | |
| High | High | 1 | |
| High | High | 1 | |
| High | High | 1 | Acetyl [N- |
| High | High | 1 | |
| Not Founc | Not Founc | 1 | |
| High | High | 1 | |
| High | High | 1 | |
| High | High | 1 | |
| High | High | 1 | |
| High | High | 1 | |
| High | High | 1 | |
| High | High | 1 | |
| High | High | 1 | |
| High | High | 1 | |
| High | High | 1 | |
| High | High | 1 | |
| High | High | 1 | |
| High | High | 1 | |
| High | High | 1 | |
| High | High | 1 | |
| High | High | 1 | |
| High | High | 1 | |
| High | High | 1 | |
| High | High | 1 | |
| High | High | 1 | |
| High | High | 1 | |
| High | High | 1 | |
| High | High | 1 | |
| High | High | 1 | |
| High | High | 1 | |
| High | High | 1 | |

| | | |
|---|---|---|
| High | High | 1 |
| High | High | 1 |
| High | High | 1 |
| High | High | 1 |
| High | High | 1 |
| High | High | 1 |
| High | High | 1 |
| High | High | 1 |
| High | High | 1 |
| High | High | 1 |
| High | High | 1 |
| High | High | 1 |
| High | High | 1 |
| High | High | 1 |
| High | High | 1 |
| High | High | 1 |
| High | High | 1 |
| High | High | 1 |
| High | High | 1 |
| High | High | 1 |
| High | High | 1 |
| High | High | 1 |
| High | High | 1 |
| High | High | 1 |
| High | High | 1 |
| High | High | 1 |
| High | High | 1 |
| High | High | 1 |
| High | High | 1 |
| High | High | 1 |
| High | High | 1 |

| | | |
|---|---|---|
| Not Founc | Not Founc | 1 |
| Not Founc | Not Founc | 1 |
| High | High | 1 |
| High | High | 1 |
| High | High | 1 |
| High | High | 1 |
| High | High | 1 |
| High | High | 1 |
| High | High | 1 |
| High | High | 1 |
| High | High | 1 |
| High | High | 1 |
| High | High | 1 |
| High | High | 1 |
| High | High | 1 |
| High | High | 1 |
| High | High | 1 |
| High | High | 1 |
| High | High | 1 |
| High | High | 1 |
| High | High | 1 |
| High | High | 1 |
| High | High | 1 |
| High | High | 1 |
| High | High | 1 |
| High | High | 1 |
| High | High | 1 |
| High | High | 1 |
| High | High | 1 |
| High | High | 1 |
| High | High | 1 |

| | | |
|---|---|---|
| Not Found | Not Found | 1 |
| Not Found | Not Found | 1 |
| High | High | 1 |
| High | High | 1 |
| Not Found | Not Found | 1 |
| High | High | 1 |
| High | High | 1 |
| High | High | 1 |
| High | High | 1 |
| High | High | 1 |
| High | High | 1 |
| High | High | 1 |
| High | High | 1 |
| High | High | 1 |
| High | High | 1 |
| High | High | 1 |
| High | High | 1 |
| High | High | 1 |
| Not Found | Not Found | 1 |
| High | High | 1 |
| High | High | 1 |
| High | High | 1 |
| High | High | 1 |
| High | High | 1 |
| High | High | 1 |
| High | High | 1 |
| High | High | 1 |
| High | High | 1 |
| High | High | 1 |
| High | High | 1 |
| High | High | 1 |

| | | | |
|---|---|---|---|
| High | High | 1 | |
| High | High | 1 | |
| High | High | 1 | |
| High | High | 1 | |
| Not Founc | Not Founc | 1 | |
| High | High | 1 | |
| High | High | 1 | |
| High | High | 1 | |
| High | High | 1 | |
| High | High | 1 | |
| High | High | 1 | |
| High | High | 1 | |
| High | High | 1 | |
| High | High | 1 | |
| High | High | 1 | |
| High | High | 1 | |
| High | High | 1 | |
| High | High | 1 | Acetyl [N- |
| High | High | 1 | |
| High | High | 1 | |
| High | High | 1 | |
| High | High | 1 | |
| High | High | 1 | |
| High | High | 1 | |
| High | High | 1 | |
| Not Founc | Not Founc | 1 | |
| High | High | 1 | |
| High | High | 1 | |
| High | High | 1 | |
| High | High | 1 | |
| High | High | 1 | |

| | | |
|---|---|---|
| High | High | 1 |
| Not Found | Not Found | 1 |
| High | High | 1 |
| High | High | 1 |
| High | High | 1 |
| High | High | 1 |
| High | High | 1 |
| High | High | 1 |
| High | High | 1 |
| High | High | 1 |
| High | High | 1 |
| High | High | 1 |
| High | High | 1 |
| High | High | 1 |
| High | High | 1 |
| High | High | 1 |
| High | High | 1 |
| High | High | 1 |
| High | High | 1 |
| High | High | 1 |
| High | High | 1 |
| High | High | 1 |
| High | High | 1 |
| High | High | 1 |
| High | High | 1 |
| High | High | 1 |
| High | High | 1 |
| High | High | 1 |
| Not Found | Not Found | 1 |
| High | High | 1 |
| High | High | 1 |

| | | |
|---|---|---|
| High | High | 1 |
| High | High | 1 |
| High | High | 1 |
| High | High | 1 |
| High | High | 1 |
| High | High | 1 |
| High | High | 1 |
| High | High | 1 |
| High | High | 1 |
| High | High | 1 |
| High | High | 1 |
| High | High | 1 |
| High | High | 1 |
| High | High | 1 |
| High | High | 1 |
| High | High | 1 |
| High | High | 1 |
| High | High | 1 |
| Not Founc | Not Founc | 1 |
| High | High | 1 |
| High | High | 1 |
| High | High | 1 |
| High | High | 1 |
| High | High | 1 |
| High | High | 1 |
| Not Founc | Not Founc | 1 |
| Not Founc | Not Founc | 1 |
| High | High | 1 |
| High | High | 1 |
| High | High | 1 |
| High | High | 1 |

| | | | |
|---|---|---|---|
| High | High | 1 | |
| High | High | 1 | |
| High | High | 1 | |
| High | High | 1 | |
| High | High | 1 | |
| High | High | 1 | |
| High | High | 1 | |
| High | High | 1 | |
| High | High | 1 | |
| Not Founc | Not Founc | 1 | |
| Not Founc | Not Founc | 1 | |
| High | High | 1 | |
| High | High | 1 | |
| High | High | 1 | |
| High | High | 1 | |
| High | High | 1 | |
| High | High | 1 | |
| Not Founc | Not Founc | 1 | |
| High | High | 1 | |
| High | High | 1 | |
| High | High | 1 | Acetyl [N- |
| High | High | 1 | |
| High | High | 1 | |
| High | High | 1 | Acetyl [N- |
| High | High | 1 | |
| High | High | 1 | |
| High | High | 1 | |
| High | High | 1 | |
| High | High | 1 | |
| High | High | 1 | |
| High | High | 1 | |

| | | | |
|---|---|---|---|
| High | High | 1 | |
| High | High | 1 | |
| High | High | 1 | |
| High | High | 1 | |
| High | High | 1 | |
| High | High | 1 | |
| High | High | 1 | |
| High | High | 1 | |
| High | High | 1 | |
| High | High | 1 | |
| Not Founc | Not Founc | 1 | |
| High | High | 1 | |
| High | High | 1 | |
| High | High | 1 | |
| High | High | 1 | |
| High | High | 1 | |
| High | High | 1 | |
| Not Founc | Not Founc | 1 | |
| High | High | 1 | |
| High | High | 1 | |
| High | High | 1 | |
| High | High | 1 | |
| High | High | 1 | Acetyl [N- |
| High | High | 1 | |
| High | High | 1 | |
| High | High | 1 | |
| High | High | 1 | |
| High | High | 1 | |
| High | High | 1 | |
| High | High | 1 | |
| High | High | 1 | |

| | | | |
|---|---|---|---|
| High | High | 1 | |
| High | High | 1 | |
| High | High | 1 | |
| High | High | 1 | |
| High | High | 1 | |
| High | High | 1 | |
| High | High | 1 | |
| High | High | 1 | |
| High | High | 1 | |
| High | High | 1 | |
| High | High | 1 | |
| Not Found | Not Found | 1 | |
| High | High | 1 | |
| High | High | 1 | |
| High | High | 1 | |
| Not Found | Not Found | 1 | |
| High | High | 1 | Acetyl [N- |
| High | High | 1 | |
| High | High | 1 | |
| High | High | 1 | |
| High | High | 1 | |
| High | High | 1 | |
| High | High | 1 | |
| High | High | 1 | |
| High | High | 1 | |
| High | High | 1 | |
| High | High | 1 | |
| High | High | 1 | |
| High | High | 1 | |
| High | High | 1 | |
| High | High | 1 | |

| | | |
|---|---|---|
| Not Founc | Not Founc | 1 |
| High | High | 1 |
| High | High | 1 |
| High | High | 1 |
| High | High | 1 |
| Not Founc | Not Founc | 1 |
| High | High | 1 |
| High | High | 1 |
| High | High | 1 |
| High | High | 1 |
| Not Founc | Not Founc | 1 |
| High | High | 1 |
| High | High | 1 |
| High | High | 1 |
| High | High | 1 |
| High | High | 1 |
| High | High | 1 |
| High | High | 1 |
| High | High | 1 |
| High | High | 1 |
| Not Founc | Not Founc | 1 |
| High | High | 1 |
| High | High | 1 |
| High | High | 1 |
| High | High | 1 |
| High | High | 1 |
| High | High | 1 |
| High | High | 1 |
| High | High | 1 |
| High | High | 1 |
| High | High | 1 |

| | | |
|---|---|---|
| High | High | 1 |
| High | High | 1 |
| High | High | 1 |
| High | High | 1 |
| High | High | 1 |
| High | High | 1 |
| High | High | 1 |
| High | High | 1 |
| High | High | 1 |
| High | High | 1 |
| High | High | 1 |
| High | High | 1 |
| High | High | 1 |
| High | High | 1 |
| High | High | 1 |
| High | High | 1 |
| High | High | 1 |
| Not Founc | Not Founc | 1 |
| Not Founc | Not Founc | 1 |
| High | High | 1 |
| High | High | 1 |
| High | High | 1 |
| High | High | 1 |
| High | High | 1 |
| High | High | 1 |
| Not Founc | Not Founc | 1 |
| High | High | 1 |
| High | High | 1 |
| High | High | 1 |
| High | High | 1 |
| High | High | 1 |

| | | |
|---|---|---|
| Not Found | Not Found | 1 |
| High | High | 1 |
| High | High | 1 |
| High | High | 1 |
| High | High | 1 |
| High | High | 1 |
| High | High | 1 |
| High | High | 1 |
| High | High | 1 |
| High | High | 1 |
| High | High | 1 |
| High | High | 1 |
| High | High | 1 |
| High | High | 1 |
| High | High | 1 |
| Not Found | Not Found | 1 |
| High | High | 1 |
| High | High | 1 |
| High | High | 1 |
| High | High | 1 |
| High | High | 1 |
| High | High | 1 |
| Not Found | Not Found | 1 |
| High | High | 1 |
| High | High | 1 |
| High | High | 1 |
| Not Found | Not Found | 1 |
| High | High | 1 |
| High | High | 1 |
| High | High | 1 |
| High | High | 1 |

| | | |
|---|---|---|
| High | High | 1 |
| High | High | 1 |
| High | High | 1 |
| Not Founc | Not Founc | 1 |
| High | High | 1 |
| High | High | 1 |
| High | High | 1 |
| High | High | 1 |
| High | High | 1 |
| Not Founc | Not Founc | 1 |
| High | High | 1 |
| High | High | 1 |
| High | High | 1 |
| High | High | 1 |
| High | High | 1 |
| High | High | 1 |
| High | High | 1 |
| High | High | 1 |
| High | High | 1 |
| Not Founc | Not Founc | 1 |
| High | High | 1 |
| High | High | 1 |
| High | High | 1 |
| High | High | 1 |
| High | High | 1 |
| High | High | 1 |
| High | High | 1 |
| High | High | 1 |
| Not Founc | Not Founc | 1 |
| High | High | 1 |
| High | High | 1 |

| | | |
|---|---|---|
| High | High | 1 |
| High | High | 1 |
| High | High | 1 |
| Not Found | Not Found | 1 |
| High | High | 1 |
| High | High | 1 |
| High | High | 1 |
| High | High | 1 |
| High | High | 1 |
| Not Found | Not Found | 1 |
| High | High | 1 |
| High | High | 1 |
| High | High | 1 |
| High | High | 1 |
| High | High | 1 |
| High | High | 1 |
| High | High | 1 |
| Not Found | Not Found | 1 |
| High | High | 1 |
| High | High | 1 |
| High | High | 1 |
| High | High | 1 |
| High | High | 1 |
| High | High | 1 |
| Not Found | Not Found | 1 |
| High | High | 1 |
| High | High | 1 |
| High | High | 1 |
| High | High | 1 |
| Not Found | Not Found | 1 |
| High | High | 1 |

| | | | |
|---|---|---|---|
| Not Founc | Not Founc | 1 | |
| High | High | 1 | |
| High | High | 1 | |
| Not Founc | Not Founc | 1 | |
| Not Founc | Not Founc | 1 | |
| High | High | 1 | |
| High | High | 1 | |
| Not Founc | Not Founc | 1 | |
| High | High | 1 | |
| High | High | 1 | |
| High | High | 1 | |
| High | High | 1 | Acetyl [N- |
| High | High | 1 | |
| High | High | 1 | |
| High | High | 1 | |
| High | High | 1 | |
| Not Founc | Not Founc | 1 | |
| High | High | 1 | |
| High | High | 1 | Acetyl [N- |
| Not Founc | Not Founc | 1 | |
| Not Founc | Not Founc | 1 | |
| High | High | 1 | |
| High | High | 1 | |
| High | High | 1 | |
| Not Founc | Not Founc | 1 | |
| High | High | 1 | |
| High | High | 1 | |
| High | High | 1 | |
| High | High | 1 | |
| High | High | 1 | |
| Not Founc | Not Founc | 1 | |

| | | | |
|---|---|---|---|
| High | High | 1 | |
| High | High | 1 | |
| High | High | 1 | |
| High | High | 1 | |
| High | High | 1 | |
| Not Founc | Not Founc | 1 | |
| Not Founc | Not Founc | 1 | |
| High | High | 1 | |
| High | High | 1 | |
| High | High | 1 | |
| High | High | 1 | |
| High | High | 1 | |
| High | High | 1 | |
| High | High | 1 | Acetyl [N- |
| High | High | 1 | |
| High | High | 1 | |
| Not Founc | Not Founc | 1 | |
| High | High | 1 | |
| High | High | 1 | |
| High | High | 1 | |
| High | High | 1 | |
| High | High | 1 | |
| High | High | 1 | |
| High | High | 1 | |
| High | High | 1 | |
| Not Founc | Not Founc | 1 | |
| High | High | 1 | |
| Not Founc | Not Founc | 1 | |
| High | High | 1 | |
| High | High | 1 | |
| High | High | 1 | |

| | | |
|---|---|---|
| High | High | 1 |
| Not Founc | Not Founc | 1 |
| High | High | 1 |
| High | High | 1 |
| Not Founc | Not Founc | 1 |
| High | High | 1 |
| Not Founc | Not Founc | 1 |
| Not Founc | Not Founc | 1 |
| High | High | 1 |
| High | High | 1 |
| High | High | 1 |
| High | High | 1 |
| High | High | 1 |
| High | High | 1 |
| High | High | 1 |
| High | High | 1 |
| High | High | 1 |
| High | High | 1 |
| High | High | 1 |
| Not Founc | Not Founc | 1 |
| High | High | 1 |
| High | High | 1 |
| High | High | 1 |
| High | High | 1 |
| High | High | 1 |
| High | High | 1 |
| High | High | 1 |
| Not Founc | Not Founc | 1 |
| High | High | 1 |
| High | High | 1 |
| High | High | 1 |

| | | | |
|---|---|---|---|
| Not Founc | Not Founc | 1 | |
| High | High | 1 | |
| Not Founc | Not Founc | 1 | |
| High | High | 1 | |
| High | High | 1 | |
| High | High | 1 | |
| Not Founc | Not Founc | 1 | |
| High | High | 1 | |
| High | High | 1 | |
| Not Founc | Not Founc | 1 | |
| High | High | 1 | |
| High | High | 1 | |
| High | High | 1 | |
| High | High | 1 | |
| High | High | 1 | |
| High | High | 1 | |
| High | High | 1 | |
| High | High | 1 | |
| High | High | 1 | |
| Not Founc | Not Founc | 1 | |
| High | High | 1 | |
| High | High | 1 | |
| Not Founc | Not Founc | 1 | |
| High | High | 1 | |
| High | High | 1 | Acetyl [N- |
| Not Founc | Not Founc | 1 | |
| Not Founc | Not Founc | 1 | |
| Not Founc | Not Founc | 1 | |
| High | High | 1 | |
| High | High | 1 | |
| High | High | 1 | |

| | | |
|---|---|---|
| High | High | 1 |
| Not Found | Not Found | 1 |
| Not Found | Not Found | 1 |
| High | High | 1 |
| Not Found | Not Found | 1 |
| High | High | 1 |
| High | High | 1 |
| High | High | 1 |
| High | High | 1 |
| High | High | 1 |
| Not Found | Not Found | 1 |
| High | High | 1 |
| High | High | 1 |
| High | High | 1 |
| High | High | 1 |
| High | High | 1 |
| High | High | 1 |
| High | High | 1 |
| High | High | 1 |
| Not Found | Not Found | 1 |
| High | High | 1 |
| High | High | 1 |
| Not Found | Not Found | 1 |
| High | High | 1 |
| High | High | 1 |
| High | High | 1 |
| High | High | 1 |
| High | High | 1 |
| High | High | 1 |
| High | High | 1 |
| Not Found | Not Found | 1 |

| | | | |
|---|---|---|---|
| High | High | 1 | |
| High | High | 1 | |
| High | High | 1 | |
| High | High | 1 | |
| High | High | 1 | |
| High | High | 1 | |
| High | High | 1 | |
| Not Founc | Not Founc | 1 | |
| Not Founc | Not Founc | 1 | |
| High | High | 1 | |
| High | High | 1 | |
| High | High | 1 | |
| High | High | 1 | |
| High | High | 1 | |
| Not Founc | Not Founc | 1 | Acetyl [N- |
| Not Founc | Not Founc | 1 | |
| High | High | 1 | |
| High | High | 1 | |
| High | High | 1 | |
| High | High | 1 | |
| Not Founc | Not Founc | 1 | |
| High | High | 1 | |
| High | High | 1 | |
| Not Founc | Not Founc | 1 | |
| High | High | 1 | |
| High | High | 1 | |
| High | High | 1 | |
| High | High | 1 | |
| High | High | 1 | |
| High | High | 1 | |
| High | High | 1 | |

| | | | |
|---|---|---|---|
| High | High | 1 | |
| High | High | 1 | |
| High | High | 1 | |
| Not Found | Not Found | 1 | |
| High | High | 1 | |
| High | High | 1 | |
| High | High | 1 | |
| Not Found | Not Found | 1 | |
| High | High | 1 | |
| High | High | 1 | |
| Not Found | Not Found | 1 | Acetyl [N- |
| High | High | 1 | |
| Not Found | Not Found | 1 | |
| High | High | 1 | |
| High | High | 1 | |
| High | High | 1 | |
| High | High | 1 | |
| Not Found | Not Found | 1 | |
| Not Found | Not Found | 1 | |
| High | High | 1 | |
| High | High | 1 | |
| Not Found | Not Found | 1 | |
| High | High | 1 | |
| High | High | 1 | |
| Not Found | Not Found | 1 | |
| Not Found | Not Found | 1 | |
| High | High | 1 | |
| Not Found | Not Found | 1 | |
| High | High | 1 | |
| Not Found | Not Found | 1 | |
| High | High | 1 | |

| | | | |
|---|---|---|---|
| Not Found | Not Found | 1 | |
| High | High | 1 | |
| Not Found | Not Found | 1 | |
| High | High | 1 | |
| High | High | 1 | |
| High | High | 1 | |
| High | High | 1 | |
| High | High | 1 | |
| High | High | 1 | |
| High | High | 1 | |
| High | High | 1 | |
| High | High | 1 | |
| High | High | 1 | |
| High | High | 1 | |
| High | High | 1 | |
| High | High | 1 | |
| High | High | 1 | |
| High | High | 1 | |
| High | High | 1 | |
| High | High | 1 | Acetyl [N- |
| Not Found | Not Found | 1 | |
| High | High | 1 | |
| Not Found | Not Found | 1 | |
| High | High | 1 | |
| High | High | 1 | |
| Not Found | Not Found | 1 | |
| High | High | 1 | |
| High | High | 1 | |
| High | High | 1 | |
| High | High | 1 | |
| High | High | 1 | |

| | | |
|---|---|---|
| High | High | 1 |
| Not Founc | Not Founc | 1 |
| High | High | 1 |
| High | High | 1 |
| Not Founc | Not Founc | 1 |
| Not Founc | Not Founc | 1 |
| Not Founc | Not Founc | 1 |
| High | High | 1 |
| High | High | 1 |
| High | High | 1 |
| High | High | 1 |
| High | High | 1 |
| High | High | 1 |
| High | High | 1 |
| High | High | 1 |
| High | High | 1 |
| High | High | 1 |
| High | High | 1 |
| High | High | 1 |
| High | High | 1 |
| High | High | 1 |
| High | High | 1 |
| High | High | 1 |
| High | High | 1 |
| High | High | 1 |
| Not Founc | Not Founc | 1 |
| High | High | 1 |
| High | High | 1 |
| High | High | 1 |
| Not Founc | Not Founc | 1 |
| High | High | 1 |

| | | |
|---|---|---|
| Not Founc | Not Founc | 1 |
| High | High | 1 |
| High | High | 1 |
| High | High | 1 |
| High | High | 1 |
| Not Founc | Not Founc | 1 |
| High | High | 1 |
| High | High | 1 |
| High | High | 1 |
| Not Founc | Not Founc | 1 |
| High | High | 1 |
| Not Founc | Not Founc | 1 |
| Not Founc | Not Founc | 1 |
| High | High | 1 |
| High | High | 1 |
| High | High | 1 |
| High | High | 1 |
| High | High | 1 |
| High | High | 1 |
| Not Founc | Not Founc | 1 |
| Not Founc | Not Founc | 1 |
| High | High | 1 |
| High | High | 1 |
| High | High | 1 |
| High | High | 1 |
| Not Founc | Not Founc | 1 |
| High | High | 1 |
| High | High | 1 |
| High | High | 1 |
| Not Founc | Not Founc | 1 |
| High | High | 1 |

| | | |
|---|---|---|
| High | High | 1 |
| Not Founc | Not Founc | 1 |
| High | High | 1 |
| High | High | 1 |
| High | High | 1 |
| High | High | 1 |
| High | High | 1 |
| Not Founc | Not Founc | 1 |
| High | High | 1 |
| Not Founc | Not Founc | 1 |
| Not Founc | Not Founc | 1 |
| High | High | 1 |
| High | High | 1 |
| Not Founc | Not Founc | 1 |
| Not Founc | Not Founc | 1 |
| High | High | 1 |
| Not Founc | Not Founc | 1 |
| High | High | 1 |
| High | High | 1 |
| High | High | 1 |
| Not Founc | Not Founc | 1 |
| High | High | 1 |
| High | High | 1 |
| High | High | 1 |
| High | High | 1 |
| High | High | 1 |
| Not Founc | Not Founc | 1 |
| High | High | 1 |
| Not Founc | Not Founc | 1 |
| High | High | 1 |
| Not Founc | Not Founc | 1 |

| | | |
|---|---|---|
| High | High | 1 |
| Not Founc | Not Founc | 1 |
| High | High | 1 |
| High | High | 1 |
| High | High | 1 |
| High | High | 1 |
| High | High | 1 |
| High | High | 1 |
| Not Founc | Not Founc | 1 |
| High | High | 1 |
| Not Founc | Not Founc | 1 |
| High | High | 1 |
| High | High | 1 |
| Not Founc | Not Founc | 1 |
| High | High | 1 |
| Not Founc | Not Founc | 1 |
| Not Founc | Not Founc | 1 |
| High | High | 1 |
| High | High | 1 |
| High | High | 1 |
| High | High | 1 |
| Not Founc | Not Founc | 1 |
| High | High | 1 |
| High | High | 1 |
| High | High | 1 |
| High | High | 1 |
| Not Founc | Not Founc | 1 |
| Not Founc | Not Founc | 1 |
| Not Founc | Not Founc | 1 |
| Not Founc | Not Founc | 1 |
| High | High | 1 |

| | | |
|---|---|---|
| High | High | 1 |
| High | High | 1 |
| Not Found | Not Found | 1 |
| High | High | 1 |
| High | High | 1 |
| Not Found | Not Found | 1 |
| Not Found | Not Found | 1 |
| High | High | 1 |
| Not Found | Not Found | 1 |
| High | High | 1 |
| High | High | 1 |
| High | High | 1 |
| High | High | 1 |
| High | High | 1 |
| High | High | 1 |
| High | High | 1 |
| Not Found | Not Found | 1 |
| High | High | 1 |
| High | High | 1 |
| High | High | 1 |
| Not Found | Not Found | 1 |
| High | High | 1 |
| High | High | 1 |
| Not Found | Not Found | 1 |
| High | High | 1 |
| Not Found | Not Found | 1 |
| High | High | 1 |
| Not Found | Not Found | 1 |
| Not Found | Not Found | 1 |
| High | High | 1 |
| High | High | 1 |

| | | |
|---|---|---|
| High | High | 1 |
| High | High | 1 |
| High | High | 1 |
| Not Founc | Not Founc | 1 |
| Not Founc | Not Founc | 1 |
| High | High | 1 |
| High | High | 1 |
| Not Founc | Not Founc | 1 |
| High | High | 1 |
| High | High | 1 |
| Not Founc | Not Founc | 1 |
| High | High | 1 |
| High | High | 1 |
| High | High | 1 |
| High | High | 1 |
| Not Founc | Not Founc | 1 |
| High | High | 1 |
| Not Founc | Not Founc | 1 |
| High | High | 1 |
| High | High | 1 |
| Not Founc | Not Founc | 1 |
| High | High | 1 |
| High | High | 1 |
| High | High | 1 |
| High | High | 1 |
| High | High | 1 |
| High | High | 1 |
| High | High | 1 |
| Not Founc | Not Founc | 1 |
| High | High | 1 |
| High | High | 1 |

| | | |
|---|---|---|
| High | High | 1 |
| Not Founc | Not Founc | 1 |
| High | High | 1 |
| Not Founc | Not Founc | 1 |
| High | High | 1 |
| Not Founc | Not Founc | 1 |
| High | High | 1 |
| High | High | 1 |
| Not Founc | Not Founc | 1 |
| High | High | 1 |
| High | High | 1 |
| High | High | 1 |
| Not Founc | Not Founc | 1 |
| High | High | 1 |
| High | High | 1 |
| High | High | 1 |
| High | High | 1 |
| High | High | 1 |
| High | High | 1 |
| Not Founc | Not Founc | 1 |
| High | High | 1 |
| High | High | 1 |
| Not Founc | Not Founc | 1 |
| High | High | 1 |
| High | High | 1 |
| Not Founc | Not Founc | 1 |
| High | High | 1 |
| High | High | 1 |
| Not Founc | Not Founc | 1 |
| Not Founc | Not Founc | 1 |
| High | High | 1 |

| | | |
|---|---|---|
| High | High | 1 |
| Not Founc | Not Founc | 1 |
| High | High | 1 |
| Not Founc | Not Founc | 1 |
| Not Founc | Not Founc | 1 |
| Not Founc | Not Founc | 1 |
| Not Founc | Not Founc | 1 |
| Not Founc | Not Founc | 1 |
| High | High | 1 |
| Not Founc | Not Founc | 1 |
| High | High | 1 |
| High | High | 1 |
| Not Founc | Not Founc | 1 |
| High | High | 1 |
| High | High | 1 |
| High | High | 1 |
| Not Founc | Not Founc | 1 |
| High | High | 1 |
| High | High | 1 |
| High | High | 1 |
| Not Founc | Not Founc | 1 |
| Not Founc | Not Founc | 1 |
| High | High | 1 |
| High | High | 1 |
| High | High | 1 |
| Not Founc | Not Founc | 1 |
| Not Founc | Not Founc | 1 |
| Not Founc | Not Founc | 1 |
| High | High | 1 |
| High | High | 1 |
| High | High | 1 |

| | | |
|---|---|---|
| High | High | 1 |
| Not Founc | Not Founc | 1 |
| High | High | 1 |
| High | High | 1 |
| High | High | 1 |
| Not Founc | Not Founc | 1 |
| Not Founc | Not Founc | 1 |
| High | High | 1 |
| High | High | 1 |
| Not Founc | Not Founc | 1 |
| Not Founc | Not Founc | 1 |
| Not Founc | Not Founc | 1 |
| Not Founc | Not Founc | 1 |
| Not Founc | Not Founc | 1 |
| Not Founc | Not Founc | 1 |
| High | High | 1 |
| High | High | 1 |
| High | High | 1 |
| High | High | 1 |
| High | High | 1 |
| High | High | 1 |
| High | High | 1 |
| High | High | 1 |
| High | High | 1 |
| Not Founc | Not Founc | 1 |
| High | High | 1 |
| Not Founc | Not Founc | 1 |
| High | High | 1 |
| High | High | 1 |
| Not Founc | Not Founc | 1 |
| High | High | 1 |

| | | | |
|---|---|---|---|
| High | High | 1 | |
| High | High | 1 | |
| Not Founc | Not Founc | 1 | |
| High | High | 1 | |
| High | High | 1 | |
| High | High | 1 | |
| High | High | 1 | |
| Not Founc | Not Founc | 1 | |
| High | High | 1 | |
| Not Founc | Not Founc | 1 | |
| High | High | 1 | |
| High | High | 1 | |
| Not Founc | Not Founc | 1 | Acetyl [N- |
| Not Founc | Not Founc | 1 | |
| High | High | 1 | |
| High | High | 1 | |
| High | High | 1 | |
| Not Founc | Not Founc | 1 | |
| High | High | 1 | |
| High | High | 1 | |
| High | High | 1 | |
| High | High | 1 | |
| High | High | 1 | |
| Not Founc | Not Founc | 1 | |
| High | High | 1 | |
| Not Founc | Not Founc | 1 | |
| High | High | 1 | |
| High | High | 1 | |
| High | High | 1 | |
| Not Founc | Not Founc | 1 | |
| Not Founc | Not Founc | 1 | |

| | | | |
|---|---|---|---|
| Not Found | Not Found | 1 | |
| Not Found | Not Found | 1 | |
| Not Found | Not Found | 1 | |
| Not Found | Not Found | 1 | |
| High | High | 1 | |
| Not Found | Not Found | 1 | |
| Not Found | Not Found | 1 | |
| High | High | 1 | |
| High | High | 1 | Acetyl [N- |
| High | High | 1 | |
| Not Found | Not Found | 1 | |
| High | High | 1 | |
| High | High | 1 | |
| High | High | 1 | |
| High | High | 1 | |
| High | High | 1 | |
| High | High | 1 | |
| Not Found | Not Found | 1 | |
| Not Found | Not Found | 1 | |
| Not Found | Not Found | 1 | |
| Not Found | Not Found | 1 | |
| High | High | 1 | |
| High | High | 1 | |
| High | High | 1 | |
| High | High | 1 | |
| High | High | 1 | |
| High | High | 1 | Acetyl [N- |
| Not Found | Not Found | 1 | |
| High | High | 1 | |
| High | High | 1 | |
| Not Found | Not Found | 1 | |

| | | | |
|---|---|---|---|
| Not Founc | Not Founc | 1 | Acetyl [N- |
| Not Founc | Not Founc | 1 | |
| Not Founc | Not Founc | 1 | |
| Not Founc | Not Founc | 1 | |
| Not Founc | Not Founc | 1 | |
| Not Founc | Not Founc | 1 | |
| High | High | 1 | |
| High | High | 1 | Acetyl [N- |
| High | High | 1 | |
| High | High | 1 | |
| Not Founc | Not Founc | 1 | |
| High | High | 1 | |
| High | High | 1 | |
| Not Founc | Not Founc | 1 | |
| High | High | 1 | |
| Not Founc | Not Founc | 1 | |
| Not Founc | Not Founc | 1 | |
| High | High | 1 | |
| Not Founc | Not Founc | 1 | |
| High | High | 1 | |
| Not Founc | Not Founc | 1 | |
| Not Founc | Not Founc | 1 | |
| High | High | 1 | |
| High | High | 1 | |
| High | High | 1 | |
| High | High | 1 | |
| Not Founc | Not Founc | 1 | |
| High | High | 1 | |
| High | High | 1 | |
| Not Founc | Not Founc | 1 | |
| Not Founc | Not Founc | 1 | |

| | | |
|---|---|---|
| Not Founc | Not Founc | 1 |
| Not Founc | Not Founc | 1 |
| High | High | 1 |
| Not Founc | Not Founc | 1 |
| Not Founc | Not Founc | 1 |
| High | High | 1 |
| Not Founc | Not Founc | 1 |
| High | High | 1 |
| High | High | 1 |
| Not Founc | Not Founc | 1 |
| High | High | 1 |
| High | High | 1 |
| High | High | 1 |
| High | High | 1 |
| High | High | 1 |
| Not Founc | Not Founc | 1 |
| Not Founc | Not Founc | 1 |
| Not Founc | Not Founc | 1 |
| High | High | 1 |
| Not Founc | Not Founc | 1 |
| High | High | 1 |
| Not Founc | Not Founc | 1 |
| Not Founc | Not Founc | 1 |
| High | High | 1 |
| High | High | 1 |
| Not Founc | Not Founc | 1 |
| High | High | 1 |
| High | High | 1 |
| Not Founc | Not Founc | 1 |
| High | High | 1 |
| Not Founc | Not Founc | 1 |

| | | | |
|---|---|---|---|
| Not Founc | Not Founc | 1 | |
| Not Founc | Not Founc | 1 | |
| High | High | 1 | |
| High | High | 1 | |
| Not Founc | Not Founc | 1 | |
| High | High | 1 | |
| Not Founc | Not Founc | 1 | |
| High | High | 1 | |
| High | High | 1 | |
| High | High | 1 | |
| High | High | 1 | |
| Not Founc | Not Founc | 1 | |
| High | High | 1 | |
| Not Founc | Not Founc | 1 | |
| Not Founc | Not Founc | 1 | |
| High | High | 1 | |
| Not Founc | Not Founc | 1 | |
| Not Founc | Not Founc | 1 | |
| Not Founc | Not Founc | 1 | |
| Not Founc | Not Founc | 1 | |
| Not Founc | Not Founc | 1 | |
| High | High | 1 | |
| Not Founc | Not Founc | 1 | |
| High | High | 1 | |
| Not Founc | Not Founc | 1 | Acetyl [N- |
| High | High | 1 | |
| Not Founc | Not Founc | 1 | |
| Not Founc | Not Founc | 1 | |
| Not Founc | Not Founc | 1 | |
| Not Founc | Not Founc | 1 | |
| Not Founc | Not Founc | 1 | |

| | | | |
|---|---|---|---|
| High | High | 1 | |
| Not Found | Not Found | 1 | |
| Not Found | Not Found | 1 | |
| High | High | 1 | |
| High | High | 1 | |
| Not Found | Not Found | 1 | |
| High | High | 1 | Acetyl [N- |
| Not Found | Not Found | 1 | |
| High | High | 1 | |
| High | High | 1 | |
| Not Found | Not Found | 1 | |
| Not Found | Not Found | 1 | |
| High | High | 1 | |
| Not Found | Not Found | 1 | |
| High | High | 1 | |
| Not Found | Not Found | 1 | |
| High | High | 1 | |
| High | High | 1 | |
| Not Found | Not Found | 1 | |
| High | High | 1 | |
| Not Found | Not Found | 1 | |
| High | High | 1 | |
| High | High | 1 | |
| Not Found | Not Found | 1 | |
| High | High | 1 | |
| Not Found | Not Found | 1 | |
| High | High | 1 | |
| Not Found | Not Found | 1 | |
| Not Found | Not Found | 1 | |
| Not Found | Not Found | 1 | |
| Not Found | Not Found | 1 | |

| | | | |
|---|---|---|---|
| Not Founc | Not Founc | 1 | |
| High | High | 1 | |
| High | High | 1 | |
| High | High | 1 | |
| Not Founc | Not Founc | 1 | |
| Not Founc | Not Founc | 1 | |
| Not Founc | Not Founc | 1 | |
| High | High | 1 | |
| Not Founc | Not Founc | 1 | |
| Not Founc | Not Founc | 1 | |
| High | High | 1 | |
| Not Founc | Not Founc | 1 | |
| Not Founc | Not Founc | 1 | |
| High | High | 1 | |
| Not Founc | Not Founc | 1 | |
| Not Founc | Not Founc | 1 | |
| High | High | 1 | |
| Not Founc | Not Founc | 1 | |
| Not Founc | Not Founc | 1 | |
| High | High | 1 | |
| High | High | 1 | |
| High | High | 1 | Acetyl [N- |
| Not Founc | Not Founc | 1 | |
| Not Founc | Not Founc | 1 | |
| High | High | 1 | |
| High | High | 1 | |
| Not Founc | Not Founc | 1 | |
| Not Founc | Not Founc | 1 | |
| High | High | 1 | |
| High | High | 1 | |
| Not Founc | Not Founc | 1 | |

| | | |
|---|---|---|
| High | High | 1 |
| High | High | 1 |
| Not Found | Not Found | 1 |
| Not Found | Not Found | 1 |
| Not Found | Not Found | 1 |
| High | High | 1 |
| High | High | 1 |
| High | High | 1 |
| High | High | 1 |
| High | High | 1 |
| Not Found | Not Found | 1 |
| Not Found | Not Found | 1 |
| Not Found | Not Found | 1 |
| High | High | 1 |
| Not Found | Not Found | 1 |
| Not Found | Not Found | 1 |
| Not Found | Not Found | 1 |
| High | High | 1 |
| High | High | 1 |
| Not Found | Not Found | 1 |
| Not Found | Not Found | 1 |
| High | High | 1 |
| Not Found | Not Found | 1 |
| Not Found | Not Found | 1 |
| High | High | 1 |
| High | High | 1 |
| High | High | 1 |
| High | High | 1 |
| Not Found | Not Found | 1 |
| Not Found | Not Found | 1 |
| Not Found | Not Found | 1 |

| | | |
|---|---|---|
| High | High | 1 |
| Not Founc | Not Founc | 1 |
| Not Founc | Not Founc | 1 |
| High | High | 1 |
| Not Founc | Not Founc | 1 |
| High | High | 1 |
| High | High | 1 |
| High | High | 1 |
| Not Founc | Not Founc | 1 |
| High | High | 1 |
| Not Founc | Not Founc | 1 |
| Not Founc | Not Founc | 1 |
| Not Founc | Not Founc | 1 |
| Not Founc | Not Founc | 1 |
| Not Founc | Not Founc | 1 |
| Not Founc | Not Founc | 1 |
| Not Founc | Not Founc | 1 |
| Not Founc | Not Founc | 1 |
| High | High | 1 |
| Not Founc | Not Founc | 1 |
| Not Founc | Not Founc | 1 |
| High | High | 1 |
| Not Founc | Not Founc | 1 |
| High | High | 1 |
| High | High | 1 |
| High | High | 1 |
| High | High | 1 |
| Not Founc | Not Founc | 1 |
| Not Founc | Not Founc | 1 |
| Not Founc | Not Founc | 1 |
| High | High | 1 |

| | | |
|---|---|---|
| High | High | 1 |
| High | High | 1 |
| Not Founc | Not Founc | 1 |
| High | High | 1 |
| High | High | 1 |
| Not Founc | Not Founc | 1 |
| Not Founc | Not Founc | 1 |
| Not Founc | Not Founc | 1 |
| High | High | 1 |
| High | High | 1 |
| Not Founc | Not Founc | 1 |
| High | High | 1 |
| Not Founc | Not Founc | 1 |
| Not Founc | Not Founc | 1 |
| Not Founc | Not Founc | 1 |
| High | High | 1 |
| Not Founc | Not Founc | 1 |
| High | High | 1 |
| Not Founc | Not Founc | 1 |
| Not Founc | Not Founc | 1 |
| Not Founc | Not Founc | 1 |
| High | High | 1 |
| High | High | 1 |
| High | High | 1 |
| Not Founc | Not Founc | 1 |
| High | High | 1 |
| High | High | 1 |
| Not Founc | Not Founc | 1 |
| Not Founc | Not Founc | 1 |
| Not Founc | Not Founc | 1 |
| Not Founc | Not Founc | 1 |

| | | |
|---|---|---|
| Not Founc | Not Founc | 1 |
| High | High | 1 |
| Not Founc | Not Founc | 1 |
| High | High | 1 |
| Not Founc | Not Founc | 1 |
| Not Founc | Not Founc | 1 |
| Not Founc | Not Founc | 1 |
| High | High | 1 |
| Not Founc | Not Founc | 1 |
| Not Founc | Not Founc | 1 |
| Not Founc | Not Founc | 1 |
| High | High | 1 |
| Not Founc | Not Founc | 1 |
| High | High | 1 |
| Not Founc | Not Founc | 1 |
| High | High | 1 |
| Not Founc | Not Founc | 1 |
| Not Founc | Not Founc | 1 |
| Not Founc | Not Founc | 1 |
| Not Founc | Not Founc | 1 |
| Not Founc | Not Founc | 1 |
| Not Founc | Not Founc | 1 |
| Not Founc | Not Founc | 1 |
| High | High | 1 |
| Not Founc | Not Founc | 1 |
| Not Founc | Not Founc | 1 |
| Not Founc | Not Founc | 1 |
| Not Founc | Not Founc | 1 |
| High | High | 1 |
| Not Founc | Not Founc | 1 |
| High | High | 1 |

| | | |
|---|---|---|
| High | High | 1 |
| Not Found | Not Found | 1 |
| Not Found | Not Found | 1 |
| High | High | 1 |
| Not Found | Not Found | 1 |
| Not Found | Not Found | 1 |
| Not Found | Not Found | 1 |
| High | High | 1 |
| Not Found | Not Found | 1 |
| Not Found | Not Found | 1 |
| Not Found | Not Found | 1 |
| Not Found | Not Found | 1 |
| Not Found | Not Found | 1 |
| High | High | 1 |
| Not Found | Not Found | 1 |
| Not Found | Not Found | 1 |
| Not Found | Not Found | 1 |
| Not Found | Not Found | 1 |
| High | High | 1 |
| Not Found | Not Found | 1 |
| Not Found | Not Found | 1 |
| High | High | 1 |
| High | High | 1 |
| Not Found | Not Found | 1 |
| Not Found | Not Found | 1 |
| Not Found | Not Found | 1 |
| High | High | 1 |
| Not Found | Not Found | 1 |
| High | High | 1 |
| Not Found | Not Found | 1 |
| High | High | 1 |

| | | |
|---|---|---|
| High | High | 1 |
| Not Founc | Not Founc | 1 |
| High | High | 1 |
| High | High | 1 |
| Not Founc | Not Founc | 1 |
| High | High | 1 |
| Not Founc | Not Founc | 1 |
| Not Founc | Not Founc | 1 |
| Not Founc | Not Founc | 1 |
| High | High | 1 |
| Not Founc | Not Founc | 1 |
| Not Founc | Not Founc | 1 |
| Not Founc | Not Founc | 1 |
| Not Founc | Not Founc | 1 |
| Not Founc | Not Founc | 1 |
| Not Founc | Not Founc | 1 |
| Not Founc | Not Founc | 1 |
| High | High | 1 |
| High | High | 1 |
| High | High | 1 |
| Not Founc | Not Founc | 1 |
| Not Founc | Not Founc | 1 |
| Not Founc | Not Founc | 1 |
| Not Founc | Not Founc | 1 |
| Not Founc | Not Founc | 1 |
| Not Founc | Not Founc | 1 |
| Not Founc | Not Founc | 1 |
| Not Founc | Not Founc | 1 |
| High | High | 1 |
| Not Founc | Not Founc | 1 |
| High | High | 1 |

| | | |
|---|---|---|
| Not Found | Not Found | 1 |
| High | High | 1 |
| Not Found | Not Found | 1 |
| High | High | 1 |
| High | High | 1 |
| Not Found | Not Found | 1 |
| Not Found | Not Found | 1 |
| Not Found | Not Found | 1 |
| High | High | 1 |
| High | High | 1 |
| Not Found | Not Found | 1 |
| Not Found | Not Found | 1 |
| Not Found | Not Found | 1 |
| Not Found | Not Found | 1 |
| Not Found | Not Found | 1 |
| Not Found | Not Found | 1 |
| Not Found | Not Found | 1 |
| High | High | 1 |
| High | High | 1 |
| Not Found | Not Found | 1 |
| Not Found | Not Found | 1 |
| Not Found | Not Found | 1 |
| Not Found | Not Found | 1 |
| Not Found | Not Found | 1 |
| Not Found | Not Found | 1 |
| Not Found | Not Found | 1 |
| High | High | 1 |
| Not Found | Not Found | 1 |
| Not Found | Not Found | 1 |
| Not Found | Not Found | 1 |
| Not Found | Not Found | 1 |

| | | |
|---|---|---|
| Not Found | Not Found | 1 |
| Not Found | Not Found | 1 |
| Not Found | Not Found | 1 |
| Not Found | Not Found | 1 |
| Not Found | Not Found | 1 |
| Not Found | Not Found | 1 |
| Not Found | Not Found | 1 |
| Not Found | Not Found | 1 |
| High | High | 1 |
| Not Found | Not Found | 1 |
| Not Found | Not Found | 1 |
| Not Found | Not Found | 1 |
| Not Found | Not Found | 1 |
| Not Found | Not Found | 1 |
| Not Found | Not Found | 1 |
| High | High | 1 |
| Not Found | Not Found | 1 |
| Not Found | Not Found | 1 |
| Not Found | Not Found | 1 |
| Not Found | Not Found | 1 |
| Not Found | Not Found | 1 |
| Not Found | Not Found | 1 |
| Not Found | Not Found | 1 |
| Not Found | Not Found | 1 |
| High | High | 1 |
| High | High | 1 |
| High | High | 1 |
| High | High | 1 |
| Not Found | Not Found | 1 |
| Not Found | Not Found | 1 |
| High | High | 1 |

| | | |
|---|---|---|
| Not Founc | Not Founc | 1 |
| Not Founc | Not Founc | 1 |
| Not Founc | Not Founc | 1 |
| Not Founc | Not Founc | 1 |
| High | High | 1 |
| Not Founc | Not Founc | 1 |
| Not Founc | Not Founc | 1 |
| Not Founc | Not Founc | 1 |
| Not Founc | Not Founc | 1 |
| Not Founc | Not Founc | 1 |
| Not Founc | Not Founc | 1 |
| Not Founc | Not Founc | 1 |
| High | High | 1 |
| Not Founc | Not Founc | 1 |
| Not Founc | Not Founc | 1 |
| Not Founc | Not Founc | 1 |
| Not Founc | Not Founc | 1 |
| High | High | 1 |
| Not Founc | Not Founc | 1 |
| Not Founc | Not Founc | 1 |
| Not Founc | Not Founc | 1 |
| Not Founc | Not Founc | 1 |
| Not Founc | Not Founc | 1 |
| Not Founc | Not Founc | 1 |
| Not Founc | Not Founc | 1 |
| Not Founc | Not Founc | 1 |
| High | High | 1 |
| Not Founc | Not Founc | 1 |
| Not Founc | Not Founc | 1 |
| Not Founc | Not Founc | 1 |
| Not Founc | Not Founc | 1 |

| | | |
|---|---|---|
| Not Founc | Not Founc | 1 |
| Not Founc | Not Founc | 1 |
| Not Founc | Not Founc | 1 |
| Not Founc | Not Founc | 1 |
| Not Founc | Not Founc | 1 |
| Not Founc | Not Founc | 1 |
| Not Founc | Not Founc | 1 |
| Not Founc | Not Founc | 1 |
| Not Founc | Not Founc | 1 |
| High | High | 1 |
| Not Founc | Not Founc | 1 |
| Not Founc | Not Founc | 1 |
| Not Founc | Not Founc | 1 |
| Not Founc | Not Founc | 1 |
| High | High | 1 |
| Not Founc | Not Founc | 1 |
| Not Founc | Not Founc | 1 |
| High | High | 1 |
| Not Founc | Not Founc | 1 |
| High | High | 1 |
| Not Founc | Not Founc | 1 |
| High | High | 1 |
| Not Founc | Not Founc | 1 |
| High | High | 1 |
| Not Founc | Not Founc | 1 |
| Not Founc | Not Founc | 1 |
| Not Founc | Not Founc | 1 |
| High | High | 1 |
| Not Founc | Not Founc | 1 |
| Not Founc | Not Founc | 1 |
| Not Founc | Not Founc | 1 |

| | | |
|---|---|---|
| Not Found | Not Found | 1 |
| Not Found | Not Found | 1 |
| High | High | 1 |
| Not Found | Not Found | 1 |
| Not Found | Not Found | 1 |
| High | High | 1 |
| High | High | 1 |
| High | High | 1 |
| Not Found | Not Found | 1 |
| High | High | 1 |
| Not Found | Not Found | 1 |
| High | High | 1 |
| High | High | 1 |
| Not Found | Not Found | 1 |
| Not Found | Not Found | 1 |
| Not Found | Not Found | 1 |
| Not Found | Not Found | 1 |
| High | High | 1 |
| Not Found | Not Found | 1 |
| Not Found | Not Found | 1 |
| Not Found | Not Found | 1 |
| Not Found | Not Found | 1 |
| Not Found | Not Found | 1 |
| Not Found | Not Found | 1 |
| Not Found | Not Found | 1 |
| Not Found | Not Found | 1 |
| Not Found | Not Found | 1 |
| Not Found | Not Found | 1 |
| Not Found | Not Found | 1 |
| Not Found | Not Found | 1 |
| Not Found | Not Found | 1 |

| | | | |
|---|---|---|---|
| Not Found | Not Found | 1 | |
| Not Found | Not Found | 1 | |
| High | High | 1 | |
| High | High | 1 | |
| High | High | 1 | |
| Not Found | Not Found | 1 | |
| Not Found | Not Found | 1 | |
| Not Found | Not Found | 1 | |
| Not Found | Not Found | 1 | |
| High | High | 1 | |
| Not Found | Not Found | 1 | |
| High | High | 1 | |
| Not Found | Not Found | 1 | Acetyl [N- |
| High | High | 1 | |
| Not Found | Not Found | 1 | |
| Not Found | Not Found | 1 | |
| Not Found | Not Found | 1 | |
| Not Found | Not Found | 1 | |
| High | High | 1 | |
| Not Found | Not Found | 1 | |
| Not Found | Not Found | 1 | |
| Not Found | Not Found | 1 | |
| Not Found | Not Found | 1 | |
| Not Found | Not Found | 1 | |
| Not Found | Not Found | 1 | |
| Not Found | Not Found | 1 | |
| Not Found | Not Found | 1 | |
| Not Found | Not Found | 1 | |
| Not Found | Not Found | 1 | |
| Not Found | Not Found | 1 | |
| Not Found | Not Found | 1 | |

| | | |
|---|---|---|
| Not Founc | Not Founc | 1 |
| Not Founc | Not Founc | 1 |
| Not Founc | Not Founc | 1 |
| High | High | 1 |
| Not Founc | Not Founc | 1 |
| Not Founc | Not Founc | 1 |
| High | High | 1 |
| Not Founc | Not Founc | 1 |
| Not Founc | Not Founc | 1 |
| Not Founc | Not Founc | 1 |
| High | High | 1 |
| Not Founc | Not Founc | 1 |
| Not Founc | Not Founc | 1 |
| Not Founc | Not Founc | 1 |
| Not Founc | Not Founc | 1 |
| High | High | 1 |
| High | High | 1 |
| Not Founc | Not Founc | 1 |
| High | High | 1 |
| High | High | 1 |
| Not Founc | Not Founc | 1 |
| High | High | 1 |
| Not Founc | Not Founc | 1 |
| Not Founc | Not Founc | 1 |
| High | High | 1 |
| Not Founc | Not Founc | 1 |
| Not Founc | Not Founc | 1 |
| High | High | 1 |
| Not Founc | Not Founc | 1 |
| Not Founc | Not Founc | 1 |
| Not Founc | Not Founc | 1 |

| | | | |
|---|---|---|---|
| High | High | 1 | |
| High | High | 1 | |
| High | High | 1 | |
| Not Founc | Not Founc | 1 | |
| Not Founc | Not Founc | 1 | |
| Not Founc | Not Founc | 1 | |
| Not Founc | Not Founc | 1 | |
| High | High | 1 | |
| High | High | 1 | |
| Not Founc | Not Founc | 1 | |
| High | High | 1 | |
| Not Founc | Not Founc | 1 | |
| Not Founc | Not Founc | 1 | |
| Not Founc | Not Founc | 1 | |
| High | High | 1 | |
| Not Founc | Not Founc | 1 | |
| High | High | 1 | |
| Not Founc | Not Founc | 1 | |
| Not Founc | Not Founc | 1 | |
| Not Founc | Not Founc | 1 | |
| Not Founc | Not Founc | 1 | |
| High | High | 1 | |
| High | High | 1 | |
| Not Founc | Not Founc | 1 | |
| Not Founc | Not Founc | 1 | |
| High | High | 1 | |
| High | High | 1 | |
| High | High | 1 | Acetyl [N- |
| Not Founc | Not Founc | 1 | |
| High | High | 1 | |
| Not Founc | Not Founc | 1 | |

| | | |
|---|---|---|
| High | High | 1 |
| Not Founc | Not Founc | 1 |
| Not Founc | Not Founc | 1 |
| Not Founc | Not Founc | 1 |
| Not Founc | Not Founc | 1 |
| Not Founc | Not Founc | 1 |
| High | High | 1 |
| High | High | 1 |
| High | High | 1 |
| Not Founc | Not Founc | 1 |
| Not Founc | Not Founc | 1 |
| Not Founc | Not Founc | 1 |
| Not Founc | Not Founc | 1 |
| Not Founc | Not Founc | 1 |
| High | High | 1 |
| Not Founc | Not Founc | 1 |
| Not Founc | Not Founc | 1 |
| Not Founc | Not Founc | 1 |
| Not Founc | Not Founc | 1 |
| Not Founc | Not Founc | 1 |
| Not Founc | Not Founc | 1 |
| Not Founc | Not Founc | 1 |
| High | High | 1 |
| Not Founc | Not Founc | 1 |
| Not Founc | Not Founc | 1 |
| Not Founc | Not Founc | 1 |
| High | High | 1 |
| High | High | 1 |
| High | High | 1 |
| High | High | 1 |
| Not Founc | Not Founc | 1 |

| | | |
|---|---|---|
| Not Founc | Not Founc | 1 |
| Not Founc | Not Founc | 1 |
| Not Founc | Not Founc | 1 |
| High | High | 1 |
| High | High | 1 |
| Not Founc | Not Founc | 1 |
| Not Founc | Not Founc | 1 |
| Not Founc | Not Founc | 1 |
| High | High | 1 |
| Not Founc | Not Founc | 1 |
| Not Founc | Not Founc | 1 |
| Not Founc | Not Founc | 1 |
| Not Founc | Not Founc | 1 |
| Not Founc | Not Founc | 1 |
| High | High | 1 |
| Not Founc | Not Founc | 1 |
| Not Founc | Not Founc | 1 |
| High | High | 1 |
| Not Founc | Not Founc | 1 |
| Not Founc | Not Founc | 1 |
| Not Founc | Not Founc | 1 |
| High | High | 1 |
| High | High | 1 |
| Not Founc | Not Founc | 1 |
| High | High | 1 |
| Not Founc | Not Founc | 1 |
| Not Founc | Not Founc | 1 |
| Not Founc | Not Founc | 1 |
| Not Founc | Not Founc | 1 |
| Not Founc | Not Founc | 1 |
| Not Founc | Not Founc | 1 |

| | | | |
|---|---|---|---|
| High | High | 1 | |
| Not Founc | Not Founc | 1 | |
| Not Founc | Not Founc | 1 | |
| Not Founc | Not Founc | 1 | |
| Not Founc | Not Founc | 1 | |
| Not Founc | Not Founc | 1 | |
| Not Founc | Not Founc | 1 | |
| High | High | 1 | |
| Not Founc | Not Founc | 1 | |
| Not Founc | Not Founc | 1 | |
| Not Founc | Not Founc | 1 | |
| Not Founc | Not Founc | 1 | |
| High | High | 1 | |
| Not Founc | Not Founc | 1 | |
| High | High | 1 | |
| Not Founc | Not Founc | 1 | |
| Not Founc | Not Founc | 1 | Acetyl [N- |
| Not Founc | Not Founc | 1 | |
| High | High | 1 | |
| Not Founc | Not Founc | 1 | |
| High | High | 1 | |
| Not Founc | Not Founc | 1 | |
| High | High | 1 | |
| High | High | 1 | |
| Not Founc | Not Founc | 1 | Acetyl [N- |
| High | High | 1 | |
| High | High | 1 | |
| Not Founc | Not Founc | 1 | |
| High | High | 1 | Acetyl [N- |
| Not Founc | Not Founc | 1 | |
| Not Founc | Not Founc | 1 | |

| | | |
|---|---|---|
| High | High | 1 |
| High | High | 1 |
| Not Found | Not Found | 1 |
| Not Found | Not Found | 1 |
| Not Found | Not Found | 1 |
| Not Found | Not Found | 1 |
| High | High | 1 |
| Not Found | Not Found | 1 |
| Not Found | Not Found | 1 |
| Not Found | Not Found | 1 |
| Not Found | Not Found | 1 |
| Not Found | Not Found | 1 |
| High | High | 1 |
| Not Found | Not Found | 1 |
| High | High | 1 |
| Not Found | Not Found | 1 |
| Not Found | Not Found | 1 |
| High | High | 1 |
| Not Found | Not Found | 1 |
| Not Found | Not Found | 1 |
| Not Found | Not Found | 1 |
| High | High | 1 |
| High | High | 1 |
| High | High | 1 |
| Not Found | Not Found | 1 |
| High | High | 1 |
| Not Found | Not Found | 1 |
| Not Found | Not Found | 1 |
| High | High | 1 |
| Not Found | Not Found | 1 |
| High | High | 1 |

| | | |
|---|---|---|
| Not Founc | Not Founc | 1 |
| Not Founc | Not Founc | 1 |
| Not Founc | Not Founc | 1 |
| Not Founc | Not Founc | 1 |
| High | High | 1 |
| Not Founc | Not Founc | 1 |
| Not Founc | Not Founc | 1 |
| High | High | 1 |
| Not Founc | Not Founc | 1 |
| Not Founc | Not Founc | 1 |
| Not Founc | Not Founc | 1 |
| Not Founc | Not Founc | 1 |
| High | High | 1 |
| High | High | 1 |
| High | High | 1 |
| Not Founc | Not Founc | 1 |
| High | High | 1 |
| Not Founc | Not Founc | 1 |
| Not Founc | Not Founc | 1 |
| High | High | 1 |
| High | High | 1 |
| High | High | 1 |
| Not Founc | Not Founc | 1 |
| High | High | 1 |
| Not Founc | Not Founc | 1 |
| Not Founc | Not Founc | 1 |
| High | High | 1 |
| Not Founc | Not Founc | 1 |
| Not Founc | Not Founc | 1 |
| High | High | 1 |
| Not Founc | Not Founc | 1 |

| | | | |
|---|---|---|---|
| High | High | 1 | |
| Not Founc | Not Founc | 1 | |
| Not Founc | Not Founc | 1 | |
| High | High | 1 | |
| Not Founc | Not Founc | 1 | |
| Not Founc | Not Founc | 1 | |
| High | High | 1 | |
| High | High | 1 | |
| Not Founc | Not Founc | 1 | |
| Not Founc | Not Founc | 1 | |
| Not Founc | Not Founc | 1 | |
| Not Founc | Not Founc | 1 | |
| Not Founc | Not Founc | 1 | |
| Not Founc | Not Founc | 1 | |
| High | High | 1 | |
| Not Founc | Not Founc | 1 | |
| High | High | 1 | |
| High | High | 1 | |
| High | High | 1 | |
| Not Founc | Not Founc | 1 | |
| Not Founc | Not Founc | 1 | |
| High | High | 1 | |
| High | High | 1 | |
| Not Founc | Not Founc | 1 | |
| Not Founc | Not Founc | 1 | |
| Not Founc | Not Founc | 1 | |
| Not Founc | Not Founc | 1 | |
| Not Founc | Not Founc | 1 | |
| Not Founc | Not Founc | 1 | |
| Not Founc | Not Founc | 1 | Acetyl [N- |
| Not Founc | Not Founc | 1 | |

| | | |
|---|---|---|
| Not Founc | Not Founc | 1 |
| High | High | 1 |
| Not Founc | Not Founc | 1 |
| High | High | 1 |
| Not Founc | Not Founc | 1 |
| Not Founc | Not Founc | 1 |
| Not Founc | Not Founc | 1 |
| High | High | 1 |
| High | High | 1 |
| Not Founc | Not Founc | 1 |
| High | High | 1 |
| Not Founc | Not Founc | 1 |
| Not Founc | Not Founc | 1 |
| High | High | 1 |
| Not Founc | Not Founc | 1 |
| Not Founc | Not Founc | 1 |
| Not Founc | Not Founc | 1 |
| Not Founc | Not Founc | 1 |
| Not Founc | Not Founc | 1 |
| High | High | 1 |
| Not Founc | Not Founc | 1 |
| Not Founc | Not Founc | 1 |
| High | High | 1 |
| Not Founc | Not Founc | 1 |
| High | High | 1 |
| Not Founc | Not Founc | 1 |
| Not Founc | Not Founc | 1 |
| Not Founc | Not Founc | 1 |
| High | High | 1 |
| High | High | 1 |
| Not Founc | Not Founc | 1 |

| | | |
|---|---|---|
| High | High | 1 |
| Not Found | Not Found | 1 |
| Not Found | Not Found | 1 |
| Not Found | Not Found | 1 |
| Not Found | Not Found | 1 |
| Not Found | Not Found | 1 |
| High | High | 1 |
| Not Found | Not Found | 1 |
| Not Found | Not Found | 1 |
| Not Found | Not Found | 1 |
| High | High | 1 |
| High | High | 1 |
| Not Found | Not Found | 1 |
| Not Found | Not Found | 1 |
| High | High | 1 |
| Not Found | Not Found | 1 |
| Not Found | Not Found | 1 |
| Not Found | Not Found | 1 |
| Not Found | Not Found | 1 |
| Not Found | Not Found | 1 |
| Not Found | Not Found | 1 |
| Not Found | Not Found | 1 |
| Not Found | Not Found | 1 |
| Not Found | Not Found | 1 |
| High | High | 1 |
| Not Found | Not Found | 1 |
| High | High | 1 |
| Not Found | Not Found | 1 |
| Not Found | Not Found | 1 |
| Not Found | Not Found | 1 |
| Not Found | Not Found | 1 |

| | | |
|---|---|---|
| Not Founc | Not Founc | 1 |
| Not Founc | Not Founc | 1 |
| Not Founc | Not Founc | 1 |
| Not Founc | Not Founc | 1 |
| High | High | 1 |
| Not Founc | Not Founc | 1 |
| Not Founc | Not Founc | 1 |
| Not Founc | Not Founc | 1 |
| Not Founc | Not Founc | 1 |
| Not Founc | Not Founc | 1 |
| High | High | 1 |
| Not Founc | Not Founc | 1 |
| Not Founc | Not Founc | 1 |
| Not Founc | Not Founc | 1 |
| Not Founc | Not Founc | 1 |
| Not Founc | Not Founc | 1 |
| Not Founc | Not Founc | 1 |
| Not Founc | Not Founc | 1 |
| High | High | 1 |
| Not Founc | Not Founc | 1 |
| Not Founc | Not Founc | 1 |
| High | High | 1 |
| Not Founc | Not Founc | 1 |
| High | High | 1 |
| Not Founc | Not Founc | 1 |
| Not Founc | Not Founc | 1 |
| Not Founc | Not Founc | 1 |
| High | High | 1 |
| Not Founc | Not Founc | 1 |
| Not Founc | Not Founc | 1 |
| Not Founc | Not Founc | 1 |

| | | | |
|---|---|---|---|
| Not Founc | Not Founc | 1 | |
| Not Founc | Not Founc | 1 | |
| High | High | 1 | |
| Not Founc | Not Founc | 1 | |
| High | High | 1 | |
| High | High | 1 | |
| Not Founc | Not Founc | 1 | |
| Not Founc | Not Founc | 1 | |
| Not Founc | Not Founc | 1 | |
| High | High | 1 | |
| High | High | 1 | |
| High | High | 1 | |
| Not Founc | Not Founc | 1 | |
| High | High | 1 | |
| Not Founc | Not Founc | 1 | |
| High | High | 1 | |
| Not Founc | Not Founc | 1 | |
| Not Founc | Not Founc | 1 | |
| Not Founc | Not Founc | 1 | |
| Not Founc | Not Founc | 1 | |
| Not Founc | Not Founc | 1 | |
| High | High | 1 | |
| Not Founc | Not Founc | 1 | |
| High | High | 1 | |
| High | High | 1 | |
| Not Founc | Not Founc | 1 | |
| Not Founc | Not Founc | 1 | |
| Not Founc | Not Founc | 1 | |
| High | High | 1 | |
| Not Founc | Not Founc | 1 | Acetyl [N- |
| Not Founc | Not Founc | 1 | |

| | | |
|---|---|---|
| High | High | 1 |
| Not Found | Not Found | 1 |
| High | High | 1 |
| Not Found | Not Found | 1 |
| High | High | 1 |
| High | High | 1 |
| Not Found | Not Found | 1 |
