## Supplementary figures and images for "Interspecies isobaric labeling-based quantitative proteomics reveals protein changes in the ovary of *Aedes aegypti* co-infected with ZIKV and Wolbachia"

### zika peptides.png

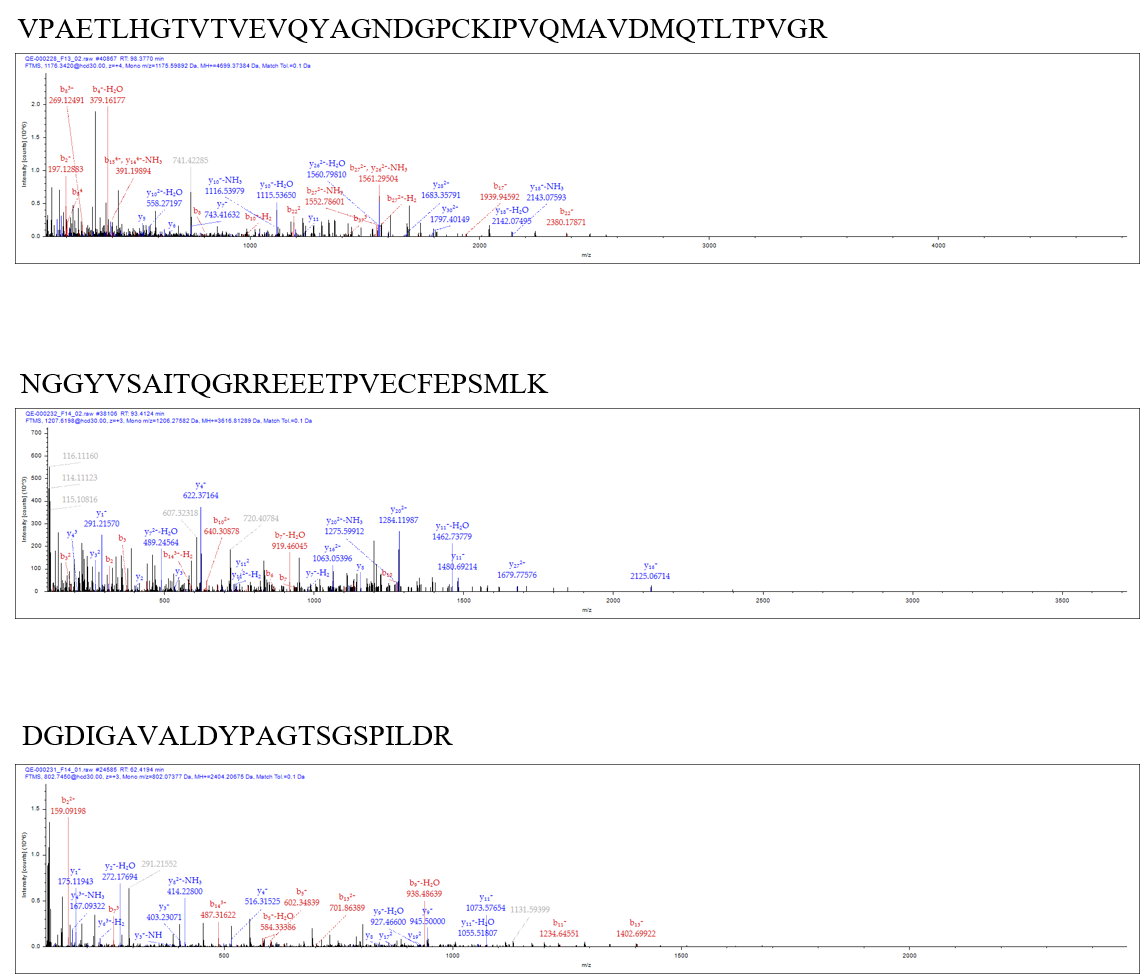
